# The mouse HP1 proteins are essential for preventing liver tumorigenesis

**DOI:** 10.1101/441279

**Authors:** Nehmé Saksouk, Shefqet Hajdari, Marine Pratlong, Célia Barrachina, Céline Graber, Aliki Zavoriti, Amélie Sarrazin, Nelly Pirot, Jean-Yohan Noël, Lakhdar Khellaf, Eric Fabbrizio, Eric Julien, Florence M. Cammas

**Affiliations:** IRCM, Institut de Recherche en Cancérologie de Montpellier, Montpellier, F-34298, France; INSERM, U1194, Montpellier, F-34298, France; Université de Montpellier, Montpellier, F-34090, France; Institut régional du Cancer de Montpellier, Montpellier, F-34298, France; MGX, Biocampus Montpellier, CNRS, INSERM, Univ Montpellier, Montpellier, France; IGBMC, Institut de Génétique, de Biologie Moléculaire et Cellulaire, Illkirch, France; MRI, BioCampus Montpellier, CNRS, INSERM, Univ Montpellier, Montpellier, France; CNRS, Route de Mende, Montpellier, France

**Author notes:** Corresponding author: Florence Cammas. Contributed equally to the work (Nehmé Saksouk, Shefqet Hajdari).

**Keywords:** chromatin, HP1, cancer, liver, transcriptional silencing, endogenous retrovirus

## Abstract

Chromatin organization is essential for appropriate interpretation of the genetic information. Here, we demonstrated that the chromatin associated proteins HP1 are dispensable for cell survival but are essential within hepatocytes to prevent liver tumor development. Molecular characterization of pre-malignant HP1-Triple KO livers revealed that HP1 are essential for the maintenance of the structural organization of heterochromatin but surprisingly, not for several well known heterochromatin functions such as the maintenance of the genome stability nor the regulation of major satellite repeat expression within liver. We further show that some specific retrotransposons, mainly of the ERV family, get reactivated in HP1-TKO livers correlating, in some cases, with the activation of the adjacent genes. We present evidence that this reactivation of ERV relies on the HP1-dependent ability of the corepressor TRIM28 to regulate KRAB-ZFP repressive activity. Intriguingly, we found that in contrast to the observation in young animals, the HP1-dependent maintenance of ERV silencing becomes independent of TRIM28 in old animals. Finally, we showed that HP1 are also essential directly or indirectly for the regulation of single genes with most of them having well characterized functions in liver homeostasis such as regulation of the redox and endoplasmic reticulum equilibrium, lipid metabolism and steroid biosynthesis.

Altogether, our findings indicate that HP1 proteins, through the modulation of multiple chromatin-associated events both within the heterochromatic and euchromatic compartments, act as guardians of liver homeostasis to prevent tumor development.

## Introduction

Chromatin dynamic organization is essential for the interpretation of genetic information in a cell-type and tissue-specific manner ^1^. Alteration of this organization can have devastating consequences, as evidenced by the large number of diseases induced by mutations in chromatin-associated proteins ^2,3^, as well as by the dramatic changes in chromatin organization observed in cancer cells ^4^. Although extensively studied in the past three decades, it is still largely unknown how chromatin organization is regulated and involved in whole organism homeostasis.

Chromatin can be divided according to its structural and functional features in euchromatin and heterochromatin. Euchromatin displays low level of compaction, is highly enriched in genes, and is transcriptionally competent. Conversely heterochromatin is highly compacted, enriched in repetitive DNA sequences, and mostly silent ^5^. Heterochromatin Protein 1 (HP1) proteins were first isolated as major heterochromatin components in *Drosophila* ^6^. These proteins are highly conserved from yeast to mammals which express three isoforms (HP1α, HP1β and HP1γ) that are distributed in both eu- and heterochromatin. These proteins are characterized by a N-terminal chromodomain (CD) involved in recognition of the heterochromatin-associated histone marks H3 lysine-9 di- or trimethylated (H3K9me2/3), and a C-terminal chromoshadow domain (CSD), which, through dimerization, constitutes a platform for interaction with many protein partners. These two domains are separated by the hinge domain that is crucial for HP1 association with RNA and recruitment to heterochromatin ^7,8^. Thus, HP1 proteins, through this structural organization, are at the crossroads of the structural and functional organization of chromatin. Accordingly, HP1 are important for heterochromatin organization and silencing, chromosome segregation, regulation of gene expression, DNA repair and DNA replication ^9–11^. Functionally, HP1 proteins are essential for embryonic development in several organisms, including *Drosophila* ^12^, C. *elegans* ^13^ and the mouse (our unpublished data). HP1α is essential for the plasticity of T helper (Th2) lymphocytes ^14^, HP1β for neuro-muscular junctions ^15^ and HP1γ for spermatogenesis ^16,17^. Several studies also suggested a correlation between the level of HP1 expression and cancer development and/or metastasis; however, how HP1 are involved in these processes remains largely to be clarified ^18,19^.

Liver chromatin organization has been well characterized in several physio-pathological conditions ^20^. In addition, several known HP1 partners, including the transcription cofactors TRIM24 and TRIM28, and the histone-lysine N-methyltransferase SUV39H1 have been shown to play key roles in hepatocytes ^21–25^. Together, this prompted us to further characterize chromatin organization and regulation in liver functions through the inactivation of all HP1 encoding genes specifically in mouse hepatocytes (HP1-TKO mice). We found that in mice, HP1 are critically required for preventing tumor development. We further identified several altered chromatin features in HP1-TKO animals, including heterochromatin organization, silencing of specific ERVs and gene expression that most likely are key players in the process of tumorigenesis. These data highlighted a new function of HP1 proteins as guardians of liver homeostasis through the modulation of various chromatin-associated events.

## RESULTS

### HP1 proteins are dispensable for hepatocyte proliferation and survival

To unravel HP1 *in vivo* functions, the HP1β and HP1γ encoding-genes (*Cbx1* and *Cbx3,* respectively) were inactivated in the liver of HP1αKO mice ^14^ using the Cre recombinase expressed under the control of the hepatocyte-specific albumin promoter ^26,27^ (Fig. 1A). Liver-specific excision of the *Cbx1* and *Cbx3* alleles was confirmed by PCR (Fig. 1B), and the level of HP1β and HP1γ protein expression was checked by western blotting. At 7 weeks post-partum as well as at middle-aged (3-6 months), the overall level of HP1β and HP1γ was decreased by about 60% in mutant as compared to controls livers (Fig. 1C). Immunofluorescence (IF) analysis of liver cryo-sections showed that HP1β and HP1γ expression was absent in about 60% of liver cells in mutant mice (Fig. 1D). As this percentage is similar to the estimated 60-70% hepatocyte fraction within liver, these findings indicated that both proteins were concomitantly depleted in most hepatocytes ^28^. As expected, HP1α was not expressed in mutant livers (Fig. 1B-C). These animals were thereafter called HP1-triple knockout (HP1-TKO). Histological analysis of liver sections from 7-week-old (young) and 3-6-month-old (middle-aged) control and HP1-TKO animals did not reveal any significant alteration of the structural organization of hepatocytes nor of the liver parenchyma (Supplementary Figure 1). In agreement with this observation, analysis of proliferation (Ki67) and apoptosis (Activated caspase 3) by immuno-histochemistry (IHC) of Tissue Micro Arrays (TMA) containing liver sections from young and middle-aged control and HP1-TKO mice did not reveal any significant difference between mutant and control animals (Fig. 1E). As HP1 have been shown to play critical roles in genome stability ^29,30^, IHC was also performed with an antibody against the phosphorylated form of H2AX (γH2AX), a marker of DNA damage ^31^. The number of γH2AX-positive cells was slightly higher, although not statistically significantly different in livers of HP1-TKO and control animals, suggesting that HP1 proteins depletion did not lead to major genomic instability within hepatocytes (Fig. 1E).

**Figure 1:**
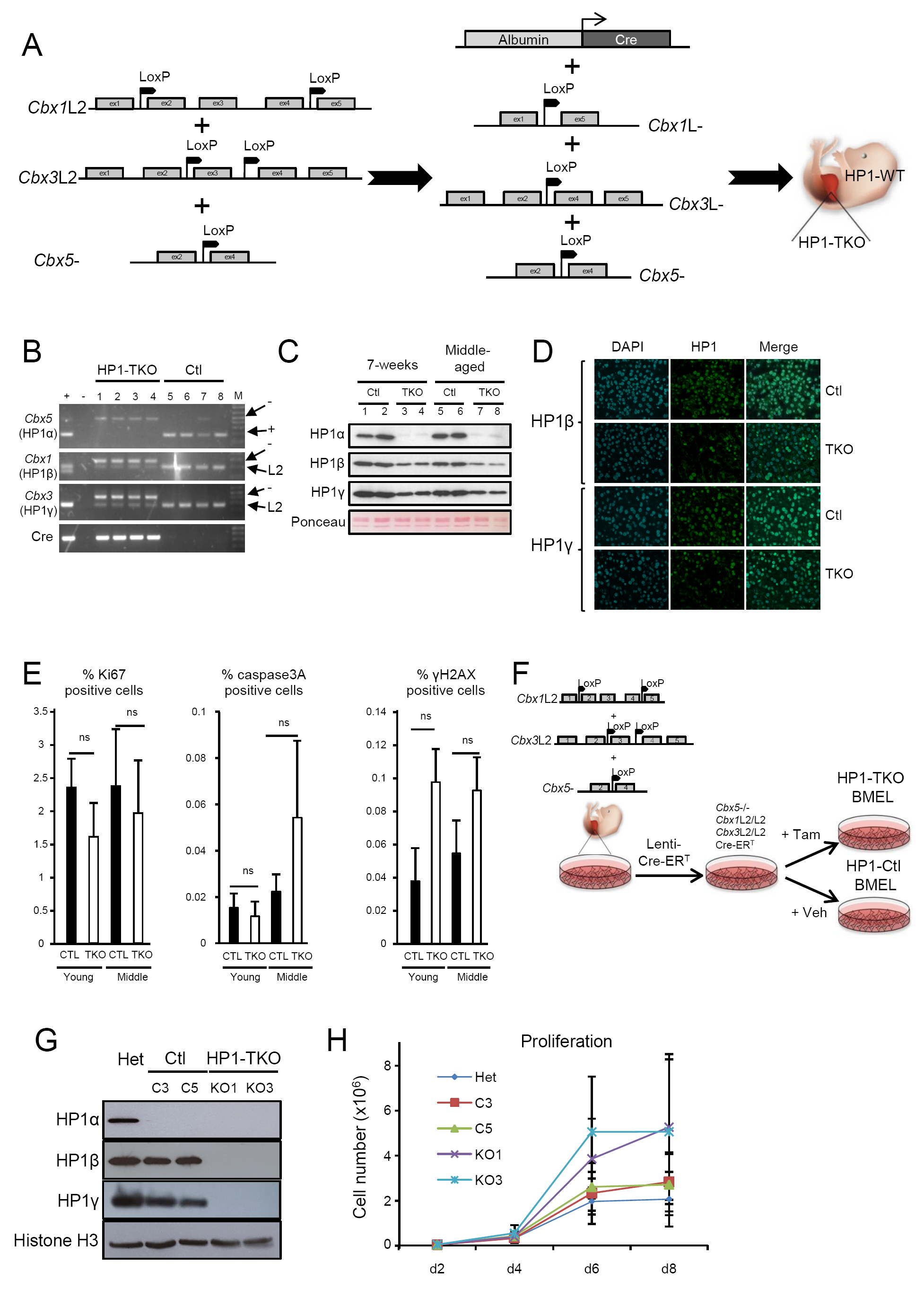
HP1 are not required for hepatocyte survival nor for liver organization and function. (A) Schematic representation of the strategy to inactivate the three HP1-encoding genes (*Cbx1, 3* and *5*) specifically in hepatocytes using the recombinase Cre expressed under the control of the albumin promoter. (B) The hepatocyte-specific excision (L-alleles) of the *Cbx1* (HP1β) and *Cbx3* (HP1γ) genes and ubiquitous excision of *Cbx5* (HP1α) in HP1-TKO (1-4) mice was verified by PCR. Controls were littermates with either L2 (*Cbx*1 and *Cbx*3) or “+” (*Cbx*5) alleles (5-8). Note that control #7 is *Cbx5*^L-/+^. (C) Western blot analysis of whole-cell extracts from liver samples confirmed the absence of HP1α and the decreased expression of HP1β and HP1γ, due to the hepatocytes-specific excision of the corresponding genes in HP1-TKO as compared to age-matched control mice. Ponceau staining was used as loading control. (D) Immuno-fluorescence analysis of liver tissue cryosections confirmed the absence of HP1β and HP1γ expression in about 60% of cells in 7-weeks old HP1-TKO mice compared with controls. (E) Immuno-histochemistry analysis of paraffin-embedded liver sections revealed no significant difference in proliferation (Ki67), apoptosis (caspase 3A), and DNA damage (γH2AX) between 7-week-old (n=5) and 3-6month-old mice (n= 5) HP1-TKO and control mice (n=7 and n= 4 respectively). The number of positive cells were normalized to the total number of cells in each section and graphs recapitulating these data are shown as the mean ± SEM. ns, no significant difference (Student’s t-test). (F) Schematic representation of the strategy to establish BMEL cells from *Cbx5*-/-; *Cbx1*L2/L2; *Cbx3*L2/L2 fetal livers and to inactivate the three HP1-encoding genes. (G) Western blot analysis of whole-cell extracts from BMEL cells confirmed the absence of all HP1 isoforms in HP1-TKO (*Cbx5*-/-; *Cbx1*L2/L2, *Cbx3*L2/L2; Cre-ERT treated with tamoxifen). “Het” were *Cbx5*+/-; *Cbx1*L2/L2 BMEL cells and “Ctl” were *Cbx5*-/-; *Cbx1*L2/L2; *Cbx3*L2/L2; Cre-ERT non treated with tamoxifen. (H) Proliferation curves of “Het", 2 Control clones (C3 and C5) and 2 HP1-TKO clones (KO1 and KO3). The graph represent the average of three independent experiments done in triplicates.

To unambiguously test the viability of hepatic cells in absence of any HP1 isoform, we established bipotential hepatic BMEL (Bipotential Mouse Embryonic Liver) cell lines according to the protocol described by Strick-Marchand & Weiss ^32^ and inactivated all HP1 encoding genes as illustrated on figure 1F-G. These cells, thereafter called HP1-TKO, were morphologically similar to control cells and had a tendency to proliferate faster than control cells (Fig. 1H). Altogether, these data demonstrated that in mouse, the three HP1 proteins are dispensable for hepatocyte survival both *in vivo* and *ex vivo* as well as for the appropriate structural organization of the liver parenchyma throughout life.

### HP1 proteins prevent tumor development in liver

Analysis of old mice (>44 weeks of age) showed that although HP1-TKO animals were morphologically indistinguishable from controls, 72.7% of females (n=11) and 87.5% of males (n=8) had developed liver tumors whereas none of the female (n=24) and 9.1% of male (n=44) controls did so (Fig. 2A-B). Analysis of the excision of the floxed *Cbx*1 and *Cbx*3 genes showed that tumors originated from HP1-TKO hepatocytes (Supplementary Fig. 2A). Histological analysis revealed that most old male (M-TKO) and female (F-TKO) HP1-TKO animals developed tumor nodules that could easily be distinguished from the rest of the liver parenchyma (Fig. 2C). These nodules were characterized by the presence of well-differentiated hepatocytes but without their specific trabecular organization, and thus, were identified as typical hepatocellular carcinoma (HCC). Analysis of cell proliferation (Ki67), apoptosis (activated caspase 3) and global response to DNA damage (γH2AX) showed a two-fold increase of cell proliferation in both the tumor (TKOT) and non-tumor (TKON) parts of HP1-TKO liver samples compared with control parts whereas no change was detected in the number of apoptotic-positive cells nor of positive γH2AX cells (Supplementary Fig. 2B). We then tested by RT-qPCR the expression of some genes frequently altered in human HCC ^33^. Although the expression of most of the tested genes was remarkably similar between controls and HP1-TKO livers both in the normal (TKON) and the tumoral parts (TKOT), *Arid1A, Trp53, E2f1* and *E2f7* were significantly over-expressed in HP1-TKO (Fig.2D and data not shown). α-fetoprotein (*Afp*), a marker of human HCC ^34^, was strongly over-expressed exclusively in the tumor tissue of three of the five tested tumors (Fig. 2D).

**Figure 2:**
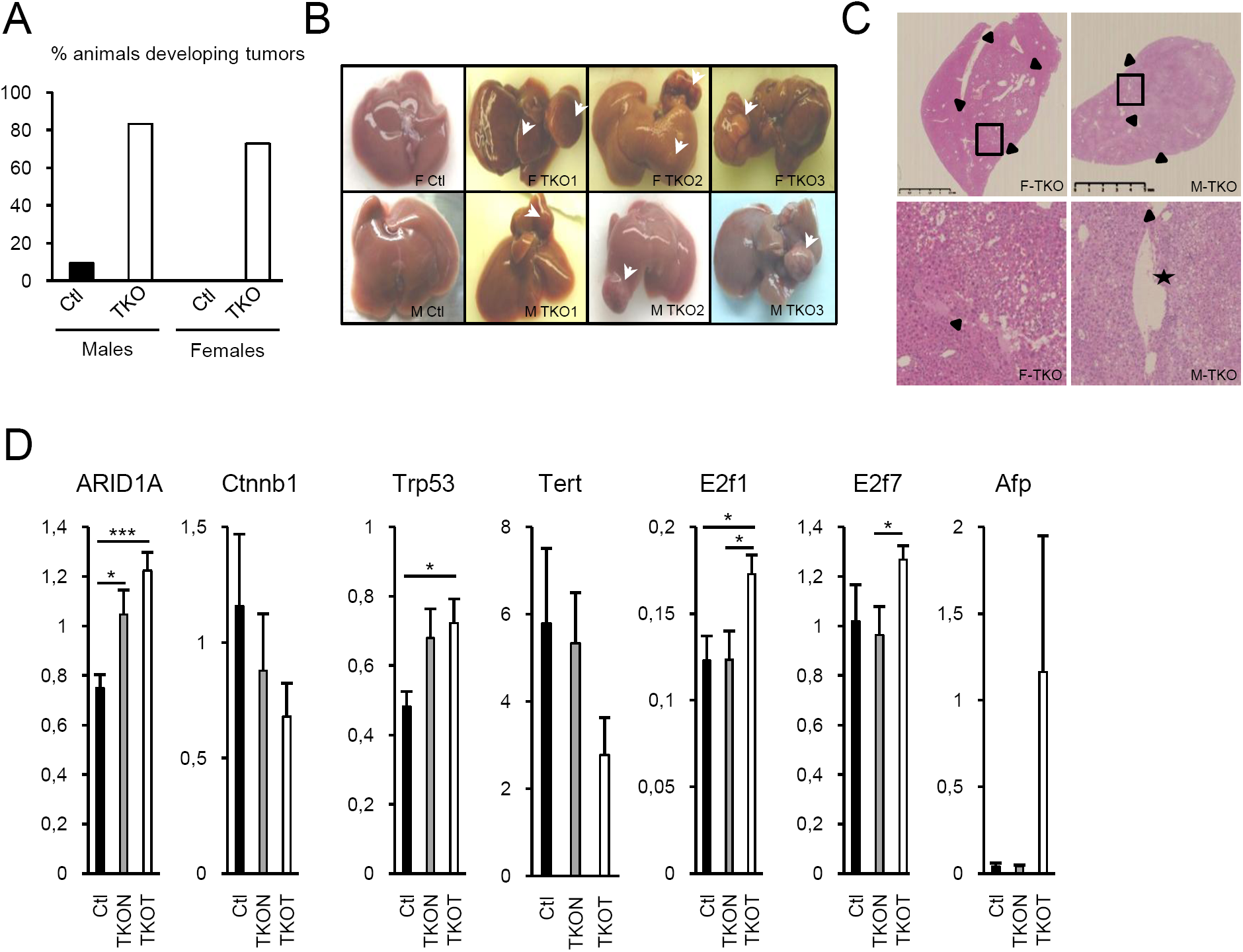
HP1 prevent tumor development in liver. (A) Controls (Ctl, n=67) and HP1-TKO (TKO, n=17) animals older than one year of age were sacrificed and the percentage of animals with tumors (morphological and histological analysis) was calculated. (B) Morphology of the livers with tumors (arrows) in three HP1-TKO females (F TKO1, 2 and 3) and three HP1-TKO males (M TKO1, 2 and 3) older than one year of age. The liver morphology of one age-matched control female and male is also shown (F Ctl and M Ctl, respectively). (C) Liver histological analysis (hematoxylin-eosin-Safran staining) of one representative HP1-TKO female (F-TKO) and one representative HP1-TKO male (M-TKO). Upper panels: tumor/liver parenchyma interface highlighted by arrowheads (low magnifications). Bottom panels: magnification (x 100) of the boxes in the upper panels showing the tumor in the right part of the images (thick plates of atypical hepatocytes). A venous tumor thrombus is also present (asterisk). (D) RT-qPCR analysis of the expression of the indicated genes in control (Ctl, n=5) and HP1-TKO (TKON: normal liver, n=6; TKOT: tumor, n=5) livers of animals older that one year.

Altogether these data clearly indicated that HP1 are essential within hepatocytes to prevent tumor development. To gain more insights into the cellular and molecular properties of HP1 underlying their protective functions against tumorigenesis, we initiated the characterization of HP1-TKO pre-malignant livers.

### Heterochromatin organization is altered in HP1-TKO hepatocytes

First, the level of different heterochromatin-associated histone marks was investigated by western blotting. H3K9me3 and H4K20me3, two marks of constitutive heterochromatin, were strongly decreased in the liver of 7-week-old and middle-aged HP1-TKO mice compared with age-matched controls. Conversely, no change of H3K27me3, a facultative heterochromatin mark, nor of H3K9me2, H4K20me2 and H4K20me1 was observed in these same samples (Fig. 3A). The decrease of H3K9me3 without any significant change of H3K27me3 nor of H3K4me3 was also observed in HP1-TKO BMEL cells (Fig. 3B). IF analysis in BMEL cells, indicated that only H3K9me3 associated with chromocenters (i.e., DAPI-dense structures that contain structural components of heterochromatin) was drastically reduced in HP1-TKO cells, whereas the labeling within euchromatin was not significantly affected (Fig. 3C). The level and distribution of 5-methyl cytosine (5mC) were not altered in HP1-TKO BMEL cells (Fig. 3C). Although these results suggested that the absence of the three HP1 proteins affected heterochromatin organization, chromocenters still clustered, but tended to be more associated with the nuclear periphery in HP1-TKO hepatocytes than in control cells (Fig. 3D). To more precisely quantify this observation, nuclei were divided in four co-centric areas in which the intensity of DAPI staining was measured using the cell profiler software (schema in Fig. 3D). In control nuclei, DAPI staining was roughly homogeneously distributed throughout the four areas whereas, conversely, in HP1-TKO nuclei, DAPI intensity increased progressively from the inner part to the external part (Fig. 3D). This indicated that in absence of HP1, heterochromatin tended to be more associated with the nuclear periphery than in control nuclei. This was not associated with any significant change of the level nor distribution of laminB1 (LamB1) as assessed by IF (Fig. 3C). We then measured the expression and the number of major satellite repeats that represent the main component of pericentromeric heterochromatin. Surprisingly this analysis revealed no significant alteration and even a tendency of these repeats to be down-regulated in absence of HP1 in both liver and BMEL cells as well as an unchanged number of these repeats within the genome (Fig. 3E-F and data not shown). These data demonstrated that in hepatocytes, HP1 proteins are essential for the maintenance of constitutive heterochromatin-associated histone marks and for the sub-nuclear organization of chromocenters but not for neither the expression nor the stability of major satellites.

**Figure 3:**
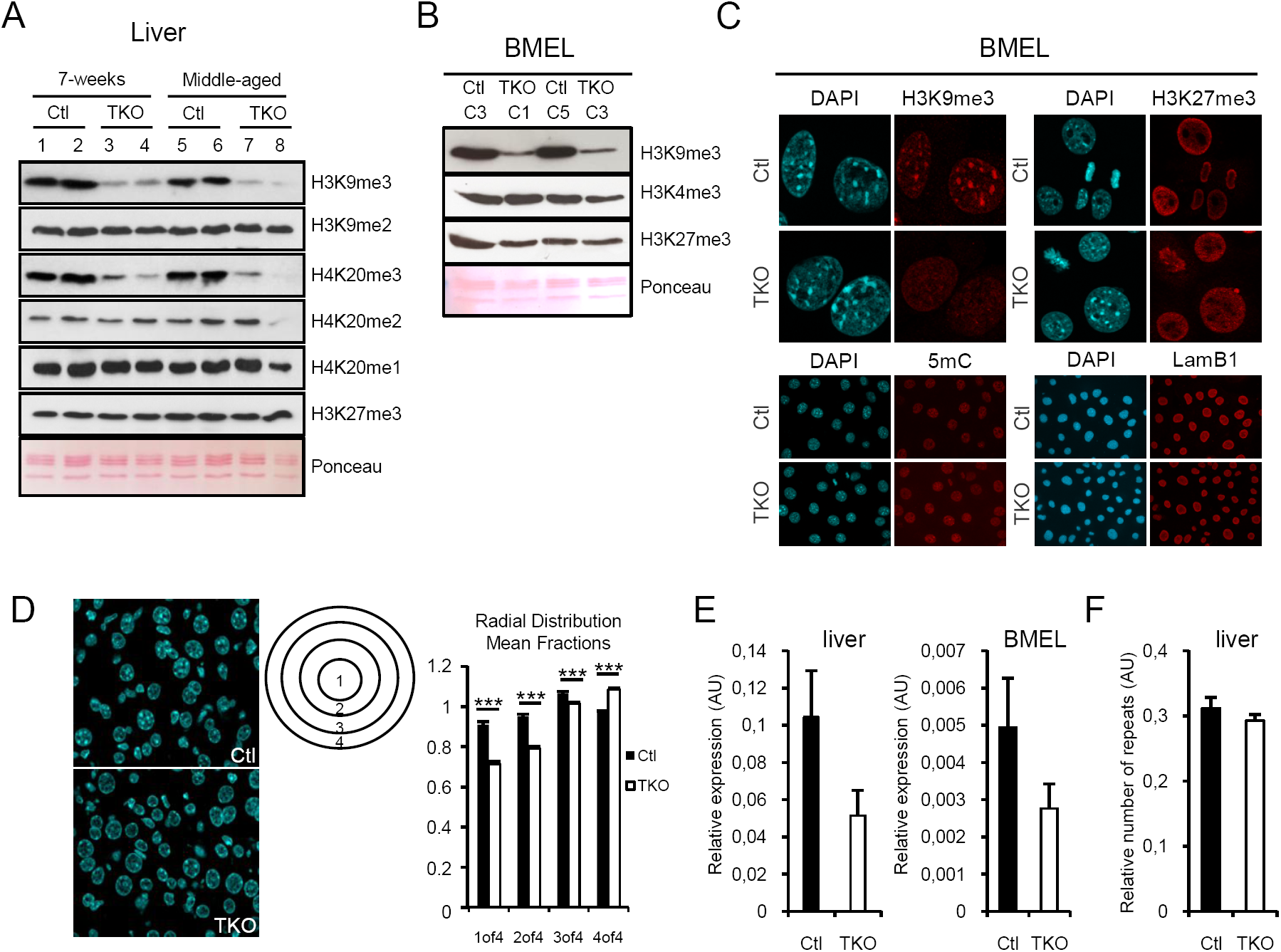
HP1 are essential for heterochromatin organization but not to regulate the expression of major satellites. (A) HP1 are essential for the maintenance of the two heterochromatin hallmarks H3K9me3 and H4K20me3 in hepatocytes. Western blot analysis of nuclear extracts from liver of 7-week-old and middle-aged (3-6-month-old) controls (Ctl: 1; 2; 5; 6) and HP1-TKO (TKO: 3; 4; 7; 8) mice with antibodies against the indicated histone marks. Ponceau staining was used as loading control. (B) Western blot analysis of the indicated marks in BMEL cells. (C) IF analysis of H3K9me3, H3K27me3, 5mC and LamB1 in BMEL cells. (D) Loss of HP1 leads to a partial relocation of DAPI-dense regions towards the nuclear periphery. Representative images of paraffin-embedded liver tissue sections from 7-week-old control (Ctl) and HP1-TKO (TKO) mice stained with DAPI (63x magnification). To select mostly hepatocytes, only the largest nuclei with a size comprised between 70 and 150 µm^2^ and with a circular shape were selected for this analysis. 2D sections of nuclei were divided in four concentric areas (1 to 4) and DAPI staining intensity was quantified using the cell profiler software. The mean fractional intensity at a given radius was calculated as the fraction of the total intensity normalized to the fraction of pixels at a given radius in n=584 control and n=762 HP1-TKO (TKO) nuclei. Data are the mean ± SEM. ***p value <0.001. (E) Loss of the three HP1 proteins in hepatocytes did not affect the expression of major satellites. qPCR assays were performed using total RNA from livers of 7-week-old control (n=4) and HP1-TKO mice (n=4) and on control (Ctl) and HP1-TKO (TKO) BMEL. (F) Satellite repeats were quantified by qPCR on genomic DNA from the same animals as those used for (E).

### HP1 proteins are involved in the regulation of liver-specific gene expression programs

We then investigated the impact of HP1 loss on gene expression. An unbiased RNA-seq transcriptomic analysis was performed on libraries prepared from 7 week-old control and HP1-TKO liver RNA. This analysis showed that 1215 genes were differentially expressed (730 up-regulated and 485 down-regulated) between control and HP1-TKO liver samples (with a 1.5-fold threshold difference and an adjusted *P* ≤0.05) (Fig. 4A and supplementary Table 1).

**Table 1:** HP1-dependent p450 genes.

| Gene name | Log2 fold-change | padj | Redox | Endoplasmic reticulum | Drug metabolism | Lipid metabolism | Steroid synthesis |
| --- | --- | --- | --- | --- | --- | --- | --- |
| Cyp2b10 | 4.06 | 1.41E-38 | 1 | 1 | 0 | 0 | 1 |
| Cyp2b9 | 2.68 | 4.51E-10 | 1 | 1 | 0 | 0 | 1 |
| Cyp2b13 | 1.80 | 0.000120 | 0 | 0 | 0 | 0 | 1 |
| Cyp4f16 | 1.63 | 1.36E-09 | 0 | 0 | 0 | 0 | 0 |
| Cyp2d12 | 1.38 | 0.000468 | 0 | 0 | 0 | 0 | 1 |
| Cyp2a4 | 1.09 | 0.0342 | 0 | 0 | 0 | 0 | 0 |
| Cyp2a22 | 1.06 | 0.000559 | 0 | 0 | 0 | 0 | 0 |
| Cyp2f2 | -0.65 | 0.00318 | 1 | 1 | 0 | 0 | 0 |
| Cyp4f13 | -0.67 | 0.0151 | 0 | 0 | 0 | 0 | 0 |
| Cyp2r1 | -0.72 | 0.0173 | 1 | 1 | 0 | 0 | 0 |
| Cyp27a1 | -0.83 | 1.30E-05 | 1 | 0 | 0 | 0 | 0 |
| Cyp2d37-ps | -0.83 | 0.0319 | 0 | 0 | 0 | 0 | 0 |
| Cyp3a25 | -0.85 | 0.00222 | 1 | 1 | 0 | 0 | 1 |
| Cyp39a1 | -0.88 | 0.00164 | 1 | 1 | 0 | 1 | 0 |
| Cyp2e1 | -0.94 | 7.81E-05 | 1 | 1 | 1 | 0 | 1 |
| Cyp2d26 | -0.95 | 2.57E-05 | 1 | 1 | 0 | 0 | 1 |
| Cyp2a5 | -0.97 | 0.00129 | 0 | 0 | 0 | 0 | 0 |
| Cyp2d13 | -1.00 | 0.00445 | 0 | 0 | 0 | 0 | 0 |
| Cyp1a2 | -1.25 | 3.65E-12 | 1 | 1 | 1 | 1 | 1 |
| Cyp2c53-ps | -1.31 | 0.01178 | 0 | 0 | 0 | 0 | 0 |
| Cyp2d40 | -1.42 | 6.83E-06 | 0 | 0 | 0 | 0 | 1 |
| Cyp46a1 | -1.64 | 0.000629 | 1 | 1 | 0 | 1 | 0 |
| Cyp3a59 | -2.15 | 3.62E-17 | 1 | 0 | 0 | 0 | 0 |
| Cyp2c44 | -2.20 | 2.18E-20 | 0 | 0 | 0 | 0 | 1 |
| Cyp2c29 | -2.60 | 7.65E-25 | 1 | 1 | 0 | 0 | 1 |

(1) found and (0) not found according to the David Gene Ontology software.

**Figure 4:**
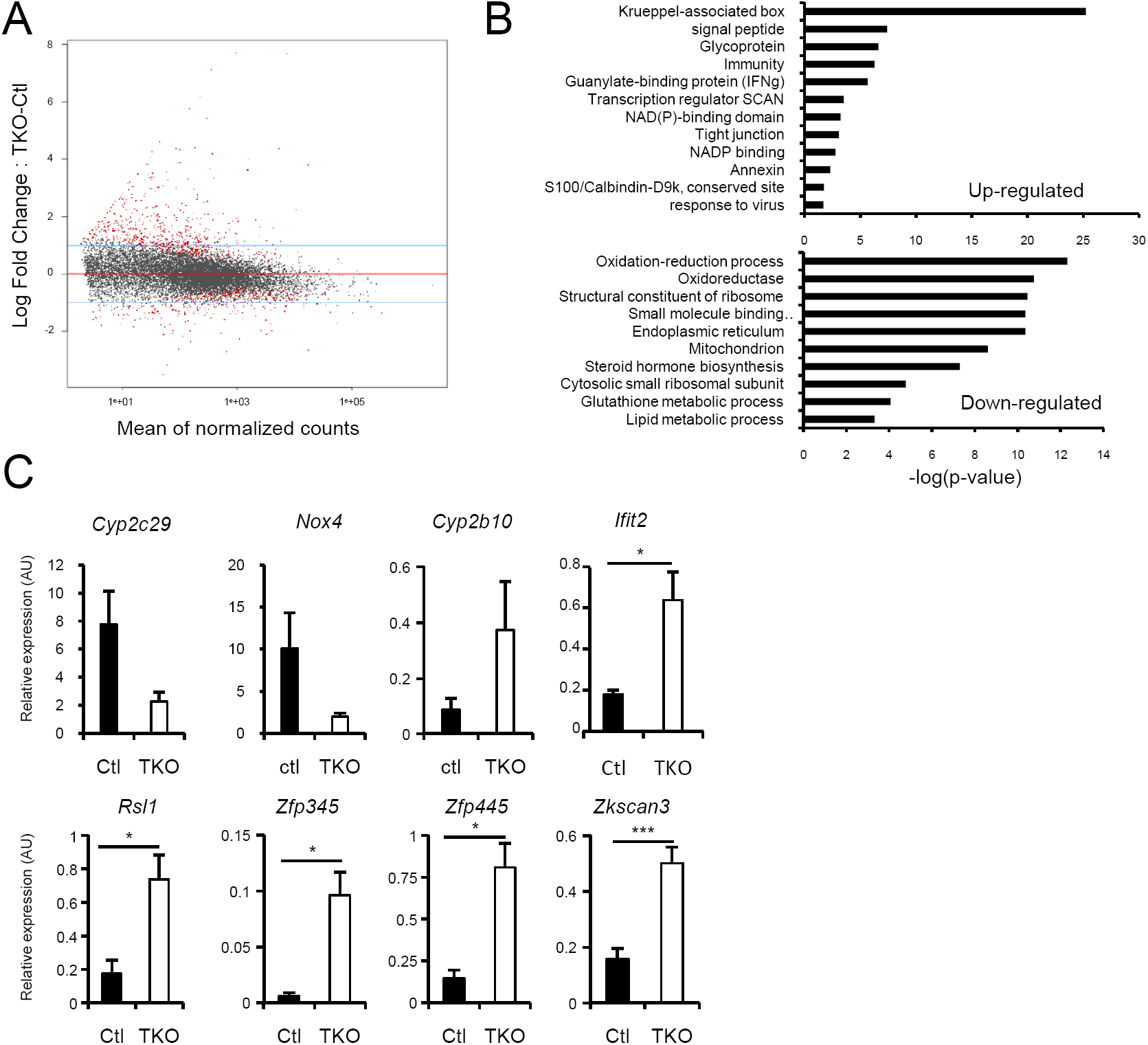
HP1 are essential regulators of gene expression in liver. (A) MA plot after DSeq2 normalization of RNA-seq data from 7-week-old control (n=3) and HP1-TKO (n=4) liver RNA samples. Red dots represent genes that are differentially expressed between control and HP1-TKO mice (adjusted p-value p <0.05). (B) Functional clustering of HP1-dependent genes using the DAVID Gene Ontology software. (C) Validation by RT-qPCR of the altered expression of the indicated genes. RNA was extracted from livers of 7-weeks control (Ctl, n=4) and HP1-TKO (TKO, n=4) animals. Data were normalized to *Hprt* expression and are shown as the mean ± SEM. *p value <0.05; ***p value <0.001 (Student’s t-test).

Analysis of differentially expressed genes (HP1-dependent genes) using David Gene Ontology (https://david.ncifcrf.gov/) and Gene Set Enrichment Analysis (GSEA; http://software.broadinstitute.org/gsea/index.jsp) programs revealed that several biological processes were significantly affected in HP1-TKO livers. The most striking feature of this analysis was the very high enrichment of genes encoding for the Krüppel Associated Box (KRAB) domain within up-regulated genes (*P* = 5.8E-26) (Fig. 4B & Supplementary Tables 1; 2 & 3). The up-regulation of several of these genes (*Rsl1, Zfp345, Zfp445* and *Zkscan3*) was validated by RT-qPCR in 7-week-old HP1-TKO and control livers (Fig. 4C). Beside these KRAB domain encoding genes, up-regulated genes were also enriched in genes belonging to the GO terms signal peptide, immunity, guanylate-binding protein, response to virus, etc… (Fig. 4B), strongly suggesting activation of an inflammatory response in HP1-TKO livers (Fig. 4B-C & Supplementary Table 4). Genes encoding for members of the p450 cytochrome (CYP) family were also strongly enriched in HP1-dependent genes with 7 up-regulated and 18 down-regulated amongst the 79 CYP genes detected in the present RNAseq analysis. In particular, 11 HP1-dependent genes encode for members of the CYP2 family were involved in Endoplasmic Reticulum (ER) and redox functions that are known to be particularly important for liver homeostasis ^35,36,37^ (Table 1). Moreover, *Nox4*, the gene encoding the nicotinamide adenine dinucleotide phosphate (NADPH) oxidase isoform most consistently associated with ER and ROS in liver ^38^, was significantly down-regulated in HP1-TKO as compared with control livers (Fig. 4C & Supplementary Table 1). It was thus not surprising that oxidation-reduction, ER, steroid hormone biosynthesis, lipid metabolic process were amongst the most affected functions in HP1-TKO livers (Fig. 4B & Supplementary Tables 2; 3; 5 and 6). The differential expression of several of these genes such as *Cyp2c29* and *Cyp2b10* (ER and redox), *Ifit2* (interferon γ signature) and *Nox4* (ROS production) was validated by RT-qPCR in 7 week-old HP1-TKO and control livers (Fig. 4C).

### HP1 loss leads to reactivation of specific endogenous retroviruses and over-expression of associated genes

As mentioned above, genes encoding for the KRAB domain were highly enriched in up-regulated genes in HP1-TKO livers. The KRAB domain is almost exclusively present in the KRAB-Zinc Finger Protein (KRAB-ZFP) family of transcriptional repressors ^39^. The best characterized genomic target of these repressors are themselves and retrotransposons of the endogenous retroviruses (ERV) family ^39,40^. We therefore investigated the expression of DNA repeats in our RNA-seq dataset. To this end, the coordinates of all annotated DNA repeats of the RepeatMasker database (mm10 assembly) were aligned against the RNA-seq reads and only those that could be assigned unambiguously to a specific genomic locus were analyzed. In total, 846 such repeats were deregulated in HP1-TKO livers compared with control livers with 71.3% being up-regulated and 28.7% down-regulated (Fig. 5A & Supplementary Table 7). Among up-regulated repeats, 59.4% were ERV, 19.2% long interspersed nuclear elements (LINEs) and 9.3% short interspersed elements (SINEs) supporting the hypothesis that HP1 were preferentially involved in ERV silencing (Fig. 5B & Supplementary Table 7).

**Figure 5:**
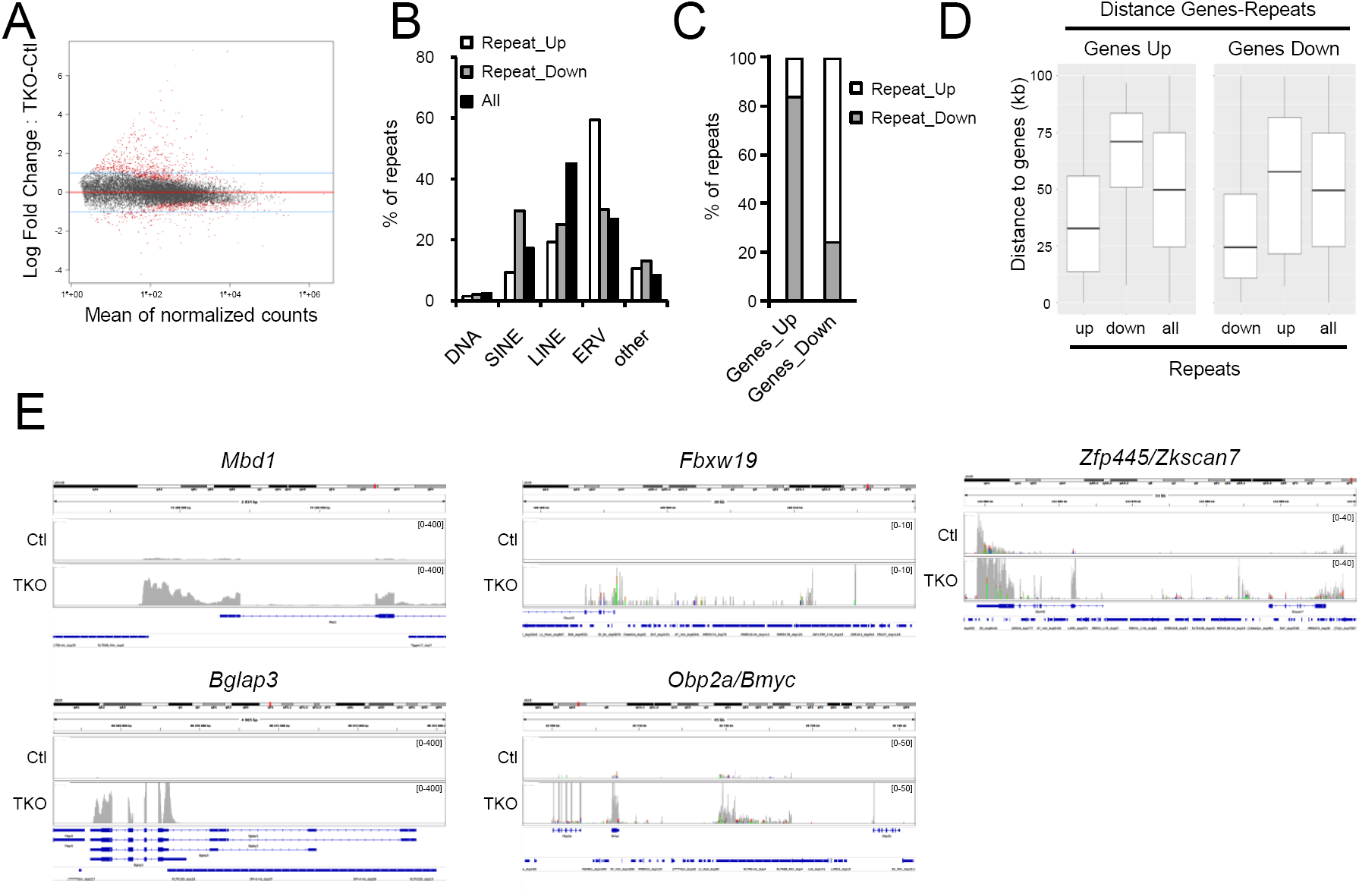
HP1 are required for silencing specific endogenous retroviruses (ERVs) in hepatocytes. (A) MA-plot after DSeq2 normalization of RNA-seq reads including repeats aligned against the Repbase database. Red dots represent genes and repeats that are differentially expressed between controls and HP1-TKO liver samples (p<0.05). (B) ERVs are over-represented in repeats that are up-regulated upon loss of HP1 (Repeat_Up) compared to repeats that are down-regulated (Repeat_Down) and to the genome-wide distribution of repeats according to the RepeatMasker database (All). (C) Repeats over-expressed in HP1-TKO liver samples compared with controls (Repeat_Up) are over-represented in regions (± 100kb) around genes that are over-expressed in HP1-TKO (genes_up). Conversely, repeats down-regulated in HP1-TKO liver samples compared with controls (Repeat_Down) are over-represented in regions (± 100kb) around genes repressed in HP1-TKO (genes_down). (D) Repeats that are up-regulated or down-regulated upon HP1 loss tend to be closer to genes that are up- or down-regulated in HP1-TKO, respectively. The absolute distance (in base pairs) was measured between the gene transcriptional start site and the beginning of the repeat, according to the RepeatMasker annotation. (E) Representative Integrative Genomic Viewer snapshots of the indicated up-regulated genes associated with up-regulated repeat sequences.

To determine whether the differential expression of these repeats was associated with deregulation of gene expression, we first generated a map of HP1-dependent repeats located in the vicinity of HP1-dependent genes. To this end, 100kb were added on both sides of each HP1-dependent gene and the HP1-dependent repeats present in these regions were scored. This analysis showed that a fraction of HP1-dependent genes (138 up-regulated and 94 down-regulated) was associated with HP1-dependent repeats. Interestingly, this physical association correlated with a functional association since 84% of repeats associated with up-regulated genes were also up-regulated and 75.5% of repeats associated with down-regulated genes were down-regulated (Fig. 5C & Supplementary Tables 8 & 9). Furthermore, up-regulated repeats tended to be located closer to up-regulated genes rather than to down-regulated genes, whereas inversely down-regulated repeats tended to be located closer to down-than up-regulated genes (Fig. 5D). Altogether, this analysis strongly suggested a link between loss of HP1, reactivation of some ERV and up-regulation of genes in their neighborhood. In agreement with this conclusion, several deregulated genes associated with deregulated repeats such as *Mbd1, Bglap3, Obpa, Bmyc, Fbxw19* and *Zfp445* have already been shown to be controlled by ERVs (Fig. 5E) ^41,42^.

### HP1 is necessary for TRIM28 activity within liver

KRAB-ZFP are known to require their interaction with the corepressor TRIM28 to sustain their repressive activity ^43^. RNA-seq, RT-qPCR and western blot assays showed that neither TRIM28 mRNA nor protein expression were significantly altered in HP1-TKO as compared to control livers (Fig. 6A-B). To investigate the relationship between HP1, KRAB-ZFPs, TRIM28 and ERVs in liver, we used the previously described mouse models in which either a mutated TRIM28 protein that cannot interact with HP1 (T28HP1box) is expressed instead of the WT TRIM28 protein or in which TRIM28 is depleted (T28KO) specifically within liver ^21,44^. As expected, western-blot analysis indicated that TRIM28 expression was strongly decreased in T28KO livers, whereas it was only marginally decreased in T28HP1box livers (this mutation is present only on one *Trim28* allele, and the other one is inactivated) (Fig. 6C). The level of the three HP1 was not affected in these mouse strains (Fig. 6C). RT-qPCR analysis showed that several HP1-dependent genes including *Nox4*; *Cypc29* and *Rsl1* were not affected in T28HP1box and T28KO livers (Fig. 6D). Conversely, *Cyp2b10; Ifit2*; *Zfp345* and *Zfp445* that were all over-expressed in HP1-TKO liver were also up-regulated in T28HP1box and T28KO livers (Fig. 6D). Altogether, these data demonstrated that HP1 proteins regulate gene expression through TRIM28-dependent and -independent mechanisms. To test whether the HP1-dependent ERV-associated genes also required TRIM28, the expression of *Mbd1* and *Bglap3* was assessed in T28KO and T28HP1box livers. Like in HP1-TKO livers, both genes were over-expressed in T28KO and T28HP1box livers, although to a lesser extent as compared to HP1-TKO livers suggesting that HP1 were able to partially repress the expression of these two genes even in the absence of TRIM28 (Fig. 6E).

**Figure 6:**
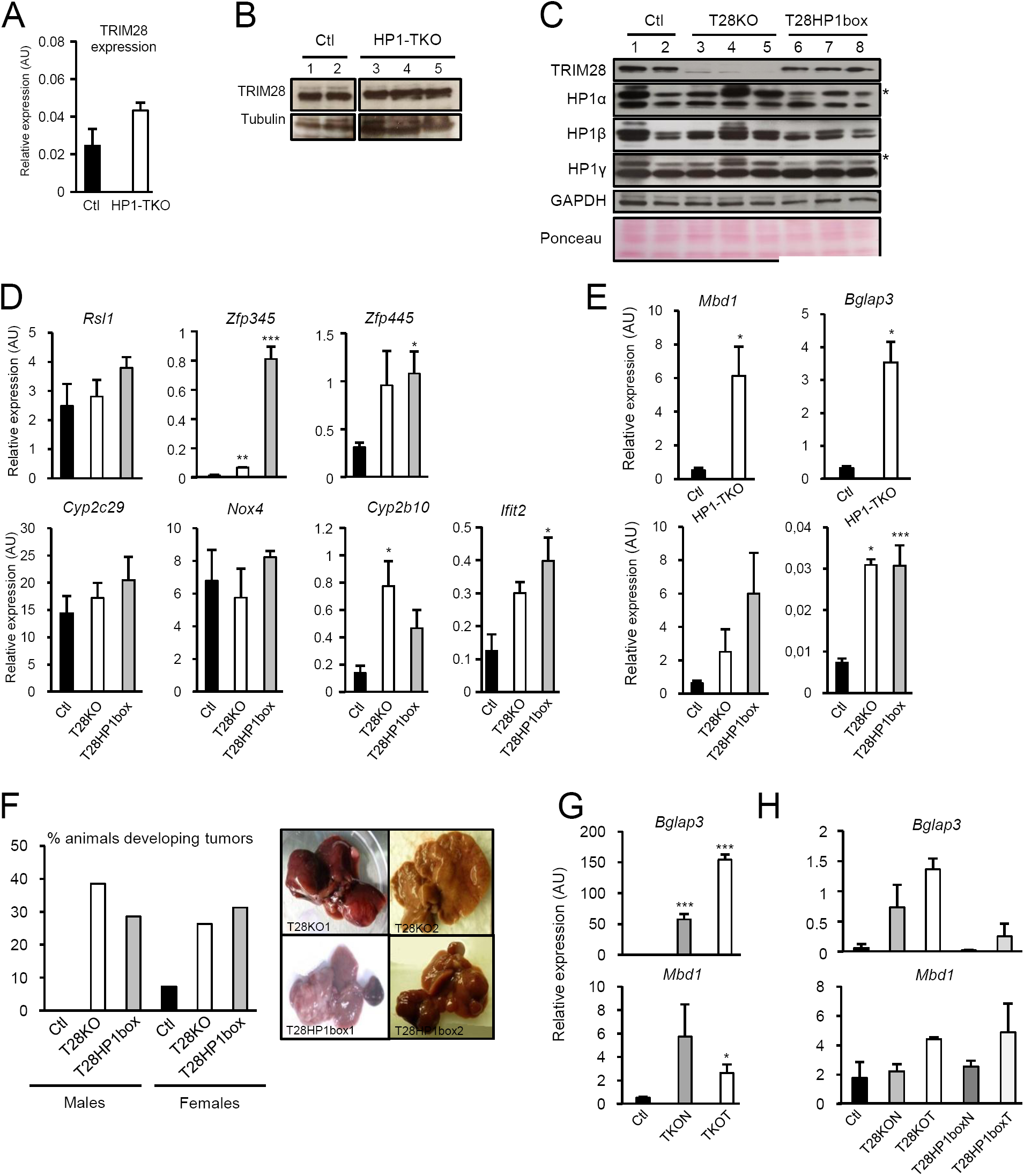
The loss of association between HP1 and TRIM28 partially recapitulates the phenotypes induced by the loss of HP1. (A) TRIM28 expression is independent of HP1 proteins. RT-qPCR quantification of TRIM28 expression in total RNA from livers of 7-week-old control (Ctl; n=4) and HP1-TKO (TKO; n=4) mice. Data were normalized to *Hprt* expression and are shown as the mean ± SEM. (B) Western blot analysis of 50µg of whole cell extracts from 7-week-old control (1 and 2) and HP1-TKO (3 to 5) livers using an anti-TRIM28 polyclonal antibody. Tubulin was used as loading control. (C) The loss of interaction between TRIM28 and HP1 does not significantly alter the level of expression of neither TRIM28 nor HP1. 50 µg of whole liver extracts from 7-week-old controls (1; 2), TRIM28KO (T28KO; 3-5) and TRIM29HP1box (T28HP1box; 6-8) mice were analyzed by western blotting using the anti-TRIM28 polyclonal and anti-HP1α, β and γ monoclonal antibodies. GAPDH and Ponceau staining were used as loading controls. (D) TRIM28 is involved in the regulation of the expression of some but not all HP1-dependent genes. RT-qPCR analysis using liver RNA samples from 5 week-old control (n=5), T28KO (n=5) and T28HP1box (n=5) mice. (E) TRIM28 is involved in the regulated expression of HP1- and ERV-dependent genes. Analysis of *Mbd1* and *Bglap3* expression by RT-qPCR using liver RNA samples from 7-week-old control (Ctl) and HP1-TKO (TKO) mice, and 5-week-old control (n=5), T28KO (n=5) and T28HP1box (n=5) mice. (F) The association between TRIM28 and HP1 is essential in hepatocytes to prevent liver tumor development. Control (n=42), T28KO (n=32) and T28HP1box (n=30) mice older than one year were sacrificed and the percentage of animals with tumors (morphological and histological analysis) was calculated. Representative morphological aspect of TRIM28 mutant livers. (G) *Bglap3* and *Mbd1* are over-expressed in HP1-TKO livers of old (>1year) mice. RT-qPCR was performed using RNA from old control (n=7), and HP1-TKO liver samples (TKON for normal part, TKOT for tumor part) (n=7). (H) The alteration of *Mbd1* and *Bglap3* expression upon loss of the association between TRIM28 and HP1 proteins was not maintained in old animals. RT-qPCR analysis using RNA from control (n=5), T28KO (T28KON for normal part, T28KOT for tumor part) (n=5) and T28HP1box (T28HP1boxN for normal part, T28HP1boxT for tumor part) livers (n=5). All expression data were normalized to *Hprt* expression and are shown as the mean ± SEM. ns, no significant difference *p value <0.05; **p value <0.01; ***p value <0.001 (Student’s t-test).

Similarly to HP1-TKO mice, T28KO and T28HP1box mice older than 42 weeks of age developed more frequently tumors in livers than controls, although with a lower penetrancy than HP1-TKO animals (38.5% and 35.7% for T28KO and T28HP1box males and 26.3% and 31.2% for T28KO and T28HP1box females, respectively, Fig. 6F) strengthening the mechanistic link between HP1 proteins and TRIM28 for liver tumor prevention.

Finally, analysis of the expression of the ERV-associated genes *Mbd1* and *Bglap3* in old animals (Fig. 6G-H) showed that both genes were over-expressed in both normal (TKON) and tumor (TKOT) liver parts from old HP1-TKO animals. In contrast, *Mbd1* was no longer over-expressed in the liver (normal and tumor parts) of old TRIM28 mutant mice. For *Bglap3*, a slight over-expression was observed in T28KO but not in T28HP1box old animals, and in both cases the level of expression was very low as compared to HP1-TKO mice (Fig. 6G-H).

Altogether, these data indicate that one important mechanism by which HP1 prevent tumor development within liver is by sustaining TRIM28 repressive activity. Moreover, we also highlight some TRIM28-independent HP1 functions in the maintenance of ERV silencing in old animals that might explained the higher incidence of tumor development in HP1-TKO animals as compared to TRIM28 mutant animals.

## DISCUSSION

In this study, we demonstrated that the hepatocyte-specific loss of HP1 proteins lead to spontaneous development of hepatocellular carcinoma (HCC). Further analysis of pre-malignant livers showed that well before tumor development livers lacking HP1 were characterized by heterochromatin disorganization, alteration of expression of many genes involved in liver specific functions as well as reactivation of specific retrotransposons. Finally, we demonstrated that this tumor suppressive function HP1 relies partially on their ability to interact with the corepressor TRIM28 to regulate its repressive activity.

The finding that in the mouse, HP1 proteins were not essential for neither cell viability nor liver function was in contrast with many studies showing the fundamental functions of each HP1 isoform in various pluripotent and differentiated cellular systems ^45,46^ as well as during embryonic development in various species, such as *Drosophila* ^12^, C. *elegans* ^13^ and the mouse (our unpublished data). One can hypothesize that liver chromatin organization and functions are highly specific and mostly independent of HP1 and/or that some compensatory mechanisms through yet unknown factors, take place specifically in mouse liver ^7,11,47^. In favor of the hypothesis of a specific liver chromatin organization, it is important to note that liver is mostly quiescent throughout life but is able to regenerate upon stress (e.g., partial hepatectomy) essentially through the re-entry of quiescent and fully differentiated hepatocytes into cell cycle rather than via stem cell proliferation, as it is the case in other tissues ^48,49^. This specific ability of differentiated hepatocytes to enter/exit quiescence could rely on a peculiar loose chromatin organization that might be less sensitive to the loss of HP1 as compared to other cell types.

We showed that HP1 loss was accompanied by a drastic reduction of the two heterochromatin marks H3K9me3 and H4K20me3 and a partial re-localization of DAPI-dense structures towards the nucleus periphery. However, in contrast to the results reported upon loss of H3K9me3 induced by inactivation of the histone methyltransferases SUV39H1 and SUV39H2, the loss of H3K9me3 in HP1-TKO hepatocytes did not result in neither decrease of H3K9me2 nor over-expression of major satellite repeats, but rather in their slight down-regulation ^50,51^. This observation supports the conclusion that HP1 are essential to maintain H3K9me3 but not H3K9me2 and that this latter histone modification is sufficient to keep major satellite sequences at a low level of transcription. It has been reported that SUV39H1 over-expression is associated with HCC development ^22^ and that HCC induced by a methyl-free diet is also characterized by elevated SUV39H1 expression and increased H3K9me3 but with reduced H4K20me3 deposition ^52^. This suggests that decreased level of H4K20me3 rather than of H3K9me3 in HP1-TKO mice could be a key determinant of tumorigenesis. In support of this hypothesis, H4K20me3 has been reported to be essential for genome integrity and for proper timing of heterochromatin replication whose deregulation has recently been proposed to be involved in cancers ^53–55^. Our indicate that

HP1 ablation also led to the deregulation (both up- and down-regulation) of many genes, strongly suggesting that HP1 are involved in both repression and activation of gene expression, as reported by others ^7,56–58^. Many of these genes are involved in liver specific functions and it will be interesting to identify the determinant for their responsiveness to HP1 depletion. Of particular interest, we found that many genes encoding for the p450 cytochrome family (Cyp) were deregulated in HP1 mutant mice. Several of these proteins are involved in the detoxification of the liver and in oxidative stress that are two key factors in hepatocarcinogenesis ^59^. How these genes are regulated by HP1 remains to be determined, however nuclear receptors of the Peroxisome Proliferation-Activated Receptors (PPAR) have been shown to be important in this process ^60^. Interestingly, our RNAseq analysis indicated that PPARγ was strongly down-regulated in HP1 mutant mice and it is tempting to speculate that this low expression of PPARγ underlies the deregulation of several *Cyp* genes. Furthermore, HP1-TKO livers were also characterized by a transcriptional signature of an interferon γ response strongly suggesting liver inflammation, a factor associated with 90% of hepatocarcinogenesis ^61^. Although the intrinsic factors involved in this inflammation remain to be discovered, our results could explained why liver is particularly prone to develop tumors in response chromatin alterations ^62^. Finally, one of the most striking result in the present study was the enrichment in genes encoding members of the KRAB-ZFP family of transcriptional co-repressors. The KRAB domain is almost exclusively present in the KRAB-Zinc Finger Protein (KRAB-ZFP) family of transcriptional repressors that have the particularity to be still actively evolving in mammals ^39^. Little information is available about the functions of most of these transcription factors, however it is now well recognized that transposable elements of the ERV family are one of their main targets through the recruitment of the TRIM28 corepressor ^39,63,64^. These mobile genetic elements constitute a threat for the genome stability and/or expression because of their ability to insert at any genomic location. Thus, an important challenge for the genome is to keep all these elements silent and unable to get transposed. However and paradoxically, increasing evidence suggests that they have been co-opted to serve as regulatory sequences in the host genome ^65^. Here, we found that, although HP1 proteins were shown to be dispensable for ERV silencing in ES cells ^66^, they are involved in silencing of specific ERVs in liver. Our data strongly suggest that the reactivation of some ERVs induce the over-expression of genes in their vicinity acting either as enhancer-like or as alternative promoters as proposed by others ^42,67^. These results could seem paradoxical with the increased expression of KRAB-ZFPs however, they are in agreement with the proposed mechanism of auto-regulation of KRAB-ZFP-encoding genes ^40^. According to this model, KRAB-ZFPs can self-inhibit their expression through interaction with the TRIM28 corepressor complex that needs to interact with HP1 for some, but not all of its functions ^40,44^. Therefore, it is very likely that in HP1-TKO livers, KRAB-ZPF-encoding genes are over-expressed because of the loss of TRIM28 activity and that, for the same reason, KRAB-ZFPs cannot repress their ERV targets. Interestingly, although our data showed that silencing of ERV in young animals relies on the activity of the functionally competent KRAB-ZFP/TRIM28/HP1 complex, TRIM28 but not HP1 becomes dispensable for this silencing in old animals highlighting complex dynamic chromatin organization and regulation throughout life.

Our study identified HP1 proteins as key players to prevent liver tumorigenesis. We highlighted major *in vivo* functions of mammalian HP1 in heterochromatin organization, regulation of gene expression and ERV silencing which most likely all contribute to the tumorigenesis process observed in livers lacking HP1.

## MATERIALS AND METHODS

### Mouse models

The Cbx5KO, T28KO (TRIM28KO) and T28HP1box (TRIM28-L2/HP1box) mouse strains were described previously ^14,44,68^. Exons 2 to 4 within the *Cbx1* gene (HP1β), and exon 3 within the *Cbx3* gene (HP1γ) were surrounded by LoxP sites. Excision of the floxed exons exclusively in hepatocytes by using mice that express the Cre recombinase under the control of the albumin promoter (Alb-Cre mice, ^26^) led to the removal of the starting ATG codon of the two genes, as well as to a frameshift within the CSD-encoding sequence of *Cbx1* and the CD-encoding sequence of *Cbx3. Cbx5,* the gene encoding HP1α, was inactivated in all body cells by removing exon 3 using the Cre recombinase under the control of the cytomegalovirus (CMV) promoter, as described previously (Cbx5KO mice) ^14^. TRIM28L2/HP1box were crossed with Alb-Cre transgenic mice to produce mice that express TRIM28HP1box as the only TRIM28 protein in hepatocytes (TRIM28-liverL-/HP1box, called T28HP1box mice in this article).

Mice were housed in a pathogen-free barrier facility, and experiments were approved by the national ethics committee for animal warfare (n°CEEA-36).

### Antibodies/oligonucleotides

The antibodies used in this study were: the rabbit anti-TRIM28 polyclonal antibody PF64, raised against amino acids 141–155 of TRIM28^64^; the anti-HP1α, anti-HP1β and anti-HP1γ monoclonal antibodies 2HP2G9, 1MOD1A9, and 2MOD1G6^65^, respectively. Anti-Casp3A (9661, Cell Signaling); anti-γH2AX (Ab11174, Abcam), anti-Ki67 (M3064, Spring Bioscience). Anti-5mC (NA81, Calbiochem). Oligonucleotides are described in Supplementary Table 10.

### Tissue processing for histology

For fresh frozen tissues, 3mm sections of the liver large lobe were embedded in the OCT compound (TissueTek) following standard protocols, and 18□ m-thick sections were cut using a Leica CM1850 cryostat and stored at −80 °C.

For paraffin-embedded tissues, 3mm sections of the liver large lobe were fixed in 4% neutral-buffered formalin (VWR Chemicals) at room temperature (RT) overnight, and stored in 70% ethanol at 4°C. Fixed tissues were processed using standard protocols and embedded in paraffin wax. Three-µm-thick sections were cut using a Thermo Scientific Microm HM325 microtome, dried at 37 °C overnight and stored at 4 °C.

### Immunofluorescence analysis

Cryo-sections were fixed in formaldehyde (2%) at RT for 15min air dried at RT for 20min and processed as described previously ^44^.

### Immunohistochemistry

Paraffin-embedded liver sections were processed for routine hematoxylin, eosin and Safran or reticulin staining. For immunohistochemistry, sections were processed according to standard protocols. Images were acquired with a Zeiss Apotome2 microscope and processed using ImageJ.

### RNA extraction and RT-qPCR assays

RNA was isolated from liver samples using TRIzol, according to the manufacturer’s recommendations (Life technologies). Reverse transcription was performed with Superscript III according to the manufacturer protocol (Invitrogen). 1/100 of this reaction was used for real-time qPCR amplification using SYBR Green I (SYBR Green SuperMix, Quanta).

### RNA-seq

The details are described in supplementary methods. Data are available at GEO (accession number: GSE119244).

### Statistics and reproducibility

The Microsoft Excel software was used for statistical analyses; statistical tests, number of independent experiments, and P-values are listed in the individual figure legends. All experiments were repeated at least twice unless otherwise stated.

## ACKNOWLEDGMENTS

We thank P. Chambon, C. Sardet, T. Forné, D.Fisher and C. Grimaud for helpful discussions and critical reading of the manuscript. We thank F. Bernex and L. LeCam, C. Keime and B. Jost for fruitful discussions. We thank L. Papon, H. Fontaine and C. Bonhomme for technical assistance and M. Oulad-Abdelghani and the IGBMC for the anti-HP1 and TRIM28 antibodies. We also thank the RHEM technical facility and particularly J. Simony for histological analysis and the IGBMC/ICS transgenic and animal facility for the initial establishment of the HP1 and TRIM28 mouse models. We thank C. Vincent and the IRCM animal core facility for the day to day care of the animal models. Finally, we thank S. Chamroeun for counting positive cells on TMA. We acknowledge the imaging facility MRI, member of the national infrastructure France-BioImaging and supported by the French National Research Agency (ANR-10-INBS-04, «Investments for the future»).

This work was supported by funds from the Centre National de la Recherche Scientifique (CNRS), the Institut National de la Santé et de la Recherche Médicale (INSERM), the University of Montpellier and the Institut regional de Cancérologie de Montpellier (ICM). SH was funded by an Erasmus PhD fellowship. We also thank the Ministry of Education, Science and Technology of the Republic of Kosovo for a scholarships to support SH. FC was supported by grants from ANR (ANR 2009 BLAN 021 91; ANR-16CE15-0018-03), INCa (PLBIO13-146), ARC (PJA20131200357), and La ligue Régionale contre le Cancer (128-R13021FF-RAB13006FFA). Sequencing was performed by the MGX facility. Montpellier, France.

## AUTHORS’ CONTRIBUTIONS

NS and SH performed the analysis of mice and interpreted the data. MP and CB made the libraries, generated and analyzed the RNA-seq data. AZ performed most of the RT-qPCR experiments. NP supervised the histological core facility and JYN performed the TMA. LK performed the pathological analysis of histological sections. EF performed some of the RT-qPCR analyses. EJ interpreted the data. FC designed, analyzed and interpreted the data and wrote the manuscript with input from all co-authors.

## SUPPLEMENTARY INFORMATIONS

**Supplementary Figure 1: HP1 are not required for liver structural organization.** The absence of HP1 proteins in hepatocytes did not induce any significant histological alteration in the liver of young (7-week-old) and middle-aged (3-6-month-old) HP1-TKO mice. (A-D) low magnification, (E-H) high magnification.

**Supplementary Figure 2: HP1-TKO tumors originate from hepatocytes lacking HP1.** (A) Excision of the *Cbx1* and *Cbx3* genes in the liver of old (x-week-old) HP1-TKO mice (TKON: normal part of liver; TKOT: tumor part of liver) compared with age-matched controls (CTL). (B) Expression of the α-fetoprotein (*Afp*) gene in the liver of old control (CTL) and HP1-TKO mice. (C) Quantification of Ki67-, caspase 3A- and γH2AX-positive cells on liver Tissue Micro Areas of old HP1-TKO mice and age-matched controls.

Supplementary table 1: analysis of RNAseq data comparing control and HP1-TKO liver total RNA

Supplementary table 2: functional clustering of genes up-regulated upon loss of HP1 witin hepatocytes (https://david.ncifcrf.gov/)

Supplementary table 3: functional clustering of genes down-regulated upon loss of HP1 witin hepatocytes (https://david.ncifcrf.gov/)

Supplementary table 4: HP1-dependent genes belonging to the IFNγ response pathway

Supplementary table 5: HP1-dependent genes with liver specific functions

Supplementary table 6: genes with liver-specific expression according to the Tissue Specific Gene Expression and Regulation software (bioinfo.wilmer.jhu.edu/tiger)

Supplementary table 7: Fold change of HP1-dependent repeats

Supplementary table 8: repeats with increased expression upon loss of HP1 associated with genes up-regulated upon loss of HP1

Supplementary table 9: repeats with decreased expression upon loss of HP1 associated with genes down-regulated upon loss of HP1

Supplementary table 10: list of the oligonucleotides used in this study

## LEGENDS FIGURES

Supplementary Figure 1: The absence of HP1 proteins in hepatocytes did not induce any significant histological alteration in the liver of young (7-week-old) and middle-aged (3-6-month-old) HP1-TKO mice. (A-D) low magnification, (E-H) high magnification.

Supplementary Figure 2: (A) Excision of the *Cbx1* and *Cbx3* genes in the liver of old (x-week-old) HP1-TKO mice (TKON: normal part of liver; TKOT: tumor part of liver) compared with age-matched controls (CTL). (B) Quantification of Ki67-, caspase 3A- and γH2AX-positive cells on liver Tissue Micro Areas of old HP1-TKO mice and age-matched controls.

