## Supplementary material for "The mouse HP1 proteins are essential for preventing liver tumorigenesis": suppl table1

**Supplementary Table 1** : analysis of RNAseq data comparing control and HP1-TKO liver total RNA

| Symbol | base Mean | log2 Fold Change | adjusted pvalue | Chr | Strand |
|---|---|---|---|---|---|
| Sdhaf4 | 614,11 | 0,0245 | 0,964556444 | Chr1 | - |
| Cfap126 | 34,29 | 0,3615 | 0,536603375 | Chr1 | + |
| 1700016C15Rik | 3,16 | 0,9581 | 0,068992694 | Chr1 | + |
| 1700025G04Rik | 71,68 | -0,0762 | 0,888648505 | Chr1 | - |
| 1700034H15Rik | 4,72 | -0,1701 | 0,847964947 | Chr1 | - |
| 1700034P13Rik | 27,18 | 0,3099 | 0,654583626 | Chr1 | + |
| 1700047M11Rik | 7,04 | -0,1241 | 0,89231057 | Chr1 | + |
| 1700056E22Rik | 2,79 | 0,3899 | 0,580044078 | Chr1 | - |
| 2310007B03Rik | 2,62 | 0,3230 | 0,645100453 | Chr1 | - |
| 2310009B15Rik | 70,07 | -0,1029 | 0,912010517 | Chr1 | - |
| 2310035C23Rik | 279,40 | 0,0393 | 0,945521759 | Chr1 | + |
| Platr5 | 2,61 | 0,7141 | 0,204417741 | Chr1 | + |
| 2610203C22Rik | 4,36 | 0,2290 | 0,773303025 | Chr1 | - |
| 2810025M15Rik | 96,74 | -0,3247 | 0,413323375 | Chr1 | + |
| 2810408I11Rik | 81,64 | -0,5223 | 0,375529622 | Chr1 | - |
| 2810459M11Rik | 956,19 | -0,2973 | 0,376557226 | Chr1 | + |
| 2900009J06Rik | 33,92 | -0,3108 | 0,592873282 | Chr1 | - |
| 3110009E18Rik | 12,85 | 0,0063 | 0,995142284 | Chr1 | + |
| 3110045C21Rik | 50,14 | 0,0911 | 0,909834629 | Chr1 | + |
| 4930523C07Rik | 249,86 | 0,3920 | 0,37274915 | Chr1 | + |
| Rab7b | 27,58 | -0,0086 | 0,992196557 | Chr1 | + |
| 5730559C18Rik | 2,47 | 0,8252 | NA | Chr1 | - |
| Teddm2 | 3,22 | 0,0900 | 0,918104715 | Chr1 | - |
| Stum | 3,24 | 0,0742 | 0,939351122 | Chr1 | - |
| 9130409I23Rik | 422,90 | 0,3788 | 0,512986719 | Chr1 | + |
| 9230116N13Rik | 4,89 | -0,1521 | 0,867206921 | Chr1 | - |
| Maip1 | 235,71 | -0,3070 | 0,29798446 | Chr1 | + |
| A130010J15Rik | 200,39 | -0,2453 | 0,534696593 | Chr1 | + |
| A330023F24Rik | 112,38 | -0,3280 | 0,465412028 | Chr1 | + |
| A530032D15Rik | 3,98 | 0,8661 | 0,147581068 | Chr1 | - |
| A630001G21Rik | 13,94 | -0,3217 | 0,639465027 | Chr1 | - |
| AA986860 | 35,38 | -0,5705 | 0,328388243 | Chr1 | + |
| Aamp | 2159,80 | -0,3503 | 0,151178014 | Chr1 | - |
| Abi2 | 41,38 | 0,3831 | 0,419539452 | Chr1 | + |
| Abl2 | 135,91 | 0,0210 | 0,969159464 | Chr1 | + |
| Acadl | 2816,43 | -0,3875 | 0,208914053 | Chr1 | - |
| Acbd3 | 257,97 | -0,0190 | 0,971430847 | Chr1 | + |
| Acbd6 | 355,04 | 0,4439 | 0,3349919 | Chr1 | + |
| Ackr1 | 44,27 | 0,2336 | 0,676060413 | Chr1 | - |
| Ackr3 | 23,32 | 0,5505 | 0,303750468 | Chr1 | + |
| Acmsd | 554,75 | 0,1113 | 0,811137679 | Chr1 | + |
| Acsl3 | 561,59 | -0,2478 | 0,503770133 | Chr1 | + |
| Actr1b | 487,28 | -0,2539 | 0,483930201 | Chr1 | - |
| Actr3 | 1586,63 | 0,1693 | 0,549296977 | Chr1 | - |
| Adam23 | 319,58 | 0,0377 | 0,938514017 | Chr1 | + |

| | | | | | |
|---|---|---|---|---|---|
| Adamts4 | 2,22 | 0,5062 | NA | Chr1 | + |
| Coq8a | 2790,45 | 0,3172 | 0,528306301 | Chr1 | - |
| Adhfe1 | 2392,87 | 0,2804 | 0,517546363 | Chr1 | + |
| Adipor1 | 1262,78 | -0,1824 | 0,510160643 | Chr1 | + |
| Adss | 887,61 | -0,4652 | 0,086491718 | Chr1 | - |
| Aff3 | 4,10 | -0,2482 | 0,762940356 | Chr1 | - |
| Agap1 | 131,53 | 0,1667 | 0,703944394 | Chr1 | + |
| Agfg1 | 270,04 | 0,3247 | 0,33637288 | Chr1 | + |
| Agxt | 4310,38 | 0,3930 | 0,22527304 | Chr1 | + |
| Ahctf1 | 804,44 | -0,2909 | 0,304623903 | Chr1 | - |
| AI597479 | 206,71 | 0,1596 | 0,696074872 | Chr1 | + |
| Ifi207 | 166,89 | 0,0388 | 0,952429505 | Chr1 | - |
| Aida | 180,40 | 0,4354 | 0,225966951 | Chr1 | + |
| Aim2 | 43,95 | 0,7972 | 0,075222739 | Chr1 | + |
| Akt3 | 58,51 | 0,5251 | 0,238655284 | Chr1 | - |
| Aldh9a1 | 3603,70 | -0,0315 | 0,935309554 | Chr1 | + |
| Als2 | 860,69 | 0,3006 | 0,382615794 | Chr1 | - |
| Als2cr12 | 3,16 | -0,1903 | 0,805867569 | Chr1 | - |
| Alyref2 | 67,91 | 0,1693 | 0,760699597 | Chr1 | + |
| Angel2 | 373,93 | 0,2624 | 0,53935078 | Chr1 | + |
| Angptl1 | 2,65 | 0,7540 | 0,183363484 | Chr1 | + |
| Ankrd23 | 14,62 | -0,1824 | 0,819337361 | Chr1 | - |
| Ankrd39 | 23,79 | -0,0512 | 0,952532092 | Chr1 | - |
| Ankrd44 | 71,17 | 0,6148 | 0,108667103 | Chr1 | - |
| Ankzf1 | 236,47 | -0,2168 | 0,576294037 | Chr1 | + |
| Aox3 | 10126,01 | -0,4687 | 0,234431867 | Chr1 | + |
| Ap1s3 | 28,52 | 0,3770 | 0,481381033 | Chr1 | - |
| Apcs | 3991,00 | -0,0445 | 0,945242233 | Chr1 | - |
| Apoa2 | 231294,22 | -0,3802 | 0,100885232 | Chr1 | + |
| Arfgef1 | 1381,92 | -0,0860 | 0,827346646 | Chr1 | - |
| Arhgap30 | 89,12 | 0,5592 | 0,136585635 | Chr1 | + |
| Arid5a | 24,89 | -0,1721 | 0,814156035 | Chr1 | + |
| Arl4c | 42,22 | 0,2048 | 0,725717458 | Chr1 | - |
| Arl8a | 170,40 | 0,0114 | 0,98489528 | Chr1 | + |
| Armc9 | 41,92 | -0,1324 | 0,826347896 | Chr1 | + |
| Arpc2 | 2422,64 | 0,0144 | 0,975445719 | Chr1 | + |
| Arpc5 | 1650,94 | -0,1424 | 0,670684766 | Chr1 | + |
| Asb1 | 130,72 | 0,1285 | 0,7950804 | Chr1 | + |
| Asnsd1 | 834,22 | -0,3359 | 0,294640151 | Chr1 | - |
| Aspm | 8,93 | -0,5141 | 0,441570242 | Chr1 | + |
| Atf3 | 40,56 | 0,4033 | 0,425493131 | Chr1 | - |
| Atf6 | 1212,78 | -0,3430 | 0,20028078 | Chr1 | - |
| Atg16l1 | 392,46 | -0,1289 | 0,737062808 | Chr1 | + |
| Atg4b | 583,44 | 0,0765 | 0,858858569 | Chr1 | + |
| Atg9a | 611,35 | -0,5177 | 0,131674744 | Chr1 | - |
| Atic | 277,47 | 0,1335 | 0,809065986 | Chr1 | + |
| Atp1a2 | 23,07 | 0,1454 | 0,86120828 | Chr1 | - |
| Atp1a4 | 3,08 | 0,2678 | 0,713614788 | Chr1 | - |
| Atp1b1 | 1104,58 | -0,3662 | 0,290848421 | Chr1 | - |
| Atp2b4 | 67,41 | 0,0588 | 0,92446794 | Chr1 | - |

| | | | | | |
|---|---|---|---|---|---|
| Atp6v1h | 1241,42 | -0,0968 | 0,7588734 | Chr1 | + |
| B4galt3 | 106,51 | -0,7099 | 0,10553196 | Chr1 | + |
| Bag2 | 28,05 | -0,5712 | 0,23475207 | Chr1 | - |
| Bard1 | 6,08 | 0,1223 | 0,895663283 | Chr1 | - |
| Batf3 | 5,64 | -0,0301 | 0,978280656 | Chr1 | + |
| BC003331 | 1131,55 | 0,0115 | 0,984345024 | Chr1 | - |
| Cryzl2 | 297,23 | -0,0206 | 0,976528475 | Chr1 | + |
| BC055324 | 9,50 | 0,0146 | 0,988512389 | Chr1 | - |
| Bcl2 | 31,71 | 0,6376 | 0,205871072 | Chr1 | - |
| Bcs1l | 201,53 | -0,1671 | 0,72659563 | Chr1 | + |
| Bend6 | 7,35 | 0,9598 | 0,102535589 | Chr1 | - |
| Bivm | 340,73 | 0,1434 | 0,716811839 | Chr1 | + |
| Blzf1 | 211,76 | 0,3173 | 0,416450614 | Chr1 | - |
| Bmpr2 | 705,24 | 0,0326 | 0,948314012 | Chr1 | + |
| Bok | 153,01 | 0,0976 | 0,871450361 | Chr1 | + |
| Bpnt1 | 948,20 | -0,0919 | 0,821196964 | Chr1 | + |
| Brox | 668,75 | 0,1412 | 0,705549096 | Chr1 | - |
| Bzw1 | 3348,49 | 0,2954 | 0,400330472 | Chr1 | + |
| C130026I21Rik | 2,86 | 0,7003 | 0,235271103 | Chr1 | - |
| C130074G19Rik | 3964,30 | 0,3033 | 0,22384344 | Chr1 | - |
| C4bp | 18902,95 | 0,0131 | 0,978402469 | Chr1 | - |
| C4bp-ps1 | 19,47 | -0,1273 | 0,881086817 | Chr1 | - |
| Cab39 | 1306,79 | -0,2467 | 0,376702038 | Chr1 | + |
| Cacybp | 764,04 | -0,0501 | 0,939520031 | Chr1 | - |
| Cadm3 | 60,10 | 0,0724 | 0,907270824 | Chr1 | - |
| Camsap2 | 287,91 | -0,0541 | 0,900441498 | Chr1 | - |
| Capn10 | 308,80 | 0,0146 | 0,978280656 | Chr1 | + |
| Capn2 | 868,68 | 0,1507 | 0,683804325 | Chr1 | - |
| Capn8 | 33,70 | -0,3283 | 0,659878933 | Chr1 | + |
| Carf | 51,06 | 0,0710 | 0,917470917 | Chr1 | + |
| Casp8 | 530,05 | -0,2526 | 0,521454979 | Chr1 | + |
| Casq1 | 3,69 | 0,5350 | 0,376282102 | Chr1 | - |
| Catip | 2,44 | -0,0816 | NA | Chr1 | + |
| Ccdc115 | 170,07 | -0,2479 | 0,582500211 | Chr1 | - |
| Ccdc93 | 207,45 | -0,0460 | 0,917022443 | Chr1 | + |
| Ccnt2 | 191,07 | 0,0264 | 0,966776406 | Chr1 | + |
| Ccnyl1 | 112,20 | -0,5132 | 0,161161606 | Chr1 | + |
| Cd247 | 4,16 | 0,3469 | 0,63960646 | Chr1 | + |
| Cd28 | 4,73 | 0,8669 | 0,151155593 | Chr1 | + |
| Cd48 | 111,81 | 0,3846 | 0,4132425 | Chr1 | + |
| Cd55 | 365,88 | 0,0884 | 0,81045863 | Chr1 | - |
| Cd84 | 163,46 | 0,4261 | 0,26777654 | Chr1 | + |
| Cdc42bpa | 300,93 | -0,0328 | 0,943717557 | Chr1 | + |
| Cdc73 | 171,06 | 0,1764 | 0,682099278 | Chr1 | - |
| Cdk18 | 337,95 | -0,5121 | 0,071771252 | Chr1 | - |
| Cenpf | 17,71 | 0,3780 | 0,585151924 | Chr1 | - |
| Cenpl | 207,46 | 0,2433 | 0,51722851 | Chr1 | + |
| Cep170 | 108,11 | 0,2419 | 0,618512306 | Chr1 | - |
| Cep350 | 399,10 | -0,1637 | 0,666442376 | Chr1 | - |
| Cfh | 23300,32 | -0,0903 | 0,824000947 | Chr1 | - |

| | | | | | |
|---|---|---|---|---|---|
| Cfhr1 | 5276,15 | 0,3219 | 0,325415084 | Chr1 | - |
| Cfhr2 | 13678,75 | -0,2284 | 0,510887696 | Chr1 | - |
| Cflar | 147,83 | 0,2188 | 0,632136487 | Chr1 | + |
| Chml | 45,23 | 0,3592 | 0,507474908 | Chr1 | - |
| Chpf | 16,26 | -0,0503 | 0,958248079 | Chr1 | - |
| Clasp1 | 131,99 | -0,1000 | 0,825518458 | Chr1 | + |
| Clk1 | 1563,08 | 0,0254 | 0,966776406 | Chr1 | - |
| Cnih4 | 333,26 | 0,1085 | 0,795812563 | Chr1 | + |
| Cnnm3 | 133,02 | 0,1319 | 0,813323745 | Chr1 | + |
| Cnnm4 | 60,22 | -0,1836 | 0,751866455 | Chr1 | + |
| Cnot11 | 233,49 | 0,0112 | 0,985628536 | Chr1 | + |
| Cnppd1 | 892,19 | -0,1881 | 0,600032204 | Chr1 | - |
| Cnst | 182,13 | 0,2592 | 0,636447281 | Chr1 | + |
| Cntn2 | 6,80 | 0,6356 | 0,330437986 | Chr1 | - |
| Col3a1 | 982,65 | 0,5243 | 0,238546081 | Chr1 | + |
| Col4a4 | 6,34 | 0,9297 | 0,118052347 | Chr1 | - |
| Col6a3 | 155,79 | 0,6775 | 0,123405092 | Chr1 | - |
| Colgalt2 | 11,41 | -0,9190 | 0,1027481 | Chr1 | + |
| Copa | 2808,35 | -0,1024 | 0,811137679 | Chr1 | + |
| Cops5 | 1854,55 | -0,2676 | 0,388953232 | Chr1 | - |
| Cops7b | 165,37 | -0,2171 | 0,628371029 | Chr1 | + |
| Cops8 | 423,40 | -0,1118 | 0,793608383 | Chr1 | + |
| Coq10b | 1736,06 | -0,2000 | 0,694472497 | Chr1 | + |
| Cox20 | 735,07 | -0,1403 | 0,716811839 | Chr1 | + |
| Cox5b | 3359,66 | -0,2255 | 0,788139622 | Chr1 | + |
| Cps1 | 71826,86 | -0,0623 | 0,910948764 | Chr1 | + |
| Creb1 | 337,81 | 0,2486 | 0,535573923 | Chr1 | + |
| Crp | 10280,09 | -0,1687 | 0,583107393 | Chr1 | + |
| Cspp1 | 225,77 | -0,2267 | 0,574841156 | Chr1 | + |
| Ctdsp1 | 2090,74 | -0,1112 | 0,801768605 | Chr1 | + |
| Ctse | 3,26 | -0,3788 | 0,582393956 | Chr1 | + |
| Cul3 | 1149,16 | -0,4611 | 0,072751135 | Chr1 | - |
| Cxcr2 | 69,56 | -0,1663 | 0,732573702 | Chr1 | + |
| Cxcr4 | 14,28 | 0,3583 | 0,602968232 | Chr1 | - |
| Cyb5r1 | 113,08 | -0,5932 | 0,140278812 | Chr1 | + |
| Cyp20a1 | 150,24 | 0,3525 | 0,356860747 | Chr1 | + |
| D030040B21Rik | 2,14 | 0,6070 | NA | Chr1 | + |
| D1Ertd622e | 327,96 | -0,0438 | 0,940390384 | Chr1 | - |
| D2hgdh | 426,31 | 0,0384 | 0,932815385 | Chr1 | + |
| D630023F18Rik | 2,47 | -0,3413 | NA | Chr1 | - |
| D830032E09Rik | 2,76 | -0,2257 | 0,737094181 | Chr1 | + |
| Dars | 2145,99 | 0,1109 | 0,811058993 | Chr1 | - |
| Dars2 | 383,83 | 0,0278 | 0,952989802 | Chr1 | - |
| Dbi | 32479,28 | -0,0128 | 0,9774226 | Chr1 | - |
| Dcaf6 | 555,55 | -0,4748 | 0,181569799 | Chr1 | - |
| Dcaf8 | 2013,03 | -0,1349 | 0,647914259 | Chr1 | + |
| Ddr2 | 115,25 | 0,2069 | 0,695967514 | Chr1 | - |
| Ddx18 | 371,10 | 0,0471 | 0,921565605 | Chr1 | - |
| Ddx59 | 63,00 | 0,0957 | 0,869236326 | Chr1 | + |
| Dedd | 346,56 | -0,2695 | 0,357345082 | Chr1 | + |

| | | | | | |
|---|---|---|---|---|---|
| Degs1 | 1284,44 | -0,1276 | 0,706001911 | Chr1 | - |
| Dennd1b | 303,09 | -0,2391 | 0,440094295 | Chr1 | + |
| Des | 53,09 | 0,1021 | 0,872521494 | Chr1 | + |
| Desi2 | 453,37 | 0,1929 | 0,678676127 | Chr1 | + |
| Dgkd | 318,32 | -0,1970 | 0,537208005 | Chr1 | + |
| Dhx9 | 1378,08 | 0,1193 | 0,754524942 | Chr1 | - |
| Diexf | 141,49 | -0,3232 | 0,352106106 | Chr1 | - |
| Dis3l2 | 213,85 | 0,1081 | 0,782080553 | Chr1 | + |
| Disp1 | 26,75 | 0,3394 | 0,564373302 | Chr1 | - |
| Dnah7b | 3,30 | 0,7558 | 0,199791783 | Chr1 | + |
| Dnajb2 | 705,73 | -0,1720 | 0,711159012 | Chr1 | + |
| Dnm3 | 2,72 | -0,1420 | 0,867671723 | Chr1 | - |
| Dnm3os | 12,00 | 0,3480 | 0,631914678 | Chr1 | + |
| Dnpep | 807,22 | -0,4142 | 0,178097604 | Chr1 | - |
| Dock10 | 68,88 | 0,4801 | 0,283971503 | Chr1 | - |
| Dpt | 217,11 | 0,1901 | 0,767917363 | Chr1 | + |
| Dsel | 22,61 | 0,6628 | 0,208914053 | Chr1 | - |
| Dst | 922,88 | 0,0812 | 0,85502909 | Chr1 | + |
| Dstyk | 188,57 | 0,0363 | 0,938661865 | Chr1 | + |
| Dtl | 11,81 | 0,4835 | 0,486005862 | Chr1 | - |
| Dtymk | 183,02 | -0,1691 | 0,736302391 | Chr1 | - |
| Dusp10 | 52,88 | -0,0860 | 0,896182211 | Chr1 | + |
| Dusp12 | 259,57 | 0,4896 | 0,224281576 | Chr1 | - |
| Dusp23 | 128,91 | -0,3366 | 0,538745986 | Chr1 | - |
| Dusp28 | 86,05 | -0,5570 | 0,191230937 | Chr1 | + |
| Dyrk3 | 16,78 | -0,2466 | 0,722152554 | Chr1 | - |
| E330020D12Rik | 29,14 | 0,1331 | 0,86436738 | Chr1 | - |
| Ecel1 | 2,70 | 0,8124 | 0,123443992 | Chr1 | - |
| Edem3 | 596,80 | -0,0216 | 0,969512928 | Chr1 | + |
| Eef1b2 | 4396,61 | -0,5045 | 0,052711367 | Chr1 | + |
| Efhc1 | 2,18 | 0,5294 | NA | Chr1 | + |
| Efhd1 | 3,92 | 0,1176 | 0,896013441 | Chr1 | + |
| Eif2d | 502,98 | -0,0882 | 0,812490734 | Chr1 | + |
| Eif4e2 | 873,20 | -0,2729 | 0,335264505 | Chr1 | + |
| Eif5b | 1650,83 | -0,3020 | 0,239404119 | Chr1 | + |
| Elf3 | 10,59 | 0,7264 | 0,242830621 | Chr1 | - |
| Elk4 | 129,05 | 0,2983 | 0,442222987 | Chr1 | + |
| Enah | 41,23 | 0,1791 | 0,764664236 | Chr1 | - |
| Epb41l5 | 965,93 | 0,1458 | 0,712389086 | Chr1 | - |
| Epha4 | 2,89 | 0,0159 | 0,986818215 | Chr1 | - |
| Eprs | 2525,82 | -0,1165 | 0,773900067 | Chr1 | + |
| Erbb4 | 6,16 | 0,4201 | 0,562610061 | Chr1 | - |
| Ercc5 | 262,09 | -0,0263 | 0,958903346 | Chr1 | + |
| Esrrg | 12,23 | -0,9134 | 0,104029003 | Chr1 | + |
| Etnk2 | 4547,91 | -0,2774 | 0,313447648 | Chr1 | + |
| Exo1 | 2,37 | 0,5522 | NA | Chr1 | + |
| F11r | 1378,46 | 0,2340 | 0,526620417 | Chr1 | + |
| F13b | 5882,59 | 0,3267 | 0,402107622 | Chr1 | + |
| F5 | 6839,03 | -0,0158 | 0,969159464 | Chr1 | + |
| Fcmr | 3,25 | 0,0214 | 0,98429278 | Chr1 | + |

| | | | | | |
|---|---|---|---|---|---|
| Fam117b | 135,82 | -0,4130 | 0,216003134 | Chr1 | + |
| Fam126b | 487,30 | -0,0402 | 0,946767463 | Chr1 | - |
| Retreg2 | 2099,25 | -0,3076 | 0,243455707 | Chr1 | + |
| Fam135a | 432,97 | -0,0217 | 0,967188554 | Chr1 | - |
| Fam168b | 643,84 | -0,1617 | 0,638888255 | Chr1 | - |
| Fam174a | 145,85 | -0,0239 | 0,969195493 | Chr1 | + |
| Fam20b | 311,53 | 0,1052 | 0,788139622 | Chr1 | - |
| Fam72a | 29,20 | -0,4146 | 0,465196538 | Chr1 | + |
| Fam78b | 2,43 | 0,1310 | NA | Chr1 | + |
| Farp2 | 212,68 | -0,2544 | 0,472260266 | Chr1 | + |
| Farsb | 880,22 | 0,0380 | 0,949779983 | Chr1 | - |
| Fasl | 2,78 | 0,1497 | 0,862616891 | Chr1 | - |
| Fastkd2 | 465,48 | -0,0182 | 0,979051488 | Chr1 | + |
| Fbxo28 | 186,78 | -0,4396 | 0,15392197 | Chr1 | - |
| Fbxo36 | 100,60 | -0,0947 | 0,886483612 | Chr1 | + |
| Fcer1g | 267,74 | 0,1838 | 0,679620039 | Chr1 | - |
| Fcgr2b | 1126,54 | 0,0993 | 0,819337361 | Chr1 | - |
| Fcgr3 | 233,19 | 0,3662 | 0,250762967 | Chr1 | - |
| Fcgr4 | 203,00 | 0,2125 | 0,658008155 | Chr1 | + |
| Fcrla | 7,74 | 0,4540 | 0,522768066 | Chr1 | - |
| Fh1 | 6643,77 | -0,1475 | 0,682575086 | Chr1 | - |
| Fhl2 | 9,52 | 0,8082 | 0,186556074 | Chr1 | - |
| Fmn2 | 10,77 | -0,8052 | 0,16832806 | Chr1 | + |
| Fmo1 | 3690,38 | 0,1289 | 0,775953072 | Chr1 | - |
| Fmo4 | 130,07 | 0,4614 | 0,365320897 | Chr1 | - |
| Fmod | 29,19 | 0,4088 | 0,505514525 | Chr1 | + |
| Fn1 | 14094,03 | -0,2891 | 0,345220446 | Chr1 | - |
| Fzd5 | 96,61 | -0,5016 | 0,214852266 | Chr1 | - |
| Fzd7 | 175,72 | -0,4080 | 0,314730989 | Chr1 | + |
| G0s2 | 2236,14 | -0,3129 | 0,690557571 | Chr1 | - |
| Gas5 | 2393,20 | -0,2985 | 0,462797865 | Chr1 | + |
| Gigyf2 | 614,92 | -0,0547 | 0,912811384 | Chr1 | + |
| Gin1 | 226,87 | -0,3771 | 0,215046281 | Chr1 | + |
| Glb1l | 44,23 | -0,0422 | 0,952424447 | Chr1 | - |
| Glrx2 | 402,58 | -0,2266 | 0,428649604 | Chr1 | + |
| Glul | 13424,07 | -0,4159 | 0,152130292 | Chr1 | + |
| Gm10516 | 29,98 | 0,1487 | 0,860078935 | Chr1 | + |
| Gm16548 | 293,53 | -0,1380 | 0,797871582 | Chr1 | - |
| Platr22 | 13,49 | 0,8777 | 0,142533012 | Chr1 | + |
| Gm17644 | 32,60 | -0,0832 | 0,913102981 | Chr1 | + |
| Gm19461 | 7,59 | 0,3931 | 0,596014309 | Chr1 | - |
| Gm19589 | 15,27 | 0,2591 | 0,724652712 | Chr1 | + |
| Gm20257 | 15,70 | -0,6060 | 0,31302658 | Chr1 | + |
| Gm2061 | 12,42 | -0,2835 | 0,699226411 | Chr1 | - |
| Gm4788 | 5996,43 | -0,3297 | 0,332014851 | Chr1 | - |
| Gm5069 | 10,45 | -0,1513 | 0,863264905 | Chr1 | - |
| Gm5523 | 8,59 | 0,2053 | 0,806181574 | Chr1 | - |
| Teddm1b | 5,16 | 0,2035 | 0,812543852 | Chr1 | + |
| Gm7609 | 74,38 | 0,2717 | 0,598404146 | Chr1 | + |
| Gm7694 | 72,72 | -0,0863 | 0,907233333 | Chr1 | - |

| | | | | | |
|---|---|---|---|---|---|
| Gm8883 | 339,89 | 0,4184 | 0,407448757 | Chr1 | + |
| Gmppa | 275,35 | -0,3424 | 0,370992482 | Chr1 | + |
| Golt1a | 430,01 | -0,3913 | 0,269881395 | Chr1 | + |
| Gorab | 75,74 | 0,2912 | 0,522469399 | Chr1 | - |
| Gpatch2 | 107,64 | 0,2584 | 0,535279147 | Chr1 | + |
| Gpc1 | 101,94 | -0,2216 | 0,709486692 | Chr1 | + |
| Gpr39 | 81,79 | 0,2817 | 0,617869186 | Chr1 | + |
| Gpr55 | 2,81 | 0,2475 | 0,747375272 | Chr1 | - |
| Gsta3 | 42293,12 | -0,4597 | 0,229159659 | Chr1 | + |
| Gtf3c3 | 202,74 | 0,6220 | 0,151387271 | Chr1 | - |
| Gulp1 | 6,53 | 0,0039 | 0,996945533 | Chr1 | + |
| H3f3a | 2516,72 | -0,0668 | 0,889068393 | Chr1 | - |
| Hdac4 | 24,89 | 0,2337 | 0,722152554 | Chr1 | - |
| Hdlbp | 5891,38 | -0,2374 | 0,457179044 | Chr1 | - |
| Hecw2 | 56,13 | 0,3577 | 0,522014539 | Chr1 | - |
| Hhat | 9,59 | 0,0516 | 0,957359271 | Chr1 | - |
| Hibch | 2123,29 | -0,1249 | 0,776119796 | Chr1 | + |
| Hlx | 20,65 | 0,2974 | 0,660982132 | Chr1 | - |
| Hmcn1 | 25,18 | 0,2015 | 0,776478455 | Chr1 | - |
| Hmga2-ps1 | 6,04 | 0,4605 | 0,522014539 | Chr1 | - |
| Hnrnpu | 2828,96 | -0,0583 | 0,896860788 | Chr1 | - |
| Hs6st1 | 1181,56 | -0,4872 | 0,10617067 | Chr1 | + |
| Hsd11b1 | 18184,30 | -0,1927 | 0,647914259 | Chr1 | - |
| Hsd17b7 | 776,31 | -0,0585 | 0,913102981 | Chr1 | - |
| Hspd1 | 11496,02 | -0,0177 | 0,969159464 | Chr1 | - |
| Hspe1 | 6592,73 | -0,0307 | 0,951032543 | Chr1 | + |
| Iars2 | 940,16 | 0,0666 | 0,862578493 | Chr1 | - |
| Icos | 22,29 | 0,5485 | 0,356860747 | Chr1 | + |
| Idh1 | 1141,76 | -0,1880 | 0,513384261 | Chr1 | - |
| Ifi203 | 274,28 | 0,1911 | 0,724187104 | Chr1 | - |
| Ifi204 | 239,05 | 0,2466 | 0,563374628 | Chr1 | - |
| Igfbp2 | 3387,72 | -0,3892 | 0,597328449 | Chr1 | + |
| Igfbp5 | 122,05 | 0,7090 | 0,19034543 | Chr1 | - |
| Igsf8 | 81,58 | -0,2597 | 0,695978671 | Chr1 | + |
| Igsf9 | 5,74 | 0,6852 | 0,285675542 | Chr1 | + |
| Ihh | 9,89 | -0,8204 | 0,165257806 | Chr1 | - |
| Ikbke | 292,99 | 0,4479 | 0,22361789 | Chr1 | - |
| Ikzf2 | 3,96 | 0,2182 | 0,773900067 | Chr1 | - |
| Il18r1 | 7,89 | -0,1948 | 0,816025345 | Chr1 | + |
| Il18rap | 7,36 | 0,2031 | 0,811137679 | Chr1 | + |
| Il1r1 | 544,83 | -0,4353 | 0,462501874 | Chr1 | + |
| Il1r2 | 3,61 | 0,6863 | 0,258561012 | Chr1 | + |
| Il1rl2 | 2,43 | -0,0011 | NA | Chr1 | + |
| Imp4 | 414,90 | -0,2217 | 0,578319732 | Chr1 | + |
| Ing5 | 187,37 | 0,4422 | 0,131674744 | Chr1 | + |
| Inha | 2,11 | 0,0737 | NA | Chr1 | + |
| Inhbb | 29,52 | 0,5435 | 0,380860886 | Chr1 | - |
| Ino80d | 268,66 | -0,0520 | 0,923778914 | Chr1 | - |
| Ino80dos | 5,49 | -0,2683 | 0,740538349 | Chr1 | + |
| Inpp1 | 93,67 | -0,1800 | 0,698557859 | Chr1 | - |

| | | | | | |
|---|---|---|---|---|---|
| Inpp4a | 205,86 | -0,6059 | 0,139781433 | Chr1 | + |
| Inpp5d | 50,57 | 0,2867 | 0,618215257 | Chr1 | + |
| Insig2 | 2757,03 | -0,0167 | 0,975528401 | Chr1 | - |
| Ints7 | 138,00 | 0,0923 | 0,86940991 | Chr1 | + |
| Ipo9 | 410,98 | -0,0498 | 0,913936879 | Chr1 | - |
| Irf6 | 1075,38 | 0,2433 | 0,430392697 | Chr1 | + |
| Irs1 | 367,27 | -0,3559 | 0,376984775 | Chr1 | - |
| Itm2c | 726,67 | 0,4547 | 0,133424603 | Chr1 | + |
| Itpkb | 76,57 | 0,5707 | 0,170659515 | Chr1 | + |
| Jph1 | 15,08 | 0,0095 | 0,992031585 | Chr1 | - |
| Kansl1l | 252,82 | 0,1972 | 0,635431815 | Chr1 | - |
| Kansl3 | 660,53 | -0,3444 | 0,239157449 | Chr1 | - |
| Kcne4 | 5,79 | 0,3057 | 0,695978671 | Chr1 | + |
| Kcnj10 | 42,69 | 0,4350 | 0,384370645 | Chr1 | + |
| Kcnk2 | 6,07 | 0,6439 | 0,323960783 | Chr1 | - |
| Kcnq5 | 2,43 | 0,1684 | NA | Chr1 | - |
| Kctd18 | 73,74 | 0,1581 | 0,743199461 | Chr1 | - |
| Kctd3 | 150,60 | -0,0827 | 0,860606321 | Chr1 | - |
| Kdelc1 | 189,63 | 0,3532 | 0,287640298 | Chr1 | - |
| Kdm5b | 297,88 | -0,0883 | 0,834675238 | Chr1 | + |
| Kdsr | 628,23 | -0,0821 | 0,875746336 | Chr1 | - |
| Kif1a | 4,32 | 0,7454 | 0,226803294 | Chr1 | - |
| Kif21b | 19,71 | 0,8326 | 0,10617067 | Chr1 | + |
| Kif26b | 4,30 | 0,4781 | 0,481381033 | Chr1 | + |
| Kifap3 | 123,11 | 0,0913 | 0,865391338 | Chr1 | + |
| Kiss1 | 2,73 | -0,3408 | 0,617869186 | Chr1 | + |
| Klf7 | 13,49 | 0,0683 | 0,943108135 | Chr1 | - |
| Klhdc8a | 14,29 | -0,0286 | 0,979212524 | Chr1 | + |
| Klhl12 | 269,25 | -0,1886 | 0,62632688 | Chr1 | + |
| Klhl20 | 127,61 | 0,4819 | 0,13773986 | Chr1 | - |
| Kmo | 46,97 | -0,0663 | 0,920434357 | Chr1 | + |
| Lactb2 | 3556,77 | -0,0205 | 0,976528475 | Chr1 | - |
| Lad1 | 16,25 | -0,7256 | 0,213959242 | Chr1 | + |
| Lamb3 | 48,07 | 0,3938 | 0,427107372 | Chr1 | + |
| Lamc1 | 173,49 | 0,0213 | 0,9695698 | Chr1 | - |
| Lamc2 | 13,34 | 0,9181 | 0,114593166 | Chr1 | - |
| Lancl1 | 458,37 | -0,2237 | 0,470066395 | Chr1 | - |
| Lax1 | 4,01 | 0,0037 | 0,997008467 | Chr1 | - |
| Lbr | 258,87 | -0,3582 | 0,182826019 | Chr1 | - |
| Lefty1 | 9,85 | -0,4298 | 0,54003692 | Chr1 | + |
| Lin9 | 61,54 | 0,2161 | 0,670891431 | Chr1 | + |
| Lipt1 | 135,17 | -0,1822 | 0,683464556 | Chr1 | + |
| Lman2l | 194,25 | -0,4272 | 0,246159206 | Chr1 | - |
| Lmbrd1 | 1684,20 | 0,0378 | 0,93731052 | Chr1 | + |
| Lmod1 | 319,11 | 0,4099 | 0,134260609 | Chr1 | + |
| Lrrfip1 | 529,27 | 0,4040 | 0,304390945 | Chr1 | + |
| Ly96 | 185,50 | 0,0003 | 0,999351454 | Chr1 | + |
| Lypla1 | 2735,74 | 0,1612 | 0,736071538 | Chr1 | + |
| Lyplal1 | 394,10 | 0,4866 | 0,060376975 | Chr1 | - |
| Map2 | 14,85 | 0,3950 | 0,563374628 | Chr1 | + |

| | | | | | |
|---|---|---|---|---|---|
| Map4k4 | 145,47 | 0,1635 | 0,677398032 | Chr1 | + |
| Mapkapk2 | 969,39 | -0,1370 | 0,678706984 | Chr1 | - |
| Marc2 | 6649,68 | -0,0661 | 0,869442758 | Chr1 | - |
| Marco | 319,11 | 0,2062 | 0,809981003 | Chr1 | - |
| Mark1 | 15,23 | -0,0937 | 0,912746031 | Chr1 | - |
| Mcm3 | 77,51 | 0,3173 | 0,528158636 | Chr1 | - |
| Mcm6 | 122,82 | 0,4834 | 0,248525122 | Chr1 | - |
| Mcmdc2 | 63,80 | 0,3060 | 0,491510849 | Chr1 | + |
| Mdm4 | 165,25 | 0,4978 | 0,186588546 | Chr1 | - |
| Mettl13 | 82,80 | -0,1472 | 0,76277049 | Chr1 | - |
| Mettl18 | 83,83 | 0,1369 | 0,788885866 | Chr1 | + |
| Mettl21a | 155,76 | 0,1336 | 0,809065986 | Chr1 | - |
| Mff | 527,56 | -0,4165 | 0,138593946 | Chr1 | + |
| Mfsd6 | 243,91 | 0,2969 | 0,413323375 | Chr1 | - |
| Flvcr1 | 78,57 | 0,2627 | 0,611887546 | Chr1 | - |
| Mfsd9 | 35,71 | 0,1208 | 0,867717945 | Chr1 | - |
| Mgat4a | 24,40 | 0,5260 | 0,364571202 | Chr1 | - |
| Mgat5 | 14,96 | 0,7357 | 0,212297691 | Chr1 | + |
| Mgst3 | 531,42 | 0,6065 | 0,104029003 | Chr1 | - |
| Mia3 | 2119,54 | -0,2236 | 0,522848151 | Chr1 | - |
| Mir3473f | 7,97 | -0,4674 | 0,507933204 | Chr1 | - |
| Mitd1 | 192,88 | 0,2319 | 0,640527814 | Chr1 | - |
| Mlph | 25,85 | 0,2579 | 0,690639808 | Chr1 | + |
| Ifi211 | 18,42 | 0,1794 | 0,820051729 | Chr1 | - |
| Mndal | 180,96 | 0,0810 | 0,896013441 | Chr1 | - |
| Mob4 | 655,69 | 0,0726 | 0,885869329 | Chr1 | + |
| Mogat1 | 7,12 | -0,2446 | 0,766619151 | Chr1 | + |
| Mpc2 | 3637,49 | -0,2925 | 0,705982644 | Chr1 | + |
| Mpz | 2,69 | -0,3834 | 0,587879802 | Chr1 | + |
| Mpzl1 | 55,55 | 0,4493 | 0,330765053 | Chr1 | - |
| Mr1 | 180,60 | 0,6902 | 0,083560273 | Chr1 | - |
| Mreg | 437,44 | -0,2187 | 0,515706574 | Chr1 | - |
| Mrpl15 | 1245,99 | -0,2195 | 0,544444415 | Chr1 | - |
| Mrpl30 | 1933,85 | -0,4307 | 0,110973039 | Chr1 | + |
| Mrpl44 | 293,10 | -0,0337 | 0,943108135 | Chr1 | + |
| Mrps14 | 1163,94 | -0,1821 | 0,580137509 | Chr1 | + |
| Mrps9 | 424,12 | -0,3686 | 0,250755642 | Chr1 | + |
| Msc | 3,25 | 0,5430 | 0,407836161 | Chr1 | - |
| Mterf4 | 455,93 | 0,1371 | 0,764664236 | Chr1 | - |
| Mybl1 | 9,50 | 0,9361 | 0,105088791 | Chr1 | - |
| Cops9 | 1163,00 | -0,2603 | 0,368588487 | Chr1 | - |
| Myo1b | 2915,55 | 0,2515 | 0,535475471 | Chr1 | - |
| Nab1 | 1196,64 | -0,3239 | 0,273018284 | Chr1 | - |
| Nabp1 | 362,94 | 0,2513 | 0,611914561 | Chr1 | - |
| Nav1 | 63,93 | 0,0833 | 0,893130925 | Chr1 | - |
| Ncf2 | 71,65 | 0,6065 | 0,162648376 | Chr1 | + |
| Nck2 | 36,03 | 0,2349 | 0,696436516 | Chr1 | + |
| Nckap5 | 3,50 | 0,5944 | 0,357290285 | Chr1 | - |
| Ncl | 4270,04 | -0,1187 | 0,753582185 | Chr1 | - |
| Ncoa2 | 432,93 | -0,0275 | 0,953911274 | Chr1 | - |

| | | | | | |
|---|---|---|---|---|---|
| Ncstn | 543,32 | -0,0033 | 0,995203097 | Chr1 | - |
| Ndufa10 | 1524,40 | -0,5545 | 0,071771252 | Chr1 | - |
| Ndufs1 | 3065,73 | -0,0663 | 0,878038315 | Chr1 | - |
| Ndufs2 | 3641,03 | -0,3037 | 0,253026101 | Chr1 | - |
| Nek2 | 52,70 | -0,0473 | 0,955803452 | Chr1 | + |
| Nek7 | 841,94 | 0,0500 | 0,925898178 | Chr1 | - |
| Nenf | 100,17 | 0,1238 | 0,806012703 | Chr1 | - |
| Neu2 | 29,12 | 0,8046 | 0,077825164 | Chr1 | + |
| Neurl3 | 124,75 | 0,2079 | 0,693242675 | Chr1 | - |
| Nfasc | 7,50 | 0,4935 | 0,47939388 | Chr1 | - |
| Ngef | 464,83 | -0,3238 | 0,300363865 | Chr1 | - |
| Nhej1 | 55,82 | 0,1640 | 0,781342762 | Chr1 | - |
| Nif3l1 | 302,69 | 0,2926 | 0,514892992 | Chr1 | + |
| Nifk | 648,77 | 0,1195 | 0,803874753 | Chr1 | + |
| Nit1 | 1282,64 | -0,0487 | 0,911736954 | Chr1 | - |
| Nme7 | 136,99 | 0,4318 | 0,283971503 | Chr1 | + |
| Nmnat2 | 5,58 | -0,5203 | 0,448865116 | Chr1 | + |
| Nop58 | 508,63 | -0,0432 | 0,932414445 | Chr1 | + |
| Nos1ap | 5,08 | 0,8237 | 0,170992427 | Chr1 | - |
| Npl | 41,26 | 0,4246 | 0,401255942 | Chr1 | - |
| Nr1i3 | 2027,59 | -0,1166 | 0,735365797 | Chr1 | + |
| Nr5a2 | 487,49 | 0,1998 | 0,672205411 | Chr1 | - |
| Nrp2 | 138,29 | -0,0319 | 0,958800482 | Chr1 | + |
| Nsl1 | 9,43 | 0,1828 | 0,830330662 | Chr1 | + |
| Nuak2 | 128,55 | 0,1111 | 0,862111303 | Chr1 | + |
| Nucks1 | 2106,38 | -0,0752 | 0,863048009 | Chr1 | + |
| Nuf2 | 19,44 | 0,6492 | 0,240555207 | Chr1 | - |
| Nvl | 232,73 | 0,1980 | 0,606429785 | Chr1 | - |
| Obsl1 | 7,24 | 0,0496 | 0,961078035 | Chr1 | - |
| Olfml2b | 5,78 | 0,7502 | 0,231528092 | Chr1 | + |
| Opn3 | 6,49 | -0,0208 | 0,985212135 | Chr1 | - |
| Optc | 2,05 | 0,3004 | NA | Chr1 | - |
| Orc2 | 113,22 | 0,2201 | 0,629243929 | Chr1 | - |
| Ormdl1 | 992,22 | 0,3515 | 0,334804026 | Chr1 | + |
| Osgepl1 | 482,77 | 0,0990 | 0,847736143 | Chr1 | + |
| Pam | 552,11 | 0,2741 | 0,522768066 | Chr1 | - |
| Paqr8 | 7,80 | 0,3518 | 0,642834828 | Chr1 | + |
| Pard3b | 59,57 | -0,3287 | 0,471563545 | Chr1 | + |
| Parp1 | 404,47 | 0,0405 | 0,927164918 | Chr1 | + |
| Pask | 8,08 | 0,3667 | 0,626412404 | Chr1 | - |
| Pbx1 | 266,32 | 0,3251 | 0,396094068 | Chr1 | - |
| Pcmtd1 | 2748,33 | -0,1626 | 0,691018693 | Chr1 | + |
| Pdcl3 | 428,60 | 0,2531 | 0,404820274 | Chr1 | + |
| Pde6d | 75,76 | -0,1302 | 0,808759895 | Chr1 | - |
| Pea15a | 432,04 | 0,4513 | 0,063989705 | Chr1 | - |
| Pecr | 4032,30 | -0,1527 | 0,596615196 | Chr1 | - |
| Per2 | 309,16 | -0,0623 | 0,909426562 | Chr1 | - |
| Pex19 | 2671,67 | 0,0119 | 0,98507181 | Chr1 | + |
| Pfdn2 | 1036,01 | 0,1372 | 0,726701117 | Chr1 | + |
| Pfkfb2 | 500,48 | -0,2024 | 0,661519766 | Chr1 | - |

| | | | | | |
|---|---|---|---|---|---|
| Pgap1 | 346,45 | -0,2918 | 0,47939388 | Chr1 | - |
| Phf3 | 743,85 | -0,2059 | 0,460037772 | Chr1 | - |
| Phlda3 | 13,02 | 0,5474 | 0,408202808 | Chr1 | + |
| Phlpp1 | 235,76 | -0,0658 | 0,899526069 | Chr1 | + |
| Pi15 | 10,87 | 0,2429 | 0,7588734 | Chr1 | + |
| Pid1 | 1132,48 | 0,2022 | 0,565966999 | Chr1 | - |
| Pigc | 260,69 | -0,0362 | 0,943305207 | Chr1 | + |
| Pigm | 195,00 | 0,1010 | 0,808156264 | Chr1 | + |
| Pign | 228,48 | 0,5127 | 0,075125268 | Chr1 | - |
| Pigr | 17269,57 | -0,4838 | 0,081415444 | Chr1 | + |
| Pik3c2b | 99,23 | -0,2682 | 0,540877577 | Chr1 | + |
| Pikfyve | 387,27 | 0,0042 | 0,993445259 | Chr1 | + |
| Pkp1 | 5,50 | 0,8500 | 0,161576375 | Chr1 | - |
| Pla2g4a | 68,31 | 0,5157 | 0,24579162 | Chr1 | - |
| Plcl1 | 12,32 | 0,1427 | 0,874833287 | Chr1 | + |
| Plekha6 | 616,27 | 0,2629 | 0,451636763 | Chr1 | + |
| Plekhb2 | 161,05 | -0,1054 | 0,873853413 | Chr1 | + |
| Plxna2 | 177,42 | -0,0677 | 0,905147397 | Chr1 | + |
| Pm20d1 | 1460,00 | -0,1837 | 0,557396844 | Chr1 | + |
| Pnkd | 823,96 | -0,1497 | 0,773303025 | Chr1 | + |
| Pogk | 113,16 | -0,5144 | 0,217693536 | Chr1 | - |
| Pou2f1 | 101,27 | 0,7263 | 0,079042198 | Chr1 | - |
| Ppfia4 | 20,03 | 0,2031 | 0,767446843 | Chr1 | - |
| Ppil3 | 327,25 | 0,0276 | 0,957837964 | Chr1 | - |
| Ppip5k2 | 947,43 | 0,3585 | 0,213198535 | Chr1 | - |
| Ppox | 245,21 | -0,2216 | 0,544991268 | Chr1 | - |
| Ppp1r12b | 17,72 | 0,0570 | 0,94952398 | Chr1 | - |
| Ppp1r15b | 1800,87 | -0,4039 | 0,118768849 | Chr1 | + |
| Ppp1r42 | 24,72 | -0,5457 | 0,357345082 | Chr1 | - |
| Ppp1r7 | 364,04 | 0,2102 | 0,587065518 | Chr1 | + |
| Ppp2r5a | 2996,17 | -0,1198 | 0,765061508 | Chr1 | - |
| Prdm14 | 3,05 | 0,7759 | 0,166993075 | Chr1 | - |
| Prdx6 | 7271,89 | -0,4324 | 0,027314689 | Chr1 | - |
| Prg4 | 855,92 | 0,1586 | 0,740663168 | Chr1 | - |
| Prex2 | 32,77 | 0,3971 | 0,472359592 | Chr1 | + |
| Prim2 | 65,57 | 0,2667 | 0,613235485 | Chr1 | - |
| Prox1 | 276,84 | 0,3163 | 0,455357491 | Chr1 | - |
| Prrc2c | 509,01 | -0,0399 | 0,929075751 | Chr1 | - |
| Prrx1 | 9,91 | -0,0900 | 0,922460349 | Chr1 | - |
| Psen2 | 2824,48 | 0,5276 | 0,092287841 | Chr1 | - |
| Psmd1 | 2137,65 | -0,2526 | 0,382008361 | Chr1 | + |
| Ptma | 5689,46 | 0,2811 | 0,471089317 | Chr1 | + |
| Ptpn14 | 34,31 | 0,1699 | 0,824468483 | Chr1 | + |
| Ptpn18 | 22,71 | -0,3967 | 0,587223505 | Chr1 | + |
| Ptpn4 | 45,88 | 0,1588 | 0,795401841 | Chr1 | - |
| Ptpn7 | 8,82 | 0,2473 | 0,757695155 | Chr1 | + |
| Nectin4 | 7,26 | -0,0573 | 0,953751362 | Chr1 | + |
| Pycr2 | 63,02 | 0,1742 | 0,738449568 | Chr1 | + |
| Ifi208 | 73,33 | 0,3513 | 0,501951423 | Chr1 | + |
| Ifi213 | 16,04 | 0,2678 | 0,714524796 | Chr1 | - |

| | | | | | |
|---|---|---|---|---|---|
| Ifi209 | 81,49 | 0,4139 | 0,388773392 | Chr1 | + |
| Qsox1 | 6615,81 | -0,0949 | 0,868886761 | Chr1 | - |
| R3hdm1 | 233,41 | -0,1594 | 0,679628788 | Chr1 | + |
| Rab17 | 356,40 | -0,0596 | 0,89879469 | Chr1 | - |
| Rab23 | 137,42 | 0,5724 | 0,230037893 | Chr1 | + |
| Rab3gap1 | 591,34 | 0,1057 | 0,830330662 | Chr1 | + |
| Rab3gap2 | 264,31 | -0,0047 | 0,993445259 | Chr1 | + |
| Rab29 | 232,23 | -0,1877 | 0,649219564 | Chr1 | + |
| Rabgap1l | 534,28 | -0,0168 | 0,977120448 | Chr1 | - |
| Rabif | 300,27 | 0,0658 | 0,869442758 | Chr1 | + |
| Ralb | 265,51 | 0,4395 | 0,202636202 | Chr1 | - |
| Ralgps2 | 301,54 | -0,2079 | 0,583782269 | Chr1 | - |
| Ramp1 | 80,37 | 0,4217 | 0,342277341 | Chr1 | + |
| Rasal2 | 256,47 | 0,5540 | 0,163729985 | Chr1 | - |
| Rassf5 | 100,12 | -0,2200 | 0,636260727 | Chr1 | - |
| Rb1cc1 | 919,77 | -0,1551 | 0,690557571 | Chr1 | + |
| Rbbp5 | 299,73 | 0,2718 | 0,478667859 | Chr1 | + |
| Rc3h1 | 736,89 | -0,1570 | 0,693242675 | Chr1 | + |
| Rcor3 | 178,63 | 0,3759 | 0,374870823 | Chr1 | - |
| Rcsd1 | 78,06 | 0,3039 | 0,464784356 | Chr1 | - |
| Rd3 | 15,38 | 0,2189 | 0,789064438 | Chr1 | + |
| Rdh10 | 342,68 | -0,0570 | 0,906240001 | Chr1 | + |
| Rev1 | 208,21 | -0,1603 | 0,729244932 | Chr1 | - |
| Rftn2 | 10,39 | 0,0599 | 0,952188756 | Chr1 | - |
| Cop1 | 1083,24 | -0,3993 | 0,186420794 | Chr1 | + |
| Rgl1 | 150,17 | 0,1997 | 0,604679897 | Chr1 | - |
| Rgs1 | 49,59 | 0,3655 | 0,546240378 | Chr1 | - |
| Rgs16 | 919,43 | 0,3637 | 0,631294599 | Chr1 | + |
| Rgs18 | 4,84 | 0,6757 | 0,293376172 | Chr1 | - |
| Rgs2 | 148,61 | 0,4174 | 0,297002219 | Chr1 | - |
| Rgs20 | 2,26 | 0,2136 | NA | Chr1 | - |
| Rgs4 | 29,94 | 0,2957 | 0,631867311 | Chr1 | - |
| Rgs7 | 3,38 | -0,4490 | 0,51770482 | Chr1 | - |
| Rhbdd1 | 407,88 | 0,2145 | 0,549054641 | Chr1 | + |
| Rnasel | 59,80 | 0,4524 | 0,345409527 | Chr1 | + |
| Rnf149 | 1144,57 | -0,0136 | 0,978402469 | Chr1 | - |
| Rnf2 | 369,37 | 0,1315 | 0,737062808 | Chr1 | - |
| Rnf25 | 229,64 | -0,1462 | 0,767446843 | Chr1 | - |
| Rnpep | 296,47 | -0,5478 | 0,144426924 | Chr1 | - |
| Rnpepl1 | 526,99 | -0,3788 | 0,322042324 | Chr1 | + |
| Rpe | 656,66 | 0,1415 | 0,696532876 | Chr1 | + |
| Rpl31 | 4598,03 | -0,2801 | 0,330791426 | Chr1 | + |
| Rpl37a | 7678,31 | -0,3650 | 0,124301562 | Chr1 | + |
| Rpl7 | 13230,65 | -0,2826 | 0,348561153 | Chr1 | - |
| Cnot9 | 377,87 | 0,1817 | 0,568858283 | Chr1 | + |
| Rrp15 | 224,00 | -0,1072 | 0,828012328 | Chr1 | - |
| Rrs1 | 202,50 | -0,1275 | 0,783422709 | Chr1 | + |
| Rufy4 | 12,46 | 0,9924 | 0,073430016 | Chr1 | + |
| Rxrg | 103,11 | 0,4788 | 0,390076693 | Chr1 | + |
| Satb2 | 4,71 | -0,5058 | 0,466141783 | Chr1 | - |

| | | | | | |
|---|---|---|---|---|---|
| Sccpdh | 94,28 | 0,5100 | 0,255658367 | Chr1 | + |
| Scly | 839,58 | -0,0164 | 0,978552759 | Chr1 | + |
| Scyl3 | 358,59 | -0,4060 | 0,173893038 | Chr1 | + |
| Sdccag8 | 102,54 | 0,3095 | 0,421362686 | Chr1 | + |
| Sde2 | 175,57 | -0,0967 | 0,82176873 | Chr1 | + |
| Sdhc | 3007,04 | -0,1260 | 0,703924654 | Chr1 | - |
| Cavin2 | 269,38 | 0,2048 | 0,613685815 | Chr1 | + |
| Sec16b | 1313,56 | 0,2538 | 0,382335568 | Chr1 | + |
| Sele | 1,97 | 0,4292 | NA | Chr1 | + |
| Sema4c | 29,33 | 0,0995 | 0,890887823 | Chr1 | - |
| Serpinc1 | 40791,14 | 0,0071 | 0,985797359 | Chr1 | + |
| Sertad4 | 9,99 | 0,9144 | 0,116820996 | Chr1 | - |
| Sf3b1 | 2541,68 | -0,1724 | 0,623297443 | Chr1 | - |
| Sft2d2 | 1131,72 | 0,1041 | 0,768686299 | Chr1 | - |
| Sgk3 | 308,51 | 0,3174 | 0,384501899 | Chr1 | + |
| Sgo2a | 9,98 | 0,4996 | 0,450228426 | Chr1 | + |
| Sh2d1b1 | 3,05 | 0,4518 | 0,50815767 | Chr1 | + |
| Sh3bp4 | 14,10 | 0,1148 | 0,896860788 | Chr1 | + |
| Shisa4 | 2,57 | 0,2218 | NA | Chr1 | - |
| Slamf6 | 9,72 | 0,4022 | 0,570866317 | Chr1 | + |
| Slamf7 | 10,77 | 0,7521 | 0,222670104 | Chr1 | - |
| Slamf8 | 18,69 | 0,4485 | 0,47056536 | Chr1 | - |
| Slamf9 | 27,11 | 0,6884 | 0,158896068 | Chr1 | + |
| Slc11a1 | 116,57 | 0,3331 | 0,402628031 | Chr1 | + |
| Slc19a2 | 1689,40 | -0,4839 | 0,105724638 | Chr1 | + |
| Slc23a3 | 3,11 | -0,3259 | 0,657129833 | Chr1 | - |
| Slc30a1 | 1124,62 | -0,2124 | 0,604679897 | Chr1 | + |
| Slc30a10 | 483,56 | 0,1231 | 0,769403008 | Chr1 | + |
| Slc35f5 | 1151,54 | 0,4090 | 0,129370441 | Chr1 | + |
| Slc39a10 | 87,65 | 0,4370 | 0,340944756 | Chr1 | - |
| Slc40a1 | 4045,10 | 0,3014 | 0,38301652 | Chr1 | - |
| Slc41a1 | 67,93 | 0,0251 | 0,972380647 | Chr1 | + |
| Slc45a3 | 396,59 | 0,5750 | 0,162648376 | Chr1 | + |
| Slc4a3 | 3,53 | 0,2498 | 0,7588734 | Chr1 | + |
| Smap1 | 367,53 | -0,3805 | 0,177916821 | Chr1 | - |
| Smarcal1 | 175,09 | 0,0743 | 0,886446111 | Chr1 | + |
| Smg7 | 380,77 | -0,3235 | 0,361447043 | Chr1 | - |
| Smyd2 | 256,27 | -0,3632 | 0,270141373 | Chr1 | - |
| Smyd3 | 45,45 | 0,2194 | 0,693151669 | Chr1 | - |
| Sned1 | 34,17 | -0,1043 | 0,883884316 | Chr1 | + |
| Snhg6 | 246,23 | -0,3662 | 0,62683519 | Chr1 | - |
| Snora41 | 2,32 | -0,2489 | NA | Chr1 | + |
| Snora75 | 2,27 | -0,5531 | NA | Chr1 | - |
| Snord47 | 5,54 | -0,2572 | 0,754953629 | Chr1 | + |
| Snord87 | 3,18 | -0,2001 | 0,796541442 | Chr1 | - |
| Snrpe | 829,17 | -0,3169 | 0,325839875 | Chr1 | - |
| Sntg1 | 13,78 | 0,4321 | 0,55440914 | Chr1 | - |
| Soat1 | 65,67 | 0,2317 | 0,632226809 | Chr1 | - |
| Sox13 | 49,32 | 0,1721 | 0,784087358 | Chr1 | - |
| Sox17 | 17,18 | 0,0895 | 0,919467514 | Chr1 | - |

| | | | | | |
|---|---|---|---|---|---|
| Sp100 | 48,56 | -0,1331 | 0,8294635 | Chr1 | + |
| Sp140 | 31,49 | 0,0689 | 0,935309554 | Chr1 | + |
| Spata17 | 4,17 | 0,8308 | 0,163729985 | Chr1 | - |
| Speg | 7,30 | 0,3603 | 0,632136487 | Chr1 | + |
| Srgap2 | 165,10 | 0,1912 | 0,700470531 | Chr1 | - |
| Srp9 | 1786,76 | -0,3797 | 0,380813891 | Chr1 | + |
| St8sia4 | 21,79 | 0,7357 | 0,177916821 | Chr1 | - |
| Stat1 | 933,59 | 0,2863 | 0,437620234 | Chr1 | + |
| Stat4 | 16,01 | 0,8594 | 0,119628533 | Chr1 | + |
| Stau2 | 55,09 | 0,2179 | 0,726324879 | Chr1 | - |
| Steap3 | 164,82 | -0,0055 | 0,991447068 | Chr1 | - |
| Stk11ip | 86,98 | 0,2834 | 0,528345204 | Chr1 | + |
| Stk16 | 789,84 | -0,5151 | 0,119939529 | Chr1 | + |
| Stk17b | 211,62 | 0,2428 | 0,611365326 | Chr1 | - |
| Stk25 | 598,00 | -0,3117 | 0,300188675 | Chr1 | - |
| Stk36 | 9,06 | -0,3850 | 0,603509332 | Chr1 | + |
| Stradb | 599,11 | -0,4099 | 0,112357003 | Chr1 | + |
| Stx6 | 88,74 | 0,3667 | 0,442074728 | Chr1 | + |
| Suco | 384,37 | 0,3462 | 0,317123683 | Chr1 | - |
| Sulf1 | 17,03 | 0,2866 | 0,692557703 | Chr1 | + |
| Sumo1 | 1322,97 | 0,0945 | 0,859745168 | Chr1 | - |
| Swt1 | 320,49 | 0,2338 | 0,583107393 | Chr1 | - |
| Tada1 | 208,48 | 0,0493 | 0,918534718 | Chr1 | + |
| Taf1a | 112,14 | 0,3171 | 0,396094068 | Chr1 | + |
| Tagln2 | 417,60 | 0,4934 | 0,092498123 | Chr1 | + |
| Tatdn3 | 109,67 | 0,1180 | 0,830662365 | Chr1 | - |
| Tbc1d8 | 226,51 | -0,4581 | 0,402148509 | Chr1 | - |
| Tcea1 | 1479,13 | -0,0099 | 0,985488456 | Chr1 | + |
| Eloc | 1233,78 | -0,2691 | 0,423089018 | Chr1 | - |
| Terf1 | 231,62 | -0,0364 | 0,947819654 | Chr1 | + |
| Tex30 | 177,15 | -0,1241 | 0,7950804 | Chr1 | - |
| Tfb2m | 646,80 | -0,1965 | 0,634623679 | Chr1 | - |
| Tfcp2l1 | 78,60 | 0,3660 | 0,445464761 | Chr1 | + |
| Tgfb2 | 11,62 | 0,2254 | 0,770543183 | Chr1 | - |
| Tgfbrap1 | 362,20 | -0,3809 | 0,237843125 | Chr1 | - |
| Thap4 | 153,82 | -0,4566 | 0,221788003 | Chr1 | - |
| Timm17a | 731,09 | -0,2985 | 0,462380045 | Chr1 | - |
| Tiprl | 389,08 | -0,0845 | 0,833247526 | Chr1 | - |
| Tlr5 | 78,54 | -0,1493 | 0,841012567 | Chr1 | + |
| Tmbim1 | 1071,41 | 0,2563 | 0,352372205 | Chr1 | - |
| Tmcc2 | 28,00 | 0,3940 | 0,474644284 | Chr1 | - |
| Tmco1 | 2218,06 | -0,1203 | 0,73377276 | Chr1 | + |
| Tmeff2 | 3,31 | 0,1402 | 0,869515614 | Chr1 | + |
| Tmem131 | 291,53 | -0,3745 | 0,31302658 | Chr1 | - |
| Tmem14a | 157,34 | 0,5875 | 0,172473519 | Chr1 | + |
| Tmem177 | 171,82 | 0,1902 | 0,696074872 | Chr1 | - |
| Tmem183a | 2029,36 | -0,4459 | 0,067820892 | Chr1 | - |
| Tmem185b | 143,77 | 0,1807 | 0,677867285 | Chr1 | + |
| Nemp2 | 10,25 | -0,0899 | 0,923778914 | Chr1 | + |
| Tmem206 | 37,67 | -0,2221 | 0,737094181 | Chr1 | + |

| | | | | | |
|---|---|---|---|---|---|
| Tmem37 | 1164,11 | -0,5459 | 0,146157562 | Chr1 | - |
| Tmem63a | 181,32 | 0,0329 | 0,955803452 | Chr1 | + |
| Tmem70 | 822,40 | -0,0914 | 0,846571896 | Chr1 | + |
| Tmem81 | 40,81 | -0,2937 | 0,56919074 | Chr1 | + |
| Tmem9 | 467,55 | 0,2722 | 0,437584445 | Chr1 | + |
| Tnfrsf11a | 22,70 | 0,2115 | 0,753233421 | Chr1 | + |
| Tnni1 | 2,25 | 0,4749 | NA | Chr1 | + |
| Tnnt2 | 8,22 | 0,5531 | 0,414312677 | Chr1 | + |
| Tnp1 | 3,89 | 0,1204 | NA | Chr1 | - |
| Tns1 | 1168,51 | 0,3901 | 0,364372969 | Chr1 | - |
| Tomm40l | 277,57 | 0,1045 | 0,820051729 | Chr1 | - |
| Tor1aip1 | 631,34 | -0,0802 | 0,835059108 | Chr1 | - |
| Tor1aip2 | 3491,79 | 0,2183 | 0,532019759 | Chr1 | + |
| Tor3a | 120,08 | 0,1889 | 0,655736343 | Chr1 | - |
| Tpp2 | 771,90 | 0,0114 | 0,98398644 | Chr1 | + |
| Tpr | 965,09 | 0,1691 | 0,67554326 | Chr1 | + |
| Traf3ip1 | 117,19 | -0,1991 | 0,660945685 | Chr1 | + |
| Traf3ip3 | 8,96 | 0,4483 | 0,5252592 | Chr1 | - |
| Traf5 | 39,08 | 0,2761 | 0,636260727 | Chr1 | - |
| Trak2 | 407,85 | 0,1679 | 0,664277995 | Chr1 | - |
| Tram1 | 3712,11 | 0,3055 | 0,332552911 | Chr1 | - |
| Tram2 | 22,48 | -0,1859 | 0,811367208 | Chr1 | - |
| Trmt1l | 290,42 | 0,0606 | 0,895594598 | Chr1 | + |
| Trove2 | 83,20 | 0,3271 | 0,398132968 | Chr1 | - |
| Trp53bp2 | 179,89 | 0,3573 | 0,319080738 | Chr1 | + |
| Tsen15 | 89,13 | 0,1544 | 0,75436056 | Chr1 | - |
| Tsga10 | 17,80 | -0,6409 | 0,286794062 | Chr1 | - |
| Tsn | 877,78 | -0,0094 | 0,98507181 | Chr1 | - |
| Tstd1 | 593,18 | -0,4082 | 0,148235412 | Chr1 | + |
| Ttll4 | 56,22 | 0,4552 | 0,305522088 | Chr1 | + |
| Tuba4a | 2028,07 | -0,4520 | 0,201490222 | Chr1 | - |
| Txndc9 | 1262,41 | -0,0292 | 0,958943411 | Chr1 | - |
| Uap1 | 703,98 | -0,0204 | 0,969195493 | Chr1 | - |
| Ube2t | 13,62 | -0,0735 | 0,93871506 | Chr1 | + |
| Ube2w | 676,82 | 0,2108 | 0,610005889 | Chr1 | - |
| Ubxn4 | 1237,37 | -0,1945 | 0,515706574 | Chr1 | + |
| Uchl5 | 420,84 | -0,4604 | 0,12296145 | Chr1 | + |
| Uck2 | 70,93 | 0,0136 | 0,985269339 | Chr1 | - |
| Ufc1 | 1480,04 | -0,3154 | 0,244000233 | Chr1 | - |
| Uggt1 | 1292,07 | -0,3384 | 0,264762084 | Chr1 | - |
| Ugt1a10 | 11,73 | 0,2156 | 0,78753026 | Chr1 | + |
| Ugt1a5 | 240,27 | -0,0867 | 0,896446457 | Chr1 | + |
| Ugt1a6a | 529,39 | 0,2260 | 0,560071658 | Chr1 | + |
| Ugt1a6b | 534,14 | 0,1119 | 0,78102295 | Chr1 | + |
| Ugt1a7c | 3,19 | -0,0117 | 0,990552671 | Chr1 | + |
| Uhmk1 | 44,73 | 0,8446 | 0,077825164 | Chr1 | - |
| Unc50 | 525,35 | -0,2091 | 0,583702066 | Chr1 | + |
| Usf1 | 283,75 | -0,0150 | 0,981067951 | Chr1 | + |
| Usp21 | 148,53 | -0,4118 | 0,3025413 | Chr1 | - |
| Usp37 | 138,00 | 0,0717 | 0,877942095 | Chr1 | - |

| | | | | | |
|---|---|---|---|---|---|
| Usp40 | 441,33 | 0,1481 | 0,721152074 | Chr1 | - |
| Utp14b | 38,06 | 0,6931 | 0,22459952 | Chr1 | + |
| Uxs1 | 164,79 | 0,1373 | 0,804418611 | Chr1 | - |
| Vamp4 | 140,87 | 0,0430 | 0,935309554 | Chr1 | + |
| Vangl2 | 3,28 | 0,0654 | 0,946767463 | Chr1 | - |
| Vcpip1 | 431,93 | 0,1945 | 0,526620417 | Chr1 | - |
| Vps4b | 1071,13 | 0,2135 | 0,505015502 | Chr1 | - |
| Wdfy1 | 169,88 | -0,5813 | 0,065887918 | Chr1 | - |
| Wdr12 | 388,19 | -0,3162 | 0,295089732 | Chr1 | - |
| Wdr26 | 1771,13 | -0,1722 | 0,547091369 | Chr1 | - |
| Wdr75 | 271,93 | 0,1780 | 0,691290598 | Chr1 | + |
| Xcl1 | 6,61 | 0,0670 | 0,947182472 | Chr1 | - |
| Xkr9 | 476,94 | -0,5418 | 0,075351003 | Chr1 | + |
| Xpr1 | 540,46 | 0,2605 | 0,336364632 | Chr1 | - |
| Xrcc5 | 594,53 | 0,2434 | 0,631914678 | Chr1 | + |
| Yod1 | 89,45 | -0,0654 | 0,935254011 | Chr1 | + |
| Zap70 | 171,17 | 0,0869 | 0,88286244 | Chr1 | + |
| Zbed6 | 91,74 | 0,5242 | 0,220997862 | Chr1 | - |
| Zbtb18 | 312,89 | -0,2900 | 0,330697335 | Chr1 | + |
| Zbtb37 | 40,92 | 0,6002 | 0,188129472 | Chr1 | - |
| Zbtb41 | 101,50 | -0,0742 | 0,925700434 | Chr1 | + |
| Zc3h11a | 1218,39 | -0,2515 | 0,376702038 | Chr1 | - |
| Zcchc2 | 303,93 | -0,3073 | 0,375215351 | Chr1 | + |
| Zfp142 | 45,33 | -0,2952 | 0,58281942 | Chr1 | - |
| Zfp281 | 568,24 | -0,1672 | 0,723824089 | Chr1 | + |
| Zfp451 | 124,26 | -0,1072 | 0,822666277 | Chr1 | - |
| Zranb3 | 71,85 | 0,3414 | 0,569088036 | Chr1 | - |
| Gm2026 | 6,94 | -0,6028 | 0,362428442 | chr2 | + |
| 1110008P14Rik | 199,51 | -0,6056 | 0,358287811 | chr2 | - |
| 1110034G24Rik | 176,16 | 0,0089 | 0,988512389 | chr2 | + |
| 1110051M20Rik | 37,32 | 0,5032 | 0,321339689 | chr2 | - |
| 1500011K16Rik | 377,00 | 0,0368 | 0,972880126 | chr2 | - |
| Zfas1 | 455,66 | -0,4153 | 0,33637288 | chr2 | + |
| 1700020I14Rik | 306,87 | 0,3679 | 0,385921204 | chr2 | + |
| Spaca9 | 2,01 | 0,0488 | NA | chr2 | - |
| 1700029I15Rik | 5,25 | -0,1337 | 0,884882423 | chr2 | + |
| 1700037H04Rik | 45,63 | 0,7283 | 0,078001372 | chr2 | - |
| 1700084E18Rik | 4,65 | -0,1527 | 0,865720762 | chr2 | - |
| 1810024B03Rik | 2,74 | 0,2938 | 0,654583626 | chr2 | - |
| 2500004C02Rik | 31,27 | 0,0461 | 0,951898903 | chr2 | - |
| Selenoh | 142,28 | -0,0475 | 0,956919137 | chr2 | - |
| 2810002D19Rik | 45,02 | 0,3097 | 0,591852522 | chr2 | + |
| Trp53rka | 80,99 | -0,1038 | 0,851472884 | chr2 | - |
| 2900097C17Rik | 2735,80 | 0,0776 | 0,842305558 | chr2 | - |
| 3300002I08Rik | 4,99 | -0,1971 | 0,819025086 | chr2 | - |
| 4921509C19Rik | 2,30 | 0,7649 | NA | chr2 | - |
| 4921531C22Rik | 14,67 | 0,4444 | 0,505514525 | chr2 | + |
| 4930402H24Rik | 225,83 | 0,2019 | 0,629823117 | chr2 | - |
| 4930405A21Rik | 2,76 | -0,2365 | 0,747447276 | chr2 | - |
| 4930473A02Rik | 2,52 | -0,4466 | NA | chr2 | - |

| | | | | | |
|---|---|---|---|---|---|
| 4932414N04Rik | 6,46 | -0,8073 | 0,192379403 | chr2 | + |
| 4932416H05Rik | 7,51 | 0,3165 | 0,683330505 | chr2 | - |
| 5031425F14Rik | 2,19 | 0,1002 | NA | chr2 | + |
| 5430405H02Rik | 27,88 | -0,4512 | 0,49530599 | chr2 | - |
| Inafm2 | 274,47 | 0,0382 | 0,944269942 | chr2 | + |
| 6430710C18Rik | 1,80 | 0,2993 | NA | chr2 | - |
| 6530402F18Rik | 13,62 | 0,5224 | 0,408538653 | chr2 | - |
| Nol4l | 18,77 | -0,2314 | 0,740269415 | chr2 | - |
| Snhg17 | 91,95 | 0,0869 | 0,895978567 | chr2 | - |
| A330069E16Rik | 89,71 | -0,4856 | 0,221366063 | chr2 | + |
| A430105I19Rik | 51,88 | -0,0476 | 0,945274006 | chr2 | - |
| A530013C23Rik | 16,24 | -0,6076 | 0,34455554 | chr2 | + |
| A730017L22Rik | 134,51 | -0,3341 | 0,355790779 | chr2 | - |
| A930004D18Rik | 12,92 | 0,7549 | 0,209104214 | chr2 | - |
| AA543186 | 7,36 | -0,3388 | 0,657129833 | chr2 | - |
| Aar2 | 291,37 | -0,0150 | 0,976528475 | chr2 | + |
| Abca2 | 374,17 | 0,0570 | 0,920510254 | chr2 | + |
| Abcb11 | 10843,65 | -0,2049 | 0,515706574 | chr2 | - |
| Abhd12 | 311,87 | -0,2517 | 0,444145513 | chr2 | - |
| Abi1 | 622,64 | 0,1287 | 0,708011891 | chr2 | - |
| Abl1 | 243,17 | 0,1489 | 0,682348071 | chr2 | + |
| Abtb2 | 113,22 | 0,4405 | 0,197774355 | chr2 | + |
| Acbd5 | 4146,58 | -0,3583 | 0,303602228 | chr2 | + |
| Accs | 55,14 | 0,3846 | 0,464991549 | chr2 | - |
| Acot8 | 153,49 | -0,7209 | 0,188249599 | chr2 | - |
| Acp2 | 1294,58 | 0,4859 | 0,171804571 | chr2 | + |
| Acss1 | 6,90 | 0,3590 | 0,634397047 | chr2 | - |
| Acss2 | 2834,34 | 0,2081 | 0,756909113 | chr2 | + |
| Actr5 | 61,99 | -0,3768 | 0,439437431 | chr2 | + |
| Acvr1 | 157,35 | 0,0929 | 0,859785376 | chr2 | - |
| Acvr2a | 409,56 | 0,0609 | 0,890761998 | chr2 | + |
| Ada | 26,35 | -0,0881 | 0,909412258 | chr2 | - |
| Adal | 218,04 | -0,1536 | 0,672205411 | chr2 | + |
| Adam33 | 8,94 | -0,3476 | 0,642834828 | chr2 | - |
| Adamts13 | 135,05 | -0,1449 | 0,745818781 | chr2 | + |
| Adamtsl2 | 67,29 | 0,6937 | 0,124301562 | chr2 | + |
| Adnp | 869,63 | 0,2929 | 0,25884542 | chr2 | - |
| Adra2b | 59,64 | 0,5704 | 0,236414731 | chr2 | + |
| Adrm1 | 707,11 | -0,6195 | 0,131561943 | chr2 | + |
| Agpat2 | 2372,75 | -0,0904 | 0,803874753 | chr2 | - |
| Agps | 281,66 | 0,3551 | 0,239250534 | chr2 | + |
| AI182371 | 11481,06 | -0,0003 | 0,999351454 | chr2 | - |
| Aif1l | 6,05 | -0,0632 | 0,950532633 | chr2 | + |
| Ak1 | 6,21 | 0,8412 | 0,169162729 | chr2 | + |
| Alkbh3 | 256,26 | -0,1236 | 0,773710065 | chr2 | - |
| Alx4 | 4,38 | -0,7001 | 0,271504179 | chr2 | + |
| Ambra1 | 275,76 | -0,0060 | 0,990846949 | chr2 | + |
| Anapc1 | 595,46 | 0,1028 | 0,804480962 | chr2 | - |
| Anapc2 | 452,02 | -0,1176 | 0,783264136 | chr2 | + |
| Angptl2 | 217,84 | 0,2276 | 0,569876243 | chr2 | + |

| | | | | | |
|---|---|---|---|---|---|
| Ankrd16 | 105,31 | -0,3932 | 0,400903279 | chr2 | + |
| Ap4e1 | 144,69 | 0,4711 | 0,208282312 | chr2 | + |
| Ap5s1 | 214,96 | -0,3066 | 0,50566917 | chr2 | + |
| Apbb1ip | 25,42 | 0,1209 | 0,877283616 | chr2 | + |
| Api5 | 1261,20 | -0,0763 | 0,838868977 | chr2 | - |
| Apip | 437,73 | -0,0227 | 0,962216378 | chr2 | + |
| Apmap | 938,81 | 0,0190 | 0,969466644 | chr2 | - |
| Aqr | 240,44 | 0,1332 | 0,793608383 | chr2 | - |
| Arfgap1 | 267,38 | 0,1260 | 0,77799861 | chr2 | + |
| Arfgap2 | 2338,80 | 0,2145 | 0,564849438 | chr2 | + |
| Arfgef2 | 608,74 | 0,0392 | 0,919612949 | chr2 | + |
| Arfrp1 | 273,60 | -0,1040 | 0,783778153 | chr2 | - |
| Arhgap1 | 183,55 | -0,1468 | 0,738449568 | chr2 | + |
| Arhgap11a | 31,30 | 0,4209 | 0,413323375 | chr2 | - |
| Arhgap15 | 29,52 | 0,6218 | 0,234422574 | chr2 | + |
| Arhgap21 | 316,11 | -0,2035 | 0,537999295 | chr2 | - |
| Arl14ep | 193,96 | 0,3240 | 0,407836161 | chr2 | - |
| Arl5a | 2944,63 | -0,3955 | 0,212149875 | chr2 | - |
| Arl5b | 182,23 | 0,5784 | 0,240254772 | chr2 | + |
| Arl6ip6 | 177,92 | -0,0698 | 0,877942095 | chr2 | + |
| Arpc5l | 634,44 | -0,4539 | 0,153555915 | chr2 | + |
| Arrdc1 | 36,10 | 0,0359 | 0,967912274 | chr2 | - |
| Asb6 | 83,04 | -0,3971 | 0,535475471 | chr2 | - |
| Asxl1 | 314,68 | 0,0518 | 0,901994263 | chr2 | + |
| Atf2 | 495,21 | -0,1450 | 0,687523396 | chr2 | - |
| Atg13 | 587,43 | -0,3399 | 0,378371266 | chr2 | - |
| Atp5c1 | 6949,17 | -0,4969 | 0,070271116 | chr2 | - |
| Atp5e | 3529,25 | -0,6033 | 0,082128644 | chr2 | - |
| Atp5g3 | 7957,39 | -0,4702 | 0,225613833 | chr2 | - |
| Atp8b4 | 44,99 | 0,5621 | 0,301884234 | chr2 | - |
| Atp9a | 635,84 | -0,1358 | 0,730432237 | chr2 | - |
| Atrn | 886,95 | -0,3090 | 0,306882295 | chr2 | + |
| Aurka | 11,44 | 0,1305 | 0,88569455 | chr2 | - |
| Aven | 204,79 | -0,5705 | 0,053014128 | chr2 | + |
| B230118H07Rik | 561,15 | 0,0107 | 0,986818215 | chr2 | - |
| B2m | 62344,51 | 0,3125 | 0,27013327 | chr2 | + |
| B4galt5 | 601,96 | 0,0973 | 0,868151289 | chr2 | - |
| Bahd1 | 213,17 | 0,3380 | 0,474794806 | chr2 | + |
| Bambi-ps1 | 7,68 | 0,0216 | 0,984995389 | chr2 | - |
| Baz2b | 140,46 | 0,2698 | 0,565966999 | chr2 | - |
| Bbox1 | 4373,49 | 0,0024 | 0,99658581 | chr2 | - |
| Bbs5 | 70,56 | -0,0026 | 0,997464504 | chr2 | + |
| BC005624 | 458,42 | -0,0268 | 0,953751362 | chr2 | - |
| Paxx | 143,51 | -0,4841 | 0,324925401 | chr2 | - |
| BC029722 | 253,69 | -0,0290 | 0,955518203 | chr2 | - |
| BC039771 | 8,65 | 0,1359 | 0,878605672 | chr2 | + |
| BC052040 | 56,37 | 0,5280 | 0,278905276 | chr2 | + |
| Bcl2l1 | 350,78 | -0,1778 | 0,744313101 | chr2 | - |
| Bcl2l11 | 102,45 | -0,0677 | 0,918077233 | chr2 | + |
| Bend7 | 143,85 | -0,4158 | 0,224651108 | chr2 | + |

| | | | | | |
|---|---|---|---|---|---|
| Blcap | 277,74 | -0,2836 | 0,472359592 | chr2 | - |
| Bloc1s6 | 280,43 | 0,0450 | 0,913936879 | chr2 | + |
| Blvra | 453,65 | 0,1715 | 0,665028267 | chr2 | + |
| Bmf | 342,51 | -0,1079 | 0,876828824 | chr2 | - |
| Bmi1 | 461,29 | 0,5667 | 0,05818386 | chr2 | + |
| Bmp2 | 231,41 | -0,1662 | 0,713614788 | chr2 | + |
| Brd3 | 380,12 | -0,1488 | 0,661748407 | chr2 | - |
| Btbd3 | 80,52 | -0,0150 | 0,98429278 | chr2 | + |
| Bub1 | 5,72 | 0,2947 | 0,711559683 | chr2 | - |
| Bub1b | 22,71 | 0,2334 | 0,744486882 | chr2 | + |
| C1qtnf4 | 1,90 | 0,1310 | NA | chr2 | + |
| Tmem250-ps | 820,84 | -0,4776 | 0,069527621 | chr2 | - |
| Cables2 | 317,80 | -0,4263 | 0,226302394 | chr2 | - |
| Cacfd1 | 356,52 | -0,3763 | 0,212297691 | chr2 | + |
| Cacnb2 | 3,14 | -0,0098 | 0,992452544 | chr2 | + |
| Cacnb4 | 2,93 | 0,0657 | 0,944253955 | chr2 | - |
| Camk1d | 815,96 | 0,0906 | 0,885528714 | chr2 | - |
| Camsap1 | 121,05 | -0,3515 | 0,348784567 | chr2 | - |
| Capn3 | 33,60 | -0,6604 | 0,197567244 | chr2 | + |
| Caprin1 | 2908,64 | 0,1965 | 0,505514525 | chr2 | - |
| Card9 | 7,27 | -0,2227 | 0,789524079 | chr2 | - |
| Casc4 | 181,63 | 0,1449 | 0,718777654 | chr2 | + |
| Knl1 | 3,02 | 0,0554 | 0,953751362 | chr2 | + |
| Cat | 38891,36 | -0,3568 | 0,073845576 | chr2 | - |
| Catsper2 | 28,94 | 0,0616 | 0,943305207 | chr2 | - |
| Cbfa2t2 | 197,73 | -0,1776 | 0,659265465 | chr2 | + |
| Kyat1 | 1954,65 | -0,2027 | 0,582615116 | chr2 | - |
| Ccdc148 | 43,94 | 0,1190 | 0,862616891 | chr2 | - |
| Ccdc173 | 8,40 | 0,2640 | 0,746827664 | chr2 | - |
| Ccdc3 | 26,73 | 0,0172 | 0,985269339 | chr2 | + |
| Ccdc32 | 123,26 | 0,0878 | 0,869687361 | chr2 | - |
| Ccdc34 | 33,87 | 0,6662 | 0,140539066 | chr2 | + |
| Ccm2l | 65,20 | 0,3675 | 0,513917577 | chr2 | + |
| Ccndbp1 | 276,99 | -0,3092 | 0,320305617 | chr2 | + |
| Cd302 | 7013,82 | -0,4550 | 0,085476448 | chr2 | - |
| Cd40 | 12,67 | -0,2575 | 0,735042834 | chr2 | + |
| Cd44 | 181,71 | 0,3785 | 0,311028921 | chr2 | - |
| Cd59a | 2010,68 | 0,2475 | 0,449196578 | chr2 | + |
| Cd59b | 133,08 | 0,4524 | 0,306882295 | chr2 | + |
| Cd82 | 1326,46 | -0,1258 | 0,692167613 | chr2 | - |
| Cdan1 | 83,65 | -0,1765 | 0,721569914 | chr2 | - |
| Cdc123 | 1089,62 | -0,0267 | 0,955234619 | chr2 | - |
| Cdc25b | 13,00 | 0,5056 | 0,437728282 | chr2 | + |
| Cdca7 | 23,69 | -0,7773 | 0,111086521 | chr2 | + |
| Cdh4 | 5,19 | -0,3357 | 0,658209405 | chr2 | + |
| Cdk5rap1 | 201,12 | 0,3400 | 0,507933204 | chr2 | - |
| Cdk9 | 363,07 | -0,4169 | 0,274469095 | chr2 | - |
| Cdnf | 8,61 | -0,0630 | 0,948356 | chr2 | + |
| Cds2 | 1512,65 | 0,0299 | 0,936825775 | chr2 | + |
| Celf1 | 1784,59 | 0,0060 | 0,988766352 | chr2 | + |

| | | | | | |
|---|---|---|---|---|---|
| Cenpb | 1067,80 | 0,3511 | 0,217835326 | chr2 | - |
| Cep152 | 35,15 | -0,4639 | 0,38141485 | chr2 | - |
| Cep250 | 81,11 | -0,5033 | 0,211374848 | chr2 | + |
| Cercam | 5,22 | 0,1287 | 0,888763259 | chr2 | + |
| Cerkl | 27,32 | 0,8797 | 0,054419598 | chr2 | - |
| Cers6 | 96,42 | 0,6571 | 0,077085268 | chr2 | + |
| Chac1 | 52,90 | -0,8117 | 0,07965206 | chr2 | + |
| Chchd5 | 302,31 | -0,2434 | 0,409970248 | chr2 | + |
| Chd6 | 192,35 | -0,1880 | 0,587879802 | chr2 | - |
| Chmp4b | 888,05 | -0,1375 | 0,748732136 | chr2 | + |
| Chn1os3 | 8,35 | -0,0851 | 0,931322892 | chr2 | + |
| Chp1 | 6376,21 | -0,3410 | 0,355790779 | chr2 | + |
| Chrna4 | 101,98 | 0,8176 | NA | chr2 | - |
| Chst14 | 26,07 | -0,1602 | 0,816025345 | chr2 | + |
| Ciao1 | 813,31 | -0,1154 | 0,755670612 | chr2 | - |
| Cir1 | 617,95 | 0,0861 | 0,83293655 | chr2 | - |
| Ciz1 | 156,61 | -0,2178 | 0,668382348 | chr2 | + |
| Ckap2l | 10,43 | 0,6534 | 0,301862555 | chr2 | - |
| Ckap5 | 195,40 | 0,4424 | 0,321845648 | chr2 | + |
| Ckmt1 | 2,39 | 0,1065 | NA | chr2 | + |
| Clp1 | 469,66 | 0,1291 | 0,813323745 | chr2 | - |
| Cnbd2 | 5,18 | 0,4302 | 0,557547051 | chr2 | + |
| Cntrl | 255,96 | 0,3347 | 0,505514525 | chr2 | + |
| Col20a1 | 14,22 | -0,4014 | 0,563271302 | chr2 | + |
| Col5a1 | 75,60 | 0,4927 | 0,258527696 | chr2 | + |
| Commd3 | 1515,42 | -0,1735 | 0,695978671 | chr2 | + |
| Commd7 | 256,50 | 0,0043 | 0,992874254 | chr2 | - |
| Commd9 | 328,08 | -0,4334 | 0,213994613 | chr2 | + |
| Cops2 | 1762,16 | -0,1496 | 0,739098859 | chr2 | - |
| Coq4 | 108,70 | -0,1261 | 0,818594346 | chr2 | + |
| Cox4i2 | 1,94 | -0,0676 | NA | chr2 | + |
| Cpne1 | 298,86 | -0,0685 | 0,869442758 | chr2 | - |
| Cpxm1 | 18,11 | 0,6697 | 0,264762084 | chr2 | - |
| Crat | 474,71 | -0,6376 | 0,234220487 | chr2 | - |
| Creb3l1 | 5,20 | 0,2281 | 0,786009002 | chr2 | - |
| Crls1 | 837,23 | -0,2603 | 0,517431746 | chr2 | + |
| Crnkl1 | 295,14 | -0,0445 | 0,92522408 | chr2 | - |
| Cry2 | 558,12 | -0,1316 | 0,768926477 | chr2 | - |
| Cse1l | 859,67 | 0,0829 | 0,863048009 | chr2 | + |
| Csnk2a1 | 2854,26 | 0,0639 | 0,892204133 | chr2 | + |
| Kat14 | 215,89 | -0,1950 | 0,666301331 | chr2 | + |
| Cst3 | 2667,67 | -0,2168 | 0,730063556 | chr2 | - |
| Cstf1 | 176,91 | 0,1534 | 0,735042834 | chr2 | + |
| Cstf3 | 260,75 | 0,0506 | 0,905517642 | chr2 | + |
| Ctcfl | 26,80 | -0,1137 | 0,876857496 | chr2 | - |
| Ctdspl2 | 241,37 | 0,3011 | 0,448865116 | chr2 | + |
| Ctnnbl1 | 252,42 | 0,2868 | 0,52825088 | chr2 | + |
| Ctnnd1 | 1306,94 | 0,3338 | 0,25568151 | chr2 | - |
| Ctsa | 1604,50 | -0,4845 | 0,074602744 | chr2 | + |
| Ctsz | 2892,42 | -0,4008 | 0,273617478 | chr2 | - |

| | | | | | |
|---|---|---|---|---|---|
| Cutal | 848,46 | 0,1490 | 0,738753015 | chr2 | + |
| Cwc22 | 217,79 | 0,0571 | 0,901601565 | chr2 | - |
| Cybrd1 | 3,19 | -0,3686 | 0,606775648 | chr2 | + |
| Cytip | 30,99 | 0,7273 | 0,160747995 | chr2 | - |
| Spout1 | 246,68 | 0,3778 | 0,339330658 | chr2 | - |
| D330023K18Rik | 41,05 | -0,3806 | 0,511962155 | chr2 | - |
| D630003M21Rik | 9,79 | 0,4218 | 0,557210708 | chr2 | - |
| Dbhos | 3,76 | 0,3756 | 0,605538599 | chr2 | - |
| Dcaf17 | 230,66 | -0,0995 | 0,830848779 | chr2 | + |
| Dclre1c | 168,68 | -0,4234 | 0,245479706 | chr2 | + |
| Ddb2 | 78,34 | -0,0933 | 0,884969456 | chr2 | - |
| Ddrgk1 | 1353,00 | -0,4047 | 0,357779158 | chr2 | - |
| Ddx27 | 243,33 | -0,2618 | 0,477004036 | chr2 | + |
| Ddx31 | 76,35 | -0,0225 | 0,976127317 | chr2 | + |
| Defb36 | 3,65 | 0,9851 | 0,06036212 | chr2 | + |
| Dennd1a | 155,92 | -0,2362 | 0,6077506 | chr2 | - |
| Depdc7 | 726,10 | -0,0109 | 0,985849808 | chr2 | - |
| Dgkz | 342,15 | -0,1130 | 0,805374431 | chr2 | - |
| Dhtkd1 | 224,80 | 0,1972 | 0,691018693 | chr2 | - |
| Dhx35 | 51,73 | 0,5321 | 0,28765305 | chr2 | + |
| Dido1 | 380,11 | -0,2986 | 0,342896939 | chr2 | - |
| Disp2 | 36,13 | -0,0617 | 0,940263679 | chr2 | + |
| Dlgap4 | 125,90 | -0,4998 | 0,217231457 | chr2 | + |
| Dll4 | 46,35 | -0,1649 | 0,804418611 | chr2 | + |
| Dnajc1 | 505,00 | -0,3083 | 0,270128032 | chr2 | - |
| Dnajc17 | 34,67 | 0,0103 | 0,990227395 | chr2 | - |
| Dnajc24 | 92,79 | 0,2990 | 0,611365326 | chr2 | - |
| Dnajc5 | 727,87 | -0,1183 | 0,738282454 | chr2 | + |
| Dnlz | 207,29 | -0,2734 | 0,482891215 | chr2 | - |
| Dnmt3b | 14,37 | 0,3967 | 0,575162128 | chr2 | + |
| Dnttip1 | 122,94 | -0,3573 | 0,26767579 | chr2 | + |
| Dok5 | 4,16 | 0,4887 | 0,477607804 | chr2 | + |
| Dolk | 248,58 | -0,2001 | 0,604679897 | chr2 | - |
| Dolpp1 | 146,71 | -0,2778 | 0,542884329 | chr2 | + |
| Dph6 | 78,60 | 0,3140 | 0,527609356 | chr2 | - |
| Dph7 | 68,49 | 0,4310 | 0,39647951 | chr2 | + |
| Dpm1 | 1165,30 | -0,4643 | 0,072751135 | chr2 | - |
| Dpm2 | 326,90 | -0,0286 | 0,951898903 | chr2 | + |
| Dpp4 | 3485,06 | 0,5152 | 0,102109628 | chr2 | - |
| Dpp7 | 641,70 | 0,1517 | 0,678130673 | chr2 | - |
| Dsn1 | 16,05 | 0,6381 | 0,298116176 | chr2 | - |
| Dstn | 4407,67 | -0,1640 | 0,585630763 | chr2 | + |
| Dtd1 | 163,42 | 0,2383 | 0,510705809 | chr2 | + |
| Dtwd1 | 139,99 | 0,1992 | 0,692167613 | chr2 | + |
| Dusp19 | 143,93 | 0,3358 | 0,400590554 | chr2 | + |
| Dusp2 | 4,67 | 0,0631 | 0,949779983 | chr2 | + |
| Dut | 101,28 | -0,5073 | 0,147364898 | chr2 | + |
| Dync1i2 | 844,68 | 0,1931 | 0,564849438 | chr2 | + |
| Dynlrb1 | 2431,51 | 0,0185 | 0,967498977 | chr2 | + |
| Dzank1 | 8,86 | -0,0603 | 0,951806422 | chr2 | - |

| | | | | | |
|---|---|---|---|---|---|
| E030013I19Rik | 3,56 | 0,5670 | 0,374529231 | chr2 | + |
| E130018N17Rik | 3,97 | 0,5590 | 0,394187455 | chr2 | - |
| E2f1 | 58,78 | 0,2786 | 0,585412613 | chr2 | - |
| Echdc3 | 1069,96 | 0,1887 | 0,649565939 | chr2 | - |
| Edem2 | 697,03 | -0,4737 | 0,215046281 | chr2 | - |
| Edf1 | 1673,00 | -0,7075 | 0,156728357 | chr2 | + |
| Egfl7 | 279,63 | -0,0014 | 0,997605264 | chr2 | + |
| Ehd4 | 122,04 | 0,6558 | 0,058073159 | chr2 | - |
| Ehf | 27,04 | 0,4142 | 0,49077279 | chr2 | - |
| Ehmt1 | 251,59 | 0,0157 | 0,978280656 | chr2 | - |
| Eif2ak4 | 122,59 | -0,1125 | 0,830330662 | chr2 | + |
| Eif2s2 | 2759,51 | -0,2379 | 0,501255989 | chr2 | - |
| Eif3m | 2162,67 | -0,1440 | 0,730063556 | chr2 | - |
| Eif6 | 851,23 | -0,1063 | 0,844029674 | chr2 | - |
| Ell3 | 47,15 | 0,7435 | 0,172087384 | chr2 | - |
| Elmo2 | 106,75 | -0,3036 | 0,526946981 | chr2 | - |
| Elp4 | 174,35 | 0,5517 | 0,061576732 | chr2 | - |
| Emc4 | 959,28 | -0,3135 | 0,434726768 | chr2 | - |
| Emc7 | 1403,21 | -0,2590 | 0,386872469 | chr2 | + |
| Endog | 242,81 | 0,1026 | 0,841012567 | chr2 | + |
| Eng | 480,10 | -0,1637 | 0,702583616 | chr2 | + |
| Enkur | 2,89 | 0,4228 | 0,527820373 | chr2 | - |
| Entpd2 | 50,74 | 0,1802 | 0,775945747 | chr2 | + |
| Entpd6 | 80,67 | 0,1416 | 0,797616774 | chr2 | + |
| Entpd8 | 1466,26 | 0,0003 | 0,999080228 | chr2 | + |
| Epb41l1 | 20,38 | 0,1509 | 0,836593297 | chr2 | + |
| Epc2 | 187,75 | -0,0354 | 0,943717557 | chr2 | + |
| Ergic3 | 1125,94 | -0,4328 | 0,098348401 | chr2 | + |
| Esf1 | 366,92 | 0,2981 | 0,439089539 | chr2 | - |
| Etl4 | 67,23 | 0,5851 | 0,111432961 | chr2 | + |
| Zfp971 | 242,57 | 0,3351 | 0,53490582 | chr2 | + |
| Exd1 | 41,23 | 0,7708 | 0,078316133 | chr2 | - |
| Exosc2 | 292,38 | -0,0177 | 0,978280656 | chr2 | + |
| Ext2 | 362,87 | -0,1019 | 0,78753026 | chr2 | - |
| F2 | 29920,17 | -0,1371 | 0,66780976 | chr2 | - |
| Fahd2a | 327,08 | -0,1623 | 0,718037643 | chr2 | - |
| Fam102a | 392,90 | -0,5231 | 0,217231457 | chr2 | + |
| Fam107b | 1134,52 | -0,2526 | 0,427422385 | chr2 | + |
| Fam110a | 23,98 | 0,0784 | 0,922379013 | chr2 | - |
| Fam171a1 | 83,03 | -0,1856 | 0,716169774 | chr2 | + |
| Fam171b | 9,25 | 0,2688 | 0,741224458 | chr2 | + |
| Fam209 | 5,27 | -0,7008 | 0,268883626 | chr2 | + |
| Fam210b | 1440,42 | -0,1401 | 0,669162583 | chr2 | + |
| Ripor3 | 11,59 | -0,2607 | 0,744618461 | chr2 | - |
| Fam69b | 3,20 | 0,2391 | 0,759631295 | chr2 | + |
| Miga2 | 589,98 | -0,0341 | 0,958903346 | chr2 | + |
| Fam78a | 15,00 | 0,2003 | 0,802706588 | chr2 | - |
| Fam83d | 4,16 | -0,2921 | 0,709458057 | chr2 | + |
| Fam98b | 264,83 | 0,6831 | 0,085593171 | chr2 | + |
| Fap | 4,21 | 0,3208 | 0,666980604 | chr2 | - |

| | | | | | |
|---|---|---|---|---|---|
| Fastkd1 | 295,03 | -0,4680 | 0,203863873 | chr2 | - |
| Fastkd5 | 223,86 | 0,0606 | 0,91071941 | chr2 | - |
| Fbln7 | 7,18 | 0,7482 | 0,232350203 | chr2 | + |
| Fbn1 | 68,51 | 0,6099 | 0,140465486 | chr2 | - |
| Fbxo18 | 502,42 | 0,0165 | 0,973458724 | chr2 | - |
| Fbxo3 | 2222,00 | 0,0941 | 0,831128367 | chr2 | + |
| Fbxw2 | 717,40 | -0,0452 | 0,918077233 | chr2 | - |
| Fbxw5 | 277,35 | -0,3052 | 0,442624295 | chr2 | + |
| Fcna | 1367,98 | 0,2677 | 0,413537476 | chr2 | - |
| Fibin | 7,36 | 0,1683 | 0,849664617 | chr2 | - |
| Fign | 107,11 | 0,7375 | 0,060051268 | chr2 | - |
| Fitm2 | 215,81 | 0,0266 | 0,969892522 | chr2 | - |
| Fkbp1a | 815,25 | 0,1250 | 0,696074872 | chr2 | + |
| Fkbp7 | 95,76 | 0,0739 | 0,897703859 | chr2 | - |
| Flrt3 | 3,43 | 0,4069 | 0,564849438 | chr2 | - |
| Fmnl2 | 128,29 | 0,0090 | 0,988512389 | chr2 | + |
| Fnbp1 | 147,22 | 0,1538 | 0,769825964 | chr2 | - |
| Fnbp4 | 230,99 | -0,6471 | 0,066396922 | chr2 | + |
| Foxa2 | 186,59 | 0,0721 | 0,925898178 | chr2 | - |
| Foxs1 | 1,88 | 0,2347 | NA | chr2 | - |
| Fpgs | 1605,68 | 0,2268 | 0,581489212 | chr2 | - |
| Frmd4a | 99,80 | 0,2050 | 0,657129833 | chr2 | + |
| Frzb | 34,81 | 0,6071 | 0,222670104 | chr2 | - |
| Fubp3 | 262,56 | 0,0328 | 0,950625589 | chr2 | + |
| Fut7 | 2,02 | 0,5962 | NA | chr2 | + |
| Gabpb1 | 106,01 | 0,0322 | 0,95728901 | chr2 | - |
| Gad2 | 3,40 | -0,3972 | 0,571917529 | chr2 | + |
| Galk2 | 420,50 | 0,1483 | 0,702177047 | chr2 | + |
| Galnt3 | 8,15 | 0,4039 | 0,583782269 | chr2 | - |
| Ganc | 229,44 | 0,1918 | 0,680997554 | chr2 | + |
| Gapvd1 | 364,12 | 0,1852 | 0,604579545 | chr2 | - |
| Garnl3 | 2,92 | 0,1795 | 0,830330662 | chr2 | - |
| Gata5 | 5,11 | -0,2981 | 0,707094339 | chr2 | - |
| Gatm | 192,99 | 0,1298 | 0,756319626 | chr2 | - |
| Gbgt1 | 2,53 | 0,1810 | NA | chr2 | + |
| Gca | 95,87 | 0,2839 | 0,526946981 | chr2 | + |
| Gchfr | 3018,77 | 0,3274 | 0,253832835 | chr2 | + |
| Ggt7 | 3,46 | -0,2000 | 0,803874753 | chr2 | - |
| Ggta1 | 94,14 | 0,5657 | 0,163729985 | chr2 | - |
| Gid8 | 1027,72 | -0,0596 | 0,8859355 | chr2 | + |
| Gins1 | 70,35 | 0,1175 | 0,848214837 | chr2 | + |
| Gle1 | 479,29 | -0,0909 | 0,795441927 | chr2 | + |
| Gm10804 | 9,35 | -0,8326 | 0,172087384 | chr2 | + |
| Gm13375 | 17,94 | -0,0196 | 0,98507181 | chr2 | + |
| Gm13483 | 1,81 | 0,3582 | NA | chr2 | + |
| Gm13490 | 6,31 | 0,5167 | 0,453331199 | chr2 | - |
| Gm13710 | 7,67 | 0,6141 | 0,35301802 | chr2 | - |
| Gm13889 | 9,80 | -0,3413 | 0,656897388 | chr2 | - |
| Gm14005 | 19,78 | 0,2077 | 0,766052367 | chr2 | - |
| Gm14023 | 20,05 | 0,6970 | 0,237903839 | chr2 | + |

| | | | | | |
|---|---|---|---|---|---|
| Gm14204 | 8,27 | 0,2231 | 0,792197952 | chr2 | - |
| Gm14207 | 3,33 | -0,3091 | 0,687523396 | chr2 | - |
| Gm14295 | 388,09 | 0,2229 | 0,702177047 | chr2 | + |
| Gm14305 | 11,76 | 0,7966 | 0,197263027 | chr2 | + |
| Gm14322 | 144,46 | 0,2059 | 0,73216561 | chr2 | + |
| Gm14325 | 218,10 | 0,1872 | 0,739141404 | chr2 | - |
| Gm14326 | 56,71 | 0,0855 | 0,905147397 | chr2 | - |
| Gm14327 | 8,27 | 0,8456 | 0,166586629 | chr2 | + |
| Gm14393 | 7,91 | 0,3056 | 0,695711387 | chr2 | - |
| Gm14403 | 719,05 | 0,0822 | 0,909096613 | chr2 | + |
| Zfp970 | 1372,34 | 0,6607 | 0,108917495 | chr2 | + |
| Zfp973 | 87,51 | 0,6500 | 0,073845576 | chr2 | + |
| Gm17762 | 4,62 | 0,3628 | 0,62632688 | chr2 | + |
| Gm3230 | 2,81 | 0,0088 | 0,992976309 | chr2 | - |
| Smim26 | 478,85 | -0,4032 | 0,245392711 | chr2 | + |
| Mir1a-1hg | 3,26 | 0,2511 | 0,743199461 | chr2 | + |
| Gm826 | 56,62 | -0,5138 | 0,191230937 | chr2 | - |
| Gmeb2 | 132,04 | -0,1996 | 0,696074872 | chr2 | - |
| Golga1 | 242,42 | 0,0401 | 0,947294154 | chr2 | - |
| Golga2 | 338,30 | -0,0039 | 0,993598948 | chr2 | + |
| Gorasp2 | 996,86 | -0,4338 | 0,093284325 | chr2 | + |
| Gpat2 | 40,16 | 0,0682 | 0,926972552 | chr2 | + |
| Gpcpd1 | 2208,32 | -0,7216 | 0,113373897 | chr2 | - |
| Gpd2 | 714,31 | 0,2948 | 0,230902599 | chr2 | + |
| Gpr107 | 272,44 | 0,0299 | 0,955440194 | chr2 | + |
| Gpr155 | 487,11 | -0,2988 | 0,582441358 | chr2 | - |
| Gpsm1 | 10,42 | 0,2462 | 0,764664236 | chr2 | + |
| Grb14 | 857,16 | -0,2843 | 0,376807903 | chr2 | - |
| Gsn | 487,55 | 0,3437 | 0,432202467 | chr2 | + |
| Gss | 1427,90 | -0,1783 | 0,722254721 | chr2 | - |
| Gtf3c4 | 465,04 | 0,1827 | 0,568858283 | chr2 | - |
| Gtf3c5 | 100,54 | 0,4049 | 0,372833388 | chr2 | - |
| Large2 | 4,01 | -0,1576 | 0,858880914 | chr2 | - |
| Gzf1 | 916,65 | 0,2872 | 0,289586014 | chr2 | + |
| Hao1 | 8744,83 | -0,2456 | 0,523898331 | chr2 | - |
| Harbi1 | 280,44 | 0,4309 | 0,188249599 | chr2 | + |
| Hat1 | 400,44 | 0,3823 | 0,328395075 | chr2 | + |
| Haus2 | 264,68 | 0,2447 | 0,564495903 | chr2 | + |
| Hc | 17069,22 | -0,2218 | 0,557071603 | chr2 | - |
| Hck | 83,06 | 0,0448 | 0,948314012 | chr2 | + |
| Hdc | 44,90 | 0,2592 | 0,672205411 | chr2 | - |
| Helz2 | 726,37 | -0,1255 | 0,812100099 | chr2 | - |
| Hipk3 | 1037,23 | -0,0734 | 0,871700795 | chr2 | - |
| Hnf4a | 7444,34 | -0,4082 | 0,109872624 | chr2 | + |
| Hnf4aos | 127,47 | -0,2035 | 0,692167613 | chr2 | - |
| Hnmt | 916,26 | 0,3891 | 0,395114656 | chr2 | - |
| Hnrnpa3 | 1551,35 | -0,0871 | 0,820051729 | chr2 | + |
| Hsd17b12 | 4086,49 | -0,1950 | 0,44242629 | chr2 | - |
| Hspa12b | 18,28 | 0,1668 | 0,836598726 | chr2 | + |
| Hspa14 | 238,17 | -0,0942 | 0,835615523 | chr2 | - |

| | | | | | |
|---|---|---|---|---|---|
| Hspa5 | 15094,88 | 0,0207 | 0,976991666 | chr2 | + |
| Hypk | 1815,49 | -0,2588 | 0,285552483 | chr2 | + |
| Id1 | 80,54 | 0,2065 | 0,803952241 | chr2 | + |
| Idh3b | 3039,85 | -0,2481 | 0,454992288 | chr2 | - |
| Ier5l | 2,69 | -0,0021 | 0,997549615 | chr2 | - |
| Ifih1 | 827,09 | 0,6706 | 0,073430016 | chr2 | - |
| Ift52 | 445,80 | -0,2754 | 0,333519645 | chr2 | + |
| Il15ra | 441,64 | -0,4462 | 0,175625239 | chr2 | + |
| Il1a | 157,43 | 0,5372 | 0,321599847 | chr2 | - |
| Il1b | 93,15 | 0,5169 | 0,407602327 | chr2 | - |
| Il1bos | 5,19 | -0,1979 | 0,80096016 | chr2 | + |
| Il1f9 | 31,95 | -0,1866 | 0,763580116 | chr2 | + |
| Immp1l | 838,90 | -0,1099 | 0,796541442 | chr2 | + |
| Ino80 | 197,50 | -0,0956 | 0,841012567 | chr2 | - |
| Inpp5e | 62,44 | 0,0068 | 0,992806079 | chr2 | - |
| Itch | 1039,27 | -0,2401 | 0,490133823 | chr2 | + |
| Itga4 | 97,53 | -0,3608 | 0,373653254 | chr2 | + |
| Itga6 | 70,64 | 0,6843 | 0,108040011 | chr2 | + |
| Itga8 | 121,16 | 0,4669 | 0,288287372 | chr2 | - |
| Itgav | 403,83 | -0,0677 | 0,886446111 | chr2 | + |
| Itgb6 | 2,30 | 0,1877 | NA | chr2 | - |
| Itih2 | 21023,81 | 0,2398 | 0,397488617 | chr2 | - |
| Itpa | 837,50 | -0,0946 | 0,858880859 | chr2 | + |
| Itpka | 23,08 | -0,2567 | 0,69217177 | chr2 | + |
| Itpripl1 | 115,40 | 0,4260 | 0,269410662 | chr2 | - |
| Ivd | 2413,09 | -0,1489 | 0,638353832 | chr2 | + |
| Jag1 | 53,90 | -0,4150 | 0,432202467 | chr2 | - |
| Jmjd7 | 3,90 | -0,0132 | 0,990189789 | chr2 | + |
| Katnbl1 | 132,42 | -0,1047 | 0,823578308 | chr2 | + |
| Kbtbd4 | 248,76 | 0,0977 | 0,842655851 | chr2 | + |
| Kcnb1 | 36,02 | 0,1112 | 0,869067375 | chr2 | - |
| Kcnip3 | 7,05 | 0,0396 | 0,970245243 | chr2 | - |
| Kif16b | 134,24 | -0,0134 | 0,982209929 | chr2 | - |
| Kif18a | 10,14 | -0,0443 | 0,964620526 | chr2 | + |
| Kif3b | 303,83 | 0,1850 | 0,701104843 | chr2 | + |
| Kif5c | 2,08 | 0,4642 | NA | chr2 | + |
| Kin | 177,93 | -0,4634 | 0,125541699 | chr2 | + |
| Kiz | 321,61 | -0,2440 | 0,452409282 | chr2 | + |
| Klhl23 | 23,59 | 0,4415 | 0,44242629 | chr2 | + |
| Klhl41 | 7,42 | -0,0615 | 0,950819236 | chr2 | + |
| Knstrn | 17,09 | -0,0516 | 0,955504626 | chr2 | + |
| Kynu | 4284,91 | -0,2953 | 0,343973842 | chr2 | + |
| Lama5 | 29,40 | 0,0595 | 0,949529403 | chr2 | - |
| Lamc3 | 14,98 | -0,3554 | 0,587901189 | chr2 | + |
| Lbp | 1708,60 | -0,4427 | 0,405102408 | chr2 | + |
| Lcmt2 | 178,22 | -0,0983 | 0,830422518 | chr2 | - |
| Lcn12 | 2,15 | 0,1887 | NA | chr2 | - |
| Lcn2 | 508,88 | 0,2682 | 0,709072206 | chr2 | - |
| Lcn8 | 4,02 | 0,2299 | 0,781051459 | chr2 | + |
| Ldlrad3 | 108,12 | 0,1439 | 0,784877345 | chr2 | - |

| | | | | | |
|---|---|---|---|---|---|
| Lgr4 | 1280,80 | -0,0980 | 0,811137679 | chr2 | + |
| Lhx2 | 16,21 | -0,0894 | 0,913936879 | chr2 | + |
| Lhx6 | 76,26 | 0,2355 | 0,704776957 | chr2 | - |
| Lime1 | 93,42 | -0,4325 | 0,429831394 | chr2 | + |
| Lin7c | 1023,33 | 0,1215 | 0,803874753 | chr2 | + |
| Lmo2 | 66,99 | 0,1887 | 0,700470531 | chr2 | + |
| Lnpk | 136,92 | -0,0915 | 0,868479494 | chr2 | - |
| Lpin3 | 52,60 | 0,1537 | 0,823949825 | chr2 | + |
| Lrp4 | 424,79 | -0,1926 | 0,662842062 | chr2 | + |
| Lrrc4c | 13,02 | 0,4054 | 0,580859417 | chr2 | + |
| Lrrc57 | 189,46 | 0,2022 | 0,68354255 | chr2 | - |
| Lrrc8a | 298,90 | -0,5374 | 0,103715866 | chr2 | + |
| Lrrn4 | 4,64 | 0,1279 | 0,88569455 | chr2 | - |
| Lrsam1 | 137,41 | 0,3810 | 0,296372739 | chr2 | - |
| Lsm14b | 438,67 | -0,1267 | 0,691615955 | chr2 | + |
| Ltk | 4,30 | 0,1029 | 0,912216728 | chr2 | - |
| Ly75 | 26,98 | -0,1219 | 0,886483612 | chr2 | - |
| Lzts3 | 282,90 | 0,0514 | 0,925898178 | chr2 | - |
| Macrod2 | 193,04 | -0,4520 | 0,343440412 | chr2 | + |
| Madd | 220,58 | 0,2849 | 0,497721225 | chr2 | - |
| Mafb | 316,96 | -0,4732 | 0,162272166 | chr2 | - |
| Mal | 7,52 | -0,2686 | 0,740663168 | chr2 | - |
| Mamdc4 | 7,15 | -0,6990 | 0,271992655 | chr2 | - |
| Man1b1 | 588,97 | 0,2181 | 0,54877155 | chr2 | + |
| Manbal | 175,14 | 0,2359 | 0,510160643 | chr2 | + |
| Map1a | 3,20 | -0,2567 | 0,738680713 | chr2 | + |
| Map1lc3a | 2383,98 | -0,4985 | 0,096261775 | chr2 | + |
| Mapk8ip1 | 21,41 | 0,1301 | 0,865199339 | chr2 | - |
| Mapkap1 | 528,90 | -0,3112 | 0,268604776 | chr2 | + |
| Mapkbp1 | 44,88 | 0,1972 | 0,740538349 | chr2 | + |
| Mapre1 | 609,60 | 0,1398 | 0,677429745 | chr2 | + |
| March7 | 227,35 | 0,3486 | 0,387031047 | chr2 | + |
| Mastl | 7,04 | 0,5366 | 0,43131925 | chr2 | - |
| Mavs | 346,55 | -0,0711 | 0,886446111 | chr2 | + |
| Mbd5 | 57,28 | -0,0074 | 0,99143613 | chr2 | + |
| Mcm8 | 43,45 | 0,4585 | 0,364379698 | chr2 | + |
| Mcts2 | 133,71 | -0,5058 | 0,215046281 | chr2 | + |
| Mdk | 7,36 | 0,0470 | 0,964556444 | chr2 | - |
| Med19 | 247,73 | -0,1899 | 0,592046274 | chr2 | + |
| Med22 | 449,26 | 0,0814 | 0,87093463 | chr2 | - |
| Med27 | 81,52 | -0,1449 | 0,802594059 | chr2 | + |
| Meis2 | 53,92 | 0,3794 | 0,415216302 | chr2 | - |
| Mertk | 203,20 | 0,0192 | 0,970245243 | chr2 | + |
| Metap1d | 542,22 | -0,3200 | 0,310756756 | chr2 | + |
| Mettl5 | 279,67 | -0,0646 | 0,89130564 | chr2 | - |
| Mettl8 | 207,07 | -0,1796 | 0,640132917 | chr2 | - |
| Mfap1a | 216,54 | 0,1408 | 0,771616285 | chr2 | - |
| Mfap1b | 17,98 | -0,3352 | 0,618512306 | chr2 | - |
| Mgme1 | 159,16 | 0,2020 | 0,588219472 | chr2 | + |
| Mkks | 891,45 | -0,3047 | 0,27912309 | chr2 | - |

| | | | | | |
|---|---|---|---|---|---|
| Mllt10 | 311,91 | 0,1406 | 0,735883435 | chr2 | + |
| Mmadhc | 1561,84 | -0,1626 | 0,652252449 | chr2 | - |
| Mmp9 | 5,73 | 0,1812 | 0,830422518 | chr2 | + |
| Mocs3 | 23,90 | -0,0034 | 0,996945533 | chr2 | + |
| Mrc1 | 1233,43 | 0,3225 | 0,31487946 | chr2 | + |
| Mrgbp | 26,41 | -0,0108 | 0,990233106 | chr2 | + |
| Mrpl41 | 760,06 | -0,0298 | 0,946177485 | chr2 | - |
| Mrps2 | 447,78 | 0,0990 | 0,844155615 | chr2 | + |
| Mrps26 | 164,70 | -0,4568 | 0,188249599 | chr2 | + |
| Mrps5 | 710,72 | -0,1793 | 0,592295685 | chr2 | + |
| Mrrf | 617,36 | -0,0870 | 0,823578308 | chr2 | + |
| Msrb2 | 245,14 | 0,1699 | 0,655452587 | chr2 | + |
| Mtch2 | 4081,51 | -0,3073 | 0,19121789 | chr2 | + |
| Mtg2 | 280,63 | -0,1037 | 0,825289629 | chr2 | + |
| Mtx2 | 937,15 | -0,4511 | 0,167807438 | chr2 | + |
| Mvb12b | 26,33 | -0,1467 | 0,830330662 | chr2 | - |
| Mybl2 | 2,41 | -0,5011 | NA | chr2 | + |
| Myl9 | 82,00 | 0,4815 | 0,286794062 | chr2 | + |
| Naa20 | 371,11 | 0,2450 | 0,486009467 | chr2 | + |
| Nacc2 | 267,70 | -0,0642 | 0,884882423 | chr2 | - |
| Naif1 | 11,45 | 0,1761 | 0,836754446 | chr2 | + |
| Nanp | 335,33 | 0,1075 | 0,793239112 | chr2 | - |
| Nat10 | 256,71 | -0,0144 | 0,980186108 | chr2 | - |
| Ncaph | 22,45 | -0,1118 | 0,884370894 | chr2 | - |
| Nckap1 | 1756,81 | -0,0196 | 0,967536738 | chr2 | - |
| Ncoa3 | 270,54 | -0,1587 | 0,663872588 | chr2 | + |
| Ncoa5 | 118,52 | 0,0137 | 0,984995389 | chr2 | - |
| Ncoa6 | 206,83 | -0,3025 | 0,434423892 | chr2 | - |
| Ncs1 | 3,74 | 0,5319 | 0,425926605 | chr2 | + |
| Ndor1 | 159,32 | -0,2551 | 0,489321662 | chr2 | - |
| Ndrg3 | 316,99 | -0,2134 | 0,552537331 | chr2 | - |
| Ndufa8 | 1486,53 | -0,3315 | 0,392217524 | chr2 | - |
| Ndufaf1 | 696,25 | -0,1344 | 0,691018693 | chr2 | - |
| Ndufaf5 | 93,09 | 0,0513 | 0,931322892 | chr2 | + |
| Ndufs3 | 2787,34 | -0,2023 | 0,670839945 | chr2 | - |
| Neb | 216,63 | 0,5961 | 0,143003054 | chr2 | - |
| Nek6 | 724,23 | -0,0096 | 0,988512389 | chr2 | + |
| Nelfb | 284,08 | 0,0114 | 0,984055177 | chr2 | - |
| Nelfcd | 328,80 | -0,4332 | 0,287543614 | chr2 | + |
| Neurl2 | 11,20 | -0,2350 | 0,759805352 | chr2 | - |
| Nfatc2 | 5,29 | -0,1136 | 0,904430709 | chr2 | - |
| Nfe2l2 | 1054,46 | 0,0054 | 0,990846949 | chr2 | - |
| Nfs1 | 978,20 | -0,1613 | 0,588219472 | chr2 | - |
| Ninl | 19,03 | 0,5594 | 0,358990473 | chr2 | - |
| Nkain4 | 5,24 | -0,3390 | 0,659350082 | chr2 | - |
| Nmt2 | 465,96 | -0,3383 | 0,29013293 | chr2 | + |
| Nop10 | 1911,64 | -0,4151 | 0,22384344 | chr2 | + |
| Nostrin | 612,41 | 0,4893 | 0,209849508 | chr2 | + |
| Notch1 | 153,33 | 0,2099 | 0,658533075 | chr2 | - |
| Noxa1 | 2,63 | 0,5731 | 0,338536613 | chr2 | - |

| | | | | | |
|---|---|---|---|---|---|
| Npdc1 | 48,70 | 0,6918 | 0,161576375 | chr2 | + |
| Npepl1 | 558,04 | -0,5818 | 0,120192741 | chr2 | + |
| Nr1h3 | 1863,52 | 0,0717 | 0,885528714 | chr2 | - |
| Nr4a2 | 6,06 | -0,1399 | 0,877942095 | chr2 | - |
| Nr6a1 | 118,21 | -0,1278 | 0,825518458 | chr2 | - |
| Nrarp | 13,38 | -0,2033 | 0,800238939 | chr2 | + |
| Nsfl1c | 1117,95 | -0,1669 | 0,598861459 | chr2 | + |
| Nsmf | 1039,70 | -0,3831 | 0,25884542 | chr2 | + |
| Nsun6 | 56,08 | 0,6146 | 0,215949298 | chr2 | - |
| Ntmt1 | 89,21 | -0,2553 | 0,654583626 | chr2 | + |
| Ntng2 | 4,61 | 0,1613 | 0,855874829 | chr2 | - |
| Nudt5 | 396,41 | 0,3145 | 0,334804026 | chr2 | + |
| Nup160 | 137,52 | 0,4544 | 0,266692824 | chr2 | + |
| Nup188 | 79,39 | 0,2979 | 0,526585296 | chr2 | + |
| Nup214 | 122,65 | -0,0807 | 0,871232434 | chr2 | + |
| Nup35 | 132,21 | 0,2444 | 0,513757407 | chr2 | + |
| Nusap1 | 20,58 | -0,0184 | 0,98507181 | chr2 | + |
| Nxt1 | 177,70 | -0,2490 | 0,645162115 | chr2 | + |
| Ocstamp | 10,55 | 0,5131 | 0,437402307 | chr2 | - |
| Odf2 | 116,09 | -0,4171 | 0,272679707 | chr2 | + |
| Ogfr | 598,97 | -0,1930 | 0,658008155 | chr2 | + |
| Oip5 | 10,17 | 0,9041 | 0,122100382 | chr2 | - |
| Olfm1 | 40,43 | 0,0624 | 0,93551479 | chr2 | + |
| Olfr1034 | 11,99 | 0,0923 | 0,922594928 | chr2 | + |
| Optn | 1225,16 | 0,0857 | 0,818303576 | chr2 | - |
| Orc4 | 366,52 | -0,1335 | 0,749269012 | chr2 | - |
| Osbpl2 | 463,99 | -0,2959 | 0,277831408 | chr2 | + |
| Oser1 | 563,62 | -0,2344 | 0,494351553 | chr2 | - |
| Otud1 | 76,52 | -0,1683 | 0,742638254 | chr2 | + |
| P2rx3 | 22,90 | 0,0246 | 0,981564504 | chr2 | - |
| Pabpc1l | 3,03 | 0,1879 | 0,817465502 | chr2 | + |
| Pacsin3 | 567,40 | 0,0792 | 0,877920448 | chr2 | + |
| Pamr1 | 73,69 | 0,1717 | 0,804380135 | chr2 | + |
| Pank2 | 17,90 | 0,7233 | 0,18870514 | chr2 | + |
| Pard6b | 48,57 | 0,0848 | 0,903337956 | chr2 | + |
| Pbx3 | 52,21 | -0,0678 | 0,916852736 | chr2 | - |
| Pced1a | 23,79 | 0,5892 | 0,311540287 | chr2 | - |
| Pcif1 | 112,85 | 0,0809 | 0,89147692 | chr2 | + |
| Pck1 | 41716,74 | -0,2740 | 0,5427973 | chr2 | + |
| Pcmtd2 | 766,14 | 0,3355 | 0,350490571 | chr2 | + |
| Pcna | 830,99 | 0,0738 | 0,862451616 | chr2 | - |
| Pdcl | 540,53 | 0,4054 | 0,228893066 | chr2 | - |
| Pde1a | 44,15 | 0,0862 | 0,909889201 | chr2 | - |
| Pdhx | 3306,95 | -0,0327 | 0,952532092 | chr2 | - |
| Pdia3 | 625,74 | -0,4298 | 0,123737394 | chr2 | + |
| Pdk1 | 1240,09 | 0,0704 | 0,916852736 | chr2 | + |
| Pdrg1 | 326,19 | -0,1035 | 0,804380135 | chr2 | - |
| Pdss1 | 35,93 | 0,1160 | 0,858288137 | chr2 | + |
| Pex16 | 1047,62 | -0,3265 | 0,269874551 | chr2 | + |
| Pfdn4 | 247,83 | 0,3237 | 0,546240378 | chr2 | + |

| | | | | | |
|---|---|---|---|---|---|
| Pfkfb3 | 400,39 | -0,0524 | 0,938068895 | chr2 | - |
| Phf19 | 5,89 | -0,3166 | 0,684393541 | chr2 | - |
| Phf20 | 171,75 | 0,0459 | 0,922594928 | chr2 | + |
| Phf21a | 149,97 | -0,1636 | 0,754953629 | chr2 | + |
| Phospho2 | 486,82 | -0,3787 | 0,209104214 | chr2 | + |
| Pigu | 248,23 | -0,1410 | 0,781342762 | chr2 | - |
| Pigt | 76,00 | 0,4876 | 0,216874749 | chr2 | + |
| Pip4k2a | 115,58 | 0,3023 | 0,505514525 | chr2 | - |
| Pkig | 174,23 | 0,1814 | 0,719564062 | chr2 | + |
| Pkn3 | 9,10 | -0,3603 | 0,62914039 | chr2 | + |
| Pkp4 | 961,34 | -0,5013 | 0,066396922 | chr2 | + |
| Pla2r1 | 61,94 | 0,2040 | 0,729883372 | chr2 | - |
| Plagl2 | 60,89 | 0,1750 | 0,73118289 | chr2 | - |
| Plcb1 | 685,87 | -0,4691 | 0,10978541 | chr2 | + |
| Plcb2 | 244,14 | -0,6468 | 0,075466772 | chr2 | - |
| Plcb4 | 20,07 | 0,3724 | 0,538245453 | chr2 | + |
| Plcg1 | 258,33 | -0,3273 | 0,432202467 | chr2 | + |
| Plekha3 | 315,73 | -0,2200 | 0,560024349 | chr2 | + |
| Pltp | 14,17 | 0,1530 | 0,858880914 | chr2 | - |
| Pmepa1 | 24,45 | 0,0658 | 0,940538622 | chr2 | - |
| Pmpca | 1226,85 | -0,0905 | 0,773900067 | chr2 | + |
| Pnpla7 | 2694,88 | 0,0405 | 0,943108135 | chr2 | + |
| Pofut1 | 237,53 | 0,0788 | 0,856476342 | chr2 | + |
| Polr1b | 161,09 | -0,3129 | 0,411119108 | chr2 | + |
| Polr3f | 184,33 | -0,0473 | 0,935698635 | chr2 | + |
| Polr3k | 456,94 | 0,1404 | 0,746414189 | chr2 | + |
| Pomt1 | 165,69 | 0,1033 | 0,819196734 | chr2 | + |
| Plpp7 | 8,12 | 0,1788 | 0,83796616 | chr2 | + |
| Ppdpf | 691,26 | -0,4759 | 0,25333134 | chr2 | + |
| Ppig | 552,25 | -0,0680 | 0,880810135 | chr2 | + |
| Ppip5k1 | 53,39 | 0,6427 | 0,153470778 | chr2 | - |
| Ppp1r16b | 60,58 | -0,1829 | 0,743199461 | chr2 | + |
| Ppp1r26 | 3,10 | 0,3680 | 0,610390411 | chr2 | + |
| Ppp1r3d | 4,72 | 0,2491 | 0,758873213 | chr2 | - |
| Ptpa | 547,78 | -0,1158 | 0,794186768 | chr2 | + |
| Ppp4r1l-ps | 35,82 | -0,5434 | 0,303602228 | chr2 | - |
| Ppp6c | 755,29 | -0,1087 | 0,781643256 | chr2 | - |
| Prex1 | 49,14 | 0,2613 | 0,618749555 | chr2 | - |
| Prkcq | 32,49 | -0,0346 | 0,965925759 | chr2 | + |
| Prkra | 290,01 | -0,4706 | 0,092854537 | chr2 | - |
| Prnp | 976,14 | -0,0332 | 0,944028647 | chr2 | + |
| Procr | 14,19 | 0,6370 | 0,280102462 | chr2 | + |
| Proser2 | 97,91 | -0,2461 | 0,611268902 | chr2 | - |
| Prpf18 | 823,60 | 0,1385 | 0,754249138 | chr2 | - |
| Prpf40a | 649,78 | 0,2180 | 0,612061669 | chr2 | - |
| Prpf6 | 813,29 | -0,1295 | 0,77481989 | chr2 | + |
| Prr5l | 24,67 | 0,2694 | 0,680066999 | chr2 | - |
| Prrc2b | 505,50 | -0,0932 | 0,847419446 | chr2 | + |
| Prrg4 | 12,12 | 0,8577 | 0,151178014 | chr2 | - |
| Psd4 | 36,99 | 0,4329 | 0,49028079 | chr2 | + |

| | | | | | |
|---|---|---|---|---|---|
| Psma7 | 2270,34 | -0,3439 | 0,396372043 | chr2 | - |
| Psmb7 | 5029,43 | -0,2512 | 0,411673886 | chr2 | - |
| Psmc3 | 2667,57 | -0,0982 | 0,759406597 | chr2 | + |
| Psmd14 | 1142,74 | -0,2977 | 0,317838059 | chr2 | + |
| Psmd5 | 814,19 | -0,0902 | 0,810120608 | chr2 | - |
| Psmf1 | 396,05 | -0,0349 | 0,938329173 | chr2 | - |
| Pter | 1647,91 | 0,3112 | 0,310109812 | chr2 | + |
| Ptgds | 36,67 | -0,0390 | 0,969512928 | chr2 | - |
| Ptges | 9,39 | -0,0087 | 0,993445259 | chr2 | - |
| Ptges2 | 282,39 | -0,2805 | 0,394599707 | chr2 | + |
| Ptgis | 28,58 | 0,2593 | 0,691951516 | chr2 | - |
| Ptgs1 | 245,12 | -0,1148 | 0,814331785 | chr2 | + |
| Ptk6 | 7,62 | 0,5571 | 0,402592894 | chr2 | - |
| Hacd1 | 200,72 | -0,5348 | 0,210347908 | chr2 | - |
| Ptpmt1 | 1093,70 | -0,2802 | 0,39810264 | chr2 | - |
| Ptpn1 | 264,11 | -0,2047 | 0,685921923 | chr2 | + |
| Ptpra | 466,23 | 0,1829 | 0,639465027 | chr2 | + |
| Ptprj | 968,12 | 0,1253 | 0,722057512 | chr2 | - |
| Ptprt | 7,36 | 0,4774 | 0,501682426 | chr2 | - |
| Ptprtos | 2,92 | 0,9062 | 0,08440425 | chr2 | + |
| Ptrh1 | 9,26 | 0,0283 | 0,979212524 | chr2 | + |
| Pxmp4 | 1025,62 | -0,1238 | 0,854019408 | chr2 | - |
| Qser1 | 290,15 | -0,0364 | 0,939403124 | chr2 | - |
| Qsox2 | 53,36 | -0,1121 | 0,874763133 | chr2 | - |
| Rab14 | 2333,48 | -0,0310 | 0,938499724 | chr2 | - |
| Rabepk | 160,22 | -0,3952 | 0,228728627 | chr2 | - |
| Rabgap1 | 432,97 | 0,1565 | 0,665098911 | chr2 | + |
| Rabl6 | 260,57 | -0,3733 | 0,212210133 | chr2 | - |
| Ncoa3 | 23,09 | 0,7912 | 0,117064174 | chr2 | + |
| Rad51 | 11,77 | 0,0111 | 0,9909028 | chr2 | + |
| Rae1 | 239,33 | -0,2931 | 0,331966576 | chr2 | + |
| Ralgapa2 | 544,79 | -0,1689 | 0,703510119 | chr2 | - |
| Ralgapb | 236,20 | -0,0151 | 0,979212524 | chr2 | + |
| Ralgds | 25,47 | -0,0447 | 0,959379389 | chr2 | + |
| Ralgps1 | 70,92 | 0,5726 | 0,201099521 | chr2 | - |
| Raly | 792,44 | -0,3991 | 0,15392197 | chr2 | + |
| Rapgef1 | 465,80 | -0,3940 | 0,196118975 | chr2 | + |
| Rapgef4 | 3120,37 | 0,1910 | 0,658008155 | chr2 | + |
| Rasgrp1 | 30,45 | -0,0639 | 0,941754349 | chr2 | - |
| Rassf2 | 29,38 | 0,1457 | 0,829783511 | chr2 | - |
| Rbbp9 | 472,24 | 0,3118 | 0,465906639 | chr2 | - |
| Rbck1 | 438,66 | -0,3448 | 0,352106106 | chr2 | - |
| Rbl1 | 38,91 | 0,8094 | 0,05400727 | chr2 | - |
| Rbm12 | 93,62 | 0,0443 | 0,943305207 | chr2 | - |
| Rbm17 | 513,86 | -0,3402 | 0,217693536 | chr2 | - |
| Rbm18 | 300,06 | -0,0610 | 0,888710003 | chr2 | - |
| Rbm38 | 26,67 | -0,4999 | 0,396094068 | chr2 | + |
| Rbm39 | 2007,24 | 0,1557 | 0,690557571 | chr2 | - |
| Rbm45 | 148,47 | -0,2269 | 0,596546314 | chr2 | + |
| Rbms1 | 283,09 | 0,2821 | 0,444662541 | chr2 | - |

| | | | | | |
|---|---|---|---|---|---|
| Rc3h2 | 429,96 | 0,0163 | 0,979051488 | chr2 | - |
| Rcn1 | 229,30 | 0,4621 | 0,100572892 | chr2 | - |
| Rem1 | 4,46 | -0,0465 | 0,963027434 | chr2 | + |
| Rexo4 | 358,42 | -0,1487 | 0,721569914 | chr2 | - |
| Rgs19 | 28,02 | 0,3043 | 0,642834828 | chr2 | - |
| Rif1 | 185,09 | 0,4231 | 0,288929615 | chr2 | + |
| Rin2 | 205,14 | -0,0331 | 0,951292919 | chr2 | + |
| Rmdn3 | 1137,74 | 0,0637 | 0,862616891 | chr2 | - |
| Rnd3 | 464,00 | -0,1360 | 0,767917363 | chr2 | - |
| Rnf114 | 912,20 | -0,2789 | 0,301862555 | chr2 | + |
| Rnf24 | 174,22 | -0,2366 | 0,729822171 | chr2 | - |
| Romo1 | 1141,33 | -0,1561 | 0,860107955 | chr2 | + |
| Rpap1 | 81,14 | 0,3789 | 0,32406341 | chr2 | - |
| Rpl12 | 4983,96 | -0,4926 | 0,358779319 | chr2 | + |
| Rpl7a | 8613,26 | -0,3018 | 0,247921371 | chr2 | + |
| Rpn2 | 2635,71 | -0,3195 | 0,202827316 | chr2 | + |
| Rpp38 | 66,29 | 0,3795 | 0,401045574 | chr2 | - |
| Rprd1b | 320,52 | -0,1062 | 0,764664236 | chr2 | + |
| Rps21 | 3783,48 | -0,6114 | 0,162740196 | chr2 | + |
| Rpusd2 | 52,37 | 0,3597 | 0,433006548 | chr2 | + |
| Rrbp1 | 5357,81 | -0,4276 | 0,180910407 | chr2 | - |
| Rsu1 | 494,21 | 0,0902 | 0,809065986 | chr2 | - |
| Rtel1 | 136,10 | 0,1848 | 0,670706543 | chr2 | + |
| Rtf1 | 349,41 | -0,2270 | 0,441724922 | chr2 | + |
| Rxra | 4036,36 | -0,3132 | 0,299245205 | chr2 | + |
| Samhd1 | 251,85 | 0,4585 | 0,118802947 | chr2 | - |
| Sardh | 8980,44 | 0,1876 | 0,585630763 | chr2 | - |
| Scai | 99,49 | 0,3333 | 0,441570242 | chr2 | - |
| Scand1 | 24,29 | -0,1848 | 0,801768605 | chr2 | - |
| Scn2a | 5,35 | 0,2669 | 0,743271454 | chr2 | + |
| Scn3a | 4,52 | -0,2089 | 0,806159394 | chr2 | - |
| Scn7a | 7,27 | 0,5410 | 0,427388057 | chr2 | - |
| Scrn3 | 1140,32 | 0,3710 | 0,295992229 | chr2 | + |
| Sdc4 | 7775,92 | -0,0207 | 0,961098177 | chr2 | - |
| Sdcbp2 | 2,94 | 0,3412 | 0,640132917 | chr2 | + |
| Sec16a | 886,05 | -0,1180 | 0,769677581 | chr2 | - |
| Sec23b | 718,56 | 0,2365 | 0,595821824 | chr2 | + |
| Sec61a2 | 89,13 | -0,5114 | 0,187760013 | chr2 | - |
| Secisbp2l | 817,72 | 0,1600 | 0,691615955 | chr2 | - |
| Sema6d | 137,50 | 0,3483 | 0,402475192 | chr2 | + |
| Sephs1 | 610,87 | -0,1485 | 0,702699676 | chr2 | + |
| Serf2 | 3568,00 | -0,2586 | 0,454695883 | chr2 | + |
| Serinc3 | 5876,78 | 0,0200 | 0,95463264 | chr2 | - |
| Serping1 | 19221,29 | 0,3121 | 0,267579152 | chr2 | - |
| Sestd1 | 39,59 | -0,4793 | 0,356860747 | chr2 | - |
| Set | 2141,56 | -0,0894 | 0,806401686 | chr2 | + |
| Setx | 418,81 | 0,0756 | 0,860078935 | chr2 | + |
| Sfmbt2 | 3,51 | 0,1284 | 0,885863143 | chr2 | + |
| Sgk2 | 838,87 | -0,3444 | 0,486967566 | chr2 | + |
| Sh2d3c | 7,43 | 0,8100 | 0,190969312 | chr2 | + |

| | | | | | |
|---|---|---|---|---|---|
| Sh3glb2 | 180,46 | -0,4866 | 0,23347764 | chr2 | - |
| Shc4 | 8,00 | 0,4790 | 0,499958086 | chr2 | - |
| Shf | 80,63 | 0,0062 | 0,992896932 | chr2 | - |
| Siglec1 | 111,05 | 0,3417 | 0,408202808 | chr2 | - |
| Sirpa | 286,13 | 0,2636 | 0,482787957 | chr2 | + |
| Sla2 | 2,39 | 0,4131 | NA | chr2 | - |
| Slc12a6 | 17,31 | -0,2651 | 0,711159012 | chr2 | + |
| Slc12a5 | 797,92 | -0,2432 | 0,499122607 | chr2 | + |
| Slc17a9 | 84,39 | 0,4820 | 0,330697335 | chr2 | + |
| Slc1a2 | 2688,46 | -0,4338 | 0,231622612 | chr2 | + |
| Slc20a1 | 739,07 | -0,2087 | 0,609614861 | chr2 | + |
| Slc23a2 | 387,08 | 0,2291 | 0,701094188 | chr2 | - |
| Slc24a3 | 6,03 | 0,2188 | 0,795796985 | chr2 | + |
| Slc24a5 | 14,08 | 0,6799 | 0,242789888 | chr2 | + |
| Slc25a12 | 308,35 | -0,0592 | 0,901481982 | chr2 | - |
| Slc25a25 | 3626,06 | -0,2605 | 0,655462854 | chr2 | - |
| Slc27a4 | 450,08 | -0,3854 | 0,171449805 | chr2 | + |
| Slc28a2 | 12,78 | 0,5137 | 0,42314498 | chr2 | + |
| Slc2a10 | 3,37 | -0,0903 | 0,920978973 | chr2 | + |
| Slc2a4rg-ps | 84,74 | -0,6608 | 0,174902719 | chr2 | + |
| Slc2a6 | 2,98 | 0,2807 | 0,695093611 | chr2 | - |
| Slc2a8 | 128,45 | -0,5138 | 0,213281186 | chr2 | - |
| Slc30a4 | 344,78 | 0,1227 | 0,770543183 | chr2 | - |
| Slc35c1 | 194,72 | 0,0428 | 0,948668792 | chr2 | - |
| Slc35c2 | 392,39 | -0,1597 | 0,744313101 | chr2 | - |
| Slc39a12 | 2,19 | 0,0763 | NA | chr2 | + |
| Slc39a13 | 91,90 | 0,1985 | 0,726344964 | chr2 | - |
| Slc43a1 | 125,89 | 0,5719 | 0,266630306 | chr2 | + |
| Slc43a3 | 619,75 | 0,3424 | 0,287408362 | chr2 | + |
| Slc52a3 | 7,15 | 0,4966 | 0,477715768 | chr2 | + |
| Slc9a8 | 389,96 | -0,1170 | 0,743138956 | chr2 | + |
| Slco4a1 | 3,23 | -0,4335 | 0,539958873 | chr2 | + |
| Prelid3b | 720,96 | -0,3767 | 0,225575035 | chr2 | - |
| Slx4ip | 32,93 | 0,7753 | 0,085593171 | chr2 | + |
| Smox | 43,48 | -0,3247 | 0,54877155 | chr2 | + |
| Snai1 | 5,69 | 0,0800 | 0,937247252 | chr2 | + |
| Snap23 | 726,11 | 0,3137 | 0,268604776 | chr2 | + |
| Snhg7 | 35,96 | -0,4738 | 0,396254007 | chr2 | - |
| Snora65 | 8,57 | -0,2699 | 0,737568553 | chr2 | + |
| Snord12 | 5,04 | -0,1147 | 0,903060276 | chr2 | + |
| Snord17 | 3,21 | -0,2543 | 0,743459124 | chr2 | - |
| Snord57 | 2,49 | 0,2831 | NA | chr2 | + |
| Snrnp200 | 1040,33 | 0,1237 | 0,7950804 | chr2 | + |
| Snrpb | 594,66 | -0,3873 | 0,433006548 | chr2 | - |
| Snrpb2 | 433,01 | 0,1494 | 0,781284538 | chr2 | + |
| Snta1 | 293,05 | 0,4355 | 0,235696806 | chr2 | - |
| Snx21 | 49,65 | -0,6137 | 0,186273168 | chr2 | + |
| Snx5 | 1308,35 | -0,0887 | 0,797559244 | chr2 | - |
| Soga1 | 37,60 | 0,5528 | 0,237499459 | chr2 | - |
| Sord | 24242,07 | -0,3608 | 0,158783149 | chr2 | + |

| | | | | | |
|---|---|---|---|---|---|
| Sox12 | 38,53 | -0,4969 | 0,39354827 | chr2 | - |
| Sox18 | 30,86 | 0,1737 | 0,816750599 | chr2 | - |
| Sp3os | 16,44 | -0,1333 | 0,87211718 | chr2 | + |
| Spag4 | 10,42 | 0,2626 | 0,744313101 | chr2 | + |
| Spata2 | 115,29 | -0,2479 | 0,564849438 | chr2 | - |
| Spata5l1 | 9,76 | 0,3513 | 0,644046286 | chr2 | + |
| Spef1 | 15,94 | -0,2796 | 0,717188441 | chr2 | - |
| Spg11 | 229,50 | 0,1043 | 0,804418611 | chr2 | - |
| Spi1 | 48,76 | -0,1362 | 0,854690386 | chr2 | + |
| Spint1 | 10,93 | 0,5006 | 0,439736908 | chr2 | + |
| Spopl | 85,46 | 0,1553 | 0,776119796 | chr2 | - |
| Sppl2a | 4068,03 | 0,0579 | 0,901445192 | chr2 | - |
| Spred1 | 341,13 | 0,0545 | 0,914198498 | chr2 | + |
| Sptan1 | 1483,89 | 0,3474 | 0,120869486 | chr2 | + |
| Sqor | 4470,73 | 0,1700 | 0,646617701 | chr2 | + |
| Srms | 7,14 | 0,3428 | 0,656130094 | chr2 | - |
| Srp14 | 948,38 | -0,1067 | 0,7950804 | chr2 | - |
| Srxn1 | 630,11 | 0,1552 | 0,757883527 | chr2 | + |
| Ss18l1 | 59,73 | -0,2119 | 0,666980604 | chr2 | + |
| Ssb | 1339,22 | 0,1446 | 0,766178949 | chr2 | + |
| Ssfa2 | 790,68 | 0,0679 | 0,875995735 | chr2 | + |
| Ssna1 | 287,90 | -0,2184 | 0,627875496 | chr2 | - |
| Ssrp1 | 968,34 | 0,0788 | 0,863490375 | chr2 | + |
| St6galnac4 | 25,21 | 0,7392 | 0,172087384 | chr2 | + |
| St6galnac6 | 647,68 | -0,1205 | 0,803863835 | chr2 | + |
| Stam | 313,26 | -0,0105 | 0,986691613 | chr2 | + |
| Stam2 | 243,66 | 0,0648 | 0,886483612 | chr2 | - |
| Stamos | 24,30 | 0,3200 | 0,607310966 | chr2 | - |
| Stard7 | 2723,58 | -0,2632 | 0,439033663 | chr2 | + |
| Stau1 | 724,47 | -0,3445 | 0,202636202 | chr2 | - |
| Stk35 | 242,64 | -0,2482 | 0,525454185 | chr2 | + |
| Stk39 | 40,37 | 0,3551 | 0,523898331 | chr2 | - |
| Stk4 | 372,89 | 0,4119 | 0,286794062 | chr2 | + |
| Stom | 179,71 | -0,1871 | 0,7242192 | chr2 | - |
| Stx16 | 350,62 | -0,2625 | 0,500561834 | chr2 | + |
| Stxbp1 | 17,13 | 0,3432 | 0,595821824 | chr2 | - |
| Sulf2 | 90,78 | -0,2307 | 0,617869186 | chr2 | - |
| Surf1 | 692,71 | -0,3520 | 0,336247849 | chr2 | - |
| Surf2 | 229,57 | -0,2210 | 0,519383722 | chr2 | + |
| Surf4 | 3489,95 | -0,2950 | 0,36985369 | chr2 | - |
| Surf6 | 257,78 | -0,1724 | 0,723326073 | chr2 | - |
| Suv39h2 | 12,05 | 0,5502 | 0,391001304 | chr2 | - |
| Swi5 | 3343,83 | -0,0799 | 0,817465502 | chr2 | - |
| Sys1 | 325,81 | -0,2073 | 0,587901189 | chr2 | + |
| Taf3 | 54,00 | -0,3800 | 0,403228071 | chr2 | - |
| Taf4 | 135,33 | -0,5434 | 0,089014161 | chr2 | - |
| Tanc1 | 127,44 | 0,2523 | 0,586201597 | chr2 | + |
| Tank | 342,64 | 0,5099 | 0,194249019 | chr2 | + |
| Tasp1 | 166,22 | 0,2142 | 0,631595585 | chr2 | - |
| Tbc1d13 | 380,64 | 0,1181 | 0,774082864 | chr2 | + |

| | | | | | |
|---|---|---|---|---|---|
| Tbc1d20 | 255,55 | 0,0010 | 0,998203916 | chr2 | + |
| Tcea2 | 9,70 | -0,2577 | 0,747022531 | chr2 | + |
| Tcp11l1 | 11,73 | 0,5511 | 0,384370645 | chr2 | - |
| Tfpi | 270,04 | 0,3703 | 0,273713146 | chr2 | - |
| Tgif2 | 6,09 | 0,3180 | 0,682841407 | chr2 | + |
| Tgm2 | 2131,57 | 0,5616 | 0,087353493 | chr2 | - |
| Thbd | 61,31 | 0,1974 | 0,71673612 | chr2 | - |
| Thbs1 | 24,60 | 0,6362 | 0,234220487 | chr2 | + |
| Them7 | 241,37 | 0,2127 | 0,677405328 | chr2 | + |
| Thnsl1 | 78,88 | -0,0962 | 0,873853413 | chr2 | + |
| Timm10 | 476,71 | -0,2798 | 0,470066395 | chr2 | + |
| Tlk1 | 689,86 | -0,2107 | 0,505258604 | chr2 | - |
| Tm9sf4 | 666,11 | -0,0915 | 0,824129134 | chr2 | + |
| Tmem127 | 925,92 | -0,0864 | 0,82176873 | chr2 | + |
| Tmem141 | 448,64 | 0,1288 | 0,792197952 | chr2 | - |
| Tmem189 | 84,93 | -0,2692 | 0,606429785 | chr2 | - |
| Tmem203 | 87,97 | -0,3634 | 0,428679263 | chr2 | + |
| Tmem230 | 384,43 | 0,1983 | 0,599608164 | chr2 | - |
| Tmem62 | 315,98 | -0,0726 | 0,899463828 | chr2 | + |
| Tmem87a | 221,53 | 0,3093 | 0,357345082 | chr2 | - |
| Tmem87b | 447,35 | 0,1363 | 0,765219833 | chr2 | + |
| Tmx2 | 1516,44 | 0,1175 | 0,763514396 | chr2 | - |
| Tnfaip6 | 4,81 | 0,2046 | 0,811137679 | chr2 | + |
| Tnks1bp1 | 722,47 | 0,1414 | 0,769969083 | chr2 | + |
| Tomm34 | 117,24 | 0,0794 | 0,88127014 | chr2 | - |
| Top1 | 870,54 | -0,0133 | 0,980186108 | chr2 | + |
| Tor1a | 939,11 | -0,3220 | 0,339323135 | chr2 | - |
| Tor1b | 816,11 | -0,2086 | 0,565885654 | chr2 | + |
| Tor2a | 416,24 | -0,4949 | 0,220997862 | chr2 | + |
| Tor4a | 147,01 | 0,1150 | 0,792197952 | chr2 | - |
| Tpd52l2 | 812,37 | 0,0763 | 0,851472884 | chr2 | + |
| Tprn | 41,42 | 0,1629 | 0,797559244 | chr2 | + |
| Tpx2 | 16,61 | 0,5877 | 0,324781401 | chr2 | + |
| Traf1 | 5,65 | 0,0253 | 0,981931319 | chr2 | - |
| Traf2 | 163,40 | -0,2697 | 0,476319975 | chr2 | - |
| Traf6 | 239,07 | -0,1602 | 0,636759527 | chr2 | + |
| Trib3 | 58,45 | -0,8516 | 0,09512379 | chr2 | - |
| Trim44 | 790,25 | 0,0783 | 0,869544763 | chr2 | - |
| Trmt6 | 226,06 | 0,1558 | 0,738449568 | chr2 | - |
| Trp53bp1 | 95,83 | 0,3512 | 0,564849438 | chr2 | - |
| Trp53i11 | 67,72 | -0,1938 | 0,751230284 | chr2 | + |
| Trp53inp2 | 4077,68 | -0,3384 | 0,343412532 | chr2 | + |
| Trp53rkb | 8,36 | 0,2501 | 0,756233857 | chr2 | + |
| Trpc4ap | 1076,64 | -0,2870 | 0,310259492 | chr2 | - |
| Trpm7 | 778,08 | 0,0724 | 0,871232434 | chr2 | - |
| Trub2 | 5,03 | -0,0664 | 0,947612608 | chr2 | - |
| Tsc1 | 490,62 | 0,1481 | 0,698043109 | chr2 | + |
| Tshz2 | 99,81 | 0,3088 | 0,465508094 | chr2 | + |
| Tspan18 | 25,73 | 0,2235 | 0,761512492 | chr2 | - |
| Tspyl3 | 7,77 | 0,3191 | 0,677398032 | chr2 | - |

| | | | | | |
|---|---|---|---|---|---|
| Ttbk2 | 126,96 | -0,0696 | 0,88127014 | chr2 | - |
| Ttbk2 | 441,47 | -0,2812 | 0,303750468 | chr2 | - |
| Ttc17 | 187,28 | -0,1458 | 0,764714786 | chr2 | - |
| Ttc21b | 47,56 | 0,2878 | 0,575998767 | chr2 | - |
| Ttc30a1 | 30,14 | 0,7828 | 0,128504077 | chr2 | - |
| Ttc30a2 | 476,27 | 0,2754 | 0,611914561 | chr2 | - |
| Ttf1 | 162,10 | 0,4860 | 0,112531452 | chr2 | + |
| Tti1 | 197,12 | 0,0043 | 0,994111666 | chr2 | - |
| Ttl | 79,64 | -0,3593 | 0,408496057 | chr2 | + |
| Ttpal | 333,89 | -0,3792 | 0,271418885 | chr2 | + |
| Tubb4b | 1275,97 | -0,5339 | 0,149744539 | chr2 | - |
| Tubgcp4 | 430,44 | -0,1689 | 0,602548857 | chr2 | + |
| Tyro3 | 6,66 | -0,0573 | 0,953911274 | chr2 | + |
| Ube2c | 17,52 | 0,2233 | 0,762940356 | chr2 | + |
| Ube2e3 | 486,90 | -0,2955 | 0,339404199 | chr2 | + |
| Ube2v1 | 971,13 | -0,2465 | 0,536536247 | chr2 | - |
| Ubox5 | 91,42 | -0,0528 | 0,932531696 | chr2 | - |
| Ubr1 | 284,41 | 0,2964 | 0,448865116 | chr2 | - |
| Ubr3 | 1504,96 | -0,2181 | 0,55714632 | chr2 | + |
| Uck1 | 575,40 | 0,3056 | 0,334804026 | chr2 | - |
| Uckl1 | 223,72 | -0,1910 | 0,619039366 | chr2 | - |
| Upf2 | 527,51 | -0,0727 | 0,83097362 | chr2 | + |
| Upp2 | 15919,16 | -0,4194 | 0,375529622 | chr2 | + |
| Urm1 | 200,37 | -0,3651 | 0,273593504 | chr2 | + |
| Usp20 | 78,44 | -0,1302 | 0,797548168 | chr2 | + |
| Usp6nl | 177,70 | 0,2678 | 0,553850707 | chr2 | + |
| Usp8 | 741,56 | 0,0898 | 0,798172714 | chr2 | + |
| Vav2 | 256,98 | 0,0247 | 0,96841267 | chr2 | - |
| Vim | 339,40 | -0,0308 | 0,95990578 | chr2 | + |
| Vps16 | 414,88 | -0,0674 | 0,871700795 | chr2 | + |
| Vps18 | 131,44 | -0,2162 | 0,677867285 | chr2 | + |
| Vps39 | 299,83 | 0,2665 | 0,500489603 | chr2 | - |
| Wdr34 | 96,99 | 0,0658 | 0,89879469 | chr2 | - |
| Wdr5 | 556,99 | -0,0209 | 0,967536738 | chr2 | + |
| Wdr76 | 48,98 | 0,4600 | 0,325415084 | chr2 | + |
| Wdsub1 | 72,14 | 0,2008 | 0,671272065 | chr2 | - |
| Wfdc3 | 9,27 | 0,9263 | 0,10976595 | chr2 | - |
| Wipf1 | 83,62 | 0,1889 | 0,704855641 | chr2 | - |
| Wisp2 | 6,69 | -0,1453 | 0,87328174 | chr2 | + |
| Wt1 | 3,99 | 0,1749 | 0,838278662 | chr2 | + |
| Xrn2 | 642,43 | 0,0597 | 0,907980635 | chr2 | + |
| Yme1l1 | 2828,15 | 0,1875 | 0,647295341 | chr2 | + |
| Ypel4 | 1,75 | 0,4825 | NA | chr2 | + |
| Ythdf1 | 452,66 | -0,2734 | 0,321708301 | chr2 | - |
| Ywhab | 1801,29 | -0,1330 | 0,689465734 | chr2 | + |
| Map3k20 | 213,49 | 0,5565 | 0,113502013 | chr2 | + |
| Zbtb26 | 55,09 | 0,4065 | 0,345153273 | chr2 | - |
| Zbtb34 | 68,19 | 0,1193 | 0,844618872 | chr2 | - |
| Zbtb43 | 361,86 | -0,0825 | 0,866006745 | chr2 | - |
| Zbtb46 | 23,82 | 0,1655 | 0,806932775 | chr2 | - |

| | | | | | |
|---|---|---|---|---|---|
| Zbtb6 | 223,94 | 0,3414 | 0,402107622 | chr2 | - |
| Zc3h15 | 1062,06 | -0,3429 | 0,197263027 | chr2 | + |
| Zc3h6 | 20,66 | 0,3247 | 0,621804913 | chr2 | + |
| Zcchc3 | 14,39 | 0,3603 | 0,613307346 | chr2 | - |
| Zdhhc12 | 141,85 | -0,2769 | 0,620918751 | chr2 | - |
| Zdhhc5 | 954,41 | 0,0119 | 0,98429278 | chr2 | - |
| Zeb2 | 187,20 | 0,4031 | 0,287543614 | chr2 | - |
| Zeb2os | 17,81 | 0,2714 | 0,696351441 | chr2 | + |
| Zer1 | 184,04 | 0,1191 | 0,813323745 | chr2 | - |
| Zfp106 | 1081,23 | -0,4163 | 0,126676113 | chr2 | - |
| Zfp133-ps | 4,57 | 0,3415 | 0,653708291 | chr2 | + |
| Zfp217 | 146,19 | 0,2450 | 0,604679897 | chr2 | - |
| Zfp334 | 31,41 | 0,8360 | 0,055007345 | chr2 | - |
| Zfp335 | 120,59 | -0,5331 | 0,259107808 | chr2 | - |
| Zfp341 | 17,42 | -0,1456 | 0,854357668 | chr2 | + |
| Zfp385b | 234,16 | 0,1788 | 0,658107384 | chr2 | - |
| Zfp408 | 66,85 | -0,2347 | 0,662842062 | chr2 | - |
| Zfp64 | 62,01 | 0,0402 | 0,959634809 | chr2 | - |
| Zfp661 | 68,66 | 0,1274 | 0,822343064 | chr2 | - |
| Zfp831 | 2,18 | 0,3364 | NA | chr2 | + |
| Zfp931 | 40,37 | -0,0982 | 0,890887823 | chr2 | - |
| Zfp937 | 238,79 | 0,3631 | 0,24625125 | chr2 | + |
| Zfyve19 | 223,43 | -0,1887 | 0,670891431 | chr2 | + |
| Zgpat | 100,03 | -0,1887 | 0,712581071 | chr2 | + |
| Zhx3 | 239,20 | 0,1377 | 0,770645004 | chr2 | - |
| Zmynd19 | 99,64 | -0,2885 | 0,553766864 | chr2 | + |
| Zmynd8 | 221,10 | 0,0650 | 0,905274865 | chr2 | - |
| Zfp512b | 269,49 | -0,4435 | 0,167807438 | chr2 | - |
| Znfx1 | 352,36 | 0,0854 | 0,859139694 | chr2 | - |
| Zscan29 | 265,57 | 0,1539 | 0,666391696 | chr2 | - |
| Zswim1 | 32,01 | -0,5144 | 0,397488617 | chr2 | + |
| Zswim3 | 26,09 | -0,3250 | 0,595460103 | chr2 | + |
| Glmp | 927,31 | -0,1998 | 0,598404146 | chr3 | + |
| 0610031O16Rik | 716,28 | 0,1286 | 0,862421004 | chr3 | - |
| 1700010I02Rik | 8,16 | -0,3117 | 0,686790162 | chr3 | - |
| 1700061I17Rik | 27,54 | 0,8944 | 0,115749538 | chr3 | - |
| 1700094D03Rik | 80,32 | -0,4550 | 0,298240526 | chr3 | + |
| 1700113A16Rik | 49,52 | -0,2098 | 0,740019762 | chr3 | - |
| 1810022K09Rik | 1149,61 | -0,1256 | 0,781284538 | chr3 | - |
| 1810037I17Rik | 700,64 | 0,2187 | 0,553833033 | chr3 | + |
| 1810062G17Rik | 10,27 | 0,0758 | 0,937247252 | chr3 | + |
| 2410004B18Rik | 65,26 | -0,3012 | 0,506368378 | chr3 | + |
| Platr10 | 3,73 | 0,0393 | 0,9700021 | chr3 | - |
| 2810403A07Rik | 482,36 | -0,4854 | 0,100572892 | chr3 | + |
| Abhd18 | 299,94 | -0,0856 | 0,869020534 | chr3 | + |
| 4632415L05Rik | 250,78 | 0,1539 | 0,761282414 | chr3 | + |
| Plppr5 | 3,70 | 0,2580 | 0,748519887 | chr3 | + |
| 4930539J05Rik | 4,40 | -0,2562 | 0,754953629 | chr3 | - |
| 4930577N17Rik | 2,12 | 0,0708 | NA | chr3 | + |
| 4930579G24Rik | 29,11 | -0,2731 | 0,721569914 | chr3 | + |

| | | | | | |
|---|---|---|---|---|---|
| 4932438A13Rik | 648,53 | 0,0619 | 0,886446111 | chr3 | + |
| 4933431E20Rik | 45,70 | 0,4082 | 0,428679263 | chr3 | - |
| 4933434E20Rik | 770,00 | -0,0980 | 0,803952241 | chr3 | + |
| 5330417C22Rik | 6,14 | 0,0732 | 0,942425773 | chr3 | - |
| Fam241a | 41,70 | -0,5423 | 0,268917837 | chr3 | - |
| 5830417I10Rik | 80,27 | -0,1055 | 0,846704266 | chr3 | + |
| 6330549D23Rik | 11,42 | -0,0855 | 0,930087232 | chr3 | - |
| A230001M10Rik | 6,81 | 0,8862 | 0,142156924 | chr3 | + |
| A930005H10Rik | 145,07 | -0,2626 | 0,533879942 | chr3 | - |
| Aadac | 24155,27 | -0,1521 | 0,74146296 | chr3 | + |
| Abca4 | 2,35 | 0,2692 | NA | chr3 | + |
| Abcd3 | 12486,36 | -0,2003 | 0,599872501 | chr3 | - |
| Acadm | 6774,97 | 0,0750 | 0,851472884 | chr3 | - |
| Acp6 | 468,24 | 0,0010 | 0,997745329 | chr3 | + |
| Actl6a | 349,52 | -0,2118 | 0,531231457 | chr3 | + |
| Adam15 | 22,50 | 0,2514 | 0,704548549 | chr3 | - |
| Adamtsl4 | 22,02 | -0,2672 | 0,678676127 | chr3 | - |
| Adar | 278,35 | 0,2535 | 0,55880551 | chr3 | + |
| Adh1 | 47456,51 | -0,1699 | 0,621775664 | chr3 | + |
| Adh4 | 2997,21 | 0,2417 | 0,589361744 | chr3 | + |
| Adh5 | 11162,54 | -0,0653 | 0,888763259 | chr3 | + |
| Adh6-ps1 | 693,73 | -0,2824 | 0,587913367 | chr3 | + |
| Adh7 | 112,41 | 0,0389 | 0,952532092 | chr3 | + |
| Agl | 875,54 | -0,2436 | 0,523118038 | chr3 | - |
| Ahcyl1 | 1492,72 | -0,1982 | 0,535573923 | chr3 | - |
| AI504432 | 9,34 | 0,1359 | 0,877237757 | chr3 | + |
| Aimp1 | 775,13 | -0,2430 | 0,495713543 | chr3 | - |
| Alg14 | 333,09 | -0,1217 | 0,748107818 | chr3 | + |
| Alg5 | 925,23 | -0,2679 | 0,407043096 | chr3 | + |
| Alpk1 | 19,95 | 0,6748 | 0,23655468 | chr3 | - |
| Amigo1 | 123,31 | 0,2484 | 0,587879802 | chr3 | + |
| Ampd2 | 298,99 | 0,0090 | 0,990846949 | chr3 | - |
| Ankrd13c | 296,29 | -0,0317 | 0,950561821 | chr3 | + |
| Ankrd35 | 2,64 | 0,4795 | 0,461369428 | chr3 | + |
| Ankrd50 | 74,71 | -0,2706 | 0,529147268 | chr3 | - |
| Ankub1 | 6,03 | 0,4264 | 0,561773364 | chr3 | - |
| Anp32e | 618,57 | 0,3712 | 0,293001163 | chr3 | + |
| Anxa9 | 18,03 | 0,2142 | 0,786227852 | chr3 | - |
| Ap1ar | 338,04 | 0,3797 | 0,208173201 | chr3 | - |
| Ap4b1 | 319,74 | 0,0069 | 0,990189789 | chr3 | + |
| Aph1a | 1529,23 | 0,0210 | 0,962739078 | chr3 | + |
| Naxe | 791,03 | -0,1061 | 0,796541442 | chr3 | - |
| Arfip1 | 487,59 | 0,2237 | 0,569286868 | chr3 | - |
| Arhgap29 | 1113,84 | 0,0211 | 0,970245243 | chr3 | + |
| Arhgef11 | 298,29 | -0,2656 | 0,537660048 | chr3 | + |
| Arhgef2 | 60,68 | 0,2201 | 0,683817601 | chr3 | + |
| Arhgef26 | 693,75 | 0,0250 | 0,975445719 | chr3 | + |
| Armc1 | 850,93 | -0,3383 | 0,240304485 | chr3 | - |
| Arnt | 472,92 | 0,2681 | 0,382591532 | chr3 | + |
| Ash1l | 414,50 | 0,0839 | 0,853848645 | chr3 | + |

| | | | | | |
|---|---|---|---|---|---|
| Asic5 | 125,16 | -0,5590 | 0,119628533 | chr3 | + |
| Atp11b | 870,63 | -0,0659 | 0,876498461 | chr3 | + |
| Atp5f1 | 7743,31 | -0,2142 | 0,590058791 | chr3 | - |
| Atp8b2 | 30,95 | 0,3284 | 0,568858283 | chr3 | - |
| Atxn7l2 | 12,99 | -0,3397 | 0,648869552 | chr3 | - |
| B3galnt1 | 14,58 | 0,3709 | 0,577069333 | chr3 | - |
| Bank1 | 46,53 | 0,3364 | 0,470831079 | chr3 | - |
| Bbs12 | 11,02 | 0,3738 | 0,618723125 | chr3 | + |
| Bbs7 | 77,05 | -0,2397 | 0,629968267 | chr3 | - |
| BC021767 | 36,67 | 0,3310 | 0,623523104 | chr3 | - |
| BC028528 | 39,72 | 0,4148 | 0,419707795 | chr3 | - |
| Bcar3 | 496,02 | -0,1908 | 0,622736166 | chr3 | + |
| Bcas2 | 410,62 | -0,0307 | 0,943725284 | chr3 | + |
| Bcl10 | 199,14 | 0,1964 | 0,683330505 | chr3 | + |
| Bcl2l15 | 2,30 | -0,5663 | NA | chr3 | + |
| Bcl9 | 35,42 | 0,1109 | 0,874763133 | chr3 | - |
| Bdh2 | 553,09 | -0,0574 | 0,925700434 | chr3 | + |
| Bola1 | 176,25 | -0,3580 | 0,63251737 | chr3 | - |
| Platr4 | 16,52 | 0,3856 | 0,571753961 | chr3 | - |
| C920021L13Rik | 4,21 | -0,1185 | 0,896182211 | chr3 | + |
| Camk2d | 111,20 | 0,1192 | 0,815253842 | chr3 | + |
| Capza1 | 1073,32 | 0,2808 | 0,41682625 | chr3 | - |
| Car1 | 171,30 | -0,2999 | 0,581725594 | chr3 | - |
| Car13 | 14,33 | 0,3521 | 0,605755718 | chr3 | + |
| Car2 | 131,61 | -0,2059 | 0,577347444 | chr3 | + |
| Car3 | 53616,93 | -0,6018 | 0,219639131 | chr3 | + |
| Casp6 | 475,69 | 0,1464 | 0,715912422 | chr3 | + |
| Casq2 | 2,23 | 0,6571 | NA | chr3 | + |
| Kyat3 | 6385,85 | -0,3787 | 0,605538599 | chr3 | + |
| Ccdc144b | 9,12 | -0,2547 | 0,756233857 | chr3 | - |
| Ccdc169 | 3,89 | -0,1029 | 0,910916841 | chr3 | + |
| Ccdc39 | 3,41 | 0,5524 | 0,396254007 | chr3 | - |
| Ccna2 | 26,49 | 0,2753 | 0,716578314 | chr3 | - |
| Ccnl1 | 538,31 | -0,0434 | 0,93807111 | chr3 | - |
| Noct | 536,93 | -0,5519 | 0,40679401 | chr3 | + |
| Cct3 | 1578,55 | -0,1501 | 0,724187104 | chr3 | + |
| Cd160 | 5,03 | 0,5908 | 0,376702038 | chr3 | - |
| Cd1d1 | 3196,67 | -0,4280 | 0,188597887 | chr3 | - |
| Cd1d2 | 3,27 | -0,1213 | 0,890887823 | chr3 | + |
| Cd2 | 4,93 | 0,4625 | 0,505514525 | chr3 | - |
| Cd53 | 320,49 | 0,4561 | 0,08440425 | chr3 | - |
| Cd5l | 2744,23 | 0,0937 | 0,829703001 | chr3 | + |
| Cdc14a | 21,23 | 0,3877 | 0,524328995 | chr3 | - |
| Cdc42se1 | 310,70 | 0,4422 | 0,12617458 | chr3 | + |
| Celsr2 | 3,34 | -0,3774 | 0,601786607 | chr3 | - |
| Cenpe | 13,56 | 0,4429 | 0,520016085 | chr3 | + |
| Cept1 | 1930,33 | 0,2291 | 0,583782269 | chr3 | - |
| Cers2 | 5590,48 | -0,1857 | 0,506368378 | chr3 | + |
| Cetn4 | 5,70 | 0,7072 | 0,267397666 | chr3 | - |
| Cfi | 17423,55 | 0,0714 | 0,851472884 | chr3 | + |

| | | | | | |
|---|---|---|---|---|---|
| Chd1l | 166,28 | 0,0487 | 0,93871506 | chr3 | - |
| Chil3 | 3,25 | 0,7182 | 0,221950245 | chr3 | - |
| Chmp4c | 20,92 | 0,7002 | 0,207368678 | chr3 | + |
| Chrnb2 | 14,57 | -0,6766 | 0,269410662 | chr3 | - |
| Chtop | 1178,37 | 0,0249 | 0,95728901 | chr3 | - |
| Ciart | 233,32 | -0,3076 | 0,48416799 | chr3 | - |
| Cisd2 | 583,05 | -0,4634 | 0,079398036 | chr3 | - |
| Cks1b | 113,22 | -0,5487 | 0,205620642 | chr3 | - |
| Clca2 | 5,87 | 0,1724 | 0,846012652 | chr3 | - |
| Clcc1 | 1230,33 | -0,0270 | 0,950391517 | chr3 | + |
| Clk2 | 126,75 | -0,0728 | 0,914623649 | chr3 | + |
| Clrn1 | 4,35 | -0,7096 | 0,261655059 | chr3 | - |
| Cnn3 | 2035,36 | -0,2060 | 0,504369791 | chr3 | + |
| Cog6 | 418,28 | -0,0297 | 0,951574951 | chr3 | - |
| Col24a1 | 4,70 | 0,4134 | 0,575364035 | chr3 | + |
| Commd2 | 358,29 | -0,0366 | 0,939403124 | chr3 | - |
| Cp | 31323,87 | 0,0032 | 0,994106851 | chr3 | + |
| Creb3l4 | 3,19 | 0,3463 | 0,629719515 | chr3 | - |
| Crtc2 | 217,78 | -0,3547 | 0,369534447 | chr3 | + |
| Cryz | 2156,54 | 0,2931 | 0,361776578 | chr3 | + |
| Csde1 | 4802,92 | -0,1002 | 0,784972561 | chr3 | + |
| Csf1 | 119,41 | 0,5383 | 0,1915571 | chr3 | - |
| Ctbs | 237,12 | 0,2254 | 0,521142761 | chr3 | + |
| Cth | 8197,79 | 0,1770 | 0,672205411 | chr3 | - |
| Ctsk | 23,74 | 0,1866 | 0,812943523 | chr3 | + |
| Ctso | 1532,31 | 0,4061 | 0,171022097 | chr3 | + |
| Ctss | 1171,27 | 0,2520 | 0,541556317 | chr3 | + |
| Cttnbp2nl | 71,23 | 0,3750 | 0,38079602 | chr3 | - |
| Cxxc4 | 5,07 | 0,0782 | 0,938514017 | chr3 | + |
| Cyb561d1 | 76,19 | 0,3658 | 0,361776578 | chr3 | - |
| Cym | 2,48 | -0,2376 | NA | chr3 | - |
| Cyp2u1 | 149,30 | -0,4051 | 0,298116176 | chr3 | - |
| Cyp7b1 | 6011,81 | -0,8295 | 0,080590154 | chr3 | - |
| Cyr61 | 73,68 | -0,4148 | 0,326662774 | chr3 | - |
| D3Ertd254e | 114,88 | 0,5325 | 0,110128525 | chr3 | + |
| D3Ertd751e | 192,02 | -0,0469 | 0,939306183 | chr3 | + |
| Tmem131l | 205,94 | -0,3798 | 0,270497584 | chr3 | - |
| Dap3 | 761,70 | 0,1341 | 0,671272065 | chr3 | - |
| Dapp1 | 36,23 | 0,5873 | 0,216462763 | chr3 | - |
| Dbt | 2006,41 | 0,0399 | 0,939054301 | chr3 | + |
| Dclk1 | 3,32 | 0,3666 | 0,60828654 | chr3 | + |
| Dclk2 | 5,47 | 0,0029 | 0,997464504 | chr3 | - |
| Dclre1b | 81,10 | -0,2843 | 0,523381331 | chr3 | - |
| Dcun1d1 | 1374,36 | 0,1207 | 0,822394911 | chr3 | - |
| Ddah1 | 2550,82 | 0,1454 | 0,702177047 | chr3 | + |
| Ddit4l | 12,80 | -0,3202 | 0,668081287 | chr3 | + |
| Ddx20 | 162,81 | -0,1494 | 0,693976837 | chr3 | - |
| Dennd2c | 15,16 | -0,0632 | 0,94647679 | chr3 | + |
| Dennd2d | 181,96 | 0,2785 | 0,57596001 | chr3 | + |
| Dennd4b | 48,47 | 0,1123 | 0,869544763 | chr3 | + |

| | | | | | |
|---|---|---|---|---|---|
| Depdc1a | 8,21 | 0,6347 | 0,330697335 | chr3 | + |
| Dhx36 | 709,70 | -0,0716 | 0,893437979 | chr3 | - |
| Dkk2 | 5,37 | -0,5601 | 0,407836161 | chr3 | + |
| Dnajb14 | 61,50 | -0,4809 | 0,240516297 | chr3 | + |
| Dnajb4 | 932,20 | -0,0118 | 0,986642823 | chr3 | - |
| Dnajc19 | 1006,64 | -0,2522 | 0,698110464 | chr3 | - |
| Dnase2b | 757,65 | -0,3205 | 0,228308438 | chr3 | - |
| Dnttip2 | 591,40 | 0,0018 | 0,996987906 | chr3 | + |
| Dph5 | 139,49 | -0,1392 | 0,793608383 | chr3 | + |
| Dpm3 | 478,23 | -0,2286 | 0,785373633 | chr3 | + |
| Dpyd | 7531,29 | -0,0467 | 0,932525817 | chr3 | + |
| Dram2 | 1999,41 | 0,3295 | 0,335347943 | chr3 | + |
| E2f5 | 118,89 | 0,0184 | 0,976788115 | chr3 | + |
| Ecm1 | 1052,03 | -0,2019 | 0,58928611 | chr3 | - |
| Efna1 | 600,66 | 0,1051 | 0,851988359 | chr3 | - |
| Efna4 | 2,95 | 0,0879 | 0,922904986 | chr3 | - |
| Egf | 8,65 | 0,3946 | 0,590718753 | chr3 | - |
| Egfem1 | 9,49 | -0,2757 | 0,7242192 | chr3 | + |
| Eif2a | 1018,89 | 0,0272 | 0,95728901 | chr3 | + |
| Eif4e | 951,91 | -0,1519 | 0,677975063 | chr3 | + |
| Eif5a2 | 12,30 | 0,1698 | 0,846571896 | chr3 | + |
| Elf2 | 403,66 | 0,2770 | 0,441724922 | chr3 | - |
| Elovl6 | 2510,40 | 0,1710 | 0,777444237 | chr3 | + |
| Adgrl4 | 365,93 | 0,1484 | 0,693803702 | chr3 | + |
| Emcn | 89,52 | 0,2440 | 0,621775664 | chr3 | + |
| Enpep | 1564,74 | 0,0033 | 0,995836778 | chr3 | - |
| Ensa | 851,00 | -0,0853 | 0,813441503 | chr3 | + |
| Etfdh | 5443,57 | 0,0607 | 0,899463828 | chr3 | - |
| Etv3 | 204,87 | 0,3577 | 0,297351163 | chr3 | + |
| Exosc8 | 233,27 | 0,1496 | 0,773900067 | chr3 | - |
| Exosc9 | 343,94 | 0,0935 | 0,867206921 | chr3 | + |
| Extl2 | 176,75 | 0,0520 | 0,909834629 | chr3 | + |
| F3 | 89,09 | 0,4760 | 0,278922122 | chr3 | + |
| Fabp2 | 3459,13 | -0,4481 | 0,219251779 | chr3 | + |
| Fabp4 | 303,42 | -0,0691 | 0,907233333 | chr3 | - |
| Fabp5 | 2563,23 | 0,2058 | 0,682989802 | chr3 | + |
| Fam102b | 50,50 | 0,4034 | 0,389788694 | chr3 | - |
| Fam160a1 | 52,83 | -0,1926 | 0,748107818 | chr3 | - |
| Fam189b | 12,73 | -0,6106 | 0,309392522 | chr3 | + |
| Fam198b | 13,05 | 0,3479 | 0,629919092 | chr3 | + |
| Fam212b | 3,00 | 0,6019 | 0,335528734 | chr3 | + |
| Fam46c | 52,39 | 0,7427 | 0,075466772 | chr3 | - |
| Mindy1 | 1011,54 | -0,2038 | 0,451443396 | chr3 | + |
| Miga1 | 35,54 | 0,1151 | 0,884003681 | chr3 | - |
| Fat4 | 22,68 | 0,2089 | 0,762940356 | chr3 | + |
| Fbxw7 | 160,86 | -0,3114 | 0,381800527 | chr3 | + |
| Fcgr1 | 103,98 | 0,2812 | 0,528512451 | chr3 | - |
| Fcrl1 | 7,25 | -0,0663 | 0,947612608 | chr3 | + |
| Fdps | 3626,02 | -0,5594 | 0,202636202 | chr3 | - |
| Fga | 76667,09 | -0,3545 | 0,380154872 | chr3 | + |

| | | | | | |
|---|---|---|---|---|---|
| Fgb | 125579,23 | -0,2636 | 0,471563545 | chr3 | - |
| Fgg | 89157,86 | -0,2984 | 0,348682374 | chr3 | + |
| Fhdc1 | 5,70 | 0,3413 | 0,649305665 | chr3 | - |
| Flad1 | 657,67 | -0,3037 | 0,428696179 | chr3 | - |
| Fmo5 | 12177,00 | -0,6486 | 0,073430016 | chr3 | + |
| Fnbp1l | 444,39 | 0,1427 | 0,710461911 | chr3 | - |
| Fndc3b | 343,06 | 0,0668 | 0,904806119 | chr3 | - |
| Fndc7 | 2,19 | -0,0746 | NA | chr3 | - |
| Fnip2 | 477,25 | -0,0465 | 0,933486262 | chr3 | - |
| Foxo1 | 586,37 | -0,5287 | 0,070720663 | chr3 | + |
| Fpgt | 329,21 | 0,3118 | 0,423149698 | chr3 | - |
| Frrs1 | 292,54 | -0,0320 | 0,944541717 | chr3 | + |
| Fubp1 | 421,19 | 0,0838 | 0,846571896 | chr3 | + |
| Fxr1 | 1185,21 | -0,0799 | 0,866499572 | chr3 | + |
| Gabpb2 | 482,15 | 0,3967 | 0,13640556 | chr3 | - |
| Gar1 | 108,18 | 0,2513 | 0,549054641 | chr3 | - |
| Gatad2b | 26,38 | 0,9090 | 0,052350418 | chr3 | + |
| Gba | 393,91 | -0,2529 | 0,381025284 | chr3 | + |
| Gbp2b | 44,53 | -0,4015 | 0,576265028 | chr3 | + |
| Gbp5 | 28,15 | 0,6430 | 0,19521384 | chr3 | + |
| Gclm | 1835,72 | -0,4883 | 0,06491165 | chr3 | + |
| Gdap2 | 604,50 | 0,5642 | 0,05304834 | chr3 | + |
| Gfm1 | 1495,09 | 0,4076 | 0,060740616 | chr3 | + |
| Gja5 | 24,63 | 0,7845 | 0,12697428 | chr3 | + |
| Glt28d2 | 111,17 | -0,5004 | 0,165257806 | chr3 | - |
| Gm15417 | 173,65 | -0,4092 | 0,263169902 | chr3 | + |
| Gm15441 | 52,00 | 0,5315 | NA | chr3 | - |
| Gm16740 | 25,14 | -0,4053 | 0,462469544 | chr3 | + |
| Gm19710 | 2,40 | 0,0270 | NA | chr3 | + |
| Gm5150 | 19,22 | 0,2494 | 0,71344087 | chr3 | - |
| Gm5544 | 1,95 | 0,3876 | NA | chr3 | + |
| Gm5547 | 2,04 | 0,1771 | NA | chr3 | + |
| Gm6525 | 5,42 | -0,2894 | 0,717188441 | chr3 | + |
| Gm6602 | 3,35 | -0,3917 | 0,592229341 | chr3 | + |
| Gm8234 | 38,20 | -0,3615 | 0,497721225 | chr3 | + |
| Gmps | 958,80 | -0,3296 | 0,265604797 | chr3 | + |
| Gnai3 | 2516,75 | 0,1337 | 0,727514707 | chr3 | - |
| Gnat2 | 52,82 | 0,5262 | 0,379417606 | chr3 | + |
| Gnb4 | 37,21 | 0,3648 | 0,473368945 | chr3 | - |
| Gng5 | 2204,51 | -0,2935 | 0,549054641 | chr3 | + |
| Golim4 | 1116,65 | 0,3384 | 0,319682721 | chr3 | - |
| Golph3l | 190,54 | 0,4944 | 0,217835326 | chr3 | + |
| Gon4l | 119,65 | 0,1020 | 0,822394911 | chr3 | + |
| Gpatch4 | 198,55 | -0,0419 | 0,948314012 | chr3 | + |
| Gpr160 | 9,06 | 0,3008 | 0,695967514 | chr3 | + |
| Gpr171 | 6,02 | 0,2807 | 0,72678231 | chr3 | - |
| Gpr89 | 551,24 | -0,0963 | 0,836985758 | chr3 | - |
| Gpsm2 | 229,66 | 0,1149 | 0,834675238 | chr3 | - |
| Gstcd | 38,92 | 0,4507 | 0,379104437 | chr3 | - |
| Gstm2 | 1044,01 | -0,0075 | 0,986997724 | chr3 | - |

| | | | | | |
|---|---|---|---|---|---|
| Gstm3 | 883,05 | 0,2801 | 0,511962155 | chr3 | - |
| Gstm4 | 1165,19 | -0,0913 | 0,823949825 | chr3 | - |
| Gstm5 | 387,02 | 0,0958 | 0,806159394 | chr3 | + |
| Gstm6 | 1490,04 | -0,2105 | 0,679262748 | chr3 | - |
| Gstm7 | 1323,36 | 0,2821 | 0,429327627 | chr3 | - |
| Gtf2b | 634,51 | -0,2669 | 0,437620234 | chr3 | + |
| Gucy1a1 | 105,87 | 0,3937 | 0,339424882 | chr3 | - |
| Gucy1b1 | 96,97 | 0,5606 | 0,131674744 | chr3 | - |
| Gyg | 169,31 | -0,3369 | 0,365428542 | chr3 | - |
| H2afz | 1712,84 | -0,2436 | 0,663872588 | chr3 | + |
| Hadh | 4792,23 | -0,0975 | 0,817827493 | chr3 | - |
| Hax1 | 1836,88 | -0,3297 | 0,191502263 | chr3 | - |
| Hcn3 | 4,59 | 0,5720 | 0,391831641 | chr3 | - |
| Hdgf | 1936,45 | -0,2651 | 0,335264505 | chr3 | + |
| Hey1 | 10,05 | 0,4797 | 0,483957097 | chr3 | - |
| Hfe2 | 865,33 | 0,1005 | 0,793250646 | chr3 | + |
| Mfsd14a | 546,72 | 0,1110 | 0,751958297 | chr3 | - |
| Hipk1 | 987,84 | -0,1207 | 0,698557859 | chr3 | - |
| Hist2h2be | 60,40 | 0,5264 | 0,33288309 | chr3 | + |
| Hmgcs2 | 43049,43 | 0,0052 | 0,993369412 | chr3 | + |
| Hnf4g | 7,33 | 0,5918 | 0,369534447 | chr3 | + |
| Hps3 | 150,65 | 0,4978 | 0,211590547 | chr3 | - |
| Hs2st1 | 202,21 | 0,4639 | 0,149712396 | chr3 | - |
| Hsd3b2 | 527,83 | -0,3729 | 0,320305617 | chr3 | - |
| Hsd3b3 | 3658,76 | -0,6299 | 0,065459455 | chr3 | - |
| Hspa4l | 458,49 | 0,4903 | 0,147367695 | chr3 | + |
| I830077J02Rik | 9,39 | 0,2161 | 0,790166895 | chr3 | - |
| Ift80 | 106,89 | 0,1492 | 0,791189278 | chr3 | - |
| Igsf10 | 22,89 | 0,2634 | 0,691615955 | chr3 | - |
| Igsf3 | 15,08 | 0,7184 | 0,216647084 | chr3 | + |
| Il6ra | 314,45 | 0,3416 | 0,314431944 | chr3 | - |
| Ilf2 | 398,72 | 0,4474 | 0,19689523 | chr3 | + |
| Impa1 | 1497,49 | -0,0458 | 0,907307534 | chr3 | - |
| Ints3 | 252,21 | 0,0701 | 0,875636241 | chr3 | - |
| Intu | 70,80 | -0,0079 | 0,990793693 | chr3 | + |
| Iqgap3 | 6,59 | -0,0274 | 0,980186108 | chr3 | + |
| Isg20l2 | 262,42 | -0,2385 | 0,51722851 | chr3 | + |
| Itga10 | 33,38 | -0,3505 | 0,581157554 | chr3 | + |
| Jade1 | 612,59 | 0,0284 | 0,963078727 | chr3 | + |
| Jtb | 894,36 | -0,6039 | 0,114525976 | chr3 | + |
| Kcna2 | 85,30 | 0,4439 | 0,269410662 | chr3 | + |
| Kcnab1 | 11,53 | 0,7419 | 0,228728627 | chr3 | + |
| Kirrel | 21,64 | 0,8777 | 0,074317353 | chr3 | - |
| Kpna4 | 546,75 | 0,1529 | 0,677019009 | chr3 | - |
| Krtcap2 | 1024,24 | -0,2155 | 0,732140841 | chr3 | + |
| Lamtor2 | 1275,23 | -0,1683 | 0,533824486 | chr3 | - |
| Lamtor3 | 259,91 | -0,2636 | 0,39852798 | chr3 | + |
| Lamtor5 | 1055,52 | -0,1537 | 0,680997554 | chr3 | + |
| Larp7 | 183,70 | 0,1730 | 0,695575308 | chr3 | - |
| Lekr1 | 4,59 | 0,4786 | 0,497298938 | chr3 | + |

| | | | | | |
|---|---|---|---|---|---|
| Lenep | 28,10 | -0,4841 | 0,407836161 | chr3 | - |
| Lhfp | 266,19 | 0,4945 | 0,093542539 | chr3 | + |
| Lingo4 | 3,73 | 0,2561 | 0,749603956 | chr3 | + |
| Lix1l | 51,63 | 0,2134 | 0,718823184 | chr3 | + |
| Lmna | 346,91 | -0,2213 | 0,666660307 | chr3 | - |
| Lmo4 | 3,91 | 0,0707 | 0,942982044 | chr3 | - |
| Lor | 4,73 | -0,8425 | 0,156728357 | chr3 | - |
| Lpar3 | 3,52 | 0,5265 | 0,41682625 | chr3 | + |
| Lrat | 674,02 | 0,6112 | 0,204076266 | chr3 | - |
| Lrba | 367,55 | 0,4620 | 0,093542539 | chr3 | + |
| Lrig2 | 118,96 | 0,3792 | 0,280877242 | chr3 | - |
| Lrrc39 | 16,96 | 0,7237 | 0,2044482 | chr3 | + |
| Lrrc40 | 253,90 | 0,3887 | 0,322121314 | chr3 | + |
| Lrrcc1 | 54,20 | 0,4992 | 0,370803111 | chr3 | + |
| Lrriq3 | 38,39 | 0,7315 | 0,11867102 | chr3 | + |
| Lxn | 18,27 | 0,3119 | 0,633486481 | chr3 | - |
| Lysmd1 | 195,80 | 0,2067 | 0,566893799 | chr3 | + |
| Magi3 | 337,65 | -0,2153 | 0,619691963 | chr3 | - |
| Maml3 | 25,48 | 0,4739 | 0,384370645 | chr3 | - |
| Man1a2 | 607,46 | -0,2102 | 0,48822276 | chr3 | - |
| Manba | 306,48 | -0,4097 | 0,219517679 | chr3 | + |
| Map9 | 5,89 | -0,3508 | 0,645384919 | chr3 | + |
| Mbnl1 | 1456,58 | -0,1438 | 0,674305684 | chr3 | + |
| Mccc1 | 1121,11 | -0,0079 | 0,985716259 | chr3 | - |
| Mccc1os | 14,90 | 0,1507 | 0,848794352 | chr3 | + |
| Mcl1 | 2052,28 | -0,0755 | 0,848389544 | chr3 | + |
| Mcoln2 | 9,52 | 0,3132 | 0,676060413 | chr3 | + |
| Mecom | 18,75 | 0,0981 | 0,905147397 | chr3 | - |
| Med12l | 53,64 | 0,0044 | 0,995735297 | chr3 | + |
| Mef2d | 56,92 | -0,0842 | 0,922460349 | chr3 | + |
| Metap1 | 668,83 | -0,2182 | 0,498227709 | chr3 | - |
| Mettl14 | 116,16 | -0,0474 | 0,926353842 | chr3 | - |
| Mex3a | 3,62 | 0,8090 | 0,166993075 | chr3 | + |
| Mfn1 | 992,47 | -0,2645 | 0,364573549 | chr3 | + |
| Mfsd1 | 2154,89 | 0,4369 | 0,113502013 | chr3 | + |
| Mfsd8 | 268,65 | -0,2457 | 0,512541555 | chr3 | - |
| Mlf1 | 6,04 | 0,2540 | 0,748107818 | chr3 | + |
| Mllt11 | 18,10 | 0,0168 | 0,986350715 | chr3 | - |
| Mnd1 | 94,31 | 0,2271 | 0,569869994 | chr3 | - |
| Mov10 | 268,10 | 0,4092 | 0,250526648 | chr3 | - |
| Mrpl24 | 1651,37 | -0,2573 | 0,345662712 | chr3 | + |
| Mrpl47 | 355,77 | -0,4002 | 0,266247219 | chr3 | - |
| Mrpl9 | 964,83 | -0,3179 | 0,308240497 | chr3 | + |
| Mrps21 | 1246,66 | -0,3564 | 0,284218549 | chr3 | - |
| Mrps28 | 414,83 | -0,3914 | 0,486005862 | chr3 | - |
| Msto1 | 103,09 | -0,0979 | 0,862578493 | chr3 | - |
| Mtfr1 | 1358,50 | -0,1020 | 0,804048972 | chr3 | + |
| Mttp | 5907,16 | 0,1026 | 0,764714786 | chr3 | - |
| Mtx1 | 540,82 | -0,2226 | 0,505223498 | chr3 | - |
| Muc1 | 11,87 | -0,0897 | 0,922594928 | chr3 | + |

| | | | | | |
|---|---|---|---|---|---|
| Mynn | 183,68 | 0,4192 | 0,215046281 | chr3 | + |
| Naa15 | 655,83 | -0,0784 | 0,858858569 | chr3 | + |
| Nbea | 14,98 | 0,0514 | 0,955234619 | chr3 | - |
| Nceh1 | 849,94 | 0,1541 | 0,773998967 | chr3 | + |
| Ndufb5 | 2567,65 | -0,5353 | 0,210515973 | chr3 | + |
| Ndufc1 | 1149,98 | -0,2827 | 0,564495903 | chr3 | - |
| Nes | 64,85 | 0,6773 | 0,114525976 | chr3 | + |
| Nexn | 20,69 | 0,7918 | 0,137239261 | chr3 | - |
| Nfkb1 | 264,04 | 0,2031 | 0,558489002 | chr3 | - |
| Ngf | 38,24 | 0,2462 | 0,687523396 | chr3 | + |
| Nhlh2 | 5,08 | -0,0799 | 0,93731052 | chr3 | + |
| Nhlrc3 | 84,12 | 0,3297 | 0,5252592 | chr3 | - |
| Nmd3 | 625,24 | -0,1912 | 0,6180541 | chr3 | + |
| Npnt | 3,24 | -0,5049 | 0,442165263 | chr3 | - |
| Npr1 | 121,43 | -0,1561 | 0,742230743 | chr3 | - |
| Nr1h5 | 43,86 | 0,2116 | 0,756242359 | chr3 | - |
| Nras | 588,15 | -0,0056 | 0,991184644 | chr3 | + |
| Ntrk1 | 4,23 | -0,3043 | 0,690862397 | chr3 | - |
| Nudt17 | 2,73 | 0,1125 | 0,895513146 | chr3 | - |
| Nudt6 | 99,88 | -0,5171 | 0,19521384 | chr3 | - |
| Oaz3 | 4,91 | 0,5673 | 0,368588487 | chr3 | - |
| Odf2l | 137,65 | 0,1006 | 0,877283616 | chr3 | + |
| Olfm3 | 31,53 | -0,8402 | 0,077260065 | chr3 | + |
| Olfml3 | 60,94 | 0,4539 | 0,310906399 | chr3 | - |
| Ostc | 1714,60 | 0,1652 | 0,661707091 | chr3 | - |
| Otud7b | 555,19 | 0,3597 | 0,271016733 | chr3 | + |
| Ovgp1 | 7,49 | -0,7168 | 0,254846988 | chr3 | + |
| P2ry12 | 50,49 | 0,2268 | 0,647234508 | chr3 | - |
| P2ry14 | 40,82 | 0,5909 | 0,303046806 | chr3 | - |
| Pabpc4l | 12,43 | 0,6893 | 0,251516758 | chr3 | - |
| Pag1 | 102,31 | 0,2469 | 0,607310966 | chr3 | - |
| Papss1 | 206,31 | 0,1930 | 0,665052079 | chr3 | + |
| Pcdh18 | 14,07 | 0,1545 | 0,86019237 | chr3 | - |
| Pdcd10 | 748,16 | 0,0091 | 0,987426448 | chr3 | - |
| Pde4dip | 893,22 | -0,1320 | 0,744081988 | chr3 | - |
| Pde5a | 29,27 | -0,0761 | 0,919961951 | chr3 | + |
| Pde7a | 83,19 | -0,1593 | 0,759695008 | chr3 | - |
| Pdgfc | 36,64 | -0,6128 | 0,168336507 | chr3 | + |
| Pdlim5 | 885,36 | 0,2939 | 0,263412599 | chr3 | - |
| Pdzk1 | 1289,96 | -0,0706 | 0,872476982 | chr3 | + |
| Pear1 | 63,92 | -0,1042 | 0,873391707 | chr3 | - |
| Gatb | 603,33 | -0,0152 | 0,971659063 | chr3 | + |
| Pex11b | 646,91 | 0,0501 | 0,912010517 | chr3 | + |
| Pex2 | 1121,09 | -0,0410 | 0,931322892 | chr3 | - |
| Pfn2 | 76,57 | 0,2552 | 0,678676127 | chr3 | - |
| Phgdh | 122,04 | 0,6494 | 0,058833993 | chr3 | - |
| Pgrmc2 | 842,04 | -0,1748 | 0,565966999 | chr3 | - |
| Phc3 | 304,85 | 0,2339 | 0,499976559 | chr3 | - |
| Phgdh | 7,18 | -0,1262 | 0,89040553 | chr3 | - |
| Phtf1 | 95,78 | 0,2004 | 0,656913078 | chr3 | + |

| | | | | | |
|---|---|---|---|---|---|
| Phtf1os | 6,46 | -0,1883 | 0,828012328 | chr3 | - |
| Pi4kb | 414,09 | -0,3345 | 0,287543614 | chr3 | + |
| Pias3 | 77,98 | -0,3812 | 0,432040063 | chr3 | + |
| Pigk | 735,67 | -0,0102 | 0,983544118 | chr3 | + |
| Pik3ca | 405,15 | 0,2999 | 0,343267632 | chr3 | + |
| Pip5k1a | 7028,79 | 0,1887 | 0,596014309 | chr3 | - |
| Pkia | 38,39 | 0,5288 | 0,295382635 | chr3 | + |
| Pklr | 4207,72 | 0,6727 | 0,104289738 | chr3 | + |
| Pkn2 | 757,94 | -0,0481 | 0,909834629 | chr3 | - |
| Pld1 | 93,04 | 0,0450 | 0,954161729 | chr3 | + |
| Plekho1 | 65,22 | -0,0661 | 0,92446794 | chr3 | - |
| Plk4 | 16,72 | 0,5700 | 0,336399308 | chr3 | + |
| Pmf1 | 187,70 | 0,1965 | 0,75616611 | chr3 | - |
| Pmvk | 792,45 | 0,3763 | 0,411117904 | chr3 | + |
| Pogz | 174,33 | 0,1451 | 0,716547565 | chr3 | + |
| Polr3c | 269,27 | -0,2510 | 0,54877155 | chr3 | - |
| Polr3gl | 137,01 | 0,4718 | 0,14613636 | chr3 | - |
| Postn | 304,84 | 0,6289 | 0,092854537 | chr3 | + |
| Ppa2 | 769,76 | -0,2287 | 0,443796273 | chr3 | + |
| Ppid | 681,19 | -0,0236 | 0,95645777 | chr3 | + |
| Ppm1j | 4,59 | -0,1550 | 0,862616891 | chr3 | + |
| Ppm1l | 72,65 | 0,1660 | 0,755785289 | chr3 | + |
| Ppp3ca | 305,03 | -0,0631 | 0,893706091 | chr3 | + |
| Prcc | 232,54 | -0,4873 | 0,100588364 | chr3 | - |
| Prkab2 | 30,27 | 0,0330 | 0,969414042 | chr3 | + |
| Prkacb | 644,99 | -0,0326 | 0,93645447 | chr3 | - |
| Prkci | 189,57 | 0,0468 | 0,922935681 | chr3 | + |
| Prmt6 | 86,94 | 0,1836 | 0,688199973 | chr3 | - |
| Prok1 | 12,81 | 0,1657 | 0,847964947 | chr3 | - |
| Proser1 | 78,47 | -0,1716 | 0,751230284 | chr3 | + |
| Prpf3 | 158,42 | -0,1773 | 0,666790135 | chr3 | - |
| Prpf38b | 537,97 | 0,1429 | 0,682099278 | chr3 | - |
| Prune1 | 276,67 | 0,1841 | 0,695681654 | chr3 | - |
| Psma5 | 2275,98 | -0,2029 | 0,545352656 | chr3 | + |
| Psmb4 | 2647,00 | -0,3120 | 0,438244894 | chr3 | - |
| Psmd4 | 2028,58 | -0,2792 | 0,349391987 | chr3 | - |
| Ptbp2 | 194,71 | -0,1450 | 0,738680713 | chr3 | - |
| Ptgfr | 3,98 | -0,0759 | 0,939054301 | chr3 | - |
| Ptgfrn | 60,33 | 0,3259 | 0,478251242 | chr3 | - |
| Ptpn22 | 15,78 | 0,7200 | 0,231959779 | chr3 | + |
| Pygo2 | 185,78 | -0,3877 | 0,301862555 | chr3 | + |
| Rab13 | 97,07 | -0,0457 | 0,938499724 | chr3 | + |
| Rab25 | 3,80 | 0,5718 | 0,370567139 | chr3 | - |
| Rab33b | 158,81 | -0,1394 | 0,773303025 | chr3 | + |
| Rabggtb | 1020,30 | -0,2843 | 0,396411176 | chr3 | - |
| Rap1a | 1248,13 | -0,2389 | 0,501951423 | chr3 | - |
| Rap1gds1 | 341,56 | -0,3854 | 0,166731951 | chr3 | - |
| Rap2b | 68,56 | 0,3879 | 0,37740141 | chr3 | + |
| Rapgef2 | 236,11 | 0,1124 | 0,822343064 | chr3 | - |
| Rbm15 | 121,77 | -0,4791 | 0,192450485 | chr3 | - |

| | | | | | |
|---|---|---|---|---|---|
| Rbm46 | 5,24 | 0,4649 | 0,51178833 | chr3 | - |
| Rbm46os | 2,16 | 0,0570 | NA | chr3 | + |
| Rbm8a | 733,01 | -0,2282 | 0,528512451 | chr3 | + |
| Reg4 | 3,41 | -0,2042 | 0,804418611 | chr3 | + |
| Rfxap | 102,18 | -0,4742 | 0,223515497 | chr3 | - |
| Rhbg | 182,13 | 0,1634 | 0,726344964 | chr3 | - |
| Rit1 | 248,81 | 0,1391 | 0,708319716 | chr3 | + |
| Rnf115 | 683,58 | -0,2471 | 0,413151627 | chr3 | + |
| Rnf13 | 3862,50 | 0,0350 | 0,948314012 | chr3 | + |
| Rnpc3 | 224,61 | -0,1140 | 0,816025345 | chr3 | - |
| Rorc | 772,67 | 0,0818 | 0,8681434 | chr3 | + |
| Rpf1 | 376,86 | -0,3221 | 0,26304439 | chr3 | - |
| Rpl22l1 | 1754,79 | -0,1152 | 0,790000542 | chr3 | + |
| Rpl34 | 7,72 | -0,1192 | 0,890325105 | chr3 | - |
| Rprd2 | 209,92 | -0,1344 | 0,730063556 | chr3 | - |
| Rps27 | 10,51 | -0,3699 | 0,621950561 | chr3 | - |
| Rps3a1 | 12625,06 | -0,4025 | 0,184634167 | chr3 | - |
| Rrnad1 | 92,37 | -0,4225 | 0,310761324 | chr3 | - |
| Rsbn1 | 134,58 | 0,2929 | 0,463230756 | chr3 | + |
| Rsrc1 | 288,70 | 0,0980 | 0,825895013 | chr3 | + |
| Rtca | 179,83 | -0,4605 | 0,138198087 | chr3 | - |
| Rusc1 | 73,25 | -0,0041 | 0,995663291 | chr3 | - |
| Rwdd3 | 73,49 | 0,2898 | 0,590082603 | chr3 | - |
| S100a1 | 2,91 | 0,5342 | 0,397488617 | chr3 | - |
| S100a13 | 1034,12 | 0,1046 | 0,816025345 | chr3 | + |
| S100a16 | 553,22 | 0,0497 | 0,912010517 | chr3 | + |
| S100a6 | 75,48 | 0,3814 | 0,47680357 | chr3 | + |
| S100a8 | 73,77 | 0,7900 | 0,177357355 | chr3 | + |
| S100a9 | 107,75 | 0,3824 | 0,606237187 | chr3 | - |
| S1pr1 | 511,33 | -0,2716 | 0,483930201 | chr3 | - |
| Sars | 755,91 | -0,4442 | 0,085593171 | chr3 | - |
| Sass6 | 30,64 | 0,6409 | 0,211374848 | chr3 | + |
| Scamp3 | 486,80 | -0,4558 | 0,265254075 | chr3 | + |
| Sclt1 | 95,93 | -0,5171 | 0,244197385 | chr3 | - |
| Scnm1 | 189,94 | -0,1972 | 0,640716535 | chr3 | - |
| Sec22b | 773,03 | -0,0947 | 0,840953217 | chr3 | + |
| Sec24b | 211,77 | -0,2493 | 0,535475471 | chr3 | - |
| Sec24d | 743,91 | -0,1510 | 0,732982182 | chr3 | + |
| Sec62 | 3668,62 | -0,2793 | 0,313774527 | chr3 | + |
| Selenbp1 | 3121,82 | 0,2834 | 0,446691936 | chr3 | + |
| Selenbp2 | 14463,33 | 0,5300 | 0,315897953 | chr3 | + |
| Selenot | 4026,56 | -0,0035 | 0,994676823 | chr3 | + |
| Sema6c | 7,01 | -0,0380 | 0,971699327 | chr3 | + |
| Serp1 | 6488,22 | -0,2140 | 0,628871721 | chr3 | - |
| Setd7 | 133,20 | 0,4424 | 0,228640673 | chr3 | - |
| Setdb1 | 345,73 | 0,1289 | 0,753325269 | chr3 | - |
| Sf3b4 | 133,05 | -0,2481 | 0,660583784 | chr3 | + |
| Sgms2 | 902,20 | 0,0864 | 0,871700795 | chr3 | - |
| Sh2d2a | 101,41 | 0,4987 | 0,171795261 | chr3 | + |
| Sh3glb1 | 1061,44 | -0,2179 | 0,456905085 | chr3 | - |

| | | | | | |
|---|---|---|---|---|---|
| Shc1 | 898,90 | 0,2630 | 0,433179416 | chr3 | + |
| She | 75,90 | 0,3467 | 0,394162656 | chr3 | + |
| Siah2 | 246,24 | -0,4119 | 0,195952045 | chr3 | - |
| Sike1 | 103,49 | 0,1814 | 0,688733355 | chr3 | + |
| Sirpb1a | 4,77 | 0,1618 | 0,85281531 | chr3 | - |
| Sirpb1b | 3,72 | 0,3454 | 0,638888255 | chr3 | - |
| Skil | 213,55 | 0,0317 | 0,96345382 | chr3 | + |
| Slc16a1 | 1135,56 | 0,3255 | 0,413323375 | chr3 | + |
| Slc22a15 | 68,54 | -0,0281 | 0,969414042 | chr3 | - |
| Slc25a24 | 33,93 | 0,7094 | 0,118025078 | chr3 | + |
| Slc25a44 | 926,73 | 0,2669 | 0,425926605 | chr3 | - |
| Slc27a3 | 6,67 | -0,1263 | 0,890970518 | chr3 | - |
| Slc30a7 | 490,46 | 0,1124 | 0,773303025 | chr3 | - |
| Slc33a1 | 4490,88 | -0,1922 | 0,592428959 | chr3 | - |
| Slc35a3 | 876,96 | 0,3096 | 0,41633338 | chr3 | - |
| Slc39a1 | 1484,58 | -0,0171 | 0,976116181 | chr3 | + |
| Slc39a8 | 524,32 | 0,2949 | 0,356487252 | chr3 | + |
| Slc44a3 | 123,23 | 0,0798 | 0,869544763 | chr3 | - |
| Slc50a1 | 5,48 | 0,8195 | 0,17817538 | chr3 | - |
| Slc7a11 | 2,07 | 0,2834 | NA | chr3 | - |
| Slc7a14 | 47,21 | -0,4312 | 0,339404199 | chr3 | - |
| Slc9b1 | 2,26 | 0,6649 | NA | chr3 | + |
| Slc9b2 | 6,03 | 0,1775 | 0,840322097 | chr3 | + |
| Smad9 | 166,70 | -0,5483 | 0,242907451 | chr3 | + |
| Smc4 | 438,30 | -0,0435 | 0,913017292 | chr3 | + |
| Smg5 | 365,89 | -0,5786 | 0,132079034 | chr3 | + |
| Snapin | 283,26 | -0,0621 | 0,899463828 | chr3 | - |
| Snhg8 | 364,35 | -0,4906 | 0,23475207 | chr3 | - |
| Snx16 | 99,61 | -0,1818 | 0,746827664 | chr3 | - |
| Snx27 | 398,00 | -0,1012 | 0,793608383 | chr3 | - |
| Snx7 | 285,53 | -0,2175 | 0,52825088 | chr3 | - |
| Sort1 | 340,68 | 0,0787 | 0,875636241 | chr3 | + |
| Spata1 | 16,83 | 0,0913 | 0,913936879 | chr3 | - |
| Spata5 | 86,08 | 0,0574 | 0,940390384 | chr3 | + |
| Spg20 | 494,06 | 0,4372 | 0,154183797 | chr3 | + |
| Spry1 | 33,21 | 0,0441 | 0,953751362 | chr3 | + |
| Srsf11 | 653,89 | -0,2188 | 0,525233119 | chr3 | - |
| Ssr2 | 1905,85 | 0,1154 | 0,766619151 | chr3 | + |
| Ssr3 | 7050,16 | 0,2088 | 0,589361744 | chr3 | - |
| Ssx2ip | 127,83 | -0,1348 | 0,756242359 | chr3 | + |
| St6galnac3 | 24,31 | 0,8376 | 0,102947816 | chr3 | - |
| St7l | 457,89 | -0,3031 | 0,278905276 | chr3 | + |
| Stpg2 | 14,71 | -0,0698 | 0,943108135 | chr3 | + |
| Strip1 | 171,16 | -0,1574 | 0,762940356 | chr3 | - |
| Stxbp3 | 559,33 | 0,3406 | 0,204417741 | chr3 | - |
| Syt11 | 64,66 | 0,4276 | 0,395294986 | chr3 | - |
| Taf13 | 482,73 | -0,4791 | 0,083560273 | chr3 | + |
| Tars2 | 265,02 | -0,1857 | 0,659265465 | chr3 | - |
| Tbck | 40,36 | 0,6496 | 0,151287224 | chr3 | + |
| Tbl1xr1 | 657,43 | 0,3652 | 0,268487435 | chr3 | + |

| | | | | | |
|---|---|---|---|---|---|
| Tchh | 2,79 | -0,5079 | 0,432005857 | chr3 | + |
| Tdo2 | 41327,97 | -0,1367 | 0,724652712 | chr3 | - |
| Tdrkh | 73,69 | 0,2880 | 0,559408814 | chr3 | + |
| Tet2 | 117,24 | -0,0393 | 0,94694207 | chr3 | - |
| Thbs3 | 7,35 | 0,0161 | 0,988414083 | chr3 | + |
| Them4 | 698,41 | -0,2044 | 0,621241714 | chr3 | + |
| Them5 | 3,44 | 0,2371 | 0,77097059 | chr3 | + |
| Tifa | 282,71 | 0,0682 | 0,908013895 | chr3 | + |
| Tiparp | 155,10 | 0,2673 | 0,504278019 | chr3 | + |
| Tlr2 | 70,34 | 0,5078 | 0,217231457 | chr3 | - |
| Tm4sf1 | 369,37 | 0,4316 | 0,202727769 | chr3 | - |
| Tm4sf4 | 2192,93 | 0,1264 | 0,737668661 | chr3 | + |
| Tmem144 | 70,14 | 0,6161 | 0,133406187 | chr3 | - |
| Tmem154 | 3,06 | 0,0955 | 0,918032423 | chr3 | + |
| Tmem167b | 663,24 | 0,0437 | 0,931322892 | chr3 | - |
| Tmem56 | 4175,43 | 0,0374 | 0,952576171 | chr3 | - |
| Tmem79 | 21,74 | -0,5539 | 0,321896095 | chr3 | - |
| Tmod4 | 2,94 | -0,0742 | 0,939221429 | chr3 | + |
| Tnfaip8l2 | 42,76 | 0,2754 | 0,583782269 | chr3 | - |
| Tnik | 19,17 | -0,8211 | 0,146048768 | chr3 | + |
| Tpd52 | 286,74 | 0,0021 | 0,996644671 | chr3 | - |
| Tpm3 | 997,32 | 0,2132 | 0,526620417 | chr3 | + |
| Trim2 | 336,14 | 0,3392 | 0,350368583 | chr3 | - |
| Trim33 | 337,79 | -0,1860 | 0,588219472 | chr3 | + |
| Trim45 | 14,95 | 0,4690 | 0,479598353 | chr3 | + |
| Trim59 | 13,50 | 0,0615 | 0,947294154 | chr3 | - |
| Trmt10a | 95,19 | 0,1265 | 0,794116376 | chr3 | + |
| Trmt13 | 62,56 | -0,0874 | 0,886130155 | chr3 | - |
| Trpc3 | 7,25 | -0,5447 | 0,423089018 | chr3 | - |
| Tspan2 | 46,02 | 0,2529 | 0,648706509 | chr3 | + |
| Tspan5 | 11,88 | 0,7831 | 0,186118652 | chr3 | + |
| Ttc14 | 356,12 | -0,2535 | 0,408538653 | chr3 | + |
| Ttf2 | 56,31 | 0,1041 | 0,863264905 | chr3 | - |
| Ttll7 | 24,03 | 0,6076 | 0,228941765 | chr3 | + |
| Txnip | 2523,24 | 0,1649 | 0,809356379 | chr3 | + |
| Tyw3 | 29,15 | -0,1571 | 0,810500447 | chr3 | - |
| Ubap2l | 423,83 | 0,0445 | 0,922460349 | chr3 | - |
| Ube2d3 | 4037,26 | -0,1167 | 0,776478455 | chr3 | + |
| Ube2q1 | 516,77 | -0,2601 | 0,365343011 | chr3 | + |
| Ubqln4 | 433,04 | -0,3559 | 0,303046806 | chr3 | + |
| Ufm1 | 1083,62 | -0,2934 | 0,345819394 | chr3 | - |
| Usp33 | 603,08 | -0,0014 | 0,997464504 | chr3 | + |
| Usp53 | 87,81 | 0,3070 | 0,463150497 | chr3 | - |
| Vangl1 | 76,35 | 0,1683 | 0,77333268 | chr3 | - |
| Vav3 | 23,25 | 0,0862 | 0,910351158 | chr3 | + |
| Vps45 | 165,15 | 0,0449 | 0,941757244 | chr3 | - |
| Vps72 | 133,17 | -0,1847 | 0,766619151 | chr3 | + |
| Wars2 | 89,00 | 0,0173 | 0,979212524 | chr3 | + |
| Wdr3 | 348,35 | 0,2401 | 0,524801822 | chr3 | - |
| Wdr47 | 22,11 | 0,2376 | 0,715470876 | chr3 | + |

| | | | | | |
|---|---|---|---|---|---|
| Wdr77 | 376,77 | -0,4413 | 0,125513994 | chr3 | + |
| Wls | 118,54 | 0,3294 | 0,394187455 | chr3 | + |
| Wwtr1 | 331,13 | 0,0376 | 0,942982044 | chr3 | - |
| Ythdf3 | 1134,50 | 0,1121 | 0,78398139 | chr3 | + |
| Zbtb10 | 84,08 | -0,2470 | 0,611887546 | chr3 | + |
| Zbtb7b | 28,18 | 0,1785 | 0,803663745 | chr3 | - |
| Zc2hc1a | 26,22 | 0,5982 | 0,250819066 | chr3 | + |
| Zfand1 | 436,92 | 0,2355 | 0,624956201 | chr3 | - |
| Zfhx4 | 153,69 | 0,2480 | 0,583782269 | chr3 | + |
| Zfp639 | 323,57 | -0,1019 | 0,797491699 | chr3 | + |
| Zfp687 | 121,49 | 0,0646 | 0,916852736 | chr3 | - |
| Zfp697 | 8,53 | -0,0329 | 0,976528475 | chr3 | + |
| Zfp704 | 132,57 | 0,6805 | 0,072955342 | chr3 | - |
| Zgrf1 | 10,56 | 0,7682 | 0,201620554 | chr3 | + |
| Zmat3 | 88,67 | -0,5564 | 0,169933972 | chr3 | - |
| Znhit6 | 229,32 | 0,4268 | 0,162509787 | chr3 | + |
| Zranb2 | 824,62 | 0,0724 | 0,87206597 | chr3 | + |
| Zzz3 | 471,44 | 0,0766 | 0,8432845 | chr3 | + |
| 0610037L13Rik | 517,34 | -0,2668 | 0,357779158 | chr4 | + |
| 0610043K17Rik | 33,67 | 0,0007 | 0,998994678 | chr4 | + |
| Virma | 328,64 | 0,4201 | 0,165182527 | chr4 | + |
| Stra6l | 2154,81 | 0,3486 | 0,186118652 | chr4 | + |
| 1700003M07Rik | 2,25 | -0,2570 | NA | chr4 | + |
| Cfap74 | 29,81 | 0,1253 | 0,862451616 | chr4 | + |
| 2510039O18Rik | 261,09 | -0,4352 | 0,309392522 | chr4 | + |
| Faap20 | 123,29 | 0,0691 | 0,917575068 | chr4 | + |
| 2610301B20Rik | 105,11 | 0,2259 | 0,69006728 | chr4 | + |
| 2610528J11Rik | 123,63 | 0,6600 | 0,075163457 | chr4 | + |
| 3110021N24Rik | 2,14 | 0,1396 | NA | chr4 | + |
| 3110043O21Rik | 317,62 | -0,4288 | 0,273018284 | chr4 | - |
| 4930412C18Rik | 90,98 | 0,5661 | 0,283971503 | chr4 | + |
| 4930528A17Rik | 2,88 | 0,1743 | 0,83372971 | chr4 | - |
| 4933421O10Rik | 60,16 | -0,0580 | 0,932531696 | chr4 | - |
| Spaar | 7,48 | 0,1200 | 0,897803386 | chr4 | + |
| 5730409E04Rik | 121,09 | 0,3132 | 0,476764512 | chr4 | + |
| Trmo | 49,88 | -0,1714 | 0,763198893 | chr4 | - |
| 5830444B04Rik | 31,77 | 0,6014 | 0,27072986 | chr4 | - |
| 5930403L14Rik | 3,29 | 0,0265 | 0,980186108 | chr4 | - |
| Tmem268 | 121,06 | -0,2581 | 0,634623679 | chr4 | + |
| 9430015G10Rik | 66,87 | -0,4801 | 0,268487435 | chr4 | + |
| 9530080O11Rik | 8,19 | -0,5732 | 0,38949535 | chr4 | - |
| 9930104L06Rik | 75,65 | 0,1590 | 0,757695155 | chr4 | + |
| A430005L14Rik | 555,94 | 0,0177 | 0,971611735 | chr4 | + |
| A530072M11Rik | 13,36 | -0,3772 | 0,581893426 | chr4 | + |
| Frg2f1 | 44,07 | -0,5001 | 0,29271782 | chr4 | - |
| Abca1 | 2305,20 | 0,1022 | 0,844249702 | chr4 | - |
| Acap3 | 36,48 | -0,4821 | 0,400344953 | chr4 | + |
| Acer2 | 351,77 | 0,3257 | 0,370757182 | chr4 | + |
| Acnat1 | 2130,86 | -0,2317 | 0,396411176 | chr4 | - |
| Aco1 | 3964,87 | 0,0630 | 0,883234655 | chr4 | + |

| | | | | | |
|---|---|---|---|---|---|
| Acot11 | 127,99 | 0,6970 | 0,085465646 | chr4 | - |
| Acot7 | 209,35 | -0,2789 | 0,394599707 | chr4 | + |
| Adamtsl1 | 8,80 | -0,1526 | 0,866619594 | chr4 | + |
| Azin2 | 20,39 | 0,6794 | 0,219049665 | chr4 | - |
| Adprhl2 | 113,03 | -0,4458 | 0,322192824 | chr4 | - |
| Agmat | 1827,87 | -0,4335 | 0,162509787 | chr4 | + |
| Ago1 | 347,87 | -0,1527 | 0,620439369 | chr4 | - |
| Ago3 | 31,68 | -0,2968 | 0,613297625 | chr4 | - |
| Ago4 | 70,73 | 0,0511 | 0,938244876 | chr4 | - |
| Agrn | 314,93 | -0,3428 | 0,390708488 | chr4 | - |
| Agtrap | 87,68 | 0,5583 | 0,124946133 | chr4 | - |
| Ahdc1 | 77,11 | -0,1197 | 0,88569455 | chr4 | + |
| AI464131 | 410,59 | -0,3176 | 0,31487946 | chr4 | - |
| Ak2 | 6154,27 | -0,2008 | 0,593190326 | chr4 | + |
| Ak4 | 2079,70 | -0,6084 | 0,083409017 | chr4 | + |
| Akap2 | 204,28 | 0,5848 | 0,067696785 | chr4 | + |
| Akirin1 | 360,19 | -0,2002 | 0,626127531 | chr4 | - |
| Akirin2 | 291,07 | -0,3992 | 0,200566259 | chr4 | + |
| Akna | 21,16 | 0,1083 | 0,886446111 | chr4 | - |
| Akr7a5 | 899,42 | -0,6469 | 0,06817624 | chr4 | + |
| Alad | 1672,87 | -0,0146 | 0,981931319 | chr4 | - |
| Aldh4a1 | 5180,07 | 0,1336 | 0,73417169 | chr4 | + |
| Aldob | 102792,05 | -0,1942 | 0,536431087 | chr4 | - |
| Alg2 | 160,66 | -0,1379 | 0,765219833 | chr4 | - |
| Alg6 | 216,36 | -0,0339 | 0,953911274 | chr4 | + |
| Alpl | 200,56 | 0,3919 | 0,306720802 | chr4 | - |
| Ambp | 43298,88 | -0,1623 | 0,637633057 | chr4 | - |
| Angptl3 | 17937,08 | 0,0273 | 0,969048579 | chr4 | + |
| Angptl7 | 6,69 | 0,0856 | 0,931322892 | chr4 | - |
| Ankrd6 | 7,34 | 0,6947 | 0,277170412 | chr4 | - |
| Anp32b | 757,18 | -0,1812 | 0,627447666 | chr4 | + |
| Cenps | 4,45 | 0,0711 | 0,943346405 | chr4 | - |
| Aptx | 259,29 | -0,0662 | 0,892875227 | chr4 | - |
| Arhgef10l | 206,37 | -0,1542 | 0,821346657 | chr4 | - |
| Arhgef19 | 254,41 | 0,0615 | 0,925898178 | chr4 | + |
| Arhgef39 | 5,49 | 0,1570 | 0,862451616 | chr4 | - |
| Arid1a | 312,10 | -0,2083 | 0,546882594 | chr4 | - |
| Asap3 | 181,86 | -0,1303 | 0,797548168 | chr4 | + |
| Asph | 696,45 | 0,1899 | 0,590317029 | chr4 | - |
| Astn2 | 24,41 | 0,9476 | 0,073599483 | chr4 | - |
| Atad3a | 486,47 | -0,5303 | 0,064726648 | chr4 | - |
| Atad3aos | 15,86 | 0,1145 | 0,896013441 | chr4 | + |
| Atg4c | 217,71 | 0,3081 | 0,371393306 | chr4 | + |
| Atp13a2 | 152,57 | -0,7260 | 0,139399836 | chr4 | + |
| Atp6v0b | 1315,79 | -0,4499 | 0,295255714 | chr4 | - |
| Atp6v1g1 | 1361,10 | -0,3285 | 0,413454178 | chr4 | + |
| Atpaf1 | 252,34 | 0,2953 | 0,425926605 | chr4 | + |
| Atpif1 | 591,76 | -0,0625 | 0,907233333 | chr4 | - |
| AU022252 | 366,90 | 0,1689 | 0,732146735 | chr4 | - |
| AU040320 | 303,52 | -0,0976 | 0,830451573 | chr4 | + |

| | | | | | |
|---|---|---|---|---|---|
| Aurkaip1 | 1132,42 | -0,3298 | 0,42417302 | chr4 | + |
| AV051173 | 4,22 | 0,1298 | 0,884858966 | chr4 | - |
| AW011738 | 45,10 | 0,6739 | 0,115797322 | chr4 | + |
| B330016D10Rik | 19,89 | -0,7980 | 0,136585635 | chr4 | + |
| B3galt6 | 32,35 | 0,4637 | 0,360147619 | chr4 | - |
| B4galt1 | 800,37 | -0,3505 | 0,430392697 | chr4 | - |
| B4galt2 | 4,12 | 0,0868 | 0,929690763 | chr4 | - |
| Fndc10 | 6,74 | 0,1300 | 0,887301605 | chr4 | + |
| Bach2 | 59,38 | 0,1691 | 0,817817063 | chr4 | + |
| Bach2os | 13,33 | 0,8302 | 0,155946567 | chr4 | - |
| Bsdc1 | 774,10 | -0,1780 | 0,632136487 | chr4 | + |
| Bspry | 1,84 | 0,5423 | NA | chr4 | + |
| Btbd19 | 24,97 | -0,6289 | 0,227017349 | chr4 | - |
| Btf3l4 | 1206,97 | -0,1251 | 0,766748847 | chr4 | - |
| C1qa | 943,45 | 0,0480 | 0,916852736 | chr4 | - |
| C1qb | 1173,00 | 0,1208 | 0,757695155 | chr4 | - |
| C1qc | 904,83 | -0,0106 | 0,987384829 | chr4 | - |
| C530005A16Rik | 5,30 | 0,8028 | 0,192452944 | chr4 | - |
| C630043F03Rik | 99,71 | -0,1290 | 0,863101638 | chr4 | + |
| C77080 | 658,52 | -0,4510 | 0,28160417 | chr4 | - |
| C8a | 22713,28 | 0,0684 | 0,893831725 | chr4 | - |
| C8b | 9190,64 | -0,4593 | 0,252617603 | chr4 | + |
| Caap1 | 67,25 | 0,1516 | 0,772961171 | chr4 | - |
| Cachd1 | 8,39 | 0,3091 | 0,688905597 | chr4 | + |
| Camk2n1 | 1327,31 | -0,1926 | 0,628871721 | chr4 | + |
| Camta1 | 58,63 | -0,1203 | 0,859361703 | chr4 | - |
| Cap1 | 981,46 | 0,0996 | 0,815240319 | chr4 | - |
| Capzb | 1279,80 | -0,0498 | 0,917470917 | chr4 | + |
| Car9 | 2,53 | 0,3206 | NA | chr4 | + |
| Casp8ap2 | 119,28 | 0,2768 | 0,493579462 | chr4 | + |
| Casp9 | 361,56 | 0,0604 | 0,877283616 | chr4 | + |
| Casz1 | 2,77 | 0,1604 | 0,848381384 | chr4 | + |
| Cc2d1b | 130,91 | 0,0890 | 0,872774688 | chr4 | + |
| Ccdc107 | 561,19 | -0,5054 | 0,203941589 | chr4 | + |
| Ccdc163 | 84,72 | -0,0248 | 0,968998918 | chr4 | + |
| Ccdc17 | 46,33 | 0,1185 | 0,846704266 | chr4 | + |
| Svbp | 108,20 | 0,0154 | 0,985488456 | chr4 | + |
| Ccdc24 | 3,19 | 0,6758 | 0,248915105 | chr4 | - |
| Ccdc28b | 98,15 | -0,3869 | 0,298705882 | chr4 | - |
| Ccdc30 | 50,58 | -0,5985 | 0,128799205 | chr4 | - |
| Ccl19 | 57,97 | -0,0791 | 0,909426562 | chr4 | - |
| Ccl27a | 60,71 | -0,3431 | 0,468711508 | chr4 | - |
| Ccnc | 451,83 | 0,2347 | 0,549054641 | chr4 | + |
| Ccne2 | 69,63 | 0,5927 | 0,229632803 | chr4 | + |
| Ccnl2 | 815,50 | -0,4616 | 0,345819394 | chr4 | + |
| Cd52 | 290,20 | 0,3915 | 0,357779158 | chr4 | - |
| Cd72 | 46,16 | 0,2438 | 0,717188441 | chr4 | - |
| Cda | 110,80 | 0,2388 | 0,738282454 | chr4 | - |
| Cdc20 | 11,04 | 0,6603 | 0,302745739 | chr4 | - |
| Cdc26 | 647,04 | -0,0375 | 0,943108135 | chr4 | - |

| | | | | | |
|---|---|---|---|---|---|
| Cdc42 | 3791,32 | 0,1079 | 0,742942233 | chr4 | - |
| Cdca8 | 21,29 | -0,2450 | 0,72981898 | chr4 | - |
| Cdh17 | 2,18 | 0,5484 | NA | chr4 | + |
| Cdk11b | 461,00 | -0,0786 | 0,862578493 | chr4 | + |
| Cdk5rap2 | 47,83 | 0,2813 | 0,636381742 | chr4 | - |
| Cdkn2b | 4,35 | -0,3278 | 0,670802155 | chr4 | - |
| Fyb2 | 84,89 | -0,1429 | 0,843604342 | chr4 | + |
| Cela2a | 28,07 | 0,1259 | 0,877283616 | chr4 | - |
| Cep104 | 56,82 | -0,3624 | 0,433006548 | chr4 | + |
| Cep85 | 482,81 | 0,2721 | 0,486353903 | chr4 | - |
| Chchd7 | 305,79 | -0,5160 | 0,157551186 | chr4 | + |
| Chd5 | 1,94 | 0,7126 | NA | chr4 | + |
| Chd7 | 198,91 | -0,2818 | 0,505223498 | chr4 | + |
| Chmp5 | 1776,09 | -0,0045 | 0,992874254 | chr4 | + |
| CK137956 | 4,87 | -0,0616 | 0,951574951 | chr4 | - |
| Clcn6 | 16,85 | -0,1499 | 0,850453487 | chr4 | - |
| Clic4 | 1032,61 | 0,0092 | 0,983634086 | chr4 | - |
| Clspn | 4,01 | 0,0554 | 0,955234619 | chr4 | + |
| Clstn1 | 67,87 | 0,6366 | 0,131175019 | chr4 | + |
| Clta | 1145,34 | -0,3182 | 0,357290285 | chr4 | + |
| Clvs1 | 6,14 | 0,0356 | 0,974008692 | chr4 | + |
| Cmpk1 | 2077,06 | -0,1951 | 0,523471926 | chr4 | - |
| Cnksr1 | 2,36 | 0,4569 | NA | chr4 | - |
| Cnr2 | 79,52 | 0,1504 | 0,765061508 | chr4 | + |
| Cntfr | 13,86 | -0,3110 | 0,663934341 | chr4 | - |
| Cntln | 30,23 | 0,2504 | 0,693242675 | chr4 | + |
| Coa7 | 591,87 | 0,0722 | 0,888917253 | chr4 | + |
| Col15a1 | 115,02 | 0,0688 | 0,940390384 | chr4 | + |
| Col16a1 | 33,09 | -0,1836 | 0,805820649 | chr4 | + |
| Col8a2 | 7,15 | 0,0379 | 0,971699327 | chr4 | + |
| Coq3 | 529,27 | 0,0869 | 0,856684656 | chr4 | + |
| Coro2a | 12,21 | 0,2845 | 0,705755674 | chr4 | - |
| Cpne3 | 1090,87 | 0,3986 | 0,176249471 | chr4 | - |
| Ints11 | 292,75 | -0,0270 | 0,955467654 | chr4 | + |
| Cpt2 | 2379,09 | -0,3484 | 0,291034839 | chr4 | - |
| Creb3 | 814,46 | -0,3256 | 0,339967066 | chr4 | + |
| Crocc | 7,85 | -0,0342 | 0,975445719 | chr4 | - |
| Csf3r | 25,52 | 0,1453 | 0,838412507 | chr4 | + |
| Ctnnal1 | 100,32 | -0,1852 | 0,731355824 | chr4 | - |
| Ctnnbip1 | 140,84 | -0,5738 | 0,119034471 | chr4 | + |
| Ctps | 54,81 | -0,3256 | 0,565966999 | chr4 | - |
| Cyb5rl | 87,66 | -0,2536 | 0,538235194 | chr4 | + |
| Cyp2j5 | 16822,81 | -0,1005 | 0,804839321 | chr4 | - |
| Cyp2j6 | 1722,09 | -0,4824 | 0,122125617 | chr4 | - |
| Cyp2j8 | 3,59 | -0,1931 | 0,812543852 | chr4 | - |
| Cyp2j9 | 154,30 | -0,4530 | 0,298116176 | chr4 | - |
| Cyp4a10 | 5084,51 | 0,6283 | NA | chr4 | + |
| Cyp4a12a | 6512,44 | -0,6302 | 0,187760013 | chr4 | + |
| Cyp4a12b | 1148,90 | -0,9053 | 0,125180998 | chr4 | + |
| Cyp4a14 | 5891,55 | 0,6221 | NA | chr4 | - |

| | | | | | |
|---|---|---|---|---|---|
| Cyp4a31 | 357,20 | 0,2521 | 0,754953629 | chr4 | + |
| Cyp4a32 | 615,29 | 0,0862 | 0,91260409 | chr4 | + |
| Cyp4b1 | 264,04 | 0,0720 | 0,858324263 | chr4 | - |
| Cyp7a1 | 1407,02 | 0,2676 | 0,716169774 | chr4 | - |
| D630039A03Rik | 169,59 | -0,0288 | 0,969512928 | chr4 | - |
| D830031N03Rik | 9,01 | 0,3604 | 0,632136487 | chr4 | - |
| Dab1 | 2,57 | -0,4558 | 0,476764512 | chr4 | + |
| Dcaf10 | 552,83 | -0,4283 | 0,125541699 | chr4 | + |
| Dcaf12 | 539,47 | -0,2994 | 0,261960171 | chr4 | - |
| Dcdc2b | 5,10 | 0,0105 | 0,992013362 | chr4 | - |
| Dctn3 | 438,61 | -0,3192 | 0,523898331 | chr4 | - |
| Ddi2 | 216,49 | 0,2019 | 0,622222569 | chr4 | - |
| Ddost | 1414,26 | -0,1563 | 0,635751575 | chr4 | + |
| Ddx58 | 493,04 | 0,7343 | 0,053047332 | chr4 | - |
| Decr1 | 6399,88 | -0,0071 | 0,989980239 | chr4 | - |
| Dennd4c | 498,51 | 0,0222 | 0,960969671 | chr4 | + |
| Dffa | 204,53 | -0,2350 | 0,515823024 | chr4 | + |
| Dffb | 37,38 | -0,3450 | 0,511512606 | chr4 | - |
| Dhcr24 | 7225,22 | -0,1726 | 0,724955742 | chr4 | + |
| Dhdds | 417,06 | 0,1119 | 0,82704652 | chr4 | - |
| Dhrs3 | 1637,92 | -0,0071 | 0,986661276 | chr4 | + |
| Dio1 | 5025,18 | -0,2221 | 0,480124139 | chr4 | - |
| Dmap1 | 166,08 | 0,2517 | 0,617467883 | chr4 | - |
| Dmrta1 | 4,72 | -0,6487 | 0,306882295 | chr4 | + |
| Dnaja1 | 7719,46 | 0,0609 | 0,89798229 | chr4 | + |
| Dnajb5 | 55,76 | -0,0431 | 0,948366244 | chr4 | + |
| Dnajc11 | 691,45 | 0,0390 | 0,950561821 | chr4 | + |
| Dnajc16 | 109,60 | 0,0657 | 0,909834629 | chr4 | - |
| Dnajc25 | 310,84 | -0,2154 | 0,53490582 | chr4 | + |
| Dnajc8 | 562,38 | 0,2879 | 0,424561414 | chr4 | + |
| Dock7 | 204,54 | 0,6582 | 0,093284325 | chr4 | - |
| Dph2 | 41,98 | -0,1587 | 0,820554805 | chr4 | - |
| Dpy19l4 | 81,39 | 0,4252 | 0,327939573 | chr4 | - |
| Dvl1 | 629,82 | -0,3177 | 0,492254415 | chr4 | + |
| E130102H24Rik | 116,89 | -0,4253 | 0,273018284 | chr4 | - |
| E130308A19Rik | 41,02 | -0,0399 | 0,953764743 | chr4 | + |
| E2f2 | 16,58 | 0,2654 | 0,705982644 | chr4 | + |
| Ebna1bp2 | 430,94 | -0,2074 | 0,582441358 | chr4 | + |
| Ece1 | 1780,04 | -0,2607 | 0,361573776 | chr4 | + |
| Efcab14 | 314,96 | -0,0382 | 0,935107784 | chr4 | + |
| Efcab7 | 6,94 | 0,6286 | 0,338502971 | chr4 | + |
| Efhd2 | 511,29 | -0,3706 | 0,315865928 | chr4 | - |
| Eif2b3 | 251,65 | -0,0774 | 0,909889201 | chr4 | + |
| Eif3i | 2304,38 | -0,3139 | 0,191376832 | chr4 | - |
| Eif4g3 | 506,95 | -0,2561 | 0,491855001 | chr4 | + |
| Elovl1 | 729,68 | -0,2057 | 0,636381742 | chr4 | + |
| Emc1 | 418,38 | -0,2632 | 0,479279449 | chr4 | + |
| Enho | 465,30 | 0,4123 | 0,446301474 | chr4 | - |
| Epb41 | 1066,45 | -0,0884 | 0,832450834 | chr4 | - |
| Epb41l4b | 1457,56 | -0,2746 | 0,472359592 | chr4 | - |

| | | | | | |
|---|---|---|---|---|---|
| Epha2 | 158,59 | 0,5147 | 0,147012425 | chr4 | + |
| Epha7 | 33,95 | 0,3346 | 0,535475471 | chr4 | + |
| Ephb2 | 79,00 | -0,0955 | 0,912746031 | chr4 | - |
| Eps15 | 940,99 | 0,0635 | 0,877560871 | chr4 | + |
| Eri3 | 217,63 | -0,2230 | 0,575547607 | chr4 | + |
| Ermap | 5,85 | 0,0935 | 0,922849883 | chr4 | - |
| Erp44 | 1615,26 | -0,4242 | 0,056576933 | chr4 | - |
| Espn | 11,04 | 0,5064 | 0,447343221 | chr4 | - |
| Esrp1 | 4,13 | 0,6191 | 0,320305617 | chr4 | - |
| Eva1b | 35,50 | -0,2250 | 0,782329904 | chr4 | + |
| Exosc10 | 241,53 | 0,0383 | 0,949479116 | chr4 | + |
| Exosc3 | 155,30 | 0,1546 | 0,736706859 | chr4 | - |
| Extl1 | 110,47 | 0,4039 | 0,47939388 | chr4 | - |
| Eya3 | 245,33 | 0,2454 | 0,507586107 | chr4 | + |
| Faah | 1577,20 | -0,0799 | 0,834675238 | chr4 | - |
| Faf1 | 725,29 | -0,4320 | 0,081644305 | chr4 | + |
| Fam110b | 2,26 | 0,0169 | NA | chr4 | + |
| Fam131c | 16,10 | -0,6206 | 0,334804026 | chr4 | + |
| C1qtnf12 | 20,61 | -0,3190 | 0,613341495 | chr4 | + |
| Fam167b | 147,56 | 0,0418 | 0,937853416 | chr4 | - |
| Fam206a | 58,55 | 0,3391 | 0,480812921 | chr4 | + |
| Fam213b | 269,07 | -0,5598 | 0,088731202 | chr4 | - |
| Fam214b | 194,07 | -0,5004 | 0,298705882 | chr4 | - |
| Fam219a | 31,44 | 0,2437 | 0,682099278 | chr4 | - |
| Fam46b | 3,40 | 0,2197 | 0,783547703 | chr4 | + |
| Fam76a | 298,25 | 0,0159 | 0,976528475 | chr4 | - |
| Fam92a | 263,39 | -0,0116 | 0,984995389 | chr4 | - |
| Fancg | 44,79 | -0,3230 | 0,540877577 | chr4 | - |
| Fblim1 | 29,83 | 0,6207 | 0,220904318 | chr4 | - |
| Fbxl4 | 412,47 | 0,2506 | 0,532901541 | chr4 | + |
| Fbxo10 | 9,45 | -0,2391 | 0,763209303 | chr4 | - |
| Fbxo2 | 2,37 | 0,4090 | NA | chr4 | + |
| Fbxo42 | 124,72 | -0,4527 | 0,148093291 | chr4 | + |
| Fbxo44 | 30,30 | -0,1043 | 0,899463828 | chr4 | - |
| Fbxo6 | 515,37 | 0,4382 | 0,136585635 | chr4 | - |
| Fggy | 2512,27 | 0,1177 | 0,803952241 | chr4 | + |
| Fgr | 51,03 | 0,8336 | 0,058898179 | chr4 | + |
| Fhad1 | 75,07 | 0,2382 | 0,721569914 | chr4 | - |
| Fhl3 | 2,95 | -0,0500 | 0,958859839 | chr4 | + |
| Fkbp15 | 317,30 | -0,0594 | 0,899148878 | chr4 | - |
| Fktn | 67,09 | 0,6478 | 0,215046281 | chr4 | + |
| Fndc5 | 13,75 | 0,4366 | 0,503079772 | chr4 | + |
| Focad | 141,14 | -0,0518 | 0,916910357 | chr4 | + |
| Foxj3 | 199,00 | -0,3589 | 0,276893041 | chr4 | + |
| Frem1 | 8,98 | 0,3850 | 0,600547654 | chr4 | - |
| Fsd1l | 9,62 | -0,0113 | 0,991035244 | chr4 | + |
| Fuca1 | 1155,48 | -0,4087 | 0,140308933 | chr4 | + |
| Fv1 | 24,32 | 0,1833 | 0,811367208 | chr4 | + |
| Gabrd | 4,53 | 0,0576 | 0,953751362 | chr4 | - |
| Gale | 160,04 | 0,1993 | 0,775458257 | chr4 | + |

| | | | | | |
|---|---|---|---|---|---|
| Galnt12 | 2,59 | 0,1771 | 0,815188933 | chr4 | + |
| Galt | 461,14 | -0,0161 | 0,980186108 | chr4 | + |
| Gdf6 | 13,45 | 0,4558 | 0,494059529 | chr4 | + |
| Gem | 27,49 | 0,2728 | 0,655423629 | chr4 | + |
| Ggh | 342,97 | -0,1796 | 0,636447281 | chr4 | + |
| Gja10 | 2,13 | 0,6025 | NA | chr4 | - |
| Gja4 | 66,86 | 0,1023 | 0,862616891 | chr4 | - |
| Glipr2 | 20,22 | 0,9165 | 0,064196281 | chr4 | + |
| Glis1 | 2,88 | 0,7738 | 0,139596627 | chr4 | + |
| Cptp | 232,24 | 0,1599 | 0,721406886 | chr4 | - |
| Gm10560 | 5,63 | -0,5368 | 0,432005857 | chr4 | - |
| Gm12504 | 6,53 | 0,4191 | 0,569607313 | chr4 | + |
| Gm12505 | 3,02 | -0,0083 | 0,993445259 | chr4 | - |
| Gm12657 | 2,49 | 0,3148 | NA | chr4 | + |
| Tmem35b | 163,53 | 0,3594 | 0,337280658 | chr4 | + |
| Gm12992 | 6,62 | 0,3184 | 0,683522146 | chr4 | - |
| Zfp985 | 8,35 | 0,9184 | 0,123183417 | chr4 | + |
| Gm13212 | 4,34 | 0,2827 | 0,721172264 | chr4 | + |
| Gm13238 | 2,29 | 0,2054 | NA | chr4 | + |
| Gm16023 | 2,35 | 0,0257 | NA | chr4 | - |
| Gm3893 | 20,80 | -0,6378 | 0,248800492 | chr4 | + |
| Gm5860 | 14,14 | -0,5515 | 0,356860747 | chr4 | + |
| Gm6297 | 8,65 | -0,4181 | 0,563960133 | chr4 | - |
| Gmeb1 | 76,60 | 0,1530 | 0,764714786 | chr4 | - |
| Gnb1 | 1266,95 | -0,2453 | 0,385921204 | chr4 | + |
| Gne | 2487,54 | -0,4719 | 0,15641327 | chr4 | - |
| Gng10 | 162,91 | 0,6550 | 0,056273421 | chr4 | + |
| Gnl2 | 506,79 | -0,0477 | 0,935085575 | chr4 | + |
| Gpatch3 | 38,25 | -0,3115 | 0,618723125 | chr4 | + |
| Gpbp1l1 | 918,67 | -0,0283 | 0,944651681 | chr4 | + |
| Gpn2 | 101,26 | -0,3954 | 0,286763659 | chr4 | + |
| Gpr153 | 13,90 | 0,2172 | 0,781562509 | chr4 | + |
| Gpr157 | 53,32 | -0,1698 | 0,776119796 | chr4 | + |
| Gpx7 | 35,02 | -0,4715 | 0,352106106 | chr4 | - |
| Grhl3 | 2,68 | -0,0441 | 0,963027434 | chr4 | - |
| Grhpr | 3134,43 | -0,3810 | 0,376557226 | chr4 | + |
| Grik3 | 2,19 | -0,0516 | NA | chr4 | + |
| H6pd | 6361,24 | -0,1399 | 0,771786001 | chr4 | - |
| Haus6 | 57,09 | 0,5623 | 0,192656505 | chr4 | - |
| Hdac1 | 533,49 | 0,2524 | 0,491855001 | chr4 | - |
| Hdhd3 | 320,49 | 0,0087 | 0,992196557 | chr4 | - |
| Hectd3 | 579,41 | -0,4739 | 0,069704032 | chr4 | + |
| Heyl | 20,07 | 0,3099 | 0,666898692 | chr4 | + |
| Hint2 | 1055,27 | -0,4130 | 0,46503777 | chr4 | - |
| Hmgcl | 4009,17 | -0,1920 | 0,613341495 | chr4 | + |
| Hmgn2 | 591,78 | 0,6042 | 0,069072526 | chr4 | - |
| Hnrnpr | 483,28 | 0,2758 | 0,441085355 | chr4 | + |
| Hook1 | 743,14 | -0,1548 | 0,736071538 | chr4 | + |
| Hp1bp3 | 965,53 | -0,1309 | 0,69071775 | chr4 | + |
| Hpdl | 3,73 | -0,2641 | 0,74146296 | chr4 | - |

| | | | | | |
|---|---|---|---|---|---|
| Hrct1 | 4,50 | 0,1274 | 0,888648505 | chr4 | + |
| Hsdl2 | 1942,69 | 0,0691 | 0,915344724 | chr4 | + |
| Hspb11 | 134,95 | -0,1715 | 0,699593307 | chr4 | + |
| Hspb7 | 1,98 | 0,4279 | NA | chr4 | + |
| Hspg2 | 170,67 | -0,0961 | 0,895663283 | chr4 | + |
| Hyi | 512,67 | 0,1067 | 0,855874829 | chr4 | + |
| Icmt | 458,23 | -0,4005 | 0,193696579 | chr4 | + |
| Id3 | 365,33 | -0,1415 | 0,876803328 | chr4 | + |
| Iffo2 | 396,22 | 0,4865 | 0,144426924 | chr4 | + |
| Ifnlr1 | 3,99 | 0,4904 | 0,4648339 | chr4 | + |
| Ift74 | 90,93 | 0,2389 | 0,671813639 | chr4 | + |
| Ikbkap | 167,43 | 0,1784 | 0,663692832 | chr4 | - |
| Il11ra1 | 262,64 | 0,1415 | 0,744313101 | chr4 | + |
| Il22ra1 | 191,76 | 0,5083 | 0,209733815 | chr4 | + |
| Inip | 75,12 | 0,1705 | 0,730379979 | chr4 | - |
| Inpp5b | 587,44 | 0,3944 | 0,265311253 | chr4 | + |
| Ints8 | 400,65 | -0,1796 | 0,592295685 | chr4 | - |
| Invs | 194,00 | 0,0032 | 0,996127775 | chr4 | + |
| Ipo13 | 157,01 | -0,2122 | 0,635835796 | chr4 | - |
| Ipp | 327,31 | 0,1270 | 0,804048972 | chr4 | + |
| Iqcc | 88,25 | 0,4632 | 0,235706307 | chr4 | - |
| Isg15 | 435,27 | 0,1164 | 0,862616891 | chr4 | - |
| Itgb3bp | 129,77 | 0,4717 | 0,166993075 | chr4 | - |
| Jak1 | 1903,65 | 0,0190 | 0,965898926 | chr4 | - |
| Jun | 132,14 | 0,3624 | 0,502079303 | chr4 | - |
| Kazn | 15,54 | -0,0632 | 0,946813357 | chr4 | - |
| Kcnab2 | 8,89 | 0,0172 | 0,987066007 | chr4 | - |
| Kdf1 | 30,45 | 0,4344 | 0,505223498 | chr4 | + |
| Kdm1a | 220,89 | -0,2445 | 0,481853495 | chr4 | - |
| Kdm4a | 282,88 | 0,2171 | 0,60487792 | chr4 | - |
| Kdm4c | 132,81 | 0,4088 | 0,273018284 | chr4 | + |
| Khdrbs1 | 507,53 | 0,1440 | 0,770796088 | chr4 | - |
| Kif12 | 4,66 | -0,1425 | 0,875717638 | chr4 | - |
| Kif1b | 2173,61 | -0,1145 | 0,757883527 | chr4 | - |
| Kif24 | 7,33 | 0,1008 | 0,915621354 | chr4 | - |
| Kif2c | 8,61 | 0,4528 | 0,526620417 | chr4 | - |
| Klf4 | 12,82 | -0,0560 | 0,953638766 | chr4 | - |
| Klhl17 | 81,03 | -0,1042 | 0,848794352 | chr4 | - |
| Klhl21 | 560,71 | -0,4318 | 0,250303145 | chr4 | + |
| Klhl9 | 1206,23 | 0,2479 | 0,462012946 | chr4 | - |
| Kpna6 | 719,56 | 0,0353 | 0,943108135 | chr4 | - |
| Kti12 | 169,33 | -0,4155 | 0,234261614 | chr4 | + |
| Laptm5 | 213,44 | 0,1114 | 0,814156035 | chr4 | + |
| Lck | 17,13 | 0,3060 | 0,659350082 | chr4 | - |
| Ldlrad2 | 2,35 | -0,2063 | NA | chr4 | - |
| P3h1 | 133,67 | -0,1703 | 0,698530192 | chr4 | + |
| Leprot | 497,72 | 0,2401 | 0,437867403 | chr4 | + |
| Lepr | 281,88 | 0,1121 | 0,761178054 | chr4 | + |
| Lpar1 | 22,07 | -0,0387 | 0,969130697 | chr4 | - |
| Lrp8 | 2,10 | -0,1572 | NA | chr4 | + |

| | | | | | |
|---|---|---|---|---|---|
| Lrrc41 | 409,74 | -0,0055 | 0,990858052 | chr4 | + |
| Lrrc42 | 304,59 | -0,2495 | 0,447657149 | chr4 | - |
| Lrrc47 | 8,81 | 0,2495 | 0,757888816 | chr4 | + |
| Lsm10 | 317,19 | -0,3599 | 0,330791426 | chr4 | + |
| Lurap1l | 1340,92 | -0,1573 | 0,713614788 | chr4 | + |
| Luzp1 | 271,84 | 0,1398 | 0,693242675 | chr4 | + |
| Lyn | 639,61 | 0,0208 | 0,958903346 | chr4 | + |
| Lypla2 | 438,63 | -0,3993 | 0,408859672 | chr4 | - |
| Lyrm2 | 438,48 | 0,6906 | 0,068869694 | chr4 | + |
| Lzic | 75,38 | 0,3225 | 0,479481987 | chr4 | + |
| Macf1 | 3,23 | 0,4866 | 0,466793644 | chr4 | - |
| Mad2l2 | 120,32 | -0,2070 | 0,63251737 | chr4 | + |
| Man1c1 | 42,06 | -0,0598 | 0,932012942 | chr4 | - |
| Manea | 385,40 | 0,1468 | 0,760115687 | chr4 | - |
| Map3k6 | 19,57 | -0,0647 | 0,947294154 | chr4 | + |
| Map3k7 | 607,45 | -0,2497 | 0,425562494 | chr4 | + |
| Map7d1 | 268,08 | -0,2988 | 0,482520633 | chr4 | - |
| Marcksl1 | 114,18 | -0,0258 | 0,966113955 | chr4 | + |
| Masp2 | 5503,20 | 0,0256 | 0,952400398 | chr4 | + |
| Mast2 | 148,03 | -0,3916 | 0,357290285 | chr4 | - |
| Meaf6 | 89,31 | -0,4048 | 0,322848017 | chr4 | + |
| Mecr | 449,82 | -0,4791 | 0,07229395 | chr4 | + |
| Med18 | 29,95 | 0,2840 | 0,623297443 | chr4 | - |
| Med8 | 554,66 | -0,1911 | 0,646093772 | chr4 | + |
| Megf9 | 109,17 | -0,2476 | 0,640948606 | chr4 | - |
| Melk | 3,46 | 0,1364 | 0,871793968 | chr4 | + |
| Mfn2 | 1027,46 | -0,1677 | 0,696922493 | chr4 | - |
| Mfsd2a | 857,53 | 0,1650 | 0,789064438 | chr4 | - |
| Mib2 | 161,84 | -0,0952 | 0,905274865 | chr4 | - |
| Mier1 | 691,21 | 0,1705 | 0,699549296 | chr4 | + |
| Miip | 156,80 | -0,4529 | 0,236204287 | chr4 | - |
| Minos1 | 1778,02 | -0,4117 | 0,119325906 | chr4 | - |
| Mknk1 | 251,17 | -0,1810 | 0,670139713 | chr4 | + |
| Mllt3 | 73,91 | 0,5409 | 0,202636202 | chr4 | - |
| Mmachc | 261,81 | 0,1203 | 0,841150543 | chr4 | - |
| Mmp23 | 3,46 | 0,5928 | 0,360959313 | chr4 | - |
| Mms22l | 26,18 | 0,7422 | 0,149780096 | chr4 | + |
| Mob3b | 767,99 | 0,4196 | 0,155221887 | chr4 | - |
| Mob3c | 226,60 | -0,4090 | 0,190096538 | chr4 | + |
| Morn1 | 28,51 | -0,0496 | 0,950847359 | chr4 | + |
| Mpdz | 262,08 | -0,1229 | 0,83796616 | chr4 | - |
| Mroh7 | 2,63 | -0,1309 | 0,877700558 | chr4 | - |
| Mrpl37 | 498,86 | -0,4240 | 0,13640556 | chr4 | - |
| Mrpl50 | 1552,98 | 0,2120 | 0,587223505 | chr4 | - |
| Mrps15 | 363,39 | -0,4411 | 0,197630691 | chr4 | + |
| Mrto4 | 335,13 | -0,2968 | 0,495713543 | chr4 | - |
| Msmp | 3,04 | -0,9053 | 0,104289738 | chr4 | - |
| Mtap | 346,93 | 0,2870 | 0,419942821 | chr4 | + |
| Mtf1 | 192,93 | -0,1622 | 0,661449625 | chr4 | + |
| Mtfr1l | 1207,59 | -0,1859 | 0,527820373 | chr4 | - |

| | | | | | |
|---|---|---|---|---|---|
| Mthfr | 78,54 | 0,1787 | 0,757883527 | chr4 | + |
| Mtor | 465,73 | -0,0121 | 0,981991128 | chr4 | + |
| Mul1 | 499,39 | -0,0856 | 0,860890261 | chr4 | + |
| Mup10 | 4,27 | -0,6079 | 0,343698954 | chr4 | - |
| Mup20 | 217605,80 | -0,4644 | 0,278905276 | chr4 | - |
| Mup4 | 250,24 | -0,7529 | 0,06989423 | chr4 | - |
| Mutyh | 3,93 | 0,1335 | 0,88422091 | chr4 | + |
| Mxra8 | 112,16 | -0,1256 | 0,793250646 | chr4 | + |
| Mycbp | 429,82 | -0,2348 | 0,435658245 | chr4 | + |
| Myom3 | 9,48 | 0,3168 | 0,684297941 | chr4 | + |
| Mysm1 | 226,83 | 0,1916 | 0,623572684 | chr4 | - |
| Nadk | 3677,07 | 0,0113 | 0,979186519 | chr4 | + |
| Nans | 358,12 | 0,0156 | 0,981194934 | chr4 | + |
| Nasp | 469,62 | -0,2829 | 0,293840783 | chr4 | - |
| Nbl1 | 4,33 | 0,2189 | 0,794944296 | chr4 | - |
| Nbn | 350,10 | -0,2402 | 0,482009343 | chr4 | + |
| Ncbp1 | 502,76 | -0,3098 | 0,402592894 | chr4 | + |
| Ncdn | 83,50 | -0,2410 | 0,701944163 | chr4 | - |
| Ncmap | 5,51 | -0,3396 | 0,659221885 | chr4 | - |
| Ndc1 | 108,73 | -0,1100 | 0,811137679 | chr4 | + |
| Ndufaf4 | 899,11 | -0,2605 | 0,517251824 | chr4 | + |
| Ndufaf6 | 304,27 | -0,0895 | 0,859505657 | chr4 | - |
| Ndufb6 | 1536,52 | -0,3764 | 0,437620234 | chr4 | - |
| Necab1 | 712,86 | 0,3406 | 0,317646908 | chr4 | - |
| Necap2 | 96,03 | -0,2431 | 0,671975571 | chr4 | - |
| Nfia | 749,68 | -0,3624 | 0,247584138 | chr4 | + |
| Nfib | 1673,96 | -0,3281 | 0,382591532 | chr4 | - |
| Nfx1 | 724,74 | -0,1589 | 0,596014309 | chr4 | + |
| Nfyc | 301,00 | -0,4947 | 0,05185674 | chr4 | - |
| Nipal3 | 147,34 | 0,2792 | 0,580137509 | chr4 | - |
| Nipsnap3b | 1348,61 | 0,3416 | 0,298116176 | chr4 | + |
| Nkain1 | 2,39 | 0,3816 | NA | chr4 | - |
| Nmnat1 | 161,63 | -0,1057 | 0,831777176 | chr4 | - |
| Noc2l | 413,75 | -0,1126 | 0,780418568 | chr4 | + |
| Nol6 | 151,81 | -0,1089 | 0,848214837 | chr4 | - |
| Nol9 | 200,73 | -0,0940 | 0,849477252 | chr4 | + |
| Nr0b2 | 304,64 | -0,2479 | 0,764363417 | chr4 | + |
| Nrd1 | 1212,40 | -0,2640 | 0,258561012 | chr4 | + |
| Nsmaf | 340,84 | 0,0647 | 0,887538079 | chr4 | - |
| Nsun4 | 418,63 | -0,1605 | 0,716547565 | chr4 | - |
| Nudc | 808,48 | -0,2583 | 0,523898331 | chr4 | - |
| Nudt2 | 345,07 | 0,0154 | 0,981978399 | chr4 | + |
| Oma1 | 292,58 | 0,1767 | 0,712560511 | chr4 | + |
| Orc3 | 436,63 | 0,1808 | 0,754953629 | chr4 | - |
| Orm1 | 27310,60 | -0,2287 | 0,596219566 | chr4 | + |
| Orm2 | 823,66 | 0,5113 | 0,451815929 | chr4 | + |
| Osbpl9 | 3728,13 | -0,1904 | 0,572115465 | chr4 | - |
| Oscp1 | 4,41 | 0,3006 | 0,69945599 | chr4 | + |
| Osgin2 | 41,62 | 0,0866 | 0,903337956 | chr4 | - |
| Otud3 | 51,29 | -0,7052 | 0,071000263 | chr4 | - |

| | | | | | |
|---|---|---|---|---|---|
| Otud6b | 879,77 | -0,0217 | 0,971699327 | chr4 | - |
| Pabpc4 | 232,48 | -0,5556 | 0,059261122 | chr4 | + |
| Padi2 | 8,26 | 0,6032 | 0,355134088 | chr4 | + |
| Pafah2 | 1712,16 | 0,1433 | 0,727318837 | chr4 | + |
| Pank4 | 171,80 | 0,0884 | 0,857773796 | chr4 | + |
| Park7 | 3181,87 | -0,3459 | 0,148068171 | chr4 | - |
| Pax5 | 2,34 | -0,6624 | NA | chr4 | - |
| Pcsk9 | 563,44 | -0,0687 | 0,909426562 | chr4 | - |
| Pde4b | 191,77 | -0,1186 | 0,853848645 | chr4 | + |
| Pdik1l | 222,26 | 0,2779 | 0,43159313 | chr4 | - |
| Pdp1 | 36,24 | 0,2547 | 0,656182304 | chr4 | - |
| Pdpn | 8,32 | 0,6425 | 0,321845648 | chr4 | - |
| Pdzk1ip1 | 144,04 | 0,2807 | 0,657471667 | chr4 | + |
| Pef1 | 759,88 | 0,1881 | 0,559032648 | chr4 | + |
| Per3 | 371,96 | 0,1380 | 0,790271076 | chr4 | - |
| Pex10 | 62,65 | 0,5160 | 0,217147923 | chr4 | + |
| Pex14 | 604,13 | -0,2137 | 0,636260727 | chr4 | - |
| Pgm2 | 3301,98 | -0,2847 | 0,230037893 | chr4 | + |
| Phactr4 | 239,96 | -0,0184 | 0,979212524 | chr4 | - |
| Phc2 | 249,22 | -0,2418 | 0,604679897 | chr4 | + |
| Phf13 | 85,68 | -0,1155 | 0,821443376 | chr4 | - |
| Pigo | 128,66 | -0,2291 | 0,546240378 | chr4 | - |
| Pigv | 393,00 | -0,1594 | 0,618749555 | chr4 | - |
| Pik3cd | 45,32 | 0,2811 | 0,623297443 | chr4 | - |
| Pik3r3 | 30,78 | 0,2891 | 0,610382836 | chr4 | + |
| Pink1 | 3246,71 | -0,2043 | 0,594322944 | chr4 | - |
| Pla2g5 | 3,45 | 0,5861 | 0,339421245 | chr4 | - |
| Plaa | 940,46 | -0,4140 | 0,138292882 | chr4 | - |
| Plekhf2 | 332,61 | -0,3123 | 0,402726862 | chr4 | - |
| Plekhg5 | 53,80 | -0,1122 | 0,887169146 | chr4 | + |
| Plekhm2 | 104,51 | -0,3349 | 0,437620234 | chr4 | - |
| Plekhn1 | 4,94 | 0,0976 | 0,915435 | chr4 | - |
| Plin2 | 11081,46 | -0,0336 | 0,957004103 | chr4 | - |
| Plk3 | 676,42 | -0,3828 | 0,370757182 | chr4 | - |
| Plod1 | 19,79 | 0,5961 | 0,297304489 | chr4 | - |
| Pm20d2 | 85,99 | -0,0893 | 0,908138272 | chr4 | - |
| Pnisr | 500,28 | -0,2549 | 0,54267972 | chr4 | + |
| Pnrc1 | 673,45 | -0,1648 | 0,676982843 | chr4 | - |
| Pnrc2 | 1094,92 | 0,2352 | 0,434726768 | chr4 | - |
| Podn | 41,66 | -0,6053 | 0,22527304 | chr4 | - |
| Pole3 | 220,23 | 0,0135 | 0,985488456 | chr4 | - |
| Polr1e | 72,64 | -0,2980 | 0,592406077 | chr4 | + |
| Pomgnt1 | 236,68 | -0,2055 | 0,613685815 | chr4 | + |
| Plpp3 | 1552,61 | 0,0956 | 0,829674496 | chr4 | + |
| Ppcs | 344,10 | -0,2525 | 0,399118497 | chr4 | - |
| Ppie | 323,64 | -0,0074 | 0,988512389 | chr4 | - |
| Ppih | 187,49 | -0,0610 | 0,909081586 | chr4 | - |
| Ppp1r8 | 210,37 | 0,1846 | 0,649875139 | chr4 | - |
| Ppt1 | 590,39 | 0,1512 | 0,619039366 | chr4 | + |
| Pqlc2 | 137,57 | 0,0322 | 0,95463264 | chr4 | - |

| | | | | | |
|---|---|---|---|---|---|
| Pramef8 | 139,67 | 0,5369 | 0,142156924 | chr4 | + |
| Prdm16 | 7,54 | 0,4075 | 0,577347444 | chr4 | - |
| Prdm2 | 361,53 | -0,4230 | 0,298774903 | chr4 | - |
| Prdx1 | 42589,49 | -0,0587 | 0,887839974 | chr4 | + |
| Prkaa2 | 1016,84 | -0,4164 | 0,153204397 | chr4 | - |
| Prkcz | 287,14 | -0,2575 | 0,532778528 | chr4 | - |
| Prpf38a | 291,28 | 0,3228 | 0,411552342 | chr4 | - |
| Prpf4 | 215,31 | -0,2114 | 0,606237187 | chr4 | + |
| Psip1 | 459,05 | 0,0817 | 0,839302072 | chr4 | - |
| Psmb2 | 1451,02 | -0,4287 | 0,126676113 | chr4 | + |
| Ptafr | 8,96 | 0,7167 | 0,258057977 | chr4 | + |
| Ptbp3 | 480,14 | -0,2438 | 0,451182001 | chr4 | - |
| Ptch2 | 5,22 | 0,0383 | 0,971699327 | chr4 | + |
| Ptp4a2 | 3430,28 | -0,2861 | 0,336671811 | chr4 | + |
| Hacd4 | 19,90 | 0,5358 | 0,331966576 | chr4 | - |
| Ptpn3 | 533,25 | -0,1363 | 0,804754692 | chr4 | - |
| Ptprd | 2376,77 | -0,1083 | 0,813415373 | chr4 | - |
| Ptprf | 1578,89 | 0,2575 | 0,522014539 | chr4 | - |
| Ptpru | 4,41 | -0,1127 | 0,902022501 | chr4 | - |
| Pum1 | 426,11 | -0,2121 | 0,548442171 | chr4 | + |
| Pusl1 | 68,61 | -0,5872 | 0,090895721 | chr4 | - |
| Rab2a | 2212,29 | -0,3390 | 0,224918351 | chr4 | + |
| Rab3b | 20,13 | 0,6263 | 0,293001163 | chr4 | + |
| Rad23b | 4186,12 | 0,1287 | 0,7588734 | chr4 | + |
| Rad54b | 8,49 | -0,1071 | 0,90764023 | chr4 | + |
| Rad54l | 3,57 | -0,1318 | 0,88067049 | chr4 | - |
| Rap1gap | 39,50 | 0,2759 | 0,690262514 | chr4 | + |
| Rars2 | 285,25 | 0,2167 | 0,692220176 | chr4 | + |
| Raver2 | 3,88 | -0,0347 | 0,974449819 | chr4 | + |
| Rbbp4 | 2679,69 | -0,3318 | 0,27946684 | chr4 | - |
| Rbm12b1 | 87,87 | 0,1126 | 0,815253842 | chr4 | + |
| Rbm12b2 | 53,86 | -0,2753 | 0,583782269 | chr4 | + |
| Rcan3 | 27,44 | 0,8767 | 0,09820895 | chr4 | - |
| Rcc1 | 102,69 | -0,3009 | 0,551409879 | chr4 | - |
| Reck | 42,97 | 0,0703 | 0,913749296 | chr4 | + |
| Rer1 | 2301,57 | -0,2752 | 0,425066328 | chr4 | - |
| Rere | 313,53 | -0,1964 | 0,615542082 | chr4 | + |
| Rgp1 | 369,48 | -0,4187 | 0,155039247 | chr4 | + |
| Rgs3 | 232,28 | -0,4526 | NA | chr4 | + |
| Ripk2 | 123,83 | -0,4972 | 0,131133541 | chr4 | - |
| Rlf | 259,04 | -0,1988 | 0,632136487 | chr4 | - |
| Rmdn1 | 664,02 | -0,4013 | 0,338536613 | chr4 | + |
| Rmrp | 36,11 | -0,8763 | 0,092854537 | chr4 | - |
| Rnf11 | 1562,26 | -0,0307 | 0,952429505 | chr4 | - |
| Rnf186 | 140,47 | 0,1194 | 0,895663283 | chr4 | + |
| Rnf19b | 432,66 | 0,0057 | 0,99162344 | chr4 | + |
| Rnf20 | 365,32 | 0,1362 | 0,733803092 | chr4 | + |
| Rnf220 | 158,89 | -0,0933 | 0,851472884 | chr4 | - |
| Rnf38 | 159,31 | 0,0434 | 0,939351122 | chr4 | - |
| Rngtt | 185,87 | 0,1957 | 0,636361813 | chr4 | + |

| | | | | | |
|---|---|---|---|---|---|
| Ror1 | 8,74 | 0,0650 | 0,948314012 | chr4 | + |
| Rpa2 | 117,18 | 0,0421 | 0,945127597 | chr4 | + |
| Rpl22 | 4147,72 | -0,2901 | 0,345662712 | chr4 | + |
| Rpp25l | 284,55 | 0,0037 | 0,995663291 | chr4 | - |
| Rps19-ps3 | 22,67 | -0,3633 | 0,565966999 | chr4 | - |
| Rps20 | 4256,61 | -0,4440 | 0,306882295 | chr4 | - |
| Rps6ka1 | 142,35 | -0,1862 | 0,682099278 | chr4 | - |
| Rraga | 565,68 | 0,0995 | 0,807670244 | chr4 | + |
| Rragc | 708,24 | -0,1452 | 0,63367203 | chr4 | + |
| Rragd | 42,06 | 0,4459 | 0,376078101 | chr4 | + |
| Rsg1 | 2,50 | -0,1734 | NA | chr4 | + |
| Rspo1 | 2,10 | 0,2406 | NA | chr4 | + |
| Runx1t1 | 5,39 | -0,6643 | 0,30485919 | chr4 | + |
| Runx3 | 5,60 | -0,0082 | 0,993738083 | chr4 | + |
| Rusc2 | 110,54 | -0,2564 | 0,644399192 | chr4 | + |
| S100pbp | 65,76 | 0,2920 | 0,553850707 | chr4 | - |
| Scmh1 | 88,17 | -0,3670 | 0,420100204 | chr4 | + |
| Scp2 | 114121,10 | -0,1860 | 0,55880551 | chr4 | - |
| Sdc3 | 554,01 | 0,0685 | 0,888710003 | chr4 | + |
| Sdcbp | 1754,48 | 0,2839 | 0,387366838 | chr4 | + |
| Sdf4 | 2410,47 | -0,1800 | 0,460037772 | chr4 | + |
| Sdhb | 3814,42 | -0,3461 | 0,379588771 | chr4 | + |
| Sdr16c5 | 23,09 | 0,6436 | 0,320520607 | chr4 | - |
| Sec61b | 1369,81 | -0,2509 | 0,757395951 | chr4 | + |
| Selenon | 32,27 | 0,5226 | 0,335357114 | chr4 | - |
| Serinc2 | 135,88 | 0,6671 | 0,140308933 | chr4 | - |
| Sesn2 | 63,96 | 0,0669 | 0,939351122 | chr4 | - |
| Sf3a3 | 380,22 | 0,3118 | 0,320530124 | chr4 | + |
| Sfn | 8,07 | -0,2037 | 0,811137679 | chr4 | - |
| Sfpq | 973,62 | -0,1015 | 0,777136205 | chr4 | + |
| Sh2d5 | 3,44 | -0,5090 | 0,426922409 | chr4 | + |
| Sh3bgrl3 | 131,37 | 0,0011 | 0,99829832 | chr4 | - |
| Sh3d21 | 13,36 | -0,2989 | 0,672263059 | chr4 | - |
| Sh3gl2 | 3,66 | -0,1872 | 0,823578308 | chr4 | + |
| Shb | 300,95 | -0,5101 | 0,131898706 | chr4 | - |
| Sigmar1 | 1139,26 | 0,1308 | 0,75616611 | chr4 | - |
| Ski | 216,05 | -0,2432 | 0,592428959 | chr4 | - |
| Skint2 | 6,82 | -0,8272 | 0,169324582 | chr4 | + |
| Skint3 | 3,82 | 0,9234 | 0,109852731 | chr4 | + |
| Skint4 | 3,52 | 0,1726 | 0,83796616 | chr4 | + |
| Slc25a33 | 389,78 | -0,8078 | 0,066417448 | chr4 | - |
| Slc25a34 | 155,61 | 0,0815 | 0,93538092 | chr4 | - |
| Slc2a1 | 155,88 | 0,0224 | 0,978280656 | chr4 | + |
| Slc31a2 | 196,77 | -0,2317 | 0,527820373 | chr4 | + |
| Slc31a1 | 2,72 | 0,8149 | 0,122125617 | chr4 | + |
| Slc35a1 | 595,74 | 0,1233 | 0,758873213 | chr4 | - |
| Slc35d1 | 946,93 | -0,0558 | 0,932012942 | chr4 | - |
| Slc35e2 | 1383,06 | 0,2217 | 0,500346905 | chr4 | + |
| Slc44a1 | 262,32 | 0,1912 | 0,693242675 | chr4 | + |
| Slc6a9 | 492,28 | -0,2314 | 0,640716535 | chr4 | + |

| | | | | | |
|---|---|---|---|---|---|
| Slc9a1 | 74,40 | -0,3144 | 0,529723794 | chr4 | + |
| Smap2 | 544,04 | 0,0527 | 0,890887823 | chr4 | - |
| Smc2 | 280,06 | 0,3879 | 0,168698533 | chr4 | + |
| Smim1 | 114,62 | -0,3654 | 0,498009876 | chr4 | - |
| Smim12 | 209,15 | -0,3168 | 0,393529719 | chr4 | + |
| Smim8 | 314,53 | 0,1084 | 0,811093035 | chr4 | - |
| Smpdl3b | 10,90 | 0,5766 | 0,365428542 | chr4 | - |
| Smu1 | 1354,70 | 0,0179 | 0,976991666 | chr4 | - |
| Snapc3 | 32,97 | -0,3229 | 0,574366053 | chr4 | + |
| Snhg12 | 114,05 | -0,2342 | 0,630950393 | chr4 | + |
| Snhg3 | 233,16 | -0,6546 | 0,073845576 | chr4 | - |
| Snip1 | 183,99 | -0,3357 | 0,332496065 | chr4 | + |
| Snora16a | 8,09 | -0,0679 | 0,944925159 | chr4 | + |
| Snora44 | 3,11 | -0,5851 | 0,354199455 | chr4 | + |
| Snora61 | 9,28 | -0,2838 | 0,718823184 | chr4 | + |
| Snord99 | 4,28 | 0,3655 | 0,620846456 | chr4 | + |
| Snrnp40 | 135,73 | 0,1991 | 0,598404146 | chr4 | + |
| Snx30 | 148,42 | 0,0153 | 0,979051488 | chr4 | + |
| Spag8 | 4,70 | -0,2980 | 0,702177047 | chr4 | - |
| Spata6 | 19,37 | 0,1219 | 0,8763719 | chr4 | + |
| Spen | 116,42 | 0,0833 | 0,89879469 | chr4 | - |
| Spsb1 | 11,62 | 0,9766 | 0,078273639 | chr4 | - |
| Srm | 245,53 | -0,5569 | 0,092888997 | chr4 | + |
| Srrm1 | 377,48 | -0,0479 | 0,913936879 | chr4 | - |
| Srsf10 | 624,51 | 0,0314 | 0,954308706 | chr4 | + |
| Srsf4 | 320,75 | -0,4756 | 0,250526648 | chr4 | + |
| Ssbp3 | 395,73 | -0,0583 | 0,888963178 | chr4 | + |
| Ssu72 | 1025,59 | -0,2809 | 0,391036013 | chr4 | + |
| St3gal3 | 1014,97 | -0,2775 | 0,55880551 | chr4 | - |
| Stil | 4,98 | -0,0512 | 0,958903346 | chr4 | + |
| Stmn1 | 356,71 | 0,0534 | 0,944213761 | chr4 | + |
| Stk40 | 905,01 | 0,1037 | 0,76908367 | chr4 | + |
| Stoml2 | 1075,01 | -0,0844 | 0,825855713 | chr4 | - |
| Stx12 | 500,56 | 0,0887 | 0,811093035 | chr4 | - |
| Stx17 | 426,15 | -0,0975 | 0,828304746 | chr4 | + |
| Susd1 | 126,32 | -0,3716 | 0,435658245 | chr4 | - |
| Svep1 | 14,05 | 0,3873 | 0,587966975 | chr4 | - |
| Syf2 | 758,24 | -0,2209 | 0,526620417 | chr4 | + |
| Sytl1 | 46,16 | -0,1126 | 0,864755921 | chr4 | - |
| Szrd1 | 1416,61 | 0,1000 | 0,848661267 | chr4 | - |
| Szt2 | 97,20 | -0,2505 | 0,657129833 | chr4 | - |
| Taf12 | 275,11 | 0,0131 | 0,980767643 | chr4 | + |
| Tal1 | 17,38 | 0,0069 | 0,994106851 | chr4 | + |
| Tardbp | 1948,15 | 0,0134 | 0,981025197 | chr4 | - |
| Tbc1d2 | 6,55 | -0,0894 | 0,92589889 | chr4 | - |
| Tcea3 | 776,19 | 0,4671 | 0,059974041 | chr4 | + |
| Tceanc2 | 252,06 | -0,0667 | 0,869442758 | chr4 | - |
| Eloa | 581,37 | -0,1621 | 0,583782269 | chr4 | - |
| Tdrd7 | 465,14 | 0,0494 | 0,912811384 | chr4 | + |
| Tek | 249,93 | 0,1977 | 0,609072466 | chr4 | + |

| | | | | | |
|---|---|---|---|---|---|
| Tesk1 | 149,45 | -0,6082 | 0,208201861 | chr4 | + |
| Tesk2 | 288,41 | -0,0763 | 0,854690386 | chr4 | + |
| Tex10 | 154,22 | 0,1818 | 0,647914259 | chr4 | - |
| Tgfbr1 | 510,14 | -0,1618 | 0,683464556 | chr4 | + |
| Tgs1 | 273,94 | 0,4335 | 0,135356973 | chr4 | + |
| Thap3 | 75,60 | -0,1204 | 0,869442758 | chr4 | - |
| Thrap3 | 935,20 | 0,1871 | 0,569425025 | chr4 | - |
| Tie1 | 177,47 | -0,1050 | 0,826107735 | chr4 | - |
| Tinagl1 | 232,29 | -0,1770 | 0,764363417 | chr4 | - |
| Tle1 | 550,08 | -0,4544 | 0,14157858 | chr4 | - |
| Tln1 | 568,37 | -0,1228 | 0,792197952 | chr4 | - |
| Tlr12 | 144,79 | 0,0413 | 0,960130794 | chr4 | - |
| Tlr4 | 55,73 | 0,7686 | 0,068351758 | chr4 | + |
| Tm2d1 | 293,41 | -0,3689 | 0,352135503 | chr4 | - |
| Tmco4 | 174,18 | -0,0094 | 0,987952599 | chr4 | + |
| Tmem125 | 80,27 | 0,0891 | 0,876803328 | chr4 | - |
| Tmem200b | 13,27 | 0,8510 | 0,12746076 | chr4 | + |
| Tmem201 | 133,66 | -0,3808 | 0,38079602 | chr4 | - |
| Tmem222 | 395,57 | 0,1244 | 0,752490199 | chr4 | - |
| Tmem234 | 313,02 | -0,3123 | 0,615799562 | chr4 | + |
| Tmem245 | 95,58 | 0,3111 | 0,428679263 | chr4 | - |
| Tmem246 | 6,92 | 0,3781 | 0,614668942 | chr4 | - |
| Tmem38b | 2011,52 | -0,5761 | 0,08440425 | chr4 | + |
| Tmem39b | 38,55 | -0,2530 | 0,720531276 | chr4 | - |
| Tmem50a | 2386,50 | -0,0505 | 0,911712632 | chr4 | - |
| Tmem51 | 141,49 | 0,8021 | 0,058138256 | chr4 | - |
| Tmem53 | 375,78 | 0,2971 | 0,404267039 | chr4 | + |
| Tmem54 | 4,35 | 0,8518 | 0,151646977 | chr4 | + |
| Tmem55a | 265,44 | -0,1343 | 0,717188441 | chr4 | + |
| Tmem59 | 4661,83 | -0,2942 | 0,361447043 | chr4 | + |
| Tmem64 | 794,69 | -0,0879 | 0,814156035 | chr4 | + |
| Tmem67 | 46,79 | 0,4542 | 0,377944963 | chr4 | - |
| Tmem69 | 344,29 | 0,0629 | 0,917148255 | chr4 | - |
| Tmem82 | 211,82 | -0,4694 | 0,367870281 | chr4 | - |
| Tmem88b | 19,14 | -0,7888 | 0,12296145 | chr4 | - |
| Tmem8b | 18,66 | -0,1396 | 0,860253707 | chr4 | + |
| Tmod1 | 6,42 | -0,4175 | 0,569869994 | chr4 | + |
| Tnc | 26,24 | 0,5449 | 0,341021075 | chr4 | - |
| Tnfrsf14 | 28,41 | 0,7857 | 0,154188248 | chr4 | - |
| Tnfrsf18 | 5,06 | 0,4235 | 0,559607122 | chr4 | + |
| Tnfrsf1b | 507,92 | -0,3157 | 0,494281913 | chr4 | - |
| Tnfrsf25 | 3,53 | 0,3324 | 0,656913078 | chr4 | + |
| Tnfrsf4 | 3,22 | -0,5281 | 0,411460129 | chr4 | + |
| Toe1 | 169,76 | 0,0590 | 0,90870562 | chr4 | - |
| Tomm5 | 688,82 | 0,0069 | 0,990189789 | chr4 | - |
| Topors | 516,90 | -0,1548 | 0,632136487 | chr4 | - |
| Smim27 | 168,88 | -0,4103 | 0,502772604 | chr4 | + |
| Tox | 41,80 | -0,6520 | 0,145466971 | chr4 | - |
| Tpm2 | 72,35 | -0,4862 | 0,335637765 | chr4 | - |
| Tprgl | 715,04 | -0,1581 | 0,658318626 | chr4 | - |

| | | | | | |
|---|---|---|---|---|---|
| Trabd2b | 6,62 | -0,0742 | 0,941754349 | chr4 | + |
| Trappc3 | 679,41 | -0,2440 | 0,416450614 | chr4 | + |
| Trim14 | 454,79 | 0,5689 | 0,157274894 | chr4 | - |
| Trim32 | 133,87 | 0,2411 | 0,558489002 | chr4 | + |
| Trim62 | 6,96 | 0,0589 | 0,95295436 | chr4 | + |
| Triqk | 4,07 | 0,2102 | 0,802299336 | chr4 | + |
| Trit1 | 268,11 | -0,1347 | 0,7232495 | chr4 | + |
| Trmt10b | 84,15 | 0,1392 | 0,800238939 | chr4 | + |
| Trnau1ap | 260,97 | -0,3478 | 0,374181006 | chr4 | - |
| Trp53inp1 | 2671,20 | 0,4757 | 0,390554711 | chr4 | + |
| Ttc39b | 573,78 | -0,0988 | 0,834675238 | chr4 | - |
| Ttc4 | 365,88 | 0,1148 | 0,81937394 | chr4 | - |
| Ttpa | 12463,13 | 0,5433 | 0,104006655 | chr4 | + |
| Tusc1 | 6,40 | -0,3251 | 0,672726404 | chr4 | - |
| Txlna | 373,04 | 0,0088 | 0,988766352 | chr4 | - |
| Txn1 | 7948,38 | -0,3133 | 0,237485754 | chr4 | - |
| Txndc12 | 816,94 | 0,2424 | 0,404820274 | chr4 | + |
| Ubap2 | 175,90 | -0,3495 | 0,385921204 | chr4 | - |
| Ube2j1 | 670,68 | -0,2130 | 0,430881045 | chr4 | + |
| Ube2j2 | 18,76 | -0,5289 | 0,378003797 | chr4 | + |
| Ube2r2 | 1591,67 | -0,4527 | 0,071729514 | chr4 | + |
| Ube2u | 91,94 | 0,4241 | 0,336618885 | chr4 | + |
| Ube4b | 524,75 | -0,4057 | 0,135356973 | chr4 | - |
| Ubiad1 | 143,52 | 0,2992 | 0,381639797 | chr4 | - |
| Ubr4 | 1016,95 | -0,2734 | 0,50625923 | chr4 | + |
| Ubxn10 | 3,74 | 1,0199 | 0,063989705 | chr4 | - |
| Ubxn11 | 3,72 | 0,4103 | 0,569607313 | chr4 | + |
| Ufl1 | 586,48 | 0,2048 | 0,666442376 | chr4 | - |
| Ugcg | 518,62 | -0,0759 | 0,889997259 | chr4 | + |
| Unc13b | 150,37 | -0,3129 | 0,414445273 | chr4 | + |
| Uqcrh | 8924,22 | -0,4374 | 0,118025078 | chr4 | - |
| Urod | 2119,75 | -0,4336 | 0,047204826 | chr4 | - |
| Usp1 | 324,09 | 0,3754 | 0,290028383 | chr4 | + |
| Usp24 | 998,76 | -0,2100 | 0,511512606 | chr4 | + |
| Usp45 | 271,43 | 0,3620 | 0,282672306 | chr4 | + |
| Usp48 | 450,14 | -0,1876 | 0,539569536 | chr4 | + |
| Utp11 | 317,45 | -0,0541 | 0,895780958 | chr4 | - |
| Vamp3 | 328,97 | -0,0429 | 0,931322892 | chr4 | - |
| Vps13d | 819,76 | -0,2175 | 0,526894474 | chr4 | - |
| Vwa1 | 8,58 | 0,1703 | 0,847796138 | chr4 | - |
| Wasf2 | 147,65 | 0,4078 | 0,321653279 | chr4 | + |
| Wdr31 | 4,92 | 1,0349 | 0,069200865 | chr4 | - |
| Wdr78 | 9,81 | -0,3106 | 0,688905597 | chr4 | - |
| Wrap73 | 86,13 | -0,1429 | 0,79351129 | chr4 | + |
| Wdtc1 | 722,01 | -0,6025 | 0,155520806 | chr4 | - |
| Whrn | 2,92 | -0,9842 | 0,066396922 | chr4 | - |
| Wnt4 | 30,40 | -0,1345 | 0,857560803 | chr4 | + |
| Wwp1 | 1454,76 | -0,2804 | 0,408981984 | chr4 | - |
| Xkr8 | 134,77 | 0,3796 | 0,34284412 | chr4 | - |
| Xpa | 149,10 | -0,0566 | 0,919467514 | chr4 | - |

| | | | | | |
|---|---|---|---|---|---|
| Yars | 452,75 | -0,3469 | 0,288059609 | chr4 | + |
| Ybx1 | 2355,31 | -0,3476 | 0,20031188 | chr4 | - |
| Yipf1 | 869,21 | -0,1233 | 0,760736495 | chr4 | + |
| Yrdc | 151,27 | -0,5087 | 0,086429447 | chr4 | + |
| Ythdf2 | 399,54 | -0,1843 | 0,564849438 | chr4 | - |
| Zbtb17 | 106,73 | -0,0559 | 0,943108135 | chr4 | + |
| Zbtb40 | 220,53 | 0,1932 | 0,636449345 | chr4 | - |
| Zbtb48 | 159,36 | 0,1215 | 0,842502441 | chr4 | - |
| Zbtb5 | 151,20 | 0,0543 | 0,917042614 | chr4 | - |
| Zbtb8a | 15,68 | 0,3239 | 0,677398032 | chr4 | - |
| Zbtb8os | 1087,59 | -0,2167 | 0,569876243 | chr4 | + |
| Zc3h12a | 40,94 | 0,0799 | 0,912811384 | chr4 | - |
| Zcchc11 | 269,62 | 0,3843 | 0,181899789 | chr4 | + |
| Zcchc17 | 323,78 | 0,1079 | 0,808156264 | chr4 | - |
| Zcchc7 | 191,99 | -0,1159 | 0,804403985 | chr4 | + |
| Zdhhc18 | 218,47 | -0,2038 | 0,661568683 | chr4 | - |
| Zdhhc21 | 219,94 | 0,3644 | 0,340522938 | chr4 | - |
| Zfp189 | 159,60 | -0,1370 | 0,74224943 | chr4 | + |
| Zfp292 | 214,23 | 0,3077 | 0,472408228 | chr4 | - |
| Zfp362 | 38,98 | 0,2725 | 0,640716535 | chr4 | - |
| Zfp37 | 23,56 | 0,8796 | 0,064726648 | chr4 | - |
| Zfp46 | 12,28 | 0,0039 | 0,996826205 | chr4 | + |
| Zfp462 | 47,15 | -0,0662 | 0,925100183 | chr4 | + |
| Zfp534 | 2,35 | 0,1464 | NA | chr4 | - |
| Zfp593 | 64,25 | 0,0734 | 0,900660395 | chr4 | - |
| Zfp618 | 3,07 | -0,0100 | 0,991098974 | chr4 | + |
| Zfp69 | 11,73 | 0,2177 | 0,794928847 | chr4 | - |
| Zfp691 | 71,08 | -0,1569 | 0,764714786 | chr4 | - |
| Zfyve9 | 213,48 | -0,0157 | 0,980481306 | chr4 | - |
| Zmpste24 | 1988,53 | 0,5164 | 0,069704032 | chr4 | - |
| Zmym1 | 71,33 | 0,5582 | 0,122800588 | chr4 | - |
| Zmym4 | 196,41 | 0,3760 | 0,261235845 | chr4 | - |
| Zscan20 | 34,37 | -0,4169 | 0,437620234 | chr4 | - |
| Zswim5 | 42,88 | 0,3417 | 0,527820373 | chr4 | + |
| Zyg11a | 43,77 | -0,4484 | 0,338476353 | chr4 | - |
| Zyg11b | 925,24 | 0,0586 | 0,899942442 | chr4 | - |
| 0610040B10Rik | 47,71 | -0,0669 | 0,941248754 | chr5 | + |
| 0610040J01Rik | 508,86 | -0,2927 | 0,290028383 | chr5 | + |
| 1110006O24Rik | 7,00 | -0,5011 | 0,472244842 | chr5 | - |
| 1500011B03Rik | 15,14 | 0,6937 | 0,217835326 | chr5 | - |
| 1600023N17Rik | 3,81 | -0,2035 | 0,807751236 | chr5 | - |
| 1700028E10Rik | 63,44 | 0,4322 | 0,385427218 | chr5 | + |
| 1700028K03Rik | 2,32 | 0,0876 | NA | chr5 | + |
| 1700096K18Rik | 14,05 | 0,1538 | 0,845988264 | chr5 | + |
| 1700109H08Rik | 3,48 | -0,2360 | 0,771038929 | chr5 | + |
| 2010109A12Rik | 2,09 | 0,3948 | NA | chr5 | + |
| 2210016L21Rik | 61,80 | -0,1178 | 0,828542262 | chr5 | + |
| 2210417A02Rik | 53,49 | 0,6772 | 0,140518953 | chr5 | + |
| 2410131K14Rik | 106,18 | -0,1068 | 0,869825447 | chr5 | + |
| 2610524H06Rik | 12,13 | 0,9661 | 0,087710179 | chr5 | - |

| | | | | | |
|---|---|---|---|---|---|
| 2700038G22Rik | 2,95 | 0,0035 | 0,996987906 | chr5 | + |
| 2810006K23Rik | 199,56 | 0,2441 | 0,569584928 | chr5 | + |
| 2900005J15Rik | 8,67 | -0,3558 | 0,636260727 | chr5 | + |
| 2900026A02Rik | 1163,80 | 0,0671 | 0,906325382 | chr5 | - |
| 3110082I17Rik | 117,31 | 0,5029 | 0,3899381 | chr5 | - |
| 4831440E17Rik | 21,98 | -0,4233 | 0,490509101 | chr5 | + |
| 4930431F12Rik | 26,28 | 0,8514 | 0,064726648 | chr5 | + |
| 4930515G01Rik | 4,88 | 0,2372 | 0,773900067 | chr5 | - |
| 4933404O12Rik | 60,99 | -0,6482 | 0,127207217 | chr5 | + |
| 5031425E22Rik | 73,97 | -0,2050 | 0,736302391 | chr5 | - |
| 5033403H07Rik | 697,88 | 0,0576 | 0,899463828 | chr5 | - |
| 5430416N02Rik | 66,82 | -0,5551 | 0,379104437 | chr5 | - |
| 5730480H06Rik | 21,60 | -0,3688 | 0,568858283 | chr5 | + |
| 5830416I19Rik | 5,84 | -0,2615 | 0,749691831 | chr5 | + |
| 5830473C10Rik | 718,80 | -0,0386 | 0,950532633 | chr5 | + |
| 5930430L01Rik | 23,26 | 0,4393 | 0,521815657 | chr5 | - |
| 6030443J06Rik | 21,12 | -0,1711 | 0,81812331 | chr5 | + |
| 6330418K02Rik | 29,24 | -0,0734 | 0,93056206 | chr5 | + |
| 9230114K14Rik | 11,52 | 0,1958 | 0,816325182 | chr5 | + |
| 9330182L06Rik | 6,86 | 0,0202 | 0,985488456 | chr5 | + |
| Cfap69 | 7,18 | 0,5026 | 0,470066395 | chr5 | - |
| A430033K04Rik | 69,32 | 0,3838 | 0,391701733 | chr5 | + |
| A630072M18Rik | 67,05 | 0,0389 | 0,951372792 | chr5 | + |
| A930011G23Rik | 3,03 | 0,4077 | 0,548561752 | chr5 | - |
| A930024E05Rik | 3,86 | 0,0201 | 0,98507181 | chr5 | + |
| AA792892 | 3,06 | 0,6012 | 0,335264505 | chr5 | + |
| Aacs | 602,58 | -0,1535 | 0,747375272 | chr5 | + |
| Aasdh | 69,73 | 0,1107 | 0,876498461 | chr5 | - |
| Abcb1b | 12,91 | -0,7099 | 0,222968111 | chr5 | + |
| Abcb4 | 3239,84 | 0,4520 | 0,217231457 | chr5 | + |
| Abcb8 | 370,24 | -0,0864 | 0,847736143 | chr5 | + |
| Abcb9 | 15,75 | -0,5140 | 0,396411176 | chr5 | - |
| Abcf2 | 676,58 | -0,2851 | 0,42977312 | chr5 | - |
| Abcg3 | 81,71 | 0,5957 | 0,108470987 | chr5 | - |
| Abhd1 | 48,77 | 0,0207 | 0,978280656 | chr5 | + |
| Abhd11 | 465,44 | -0,3444 | 0,382335568 | chr5 | + |
| Abhd11os | 5,79 | -0,1433 | 0,875717638 | chr5 | - |
| Ablim2 | 17,81 | 0,1391 | 0,862616891 | chr5 | + |
| Acacb | 852,88 | 0,1561 | 0,804418611 | chr5 | + |
| Acad10 | 244,66 | 0,3533 | 0,234074257 | chr5 | - |
| Acads | 2426,27 | -0,3268 | 0,4891753 | chr5 | - |
| Acox3 | 427,19 | -0,2251 | 0,645100453 | chr5 | + |
| Actb | 5587,21 | -0,2066 | 0,555913954 | chr5 | - |
| Actr3b | 2,57 | -0,0086 | 0,992806079 | chr5 | + |
| Adam1a | 6,69 | -0,1009 | 0,915227882 | chr5 | - |
| Adam22 | 5,18 | 0,6999 | 0,271504179 | chr5 | - |
| Adamts3 | 4,55 | -0,1498 | 0,869067375 | chr5 | - |
| Adap1 | 15,63 | 0,4656 | 0,465782618 | chr5 | - |
| Add1 | 365,47 | 0,2779 | 0,394148671 | chr5 | + |
| Grk3 | 90,00 | 0,7044 | 0,104029003 | chr5 | - |

| | | | | | |
|---|---|---|---|---|---|
| Afap1 | 16,14 | 0,6098 | 0,301862555 | chr5 | + |
| Aff1 | 337,52 | -0,1177 | 0,775458257 | chr5 | + |
| Afm | 5622,89 | -0,0825 | 0,834675238 | chr5 | + |
| Agap3 | 236,33 | -0,2263 | 0,695575308 | chr5 | + |
| Agbl5 | 48,66 | -0,1333 | 0,840105898 | chr5 | + |
| Agfg2 | 433,55 | -0,4049 | 0,348561153 | chr5 | - |
| Aimp2 | 178,27 | -0,2486 | 0,548818911 | chr5 | - |
| Akap9 | 451,55 | 0,0958 | 0,829308766 | chr5 | + |
| Alb | 2284647,37 | -0,1088 | 0,798875954 | chr5 | + |
| Aldh2 | 16797,26 | -0,4517 | 0,216003134 | chr5 | - |
| Alkbh2 | 66,25 | -0,4711 | 0,335264505 | chr5 | - |
| Alkbh4 | 45,06 | -0,4965 | 0,269236255 | chr5 | + |
| Alox5ap | 47,30 | 0,5154 | 0,315094061 | chr5 | + |
| Amz1 | 9,55 | 0,5883 | 0,359936311 | chr5 | + |
| Anapc4 | 301,42 | 0,2532 | 0,609310775 | chr5 | + |
| Anapc5 | 923,67 | -0,1727 | 0,565966999 | chr5 | - |
| Anapc7 | 357,91 | -0,1029 | 0,802329508 | chr5 | + |
| Ankib1 | 349,04 | -0,0353 | 0,942935841 | chr5 | - |
| Ankle2 | 467,73 | -0,0016 | 0,997464504 | chr5 | + |
| Ankrd13a | 230,96 | -0,2393 | 0,501951423 | chr5 | + |
| Ankrd17 | 791,63 | -0,0077 | 0,987508343 | chr5 | - |
| Antxr2 | 425,24 | -0,1464 | 0,732937334 | chr5 | - |
| Anxa3 | 115,76 | 0,3448 | 0,412631408 | chr5 | + |
| Ap1s1 | 675,72 | 0,0462 | 0,909834629 | chr5 | - |
| Ap4m1 | 116,34 | 0,0554 | 0,921657556 | chr5 | + |
| Ap5z1 | 100,47 | -0,3064 | 0,625823055 | chr5 | + |
| Apbb2 | 154,35 | -0,2564 | 0,572534156 | chr5 | - |
| Arap2 | 284,17 | 0,1559 | 0,692971754 | chr5 | - |
| Arhgap24 | 78,34 | 0,7367 | 0,081525882 | chr5 | + |
| Arl6ip4 | 380,53 | -0,5413 | 0,190097808 | chr5 | + |
| Armc10 | 115,87 | -0,1531 | 0,739141404 | chr5 | + |
| Arpc1a | 895,27 | -0,2994 | 0,208518411 | chr5 | + |
| Arpc1b | 723,69 | -0,1294 | 0,814052362 | chr5 | + |
| Arpc3 | 2037,39 | -0,0978 | 0,771786001 | chr5 | + |
| Art3 | 70,68 | 0,0020 | 0,997464504 | chr5 | + |
| Asic3 | 2,24 | -0,4215 | NA | chr5 | + |
| Asl | 8335,36 | -0,4544 | 0,144524949 | chr5 | - |
| Atg9b | 4,09 | -0,1080 | 0,90633584 | chr5 | - |
| Atp10d | 50,45 | 0,2236 | 0,718541916 | chr5 | + |
| Atp2a2 | 1878,89 | -0,3437 | 0,475352298 | chr5 | - |
| Atp5k | 4479,94 | -0,2267 | 0,517783511 | chr5 | - |
| Atp6v0a2 | 490,72 | -0,3935 | 0,170992427 | chr5 | + |
| Atp8a1 | 80,39 | 0,3505 | 0,449319302 | chr5 | - |
| Atraid | 1765,04 | -0,1707 | 0,669817756 | chr5 | + |
| Atxn2 | 413,57 | -0,5767 | 0,060157716 | chr5 | + |
| Auts2 | 3,18 | 0,2459 | 0,757188376 | chr5 | - |
| AW549542 | 145,04 | -0,5437 | 0,244197385 | chr5 | - |
| Azgp1 | 44720,00 | -0,2931 | 0,264796388 | chr5 | + |
| B3glct | 30,84 | -0,4556 | 0,387366838 | chr5 | + |
| Baiap2l1 | 435,49 | -0,4748 | 0,243141521 | chr5 | - |

| | | | | | |
|---|---|---|---|---|---|
| Baz1b | 909,40 | -0,0873 | 0,820554805 | chr5 | + |
| BC005561 | 59,36 | 0,6337 | 0,165172417 | chr5 | + |
| BC037034 | 110,24 | -0,2228 | 0,725248714 | chr5 | - |
| Bcl7a | 114,82 | -0,0947 | 0,860107955 | chr5 | + |
| Bcl7b | 136,55 | -0,5054 | 0,283527481 | chr5 | + |
| Bend4 | 19,57 | -0,0346 | 0,971699327 | chr5 | - |
| Bhlha15 | 9,07 | 0,4888 | 0,483930201 | chr5 | + |
| Bloc1s4 | 94,40 | -0,4003 | 0,39354827 | chr5 | - |
| Bmp2k | 77,92 | 0,1297 | 0,804091647 | chr5 | + |
| Bod1l | 295,78 | 0,2348 | 0,550266125 | chr5 | - |
| Brat1 | 89,40 | 0,0676 | 0,911491358 | chr5 | + |
| Brca2 | 35,24 | -0,1725 | 0,814771257 | chr5 | + |
| Brdt | 29,74 | 0,3118 | 0,606429785 | chr5 | + |
| Babam2 | 311,94 | -0,2830 | 0,456714923 | chr5 | + |
| Bri3 | 394,41 | -0,5693 | 0,091451194 | chr5 | + |
| Bri3bp | 443,71 | -0,0995 | 0,781678648 | chr5 | + |
| Bst1 | 11,29 | 0,4373 | 0,543927518 | chr5 | + |
| Btc | 43,78 | 0,4196 | 0,466812905 | chr5 | - |
| Bud31 | 1327,69 | -0,1073 | 0,806737261 | chr5 | + |
| C130050O18Rik | 11,76 | 0,3026 | 0,681917507 | chr5 | + |
| C1qtnf7 | 19,91 | 0,0087 | 0,992729152 | chr5 | + |
| C330024D21Rik | 3,74 | -0,5634 | 0,395294986 | chr5 | + |
| Cacna2d1 | 13,12 | -0,0973 | 0,913936879 | chr5 | + |
| Cad | 90,52 | -0,1083 | 0,873132528 | chr5 | + |
| Caln1 | 33,00 | -0,5988 | 0,226673106 | chr5 | + |
| Camkk2 | 112,52 | -0,1350 | 0,822394911 | chr5 | - |
| Card11 | 5,79 | 0,1516 | 0,867698337 | chr5 | - |
| Cc2d2a | 46,79 | 0,9192 | 0,050536418 | chr5 | + |
| Ccdc149 | 34,05 | -0,1166 | 0,87093463 | chr5 | - |
| Ccdc158 | 3,87 | 0,5968 | 0,34812896 | chr5 | - |
| Ccdc18 | 7,56 | 0,6094 | 0,34827585 | chr5 | + |
| Ccdc62 | 23,97 | -0,3613 | 0,563218321 | chr5 | + |
| Bicdl1 | 6,50 | 0,2112 | 0,804110902 | chr5 | - |
| Ccdc96 | 2,57 | 0,2040 | NA | chr5 | + |
| Ccl24 | 85,83 | -0,1323 | 0,83176435 | chr5 | - |
| Ccng2 | 269,20 | -0,0797 | 0,888763259 | chr5 | + |
| Ccni | 1937,89 | -0,2626 | 0,371976461 | chr5 | - |
| Cct6a | 2357,17 | -0,0155 | 0,973458724 | chr5 | + |
| Ccz1 | 591,44 | -0,2077 | 0,524801822 | chr5 | - |
| Cd36 | 1060,86 | 0,3971 | 0,234431867 | chr5 | - |
| Cd38 | 385,99 | 0,5444 | 0,057218287 | chr5 | + |
| Cdc7 | 4,25 | 0,2685 | 0,736302391 | chr5 | + |
| Cdk14 | 23,11 | 0,7268 | 0,178097604 | chr5 | - |
| Cdk2ap1 | 373,48 | -0,4351 | 0,181404581 | chr5 | - |
| Cdk5 | 323,35 | -0,3583 | 0,223366879 | chr5 | - |
| Cdk6 | 19,83 | -0,0323 | 0,971756379 | chr5 | + |
| Cdk8 | 471,09 | -0,4471 | 0,074111544 | chr5 | + |
| Cdkl2 | 143,36 | 0,2813 | 0,484897818 | chr5 | - |
| Cds1 | 7,56 | 0,0926 | 0,922227922 | chr5 | + |
| Cenpa | 89,61 | 0,0251 | 0,971699327 | chr5 | + |

| | | | | | |
|---|---|---|---|---|---|
| Cenpc1 | 86,46 | 0,3033 | 0,49077279 | chr5 | - |
| Cep135 | 66,26 | 0,1966 | 0,720531276 | chr5 | + |
| Cgref1 | 20,39 | -0,0479 | 0,955504626 | chr5 | - |
| Chchd2 | 3727,48 | -0,5359 | 0,096261775 | chr5 | - |
| Chek2 | 40,04 | -0,0301 | 0,969512928 | chr5 | + |
| Chfr | 261,31 | 0,0774 | 0,85496895 | chr5 | + |
| Chic2 | 137,68 | -0,6163 | 0,076420365 | chr5 | - |
| Chpf2 | 201,77 | -0,0555 | 0,938661865 | chr5 | + |
| Chst12 | 7,33 | -0,2922 | 0,712160795 | chr5 | + |
| Cit | 3,88 | 0,2249 | 0,783547703 | chr5 | + |
| Cldn12 | 1669,69 | 0,0058 | 0,990189789 | chr5 | - |
| Cldn15 | 8,88 | -0,4060 | 0,582615116 | chr5 | + |
| Cldn3 | 572,70 | 0,1295 | 0,886748924 | chr5 | + |
| Clip1 | 418,88 | -0,2705 | 0,343715756 | chr5 | - |
| Clip2 | 24,32 | 0,7260 | 0,172446714 | chr5 | - |
| Clock | 562,29 | 0,0393 | 0,930782422 | chr5 | - |
| Cmklr1 | 39,40 | 0,0197 | 0,981991128 | chr5 | - |
| Cnot6l | 847,55 | 0,1754 | 0,627992485 | chr5 | - |
| Cnpy4 | 159,16 | 0,2493 | 0,495713543 | chr5 | + |
| Commd8 | 493,73 | 0,1250 | 0,754524942 | chr5 | - |
| Cops4 | 651,77 | 0,0339 | 0,939403124 | chr5 | + |
| Cops6 | 2146,18 | -0,1083 | 0,774830664 | chr5 | + |
| Coq2 | 470,35 | -0,1595 | 0,618749555 | chr5 | - |
| Coq5 | 382,25 | -0,1062 | 0,777753633 | chr5 | + |
| Coro1c | 306,72 | 0,1209 | 0,781342762 | chr5 | - |
| Cox18 | 72,14 | -0,2206 | 0,709072206 | chr5 | - |
| Cox19 | 577,57 | -0,3805 | 0,148471484 | chr5 | - |
| Cox6a1 | 3494,49 | -0,3175 | 0,684264304 | chr5 | - |
| Cpeb2 | 549,63 | 0,2410 | 0,602938667 | chr5 | + |
| Cpsf4 | 252,58 | -0,1906 | 0,61720787 | chr5 | + |
| Cpz | 9,60 | 0,0305 | 0,977120448 | chr5 | - |
| Crcp | 760,27 | -0,1481 | 0,670139713 | chr5 | + |
| Crlf2 | 28,08 | 0,3895 | 0,596014309 | chr5 | - |
| Crot | 13351,00 | 0,2787 | 0,536603375 | chr5 | - |
| Cryba4 | 2,00 | 0,0207 | NA | chr5 | - |
| Crybb3 | 15,03 | -0,2328 | 0,77634365 | chr5 | - |
| Ctbp1 | 348,32 | -0,2679 | 0,548561752 | chr5 | - |
| Cux1 | 257,94 | -0,1210 | 0,799982428 | chr5 | - |
| Cux2 | 6,75 | 0,8086 | 0,151731441 | chr5 | - |
| Cxcl1 | 119,20 | -0,4622 | 0,519206644 | chr5 | + |
| Cxcl11 | 13,89 | 0,3056 | 0,693803702 | chr5 | - |
| Cxcl13 | 14,72 | -0,3028 | 0,693242675 | chr5 | + |
| Cxcl2 | 5,72 | 0,4222 | 0,559428875 | chr5 | + |
| Cxcl9 | 480,82 | 0,2958 | 0,52359831 | chr5 | - |
| Cyp3a11 | 143706,52 | 0,0216 | 0,962779295 | chr5 | - |
| Cyp3a13 | 5905,83 | -0,0469 | 0,936825775 | chr5 | - |
| Cyp3a16 | 43,29 | -0,1103 | 0,88569455 | chr5 | - |
| Cyp3a44 | 16,63 | 0,4040 | 0,533752885 | chr5 | - |
| Cyp3a57 | 6,97 | -1,0279 | 0,073599483 | chr5 | + |
| Cyp51 | 2889,59 | -0,4137 | 0,127901037 | chr5 | - |

| | | | | | |
|---|---|---|---|---|---|
| Cyth3 | 92,34 | 0,2313 | 0,589961159 | chr5 | + |
| D5Ertd579e | 873,29 | 0,1581 | 0,702177047 | chr5 | - |
| D5Ertd605e | 3,45 | 0,3346 | 0,64775526 | chr5 | + |
| D930016D06Rik | 184,46 | 0,2652 | 0,504369791 | chr5 | + |
| Daglb | 260,49 | -0,1507 | 0,75436056 | chr5 | + |
| Dbf4 | 53,06 | -0,5864 | 0,155221887 | chr5 | - |
| Dck | 85,17 | 0,0491 | 0,943108135 | chr5 | + |
| Dcun1d4 | 617,61 | -0,1654 | 0,612582336 | chr5 | + |
| Ddx51 | 103,32 | 0,0353 | 0,948314012 | chr5 | + |
| Ddx54 | 281,39 | -0,5050 | 0,312564069 | chr5 | + |
| Ddx55 | 47,82 | -0,4154 | 0,34456871 | chr5 | + |
| Denr | 936,66 | -0,2112 | 0,499218871 | chr5 | + |
| Depdc5 | 156,36 | -0,0490 | 0,931322892 | chr5 | + |
| Dgkq | 247,90 | -0,1832 | 0,672205411 | chr5 | - |
| Dhx15 | 1405,79 | 0,1449 | 0,716547565 | chr5 | - |
| Dhx37 | 78,65 | -0,4153 | 0,343449498 | chr5 | - |
| Diablo | 613,89 | -0,0102 | 0,985628536 | chr5 | - |
| Dmtf1 | 221,75 | 0,1449 | 0,771038929 | chr5 | - |
| Dnajb6 | 1739,54 | -0,1934 | 0,614056134 | chr5 | + |
| Dnajc2 | 326,86 | -0,5348 | 0,064283934 | chr5 | - |
| Dnajc30 | 157,20 | 0,0861 | 0,850265859 | chr5 | + |
| Dr1 | 300,89 | 0,4202 | 0,181882538 | chr5 | + |
| Dtx1 | 5,41 | -0,3894 | 0,602509602 | chr5 | - |
| Dtx2 | 42,21 | -0,2103 | 0,744313101 | chr5 | + |
| Dynll1 | 406,15 | 0,1460 | 0,775308498 | chr5 | - |
| E130309D02Rik | 106,18 | -0,5236 | 0,160011116 | chr5 | - |
| Eif2ak1 | 622,24 | -0,1602 | 0,617869186 | chr5 | + |
| Eif2b1 | 531,60 | -0,2832 | 0,39060004 | chr5 | - |
| Eif2b4 | 389,02 | -0,4436 | 0,145466971 | chr5 | - |
| Eif3b | 1093,10 | -0,5235 | 0,100145036 | chr5 | + |
| Eif4h | 1972,84 | -0,0449 | 0,93056206 | chr5 | - |
| Emilin1 | 199,63 | 0,3673 | 0,527304027 | chr5 | + |
| Enoph1 | 44,14 | 0,0817 | 0,909774279 | chr5 | + |
| Ep400 | 409,32 | -0,1792 | 0,665658933 | chr5 | - |
| Ephb4 | 293,57 | 0,0941 | 0,812943523 | chr5 | + |
| Selenoi | 825,11 | -0,2385 | 0,37601808 | chr5 | + |
| Ereg | 5,92 | 0,7457 | 0,231675652 | chr5 | + |
| Erp29 | 888,08 | -0,4969 | 0,157586062 | chr5 | - |
| Evc | 33,07 | 0,6548 | 0,162272166 | chr5 | - |
| Evc2 | 16,80 | 0,6037 | 0,339404199 | chr5 | + |
| Evi5 | 1822,20 | 0,0997 | 0,811137679 | chr5 | - |
| Exoc1 | 134,37 | 0,2051 | 0,662842062 | chr5 | + |
| Fam109a | 90,21 | 0,0381 | 0,968669888 | chr5 | + |
| Fam114a1 | 517,29 | 0,2972 | 0,499662767 | chr5 | + |
| Fam133b | 156,06 | -0,0599 | 0,911210861 | chr5 | + |
| Abraxas1 | 117,83 | 0,4143 | 0,247068524 | chr5 | - |
| Fam185a | 128,25 | -0,2122 | 0,604326523 | chr5 | + |
| Fam193a | 113,91 | -0,2175 | 0,611389077 | chr5 | + |
| Fam20c | 495,15 | -0,2969 | 0,293404423 | chr5 | + |
| Fam216a | 36,71 | -0,3101 | 0,597702953 | chr5 | - |

| | | | | | |
|---|---|---|---|---|---|
| Fam220a | 80,38 | 0,0056 | 0,992874254 | chr5 | + |
| Fam222a | 14,53 | -0,9257 | 0,089136522 | chr5 | + |
| Fam47e | 505,03 | -0,6364 | 0,191376832 | chr5 | + |
| Fam53a | 152,36 | -0,5045 | 0,207825164 | chr5 | - |
| Fam69a | 255,29 | -0,0890 | 0,853266197 | chr5 | - |
| Fastk | 305,29 | -0,4467 | 0,228519955 | chr5 | - |
| Fbrsl1 | 77,05 | -0,3313 | 0,522840311 | chr5 | - |
| Fbxl13 | 2,67 | 0,4694 | 0,468711508 | chr5 | - |
| Fbxl18 | 9,57 | -0,0477 | 0,963027434 | chr5 | - |
| Fbxl5 | 530,45 | 0,3230 | 0,328388243 | chr5 | - |
| Fbxo21 | 958,60 | -0,2645 | 0,571917529 | chr5 | + |
| Fbxw8 | 584,81 | -0,1280 | 0,762859711 | chr5 | - |
| Fgfbp1 | 1,90 | 0,4396 | NA | chr5 | - |
| Fgfr3 | 319,79 | 0,4162 | 0,19521384 | chr5 | + |
| Fgfrl1 | 335,87 | -0,0996 | 0,839302072 | chr5 | + |
| Fgl2 | 76,48 | 0,5580 | 0,175457889 | chr5 | + |
| Ficd | 95,24 | -0,1372 | 0,815253842 | chr5 | + |
| Fip1l1 | 191,64 | 0,3378 | 0,360948982 | chr5 | + |
| Fis1 | 1542,70 | -0,3679 | 0,156266936 | chr5 | + |
| Flt1 | 73,05 | 0,3687 | 0,389200964 | chr5 | - |
| Flt3 | 7,41 | 0,0572 | 0,953764743 | chr5 | - |
| Fndc4 | 385,48 | -0,1249 | 0,803874753 | chr5 | - |
| Fosl2 | 119,09 | 0,0219 | 0,973840209 | chr5 | + |
| Foxk1 | 243,42 | 0,3196 | 0,517431746 | chr5 | + |
| Fry | 19,11 | -0,1236 | 0,88127014 | chr5 | + |
| Fryl | 179,19 | 0,1650 | 0,681091125 | chr5 | - |
| Fscn1 | 25,76 | 0,0996 | 0,903918719 | chr5 | + |
| Mrm2 | 101,05 | 0,0587 | 0,913102981 | chr5 | - |
| Fzd1 | 74,40 | 0,1030 | 0,865679649 | chr5 | - |
| G3bp2 | 1539,84 | 0,4494 | 0,177916821 | chr5 | - |
| Gak | 759,02 | -0,3056 | 0,360147619 | chr5 | - |
| Galnt11 | 215,25 | 0,2054 | 0,640716535 | chr5 | + |
| Gatad1 | 874,49 | -0,1893 | 0,559634089 | chr5 | - |
| Gatc | 302,95 | -0,2735 | 0,345409527 | chr5 | - |
| Gatsl2 | 12,65 | 0,1125 | 0,899463828 | chr5 | + |
| Nipsnap2 | 195,50 | -0,0333 | 0,947819654 | chr5 | + |
| Gbp11 | 265,51 | 0,6100 | NA | chr5 | - |
| Gbp4 | 69,46 | 0,1720 | 0,793239112 | chr5 | - |
| Gc | 190075,38 | -0,0367 | 0,925034844 | chr5 | - |
| Gckr | 2550,36 | -0,0237 | 0,965770119 | chr5 | + |
| Gcn1l1 | 361,85 | -0,0409 | 0,935254011 | chr5 | + |
| Get4 | 296,40 | -0,2904 | 0,396254007 | chr5 | + |
| Gigyf1 | 185,87 | -0,1410 | 0,838868977 | chr5 | + |
| Git2 | 120,14 | -0,1500 | 0,740663168 | chr5 | - |
| Gjc3 | 38,00 | -0,2323 | 0,721569914 | chr5 | - |
| Glmn | 84,91 | -0,0380 | 0,965271905 | chr5 | - |
| Glt1d1 | 256,81 | -0,1169 | 0,86019237 | chr5 | + |
| Gltp | 72,61 | 0,3600 | 0,425493131 | chr5 | - |
| Gm10416 | 7,04 | 0,9617 | 0,10175355 | chr5 | - |
| Rpl37rt | 96,14 | -0,7404 | 0,142012892 | chr5 | - |

| | | | | | |
|---|---|---|---|---|---|
| Gm15408 | 60,28 | 0,2841 | 0,577257466 | chr5 | + |
| Gm15421 | 25,27 | 0,0519 | 0,952511874 | chr5 | - |
| Gm15446 | 13,02 | 0,5106 | 0,414126925 | chr5 | + |
| Gm15708 | 4,11 | -0,3437 | 0,644046286 | chr5 | + |
| Gm15787 | 6,07 | -0,4644 | 0,511143322 | chr5 | - |
| Hectd4 | 196,03 | -0,4039 | 0,282189277 | chr5 | + |
| Gm19619 | 54,70 | 0,6999 | 0,175474884 | chr5 | - |
| Gm20605 | 4,76 | -0,9575 | 0,103014216 | chr5 | + |
| Gm3414 | 35,96 | -0,1212 | 0,87328174 | chr5 | - |
| Gm7854 | 7,51 | -0,0062 | 0,995663291 | chr5 | - |
| Gm9958 | 13,57 | -0,6262 | 0,281688847 | chr5 | + |
| Gna12 | 419,83 | 0,1207 | 0,847419446 | chr5 | - |
| Gnai1 | 18,64 | -0,4455 | 0,469270232 | chr5 | - |
| Gnb2 | 764,09 | -0,5048 | 0,199710944 | chr5 | - |
| Gnpda2 | 76,49 | 0,6147 | 0,224946844 | chr5 | - |
| Golga3 | 242,90 | -0,0072 | 0,990189789 | chr5 | + |
| Gpn1 | 233,12 | -0,1828 | 0,695978671 | chr5 | + |
| Gpn3 | 179,19 | 0,2450 | 0,566087418 | chr5 | + |
| Gpr12 | 8,83 | 0,4237 | 0,564724076 | chr5 | - |
| Adgra3 | 345,28 | -0,1739 | 0,582630294 | chr5 | - |
| Adgrd1 | 10,45 | 0,0292 | 0,978386671 | chr5 | + |
| Gpr146 | 1247,66 | 0,1056 | 0,8232041 | chr5 | + |
| Grk4 | 131,74 | 0,1716 | 0,679461287 | chr5 | + |
| Grpel1 | 3500,57 | -0,1480 | 0,683952948 | chr5 | + |
| Grsf1 | 1896,95 | -0,3885 | 0,163596704 | chr5 | - |
| Gsap | 626,98 | 0,1057 | 0,843329386 | chr5 | + |
| Gtf2h3 | 256,13 | -0,1328 | 0,773303025 | chr5 | + |
| Gtf2i | 998,11 | -0,3139 | 0,186783528 | chr5 | - |
| Gtf2ird1 | 132,56 | -0,6049 | 0,155520806 | chr5 | - |
| Gtf3c2 | 488,36 | -0,2530 | 0,426630938 | chr5 | - |
| Gtpbp10 | 449,57 | 0,2322 | 0,598783534 | chr5 | - |
| Gtpbp6 | 81,59 | -0,3027 | 0,663086212 | chr5 | - |
| Guf1 | 332,82 | 0,0831 | 0,863829745 | chr5 | + |
| Gusb | 496,41 | -0,3043 | 0,336411059 | chr5 | - |
| Hadha | 5267,17 | -0,1493 | 0,716275354 | chr5 | - |
| Hadhb | 6868,83 | 0,0708 | 0,885409325 | chr5 | + |
| Haus3 | 124,89 | -0,0910 | 0,852862959 | chr5 | - |
| Hcar2 | 12,61 | 0,3028 | 0,6841486 | chr5 | - |
| Dnaaf5 | 68,97 | -0,2002 | 0,672205411 | chr5 | + |
| Helq | 51,40 | 0,1641 | 0,766748847 | chr5 | - |
| Hgfac | 2530,37 | 0,2481 | 0,581990132 | chr5 | + |
| Hip1 | 91,72 | 0,1638 | 0,753582185 | chr5 | - |
| Hip1r | 257,08 | -0,4309 | 0,428919534 | chr5 | + |
| Hmgb1 | 3840,45 | -0,0511 | 0,912626431 | chr5 | - |
| Hnf1a | 172,03 | -0,1382 | 0,807871939 | chr5 | - |
| Hnrnpd | 549,65 | 0,0326 | 0,957318111 | chr5 | - |
| Hnrnpdl | 886,28 | -0,1340 | 0,695978671 | chr5 | - |
| Hps4 | 127,85 | -0,2359 | 0,702094304 | chr5 | + |
| Hs3st1 | 2,23 | 0,2449 | NA | chr5 | - |
| Hscb | 54,86 | 0,0222 | 0,9774196 | chr5 | - |

| | | | | | |
|---|---|---|---|---|---|
| Hsd17b11 | 3789,76 | -0,4315 | 0,186118652 | chr5 | - |
| Hsd17b13 | 11431,72 | -0,0547 | 0,877920448 | chr5 | - |
| Hspb8 | 2636,13 | -0,4885 | 0,060685262 | chr5 | - |
| Htra3 | 15,53 | 0,0842 | 0,929225806 | chr5 | - |
| Htt | 262,08 | 0,1768 | 0,656897388 | chr5 | + |
| Hvcn1 | 25,54 | 0,7577 | 0,114525976 | chr5 | + |
| Idua | 63,46 | -0,5177 | 0,210808905 | chr5 | + |
| Ift172 | 238,46 | -0,0496 | 0,916852736 | chr5 | - |
| Ift22 | 63,40 | 0,1301 | 0,82790817 | chr5 | + |
| Ift81 | 64,90 | 0,1308 | 0,836754446 | chr5 | - |
| Igfbp7 | 1018,10 | 0,0959 | 0,837622669 | chr5 | - |
| Jchain | 43,31 | 0,7443 | 0,0686876 | chr5 | - |
| Insig1 | 7022,70 | -0,1923 | 0,636260727 | chr5 | + |
| Ints1 | 180,57 | 0,1982 | 0,63959323 | chr5 | - |
| Iqce | 49,98 | 0,4544 | 0,327583115 | chr5 | - |
| Katnal1 | 69,31 | 0,3322 | 0,430805668 | chr5 | - |
| Kcnk3 | 7,54 | 0,1113 | 0,905573242 | chr5 | + |
| Kctd10 | 216,55 | 0,1650 | 0,66477796 | chr5 | - |
| Kctd7 | 43,78 | 0,3411 | 0,515706574 | chr5 | + |
| Kdelr2 | 1618,07 | -0,4999 | 0,105630018 | chr5 | + |
| Kdm2b | 102,14 | 0,5524 | 0,152264308 | chr5 | - |
| Kdr | 459,07 | 0,1427 | 0,7242192 | chr5 | - |
| Khk | 8052,93 | 0,2499 | 0,618723125 | chr5 | + |
| Kit | 80,35 | -0,1418 | 0,834675238 | chr5 | + |
| Klb | 383,66 | -0,1993 | 0,684461361 | chr5 | + |
| Klf3 | 196,55 | -0,2857 | 0,503626727 | chr5 | + |
| Klhl5 | 385,01 | 0,1429 | 0,764363417 | chr5 | + |
| Klhl7 | 229,46 | 0,1242 | 0,783422709 | chr5 | + |
| Klhl8 | 38,55 | 0,0790 | 0,910916841 | chr5 | - |
| Kmt2c | 271,92 | -0,1217 | 0,764363417 | chr5 | - |
| Kmt2e | 272,46 | 0,0725 | 0,866039607 | chr5 | + |
| Kntc1 | 3,93 | 0,8804 | 0,12746076 | chr5 | + |
| Krit1 | 253,56 | -0,0961 | 0,84892478 | chr5 | + |
| Lamtor4 | 1480,91 | -0,4804 | 0,088624902 | chr5 | + |
| Lap3 | 6606,91 | -0,2615 | 0,337783618 | chr5 | + |
| Lat2 | 11,65 | 0,1812 | 0,83304202 | chr5 | - |
| Lcorl | 104,64 | 0,2620 | 0,541345095 | chr5 | - |
| Ldb2 | 48,88 | 0,3238 | 0,512986719 | chr5 | - |
| Letm1 | 26,14 | -0,2613 | 0,709072206 | chr5 | - |
| Lgi2 | 2,71 | 0,3553 | 0,611365326 | chr5 | - |
| Lias | 711,66 | -0,2258 | 0,507139053 | chr5 | + |
| Limch1 | 16,57 | 0,1236 | 0,881755379 | chr5 | + |
| Limk1 | 23,96 | -0,3501 | 0,587879802 | chr5 | - |
| Lin54 | 105,56 | -0,1554 | 0,726344964 | chr5 | - |
| Lmbr1 | 89,95 | 0,0088 | 0,991035244 | chr5 | - |
| Lmtk2 | 151,53 | -0,1478 | 0,73216561 | chr5 | + |
| Lnx1 | 6,94 | 0,1564 | 0,862616891 | chr5 | - |
| Lnx2 | 181,83 | -0,0422 | 0,93114272 | chr5 | - |
| Lrpap1 | 1052,24 | -0,1184 | 0,754524942 | chr5 | - |
| Lrrc17 | 3,43 | -0,0397 | 0,969512928 | chr5 | + |

| | | | | | |
|---|---|---|---|---|---|
| Lrrc8b | 8,09 | -0,0430 | 0,968614696 | chr5 | + |
| Lrrc8c | 23,84 | 0,0839 | 0,914198498 | chr5 | + |
| Lrrc8d | 376,06 | 0,0826 | 0,866039607 | chr5 | + |
| Lrwd1 | 84,20 | -0,2525 | 0,604679897 | chr5 | - |
| Lyar | 143,75 | -0,3001 | 0,411552342 | chr5 | + |
| Mad1l1 | 86,64 | -0,4979 | 0,232192464 | chr5 | - |
| Maea | 316,13 | 0,2740 | 0,425926605 | chr5 | + |
| Mafk | 220,96 | -0,4180 | 0,283414267 | chr5 | + |
| Magi2 | 2,30 | 0,1089 | NA | chr5 | + |
| Man2b2 | 395,41 | 0,2565 | 0,577347444 | chr5 | - |
| Mapre3 | 217,30 | -0,4426 | 0,361080708 | chr5 | + |
| Mblac1 | 12,58 | 0,4250 | 0,532988272 | chr5 | + |
| Mcm7 | 204,87 | 0,0112 | 0,98507181 | chr5 | - |
| Mdh2 | 3027,95 | -0,3232 | 0,343634659 | chr5 | + |
| Med13l | 131,84 | 0,1787 | 0,715912422 | chr5 | + |
| Med28 | 450,94 | 0,0545 | 0,930082891 | chr5 | + |
| Medag | 6,43 | 0,1721 | 0,8463949 | chr5 | + |
| Mepce | 410,61 | -0,3600 | 0,324324312 | chr5 | - |
| Mfsd10 | 37,24 | -0,1802 | 0,7950804 | chr5 | - |
| Mfsd7a | 56,18 | -0,3211 | 0,496763475 | chr5 | - |
| Micall2 | 17,58 | 0,2958 | 0,663810122 | chr5 | - |
| Mir25 | 3,14 | -0,0153 | 0,988025306 | chr5 | - |
| Mir703 | 49,57 | -0,7883 | 0,070271273 | chr5 | - |
| Mlec | 2254,15 | 0,0757 | 0,88569455 | chr5 | - |
| Mlxip | 134,45 | -0,1948 | 0,679628788 | chr5 | + |
| Mlxipl | 2031,03 | -0,5018 | 0,296808586 | chr5 | + |
| Mmab | 274,62 | -0,0826 | 0,877942095 | chr5 | - |
| Mmp17 | 5,30 | 0,2512 | 0,760953636 | chr5 | + |
| Mob1b | 252,49 | 0,3685 | 0,381639797 | chr5 | + |
| Mphosph9 | 54,06 | -0,3798 | 0,395886749 | chr5 | - |
| Mrfap1 | 2889,73 | -0,2447 | 0,34455554 | chr5 | - |
| Mrpl1 | 851,65 | -0,1413 | 0,719668754 | chr5 | + |
| Mrpl33 | 1310,55 | -0,0330 | 0,947294154 | chr5 | + |
| Mrps17 | 845,99 | -0,1066 | 0,773303025 | chr5 | + |
| Mrps18c | 1132,75 | -0,1214 | 0,768682625 | chr5 | + |
| Msantd1 | 5,30 | -0,5059 | 0,463892619 | chr5 | + |
| Mterf1a | 19,44 | 0,0022 | 0,997549615 | chr5 | - |
| Mterf1b | 9,64 | 0,0762 | 0,939403124 | chr5 | + |
| Mtf2 | 161,42 | 0,3428 | 0,302469512 | chr5 | + |
| Mthfd2l | 12,51 | -0,1739 | 0,831313706 | chr5 | + |
| Mtif3 | 485,84 | -0,0830 | 0,842655851 | chr5 | - |
| Mtus2 | 1991,17 | -0,0833 | 0,809343129 | chr5 | + |
| Mvk | 173,84 | -0,1813 | 0,78476189 | chr5 | + |
| Myo18b | 3,96 | 0,5399 | 0,411295797 | chr5 | - |
| N4bp2 | 109,13 | 0,2512 | 0,561165258 | chr5 | + |
| N4bp2l1 | 665,88 | -0,2290 | 0,647914259 | chr5 | - |
| N4bp2l2 | 623,65 | 0,1709 | 0,629895923 | chr5 | - |
| Naa25 | 235,15 | -0,1911 | 0,600688917 | chr5 | + |
| Naaa | 53,21 | 0,2452 | 0,667113793 | chr5 | - |
| Napepld | 14,25 | -0,1093 | 0,895961198 | chr5 | - |

| | | | | | |
|---|---|---|---|---|---|
| Ncapg | 130,81 | -0,0756 | 0,900182984 | chr5 | + |
| Ncf1 | 85,92 | 0,3686 | 0,3698416 | chr5 | - |
| Ncor2 | 169,77 | -0,3175 | 0,570561082 | chr5 | - |
| Nelfa | 79,60 | -0,4397 | 0,266692824 | chr5 | - |
| Nfxl1 | 232,01 | 0,0173 | 0,980662284 | chr5 | - |
| Noa1 | 115,45 | -0,4828 | 0,128835828 | chr5 | - |
| Nom1 | 295,07 | -0,0014 | 0,997464504 | chr5 | + |
| Nop14 | 201,65 | -0,3090 | 0,31970856 | chr5 | - |
| Nos3 | 45,18 | 0,0779 | 0,900660395 | chr5 | + |
| Nrbp1 | 752,19 | -0,3464 | 0,152130292 | chr5 | + |
| Nsg1 | 10,64 | 0,5248 | 0,419610935 | chr5 | - |
| Nsun5 | 120,85 | -0,4858 | 0,294160783 | chr5 | + |
| Nsun7 | 4,11 | 0,3131 | 0,681552027 | chr5 | + |
| Nub1 | 492,64 | -0,0685 | 0,905147397 | chr5 | + |
| Nudt1 | 450,80 | 0,1237 | 0,789064438 | chr5 | + |
| Nudt9 | 494,69 | -0,1018 | 0,813323745 | chr5 | + |
| Nup54 | 498,09 | 0,1892 | 0,640705696 | chr5 | - |
| Nupl2 | 50,96 | 0,1479 | 0,820554805 | chr5 | + |
| Nwd2 | 6,93 | 0,2819 | 0,725810792 | chr5 | + |
| Oas1b | 23,82 | -0,3402 | 0,572535452 | chr5 | + |
| Oas1c | 41,64 | 0,2416 | 0,715025856 | chr5 | - |
| Oas1g | 28,46 | 0,8740 | 0,083510154 | chr5 | - |
| Oas2 | 49,12 | 0,2081 | 0,75616611 | chr5 | - |
| Oas3 | 44,61 | 0,2925 | 0,584032361 | chr5 | - |
| Oasl2 | 97,21 | 0,4945 | 0,308762404 | chr5 | + |
| Ociad1 | 2015,51 | -0,1401 | 0,610189036 | chr5 | + |
| Ociad2 | 1200,65 | 0,1037 | 0,788428537 | chr5 | - |
| Ogfod2 | 324,24 | -0,5663 | 0,145168891 | chr5 | + |
| Orai1 | 105,06 | -0,7014 | 0,073903027 | chr5 | + |
| Orai2 | 17,84 | 0,3874 | 0,564495903 | chr5 | - |
| Orc5 | 405,56 | 0,2156 | 0,691018693 | chr5 | - |
| Ost4 | 2457,78 | -0,1149 | 0,762940356 | chr5 | - |
| P2rx4 | 1271,89 | -0,0002 | 0,999351454 | chr5 | + |
| P2rx7 | 52,33 | 0,5696 | 0,208282312 | chr5 | + |
| Pacrgl | 51,56 | 0,5488 | 0,23475207 | chr5 | + |
| Paics | 7192,91 | 0,0765 | 0,847736143 | chr5 | + |
| Pan3 | 298,40 | -0,4708 | 0,10394222 | chr5 | + |
| Paqr3 | 24,91 | 0,3104 | 0,627875496 | chr5 | - |
| Paxip1 | 188,26 | -0,4019 | 0,236693921 | chr5 | - |
| Pcdh7 | 15,06 | 0,7992 | 0,165257806 | chr5 | + |
| Pcgf3 | 160,97 | -0,0234 | 0,970419105 | chr5 | + |
| Pclo | 3,22 | -0,4189 | 0,499958086 | chr5 | + |
| Pcolce | 125,14 | 0,3423 | 0,384370645 | chr5 | - |
| Pdap1 | 1006,78 | -0,2413 | 0,582441358 | chr5 | - |
| Pdgfa | 33,44 | 0,7403 | 0,155078765 | chr5 | - |
| Pdgfra | 80,96 | 0,5242 | 0,195819213 | chr5 | + |
| Pds5a | 343,99 | -0,2946 | 0,312993461 | chr5 | - |
| Pds5b | 369,68 | 0,2883 | 0,464991549 | chr5 | + |
| Pebp1 | 2067,38 | -0,5152 | 0,104006655 | chr5 | - |
| Pex1 | 357,30 | -0,0387 | 0,937853416 | chr5 | + |

| | | | | | |
|---|---|---|---|---|---|
| Pf4 | 22,46 | 0,6159 | 0,252603799 | chr5 | + |
| Pgam5 | 240,28 | -0,1469 | 0,789666733 | chr5 | - |
| Phkg1 | 77,42 | -0,7243 | 0,061356755 | chr5 | - |
| Phtf2 | 139,91 | 0,0265 | 0,964556444 | chr5 | - |
| Pi4k2b | 744,64 | -0,4306 | 0,121401911 | chr5 | + |
| Pigg | 171,80 | 0,1334 | 0,783422709 | chr5 | + |
| Pilra | 111,50 | 0,0840 | 0,887839974 | chr5 | - |
| Pilrb1 | 53,23 | 0,1396 | 0,812490734 | chr5 | - |
| Pilrb2 | 44,70 | 0,0999 | 0,889053514 | chr5 | - |
| Pisd | 493,54 | 0,1117 | 0,744313101 | chr5 | - |
| Pitpnb | 1106,03 | -0,0281 | 0,951898903 | chr5 | + |
| Pitpnm2 | 271,03 | -0,5229 | 0,258301555 | chr5 | - |
| Pkd2 | 197,07 | 0,3035 | 0,4545859 | chr5 | + |
| Plac8 | 526,36 | 0,5131 | 0,205482823 | chr5 | - |
| Plbd2 | 18,85 | 0,2905 | 0,661627141 | chr5 | - |
| Plcxd1 | 25,22 | 0,9948 | 0,065354125 | chr5 | + |
| Plod3 | 21,46 | 0,2518 | 0,732140841 | chr5 | + |
| Pmpcb | 2313,08 | -0,2908 | 0,224946844 | chr5 | + |
| Pms2 | 34,10 | 0,6790 | 0,149536902 | chr5 | + |
| Pole | 28,71 | 0,1263 | 0,864697132 | chr5 | + |
| Polr2b | 1081,93 | -0,0332 | 0,962591292 | chr5 | + |
| Polr2j | 522,97 | -0,0552 | 0,9024844 | chr5 | + |
| Pom121 | 125,65 | -0,3136 | 0,470730504 | chr5 | - |
| Pomp | 3331,94 | -0,3014 | 0,358653761 | chr5 | + |
| Pop7 | 74,57 | -0,0244 | 0,97351573 | chr5 | - |
| Por | 3845,38 | -0,6763 | 0,071914286 | chr5 | + |
| Ppargc1a | 205,39 | 0,5886 | 0,105299254 | chr5 | - |
| Ppat | 787,61 | 0,2616 | 0,607664696 | chr5 | - |
| Ppbp | 20,33 | 0,9115 | 0,067696785 | chr5 | + |
| Ppm1g | 594,19 | -0,4017 | 0,223366879 | chr5 | - |
| Ppp1cb | 2434,30 | -0,0260 | 0,959255399 | chr5 | + |
| Ppp1cc | 1452,30 | 0,0764 | 0,842502441 | chr5 | + |
| Ppp1r35 | 54,40 | -0,3365 | 0,613341495 | chr5 | + |
| Pptc7 | 420,75 | -0,1429 | 0,743124066 | chr5 | + |
| Preb | 1113,08 | -0,1951 | 0,596546314 | chr5 | - |
| Prkab1 | 283,98 | 0,1183 | 0,793239112 | chr5 | - |
| Prkag2 | 269,50 | -0,1236 | 0,842502441 | chr5 | - |
| Prkar1b | 5,31 | -0,0910 | 0,925382443 | chr5 | - |
| Prkrip1 | 105,70 | -0,3370 | 0,349589724 | chr5 | - |
| Prom1 | 23,98 | -0,0131 | 0,988512389 | chr5 | - |
| Prr14l | 241,76 | 0,1309 | 0,732140841 | chr5 | - |
| Psmc2 | 1884,57 | -0,1760 | 0,616591847 | chr5 | + |
| Psmg3 | 218,04 | -0,4391 | 0,16215998 | chr5 | - |
| Psph | 106,05 | -0,1189 | 0,830662365 | chr5 | - |
| Ptcd1 | 1757,11 | 0,3554 | 0,279939661 | chr5 | - |
| Ptpn11 | 1050,76 | -0,1824 | 0,532901541 | chr5 | - |
| Ptpn12 | 400,69 | 0,2306 | 0,511533558 | chr5 | - |
| Ptpn13 | 12,11 | 0,1723 | 0,8332117 | chr5 | + |
| Pus1 | 177,90 | -0,1374 | 0,750261829 | chr5 | - |
| Pus7 | 157,64 | -0,0740 | 0,89468939 | chr5 | - |

| | | | | | |
|---|---|---|---|---|---|
| Pxmp2 | 2233,69 | 0,0031 | 0,994676823 | chr5 | - |
| Pxn | 473,39 | 0,1662 | 0,727329727 | chr5 | + |
| Qdpr | 6329,16 | 0,1042 | 0,845971244 | chr5 | - |
| Rab28 | 265,41 | 0,1298 | 0,773900067 | chr5 | - |
| Rab35 | 203,75 | -0,4953 | 0,135623763 | chr5 | + |
| Rabgef1 | 248,79 | -0,3165 | 0,428679263 | chr5 | + |
| Rac1 | 2044,47 | -0,3082 | 0,239040051 | chr5 | - |
| Ran | 2037,08 | -0,3297 | 0,253708477 | chr5 | + |
| Rasa4 | 18,59 | -0,0468 | 0,958859839 | chr5 | + |
| Rasal1 | 4,12 | 0,8640 | 0,147364898 | chr5 | + |
| Rasgef1b | 323,36 | 0,4397 | 0,420273711 | chr5 | - |
| Rasl11a | 2,73 | 0,3545 | 0,613341495 | chr5 | + |
| Rasl11b | 32,39 | -0,2732 | 0,647295341 | chr5 | + |
| Rassf6 | 209,42 | 0,1583 | 0,710077924 | chr5 | - |
| Rbak | 53,33 | -0,0124 | 0,988414083 | chr5 | - |
| Rbks | 257,34 | -0,1561 | 0,656102596 | chr5 | - |
| Rbm19 | 124,31 | 0,2117 | 0,670839945 | chr5 | + |
| Rbm33 | 684,90 | -0,6587 | 0,118025078 | chr5 | + |
| Rbm47 | 748,01 | -0,2885 | 0,453418644 | chr5 | - |
| Rbm48 | 97,78 | 0,0252 | 0,969159464 | chr5 | - |
| Rbpj | 211,67 | 0,2013 | 0,658008155 | chr5 | + |
| Rchy1 | 563,14 | 0,0190 | 0,971895772 | chr5 | - |
| Reln | 645,05 | 0,0660 | 0,909774279 | chr5 | - |
| Rest | 53,30 | 0,8045 | 0,066645098 | chr5 | + |
| Rfc1 | 386,89 | -0,0336 | 0,953911274 | chr5 | - |
| Rfc2 | 408,81 | 0,2960 | 0,374589794 | chr5 | + |
| Rfc3 | 102,33 | -0,1912 | 0,71673612 | chr5 | - |
| Rfc5 | 147,03 | 0,4077 | 0,277532928 | chr5 | - |
| Rhbdd2 | 216,46 | -0,2744 | 0,404820274 | chr5 | + |
| Rheb | 841,51 | -0,1540 | 0,67080798 | chr5 | - |
| Rhof | 2,21 | -0,0526 | NA | chr5 | - |
| Rhoh | 213,45 | -0,2226 | 0,697885289 | chr5 | + |
| Rilpl1 | 37,93 | 0,1682 | 0,783875547 | chr5 | - |
| Rilpl2 | 114,93 | -0,0093 | 0,987350308 | chr5 | - |
| Rint1 | 270,10 | -0,0311 | 0,952188756 | chr5 | + |
| Rita1 | 39,86 | -0,0167 | 0,98507181 | chr5 | - |
| Rnf10 | 1574,38 | -0,4879 | 0,060685262 | chr5 | - |
| Rnf216 | 239,04 | -0,0331 | 0,953911274 | chr5 | - |
| Rnf32 | 5,61 | 0,2003 | 0,815253842 | chr5 | + |
| Rnf34 | 150,96 | 0,0395 | 0,937853416 | chr5 | + |
| Rnf4 | 1729,05 | 0,1016 | 0,800998921 | chr5 | + |
| Rnf6 | 725,73 | -0,0090 | 0,985212135 | chr5 | - |
| Rpap2 | 234,58 | -0,2176 | 0,572534156 | chr5 | + |
| Rpl5 | 10416,30 | -0,3846 | 0,188860008 | chr5 | + |
| Rpl6 | 6997,04 | -0,4226 | 0,030098531 | chr5 | + |
| Rpl9 | 6748,85 | -0,4281 | 0,077452906 | chr5 | - |
| Rsbn1l | 182,96 | 0,4409 | 0,175381472 | chr5 | - |
| Rsrc2 | 718,81 | 0,1903 | 0,629969991 | chr5 | - |
| Rundc3b | 6,54 | -0,3120 | 0,690557571 | chr5 | - |
| Sart3 | 145,10 | -0,3165 | 0,434164637 | chr5 | - |

| | | | | | |
|---|---|---|---|---|---|
| Sbds | 880,90 | -0,3344 | 0,325000508 | chr5 | - |
| Sbno1 | 898,59 | -0,1040 | 0,743124066 | chr5 | - |
| Scarb1 | 1410,51 | -0,2523 | 0,546240378 | chr5 | - |
| Scarb2 | 1884,27 | -0,0264 | 0,949479116 | chr5 | - |
| Scfd2 | 111,44 | -0,3026 | 0,502231951 | chr5 | - |
| Sdad1 | 221,92 | 0,0379 | 0,935762352 | chr5 | - |
| Sds | 2664,96 | -0,3287 | 0,583782269 | chr5 | + |
| Sdsl | 153,25 | -0,2648 | 0,745938846 | chr5 | - |
| Sec31a | 993,27 | -0,1561 | 0,717576273 | chr5 | - |
| Sel1l3 | 236,96 | -0,6212 | 0,176233509 | chr5 | - |
| Selplg | 32,59 | 0,8706 | 0,055355843 | chr5 | - |
| Sema3a | 4,46 | 0,0299 | 0,978280656 | chr5 | + |
| Sema3c | 12,61 | -0,3197 | 0,679434494 | chr5 | + |
| Sepsecs | 243,68 | 0,6314 | 0,054328884 | chr5 | - |
| Setd1b | 198,96 | -0,1914 | 0,764363417 | chr5 | + |
| Kmt5a | 370,65 | -0,2148 | 0,650939157 | chr5 | + |
| Sfswap | 117,42 | -0,3370 | 0,432040063 | chr5 | + |
| Sh2b2 | 17,11 | -0,2064 | 0,803874753 | chr5 | - |
| Sh2b3 | 80,49 | 0,0186 | 0,979212524 | chr5 | - |
| Sh3bp2 | 311,92 | 0,0621 | 0,906578208 | chr5 | + |
| Sh3tc1 | 52,92 | -0,0192 | 0,979212524 | chr5 | - |
| Shisa3 | 4,83 | 0,4150 | 0,571942598 | chr5 | + |
| Shroom3 | 54,26 | 0,1367 | 0,819026397 | chr5 | + |
| Sirt4 | 100,45 | -0,6438 | 0,053298639 | chr5 | - |
| Slain2 | 437,55 | -0,0337 | 0,943108135 | chr5 | + |
| Slbp | 283,57 | -0,0675 | 0,883898229 | chr5 | - |
| Slc10a6 | 5,54 | -0,6078 | 0,357779158 | chr5 | - |
| Slc12a9 | 91,72 | -0,4702 | 0,393529719 | chr5 | - |
| Slc15a4 | 359,77 | -0,1080 | 0,827346646 | chr5 | - |
| Slc25a40 | 56,33 | 0,2054 | 0,721172264 | chr5 | + |
| Slc26a1 | 1848,44 | 0,4073 | 0,22361789 | chr5 | - |
| Slc2a9 | 786,88 | 0,1211 | 0,7950804 | chr5 | - |
| Slc30a3 | 11,31 | -0,3290 | 0,667450862 | chr5 | - |
| Slc30a9 | 1198,96 | 0,1201 | 0,754524942 | chr5 | + |
| Slc34a2 | 1640,40 | -0,1075 | 0,783875547 | chr5 | + |
| Slc35f6 | 343,60 | 0,0025 | 0,996644671 | chr5 | + |
| Slc4a1ap | 175,93 | -0,0207 | 0,969159464 | chr5 | + |
| Slc4a2 | 332,33 | -0,2549 | 0,538042836 | chr5 | + |
| Slc4a4 | 771,43 | -0,2752 | 0,438615439 | chr5 | + |
| Slc5a1 | 21,27 | 0,4654 | 0,464736787 | chr5 | + |
| Slc5a6 | 205,19 | -0,6089 | 0,056327501 | chr5 | - |
| Slc7a1 | 8,43 | 0,3587 | 0,630728761 | chr5 | - |
| Slc8b1 | 1018,98 | -0,2875 | 0,563343568 | chr5 | + |
| Smarcd3 | 7,60 | -0,0042 | 0,996823488 | chr5 | - |
| Smim14 | 861,73 | -0,0718 | 0,883164594 | chr5 | - |
| Smim20 | 500,14 | -0,3924 | 0,176233509 | chr5 | + |
| Smurf1 | 177,65 | -0,2932 | 0,425926605 | chr5 | - |
| Snrnp35 | 154,99 | 0,3055 | 0,510887696 | chr5 | + |
| Snx17 | 189,63 | -0,2184 | 0,606429785 | chr5 | + |
| Snx8 | 205,33 | -0,0121 | 0,983887477 | chr5 | - |

| | | | | | |
|---|---|---|---|---|---|
| Sod3 | 299,61 | 0,3738 | 0,245392711 | chr5 | + |
| Sorcs2 | 4,82 | 0,7733 | 0,214347061 | chr5 | - |
| Sowahb | 65,43 | 0,0863 | 0,888763259 | chr5 | - |
| Sparcl1 | 144,02 | 0,6249 | 0,124192205 | chr5 | - |
| Sppl3 | 243,77 | -0,3243 | 0,350774851 | chr5 | + |
| Ssc4d | 55,34 | 0,4284 | 0,31302658 | chr5 | - |
| Srd5a3 | 176,19 | -0,2896 | 0,427134717 | chr5 | + |
| Sri | 718,14 | 0,3574 | 0,206273473 | chr5 | + |
| Srp72 | 2089,98 | 0,0050 | 0,991035244 | chr5 | + |
| Srpk2 | 272,17 | 0,4770 | 0,180910407 | chr5 | - |
| Srrd | 42,22 | -0,2918 | 0,587223505 | chr5 | - |
| Srrm4 | 45,77 | 0,3079 | 0,528512451 | chr5 | - |
| Srrm4os | 28,03 | -0,1907 | 0,80192274 | chr5 | + |
| Srrt | 386,57 | -0,3931 | 0,344664343 | chr5 | - |
| Srsf9 | 185,45 | -0,4416 | 0,139059991 | chr5 | + |
| Ssh1 | 40,25 | -0,0521 | 0,943108135 | chr5 | - |
| Stard13 | 224,97 | -0,1328 | 0,776478455 | chr5 | - |
| Stbd1 | 2359,51 | -0,3811 | 0,29741825 | chr5 | + |
| Steap2 | 10,08 | 0,4176 | 0,561283233 | chr5 | - |
| Steap1 | 152,74 | -0,3907 | 0,387925064 | chr5 | - |
| Stim2 | 309,37 | -0,4477 | 0,252168188 | chr5 | + |
| Stx18 | 211,52 | 0,0271 | 0,962657075 | chr5 | + |
| Stx1a | 5,78 | -0,2319 | 0,781989945 | chr5 | + |
| Stx2 | 149,28 | 0,5423 | 0,093542539 | chr5 | - |
| Styxl1 | 3,86 | -0,1381 | 0,869687361 | chr5 | - |
| Sult1d1 | 4161,19 | 0,4568 | 0,33288309 | chr5 | - |
| Sult1e1 | 9,99 | 0,7389 | 0,231528092 | chr5 | - |
| Sumf2 | 122,74 | 0,0058 | 0,992812238 | chr5 | + |
| Sun1 | 183,69 | 0,2010 | 0,609861423 | chr5 | + |
| Supt7l | 126,42 | 0,1272 | 0,815549254 | chr5 | - |
| Tacc3 | 24,78 | -0,0243 | 0,979212524 | chr5 | + |
| Tada2b | 302,59 | -0,3101 | 0,381639797 | chr5 | - |
| Taf6 | 185,71 | 0,0455 | 0,940206753 | chr5 | - |
| Taok3 | 415,48 | 0,0440 | 0,943346405 | chr5 | + |
| Tapt1 | 979,63 | -0,3254 | 0,332496065 | chr5 | - |
| Tbc1d1 | 105,28 | 0,1068 | 0,840322097 | chr5 | + |
| Tbc1d14 | 294,89 | -0,0032 | 0,995598861 | chr5 | - |
| Tbl2 | 219,13 | -0,3083 | 0,360147619 | chr5 | + |
| Tbx3 | 219,03 | -0,5374 | 0,238825032 | chr5 | + |
| Tbx5 | 21,37 | 0,1239 | 0,877920448 | chr5 | + |
| Tcf23 | 2,70 | 0,2449 | 0,75378365 | chr5 | + |
| Tchp | 25,09 | 0,5194 | 0,357345082 | chr5 | + |
| Tctn1 | 38,18 | 0,2995 | 0,572115465 | chr5 | - |
| Tctn2 | 20,37 | -0,6696 | 0,204851518 | chr5 | + |
| Tec | 306,37 | -0,2583 | 0,461369428 | chr5 | - |
| Tecpr1 | 51,47 | 0,3366 | 0,576013884 | chr5 | - |
| Tfip11 | 193,42 | -0,2639 | 0,55880551 | chr5 | + |
| Tfr2 | 4387,03 | -0,4127 | 0,140524285 | chr5 | + |
| Tgfbr3 | 196,58 | -0,0436 | 0,948200011 | chr5 | - |
| Thap6 | 130,39 | 0,1647 | 0,705549096 | chr5 | + |

| | | | | | |
|---|---|---|---|---|---|
| Tlr1 | 23,60 | 0,4818 | 0,432040063 | chr5 | - |
| Tmed2 | 2501,26 | -0,2152 | 0,470905336 | chr5 | + |
| Tmed5 | 2421,86 | 0,0150 | 0,98489528 | chr5 | - |
| Tmem119 | 25,03 | 0,6700 | 0,216064568 | chr5 | - |
| Tmem120a | 836,42 | -0,1562 | 0,677299626 | chr5 | - |
| Tmem120b | 49,10 | -0,2106 | 0,692167613 | chr5 | + |
| Tmem128 | 457,36 | 0,0184 | 0,970755708 | chr5 | + |
| Tmem129 | 305,16 | -0,2817 | 0,361971351 | chr5 | - |
| Tmem165 | 177,54 | 0,2238 | 0,590058791 | chr5 | + |
| Tmem175 | 132,47 | 0,3045 | 0,48643419 | chr5 | + |
| Tmem184a | 105,13 | 0,2916 | 0,535573923 | chr5 | - |
| Tmem214 | 836,13 | -0,1607 | 0,71673612 | chr5 | + |
| Tmem243 | 1182,80 | -0,2890 | 0,266466163 | chr5 | + |
| Tmem248 | 898,93 | -0,5071 | 0,074602744 | chr5 | + |
| Tmem33 | 1769,58 | -0,1006 | 0,817848245 | chr5 | + |
| Tmem60 | 380,38 | 0,2363 | 0,537837216 | chr5 | + |
| Tmub1 | 82,01 | -0,5754 | 0,268073864 | chr5 | - |
| Tnip2 | 61,10 | -0,1783 | 0,778357029 | chr5 | - |
| Tnrc18 | 127,45 | 0,2482 | 0,67080798 | chr5 | - |
| Tomm7 | 1949,07 | -0,2734 | 0,345583636 | chr5 | - |
| Tpcn1 | 409,73 | -0,2738 | 0,507587541 | chr5 | - |
| Tpst1 | 361,83 | -0,3349 | 0,210808905 | chr5 | + |
| Tpst2 | 276,85 | -0,0355 | 0,953911274 | chr5 | + |
| Trafd1 | 492,18 | 0,1830 | 0,568266273 | chr5 | - |
| Triap1 | 305,07 | -0,0301 | 0,953911274 | chr5 | + |
| Trip6 | 171,32 | -0,0040 | 0,995735297 | chr5 | - |
| Trmt44 | 27,21 | -0,2821 | 0,634896352 | chr5 | - |
| Trpv4 | 5,20 | 0,1299 | 0,88569455 | chr5 | - |
| Trrap | 250,26 | -0,3610 | 0,39432036 | chr5 | + |
| Tsc22d4 | 86,19 | 0,2448 | 0,693242675 | chr5 | + |
| Ttyh3 | 72,21 | -0,4084 | 0,484137651 | chr5 | - |
| Txk | 4,78 | -0,4960 | 0,476764512 | chr5 | - |
| Tyms | 208,73 | 0,1430 | 0,722057512 | chr5 | - |
| Tyw1 | 322,33 | -0,2932 | 0,311625736 | chr5 | + |
| Ubc | 2369,24 | -0,6532 | 0,067943879 | chr5 | - |
| Ube2k | 978,62 | -0,2673 | 0,321371459 | chr5 | + |
| Ube3b | 517,05 | -0,2101 | 0,551170533 | chr5 | + |
| Ube3c | 673,29 | -0,0619 | 0,888648505 | chr5 | + |
| Ubl3 | 729,77 | -0,5830 | 0,056273421 | chr5 | - |
| Ufsp1 | 35,29 | -0,0751 | 0,913617288 | chr5 | + |
| Ugt2a1 | 2,15 | 0,0367 | NA | chr5 | - |
| Ugt2a3 | 7280,45 | -0,4773 | 0,175736443 | chr5 | - |
| Ugt2b34 | 11261,88 | -0,5412 | 0,125541699 | chr5 | - |
| Ugt2b36 | 25394,75 | -0,5192 | 0,121845614 | chr5 | - |
| Ugt2b37 | 178,77 | 0,2698 | 0,740538349 | chr5 | - |
| Ugt2b38 | 945,36 | -0,3612 | 0,599464204 | chr5 | - |
| Ulk1 | 1223,00 | -0,1244 | 0,811137679 | chr5 | - |
| Unc119b | 482,86 | -0,0329 | 0,947612608 | chr5 | - |
| Upk3b | 20,56 | -0,3636 | 0,60905895 | chr5 | + |
| Urad | 493,35 | 0,3764 | 0,319311067 | chr5 | - |

| | | | | | |
|---|---|---|---|---|---|
| Uso1 | 1585,39 | -0,0965 | 0,830330662 | chr5 | + |
| Usp12 | 512,44 | 0,1222 | 0,755984922 | chr5 | - |
| Usp30 | 174,71 | -0,3520 | 0,268487435 | chr5 | + |
| Usp42 | 70,14 | 0,4219 | 0,285043491 | chr5 | - |
| Usp46 | 189,81 | -0,3845 | 0,303231314 | chr5 | - |
| Uspl1 | 424,96 | 0,1264 | 0,743401455 | chr5 | + |
| Utp3 | 560,46 | -0,3793 | 0,146432004 | chr5 | + |
| Uvssa | 165,62 | -0,1110 | 0,804110902 | chr5 | + |
| Vkorc1l1 | 1040,80 | -0,2119 | 0,470066395 | chr5 | + |
| Vps29 | 959,24 | 0,0958 | 0,811533354 | chr5 | + |
| Vps33a | 398,22 | -0,1830 | 0,592406077 | chr5 | - |
| Vps37b | 145,67 | -0,1584 | 0,842320388 | chr5 | - |
| Vps37d | 2,41 | 0,0060 | NA | chr5 | - |
| Rcc1l | 189,13 | -0,5519 | 0,07840338 | chr5 | - |
| Bud23 | 326,44 | -0,3316 | 0,372630598 | chr5 | - |
| Mettl27 | 197,15 | -0,0515 | 0,916852736 | chr5 | + |
| Wdfy3 | 470,08 | 0,0425 | 0,925508783 | chr5 | - |
| Wdr1 | 659,73 | -0,0008 | 0,998195551 | chr5 | - |
| Wdr19 | 29,30 | 0,3776 | 0,536603375 | chr5 | + |
| Wdr86 | 2,48 | -0,1289 | NA | chr5 | - |
| Wfs1 | 71,68 | -0,7050 | 0,119171672 | chr5 | - |
| Nsd2 | 327,04 | 0,1288 | 0,715232158 | chr5 | + |
| Wipi2 | 1009,37 | -0,2751 | 0,363128879 | chr5 | + |
| Wsb2 | 657,61 | 0,0184 | 0,969195493 | chr5 | + |
| Xrcc2 | 22,11 | 0,1889 | 0,790595509 | chr5 | - |
| Yes1 | 514,95 | 0,1230 | 0,806401686 | chr5 | + |
| Ythdc1 | 405,64 | 0,1769 | 0,562291762 | chr5 | + |
| Ywhag | 1330,84 | -0,1514 | 0,695575308 | chr5 | - |
| Ywhah | 816,59 | 0,1281 | 0,692167613 | chr5 | + |
| Zan | 2,30 | 0,7897 | NA | chr5 | - |
| Zbed5 | 56,12 | -0,5879 | 0,215949298 | chr5 | + |
| Zbtb49 | 11,35 | 0,3393 | 0,656897388 | chr5 | - |
| Zcchc4 | 34,68 | 0,6007 | 0,22384344 | chr5 | + |
| Zcchc8 | 196,87 | 0,0876 | 0,851868245 | chr5 | - |
| Zcwpw1 | 85,29 | -0,0295 | 0,963854553 | chr5 | + |
| Zdhhc4 | 160,16 | -0,3964 | 0,242830621 | chr5 | - |
| Zfand2a | 241,26 | 0,1006 | 0,839045368 | chr5 | - |
| Zfp11 | 24,48 | 0,8020 | 0,088689123 | chr5 | - |
| Zfp113 | 145,00 | 0,4593 | 0,228280218 | chr5 | - |
| Zfp12 | 189,36 | -0,2749 | 0,43326072 | chr5 | + |
| Zfp316 | 52,49 | 0,4936 | 0,258301555 | chr5 | - |
| Zfp326 | 202,55 | 0,0598 | 0,908176736 | chr5 | + |
| Zfp512 | 148,20 | 0,3998 | 0,22942307 | chr5 | + |
| Zfp513 | 97,77 | -0,0020 | 0,997464504 | chr5 | - |
| Zfp518b | 20,54 | 0,1460 | 0,840700058 | chr5 | - |
| Zfp644 | 280,19 | 0,1924 | 0,617869186 | chr5 | - |
| Zfp655 | 674,95 | 0,0299 | 0,947612608 | chr5 | + |
| Zfp664 | 553,93 | 0,3030 | 0,341552333 | chr5 | + |
| Zfp68 | 379,45 | -0,0359 | 0,948200011 | chr5 | - |
| Zfp932 | 287,77 | 0,4726 | 0,149536902 | chr5 | + |

| | | | | | |
|---|---|---|---|---|---|
| Zkscan1 | 732,64 | -0,2553 | 0,419474305 | chr5 | + |
| Zkscan14 | 141,35 | -0,3991 | 0,228941765 | chr5 | - |
| Zkscan5 | 191,70 | -0,0882 | 0,834675238 | chr5 | + |
| Znhit1 | 407,69 | -0,2712 | 0,648706509 | chr5 | - |
| Zscan21 | 309,84 | 0,0748 | 0,850684447 | chr5 | + |
| Zscan25 | 42,02 | 0,2203 | 0,732933983 | chr5 | + |
| 0610030E20Rik | 464,23 | 0,2479 | 0,486967566 | chr6 | + |
| 0610040F04Rik | 4,30 | 0,1608 | 0,850915997 | chr6 | - |
| Fmc1 | 513,59 | -0,1623 | 0,856738348 | chr6 | + |
| 1110019D14Rik | 82,26 | -0,0525 | 0,936825775 | chr6 | + |
| Lockd | 8,43 | -0,0485 | 0,96272356 | chr6 | + |
| 1600020E01Rik | 122,51 | -0,5075 | 0,211374848 | chr6 | + |
| 1700003E16Rik | 3,62 | 0,0357 | 0,973485679 | chr6 | + |
| Nat8f4 | 144,88 | 0,3864 | 0,380878258 | chr6 | - |
| 1700025N23Rik | 18,12 | -0,3159 | 0,618512306 | chr6 | - |
| 1700101I11Rik | 3,11 | 0,2363 | 0,765231448 | chr6 | - |
| 1810044D09Rik | 77,26 | -0,5484 | 0,144177606 | chr6 | + |
| 1810058I24Rik | 1254,80 | -0,4986 | 0,074312572 | chr6 | + |
| 2210408F21Rik | 234,68 | -0,2451 | 0,663934341 | chr6 | + |
| 2310001H17Rik | 541,84 | -0,3831 | 0,37274915 | chr6 | - |
| 2310040G24Rik | 111,10 | -0,1793 | 0,736822651 | chr6 | - |
| Mkrn2os | 565,48 | 0,3347 | 0,339967066 | chr6 | - |
| 2610001J05Rik | 716,95 | 0,3802 | 0,317561996 | chr6 | - |
| 2610306M01Rik | 41,54 | 0,4395 | 0,370887749 | chr6 | - |
| Smim10l1 | 1412,29 | -0,2492 | 0,464133699 | chr6 | + |
| 2810454H06Rik | 4,49 | 0,0479 | 0,962779295 | chr6 | + |
| 2810474O19Rik | 380,16 | 0,3420 | 0,354512892 | chr6 | + |
| 3110062M04Rik | 27,38 | -0,0176 | 0,985702623 | chr6 | - |
| 4732416N19Rik | 3,06 | 0,1900 | 0,815253842 | chr6 | + |
| 4921507P07Rik | 6,16 | -0,8528 | 0,159653662 | chr6 | - |
| Particl | 6,79 | 0,3492 | 0,646093772 | chr6 | + |
| 5430402O13Rik | 14,51 | -0,1185 | 0,888917253 | chr6 | + |
| 8430408G22Rik | 840,71 | -0,2571 | 0,751866455 | chr6 | + |
| Fam234b | 642,40 | -0,0713 | 0,895961198 | chr6 | + |
| 9130221F21Rik | 13,91 | 0,4907 | 0,459906177 | chr6 | - |
| 9330102E08Rik | 10,35 | -0,2229 | 0,788000186 | chr6 | - |
| 9330158H04Rik | 2,90 | -0,0941 | 0,918077233 | chr6 | - |
| 9430018G01Rik | 5,30 | 0,8648 | 0,154257003 | chr6 | + |
| Tigar | 178,60 | -0,2621 | 0,554786054 | chr6 | - |
| A2m | 8,92 | -0,5710 | 0,391907358 | chr6 | + |
| A430035B10Rik | 27,18 | -0,0983 | 0,893437979 | chr6 | - |
| A730020E08Rik | 2,60 | -0,2720 | 0,707267324 | chr6 | - |
| Aak1 | 238,06 | -0,1174 | 0,751866455 | chr6 | + |
| Abcc9 | 252,08 | 0,4336 | 0,278037339 | chr6 | - |
| Abtb1 | 334,07 | -0,4468 | 0,302168049 | chr6 | - |
| Sdhaf3 | 141,28 | -0,1216 | 0,796541442 | chr6 | + |
| Acrbp | 26,87 | -0,7523 | 0,130063774 | chr6 | + |
| Actg2 | 4,25 | -0,1479 | 0,8672888 | chr6 | - |
| Adck2 | 126,47 | -0,1338 | 0,795226131 | chr6 | + |
| Adcyap1r1 | 23,39 | 0,1759 | 0,828259032 | chr6 | + |

| | | | | | |
|---|---|---|---|---|---|
| Adipor2 | 12976,62 | 0,1723 | 0,631855011 | chr6 | - |
| Aebp2 | 357,04 | -0,0121 | 0,979051488 | chr6 | + |
| Agbl3 | 58,57 | -0,5015 | 0,23655468 | chr6 | + |
| Ahcyl2 | 302,03 | 0,0408 | 0,949479116 | chr6 | + |
| Akr1b10 | 49,72 | 0,3699 | 0,430881045 | chr6 | + |
| Akr1b7 | 19,25 | 0,6223 | 0,283983764 | chr6 | + |
| Akr1b8 | 84,99 | 0,4961 | 0,2044482 | chr6 | + |
| Akr1d1 | 4271,10 | 0,6048 | 0,196626688 | chr6 | + |
| Aldh1l1 | 15387,41 | -0,0641 | 0,879708396 | chr6 | + |
| Alms1 | 34,53 | -0,0777 | 0,912811384 | chr6 | + |
| Amn1 | 127,32 | -0,0935 | 0,88569455 | chr6 | - |
| Ankrd26 | 57,09 | 0,5982 | 0,143109251 | chr6 | - |
| Antxr1 | 28,63 | 0,3099 | 0,636260727 | chr6 | - |
| Anxa4 | 1476,18 | 0,1212 | 0,755059918 | chr6 | - |
| Aplf | 88,90 | -0,4586 | 0,317142472 | chr6 | - |
| Apobec1 | 439,41 | 0,1212 | 0,740715013 | chr6 | - |
| Apold1 | 21,49 | 0,3633 | 0,558841064 | chr6 | + |
| Aqp1 | 2489,67 | 0,2906 | 0,277293474 | chr6 | + |
| Arf5 | 616,45 | -0,2298 | 0,67322586 | chr6 | + |
| Arhgap25 | 28,35 | 0,4136 | 0,491043827 | chr6 | - |
| Arhgdib | 246,58 | 0,3875 | 0,206194284 | chr6 | - |
| Arhgef5 | 395,42 | 0,1966 | 0,564495903 | chr6 | + |
| Arl6ip5 | 528,36 | -0,0929 | 0,814156035 | chr6 | + |
| Arl8b | 1215,96 | -0,2322 | 0,477715768 | chr6 | + |
| Arntl2 | 10,12 | 0,1392 | 0,877920448 | chr6 | + |
| Arpc4 | 675,10 | -0,2291 | 0,581941669 | chr6 | + |
| Art4 | 94,23 | 0,4834 | 0,259023596 | chr6 | - |
| Asb4 | 8,77 | 0,5204 | 0,443005632 | chr6 | + |
| Asprv1 | 2,12 | 0,0689 | NA | chr6 | + |
| Ints13 | 366,10 | -0,1931 | 0,528082877 | chr6 | - |
| Atf7ip | 118,83 | 0,5406 | 0,104006655 | chr6 | + |
| Atg7 | 429,21 | -0,3141 | 0,246698486 | chr6 | + |
| Atn1 | 142,09 | -0,3298 | 0,569607313 | chr6 | - |
| Atoh8 | 155,53 | 0,1233 | 0,850353974 | chr6 | - |
| Atp6v1e1 | 2789,61 | -0,0883 | 0,804380135 | chr6 | - |
| Atp6v1f | 1490,78 | -0,2293 | 0,442222987 | chr6 | + |
| Aup1 | 1403,50 | -0,4719 | 0,211603645 | chr6 | + |
| Avl9 | 374,20 | 0,4048 | 0,139399836 | chr6 | + |
| B230319C09Rik | 2,54 | 0,2851 | NA | chr6 | + |
| Bmt2 | 189,10 | -0,0478 | 0,925898178 | chr6 | - |
| BC035044 | 17,24 | 0,5273 | 0,39766425 | chr6 | - |
| BC049715 | 3,11 | -0,2091 | 0,792116735 | chr6 | + |
| BC064078 | 16,31 | -0,0643 | 0,943305207 | chr6 | + |
| Bcl2l13 | 637,53 | -0,0528 | 0,908013895 | chr6 | + |
| Bet1 | 1370,94 | -0,4744 | 0,139956847 | chr6 | - |
| Bhlhe40 | 1344,80 | -0,1283 | 0,676268178 | chr6 | + |
| Bhlhe41 | 61,73 | 0,2672 | 0,720470311 | chr6 | - |
| Bicd1 | 3,65 | 0,9344 | 0,090116069 | chr6 | + |
| Bid | 277,88 | 0,0908 | 0,842655851 | chr6 | - |
| Bms1 | 326,08 | -0,1103 | 0,761732176 | chr6 | - |

| | | | | | |
|---|---|---|---|---|---|
| Bola3 | 540,06 | -0,1393 | 0,834856381 | chr6 | + |
| Bpgm | 333,00 | -0,1990 | 0,649305665 | chr6 | + |
| Braf | 333,13 | -0,2363 | 0,465508094 | chr6 | - |
| Brk1 | 1458,39 | -0,0676 | 0,863445279 | chr6 | + |
| Brpf1 | 111,12 | -0,1420 | 0,795441927 | chr6 | + |
| C1galt1 | 209,21 | 0,6022 | 0,123890107 | chr6 | + |
| C1ra | 3397,89 | -0,0744 | 0,838785235 | chr6 | + |
| C1rb | 3,51 | 0,2329 | 0,773900067 | chr6 | + |
| C1rl | 2820,16 | 0,4501 | 0,211230512 | chr6 | + |
| C1s2 | 2,97 | -0,1638 | 0,844618872 | chr6 | - |
| C2cd5 | 61,74 | 0,2164 | 0,668293319 | chr6 | - |
| C3ar1 | 93,70 | 0,3967 | 0,311625736 | chr6 | - |
| C87436 | 205,72 | -0,3990 | 0,196045303 | chr6 | + |
| Cacna1c | 2,21 | 0,2161 | NA | chr6 | - |
| Cadps2 | 188,03 | -0,5351 | 0,132442871 | chr6 | - |
| Cald1 | 3802,29 | 0,2528 | 0,345819394 | chr6 | + |
| Calu | 3049,21 | 0,3637 | 0,153520353 | chr6 | + |
| Camk1 | 160,36 | -0,0851 | 0,8763719 | chr6 | - |
| Cand2 | 11,02 | 0,9437 | 0,103473805 | chr6 | + |
| Capg | 40,55 | 0,1701 | 0,804418611 | chr6 | + |
| Caprin2 | 33,67 | 0,7320 | 0,145007441 | chr6 | - |
| Capza2 | 2113,76 | 0,1383 | 0,701091698 | chr6 | + |
| Casd1 | 560,21 | 0,1692 | 0,664862994 | chr6 | + |
| Casp2 | 116,02 | 0,5407 | 0,073430016 | chr6 | + |
| Cav1 | 61,86 | -0,2124 | 0,746932573 | chr6 | + |
| Cav2 | 132,72 | 0,4826 | 0,254473076 | chr6 | + |
| Ccdc126 | 30,21 | 0,2220 | 0,736302391 | chr6 | + |
| Vps50 | 266,09 | 0,0651 | 0,899463828 | chr6 | + |
| Ccdc136 | 3,75 | 0,9092 | 0,115359796 | chr6 | + |
| Ccdc142 | 28,74 | 0,1389 | 0,855976995 | chr6 | + |
| Ccdc174 | 163,20 | -0,1285 | 0,748107818 | chr6 | + |
| Cfap100 | 7,73 | 0,5275 | 0,441085355 | chr6 | - |
| Ccdc77 | 71,92 | -0,1428 | 0,773710065 | chr6 | - |
| Ccnd2 | 171,06 | 0,6302 | 0,073599915 | chr6 | - |
| Cct7 | 1698,05 | -0,1443 | 0,670191619 | chr6 | + |
| Cd207 | 21,44 | -0,2700 | 0,717188441 | chr6 | - |
| Cd27 | 5,10 | 0,3030 | 0,700075571 | chr6 | - |
| Cd4 | 42,23 | 0,6271 | 0,286306833 | chr6 | - |
| Cd69 | 10,38 | -0,1227 | 0,892779606 | chr6 | - |
| Cd8a | 5,49 | 0,6282 | 0,334804026 | chr6 | + |
| Cd8b1 | 6,11 | 0,1243 | 0,893437979 | chr6 | + |
| Cd9 | 494,79 | 0,5244 | 0,147367695 | chr6 | - |
| Cdca3 | 22,59 | -0,2772 | 0,67311371 | chr6 | + |
| Cdkn1b | 316,59 | 0,3686 | 0,291685903 | chr6 | + |
| Cecr2 | 274,25 | -0,4122 | 0,287543614 | chr6 | + |
| Hdhd5 | 279,74 | -0,3299 | 0,288929615 | chr6 | - |
| Cep41 | 46,13 | 0,6341 | 0,146938521 | chr6 | - |
| Cftr | 3,29 | -0,1892 | 0,811137679 | chr6 | + |
| Chchd3 | 2097,78 | -0,3601 | 0,284909439 | chr6 | - |
| Chchd4 | 351,41 | -0,5064 | 0,096593506 | chr6 | - |

| | | | | | |
|---|---|---|---|---|---|
| Chchd6 | 38,61 | -0,2756 | 0,676060413 | chr6 | - |
| Chd4 | 947,58 | -0,2634 | 0,466451498 | chr6 | + |
| Chmp3 | 1693,10 | -0,1458 | 0,664226022 | chr6 | + |
| Chn2 | 680,96 | -0,3290 | 0,380031362 | chr6 | + |
| Chst13 | 66,98 | 0,2006 | 0,702487344 | chr6 | - |
| Cidec | 29,37 | 0,4628 | 0,499820761 | chr6 | - |
| Clec12a | 171,15 | 0,3846 | 0,435708717 | chr6 | + |
| Clec1a | 52,76 | 0,1190 | 0,841579427 | chr6 | - |
| Clec1b | 304,06 | 0,1347 | 0,773900067 | chr6 | + |
| Clec2d | 3687,36 | -0,4206 | 0,301101755 | chr6 | + |
| Clec2h | 400,88 | -0,4220 | 0,469270232 | chr6 | + |
| Clec2i | 7,76 | 0,0865 | 0,93056206 | chr6 | + |
| Clec4a1 | 61,75 | 0,4014 | 0,369534447 | chr6 | + |
| Clec4a2 | 47,98 | 0,3615 | 0,432797927 | chr6 | + |
| Clec4a3 | 104,25 | 0,6351 | 0,050290244 | chr6 | + |
| Clec4b1 | 15,42 | 0,2537 | 0,733541536 | chr6 | + |
| Clec4d | 9,46 | -0,0218 | 0,984995389 | chr6 | + |
| Clec4e | 4,71 | 0,6019 | 0,346086865 | chr6 | - |
| Clec4f | 3580,26 | 0,0060 | 0,991035244 | chr6 | - |
| Clec4n | 133,70 | 0,3055 | 0,454695883 | chr6 | + |
| Clec5a | 6,33 | 0,7733 | 0,215519599 | chr6 | - |
| Clec9a | 43,23 | 0,0049 | 0,995735297 | chr6 | + |
| Clstn3 | 105,40 | 0,1149 | 0,874477409 | chr6 | - |
| Nat8f1 | 8020,43 | -0,0142 | 0,981931319 | chr6 | - |
| Nat8f2 | 19077,29 | 0,2743 | 0,626272947 | chr6 | - |
| Nat8f3 | 7,89 | -0,1220 | 0,895663283 | chr6 | - |
| Nat8f5 | 89,37 | 0,1194 | 0,896013441 | chr6 | - |
| Cnbp | 5961,76 | -0,2677 | 0,430392697 | chr6 | - |
| Cnot4 | 161,96 | -0,1039 | 0,809343129 | chr6 | - |
| Col1a2 | 341,54 | 0,5204 | 0,277831408 | chr6 | + |
| Col28a1 | 5,15 | 0,6210 | 0,320959814 | chr6 | - |
| Copg1 | 1106,53 | -0,0778 | 0,863048009 | chr6 | + |
| Copg2 | 589,87 | 0,1501 | 0,737668661 | chr6 | - |
| Cops7a | 655,29 | -0,2771 | 0,36312833 | chr6 | - |
| Cped1 | 729,62 | 0,2184 | 0,612061669 | chr6 | + |
| Crbn | 916,09 | 0,1945 | 0,641315797 | chr6 | - |
| Creb3l2 | 66,13 | 0,7945 | 0,084192891 | chr6 | - |
| Crebl2 | 633,84 | 0,1286 | 0,781281681 | chr6 | + |
| Creld1 | 482,72 | -0,4461 | 0,273018284 | chr6 | + |
| Csgalnact2 | 168,36 | 0,2408 | 0,553850707 | chr6 | - |
| Cttnbp2 | 3,79 | -0,5168 | 0,433302067 | chr6 | - |
| Cul1 | 1272,49 | -0,0601 | 0,869442758 | chr6 | + |
| Cxcl12 | 5149,65 | -0,0009 | 0,998121281 | chr6 | + |
| Cycs | 7625,50 | -0,2121 | 0,549054641 | chr6 | - |
| Cyp26b1 | 137,95 | 0,4190 | 0,523438189 | chr6 | - |
| D6Wsu163e | 184,95 | 0,1244 | 0,818573937 | chr6 | + |
| Dcp1b | 21,04 | 0,3334 | 0,604355559 | chr6 | + |
| Dctn1 | 186,03 | -0,0191 | 0,980481306 | chr6 | + |
| Ddx47 | 415,11 | -0,1544 | 0,678707677 | chr6 | + |
| Dennd2a | 37,45 | 0,3110 | 0,57626953 | chr6 | - |

| | | | | | |
|---|---|---|---|---|---|
| Dennd5b | 1353,98 | 0,1517 | 0,67322586 | chr6 | - |
| Dera | 1366,96 | 0,0084 | 0,988512389 | chr6 | + |
| Gsdme | 12,18 | 0,2319 | 0,774082864 | chr6 | - |
| Dguok | 149,72 | -0,1930 | 0,647914259 | chr6 | - |
| Dok1 | 23,82 | 0,1066 | 0,885326978 | chr6 | - |
| Dusp11 | 770,00 | -0,1494 | 0,711934905 | chr6 | - |
| Dusp16 | 615,66 | -0,0554 | 0,886098675 | chr6 | - |
| Dysf | 74,32 | -0,1650 | 0,7588734 | chr6 | + |
| E130112N10Rik | 20,24 | -0,4606 | 0,447848007 | chr6 | + |
| E230016M11Rik | 11,50 | 0,0750 | 0,938072754 | chr6 | + |
| E330009J07Rik | 7,85 | 0,5445 | 0,415605593 | chr6 | - |
| Edem1 | 2539,79 | 0,0148 | 0,969892522 | chr6 | + |
| Eefsec | 288,45 | -0,1557 | 0,74785575 | chr6 | - |
| Efcab12 | 5,80 | -0,0160 | 0,988414083 | chr6 | - |
| Efcc1 | 10,90 | 0,4008 | 0,569607313 | chr6 | + |
| Eif2ak3 | 215,90 | 0,0029 | 0,996823488 | chr6 | + |
| Eif4e3 | 161,84 | 0,7570 | 0,073845576 | chr6 | - |
| Elmod3 | 254,35 | -0,0588 | 0,925039538 | chr6 | - |
| Emc3 | 995,68 | -0,2719 | 0,283172973 | chr6 | - |
| Emg1 | 1049,15 | -0,2719 | 0,298176352 | chr6 | - |
| Eogt | 60,84 | 0,1456 | 0,809065986 | chr6 | - |
| Epha1 | 606,80 | 0,3173 | 0,30944949 | chr6 | - |
| Ephb6 | 13,76 | 0,6165 | 0,315855622 | chr6 | + |
| Eps8 | 26,52 | 0,2097 | 0,74469138 | chr6 | - |
| Erc1 | 336,40 | -0,2655 | 0,466932584 | chr6 | - |
| Ergic2 | 1440,77 | -0,2273 | 0,546076999 | chr6 | - |
| Erp27 | 4,55 | -0,0084 | 0,99298417 | chr6 | - |
| Etnk1 | 1281,51 | -0,2848 | 0,317387674 | chr6 | + |
| Etv6 | 215,33 | 0,0246 | 0,969130697 | chr6 | + |
| Eva1a | 1086,80 | -0,0683 | 0,91644885 | chr6 | + |
| Exoc4 | 188,86 | 0,3676 | 0,416233558 | chr6 | + |
| Exoc6b | 124,08 | -0,2569 | 0,516883654 | chr6 | - |
| Ezh2 | 77,01 | -0,0764 | 0,897877163 | chr6 | - |
| Tcaf1 | 28,61 | -0,0069 | 0,993445259 | chr6 | - |
| Tcaf2 | 17,96 | 0,4934 | 0,402148509 | chr6 | - |
| Fam136a | 1203,40 | -0,0304 | 0,948314012 | chr6 | + |
| Fam13a | 328,61 | 0,4495 | 0,40679401 | chr6 | - |
| Fam180a | 32,44 | -0,2685 | 0,691486347 | chr6 | - |
| Mindy4 | 44,62 | 0,2820 | 0,603127139 | chr6 | + |
| Washc2 | 689,01 | -0,0974 | 0,803118336 | chr6 | + |
| Fam221a | 5,48 | 0,2009 | 0,814990243 | chr6 | + |
| Fam3c | 1001,85 | 0,5405 | 0,220525617 | chr6 | - |
| Fam60a | 61,91 | 0,1962 | 0,74146296 | chr6 | - |
| Fancd2 | 4,21 | 0,6302 | 0,330697335 | chr6 | + |
| Fbln2 | 30,39 | 0,5083 | 0,355683319 | chr6 | + |
| Fbxl14 | 113,27 | -0,2201 | 0,607677923 | chr6 | + |
| Fgd5 | 102,57 | 0,1680 | 0,748324453 | chr6 | + |
| Fgfr1op2 | 676,33 | -0,2249 | 0,413323375 | chr6 | + |
| Fkbp14 | 99,99 | 0,1868 | 0,693803702 | chr6 | - |
| Fkbp4 | 2429,35 | -0,0619 | 0,88569455 | chr6 | - |

| | | | | | |
|---|---|---|---|---|---|
| Fkbp9 | 723,62 | -0,0925 | 0,797253751 | chr6 | + |
| Flnc | 2,85 | 0,2251 | 0,764363417 | chr6 | + |
| Foxj2 | 110,34 | -0,5440 | 0,201119788 | chr6 | + |
| Foxm1 | 7,08 | -0,2240 | 0,788139622 | chr6 | + |
| Foxp1 | 269,55 | -0,0876 | 0,8763719 | chr6 | - |
| Frmd4b | 874,50 | 0,3410 | 0,413323375 | chr6 | - |
| Gabarapl1 | 8315,71 | -0,2341 | 0,549054641 | chr6 | + |
| Gadd45a | 91,88 | 0,5164 | 0,39588956 | chr6 | - |
| Gapdh | 8279,58 | -0,2553 | 0,43326072 | chr6 | - |
| Gars | 732,11 | -0,3400 | 0,267016205 | chr6 | + |
| Gata2 | 8,31 | -0,3293 | 0,669923083 | chr6 | + |
| Gcc1 | 164,51 | 0,1860 | 0,63960646 | chr6 | - |
| Gcfc2 | 53,41 | 0,4026 | 0,372385255 | chr6 | + |
| Gdf3 | 5,79 | -0,1685 | 0,849477252 | chr6 | - |
| Gfpt1 | 371,22 | 0,5524 | 0,147429065 | chr6 | + |
| Ggcx | 793,01 | 0,1543 | 0,654788851 | chr6 | + |
| Gimap1 | 39,65 | 0,2327 | 0,690747349 | chr6 | + |
| Gimap3 | 23,81 | 0,0447 | 0,95728901 | chr6 | - |
| Gimap4 | 366,33 | 0,3503 | 0,204564497 | chr6 | + |
| Gimap5 | 21,49 | 0,4290 | 0,495277308 | chr6 | + |
| Gimap6 | 384,24 | 0,2172 | 0,58887292 | chr6 | - |
| Gimap7 | 10,40 | 0,6376 | 0,318113166 | chr6 | + |
| Gimap8 | 136,60 | 0,7134 | 0,050965769 | chr6 | + |
| Gimap9 | 287,63 | -0,0305 | 0,955803452 | chr6 | + |
| Glcci1 | 74,18 | -0,2174 | 0,649305665 | chr6 | + |
| Gm10069 | 17,62 | 0,2505 | 0,724187104 | chr6 | + |
| Nat8f6 | 3,20 | 0,2208 | 0,776478455 | chr6 | - |
| Gm15401 | 17,21 | 0,5342 | 0,375767749 | chr6 | + |
| Gm15706 | 5,95 | -0,2520 | 0,759631295 | chr6 | + |
| Umad1 | 299,27 | 0,2893 | 0,413323375 | chr6 | + |
| Gm19434 | 14,46 | 0,1295 | 0,88127014 | chr6 | + |
| Gm19757 | 35,82 | -0,0531 | 0,943064069 | chr6 | - |
| Gm20362 | 40,60 | 0,4272 | 0,426610147 | chr6 | + |
| Gm21284 | 2,16 | 0,4860 | NA | chr6 | + |
| Gm5577 | 2,64 | 0,3562 | 0,611578224 | chr6 | + |
| Gm6313 | 22,98 | -0,2220 | 0,757695155 | chr6 | - |
| Gm6614 | 2,72 | -0,6245 | 0,29741825 | chr6 | - |
| Gmcl1 | 299,54 | 0,4221 | 0,221733317 | chr6 | - |
| Gng11 | 169,28 | 0,1115 | 0,835335693 | chr6 | + |
| Gng12 | 940,32 | -0,1669 | 0,654588343 | chr6 | + |
| Gngt1 | 27,15 | 0,5421 | 0,395899551 | chr6 | + |
| Golt1b | 97,42 | -0,1321 | 0,800238939 | chr6 | + |
| Gpnmb | 6,46 | 0,8044 | 0,180626552 | chr6 | + |
| Gpr162 | 9,34 | -0,4040 | 0,580569339 | chr6 | - |
| Gpr19 | 51,87 | 0,1698 | 0,78476189 | chr6 | - |
| Gprin3 | 59,86 | 0,5385 | 0,250135777 | chr6 | - |
| Grcc10 | 1162,08 | -0,3834 | 0,610022568 | chr6 | - |
| Gstk1 | 3371,79 | -0,3104 | 0,229632803 | chr6 | + |
| Gt(ROSA)26Sor | 170,43 | -0,1770 | 0,693803702 | chr6 | - |
| Gucy2c | 4,41 | -0,2597 | 0,74829088 | chr6 | - |

| | | | | | |
|---|---|---|---|---|---|
| Gys2 | 6479,11 | -0,4561 | 0,220525617 | chr6 | - |
| H2afj | 130,67 | -0,4673 | 0,513577902 | chr6 | + |
| Hdac11 | 697,41 | -0,4620 | 0,142156924 | chr6 | + |
| Hebp1 | 7417,96 | 0,0303 | 0,950078169 | chr6 | - |
| Hepacam2 | 3,95 | 0,3477 | 0,646093772 | chr6 | - |
| Herc3 | 75,17 | -0,0775 | 0,917470917 | chr6 | + |
| Hibadh | 6200,32 | -0,2578 | 0,352372205 | chr6 | - |
| Hilpda | 62,90 | 0,3540 | 0,431993683 | chr6 | + |
| Hipk2 | 40,50 | 0,0123 | 0,987025571 | chr6 | - |
| Hk2 | 22,86 | 0,3983 | 0,50154433 | chr6 | - |
| Hmces | 129,56 | 0,4059 | 0,323355495 | chr6 | + |
| Hnrnpa2b1 | 5612,34 | -0,0833 | 0,811093035 | chr6 | - |
| Hnrnpf | 3031,87 | 0,1301 | 0,712744391 | chr6 | + |
| Hoxa1 | 2,80 | 0,4120 | 0,554786054 | chr6 | - |
| Hoxa2 | 5,10 | 0,4048 | 0,58497677 | chr6 | - |
| Hoxa3 | 3,39 | -0,6518 | 0,293333978 | chr6 | - |
| Hoxa4 | 2,67 | 0,4848 | 0,44995478 | chr6 | - |
| Hoxa5 | 6,54 | -0,0096 | 0,992806079 | chr6 | - |
| Hpgds | 22,39 | 0,2536 | 0,734580366 | chr6 | - |
| Htra2 | 172,01 | 0,0456 | 0,939054301 | chr6 | - |
| Ica1 | 42,16 | -0,2580 | 0,629243929 | chr6 | - |
| Iffo1 | 54,52 | -0,3305 | 0,515706574 | chr6 | + |
| Ift122 | 65,22 | 0,3558 | 0,453845946 | chr6 | + |
| Il17ra | 263,64 | 0,0635 | 0,888648505 | chr6 | + |
| Il17rc | 222,25 | 0,0621 | 0,931322892 | chr6 | + |
| Il17re | 2,84 | -0,1000 | 0,912811384 | chr6 | + |
| Il5ra | 9,79 | 0,1177 | 0,895663283 | chr6 | - |
| Immt | 1853,64 | -0,2286 | 0,41552947 | chr6 | + |
| Impdh1 | 28,08 | -0,2153 | 0,767422402 | chr6 | - |
| Ing3 | 143,20 | 0,0309 | 0,955504626 | chr6 | + |
| Ing4 | 371,71 | -0,1678 | 0,691635748 | chr6 | + |
| Inmt | 16548,07 | -0,4570 | 0,411346711 | chr6 | - |
| Ino80b | 95,76 | -0,4286 | 0,477715768 | chr6 | - |
| Ipo8 | 766,06 | -0,0135 | 0,980154061 | chr6 | - |
| Iqsec1 | 411,58 | 0,2386 | 0,50757296 | chr6 | - |
| Irak2 | 396,57 | 0,0480 | 0,938514017 | chr6 | + |
| Irf5 | 168,94 | -0,0206 | 0,974008692 | chr6 | + |
| Isy1 | 225,57 | 0,0050 | 0,992089773 | chr6 | - |
| Itfg2 | 90,34 | -0,1966 | 0,703924654 | chr6 | - |
| Jagn1 | 491,42 | -0,3612 | 0,263412599 | chr6 | + |
| Jazf1 | 28,50 | -0,2561 | 0,721569914 | chr6 | - |
| Kbtbd12 | 7,98 | -0,1724 | 0,844029674 | chr6 | - |
| Kbtbd2 | 284,15 | 0,0432 | 0,938499724 | chr6 | - |
| Kbtbd8 | 59,41 | -0,0092 | 0,990115234 | chr6 | + |
| Kcnj8 | 45,32 | 0,2952 | 0,616552612 | chr6 | - |
| Kcp | 52,59 | -0,0300 | 0,974449819 | chr6 | - |
| Kdm3a | 333,85 | 0,1847 | 0,593645254 | chr6 | - |
| Kdm5a | 508,47 | 0,4677 | 0,15635499 | chr6 | + |
| Kdm7a | 347,88 | 0,2265 | 0,499958086 | chr6 | - |
| Klf15 | 1893,95 | -0,2990 | 0,515432158 | chr6 | + |

| | | | | | |
|---|---|---|---|---|---|
| Klhdc10 | 875,94 | -0,1707 | 0,652137083 | chr6 | + |
| Klhl42 | 254,74 | 0,1204 | 0,784505535 | chr6 | + |
| Klra17 | 3,29 | 0,3331 | 0,636260727 | chr6 | - |
| Klra2 | 59,34 | 0,6269 | 0,127555586 | chr6 | - |
| Klrb1c | 14,49 | 0,4154 | 0,523048388 | chr6 | - |
| Klrc1 | 11,01 | 0,2587 | 0,738282454 | chr6 | - |
| Klrd1 | 11,17 | 0,6719 | 0,293518157 | chr6 | + |
| Klre1 | 4,25 | 0,4981 | 0,466451498 | chr6 | + |
| Klri2 | 4,25 | 0,2346 | 0,767817776 | chr6 | - |
| Klrk1 | 185,81 | 0,0791 | 0,88127014 | chr6 | - |
| Kras | 499,42 | 0,1280 | 0,737568553 | chr6 | - |
| Krba1 | 77,65 | -0,5450 | 0,192260732 | chr6 | + |
| Krcc1 | 1023,73 | 0,1141 | 0,792116735 | chr6 | + |
| Lag3 | 32,83 | -0,2647 | 0,70880025 | chr6 | - |
| Lancl2 | 113,50 | 0,3175 | 0,466698801 | chr6 | + |
| P3h3 | 41,40 | 0,2675 | 0,624005041 | chr6 | - |
| Lmcd1 | 4,22 | -0,0262 | 0,981025197 | chr6 | + |
| Lmod2 | 4,38 | -0,6113 | 0,333257293 | chr6 | + |
| Lmod3 | 2,79 | 0,1532 | 0,854019408 | chr6 | - |
| Borcs5 | 84,64 | -0,5780 | 0,129193513 | chr6 | + |
| Loxl3 | 29,05 | 0,3426 | 0,577347444 | chr6 | + |
| Lrig1 | 337,62 | -0,1835 | 0,672907874 | chr6 | - |
| Lrmp | 41,20 | 0,0899 | 0,888763259 | chr6 | + |
| Lrp6 | 1069,10 | -0,0136 | 0,978402469 | chr6 | - |
| Lrrc61 | 109,67 | 0,4042 | 0,423149698 | chr6 | + |
| Lrtm2 | 2,65 | 0,9062 | 0,085785362 | chr6 | - |
| Lsm3 | 187,15 | 0,4087 | 0,212661076 | chr6 | + |
| Lsm5 | 258,52 | -0,0766 | 0,890887823 | chr6 | - |
| Lsm8 | 213,81 | 0,3273 | 0,445567031 | chr6 | + |
| Ltbr | 775,00 | -0,5522 | 0,07965206 | chr6 | - |
| Luc7l2 | 1667,76 | -0,0122 | 0,98489528 | chr6 | + |
| Etfrf1 | 1114,05 | 0,0408 | 0,945130695 | chr6 | + |
| M6pr | 2648,64 | 0,0859 | 0,804460185 | chr6 | + |
| Mad2l1 | 127,18 | -0,3411 | 0,38301652 | chr6 | + |
| Magi1 | 274,57 | -0,0767 | 0,863460985 | chr6 | - |
| Magohb | 74,45 | 0,5815 | 0,224157717 | chr6 | - |
| Malsu1 | 207,45 | -0,5813 | 0,050838368 | chr6 | + |
| March8 | 2317,24 | -0,1280 | 0,749603956 | chr6 | + |
| Mat2a | 1331,49 | -0,3060 | 0,357290285 | chr6 | - |
| Mbd4 | 27,30 | 0,8279 | 0,110745335 | chr6 | - |
| Mcm2 | 34,17 | -0,0560 | 0,950532633 | chr6 | - |
| Mdfic | 475,48 | 0,1096 | 0,797252833 | chr6 | + |
| Med21 | 640,23 | 0,0638 | 0,895594598 | chr6 | + |
| Mest | 39,32 | 0,4780 | 0,354338694 | chr6 | + |
| Met | 1341,25 | -0,3074 | 0,335347943 | chr6 | + |
| Etfbkmt | 1755,80 | -0,0585 | 0,924363403 | chr6 | + |
| Mgam | 1810,73 | 0,0163 | 0,976528475 | chr6 | + |
| Mgll | 2406,30 | -0,2115 | 0,674305684 | chr6 | + |
| Mgp | 129,24 | 0,6451 | 0,116363934 | chr6 | - |
| Mgst1 | 116995,16 | -0,3118 | 0,39588956 | chr6 | + |

| | | | | | |
|---|---|---|---|---|---|
| Mical3 | 150,40 | -0,2109 | 0,606429785 | chr6 | - |
| Mios | 226,71 | 0,0798 | 0,88333112 | chr6 | + |
| Mir8112 | 6,91 | 0,1833 | 0,830848779 | chr6 | + |
| Mitf | 65,33 | -0,2254 | 0,651354867 | chr6 | + |
| Mkln1 | 303,61 | -0,1737 | 0,698825001 | chr6 | + |
| Mkln1os | 1187,47 | -0,1795 | 0,561210905 | chr6 | - |
| Mkrn1 | 276,85 | 0,1057 | 0,817637582 | chr6 | - |
| Mkrn2 | 222,63 | 0,0528 | 0,913877005 | chr6 | + |
| Mlf2 | 1155,57 | -0,6042 | 0,098598649 | chr6 | + |
| Mmrn1 | 11,28 | -0,2967 | 0,691018693 | chr6 | + |
| Mogs | 528,90 | 0,1277 | 0,760028388 | chr6 | + |
| Mpp6 | 454,24 | 0,0008 | 0,998482911 | chr6 | + |
| Mrpl35 | 941,11 | -0,0698 | 0,869442758 | chr6 | - |
| Mrpl51 | 1412,63 | -0,3659 | 0,104006655 | chr6 | + |
| Mrpl53 | 640,36 | -0,3173 | 0,591091892 | chr6 | + |
| Mrps25 | 222,43 | -0,4324 | 0,197774355 | chr6 | - |
| Mrps33 | 1220,31 | 0,0934 | 0,820051729 | chr6 | - |
| Mrps35 | 397,64 | -0,2609 | 0,408202808 | chr6 | + |
| Mtmr14 | 372,65 | -0,2547 | 0,437265048 | chr6 | + |
| Mtpn | 1997,66 | -0,0973 | 0,7950804 | chr6 | - |
| Mug2 | 7259,33 | -0,3556 | 0,295255714 | chr6 | + |
| Mug-ps1 | 795,95 | -0,3961 | 0,462469544 | chr6 | + |
| Mxd1 | 109,53 | -0,2817 | 0,522768066 | chr6 | - |
| Nagk | 308,81 | 0,0525 | 0,891521879 | chr6 | + |
| Ncapd2 | 34,52 | -0,1586 | 0,81172751 | chr6 | - |
| Ndufa5 | 2384,10 | -0,3774 | 0,217849962 | chr6 | - |
| Ndufa9 | 3604,49 | -0,0885 | 0,807278226 | chr6 | - |
| Ndufb2 | 1189,44 | -0,2943 | 0,707267324 | chr6 | + |
| Necap1 | 740,28 | 0,2004 | 0,535026765 | chr6 | + |
| Nfe2l3 | 10,65 | 0,3408 | 0,656913078 | chr6 | + |
| Nfu1 | 646,52 | -0,0656 | 0,88569455 | chr6 | + |
| Nod1 | 104,24 | -0,0467 | 0,93807111 | chr6 | - |
| Nop2 | 239,21 | -0,2625 | 0,433302067 | chr6 | + |
| Npm3-ps1 | 4,56 | 0,0554 | 0,955328889 | chr6 | + |
| Nr2c2 | 569,81 | -0,0761 | 0,862421004 | chr6 | + |
| Nrf1 | 103,04 | -0,0555 | 0,917095291 | chr6 | + |
| Nrip2 | 3,82 | 0,4203 | 0,557606796 | chr6 | + |
| Nt5c3 | 1001,76 | 0,5089 | 0,068159818 | chr6 | - |
| Ntf3 | 8,75 | 0,3762 | 0,609072466 | chr6 | - |
| Nup205 | 247,84 | 0,2330 | 0,558841064 | chr6 | + |
| Ogg1 | 64,14 | 0,2040 | 0,680618596 | chr6 | + |
| Paip2b | 530,95 | 0,0622 | 0,889791056 | chr6 | - |
| Pcbp1 | 1238,89 | -0,3396 | 0,243000505 | chr6 | - |
| Pcgf1 | 109,67 | 0,0769 | 0,888964395 | chr6 | + |
| Pcyox1 | 2754,93 | 0,0169 | 0,965271905 | chr6 | - |
| Pde1c | 2,72 | 0,1132 | 0,898948254 | chr6 | - |
| Pde3a | 4,65 | 0,6286 | 0,336949509 | chr6 | + |
| Pdia4 | 3697,77 | 0,2759 | 0,49878857 | chr6 | - |
| Pdk4 | 410,13 | 0,4166 | 0,505514525 | chr6 | - |
| Pdzrn3 | 46,87 | 0,0510 | 0,945274006 | chr6 | - |

| | | | | | |
|---|---|---|---|---|---|
| Peg10 | 12,92 | -0,7134 | 0,245392711 | chr6 | + |
| Pex26 | 433,87 | -0,0823 | 0,87574113 | chr6 | + |
| Pex5 | 1304,48 | -0,3810 | 0,169413208 | chr6 | - |
| Phb2 | 1586,21 | -0,4370 | 0,119147532 | chr6 | + |
| Phc1 | 28,96 | 0,0874 | 0,91071941 | chr6 | - |
| Phf14 | 439,94 | -0,3155 | 0,357345082 | chr6 | + |
| Pianp | 2,19 | 0,0846 | NA | chr6 | + |
| Pik3c2g | 716,07 | -0,0840 | 0,8112244 | chr6 | + |
| Plbd1 | 699,40 | 0,1560 | 0,621474574 | chr6 | - |
| Plekha5 | 174,06 | -0,1181 | 0,813185941 | chr6 | + |
| Plekha8 | 319,01 | -0,4042 | 0,299239961 | chr6 | + |
| Plekhg6 | 92,31 | -0,0189 | 0,980767643 | chr6 | - |
| Plxna1 | 81,23 | 0,2237 | 0,681592614 | chr6 | - |
| Plxna4 | 31,05 | 0,1064 | 0,88569455 | chr6 | - |
| Plxna4os1 | 13,41 | 0,6070 | 0,33288309 | chr6 | + |
| Plxnd1 | 247,75 | -0,1587 | 0,724652712 | chr6 | - |
| Podxl | 146,12 | 0,1561 | 0,69847197 | chr6 | - |
| Podxl2 | 4,48 | -0,2516 | 0,760045267 | chr6 | - |
| Pole4 | 498,93 | -0,0034 | 0,995750189 | chr6 | - |
| Polr1a | 235,81 | 0,1866 | 0,678130673 | chr6 | + |
| Pon1 | 28512,73 | -0,0658 | 0,906333899 | chr6 | - |
| Pon2 | 1094,73 | -0,4500 | 0,072050071 | chr6 | - |
| Pon3 | 1484,58 | -0,1439 | 0,704334789 | chr6 | - |
| Ppfibp1 | 397,70 | 0,0672 | 0,903337956 | chr6 | + |
| Ppm1k | 2689,79 | 0,1296 | 0,783264136 | chr6 | - |
| Ppp4r2 | 736,67 | -0,0352 | 0,947294154 | chr6 | + |
| Pradc1 | 131,47 | -0,6617 | 0,051590044 | chr6 | - |
| Prdm5 | 2,21 | -0,0485 | NA | chr6 | + |
| Prickle2 | 129,96 | 0,3795 | 0,446054708 | chr6 | - |
| Prr15 | 3,64 | 0,2639 | 0,739141404 | chr6 | + |
| Ptcd3 | 996,26 | 0,0081 | 0,988512389 | chr6 | - |
| Ptms | 2536,88 | -0,3250 | 0,432005857 | chr6 | - |
| Ptpn6 | 269,81 | 0,0080 | 0,987508343 | chr6 | - |
| Ptpro | 14,53 | 0,8392 | 0,132914838 | chr6 | + |
| Pyroxd1 | 226,40 | 0,2235 | 0,615820434 | chr6 | + |
| Pyurf | 467,01 | -0,6041 | 0,141518635 | chr6 | - |
| Pzp | 105546,05 | -0,3364 | 0,286763659 | chr6 | - |
| Rab11fip5 | 13,07 | 0,5669 | 0,362631895 | chr6 | - |
| Rab19 | 8,32 | 0,3408 | 0,651354867 | chr6 | + |
| Rab43 | 2878,00 | 0,0059 | 0,991295673 | chr6 | - |
| Rab7 | 802,97 | 0,1147 | 0,816133189 | chr6 | - |
| Rad51ap1 | 5,34 | -0,1667 | 0,851158558 | chr6 | - |
| Rad52 | 68,11 | 0,0469 | 0,939403124 | chr6 | + |
| Raf1 | 1113,21 | -0,0620 | 0,869067375 | chr6 | - |
| Rarres2 | 17786,26 | 0,0636 | 0,885542101 | chr6 | - |
| Rassf4 | 92,15 | 0,1089 | 0,85281531 | chr6 | - |
| Rassf8 | 81,98 | 0,7180 | 0,089333342 | chr6 | + |
| Rbm28 | 392,52 | -0,0149 | 0,978280656 | chr6 | - |
| Recql | 106,87 | 0,1575 | 0,792197952 | chr6 | - |
| Reep1 | 14,38 | 0,2105 | 0,79002616 | chr6 | + |

| | | | | | |
|---|---|---|---|---|---|
| Repin1 | 439,66 | -0,2808 | 0,584970164 | chr6 | + |
| Rerg | 50,27 | 0,2898 | 0,607182269 | chr6 | - |
| Retsat | 6896,37 | -0,0738 | 0,909774279 | chr6 | + |
| Rhno1 | 54,74 | -0,0392 | 0,95728901 | chr6 | - |
| Rmnd5a | 2846,65 | 0,0375 | 0,939521529 | chr6 | - |
| Rnf103 | 1548,97 | -0,4760 | 0,127213067 | chr6 | + |
| Rnf181 | 1474,91 | -0,0170 | 0,965982099 | chr6 | - |
| Rpa3 | 213,16 | 0,1057 | 0,851158558 | chr6 | - |
| Rpia | 101,72 | 0,6291 | 0,078273639 | chr6 | - |
| Rpl32 | 5729,27 | -0,3692 | 0,135247543 | chr6 | - |
| Rpn1 | 2980,97 | -0,1210 | 0,77350788 | chr6 | + |
| Rps4l | 221,58 | 0,3488 | 0,596499464 | chr6 | + |
| Rpusd3 | 124,08 | -0,6622 | 0,180386456 | chr6 | - |
| Rtkn | 136,23 | 0,0331 | 0,968669888 | chr6 | + |
| Ruvbl1 | 126,51 | -0,1266 | 0,853027043 | chr6 | + |
| Rybp | 332,93 | -0,2812 | 0,357290285 | chr6 | - |
| Scnn1a | 280,60 | -0,0380 | 0,949479116 | chr6 | + |
| Sec13 | 1282,06 | -0,1734 | 0,627875496 | chr6 | - |
| Sec61a1 | 2824,67 | -0,2369 | 0,549054641 | chr6 | - |
| Sema4f | 8,38 | 0,4216 | 0,407448757 | chr6 | - |
| Serbp1 | 2515,90 | -0,0909 | 0,848794352 | chr6 | + |
| Setd5 | 330,57 | -0,1431 | 0,75379086 | chr6 | + |
| Setmar | 91,08 | 0,0484 | 0,939221429 | chr6 | + |
| Sfxn5 | 508,54 | 0,1466 | 0,701094188 | chr6 | - |
| Sgce | 143,51 | 0,3152 | 0,413323375 | chr6 | - |
| Shq1 | 60,95 | 0,1935 | 0,723157997 | chr6 | - |
| Skap2 | 551,65 | 0,0557 | 0,912010517 | chr6 | - |
| Slc13a4 | 7,53 | -0,7517 | 0,224157717 | chr6 | - |
| Slc25a13 | 3367,75 | -0,2091 | 0,582968859 | chr6 | - |
| Slc25a26 | 114,96 | -0,1360 | 0,772309251 | chr6 | + |
| Slc2a3 | 8,99 | 0,4044 | 0,576013884 | chr6 | - |
| Slc35b4 | 219,57 | -0,2518 | 0,441085355 | chr6 | - |
| Slc37a3 | 504,07 | 0,0009 | 0,998195551 | chr6 | - |
| Slc41a3 | 50,88 | 0,8135 | 0,068521927 | chr6 | + |
| Slc4a5 | 5,17 | 0,6139 | 0,352381684 | chr6 | + |
| Slc6a12 | 1436,00 | 0,2766 | 0,406787501 | chr6 | + |
| Slc6a13 | 4220,04 | -0,2159 | 0,535475471 | chr6 | + |
| Slc6a6 | 1131,04 | 0,1310 | 0,721177052 | chr6 | + |
| Slco1a4 | 1020,09 | -0,0331 | 0,969892522 | chr6 | - |
| Slco1b2 | 25947,74 | -0,3750 | 0,320810739 | chr6 | + |
| Smarcad1 | 235,01 | 0,3738 | 0,330122535 | chr6 | + |
| Smyd1 | 56,18 | -0,1503 | 0,836985758 | chr6 | - |
| Smyd5 | 120,80 | 0,0021 | 0,997464504 | chr6 | + |
| Snca | 12,43 | -0,1501 | 0,860606321 | chr6 | - |
| Snd1 | 1692,51 | -0,2604 | 0,438615439 | chr6 | + |
| Snora7a | 2,28 | -0,1326 | NA | chr6 | - |
| Snrnp27 | 487,33 | 0,2264 | 0,524801822 | chr6 | - |
| Snrpg | 771,69 | -0,1262 | 0,704789011 | chr6 | + |
| Snx10 | 1167,35 | 0,5027 | 0,136970896 | chr6 | + |
| Sox5 | 201,92 | 0,2868 | 0,592873282 | chr6 | - |

| | | | | | |
|---|---|---|---|---|---|
| Spr | 1174,24 | -0,4265 | 0,388402325 | chr6 | - |
| Spsb2 | 182,69 | 0,0382 | 0,952576171 | chr6 | + |
| Srgap3 | 7,60 | -0,2240 | 0,788691634 | chr6 | - |
| Ssbp1 | 795,78 | -0,0245 | 0,969159464 | chr6 | + |
| Sspn | 19,84 | 0,7523 | 0,176771112 | chr6 | + |
| Sspo | 9,21 | -0,0840 | 0,932414445 | chr6 | + |
| St3gal5 | 3366,69 | -0,4472 | 0,178097604 | chr6 | + |
| St7 | 124,50 | -0,1364 | 0,755748252 | chr6 | + |
| St8sia1 | 7,31 | 0,0251 | 0,981991128 | chr6 | - |
| Stambp | 178,72 | 0,0098 | 0,988512389 | chr6 | - |
| Stk38l | 429,86 | 0,1450 | 0,698557859 | chr6 | + |
| Strip2 | 2,72 | 0,2356 | 0,761178054 | chr6 | + |
| Suclg1 | 4681,52 | -0,3078 | 0,250303145 | chr6 | + |
| Suclg2 | 5175,51 | -0,1141 | 0,692773748 | chr6 | - |
| Sumf1 | 470,41 | -0,3239 | 0,370362371 | chr6 | - |
| Tac1 | 6,27 | 0,2211 | 0,584602378 | chr6 | + |
| Tada3 | 80,73 | -0,0076 | 0,992196557 | chr6 | - |
| Tamm41 | 36,85 | 0,0816 | 0,912811384 | chr6 | - |
| Tapbpl | 365,94 | 0,3333 | 0,394187455 | chr6 | - |
| Tax1bp1 | 3058,18 | -0,0532 | 0,89879469 | chr6 | + |
| Tbxas1 | 54,84 | 0,5102 | 0,262757002 | chr6 | + |
| Tcf7l1 | 42,76 | -0,5243 | 0,296117154 | chr6 | - |
| Tead4 | 4,78 | 0,6079 | 0,354512892 | chr6 | - |
| Tes | 54,32 | 0,0618 | 0,932531696 | chr6 | + |
| Tet3 | 245,60 | -0,1965 | 0,618414691 | chr6 | - |
| Tex261 | 590,83 | 0,1149 | 0,777924651 | chr6 | - |
| Tfec | 62,65 | 0,5024 | 0,190097808 | chr6 | - |
| Tfpi2 | 1534,05 | 0,2123 | 0,636361813 | chr6 | - |
| Tgfa | 199,94 | -0,2150 | 0,612061669 | chr6 | + |
| Tgoln1 | 2838,90 | -0,1534 | 0,723824089 | chr6 | - |
| Thnsl2 | 776,93 | 0,0259 | 0,953751362 | chr6 | - |
| Thsd7a | 5,09 | -0,0400 | 0,969892522 | chr6 | - |
| Thumpd3 | 374,18 | 0,1547 | 0,748511903 | chr6 | + |
| Tia1 | 102,58 | 0,0383 | 0,951806373 | chr6 | + |
| Tigd2 | 219,63 | -0,2724 | 0,409195505 | chr6 | + |
| Tm7sf3 | 1114,58 | 0,1353 | 0,744313101 | chr6 | - |
| Tmem106b | 3879,96 | 0,0062 | 0,990793693 | chr6 | + |
| Tmem139 | 2,14 | 0,5146 | NA | chr6 | + |
| Tmem140 | 707,00 | 0,4152 | 0,18299432 | chr6 | + |
| Tmem150a | 1045,71 | -0,4753 | 0,145049222 | chr6 | + |
| Tmem168 | 448,59 | 0,2205 | 0,543927518 | chr6 | - |
| Tmem176a | 3921,82 | 0,3989 | 0,210391102 | chr6 | + |
| Tmem176b | 4720,72 | 0,1698 | 0,718823184 | chr6 | - |
| Tmem229a | 20,07 | 0,7879 | 0,127722698 | chr6 | - |
| Tmem43 | 171,21 | 0,3839 | 0,19689523 | chr6 | + |
| Tmf1 | 436,59 | -0,3006 | 0,311651716 | chr6 | - |
| Tmsb10 | 266,98 | 0,5994 | 0,064991394 | chr6 | - |
| Tmtc1 | 14,93 | 0,1506 | 0,851076156 | chr6 | - |
| Tnfrsf1a | 955,16 | 0,4224 | 0,138850505 | chr6 | + |
| Tnip3 | 2,49 | 0,2336 | NA | chr6 | + |

| | | | | | |
|---|---|---|---|---|---|
| Tnpo3 | 787,21 | -0,2267 | 0,452275571 | chr6 | - |
| Tpi1 | 7719,84 | -0,3986 | 0,169324582 | chr6 | - |
| Tpk1 | 480,82 | 0,4829 | 0,164406774 | chr6 | - |
| Tpra1 | 113,20 | -0,3292 | 0,477326847 | chr6 | + |
| Tprkb | 3799,58 | -0,1930 | 0,636447281 | chr6 | + |
| Tril | 6,44 | -0,3817 | 0,611268902 | chr6 | - |
| Trnt1 | 389,95 | 0,1897 | 0,600032204 | chr6 | + |
| Tsen2 | 77,83 | 0,0725 | 0,911210861 | chr6 | + |
| Tspan12 | 3456,72 | -0,1412 | 0,748474371 | chr6 | - |
| Ttc26 | 199,09 | 0,0775 | 0,899148878 | chr6 | + |
| Ttll3 | 9,24 | -1,0047 | 0,079114437 | chr6 | + |
| Tuba8 | 39,36 | -0,0927 | 0,913936879 | chr6 | + |
| Tulp3 | 18,48 | 0,5402 | 0,33637288 | chr6 | - |
| Uba3 | 861,87 | 0,0942 | 0,840953217 | chr6 | - |
| Ube2h | 1587,52 | -0,2053 | 0,533104281 | chr6 | - |
| Ubn2 | 366,89 | -0,1574 | 0,764363417 | chr6 | + |
| Uroc1 | 5772,27 | 0,0333 | 0,935762352 | chr6 | + |
| Usp18 | 441,29 | 0,4236 | 0,210335339 | chr6 | + |
| Usp39 | 367,15 | -0,3271 | 0,383392162 | chr6 | - |
| Usp5 | 656,28 | -0,2202 | 0,443862531 | chr6 | - |
| Vamp1 | 38,12 | -0,5696 | 0,279939661 | chr6 | + |
| Vamp8 | 4549,73 | -0,0359 | 0,929973025 | chr6 | - |
| Vgll4 | 102,46 | 0,4761 | 0,336824399 | chr6 | - |
| Vhl | 265,71 | -0,2558 | 0,510887696 | chr6 | + |
| Vmn1r53 | 14,60 | 0,0236 | 0,981025197 | chr6 | - |
| Vopp1 | 95,60 | -0,3591 | 0,34745302 | chr6 | - |
| Vwf | 67,33 | 0,0534 | 0,943305207 | chr6 | + |
| Wbp1 | 489,27 | -0,2630 | 0,505514525 | chr6 | - |
| Wbp11 | 272,87 | 0,0723 | 0,876272552 | chr6 | - |
| Wdr54 | 3,40 | -0,1865 | 0,823537093 | chr6 | - |
| Wdr91 | 162,80 | -0,5504 | 0,140308933 | chr6 | - |
| Wipf3 | 123,64 | -0,0349 | 0,952076682 | chr6 | + |
| Wnk1 | 1064,90 | 0,0214 | 0,960688549 | chr6 | - |
| Wnt2 | 62,39 | -0,1545 | 0,81349332 | chr6 | - |
| Wnt5b | 82,74 | 0,6357 | 0,253908866 | chr6 | - |
| Xpc | 311,95 | 0,0340 | 0,94647679 | chr6 | - |
| Ybx3 | 262,38 | -0,2944 | 0,371556359 | chr6 | - |
| Zc3hav1 | 296,00 | 0,7069 | 0,086339532 | chr6 | - |
| Zc3hav1l | 9,01 | 0,4166 | 0,563374628 | chr6 | - |
| Zc3hc1 | 157,28 | -0,2216 | 0,570187994 | chr6 | - |
| Zfand4 | 394,95 | -0,0925 | 0,855976995 | chr6 | + |
| Zfp638 | 789,69 | 0,2863 | 0,407448757 | chr6 | + |
| Zfp212 | 93,34 | 0,2163 | 0,677398032 | chr6 | + |
| Zfp239 | 2,61 | 0,4531 | 0,484137651 | chr6 | + |
| Zfp248 | 13,23 | 0,1510 | 0,858112512 | chr6 | - |
| Zfp282 | 53,43 | 0,1724 | 0,764363417 | chr6 | + |
| Zfp384 | 276,02 | -0,2649 | 0,465196538 | chr6 | + |
| Zfp398 | 23,91 | 0,1285 | 0,863460985 | chr6 | + |
| Zfp422 | 265,77 | 0,3029 | 0,399722761 | chr6 | - |
| Zfp467 | 214,44 | -0,4074 | 0,2044482 | chr6 | - |

| | | | | | |
|---|---|---|---|---|---|
| Zfp637 | 201,85 | -0,3454 | 0,277831408 | chr6 | + |
| Zfp746 | 166,53 | -0,2094 | 0,580137509 | chr6 | - |
| Zfp775 | 27,81 | 0,0923 | 0,906383904 | chr6 | + |
| Zfp777 | 62,03 | -0,1694 | 0,768847925 | chr6 | - |
| Zfp783 | 1,98 | 0,3128 | NA | chr6 | + |
| Zfp786 | 3,13 | 0,4797 | 0,465841161 | chr6 | - |
| Zfp800 | 120,97 | 0,4354 | 0,247554818 | chr6 | - |
| Zfp862-ps | 29,12 | -0,0054 | 0,995598861 | chr6 | + |
| Zfp9 | 24,09 | -0,1794 | 0,803874753 | chr6 | - |
| Zfp956 | 41,15 | 0,0658 | 0,930684299 | chr6 | + |
| Rbsn | 208,62 | 0,2337 | 0,486771962 | chr6 | - |
| Znrf2 | 645,42 | 0,2664 | 0,402107622 | chr6 | + |
| Zxdc | 184,72 | -0,0083 | 0,9881871 | chr6 | + |
| Zyx | 296,79 | -0,2965 | 0,592428959 | chr6 | + |
| 1110004F10Rik | 722,93 | 0,0805 | 0,824866643 | chr7 | + |
| 1600010M07Rik | 8,06 | 0,1691 | 0,847419446 | chr7 | - |
| 1600014C10Rik | 3879,13 | 0,2200 | 0,449823999 | chr7 | + |
| Lmntd2 | 17,43 | -0,2490 | 0,755670612 | chr7 | - |
| 1700008J07Rik | 15,77 | 0,1036 | 0,899463828 | chr7 | - |
| 1700012D14Rik | 5,11 | 0,3827 | 0,611268902 | chr7 | - |
| 1700028J19Rik | 7,08 | 0,0666 | 0,947566465 | chr7 | - |
| Zfp974 | 43,06 | 0,3986 | 0,432005857 | chr7 | - |
| 1810008I18Rik | 1654,04 | 0,3904 | 0,476764512 | chr7 | + |
| 1810026B05Rik | 114,61 | -0,0511 | 0,925898178 | chr7 | - |
| 2200002D01Rik | 246,59 | -0,0111 | 0,988512389 | chr7 | - |
| Emsy | 101,27 | -0,1275 | 0,817265839 | chr7 | - |
| 2310010J17Rik | 56,33 | -0,3608 | 0,467424704 | chr7 | - |
| Rnf225 | 4,50 | -0,2070 | 0,808537363 | chr7 | + |
| 2310022A10Rik | 46,20 | -0,2020 | 0,709191439 | chr7 | + |
| 2310057M21Rik | 31,88 | -0,1348 | 0,834774375 | chr7 | - |
| 2410002F23Rik | 94,80 | -0,2023 | 0,649851928 | chr7 | + |
| Rexo5 | 39,27 | -0,0722 | 0,919961951 | chr7 | + |
| Arpin | 103,08 | 0,3128 | 0,538235194 | chr7 | - |
| Sult2a8 | 27171,31 | -0,2136 | 0,642313254 | chr7 | - |
| 2900076A07Rik | 39,32 | -0,4281 | 0,394162656 | chr7 | + |
| 3110040N11Rik | 229,45 | 0,0988 | 0,811137679 | chr7 | - |
| 4632427E13Rik | 2,46 | 0,0647 | NA | chr7 | - |
| Ddias | 16,09 | 0,6438 | 0,292182106 | chr7 | - |
| 4732471J01Rik | 8,52 | -0,7945 | 0,197796382 | chr7 | + |
| 4931406B18Rik | 5,09 | 0,2695 | 0,740663168 | chr7 | - |
| 4931406P16Rik | 175,89 | 0,0921 | 0,840322097 | chr7 | - |
| 4933406I18Rik | 5,22 | 0,3390 | 0,658755139 | chr7 | - |
| Zswim9 | 38,02 | 0,5989 | 0,213994613 | chr7 | - |
| 9030624J02Rik | 303,47 | -0,2069 | 0,62632688 | chr7 | + |
| 9130019O22Rik | 21,82 | 0,1305 | 0,863048009 | chr7 | - |
| 9130023H24Rik | 49,55 | 0,4207 | 0,401220967 | chr7 | - |
| 9130221H12Rik | 49,96 | -0,1551 | 0,784369718 | chr7 | + |
| 9430038I01Rik | 103,12 | -0,0128 | 0,987350308 | chr7 | - |
| Zfp976 | 67,75 | -0,3574 | 0,420538313 | chr7 | - |
| A230056P14Rik | 15,73 | 0,1096 | 0,896013441 | chr7 | + |

| | | | | | |
|---|---|---|---|---|---|
| A230077H06Rik | 3,29 | -0,3898 | 0,584288998 | chr7 | - |
| Aamdc | 641,42 | -0,1185 | 0,777753633 | chr7 | - |
| Abca14 | 5,24 | -0,1081 | 0,906333899 | chr7 | + |
| Abcc6 | 1250,53 | -0,1831 | 0,733944254 | chr7 | - |
| Abhd17c | 929,38 | -0,4130 | 0,119628533 | chr7 | - |
| Acadsb | 3133,21 | -0,0766 | 0,857773796 | chr7 | + |
| Acer3 | 337,43 | 0,1379 | 0,75436056 | chr7 | - |
| Acsm1 | 9043,63 | -0,3329 | 0,208518411 | chr7 | + |
| Acsm3 | 2831,54 | -0,4032 | 0,283855485 | chr7 | + |
| Acsm5 | 2437,44 | -0,0373 | 0,947819654 | chr7 | + |
| Actn4 | 1542,35 | -0,2110 | 0,585574166 | chr7 | - |
| Adam12 | 4,44 | 0,0360 | 0,973485679 | chr7 | - |
| Adam8 | 6,90 | 0,0865 | 0,93056206 | chr7 | - |
| Adamtsl3 | 2,21 | 0,4958 | NA | chr7 | + |
| Coq8b | 227,59 | 0,4551 | 0,289820918 | chr7 | + |
| Adm | 26,30 | -0,0294 | 0,974008692 | chr7 | + |
| Aen | 240,57 | -0,0959 | 0,830330662 | chr7 | + |
| AI314278 | 2,92 | 0,3042 | 0,683931222 | chr7 | + |
| AI467606 | 4,82 | 0,2622 | 0,741617215 | chr7 | + |
| AI987944 | 291,08 | 0,0732 | 0,88569455 | chr7 | - |
| Akap13 | 564,96 | -0,1668 | 0,711258571 | chr7 | + |
| Akip1 | 371,72 | -0,0555 | 0,92861245 | chr7 | + |
| Akt1s1 | 266,32 | -0,5857 | 0,273713146 | chr7 | + |
| Akt2 | 1285,39 | -0,1669 | 0,633994747 | chr7 | + |
| Aldh16a1 | 697,95 | -0,3041 | 0,586201597 | chr7 | - |
| Aldh1a3 | 11,06 | 0,5517 | 0,411158799 | chr7 | - |
| Aldoa | 1398,62 | -0,5023 | 0,128136319 | chr7 | - |
| Alg8 | 157,13 | -0,1524 | 0,755670612 | chr7 | + |
| Alkbh6 | 179,47 | 0,0150 | 0,981991128 | chr7 | + |
| Ampd3 | 13,89 | 0,4840 | 0,453418644 | chr7 | + |
| Anapc15 | 206,74 | -0,3548 | 0,299089342 | chr7 | + |
| Ankrd27 | 278,58 | -0,2048 | 0,587901189 | chr7 | + |
| Ankrd42 | 17,86 | 0,4332 | 0,513351366 | chr7 | - |
| Anks4b | 1307,98 | 0,2650 | 0,416860553 | chr7 | + |
| Ano1 | 25,44 | 0,5128 | 0,391701733 | chr7 | - |
| Ano5 | 4,52 | 0,5425 | 0,417918742 | chr7 | + |
| Anpep | 1610,58 | 0,1852 | 0,670684766 | chr7 | - |
| Ap2a1 | 216,07 | -0,3935 | 0,432040063 | chr7 | - |
| Ap2a2 | 1072,45 | -0,0682 | 0,872020846 | chr7 | + |
| Ap2s1 | 728,72 | -0,4984 | 0,23305528 | chr7 | + |
| Ap3s2 | 547,55 | 0,0598 | 0,89879469 | chr7 | - |
| Apba2 | 3,56 | 0,2859 | 0,703944394 | chr7 | + |
| Apbb1 | 11,37 | -0,1573 | 0,849606227 | chr7 | - |
| Aplp1 | 5,58 | -0,1307 | 0,886483612 | chr7 | - |
| Apobr | 39,00 | 0,5054 | 0,250762967 | chr7 | + |
| Apoc1 | 108463,87 | -0,2704 | 0,354665719 | chr7 | - |
| Apoc2 | 9384,57 | 0,3352 | 0,293840783 | chr7 | - |
| Apoc4 | 20741,83 | -0,3275 | 0,40070132 | chr7 | - |
| Apoe | 322227,79 | -0,3142 | 0,411104454 | chr7 | - |
| Aqp11 | 328,11 | -0,1688 | 0,599047998 | chr7 | - |

| | | | | | |
|---|---|---|---|---|---|
| Arap1 | 540,89 | 0,1085 | 0,804839321 | chr7 | + |
| Arfip2 | 170,48 | -0,5514 | 0,126676113 | chr7 | - |
| Arhgap17 | 116,13 | -0,1093 | 0,827378223 | chr7 | - |
| Arhgap33 | 3,03 | -0,1461 | 0,8636022 | chr7 | - |
| Arhgap35 | 1021,67 | -0,3334 | 0,268487435 | chr7 | - |
| Arhgef1 | 232,83 | -0,3682 | 0,496449986 | chr7 | + |
| Arhgef17 | 35,23 | 0,3781 | 0,469029897 | chr7 | - |
| Arl6ip1 | 4598,78 | -0,1929 | 0,610005889 | chr7 | - |
| Armc5 | 61,35 | -0,2149 | 0,765636461 | chr7 | + |
| Arntl | 27,65 | 0,3144 | 0,636381742 | chr7 | + |
| Arrb1 | 208,09 | 0,4584 | 0,134746323 | chr7 | + |
| Arrdc4 | 184,77 | -0,0384 | 0,950625589 | chr7 | - |
| Art2b | 3,60 | -0,0224 | 0,981991128 | chr7 | - |
| Asb7 | 155,89 | 0,1240 | 0,775026955 | chr7 | - |
| Aspdh | 1198,25 | -0,3978 | 0,390709604 | chr7 | + |
| Ate1 | 558,84 | 0,2844 | 0,408538653 | chr7 | - |
| Atf5 | 1647,44 | -0,5055 | 0,288486977 | chr7 | - |
| Atg16l2 | 129,85 | -0,2124 | 0,75379086 | chr7 | - |
| Pgghg | 392,90 | -0,1173 | 0,774324297 | chr7 | + |
| Atp10a | 7,28 | -0,1501 | 0,868151289 | chr7 | + |
| Atp2a1 | 2,37 | 0,3931 | NA | chr7 | - |
| Atp4a | 23,16 | 0,0942 | 0,921179542 | chr7 | + |
| Dmac2 | 443,03 | -0,4740 | 0,090472396 | chr7 | + |
| Atxn2l | 331,41 | -0,5381 | 0,298116176 | chr7 | - |
| AW146154 | 120,77 | 0,4387 | 0,332709151 | chr7 | - |
| Axl | 380,64 | 0,4688 | 0,098474583 | chr7 | - |
| B130024G19Rik | 10,76 | -0,2574 | 0,737374922 | chr7 | + |
| B3gnt8 | 3,68 | -0,1252 | 0,887839974 | chr7 | + |
| B9d2 | 81,38 | -0,4543 | 0,352372205 | chr7 | + |
| Bax | 327,46 | -0,4264 | 0,323328805 | chr7 | - |
| Bbc3 | 65,24 | 0,3558 | 0,631914678 | chr7 | + |
| BC017158 | 167,29 | -0,1625 | 0,738282454 | chr7 | - |
| BC024978 | 71,16 | -0,2259 | 0,691615955 | chr7 | + |
| BC030336 | 346,06 | 0,0306 | 0,951898903 | chr7 | + |
| Proser3 | 29,04 | -0,2697 | 0,677867285 | chr7 | - |
| Bc1 | 49,53 | -0,1506 | 0,814156035 | chr7 | + |
| Bcam | 37,13 | -0,1140 | 0,888917253 | chr7 | - |
| Bcat2 | 207,07 | -0,0167 | 0,98099384 | chr7 | + |
| Bccip | 1690,33 | -0,2302 | 0,430392697 | chr7 | + |
| Bckdha | 3384,31 | -0,6118 | 0,10394222 | chr7 | - |
| Bckdk | 143,57 | -0,0584 | 0,918077233 | chr7 | + |
| Bcl2l12 | 31,90 | -0,2084 | 0,753582185 | chr7 | - |
| Bcl3 | 91,35 | -0,9154 | 0,060685262 | chr7 | - |
| Bcl7c | 89,90 | -0,8739 | 0,062791791 | chr7 | - |
| Bet1l | 446,76 | -0,1984 | 0,47939388 | chr7 | - |
| Blm | 34,35 | 0,5490 | 0,331756559 | chr7 | - |
| Bloc1s3 | 73,05 | -0,1238 | 0,822394911 | chr7 | - |
| Blvrb | 1872,12 | -0,5281 | 0,252815255 | chr7 | + |
| Bnc1 | 2,94 | 0,4464 | 0,432704831 | chr7 | - |
| Bnip3 | 5394,47 | -0,2087 | 0,574724907 | chr7 | - |

| | | | | | |
|---|---|---|---|---|---|
| Bola2 | 401,91 | -0,5212 | 0,325000508 | chr7 | + |
| Btbd1 | 728,94 | -0,3405 | 0,25484255 | chr7 | - |
| Btbd10 | 42,90 | 0,0595 | 0,938514017 | chr7 | - |
| Btbd16 | 26,04 | -0,3601 | 0,518563051 | chr7 | + |
| Bub3 | 536,93 | 0,0188 | 0,967912274 | chr7 | + |
| Zfp607b | 10,48 | 0,4254 | 0,537545925 | chr7 | + |
| Faap24 | 29,67 | 0,3583 | 0,508698574 | chr7 | - |
| Snhg14 | 3,78 | 0,3021 | 0,692167613 | chr7 | - |
| C2cd3 | 122,33 | 0,1758 | 0,734502373 | chr7 | + |
| C5ar1 | 25,61 | 0,7358 | 0,152130292 | chr7 | - |
| C5ar2 | 48,32 | 0,5707 | 0,208577471 | chr7 | - |
| Calm3 | 757,42 | 0,4009 | 0,245990952 | chr7 | - |
| Capn5 | 34,74 | 0,4862 | 0,330697335 | chr7 | - |
| Capns1 | 1646,44 | -0,2520 | 0,378311758 | chr7 | - |
| Car11 | 7,78 | -0,5662 | 0,392238586 | chr7 | + |
| Cars | 371,40 | -0,0602 | 0,903337956 | chr7 | - |
| Catsperg1 | 2,42 | 0,5235 | NA | chr7 | - |
| Cblc | 264,36 | 0,3471 | 0,377944963 | chr7 | - |
| Sgf29 | 201,64 | -0,7255 | 0,052946934 | chr7 | + |
| Ccdc114 | 8,05 | 0,7028 | 0,268883626 | chr7 | + |
| Ccdc61 | 35,00 | -0,1762 | 0,826107735 | chr7 | - |
| Ccdc8 | 6,73 | 0,2129 | 0,803874753 | chr7 | + |
| Ccdc89 | 3,07 | 0,8541 | 0,126723342 | chr7 | + |
| Ccdc9 | 196,79 | -0,2336 | 0,579355496 | chr7 | - |
| Ccdc90b | 183,72 | 0,2020 | 0,620120518 | chr7 | + |
| Ccdc97 | 163,37 | -0,2519 | 0,541556317 | chr7 | - |
| Ccnd1 | 606,18 | 0,6696 | 0,141463153 | chr7 | - |
| Ccne1 | 63,90 | -0,1499 | 0,844155615 | chr7 | - |
| Ccp110 | 91,10 | -0,0318 | 0,959902994 | chr7 | + |
| Cd151 | 1231,79 | 0,2013 | 0,510735507 | chr7 | + |
| Cd177 | 3,25 | 0,9702 | 0,065752252 | chr7 | - |
| Cd19 | 7,87 | -0,2903 | 0,714765471 | chr7 | - |
| Cd22 | 20,61 | -0,4441 | 0,419942821 | chr7 | - |
| Cd2bp2 | 331,62 | -0,1454 | 0,720926726 | chr7 | - |
| Cd33 | 11,18 | 0,6075 | 0,341871878 | chr7 | - |
| Cd37 | 19,82 | 0,2802 | 0,666442376 | chr7 | - |
| Cd3eap | 79,23 | -0,1134 | 0,83176435 | chr7 | - |
| Cd79a | 10,40 | -0,0226 | 0,983780811 | chr7 | + |
| Cd81 | 3976,86 | -0,3871 | 0,165257806 | chr7 | + |
| Cdc42ep5 | 24,98 | -0,1258 | 0,869067375 | chr7 | - |
| Cdhr5 | 749,30 | -0,1148 | 0,803874753 | chr7 | - |
| Cdipt | 552,66 | -0,4363 | 0,10640307 | chr7 | + |
| Cdkn1c | 18,92 | -0,5725 | 0,378289895 | chr7 | - |
| Cdr2 | 14,56 | 0,4934 | 0,449778593 | chr7 | - |
| Ceacam1 | 1349,64 | 0,1572 | 0,683804325 | chr7 | - |
| Ceacam2 | 3,61 | 0,3547 | 0,630381755 | chr7 | - |
| Cebpa | 2620,12 | -0,4440 | 0,156475121 | chr7 | + |
| Cebpg | 1190,79 | -0,3204 | 0,290714093 | chr7 | - |
| Cend1 | 2,20 | 0,0815 | NA | chr7 | - |
| Cep89 | 94,86 | 0,3974 | 0,29741825 | chr7 | + |

| | | | | | |
|---|---|---|---|---|---|
| Chd2 | 572,11 | 0,3638 | 0,247432631 | chr7 | - |
| Chid1 | 983,47 | 0,0080 | 0,98867699 | chr7 | - |
| Chmp2a | 2158,43 | -0,1295 | 0,657471667 | chr7 | - |
| Chst15 | 113,48 | 0,1719 | 0,690557571 | chr7 | - |
| Chst8 | 1,87 | 0,2244 | NA | chr7 | - |
| Chsy1 | 67,89 | 0,1948 | 0,70880025 | chr7 | + |
| Cib1 | 1177,35 | -0,3651 | 0,319311067 | chr7 | - |
| Cic | 145,15 | -0,4314 | 0,401045574 | chr7 | + |
| Clasrp | 47,79 | -0,3574 | 0,547268381 | chr7 | - |
| Clcn4 | 1270,26 | -0,2660 | 0,328388243 | chr7 | - |
| Clec11a | 34,38 | 0,2256 | 0,731011847 | chr7 | - |
| Clip3 | 4,29 | 0,5657 | 0,386872469 | chr7 | + |
| Cln3 | 204,18 | -0,1477 | 0,712500177 | chr7 | - |
| Clptm1 | 861,71 | -0,4824 | 0,131175019 | chr7 | - |
| Cnot3 | 129,83 | -0,5848 | 0,22361789 | chr7 | + |
| Coa4 | 82,52 | 0,3794 | 0,369664628 | chr7 | + |
| Cog7 | 255,70 | -0,2270 | 0,49577637 | chr7 | - |
| Copb1 | 2383,22 | 0,0311 | 0,950795699 | chr7 | - |
| Coq7 | 409,23 | -0,2874 | 0,537078872 | chr7 | - |
| Coro1a | 100,81 | 0,1505 | 0,760045267 | chr7 | - |
| Cox6a2 | 2,09 | 0,1951 | NA | chr7 | - |
| Cox6b1 | 8942,14 | -0,3542 | 0,195669979 | chr7 | - |
| Cox6b2 | 18,20 | 0,9076 | 0,118025078 | chr7 | - |
| Cox7a1 | 47,39 | 0,0057 | 0,993991445 | chr7 | + |
| Cpt1c | 4,50 | 0,5505 | 0,410110272 | chr7 | - |
| Cpxm2 | 8,36 | 0,4991 | 0,474644284 | chr7 | - |
| Crebzf | 357,72 | 0,0434 | 0,930684299 | chr7 | + |
| Crtc3 | 116,13 | 0,1029 | 0,828836407 | chr7 | - |
| Crx | 2,89 | 0,2912 | 0,695093611 | chr7 | - |
| Ctbp2 | 30,49 | -0,0029 | 0,996987906 | chr7 | - |
| Ctf1 | 60,66 | 0,1266 | 0,827883692 | chr7 | + |
| Ctr9 | 430,50 | 0,0731 | 0,88569455 | chr7 | + |
| Ctsd | 7939,41 | -0,1064 | 0,69294371 | chr7 | - |
| Cttn | 710,52 | 0,2150 | 0,471089317 | chr7 | - |
| Ctu1 | 56,77 | -0,1557 | 0,773900067 | chr7 | + |
| Cyfip1 | 687,09 | 0,2114 | 0,553328477 | chr7 | + |
| Cyp2a12 | 11000,53 | 0,0390 | 0,920434357 | chr7 | + |
| Cyp2g1 | 17,91 | -0,4342 | 0,53490582 | chr7 | + |
| Cyp2s1 | 7,70 | 0,5997 | 0,367971555 | chr7 | - |
| Cyth2 | 204,43 | -0,4865 | 0,08440425 | chr7 | - |
| D430042O09Rik | 134,75 | -0,0962 | 0,840418613 | chr7 | + |
| Snhg14 | 24,96 | 0,3113 | 0,642318814 | chr7 | - |
| Dact3 | 3,36 | 0,0624 | 0,950099786 | chr7 | + |
| Dbp | 1520,51 | -0,2209 | 0,657774838 | chr7 | + |
| Dchs1 | 23,34 | 0,2416 | 0,740024898 | chr7 | - |
| Dctn5 | 604,60 | 0,2612 | 0,44820883 | chr7 | + |
| Dctpp1 | 746,98 | -0,3736 | 0,354876286 | chr7 | - |
| Dcun1d3 | 189,74 | -0,3517 | 0,272642718 | chr7 | - |
| Deaf1 | 112,40 | -0,4102 | 0,343806877 | chr7 | - |
| Dedd2 | 230,40 | -0,4503 | 0,332014851 | chr7 | - |

| | | | | | |
|---|---|---|---|---|---|
| Dennd5a | 455,62 | 0,1464 | 0,631914678 | chr7 | - |
| Dgat2 | 6655,01 | -0,1963 | 0,440917912 | chr7 | - |
| Dhcr7 | 947,35 | 0,1311 | 0,843329386 | chr7 | + |
| Dhdh | 1693,75 | 0,0329 | 0,947294154 | chr7 | - |
| Dhx32 | 338,58 | -0,3752 | 0,220562836 | chr7 | - |
| Dhx34 | 66,73 | 0,3913 | 0,394399177 | chr7 | - |
| Dkk3 | 38,89 | 0,7749 | 0,083730864 | chr7 | - |
| Dmpk | 134,30 | -0,2723 | 0,652658393 | chr7 | + |
| Dmwd | 140,05 | -0,2210 | 0,705982644 | chr7 | + |
| Dnajb13 | 4,35 | 0,5239 | 0,437620234 | chr7 | - |
| Dock1 | 192,06 | 0,0179 | 0,9774196 | chr7 | + |
| Dpysl4 | 2,48 | 0,2995 | NA | chr7 | + |
| Dusp8 | 17,13 | -0,3866 | 0,602175054 | chr7 | - |
| Dyrk1b | 56,45 | -0,4062 | 0,508268404 | chr7 | + |
| E030018B13Rik | 7,93 | -1,0110 | 0,077224204 | chr7 | + |
| E230029C05Rik | 8,52 | 0,1160 | 0,899497201 | chr7 | + |
| E2f8 | 57,88 | -0,4952 | 0,403302288 | chr7 | - |
| E430018J23Rik | 45,79 | 0,1389 | 0,816359983 | chr7 | - |
| Ears2 | 110,45 | 0,2667 | 0,528469822 | chr7 | - |
| Ech1 | 7600,98 | -0,2176 | 0,576231821 | chr7 | + |
| Echs1 | 4833,60 | 0,0465 | 0,899463828 | chr7 | - |
| Edrf1 | 327,83 | -0,3890 | 0,194537528 | chr7 | + |
| Eed | 438,45 | 0,1473 | 0,707968435 | chr7 | - |
| Eef2k | 176,93 | 0,3799 | 0,224157717 | chr7 | + |
| Cracr2b | 29,19 | -0,1005 | 0,88569455 | chr7 | + |
| Efl1 | 147,54 | -0,0854 | 0,854019408 | chr7 | + |
| Egln2 | 784,28 | -0,2169 | 0,737062808 | chr7 | - |
| Ehd2 | 40,44 | 0,6193 | 0,13969188 | chr7 | - |
| Eid2 | 8,10 | 0,2017 | 0,813415373 | chr7 | + |
| Eid2b | 35,88 | 0,0911 | 0,907201495 | chr7 | + |
| Eif3c | 2445,54 | -0,0566 | 0,86436738 | chr7 | - |
| Eif3f | 1043,64 | -0,5422 | 0,080590154 | chr7 | + |
| Eif3k | 1042,95 | -0,3883 | 0,19121789 | chr7 | - |
| Eif4g2 | 8455,81 | -0,2234 | 0,438201451 | chr7 | - |
| Emc10 | 473,45 | -0,4177 | 0,438263068 | chr7 | - |
| Eml2 | 109,94 | 0,5373 | 0,231030075 | chr7 | + |
| Emp3 | 72,63 | 0,4053 | 0,369147908 | chr7 | - |
| Epn1 | 876,35 | -0,6064 | 0,142312129 | chr7 | + |
| Eps8l1 | 21,04 | 0,4039 | 0,569607313 | chr7 | + |
| Eps8l2 | 490,54 | -0,1801 | 0,775308498 | chr7 | + |
| Ercc1 | 40,97 | -0,4917 | 0,310647489 | chr7 | + |
| Ercc2 | 97,50 | -0,3421 | 0,430881045 | chr7 | + |
| Erf | 208,15 | 0,0379 | 0,964620526 | chr7 | - |
| Eri2 | 113,59 | 0,5375 | 0,148471484 | chr7 | - |
| Etfb | 7401,32 | -0,2935 | 0,312157245 | chr7 | + |
| Ethe1 | 900,44 | 0,0061 | 0,993598948 | chr7 | + |
| Exosc5 | 251,06 | -0,3773 | 0,436827862 | chr7 | + |
| Fadd | 113,68 | 0,0246 | 0,967878224 | chr7 | - |
| Fah | 10430,10 | -0,2618 | 0,370803111 | chr7 | - |
| Fam103a1 | 975,65 | 0,2454 | 0,546712084 | chr7 | + |

| | | | | | |
|---|---|---|---|---|---|
| Saxo2 | 4,17 | -0,0588 | 0,951934337 | chr7 | - |
| Fam160a2 | 224,08 | -0,2606 | 0,468217847 | chr7 | - |
| Fam168a | 453,94 | -0,1657 | 0,648165583 | chr7 | + |
| Fam169b | 367,93 | -0,0422 | 0,93731052 | chr7 | + |
| Fam174b | 66,59 | -0,5605 | 0,151287224 | chr7 | + |
| Abraxas2 | 495,19 | -0,0484 | 0,912136456 | chr7 | + |
| Fam187b | 87,74 | 0,3484 | 0,448256963 | chr7 | + |
| Fam189a1 | 3,08 | 0,3560 | 0,596219566 | chr7 | - |
| Fam71e1 | 8,75 | -0,3736 | 0,619691963 | chr7 | + |
| Fan1 | 51,19 | 0,4552 | 0,33288309 | chr7 | - |
| Fancf | 23,30 | -0,1530 | 0,846843052 | chr7 | - |
| Fanci | 4,64 | 0,2371 | 0,772961171 | chr7 | + |
| Fank1 | 2,77 | 0,8094 | 0,123940582 | chr7 | + |
| Fbl | 880,64 | -0,1119 | 0,802594059 | chr7 | + |
| Fbrs | 144,06 | -0,6917 | 0,094821256 | chr7 | + |
| Fbxl19 | 42,87 | -0,4771 | 0,408372125 | chr7 | + |
| Fbxo17 | 23,84 | -0,2093 | 0,754524942 | chr7 | + |
| Fbxo46 | 43,71 | -0,1976 | 0,751034711 | chr7 | + |
| Fcgbp | 3,02 | 0,5185 | 0,404187666 | chr7 | + |
| Fcgrt | 2610,63 | -0,2343 | 0,612061669 | chr7 | - |
| Fchsd2 | 382,47 | 0,1763 | 0,724295112 | chr7 | + |
| Fes | 83,02 | -0,4678 | 0,283177097 | chr7 | - |
| Ffar2 | 3,41 | 0,5984 | 0,330791426 | chr7 | - |
| Fgfr2 | 495,40 | -0,5576 | 0,078099222 | chr7 | - |
| Fiz1 | 176,49 | -0,3991 | 0,318113166 | chr7 | - |
| Fkrp | 156,79 | 0,0228 | 0,966233054 | chr7 | - |
| Flt3l | 54,47 | -0,3058 | 0,618723125 | chr7 | - |
| Folr1 | 2,84 | 0,1607 | 0,849664617 | chr7 | - |
| Folr2 | 389,89 | 0,0597 | 0,909834629 | chr7 | - |
| Foxa3 | 1113,44 | 0,0813 | 0,901280629 | chr7 | - |
| Fuom | 582,80 | -0,1757 | 0,676716141 | chr7 | - |
| Furin | 1395,98 | -0,3711 | 0,402148509 | chr7 | - |
| Fus | 1197,75 | 0,3189 | 0,450112767 | chr7 | + |
| Fuz | 121,77 | -0,0935 | 0,869544763 | chr7 | + |
| Fxyd1 | 5261,56 | -0,1296 | 0,645262633 | chr7 | - |
| Fxyd7 | 2,07 | 0,3261 | NA | chr7 | - |
| Fzd4 | 189,90 | 0,5095 | 0,217175999 | chr7 | + |
| Gab2 | 36,18 | 0,5122 | 0,278037339 | chr7 | + |
| Gabrb3 | 23,88 | -0,7247 | 0,18870514 | chr7 | + |
| Galnt18 | 24,59 | -0,4208 | 0,517431746 | chr7 | - |
| Gas2 | 1050,59 | -0,0178 | 0,976991666 | chr7 | + |
| Gdpd3 | 4,87 | 0,0300 | 0,978280656 | chr7 | + |
| Gdpd5 | 9,23 | -0,0294 | 0,978280656 | chr7 | + |
| Gdpgp1 | 46,82 | 0,0572 | 0,932531696 | chr7 | + |
| Gemin7 | 250,52 | -0,4321 | 0,228728627 | chr7 | - |
| Gga2 | 267,78 | 0,1202 | 0,795796985 | chr7 | - |
| Glrx3 | 316,18 | -0,2867 | 0,514682351 | chr7 | + |
| Bicra | 22,49 | 0,4552 | 0,485418468 | chr7 | - |
| Nop53 | 726,35 | -0,4008 | 0,499301366 | chr7 | - |
| Gm15133 | 3,80 | 0,1149 | 0,899463828 | chr7 | + |

| | | | | | |
|---|---|---|---|---|---|
| Gm15545 | 14,01 | 0,0710 | 0,938072754 | chr7 | + |
| Gm15645 | 70,40 | -0,1185 | 0,857665028 | chr7 | + |
| Gm15880 | 3,04 | -0,3857 | 0,58887292 | chr7 | + |
| Gm16157 | 457,82 | 0,0046 | 0,994676823 | chr7 | - |
| Ccdc189 | 13,56 | 0,0541 | 0,952576171 | chr7 | - |
| Gm16938 | 8,28 | -0,0195 | 0,985851274 | chr7 | + |
| Gm1966 | 81,70 | 0,7676 | 0,069527621 | chr7 | - |
| Gm2381 | 6,99 | 0,5493 | 0,416860553 | chr7 | - |
| Gm4532 | 4,06 | -0,3788 | 0,602123385 | chr7 | - |
| Tpbgl | 97,97 | -0,0352 | 0,956605007 | chr7 | - |
| Gm5113 | 24,26 | 0,8845 | 0,064500269 | chr7 | + |
| Zfp975 | 95,95 | 0,6447 | 0,089779423 | chr7 | - |
| Gm8979 | 2,50 | 0,4175 | NA | chr7 | - |
| Gmfg | 270,26 | 0,6565 | 0,05081654 | chr7 | + |
| Gpatch1 | 90,42 | 0,3679 | 0,393345568 | chr7 | - |
| Gpi1 | 2429,40 | 0,2480 | 0,500561834 | chr7 | - |
| Gpr4 | 11,17 | 0,2518 | 0,753971986 | chr7 | + |
| Gprc5b | 10,50 | 0,5064 | 0,443005632 | chr7 | - |
| Gramd1a | 94,22 | -0,3585 | 0,468283516 | chr7 | - |
| Grik5 | 59,02 | -0,0335 | 0,971618406 | chr7 | - |
| Grwd1 | 90,70 | -0,2943 | 0,466793644 | chr7 | - |
| Gsg1l | 3,95 | 0,1043 | 0,911210861 | chr7 | - |
| Gsk3a | 442,92 | -0,4649 | 0,227017349 | chr7 | - |
| Gtf2h1 | 619,96 | -0,3970 | 0,149780096 | chr7 | + |
| Gtf3c1 | 331,80 | -0,0650 | 0,909774279 | chr7 | - |
| Gys1 | 21,28 | -0,1758 | 0,803690485 | chr7 | + |
| Hamp | 12083,80 | 0,0885 | 0,925700434 | chr7 | - |
| Hamp2 | 12302,77 | 1,3452 | NA | chr7 | - |
| Haus5 | 72,88 | -0,4113 | 0,39766425 | chr7 | - |
| Hcst | 15,53 | 0,3168 | 0,644195217 | chr7 | - |
| Hdgfl3 | 22,37 | 0,3517 | 0,613235485 | chr7 | - |
| Herc2 | 464,69 | 0,0200 | 0,971506315 | chr7 | + |
| Hif3a | 2,84 | -0,1528 | 0,847736143 | chr7 | - |
| Hirip3 | 66,04 | -0,1544 | 0,789064438 | chr7 | + |
| Hnrnpl | 1070,30 | -0,4045 | 0,155785005 | chr7 | + |
| Hnrnpul1 | 335,62 | 0,0033 | 0,995598861 | chr7 | - |
| Homer2 | 445,80 | -0,5424 | 0,051874818 | chr7 | - |
| Hpn | 3839,01 | -0,3484 | 0,355134088 | chr7 | - |
| Hps5 | 193,47 | 0,0530 | 0,928682341 | chr7 | - |
| Hpx | 88,06 | 0,4360 | 0,319080738 | chr7 | - |
| Hsd3b7 | 7815,29 | 0,0377 | 0,928423971 | chr7 | + |
| Hspbp1 | 250,86 | -0,3198 | 0,547712688 | chr7 | - |
| Htatip2 | 644,07 | -0,2756 | 0,392464224 | chr7 | + |
| Htra1 | 47,92 | 0,2827 | 0,644195217 | chr7 | + |
| Idh2 | 13050,78 | 0,0846 | 0,845013263 | chr7 | - |
| Ifitm1 | 280,03 | 0,5086 | 0,119628533 | chr7 | + |
| Ifitm2 | 5647,52 | -0,0139 | 0,981991128 | chr7 | - |
| Ifitm3 | 10859,95 | 0,1983 | 0,585908204 | chr7 | - |
| Ifitm5 | 4,30 | 0,0188 | 0,985851274 | chr7 | - |
| Ifitm6 | 19,54 | 0,0501 | 0,953578516 | chr7 | - |

| | | | | | |
|---|---|---|---|---|---|
| Igf1r | 35,73 | 0,0813 | 0,915286446 | chr7 | + |
| Igf2 | 13,99 | 0,2240 | 0,790595509 | chr7 | - |
| Igflr1 | 22,34 | -0,5442 | 0,391701733 | chr7 | + |
| Iglon5 | 3,10 | -0,0967 | 0,913936879 | chr7 | - |
| Igsf23 | 10,04 | -0,1270 | 0,88569455 | chr7 | - |
| Igsf6 | 76,19 | 0,3784 | 0,34427921 | chr7 | - |
| Ikzf5 | 119,27 | -0,0954 | 0,840753129 | chr7 | - |
| Il16 | 79,27 | 0,2374 | 0,60487792 | chr7 | - |
| Il21r | 10,00 | 0,8590 | 0,145912941 | chr7 | + |
| Il4ra | 423,15 | 0,1827 | 0,577257466 | chr7 | + |
| Ilk | 761,57 | -0,3247 | 0,269336293 | chr7 | + |
| Ino80e | 57,22 | -0,3230 | 0,505514525 | chr7 | - |
| Inpp5a | 167,51 | -0,5746 | 0,07494103 | chr7 | + |
| Inpp5f | 387,40 | -0,1784 | 0,707581291 | chr7 | + |
| Inppl1 | 317,13 | -0,2885 | 0,546240378 | chr7 | - |
| Insc | 213,98 | -0,3854 | 0,380860886 | chr7 | + |
| Ints4 | 390,71 | 0,0975 | 0,849664617 | chr7 | + |
| Ipo7 | 1297,07 | 0,2891 | 0,470905336 | chr7 | + |
| Iqgap1 | 215,08 | 0,3062 | 0,339404199 | chr7 | - |
| Irf2bp1 | 95,24 | -0,0781 | 0,918946301 | chr7 | + |
| Irf3 | 588,84 | -0,3676 | 0,302013909 | chr7 | + |
| Irf7 | 937,26 | 0,4140 | 0,225253123 | chr7 | - |
| Irgq | 169,17 | -0,0516 | 0,922379013 | chr7 | + |
| Isg20 | 58,51 | 0,3750 | 0,437072528 | chr7 | + |
| Isoc2a | 2648,00 | -0,5434 | 0,073845576 | chr7 | + |
| Isoc2b | 428,04 | 0,2898 | 0,383819674 | chr7 | - |
| Itgal | 157,90 | 0,0120 | 0,987952599 | chr7 | + |
| Itgam | 19,81 | 0,9526 | 0,062331577 | chr7 | + |
| Itgax | 17,45 | 0,6887 | 0,244197385 | chr7 | + |
| Itpkc | 78,97 | 0,0124 | 0,986642823 | chr7 | - |
| Itpripl2 | 188,78 | 0,2986 | 0,402592894 | chr7 | - |
| Jakmip3 | 2,11 | 0,7463 | NA | chr7 | + |
| Josd2 | 216,51 | -0,5980 | 0,274683426 | chr7 | + |
| Kat8 | 284,75 | -0,2914 | 0,46093503 | chr7 | + |
| Kcnk6 | 44,34 | 0,0597 | 0,931322892 | chr7 | - |
| Kcnn4 | 3,03 | -0,2340 | 0,763514396 | chr7 | + |
| Kcnq1 | 22,39 | -0,3451 | 0,580044078 | chr7 | + |
| Kcnq1ot1 | 664,56 | -0,5343 | 0,059117961 | chr7 | - |
| Kctd13 | 20,61 | -0,0852 | 0,920386232 | chr7 | + |
| Kctd14 | 3,06 | 0,1798 | 0,825674386 | chr7 | + |
| Kctd15 | 128,89 | -0,3552 | 0,381800527 | chr7 | - |
| Kctd21 | 132,98 | -0,1296 | 0,783264136 | chr7 | + |
| Kdelr1 | 1425,32 | -0,3139 | 0,178387119 | chr7 | + |
| Kdm8 | 24,62 | -0,1555 | 0,83097362 | chr7 | + |
| Kif22 | 9,73 | -0,0856 | 0,927365453 | chr7 | - |
| Kif7 | 4,94 | -0,5247 | 0,440094295 | chr7 | - |
| Klc3 | 1,97 | 0,3013 | NA | chr7 | - |
| Klf13 | 1004,17 | -0,1801 | 0,644822613 | chr7 | - |
| Klhl25 | 262,21 | 0,2661 | 0,547091369 | chr7 | + |
| Klk1 | 5,90 | 0,9126 | 0,125787957 | chr7 | + |

| | | | | | |
|---|---|---|---|---|---|
| Klk1b11 | 2,85 | 0,3058 | 0,655665624 | chr7 | + |
| Klk1b3 | 4,22 | 0,3539 | 0,634896352 | chr7 | + |
| Klk8 | 4,58 | -0,1406 | 0,877237757 | chr7 | + |
| Kmt2b | 191,38 | -0,1302 | 0,795853157 | chr7 | - |
| Knop1 | 528,77 | -0,0964 | 0,823659455 | chr7 | - |
| Kptn | 166,12 | -0,2911 | 0,463230756 | chr7 | + |
| Hikeshi | 427,69 | -0,2119 | 0,628656316 | chr7 | - |
| Lair1 | 96,67 | 0,6230 | 0,169324582 | chr7 | - |
| Lamtor1 | 670,94 | -0,1925 | 0,590763774 | chr7 | + |
| Lat | 10,15 | 0,0059 | 0,995663291 | chr7 | - |
| Lcmt1 | 205,30 | -0,1496 | 0,698110464 | chr7 | + |
| Ldha | 13615,14 | -0,1464 | 0,781443303 | chr7 | + |
| Ldhc | 3,68 | 0,1201 | NA | chr7 | + |
| Leng1 | 116,84 | -0,1952 | 0,745498647 | chr7 | - |
| Leng9 | 45,63 | -0,2328 | 0,691395693 | chr7 | - |
| Lgals4 | 4,49 | -0,4261 | 0,561210905 | chr7 | + |
| Lhpp | 731,43 | -0,3486 | 0,39766425 | chr7 | + |
| Lig1 | 37,93 | 0,6430 | 0,182069448 | chr7 | + |
| Lilra5 | 107,13 | 0,4947 | 0,211100165 | chr7 | + |
| Lilra6 | 3,15 | 0,3116 | 0,670139713 | chr7 | - |
| Lin37 | 99,58 | -0,3683 | 0,417486404 | chr7 | - |
| Lins1 | 59,19 | -0,2429 | 0,620367088 | chr7 | + |
| Lipe | 180,23 | -0,3001 | 0,549701484 | chr7 | - |
| Lrfn3 | 36,29 | -0,1071 | 0,905147397 | chr7 | - |
| Lrp3 | 190,44 | -0,2960 | 0,53490582 | chr7 | - |
| Lrrc27 | 6,88 | 0,2003 | 0,814733578 | chr7 | + |
| Lrrc28 | 566,74 | -0,1330 | 0,754953629 | chr7 | - |
| Lrrc32 | 117,62 | 0,1777 | 0,713058621 | chr7 | + |
| Lrrc51 | 62,28 | -0,1231 | 0,848794352 | chr7 | - |
| Lrrc56 | 40,68 | -0,4285 | 0,368642366 | chr7 | + |
| Lrrk1 | 149,43 | 0,2721 | 0,449196578 | chr7 | - |
| Lsm14a | 486,72 | -0,0160 | 0,971756379 | chr7 | - |
| Lsp1 | 57,58 | 0,5851 | 0,248279698 | chr7 | + |
| Lsr | 2472,62 | 0,4324 | 0,235960821 | chr7 | - |
| Ltbp4 | 227,93 | -0,4824 | 0,380860886 | chr7 | - |
| Lyrm1 | 170,44 | -0,4713 | 0,413156981 | chr7 | + |
| Lysmd4 | 198,29 | 0,0383 | 0,941754349 | chr7 | + |
| Lyve1 | 242,90 | 0,4120 | 0,285499423 | chr7 | - |
| Mag | 168,32 | 0,1088 | 0,876857496 | chr7 | - |
| Mtag2 | 2,71 | 0,2396 | 0,757695155 | chr7 | + |
| Man2a2 | 389,89 | 0,0659 | 0,882318627 | chr7 | - |
| Map3k10 | 79,12 | -0,6519 | 0,181882538 | chr7 | - |
| Map4k1 | 6,58 | 0,2195 | 0,795600492 | chr7 | + |
| Mapk1ip1 | 90,25 | -0,0211 | 0,976528475 | chr7 | - |
| Mapk3 | 420,73 | 0,0775 | 0,827603438 | chr7 | + |
| Mark4 | 37,91 | -0,3812 | 0,550015683 | chr7 | - |
| Maz | 421,10 | -0,6150 | 0,063577348 | chr7 | - |
| Mboat7 | 381,54 | -0,3627 | 0,271992655 | chr7 | - |
| Mcee | 1282,98 | -0,3972 | 0,209104214 | chr7 | + |
| Mcmbp | 405,16 | -0,1584 | 0,604579545 | chr7 | - |

| | | | | | |
|---|---|---|---|---|---|
| Mctp2 | 206,26 | -0,3488 | 0,441724922 | chr7 | - |
| Me3 | 4,45 | 0,0061 | 0,995663291 | chr7 | + |
| Med25 | 145,71 | -0,6097 | 0,223366879 | chr7 | - |
| Med29 | 353,20 | -0,2908 | 0,455260576 | chr7 | - |
| Mef2a | 457,29 | -0,1506 | 0,644046286 | chr7 | - |
| Meis3 | 9,96 | -0,1559 | 0,857478182 | chr7 | + |
| Tlnrd1 | 42,16 | 0,1927 | 0,764664236 | chr7 | - |
| Mesd | 541,89 | -0,0504 | 0,907233333 | chr7 | + |
| Mesp1 | 1,88 | 0,3752 | NA | chr7 | - |
| Mesp2 | 2,45 | -0,4844 | NA | chr7 | + |
| Eef1akmt2 | 115,26 | 0,4355 | 0,260028246 | chr7 | - |
| Mettl9 | 747,51 | -0,1695 | 0,617391963 | chr7 | + |
| Mex3b | 7,39 | 0,2929 | 0,70880025 | chr7 | + |
| Mgmt | 335,31 | 0,2920 | 0,41977893 | chr7 | + |
| Mia | 7,46 | -0,0405 | 0,969466644 | chr7 | - |
| Micalcl | 12,33 | 0,5332 | 0,400903279 | chr7 | + |
| Mill2 | 7,92 | 0,0628 | 0,950391517 | chr7 | + |
| Mir3101 | 2,81 | 0,1280 | 0,87867141 | chr7 | - |
| Mir7051 | 2,42 | 0,0072 | NA | chr7 | + |
| Mki67 | 59,37 | 0,1519 | 0,820577263 | chr7 | - |
| Mkrn3 | 6,26 | 1,0117 | 0,079042198 | chr7 | - |
| Mob2 | 367,95 | -0,3623 | 0,242830621 | chr7 | - |
| Mphosph10 | 292,96 | -0,0728 | 0,913017292 | chr7 | - |
| Mrgpre | 51,34 | 0,2648 | 0,681592614 | chr7 | - |
| Mrgprf | 2,76 | 0,2360 | 0,760825768 | chr7 | + |
| Mrpl17 | 729,55 | -0,3100 | 0,463314602 | chr7 | - |
| Mrpl23 | 805,07 | 0,1979 | 0,479398672 | chr7 | + |
| Mrpl46 | 433,58 | -0,1046 | 0,812490734 | chr7 | - |
| Mrpl48 | 768,19 | -0,3649 | 0,229925363 | chr7 | - |
| Mrps11 | 303,21 | -0,3530 | 0,282189277 | chr7 | + |
| Mrps12 | 379,42 | -0,2093 | 0,806012703 | chr7 | - |
| Mrvi1 | 22,80 | 0,2087 | 0,754953629 | chr7 | - |
| Mtg1 | 346,09 | -0,0986 | 0,797559244 | chr7 | + |
| Mtmr10 | 588,22 | -0,1672 | 0,624956201 | chr7 | + |
| Mvp | 502,01 | -0,2172 | 0,623104286 | chr7 | - |
| Myh14 | 108,51 | 0,0510 | 0,941474427 | chr7 | - |
| Mylpf | 23,86 | 0,5792 | 0,292588112 | chr7 | + |
| Myo7a | 165,27 | 0,1513 | 0,764363417 | chr7 | - |
| Mypop | 13,22 | 0,1050 | 0,912010517 | chr7 | + |
| Mzf1 | 2,16 | 0,0877 | NA | chr7 | - |
| Nadsyn1 | 320,75 | 0,0608 | 0,906066614 | chr7 | - |
| Nap1l4 | 830,43 | 0,2385 | 0,392238586 | chr7 | - |
| Napa | 625,77 | -0,4736 | 0,075686715 | chr7 | + |
| Napsa | 11,20 | 0,4529 | 0,516624153 | chr7 | + |
| Nars2 | 327,02 | -0,5240 | 0,073430016 | chr7 | + |
| Nat14 | 5,39 | -0,5148 | 0,452418751 | chr7 | + |
| Nav2 | 142,70 | 0,5324 | 0,115172572 | chr7 | + |
| Ncr1 | 5,26 | 0,6412 | 0,326858708 | chr7 | + |
| Ndn | 17,32 | -0,1593 | 0,834675238 | chr7 | + |
| Nsmce3 | 175,23 | -0,1714 | 0,722794314 | chr7 | - |

| | | | | | |
|---|---|---|---|---|---|
| Ndufa3 | 3675,57 | -0,0986 | 0,795600492 | chr7 | + |
| Ndufab1 | 577,64 | -0,5818 | 0,178097604 | chr7 | - |
| Ndufc2 | 1975,35 | -0,0489 | 0,950532633 | chr7 | + |
| Nfatc2ip | 58,04 | -0,0925 | 0,881325856 | chr7 | - |
| Nfkbib | 131,19 | -0,3883 | 0,528469822 | chr7 | - |
| Nfkbid | 4,21 | 0,6184 | 0,342224382 | chr7 | + |
| Ngrn | 268,79 | -0,1960 | 0,572214131 | chr7 | + |
| Nipa1 | 29,88 | 0,6589 | 0,180386456 | chr7 | - |
| Nipa2 | 659,80 | -0,1978 | 0,54877155 | chr7 | - |
| Nkg7 | 12,95 | 0,0434 | 0,965770119 | chr7 | + |
| Nlrp10 | 4,01 | 0,4600 | 0,515706574 | chr7 | - |
| Nlrp12 | 1241,08 | 0,1284 | 0,793608383 | chr7 | - |
| Nlrp6 | 1631,11 | -0,4691 | 0,116833055 | chr7 | + |
| Nmb | 6,76 | 0,3201 | 0,681592614 | chr7 | - |
| Nomo1 | 412,91 | -0,5195 | 0,072050071 | chr7 | + |
| Nosip | 277,31 | -0,1160 | 0,787073094 | chr7 | + |
| Nova2 | 8,96 | 0,1577 | 0,860107955 | chr7 | + |
| Nphs1 | 2,03 | 0,5653 | NA | chr7 | + |
| Nr1h2 | 315,80 | -0,0836 | 0,907233333 | chr7 | - |
| Nr2f2 | 504,54 | -0,0740 | 0,881809902 | chr7 | - |
| Nsmce1 | 281,73 | -0,0290 | 0,949619489 | chr7 | - |
| Nsmce4a | 612,04 | -0,3944 | 0,264005278 | chr7 | - |
| Nucb1 | 1610,71 | -0,0949 | 0,86120828 | chr7 | - |
| Nucb2 | 142,17 | -0,1343 | 0,868115228 | chr7 | + |
| Nudt19 | 793,77 | -0,3106 | 0,383392162 | chr7 | - |
| Numa1 | 430,68 | 0,0093 | 0,988766352 | chr7 | + |
| Numbl | 6,37 | 0,3090 | 0,693242675 | chr7 | + |
| Nup62 | 181,56 | -0,1621 | 0,757565313 | chr7 | + |
| Nup98 | 347,10 | -0,1084 | 0,771786001 | chr7 | - |
| Nupr1 | 75,32 | -0,0125 | 0,98797801 | chr7 | - |
| Oat | 12600,33 | -0,3631 | 0,323355495 | chr7 | - |
| Olfml1 | 392,68 | 0,2838 | 0,62811896 | chr7 | + |
| Olfr613 | 47,54 | -0,0032 | 0,996644671 | chr7 | + |
| Omp | 2,52 | 0,5779 | NA | chr7 | - |
| Opa3 | 244,21 | 0,2989 | 0,48643419 | chr7 | + |
| Orai3 | 238,53 | 0,0077 | 0,988912705 | chr7 | + |
| Oraov1 | 179,22 | -0,3165 | 0,375935576 | chr7 | + |
| Osbpl5 | 38,35 | -0,0885 | 0,896013441 | chr7 | - |
| Ovol3 | 5,19 | -0,3745 | 0,618723125 | chr7 | - |
| P2ry2 | 217,95 | -0,1773 | 0,720291746 | chr7 | - |
| P2ry6 | 22,51 | 0,6441 | 0,218722998 | chr7 | - |
| P4ha3 | 2,00 | 0,3034 | NA | chr7 | + |
| Paf1 | 320,75 | -0,3963 | 0,182208897 | chr7 | + |
| Pafah1b3 | 38,97 | -0,2232 | 0,757695155 | chr7 | - |
| Pagr1a | 203,07 | -0,2176 | 0,606429785 | chr7 | - |
| Pak4 | 100,44 | -0,1616 | 0,813323745 | chr7 | - |
| Palb2 | 13,67 | -0,1421 | 0,872117566 | chr7 | - |
| Paox | 651,65 | 0,0715 | 0,906578208 | chr7 | + |
| Parva | 1306,51 | -0,1632 | 0,603256989 | chr7 | + |
| Pcf11 | 451,94 | -0,0592 | 0,905147397 | chr7 | - |

| | | | | | |
|---|---|---|---|---|---|
| Pdcd2l | 100,38 | -0,0964 | 0,850453487 | chr7 | - |
| Pdcd5 | 420,05 | -0,5067 | 0,138593946 | chr7 | - |
| Gatd1 | 241,49 | 0,0283 | 0,960906846 | chr7 | - |
| Pde2a | 319,54 | 0,1171 | 0,762922832 | chr7 | + |
| Pde8a | 182,61 | 0,0114 | 0,98429278 | chr7 | + |
| Peg12 | 2,86 | 0,2745 | 0,717851252 | chr7 | - |
| Peg3 | 40,35 | 0,6525 | 0,177916821 | chr7 | - |
| Pepd | 814,07 | -0,3088 | 0,215046281 | chr7 | + |
| Pgap2 | 501,78 | 0,2740 | 0,482787957 | chr7 | + |
| Pglyrp1 | 1249,07 | 0,0465 | 0,929917979 | chr7 | + |
| Pgpep1l | 19,01 | 0,6824 | 0,217231457 | chr7 | - |
| Phlda2 | 5,30 | 0,5178 | 0,449196578 | chr7 | - |
| Phldb3 | 35,40 | -0,1167 | 0,87618436 | chr7 | + |
| Picalm | 2306,97 | -0,2960 | 0,277831408 | chr7 | + |
| Pidd1 | 15,68 | -0,1230 | 0,889997259 | chr7 | - |
| Pih1d1 | 532,46 | -0,4653 | 0,222670104 | chr7 | + |
| Pik3c2a | 792,24 | 0,0698 | 0,886748924 | chr7 | - |
| Pira2 | 11,72 | 0,7411 | 0,205871072 | chr7 | - |
| Pirb | 56,62 | 0,1104 | 0,877920448 | chr7 | - |
| Pkp3 | 9,92 | 0,1094 | 0,906333899 | chr7 | + |
| Plaur | 7,40 | -0,0706 | 0,943593159 | chr7 | + |
| Pld3 | 221,94 | -0,2601 | 0,580859417 | chr7 | - |
| Plekha4 | 6,42 | 0,3871 | 0,606323442 | chr7 | + |
| Plekha7 | 33,86 | 0,3167 | 0,634237396 | chr7 | - |
| Plekhg2 | 42,37 | 0,2921 | 0,624676895 | chr7 | - |
| Plk1 | 2,42 | 0,4279 | NA | chr7 | + |
| Pnkp | 215,68 | -0,2065 | 0,6180541 | chr7 | + |
| Pnpla2 | 743,77 | -0,3828 | 0,430842469 | chr7 | + |
| Pold1 | 25,81 | -0,2489 | 0,744313101 | chr7 | - |
| Pold3 | 151,23 | 0,4128 | 0,296117154 | chr7 | - |
| Polg | 469,40 | -0,1358 | 0,679262748 | chr7 | - |
| Polr2i | 566,19 | -0,3898 | 0,141463153 | chr7 | + |
| Polr2l | 897,49 | -0,3412 | 0,52825088 | chr7 | - |
| Polr3e | 142,48 | -0,3413 | 0,357345082 | chr7 | + |
| Pop4 | 328,46 | 0,0078 | 0,991035244 | chr7 | - |
| Pou2f2 | 6,72 | 0,4306 | 0,556416148 | chr7 | - |
| Ppfia1 | 273,84 | -0,2800 | 0,359146889 | chr7 | - |
| Ppfibp2 | 348,57 | -0,0628 | 0,884712754 | chr7 | + |
| Ppme1 | 386,02 | -0,0973 | 0,839302072 | chr7 | - |
| Ppp1r12c | 200,44 | -0,5886 | 0,222670104 | chr7 | - |
| Ppp1r13l | 16,77 | -0,7407 | 0,175457889 | chr7 | + |
| Ppp1r14a | 10,36 | 0,5285 | 0,425926605 | chr7 | + |
| Ppp1r15a | 115,89 | -0,4716 | 0,267895481 | chr7 | - |
| Ppp1r37 | 468,36 | -0,5369 | 0,157077211 | chr7 | - |
| Ppp2r2d | 469,34 | -0,2329 | 0,647295341 | chr7 | + |
| Ppp4c | 487,00 | -0,4250 | 0,165182527 | chr7 | - |
| Ppp5c | 467,90 | -0,2264 | 0,539194011 | chr7 | - |
| Ppp6r1 | 948,85 | 0,0458 | 0,916852736 | chr7 | - |
| Prc1 | 26,11 | 0,1376 | 0,848887287 | chr7 | + |
| Prcp | 193,05 | 0,0873 | 0,838412507 | chr7 | + |

| | | | | | |
|---|---|---|---|---|---|
| Prkcb | 70,17 | 0,2536 | 0,595133363 | chr7 | + |
| Cavin3 | 10,77 | 0,0892 | 0,922904986 | chr7 | - |
| Prkcg | 2,37 | -0,1108 | NA | chr7 | + |
| Prkd2 | 66,76 | 0,2387 | 0,614721295 | chr7 | + |
| Thap12 | 604,51 | -0,1666 | 0,653708291 | chr7 | + |
| Prmt1 | 368,74 | -0,1043 | 0,796541442 | chr7 | - |
| Prmt3 | 140,65 | -0,2328 | 0,612582336 | chr7 | + |
| Prodh2 | 2368,69 | -0,1214 | 0,842502441 | chr7 | + |
| Prpf31 | 269,84 | 0,0855 | 0,882354365 | chr7 | + |
| Prr12 | 50,49 | 0,1989 | 0,756319626 | chr7 | - |
| Prr14 | 298,31 | -0,5281 | 0,173352491 | chr7 | + |
| Inafm1 | 56,97 | -0,0084 | 0,991098974 | chr7 | - |
| Prrg2 | 365,46 | -0,5795 | 0,116324739 | chr7 | - |
| Prrt2 | 2,49 | -0,0661 | NA | chr7 | - |
| Prss23 | 83,58 | 0,4173 | 0,35583041 | chr7 | - |
| Prss36 | 115,62 | -0,0384 | 0,948200011 | chr7 | - |
| Prss53 | 88,82 | -0,6784 | 0,196659616 | chr7 | - |
| Prss8 | 147,54 | 0,2110 | 0,636381742 | chr7 | - |
| Prx | 4,58 | -0,6522 | 0,314703919 | chr7 | + |
| Psenen | 1172,22 | -0,3753 | 0,194460708 | chr7 | - |
| Psg23 | 92,90 | -0,6945 | 0,10617067 | chr7 | - |
| Psg28 | 2,35 | -0,1288 | NA | chr7 | - |
| Psg29 | 2,99 | -0,6265 | 0,297002219 | chr7 | + |
| Psma1 | 2683,91 | -0,0856 | 0,838412507 | chr7 | - |
| Psmc4 | 1319,31 | -0,2655 | 0,464131878 | chr7 | - |
| Psmd13 | 1439,02 | -0,3358 | 0,258527696 | chr7 | + |
| Psmd8 | 2192,23 | -0,3952 | 0,19689523 | chr7 | - |
| Pstk | 138,06 | -0,2551 | 0,624005041 | chr7 | + |
| Ptdss2 | 138,35 | -0,4607 | 0,197777944 | chr7 | + |
| Ptgir | 2,93 | 0,4027 | 0,564495903 | chr7 | + |
| Ptpre | 26,90 | 0,2480 | 0,687001462 | chr7 | + |
| Pvr | 204,99 | 0,2043 | 0,587901189 | chr7 | - |
| Nectin2 | 566,08 | -0,4098 | 0,211038545 | chr7 | - |
| Pwwp2b | 28,69 | -0,2707 | 0,640132917 | chr7 | + |
| Pycard | 41,68 | 0,6510 | 0,160487212 | chr7 | - |
| Qpctl | 90,42 | -0,5331 | 0,208381949 | chr7 | - |
| Qprt | 2299,16 | 0,2369 | 0,606572668 | chr7 | - |
| R74862 | 7,77 | -0,3980 | 0,588663593 | chr7 | - |
| Rab30 | 395,13 | -0,0173 | 0,987282322 | chr7 | + |
| Rab4b | 120,27 | -0,3358 | 0,496763475 | chr7 | - |
| Rab6a | 1082,77 | 0,0382 | 0,932012942 | chr7 | + |
| Rabac1 | 839,51 | -0,7182 | 0,160195268 | chr7 | - |
| Rabep2 | 88,56 | -0,4794 | 0,414445273 | chr7 | + |
| Rasgrp4 | 5,00 | -0,0439 | 0,967498977 | chr7 | + |
| Rasip1 | 40,63 | -0,1994 | 0,765219833 | chr7 | + |
| Rasl2-9 | 21,70 | 0,8595 | 0,129071791 | chr7 | - |
| Rassf7 | 260,29 | -0,1689 | 0,755785289 | chr7 | + |
| Rbbp6 | 578,85 | -0,2455 | 0,37347595 | chr7 | + |
| Rbm42 | 252,06 | -0,4233 | 0,425523334 | chr7 | - |
| Rccd1 | 17,69 | 0,2475 | 0,743508016 | chr7 | - |

| | | | | | |
|---|---|---|---|---|---|
| Rcn3 | 74,56 | -0,0288 | 0,968759875 | chr7 | - |
| Rdh13 | 184,82 | -0,4867 | 0,242907451 | chr7 | - |
| Relb | 24,62 | -0,0729 | 0,932525817 | chr7 | - |
| Relt | 5,48 | -0,0999 | 0,916757648 | chr7 | - |
| Rgma | 3,13 | 0,1278 | 0,88306163 | chr7 | + |
| Rgs10 | 41,38 | 0,2186 | 0,699742099 | chr7 | - |
| Rgs9bp | 3,32 | 0,2296 | 0,765061508 | chr7 | - |
| Rhog | 54,25 | 0,1491 | 0,777579 | chr7 | - |
| Rhpn2 | 223,14 | -0,2932 | 0,479837755 | chr7 | + |
| Ric8a | 275,20 | -0,2576 | 0,468217847 | chr7 | + |
| Rinl | 23,06 | 0,6614 | 0,208911872 | chr7 | + |
| Rnf121 | 325,26 | 0,2052 | 0,623297443 | chr7 | - |
| Rnf169 | 748,20 | -0,1716 | 0,681752477 | chr7 | - |
| Rnf40 | 271,61 | -0,0543 | 0,913102981 | chr7 | + |
| Rnh1 | 1762,22 | 0,2196 | 0,407206512 | chr7 | - |
| Rpl13a | 7860,95 | -0,6305 | 0,108667103 | chr7 | - |
| Rpl14-ps1 | 9,23 | -0,7154 | 0,260028246 | chr7 | + |
| Rpl18 | 8598,56 | -0,3832 | 0,152380315 | chr7 | + |
| Rpl27a | 5506,78 | -0,5941 | 0,243141521 | chr7 | + |
| Rplp2 | 1270,50 | -0,5525 | 0,413537476 | chr7 | + |
| Rps11 | 4794,34 | -0,3568 | 0,212210133 | chr7 | - |
| Rps13 | 2353,87 | -0,5389 | 0,227528325 | chr7 | - |
| Rps15a | 3518,55 | -0,5279 | 0,061053776 | chr7 | - |
| Rps16 | 3864,98 | -0,3659 | 0,458689654 | chr7 | + |
| Rps17 | 7853,69 | -0,5003 | 0,050030275 | chr7 | - |
| Rps19 | 4723,09 | -0,4745 | 0,062190624 | chr7 | + |
| Rps3 | 9640,86 | -0,3314 | 0,224150405 | chr7 | - |
| Rps5 | 6190,86 | -0,5987 | 0,13613671 | chr7 | + |
| Rps9 | 6231,91 | -0,4799 | 0,054141872 | chr7 | + |
| Rras | 284,80 | 0,0314 | 0,951574951 | chr7 | + |
| Rras2 | 385,64 | 0,2121 | 0,567161852 | chr7 | - |
| Rrm1 | 279,88 | 0,4729 | 0,237120323 | chr7 | + |
| Rrp8 | 314,10 | -0,0075 | 0,988512389 | chr7 | - |
| Rsf1 | 262,66 | -0,0876 | 0,848140004 | chr7 | + |
| Rtn2 | 6,31 | -0,0491 | 0,962251592 | chr7 | + |
| Ruvbl2 | 241,97 | -0,0136 | 0,979212524 | chr7 | - |
| Ryr1 | 618,68 | -0,2139 | 0,465412028 | chr7 | - |
| Saa1 | 5275,39 | -0,4316 | 0,518617521 | chr7 | - |
| Saa2 | 2638,10 | -0,3081 | 0,648706509 | chr7 | + |
| Saa3 | 282,78 | -0,7076 | 0,266247219 | chr7 | - |
| Saa4 | 8073,53 | -0,8365 | 0,079609563 | chr7 | - |
| Saal1 | 119,54 | 0,2548 | 0,532778528 | chr7 | - |
| Sae1 | 330,15 | 0,1061 | 0,794631983 | chr7 | - |
| Samd4b | 137,20 | -0,4891 | 0,284114812 | chr7 | - |
| Sars2 | 89,05 | -0,5802 | 0,244584613 | chr7 | + |
| Sbk1 | 342,47 | 0,2263 | 0,538146816 | chr7 | + |
| Sbsn | 7,17 | 0,5621 | 0,405647089 | chr7 | + |
| Scaf1 | 182,19 | -0,4055 | 0,509118004 | chr7 | - |
| Scgb1c1 | 3,75 | 0,3037 | 0,695978671 | chr7 | + |
| Scn1b | 37,17 | 0,3491 | 0,564849438 | chr7 | - |

| | | | | | |
|---|---|---|---|---|---|
| Sdhaf1 | 120,49 | -0,4949 | 0,187760013 | chr7 | - |
| Sec11a | 1498,85 | -0,0335 | 0,93551479 | chr7 | - |
| Sec23ip | 335,75 | -0,1865 | 0,599464204 | chr7 | + |
| Sema4b | 59,24 | 0,1193 | 0,874485842 | chr7 | + |
| Selenow | 1670,12 | -0,4608 | 0,138593946 | chr7 | - |
| Sergef | 73,36 | -0,0678 | 0,902760092 | chr7 | - |
| Sertad1 | 138,73 | 0,0217 | 0,979212524 | chr7 | + |
| Sertad3 | 90,05 | -0,0630 | 0,932528423 | chr7 | + |
| Setd1a | 217,85 | -0,6253 | 0,169529667 | chr7 | + |
| Sez6l2 | 2,77 | 0,1353 | 0,876857496 | chr7 | + |
| Sh2b1 | 297,81 | -0,5169 | 0,176392509 | chr7 | - |
| Shank2 | 64,27 | 0,3748 | 0,570235502 | chr7 | + |
| Shkbp1 | 64,95 | -0,3538 | 0,463892619 | chr7 | - |
| Sigirr | 254,16 | -0,2119 | 0,669817756 | chr7 | - |
| Siglece | 55,41 | 0,3542 | 0,546240378 | chr7 | - |
| Siglecg | 2,76 | -0,4941 | 0,456471434 | chr7 | + |
| Siglech | 24,26 | 0,1616 | 0,833762767 | chr7 | + |
| Sipa1l3 | 43,38 | -0,4301 | 0,444145513 | chr7 | - |
| Sirt3 | 1832,31 | -0,1207 | 0,740352128 | chr7 | - |
| Six5 | 19,80 | -0,6453 | 0,271504179 | chr7 | + |
| Slc1a5 | 45,35 | -0,1930 | 0,748519887 | chr7 | + |
| Slc22a18 | 1573,77 | 0,0685 | 0,840817476 | chr7 | + |
| Slc25a22 | 1941,92 | 0,0405 | 0,953911274 | chr7 | - |
| Slc27a5 | 19921,76 | -0,2823 | 0,29082501 | chr7 | - |
| Slc28a1 | 5,03 | 0,7488 | 0,217231457 | chr7 | + |
| Slc5a11 | 2,76 | 0,1457 | 0,86698964 | chr7 | + |
| Slco2b1 | 1948,13 | 0,0007 | 0,998514705 | chr7 | - |
| Slco3a1 | 62,34 | -0,2490 | 0,610005889 | chr7 | - |
| Slx1b | 125,80 | -0,3119 | 0,447359916 | chr7 | - |
| Smg1 | 691,87 | 0,0923 | 0,84862368 | chr7 | - |
| Smg9 | 73,72 | -0,5599 | 0,233364856 | chr7 | + |
| Snora3 | 3,30 | 0,1795 | 0,83293655 | chr7 | + |
| Snord15b | 6,03 | -0,5595 | 0,40679401 | chr7 | - |
| Snord32a | 2,44 | -0,3139 | NA | chr7 | - |
| Snord33 | 8,16 | -0,3747 | 0,618723125 | chr7 | - |
| Snord34 | 2,54 | -0,0316 | NA | chr7 | - |
| Snord35a | 2,82 | -0,1061 | 0,906066614 | chr7 | - |
| Snrnp70 | 671,57 | -0,2550 | 0,666980604 | chr7 | - |
| Snrpa | 220,24 | -0,4519 | 0,357200836 | chr7 | - |
| Snrpa1 | 276,47 | -0,1506 | 0,705982644 | chr7 | + |
| Snrpd2 | 849,83 | -0,1970 | 0,581990132 | chr7 | + |
| Sox6 | 184,30 | -0,2072 | 0,627992485 | chr7 | - |
| Spcs2 | 2038,10 | -0,0332 | 0,957758516 | chr7 | - |
| Sphk2 | 271,12 | -0,2677 | 0,604491975 | chr7 | - |
| Spib | 9,70 | 0,3199 | 0,670881738 | chr7 | - |
| Spint2 | 561,18 | -0,4213 | 0,259978915 | chr7 | - |
| Spn | 12,67 | 0,7029 | 0,248800492 | chr7 | - |
| Spns1 | 235,63 | -0,4598 | 0,29259208 | chr7 | - |
| Spon1 | 17,45 | 0,2636 | 0,736998711 | chr7 | + |
| Spred3 | 4,21 | -0,3687 | 0,616161968 | chr7 | - |

| | | | | | |
|---|---|---|---|---|---|
| Sprn | 20,66 | -0,3223 | 0,649651145 | chr7 | - |
| Spty2d1 | 277,04 | 0,0494 | 0,921261454 | chr7 | - |
| Ssc5d | 2,68 | 0,2451 | 0,753233421 | chr7 | + |
| St5 | 237,63 | 0,5153 | 0,10394222 | chr7 | - |
| Stard5 | 2452,51 | 0,1438 | 0,683592841 | chr7 | + |
| Stim1 | 281,43 | -0,3279 | 0,315414722 | chr7 | + |
| Strn4 | 308,86 | -0,1691 | 0,646093772 | chr7 | + |
| Stx1b | 10,35 | 0,6708 | 0,280299373 | chr7 | - |
| Stx4a | 346,52 | -0,3422 | 0,306882295 | chr7 | + |
| Sult1a1 | 8443,88 | 0,5512 | 0,057769579 | chr7 | - |
| Sult2a7 | 38,82 | -0,5533 | 0,2894527 | chr7 | - |
| Supt5 | 653,55 | -0,0521 | 0,903337956 | chr7 | - |
| Kmt5c | 101,97 | -0,0738 | 0,918032423 | chr7 | + |
| Svip | 295,15 | -0,4030 | 0,202636202 | chr7 | - |
| Swap70 | 125,45 | -0,0440 | 0,938364545 | chr7 | + |
| Sympk | 441,74 | -0,2031 | 0,631294599 | chr7 | + |
| Syne4 | 34,15 | -0,1420 | 0,853848645 | chr7 | + |
| Synm | 9,31 | 0,4432 | 0,532313185 | chr7 | - |
| Syt9 | 10,41 | 0,0452 | 0,965898926 | chr7 | + |
| Sytl2 | 19,82 | 0,4047 | 0,546882594 | chr7 | + |
| Tacc2 | 241,58 | 0,4105 | 0,42574908 | chr7 | + |
| Taf10 | 372,61 | -0,5754 | 0,081609555 | chr7 | - |
| Taldo1 | 1949,96 | -0,3288 | 0,264796388 | chr7 | + |
| Taok2 | 491,56 | -0,2015 | 0,651798423 | chr7 | - |
| Tarsl2 | 215,59 | -0,2967 | 0,379622702 | chr7 | + |
| Tbc1d10b | 105,81 | -0,2889 | 0,501991823 | chr7 | - |
| Tbc1d17 | 328,00 | -0,2644 | 0,500489603 | chr7 | - |
| Tbcb | 364,92 | -0,3298 | 0,477004036 | chr7 | - |
| Tbx6 | 7,04 | -0,1380 | 0,880056544 | chr7 | + |
| Tead1 | 132,10 | 0,1044 | 0,856916818 | chr7 | + |
| Tead2 | 16,33 | 0,6209 | 0,296372739 | chr7 | + |
| Tfpt | 111,89 | -0,4979 | 0,220525617 | chr7 | - |
| Tgfb1 | 52,70 | -0,3152 | 0,605538599 | chr7 | + |
| Tgfb1i1 | 22,84 | -0,0377 | 0,969159464 | chr7 | + |
| Thrsp | 8995,63 | -0,3125 | 0,590082603 | chr7 | - |
| Thumpd1 | 354,60 | 0,1539 | 0,715025856 | chr7 | - |
| Tial1 | 704,64 | -0,2456 | 0,393059795 | chr7 | - |
| Timm10b | 301,32 | -0,4733 | 0,358287811 | chr7 | + |
| Timm50 | 342,46 | -0,4008 | 0,306398925 | chr7 | - |
| Tjp1 | 571,08 | 0,3823 | 0,151573825 | chr7 | - |
| Tm2d3 | 550,98 | -0,5766 | 0,090019039 | chr7 | + |
| Tm6sf1 | 46,26 | 0,2616 | 0,612061669 | chr7 | + |
| Tmc7 | 92,80 | -0,1462 | 0,811801336 | chr7 | - |
| Tmc4 | 105,25 | -0,4378 | 0,321371459 | chr7 | - |
| Tmem126a | 902,55 | -0,3812 | 0,22384344 | chr7 | - |
| Tmem126b | 544,43 | -0,0587 | 0,912811384 | chr7 | - |
| Tmem135 | 1541,25 | -0,1792 | 0,576013884 | chr7 | - |
| Tmem143 | 217,97 | -0,3006 | 0,476997063 | chr7 | + |
| Tmem150b | 3,42 | -0,1573 | 0,85511764 | chr7 | - |
| Tmem159 | 71,50 | 0,5371 | 0,173484224 | chr7 | + |

| | | | | | |
|---|---|---|---|---|---|
| Tmem238 | 27,30 | -0,0516 | 0,955496112 | chr7 | - |
| Tmem41b | 612,74 | -0,0270 | 0,955803452 | chr7 | - |
| Tmem80 | 145,47 | -0,0312 | 0,951574951 | chr7 | + |
| Tmem86a | 282,11 | 0,5671 | 0,110056031 | chr7 | + |
| Tmem86b | 1179,21 | 0,0100 | 0,986997724 | chr7 | - |
| Tmem9b | 419,53 | -0,2856 | 0,377748546 | chr7 | - |
| Tnfrsf23 | 18,38 | 0,4504 | 0,472886294 | chr7 | - |
| Tnfrsf26 | 19,00 | 0,4639 | 0,43326072 | chr7 | - |
| Tnni2 | 2,68 | 0,3350 | 0,634288113 | chr7 | + |
| Tnnt1 | 35,83 | 0,6818 | 0,1598616 | chr7 | - |
| Tnrc6a | 715,99 | 0,1363 | 0,714472048 | chr7 | + |
| Tollip | 784,59 | -0,2178 | 0,449891525 | chr7 | - |
| Tomm40 | 232,35 | -0,5127 | 0,142377623 | chr7 | - |
| Tpcn2 | 37,36 | -0,1424 | 0,823321555 | chr7 | - |
| Tpp1 | 1374,34 | -0,3045 | 0,235102108 | chr7 | - |
| Trappc6a | 128,31 | -0,3446 | 0,615847246 | chr7 | + |
| Trim12a | 157,46 | 0,3066 | 0,562610061 | chr7 | - |
| Trim12c | 427,74 | 0,5667 | 0,218750305 | chr7 | - |
| Trim28 | 693,85 | -0,3431 | 0,258287939 | chr7 | + |
| Trim3 | 80,78 | -0,3647 | 0,418748096 | chr7 | - |
| Trim30b | 6,29 | 0,2221 | 0,793415854 | chr7 | - |
| Trim34a | 136,19 | 0,5719 | 0,234863974 | chr7 | + |
| Trim6 | 3,23 | 0,7702 | 0,190969312 | chr7 | + |
| Trim72 | 7,44 | -0,6224 | 0,343973842 | chr7 | + |
| Tsen34 | 338,35 | -0,0501 | 0,932531696 | chr7 | + |
| Tsg101 | 642,19 | -0,2482 | 0,371976461 | chr7 | - |
| Tshz3 | 13,74 | -0,2699 | 0,715436191 | chr7 | + |
| Tspan32 | 6,56 | 0,0667 | 0,947294154 | chr7 | + |
| Tspan4 | 1042,27 | -0,5083 | 0,166993075 | chr7 | + |
| Tssc4 | 162,55 | -0,0789 | 0,883885377 | chr7 | + |
| Ttc23 | 1098,76 | 0,6001 | 0,069200865 | chr7 | + |
| Ttll13 | 6,83 | 0,5419 | 0,424735065 | chr7 | + |
| Ttyh1 | 26,23 | 0,4628 | 0,512986719 | chr7 | + |
| Tubgcp2 | 121,32 | -0,0260 | 0,966776406 | chr7 | - |
| Tubgcp5 | 90,29 | 0,5396 | 0,138971192 | chr7 | + |
| Tufm | 1320,93 | -0,3794 | 0,249205625 | chr7 | + |
| Tulp2 | 2,86 | 0,4173 | 0,541556317 | chr7 | + |
| Tyr | 6,85 | -0,5547 | 0,413323375 | chr7 | - |
| Tyrobp | 760,69 | 0,4622 | 0,063989705 | chr7 | + |
| U2af1l4 | 216,23 | -0,2538 | 0,433006548 | chr7 | + |
| U2af2 | 511,78 | -0,2223 | 0,552537331 | chr7 | + |
| Uba2 | 502,61 | 0,0671 | 0,860606321 | chr7 | - |
| Ube2m | 420,48 | -0,6428 | 0,083831145 | chr7 | - |
| Ube3a | 623,04 | 0,1526 | 0,747375272 | chr7 | + |
| Ubfd1 | 589,96 | -0,1078 | 0,77991551 | chr7 | + |
| Uevld | 283,79 | 0,2552 | 0,494267596 | chr7 | - |
| Unc45a | 116,46 | -0,4395 | 0,39810264 | chr7 | - |
| Uqcrc2 | 7915,89 | -0,0472 | 0,907233333 | chr7 | + |
| Urah | 5304,24 | -0,4419 | 0,33288309 | chr7 | + |
| Uros | 277,91 | 0,0206 | 0,976528475 | chr7 | - |

| | | | | | |
|---|---|---|---|---|---|
| Usf2 | 575,50 | -0,5806 | 0,092854537 | chr7 | - |
| Usp17la | 14,06 | -0,0824 | 0,925042168 | chr7 | + |
| Usp31 | 107,76 | -0,2724 | 0,586201597 | chr7 | - |
| Usp35 | 17,16 | -0,5174 | 0,413537476 | chr7 | - |
| Usp47 | 1641,76 | -0,2475 | 0,434117948 | chr7 | + |
| Uvrag | 433,12 | -0,1731 | 0,635835796 | chr7 | - |
| Vasp | 200,07 | 0,3126 | 0,414291323 | chr7 | - |
| Selenos | 1314,28 | -0,2308 | 0,638888255 | chr7 | + |
| Vkorc1 | 3334,66 | -0,5260 | 0,245392711 | chr7 | - |
| Vmn1r58 | 180,66 | 0,1334 | 0,762940356 | chr7 | - |
| Vmn1r65 | 53,70 | -0,8247 | 0,068187551 | chr7 | - |
| Vmn2r29 | 18,44 | 0,4399 | 0,453300339 | chr7 | - |
| Vmn2r57 | 3,01 | 0,3458 | 0,629873977 | chr7 | - |
| Vmn2r78 | 1,78 | 0,6548 | NA | chr7 | + |
| Vps33b | 109,72 | -0,1374 | 0,793250646 | chr7 | + |
| Vrk3 | 364,55 | -0,1001 | 0,780680545 | chr7 | + |
| Vsig10l | 5,17 | -0,2999 | 0,704347771 | chr7 | + |
| Wdr11 | 450,71 | 0,1284 | 0,763580116 | chr7 | + |
| Wdr62 | 20,74 | -0,5362 | 0,372630598 | chr7 | - |
| Wdr73 | 268,89 | 0,3058 | 0,417918742 | chr7 | - |
| Wdr93 | 23,50 | 0,5867 | 0,277831408 | chr7 | + |
| Wee1 | 249,17 | -0,2490 | 0,572666827 | chr7 | + |
| Whamm | 460,20 | -0,1604 | 0,651796722 | chr7 | + |
| Wtip | 765,80 | -0,0993 | 0,813323745 | chr7 | - |
| Xndc1 | 23,46 | 0,3897 | 0,523898331 | chr7 | + |
| Xpo6 | 277,77 | -0,2407 | 0,437620234 | chr7 | - |
| Xrcc1 | 78,13 | -0,3187 | 0,51722851 | chr7 | + |
| Yif1b | 389,17 | -0,3198 | 0,558489002 | chr7 | + |
| Ypel3 | 313,53 | -0,5805 | 0,135056018 | chr7 | + |
| Zbtb32 | 3,37 | 0,0814 | 0,933514578 | chr7 | - |
| Zbtb45 | 6,87 | 0,0079 | 0,994106851 | chr7 | - |
| Zc3h4 | 218,21 | 0,1126 | 0,77333268 | chr7 | + |
| Zdhhc13 | 139,26 | 0,0246 | 0,972238068 | chr7 | + |
| Zfand6 | 2008,52 | -0,3023 | 0,482516058 | chr7 | - |
| Zfp108 | 27,84 | 0,4373 | 0,423089018 | chr7 | + |
| Zfp109 | 22,21 | 0,8991 | 0,067543992 | chr7 | - |
| Zfp110 | 659,33 | 0,4215 | 0,099591501 | chr7 | + |
| Zfp111 | 70,55 | 0,6998 | 0,075163457 | chr7 | - |
| Zfp112 | 13,46 | 0,5199 | 0,401382558 | chr7 | + |
| Zfp128 | 37,12 | 0,2172 | 0,710225448 | chr7 | + |
| Zfp14 | 23,45 | -0,2110 | 0,761282414 | chr7 | - |
| Zfp143 | 86,79 | 0,2863 | 0,516624153 | chr7 | + |
| Zfp146 | 219,99 | 0,2165 | 0,596546314 | chr7 | - |
| Zfp180 | 280,16 | -0,1234 | 0,764664236 | chr7 | + |
| Zfp235 | 67,52 | 0,3723 | 0,417918742 | chr7 | + |
| Zfp260 | 597,52 | 0,2796 | 0,436827862 | chr7 | + |
| Zfp27 | 71,13 | 0,3423 | 0,465196538 | chr7 | - |
| Zfp28 | 27,46 | 0,4720 | 0,391807184 | chr7 | + |
| Zfp296 | 3,15 | 0,0275 | 0,979212524 | chr7 | + |
| Zfp30 | 29,75 | 0,5096 | 0,314431944 | chr7 | + |

| | | | | | |
|---|---|---|---|---|---|
| Zfp324 | 51,16 | 0,3015 | 0,515973336 | chr7 | + |
| Zfp329 | 89,63 | 0,4523 | 0,228728627 | chr7 | - |
| Zfp36 | 611,29 | -0,3313 | 0,483882557 | chr7 | - |
| Zfp382 | 2,24 | 0,0885 | NA | chr7 | + |
| Zfp383 | 71,32 | 0,1705 | 0,737575031 | chr7 | + |
| Zfp418 | 30,35 | 0,4880 | 0,352135503 | chr7 | + |
| Zfp420 | 51,21 | 0,1669 | 0,776467957 | chr7 | + |
| Zfp428 | 7,34 | -0,5284 | 0,439137219 | chr7 | + |
| Zfp444 | 146,59 | -0,0884 | 0,886603157 | chr7 | + |
| Zfp446 | 31,49 | 0,1336 | 0,843934425 | chr7 | + |
| Zfp507 | 164,92 | 0,2706 | 0,423149698 | chr7 | - |
| Zfp524 | 136,85 | -0,5960 | 0,099333119 | chr7 | + |
| Zfp551 | 13,11 | 0,7410 | 0,212049289 | chr7 | - |
| Zfp553 | 75,58 | -0,0709 | 0,910351158 | chr7 | + |
| Zfp566 | 9,61 | -0,1036 | 0,909889201 | chr7 | - |
| Zfp568 | 205,43 | -0,0829 | 0,849424132 | chr7 | + |
| Zfp574 | 89,82 | -0,1603 | 0,830330662 | chr7 | + |
| Zfp579 | 31,29 | -0,0573 | 0,949479116 | chr7 | - |
| Zfp580 | 4,20 | 0,3461 | 0,636759527 | chr7 | + |
| Zfp583 | 4,47 | 0,2609 | 0,748519887 | chr7 | - |
| Zfp59 | 73,16 | 0,3185 | 0,469528266 | chr7 | + |
| Zfp60 | 201,12 | 0,2663 | 0,549296977 | chr7 | + |
| Zfp606 | 134,22 | 0,3448 | 0,342841028 | chr7 | + |
| Zfp607a | 37,24 | 0,2974 | 0,57838597 | chr7 | + |
| Zfp61 | 6,94 | 0,2110 | 0,804380135 | chr7 | - |
| Zfp619 | 18,43 | 0,0660 | 0,941754349 | chr7 | + |
| Zfp628 | 33,36 | -0,4280 | 0,520159157 | chr7 | + |
| Zfp629 | 153,44 | 0,1707 | 0,736302391 | chr7 | - |
| Zfp646 | 148,41 | -0,1814 | 0,636447281 | chr7 | + |
| Zfp658 | 21,10 | -0,1164 | 0,880273946 | chr7 | + |
| Zfp668 | 21,32 | -0,4471 | 0,485983895 | chr7 | - |
| Zfp688 | 32,18 | -0,6422 | 0,246948142 | chr7 | - |
| Zfp689 | 24,29 | -0,1328 | 0,869442758 | chr7 | - |
| Zfp710 | 126,85 | 0,0771 | 0,88569455 | chr7 | + |
| Zfp715 | 218,88 | 0,0824 | 0,854690386 | chr7 | - |
| Zfp74 | 52,14 | 0,2351 | 0,672907874 | chr7 | - |
| Zfp747 | 49,51 | -0,2669 | 0,580218822 | chr7 | - |
| Zfp764 | 26,64 | 0,2975 | 0,625823055 | chr7 | - |
| Zfp768 | 132,41 | -0,1222 | 0,841053685 | chr7 | - |
| Zfp771 | 58,82 | -0,6872 | 0,170659515 | chr7 | + |
| Zfp772 | 53,65 | -0,0006 | 0,998994678 | chr7 | - |
| Zfp780b | 116,18 | 0,5619 | 0,073845576 | chr7 | - |
| Zfp784 | 33,60 | 0,1078 | 0,876803328 | chr7 | - |
| Zfp787 | 234,45 | -0,0769 | 0,906333899 | chr7 | - |
| Zfp788 | 91,31 | 0,5231 | 0,217231457 | chr7 | + |
| Zfp790 | 194,13 | 0,1270 | 0,775458257 | chr7 | + |
| Zfp82 | 6,59 | 0,0221 | 0,98489528 | chr7 | - |
| Zfp865 | 40,69 | -0,6790 | 0,191230937 | chr7 | + |
| Zfp93 | 70,96 | 0,5848 | 0,119147532 | chr7 | + |
| Zfp939 | 23,33 | 0,2977 | 0,646093772 | chr7 | + |

| | | | | | |
|---|---|---|---|---|---|
| Zfp94 | 22,97 | -0,1726 | 0,827498825 | chr7 | - |
| Zfp940 | 16,54 | -0,8306 | 0,136585635 | chr7 | - |
| Zfp954 | 169,87 | 0,0432 | 0,932815385 | chr7 | - |
| Zranb1 | 840,40 | -0,1444 | 0,66525601 | chr7 | + |
| Zscan2 | 25,68 | -0,3009 | 0,640948606 | chr7 | + |
| Zscan22 | 58,20 | 0,1607 | 0,769969083 | chr7 | + |
| 1700030J22Rik | 5,63 | -0,2935 | 0,712967109 | chr8 | - |
| 1700030K09Rik | 55,18 | 0,0240 | 0,974449819 | chr8 | + |
| 1700067K01Rik | 3,15 | -0,1213 | 0,894408669 | chr8 | + |
| Tcim | 740,48 | 0,4551 | 0,403301059 | chr8 | - |
| 2010320M18Rik | 156,43 | -0,2756 | 0,526620417 | chr8 | + |
| 2310022B05Rik | 146,32 | 0,2388 | 0,550064422 | chr8 | - |
| Trir | 465,02 | -0,5321 | 0,23347764 | chr8 | + |
| Borcs8 | 235,94 | -0,4126 | 0,286763659 | chr8 | + |
| 2610005L07Rik | 115,72 | -0,0581 | 0,93551479 | chr8 | - |
| 2610044O15Rik8 | 120,13 | 0,6015 | 0,095464405 | chr8 | - |
| Hpf1 | 146,93 | -0,0033 | 0,995750189 | chr8 | + |
| 2810004N23Rik | 328,45 | -0,0310 | 0,953911274 | chr8 | - |
| 2810013P06Rik | 95,55 | -0,2963 | 0,47939388 | chr8 | + |
| Rex1bd | 438,02 | -0,3345 | 0,665144558 | chr8 | - |
| Lncbate1 | 4,97 | 0,5136 | 0,450460396 | chr8 | - |
| 3930402G23Rik | 7,99 | -0,5048 | 0,466451498 | chr8 | - |
| 4921524J17Rik | 215,88 | 0,5219 | 0,064539659 | chr8 | - |
| 4930513N10Rik | 2,54 | 0,0602 | NA | chr8 | + |
| 4930563M20Rik | 2,19 | 0,5167 | NA | chr8 | - |
| 4931428F04Rik | 26,75 | 0,3613 | 0,53757231 | chr8 | - |
| Cfap97 | 290,77 | 0,2879 | 0,371011913 | chr8 | + |
| 4933417D19Rik | 2,05 | -0,0649 | NA | chr8 | + |
| 6430548M08Rik | 83,48 | 0,0107 | 0,987508343 | chr8 | + |
| 6430573F11Rik | 35,29 | -0,1657 | 0,834675238 | chr8 | + |
| 6820431F20Rik | 979,53 | 0,0192 | 0,975879219 | chr8 | - |
| 9330133O14Rik | 14,05 | 0,1169 | 0,897817474 | chr8 | + |
| 9330175E14Rik | 3,57 | -0,0721 | 0,941553259 | chr8 | - |
| 9430091E24Rik | 7,01 | -0,5239 | 0,444432548 | chr8 | - |
| A230103J11Rik | 2,15 | 0,2065 | NA | chr8 | - |
| A430078G23Rik | 25,85 | -0,6895 | 0,192736991 | chr8 | + |
| A930013F10Rik | 7,64 | -0,5201 | 0,44242629 | chr8 | - |
| Aadat | 3790,06 | -0,2443 | 0,603818311 | chr8 | + |
| Aars | 2477,35 | -0,4507 | 0,235355755 | chr8 | + |
| Abce1 | 1736,73 | -0,1239 | 0,791579157 | chr8 | - |
| Abhd13 | 467,46 | -0,1135 | 0,780418568 | chr8 | + |
| Abhd8 | 34,75 | -0,3941 | 0,564724076 | chr8 | - |
| Acd | 119,08 | -0,3676 | 0,298116176 | chr8 | - |
| Acsf3 | 279,63 | 0,0732 | 0,894408669 | chr8 | + |
| Acsl1 | 13509,32 | -0,4045 | 0,273018284 | chr8 | + |
| Acta1 | 5,01 | 0,1079 | NA | chr8 | - |
| Adam32 | 2,56 | -0,3490 | NA | chr8 | - |
| Adam9 | 1210,46 | -0,1379 | 0,66529561 | chr8 | - |
| Adat1 | 60,61 | -0,1489 | 0,793250646 | chr8 | - |
| Adcy7 | 60,25 | 0,4442 | 0,34427921 | chr8 | + |

| | | | | | |
|---|---|---|---|---|---|
| Adrb3 | 162,74 | 0,0171 | 0,98507181 | chr8 | - |
| Afg3l1 | 872,53 | -0,2476 | 0,404889595 | chr8 | + |
| Aga | 452,95 | 0,2295 | 0,500489603 | chr8 | + |
| Agpat5 | 239,04 | -0,2097 | 0,57626953 | chr8 | + |
| Agt | 8988,58 | -0,0623 | 0,887518458 | chr8 | - |
| AI429214 | 24,63 | 0,7214 | 0,211038545 | chr8 | + |
| Aktip | 249,30 | -0,2793 | 0,355790779 | chr8 | - |
| Alg11 | 718,83 | 0,1845 | 0,598837173 | chr8 | + |
| Amfr | 4530,63 | -0,3018 | 0,27946684 | chr8 | - |
| Anapc10 | 109,53 | 0,2137 | 0,680674843 | chr8 | + |
| Angpt2 | 29,35 | 0,1836 | 0,803874753 | chr8 | - |
| Ankle1 | 11,87 | -0,2186 | 0,784034683 | chr8 | + |
| Ankrd10 | 106,61 | -0,3253 | 0,451182001 | chr8 | - |
| Ankrd11 | 286,94 | -0,0111 | 0,986818215 | chr8 | - |
| Ankrd37 | 21,49 | -0,1265 | 0,883681865 | chr8 | - |
| Ano8 | 31,61 | 0,2597 | 0,701007919 | chr8 | - |
| Ap1g1 | 1359,98 | -0,0409 | 0,920651925 | chr8 | + |
| Ap1m1 | 316,90 | 0,2001 | 0,685587331 | chr8 | + |
| Ap3m2 | 47,47 | 0,4952 | 0,258561012 | chr8 | - |
| Aprt | 523,54 | -0,2144 | 0,628656316 | chr8 | - |
| Arglu1 | 457,29 | -0,4404 | 0,12234836 | chr8 | - |
| Arhgap10 | 24,78 | 0,1608 | 0,817848245 | chr8 | - |
| Arhgef10 | 20,83 | 0,4936 | 0,399118497 | chr8 | + |
| Arhgef18 | 245,17 | -0,1642 | 0,712238951 | chr8 | + |
| Arhgef7 | 227,60 | 0,1393 | 0,723157997 | chr8 | + |
| Arl2bp | 98,39 | -0,3059 | 0,60974348 | chr8 | + |
| Armc6 | 64,70 | -0,2791 | 0,654151849 | chr8 | - |
| Arrdc2 | 121,03 | -0,5912 | 0,131674744 | chr8 | - |
| Arv1 | 22,91 | 0,3139 | 0,627447666 | chr8 | + |
| Asah1 | 879,90 | 0,4736 | 0,193343482 | chr8 | - |
| Asf1b | 15,22 | -0,2646 | 0,703944394 | chr8 | + |
| Ash2l | 250,51 | -0,2625 | 0,492833155 | chr8 | - |
| Asna1 | 359,56 | -0,4550 | 0,12746076 | chr8 | - |
| Atmin | 110,91 | -0,0081 | 0,98947313 | chr8 | + |
| Atp11a | 142,48 | 0,3767 | 0,528985529 | chr8 | + |
| Atp13a1 | 387,89 | -0,6265 | 0,066396922 | chr8 | + |
| Atp6v0d1 | 778,83 | -0,5304 | 0,107246729 | chr8 | - |
| Atp6v1b2 | 727,28 | 0,1641 | 0,629895923 | chr8 | + |
| AY512931 | 13,05 | -0,1177 | 0,894984867 | chr8 | - |
| B3gnt3 | 14,06 | 0,1326 | 0,879201267 | chr8 | - |
| B3gnt9 | 13,84 | 0,4323 | 0,535041318 | chr8 | - |
| Babam1 | 724,40 | -0,3849 | 0,308448866 | chr8 | + |
| Bag4 | 531,29 | -0,2196 | 0,581164752 | chr8 | - |
| Banp | 126,93 | -0,0701 | 0,931864162 | chr8 | + |
| Bbs2 | 34,28 | -0,0304 | 0,969195493 | chr8 | - |
| Map3k21 | 114,95 | -0,1142 | 0,87618436 | chr8 | + |
| Prr36 | 10,03 | 0,1548 | 0,860890261 | chr8 | - |
| Bcar1 | 238,67 | -0,6020 | 0,227023515 | chr8 | - |
| Bco1 | 175,69 | -0,5573 | 0,149780096 | chr8 | + |
| Brd7 | 514,97 | -0,0868 | 0,843329386 | chr8 | - |

| | | | | | |
|---|---|---|---|---|---|
| Brf2 | 43,31 | 0,4187 | 0,470066395 | chr8 | - |
| Bst2 | 3424,58 | -0,1500 | 0,818573937 | chr8 | - |
| C330021F23Rik | 54,00 | 0,4302 | 0,501951423 | chr8 | + |
| Cacna1a | 17,47 | 0,4353 | 0,550044971 | chr8 | + |
| Calr | 11718,85 | -0,1861 | 0,673822592 | chr8 | - |
| Calr3 | 48,42 | 0,0176 | 0,981991128 | chr8 | - |
| Camsap3 | 109,78 | -0,6989 | 0,071771252 | chr8 | + |
| Car5a | 963,02 | -0,0620 | 0,891830745 | chr8 | - |
| Car7 | 10,44 | 0,4181 | 0,566893799 | chr8 | + |
| Naxd | 749,14 | -0,4095 | 0,147022766 | chr8 | + |
| Cars2 | 143,36 | -0,4747 | 0,189223209 | chr8 | - |
| Casp3 | 644,82 | 0,0614 | 0,88882432 | chr8 | + |
| Cbfa2t3 | 24,45 | -0,0488 | 0,953751362 | chr8 | - |
| Cbfb | 298,67 | 0,1777 | 0,576265028 | chr8 | + |
| Cbr4 | 725,50 | -0,0712 | 0,872238089 | chr8 | + |
| Cc2d1a | 83,40 | -0,3648 | 0,540815736 | chr8 | - |
| Ccdc102a | 8,17 | 0,0322 | 0,9766348 | chr8 | - |
| Ccdc124 | 363,15 | -0,5685 | 0,293001163 | chr8 | - |
| Ccdc130 | 62,27 | -0,1302 | 0,811137679 | chr8 | - |
| Ccl17 | 1,99 | 0,7018 | NA | chr8 | + |
| Ccl22 | 5,71 | 0,1179 | 0,899463828 | chr8 | + |
| Ccl25 | 69,23 | -0,3457 | 0,460634874 | chr8 | + |
| Ccsap | 2,87 | 0,1632 | 0,844155615 | chr8 | - |
| Cd209a | 13,01 | 0,7087 | 0,260767757 | chr8 | - |
| Cd209b | 9,26 | 0,1470 | 0,869067375 | chr8 | - |
| Cd209d | 2,72 | 0,5010 | 0,435857678 | chr8 | - |
| Cd209f | 14,37 | 0,7591 | 0,22384344 | chr8 | - |
| Cd209g | 2,62 | 0,3955 | 0,524328995 | chr8 | + |
| Adgre5 | 262,85 | -0,0658 | 0,906998011 | chr8 | - |
| Cdc16 | 406,89 | -0,0607 | 0,895961198 | chr8 | + |
| Cdh11 | 19,66 | 0,2047 | 0,770917245 | chr8 | - |
| Cdh13 | 20,32 | 0,2709 | 0,681592614 | chr8 | + |
| Cdh3 | 5,62 | 0,3093 | 0,693242675 | chr8 | + |
| Cdh5 | 231,23 | 0,2441 | 0,563218321 | chr8 | + |
| Cdk10 | 246,77 | -0,0123 | 0,984931953 | chr8 | + |
| Cdkn2aip | 250,08 | 0,4229 | 0,162891445 | chr8 | - |
| Cdt1 | 30,08 | -0,0766 | 0,925898178 | chr8 | + |
| Cdyl2 | 42,44 | -0,4463 | 0,341412029 | chr8 | - |
| Cenpn | 11,69 | -0,1426 | 0,869067375 | chr8 | + |
| Cenpt | 20,95 | -0,0862 | 0,924557072 | chr8 | - |
| Cenpu | 7,70 | -0,1120 | 0,903544399 | chr8 | + |
| Cep44 | 140,44 | -0,0739 | 0,888917253 | chr8 | - |
| Cers4 | 41,25 | 0,0393 | 0,961098177 | chr8 | + |
| Ces1a | 5,47 | -0,0912 | 0,92522408 | chr8 | - |
| Ces1b | 5493,65 | -0,4811 | 0,497298938 | chr8 | - |
| Ces1c | 47464,89 | -0,3069 | 0,355918478 | chr8 | - |
| Ces1d | 15772,16 | -0,6687 | 0,093882396 | chr8 | - |
| Ces1f | 13148,13 | -0,6087 | 0,096308851 | chr8 | - |
| Ces1g | 3797,40 | 0,0791 | 0,912746031 | chr8 | - |
| Ces2b | 180,84 | -0,3039 | 0,66209482 | chr8 | + |

| | | | | | |
|---|---|---|---|---|---|
| Ces2c | 874,52 | 0,2317 | 0,704789011 | chr8 | + |
| Ces2d-ps | 9,45 | -0,0334 | 0,976155242 | chr8 | + |
| Ces2e | 6680,35 | -0,0329 | 0,950539822 | chr8 | + |
| Ces2g | 818,08 | -0,3245 | 0,308229345 | chr8 | + |
| Ces2h | 3,34 | 0,0673 | 0,944511666 | chr8 | + |
| Ces3b | 9256,30 | -0,3642 | 0,320153548 | chr8 | + |
| Ces4a | 102,06 | -1,6733 | NA | chr8 | + |
| Cfdp1 | 1208,07 | -0,0539 | 0,89130564 | chr8 | - |
| Chd9 | 379,04 | 0,0310 | 0,955234619 | chr8 | + |
| Cherp | 175,46 | -0,3603 | 0,330791426 | chr8 | - |
| Chmp1a | 871,16 | -0,2213 | 0,552474789 | chr8 | - |
| Chst4 | 2,22 | 0,0617 | NA | chr8 | - |
| Chtf8 | 593,40 | -0,0328 | 0,943064069 | chr8 | - |
| Ciapin1 | 689,25 | -0,2327 | 0,716547565 | chr8 | - |
| Utp4 | 280,71 | -0,2871 | 0,367343436 | chr8 | + |
| Ckap2 | 13,80 | 0,4115 | 0,564495903 | chr8 | - |
| Cklf | 17,79 | -0,4894 | 0,401255942 | chr8 | + |
| Clcn3 | 880,07 | 0,0832 | 0,877942095 | chr8 | - |
| Cldn23 | 2,03 | 0,6799 | NA | chr8 | - |
| Clec4g | 1992,60 | 0,0341 | 0,938131508 | chr8 | - |
| Cln8 | 818,79 | 0,0781 | 0,885187516 | chr8 | + |
| Cmip | 328,24 | -0,3708 | 0,29271782 | chr8 | + |
| Cmtm3 | 148,26 | 0,0810 | 0,87246283 | chr8 | + |
| Cmtm4 | 716,34 | -0,1882 | 0,626400494 | chr8 | - |
| Cmtr2 | 61,90 | 0,3066 | 0,502231951 | chr8 | + |
| Cnep1r1 | 316,43 | 0,3145 | 0,37928598 | chr8 | + |
| Cngb1 | 2,01 | 0,2522 | NA | chr8 | - |
| Cnot1 | 1314,57 | -0,0673 | 0,873325649 | chr8 | - |
| Cnot7 | 466,25 | -0,0715 | 0,869442758 | chr8 | - |
| Coa6 | 379,92 | -0,0529 | 0,943064069 | chr8 | + |
| Cog2 | 291,48 | 0,0254 | 0,963845098 | chr8 | + |
| Cog4 | 1415,76 | 0,0592 | 0,899853759 | chr8 | + |
| Cog8 | 153,52 | -0,2515 | 0,562418905 | chr8 | - |
| Col4a1 | 513,89 | 0,2288 | 0,581990132 | chr8 | - |
| Col4a2 | 381,45 | 0,0436 | 0,943108135 | chr8 | + |
| Cope | 1269,61 | -0,4391 | 0,217231457 | chr8 | + |
| Coprs | 38,53 | -0,7845 | 0,119628533 | chr8 | - |
| Coq9 | 1464,75 | -0,1234 | 0,730989713 | chr8 | + |
| Cotl1 | 152,66 | 0,0537 | 0,941553175 | chr8 | - |
| Cpe | 11,97 | 0,7587 | 0,217231457 | chr8 | - |
| Cpne2 | 102,46 | -0,4612 | 0,337280658 | chr8 | + |
| Crispld2 | 16,28 | 0,5405 | 0,398198084 | chr8 | + |
| Crtc1 | 62,12 | -0,1140 | 0,854019408 | chr8 | - |
| Csgalnact1 | 14,65 | 0,4181 | 0,522768066 | chr8 | - |
| Csnk2a2 | 194,18 | -0,4676 | 0,135427143 | chr8 | - |
| Ctcf | 417,75 | 0,2269 | 0,524298814 | chr8 | + |
| Ctrl | 3,55 | -0,2184 | 0,794928847 | chr8 | - |
| Ctu2 | 73,64 | -0,5404 | 0,295666715 | chr8 | + |
| Ctxn1 | 4,18 | 0,1071 | 0,90870562 | chr8 | - |
| Cul4a | 1451,31 | -0,0735 | 0,854019408 | chr8 | + |

| | | | | | |
|---|---|---|---|---|---|
| Cx3cl1 | 13,61 | 0,3947 | 0,578442407 | chr8 | + |
| Cyba | 143,63 | 0,1376 | 0,814156035 | chr8 | - |
| Cyld | 431,73 | 0,0336 | 0,952576171 | chr8 | + |
| Cyp4f18 | 7,44 | 1,0574 | 0,064088986 | chr8 | - |
| Cyp4v3 | 13520,10 | 0,2525 | 0,46846179 | chr8 | - |
| D130040H23Rik | 4,27 | 0,6681 | 0,298116176 | chr8 | + |
| D230025D16Rik | 852,36 | -0,5508 | 0,119147532 | chr8 | + |
| D8Ertd738e | 1402,85 | -0,4554 | 0,426572731 | chr8 | - |
| Prag1 | 6,15 | 0,1421 | 0,87623846 | chr8 | + |
| Dand5 | 32,03 | -0,7700 | 0,080271426 | chr8 | - |
| Dcaf15 | 59,95 | -0,0107 | 0,990115234 | chr8 | - |
| Dctn6 | 235,32 | 0,1715 | 0,633566893 | chr8 | - |
| Dcun1d2 | 235,32 | -0,0963 | 0,823578308 | chr8 | - |
| Dda1 | 166,58 | -0,5460 | 0,223737597 | chr8 | + |
| Ddhd2 | 484,86 | -0,0874 | 0,833053102 | chr8 | - |
| Ddx19a | 188,14 | -0,1510 | 0,72678231 | chr8 | - |
| Ddx19b | 374,20 | -0,1278 | 0,797253751 | chr8 | - |
| Ddx28 | 206,99 | -0,2774 | 0,470730504 | chr8 | - |
| Ddx39 | 204,97 | -0,3495 | 0,376984775 | chr8 | + |
| Ddx49 | 234,10 | -0,5908 | 0,093044072 | chr8 | - |
| Ddx60 | 159,81 | 0,8934 | 0,053192858 | chr8 | + |
| Def8 | 712,04 | 0,1881 | 0,70245129 | chr8 | + |
| Dhodh | 111,98 | -0,1445 | 0,754913678 | chr8 | - |
| Dhps | 104,13 | -0,2681 | 0,602938667 | chr8 | + |
| Dhx38 | 183,40 | -0,0362 | 0,9455627 | chr8 | - |
| Dkk4 | 10,11 | 0,2952 | 0,703067947 | chr8 | + |
| Dlc1 | 626,67 | -0,2253 | 0,590058791 | chr8 | - |
| Dnaja2 | 2583,37 | -0,4037 | 0,169238731 | chr8 | - |
| Dnajb1 | 427,75 | -0,4738 | 0,208173201 | chr8 | + |
| Dnase2a | 227,88 | -0,4217 | 0,183137332 | chr8 | + |
| Dok4 | 22,98 | 0,0245 | 0,978280656 | chr8 | - |
| Dpep1 | 20,90 | 0,6598 | 0,258301555 | chr8 | + |
| Dus2 | 124,97 | 0,0020 | 0,997464504 | chr8 | + |
| Dusp4 | 10,23 | -0,6423 | 0,323355495 | chr8 | + |
| Dync1li2 | 474,45 | 0,1303 | 0,692167613 | chr8 | - |
| E2f4 | 142,40 | -0,2410 | 0,577069333 | chr8 | + |
| Edc4 | 172,17 | -0,3141 | 0,396254007 | chr8 | + |
| Ednra | 46,31 | 0,0052 | 0,994765486 | chr8 | - |
| Efnb2 | 55,81 | 0,0088 | 0,990189789 | chr8 | - |
| Egln1 | 366,29 | -0,4628 | 0,107982926 | chr8 | - |
| Eif4ebp1 | 621,62 | -0,4748 | 0,283414267 | chr8 | + |
| Elavl1 | 481,98 | -0,2199 | 0,4345327 | chr8 | - |
| Ell | 302,45 | -0,2447 | 0,592046274 | chr8 | + |
| Elmo3 | 291,81 | -0,1817 | 0,625365358 | chr8 | + |
| Elmod2 | 383,13 | 0,5613 | 0,118961036 | chr8 | - |
| Emc8 | 277,20 | -0,4776 | 0,157491037 | chr8 | - |
| Enkd1 | 3,92 | -0,1049 | 0,907233333 | chr8 | - |
| Enpp6 | 18,28 | -0,2202 | 0,766288203 | chr8 | + |
| Eps15l1 | 145,80 | -0,2488 | 0,592295685 | chr8 | - |
| Eri1 | 262,35 | 0,2210 | 0,576265028 | chr8 | - |

| | | | | | |
|---|---|---|---|---|---|
| Erich1 | 38,22 | 0,0011 | 0,998482911 | chr8 | - |
| Erlin2 | 475,32 | 0,1780 | 0,561773364 | chr8 | + |
| Esrp2 | 455,33 | 0,0872 | 0,865418807 | chr8 | - |
| Evi5l | 13,29 | 0,0432 | 0,966233054 | chr8 | + |
| Exoc3l | 27,94 | -0,4146 | 0,470196942 | chr8 | - |
| Exoc8 | 131,52 | 0,1027 | 0,823537093 | chr8 | - |
| Exosc6 | 44,81 | -0,1898 | 0,784505535 | chr8 | + |
| F11 | 1392,20 | -0,5433 | 0,076519476 | chr8 | - |
| F7 | 1659,59 | 0,0939 | 0,788002554 | chr8 | + |
| Fam129c | 7,89 | -0,2519 | 0,758743175 | chr8 | + |
| Fam149a | 315,00 | 0,1518 | 0,718060307 | chr8 | - |
| Fam192a | 299,25 | -0,2778 | 0,476631852 | chr8 | - |
| Fam32a | 1170,62 | -0,0813 | 0,869442758 | chr8 | + |
| Ripor1 | 101,75 | -0,3159 | 0,515706574 | chr8 | + |
| Fam89a | 41,53 | 0,6121 | 0,152130292 | chr8 | - |
| Fam96b | 222,98 | -0,1985 | 0,716359853 | chr8 | - |
| Fanca | 7,08 | -0,2022 | 0,813415373 | chr8 | - |
| Farsa | 205,67 | -0,3665 | 0,259978915 | chr8 | + |
| Fat1 | 255,16 | 0,3845 | 0,263245957 | chr8 | + |
| Fbxl8 | 27,24 | -0,1055 | 0,899463828 | chr8 | + |
| Fbxo25 | 114,99 | 0,0292 | 0,959145268 | chr8 | + |
| Fbxo8 | 996,68 | -0,1042 | 0,806401686 | chr8 | + |
| Fbxw9 | 265,37 | -0,1482 | 0,758077598 | chr8 | + |
| Fcer2a | 5,18 | -0,4581 | 0,52377705 | chr8 | - |
| Fcho1 | 2,36 | -0,2231 | NA | chr8 | - |
| Fendrr | 14,97 | -0,5893 | 0,354338694 | chr8 | - |
| Fgfr1 | 48,53 | -0,1221 | 0,841012567 | chr8 | + |
| Fgl1 | 12403,98 | 0,0143 | 0,986661276 | chr8 | - |
| Fhod1 | 39,05 | -0,1765 | 0,825895013 | chr8 | - |
| Fkbp8 | 2402,39 | -0,4235 | 0,423755863 | chr8 | + |
| Fnta | 643,28 | -0,2867 | 0,407858893 | chr8 | - |
| Foxf1 | 23,40 | 0,2132 | 0,761178054 | chr8 | + |
| Frem3 | 2,48 | 0,0215 | NA | chr8 | + |
| Frg1 | 349,37 | -0,0733 | 0,881044464 | chr8 | - |
| Fto | 332,39 | 0,1399 | 0,731081367 | chr8 | + |
| Fuk | 98,33 | -0,2836 | 0,449506818 | chr8 | - |
| Fut10 | 33,05 | 0,0869 | 0,918104715 | chr8 | + |
| Gab1 | 205,04 | -0,2118 | 0,52825088 | chr8 | - |
| Gabarapl2 | 1556,35 | -0,3494 | 0,202636202 | chr8 | + |
| Gadd45gip1 | 263,13 | -0,4675 | 0,376702038 | chr8 | + |
| Galns | 86,02 | -0,2000 | 0,741224458 | chr8 | - |
| Galnt2 | 369,56 | -0,1466 | 0,693803702 | chr8 | + |
| Galnt7 | 53,00 | -0,4811 | 0,277423579 | chr8 | - |
| Gan | 10,59 | -0,3785 | 0,611603353 | chr8 | + |
| Gas6 | 486,13 | 0,4090 | 0,293404423 | chr8 | - |
| Gas8 | 23,98 | 0,0804 | 0,920651925 | chr8 | + |
| Gatad2a | 444,65 | -0,4922 | 0,169162729 | chr8 | - |
| Gcdh | 10033,70 | -0,1905 | 0,449196578 | chr8 | - |
| Gcsh | 1687,26 | -0,3818 | 0,138593946 | chr8 | - |
| Gdf15 | 19,68 | -1,0300 | 0,072934191 | chr8 | - |

| | | | | | |
|---|---|---|---|---|---|
| Gfod2 | 74,69 | -0,1368 | 0,841150543 | chr8 | - |
| Gins2 | 13,30 | 0,2327 | 0,765588119 | chr8 | - |
| Gins3 | 16,76 | 0,6904 | 0,246966539 | chr8 | + |
| Gins4 | 323,85 | -0,3892 | 0,301759036 | chr8 | - |
| Gipc1 | 125,33 | -0,1564 | 0,823537093 | chr8 | + |
| Colgalt1 | 647,29 | 0,2292 | 0,544838491 | chr8 | + |
| Gm10033 | 97,51 | -0,2206 | 0,702177047 | chr8 | - |
| Tgfbr3l | 18,82 | -0,0130 | 0,990189789 | chr8 | + |
| Gm15348 | 8,59 | 0,2379 | 0,658257248 | chr8 | - |
| Gm16675 | 12,95 | -0,0320 | 0,976116181 | chr8 | - |
| Gm16793 | 31,37 | -0,5332 | 0,375032546 | chr8 | - |
| Gm1943 | 80,60 | -0,1873 | 0,680798323 | chr8 | - |
| Gm3336 | 2,61 | 0,0783 | 0,931322892 | chr8 | + |
| Gm4890 | 17,15 | -0,3580 | 0,615542082 | chr8 | + |
| Sox1ot | 13,67 | -0,0873 | 0,922849883 | chr8 | + |
| Gm5741 | 3,79 | -0,4656 | 0,484348421 | chr8 | - |
| Gmip | 41,76 | -0,1506 | 0,802329508 | chr8 | + |
| Gnpat | 607,91 | -0,1310 | 0,680276822 | chr8 | + |
| Golga7 | 1115,83 | -0,0616 | 0,885187516 | chr8 | - |
| Got2 | 8592,63 | -0,0464 | 0,895961198 | chr8 | - |
| Gpm6a | 36,10 | 0,6443 | 0,212649167 | chr8 | + |
| Adgrg5 | 4,77 | -0,3103 | 0,681592614 | chr8 | + |
| Adgra2 | 86,28 | 0,2705 | 0,55911685 | chr8 | + |
| Adgrg1 | 56,17 | 0,5762 | 0,206933423 | chr8 | + |
| Adgrg3 | 86,83 | 0,1808 | 0,742570557 | chr8 | + |
| Gpt2 | 4480,98 | -0,3890 | 0,348682374 | chr8 | + |
| Grtp1 | 357,71 | -0,3523 | 0,41612591 | chr8 | - |
| Gse1 | 107,75 | 0,0809 | 0,88569455 | chr8 | + |
| Gsr | 867,35 | -0,3703 | 0,156728357 | chr8 | + |
| Gtf2e2 | 258,54 | 0,1125 | 0,807623197 | chr8 | + |
| Cfap20 | 421,69 | -0,0683 | 0,880695141 | chr8 | - |
| Gtpbp3 | 94,71 | 0,0230 | 0,9774226 | chr8 | + |
| Hand2 | 33,88 | -0,0335 | 0,969512928 | chr8 | + |
| Haus8 | 63,60 | -0,0834 | 0,899497201 | chr8 | - |
| Heatr3 | 407,65 | -0,1761 | 0,666348577 | chr8 | + |
| Herpud1 | 4911,30 | -0,5603 | 0,08467489 | chr8 | + |
| Hgsnat | 804,98 | -0,1240 | 0,731048653 | chr8 | - |
| Hhip | 44,64 | 0,4379 | 0,381601977 | chr8 | - |
| Hmgb2 | 189,47 | 0,5500 | 0,070271273 | chr8 | + |
| Hmgxb4 | 116,61 | 0,6134 | 0,150902828 | chr8 | + |
| Hmox1 | 423,91 | 0,0145 | 0,986692637 | chr8 | + |
| Homer3 | 20,10 | -0,2044 | 0,78753026 | chr8 | + |
| Hook2 | 269,39 | -0,6275 | 0,184634167 | chr8 | + |
| Hook3 | 866,72 | 0,0355 | 0,941553259 | chr8 | - |
| Hpgd | 2811,76 | 0,0394 | 0,939983299 | chr8 | + |
| Hsbp1 | 1603,95 | -0,2963 | 0,201904156 | chr8 | + |
| Hsdl1 | 142,62 | 0,1754 | 0,658779108 | chr8 | - |
| Hsf4 | 49,68 | -0,5639 | 0,284599004 | chr8 | + |
| Htra4 | 9,84 | 0,1749 | 0,842940743 | chr8 | - |
| Ido2 | 1284,37 | -0,0141 | 0,98507181 | chr8 | - |

| | | | | | |
|---|---|---|---|---|---|
| Ier2 | 231,42 | 0,1202 | 0,879708396 | chr8 | - |
| Ifi30 | 414,33 | 0,0239 | 0,969433358 | chr8 | - |
| Ikbkb | 566,27 | 0,0382 | 0,931322892 | chr8 | - |
| Il12rb1 | 12,78 | -0,9740 | 0,073845576 | chr8 | + |
| Il15 | 34,68 | 0,1337 | 0,859638359 | chr8 | - |
| Il27ra | 26,58 | -0,3170 | 0,599069696 | chr8 | - |
| Il34 | 29,56 | 0,3092 | 0,629968267 | chr8 | - |
| Ing1 | 226,76 | -0,2318 | 0,517431746 | chr8 | + |
| Ing2 | 129,91 | -0,3556 | 0,370803111 | chr8 | - |
| Inpp4b | 3,97 | 0,1201 | 0,894984867 | chr8 | + |
| Insr | 2333,13 | -0,0553 | 0,89879469 | chr8 | - |
| Ints10 | 126,34 | -0,2555 | 0,701094188 | chr8 | + |
| Irf8 | 102,26 | 0,0815 | 0,86940991 | chr8 | + |
| Irs2 | 28,60 | -0,2327 | 0,762940356 | chr8 | - |
| Irx3 | 2,83 | -0,0782 | 0,933235928 | chr8 | - |
| Ist1 | 1027,51 | 0,0162 | 0,979212524 | chr8 | - |
| Itfg1 | 1524,87 | -0,2933 | 0,359146889 | chr8 | - |
| Itgb1 | 3892,94 | 0,0214 | 0,963296265 | chr8 | + |
| Jak3 | 47,19 | -0,1857 | 0,768327709 | chr8 | + |
| Junb | 218,19 | 0,0774 | 0,932528423 | chr8 | - |
| Jund | 144,89 | -0,2587 | 0,709746407 | chr8 | + |
| Kars | 1040,17 | -0,0125 | 0,978062015 | chr8 | - |
| Kat6a | 308,61 | 0,1367 | 0,685329957 | chr8 | + |
| Katnb1 | 17,27 | 0,1834 | 0,824913702 | chr8 | + |
| Kbtbd11 | 26,58 | 0,6932 | 0,209412518 | chr8 | + |
| Kcnk1 | 9,23 | 0,0783 | 0,93807111 | chr8 | + |
| Kcnn1 | 41,28 | -0,5955 | 0,187813678 | chr8 | - |
| Kifc3 | 176,25 | 0,5842 | 0,182663462 | chr8 | - |
| Klf1 | 15,11 | 0,3121 | 0,682841407 | chr8 | + |
| Klf2 | 58,42 | -0,2853 | 0,597328449 | chr8 | + |
| Klhdc4 | 199,65 | -0,5041 | 0,118025078 | chr8 | - |
| Klhl2 | 512,37 | -0,3736 | 0,240317215 | chr8 | - |
| Klhl26 | 63,46 | 0,4347 | 0,34745302 | chr8 | - |
| Klhl36 | 25,09 | -0,1587 | 0,832117263 | chr8 | + |
| Klkb1 | 3967,81 | -0,0935 | 0,820554805 | chr8 | - |
| Kxd1 | 563,38 | -0,0867 | 0,824028899 | chr8 | - |
| Lamp1 | 11150,88 | -0,4081 | 0,051853248 | chr8 | + |
| Large1 | 113,43 | 0,6067 | 0,05081654 | chr8 | - |
| Lcat | 4958,37 | -0,3034 | 0,496449986 | chr8 | - |
| Ldhd | 2191,43 | -0,0266 | 0,959145268 | chr8 | - |
| Leprotl1 | 885,77 | -0,5351 | 0,068789213 | chr8 | - |
| Letm2 | 287,14 | -0,3941 | 0,219049665 | chr8 | - |
| Lig4 | 91,44 | -0,0938 | 0,880649922 | chr8 | - |
| Lonp2 | 5926,83 | -0,3784 | 0,141295617 | chr8 | + |
| Lonrf1 | 112,77 | -0,6555 | 0,067696785 | chr8 | - |
| Lpar2 | 15,32 | -0,4667 | 0,505015502 | chr8 | + |
| Lpl | 1374,12 | 0,6731 | 0,053748254 | chr8 | + |
| Lrp2bp | 49,16 | 0,2909 | 0,598404146 | chr8 | + |
| Lrrc25 | 39,25 | -0,2259 | 0,712696717 | chr8 | + |
| Lrrc29 | 38,22 | -0,4577 | 0,423089018 | chr8 | - |

| | | | | | |
|---|---|---|---|---|---|
| Lrrc8e | 6,59 | 0,3800 | 0,613341495 | chr8 | + |
| Lsm1 | 287,33 | -0,2626 | 0,594896067 | chr8 | + |
| Lsm4 | 215,48 | -0,3561 | 0,470066395 | chr8 | + |
| Lsm6 | 1055,28 | -0,3248 | 0,356028986 | chr8 | - |
| Lyl1 | 6,69 | 0,4506 | 0,528261235 | chr8 | + |
| Maf | 449,10 | -0,0311 | 0,952640742 | chr8 | - |
| Mak16 | 273,03 | 0,0619 | 0,882318627 | chr8 | - |
| Man2b1 | 1187,49 | -0,0824 | 0,840953217 | chr8 | + |
| Map1s | 26,17 | -0,1728 | 0,83796616 | chr8 | + |
| Map2k7 | 337,33 | -0,1344 | 0,766178949 | chr8 | + |
| Marveld3 | 20,82 | 0,6521 | 0,263412599 | chr8 | - |
| Mast1 | 5,00 | -0,4115 | 0,540877577 | chr8 | - |
| Mast3 | 261,50 | -0,0343 | 0,968644508 | chr8 | - |
| Mau2 | 355,60 | -0,0582 | 0,895663283 | chr8 | - |
| Mbtps1 | 659,96 | 0,0470 | 0,916162095 | chr8 | - |
| Mcemp1 | 4,37 | 0,6100 | 0,341009315 | chr8 | + |
| Mcf2l | 4,47 | 0,5590 | 0,402148509 | chr8 | + |
| Mcm5 | 49,98 | 0,0356 | 0,967188554 | chr8 | + |
| Mcph1 | 44,29 | 0,2849 | 0,593524501 | chr8 | + |
| Med26 | 38,76 | -0,7122 | 0,079251671 | chr8 | - |
| Mfap3l | 7,81 | -0,8506 | 0,152264308 | chr8 | + |
| Mfhas1 | 399,36 | -0,5363 | 0,117305849 | chr8 | + |
| Micu3 | 35,78 | 0,3002 | 0,588219472 | chr8 | + |
| Mlycd | 1137,86 | -0,2809 | 0,425493131 | chr8 | + |
| Mmaa | 319,18 | -0,1207 | 0,795812563 | chr8 | - |
| Mmp2 | 53,38 | 0,8486 | 0,06626141 | chr8 | + |
| Mon1b | 118,95 | 0,5229 | 0,112905626 | chr8 | + |
| Mphosph6 | 83,87 | 0,6300 | 0,175457889 | chr8 | - |
| Mpv17l2 | 116,94 | -0,3564 | 0,42509819 | chr8 | - |
| Mri1 | 100,99 | 0,0456 | 0,950795699 | chr8 | - |
| Mrpl34 | 325,41 | -0,6898 | 0,12746076 | chr8 | + |
| Mrps31 | 903,69 | -0,0443 | 0,92066291 | chr8 | + |
| Msmo1 | 4631,83 | 0,1191 | 0,865199339 | chr8 | - |
| Msr1 | 565,56 | 0,4412 | 0,250762967 | chr8 | - |
| Mt2 | 1388,30 | -0,7822 | 0,201435122 | chr8 | + |
| Mthfsd | 162,22 | -0,1037 | 0,838806042 | chr8 | - |
| Mtmr7 | 255,52 | 0,3756 | 0,544991268 | chr8 | - |
| Mtnr1a | 18,39 | 0,1334 | 0,87206597 | chr8 | + |
| Mtss1l | 21,97 | 0,2380 | 0,725433565 | chr8 | + |
| Mtus1 | 43,00 | 0,2591 | 0,649875139 | chr8 | - |
| Mvb12a | 557,16 | -0,5663 | 0,190942933 | chr8 | + |
| Mvd | 179,70 | -0,1777 | 0,788923604 | chr8 | - |
| Mylk3 | 10,31 | 0,1297 | 0,882124635 | chr8 | - |
| Myo9b | 233,59 | 0,0360 | 0,94647679 | chr8 | + |
| Myom2 | 125,65 | -0,1150 | 0,814990243 | chr8 | + |
| N4bp1 | 203,83 | -0,0036 | 0,995142284 | chr8 | - |
| Nacc1 | 244,42 | -0,0836 | 0,888763259 | chr8 | - |
| Nae1 | 495,31 | -0,0345 | 0,953911274 | chr8 | - |
| Naf1 | 55,12 | -0,5316 | 0,178782041 | chr8 | + |
| Nat1 | 57,82 | 0,6054 | 0,208201861 | chr8 | + |

| | | | | | |
|---|---|---|---|---|---|
| Nat2 | 501,74 | 0,2461 | 0,510690791 | chr8 | + |
| Ndrg4 | 3,75 | -0,2560 | 0,740352128 | chr8 | + |
| Ndufa13 | 3595,91 | -0,2563 | 0,658533075 | chr8 | - |
| Ndufb7 | 1543,96 | -0,3836 | 0,495256859 | chr8 | + |
| Neil3 | 2,48 | 0,1825 | NA | chr8 | - |
| Nek1 | 122,76 | 0,4272 | 0,331535469 | chr8 | + |
| Nek3 | 29,22 | 0,1148 | 0,8763719 | chr8 | - |
| Nek5 | 3,16 | 0,2616 | 0,716169774 | chr8 | - |
| Neto2 | 8,95 | 0,2539 | 0,75616611 | chr8 | - |
| Nfat5 | 409,46 | -0,4206 | 0,217231457 | chr8 | + |
| Nfatc3 | 421,94 | -0,3557 | 0,233524306 | chr8 | + |
| Nfix | 407,80 | -0,0675 | 0,903060276 | chr8 | - |
| Nip7 | 433,53 | -0,2044 | 0,581990132 | chr8 | + |
| Nkd1 | 38,27 | 0,1214 | 0,859896838 | chr8 | + |
| Nlrc5 | 205,22 | 0,6163 | 0,118025078 | chr8 | + |
| Nob1 | 161,49 | -0,1420 | 0,763514396 | chr8 | - |
| Nod2 | 4,14 | 0,2903 | 0,707969381 | chr8 | + |
| Nol3 | 19,56 | -0,2145 | 0,767809246 | chr8 | + |
| Npy1r | 5,24 | 0,2766 | 0,732392788 | chr8 | + |
| Nqo1 | 187,70 | 0,7103 | 0,10976595 | chr8 | - |
| Nr2c2ap | 229,90 | -0,4222 | 0,169622866 | chr8 | + |
| Nr2f6 | 455,85 | -0,4263 | 0,312403089 | chr8 | - |
| Nr3c2 | 69,63 | -0,1563 | 0,82176873 | chr8 | + |
| Nrp1 | 1887,31 | 0,3782 | 0,364595998 | chr8 | + |
| Ntpcr | 98,24 | 0,2360 | 0,581394499 | chr8 | + |
| Nudt21 | 261,67 | 0,1452 | 0,714657949 | chr8 | - |
| Nudt7 | 19444,33 | -0,5210 | 0,215046281 | chr8 | + |
| Nup133 | 182,75 | 0,1495 | 0,746751926 | chr8 | - |
| Nup93 | 67,49 | 0,1203 | 0,855245012 | chr8 | + |
| Nutf2 | 31,43 | 0,0381 | 0,962446025 | chr8 | + |
| Ocel1 | 108,61 | -0,2936 | 0,662842062 | chr8 | + |
| Ogfod1 | 204,27 | 0,0931 | 0,830451573 | chr8 | + |
| Olfr372 | 2,48 | 0,8833 | NA | chr8 | + |
| Orc6 | 107,60 | 0,0205 | 0,976528475 | chr8 | + |
| Otud4 | 597,30 | 0,1587 | 0,68001254 | chr8 | + |
| Palld | 88,56 | 0,4867 | 0,335264505 | chr8 | - |
| Papd5 | 604,69 | -0,1993 | 0,572283367 | chr8 | + |
| Pard3 | 742,89 | -0,3655 | 0,209104214 | chr8 | + |
| Pard6a | 52,05 | -0,2774 | 0,581725594 | chr8 | + |
| Pbx4 | 2,57 | 0,1992 | 0,767810431 | chr8 | + |
| Pcid2 | 126,59 | -0,0945 | 0,844155615 | chr8 | - |
| Pcm1 | 360,32 | 0,2341 | 0,505514525 | chr8 | + |
| Pde4c | 293,84 | -0,3151 | 0,564724076 | chr8 | + |
| Pdf | 321,12 | -0,4305 | 0,2044482 | chr8 | - |
| Pdlim3 | 3,13 | -0,0908 | 0,921388882 | chr8 | + |
| Pdp2 | 419,26 | 0,2730 | 0,583782269 | chr8 | + |
| Pdpr | 62,87 | 0,4157 | 0,499682405 | chr8 | + |
| Pet100 | 620,77 | -0,2933 | 0,390554711 | chr8 | + |
| Pex11g | 775,88 | -0,2819 | 0,362631895 | chr8 | - |
| Pgls | 3,80 | 0,2981 | 0,701217751 | chr8 | + |

| | | | | | |
|---|---|---|---|---|---|
| Pgpep1 | 1334,71 | 0,4474 | 0,118740672 | chr8 | - |
| Phkb | 482,15 | -0,2925 | 0,449196578 | chr8 | + |
| Phlpp2 | 81,56 | 0,4826 | 0,239250534 | chr8 | + |
| Piezo1 | 49,62 | -0,1088 | 0,886446111 | chr8 | - |
| Pik3r2 | 119,60 | -0,4187 | 0,403228071 | chr8 | - |
| Pkn1 | 164,44 | -0,1185 | 0,816025345 | chr8 | - |
| Pla2g15 | 204,03 | -0,0115 | 0,985628536 | chr8 | + |
| Plat | 14,19 | 0,8259 | 0,131674744 | chr8 | + |
| Plcg2 | 27,46 | 0,3132 | 0,603509332 | chr8 | + |
| Plekha2 | 92,08 | 0,2534 | 0,564849438 | chr8 | - |
| Pllp | 725,60 | 0,0618 | 0,881809902 | chr8 | - |
| Plvap | 275,69 | 0,2052 | 0,696074872 | chr8 | - |
| Pnpla6 | 183,92 | -0,5523 | 0,166993075 | chr8 | + |
| Polb | 382,24 | -0,2382 | 0,463892619 | chr8 | - |
| Polr2c | 332,76 | -0,1257 | 0,765801166 | chr8 | + |
| Plpp5 | 179,33 | 0,2418 | 0,658008155 | chr8 | + |
| Ppp1r3b | 2407,50 | -0,3274 | 0,581725594 | chr8 | + |
| Ppp2cb | 466,01 | -0,2698 | 0,382335568 | chr8 | + |
| Prdx2 | 2727,19 | -0,3013 | 0,313578168 | chr8 | + |
| Primpol | 102,50 | 0,3091 | 0,438110291 | chr8 | - |
| Prkaca | 557,20 | 0,1578 | 0,695575308 | chr8 | + |
| Prmt9 | 247,12 | 0,1523 | 0,775332905 | chr8 | + |
| Prmt7 | 196,78 | 0,1113 | 0,829703001 | chr8 | + |
| Plpbp | 968,14 | -0,1792 | 0,540815736 | chr8 | + |
| Proz | 4288,85 | 0,2008 | 0,500489603 | chr8 | + |
| Psd3 | 634,81 | 0,0656 | 0,88569455 | chr8 | - |
| Pskh1 | 424,89 | 0,0996 | 0,816025345 | chr8 | + |
| Psmb10 | 860,91 | 0,1013 | 0,793030656 | chr8 | - |
| Psmd7 | 1190,07 | -0,1993 | 0,518555479 | chr8 | - |
| Purg | 29,18 | -0,5898 | 0,217231457 | chr8 | + |
| Rab11fip1 | 131,13 | 0,4015 | 0,267895481 | chr8 | - |
| Rab20 | 107,66 | -0,2218 | 0,76049745 | chr8 | - |
| Rab3a | 33,78 | -0,3179 | 0,585240995 | chr8 | + |
| Rab4a | 331,57 | -0,3745 | 0,352106106 | chr8 | + |
| Rab8a | 1418,24 | -0,0084 | 0,986027421 | chr8 | + |
| Rad23a | 385,52 | -0,3088 | 0,278686919 | chr8 | - |
| Ranbp10 | 953,94 | -0,2221 | 0,537078872 | chr8 | - |
| Rasa3 | 118,88 | 0,0041 | 0,995663291 | chr8 | - |
| Rbl2 | 487,68 | 0,0164 | 0,978552759 | chr8 | + |
| Rbm34 | 176,38 | 0,0407 | 0,941757244 | chr8 | - |
| Rbmxl1 | 304,25 | 0,2276 | 0,503936414 | chr8 | - |
| Rbpms | 434,77 | -0,4490 | 0,200938023 | chr8 | - |
| Rfwd3 | 529,24 | 0,5747 | 0,050829111 | chr8 | - |
| Rfx1 | 62,47 | -0,5410 | 0,376078101 | chr8 | + |
| Rfxank | 147,85 | 0,4875 | 0,155520806 | chr8 | - |
| Rhou | 1315,89 | 0,1185 | 0,743199461 | chr8 | + |
| Rnaseh2a | 420,07 | -0,0216 | 0,972360063 | chr8 | - |
| Rnf122 | 10,79 | 0,6202 | 0,329981156 | chr8 | + |
| Rnf150 | 26,90 | 0,5233 | 0,388785569 | chr8 | + |
| Rnf166 | 97,67 | -0,1906 | 0,684393541 | chr8 | - |

| | | | | | |
|---|---|---|---|---|---|
| Rnf170 | 464,45 | -0,2662 | 0,392238586 | chr8 | + |
| Rpgrip1l | 44,56 | 0,5795 | 0,232658696 | chr8 | - |
| Rpl13 | 3397,51 | -0,7204 | 0,135452853 | chr8 | + |
| Rpl18a | 3587,23 | -0,5257 | 0,082179991 | chr8 | - |
| Rrad | 2,61 | -0,2972 | 0,672218442 | chr8 | - |
| Rspry1 | 339,75 | 0,1852 | 0,663934341 | chr8 | + |
| Rwdd4a | 331,81 | -0,3392 | 0,238655284 | chr8 | + |
| Sall1 | 237,18 | 0,1772 | 0,7009274 | chr8 | - |
| Samd1 | 70,45 | -0,6175 | 0,185276108 | chr8 | + |
| Sap30 | 53,55 | -0,2020 | 0,705106049 | chr8 | - |
| Scoc | 320,77 | 0,2530 | 0,615783353 | chr8 | - |
| Sdr42e1 | 1628,67 | -0,0168 | 0,98429278 | chr8 | - |
| Setd6 | 217,37 | -0,4625 | 0,103014216 | chr8 | + |
| Sf3b3 | 476,77 | -0,0081 | 0,990189789 | chr8 | - |
| Sfrp1 | 13,19 | 0,1992 | 0,815176408 | chr8 | + |
| Sh2d4a | 76,54 | -0,0373 | 0,961556353 | chr8 | + |
| Sh3rf1 | 42,43 | 0,0688 | 0,92369436 | chr8 | + |
| Shcbp1 | 11,44 | -0,0250 | 0,981991128 | chr8 | - |
| Siah1a | 86,89 | 0,2229 | 0,656897388 | chr8 | - |
| Sin3b | 624,48 | -0,3473 | 0,419539452 | chr8 | + |
| Sipa1l2 | 65,45 | 0,1355 | 0,838041793 | chr8 | - |
| Slc10a7 | 158,40 | 0,3636 | 0,40906746 | chr8 | + |
| Slc12a4 | 198,01 | -0,0465 | 0,942982044 | chr8 | - |
| Slc18a1 | 37,36 | -0,0688 | 0,938364545 | chr8 | - |
| Slc20a2 | 790,19 | -0,0602 | 0,869825447 | chr8 | + |
| Slc25a15 | 6953,00 | -0,0911 | 0,81937394 | chr8 | - |
| Slc25a42 | 790,34 | -0,3658 | 0,47972727 | chr8 | - |
| Slc27a1 | 50,40 | 0,2294 | 0,720291746 | chr8 | + |
| Slc35e1 | 664,74 | -0,2942 | 0,377944963 | chr8 | - |
| Slc38a7 | 236,17 | -0,1141 | 0,802632263 | chr8 | - |
| Slc7a2 | 7810,09 | -0,5423 | 0,115749538 | chr8 | + |
| Slc7a5 | 24,60 | -0,5130 | 0,339424882 | chr8 | - |
| Slc7a6 | 10,54 | 0,5253 | 0,41633338 | chr8 | + |
| Slc9a5 | 5,18 | -0,2377 | 0,761836557 | chr8 | + |
| Smad1 | 190,34 | -0,1066 | 0,826347896 | chr8 | - |
| Smarca5 | 660,34 | -0,0708 | 0,877062109 | chr8 | - |
| Smim19 | 289,38 | 0,0002 | 0,999426132 | chr8 | - |
| Smim7 | 1505,20 | -0,1882 | 0,666980604 | chr8 | - |
| Smpd3 | 66,02 | 0,6844 | 0,226302394 | chr8 | - |
| Snai3 | 3,98 | 0,4705 | 0,501000077 | chr8 | - |
| Snapc2 | 100,89 | -0,4873 | 0,25884542 | chr8 | + |
| Snora68 | 4,98 | -0,4006 | 0,588589342 | chr8 | - |
| Sntb2 | 49,11 | 0,5990 | 0,170076715 | chr8 | + |
| Snx20 | 8,24 | 0,3175 | 0,680997554 | chr8 | - |
| Snx25 | 314,06 | -0,1556 | 0,721569914 | chr8 | - |
| Sorbs2 | 785,34 | 0,0917 | 0,815421548 | chr8 | + |
| Spata2l | 230,51 | 0,1922 | 0,683804325 | chr8 | - |
| Spcs3 | 2252,34 | 0,1296 | 0,748732136 | chr8 | - |
| Spg7 | 409,39 | -0,4080 | 0,169162729 | chr8 | + |
| Spire2 | 2,60 | 0,1147 | 0,895663283 | chr8 | + |

| | | | | | |
|---|---|---|---|---|---|
| Sprtn | 30,19 | 0,3570 | 0,531354472 | chr8 | + |
| Ssbp4 | 12,20 | -0,1176 | 0,89773876 | chr8 | - |
| St3gal2 | 26,86 | 0,4871 | 0,381639797 | chr8 | + |
| Star | 18,29 | -0,6439 | 0,220418266 | chr8 | + |
| Stox2 | 125,90 | 0,1354 | 0,788459223 | chr8 | - |
| Stxbp2 | 628,75 | -0,0121 | 0,979186519 | chr8 | + |
| Sugp1 | 167,39 | -0,1713 | 0,678735998 | chr8 | + |
| Sugp2 | 52,95 | -0,3473 | 0,509826948 | chr8 | + |
| Sult5a1 | 204,76 | -0,4418 | 0,390832005 | chr8 | - |
| Syce2 | 26,63 | -0,2238 | 0,767446843 | chr8 | + |
| Tacc1 | 693,81 | -0,0173 | 0,978378397 | chr8 | - |
| Taf1c | 20,36 | -0,0916 | 0,916852736 | chr8 | - |
| Taf5l | 200,80 | -0,0911 | 0,848529459 | chr8 | - |
| Tango6 | 50,45 | -0,0286 | 0,969159464 | chr8 | + |
| Tat | 30695,35 | -0,4824 | 0,314000124 | chr8 | + |
| Tbc1d9 | 20,36 | 0,3568 | 0,590680988 | chr8 | + |
| Tcf25 | 1547,07 | -0,3997 | 0,149536902 | chr8 | + |
| Tecr | 2374,33 | -0,4609 | 0,172265665 | chr8 | - |
| Tenm3 | 116,57 | 0,0271 | 0,960225359 | chr8 | - |
| Terf2 | 103,54 | 0,0121 | 0,98429278 | chr8 | - |
| Terf2ip | 205,32 | -0,2852 | 0,391497936 | chr8 | + |
| Tfdp1 | 286,06 | -0,1556 | 0,723157997 | chr8 | + |
| Thap1 | 91,58 | -0,1769 | 0,691615955 | chr8 | + |
| Thap11 | 180,09 | -0,3767 | 0,266630306 | chr8 | + |
| Thsd1 | 37,49 | -0,0750 | 0,916757648 | chr8 | + |
| Timm44 | 848,74 | -0,3137 | 0,211650242 | chr8 | - |
| Tk2 | 237,14 | -0,4742 | 0,112204955 | chr8 | - |
| Tldc1 | 119,62 | 0,4165 | 0,345715253 | chr8 | - |
| Tm2d2 | 749,99 | -0,4860 | 0,050838368 | chr8 | + |
| Tm6sf2 | 368,24 | 0,5450 | 0,244584613 | chr8 | + |
| Tma16 | 104,95 | 0,0304 | 0,957956679 | chr8 | - |
| Tmco3 | 80,82 | -0,0729 | 0,905726602 | chr8 | + |
| Tmed6 | 10,90 | 0,6101 | 0,352830402 | chr8 | - |
| Tmem161a | 225,40 | -0,0852 | 0,862964916 | chr8 | + |
| Tmem170 | 75,99 | 0,3010 | 0,502106225 | chr8 | - |
| Tmem192 | 133,83 | -0,2722 | 0,485023704 | chr8 | + |
| Tmem208 | 1182,08 | -0,4376 | 0,065659595 | chr8 | + |
| Tmem221 | 6,22 | 0,3884 | 0,604679897 | chr8 | - |
| Tmem231 | 4,52 | 0,2779 | 0,727360927 | chr8 | - |
| Tmem38a | 31,98 | -0,5379 | 0,29105938 | chr8 | + |
| Saraf | 2029,83 | -0,2181 | 0,49077279 | chr8 | + |
| Tnfsf13b | 7,58 | 0,1592 | 0,859286024 | chr8 | + |
| Tnks | 113,46 | 0,4933 | 0,220525617 | chr8 | - |
| Tnpo2 | 662,90 | 0,1026 | 0,775454442 | chr8 | + |
| Tom1 | 295,34 | -0,3552 | 0,283447934 | chr8 | + |
| Tomm20 | 1058,04 | -0,3001 | 0,37928598 | chr8 | - |
| Tppp3 | 5,76 | 0,1198 | 0,895663283 | chr8 | - |
| Tradd | 129,24 | 0,0906 | 0,862578493 | chr8 | - |
| Trappc11 | 506,91 | 0,0545 | 0,918077233 | chr8 | - |
| Trappc2l | 404,49 | 0,0113 | 0,981991128 | chr8 | + |

| | | | | | |
|---|---|---|---|---|---|
| Trappc5 | 268,71 | -0,2555 | 0,648165583 | chr8 | + |
| Trmt1 | 260,33 | -0,5164 | 0,192272022 | chr8 | + |
| Tsnax | 513,77 | 0,0177 | 0,970419105 | chr8 | + |
| Ttc13 | 313,00 | -0,1387 | 0,725810792 | chr8 | - |
| Ttc29 | 391,36 | 0,1298 | 0,726240854 | chr8 | + |
| Tti2 | 163,41 | 0,0153 | 0,978280656 | chr8 | + |
| Tubgcp3 | 153,09 | 0,1603 | 0,706001911 | chr8 | - |
| Tusc3 | 127,01 | 0,4978 | 0,166136649 | chr8 | + |
| Txnl4b | 222,29 | -0,0591 | 0,903638021 | chr8 | + |
| Uba52 | 12187,53 | -0,3446 | 0,253392997 | chr8 | - |
| Ubxn8 | 380,01 | -0,1139 | 0,760699597 | chr8 | - |
| Ufsp2 | 1101,33 | 0,0028 | 0,995836778 | chr8 | + |
| Unc13a | 4,31 | 0,6438 | 0,31487946 | chr8 | - |
| Upf1 | 320,28 | -0,6060 | 0,094600644 | chr8 | - |
| Upf3a | 146,43 | -0,5091 | 0,163729985 | chr8 | + |
| Urb2 | 122,61 | -0,1865 | 0,657858101 | chr8 | + |
| Usb1 | 156,46 | -0,0412 | 0,940188887 | chr8 | + |
| Use1 | 922,83 | -0,5136 | 0,05541471 | chr8 | + |
| Ushbp1 | 119,59 | -0,1451 | 0,820554805 | chr8 | - |
| Usp10 | 604,81 | -0,5773 | 0,10394222 | chr8 | + |
| Usp38 | 440,13 | -0,2240 | 0,633994747 | chr8 | - |
| Vac14 | 245,42 | 0,0653 | 0,922460349 | chr8 | + |
| Vdac3 | 2508,56 | -0,0146 | 0,974449819 | chr8 | - |
| Vegfc | 59,36 | -0,4228 | 0,345491882 | chr8 | + |
| Vps35 | 2332,01 | 0,1801 | 0,615783353 | chr8 | - |
| Vps36 | 344,78 | 0,3360 | 0,279486542 | chr8 | + |
| Vps37a | 742,38 | -0,3628 | 0,297002219 | chr8 | + |
| Vps4a | 307,49 | -0,2930 | 0,332496065 | chr8 | + |
| Vps9d1 | 148,92 | -0,2188 | 0,629719515 | chr8 | - |
| Wdr17 | 1,95 | 0,5244 | NA | chr8 | - |
| Wdr59 | 182,06 | -0,0128 | 0,982521526 | chr8 | - |
| Wdr83 | 193,34 | -0,1675 | 0,704653924 | chr8 | - |
| Wdr83os | 783,84 | 0,0643 | 0,905147397 | chr8 | + |
| Wfdc1 | 65,09 | 0,4907 | 0,394162656 | chr8 | + |
| Nsd3 | 664,33 | -0,3027 | 0,413615196 | chr8 | + |
| Wrn | 189,40 | 0,3859 | 0,357345082 | chr8 | - |
| Wwc2 | 105,13 | -0,0576 | 0,917042614 | chr8 | - |
| Wwox | 71,24 | 0,0768 | 0,9024844 | chr8 | + |
| Wwp2 | 263,43 | 0,2792 | 0,549296977 | chr8 | + |
| Xab2 | 188,27 | -0,1138 | 0,793608383 | chr8 | - |
| Zc3h18 | 346,04 | -0,1840 | 0,710761695 | chr8 | + |
| Zcchc14 | 298,17 | -0,2206 | 0,480021428 | chr8 | - |
| Zdhhc1 | 18,56 | -0,4016 | 0,561310046 | chr8 | - |
| Zdhhc2 | 37,41 | 0,0769 | 0,918104715 | chr8 | + |
| Zdhhc7 | 159,61 | -0,4151 | 0,25148816 | chr8 | - |
| Zfhx3 | 123,93 | -0,2761 | 0,536603375 | chr8 | + |
| Zfp1 | 86,65 | 0,0334 | 0,955496112 | chr8 | + |
| Zfp276 | 220,98 | 0,0168 | 0,978552759 | chr8 | + |
| Zfp319 | 49,31 | 0,1548 | 0,835335848 | chr8 | - |
| Zfp330 | 318,97 | 0,2427 | 0,521815657 | chr8 | - |

| | | | | | |
|---|---|---|---|---|---|
| Zfp358 | 82,64 | -0,3729 | 0,440094295 | chr8 | + |
| Zfp612 | 141,78 | 0,3444 | 0,491855001 | chr8 | + |
| Zfp617 | 402,04 | 0,4340 | 0,115749538 | chr8 | + |
| Zfp821 | 39,17 | -0,4697 | 0,356860747 | chr8 | + |
| Zfp827 | 4,64 | 0,3620 | 0,627992485 | chr8 | + |
| Zfp866 | 175,41 | 0,1060 | 0,793076009 | chr8 | - |
| Zfp868 | 212,38 | -0,1067 | 0,851377761 | chr8 | - |
| Zfp869 | 393,21 | 0,1660 | 0,65809882 | chr8 | - |
| Zfp90 | 26,31 | 0,1989 | 0,776345685 | chr8 | + |
| Zfp930 | 160,95 | 0,1928 | 0,688733355 | chr8 | + |
| Zfp958 | 148,04 | 0,3170 | 0,357345082 | chr8 | + |
| Zfp961 | 176,79 | 0,2424 | 0,629075916 | chr8 | + |
| Zfp963 | 91,83 | -0,2122 | 0,658008155 | chr8 | - |
| Zfp964 | 10,36 | -0,1644 | 0,849477252 | chr8 | + |
| Zfpm1 | 187,38 | -0,2683 | 0,740715013 | chr8 | + |
| Znrf1 | 323,54 | -0,6369 | 0,073845576 | chr8 | + |
| Zswim4 | 79,00 | -0,3269 | 0,532105996 | chr8 | - |
| 1110032A03Rik | 851,47 | -0,2055 | 0,581164752 | chr9 | - |
| 1110059G10Rik | 305,31 | 0,5103 | 0,181324183 | chr9 | - |
| 1300017J02Rik | 12573,61 | -0,0988 | 0,789064438 | chr9 | - |
| 1700017B05Rik | 100,52 | -0,0232 | 0,969301796 | chr9 | - |
| 1700063D05Rik | 6,76 | -0,3103 | 0,692557703 | chr9 | + |
| 1700102P08Rik | 6,79 | -1,0268 | 0,073903027 | chr9 | + |
| Timm29 | 627,61 | -0,3120 | 0,358990473 | chr9 | + |
| 1810046K07Rik | 27,26 | 1,0458 | 0,062791791 | chr9 | - |
| 2300009A05Rik | 108,08 | -0,5226 | 0,282931396 | chr9 | - |
| 2310009A05Rik | 291,88 | -0,2354 | 0,665028267 | chr9 | + |
| 2310030G06Rik | 380,95 | 0,1604 | 0,678853818 | chr9 | - |
| Rec114 | 35,14 | 1,0101 | 0,071771252 | chr9 | - |
| 2610203C20Rik | 8,95 | -1,0689 | 0,05009592 | chr9 | + |
| Pclaf | 21,37 | 0,1428 | 0,84862368 | chr9 | + |
| 3000002C10Rik | 66,33 | -0,0113 | 0,986889999 | chr9 | - |
| 4930429F24Rik | 2,81 | 0,3016 | 0,663873104 | chr9 | + |
| 4930433N12Rik | 2,49 | 0,2848 | NA | chr9 | - |
| 4930509E16Rik | 4,57 | 0,9605 | 0,086187166 | chr9 | - |
| 4930526I15Rik | 12,91 | -0,4053 | 0,582699969 | chr9 | + |
| 4930550C14Rik | 4,27 | 0,4725 | 0,497298938 | chr9 | + |
| 4930579K19Rik | 3,57 | 0,3737 | 0,612061669 | chr9 | - |
| 4930581F22Rik | 236,51 | -0,1413 | 0,792197952 | chr9 | + |
| 4931406C07Rik | 7418,98 | 0,1936 | 0,617730194 | chr9 | - |
| Cep295 | 83,03 | 0,2985 | 0,572115465 | chr9 | - |
| 5830454E08Rik | 11,05 | 0,0397 | 0,969159464 | chr9 | + |
| 6430571L13Rik | 9,96 | 0,2968 | 0,7023898 | chr9 | + |
| 9230110C19Rik | 8,99 | 0,1576 | 0,860606321 | chr9 | - |
| 9330159M07Rik | 27,49 | 0,6380 | 0,179897103 | chr9 | - |
| 9530077C05Rik | 3,45 | -0,4277 | 0,546240378 | chr9 | + |
| A230050P20Rik | 396,62 | -0,1566 | 0,795157766 | chr9 | + |
| A330041J22Rik | 4,03 | 0,2734 | 0,73216561 | chr9 | + |
| A4gnt | 5,12 | 0,6111 | NA | chr9 | + |
| Aagab | 445,09 | -0,2150 | 0,524801822 | chr9 | + |

| | | | | | |
|---|---|---|---|---|---|
| Aasdhppt | 718,76 | -0,1743 | 0,695978671 | chr9 | - |
| AB124611 | 79,07 | 0,3924 | 0,376349515 | chr9 | + |
| Abhd14a | 189,70 | 0,2637 | 0,514316163 | chr9 | - |
| Abhd14b | 2286,11 | -0,3701 | 0,184518314 | chr9 | + |
| Abhd5 | 680,35 | -0,4858 | 0,066790906 | chr9 | + |
| Acaa1a | 1251,98 | -0,6186 | 0,092096287 | chr9 | + |
| Acad11 | 8014,48 | -0,1139 | 0,809790857 | chr9 | + |
| Acad8 | 1468,79 | -0,0981 | 0,783418712 | chr9 | - |
| Acat1 | 8797,71 | 0,0619 | 0,895594598 | chr9 | - |
| Ackr2 | 27,30 | 0,5972 | 0,217231457 | chr9 | + |
| Ackr4 | 27,35 | 0,2330 | 0,732982182 | chr9 | - |
| Acpp | 68,84 | 0,1168 | 0,878605672 | chr9 | - |
| Acvr2b | 12,49 | 0,0435 | 0,9651628 | chr9 | + |
| Acy1 | 567,68 | -0,0932 | 0,812943523 | chr9 | - |
| Adam10 | 646,84 | 0,1053 | 0,782080553 | chr9 | + |
| Adamts15 | 12,92 | 0,2052 | 0,79991265 | chr9 | - |
| Adamts7 | 67,16 | 0,4076 | 0,404238051 | chr9 | + |
| Adpgk | 170,22 | -0,4014 | 0,173352491 | chr9 | + |
| Tmem266 | 9,20 | 0,7279 | 0,242907451 | chr9 | + |
| Cep126 | 2,79 | -0,1013 | 0,907779947 | chr9 | - |
| Alas1 | 4889,40 | -0,4280 | 0,397488617 | chr9 | - |
| Aldh1a2 | 2,26 | -0,1650 | NA | chr9 | + |
| Alg9 | 276,10 | 0,0185 | 0,970419105 | chr9 | + |
| Alkbh8 | 53,38 | 0,5170 | 0,339424882 | chr9 | + |
| Als2cl | 298,60 | -0,2210 | 0,658008155 | chr9 | + |
| Jaml | 51,92 | -0,1765 | 0,834553481 | chr9 | + |
| Amigo3 | 5,32 | 0,1699 | 0,847964947 | chr9 | + |
| Amotl1 | 140,44 | 0,2175 | 0,577838565 | chr9 | - |
| Amotl2 | 269,77 | 0,1525 | 0,799805793 | chr9 | + |
| Amt | 424,59 | 0,3195 | 0,41633338 | chr9 | + |
| Anapc13 | 1060,96 | -0,3216 | 0,215519599 | chr9 | + |
| Angptl6 | 153,35 | -0,1553 | 0,804418611 | chr9 | - |
| Ankrd49 | 136,17 | 0,5897 | 0,098663118 | chr9 | - |
| Anln | 19,75 | 0,0893 | 0,918946301 | chr9 | - |
| Ano10 | 217,79 | 0,0070 | 0,99143613 | chr9 | - |
| Anp32a | 2915,43 | 0,4763 | 0,211603645 | chr9 | + |
| Ap1m2 | 4,72 | 0,9635 | 0,094281875 | chr9 | - |
| Apeh | 1027,52 | -0,1093 | 0,753532095 | chr9 | - |
| Aph1c | 23,20 | -0,1589 | 0,845257983 | chr9 | - |
| Aplp2 | 3746,39 | -0,1799 | 0,49917227 | chr9 | - |
| Apoa1 | 169709,28 | -0,2411 | 0,505223498 | chr9 | + |
| Apoa4 | 4969,62 | 0,2149 | 0,662842062 | chr9 | + |
| Apoa5 | 25844,90 | -0,3172 | 0,375836696 | chr9 | + |
| Apoc3 | 60866,54 | -0,0980 | 0,765061508 | chr9 | - |
| Aqp9 | 6818,27 | -0,2665 | 0,483638076 | chr9 | - |
| Arcn1 | 2718,15 | -0,0149 | 0,978160703 | chr9 | - |
| Arhgap32 | 126,88 | -0,0280 | 0,971664369 | chr9 | + |
| Arhgap42 | 913,04 | -0,1675 | 0,546813684 | chr9 | - |
| Arhgef12 | 1451,20 | -0,0899 | 0,793608383 | chr9 | - |
| Arid3b | 27,18 | 0,2010 | 0,76530222 | chr9 | - |

| | | | | | |
|---|---|---|---|---|---|
| Arih1 | 1015,32 | -0,2370 | 0,41272264 | chr9 | - |
| Armc8 | 281,91 | 0,1882 | 0,68361653 | chr9 | - |
| Arpp19 | 1326,48 | -0,1671 | 0,588219472 | chr9 | + |
| Aste1 | 42,88 | -0,3624 | 0,501951423 | chr9 | + |
| Atm | 239,76 | 0,1732 | 0,701666523 | chr9 | - |
| Atp1b3 | 605,42 | -0,4008 | 0,257021621 | chr9 | - |
| Atp2c1 | 743,58 | -0,1331 | 0,677299626 | chr9 | - |
| Atp5l | 9295,11 | -0,4352 | 0,129469509 | chr9 | - |
| Atr | 334,57 | 0,3890 | 0,302314196 | chr9 | + |
| Atrip | 65,88 | -0,0373 | 0,952576171 | chr9 | - |
| AU019823 | 189,20 | 0,1444 | 0,736631536 | chr9 | - |
| AU023762 | 3,66 | 0,1282 | 0,886098212 | chr9 | - |
| Azi2 | 847,72 | -0,0057 | 0,99137621 | chr9 | + |
| Bace1 | 259,89 | 0,1841 | 0,720000255 | chr9 | + |
| Bbs9 | 85,33 | 0,1336 | 0,841053685 | chr9 | + |
| Plppr2 | 30,35 | -0,2423 | 0,740024898 | chr9 | + |
| BC049352 | 3,13 | -0,4443 | 0,498484848 | chr9 | + |
| Bckdhb | 1559,76 | -0,0136 | 0,979212524 | chr9 | + |
| Bcl2a1a | 3,85 | 0,2173 | 0,783418712 | chr9 | + |
| Bcl2a1b | 21,04 | 0,3032 | 0,661748407 | chr9 | + |
| Bcl9l | 155,97 | 0,0250 | 0,974449819 | chr9 | + |
| Bco2 | 449,70 | 0,2039 | 0,687523396 | chr9 | - |
| Birc2 | 466,80 | 0,3729 | 0,245076229 | chr9 | - |
| Birc3 | 272,06 | 0,2316 | 0,613297625 | chr9 | - |
| Bmper | 7,84 | 0,5448 | 0,41552947 | chr9 | + |
| Bnip2 | 858,51 | -0,3008 | 0,229410024 | chr9 | + |
| C920006O11Rik | 45,02 | -0,1834 | 0,777915388 | chr9 | + |
| Cadm1 | 320,31 | -0,5561 | 0,067696785 | chr9 | + |
| Carm1 | 151,74 | -0,2080 | 0,657858101 | chr9 | + |
| Casp1 | 95,15 | 0,3213 | 0,415700483 | chr9 | + |
| Casp4 | 49,82 | 0,1237 | 0,848389544 | chr9 | + |
| Cbl | 17,87 | 0,6567 | 0,252815255 | chr9 | - |
| Ccdc12 | 345,22 | -0,2727 | 0,522942093 | chr9 | + |
| Ccdc15 | 6,62 | 0,6154 | 0,349661709 | chr9 | - |
| Ccdc151 | 19,99 | -0,5491 | 0,396182687 | chr9 | - |
| Ccdc159 | 54,26 | -0,0678 | 0,914198498 | chr9 | + |
| Ccdc36 | 2,35 | 0,3318 | NA | chr9 | - |
| Ccdc51 | 164,74 | 0,3085 | 0,396447904 | chr9 | + |
| Deup1 | 8,76 | 0,6358 | 0,330471707 | chr9 | - |
| Ccdc71 | 238,78 | 0,0660 | 0,896037411 | chr9 | + |
| Ccdc82 | 683,27 | 0,1686 | 0,627637264 | chr9 | + |
| Ccnb2 | 19,48 | -0,2822 | 0,668257037 | chr9 | - |
| Ccpg1 | 1237,59 | -0,0799 | 0,825839229 | chr9 | + |
| Ccpg1os | 207,11 | -0,0562 | 0,927339856 | chr9 | - |
| Ccr1 | 9,92 | 0,7570 | 0,22361789 | chr9 | - |
| Ccr3 | 58,54 | 1,0412 | 0,065247775 | chr9 | + |
| Ccr9 | 5,36 | 0,7478 | 0,231140315 | chr9 | + |
| Ccrl2 | 32,19 | 0,5127 | 0,368196526 | chr9 | - |
| Cd3d | 15,17 | 0,3412 | 0,621241714 | chr9 | + |
| Cd3e | 7,07 | 0,0120 | 0,990846949 | chr9 | - |

| | | | | | |
|---|---|---|---|---|---|
| Cd3g | 21,33 | 0,3167 | 0,636447281 | chr9 | - |
| Cdc25a | 62,36 | -0,2804 | 0,589961159 | chr9 | + |
| Cdc37 | 1015,59 | -0,4657 | 0,258561012 | chr9 | - |
| Cdcp1 | 7,14 | 0,5223 | 0,431872926 | chr9 | - |
| Cdkn2d | 41,57 | -0,0834 | 0,901445192 | chr9 | - |
| Cdon | 22,79 | 0,0366 | 0,966608842 | chr9 | + |
| Cdv3 | 3188,80 | -0,4928 | 0,067696785 | chr9 | - |
| Cep162 | 37,22 | 0,1464 | 0,834675238 | chr9 | - |
| Cep164 | 108,32 | -0,3557 | 0,437914468 | chr9 | - |
| Cep57 | 206,86 | 0,1652 | 0,667633531 | chr9 | - |
| Cep63 | 174,11 | 0,0142 | 0,981991128 | chr9 | - |
| Cep70 | 54,50 | 0,2490 | 0,613341495 | chr9 | + |
| Cgnl1 | 1299,74 | -0,2183 | 0,578442407 | chr9 | - |
| Chek1 | 16,00 | 0,3663 | 0,584602378 | chr9 | - |
| Chordc1 | 964,37 | 0,4136 | 0,302314196 | chr9 | + |
| Chst2 | 29,76 | -0,2047 | 0,757883527 | chr9 | - |
| Cib2 | 57,83 | -0,2533 | 0,622736166 | chr9 | - |
| Cilp | 7,71 | -0,0734 | 0,942585501 | chr9 | + |
| Cish | 168,70 | 0,0577 | 0,952511874 | chr9 | + |
| Clasp2 | 172,90 | 0,4943 | 0,203743201 | chr9 | + |
| Clec3b | 11,48 | -0,1673 | 0,84933342 | chr9 | + |
| Clk3 | 498,80 | -0,1879 | 0,689392185 | chr9 | - |
| Cln6 | 62,56 | -0,2775 | 0,604682319 | chr9 | + |
| Clpx | 1460,73 | 0,1953 | 0,44242629 | chr9 | + |
| Clstn2 | 3,21 | -0,4201 | 0,546240378 | chr9 | - |
| Cmc1 | 491,97 | 0,3257 | 0,347246285 | chr9 | - |
| Cmtm6 | 2050,83 | -0,0297 | 0,953751362 | chr9 | + |
| Cmtm7 | 41,30 | 0,1434 | 0,814990243 | chr9 | - |
| Cmtm8 | 959,79 | -0,3601 | 0,437620234 | chr9 | - |
| Cnn1 | 2,55 | 0,2448 | NA | chr9 | + |
| Cnot10 | 325,23 | -0,1060 | 0,760736495 | chr9 | - |
| Col12a1 | 22,50 | -0,6655 | 0,206831453 | chr9 | - |
| Col5a3 | 314,78 | -0,3050 | 0,62632688 | chr9 | - |
| Col6a5 | 10,80 | 0,4430 | 0,536603375 | chr9 | - |
| Col6a6 | 4,82 | 0,4579 | 0,523689387 | chr9 | - |
| Col7a1 | 3,01 | -0,3701 | 0,608287564 | chr9 | + |
| Commd4 | 574,63 | -0,3869 | 0,086066172 | chr9 | - |
| Copb2 | 3034,43 | -0,0438 | 0,923778914 | chr9 | + |
| Coro2b | 21,70 | 0,2043 | 0,783216386 | chr9 | - |
| Crtap | 97,35 | -0,6294 | 0,076373799 | chr9 | - |
| Csk | 571,28 | -0,3680 | 0,341021075 | chr9 | - |
| Csnk1g1 | 75,94 | 0,0187 | 0,978280656 | chr9 | + |
| Csrnp1 | 160,55 | -0,0144 | 0,981931319 | chr9 | - |
| Ctdspl | 68,98 | -0,1335 | 0,802706588 | chr9 | + |
| Ctnnb1 | 3617,45 | -0,1217 | 0,744078898 | chr9 | + |
| Ctsh | 6468,04 | -0,2042 | 0,614209897 | chr9 | + |
| Cul5 | 423,41 | 0,1266 | 0,794944296 | chr9 | - |
| Cwc15 | 1051,03 | -0,2740 | 0,377944963 | chr9 | + |
| Cwf19l2 | 314,64 | 0,5537 | 0,2163009 | chr9 | + |
| Cx3cr1 | 22,70 | 0,9275 | 0,066915827 | chr9 | - |

| | | | | | |
|---|---|---|---|---|---|
| Cxcr5 | 3,11 | 0,0122 | 0,990472463 | chr9 | - |
| Cxcr6 | 28,16 | 0,2569 | 0,714302449 | chr9 | + |
| Cyb561d2 | 139,81 | -0,2844 | 0,524328995 | chr9 | - |
| Cyb5r4 | 642,68 | -0,1034 | 0,803874753 | chr9 | + |
| Cyp1a1 | 19,23 | -0,5501 | 0,351130787 | chr9 | + |
| Cyp8b1 | 2674,01 | 0,0154 | 0,985212135 | chr9 | - |
| D630033O11Rik | 9,79 | -0,4309 | 0,546917025 | chr9 | + |
| Dag1 | 846,13 | 0,1703 | 0,593932312 | chr9 | - |
| Dalrd3 | 320,43 | -0,3494 | 0,381800527 | chr9 | + |
| Dapk2 | 99,51 | 0,2288 | 0,677867285 | chr9 | + |
| Dbr1 | 152,47 | 0,0188 | 0,974621894 | chr9 | + |
| Dclk3 | 169,34 | 0,0243 | 0,969512928 | chr9 | + |
| Dcps | 356,28 | -0,1464 | 0,696615722 | chr9 | - |
| Dcun1d5 | 446,84 | -0,3708 | 0,281377199 | chr9 | + |
| Ddx10 | 270,76 | 0,2226 | 0,543578711 | chr9 | - |
| Ddx6 | 2583,42 | 0,1271 | 0,736631536 | chr9 | + |
| Dennd4a | 519,68 | 0,1592 | 0,688905597 | chr9 | + |
| Dhx30 | 268,14 | -0,0299 | 0,952188756 | chr9 | - |
| Dis3l | 257,25 | -0,2809 | 0,428001468 | chr9 | - |
| Dixdc1 | 315,73 | 0,5772 | 0,190066764 | chr9 | - |
| Dlat | 658,29 | -0,2102 | 0,598404146 | chr9 | - |
| Dlec1 | 3,45 | 0,2117 | 0,798128487 | chr9 | + |
| Dmxl2 | 326,39 | -0,2127 | 0,577635343 | chr9 | - |
| Dnaja4 | 110,46 | 0,0874 | 0,875717638 | chr9 | + |
| Dnajc13 | 490,28 | 0,0061 | 0,991295673 | chr9 | - |
| Dnm2 | 414,16 | -0,0180 | 0,977170685 | chr9 | + |
| Dnmt1 | 113,42 | 0,1901 | 0,665658933 | chr9 | - |
| Dock6 | 137,12 | -0,0812 | 0,89879469 | chr9 | - |
| Dopey1 | 178,43 | 0,4574 | 0,349326231 | chr9 | + |
| Dpagt1 | 477,31 | -0,3502 | 0,20028078 | chr9 | + |
| Dpp8 | 666,52 | 0,1105 | 0,797919644 | chr9 | + |
| Dpy19l1 | 1095,77 | -0,0428 | 0,931322892 | chr9 | - |
| Dusp7 | 48,63 | -0,1144 | 0,864500236 | chr9 | + |
| Dync1li1 | 176,50 | -0,1422 | 0,762940356 | chr9 | + |
| Dync2h1 | 65,30 | 0,0856 | 0,899188684 | chr9 | - |
| Dyx1c1 | 6,25 | 0,7394 | 0,238321283 | chr9 | + |
| Dzip1l | 7,46 | 0,6891 | 0,282189277 | chr9 | + |
| E330023G01Rik | 11,84 | 0,3338 | 0,650328344 | chr9 | - |
| Ecsit | 505,84 | -0,3774 | 0,248800492 | chr9 | - |
| Edc3 | 258,55 | 0,0672 | 0,880529758 | chr9 | + |
| Eef1a1 | 111792,64 | -0,2580 | 0,361447043 | chr9 | - |
| Eepd1 | 259,15 | -0,5077 | 0,163659381 | chr9 | + |
| Ei24 | 4284,57 | 0,1183 | 0,743138956 | chr9 | - |
| Eif1b | 263,18 | -0,1721 | 0,656913078 | chr9 | + |
| Eif3g | 855,40 | -0,4542 | 0,115749538 | chr9 | - |
| Elavl3 | 4,14 | -0,8286 | 0,154269521 | chr9 | - |
| Elof1 | 844,07 | -0,1305 | 0,740538349 | chr9 | - |
| Elovl5 | 6887,08 | 0,2922 | 0,395114656 | chr9 | + |
| Elp6 | 78,19 | -0,2254 | 0,632136487 | chr9 | + |
| Epm2aip1 | 324,06 | 0,2349 | 0,618723125 | chr9 | + |

| | | | | | |
|---|---|---|---|---|---|
| Epor | 12,63 | 0,1478 | 0,862965009 | chr9 | - |
| Esam | 199,67 | -0,0808 | 0,866934682 | chr9 | + |
| Esyt3 | 2,04 | 0,5717 | NA | chr9 | - |
| Etfa | 7779,42 | -0,0820 | 0,83081648 | chr9 | - |
| Ets1 | 344,72 | 0,4230 | 0,187852299 | chr9 | + |
| Exog | 30,13 | -0,3035 | 0,649851928 | chr9 | + |
| Exosc7 | 718,62 | 0,2160 | 0,631200781 | chr9 | + |
| Exph5 | 280,76 | 0,1047 | 0,81194417 | chr9 | + |
| Faim | 84,17 | 0,4151 | 0,380860886 | chr9 | + |
| Fam118b | 120,45 | -0,2085 | 0,654583626 | chr9 | - |
| Fam212a | 5,43 | -0,0022 | 0,997745329 | chr9 | - |
| Fam214a | 765,54 | -0,4210 | 0,250533705 | chr9 | + |
| Fam219b | 149,51 | 0,2650 | 0,623104286 | chr9 | + |
| Fam46a | 344,25 | 0,1742 | 0,617869186 | chr9 | - |
| Mindy2 | 205,19 | 0,3564 | 0,399439692 | chr9 | - |
| Fam76b | 142,20 | -0,1901 | 0,646093772 | chr9 | + |
| Fam96a | 1124,53 | 0,1183 | 0,740024898 | chr9 | + |
| Fbxl12 | 56,76 | -0,1137 | 0,847419446 | chr9 | - |
| Fbxl12os | 42,82 | 0,5461 | 0,211542715 | chr9 | + |
| Fbxl22 | 9,22 | -1,0131 | 0,077273342 | chr9 | - |
| Fbxo22 | 903,93 | 0,0022 | 0,995735598 | chr9 | + |
| Fbxo9 | 835,29 | -0,4153 | 0,083831145 | chr9 | - |
| Fbxw15 | 5,72 | 0,9493 | 0,096645255 | chr9 | - |
| Fdx1 | 1112,05 | -0,4413 | 0,065752252 | chr9 | - |
| Fdx1l | 6,88 | 0,0027 | 0,996645926 | chr9 | - |
| Fdxacb1 | 90,31 | -0,1475 | 0,773303025 | chr9 | + |
| Fem1b | 444,60 | 0,1306 | 0,739141404 | chr9 | - |
| Filip1 | 10,78 | 0,5575 | 0,385438359 | chr9 | - |
| Fli1 | 62,83 | 0,4817 | 0,273713146 | chr9 | - |
| Foxr1 | 3,52 | 0,1850 | 0,823949825 | chr9 | - |
| Foxred1 | 643,35 | -0,4041 | 0,144604465 | chr9 | - |
| Fut4 | 4,34 | -0,2867 | 0,712237567 | chr9 | - |
| Fxyd2 | 2,17 | 0,2110 | NA | chr9 | + |
| Fxyd6 | 10,52 | 0,3467 | 0,637627461 | chr9 | + |
| Fyco1 | 603,68 | 0,0083 | 0,990189789 | chr9 | - |
| Gclc | 3176,26 | -0,4363 | 0,175625239 | chr9 | + |
| Gcm1 | 2,05 | -0,1899 | NA | chr9 | + |
| Glb1 | 214,54 | -0,2312 | 0,533151056 | chr9 | + |
| Glce | 491,85 | 0,3958 | 0,227218378 | chr9 | - |
| Gldn | 2,19 | 0,6815 | NA | chr9 | + |
| Glyctk | 3208,25 | 0,0580 | 0,904903021 | chr9 | - |
| Gm10653 | 22,23 | -0,2030 | 0,761732176 | chr9 | - |
| Gm16845 | 5,98 | -0,2398 | 0,772705111 | chr9 | + |
| Mthfsl | 296,28 | 0,4052 | 0,354512892 | chr9 | - |
| Gm4791 | 10,29 | 0,4226 | 0,559634089 | chr9 | + |
| Gm5617 | 184,74 | 0,2618 | 0,701094188 | chr9 | + |
| Angptl8 | 401,82 | -0,1141 | 0,9024844 | chr9 | + |
| Colca2 | 3,57 | 0,3097 | 0,681592614 | chr9 | - |
| Gm6981 | 10,35 | -0,1711 | 0,841326785 | chr9 | - |
| Gmppb | 248,99 | -0,0188 | 0,9774226 | chr9 | + |

| | | | | | |
|---|---|---|---|---|---|
| Gnai2 | 776,48 | 0,1995 | 0,603013282 | chr9 | - |
| Gnat1 | 106,82 | 0,1555 | 0,862421004 | chr9 | - |
| Gnb5 | 12,47 | -0,3303 | 0,633851047 | chr9 | + |
| Golga4 | 772,09 | -0,0895 | 0,811137679 | chr9 | + |
| Gorasp1 | 373,78 | -0,0652 | 0,89147692 | chr9 | - |
| Gpd1l | 636,48 | -0,0677 | 0,896013441 | chr9 | - |
| Gramd1b | 11,04 | 0,9599 | 0,090916651 | chr9 | - |
| Gsta1 | 8,57 | -0,6202 | 0,346086865 | chr9 | + |
| Gsta2 | 136,63 | -0,6589 | 0,184748288 | chr9 | - |
| Gtf2a2 | 678,61 | -0,0502 | 0,891236226 | chr9 | + |
| Gucy1a2 | 35,49 | 0,3925 | 0,536622488 | chr9 | + |
| H2afx | 49,11 | -0,6352 | 0,225613833 | chr9 | + |
| Hcrtr2 | 20,34 | 0,6277 | 0,337436925 | chr9 | - |
| Hemk1 | 205,50 | 0,2509 | 0,559032648 | chr9 | - |
| Herc1 | 394,42 | -0,1199 | 0,738282454 | chr9 | + |
| Herpud2 | 320,68 | -0,3071 | 0,33288309 | chr9 | - |
| Hexa | 783,20 | -0,1099 | 0,785583218 | chr9 | + |
| Higd1a | 2148,79 | -0,2940 | 0,340289195 | chr9 | - |
| Hinfp | 45,01 | -0,0841 | 0,903337956 | chr9 | - |
| Hmbs | 655,36 | -0,5137 | 0,073430016 | chr9 | - |
| Hmg20a | 178,38 | 0,1103 | 0,808864467 | chr9 | + |
| Hmgn3 | 11,45 | -0,0762 | 0,93871506 | chr9 | - |
| Hspa8 | 25636,42 | -0,1099 | 0,844029674 | chr9 | + |
| Hspb2 | 3,14 | 0,0934 | 0,918104715 | chr9 | - |
| Hyal1 | 230,62 | 0,1203 | 0,820554805 | chr9 | + |
| Hyal2 | 585,24 | -0,0705 | 0,913936879 | chr9 | + |
| Hyal3 | 22,44 | 0,1858 | 0,806012703 | chr9 | + |
| Hykk | 1377,08 | -0,0276 | 0,948314012 | chr9 | + |
| Hyls1 | 34,74 | -0,6631 | 0,158831698 | chr9 | - |
| Hyou1 | 2008,63 | 0,0758 | 0,916852736 | chr9 | + |
| Ibtk | 920,31 | -0,0250 | 0,95728901 | chr9 | - |
| Icam1 | 152,19 | 0,1077 | 0,842655851 | chr9 | + |
| Icam4 | 9,56 | -0,6900 | 0,258301555 | chr9 | + |
| Ick | 178,30 | -0,2894 | 0,423089018 | chr9 | + |
| Idh3a | 474,42 | -0,2510 | 0,540223765 | chr9 | + |
| Ifrd2 | 394,97 | -0,5098 | 0,208914053 | chr9 | + |
| Ift46 | 606,13 | -0,2661 | 0,515706574 | chr9 | + |
| Igdcc4 | 3,61 | 0,3556 | 0,618749555 | chr9 | + |
| Il10ra | 60,66 | 0,7734 | 0,066591982 | chr9 | - |
| Il18 | 355,29 | -0,2738 | 0,510450019 | chr9 | + |
| Il20rb | 11,78 | -0,2814 | 0,706352211 | chr9 | - |
| Ilf3 | 263,44 | 0,3074 | 0,449778593 | chr9 | + |
| Imp3 | 425,86 | -0,3513 | 0,277831408 | chr9 | + |
| Impdh2 | 111,90 | 0,3078 | 0,437750207 | chr9 | + |
| Ip6k1 | 939,37 | -0,0820 | 0,809686855 | chr9 | + |
| Ip6k2 | 504,31 | -0,5870 | 0,064539659 | chr9 | + |
| Irak1bp1 | 28,57 | 0,3689 | 0,553294406 | chr9 | + |
| Ireb2 | 1748,68 | -0,1349 | 0,736302391 | chr9 | + |
| Islr | 51,28 | 0,1101 | 0,863048009 | chr9 | - |
| Itga11 | 2,66 | 0,0767 | 0,932531696 | chr9 | + |

| | | | | | |
|---|---|---|---|---|---|
| Itga9 | 235,18 | 0,2571 | 0,505965828 | chr9 | + |
| Jam3 | 23,18 | 0,1588 | 0,822514387 | chr9 | - |
| Jrkl | 69,63 | 0,6218 | 0,119805774 | chr9 | - |
| Kank2 | 463,80 | 0,2414 | 0,501991823 | chr9 | - |
| Kbtbd3 | 87,66 | 0,0475 | 0,952532092 | chr9 | + |
| Kcnj5 | 4,60 | 0,2929 | 0,711559683 | chr9 | - |
| Kdelc2 | 158,69 | 0,1012 | 0,83956294 | chr9 | + |
| Keap1 | 649,13 | -0,1520 | 0,70612403 | chr9 | - |
| Kif15 | 9,88 | 0,6631 | 0,290981086 | chr9 | + |
| Kif23 | 16,16 | -0,0457 | 0,961556353 | chr9 | - |
| Kif9 | 13,56 | -0,1728 | 0,830330662 | chr9 | + |
| Kirrel3 | 2,67 | -0,4593 | 0,422047896 | chr9 | + |
| Klhdc8b | 87,59 | 0,0504 | 0,951372792 | chr9 | - |
| Klhl18 | 82,42 | -0,0433 | 0,952639267 | chr9 | - |
| Kmt2a | 9857,87 | 0,3061 | 0,415252477 | chr9 | - |
| Kri1 | 152,02 | 0,0018 | 0,997464504 | chr9 | - |
| Lamb2 | 59,35 | -0,1656 | 0,809981003 | chr9 | + |
| Lars2 | 40098,13 | -1,0166 | 0,073845576 | chr9 | + |
| Layn | 10,00 | 0,8389 | 0,157586062 | chr9 | - |
| Lca5 | 3,46 | 0,3179 | 0,671272065 | chr9 | - |
| Ldlr | 1783,14 | 0,0264 | 0,95990578 | chr9 | + |
| Leo1 | 130,98 | 0,1403 | 0,788139622 | chr9 | + |
| Limd1 | 499,31 | 0,0220 | 0,96345382 | chr9 | + |
| Lipc | 5380,30 | 0,1647 | 0,664082981 | chr9 | - |
| Loxl1 | 16,86 | 0,1960 | 0,806012703 | chr9 | - |
| Lrrc1 | 6,04 | 0,1442 | 0,874583899 | chr9 | - |
| Lrrc2 | 23,84 | 0,0268 | 0,976788115 | chr9 | + |
| Lrrc49 | 8,22 | 0,7357 | 0,237903839 | chr9 | - |
| Lrrfip2 | 569,29 | -0,5071 | 0,067117612 | chr9 | + |
| Lysmd2 | 13,54 | 0,5264 | 0,415252477 | chr9 | + |
| Lztfl1 | 322,66 | 0,0233 | 0,973932327 | chr9 | - |
| Maml2 | 36,89 | -0,0706 | 0,917614646 | chr9 | + |
| Man2c1 | 264,20 | -0,0510 | 0,938244876 | chr9 | + |
| Man2c1os | 11,08 | -0,2418 | 0,764664236 | chr9 | - |
| Manf | 1979,77 | 0,1699 | 0,847419446 | chr9 | - |
| Map2k1 | 738,29 | -0,0851 | 0,796014393 | chr9 | - |
| Map2k5 | 257,87 | -0,3812 | 0,175457889 | chr9 | - |
| Map4 | 504,87 | 0,0223 | 0,962739078 | chr9 | + |
| Mapk6 | 634,62 | 0,1286 | 0,773710065 | chr9 | - |
| Mapkapk3 | 80,84 | -0,1338 | 0,820554805 | chr9 | - |
| Mb21d1 | 6,88 | -0,0975 | 0,918077233 | chr9 | - |
| Mcam | 100,69 | 0,4650 | 0,208518411 | chr9 | + |
| Me1 | 4556,17 | -0,2942 | 0,570446186 | chr9 | - |
| Med17 | 200,31 | -0,1149 | 0,765219833 | chr9 | - |
| Mir6236 | 3916,18 | -0,9951 | 0,085593171 | chr9 | + |
| Mlh1 | 243,96 | 0,4220 | 0,172087384 | chr9 | - |
| Mmp13 | 10,95 | 0,7551 | 0,217465538 | chr9 | + |
| Mmp27 | 13,42 | -0,5889 | 0,314452617 | chr9 | + |
| Mmp8 | 19,27 | 0,2349 | 0,742089796 | chr9 | + |
| Mon1a | 151,00 | -0,6203 | 0,147003641 | chr9 | + |

| | | | | | |
|---|---|---|---|---|---|
| Morf4l1 | 2296,62 | 0,0892 | 0,827995984 | chr9 | - |
| Mpi | 255,67 | -0,1071 | 0,821346657 | chr9 | - |
| Mpzl2 | 715,13 | 0,2745 | 0,47939388 | chr9 | + |
| Mpzl3 | 45,03 | 0,4640 | 0,3698416 | chr9 | + |
| Mrap2 | 2,23 | -0,2355 | NA | chr9 | + |
| Mras | 42,67 | -0,1886 | 0,743508016 | chr9 | - |
| Mre11a | 47,97 | 0,4612 | 0,349661709 | chr9 | + |
| Mrpl3 | 292,41 | -0,0339 | 0,952710435 | chr9 | + |
| Mrpl4 | 292,44 | -0,5276 | 0,103115599 | chr9 | + |
| Mrps22 | 493,50 | -0,2674 | 0,465196538 | chr9 | - |
| Msantd2 | 96,30 | -0,3270 | 0,42048967 | chr9 | + |
| Msantd4 | 499,20 | 0,1011 | 0,831759716 | chr9 | + |
| Msl2 | 445,58 | -0,1326 | 0,695967514 | chr9 | + |
| Mst1 | 1989,52 | -0,2438 | 0,455499332 | chr9 | + |
| Mst1r | 3,54 | 0,3567 | 0,629873977 | chr9 | + |
| Mtfmt | 103,86 | 0,0343 | 0,953805583 | chr9 | + |
| Mthfs | 220,95 | 0,3312 | 0,52853113 | chr9 | + |
| Mtmr2 | 400,94 | -0,0653 | 0,895663283 | chr9 | + |
| Myd88 | 193,13 | -0,3927 | 0,226113234 | chr9 | - |
| Myo1e | 521,86 | 0,2315 | 0,549054641 | chr9 | + |
| Myo5a | 13,36 | 0,6941 | 0,244197385 | chr9 | + |
| Myo5c | 8,01 | 0,8276 | 0,17526415 | chr9 | + |
| Myo6 | 754,94 | -0,4510 | 0,148467563 | chr9 | + |
| Myo9a | 128,52 | 0,2626 | 0,499808492 | chr9 | + |
| Myrip | 3,19 | 0,2304 | 0,7599238 | chr9 | + |
| Myzap | 32,92 | 0,0808 | 0,909774279 | chr9 | - |
| Naalad2 | 22,30 | 0,1450 | 0,850270865 | chr9 | - |
| Ice2 | 212,06 | 0,4848 | 0,112715106 | chr9 | + |
| Nat6 | 531,79 | -0,3364 | 0,343973842 | chr9 | + |
| Nbeal2 | 55,05 | -0,4572 | 0,428649604 | chr9 | - |
| Ncam1 | 6,78 | 0,3507 | 0,644687571 | chr9 | - |
| Ncapd3 | 29,65 | 0,4814 | 0,379104437 | chr9 | + |
| Nck1 | 463,11 | -0,2091 | 0,598404146 | chr9 | - |
| Nckipsd | 40,17 | -0,1904 | 0,775458257 | chr9 | + |
| Ndufaf3 | 224,62 | -0,1231 | 0,810346289 | chr9 | - |
| Nedd4 | 3034,89 | 0,2098 | 0,420273711 | chr9 | + |
| Neil1 | 63,78 | -0,0432 | 0,949779983 | chr9 | - |
| Neo1 | 264,48 | -0,0159 | 0,9774226 | chr9 | - |
| Nfrkb | 111,00 | -0,2581 | 0,499958086 | chr9 | + |
| Ngp | 3,84 | 0,6783 | 0,261778745 | chr9 | + |
| Nicn1 | 214,71 | -0,0985 | 0,843102154 | chr9 | + |
| Nktr | 582,22 | -0,1274 | 0,736302391 | chr9 | + |
| Nlrx1 | 93,75 | 0,2133 | 0,625413559 | chr9 | - |
| Nme6 | 412,34 | -0,1133 | 0,825918365 | chr9 | + |
| Nmnat3 | 77,18 | 0,5063 | 0,327217799 | chr9 | + |
| Nnmt | 4065,62 | 0,0899 | 0,895663283 | chr9 | - |
| Npat | 124,68 | 0,2949 | 0,411158799 | chr9 | + |
| Nphp3 | 69,10 | -0,0941 | 0,871122558 | chr9 | + |
| Nprl2 | 242,53 | -0,2610 | 0,501951423 | chr9 | + |
| Nptn | 1196,18 | 0,1300 | 0,715232158 | chr9 | + |

| | | | | | |
|---|---|---|---|---|---|
| Nradd | 10,49 | -0,5917 | 0,36985369 | chr9 | - |
| Nrg4 | 157,93 | -0,3459 | 0,361776578 | chr9 | - |
| Ntm | 15,97 | 0,0665 | 0,942216943 | chr9 | - |
| Nudt16 | 123,45 | 0,3332 | 0,449506818 | chr9 | - |
| Nxpe2 | 332,60 | -0,4311 | 0,300779985 | chr9 | - |
| Nxpe4 | 18,95 | 0,4781 | 0,459906177 | chr9 | + |
| Oaf | 4419,30 | -0,0976 | 0,823949825 | chr9 | - |
| Oaz2 | 565,76 | -0,0267 | 0,955234619 | chr9 | + |
| Olfm2 | 2,85 | -0,0004 | 0,999351454 | chr9 | - |
| Olfr856-ps1 | 23,81 | -0,2356 | 0,722057512 | chr9 | - |
| Olfr920 | 2,33 | 0,1187 | NA | chr9 | + |
| Onecut1 | 32,04 | 0,0952 | 0,906333899 | chr9 | + |
| Osbpl10 | 3,25 | -0,1913 | 0,815188933 | chr9 | + |
| Oxsr1 | 494,80 | -0,2159 | 0,528345204 | chr9 | - |
| P4htm | 2,20 | -0,0965 | NA | chr9 | - |
| Pafah1b2 | 529,81 | -0,1550 | 0,635764582 | chr9 | - |
| Paqr9 | 1400,58 | -0,1884 | 0,636447281 | chr9 | + |
| Parp3 | 313,16 | 0,0665 | 0,912418045 | chr9 | - |
| Parp6 | 152,46 | -0,5057 | 0,100266579 | chr9 | + |
| Pcbp4 | 154,41 | -0,3042 | 0,558926649 | chr9 | + |
| Pccb | 90,83 | 0,2978 | 0,449196578 | chr9 | - |
| Pcolce2 | 45,50 | -0,1819 | 0,739141404 | chr9 | + |
| Pcsk7 | 494,15 | -0,3331 | 0,265239441 | chr9 | + |
| Pdcd6ip | 951,90 | -0,1917 | 0,504278019 | chr9 | - |
| Pdcd7 | 93,77 | -0,0993 | 0,842502441 | chr9 | + |
| Pde4a | 48,97 | -0,1966 | 0,786227852 | chr9 | + |
| Pdgfd | 13,40 | 0,9597 | 0,076968799 | chr9 | + |
| Peak1 | 186,39 | 0,0556 | 0,916852736 | chr9 | - |
| Pfkfb4 | 15,28 | -0,3636 | 0,606429785 | chr9 | + |
| Pgm3 | 371,66 | 0,5166 | 0,227796217 | chr9 | - |
| Phip | 285,79 | 0,3631 | 0,336364632 | chr9 | - |
| Phldb1 | 56,38 | 0,7185 | 0,083560273 | chr9 | - |
| Pias1 | 198,12 | 0,0020 | 0,996823488 | chr9 | - |
| Pigb | 41,11 | 0,2906 | 0,632136487 | chr9 | - |
| Pigyl | 230,94 | -0,4610 | 0,517431746 | chr9 | + |
| Pih1d2 | 13,62 | 0,5671 | 0,35583041 | chr9 | + |
| Pik3cb | 193,29 | -0,2108 | 0,658107384 | chr9 | - |
| Pik3r4 | 234,57 | -0,0936 | 0,851472884 | chr9 | + |
| Pin1 | 267,68 | -0,2023 | 0,708761088 | chr9 | + |
| Pkm | 284,47 | 0,3491 | 0,339404199 | chr9 | + |
| Pknox2 | 6,37 | -0,3552 | 0,63960646 | chr9 | - |
| Plcd1 | 4,20 | -0,0306 | 0,977590816 | chr9 | - |
| Plekho2 | 79,27 | 0,3009 | 0,522722304 | chr9 | - |
| Plet1 | 75,92 | 0,5176 | 0,229293047 | chr9 | + |
| Plet1os | 40881,16 | -0,0636 | 0,831759716 | chr9 | - |
| Plod2 | 264,72 | -0,3646 | 0,321255221 | chr9 | + |
| Pls1 | 1853,88 | 0,1259 | 0,773900067 | chr9 | - |
| Plscr1 | 414,15 | -0,1856 | 0,724652712 | chr9 | + |
| Plscr2 | 80,25 | 0,0294 | 0,963027434 | chr9 | + |
| Pml | 195,58 | 0,1945 | 0,67554326 | chr9 | - |

| | | | | | |
|---|---|---|---|---|---|
| Poc1a | 33,29 | 0,2713 | 0,629243929 | chr9 | + |
| Polr2m | 998,02 | -0,1220 | 0,765061508 | chr9 | - |
| Pomgnt2 | 26,08 | 0,3833 | 0,491510849 | chr9 | - |
| Pou2af1 | 4,52 | 0,0578 | 0,953638766 | chr9 | + |
| Ppan | 85,46 | -0,6453 | 0,181404581 | chr9 | + |
| Ppcdc | 193,73 | -0,0381 | 0,94694207 | chr9 | - |
| Ppib | 8369,01 | -0,4113 | 0,189774146 | chr9 | + |
| Ppm1m | 190,93 | -0,3414 | 0,313893111 | chr9 | - |
| Ppp2r1b | 288,36 | -0,0779 | 0,866934682 | chr9 | + |
| Ppp2r3a | 213,20 | -0,0693 | 0,899643166 | chr9 | - |
| Ppp2r3d | 25,39 | 0,2258 | 0,753233421 | chr9 | - |
| Prdm10 | 12,65 | 0,0147 | 0,988512389 | chr9 | + |
| Prkar2a | 698,91 | -0,0112 | 0,98507181 | chr9 | + |
| Prkcsh | 1318,58 | -0,2546 | 0,562610061 | chr9 | + |
| Prss35 | 2,97 | -0,0311 | 0,976788115 | chr9 | + |
| Psma4 | 3968,04 | -0,0669 | 0,88569455 | chr9 | + |
| Pstpip1 | 6,11 | 0,6498 | 0,319080738 | chr9 | + |
| Pth1r | 92,19 | -0,3826 | 0,39354827 | chr9 | - |
| Hacd3 | 3631,62 | -0,1244 | 0,757883527 | chr9 | - |
| Ptpn23 | 134,61 | -0,3559 | 0,507875164 | chr9 | - |
| Ptpn9 | 263,93 | 0,1870 | 0,61737806 | chr9 | + |
| Pts | 664,49 | -0,1205 | 0,842655851 | chr9 | - |
| Pus3 | 103,80 | 0,2243 | 0,633486481 | chr9 | + |
| Nectin1 | 297,18 | 0,1302 | 0,814990243 | chr9 | + |
| Pxylp1 | 98,72 | 0,3882 | 0,343554244 | chr9 | - |
| Pygo1 | 2,30 | 0,3363 | NA | chr9 | + |
| Qars | 945,89 | -0,3507 | 0,208381949 | chr9 | + |
| Qrich1 | 543,16 | -0,1198 | 0,710761695 | chr9 | + |
| Qtrt1 | 41,95 | 0,0215 | 0,979212524 | chr9 | + |
| Rab11a | 1110,89 | -0,2201 | 0,531668406 | chr9 | - |
| Rab39 | 14,10 | 0,4014 | 0,54375285 | chr9 | - |
| Rab3d | 134,10 | -0,2489 | 0,540699632 | chr9 | - |
| Rab6b | 19,89 | -0,1229 | 0,882318627 | chr9 | + |
| Rab8b | 121,36 | 0,4522 | 0,223366879 | chr9 | - |
| Rad54l2 | 530,86 | 0,2538 | 0,563960133 | chr9 | - |
| Rasl12 | 2,95 | 0,2797 | 0,708319716 | chr9 | + |
| Rassf1 | 111,27 | -0,3628 | 0,419953461 | chr9 | + |
| Raver1 | 101,24 | -0,5969 | 0,164406774 | chr9 | - |
| Rbm15b | 71,61 | 0,1216 | 0,850453487 | chr9 | - |
| Rbm5 | 510,12 | 0,0592 | 0,8859355 | chr9 | - |
| Rbm6 | 410,27 | 0,0231 | 0,961798305 | chr9 | - |
| Rbm7 | 876,73 | 0,1205 | 0,796765648 | chr9 | - |
| Rbms3 | 97,36 | 0,4573 | 0,339404075 | chr9 | - |
| Rbpms2 | 618,45 | -0,4556 | 0,137147675 | chr9 | + |
| Rcn2 | 121,87 | -0,2176 | 0,663934341 | chr9 | + |
| Rdx | 3162,13 | -0,0476 | 0,900660395 | chr9 | + |
| Rexo2 | 1137,79 | -0,4790 | 0,069540078 | chr9 | - |
| Rfx7 | 214,06 | -0,0967 | 0,814373908 | chr9 | + |
| Rgl3 | 141,06 | -0,2032 | 0,741074355 | chr9 | - |
| Rhoa | 3146,55 | -0,1044 | 0,704653924 | chr9 | + |

| | | | | | |
|---|---|---|---|---|---|
| Rnf111 | 287,37 | -0,4060 | 0,204417741 | chr9 | - |
| Rnf123 | 518,19 | -0,3363 | 0,366528137 | chr9 | - |
| Rnf214 | 205,76 | -0,2723 | 0,527966188 | chr9 | - |
| Rnf26 | 128,20 | -0,2001 | 0,672801255 | chr9 | - |
| Rnf7 | 618,84 | -0,1804 | 0,558841064 | chr9 | - |
| Robo4 | 131,65 | -0,1722 | 0,701666523 | chr9 | + |
| Rora | 1355,37 | -0,2065 | 0,571686775 | chr9 | + |
| Rpl29 | 3329,93 | -0,3159 | 0,395294986 | chr9 | + |
| Rpl4 | 15025,65 | -0,2233 | 0,444852563 | chr9 | + |
| Rplp1 | 12661,86 | -0,2555 | 0,755785289 | chr9 | - |
| Rps25 | 9242,02 | -0,3519 | 0,234431867 | chr9 | + |
| Rps27l | 5216,09 | -0,5491 | 0,116602928 | chr9 | + |
| Rpusd4 | 139,38 | 0,3950 | 0,283971503 | chr9 | + |
| Rrp9 | 89,40 | -0,2385 | 0,736631536 | chr9 | + |
| Rsl24d1 | 590,64 | -0,1330 | 0,743124066 | chr9 | + |
| Rtp3 | 817,93 | -0,3202 | 0,31302658 | chr9 | - |
| Rwdd2a | 2,40 | 0,2537 | NA | chr9 | + |
| Ryk | 3432,03 | -0,3624 | 0,154257003 | chr9 | + |
| S1pr2 | 82,14 | 0,2428 | 0,585752237 | chr9 | - |
| S1pr5 | 60,51 | -0,2356 | 0,713576775 | chr9 | - |
| Sacm1l | 1455,89 | -0,0199 | 0,969512928 | chr9 | + |
| Sc5d | 7159,22 | 0,2524 | 0,55986821 | chr9 | - |
| Scamp2 | 304,05 | 0,0090 | 0,985488456 | chr9 | + |
| Scamp5 | 326,19 | 0,4101 | 0,205100332 | chr9 | - |
| Scap | 1104,95 | -0,2124 | 0,577347444 | chr9 | + |
| Scaper | 105,48 | 0,3964 | 0,481381033 | chr9 | - |
| Scn2b | 2,83 | -0,2173 | 0,777753633 | chr9 | + |
| Sdhd | 7787,13 | -0,1796 | 0,606323442 | chr9 | - |
| Sec22c | 193,37 | -0,4092 | 0,266061606 | chr9 | - |
| Sema3b | 6,57 | 0,6903 | 0,281403962 | chr9 | - |
| Sema3f | 42,60 | -0,0903 | 0,918946301 | chr9 | - |
| Senp6 | 709,28 | 0,1503 | 0,651354867 | chr9 | + |
| Senp8 | 75,35 | 0,4914 | 0,271262155 | chr9 | - |
| Sesn3 | 36,82 | 0,7033 | 0,138593946 | chr9 | + |
| Setd2 | 468,22 | -0,2202 | 0,470196942 | chr9 | + |
| Sh3bgrl2 | 682,63 | -0,4818 | 0,152264308 | chr9 | + |
| Shisa5 | 635,59 | 0,1774 | 0,55219958 | chr9 | + |
| Siae | 849,95 | 0,4346 | 0,187852299 | chr9 | + |
| Sidt2 | 1412,35 | -0,3313 | 0,29013293 | chr9 | - |
| Sik2 | 55,53 | 0,2296 | 0,657542517 | chr9 | - |
| Sik3 | 451,19 | -0,0490 | 0,936911918 | chr9 | + |
| Sin3a | 364,06 | 0,1149 | 0,754249138 | chr9 | + |
| Slc17a5 | 485,24 | -0,3214 | 0,281016665 | chr9 | - |
| Slc22a13b-ps | 2,50 | -0,4918 | NA | chr9 | - |
| Slc25a20 | 2415,80 | -0,2981 | 0,479398672 | chr9 | + |
| Slc25a36 | 34,30 | 0,7371 | 0,117947615 | chr9 | - |
| Slc26a6 | 24,87 | -0,4493 | 0,446707429 | chr9 | + |
| Slc35g2 | 6,81 | 0,0906 | 0,925700434 | chr9 | - |
| Slc36a4 | 24,26 | 0,8139 | 0,094901224 | chr9 | + |
| Slc37a2 | 32,93 | 0,4829 | 0,333905884 | chr9 | - |

| | | | | | |
|---|---|---|---|---|---|
| Slc37a4 | 5637,67 | 0,0605 | 0,90199272 | chr9 | + |
| Slc38a3 | 19896,75 | -0,2272 | 0,443146136 | chr9 | - |
| Slc44a2 | 125,39 | 0,1797 | 0,67848774 | chr9 | + |
| Slc51b | 200,16 | 0,3191 | 0,343973842 | chr9 | - |
| Slc9a9 | 110,62 | 0,4582 | 0,190969312 | chr9 | + |
| Slco2a1 | 979,12 | -0,3323 | 0,36387652 | chr9 | + |
| Sltm | 535,62 | -0,2051 | 0,522874974 | chr9 | + |
| Smad3 | 198,27 | 0,1306 | 0,800238939 | chr9 | - |
| Smad6 | 25,07 | 0,2163 | 0,776246447 | chr9 | - |
| Smarca4 | 353,44 | 0,0338 | 0,95463264 | chr9 | + |
| Smarcc1 | 303,33 | -0,0741 | 0,866934682 | chr9 | + |
| Smco4 | 148,25 | 0,5810 | 0,094600644 | chr9 | + |
| Snapc5 | 703,65 | -0,0314 | 0,945274006 | chr9 | + |
| Snhg5 | 143,99 | -0,4505 | 0,324242521 | chr9 | - |
| Snord16a | 2,81 | -0,0053 | 0,995750189 | chr9 | + |
| Snrk | 772,74 | -0,1185 | 0,75378365 | chr9 | + |
| Snupn | 127,59 | 0,2762 | 0,588008527 | chr9 | + |
| Snx1 | 344,36 | 0,3805 | 0,228941765 | chr9 | - |
| Snx14 | 467,76 | -0,0321 | 0,948937066 | chr9 | - |
| Snx19 | 382,67 | 0,1477 | 0,769677581 | chr9 | + |
| Snx22 | 14,18 | -0,0517 | 0,955440194 | chr9 | - |
| Snx33 | 191,55 | 0,4723 | 0,198455103 | chr9 | - |
| Spa17 | 28,75 | 0,6016 | 0,246435345 | chr9 | - |
| Spc24 | 140,43 | -0,1245 | 0,814733578 | chr9 | - |
| Spg21 | 452,20 | 0,0940 | 0,827739239 | chr9 | + |
| Spsb4 | 57,76 | 0,4003 | 0,506539539 | chr9 | - |
| Srpr | 2894,11 | -0,2784 | 0,437263599 | chr9 | + |
| Srprb | 425,36 | -0,1816 | 0,732392788 | chr9 | - |
| St14 | 11,77 | 0,6711 | 0,286608409 | chr9 | - |
| St3gal4 | 1446,19 | 0,4236 | 0,18870514 | chr9 | - |
| Stoml1 | 45,97 | 0,1442 | 0,80096016 | chr9 | + |
| Stt3a | 2661,99 | 0,2289 | 0,462174993 | chr9 | - |
| Stt3b | 1520,21 | -0,1419 | 0,64665416 | chr9 | - |
| Swsap1 | 90,34 | 0,3811 | 0,360147619 | chr9 | + |
| Syncrip | 800,68 | 0,0612 | 0,888963178 | chr9 | - |
| Taf1d | 212,49 | -0,4310 | 0,238321283 | chr9 | + |
| Tagln | 219,53 | 0,7025 | 0,07041 | chr9 | - |
| Tbc1d2b | 355,99 | -0,0785 | 0,869067375 | chr9 | - |
| Tbcel | 869,07 | -0,4351 | 0,166993075 | chr9 | - |
| Tbrg1 | 715,29 | 0,4441 | 0,162648376 | chr9 | - |
| Tbx20 | 40,13 | 0,2809 | 0,651757298 | chr9 | - |
| Tcaim | 372,95 | 0,5186 | 0,154183797 | chr9 | + |
| Tcf12 | 293,81 | 0,2306 | 0,566893799 | chr9 | - |
| Tcta | 272,94 | -0,1778 | 0,636447281 | chr9 | - |
| Tex12 | 7,27 | -0,9113 | 0,127029796 | chr9 | - |
| Tex264 | 1033,01 | -0,1210 | 0,793247794 | chr9 | - |
| Tex9 | 11,55 | 0,2448 | 0,764664236 | chr9 | - |
| Tfdp2 | 268,24 | -0,3634 | 0,174553196 | chr9 | + |
| Tgfbr2 | 393,97 | 0,0149 | 0,98429278 | chr9 | - |
| Thsd4 | 12,32 | 0,2090 | 0,804110902 | chr9 | - |

| | | | | | |
|---|---|---|---|---|---|
| Thy1 | 15,01 | 0,0518 | 0,955504626 | chr9 | + |
| Thyn1 | 390,00 | 0,0280 | 0,960906846 | chr9 | + |
| Timm8b | 1379,79 | -0,4300 | 0,145508611 | chr9 | + |
| Tipin | 132,20 | -0,0114 | 0,988512389 | chr9 | + |
| Tirap | 411,35 | 0,4485 | 0,080356911 | chr9 | - |
| Tle3 | 100,23 | -0,2629 | 0,570187994 | chr9 | + |
| Tln2 | 83,90 | 0,0193 | 0,979917083 | chr9 | - |
| Tlr9 | 2,83 | 0,7696 | 0,172473519 | chr9 | + |
| Tma7 | 1843,05 | 0,0772 | 0,823537093 | chr9 | - |
| Tmed1 | 239,33 | -0,3882 | 0,354116406 | chr9 | - |
| Tmed3 | 4354,95 | -0,2289 | 0,442976355 | chr9 | - |
| Tmem108 | 2,11 | 0,3795 | NA | chr9 | - |
| Tmem115 | 262,21 | -0,3841 | 0,319381722 | chr9 | + |
| Tmem123 | 789,33 | 0,5689 | 0,077346873 | chr9 | + |
| Tmem136 | 14,57 | 0,4340 | 0,507587541 | chr9 | - |
| Tmem158 | 3,50 | -0,1087 | 0,905147397 | chr9 | - |
| Tmem205 | 11758,47 | -0,4479 | 0,067117612 | chr9 | - |
| Tmem218 | 66,33 | 0,0820 | 0,890761998 | chr9 | + |
| Tmem25 | 398,84 | -0,2705 | 0,568215867 | chr9 | - |
| Tmem30a | 3436,53 | -0,2310 | 0,570516869 | chr9 | - |
| Tmem42 | 92,12 | -0,3631 | 0,361776578 | chr9 | + |
| Tmod3 | 415,51 | 0,2190 | 0,464131878 | chr9 | - |
| Tmppe | 86,67 | -0,4262 | 0,217530209 | chr9 | + |
| Tnfaip8l3 | 7,84 | 0,2703 | 0,739874656 | chr9 | - |
| Topbp1 | 138,70 | 0,1717 | 0,693432851 | chr9 | + |
| Tpbg | 2,26 | 0,0919 | NA | chr9 | + |
| Tpm1 | 496,67 | 0,2723 | 0,402781375 | chr9 | - |
| Traip | 5,80 | 0,2192 | 0,795812563 | chr9 | + |
| Trak1 | 533,18 | -0,2394 | 0,528306301 | chr9 | + |
| Trappc4 | 501,64 | 0,1310 | 0,695288565 | chr9 | - |
| Treh | 19,61 | 0,8721 | 0,10357863 | chr9 | + |
| Trex1 | 99,14 | -0,1206 | 0,838412507 | chr9 | - |
| Trf | 245494,57 | -0,3811 | 0,162388298 | chr9 | - |
| Trim43b | 5,40 | 0,2553 | 0,756531051 | chr9 | - |
| Trip4 | 1279,88 | -0,3019 | 0,430392697 | chr9 | - |
| Trpc1 | 2,23 | -0,2395 | NA | chr9 | - |
| Trpc6 | 21,57 | -0,2794 | 0,684419282 | chr9 | + |
| Tspan3 | 74,22 | -0,1854 | 0,721569914 | chr9 | - |
| Ttc12 | 6,21 | 0,8428 | 0,16832806 | chr9 | - |
| Ttc36 | 9017,52 | -0,2260 | 0,378100794 | chr9 | - |
| Ttk | 6,55 | 0,5364 | 0,432040063 | chr9 | + |
| Tusc2 | 415,29 | -0,4607 | 0,116023315 | chr9 | + |
| Twf2 | 42,13 | 0,3673 | 0,511600714 | chr9 | + |
| Tyk2 | 267,82 | -0,0040 | 0,994676823 | chr9 | - |
| U2surp | 517,36 | 0,4349 | 0,106945732 | chr9 | - |
| Uaca | 132,39 | 0,1522 | 0,764363417 | chr9 | + |
| Uba5 | 545,07 | -0,1396 | 0,732393059 | chr9 | - |
| Ubash3b | 16,84 | 0,3527 | 0,613341495 | chr9 | - |
| Ube2cbp | 28,53 | -0,0975 | 0,890761998 | chr9 | - |
| Ube2q2 | 538,23 | -0,2250 | 0,557507087 | chr9 | + |

| | | | | | |
|---|---|---|---|---|---|
| Ube4a | 1128,86 | -0,1436 | 0,726344964 | chr9 | - |
| Ubl5 | 2354,30 | -0,3023 | 0,308448866 | chr9 | + |
| Ubl7 | 390,02 | -0,2374 | 0,553294406 | chr9 | + |
| Ubp1 | 672,82 | -0,4088 | 0,129193513 | chr9 | + |
| Ulk3 | 90,34 | -0,2899 | 0,577036779 | chr9 | + |
| Uqcrc1 | 2562,76 | -0,4812 | 0,192926688 | chr9 | + |
| Usp19 | 505,22 | -0,2381 | 0,607182269 | chr9 | + |
| Usp28 | 113,78 | -0,3287 | 0,335264505 | chr9 | + |
| Usp3 | 371,01 | -0,3959 | 0,203863873 | chr9 | - |
| Usp4 | 1292,05 | -0,3268 | 0,31487946 | chr9 | + |
| Vill | 36,57 | 0,7996 | 0,068748156 | chr9 | + |
| Vipr1 | 433,09 | 0,1010 | 0,804418611 | chr9 | + |
| Dcaf1 | 354,66 | -0,1474 | 0,71822878 | chr9 | + |
| Vps11 | 322,11 | -0,2569 | 0,386872469 | chr9 | - |
| Vps13c | 291,18 | 0,1659 | 0,705258097 | chr9 | + |
| Vps26b | 427,94 | -0,2824 | 0,314431944 | chr9 | - |
| Vsig2 | 6,08 | -0,1216 | 0,896013441 | chr9 | + |
| Vstm5 | 2,55 | 0,1807 | NA | chr9 | + |
| Vwa5a | 707,00 | 0,5802 | 0,072751135 | chr9 | + |
| Ints14 | 199,28 | 0,0183 | 0,974449819 | chr9 | + |
| Wdr48 | 495,74 | 0,0700 | 0,855566249 | chr9 | + |
| Wdr6 | 206,43 | -0,2856 | 0,496113427 | chr9 | - |
| Wdr61 | 1447,33 | -0,0191 | 0,969870909 | chr9 | - |
| Wdr82 | 813,69 | -0,0741 | 0,861046652 | chr9 | + |
| Xcr1 | 22,10 | -0,0534 | 0,953547526 | chr9 | - |
| Xirp1 | 6,12 | 0,7791 | 0,208828318 | chr9 | - |
| Xrn1 | 50,26 | 0,5252 | 0,333857286 | chr9 | + |
| Yap1 | 1043,60 | -0,0922 | 0,788512632 | chr9 | - |
| Yipf2 | 435,43 | 0,0632 | 0,87553708 | chr9 | - |
| Zbtb16 | 258,84 | 0,4478 | 0,5294919 | chr9 | - |
| Zbtb38 | 72,72 | 0,4068 | 0,302314196 | chr9 | - |
| Zbtb44 | 784,50 | -0,2192 | 0,541556317 | chr9 | + |
| Zc3h12c | 97,57 | 0,2165 | 0,637958595 | chr9 | - |
| Zdhhc3 | 335,56 | -0,0289 | 0,959186125 | chr9 | - |
| Zfp202 | 21,80 | 0,7660 | 0,125190331 | chr9 | + |
| Zfp280d | 182,71 | 0,4654 | 0,258527696 | chr9 | + |
| Zfp317 | 241,17 | 0,0877 | 0,865984359 | chr9 | + |
| Zfp560 | 78,62 | -0,0482 | 0,941754349 | chr9 | - |
| Zfp599 | 9,90 | 0,5448 | 0,420273711 | chr9 | - |
| Zfp609 | 138,32 | -0,0901 | 0,884364382 | chr9 | - |
| Zfp651 | 9,88 | 0,1808 | 0,830130691 | chr9 | + |
| Zfp653 | 43,04 | -0,3977 | 0,523898331 | chr9 | - |
| Zfp810 | 100,91 | 0,0542 | 0,937830007 | chr9 | - |
| Zfp846 | 55,71 | 0,1121 | 0,848745988 | chr9 | + |
| Zfp949 | 170,16 | -0,0633 | 0,89879469 | chr9 | + |
| Zmynd10 | 5,30 | -0,7172 | 0,258561012 | chr9 | + |
| Zw10 | 238,06 | -0,2992 | 0,41552947 | chr9 | + |
| Zwilch | 13,80 | 0,6376 | 0,283539488 | chr9 | - |
| 1190007I07Rik | 111,95 | -0,0524 | 0,949680678 | chr10 | - |
| 1600002K03Rik | 75,24 | -0,5087 | 0,184081758 | chr10 | + |

| | | | | | |
|---|---|---|---|---|---|
| 1700012D01Rik | 87,86 | -0,1346 | 0,84357068 | chr10 | + |
| 1700021F05Rik | 378,92 | 0,1422 | 0,716555628 | chr10 | - |
| Cabcoco1 | 25,92 | 0,3723 | 0,597620615 | chr10 | - |
| Armt1 | 245,98 | -0,2566 | 0,532105996 | chr10 | + |
| 1810043G02Rik | 49,95 | -0,3439 | 0,467424704 | chr10 | + |
| 2310011J03Rik | 196,64 | -0,2010 | 0,664063523 | chr10 | - |
| 2310039L15Rik | 27,60 | 0,6709 | 0,158861131 | chr10 | + |
| Kif1bp | 199,52 | -0,0216 | 0,972880126 | chr10 | - |
| 2610008E11Rik | 167,10 | 0,4616 | 0,236644041 | chr10 | - |
| 2610028H24Rik | 3,22 | 0,4100 | 0,543074584 | chr10 | + |
| Vsir | 156,88 | 0,5785 | 0,165273812 | chr10 | + |
| 4930430F08Rik | 98,83 | 0,1951 | 0,679628788 | chr10 | - |
| 4930519F09Rik | 8,20 | -0,1911 | 0,824901134 | chr10 | - |
| 4933412E12Rik | 9,82 | 0,3074 | 0,692557703 | chr10 | + |
| 5730420D15Rik | 2,33 | 0,2542 | NA | chr10 | + |
| 9330159F19Rik | 2,70 | 0,0229 | 0,981991128 | chr10 | + |
| A130077B15Rik | 445,71 | -0,1337 | 0,764893829 | chr10 | - |
| Washc4 | 434,13 | 0,5926 | 0,089263853 | chr10 | + |
| A330049N07Rik | 8,44 | 1,0003 | 0,083935391 | chr10 | - |
| A630066F11Rik | 13,29 | -0,3115 | 0,661313319 | chr10 | + |
| A830082N09Rik | 10,32 | -0,3087 | 0,67848774 | chr10 | - |
| Mfsd4b4 | 4,42 | 0,5615 | 0,398449964 | chr10 | - |
| Abca7 | 64,35 | -0,3330 | 0,570187994 | chr10 | + |
| Abhd17a | 363,05 | -0,4965 | 0,301759036 | chr10 | - |
| Abracl | 224,55 | -0,1985 | 0,62811896 | chr10 | - |
| Actr6 | 405,28 | 0,4725 | 0,217231457 | chr10 | - |
| Adamts14 | 14,54 | -0,0106 | 0,991447068 | chr10 | - |
| Adamtsl5 | 6,03 | -0,0967 | 0,919467514 | chr10 | - |
| Adarb1 | 165,59 | -0,1886 | 0,660583784 | chr10 | - |
| Adat2 | 50,14 | 0,1338 | 0,818837072 | chr10 | + |
| Adat3 | 5,62 | -0,0685 | 0,943064069 | chr10 | + |
| Adgb | 17,76 | -0,2624 | 0,712500177 | chr10 | - |
| Ado | 263,80 | -0,0412 | 0,938514017 | chr10 | - |
| Adora2a | 14,78 | 0,5749 | 0,357703576 | chr10 | + |
| Aes | 3425,48 | -0,4915 | 0,178097604 | chr10 | + |
| Agap2 | 33,35 | 0,6317 | 0,303602993 | chr10 | + |
| Agpat3 | 1930,89 | -0,2133 | 0,58887292 | chr10 | - |
| Ahi1 | 21,07 | 0,3316 | 0,606237187 | chr10 | + |
| Mfsd4b1 | 840,47 | -0,5165 | 0,074290268 | chr10 | - |
| Aifm2 | 483,17 | -0,1145 | 0,792197952 | chr10 | + |
| Aig1 | 100,81 | 0,5271 | 0,157879911 | chr10 | - |
| Akap12 | 69,49 | 0,2475 | 0,647914259 | chr10 | + |
| Akap7 | 111,72 | 0,6055 | 0,213281186 | chr10 | - |
| Aldh1l2 | 3,82 | -0,6234 | 0,314431944 | chr10 | - |
| Aldh8a1 | 6889,31 | -0,0292 | 0,947819654 | chr10 | + |
| Amdhd1 | 2341,65 | 0,0224 | 0,974809304 | chr10 | - |
| Anapc16 | 509,23 | -0,3771 | 0,356514018 | chr10 | - |
| Ank3 | 315,87 | 0,1002 | 0,860639142 | chr10 | + |
| Ankrd24 | 46,25 | 0,1581 | 0,803027202 | chr10 | + |
| Ankrd52 | 171,23 | -0,3754 | 0,430881045 | chr10 | + |

| | | | | | |
|---|---|---|---|---|---|
| Ap3d1 | 607,40 | -0,3016 | 0,352106106 | chr10 | - |
| Apaf1 | 56,76 | 0,5936 | 0,139399836 | chr10 | - |
| Apba3 | 271,93 | -0,3443 | 0,572115465 | chr10 | + |
| Apc2 | 2,43 | 0,0496 | NA | chr10 | + |
| Apof | 23406,81 | 0,3951 | 0,109852731 | chr10 | + |
| Apon | 5596,20 | -0,1916 | 0,74146296 | chr10 | + |
| Appl2 | 252,69 | 0,0188 | 0,973568634 | chr10 | - |
| Arg1 | 31191,94 | -0,0979 | 0,828120899 | chr10 | - |
| Arhgap18 | 250,59 | -0,0769 | 0,873325649 | chr10 | + |
| Arhgap9 | 33,25 | -0,0463 | 0,952541585 | chr10 | + |
| Arhgef25 | 20,00 | 0,2064 | 0,771616285 | chr10 | - |
| Arid3a | 14,02 | 0,1309 | 0,879637126 | chr10 | + |
| Arid5b | 170,00 | 0,1803 | 0,793250646 | chr10 | - |
| Arl1 | 1322,12 | 0,0545 | 0,910649603 | chr10 | + |
| Ascc1 | 202,44 | -0,0905 | 0,841053685 | chr10 | + |
| Ascc3 | 592,87 | -0,0868 | 0,839451425 | chr10 | + |
| Asf1a | 154,29 | -0,2203 | 0,623207643 | chr10 | + |
| Atg5 | 986,05 | -0,1994 | 0,543779254 | chr10 | + |
| Atp2b1 | 717,43 | 0,0654 | 0,881809902 | chr10 | + |
| Atp5b | 20996,99 | -0,3231 | 0,190441293 | chr10 | + |
| Atp5d | 2496,65 | -0,6268 | 0,198686541 | chr10 | + |
| Atxn7l3b | 1750,75 | 0,1675 | 0,630728761 | chr10 | - |
| Avil | 3,78 | -0,5283 | 0,423412429 | chr10 | + |
| B230208H11Rik | 12,01 | -0,1546 | 0,857478182 | chr10 | - |
| Baz2a | 402,53 | -0,1633 | 0,681592614 | chr10 | + |
| Bbs10 | 82,32 | 0,3378 | 0,525561659 | chr10 | + |
| BC020402 | 15,54 | -0,1618 | 0,834822458 | chr10 | - |
| Mfsd4b5 | 7,87 | 0,1185 | 0,89879469 | chr10 | - |
| BC025920 | 14,52 | 0,6550 | 0,287271906 | chr10 | + |
| Ttc41 | 38,39 | -0,3075 | 0,592295685 | chr10 | + |
| BC048403 | 42,25 | -0,0097 | 0,990189789 | chr10 | + |
| Bclaf1 | 913,26 | 0,2573 | 0,541556317 | chr10 | + |
| Bcr | 245,85 | -0,1220 | 0,801768605 | chr10 | + |
| Bend3 | 85,26 | -0,4765 | 0,17588728 | chr10 | + |
| Bloc1s1 | 1748,81 | -0,2972 | 0,357703576 | chr10 | - |
| Bsg | 5309,39 | -0,2339 | 0,645264604 | chr10 | + |
| Btbd11 | 8,70 | -0,4957 | 0,479068232 | chr10 | + |
| Btbd2 | 188,02 | -0,1875 | 0,655604557 | chr10 | - |
| Btg1 | 873,56 | -0,2451 | 0,471893954 | chr10 | + |
| C920009B18Rik | 2,58 | 0,3055 | 0,655452587 | chr10 | + |
| Cabin1 | 145,08 | 0,0570 | 0,916359044 | chr10 | - |
| Cactin | 90,28 | -0,0886 | 0,906081073 | chr10 | + |
| Cand1 | 853,64 | -0,1177 | 0,746827664 | chr10 | - |
| Ccar1 | 837,20 | -0,0580 | 0,891830745 | chr10 | - |
| Ccdc138 | 53,63 | 0,3971 | 0,444291084 | chr10 | + |
| Ccdc162 | 75,19 | 0,1155 | 0,851472884 | chr10 | - |
| Ccdc28a | 182,14 | -0,0202 | 0,968669888 | chr10 | - |
| Ccdc38 | 5,48 | -0,3701 | 0,623592138 | chr10 | + |
| Washc3 | 761,57 | -0,0793 | 0,85502909 | chr10 | + |
| Ccdc59 | 178,90 | 0,0029 | 0,995735598 | chr10 | + |

| | | | | | |
|---|---|---|---|---|---|
| Ccdc6 | 360,06 | -0,2891 | 0,277366051 | chr10 | + |
| Cct2 | 2014,00 | -0,2494 | 0,429633784 | chr10 | - |
| Cd164 | 6100,92 | -0,0762 | 0,833056752 | chr10 | + |
| Cd24a | 106,65 | 0,8290 | 0,058961472 | chr10 | + |
| Cdc40 | 547,10 | -0,0191 | 0,975445719 | chr10 | - |
| Cdk1 | 80,42 | 0,0074 | 0,992625586 | chr10 | - |
| Cdk17 | 330,00 | -0,1735 | 0,636260727 | chr10 | + |
| Cdk19 | 157,31 | -0,1639 | 0,683522146 | chr10 | + |
| Cdk2 | 205,57 | -0,0381 | 0,942982044 | chr10 | - |
| Cdk4 | 580,69 | 0,0072 | 0,988990696 | chr10 | + |
| Cenpw | 38,72 | -0,5006 | 0,41633338 | chr10 | - |
| Cep290 | 55,39 | 0,6019 | 0,19691213 | chr10 | + |
| Cep57l1 | 99,74 | 0,3932 | 0,35583041 | chr10 | - |
| Cep85l | 51,17 | 0,5581 | 0,273224953 | chr10 | - |
| Chchd10 | 2252,48 | -0,7468 | 0,093044072 | chr10 | + |
| Chst11 | 5,55 | 0,5559 | 0,411104454 | chr10 | + |
| Chst3 | 5,15 | 0,3179 | 0,676716141 | chr10 | - |
| Cirbp | 200,49 | -0,1027 | 0,851472884 | chr10 | + |
| Cisd1 | 2504,63 | -0,4317 | 0,233121751 | chr10 | - |
| Cited2 | 160,31 | -0,1070 | 0,826227466 | chr10 | + |
| Ckap4 | 34,15 | 0,1549 | 0,817465502 | chr10 | - |
| Cnksr3 | 70,61 | 0,2142 | 0,660583784 | chr10 | - |
| Cnn2 | 86,33 | -0,0217 | 0,976991666 | chr10 | + |
| Cnot2 | 619,58 | 0,0629 | 0,865984359 | chr10 | - |
| Cnpy2 | 2128,99 | -0,1975 | 0,517431746 | chr10 | + |
| Col13a1 | 54,23 | 0,1026 | 0,909142576 | chr10 | - |
| Col18a1 | 6579,02 | -0,1409 | 0,795796985 | chr10 | - |
| Col6a1 | 117,34 | 0,0291 | 0,965770119 | chr10 | - |
| Col6a2 | 47,63 | -0,1464 | 0,834002746 | chr10 | - |
| Cpsf6 | 530,77 | -0,1545 | 0,741076165 | chr10 | - |
| Cradd | 281,17 | 0,5079 | 0,103928027 | chr10 | - |
| Creb3l3 | 5016,65 | -0,1526 | 0,76563861 | chr10 | - |
| Cry1 | 120,87 | -0,4946 | 0,268487435 | chr10 | - |
| Cs | 1498,06 | -0,2102 | 0,47056536 | chr10 | + |
| Csnk1g2 | 512,70 | -0,4024 | 0,453845946 | chr10 | + |
| Cstb | 777,54 | -0,0660 | 0,901994263 | chr10 | + |
| Ctdsp2 | 788,49 | -0,1627 | 0,633994747 | chr10 | + |
| Ctgf | 131,06 | -0,1540 | 0,811137679 | chr10 | + |
| D10Jhu81e | 1538,57 | -0,2541 | 0,526350762 | chr10 | - |
| D10Wsu102e | 235,86 | 0,0706 | 0,884858966 | chr10 | + |
| D830005E20Rik | 2,66 | 0,3016 | 0,665028267 | chr10 | - |
| Dapk3 | 173,31 | -0,4649 | 0,212915792 | chr10 | + |
| Dazap1 | 305,71 | -0,3145 | 0,582441358 | chr10 | + |
| Dcbld1 | 50,99 | -0,0511 | 0,940390384 | chr10 | + |
| Dctn2 | 1335,00 | -0,4084 | 0,149221118 | chr10 | + |
| Ddit3 | 156,05 | -0,2587 | 0,653280861 | chr10 | + |
| Ddit4 | 160,50 | -0,3371 | 0,661627141 | chr10 | - |
| Ddo | 443,63 | 0,2759 | 0,487920673 | chr10 | + |
| Ddt | 6553,26 | -0,0660 | 0,903337956 | chr10 | - |
| Ddx21 | 541,83 | -0,2253 | 0,51294059 | chr10 | - |

| | | | | | |
|---|---|---|---|---|---|
| Ddx50 | 1114,20 | -0,2846 | 0,395294986 | chr10 | - |
| Derl3 | 177,66 | -0,0899 | 0,91260409 | chr10 | + |
| Dgka | 148,12 | -0,0718 | 0,90065927 | chr10 | - |
| Dip2a | 44,37 | -0,1862 | 0,754524942 | chr10 | - |
| Dna2 | 129,22 | -0,0114 | 0,98507181 | chr10 | + |
| Dnajb12 | 286,43 | 0,1638 | 0,760045267 | chr10 | + |
| Dnajc12 | 349,67 | -0,2125 | 0,70612403 | chr10 | + |
| Dnajc14 | 621,89 | 0,2822 | 0,393993287 | chr10 | + |
| Dohh | 273,40 | -0,3876 | 0,3899381 | chr10 | + |
| Cbarp | 10,98 | -0,1088 | 0,907233333 | chr10 | - |
| Dot1l | 89,00 | -0,3125 | 0,630199445 | chr10 | + |
| Dram1 | 17,10 | 0,1612 | 0,841053685 | chr10 | - |
| Dse | 199,78 | 0,3760 | 0,277831408 | chr10 | - |
| Dtx3 | 58,11 | 0,4549 | 0,364372969 | chr10 | - |
| Dusp6 | 425,03 | -0,4867 | 0,29259208 | chr10 | + |
| Dyrk2 | 91,22 | -0,4204 | 0,332014851 | chr10 | - |
| E030030I06Rik | 30,71 | -0,7444 | 0,196045303 | chr10 | - |
| E130307A14Rik | 5,50 | -0,0058 | 0,995735598 | chr10 | - |
| E130317F20Rik | 7,34 | -0,1443 | 0,873391707 | chr10 | - |
| E2f7 | 3,08 | -0,1192 | 0,893538095 | chr10 | + |
| Echdc1 | 2034,70 | 0,3694 | 0,321487973 | chr10 | + |
| Eea1 | 503,14 | -0,2609 | 0,432040063 | chr10 | + |
| Eef2 | 13501,69 | -0,4598 | 0,220525617 | chr10 | + |
| Egr2 | 4,51 | 0,4585 | 0,517431746 | chr10 | + |
| Eif4ebp2 | 387,15 | 0,2043 | 0,649875139 | chr10 | - |
| Elk3 | 183,60 | 0,1059 | 0,815188933 | chr10 | - |
| Enpp1 | 352,49 | 0,1194 | 0,795796985 | chr10 | - |
| Enpp3 | 829,72 | 0,1394 | 0,692167613 | chr10 | - |
| Epb41l2 | 221,89 | 0,4486 | 0,141463153 | chr10 | + |
| Epm2a | 9,92 | -0,0652 | 0,948314012 | chr10 | + |
| Erbb3 | 680,39 | 0,6180 | 0,134272248 | chr10 | - |
| Esyt1 | 142,15 | -0,1765 | 0,74829088 | chr10 | - |
| Fabp7 | 147,53 | -0,2101 | 0,777753633 | chr10 | + |
| Fam13c | 4,41 | 0,0857 | 0,92861245 | chr10 | + |
| Fam207a | 337,04 | 0,0886 | 0,856476342 | chr10 | - |
| Fam229b | 15,96 | 0,8401 | 0,117246861 | chr10 | - |
| Fam26e | 10,82 | 0,5398 | 0,404820274 | chr10 | - |
| Fam26f | 27,00 | -0,0722 | 0,925042168 | chr10 | - |
| Fbxo30 | 342,15 | -0,0900 | 0,881086817 | chr10 | + |
| Fbxo5 | 16,40 | -0,1693 | 0,830330662 | chr10 | - |
| Fbxo7 | 303,40 | -0,0683 | 0,860890261 | chr10 | + |
| Fgd6 | 577,77 | -0,1819 | 0,599811615 | chr10 | + |
| Fhl4 | 2,37 | 0,3800 | NA | chr10 | - |
| Fig4 | 216,22 | 0,0921 | 0,86019237 | chr10 | - |
| Foxo3 | 109,60 | 0,2324 | 0,615847246 | chr10 | - |
| Frk | 119,33 | -0,0562 | 0,932525817 | chr10 | + |
| Frs2 | 200,65 | 0,2609 | 0,491855001 | chr10 | - |
| Fstl3 | 3,11 | 0,2372 | 0,764664236 | chr10 | + |
| Ftcd | 6216,02 | -0,3236 | 0,306882295 | chr10 | + |
| Fyn | 119,41 | 0,2324 | 0,55880551 | chr10 | + |

| | | | | | |
|---|---|---|---|---|---|
| Fzr1 | 97,87 | -0,0911 | 0,868039727 | chr10 | - |
| Mfsd4b3 | 227,15 | -0,1475 | 0,759265283 | chr10 | - |
| Gadd45b | 23,45 | -0,1884 | 0,825518458 | chr10 | + |
| Galnt4 | 234,76 | 0,0955 | 0,839302072 | chr10 | + |
| Gamt | 1499,79 | 0,1087 | 0,740596839 | chr10 | - |
| Gas2l3 | 43,52 | -0,0664 | 0,932012942 | chr10 | - |
| Gcc2 | 485,04 | 0,4296 | 0,290238703 | chr10 | + |
| Ggt5 | 39,87 | -0,0910 | 0,900342259 | chr10 | + |
| Ginm1 | 885,29 | -0,3520 | 0,267303042 | chr10 | - |
| Gja1 | 141,12 | 0,4721 | 0,208540118 | chr10 | + |
| Gje1 | 2,11 | 0,0301 | NA | chr10 | - |
| Glipr1 | 33,81 | 0,3923 | 0,484137651 | chr10 | - |
| Gls2 | 4869,63 | -0,0676 | 0,887518458 | chr10 | + |
| Gm10941 | 2,22 | -0,0361 | NA | chr10 | - |
| Gm15663 | 6,65 | 0,2491 | 0,761512492 | chr10 | + |
| Gm16998 | 2,13 | 0,0596 | NA | chr10 | + |
| Gm20300 | 499,67 | -0,0335 | 0,943108135 | chr10 | - |
| Gm20597 | 2,54 | -0,2209 | NA | chr10 | - |
| Gm4925 | 4,84 | 0,4106 | 0,574181008 | chr10 | - |
| Gm5177 | 2,59 | -0,0523 | 0,955066061 | chr10 | + |
| Gm5420 | 4,32 | 0,9959 | 0,068187551 | chr10 | + |
| Rpl32l | 6,88 | -0,2833 | 0,720000255 | chr10 | + |
| Gm8580 | 19,00 | -0,5892 | 0,323355495 | chr10 | - |
| Gm8709 | 14,66 | 0,9484 | 0,075466772 | chr10 | - |
| Gm9079 | 3,86 | 0,2034 | 0,811137679 | chr10 | - |
| Gna11 | 575,54 | -0,2784 | 0,442032524 | chr10 | - |
| Gna15 | 6,75 | 0,1165 | 0,90065927 | chr10 | - |
| Gng7 | 6,95 | -0,3057 | 0,693803702 | chr10 | - |
| Gnptab | 145,96 | 0,2825 | 0,485968266 | chr10 | + |
| Gns | 1812,29 | -0,3310 | 0,220525617 | chr10 | + |
| Gopc | 453,24 | 0,2385 | 0,543927518 | chr10 | - |
| Lilr4b | 20,23 | 0,3080 | 0,655774674 | chr10 | + |
| Adgrg6 | 23,04 | 0,0618 | 0,943108135 | chr10 | - |
| Gpr182 | 328,55 | 0,1054 | 0,865199339 | chr10 | - |
| Gpx4 | 4635,80 | -0,5460 | 0,169210008 | chr10 | + |
| Gstt1 | 7488,04 | 0,4621 | 0,05818386 | chr10 | - |
| Gstt2 | 1571,74 | 0,3035 | 0,490509101 | chr10 | - |
| Gstt3 | 483,99 | -0,2421 | 0,460761576 | chr10 | - |
| Gtf3c6 | 450,96 | -0,1450 | 0,658257248 | chr10 | - |
| Gucd1 | 2302,78 | -0,1065 | 0,83176435 | chr10 | - |
| H2afy2 | 10,57 | 0,2122 | 0,792197952 | chr10 | - |
| Hace1 | 60,76 | 0,2658 | 0,601591922 | chr10 | + |
| Hal | 4285,83 | 0,5014 | 0,172087384 | chr10 | + |
| Hbs1l | 1252,21 | -0,3571 | 0,121757173 | chr10 | + |
| Hcfc2 | 129,09 | 0,0271 | 0,968744215 | chr10 | + |
| Hcn2 | 4,03 | -0,2443 | 0,757395951 | chr10 | + |
| Hdac2 | 375,20 | -0,3486 | 0,319080738 | chr10 | + |
| Hddc2 | 32,93 | -0,5885 | 0,212149875 | chr10 | + |
| Heca | 434,26 | -0,2428 | 0,569876243 | chr10 | - |
| Helb | 137,45 | -0,3711 | 0,246966539 | chr10 | - |

| | | | | | |
|---|---|---|---|---|---|
| Herc4 | 961,50 | 0,0159 | 0,9774226 | chr10 | + |
| Hey2 | 2,65 | -0,3207 | 0,637721381 | chr10 | - |
| Hint3 | 555,11 | -0,3627 | 0,462995924 | chr10 | - |
| Hivep2 | 105,91 | 0,2020 | 0,663924988 | chr10 | + |
| Hk1 | 38,66 | 0,1709 | 0,775458257 | chr10 | - |
| Hmg20b | 131,98 | -0,7082 | 0,158793576 | chr10 | - |
| Hmga2 | 3,67 | 0,3664 | 0,617869186 | chr10 | - |
| Arhgap45 | 95,70 | -0,2828 | 0,646342923 | chr10 | + |
| Hnrnph3 | 266,33 | -0,3108 | 0,417918742 | chr10 | - |
| Hsd17b6 | 4894,51 | -1,0341 | 0,067696785 | chr10 | - |
| Hsf2 | 603,99 | -0,2312 | 0,629873977 | chr10 | + |
| Hsp90b1 | 26838,91 | 0,2119 | 0,641854125 | chr10 | - |
| Icosl | 15,11 | 0,7907 | 0,155946567 | chr10 | + |
| Ifngr1 | 417,60 | 0,0352 | 0,943346405 | chr10 | + |
| Igf1 | 19281,29 | 0,0708 | 0,875717638 | chr10 | + |
| Ikbip | 64,90 | 0,0865 | 0,896614435 | chr10 | + |
| Ikzf4 | 4,17 | 0,8032 | 0,177916821 | chr10 | - |
| Ilvbl | 416,71 | -0,2562 | 0,537966644 | chr10 | + |
| Inhbc | 1033,53 | -0,6709 | 0,135447688 | chr10 | - |
| Ipmk | 1145,02 | -0,1022 | 0,793250646 | chr10 | + |
| Irak3 | 12,64 | -0,0717 | 0,941553175 | chr10 | - |
| Itga7 | 90,75 | 0,1608 | 0,81172751 | chr10 | + |
| Itgb2 | 116,91 | 0,3337 | 0,367689033 | chr10 | + |
| Iyd | 1996,23 | 0,3408 | 0,213281186 | chr10 | + |
| Izumo4 | 117,80 | 0,1592 | 0,773763312 | chr10 | + |
| Jmjd1c | 639,32 | 0,3275 | 0,260028246 | chr10 | + |
| Katna1 | 290,45 | -0,2946 | 0,370803111 | chr10 | + |
| Kif5a | 9,62 | -0,2498 | 0,751956685 | chr10 | - |
| Kitl | 67,77 | 0,1894 | 0,718689474 | chr10 | + |
| Krr1 | 459,38 | 0,1037 | 0,804839321 | chr10 | + |
| L3mbtl3 | 42,57 | 0,5660 | 0,311625736 | chr10 | - |
| Afg1l | 261,05 | -0,2314 | 0,488661764 | chr10 | - |
| Lama2 | 53,36 | 0,2850 | 0,637136777 | chr10 | - |
| Lama4 | 87,17 | -0,0527 | 0,93056206 | chr10 | + |
| Lats1 | 359,76 | 0,0389 | 0,938152195 | chr10 | + |
| Lemd3 | 149,58 | 0,1204 | 0,799384382 | chr10 | - |
| Lgr5 | 56,70 | -0,5072 | 0,373204176 | chr10 | - |
| Lilrb4a | 89,05 | 0,0955 | 0,87623846 | chr10 | + |
| Lims1 | 326,94 | 0,6013 | 0,078316133 | chr10 | + |
| Lin7a | 765,02 | -0,3314 | 0,528469822 | chr10 | + |
| Llph | 673,17 | -0,3524 | 0,319080738 | chr10 | + |
| Lmnb2 | 102,22 | -0,0456 | 0,947707681 | chr10 | - |
| Lrig3 | 95,62 | 0,1845 | 0,691018693 | chr10 | + |
| Lrp1 | 4001,17 | -0,1873 | 0,667325588 | chr10 | - |
| Lrrc20 | 65,06 | 0,1023 | 0,867671723 | chr10 | + |
| Lrrc3 | 1634,79 | 0,2224 | 0,613341495 | chr10 | - |
| Lrrc75b | 4,62 | -0,4530 | 0,52825088 | chr10 | - |
| Lsm7 | 152,96 | 0,0545 | 0,932528423 | chr10 | - |
| Lss | 452,55 | -0,0451 | 0,949680678 | chr10 | + |
| Lta4h | 529,17 | -0,1149 | 0,762940356 | chr10 | + |

| | | | | | |
|---|---|---|---|---|---|
| Ltv1 | 251,71 | -0,0588 | 0,911294941 | chr10 | - |
| Lyz1 | 322,32 | -0,0760 | 0,907865203 | chr10 | - |
| Lyz2 | 1784,28 | 0,1477 | 0,792197952 | chr10 | - |
| Man1a | 2339,80 | -0,0754 | 0,823828114 | chr10 | - |
| Map3k5 | 518,43 | -0,2359 | 0,61720787 | chr10 | + |
| Map7 | 293,90 | -0,0382 | 0,943108135 | chr10 | + |
| March9 | 4,52 | -0,6415 | 0,324324312 | chr10 | - |
| Marcks | 443,13 | 0,1422 | 0,731048653 | chr10 | - |
| Matk | 4,33 | -0,2730 | 0,732140841 | chr10 | + |
| Mbd3 | 422,11 | -0,7408 | 0,062031449 | chr10 | - |
| Mbd6 | 184,14 | -0,4832 | 0,389499877 | chr10 | - |
| Mcm3ap | 232,28 | -0,0341 | 0,945407007 | chr10 | + |
| Mcm9 | 28,05 | -0,0907 | 0,89879469 | chr10 | - |
| Mcu | 144,09 | -0,2172 | 0,600001528 | chr10 | - |
| Mdm1 | 12,95 | -0,0109 | 0,991035244 | chr10 | + |
| Mdm2 | 303,90 | -0,4494 | 0,081609555 | chr10 | - |
| Med16 | 113,47 | -0,1036 | 0,869442758 | chr10 | - |
| Med23 | 140,18 | 0,3033 | 0,393345568 | chr10 | + |
| Metap2 | 1520,75 | -0,1572 | 0,612061669 | chr10 | - |
| Mettl1 | 78,59 | 0,1759 | 0,772262933 | chr10 | + |
| Mettl25 | 43,79 | 0,0883 | 0,896013441 | chr10 | - |
| Mettl7b | 8021,82 | -0,4002 | 0,252815255 | chr10 | - |
| Mex3d | 18,82 | 0,1212 | 0,880810135 | chr10 | - |
| Mfsd12 | 24,54 | 0,8345 | 0,141463153 | chr10 | + |
| Mical1 | 20,64 | 0,3046 | 0,627875496 | chr10 | + |
| Micu1 | 396,23 | 0,3000 | 0,34997493 | chr10 | + |
| Midn | 319,79 | -0,0972 | 0,85281531 | chr10 | + |
| Mier2 | 62,72 | -0,1499 | 0,823537093 | chr10 | - |
| Mif | 1218,04 | -0,4843 | 0,487542473 | chr10 | - |
| Misp | 5,55 | 0,4416 | 0,54003692 | chr10 | + |
| Mknk2 | 1976,38 | -0,1797 | 0,721557952 | chr10 | - |
| Mmp11 | 9,09 | 0,2303 | 0,773303025 | chr10 | - |
| Mmp19 | 1251,32 | 0,3243 | 0,44461139 | chr10 | + |
| Mob3a | 66,32 | 0,2487 | 0,599980293 | chr10 | - |
| Mon2 | 676,93 | -0,2291 | 0,423089018 | chr10 | - |
| Moxd1 | 4,75 | -0,1623 | 0,605538599 | chr10 | + |
| Mrpl42 | 1589,37 | -0,1153 | 0,743336656 | chr10 | - |
| Mrpl54 | 335,97 | -0,3320 | 0,60249871 | chr10 | - |
| Msl3l2 | 13,24 | 0,4723 | 0,479279449 | chr10 | + |
| Msrb3 | 270,34 | 0,2022 | 0,56720184 | chr10 | - |
| Mterf2 | 79,72 | 0,7402 | 0,175457889 | chr10 | - |
| Mtfr2 | 5,66 | -0,9409 | 0,110056031 | chr10 | + |
| Mthfd1l | 49,63 | 0,4876 | 0,381169644 | chr10 | + |
| Mtrf1l | 77,41 | -0,0436 | 0,941192433 | chr10 | - |
| Mum1 | 81,03 | 0,0318 | 0,962657075 | chr10 | + |
| Myct1 | 107,68 | 0,2139 | 0,652294111 | chr10 | + |
| Myl6 | 8197,87 | -0,0584 | 0,872521494 | chr10 | - |
| Myl6b | 7,42 | -0,4830 | 0,491855001 | chr10 | - |
| Nabp2 | 305,12 | -0,4730 | 0,228893066 | chr10 | - |
| Naca | 6805,85 | -0,4607 | 0,075945592 | chr10 | + |

| | | | | | |
|---|---|---|---|---|---|
| Nap1l1 | 1294,77 | -0,4297 | 0,118025078 | chr10 | + |
| Ncln | 388,26 | -0,4929 | 0,279505776 | chr10 | - |
| Ncoa7 | 38,49 | 0,1092 | 0,884297082 | chr10 | - |
| Ndufa12 | 1768,42 | -0,3340 | 0,449238443 | chr10 | + |
| Ndufs7 | 1032,44 | -0,3992 | 0,581725594 | chr10 | + |
| Nedd1 | 395,65 | 0,4307 | 0,268870835 | chr10 | - |
| Nfic | 588,22 | -0,2373 | 0,600688917 | chr10 | - |
| Nfyb | 557,10 | 0,0728 | 0,867698337 | chr10 | - |
| Nkain2 | 5,01 | 0,6663 | 0,300779985 | chr10 | - |
| Nodal | 6,97 | 0,1733 | 0,843329386 | chr10 | + |
| Nr1h4 | 2715,35 | -0,0551 | 0,91260409 | chr10 | - |
| Nrbf2 | 343,22 | -0,4124 | 0,160487212 | chr10 | - |
| Nt5dc1 | 312,65 | 0,0417 | 0,938514017 | chr10 | - |
| Ntn4 | 44,74 | 0,4222 | 0,396411176 | chr10 | + |
| Nuak1 | 37,01 | 0,2672 | 0,63960646 | chr10 | - |
| Nudt4 | 2982,74 | -0,3442 | 0,242907451 | chr10 | - |
| Nup107 | 73,55 | 0,3531 | 0,441570242 | chr10 | - |
| Nup37 | 117,23 | 0,4105 | 0,305522088 | chr10 | + |
| Nup43 | 94,78 | 0,0918 | 0,869481228 | chr10 | + |
| Nus1 | 1217,75 | -0,2039 | 0,573066248 | chr10 | + |
| Oaz1 | 83,50 | -0,6186 | 0,116688134 | chr10 | + |
| Oit3 | 563,66 | 0,1038 | 0,78044956 | chr10 | - |
| Ormdl2 | 1473,54 | 0,0085 | 0,986661276 | chr10 | - |
| Os9 | 1552,35 | -0,2488 | 0,514431803 | chr10 | - |
| Osbpl8 | 632,40 | -0,0739 | 0,88569455 | chr10 | + |
| Ostm1 | 389,70 | -0,0497 | 0,912811384 | chr10 | + |
| P4ha1 | 178,70 | -0,2877 | 0,518493935 | chr10 | + |
| Pa2g4 | 1597,10 | -0,4078 | 0,183363484 | chr10 | - |
| Pald1 | 158,65 | -0,1922 | 0,60828654 | chr10 | - |
| Palm | 15,99 | 0,1436 | 0,8672888 | chr10 | + |
| Pan2 | 331,51 | 0,1619 | 0,715448456 | chr10 | + |
| Parpbp | 6,33 | -0,0227 | 0,98429278 | chr10 | - |
| Pawr | 122,46 | 0,1824 | 0,712389086 | chr10 | + |
| Pbld1 | 2609,19 | -0,1729 | 0,62208121 | chr10 | + |
| Pcbd1 | 3999,11 | -0,1533 | 0,645162115 | chr10 | + |
| Pcbp3 | 10,45 | -0,2611 | 0,736071538 | chr10 | - |
| Pcmt1 | 575,11 | 0,0606 | 0,889053514 | chr10 | - |
| Pcnt | 79,37 | -0,0405 | 0,953751362 | chr10 | - |
| Pde7b | 48,78 | -0,1776 | 0,801768605 | chr10 | - |
| Pdss2 | 124,62 | 0,0563 | 0,908913202 | chr10 | + |
| Pdxk | 1186,97 | -0,4202 | 0,248982001 | chr10 | - |
| Perp | 1661,31 | -0,4666 | 0,058863433 | chr10 | + |
| Pex3 | 625,68 | -0,0174 | 0,971747809 | chr10 | - |
| Pex7 | 1256,64 | -0,0815 | 0,862616891 | chr10 | - |
| Pfkl | 419,78 | -0,5697 | 0,119628533 | chr10 | - |
| Phactr2 | 89,73 | -0,0283 | 0,965007096 | chr10 | - |
| Phlda1 | 334,60 | -0,1470 | 0,871232434 | chr10 | + |
| Pias4 | 84,00 | -0,2643 | 0,627875496 | chr10 | - |
| Pip4k2c | 396,28 | 0,1337 | 0,722057512 | chr10 | - |
| Pip5k1c | 107,40 | 0,1553 | 0,743124066 | chr10 | + |

| | | | | | |
|---|---|---|---|---|---|
| Pkib | 24,38 | 0,2480 | 0,712581071 | chr10 | + |
| Pla2g12b | 1277,23 | 0,0826 | 0,821346657 | chr10 | + |
| Plekhg1 | 68,05 | 0,0935 | 0,879380574 | chr10 | + |
| Plekhj1 | 186,11 | -0,1942 | 0,722152554 | chr10 | - |
| Plk5 | 168,18 | -0,0100 | 0,990189789 | chr10 | + |
| Plxnc1 | 459,97 | 0,3543 | 0,230037893 | chr10 | - |
| Pmel | 5,12 | -0,0836 | 0,93189316 | chr10 | + |
| Poc1b | 67,92 | -0,1038 | 0,85586037 | chr10 | + |
| Pofut2 | 147,18 | -0,2262 | 0,634237396 | chr10 | + |
| Polr2e | 346,43 | -0,3406 | 0,408981984 | chr10 | - |
| Polrmt | 139,88 | -0,5264 | 0,331535469 | chr10 | - |
| Ppa1 | 2813,82 | 0,0218 | 0,969414042 | chr10 | + |
| Plpp2 | 388,27 | 0,0266 | 0,968669888 | chr10 | - |
| Ppfia2 | 10,57 | 0,1387 | 0,87652112 | chr10 | + |
| Ppil4 | 305,63 | -0,0421 | 0,942935841 | chr10 | + |
| Ppil6 | 8,93 | 0,0919 | 0,923778914 | chr10 | + |
| Ppm1h | 55,66 | -0,0854 | 0,893437979 | chr10 | + |
| Ppp1r12a | 148,93 | 0,4760 | 0,166380044 | chr10 | + |
| Prdm1 | 9,94 | 0,1264 | 0,888648505 | chr10 | - |
| Prdm4 | 129,31 | -0,4009 | 0,283332998 | chr10 | - |
| Prep | 435,40 | -0,4560 | 0,129937883 | chr10 | + |
| Prim1 | 109,03 | 0,2636 | 0,62632688 | chr10 | + |
| Prmt2 | 11,84 | 0,1964 | 0,806159394 | chr10 | - |
| Prtn3 | 46,65 | 0,3117 | 0,665829499 | chr10 | + |
| Psap | 7077,36 | -0,0999 | 0,804244364 | chr10 | + |
| Ptbp1 | 870,18 | -0,1102 | 0,807413895 | chr10 | + |
| Ptges3 | 2612,00 | -0,3075 | 0,354338694 | chr10 | + |
| Ptprb | 1203,83 | 0,1106 | 0,741411104 | chr10 | + |
| Ptprk | 882,70 | 0,1065 | 0,746941515 | chr10 | + |
| Pttg1ip | 3785,53 | 0,4032 | 0,073076939 | chr10 | + |
| Pwp1 | 339,08 | 0,2362 | 0,512623736 | chr10 | + |
| Pwp2 | 158,35 | 0,0189 | 0,973455888 | chr10 | - |
| Qrsl1 | 73,05 | 0,0583 | 0,935309554 | chr10 | - |
| R3hdm2 | 548,91 | -0,0508 | 0,909774279 | chr10 | + |
| R3hdm4 | 439,84 | -0,3903 | 0,273864382 | chr10 | - |
| Rab21 | 283,95 | -0,0857 | 0,859286024 | chr10 | - |
| Rab32 | 486,64 | -0,4538 | 0,106945732 | chr10 | - |
| Rab36 | 4,54 | 0,5584 | 0,398141536 | chr10 | + |
| Rab3ip | 327,48 | -0,3718 | 0,162719431 | chr10 | - |
| Rab5b | 1639,36 | 0,2095 | 0,465196538 | chr10 | - |
| Raet1d | 29,46 | 0,4494 | 0,505514525 | chr10 | + |
| Raet1e | 5,03 | 0,4186 | 0,567249933 | chr10 | + |
| Ranbp2 | 1617,47 | 0,0986 | 0,846704266 | chr10 | + |
| Rap1b | 965,83 | -0,1043 | 0,78476189 | chr10 | - |
| Rassf3 | 1793,42 | 0,0790 | 0,835615523 | chr10 | - |
| Rassf9 | 3,30 | 0,4524 | 0,484137651 | chr10 | + |
| Rbms2 | 504,39 | 0,3946 | 0,130797733 | chr10 | - |
| Rdh1 | 21,79 | -0,6126 | 0,352573889 | chr10 | + |
| Rdh16 | 2307,60 | 0,0506 | 0,935309554 | chr10 | + |
| Rdh18-ps | 71,86 | -0,1687 | 0,726240854 | chr10 | + |

| | | | | | |
|---|---|---|---|---|---|
| Rdh19 | 16,00 | 0,1918 | 0,783264136 | chr10 | + |
| Rdh7 | 21244,96 | -0,3268 | 0,223523653 | chr10 | - |
| Reep3 | 1533,41 | -0,1033 | 0,760699597 | chr10 | - |
| Reps1 | 202,59 | -0,3086 | 0,324418269 | chr10 | + |
| Rev3l | 242,77 | -0,0222 | 0,963557252 | chr10 | + |
| Rexo1 | 130,12 | -0,1562 | 0,824129134 | chr10 | - |
| Rfx4 | 19,02 | -0,3766 | 0,547964222 | chr10 | + |
| Rgs17 | 3,93 | 0,4225 | 0,557112397 | chr10 | - |
| Rhobtb1 | 976,05 | -0,2551 | 0,537545925 | chr10 | + |
| Ric8b | 167,01 | -0,3247 | 0,340674125 | chr10 | + |
| Rnf126 | 100,71 | -0,6838 | 0,239683068 | chr10 | - |
| Rnf146 | 574,86 | 0,0622 | 0,8859355 | chr10 | - |
| Rnf217 | 217,03 | -0,4034 | 0,276877002 | chr10 | - |
| Rnf41 | 104,08 | -0,0952 | 0,893437979 | chr10 | + |
| Rpf2 | 280,45 | 0,1951 | 0,64281347 | chr10 | - |
| Rps12 | 4966,29 | -0,3994 | 0,28665528 | chr10 | - |
| Rps26 | 2275,37 | -0,6482 | 0,191230937 | chr10 | - |
| Rrp1 | 819,58 | -0,1484 | 0,653280861 | chr10 | - |
| Rspo3 | 80,62 | -0,0984 | 0,863490375 | chr10 | - |
| Rtcb | 1466,68 | 0,0987 | 0,812490734 | chr10 | - |
| Rtn4ip1 | 414,24 | 0,0461 | 0,93538092 | chr10 | + |
| Rufy2 | 78,85 | 0,1753 | 0,760825768 | chr10 | + |
| Rwdd1 | 569,56 | -0,4318 | 0,070720663 | chr10 | - |
| S1pr4 | 4,34 | 0,1095 | 0,906773634 | chr10 | - |
| Samd5 | 2,81 | 0,2262 | 0,767446843 | chr10 | - |
| Sar1a | 1901,16 | 0,0056 | 0,990189789 | chr10 | + |
| Sarnp | 956,47 | 0,0417 | 0,938786754 | chr10 | + |
| Sash1 | 490,53 | 0,1835 | 0,649305665 | chr10 | - |
| Sbno2 | 161,91 | -0,1612 | 0,8044742 | chr10 | - |
| Scamp4 | 313,78 | -0,2449 | 0,564495903 | chr10 | + |
| Scml4 | 3,30 | 0,1723 | 0,841150543 | chr10 | + |
| Scyl2 | 383,48 | 0,4899 | 0,130714829 | chr10 | - |
| Sdr9c7 | 906,63 | -0,0200 | 0,973516404 | chr10 | + |
| Sec63 | 3338,63 | -0,3275 | 0,223776633 | chr10 | + |
| Serinc1 | 4923,71 | 0,2206 | 0,577838565 | chr10 | - |
| Sesn1 | 275,43 | 0,4025 | 0,253026101 | chr10 | + |
| Sf3a2 | 77,99 | -0,4115 | 0,434423892 | chr10 | + |
| Sf3b5 | 268,63 | -0,2322 | 0,781284538 | chr10 | + |
| Sgk1 | 357,67 | -0,1738 | 0,789064438 | chr10 | + |
| Sgpl1 | 1020,06 | -0,0414 | 0,907233333 | chr10 | - |
| Sgta | 540,90 | -0,4542 | 0,183436041 | chr10 | - |
| Shc2 | 7,74 | 0,6109 | 0,351830969 | chr10 | - |
| Shmt2 | 1729,64 | -0,3554 | 0,323542972 | chr10 | - |
| Shprh | 222,13 | 0,2721 | 0,589817385 | chr10 | + |
| Sirt1 | 226,20 | 0,0511 | 0,913936879 | chr10 | - |
| Sirt6 | 30,79 | -0,4580 | 0,462039583 | chr10 | - |
| Slc16a10 | 1610,49 | -0,1493 | 0,629075916 | chr10 | - |
| Slc16a7 | 1071,53 | 0,2036 | 0,690021525 | chr10 | - |
| Slc16a9 | 60,05 | -0,0024 | 0,997302965 | chr10 | + |
| Slc18b1 | 39,95 | 0,1048 | 0,879201267 | chr10 | + |

| | | | | | |
|---|---|---|---|---|---|
| Slc25a16 | 962,63 | -0,2205 | 0,536603375 | chr10 | + |
| Slc26a10 | 24,56 | -0,0144 | 0,988512389 | chr10 | - |
| Slc29a3 | 74,83 | 0,5706 | 0,152264308 | chr10 | - |
| Slc39a3 | 102,81 | 0,0088 | 0,990189789 | chr10 | - |
| Slc39a5 | 11,95 | 0,4183 | 0,569607313 | chr10 | - |
| Slc41a2 | 295,79 | -0,4905 | 0,47979937 | chr10 | - |
| Smarcb1 | 169,61 | -0,0519 | 0,925898178 | chr10 | - |
| Smarcc2 | 644,88 | -0,3773 | 0,208282312 | chr10 | + |
| Smlr1 | 942,44 | -0,4538 | 0,167807438 | chr10 | - |
| Smpd2 | 524,19 | -0,4294 | 0,110973039 | chr10 | - |
| Smpdl3a | 1362,87 | 0,2897 | 0,388773392 | chr10 | + |
| Snora33 | 4,23 | 0,0659 | 0,945274006 | chr10 | - |
| Snord100 | 3,66 | 0,3880 | 0,592873282 | chr10 | - |
| Snord37 | 2,30 | 0,2302 | NA | chr10 | + |
| Snrpd3 | 1678,37 | -0,0207 | 0,969130697 | chr10 | + |
| Snrpf | 183,69 | -0,1893 | 0,804418611 | chr10 | - |
| Snx3 | 845,99 | -0,4336 | 0,165257806 | chr10 | + |
| Socs2 | 250,08 | -0,4238 | NA | chr10 | - |
| Sowahc | 134,75 | 0,2357 | 0,581990132 | chr10 | + |
| Specc1l | 269,49 | 0,0506 | 0,906652509 | chr10 | + |
| Spic | 62,69 | 0,4279 | 0,375672258 | chr10 | - |
| Spock2 | 4,59 | 0,4378 | 0,537078872 | chr10 | + |
| Sppl2b | 109,64 | -0,5719 | 0,264341661 | chr10 | + |
| Spryd4 | 612,30 | 0,1968 | 0,680997554 | chr10 | - |
| Srgap1 | 8,74 | 0,4478 | 0,526958509 | chr10 | - |
| Srgn | 327,21 | 0,5297 | 0,106121919 | chr10 | - |
| Stab2 | 530,92 | -0,2020 | 0,563218321 | chr10 | - |
| Stac3 | 9,37 | -0,3881 | 0,586349271 | chr10 | + |
| Stat2 | 843,85 | -0,0901 | 0,871232434 | chr10 | + |
| Stat6 | 784,18 | 0,0064 | 0,990552671 | chr10 | + |
| Stk11 | 733,42 | -0,2385 | 0,604679897 | chr10 | + |
| Stx11 | 10,33 | 0,3957 | 0,577347444 | chr10 | - |
| Stx7 | 362,04 | 0,3643 | 0,193343482 | chr10 | + |
| Stxbp5 | 187,41 | -0,4291 | 0,195952045 | chr10 | - |
| Sult3a1 | 8,17 | 0,1818 | NA | chr10 | + |
| Sumo3 | 1664,95 | 0,4099 | 0,068097992 | chr10 | + |
| Suox | 1997,25 | -0,0077 | 0,988512389 | chr10 | - |
| Supv3l1 | 239,21 | -0,4940 | 0,136585635 | chr10 | - |
| Susd2 | 29,67 | 0,1713 | 0,803690485 | chr10 | - |
| Sycp3 | 2,33 | 0,4081 | NA | chr10 | + |
| Syde1 | 35,26 | -0,3287 | 0,563374628 | chr10 | - |
| Syn3 | 14,88 | 0,1912 | 0,804418611 | chr10 | - |
| Syne1 | 333,87 | 0,3390 | 0,383467733 | chr10 | - |
| Tab2 | 661,83 | -0,3241 | 0,188129472 | chr10 | - |
| Tbc1d15 | 1252,17 | 0,1627 | 0,696532876 | chr10 | - |
| Tbc1d30 | 174,99 | 0,4081 | 0,266382194 | chr10 | - |
| Tbc1d32 | 200,69 | 0,6722 | 0,154188248 | chr10 | - |
| Tbk1 | 424,39 | 0,0650 | 0,887518458 | chr10 | - |
| Tbpl1 | 180,13 | -0,0943 | 0,830330662 | chr10 | - |
| Tbxa2r | 15,89 | -0,1564 | 0,851988359 | chr10 | + |

| | | | | | |
|---|---|---|---|---|---|
| Tcf21 | 83,00 | 0,4629 | 0,217147923 | chr10 | - |
| Tcf3 | 94,14 | -0,2700 | 0,609664949 | chr10 | - |
| Tcp11l2 | 575,22 | -0,2101 | 0,621474574 | chr10 | + |
| Tet1 | 3,14 | 0,1491 | 0,866208949 | chr10 | - |
| Tfam | 834,73 | -0,1296 | 0,727618101 | chr10 | - |
| Thap2 | 104,31 | 0,1814 | 0,682099278 | chr10 | - |
| Thop1 | 76,02 | -0,2423 | 0,661493297 | chr10 | + |
| Timeless | 34,17 | 0,4864 | 0,310109812 | chr10 | + |
| Timm13 | 965,68 | -0,4253 | 0,549838805 | chr10 | - |
| Timp3 | 648,22 | 0,0193 | 0,976286111 | chr10 | + |
| Tjp3 | 87,28 | 0,1032 | 0,896013441 | chr10 | - |
| Tle6 | 19,21 | -0,3948 | 0,53490582 | chr10 | - |
| Tmbim4 | 1265,31 | -0,4413 | 0,253908866 | chr10 | + |
| Tmcc3 | 210,81 | 0,1074 | 0,83512246 | chr10 | + |
| Tmem19 | 1945,52 | -0,5053 | 0,10663065 | chr10 | - |
| Nemp1 | 63,97 | 0,6397 | 0,160747995 | chr10 | + |
| Tmem198b | 50,91 | -0,0165 | 0,981991128 | chr10 | - |
| Tmem259 | 1682,63 | -0,0677 | 0,86698964 | chr10 | - |
| Tmem26 | 526,88 | -0,6419 | 0,125047679 | chr10 | + |
| Tmem263 | 177,96 | -0,1468 | 0,715185369 | chr10 | + |
| Tmpo | 591,85 | 0,4141 | 0,271016733 | chr10 | - |
| Tmtc2 | 163,59 | 0,2237 | 0,684393541 | chr10 | - |
| Tmtc3 | 160,94 | 0,5421 | 0,155520806 | chr10 | - |
| Tnfaip3 | 34,61 | 0,4765 | 0,334804026 | chr10 | - |
| Tpd52l1 | 141,01 | -0,5303 | 0,188855714 | chr10 | - |
| Tpgs1 | 109,12 | -0,3516 | 0,633049694 | chr10 | + |
| Traf3ip2 | 20,14 | 0,3020 | 0,635543228 | chr10 | + |
| Trappc10 | 244,58 | -0,3011 | 0,309392522 | chr10 | - |
| Trhde | 12,25 | -0,5336 | 0,416860553 | chr10 | - |
| Trmt11 | 146,97 | 0,0244 | 0,973485679 | chr10 | - |
| Trpm2 | 67,10 | 0,0995 | 0,882124635 | chr10 | - |
| Tsfm | 153,43 | -0,2675 | 0,587879802 | chr10 | - |
| Tspan15 | 83,33 | 0,4725 | 0,222670104 | chr10 | - |
| Tspan31 | 4721,57 | 0,0334 | 0,950819236 | chr10 | - |
| Tspan8 | 18,78 | -0,0843 | 0,923613432 | chr10 | + |
| Tspear | 4,40 | 0,0566 | 0,953805583 | chr10 | + |
| Tspyl1 | 321,62 | 0,3060 | 0,283172973 | chr10 | + |
| Tspyl4 | 25,50 | 0,3551 | 0,602610203 | chr10 | + |
| Tube1 | 12,64 | -0,0849 | 0,926972552 | chr10 | + |
| Txnrd1 | 1263,68 | -0,1212 | 0,776776996 | chr10 | + |
| Tysnd1 | 320,54 | -0,4226 | 0,299584947 | chr10 | + |
| Ube2d1 | 100,33 | -0,0727 | 0,88569455 | chr10 | - |
| Ube2g2 | 556,53 | 0,0303 | 0,953911274 | chr10 | + |
| Ube2n | 977,47 | 0,1028 | 0,804380135 | chr10 | + |
| Uhrf1bp1l | 573,56 | 0,0995 | 0,81907195 | chr10 | + |
| Ulbp1 | 2,20 | -0,0611 | NA | chr10 | - |
| Upb1 | 5955,97 | -0,1935 | 0,462967038 | chr10 | + |
| Uqcr11 | 4260,30 | -0,4295 | 0,430392697 | chr10 | - |
| Usp15 | 874,36 | -0,2944 | 0,364379698 | chr10 | - |
| Utp20 | 191,05 | 0,0519 | 0,930087232 | chr10 | - |

| | | | | | |
|---|---|---|---|---|---|
| Utrn | 361,85 | -0,0244 | 0,958859839 | chr10 | - |
| Vezt | 294,73 | -0,1665 | 0,631200781 | chr10 | - |
| Vnn1 | 1385,99 | 0,0041 | 0,995884982 | chr10 | + |
| Vnn3 | 1580,83 | 0,4619 | 0,096026608 | chr10 | + |
| Vpreb3 | 3,96 | -0,5747 | 0,378055718 | chr10 | + |
| Vps26a | 906,02 | 0,0418 | 0,932968636 | chr10 | - |
| Vta1 | 297,67 | -0,1798 | 0,596014309 | chr10 | - |
| Wasf1 | 21,22 | -0,8603 | 0,083831145 | chr10 | + |
| Wdr18 | 568,81 | -0,1635 | 0,715542157 | chr10 | + |
| Pym1 | 148,03 | -0,5132 | 0,227017349 | chr10 | + |
| Wif1 | 2,72 | -0,3525 | 0,602407167 | chr10 | + |
| Wisp3 | 3,18 | -0,1358 | 0,871232434 | chr10 | - |
| Xpot | 1113,81 | -0,1864 | 0,532105996 | chr10 | - |
| Atp23 | 48,26 | -0,5784 | 0,148762588 | chr10 | - |
| Ybey | 82,07 | 0,0055 | 0,993738083 | chr10 | - |
| Yeats4 | 214,70 | -0,0109 | 0,985628536 | chr10 | - |
| Zbtb2 | 87,40 | -0,5156 | 0,179535584 | chr10 | - |
| Zbtb24 | 149,86 | -0,3829 | 0,285255073 | chr10 | + |
| Zbtb39 | 77,57 | -0,0369 | 0,958903346 | chr10 | + |
| Zbtb7a | 237,67 | -0,5903 | 0,25333134 | chr10 | + |
| Zc3h10 | 88,82 | -0,5950 | 0,168950224 | chr10 | - |
| Zdhhc17 | 232,81 | 0,5148 | 0,06162809 | chr10 | - |
| Zfp280b | 111,62 | -0,3376 | 0,46279986 | chr10 | + |
| Zfp433 | 6,22 | 0,6259 | 0,340674125 | chr10 | + |
| Zufsp | 70,28 | 0,5141 | 0,269236255 | chr10 | - |
| Zwint | 948,52 | 0,0943 | 0,848381384 | chr10 | + |
| 0610009B22Rik | 678,23 | -0,1592 | 0,684077978 | chr11 | - |
| 0610010F05Rik | 334,14 | -0,0935 | 0,867809741 | chr11 | - |
| 0610010K14Rik | 181,33 | 0,3503 | 0,378627963 | chr11 | - |
| Wfdc21 | 14130,54 | -0,2531 | 0,45607208 | chr11 | + |
| 1110028F11Rik | 35,35 | -0,0610 | 0,937303754 | chr11 | + |
| Ncbp3 | 341,09 | 0,0808 | 0,847796138 | chr11 | + |
| 1700007J10Rik | 7,34 | -0,1407 | 0,876803328 | chr11 | - |
| 1700052K11Rik | 25,93 | -0,0635 | 0,938514017 | chr11 | - |
| 1810010H24Rik | 31,68 | -0,0324 | 0,97094427 | chr11 | + |
| 1810032O08Rik | 50,00 | -0,1465 | 0,827995984 | chr11 | + |
| Ndufaf8 | 172,66 | -0,1883 | 0,825889246 | chr11 | + |
| Ube2d-ps | 182,89 | -0,2467 | 0,569607313 | chr11 | + |
| Faap100 | 38,49 | 0,0707 | 0,93056206 | chr11 | - |
| 2310033P09Rik | 40,05 | 0,2799 | 0,603142371 | chr11 | + |
| Borcs6 | 78,41 | -0,6558 | 0,10210139 | chr11 | + |
| Tmem94 | 231,08 | 0,1436 | 0,776425522 | chr11 | + |
| 2410003L11Rik | 6,67 | -0,9490 | 0,108470987 | chr11 | + |
| 2410006H16Rik | 307,28 | -0,3255 | 0,676239605 | chr11 | + |
| 2610035D17Rik | 12,53 | 0,3306 | 0,639465027 | chr11 | - |
| 2610507B11Rik | 1980,08 | -0,0455 | 0,936483818 | chr11 | + |
| 2610507I01Rik | 101,78 | 0,1256 | NA | chr11 | - |
| 2810001G20Rik | 32,27 | -0,3102 | 0,592940467 | chr11 | + |
| Snhg20 | 62,85 | -1,0464 | 0,05129496 | chr11 | + |
| 2810021J22Rik | 53,51 | 0,4529 | 0,40679401 | chr11 | + |

| | | | | | |
|---|---|---|---|---|---|
| 2810410L24Rik | 27,19 | -0,6376 | 0,322847917 | chr11 | - |
| Eldr | 2,52 | 0,4516 | NA | chr11 | - |
| Mrnip | 113,97 | -0,6250 | 0,057877889 | chr11 | + |
| 4833417C18Rik | 10,18 | 0,0662 | 0,945274006 | chr11 | + |
| 4930563E22Rik | 4,49 | 0,0245 | 0,981991128 | chr11 | + |
| Trim80 | 8,96 | -0,0767 | 0,939190798 | chr11 | + |
| 4933427D14Rik | 38,08 | 0,0270 | 0,974621894 | chr11 | - |
| 4933439C10Rik | 39,97 | -0,7817 | 0,136970896 | chr11 | + |
| 5730455P16Rik | 285,78 | 0,2190 | 0,611914561 | chr11 | - |
| 8430429K09Rik | 14,26 | 0,2056 | 0,790000542 | chr11 | + |
| 9530068E07Rik | 1903,39 | -0,3592 | 0,240304485 | chr11 | + |
| AA465934 | 129,36 | 0,0299 | 0,978280656 | chr11 | + |
| Aarsd1 | 325,47 | -0,2523 | 0,481464342 | chr11 | - |
| Aatf | 302,23 | 0,1293 | 0,720926726 | chr11 | - |
| Abca5 | 97,68 | 0,3853 | 0,347104845 | chr11 | - |
| Abca6 | 3363,80 | 0,4179 | 0,220525617 | chr11 | - |
| Abca8a | 2353,83 | -0,3462 | 0,47339087 | chr11 | - |
| Abca8b | 820,24 | 0,2271 | 0,588219472 | chr11 | - |
| Abca9 | 76,86 | 0,6408 | 0,078316133 | chr11 | - |
| Abcc3 | 2737,53 | -0,1590 | 0,661748407 | chr11 | - |
| Abhd15 | 375,34 | 0,1992 | 0,588008527 | chr11 | + |
| Abi3 | 53,08 | -0,0644 | 0,931474598 | chr11 | - |
| Abr | 66,26 | 0,3720 | 0,407448757 | chr11 | - |
| Acadvl | 7982,62 | -0,3648 | 0,173113949 | chr11 | - |
| Acap1 | 28,64 | -0,2853 | 0,63960646 | chr11 | - |
| Acbd4 | 524,57 | -0,1466 | 0,715690739 | chr11 | + |
| Acox1 | 32613,05 | -0,4334 | 0,174173036 | chr11 | - |
| Acsf2 | 2362,87 | -0,1122 | 0,841579427 | chr11 | - |
| Actr2 | 2897,38 | 0,1523 | 0,683019231 | chr11 | - |
| Acyp2 | 62,49 | -0,6620 | 0,073430016 | chr11 | - |
| Adam11 | 62,57 | -0,0116 | 0,988766352 | chr11 | + |
| Adam19 | 19,59 | 0,3886 | 0,531668406 | chr11 | + |
| Adamts2 | 73,23 | 0,6996 | 0,080350582 | chr11 | + |
| Adap2 | 230,75 | 0,2719 | 0,570235502 | chr11 | + |
| Adprm | 383,49 | 0,0216 | 0,971756379 | chr11 | - |
| Adra1b | 502,62 | -0,1367 | 0,761732176 | chr11 | - |
| Aebp1 | 22,43 | 0,5545 | 0,296808586 | chr11 | + |
| Cd300c2 | 137,70 | 0,2052 | 0,636447281 | chr11 | - |
| Aff4 | 1169,39 | -0,2148 | 0,451241215 | chr11 | + |
| Afmid | 1695,80 | 0,0304 | 0,955834914 | chr11 | + |
| Aftph | 693,80 | -0,2067 | 0,512639686 | chr11 | - |
| Ahsa2 | 917,12 | -0,2995 | 0,379104437 | chr11 | - |
| AI450353 | 10,42 | -0,0944 | 0,92066291 | chr11 | + |
| AI662270 | 60,73 | 0,5041 | 0,277831408 | chr11 | + |
| Akap1 | 1172,77 | -0,0001 | 0,999705919 | chr11 | - |
| Akap10 | 116,19 | -0,2343 | 0,544069376 | chr11 | - |
| Aldh3a2 | 4027,47 | -0,5451 | 0,073412412 | chr11 | - |
| Aldoc | 245,80 | 0,0059 | 0,992874254 | chr11 | + |
| Alkbh5 | 1312,68 | -0,4056 | 0,108917495 | chr11 | + |
| Alox12 | 14,47 | 0,6284 | 0,295186082 | chr11 | - |

| | | | | | |
|---|---|---|---|---|---|
| Alox8 | 6,70 | -0,3927 | 0,599856218 | chr11 | - |
| Alyref | 141,40 | -0,0382 | 0,947294154 | chr11 | - |
| Amz2 | 359,59 | -0,0216 | 0,96651032 | chr11 | + |
| Anapc11 | 174,43 | -0,3595 | 0,522768066 | chr11 | + |
| Ankfy1 | 615,66 | 0,1650 | 0,635431815 | chr11 | + |
| Ankrd13b | 25,03 | -0,5357 | 0,413323375 | chr11 | - |
| Ankrd40 | 458,04 | -0,1525 | 0,63960646 | chr11 | + |
| Anxa6 | 1457,77 | 0,2369 | 0,402726862 | chr11 | - |
| Aoc2 | 10,98 | -0,4682 | 0,501000077 | chr11 | + |
| Aoc3 | 3,00 | -0,2723 | 0,720291746 | chr11 | + |
| Ap1b1 | 582,67 | -0,0687 | 0,884712754 | chr11 | + |
| Ap2b1 | 774,10 | -0,0113 | 0,984995389 | chr11 | + |
| Apoh | 59548,04 | 0,1104 | 0,749603956 | chr11 | + |
| Appbp2 | 204,27 | 0,5602 | 0,074290268 | chr11 | - |
| Arf1 | 4005,75 | 0,0531 | 0,888964395 | chr11 | - |
| Arf2 | 244,26 | -0,1627 | 0,699226411 | chr11 | + |
| Arhgap23 | 139,16 | 0,0031 | 0,99636982 | chr11 | + |
| Arhgap27 | 16,90 | 0,5208 | 0,396254007 | chr11 | - |
| Arhgap44 | 10,74 | -0,0281 | 0,979212524 | chr11 | - |
| Arhgdia | 1046,01 | -0,0488 | 0,939306183 | chr11 | - |
| Arhgef15 | 97,34 | -0,1898 | 0,674211854 | chr11 | - |
| Arl16 | 132,03 | -0,2110 | 0,614056134 | chr11 | - |
| Arl4d | 472,41 | -0,6605 | 0,157586062 | chr11 | + |
| Arl5c | 11,65 | 0,0281 | 0,978552759 | chr11 | - |
| Armc7 | 27,28 | 0,2585 | 0,672918705 | chr11 | + |
| Arrb2 | 189,00 | -0,1372 | 0,792197952 | chr11 | + |
| Asb3 | 161,26 | 0,0585 | 0,925898178 | chr11 | + |
| Ascc2 | 223,88 | -0,0663 | 0,906031026 | chr11 | + |
| Asgr1 | 11694,21 | -0,2479 | 0,35583041 | chr11 | + |
| Asgr2 | 3893,57 | 0,1277 | 0,666980604 | chr11 | + |
| Aspa | 429,73 | 0,0928 | 0,868187243 | chr11 | - |
| Aspscr1 | 726,74 | -0,3344 | 0,249470803 | chr11 | + |
| Atad5 | 23,00 | 0,2783 | 0,6711123 | chr11 | + |
| Atox1 | 1671,34 | -0,3820 | 0,340130203 | chr11 | - |
| Atp1b2 | 5,84 | 0,2525 | 0,759556393 | chr11 | - |
| Atp2a3 | 11,08 | 0,2678 | 0,73474321 | chr11 | + |
| Atp5g1 | 5165,94 | -0,3741 | 0,137166568 | chr11 | - |
| Atp5h | 6277,79 | -0,2034 | 0,419942821 | chr11 | - |
| Atp6v0a1 | 334,02 | -0,1428 | 0,757805775 | chr11 | + |
| Atpaf2 | 264,72 | -0,1938 | 0,633071535 | chr11 | - |
| Atxn7l3 | 246,87 | -0,1130 | 0,816025345 | chr11 | - |
| Aurkb | 8,81 | 0,1115 | 0,905147397 | chr11 | + |
| Axin2 | 36,19 | -0,7552 | 0,147012425 | chr11 | + |
| B230217C12Rik | 6,77 | 0,6813 | 0,288929615 | chr11 | + |
| B3gnt2 | 212,81 | -0,3609 | 0,273018284 | chr11 | - |
| B3gnt2 | 403,87 | 0,0685 | 0,877283616 | chr11 | - |
| B3gntl1 | 34,58 | 0,0345 | 0,965770119 | chr11 | - |
| B9d1 | 9,90 | 0,3440 | 0,646093772 | chr11 | + |
| Bahcc1 | 43,08 | -0,0249 | 0,97520712 | chr11 | + |
| Baiap2 | 244,29 | 0,1539 | 0,774841063 | chr11 | + |

| | | | | | |
|---|---|---|---|---|---|
| BC006965 | 2,79 | -0,1800 | 0,820837596 | chr11 | - |
| BC017643 | 208,46 | 0,0807 | 0,854357668 | chr11 | - |
| BC018473 | 3,01 | 0,0657 | 0,944269942 | chr11 | + |
| BC030499 | 3,04 | 0,0631 | 0,947819654 | chr11 | + |
| BC030867 | 3,37 | 0,1338 | 0,876803328 | chr11 | + |
| BC049762 | 5,19 | 0,2416 | 0,769677581 | chr11 | - |
| Bcas3 | 179,68 | -0,5230 | 0,147003641 | chr11 | + |
| Bcl6b | 38,96 | -0,0878 | 0,918946301 | chr11 | - |
| Becn1 | 1013,76 | -0,1795 | 0,52377705 | chr11 | - |
| Birc5 | 35,21 | 0,4086 | 0,458689654 | chr11 | + |
| Blmh | 873,79 | 0,1658 | 0,636447281 | chr11 | + |
| Bod1 | 369,69 | -0,4814 | 0,092498123 | chr11 | - |
| Bptf | 336,01 | -0,1371 | 0,736706859 | chr11 | - |
| Brca1 | 65,85 | 0,6498 | 0,166136649 | chr11 | - |
| Brip1 | 105,66 | 0,1246 | 0,830330662 | chr11 | - |
| Btnl9 | 15,29 | 0,8470 | 0,124946133 | chr11 | - |
| Tspoap1 | 2,78 | 0,0070 | 0,994334377 | chr11 | + |
| C130046K22Rik | 40,61 | -0,2558 | 0,651798423 | chr11 | + |
| C1d | 1827,72 | 0,1548 | 0,715542157 | chr11 | + |
| C1qbp | 912,13 | -0,4241 | 0,098620367 | chr11 | - |
| C1qtnf1 | 334,27 | 0,2755 | 0,383392162 | chr11 | + |
| C1qtnf2 | 18,56 | 0,2398 | 0,731047462 | chr11 | + |
| Cacnb1 | 4,72 | 0,3630 | 0,62811896 | chr11 | - |
| Camk2b | 59,54 | 0,1055 | 0,896013441 | chr11 | - |
| Camkk1 | 6,35 | 0,5584 | 0,408859672 | chr11 | + |
| Camta2 | 175,56 | -0,3581 | 0,436127277 | chr11 | - |
| Cant1 | 299,28 | -0,1187 | 0,813358947 | chr11 | - |
| Canx | 8872,38 | -0,0617 | 0,873391707 | chr11 | - |
| Card14 | 5,66 | 0,9950 | 0,084737957 | chr11 | + |
| Casc3 | 243,56 | -0,1758 | 0,610022568 | chr11 | + |
| Caskin2 | 80,64 | 0,3072 | 0,60487792 | chr11 | - |
| Cbr2 | 2,09 | 0,0508 | NA | chr11 | - |
| Cbx2 | 87,30 | 0,5402 | 0,212649167 | chr11 | + |
| Cbx4 | 147,49 | 0,2793 | 0,51447296 | chr11 | - |
| Cbx8 | 75,85 | 0,4755 | 0,319175548 | chr11 | - |
| Cfap36 | 297,33 | 0,2504 | 0,402148509 | chr11 | - |
| Ccdc117 | 504,71 | -0,1241 | 0,757888816 | chr11 | - |
| Ccdc137 | 134,83 | -0,2389 | 0,536536247 | chr11 | + |
| Ccdc43 | 221,64 | 0,2727 | 0,433006548 | chr11 | - |
| Ccdc47 | 2173,99 | 0,0702 | 0,877283616 | chr11 | - |
| Nsrp1 | 327,87 | -0,0619 | 0,893432234 | chr11 | - |
| Ccdc57 | 27,95 | 0,1951 | 0,783778153 | chr11 | - |
| Ccdc69 | 9,17 | 0,4827 | 0,489636271 | chr11 | - |
| Ccdc88a | 20,17 | 0,6739 | 0,19689523 | chr11 | + |
| Ccl12 | 2,10 | 0,2929 | NA | chr11 | + |
| Ccl3 | 3,06 | 0,2903 | 0,693097924 | chr11 | - |
| Ccl4 | 4,45 | 0,5348 | 0,421823262 | chr11 | + |
| Ccl5 | 19,41 | 0,5517 | 0,34827585 | chr11 | - |
| Ccl6 | 436,75 | 0,4182 | 0,444145513 | chr11 | - |
| Ccl7 | 7,72 | 0,6019 | 0,365320897 | chr11 | + |

| | | | | | |
|---|---|---|---|---|---|
| Ccl8 | 4,26 | -0,0465 | 0,963027434 | chr11 | + |
| Ccm2 | 289,66 | -0,3343 | 0,412631408 | chr11 | + |
| Ccng1 | 771,60 | 0,3244 | 0,320413698 | chr11 | - |
| Ccr7 | 4,55 | 0,2316 | 0,780303812 | chr11 | - |
| Cct4 | 2137,49 | -0,2684 | 0,390708488 | chr11 | + |
| Cct6b | 8,23 | -0,4457 | 0,534939703 | chr11 | - |
| Cd300a | 122,14 | 0,3724 | 0,278443756 | chr11 | + |
| Cd300c | 2,97 | -0,3050 | 0,677867285 | chr11 | - |
| Cd300e | 57,28 | 0,0575 | 0,947612608 | chr11 | - |
| Cd300ld | 191,93 | 0,0603 | 0,896013441 | chr11 | - |
| Cd300lf | 19,99 | 0,0507 | 0,952640742 | chr11 | - |
| Cd300lg | 303,40 | 0,0246 | 0,960164278 | chr11 | + |
| Cd68 | 194,48 | 0,1492 | 0,738680713 | chr11 | - |
| Cd7 | 9,23 | 0,4755 | 0,483930201 | chr11 | - |
| Cd79b | 24,73 | 0,2378 | 0,717851252 | chr11 | - |
| Cdc27 | 515,52 | 0,1087 | 0,814156035 | chr11 | - |
| Cdc42ep4 | 600,53 | -0,0252 | 0,959753478 | chr11 | - |
| Cdc42se2 | 213,13 | 0,0076 | 0,988766352 | chr11 | - |
| Cdc6 | 6,62 | 0,2234 | 0,791357694 | chr11 | + |
| Cdk12 | 181,48 | 0,2564 | 0,514431803 | chr11 | + |
| Cdk3-ps | 3,17 | 0,1478 | 0,863048009 | chr11 | + |
| Cdk5r1 | 4,29 | 0,1194 | 0,898098997 | chr11 | + |
| Cdk5rap3 | 1160,28 | -0,0928 | 0,803874753 | chr11 | - |
| Cdkl3 | 5,66 | -0,1075 | 0,909774279 | chr11 | + |
| Cdkn2aipnl | 226,25 | -0,1082 | 0,811093035 | chr11 | + |
| Cdr2l | 28,08 | 0,0134 | 0,987350308 | chr11 | + |
| Cenpv | 144,38 | -0,5922 | 0,074214969 | chr11 | - |
| Cep112 | 7,93 | 0,2530 | 0,754106401 | chr11 | + |
| Cep131 | 23,45 | -0,3085 | 0,642834828 | chr11 | - |
| Cep68 | 148,80 | -0,2615 | 0,491342637 | chr11 | - |
| Cep95 | 107,06 | -0,0243 | 0,96674257 | chr11 | + |
| Chac2 | 857,96 | 0,3137 | 0,385497664 | chr11 | - |
| Chd3 | 813,31 | -0,3630 | 0,395114656 | chr11 | - |
| Chd3os | 27,46 | -0,0098 | 0,990846949 | chr11 | + |
| Chmp6 | 196,52 | -0,0451 | 0,943291142 | chr11 | + |
| Chrnb1 | 9,05 | 0,3947 | 0,594896067 | chr11 | - |
| Chrne | 2,96 | 0,5513 | 0,382591532 | chr11 | - |
| Cisd3 | 897,79 | 0,3232 | 0,235271103 | chr11 | + |
| Clec10a | 10,90 | 0,5470 | 0,396254007 | chr11 | + |
| Clk4 | 801,01 | -0,1665 | 0,646093772 | chr11 | + |
| Cltc | 3359,76 | 0,0302 | 0,947294154 | chr11 | - |
| Cluh | 2135,70 | -0,3576 | 0,171869196 | chr11 | + |
| Cnot6 | 715,65 | 0,2683 | 0,437620234 | chr11 | - |
| Cnot8 | 1058,84 | -0,1692 | 0,771616285 | chr11 | + |
| Cnrip1 | 16,48 | -0,2103 | 0,77333268 | chr11 | + |
| Cntnap1 | 10,66 | -0,3131 | 0,684419282 | chr11 | + |
| Cntrob | 52,69 | 0,1428 | 0,799982428 | chr11 | - |
| Coa3 | 820,78 | -0,0816 | 0,859638359 | chr11 | - |
| Cobl | 273,20 | -0,7103 | 0,06498792 | chr11 | - |
| Cog1 | 140,87 | -0,0781 | 0,869825447 | chr11 | + |

| | | | | | |
|---|---|---|---|---|---|
| Coil | 59,10 | 0,3908 | 0,3845875 | chr11 | + |
| Col1a1 | 98,28 | 0,7532 | 0,150815507 | chr11 | + |
| Col23a1 | 3,06 | -0,2499 | 0,740663168 | chr11 | + |
| Commd1 | 935,41 | -0,1733 | 0,714195185 | chr11 | - |
| Cops3 | 1151,35 | -0,2139 | 0,556564028 | chr11 | - |
| Copz2 | 144,50 | -0,5788 | 0,11507047 | chr11 | + |
| Cox10 | 144,31 | -0,3802 | 0,317453963 | chr11 | - |
| Cox11 | 208,23 | -0,4319 | 0,211038545 | chr11 | + |
| Cpd | 611,94 | 0,3074 | 0,394599707 | chr11 | - |
| Cpeb4 | 575,02 | 0,0160 | 0,981931319 | chr11 | + |
| Cpsf4l | 79,80 | 0,1605 | 0,839302072 | chr11 | - |
| Crk | 1595,95 | -0,1726 | 0,621693318 | chr11 | + |
| Crlf3 | 128,11 | 0,0968 | 0,866934682 | chr11 | - |
| Csnk1d | 707,92 | -0,1339 | 0,740605363 | chr11 | - |
| Ctc1 | 69,43 | 0,2354 | 0,633486481 | chr11 | + |
| Ctdnep1 | 351,18 | -0,3867 | 0,268690208 | chr11 | + |
| Ctns | 265,52 | -0,0598 | 0,890881677 | chr11 | - |
| Cuedc1 | 17,49 | 0,1842 | 0,82773591 | chr11 | + |
| Cwc25 | 142,27 | 0,0705 | 0,875636241 | chr11 | - |
| Cxcl16 | 224,13 | 0,1308 | 0,769677581 | chr11 | - |
| Cyb561 | 141,26 | 0,4670 | 0,413323375 | chr11 | - |
| Cyb5d1 | 120,07 | 0,5622 | 0,181882538 | chr11 | - |
| Cyb5d2 | 226,92 | -0,2291 | 0,511056976 | chr11 | - |
| Cyfip2 | 27,88 | 0,7653 | 0,118025078 | chr11 | - |
| Cygb | 75,79 | 0,7160 | 0,105993721 | chr11 | - |
| Cyth1 | 679,39 | 0,2252 | 0,651354867 | chr11 | - |
| D030028A08Rik | 7,22 | 0,3380 | 0,661519766 | chr11 | + |
| D11Wsu47e | 71,23 | 0,4351 | 0,295152651 | chr11 | + |
| Appbp2os | 11,46 | -0,0245 | 0,981051811 | chr11 | + |
| D930048N14Rik | 42,29 | -0,2920 | 0,673076056 | chr11 | + |
| Dbil5 | 4,42 | 0,1372 | 0,880478973 | chr11 | + |
| Dbnl | 330,76 | 0,0606 | 0,90199272 | chr11 | + |
| Dcaf7 | 608,90 | 0,3258 | 0,210808905 | chr11 | + |
| Dcakd | 226,13 | -0,1805 | 0,768481763 | chr11 | - |
| Dcxr | 1038,74 | -0,0964 | 0,872326129 | chr11 | - |
| Ddc | 996,01 | -0,4009 | 0,500561834 | chr11 | - |
| Ddx42 | 510,42 | 0,0432 | 0,917575068 | chr11 | + |
| Ddx5 | 6269,05 | -0,1723 | 0,617219405 | chr11 | - |
| Ddx52 | 470,09 | 0,1276 | 0,806212175 | chr11 | + |
| Ddx56 | 364,41 | -0,0896 | 0,852975664 | chr11 | - |
| Derl2 | 2084,21 | -0,0869 | 0,879356682 | chr11 | - |
| Dgke | 19,46 | 0,3057 | 0,639465027 | chr11 | - |
| Dhrs11 | 278,99 | -0,3129 | 0,435611308 | chr11 | - |
| Dhrs7b | 646,89 | 0,1387 | 0,662842062 | chr11 | + |
| Dhx33 | 289,55 | -0,0714 | 0,869067375 | chr11 | - |
| Dhx40 | 480,69 | 0,0329 | 0,956122897 | chr11 | - |
| Dhx58 | 197,55 | 0,4874 | 0,287543614 | chr11 | - |
| Dhx8 | 207,93 | -0,0845 | 0,846704266 | chr11 | + |
| Dlg4 | 6,41 | 0,2884 | 0,717851252 | chr11 | + |
| Dnajc7 | 716,95 | -0,2680 | 0,372355701 | chr11 | - |

| | | | | | |
|---|---|---|---|---|---|
| Dock2 | 45,53 | 0,4168 | 0,357290285 | chr11 | - |
| Dph1 | 163,32 | -0,0853 | 0,863460985 | chr11 | - |
| Drg1 | 1170,22 | -0,0955 | 0,797559244 | chr11 | - |
| Drg2 | 559,13 | -0,1261 | 0,792197952 | chr11 | + |
| Dusp14 | 20,33 | 0,9036 | 0,112357003 | chr11 | - |
| Dusp3 | 900,71 | -0,4104 | 0,22942307 | chr11 | - |
| Dvl2 | 34,82 | -0,1488 | 0,840322097 | chr11 | + |
| Dynll2 | 4435,15 | -0,3032 | 0,267895481 | chr11 | - |
| E230016K23Rik | 2,59 | 0,2179 | 0,783418712 | chr11 | + |
| Ebf1 | 2,62 | -0,1420 | 0,865418807 | chr11 | + |
| Efemp1 | 177,65 | 0,5401 | 0,138593946 | chr11 | + |
| Eftud2 | 301,79 | -0,0439 | 0,939351122 | chr11 | - |
| Ehbp1 | 133,76 | -0,1808 | 0,693280266 | chr11 | - |
| Eif1 | 6656,28 | -0,3969 | 0,117722866 | chr11 | + |
| Eif4a1 | 8605,15 | -0,2747 | 0,434864736 | chr11 | - |
| Eif4a3 | 319,13 | -0,3131 | 0,428679263 | chr11 | - |
| Eif4enif1 | 316,01 | 0,0339 | 0,950532633 | chr11 | + |
| Eif5a | 4706,07 | -0,3821 | 0,203633702 | chr11 | - |
| Elac2 | 214,22 | -0,3526 | 0,273018284 | chr11 | + |
| Elp5 | 586,39 | -0,1581 | 0,715135442 | chr11 | - |
| Eme1 | 3,01 | 0,0963 | 0,916852736 | chr11 | - |
| Eml6 | 8,30 | 0,3329 | 0,665242648 | chr11 | - |
| Endov | 83,23 | -0,1517 | 0,756319626 | chr11 | + |
| Engase | 56,13 | -0,4028 | 0,41272264 | chr11 | + |
| Eno3 | 48,69 | -0,6808 | 0,10872394 | chr11 | + |
| Tepsin | 82,31 | -0,4350 | 0,336399308 | chr11 | - |
| Epn2 | 109,95 | 0,2712 | 0,483930201 | chr11 | - |
| Eral1 | 249,61 | -0,1341 | 0,751909492 | chr11 | - |
| Erbb2 | 12,24 | 0,5531 | 0,37980999 | chr11 | + |
| Erlec1 | 623,82 | -0,2763 | 0,403718537 | chr11 | - |
| Ern1 | 192,67 | 0,2085 | 0,615799562 | chr11 | - |
| Etaa1 | 115,45 | 0,2460 | 0,568858283 | chr11 | - |
| Etv4 | 2,62 | 0,3370 | 0,628656316 | chr11 | - |
| Evi2a | 35,14 | 0,1519 | 0,814771257 | chr11 | - |
| Evpl | 3,17 | 0,3501 | 0,617869186 | chr11 | - |
| Ewsr1 | 1301,34 | -0,0137 | 0,981067951 | chr11 | - |
| Exoc7 | 385,86 | -0,0524 | 0,909426562 | chr11 | - |
| Ezh1 | 220,80 | 0,4619 | 0,166993075 | chr11 | - |
| Fads6 | 205,98 | 0,5157 | 0,209412299 | chr11 | - |
| Rflnb | 15,30 | -0,7059 | 0,222670104 | chr11 | - |
| Fam104a | 191,52 | -0,4402 | 0,265239441 | chr11 | - |
| Fam114a2 | 1067,76 | 0,1785 | 0,614056134 | chr11 | - |
| Fam117a | 30,51 | -0,3073 | 0,63959323 | chr11 | + |
| Retreg3 | 1495,93 | -0,3031 | 0,329985059 | chr11 | - |
| Fam161a | 30,83 | -0,3008 | 0,618381442 | chr11 | + |
| Fam171a2 | 5,56 | -0,1059 | 0,910138296 | chr11 | - |
| Mcrip1 | 211,92 | -0,0548 | 0,909519681 | chr11 | - |
| Fam20a | 331,22 | -0,1681 | 0,743336656 | chr11 | - |
| Fam222b | 182,65 | -0,4739 | 0,162272166 | chr11 | + |
| Fam57a | 10,43 | 0,5509 | 0,394187455 | chr11 | + |

| | | | | | |
|---|---|---|---|---|---|
| Fam58b | 103,42 | 0,0241 | 0,969512928 | chr11 | - |
| Pimreg | 3,75 | 0,0370 | 0,971756379 | chr11 | + |
| Fam83g | 22,91 | -0,2777 | 0,695967514 | chr11 | + |
| Fasn | 3506,62 | 0,3408 | 0,646273602 | chr11 | - |
| Fbf1 | 173,95 | -0,8228 | 0,107501807 | chr11 | - |
| Fbxl20 | 585,64 | -0,1410 | 0,681592614 | chr11 | - |
| Fbxw11 | 536,48 | 0,0656 | 0,862616891 | chr11 | + |
| Fdxr | 191,96 | -0,3021 | 0,563218321 | chr11 | - |
| Fgf11 | 5,70 | 0,1357 | 0,882801741 | chr11 | - |
| Fignl1 | 11,03 | 0,9873 | 0,084779896 | chr11 | - |
| Fkbp10 | 14,44 | 0,5900 | 0,341412029 | chr11 | + |
| Flcn | 543,36 | -0,1920 | 0,683128829 | chr11 | - |
| Flii | 434,43 | -0,1585 | 0,692352484 | chr11 | - |
| Flot2 | 367,28 | -0,2020 | 0,630381755 | chr11 | + |
| Fmnl1 | 12,82 | 0,5328 | 0,413529686 | chr11 | + |
| Fn3k | 321,61 | -0,5236 | 0,087710179 | chr11 | + |
| Fn3krp | 134,00 | 0,5137 | 0,260028246 | chr11 | + |
| Fnip1 | 939,32 | -0,0193 | 0,976788115 | chr11 | + |
| Foxj1 | 5,39 | 0,4860 | 0,485480319 | chr11 | - |
| Foxk2 | 286,17 | -0,3776 | 0,227528657 | chr11 | + |
| Ftsj3 | 820,86 | -0,2152 | 0,584765621 | chr11 | - |
| Fxr2 | 415,31 | -0,3597 | 0,334804026 | chr11 | + |
| Fzd2 | 2,40 | -0,2971 | NA | chr11 | + |
| G3bp1 | 717,83 | 0,1934 | 0,611268902 | chr11 | + |
| G630025P09Rik | 2,14 | 0,1886 | NA | chr11 | + |
| G6pc | 9541,43 | -0,4237 | 0,553397688 | chr11 | + |
| G6pc3 | 59,60 | -0,1835 | 0,743253299 | chr11 | + |
| Gaa | 572,64 | -0,4229 | 0,331535469 | chr11 | + |
| Gabarap | 4999,23 | -0,4150 | 0,095464405 | chr11 | + |
| Galk1 | 579,98 | -0,0124 | 0,988512389 | chr11 | - |
| Galnt10 | 104,45 | 0,7063 | 0,065354125 | chr11 | + |
| Gas2l1 | 106,87 | -0,7917 | 0,111722947 | chr11 | - |
| Gas7 | 58,12 | 0,1961 | 0,744313101 | chr11 | + |
| Gatsl3 | 20,60 | 0,3964 | 0,52307695 | chr11 | + |
| Gck | 2430,27 | -0,2674 | 0,618512306 | chr11 | - |
| Gdf9 | 6,52 | -0,9883 | 0,081189385 | chr11 | + |
| Gdpd1 | 19,65 | 0,2676 | 0,695978671 | chr11 | - |
| Gemin4 | 119,75 | -0,4383 | 0,217231457 | chr11 | - |
| Gemin5 | 127,52 | 0,0269 | 0,964556444 | chr11 | - |
| Gfap | 1,99 | 0,1490 | NA | chr11 | - |
| Gfpt2 | 4,55 | 0,4000 | 0,58928611 | chr11 | + |
| Gga3 | 79,62 | 0,1742 | 0,761732176 | chr11 | - |
| Ggnbp2 | 1577,07 | -0,0966 | 0,816025345 | chr11 | - |
| Ggt6 | 220,19 | -0,0403 | 0,952576171 | chr11 | + |
| Ghdc | 209,92 | 0,0035 | 0,995735598 | chr11 | - |
| Gid4 | 520,63 | -0,3184 | 0,217419188 | chr11 | + |
| Git1 | 76,13 | -0,4086 | 0,462469544 | chr11 | + |
| Gjc1 | 5,21 | 0,6773 | 0,290714093 | chr11 | - |
| Gjc2 | 4,43 | 0,3784 | 0,605538599 | chr11 | - |
| Glod4 | 1771,62 | 0,1582 | 0,666980604 | chr11 | - |

| | | | | | |
|---|---|---|---|---|---|
| Gltpd2 | 566,05 | -0,1718 | 0,754460745 | chr11 | + |
| Gm11517 | 2,92 | 0,3290 | 0,656897388 | chr11 | - |
| Gm11545 | 6,49 | 0,3490 | 0,647914259 | chr11 | - |
| Gm11696 | 5,01 | -0,2839 | 0,723425525 | chr11 | - |
| Prcd | 3,00 | 0,5584 | 0,374674288 | chr11 | + |
| Snhg15 | 22,72 | -0,0890 | 0,914454715 | chr11 | - |
| Gm11992 | 14,45 | 0,4023 | 0,549054641 | chr11 | + |
| Gm12070 | 111,81 | 0,2837 | 0,499122607 | chr11 | + |
| Gm12185 | 9,60 | 0,0474 | 0,961393918 | chr11 | - |
| Gm12216 | 6,07 | -0,2856 | 0,721172264 | chr11 | - |
| Gm12250 | 203,60 | 0,4609 | 0,521462295 | chr11 | + |
| Gm12338 | 30,53 | -0,5225 | 0,325442999 | chr11 | - |
| Natd1 | 100,82 | -0,0841 | 0,869067375 | chr11 | - |
| Gm4926 | 2,33 | -0,0176 | NA | chr11 | - |
| Gm5431 | 29,51 | 0,1013 | 0,905147397 | chr11 | - |
| Gna13 | 504,57 | 0,1062 | 0,7588734 | chr11 | + |
| Rack1 | 7124,19 | -0,4741 | 0,163498128 | chr11 | + |
| Gngt2 | 259,62 | 0,1805 | 0,658260611 | chr11 | + |
| Gosr1 | 579,38 | 0,3105 | 0,253291899 | chr11 | - |
| Gosr2 | 1404,89 | -0,1713 | 0,664283993 | chr11 | - |
| Gpatch8 | 217,13 | 0,1160 | 0,816914587 | chr11 | - |
| Gprc5c | 1068,85 | 0,3092 | 0,247554818 | chr11 | + |
| Gps1 | 1071,58 | -0,2016 | 0,523118038 | chr11 | + |
| Gps2 | 126,44 | -0,1951 | 0,690373216 | chr11 | + |
| Gpx3 | 60,21 | 0,1470 | 0,793250646 | chr11 | + |
| Grap | 78,12 | 0,1388 | 0,773900067 | chr11 | + |
| Grb10 | 109,06 | 0,4741 | 0,274268187 | chr11 | - |
| Grb2 | 512,60 | 0,0356 | 0,947566465 | chr11 | - |
| Grb7 | 939,00 | -0,0462 | 0,948314012 | chr11 | + |
| Grn | 2256,48 | 0,0094 | 0,988603143 | chr11 | + |
| Haspin | 7,44 | 0,3796 | 0,613341495 | chr11 | - |
| Guk1 | 399,53 | -0,4750 | 0,336298097 | chr11 | - |
| H2afv | 469,43 | -0,4267 | 0,317123683 | chr11 | - |
| H3f3b | 3739,45 | 0,1250 | 0,719953965 | chr11 | - |
| Hap1 | 20,58 | -0,1913 | 0,803952241 | chr11 | - |
| Havcr1 | 2,84 | 0,1533 | 0,860044836 | chr11 | + |
| Havcr2 | 10,18 | 0,8952 | 0,13621142 | chr11 | + |
| Hdac5 | 377,41 | -0,0771 | 0,906240001 | chr11 | - |
| Heatr6 | 132,90 | 0,0643 | 0,894495595 | chr11 | + |
| Helz | 36,49 | 0,0815 | 0,910351158 | chr11 | + |
| Hexdc | 104,78 | 0,4483 | 0,240312845 | chr11 | + |
| Hexim1 | 476,20 | -0,3150 | 0,527966188 | chr11 | + |
| Hexim2 | 23,36 | 0,4876 | 0,405980286 | chr11 | + |
| Hgs | 287,88 | -0,0449 | 0,941754349 | chr11 | + |
| Hic1 | 19,34 | 0,3892 | 0,575646447 | chr11 | - |
| Hid1 | 14,69 | 0,1365 | 0,868723893 | chr11 | - |
| Hint1 | 3361,53 | -0,4065 | 0,419474305 | chr11 | + |
| Hist3h2a | 19,14 | 0,0232 | 0,982554344 | chr11 | + |
| Hlf | 770,27 | -0,0791 | 0,87618436 | chr11 | - |
| Hmmr | 22,49 | 0,4717 | 0,42048967 | chr11 | - |

| | | | | | |
|---|---|---|---|---|---|
| Jpt1 | 258,68 | -0,1825 | 0,691519383 | chr11 | - |
| Hnf1b | 95,87 | 0,2465 | 0,63251737 | chr11 | + |
| Hnrnpab | 1813,81 | -0,3531 | 0,188249599 | chr11 | - |
| Hnrnph1 | 1159,86 | -0,0854 | 0,841579427 | chr11 | + |
| Hormad2 | 2,41 | 0,5406 | NA | chr11 | - |
| Hoxb2 | 4,95 | 0,2134 | 0,797919644 | chr11 | + |
| Hoxb3 | 8,95 | 0,5084 | 0,464378372 | chr11 | + |
| Hoxb4 | 11,95 | 0,2433 | 0,766748847 | chr11 | + |
| Hoxb5 | 8,85 | 0,0193 | 0,985716259 | chr11 | + |
| Hoxb7 | 3,72 | 0,6598 | 0,299245205 | chr11 | + |
| Hs3st3a1 | 7,54 | 0,0547 | 0,955496112 | chr11 | + |
| Hs3st3b1 | 1501,05 | 0,1087 | 0,78569213 | chr11 | - |
| Hsd17b1 | 2,59 | -0,3075 | 0,673076056 | chr11 | + |
| Hsf5 | 2,47 | -0,3182 | NA | chr11 | + |
| Hspa4 | 2486,87 | -0,3027 | 0,365218668 | chr11 | - |
| Hus1 | 90,04 | 0,5185 | 0,172076776 | chr11 | - |
| Iba57 | 164,39 | 0,1069 | 0,853363525 | chr11 | - |
| Icam2 | 64,05 | -0,0646 | 0,925898178 | chr11 | - |
| Mrpl58 | 398,34 | -0,2011 | 0,697779102 | chr11 | + |
| Ifi35 | 699,02 | 0,1851 | 0,715419044 | chr11 | + |
| Ifi47 | 854,18 | 0,8437 | 0,1598616 | chr11 | + |
| Ift20 | 868,18 | -0,5602 | 0,176249471 | chr11 | + |
| Igfbp1 | 1203,92 | -0,6208 | 0,201435122 | chr11 | + |
| Igfbp3 | 390,10 | 0,1845 | 0,739307754 | chr11 | - |
| Igfbp4 | 16921,57 | -0,4742 | 0,066490233 | chr11 | + |
| Igtp | 1727,26 | 0,2438 | 0,767097454 | chr11 | + |
| Ikzf1 | 41,11 | 0,3380 | 0,522722304 | chr11 | + |
| Ikzf3 | 5,00 | 0,5348 | 0,430392697 | chr11 | - |
| Inca1 | 300,64 | 0,3120 | 0,480124139 | chr11 | - |
| Inpp5k | 239,51 | -0,0875 | 0,862451616 | chr11 | + |
| Ints2 | 89,68 | 0,5501 | 0,151178014 | chr11 | - |
| Irf1 | 546,79 | 0,3059 | 0,546270816 | chr11 | + |
| Irgm1 | 1357,48 | 0,3340 | 0,515706574 | chr11 | - |
| Irgm2 | 842,75 | 0,6370 | 0,096645255 | chr11 | + |
| Itga2b | 36,58 | -0,3884 | 0,465196538 | chr11 | - |
| Itga3 | 64,95 | 0,1125 | 0,872521494 | chr11 | - |
| Itgae | 11,97 | 0,2525 | 0,740716918 | chr11 | + |
| Itgb3 | 79,47 | 0,5986 | 0,209104214 | chr11 | + |
| Itgb4 | 6,37 | 0,1023 | 0,913936879 | chr11 | + |
| Itk | 5,02 | 0,3335 | 0,665028267 | chr11 | - |
| Jade2 | 85,15 | 0,5062 | 0,274268187 | chr11 | - |
| Jmjd4 | 92,67 | 0,3407 | 0,432704831 | chr11 | + |
| Jmjd6 | 196,32 | -0,3183 | 0,301759036 | chr11 | - |
| Jup | 255,14 | -0,0910 | 0,875746336 | chr11 | - |
| Kansl1 | 269,23 | -0,1254 | 0,779818054 | chr11 | - |
| Kat2a | 182,50 | -0,4649 | 0,202636202 | chr11 | - |
| Kat7 | 328,13 | -0,2392 | 0,463462348 | chr11 | - |
| Kcnab3 | 4,94 | -0,2970 | 0,704548549 | chr11 | + |
| Kcnj16 | 39,21 | 0,0198 | 0,981991128 | chr11 | + |
| Kcnj2 | 20,29 | 0,0703 | 0,9368702 | chr11 | + |

| | | | | | |
|---|---|---|---|---|---|
| Kcnmb1 | 3,72 | -0,1484 | 0,862616891 | chr11 | + |
| Kctd11 | 19,61 | 0,3400 | 0,60487792 | chr11 | - |
| Kctd2 | 301,21 | 0,0134 | 0,979212524 | chr11 | + |
| Kdm6b | 185,79 | -0,1456 | 0,822067344 | chr11 | - |
| Kif19a | 2,10 | 0,4435 | NA | chr11 | + |
| Kif1c | 1646,72 | -0,1976 | 0,613341495 | chr11 | + |
| Kif3a | 104,87 | 0,2370 | 0,631417604 | chr11 | + |
| Klhl11 | 16,88 | 0,6035 | 0,297351163 | chr11 | - |
| Kpna2 | 1430,71 | -0,1861 | 0,688062524 | chr11 | - |
| Kpnb1 | 1649,64 | -0,1861 | 0,523048388 | chr11 | - |
| Kremen1 | 117,37 | -0,1389 | 0,783422709 | chr11 | - |
| Krt10 | 50,76 | 0,0182 | 0,980186108 | chr11 | - |
| Krt19 | 25,13 | 0,6531 | 0,213994613 | chr11 | - |
| Krt222 | 2,38 | 0,0226 | NA | chr11 | - |
| Ksr1 | 11,03 | 0,2037 | 0,803874753 | chr11 | - |
| Larp1 | 1194,61 | -0,2350 | 0,421803323 | chr11 | + |
| Lasp1 | 1262,60 | 0,1654 | 0,713100089 | chr11 | + |
| Lcp2 | 1645,01 | -0,3394 | 0,273018284 | chr11 | + |
| Leap2 | 1268,80 | -0,3507 | 0,438190165 | chr11 | - |
| P3h4 | 12,33 | -0,4021 | 0,554786054 | chr11 | - |
| Lgals3bp | 30,66 | -0,4977 | 0,421924411 | chr11 | - |
| Lgals9 | 9580,22 | 0,2906 | 0,328388243 | chr11 | - |
| Lgalsl | 224,49 | 0,5454 | 0,075380094 | chr11 | - |
| Lif | 1,92 | 0,5497 | NA | chr11 | + |
| Lig3 | 308,30 | 0,1316 | 0,78387537 | chr11 | + |
| Limk2 | 88,85 | 0,6165 | 0,151731441 | chr11 | - |
| Llgl1 | 78,43 | -0,0090 | 0,991298694 | chr11 | + |
| Llgl2 | 195,80 | -0,0804 | 0,924573117 | chr11 | + |
| Lrrc45 | 79,46 | -0,5945 | 0,235271103 | chr11 | + |
| Lrrc46 | 16,74 | 0,0990 | 0,905147397 | chr11 | - |
| Drc3 | 341,67 | -0,1236 | 0,780472013 | chr11 | + |
| Lrrc59 | 889,33 | -0,1199 | 0,783373467 | chr11 | + |
| Lrrc75a | 9,20 | -0,2503 | 0,756482319 | chr11 | - |
| Lsm11 | 37,05 | -0,2374 | 0,669131027 | chr11 | - |
| Lsm12 | 212,83 | -0,3281 | 0,447848007 | chr11 | - |
| Ltc4s | 8,30 | 0,1056 | 0,90993422 | chr11 | - |
| Luc7l3 | 876,02 | -0,1716 | 0,579010212 | chr11 | - |
| Lyrm7 | 21,16 | 0,0007 | 0,999080228 | chr11 | - |
| Lyrm9 | 107,56 | 0,1756 | 0,716547565 | chr11 | + |
| Maml1 | 140,56 | 0,1327 | 0,769969083 | chr11 | - |
| Map2k3 | 943,79 | -0,1903 | 0,688905597 | chr11 | + |
| Map2k3os | 19,04 | -0,3312 | 0,640132917 | chr11 | - |
| Map2k4 | 581,61 | -0,1863 | 0,619935309 | chr11 | - |
| Map2k6 | 112,01 | -0,4195 | 0,434177785 | chr11 | + |
| Map3k14 | 30,43 | -0,0688 | 0,932528423 | chr11 | - |
| Map3k3 | 149,35 | 0,1454 | 0,721615343 | chr11 | + |
| Mapk7 | 22,31 | -0,2662 | 0,709170968 | chr11 | - |
| Mapk9 | 652,80 | -0,0015 | 0,997464504 | chr11 | + |
| Mapt | 35,16 | 0,1147 | 0,86120828 | chr11 | + |
| Mat2b | 819,03 | 0,0004 | 0,998994678 | chr11 | - |

| | | | | | |
|---|---|---|---|---|---|
| Mbtd1 | 321,75 | 0,0677 | 0,87328174 | chr11 | + |
| Mdh1 | 21771,06 | -0,1674 | 0,650939157 | chr11 | - |
| Med1 | 419,79 | 0,0035 | 0,994315523 | chr11 | - |
| Med11 | 155,12 | 0,0343 | 0,96674257 | chr11 | + |
| Med13 | 518,93 | -0,0640 | 0,876857496 | chr11 | - |
| Med24 | 130,20 | -0,1139 | 0,821196964 | chr11 | - |
| Med31 | 114,11 | 0,2418 | 0,581725594 | chr11 | - |
| Med7 | 270,31 | -0,2100 | 0,583782269 | chr11 | + |
| Med9 | 65,95 | 0,1422 | 0,792197952 | chr11 | + |
| Meis1 | 45,05 | 0,6466 | 0,165257806 | chr11 | - |
| Metrnl | 54,54 | -0,6800 | 0,15338656 | chr11 | + |
| Mettl16 | 244,86 | 0,0018 | 0,997008467 | chr11 | + |
| Mettl2 | 166,18 | -0,0069 | 0,989980239 | chr11 | + |
| Mettl23 | 475,08 | -0,1907 | 0,564432176 | chr11 | + |
| Mfap3 | 238,23 | 0,2567 | 0,490386392 | chr11 | + |
| Mfap4 | 38,81 | 0,4797 | 0,399877545 | chr11 | + |
| Mfsd11 | 197,49 | -0,5463 | 0,061534399 | chr11 | + |
| Mgat1 | 1016,32 | -0,0015 | 0,997464504 | chr11 | + |
| Mgat4b | 230,95 | -0,2763 | 0,471103626 | chr11 | + |
| Mgl2 | 7,90 | 0,1188 | 0,89879469 | chr11 | + |
| Mief2 | 260,15 | -0,5304 | 0,172578291 | chr11 | + |
| Mif4gd | 482,93 | -0,1830 | 0,732982182 | chr11 | - |
| Milr1 | 21,34 | 0,3231 | 0,619691963 | chr11 | + |
| Mink1 | 364,59 | -0,1651 | 0,730063556 | chr11 | + |
| Mir3061 | 3,36 | -0,5649 | 0,382591532 | chr11 | + |
| Mir3064 | 59,57 | -0,0713 | 0,924731513 | chr11 | - |
| Mis12 | 423,06 | 0,3495 | 0,232472267 | chr11 | + |
| Mks1 | 17,16 | 0,8766 | 0,10921447 | chr11 | + |
| Mllt6 | 262,10 | 0,1052 | 0,793705416 | chr11 | + |
| Mlx | 749,40 | -0,0455 | 0,925034844 | chr11 | + |
| Mmd | 2913,70 | 0,2495 | 0,520016085 | chr11 | + |
| Mmgt2 | 21,19 | 0,1422 | 0,847964947 | chr11 | + |
| Mnt | 99,94 | 0,0809 | 0,920355471 | chr11 | + |
| Morc2a | 237,60 | -0,0864 | 0,830330662 | chr11 | + |
| Mpdu1 | 943,85 | -0,1816 | 0,618381442 | chr11 | - |
| Mpg | 122,79 | 0,5139 | 0,122950106 | chr11 | + |
| Mpp2 | 5,13 | -0,0090 | 0,99298417 | chr11 | - |
| Mprip | 922,25 | -0,3134 | 0,390793229 | chr11 | + |
| Mrc2 | 18,63 | 0,6162 | 0,303602228 | chr11 | + |
| Mrm1 | 51,73 | 0,1476 | 0,806159394 | chr11 | - |
| Mrpl10 | 738,41 | -0,2449 | 0,342972129 | chr11 | + |
| Mrpl12 | 2032,91 | -0,5244 | 0,183699238 | chr11 | + |
| Mrpl22 | 577,38 | -0,0947 | 0,818573937 | chr11 | + |
| Mrpl27 | 1042,35 | -0,0685 | 0,869687361 | chr11 | + |
| Mrpl38 | 686,44 | -0,6891 | 0,056627138 | chr11 | - |
| Mrpl45 | 552,95 | -0,1381 | 0,744322442 | chr11 | + |
| Mrpl55 | 260,48 | -0,3143 | 0,39766425 | chr11 | + |
| Mrps23 | 403,82 | -0,2866 | 0,420955645 | chr11 | + |
| Mrps7 | 923,23 | -0,2870 | 0,377944963 | chr11 | + |
| Msi2 | 337,60 | 0,3305 | 0,3025413 | chr11 | - |

| | | | | | |
|---|---|---|---|---|---|
| Msl1 | 596,73 | -0,2684 | 0,379104437 | chr11 | + |
| Mtfp1 | 274,20 | -0,2822 | 0,526958509 | chr11 | - |
| Mtif2 | 615,97 | 0,0554 | 0,909774279 | chr11 | + |
| Mtmr3 | 449,04 | -0,2924 | 0,358990473 | chr11 | - |
| Mtmr4 | 677,91 | -0,2307 | 0,55237484 | chr11 | + |
| Mxra7 | 38,50 | 0,4241 | 0,380860886 | chr11 | - |
| Mybbp1a | 369,80 | -0,1271 | 0,754953629 | chr11 | + |
| Myh1 | 2,40 | 0,4865 | NA | chr11 | + |
| Myh8 | 2,28 | 0,0982 | NA | chr11 | + |
| Myl4 | 6,48 | 0,0172 | 0,987025571 | chr11 | + |
| Myl7 | 5,95 | 0,2910 | 0,712581071 | chr11 | - |
| Myo18a | 818,08 | -0,1090 | 0,815188933 | chr11 | + |
| Myo19 | 76,60 | -0,4827 | 0,227017349 | chr11 | + |
| Myo1c | 544,33 | 0,0920 | 0,810525833 | chr11 | + |
| Myo1d | 108,90 | 0,1798 | 0,692167613 | chr11 | - |
| Myo1g | 18,11 | 0,1373 | 0,868479494 | chr11 | - |
| N4bp3 | 37,92 | 0,4924 | 0,400330472 | chr11 | - |
| Naa38 | 670,97 | -0,4665 | 0,195865369 | chr11 | + |
| Naglu | 314,88 | 0,1148 | 0,772705111 | chr11 | + |
| Nags | 665,29 | -0,0427 | 0,943108135 | chr11 | + |
| Narf | 1237,11 | 0,0361 | 0,940450964 | chr11 | + |
| Nat9 | 505,92 | -0,0419 | 0,921657556 | chr11 | - |
| Nbr1 | 3877,20 | 0,2678 | 0,357345082 | chr11 | + |
| Ncor1 | 843,13 | -0,2359 | 0,390832005 | chr11 | - |
| Ndel1 | 687,14 | 0,0025 | 0,996644671 | chr11 | - |
| Nek8 | 78,52 | -0,1643 | 0,751184537 | chr11 | - |
| Neurl4 | 152,88 | -0,1187 | 0,805374431 | chr11 | + |
| Nf1 | 228,18 | 0,0374 | 0,951934337 | chr11 | + |
| Nf2 | 307,00 | 0,0030 | 0,995735598 | chr11 | - |
| Nfe2l1 | 1758,47 | 0,0116 | 0,986661276 | chr11 | - |
| Ngfr | 21,08 | 0,6460 | 0,260755195 | chr11 | - |
| Nhp2 | 409,52 | -0,4985 | 0,263764485 | chr11 | + |
| Nipsnap1 | 3923,51 | -0,1395 | 0,666630958 | chr11 | + |
| Nkiras2 | 225,84 | 0,0098 | 0,986661276 | chr11 | + |
| Nle1 | 30,14 | 0,0046 | 0,995836778 | chr11 | - |
| Nlgn2 | 19,29 | -0,3329 | 0,614209897 | chr11 | - |
| Nlk | 193,73 | -0,4506 | 0,169162729 | chr11 | - |
| Nlrp1a | 38,01 | 0,1722 | 0,803027202 | chr11 | - |
| Nlrp1b | 12,27 | 0,2506 | 0,75616611 | chr11 | - |
| Nlrp3 | 9,63 | 0,3556 | 0,632226809 | chr11 | + |
| Nmt1 | 1016,06 | -0,1348 | 0,687892476 | chr11 | + |
| Nol11 | 200,46 | 0,1252 | 0,77892794 | chr11 | - |
| Notum | 263,49 | -0,2173 | 0,621479923 | chr11 | - |
| Npb | 3,97 | -0,2129 | 0,796014393 | chr11 | + |
| Npepps | 365,37 | -0,1287 | 0,74146296 | chr11 | - |
| Nploc4 | 1049,84 | -0,3733 | 0,162509787 | chr11 | - |
| Npm1 | 4464,09 | -0,0468 | 0,933235928 | chr11 | - |
| Nprl3 | 111,55 | -0,2210 | 0,61720787 | chr11 | - |
| Nr1d1 | 751,32 | -0,1108 | 0,906998011 | chr11 | - |
| Nsf | 176,72 | 0,0776 | 0,879335212 | chr11 | - |

| | | | | | |
|---|---|---|---|---|---|
| Nt5c | 295,64 | -0,0913 | 0,870119976 | chr11 | - |
| Nt5c3b | 51,84 | 0,0065 | 0,993445259 | chr11 | - |
| Nt5m | 87,09 | -0,1070 | 0,847964947 | chr11 | + |
| Ntn1 | 121,10 | 0,2791 | 0,516675984 | chr11 | - |
| Nudcd2 | 497,94 | -0,0384 | 0,941553259 | chr11 | + |
| Nudcd3 | 412,09 | 0,1167 | 0,789524079 | chr11 | - |
| Nufip2 | 952,26 | -0,2189 | 0,464998681 | chr11 | + |
| Nup85 | 245,11 | 0,0922 | 0,834507118 | chr11 | + |
| Nup88 | 502,75 | -0,2117 | 0,474794806 | chr11 | - |
| Nxn | 28,02 | -0,3979 | 0,475870181 | chr11 | - |
| Ogdh | 1817,79 | -0,4633 | 0,147367695 | chr11 | + |
| Ogfod3 | 118,93 | -0,1152 | 0,799982428 | chr11 | - |
| Olfr1372-ps1 | 8,03 | -0,2939 | 0,712160795 | chr11 | - |
| Olfr56 | 2,44 | 0,3711 | NA | chr11 | + |
| Ormdl3 | 741,10 | 0,0423 | 0,918077233 | chr11 | - |
| Osbp2 | 9,39 | 0,9627 | 0,098009661 | chr11 | - |
| Osbpl7 | 12,12 | 0,3593 | 0,60933115 | chr11 | + |
| Ovca2 | 138,45 | -0,0232 | 0,977112784 | chr11 | - |
| Oxld1 | 100,02 | -0,5053 | 0,228424355 | chr11 | - |
| P4ha2 | 96,07 | -0,3759 | 0,473786638 | chr11 | + |
| P4hb | 22754,41 | -0,3454 | 0,235063176 | chr11 | - |
| Pafah1b1 | 1684,15 | -0,0302 | 0,943305207 | chr11 | - |
| Pank3 | 1198,60 | -0,1347 | 0,720291746 | chr11 | + |
| Papolg | 103,43 | 0,6051 | 0,077385039 | chr11 | - |
| Patz1 | 29,80 | -0,1991 | 0,760045267 | chr11 | + |
| Pcgf2 | 29,72 | -0,5345 | 0,39588956 | chr11 | - |
| Pctp | 790,23 | 0,0474 | 0,941757244 | chr11 | - |
| Pcyt2 | 3,83 | -0,1190 | 0,884450766 | chr11 | - |
| Pde6g | 5,63 | 0,1768 | 0,841053685 | chr11 | - |
| Pdk2 | 837,26 | -0,3021 | 0,411158799 | chr11 | - |
| Pdlim4 | 8,06 | -0,0316 | 0,977120448 | chr11 | - |
| Pecam1 | 344,11 | 0,3523 | 0,174738595 | chr11 | - |
| Peli1 | 324,43 | 0,2095 | 0,606534862 | chr11 | + |
| Pelp1 | 101,96 | 0,2537 | 0,607935435 | chr11 | - |
| Pemt | 8473,55 | -0,1131 | 0,757695155 | chr11 | - |
| Per1 | 307,19 | -0,2189 | 0,656897388 | chr11 | + |
| Pes1 | 700,48 | 0,0954 | 0,838879513 | chr11 | + |
| Pex12 | 191,77 | -0,1148 | 0,797252833 | chr11 | - |
| Pex13 | 1246,88 | -0,3021 | 0,317045416 | chr11 | - |
| Pfas | 74,52 | 0,1809 | 0,746414189 | chr11 | - |
| Pfn1 | 1709,12 | -0,4812 | 0,202636202 | chr11 | - |
| Pgap3 | 13,78 | -0,2859 | 0,696436516 | chr11 | - |
| Pgs1 | 94,51 | -0,2890 | 0,572283367 | chr11 | + |
| Phb | 4073,16 | -0,1965 | 0,466056885 | chr11 | + |
| Phf12 | 160,04 | -0,4131 | 0,276399425 | chr11 | + |
| Phf23 | 247,57 | -0,1056 | 0,841053685 | chr11 | + |
| Phospho1 | 42,11 | -0,0655 | 0,940390384 | chr11 | + |
| Phykpl | 174,44 | -0,0413 | 0,941553259 | chr11 | + |
| Pigl | 42,25 | -0,2613 | 0,612839776 | chr11 | + |
| Pigs | 246,16 | -0,2849 | 0,536983222 | chr11 | + |

| | | | | | |
|---|---|---|---|---|---|
| Pigw | 89,31 | -0,1651 | 0,764664236 | chr11 | - |
| Pik3ip1 | 16,02 | -0,4604 | 0,468711508 | chr11 | + |
| Pik3r5 | 16,67 | 0,3197 | 0,658260611 | chr11 | + |
| Pik3r6 | 11,88 | -0,1527 | 0,854690386 | chr11 | + |
| Pip4k2b | 284,14 | 0,1516 | 0,700075571 | chr11 | - |
| Pipox | 220,18 | 0,3052 | 0,470066395 | chr11 | - |
| Pisd-ps1 | 821,23 | 0,3762 | 0,37315636 | chr11 | + |
| Pitpna | 888,37 | -0,0103 | 0,984655656 | chr11 | + |
| Pitpnc1 | 524,16 | 0,0259 | 0,971756379 | chr11 | - |
| Plcd3 | 8,81 | 0,3110 | 0,686794434 | chr11 | - |
| Pld2 | 78,59 | -0,1226 | 0,823537093 | chr11 | + |
| Pld6 | 24,03 | -0,0750 | 0,93238328 | chr11 | - |
| Plekhh3 | 51,52 | -0,3383 | 0,631591513 | chr11 | - |
| Plekhm1 | 176,40 | -0,2800 | 0,559525694 | chr11 | - |
| Plscr3 | 52,70 | -0,1742 | 0,77333268 | chr11 | + |
| Plxdc1 | 15,52 | 0,2304 | 0,762940356 | chr11 | - |
| Pmp22 | 82,80 | 0,0980 | 0,85502909 | chr11 | + |
| Pno1 | 613,73 | -0,2730 | 0,408981984 | chr11 | - |
| Pnpt1 | 537,58 | 0,0240 | 0,955504626 | chr11 | + |
| Pold2 | 107,38 | -0,4960 | 0,298176352 | chr11 | - |
| Poldip2 | 1131,71 | -0,2115 | 0,468283516 | chr11 | + |
| Polg2 | 163,06 | -0,3708 | 0,342277341 | chr11 | - |
| Polm | 38,25 | 0,6920 | 0,166586629 | chr11 | - |
| Polr2a | 417,40 | -0,1772 | 0,738449568 | chr11 | - |
| Ppm1d | 80,94 | -0,3243 | 0,437167777 | chr11 | + |
| Ppp1r1b | 122,71 | 0,0791 | 0,919149053 | chr11 | + |
| Ppp1r9b | 74,50 | 0,4680 | 0,211374848 | chr11 | + |
| Ppp2ca | 2064,09 | -0,2842 | 0,298774903 | chr11 | + |
| Ppp3r1 | 759,22 | -0,1152 | 0,79999348 | chr11 | + |
| Prkar1a | 2143,19 | 0,1017 | 0,751702652 | chr11 | + |
| Proca1 | 77,80 | -0,4412 | 0,41977893 | chr11 | + |
| Prorsd1 | 473,26 | -0,1644 | 0,665256143 | chr11 | - |
| Prpf8 | 1918,73 | -0,0650 | 0,868151289 | chr11 | + |
| Prpsap1 | 662,87 | -0,3731 | 0,333078468 | chr11 | - |
| Prpsap2 | 251,94 | 0,2162 | 0,570235502 | chr11 | - |
| Prr11 | 10,72 | -0,0964 | 0,916359044 | chr11 | - |
| Prr15l | 11,84 | 0,1962 | 0,806159394 | chr11 | + |
| Psmb3 | 2850,16 | -0,4522 | 0,060451306 | chr11 | + |
| Psmb6 | 3138,31 | -0,2538 | 0,539172259 | chr11 | + |
| Psmc3ip | 6,68 | 0,4340 | 0,549586708 | chr11 | - |
| Psmc5 | 1399,78 | -0,3035 | 0,342224382 | chr11 | + |
| Psmd11 | 1494,06 | -0,1286 | 0,66076849 | chr11 | + |
| Psmd12 | 1561,64 | -0,3703 | 0,157491037 | chr11 | + |
| Psmd3 | 697,02 | -0,5308 | 0,129193513 | chr11 | + |
| Psme2b | 5,26 | 0,3244 | 0,676312675 | chr11 | - |
| Psme3 | 622,15 | -0,4125 | 0,212297691 | chr11 | + |
| Psme4 | 2934,50 | -0,3656 | 0,168986672 | chr11 | + |
| Ptges3l | 12,79 | -0,6929 | 0,234863974 | chr11 | - |
| Cavin1 | 147,07 | 0,2784 | 0,506368378 | chr11 | - |
| Ptrh2 | 437,61 | 0,1327 | 0,816025345 | chr11 | + |

| | | | | | |
|---|---|---|---|---|---|
| Pttg1 | 108,45 | -0,0984 | 0,8763719 | chr11 | - |
| Purb | 803,54 | 0,1736 | 0,603153385 | chr11 | - |
| Pus10 | 416,81 | 0,0091 | 0,986661276 | chr11 | + |
| Pycr1 | 3,45 | 0,2191 | 0,776478455 | chr11 | - |
| Rab1a | 2828,33 | -0,0223 | 0,961078035 | chr11 | + |
| Rab11fip4 | 18,85 | 0,6334 | 0,268872783 | chr11 | + |
| Rab34 | 71,23 | -0,2935 | 0,55714632 | chr11 | + |
| Rab37 | 6,77 | 0,0112 | 0,991298694 | chr11 | + |
| Rab40b | 4,96 | -0,1270 | 0,89040553 | chr11 | - |
| Rab5c | 1349,04 | -0,0697 | 0,89040553 | chr11 | - |
| Rabep1 | 1125,49 | 0,0146 | 0,977590816 | chr11 | + |
| Rac3 | 3,66 | -0,2144 | 0,790692717 | chr11 | + |
| Rad50 | 159,56 | 0,4567 | 0,161186563 | chr11 | - |
| Rad51c | 19,60 | 0,1214 | 0,888710003 | chr11 | - |
| Rad51d | 167,42 | -0,4099 | 0,212915792 | chr11 | - |
| Rai1 | 63,89 | 0,2907 | 0,581237396 | chr11 | + |
| Ramp2 | 109,62 | -0,2971 | 0,569425025 | chr11 | + |
| Ranbp17 | 2,78 | 0,2648 | 0,726240854 | chr11 | - |
| Rangrf | 52,28 | -0,2066 | 0,705982644 | chr11 | - |
| Rap1gap2 | 11,94 | 0,1735 | 0,830330662 | chr11 | - |
| Rapgef6 | 268,32 | 0,0325 | 0,953751362 | chr11 | + |
| Rapgefl1 | 6,25 | 0,0420 | 0,969159464 | chr11 | + |
| Rara | 143,36 | -0,4277 | 0,390708488 | chr11 | + |
| Rars | 755,13 | 0,0007 | 0,998789109 | chr11 | - |
| Rasd1 | 13,54 | 0,5478 | 0,383567085 | chr11 | - |
| Rbfox3 | 10,90 | -0,0427 | 0,967019378 | chr11 | - |
| Rdm1 | 233,65 | 0,2506 | 0,438176462 | chr11 | + |
| Recql5 | 126,11 | -0,0667 | 0,909289886 | chr11 | - |
| Rel | 18,65 | 0,4451 | 0,456560888 | chr11 | - |
| Rffl | 371,82 | -0,3924 | 0,254608455 | chr11 | - |
| Rfng | 88,36 | -0,6724 | 0,062782484 | chr11 | - |
| Rgs9 | 9,48 | 0,2028 | 0,806012703 | chr11 | - |
| Rhbdd3 | 187,31 | -0,6072 | 0,124458443 | chr11 | + |
| Rhbdf1 | 61,89 | 0,7528 | 0,08440425 | chr11 | - |
| Rhbdf2 | 78,84 | -0,1342 | 0,858880859 | chr11 | - |
| Rhbdl3 | 2,75 | -0,3776 | 0,594322944 | chr11 | + |
| Rhot1 | 705,36 | -0,1958 | 0,523471926 | chr11 | + |
| Rilp | 208,83 | -0,6867 | 0,057654134 | chr11 | + |
| Rmnd5b | 148,41 | 0,1924 | 0,642867567 | chr11 | - |
| Rnasek | 671,97 | -0,6486 | 0,191230937 | chr11 | - |
| Rnd2 | 134,08 | -0,0808 | 0,88569455 | chr11 | + |
| Rnf130 | 1170,50 | -0,3975 | 0,124301562 | chr11 | + |
| Rnf135 | 167,98 | 0,0287 | 0,95728901 | chr11 | + |
| Rnf145 | 503,60 | 0,1087 | 0,828422381 | chr11 | + |
| Rnf157 | 18,50 | -0,2885 | 0,671975571 | chr11 | - |
| Rnf167 | 680,23 | 0,0513 | 0,890293696 | chr11 | + |
| Rnf185 | 470,20 | -0,2331 | 0,437167777 | chr11 | - |
| Rnf187 | 635,13 | -0,3310 | 0,448865116 | chr11 | - |
| Rnf215 | 129,59 | -0,6273 | 0,056429998 | chr11 | + |
| Rnf222 | 3,21 | 0,0859 | 0,923931243 | chr11 | + |

| | | | | | |
|---|---|---|---|---|---|
| Rnf43 | 174,30 | -0,3196 | 0,335528734 | chr11 | + |
| Rnft1 | 853,35 | 0,3131 | 0,305522088 | chr11 | + |
| Mrm3 | 121,42 | -0,0192 | 0,973290912 | chr11 | + |
| Rpain | 319,74 | -0,5149 | 0,108917495 | chr11 | + |
| Rph3al | 38,84 | 0,5104 | 0,287271906 | chr11 | - |
| Rpl19 | 9685,01 | -0,2475 | 0,41272264 | chr11 | + |
| Rpl23 | 10845,79 | -0,5113 | 0,076519476 | chr11 | - |
| Rpl23a | 8626,67 | -0,3675 | 0,213994613 | chr11 | - |
| Rpl27 | 3568,02 | -0,5854 | 0,093542539 | chr11 | + |
| Rps15a-ps6 | 19,09 | -0,2560 | 0,739631507 | chr11 | - |
| Rps27a | 9686,93 | -0,4253 | 0,151155593 | chr11 | - |
| Rps6kb1 | 626,93 | -0,4043 | 0,181882538 | chr11 | - |
| Rptor | 318,45 | -0,2584 | 0,433615866 | chr11 | + |
| Rptoros | 3,23 | 0,7872 | 0,181940259 | chr11 | - |
| Rsad1 | 77,28 | -0,5867 | 0,207095904 | chr11 | - |
| Rufy1 | 253,94 | -0,0809 | 0,860606321 | chr11 | - |
| Rundc1 | 145,84 | -0,2469 | 0,522085068 | chr11 | + |
| Rundc3a | 17,75 | 0,7113 | 0,214852266 | chr11 | + |
| Samd14 | 2,72 | 0,4932 | 0,443670281 | chr11 | + |
| Sap30bp | 302,92 | 0,0097 | 0,985361609 | chr11 | + |
| Sap30l | 187,01 | -0,3549 | 0,263665648 | chr11 | + |
| Sar1b | 4676,20 | -0,3092 | 0,358990473 | chr11 | + |
| Sarm1 | 4,98 | 0,5360 | 0,430392697 | chr11 | - |
| Sat2 | 264,29 | 0,2094 | 0,626400494 | chr11 | + |
| Scarf1 | 175,08 | -0,5492 | 0,210335339 | chr11 | + |
| Scimp | 34,36 | -0,0190 | 0,981991128 | chr11 | - |
| Sco1 | 234,60 | -0,0339 | 0,947612608 | chr11 | + |
| Scpep1 | 801,13 | -0,2571 | 0,465196538 | chr11 | - |
| Scpep1os | 5,88 | 0,4115 | 0,577635343 | chr11 | + |
| Scrn2 | 413,60 | -0,3493 | 0,435263224 | chr11 | + |
| Sdf2 | 1226,93 | -0,3406 | 0,228941765 | chr11 | + |
| Sebox | 16,26 | 0,0078 | 0,993598948 | chr11 | + |
| Sec14l1 | 136,48 | -0,0910 | 0,876030136 | chr11 | + |
| Sec14l2 | 7381,62 | 0,1458 | 0,678676127 | chr11 | - |
| Sec14l4 | 1759,92 | 0,2096 | 0,577347444 | chr11 | + |
| Sec24a | 1594,05 | -0,2055 | 0,55335046 | chr11 | - |
| Selenom | 15,23 | -0,0861 | 0,925898178 | chr11 | + |
| Senp3 | 509,90 | -0,3283 | 0,237903839 | chr11 | - |
| Serpinf1 | 7051,80 | -0,4411 | 0,19521384 | chr11 | - |
| Sertad2 | 324,79 | -0,4070 | 0,171331921 | chr11 | + |
| Sf3a1 | 215,38 | -0,2050 | 0,573421549 | chr11 | + |
| Sfi1 | 153,93 | -0,1277 | 0,77079779 | chr11 | - |
| Sgsh | 57,73 | 0,2220 | 0,723824089 | chr11 | - |
| Sgsm2 | 55,50 | -0,7533 | 0,073845576 | chr11 | - |
| Sh3bp5l | 280,74 | 0,2046 | 0,581252766 | chr11 | + |
| Sh3pxd2b | 14,85 | 0,6488 | 0,288798737 | chr11 | + |
| Shmt1 | 2293,16 | -0,1061 | 0,809686855 | chr11 | - |
| Shpk | 1106,76 | 0,0429 | 0,92369436 | chr11 | + |
| Shroom1 | 146,34 | 0,0928 | 0,875746336 | chr11 | + |
| Sirt7 | 280,91 | -0,4779 | 0,262930762 | chr11 | - |

| | | | | | |
|---|---|---|---|---|---|
| Ska2 | 34,89 | 0,4518 | 0,358779319 | chr11 | + |
| Skap1 | 7,38 | -0,3470 | 0,64594561 | chr11 | + |
| Skp1a | 6479,36 | -0,1469 | 0,692167613 | chr11 | + |
| Slc13a2 | 26,16 | -0,7234 | 0,161186563 | chr11 | - |
| Slc16a11 | 131,40 | -0,0437 | 0,959428625 | chr11 | + |
| Slc16a13 | 26,43 | -0,2685 | 0,66477796 | chr11 | - |
| Slc16a3 | 5,12 | 0,2535 | 0,75378365 | chr11 | + |
| Slc16a6 | 79,72 | 0,5690 | 0,213150875 | chr11 | - |
| Slc1a4 | 55,66 | 0,4819 | 0,296228386 | chr11 | - |
| Slc22a21 | 6,80 | 0,4632 | 0,514682351 | chr11 | - |
| Slc22a4 | 44,31 | -0,4252 | 0,451815929 | chr11 | - |
| Slc22a5 | 552,85 | -0,6359 | 0,154768955 | chr11 | - |
| Slc25a10 | 1719,66 | -0,1784 | 0,701094188 | chr11 | + |
| Slc25a11 | 1323,15 | 0,0491 | 0,896860788 | chr11 | - |
| Slc25a19 | 172,33 | -0,2369 | 0,542534449 | chr11 | - |
| Slc25a35 | 12,40 | -0,0303 | 0,977120448 | chr11 | + |
| Slc25a39 | 3804,41 | -0,0334 | 0,95463264 | chr11 | - |
| Slc26a11 | 103,86 | -0,3992 | 0,302221069 | chr11 | + |
| Slc35b1 | 1064,35 | -0,3744 | 0,338536613 | chr11 | + |
| Slc35e4 | 7,29 | -0,2870 | 0,719953965 | chr11 | - |
| Slc36a1 | 318,82 | 0,1795 | 0,732140841 | chr11 | + |
| Slc36a1os | 7,03 | 0,1003 | 0,915071511 | chr11 | - |
| Slc38a10 | 1378,44 | -0,1686 | 0,670891431 | chr11 | - |
| Slc39a11 | 352,64 | -0,0708 | 0,85848279 | chr11 | - |
| Slc46a1 | 569,38 | -0,5004 | 0,115557884 | chr11 | + |
| Slc47a1 | 1419,86 | 0,4227 | 0,173746653 | chr11 | - |
| Slc4a1 | 2,51 | -0,2821 | NA | chr11 | - |
| Slc6a4 | 10,97 | 0,9339 | 0,105257346 | chr11 | + |
| Slc9a3r1 | 2226,67 | -0,4417 | 0,193343482 | chr11 | + |
| Slfn1 | 8,80 | 0,4344 | 0,550885834 | chr11 | + |
| Slfn2 | 137,31 | 0,3909 | 0,306398925 | chr11 | + |
| Slfn3 | 5,25 | 0,0155 | 0,988512389 | chr11 | + |
| Slfn4 | 10,30 | 0,6472 | 0,316622965 | chr11 | + |
| Slfn5 | 381,26 | 0,3865 | 0,301759036 | chr11 | + |
| Slfn8 | 13,40 | -0,3734 | 0,619039366 | chr11 | - |
| Slfn9 | 19,84 | 0,8335 | 0,112275385 | chr11 | - |
| Slu7 | 577,03 | 0,1907 | 0,575705463 | chr11 | + |
| Smarcd2 | 686,00 | -0,1740 | 0,633851047 | chr11 | - |
| Smarce1 | 793,67 | 0,2316 | 0,527966188 | chr11 | - |
| Smcr8 | 551,88 | 0,1560 | 0,668264827 | chr11 | + |
| Ppp4r3b | 889,33 | 0,1018 | 0,827441247 | chr11 | + |
| Smg6 | 296,71 | -0,2595 | 0,360023268 | chr11 | + |
| Smg8 | 140,23 | 0,0970 | 0,83003778 | chr11 | - |
| Smim5 | 4,31 | 0,2183 | 0,795976629 | chr11 | + |
| Smim6 | 41,91 | 0,4919 | 0,377944963 | chr11 | + |
| Smtn | 46,17 | 0,0388 | 0,955234619 | chr11 | - |
| Smurf2 | 805,36 | 0,0675 | 0,866619594 | chr11 | - |
| Snap47 | 1550,61 | 0,0251 | 0,969159464 | chr11 | - |
| Snf8 | 617,23 | -0,1876 | 0,611914561 | chr11 | + |
| Snora5c | 2,22 | -0,2557 | NA | chr11 | - |

| | | | | | |
|---|---|---|---|---|---|
| Snord104 | 51,82 | -0,5393 | 0,265239441 | chr11 | + |
| Snord49a | 2,64 | -0,4110 | 0,550118606 | chr11 | + |
| Snord96a | 10,61 | -0,3758 | 0,604876098 | chr11 | + |
| Snrnp25 | 100,28 | 0,1939 | 0,692167613 | chr11 | + |
| Snx11 | 87,30 | -0,2712 | 0,575906025 | chr11 | - |
| Socs3 | 112,32 | -0,0213 | 0,9827764 | chr11 | - |
| Socs7 | 285,79 | 0,0802 | 0,850453487 | chr11 | + |
| Sox9 | 692,54 | 0,3530 | 0,247573115 | chr11 | + |
| Sp2 | 696,20 | 0,1561 | 0,743508016 | chr11 | - |
| Spag5 | 13,10 | 0,3168 | 0,668293319 | chr11 | + |
| Spag7 | 625,71 | -0,1756 | 0,583782269 | chr11 | - |
| Spag9 | 1009,32 | 0,0612 | 0,88099064 | chr11 | + |
| Sparc | 1977,45 | 0,2906 | 0,382780136 | chr11 | - |
| Spata22 | 80,07 | 0,4306 | 0,385921204 | chr11 | + |
| Spdl1 | 9,62 | 0,6986 | 0,265239441 | chr11 | - |
| Specc1 | 36,88 | 0,4500 | 0,357779158 | chr11 | + |
| Sphk1 | 3,12 | 0,3671 | 0,610537466 | chr11 | + |
| Spns2 | 297,12 | 0,7286 | 0,067117612 | chr11 | - |
| Spop | 2297,94 | -0,0509 | 0,89928817 | chr11 | + |
| Spred2 | 45,09 | 0,4532 | 0,367954693 | chr11 | + |
| Sptbn1 | 1175,17 | 0,3971 | 0,166077306 | chr11 | - |
| Sqstm1 | 4092,14 | -0,2836 | 0,442624295 | chr11 | - |
| Srebf1 | 1380,45 | 0,0515 | 0,93700068 | chr11 | - |
| Srp68 | 532,16 | -0,2124 | 0,520065437 | chr11 | - |
| Srr | 1624,36 | -0,2427 | 0,340229346 | chr11 | - |
| Srsf1 | 1456,33 | 0,0280 | 0,95728901 | chr11 | + |
| Srsf2 | 950,12 | -0,3670 | 0,258301555 | chr11 | - |
| Ssh2 | 438,19 | -0,3271 | 0,401382558 | chr11 | + |
| St6galnac2 | 20,92 | 0,7007 | 0,217147923 | chr11 | - |
| Stac2 | 3,26 | 0,1610 | 0,851472884 | chr11 | - |
| Stard3 | 231,85 | -0,2627 | 0,554660735 | chr11 | + |
| Stat3 | 1304,96 | -0,0251 | 0,969130697 | chr11 | - |
| Stat5a | 153,50 | 0,3775 | 0,340522938 | chr11 | + |
| Stat5b | 656,11 | 0,3290 | 0,356901218 | chr11 | - |
| Cenpx | 737,71 | -0,1450 | 0,672907874 | chr11 | - |
| Strada | 27,73 | -0,2000 | 0,773900067 | chr11 | - |
| Stx8 | 494,14 | 0,0690 | 0,897982563 | chr11 | + |
| Stxbp4 | 64,18 | 0,1508 | 0,802594059 | chr11 | - |
| Sumo2 | 1554,22 | 0,1655 | 0,647914259 | chr11 | - |
| Sun3 | 7,62 | -0,0318 | 0,976788115 | chr11 | - |
| Supt4a | 754,69 | -0,2781 | 0,335264505 | chr11 | + |
| Supt6 | 498,06 | -0,1924 | 0,598404146 | chr11 | - |
| Suz12 | 585,34 | 0,0965 | 0,816359983 | chr11 | + |
| Syngr2 | 714,81 | -0,2199 | 0,544991268 | chr11 | + |
| Synrg | 221,77 | 0,0243 | 0,962251592 | chr11 | + |
| Taco1 | 265,24 | 0,2238 | 0,564724076 | chr11 | + |
| Tada2a | 71,67 | 0,1943 | 0,717188441 | chr11 | - |
| Taf15 | 1470,36 | -0,0289 | 0,948314012 | chr11 | + |
| Tanc2 | 66,96 | -0,0001 | 0,999705919 | chr11 | + |
| Taok1 | 815,90 | -0,0460 | 0,923778914 | chr11 | - |

| | | | | | |
|---|---|---|---|---|---|
| Tax1bp3 | 396,38 | 0,1024 | 0,830330662 | chr11 | + |
| Tbc1d10a | 139,07 | 0,2003 | 0,742967744 | chr11 | + |
| Tbc1d16 | 381,42 | 0,1965 | 0,629895923 | chr11 | - |
| Tbc1d9b | 597,14 | 0,0961 | 0,78476189 | chr11 | + |
| Tbcd | 440,97 | 0,1065 | 0,804480962 | chr11 | + |
| Tbkbp1 | 55,70 | 0,1139 | 0,843171378 | chr11 | - |
| Tbrg4 | 313,65 | -0,4624 | 0,093882396 | chr11 | - |
| Tbx2 | 22,62 | -0,2082 | 0,762940356 | chr11 | + |
| Tcf7 | 31,04 | -0,2799 | 0,637139802 | chr11 | - |
| Tcn2 | 1290,17 | -0,0170 | 0,972360847 | chr11 | - |
| Tefm | 269,92 | 0,5785 | 0,099827714 | chr11 | - |
| Ten1 | 75,80 | -0,0572 | 0,941004321 | chr11 | + |
| Tex14 | 2,11 | 0,1719 | NA | chr11 | + |
| Tex19.2 | 1,94 | 0,5470 | NA | chr11 | - |
| Tex2 | 1877,79 | -0,1354 | 0,636381742 | chr11 | - |
| Tgtp2 | 22,53 | 0,7920 | 0,119814602 | chr11 | - |
| Tha1 | 46,41 | -0,1636 | 0,802706588 | chr11 | - |
| Thg1l | 32,05 | 0,2056 | 0,775953072 | chr11 | - |
| Thoc5 | 202,88 | -0,0779 | 0,88569455 | chr11 | + |
| Timd2 | 4884,91 | -0,0769 | 0,854019408 | chr11 | - |
| Timd4 | 258,35 | -0,0496 | 0,930782422 | chr11 | + |
| Timm22 | 265,90 | -0,1794 | 0,627875496 | chr11 | + |
| Timp2 | 182,94 | 0,3072 | 0,402148509 | chr11 | - |
| Tk1 | 545,37 | -0,1801 | 0,677867285 | chr11 | - |
| Tlcd1 | 455,19 | -0,1598 | 0,770487872 | chr11 | + |
| Tlcd2 | 2307,70 | 0,2009 | 0,559428875 | chr11 | + |
| Tlk2 | 373,52 | -0,3208 | 0,208173201 | chr11 | + |
| Tmc6 | 39,76 | -0,5261 | 0,344375259 | chr11 | - |
| Tmc8 | 14,37 | -0,1512 | 0,855976995 | chr11 | + |
| Tmed4 | 1097,43 | -0,2053 | 0,52218039 | chr11 | - |
| Tmem100 | 9,32 | 0,4086 | 0,581394499 | chr11 | + |
| Tmem101 | 77,18 | 0,1034 | 0,872326129 | chr11 | - |
| Tmem102 | 96,28 | -0,1599 | 0,768686299 | chr11 | - |
| Tmem104 | 79,97 | -0,1954 | 0,727215824 | chr11 | + |
| Tmem106a | 158,30 | 0,0820 | 0,867747013 | chr11 | + |
| Tmem107 | 24,07 | 0,3161 | 0,611365326 | chr11 | + |
| Tmem11 | 394,43 | -0,1240 | 0,793250646 | chr11 | - |
| Tmem132e | 31,11 | -0,4501 | 0,370219404 | chr11 | + |
| Tmem17 | 12,69 | -0,0531 | 0,955504626 | chr11 | + |
| Tmem199 | 190,21 | -0,1777 | 0,656955579 | chr11 | - |
| Tmem220 | 367,92 | 0,1075 | 0,814156035 | chr11 | + |
| Tmem256 | 1976,15 | -0,4704 | 0,198624966 | chr11 | + |
| Tmem88 | 33,67 | 0,1606 | 0,825518458 | chr11 | - |
| Tmem97 | 2026,47 | -0,1841 | 0,610005889 | chr11 | - |
| Tmem98 | 78,67 | -0,3178 | 0,47939388 | chr11 | + |
| Tmigd1 | 1,58 | 0,4613 | NA | chr11 | + |
| Tmub2 | 355,04 | 0,0321 | 0,945274006 | chr11 | + |
| Tnfaip1 | 559,19 | -0,1694 | 0,593279907 | chr11 | - |
| Tnfsf12 | 69,79 | 0,5699 | 0,126353783 | chr11 | - |
| Tnfsf13 | 5,32 | 0,1637 | 0,852511194 | chr11 | - |

| | | | | | |
|---|---|---|---|---|---|
| Tnip1 | 128,18 | 0,5042 | 0,258301555 | chr11 | - |
| Tnrc6c | 164,82 | -0,0087 | 0,988512389 | chr11 | + |
| Tns3 | 625,30 | 0,3030 | 0,378100794 | chr11 | - |
| Tns4 | 11,63 | 0,2180 | 0,786277754 | chr11 | - |
| Tob1 | 1392,63 | 0,0630 | 0,910775679 | chr11 | + |
| Tom1l1 | 536,39 | -0,0852 | 0,860253707 | chr11 | - |
| Tom1l2 | 358,97 | -0,1972 | 0,644046286 | chr11 | - |
| Top2a | 55,61 | 0,4746 | 0,311319252 | chr11 | - |
| Top3a | 71,61 | -0,1519 | 0,783422709 | chr11 | - |
| Trappc1 | 1030,26 | -0,1225 | 0,764363417 | chr11 | + |
| Trim11 | 123,91 | -0,1312 | 0,862616891 | chr11 | + |
| Trim16 | 3,55 | 0,3540 | 0,627875496 | chr11 | + |
| Trim25 | 1450,75 | 0,1768 | 0,633578361 | chr11 | + |
| Trim37 | 224,80 | 0,0741 | 0,906843491 | chr11 | + |
| Trim41 | 480,47 | -0,1842 | 0,636381742 | chr11 | - |
| Trim47 | 51,75 | -0,6609 | 0,1598616 | chr11 | - |
| Trim65 | 22,80 | -0,4467 | 0,45239822 | chr11 | - |
| Trim7 | 47,41 | 0,1321 | 0,846704266 | chr11 | + |
| Trp53 | 176,31 | 0,2438 | 0,556131788 | chr11 | + |
| Trp53i13 | 10,16 | -0,0747 | 0,941553175 | chr11 | - |
| Trpv2 | 27,48 | 0,2526 | 0,672321306 | chr11 | + |
| Tsen54 | 64,28 | 0,4126 | 0,335637765 | chr11 | + |
| Tsr1 | 306,55 | -0,0493 | 0,929554961 | chr11 | + |
| Ttc1 | 551,73 | -0,2490 | 0,442032524 | chr11 | - |
| Ttc19 | 14,43 | 0,6806 | 0,282955582 | chr11 | + |
| Ttc25 | 22,07 | 0,2699 | 0,696074872 | chr11 | + |
| Ttyh2 | 843,44 | -0,2919 | 0,407448757 | chr11 | + |
| Tubd1 | 83,72 | -0,3299 | 0,528469822 | chr11 | + |
| Tug1 | 846,33 | 0,1362 | 0,731011847 | chr11 | - |
| Tusc5 | 6,63 | 0,2783 | 0,729883372 | chr11 | + |
| Tvp23b | 551,67 | 0,3406 | 0,299860814 | chr11 | + |
| Txndc17 | 1438,88 | -0,3857 | 0,135274149 | chr11 | + |
| Ubb | 832,01 | -0,2821 | 0,474794806 | chr11 | + |
| Ube2b | 2290,81 | -0,0765 | 0,848041213 | chr11 | - |
| Ube2g1 | 1113,69 | -0,1629 | 0,632136487 | chr11 | + |
| Ube2o | 67,39 | 0,0072 | 0,992023245 | chr11 | - |
| Ube2z | 806,20 | -0,1366 | 0,682099278 | chr11 | - |
| Ublcp1 | 524,75 | -0,1359 | 0,7588734 | chr11 | - |
| Ubtd2 | 55,69 | 0,3444 | 0,511143322 | chr11 | + |
| Ubtf | 599,08 | -0,2299 | 0,402592894 | chr11 | - |
| Ugp2 | 11006,21 | -0,4424 | 0,306882295 | chr11 | - |
| Ulk2 | 1534,59 | -0,2691 | 0,480021428 | chr11 | - |
| Unc119 | 86,54 | -0,2254 | 0,677762406 | chr11 | + |
| Unc13d | 11,72 | -0,0920 | 0,919467514 | chr11 | - |
| Unc45b | 19,83 | -0,2502 | 0,740022371 | chr11 | + |
| Unk | 57,76 | -0,0277 | 0,972909166 | chr11 | + |
| Uqcr10 | 3879,63 | -0,5454 | 0,217835326 | chr11 | - |
| Uqcrq | 6657,44 | -0,3888 | 0,603074159 | chr11 | - |
| Urgcp | 282,65 | -0,2407 | 0,505514525 | chr11 | - |
| Ush1g | 4,54 | -0,0312 | 0,977335689 | chr11 | - |

| | | | | | |
|---|---|---|---|---|---|
| Usp22 | 333,22 | 0,0321 | 0,943108135 | chr11 | - |
| Usp32 | 488,43 | 0,0426 | 0,922594928 | chr11 | - |
| Usp34 | 834,55 | -0,1270 | 0,705870256 | chr11 | + |
| Usp36 | 161,96 | -0,4810 | 0,144524949 | chr11 | - |
| Utp18 | 220,22 | -0,2458 | 0,566893799 | chr11 | - |
| Utp6 | 681,96 | 0,0690 | 0,888763259 | chr11 | - |
| Vamp2 | 179,55 | -0,3386 | 0,333257293 | chr11 | + |
| Vdac1 | 3791,45 | -0,2987 | 0,208577471 | chr11 | + |
| Vezf1 | 219,12 | -0,1394 | 0,740269415 | chr11 | + |
| Vmo1 | 144,24 | -0,5754 | 0,220525617 | chr11 | - |
| Vmp1 | 3143,73 | -0,4337 | 0,14743063 | chr11 | - |
| Vps25 | 992,75 | -0,3311 | 0,254577388 | chr11 | + |
| Vps53 | 340,95 | -0,1763 | 0,666442376 | chr11 | - |
| Vrk2 | 94,48 | 0,0207 | 0,974809304 | chr11 | - |
| Vtn | 36687,12 | -0,3417 | 0,240427136 | chr11 | + |
| Vwc2 | 2,12 | 0,7256 | NA | chr11 | + |
| Wbp2 | 1134,55 | -0,0806 | 0,862667917 | chr11 | - |
| Wdpcp | 68,98 | 0,2643 | 0,617467883 | chr11 | + |
| Wdr45b | 810,48 | 0,4106 | 0,187487434 | chr11 | - |
| Wdr81 | 371,96 | 0,0024 | 0,997464504 | chr11 | - |
| Wdr92 | 4658,14 | 0,1929 | 0,557396844 | chr11 | + |
| Wfdc17 | 877,04 | 0,3198 | 0,374589794 | chr11 | + |
| Wfdc18 | 2,78 | 0,2653 | 0,723157997 | chr11 | + |
| Wipf2 | 138,27 | -0,1562 | 0,74770519 | chr11 | + |
| Wipi1 | 231,64 | 0,0923 | 0,843171378 | chr11 | - |
| Wnk4 | 39,26 | -0,1619 | 0,844029674 | chr11 | + |
| Wnt9a | 11,04 | 0,0126 | 0,990189789 | chr11 | + |
| Wnt9b | 13,17 | -0,0953 | 0,916852736 | chr11 | - |
| Wrap53 | 92,78 | -0,2162 | 0,617767472 | chr11 | - |
| Wsb1 | 727,44 | 0,1571 | 0,776478455 | chr11 | - |
| Wscd1 | 3,84 | -0,1478 | 0,869544763 | chr11 | + |
| Wwc1 | 25,03 | 0,0521 | 0,950625589 | chr11 | - |
| Xaf1 | 684,04 | 0,8095 | 0,058286811 | chr11 | + |
| Xbp1 | 3275,00 | -0,1121 | 0,772705111 | chr11 | + |
| Xpo1 | 831,88 | 0,4848 | 0,086518104 | chr11 | + |
| Xylt2 | 186,07 | 0,0106 | 0,986661276 | chr11 | - |
| Ykt6 | 1414,16 | -0,1945 | 0,579355496 | chr11 | + |
| Ypel2 | 314,95 | -0,1119 | 0,841579427 | chr11 | - |
| Ywhae | 3320,68 | -0,0874 | 0,824806669 | chr11 | + |
| Zbtb4 | 336,21 | 0,0293 | 0,963400641 | chr11 | + |
| Zcchc10 | 70,93 | 0,0392 | 0,951733365 | chr11 | + |
| Zfp2 | 94,67 | -0,2121 | 0,636260727 | chr11 | - |
| Zfp207 | 1132,21 | 0,1281 | 0,800238939 | chr11 | + |
| Zfp287 | 4,83 | 0,5369 | 0,430392697 | chr11 | - |
| Zfp3 | 19,78 | 0,0616 | 0,943108135 | chr11 | + |
| Zfp354b | 33,58 | 0,1429 | 0,827353667 | chr11 | - |
| Zfp354c | 12,71 | 0,0160 | 0,987350308 | chr11 | - |
| Zfp39 | 58,61 | 0,0901 | 0,924834619 | chr11 | - |
| Zfp454 | 10,39 | -0,6401 | 0,323960783 | chr11 | - |
| Zfp62 | 345,06 | 0,5730 | 0,056273421 | chr11 | + |

| | | | | | |
|---|---|---|---|---|---|
| Zfp652 | 46,14 | 0,3723 | 0,464680168 | chr11 | + |
| Zfp652os | 2,26 | 0,1100 | NA | chr11 | - |
| Zfp672 | 174,04 | -0,0190 | 0,979051488 | chr11 | - |
| Zfp692 | 56,93 | -0,7370 | 0,101234862 | chr11 | + |
| Zfp750 | 151,62 | 0,5530 | 0,132625172 | chr11 | - |
| Zfp830 | 154,03 | -0,1465 | 0,722057512 | chr11 | + |
| Zfp867 | 112,80 | -0,0950 | 0,867698337 | chr11 | - |
| Zfp879 | 6,34 | 0,5679 | 0,398344449 | chr11 | - |
| Zkscan17 | 236,63 | -0,4194 | 0,147012425 | chr11 | - |
| Zkscan6 | 77,31 | 0,1061 | 0,847964947 | chr11 | + |
| Zmat5 | 111,45 | 0,1783 | 0,743138956 | chr11 | + |
| Zmiz2 | 182,43 | -0,3097 | 0,569246374 | chr11 | + |
| Zmynd15 | 40,79 | -0,1571 | 0,8044742 | chr11 | + |
| Znhit3 | 142,78 | -0,0142 | 0,985716259 | chr11 | - |
| Znrf3 | 134,25 | -0,3775 | 0,320413698 | chr11 | - |
| Zpbp | 2,86 | 0,4787 | 0,448454179 | chr11 | - |
| Zrsr1 | 288,80 | -0,1801 | 0,693158302 | chr11 | + |
| Zswim7 | 216,40 | 0,4454 | 0,293001163 | chr11 | - |
| Zzef1 | 351,74 | -0,1349 | 0,688905597 | chr11 | + |
| Erg28 | 2011,70 | 0,2313 | 0,537208005 | chr12 | - |
| 1110002L01Rik | 91,52 | -0,4130 | 0,316622965 | chr12 | - |
| 1110008L16Rik | 215,75 | -0,1141 | 0,809356379 | chr12 | + |
| Ldah | 1198,17 | -0,0609 | 0,88726005 | chr12 | + |
| 1700030C10Rik | 7,84 | 0,2008 | 0,814990243 | chr12 | - |
| 2010107E04Rik | 4139,68 | -0,3038 | 0,327028954 | chr12 | - |
| 2210039B01Rik | 30,22 | 0,0857 | 0,912811384 | chr12 | + |
| 2310015A10Rik | 31,38 | 0,2754 | 0,660827073 | chr12 | - |
| Riox1 | 65,01 | -0,3524 | 0,497114946 | chr12 | + |
| 2700049A03Rik | 40,17 | 0,2428 | 0,642393002 | chr12 | + |
| 2700097O09Rik | 80,15 | -0,1296 | 0,811367208 | chr12 | - |
| 2810029C07Rik | 3,82 | -0,0539 | 0,955504626 | chr12 | - |
| 3110056K07Rik | 32,14 | 0,0373 | 0,963400641 | chr12 | - |
| Tedc1 | 3,71 | 0,1754 | 0,838806042 | chr12 | + |
| Susd6 | 763,01 | 0,1876 | 0,592763574 | chr12 | + |
| 5033406O09Rik | 202,34 | 0,6219 | 0,132331753 | chr12 | - |
| Dglucy | 559,56 | 0,1075 | 0,840753129 | chr12 | + |
| 9330151L19Rik | 59,89 | -0,5723 | 0,220069886 | chr12 | + |
| Abcd4 | 123,43 | -0,1814 | 0,734571281 | chr12 | - |
| Acot1 | 250,92 | -0,3933 | 0,58887292 | chr12 | + |
| Acot2 | 78,26 | 0,2021 | 0,804214192 | chr12 | + |
| Acot3 | 402,90 | 0,2736 | 0,713100089 | chr12 | + |
| Acot4 | 329,53 | 0,3613 | 0,57157335 | chr12 | + |
| Acot6 | 7,88 | 0,4641 | 0,514753779 | chr12 | + |
| Acp1 | 2103,03 | -0,0117 | 0,981991128 | chr12 | - |
| Actn1 | 263,31 | 0,0704 | 0,88569455 | chr12 | - |
| Actr10 | 708,49 | 0,0328 | 0,943064069 | chr12 | + |
| Acyp1 | 211,50 | -0,2559 | 0,435658245 | chr12 | - |
| Adam17 | 497,46 | 0,1440 | 0,669280735 | chr12 | - |
| Adam4 | 5,43 | 0,0083 | 0,993727229 | chr12 | - |
| Adck1 | 202,56 | 0,0652 | 0,895663283 | chr12 | + |

| | | | | | |
|---|---|---|---|---|---|
| Adcy3 | 18,47 | 0,4085 | 0,526620417 | chr12 | + |
| Adi1 | 2305,27 | -0,2684 | 0,343230611 | chr12 | + |
| Adssl1 | 184,14 | 0,0072 | 0,990846949 | chr12 | + |
| Agmo | 1929,43 | 0,3037 | 0,464736787 | chr12 | + |
| Ahr | 329,47 | -0,4877 | 0,220525617 | chr12 | - |
| Ahsa1 | 1937,32 | -0,1096 | 0,802299336 | chr12 | + |
| AI463170 | 21,10 | -0,2777 | 0,71673612 | chr12 | - |
| Gon7 | 68,57 | -0,2189 | 0,680997554 | chr12 | - |
| Ak7 | 11,76 | 0,0072 | 0,994676823 | chr12 | + |
| Akap5 | 19,60 | -0,2198 | 0,774324297 | chr12 | + |
| Akap6 | 3,57 | 0,2404 | 0,766619151 | chr12 | + |
| Akt1 | 491,27 | -0,3495 | 0,374589794 | chr12 | - |
| Aldh6a1 | 7173,03 | 0,0337 | 0,944925159 | chr12 | - |
| Alkbh1 | 66,96 | -0,2090 | 0,663089738 | chr12 | - |
| Angel1 | 58,21 | 0,3944 | 0,368428658 | chr12 | - |
| Ankmy2 | 174,18 | 0,2702 | 0,579010212 | chr12 | + |
| Ankrd9 | 13,25 | -0,3838 | 0,59495419 | chr12 | - |
| Ap4s1 | 127,73 | -0,2149 | 0,648869552 | chr12 | + |
| Apob | 97754,23 | -0,4212 | 0,118153597 | chr12 | + |
| Apopt1 | 258,63 | -0,1335 | 0,783422709 | chr12 | + |
| Arel1 | 656,23 | 0,0646 | 0,89879469 | chr12 | - |
| Arf6 | 1224,20 | -0,1564 | 0,636260727 | chr12 | + |
| Arg2 | 4,36 | -0,4735 | 0,497745835 | chr12 | + |
| Arhgap5 | 644,16 | 0,5586 | 0,217175999 | chr12 | + |
| Arid4a | 210,37 | 0,0944 | 0,836737256 | chr12 | + |
| Arl4a | 630,36 | -0,4640 | 0,115749538 | chr12 | - |
| Asap2 | 50,10 | 0,5456 | 0,282672306 | chr12 | + |
| Asb2 | 13,89 | -0,1102 | 0,896860788 | chr12 | - |
| Aspg | 2534,31 | -0,0018 | 0,997464504 | chr12 | + |
| Asxl2 | 305,39 | 0,2091 | 0,636447281 | chr12 | + |
| Atad2b | 198,17 | 0,2074 | 0,588008527 | chr12 | + |
| Atg2b | 405,66 | -0,4220 | 0,196513229 | chr12 | - |
| Atl1 | 16,15 | 0,6116 | 0,318113166 | chr12 | + |
| Atp5s | 94,98 | 0,3202 | 0,492254415 | chr12 | + |
| Atp6v1d | 988,75 | -0,2691 | 0,438879622 | chr12 | - |
| Atxn3 | 347,40 | 0,1407 | 0,761282414 | chr12 | - |
| Atxn7l1 | 91,41 | -0,2686 | 0,508958442 | chr12 | + |
| B930059L03Rik | 7,93 | -0,5786 | 0,385049299 | chr12 | + |
| Bag5 | 249,63 | -0,2569 | 0,471443874 | chr12 | - |
| Batf | 9,79 | 0,3185 | 0,681592614 | chr12 | + |
| Baz1a | 236,29 | -0,1272 | 0,771142222 | chr12 | - |
| BC022687 | 8,30 | 0,9542 | 0,104667853 | chr12 | + |
| Bcap29 | 227,18 | 0,2153 | 0,676239605 | chr12 | - |
| Bcl11b | 3,66 | 0,1303 | 0,88569455 | chr12 | - |
| Brf1 | 120,09 | -0,3541 | 0,32697062 | chr12 | - |
| Brms1l | 71,52 | -0,1158 | 0,835059108 | chr12 | + |
| Btbd6 | 144,63 | -0,1414 | 0,764664236 | chr12 | + |
| Btbd7 | 66,59 | 0,6372 | 0,120869486 | chr12 | - |
| Bzw2 | 128,35 | -0,1851 | 0,754249138 | chr12 | - |
| Calm1 | 4328,74 | -0,0767 | 0,874515232 | chr12 | + |

| | | | | | |
|---|---|---|---|---|---|
| Cbll1 | 114,23 | 0,0518 | 0,924834619 | chr12 | - |
| Bbof1 | 10,48 | -0,3806 | 0,610022568 | chr12 | + |
| Ccdc71l | 102,92 | 0,4459 | 0,214058621 | chr12 | + |
| Ccdc85c | 8,29 | -0,1408 | 0,87618436 | chr12 | - |
| Ccdc88c | 10,34 | -0,0151 | 0,988414083 | chr12 | - |
| Ccnk | 203,06 | -0,2912 | 0,417585972 | chr12 | + |
| Cdc42bpb | 452,81 | -0,1567 | 0,751657602 | chr12 | - |
| Cdca4 | 63,87 | 0,1664 | 0,747375272 | chr12 | - |
| Cdca7l | 10,65 | 0,2186 | 0,78476189 | chr12 | + |
| Cdkl1 | 3,82 | 0,7439 | 0,217231457 | chr12 | - |
| Cenpo | 55,10 | 0,5277 | 0,214347061 | chr12 | - |
| Cep128 | 18,14 | 0,4162 | 0,539194011 | chr12 | - |
| Cep170b | 267,51 | -0,0398 | 0,953751362 | chr12 | + |
| Cfl2 | 1289,46 | -0,0442 | 0,927365453 | chr12 | - |
| Churc1 | 940,75 | 0,1232 | 0,757695155 | chr12 | + |
| Cinp | 226,45 | 0,0322 | 0,945127597 | chr12 | - |
| Cipc | 439,99 | -0,4441 | 0,131561943 | chr12 | + |
| Ckb | 55,68 | 0,1260 | 0,87093463 | chr12 | - |
| Clec14a | 309,32 | 0,0468 | 0,922379013 | chr12 | - |
| Clmn | 1723,80 | -0,3334 | 0,246698486 | chr12 | - |
| Cmpk2 | 99,29 | 0,6098 | 0,07787425 | chr12 | + |
| Cog5 | 133,26 | 0,5550 | 0,165182527 | chr12 | + |
| Colec11 | 684,35 | 0,2659 | 0,367954693 | chr12 | - |
| Coq6 | 218,46 | -0,1059 | 0,816325182 | chr12 | + |
| Cox16 | 649,00 | -0,5375 | 0,280066015 | chr12 | - |
| Cpsf2 | 476,89 | 0,0502 | 0,924363403 | chr12 | + |
| Cpsf3 | 596,74 | -0,0084 | 0,986997724 | chr12 | + |
| Crip1 | 158,97 | 0,4414 | 0,224150405 | chr12 | + |
| Crip2 | 459,65 | -0,1266 | 0,846816271 | chr12 | + |
| Mia2 | 3586,77 | -0,0558 | 0,913017292 | chr12 | + |
| Cys1 | 74,40 | -0,4708 | 0,271131834 | chr12 | - |
| D130020L05Rik | 163,97 | -0,6781 | 0,086587954 | chr12 | + |
| D430019H16Rik | 23,60 | 0,5612 | 0,318214275 | chr12 | + |
| Daam1 | 1045,84 | 0,2412 | 0,568858283 | chr12 | + |
| Dact1 | 15,96 | -0,6369 | 0,287543614 | chr12 | + |
| Dcaf5 | 449,30 | -0,3695 | 0,271455466 | chr12 | - |
| Dcdc2c | 33,42 | 0,4057 | 0,482787957 | chr12 | - |
| Ddx1 | 1486,81 | 0,1504 | 0,709716065 | chr12 | - |
| Ddx24 | 486,19 | 0,0069 | 0,990189789 | chr12 | - |
| Degs2 | 2,09 | -0,0685 | NA | chr12 | - |
| Dhrs7 | 639,59 | 0,5793 | 0,209104214 | chr12 | - |
| Dicer1 | 407,41 | -0,1991 | 0,561041153 | chr12 | - |
| Dld | 4072,24 | 0,2528 | 0,460946278 | chr12 | - |
| Dlst | 2975,37 | 0,0866 | 0,807278226 | chr12 | + |
| Dnaaf2 | 54,76 | 0,1901 | 0,74224943 | chr12 | - |
| Dnah11 | 2,99 | 0,0943 | 0,918617345 | chr12 | - |
| Dnajb9 | 785,81 | 0,1994 | 0,78102295 | chr12 | - |
| Dnajc27 | 10,77 | 0,3243 | 0,662339463 | chr12 | + |
| Dnal1 | 26,74 | 0,3699 | 0,522917242 | chr12 | + |
| Dnmt3a | 286,98 | 0,1687 | 0,621868755 | chr12 | + |

| | | | | | |
|---|---|---|---|---|---|
| Dock4 | 227,56 | 0,0587 | 0,923778914 | chr12 | + |
| Dtd2 | 232,92 | 0,4084 | 0,250526648 | chr12 | - |
| Dus4l | 64,56 | -0,0188 | 0,980379461 | chr12 | - |
| Dync1h1 | 1096,80 | -0,4894 | 0,100767966 | chr12 | + |
| E2f6 | 228,78 | -0,1487 | 0,692167613 | chr12 | + |
| Eapp | 525,93 | -0,1630 | 0,660945685 | chr12 | - |
| Efr3b | 10,00 | 0,1254 | 0,892347263 | chr12 | - |
| Egln3 | 83,78 | -0,4898 | 0,225575035 | chr12 | - |
| Eif2b2 | 233,74 | -0,2874 | 0,470637382 | chr12 | + |
| Eif2s1 | 1263,14 | -0,0294 | 0,954781536 | chr12 | + |
| Elmsan1 | 93,41 | -0,1656 | 0,743336656 | chr12 | - |
| Eml1 | 51,57 | 0,2224 | 0,670139713 | chr12 | + |
| Eml5 | 51,02 | -0,3622 | 0,468248856 | chr12 | - |
| Entpd5 | 3774,74 | -0,2481 | 0,505514525 | chr12 | - |
| Erh | 966,99 | -0,0665 | 0,872774688 | chr12 | - |
| Esr2 | 1,90 | 0,3636 | NA | chr12 | - |
| Esrrb | 5,27 | 0,9636 | 0,099247572 | chr12 | + |
| Esyt2 | 512,04 | 0,1043 | 0,804380135 | chr12 | + |
| Etv1 | 10,92 | 0,4019 | 0,572283367 | chr12 | + |
| Evl | 53,38 | 0,1890 | 0,760028388 | chr12 | + |
| Exd2 | 155,86 | 0,0542 | 0,91260409 | chr12 | + |
| Exoc3l4 | 23,54 | 0,9621 | 0,058032462 | chr12 | + |
| F730035M05Rik | 4,64 | 0,0587 | 0,952576171 | chr12 | + |
| Fam110c | 10,21 | 0,5904 | 0,376557226 | chr12 | + |
| Fam161b | 9,46 | -0,1839 | 0,830848779 | chr12 | - |
| Togaram1 | 225,63 | -0,0145 | 0,981025197 | chr12 | + |
| Fam228b | 7,98 | 0,2570 | 0,754953629 | chr12 | - |
| Fam49a | 83,75 | 0,5522 | 0,15607681 | chr12 | + |
| Fam84a | 12,77 | 0,0926 | 0,922379013 | chr12 | - |
| Fancm | 67,17 | 0,3214 | 0,472359592 | chr12 | + |
| Fbln5 | 106,93 | 0,1754 | 0,78755647 | chr12 | - |
| Fbxo33 | 293,90 | -0,1638 | 0,718541916 | chr12 | - |
| Fcf1 | 770,20 | -0,1710 | 0,636260727 | chr12 | + |
| Fkbp1b | 2,63 | 0,0008 | 0,998994678 | chr12 | - |
| Fkbp3 | 583,59 | -0,3381 | 0,368448147 | chr12 | - |
| Flrt2 | 9,17 | 0,7221 | 0,242678725 | chr12 | + |
| Fntb | 32,38 | 0,4197 | 0,458688593 | chr12 | + |
| Fos | 18,10 | 0,4205 | 0,549753626 | chr12 | + |
| Foxa1 | 205,87 | -0,4931 | 0,318491559 | chr12 | - |
| Foxn3 | 108,38 | -0,0432 | 0,948314012 | chr12 | - |
| Frmd6 | 71,59 | 0,3791 | 0,336411059 | chr12 | + |
| Fut8 | 109,10 | 0,3374 | 0,462380045 | chr12 | + |
| Galc | 83,29 | -0,2559 | 0,567666412 | chr12 | - |
| Gdap10 | 7,78 | -0,3096 | 0,685542591 | chr12 | + |
| Gemin2 | 329,49 | -0,2755 | 0,466932584 | chr12 | + |
| Gen1 | 12,74 | 0,4747 | 0,48822276 | chr12 | - |
| Glrx5 | 597,81 | -0,5247 | 0,095514195 | chr12 | + |
| Gm10433 | 18,75 | 0,2175 | 0,766052367 | chr12 | + |
| Gm10451 | 4,00 | 0,2716 | 0,718689474 | chr12 | + |
| Gm17746 | 3,80 | 0,6340 | 0,316622965 | chr12 | + |

| | | | | | |
|---|---|---|---|---|---|
| Gm17821 | 693,16 | 0,0779 | 0,841150543 | chr12 | - |
| Gm20187 | 3,46 | 0,2313 | 0,774324297 | chr12 | - |
| Gm4787 | 8,63 | 0,3456 | 0,648082042 | chr12 | - |
| Gm527 | 8,30 | -0,2032 | 0,806976166 | chr12 | + |
| Golga5 | 381,84 | -0,3689 | 0,22384344 | chr12 | + |
| Gpatch2l | 128,61 | 0,5275 | 0,177916821 | chr12 | + |
| Gphn | 1069,90 | 0,1317 | 0,716005335 | chr12 | + |
| Gpr132 | 5,14 | 0,6962 | 0,274683426 | chr12 | - |
| Gpr65 | 75,48 | 0,5920 | 0,136585635 | chr12 | + |
| Gpr68 | 3,00 | 0,2027 | 0,805867569 | chr12 | - |
| Gpx2 | 3,46 | 0,7018 | 0,227023515 | chr12 | - |
| Grhl1 | 12,86 | 0,2502 | 0,747576832 | chr12 | + |
| Gskip | 275,31 | 0,1973 | 0,723824089 | chr12 | + |
| Gstz1 | 19172,77 | -0,0104 | 0,98489528 | chr12 | + |
| Gtf2a1 | 539,29 | 0,4291 | 0,211349462 | chr12 | - |
| Hbp1 | 2523,30 | 0,1432 | 0,709458057 | chr12 | - |
| Hdac9 | 12,92 | 0,4227 | 0,552538285 | chr12 | - |
| Heatr5a | 212,78 | -0,2991 | 0,312022245 | chr12 | - |
| Hectd1 | 2579,93 | -0,0785 | 0,859139694 | chr12 | - |
| Hif1a | 649,25 | 0,1231 | 0,754524942 | chr12 | + |
| Hpcal1 | 104,82 | -0,2959 | 0,517082232 | chr12 | + |
| Hs1bp3 | 206,14 | 0,3450 | 0,345662712 | chr12 | + |
| Hsp90aa1 | 7992,07 | 0,4613 | 0,244548585 | chr12 | - |
| Iah1 | 2226,87 | 0,0614 | 0,890720786 | chr12 | + |
| Ifi27 | 3705,32 | 0,1401 | 0,653474875 | chr12 | + |
| Ifi27l2a | 428,83 | 0,0838 | 0,84862368 | chr12 | - |
| Ifrd1 | 292,63 | -0,4838 | 0,191230937 | chr12 | - |
| Ift43 | 39,14 | 0,3594 | 0,479068232 | chr12 | + |
| Immp2l | 759,74 | 0,1899 | 0,66209482 | chr12 | + |
| Inf2 | 261,18 | -0,1126 | 0,829023073 | chr12 | + |
| Irf2bpl | 219,10 | 0,2673 | 0,579028345 | chr12 | - |
| Isca2 | 535,83 | 0,0275 | 0,951372792 | chr12 | + |
| Ispd | 51,50 | 0,6008 | 0,162687791 | chr12 | + |
| Itgb1bp1 | 336,29 | -0,0428 | 0,941553259 | chr12 | - |
| Itpk1 | 174,60 | 0,1549 | 0,781281681 | chr12 | - |
| Itsn2 | 210,43 | 0,2729 | 0,480812921 | chr12 | + |
| Jag2 | 2,28 | 0,1727 | NA | chr12 | - |
| Jdp2 | 14,15 | 0,3215 | 0,652658393 | chr12 | + |
| Jkamp | 472,94 | -0,0179 | 0,971699327 | chr12 | + |
| Kcnk10 | 27,30 | 0,0386 | 0,969414042 | chr12 | - |
| Kcnk13 | 10,65 | 0,3839 | 0,587007634 | chr12 | + |
| Kcns3 | 2,66 | 0,1184 | 0,891868197 | chr12 | - |
| Kidins220 | 801,71 | 0,2753 | 0,332496065 | chr12 | + |
| Kif26a | 11,27 | -0,0532 | 0,955234619 | chr12 | + |
| Kif3c | 11,94 | -0,2284 | 0,769963557 | chr12 | + |
| Klc1 | 105,72 | 0,0115 | 0,986360395 | chr12 | + |
| Klf11 | 146,12 | 0,2915 | 0,620918751 | chr12 | + |
| Klhdc1 | 3,86 | 0,3171 | 0,680997554 | chr12 | + |
| Klhdc2 | 716,03 | 0,0165 | 0,975496727 | chr12 | + |
| Klhl28 | 18,48 | 0,2504 | 0,722057512 | chr12 | - |

| | | | | | |
|---|---|---|---|---|---|
| Klhl29 | 2,99 | -0,2279 | 0,770476204 | chr12 | - |
| L2hgdh | 968,67 | -0,2175 | 0,567046015 | chr12 | - |
| L3hypdh | 374,29 | -0,2683 | 0,494370662 | chr12 | - |
| Lamb1 | 140,12 | 0,2193 | 0,636361813 | chr12 | + |
| Laptm4a | 5199,81 | 0,0280 | 0,952989802 | chr12 | + |
| Lgmn | 1114,07 | -0,0115 | 0,980186108 | chr12 | - |
| Lin52 | 143,30 | -0,1997 | 0,63367203 | chr12 | + |
| Lrrn3 | 10,94 | -0,0883 | 0,925898178 | chr12 | - |
| Ltbp2 | 11,34 | -0,0056 | 0,995735598 | chr12 | - |
| Macc1 | 4,65 | 0,5672 | 0,385921204 | chr12 | + |
| Map4k5 | 362,33 | 0,0172 | 0,976528475 | chr12 | - |
| Mark3 | 513,08 | -0,4119 | 0,119171672 | chr12 | + |
| Max | 1045,90 | -0,2707 | 0,464377274 | chr12 | - |
| Mbip | 77,30 | 0,3601 | 0,427107372 | chr12 | - |
| Mboat2 | 2,54 | 0,1877 | NA | chr12 | + |
| Med6 | 284,57 | 0,0377 | 0,943305207 | chr12 | - |
| Meg3 | 9,38 | 0,1478 | 0,869687361 | chr12 | + |
| Mfsd2b | 17,75 | -0,7417 | 0,171871417 | chr12 | - |
| Mgat2 | 752,16 | 0,0326 | 0,950819236 | chr12 | + |
| Mia2 | 1017,71 | 0,5543 | 0,170839779 | chr12 | + |
| Mipol1 | 30,82 | -0,1403 | 0,854690386 | chr12 | + |
| Mis18bp1 | 7,49 | 0,4986 | 0,468682808 | chr12 | - |
| Mlh3 | 147,36 | 0,6069 | 0,120869486 | chr12 | - |
| Mnat1 | 179,19 | 0,4404 | 0,309263705 | chr12 | + |
| Mok | 3,40 | 0,3175 | 0,669958697 | chr12 | - |
| Mpp5 | 30,24 | -0,7226 | 0,177916821 | chr12 | + |
| Mta1 | 127,57 | -0,3383 | 0,483930201 | chr12 | + |
| Mthfd1 | 3040,69 | -0,0372 | 0,931322892 | chr12 | + |
| Nbas | 249,78 | 0,1060 | 0,830330662 | chr12 | + |
| Ncapg2 | 11,93 | 0,2301 | 0,771038929 | chr12 | + |
| Ncoa1 | 170,48 | -0,2708 | 0,416450614 | chr12 | - |
| Nek9 | 1009,86 | -0,0504 | 0,911892071 | chr12 | - |
| Nemf | 488,20 | 0,1461 | 0,678853818 | chr12 | - |
| Nfkbia | 596,33 | -0,4638 | 0,168698533 | chr12 | - |
| Ngb | 5,44 | 0,7765 | 0,213663916 | chr12 | - |
| Nin | 54,33 | 0,0477 | 0,943719859 | chr12 | - |
| Nkx2-1 | 8,15 | -0,4061 | 0,583782269 | chr12 | - |
| Nol10 | 72,06 | -0,0739 | 0,903337956 | chr12 | + |
| Noxred1 | 6,87 | -0,5204 | 0,449238443 | chr12 | - |
| Npc2 | 2005,61 | 0,3566 | 0,169162729 | chr12 | - |
| Nrde2 | 107,83 | 0,1794 | 0,665256143 | chr12 | - |
| Alkbh1 | 4,43 | -0,6470 | 0,316161394 | chr12 | - |
| Nrxn3 | 5,32 | -0,0265 | 0,981025197 | chr12 | + |
| Nubpl | 307,73 | 0,3919 | 0,295500275 | chr12 | + |
| Nudt14 | 165,73 | -0,4321 | 0,423813652 | chr12 | - |
| Numb | 552,07 | -0,0375 | 0,937853416 | chr12 | - |
| Odc1 | 1197,87 | -0,4111 | 0,230153033 | chr12 | + |
| Otub2 | 34,66 | -0,0148 | 0,987050646 | chr12 | + |
| Pacs2 | 182,98 | 0,0450 | 0,933203734 | chr12 | + |
| Papln | 11,27 | 0,6944 | 0,262601683 | chr12 | + |

| | | | | | |
|---|---|---|---|---|---|
| Papola | 1582,77 | -0,0367 | 0,918513859 | chr12 | + |
| Pcnx | 302,91 | -0,0280 | 0,950561821 | chr12 | + |
| Pcnx4 | 107,94 | -0,1049 | 0,816914587 | chr12 | + |
| Pdia6 | 9500,95 | 0,0837 | 0,855375466 | chr12 | + |
| Pfn4 | 4,80 | 0,6017 | 0,344366002 | chr12 | + |
| Pigh | 82,27 | 0,3640 | 0,385655922 | chr12 | - |
| Pik3cg | 21,31 | 0,0854 | 0,914198498 | chr12 | - |
| Pld4 | 141,13 | 0,2253 | 0,581394499 | chr12 | + |
| Plek2 | 11,54 | 0,1857 | 0,823828114 | chr12 | - |
| Plekhg3 | 291,02 | -0,0777 | 0,905274865 | chr12 | + |
| Plekhh1 | 6,08 | 0,6447 | 0,30944949 | chr12 | + |
| Pnn | 506,53 | -0,4036 | 0,156728357 | chr12 | + |
| Pnpla8 | 2630,67 | -0,0816 | 0,860835544 | chr12 | + |
| Pole2 | 17,30 | 0,0271 | 0,978910193 | chr12 | - |
| Pomc | 4,09 | 0,6045 | 0,331535469 | chr12 | + |
| Ppm1a | 1798,87 | -0,3905 | 0,212297691 | chr12 | + |
| Ppp1r13b | 228,12 | 0,4059 | 0,419539452 | chr12 | - |
| Ppp2r3c | 380,44 | 0,1428 | 0,764363417 | chr12 | - |
| Ppp2r5c | 717,75 | -0,3923 | 0,10553196 | chr12 | + |
| Ppp2r5e | 770,14 | -0,1563 | 0,688344487 | chr12 | - |
| Ppp4r4 | 90,88 | -0,2271 | 0,62632688 | chr12 | + |
| Pqlc3 | 13,38 | 0,8635 | 0,128835828 | chr12 | - |
| Prkar2b | 11,22 | 0,4020 | 0,581157554 | chr12 | - |
| Prkch | 63,22 | 0,2939 | 0,62632688 | chr12 | + |
| Prox2 | 10,04 | 0,3060 | 0,686977938 | chr12 | - |
| Prpf39 | 372,98 | -0,2601 | 0,537545925 | chr12 | + |
| Prps1l3 | 679,47 | 0,1375 | 0,761178054 | chr12 | + |
| Psen1 | 444,35 | -0,1356 | 0,738098909 | chr12 | + |
| Psma3 | 5039,54 | -0,1096 | 0,783418712 | chr12 | + |
| Psma6 | 2816,86 | -0,2382 | 0,413537476 | chr12 | + |
| Psmc1 | 2152,46 | -0,2801 | 0,368173309 | chr12 | + |
| Ptgr2 | 1656,46 | 0,0401 | 0,924680442 | chr12 | + |
| Ptpn21 | 160,04 | -0,0008 | 0,998482911 | chr12 | - |
| Ptprn2 | 5,99 | -0,2253 | 0,787663684 | chr12 | + |
| Ptrhd1 | 140,30 | -0,2368 | 0,587879802 | chr12 | + |
| Pum2 | 1091,08 | 0,0529 | 0,884712754 | chr12 | + |
| Pxdn | 129,10 | 0,1734 | 0,740269415 | chr12 | + |
| Pygl | 9325,61 | 0,0958 | 0,823949825 | chr12 | - |
| Rab10 | 1289,74 | -0,1582 | 0,589903004 | chr12 | - |
| Rab10os | 51,37 | -0,1845 | 0,754953629 | chr12 | + |
| Ralgapa1 | 147,92 | 0,1582 | 0,707094339 | chr12 | - |
| Rbm25 | 1110,95 | -0,3353 | 0,187852299 | chr12 | + |
| Rcor1 | 61,62 | 0,3045 | 0,568857865 | chr12 | + |
| Rdh11 | 800,33 | 0,4224 | 0,27946684 | chr12 | - |
| Rdh14 | 452,72 | -0,3426 | 0,253291899 | chr12 | + |
| Rhob | 212,41 | 0,1615 | 0,683658211 | chr12 | - |
| Rhoj | 15,59 | 0,1472 | 0,861772283 | chr12 | + |
| Rin3 | 143,01 | -0,4298 | 0,173484224 | chr12 | + |
| Rnaseh1 | 81,45 | 0,1466 | 0,774453704 | chr12 | + |
| Rnf113a2 | 135,94 | -0,4815 | 0,225581169 | chr12 | + |

| | | | | | |
|---|---|---|---|---|---|
| Rnf144a | 36,70 | 0,6350 | 0,157586062 | chr12 | - |
| Rock2 | 480,11 | 0,0784 | 0,871700795 | chr12 | + |
| Rpl36al | 6893,98 | -0,4836 | 0,069013539 | chr12 | - |
| Rps29 | 8555,52 | -0,4384 | 0,233901274 | chr12 | - |
| Rps6ka5 | 23,93 | -0,2515 | 0,690393892 | chr12 | - |
| Rps6kl1 | 16,28 | 1,0025 | 0,077346873 | chr12 | - |
| Rps7 | 7999,56 | -0,5013 | 0,141730165 | chr12 | - |
| Rrm2 | 59,93 | 0,1381 | 0,855245012 | chr12 | + |
| Rsad2 | 250,54 | 0,4963 | 0,133008375 | chr12 | - |
| Sav1 | 151,82 | -0,0303 | 0,95728901 | chr12 | - |
| Scfd1 | 1214,82 | 0,2443 | 0,534696593 | chr12 | + |
| Sec23a | 2025,53 | 0,1405 | 0,722057512 | chr12 | - |
| Sel1l | 1641,79 | -0,2312 | 0,510160643 | chr12 | - |
| Serpina10 | 9392,88 | -0,1027 | 0,872774688 | chr12 | - |
| Serpina11 | 2065,00 | 0,1349 | 0,834675238 | chr12 | - |
| Serpina1a | 36245,40 | -0,3171 | 0,279461687 | chr12 | - |
| Serpina1b | 85937,11 | -0,2298 | 0,420035183 | chr12 | - |
| Serpina1d | 49304,67 | -0,4064 | 0,152264308 | chr12 | - |
| Serpina1e | 132654,85 | -0,5600 | 0,242257968 | chr12 | - |
| Serpina3a | 7,27 | -0,3371 | 0,657129833 | chr12 | + |
| Serpina3b | 2,95 | -0,0501 | 0,958483224 | chr12 | + |
| Serpina3c | 12,11 | -0,8616 | 0,135521325 | chr12 | - |
| Serpina3f | 83,07 | 0,0914 | 0,860078935 | chr12 | + |
| Serpina3g | 270,24 | 0,3601 | 0,364571202 | chr12 | + |
| Serpina3h | 63,21 | 0,7039 | 0,239971296 | chr12 | + |
| Serpina3m | 6870,28 | 0,1632 | 0,747576832 | chr12 | + |
| Serpina3n | 8496,00 | 0,0179 | 0,981067951 | chr12 | + |
| Serpina4-ps1 | 2117,97 | 0,0749 | 0,939403124 | chr12 | + |
| Serpina5 | 6,12 | -0,0629 | 0,948314012 | chr12 | + |
| Serpina9 | 87,40 | 0,4464 | 0,490321719 | chr12 | - |
| Setd3 | 1538,32 | -0,2834 | 0,286763659 | chr12 | - |
| Sf3b6 | 1068,61 | -0,1130 | 0,825855713 | chr12 | + |
| Sgpp1 | 915,34 | -0,5378 | 0,063989705 | chr12 | - |
| Sh3yl1 | 126,29 | 0,3691 | 0,339404199 | chr12 | + |
| Sipa1l1 | 210,52 | -0,4052 | 0,208540118 | chr12 | + |
| Siva1 | 56,40 | -0,0626 | 0,939351122 | chr12 | + |
| Slc10a1 | 16112,52 | -0,4785 | 0,130822929 | chr12 | - |
| Slc25a21 | 176,11 | 0,0388 | 0,953751362 | chr12 | - |
| Slc25a29 | 15,88 | -0,0816 | 0,927608748 | chr12 | - |
| Slc25a47 | 11846,86 | -0,3036 | 0,581157554 | chr12 | + |
| Slc26a4 | 10,66 | 0,1349 | 0,88333112 | chr12 | - |
| Slc38a6 | 109,60 | -0,1726 | 0,701094188 | chr12 | + |
| Slc39a9 | 909,89 | 0,0026 | 0,995750189 | chr12 | + |
| Slc7a15 | 2,48 | 0,0343 | NA | chr12 | - |
| Slirp | 1117,63 | -0,1902 | 0,613685815 | chr12 | + |
| Smc6 | 494,83 | 0,0010 | 0,997623223 | chr12 | + |
| Ppp4r3a | 899,69 | -0,2712 | 0,417360803 | chr12 | - |
| Smoc1 | 1351,94 | -0,0484 | 0,931322892 | chr12 | + |
| Snapc1 | 99,61 | -0,0427 | 0,942571023 | chr12 | + |
| Snhg10 | 7,11 | -0,0591 | 0,952989802 | chr12 | - |

| | | | | | |
|---|---|---|---|---|---|
| Snora28 | 8,30 | 0,3747 | 0,614020796 | chr12 | + |
| Sntg2 | 341,04 | 0,3082 | 0,441888713 | chr12 | - |
| Snw1 | 1084,27 | 0,1266 | 0,761282414 | chr12 | - |
| Snx13 | 586,46 | 0,2137 | 0,517431746 | chr12 | + |
| Snx6 | 399,23 | 0,4108 | 0,173484224 | chr12 | - |
| Sos2 | 173,74 | 0,0403 | 0,941250497 | chr12 | - |
| Sp4 | 127,59 | 0,2207 | 0,642393002 | chr12 | - |
| Spata7 | 26,46 | 0,6028 | 0,271992655 | chr12 | + |
| Sptssa | 1364,54 | 0,3822 | 0,277831408 | chr12 | - |
| Srp54a | 97,36 | 0,3235 | 0,430134015 | chr12 | + |
| Srsf5 | 2155,75 | -0,2770 | 0,483930201 | chr12 | + |
| Strn3 | 580,65 | 0,1233 | 0,74146296 | chr12 | - |
| Stxbp6 | 249,77 | 0,3966 | 0,370887749 | chr12 | - |
| Syne2 | 303,14 | 0,0581 | 0,89879469 | chr12 | + |
| Syne3 | 86,35 | 0,6242 | 0,197796382 | chr12 | - |
| Synj2bp | 2103,07 | 0,0514 | 0,91371333 | chr12 | - |
| Sypl | 1523,30 | 0,1334 | 0,696074872 | chr12 | + |
| Tdp1 | 39,86 | 0,5062 | 0,279290314 | chr12 | + |
| Tecpr2 | 99,74 | -0,1551 | 0,786477935 | chr12 | + |
| Tgfb3 | 16,53 | 0,3986 | 0,543779254 | chr12 | - |
| Tmed10 | 322,97 | 0,1178 | 0,814156035 | chr12 | - |
| Tmed8 | 69,89 | 0,1754 | 0,743336656 | chr12 | - |
| Tmem18 | 356,70 | 0,1962 | 0,668257037 | chr12 | + |
| Tmem251 | 145,04 | -0,2698 | 0,510887696 | chr12 | + |
| Tmem30b | 495,35 | 0,4375 | 0,139399836 | chr12 | - |
| Tmem63c | 2,39 | 0,8196 | NA | chr12 | + |
| Tmx1 | 993,19 | 0,4430 | 0,171297118 | chr12 | + |
| Tnfaip2 | 265,45 | 0,4115 | 0,168369935 | chr12 | + |
| Traf3 | 83,62 | -0,0910 | 0,888917253 | chr12 | + |
| Trappc12 | 346,32 | -0,0083 | 0,988512389 | chr12 | - |
| Trappc6b | 570,74 | -0,2051 | 0,51143238 | chr12 | - |
| Trib2 | 31,85 | 0,3485 | 0,554660735 | chr12 | - |
| Trip11 | 100,19 | -0,3026 | 0,615826521 | chr12 | - |
| Trmt5 | 142,95 | 0,0287 | 0,965925759 | chr12 | - |
| Trmt61a | 61,40 | -0,1640 | 0,771616285 | chr12 | + |
| Tspan13 | 27,66 | -0,1723 | 0,801614342 | chr12 | - |
| Eipr1 | 145,84 | 0,0386 | 0,951898903 | chr12 | + |
| Ttc32 | 266,40 | -0,0155 | 0,98507181 | chr12 | + |
| Ttc8 | 66,91 | -0,1610 | 0,808947091 | chr12 | + |
| Ttll5 | 95,24 | 0,2293 | 0,575547607 | chr12 | + |
| Twist1 | 7,14 | -0,4486 | 0,532901541 | chr12 | + |
| Twistnb | 298,57 | -0,1099 | 0,804091647 | chr12 | + |
| Ubr7 | 321,91 | -0,0829 | 0,845013263 | chr12 | + |
| Ubxn2a | 366,60 | 0,0147 | 0,981067951 | chr12 | - |
| Unc79 | 9,80 | 0,2494 | 0,758057991 | chr12 | + |
| Vash1 | 60,20 | 0,5251 | 0,181404581 | chr12 | + |
| Vcpkmt | 92,09 | 0,0470 | 0,948314012 | chr12 | - |
| Vipas39 | 223,09 | 0,0119 | 0,984995389 | chr12 | - |
| Vrk1 | 125,51 | 0,1972 | 0,689884608 | chr12 | + |
| Vti1b | 745,92 | -0,4058 | 0,266518808 | chr12 | - |

| | | | | | |
|---|---|---|---|---|---|
| Wars | 239,00 | -0,0516 | 0,921903116 | chr12 | - |
| Wdr20 | 182,64 | 0,1108 | 0,793247794 | chr12 | + |
| Wdr25 | 52,29 | 0,2903 | 0,611914561 | chr12 | + |
| Wdr35 | 81,06 | 0,6127 | 0,143715786 | chr12 | + |
| Wdr60 | 51,37 | -0,1399 | 0,816025345 | chr12 | - |
| Wdr89 | 18,12 | 0,6045 | 0,342841028 | chr12 | - |
| Xrcc3 | 22,47 | -0,3886 | 0,505514525 | chr12 | - |
| Ylpm1 | 193,67 | -0,0198 | 0,976528475 | chr12 | + |
| Ywhaq | 1898,61 | -0,2027 | 0,52825088 | chr12 | - |
| Yy1 | 396,66 | -0,1773 | 0,677398032 | chr12 | + |
| Zbtb1 | 163,62 | 0,0498 | 0,935938583 | chr12 | + |
| Zbtb25 | 25,43 | 0,0599 | 0,943108135 | chr12 | - |
| Zbtb42 | 60,46 | -0,6458 | 0,164406774 | chr12 | + |
| Zc2hc1c | 34,46 | -0,1527 | 0,809790857 | chr12 | + |
| Zc3h14 | 969,13 | -0,3408 | 0,212297691 | chr12 | + |
| Zfp277 | 366,07 | -0,2746 | 0,399950473 | chr12 | - |
| Zfp36l1 | 1711,10 | 0,2845 | 0,546240378 | chr12 | - |
| Zfp386 | 267,24 | 0,5883 | 0,160369993 | chr12 | + |
| Zfp410 | 331,30 | -0,1896 | 0,603818311 | chr12 | + |
| Zfp839 | 103,51 | 0,0808 | 0,888839251 | chr12 | + |
| Zfyve1 | 852,95 | -0,0718 | 0,904806119 | chr12 | - |
| Zfyve21 | 487,78 | -0,0443 | 0,948356 | chr12 | + |
| Zfyve26 | 334,03 | -0,0773 | 0,8636022 | chr12 | - |
| Card19 | 85,28 | -0,2626 | 0,68001254 | chr13 | - |
| 1110046J04Rik | 9,91 | -0,3316 | 0,653280861 | chr13 | + |
| 1810034E14Rik | 130,23 | -0,3381 | 0,414126925 | chr13 | + |
| 2010111I01Rik | 534,17 | -0,5210 | 0,068187551 | chr13 | + |
| 2210016F16Rik | 509,19 | 0,3713 | 0,186036343 | chr13 | - |
| 2210408I21Rik | 51,77 | 0,6131 | 0,200225987 | chr13 | + |
| 2610307P16Rik | 2,42 | 0,6771 | NA | chr13 | - |
| 3110070M22Rik | 4,76 | 1,0763 | 0,052145393 | chr13 | - |
| 4833420G17Rik | 332,05 | -0,2732 | 0,578442407 | chr13 | + |
| 4833422C13Rik | 19,25 | -0,3523 | 0,611887546 | chr13 | - |
| 4833439L19Rik | 923,14 | -0,4019 | 0,208518411 | chr13 | - |
| 4930486L24Rik | 9,97 | -0,1325 | 0,88306163 | chr13 | - |
| 4930525G20Rik | 8,57 | 0,6575 | 0,300188675 | chr13 | - |
| 4930528D03Rik | 2,77 | 0,0082 | 0,993487304 | chr13 | + |
| 4933433G19Rik | 2,86 | -0,8130 | 0,147367695 | chr13 | - |
| 6430562O15Rik | 2,68 | -0,0918 | 0,917042614 | chr13 | - |
| 9330162012Rik | 33,68 | 0,0799 | 0,913102981 | chr13 | + |
| 9430083A17Rik | 7,86 | -0,1000 | 0,915977656 | chr13 | - |
| Zfp729a | 90,68 | 0,5994 | 0,128136319 | chr13 | - |
| A530099J19Rik | 3,19 | 0,2339 | 0,76908367 | chr13 | - |
| A630019I02Rik | 2,20 | 0,3380 | NA | chr13 | + |
| A830082K12Rik | 6,70 | -0,4459 | 0,536622488 | chr13 | + |
| Aaed1 | 349,13 | 0,2261 | 0,602968232 | chr13 | - |
| Abt1 | 165,57 | -0,0535 | 0,922855195 | chr13 | - |
| Acot12 | 1851,05 | -0,2210 | 0,572283367 | chr13 | + |
| Acot13 | 1448,21 | -0,2346 | 0,614056134 | chr13 | - |
| Adarb2 | 5,17 | -0,1561 | 0,862616891 | chr13 | + |

| | | | | | |
|---|---|---|---|---|---|
| Adcy2 | 4,51 | -0,0323 | 0,97669908 | chr13 | - |
| Adtrp | 3909,70 | 0,1203 | 0,81172751 | chr13 | - |
| Aggf1 | 544,99 | 0,0498 | 0,922379013 | chr13 | - |
| Agtpbp1 | 133,82 | -0,6804 | 0,05009592 | chr13 | - |
| Agtr1a | 786,92 | 0,2844 | 0,286737425 | chr13 | + |
| Ahrr | 2,10 | 0,0741 | NA | chr13 | - |
| AI197445 | 2,63 | 0,0778 | 0,932839392 | chr13 | + |
| Akr1c14 | 9869,72 | -0,5440 | 0,204823354 | chr13 | + |
| Akr1c18 | 2,67 | -0,1596 | 0,848140004 | chr13 | - |
| Akr1c20 | 1824,37 | -0,6015 | 0,078201192 | chr13 | - |
| Akr1c21 | 4,73 | -0,9581 | 0,07965206 | chr13 | + |
| Akr1c6 | 39964,45 | -0,2968 | 0,480021428 | chr13 | + |
| Akr1e1 | 1178,10 | -0,2770 | 0,470730504 | chr13 | - |
| Aldh5a1 | 970,73 | 0,0333 | 0,938364545 | chr13 | - |
| Ankra2 | 173,37 | -0,0292 | 0,953751362 | chr13 | + |
| Slf1 | 85,75 | 0,6916 | 0,089333342 | chr13 | - |
| Ankrd55 | 5,01 | 0,6432 | 0,321106759 | chr13 | + |
| Aoah | 40,71 | 0,2777 | 0,60905895 | chr13 | + |
| Ap3b1 | 690,48 | 0,0741 | 0,838868977 | chr13 | + |
| Arid4b | 297,27 | 0,2117 | 0,614390354 | chr13 | + |
| Arl10 | 16,12 | 0,1265 | 0,877920448 | chr13 | + |
| Arl15 | 290,06 | 0,6193 | 0,075686715 | chr13 | + |
| Arrdc3 | 1557,42 | -0,4559 | 0,522848151 | chr13 | + |
| Arsb | 376,27 | 0,0936 | 0,847419446 | chr13 | + |
| Arsk | 160,26 | 0,3515 | 0,349661709 | chr13 | - |
| Asb13 | 727,15 | -0,0175 | 0,971522662 | chr13 | + |
| Aspn | 4,93 | 0,6326 | 0,332668588 | chr13 | + |
| Atg10 | 121,62 | 0,3225 | 0,357345082 | chr13 | - |
| Atxn1 | 435,63 | 0,0757 | 0,92861245 | chr13 | - |
| Auh | 325,00 | -0,4800 | 0,053880988 | chr13 | - |
| AW209491 | 328,11 | 0,0425 | 0,933514578 | chr13 | + |
| AW495222 | 3,68 | -0,3706 | 0,613341495 | chr13 | - |
| B230219D22Rik | 582,93 | 0,0225 | 0,969048579 | chr13 | + |
| B3galnt2 | 361,59 | -0,3651 | 0,325839875 | chr13 | + |
| B4galt7 | 87,97 | -0,2448 | 0,692272065 | chr13 | + |
| BC005537 | 1671,40 | -0,2451 | 0,356860747 | chr13 | + |
| Ice1 | 199,94 | 0,0548 | 0,911892071 | chr13 | - |
| BC147527 | 5,42 | 0,5797 | 0,369664628 | chr13 | + |
| Bdp1 | 242,84 | 0,0194 | 0,969159464 | chr13 | - |
| Bhmt2 | 3956,54 | 0,3148 | 0,374761697 | chr13 | - |
| Bicd2 | 239,74 | -0,2265 | 0,589903004 | chr13 | + |
| Bloc1s5 | 329,53 | 0,3621 | 0,182985049 | chr13 | - |
| Bmp6 | 37,69 | 0,3848 | 0,498227709 | chr13 | + |
| Bphl | 2343,96 | -0,1714 | 0,607310966 | chr13 | + |
| Brd9 | 176,32 | -0,2907 | 0,37740141 | chr13 | + |
| Btf3 | 2793,19 | -0,2853 | 0,253392997 | chr13 | - |
| C230035I16Rik | 3,36 | 0,2826 | 0,713614788 | chr13 | - |
| Fam8a1 | 148,13 | 0,4901 | 0,172087384 | chr13 | + |
| Cage1 | 7,77 | -0,2325 | 0,77991551 | chr13 | - |
| Caml | 224,94 | -0,3847 | 0,295398429 | chr13 | + |

| | | | | | |
|---|---|---|---|---|---|
| Cap2 | 2,07 | 0,3192 | NA | chr13 | + |
| Cast | 554,06 | 0,2622 | 0,58308892 | chr13 | - |
| Catsper3 | 2,40 | -0,4588 | NA | chr13 | + |
| Ccdc125 | 111,47 | 0,3958 | 0,293336152 | chr13 | + |
| Ccdc127 | 419,09 | -0,1605 | 0,59511312 | chr13 | + |
| Ccl28 | 2,33 | -0,1719 | NA | chr13 | + |
| Ccnb1 | 20,33 | 0,4768 | 0,456715032 | chr13 | - |
| Ccnh | 317,79 | 0,0145 | 0,978402469 | chr13 | + |
| Ccno | 5,87 | 0,1234 | 0,89468939 | chr13 | + |
| Cd180 | 30,65 | 0,8877 | 0,068216978 | chr13 | + |
| Cd83 | 48,44 | 0,2558 | 0,688193661 | chr13 | + |
| Cdc14b | 548,01 | 0,0066 | 0,990846949 | chr13 | - |
| Cdhr2 | 4,42 | 0,5009 | 0,45172307 | chr13 | + |
| Cdk13 | 318,39 | -0,0569 | 0,906998011 | chr13 | - |
| Cdk20 | 91,10 | 0,4706 | 0,286763659 | chr13 | + |
| Cdk7 | 393,68 | 0,1677 | 0,680997554 | chr13 | - |
| Cdkal1 | 240,99 | 0,1409 | 0,764664236 | chr13 | - |
| Cdyl | 54,04 | -0,5312 | 0,178097604 | chr13 | + |
| Cenph | 4,57 | 0,3786 | 0,612916595 | chr13 | - |
| Cenpk | 20,77 | -0,4498 | 0,4891753 | chr13 | + |
| Cenpp | 5,80 | 0,3426 | 0,656332932 | chr13 | - |
| Cep72 | 13,43 | -0,1673 | 0,835640396 | chr13 | - |
| Cetn3 | 920,38 | 0,1092 | 0,842035655 | chr13 | + |
| Chrm3 | 2,47 | -0,5021 | NA | chr13 | - |
| Cks2 | 528,43 | 0,0873 | 0,842320388 | chr13 | + |
| Clptm1l | 1177,24 | -0,2211 | 0,524169625 | chr13 | + |
| Cltb | 476,34 | -0,7185 | 0,081343641 | chr13 | - |
| Cmah | 2751,98 | 0,1216 | 0,741224458 | chr13 | + |
| Cmya5 | 2,06 | 0,7632 | NA | chr13 | - |
| Col4a3bp | 747,66 | 0,2167 | 0,540815736 | chr13 | + |
| Cplx2 | 21,98 | 0,0884 | 0,909774279 | chr13 | + |
| Ctla2a | 158,18 | 0,3527 | 0,440094295 | chr13 | - |
| Ctla2b | 134,27 | -0,0370 | 0,953638766 | chr13 | - |
| Ctsl | 13790,11 | 0,0854 | 0,851472884 | chr13 | - |
| Cwc27 | 148,44 | 0,0881 | 0,887119817 | chr13 | - |
| D130043K22Rik | 84,97 | 0,1471 | 0,805831145 | chr13 | + |
| Dapk1 | 544,82 | 0,3344 | 0,394187455 | chr13 | + |
| Dbn1 | 15,52 | 0,2370 | 0,74829088 | chr13 | - |
| Dcdc2a | 42,79 | 0,7312 | 0,132461478 | chr13 | + |
| Ddx41 | 428,77 | -0,2696 | 0,385752804 | chr13 | - |
| Ddx46 | 612,18 | -0,0560 | 0,891236226 | chr13 | + |
| Dek | 911,49 | -0,0564 | 0,900327088 | chr13 | - |
| Dhfr | 1449,94 | 0,0868 | 0,85502909 | chr13 | + |
| Dhx29 | 166,43 | 0,2380 | 0,608025657 | chr13 | + |
| Dimt1 | 67,94 | 0,4020 | 0,355918478 | chr13 | + |
| Dip2c | 245,00 | -0,1578 | 0,657774838 | chr13 | + |
| Dmgdh | 7158,45 | 0,1792 | 0,58497677 | chr13 | + |
| Dok3 | 16,86 | -0,4823 | 0,473519111 | chr13 | - |
| Dsp | 596,68 | 0,3175 | 0,28060505 | chr13 | + |
| Dusp22 | 273,45 | -0,1785 | 0,611941622 | chr13 | + |

| | | | | | |
|---|---|---|---|---|---|
| E2f3 | 102,42 | -0,6635 | 0,069200865 | chr13 | - |
| Eci2 | 4932,37 | -0,5521 | 0,079955924 | chr13 | - |
| Eci3 | 5,64 | 0,3994 | 0,590951254 | chr13 | - |
| Ecm2 | 28,00 | 0,2591 | 0,704347771 | chr13 | + |
| Edn1 | 7,98 | -0,2033 | 0,812820227 | chr13 | + |
| Eef1e1 | 222,26 | 0,1865 | 0,640705696 | chr13 | - |
| Ell2 | 1125,94 | -0,4017 | 0,226610828 | chr13 | + |
| Elmo1 | 25,01 | -0,0051 | 0,995660674 | chr13 | + |
| Elovl2 | 9579,34 | -0,3540 | 0,300363865 | chr13 | - |
| Elovl7 | 17,17 | 0,5647 | 0,365500962 | chr13 | + |
| Emb | 17,61 | 0,7352 | 0,217231457 | chr13 | + |
| Enc1 | 108,93 | 0,6121 | 0,065752252 | chr13 | + |
| Epdr1 | 15,15 | 0,8123 | 0,158489444 | chr13 | - |
| Erap1 | 1801,41 | 0,5803 | 0,074073488 | chr13 | + |
| Erbin | 1133,56 | -0,1466 | 0,654788851 | chr13 | - |
| Ercc6l2 | 216,66 | 0,2071 | 0,633566893 | chr13 | + |
| Ercc8 | 53,19 | 0,1086 | 0,860890261 | chr13 | + |
| Ero1lb | 1510,23 | -0,4687 | 0,140278812 | chr13 | + |
| Esm1 | 30,65 | 0,2711 | 0,703924654 | chr13 | + |
| Etohd2 | 12,85 | -0,6411 | 0,273713146 | chr13 | + |
| Exoc2 | 161,12 | 0,4201 | 0,319080738 | chr13 | - |
| Exoc3 | 508,38 | 0,0830 | 0,869687361 | chr13 | - |
| F12 | 6723,44 | -0,1051 | 0,842925053 | chr13 | - |
| F13a1 | 8,58 | 0,7316 | 0,246159206 | chr13 | - |
| F2r | 666,26 | 0,2390 | 0,549054641 | chr13 | - |
| F2rl1 | 7,11 | 0,7926 | 0,201324217 | chr13 | - |
| Faf2 | 746,17 | 0,0543 | 0,885187516 | chr13 | + |
| Fam120a | 1721,24 | -0,2180 | 0,432005857 | chr13 | - |
| Fam120aos | 125,98 | -0,0596 | 0,909703206 | chr13 | + |
| Fam151b | 95,03 | 0,2442 | 0,565929347 | chr13 | - |
| Fam172a | 198,06 | 0,2737 | 0,43326072 | chr13 | + |
| Fam193b | 367,36 | -0,5971 | 0,228930646 | chr13 | - |
| Ripor2 | 38,52 | 0,3232 | 0,663082496 | chr13 | + |
| Fancc | 63,31 | -0,0541 | 0,933514578 | chr13 | - |
| Fastkd3 | 97,86 | 0,1240 | 0,818405803 | chr13 | + |
| Fbp1 | 17129,87 | -0,2485 | 0,527966188 | chr13 | - |
| Fbp2 | 4,65 | 0,4873 | 0,463892619 | chr13 | - |
| Fbxl21 | 21,26 | 0,1738 | 0,811801336 | chr13 | + |
| Fbxw17 | 17,94 | 0,6159 | 0,284930153 | chr13 | + |
| Fcho2 | 508,53 | 0,4009 | 0,218565307 | chr13 | - |
| Fgd3 | 7,81 | 0,7106 | 0,258061915 | chr13 | - |
| Fgfr4 | 300,19 | -0,0344 | 0,957017397 | chr13 | + |
| Fst | 84,99 | -0,3035 | 0,606237187 | chr13 | - |
| Gas1 | 177,53 | -0,0090 | 0,987952599 | chr13 | - |
| Gcnt2 | 579,85 | 0,1547 | 0,686854707 | chr13 | + |
| Gdi2 | 5083,95 | -0,2350 | 0,372878192 | chr13 | + |
| Gfm2 | 601,89 | -0,1174 | 0,764363417 | chr13 | + |
| Gfod1 | 29,31 | 0,4666 | 0,408167016 | chr13 | - |
| Ggps1 | 164,79 | 0,3484 | 0,286640164 | chr13 | - |
| Gkap1 | 317,77 | -0,1396 | 0,732140841 | chr13 | - |

| | | | | | |
|---|---|---|---|---|---|
| Gli3 | 8,08 | 0,2521 | 0,756319626 | chr13 | + |
| Gm10336 | 134,37 | 0,4368 | 0,263823735 | chr13 | - |
| Gm10767 | 17,57 | -0,3168 | 0,62683519 | chr13 | + |
| Gm5086 | 3,54 | 0,3966 | 0,571942598 | chr13 | + |
| Gm5141 | 12,23 | 1,0739 | 0,053014128 | chr13 | - |
| Tmem267 | 148,18 | -0,1547 | 0,820554805 | chr13 | + |
| Gm904 | 1,90 | 0,6937 | NA | chr13 | + |
| Gm9776 | 8,24 | -0,1338 | 0,881809902 | chr13 | - |
| Gmds | 45,60 | -0,6230 | 0,24579162 | chr13 | - |
| Gmpr | 13,42 | 0,1999 | 0,808526009 | chr13 | + |
| Gpbp1 | 1314,81 | 0,2038 | 0,581990132 | chr13 | - |
| Gpld1 | 4113,48 | 0,0799 | 0,804380135 | chr13 | + |
| Gpr137b | 10,60 | 0,0962 | 0,917470917 | chr13 | - |
| Gpr137b-ps | 11,22 | -0,0194 | 0,985628536 | chr13 | - |
| Gpr141 | 3,16 | 0,9245 | 0,098598649 | chr13 | - |
| Gpx6 | 25,88 | 0,6581 | 0,202636202 | chr13 | + |
| Gpx8 | 38,59 | 0,1561 | 0,842655851 | chr13 | - |
| Grk6 | 179,82 | -0,2271 | 0,522014539 | chr13 | + |
| Gtf2h2 | 201,84 | 0,3166 | 0,376078101 | chr13 | - |
| Gtpbp4 | 760,49 | 0,0003 | 0,9993428 | chr13 | - |
| Gzma | 18,63 | 0,3350 | 0,632226809 | chr13 | - |
| H2afy | 868,74 | -0,1582 | 0,588219472 | chr13 | - |
| Habp4 | 805,35 | -0,5457 | 0,050536418 | chr13 | + |
| Hapln1 | 15,95 | 0,8279 | 0,151731441 | chr13 | + |
| Heatr1 | 362,65 | 0,2706 | 0,585752237 | chr13 | + |
| Hfe | 777,61 | -0,4276 | 0,109794272 | chr13 | - |
| Mfsd14b | 725,65 | 0,2026 | 0,625710581 | chr13 | - |
| Higd2a | 1351,03 | -0,2479 | 0,422960414 | chr13 | + |
| Hist1h1c | 72,91 | -0,0437 | 0,95463264 | chr13 | + |
| Hist1h1e | 2,20 | 0,2111 | NA | chr13 | - |
| Hist1h2ba | 4,54 | 0,1209 | NA | chr13 | - |
| Hist1h2be | 4,93 | 0,2275 | 0,786009002 | chr13 | - |
| Hist1h4h | 13,00 | 0,0745 | 0,940538622 | chr13 | + |
| Hist1h4i | 8,36 | 0,1993 | 0,816133189 | chr13 | - |
| Hivep1 | 87,75 | 0,0355 | 0,953911274 | chr13 | + |
| Hk3 | 54,18 | -0,1771 | 0,754953629 | chr13 | - |
| Hmgcr | 1999,74 | -0,0017 | 0,997549615 | chr13 | - |
| Hmgcs1 | 3251,75 | 0,3284 | 0,596014309 | chr13 | + |
| Hnrnpa0 | 620,60 | 0,1119 | 0,766619151 | chr13 | - |
| Hnrnpk | 3654,19 | 0,1150 | 0,714323789 | chr13 | - |
| Homer1 | 117,32 | 0,2144 | 0,639465027 | chr13 | + |
| Iars | 530,06 | -0,3496 | 0,278686919 | chr13 | + |
| Id4 | 11,63 | 0,2018 | 0,804418611 | chr13 | + |
| Idi1 | 4320,92 | -0,2400 | 0,582449728 | chr13 | + |
| Idnk | 516,46 | 0,1224 | 0,771736977 | chr13 | + |
| Il6st | 1395,62 | 0,2114 | 0,449196578 | chr13 | + |
| Ipo11 | 319,48 | 0,3372 | 0,419095885 | chr13 | - |
| Ippk | 158,60 | -0,2886 | 0,492254415 | chr13 | + |
| Iqgap2 | 4047,86 | -0,1926 | 0,583654143 | chr13 | - |
| Irf4 | 4,63 | 0,6820 | 0,264762084 | chr13 | + |

| | | | | | |
|---|---|---|---|---|---|
| Isca1 | 1236,21 | -0,3737 | 0,248525122 | chr13 | - |
| Itga1 | 333,20 | 0,3974 | 0,180060786 | chr13 | - |
| Itga2 | 2,60 | -0,0069 | 0,994384173 | chr13 | - |
| Jarid2 | 168,89 | -0,2160 | 0,633248858 | chr13 | + |
| Kdm1b | 415,86 | 0,4265 | 0,149712396 | chr13 | + |
| Kif13a | 164,26 | -0,0593 | 0,926353842 | chr13 | - |
| Kif2a | 160,66 | 0,3638 | 0,24460589 | chr13 | - |
| Klhl3 | 24,09 | 0,4032 | 0,501991823 | chr13 | - |
| Larp4b | 587,15 | 0,0706 | 0,866934682 | chr13 | + |
| Lect2 | 4141,14 | 0,3325 | 0,383568565 | chr13 | - |
| Lgals8 | 2956,53 | 0,4215 | 0,254608455 | chr13 | - |
| Lhfpl2 | 204,79 | 0,3266 | 0,413323375 | chr13 | + |
| Lman2 | 2950,40 | -0,2753 | 0,365123284 | chr13 | - |
| Lrrc14b | 7,83 | -0,5853 | 0,380878258 | chr13 | - |
| Carmil1 | 217,27 | -0,4298 | 0,294160783 | chr13 | - |
| Ly86 | 212,19 | 0,1642 | 0,746827664 | chr13 | + |
| Lysmd3 | 341,96 | 0,5558 | 0,217835326 | chr13 | + |
| Lyst | 176,37 | 0,2331 | 0,713614788 | chr13 | + |
| Map3k1 | 165,86 | 0,1675 | 0,701666523 | chr13 | - |
| Marveld2 | 175,19 | 0,4097 | 0,293404423 | chr13 | - |
| Mast4 | 25,84 | 0,1968 | 0,797559244 | chr13 | - |
| Mblac2 | 138,27 | 0,0147 | 0,98429278 | chr13 | + |
| Mboat1 | 6,33 | 0,6333 | 0,3349919 | chr13 | + |
| Mccc2 | 1146,48 | 0,0578 | 0,880810135 | chr13 | - |
| Mctp1 | 1,89 | 0,3828 | NA | chr13 | + |
| Mcur1 | 541,13 | -0,3344 | 0,280519509 | chr13 | - |
| Med10 | 234,36 | -0,0874 | 0,851988359 | chr13 | + |
| Mef2c | 61,84 | -0,0154 | 0,983887477 | chr13 | + |
| Mier3 | 255,58 | 0,1913 | 0,61772121 | chr13 | + |
| Mirlet7d | 4,69 | -0,5105 | 0,46093503 | chr13 | - |
| Mocs2 | 3040,34 | -0,2575 | 0,394399177 | chr13 | + |
| Mplkip | 125,63 | -0,1504 | 0,765219833 | chr13 | + |
| Mrpl32 | 614,42 | -0,0825 | 0,849664617 | chr13 | - |
| Mrpl36 | 394,75 | -0,2784 | 0,526620417 | chr13 | + |
| Mrps27 | 482,65 | 0,0717 | 0,887468345 | chr13 | + |
| Mrps30 | 315,19 | -0,2449 | 0,565966999 | chr13 | - |
| Mrps36 | 303,20 | -0,3302 | 0,256223204 | chr13 | - |
| Mrs2 | 88,40 | -0,1897 | 0,73161502 | chr13 | - |
| Msh3 | 223,75 | -0,4703 | 0,19521384 | chr13 | - |
| Mterf3 | 382,45 | -0,2856 | 0,352106106 | chr13 | - |
| Mtrr | 100,44 | 0,2647 | 0,554974606 | chr13 | - |
| Mtx3 | 279,70 | 0,2273 | 0,582441358 | chr13 | + |
| Mxd3 | 3,72 | 0,2195 | 0,77892794 | chr13 | - |
| Mylip | 190,18 | -0,0496 | 0,914198498 | chr13 | + |
| Mylk4 | 28,28 | -0,6426 | 0,197447777 | chr13 | - |
| Naa35 | 558,86 | -0,1189 | 0,77892794 | chr13 | + |
| Naip1 | 10,58 | -0,3324 | 0,66477796 | chr13 | - |
| Naip5 | 31,72 | -0,1407 | 0,836394024 | chr13 | - |
| Naip6 | 6,64 | -0,0045 | 0,996644671 | chr13 | - |
| Ndufaf2 | 317,94 | -0,1761 | 0,674211854 | chr13 | - |

| | | | | | |
|---|---|---|---|---|---|
| Ndufs4 | 2975,60 | -0,4085 | 0,149780096 | chr13 | - |
| Ndufs6 | 2757,99 | -0,4906 | 0,110409761 | chr13 | - |
| Nedd9 | 59,94 | 0,2264 | 0,682032071 | chr13 | - |
| Net1 | 917,37 | -0,2628 | 0,348561153 | chr13 | - |
| Nfil3 | 280,04 | -0,3585 | 0,402628031 | chr13 | - |
| Nim1k | 39,85 | 0,1851 | 0,802632263 | chr13 | - |
| Ninj1 | 1533,75 | -0,3961 | 0,466018496 | chr13 | + |
| Nkapl | 4,28 | 0,7577 | 0,217779126 | chr13 | - |
| Nln | 597,57 | 0,0467 | 0,931322892 | chr13 | - |
| Nnt | 1201,39 | 0,2311 | 0,53935078 | chr13 | - |
| Nol7 | 626,54 | -0,0727 | 0,877920448 | chr13 | + |
| Nol8 | 311,03 | -0,0752 | 0,88569455 | chr13 | + |
| Nop16 | 541,89 | -0,4001 | 0,224157717 | chr13 | - |
| Nqo2 | 2909,03 | 0,0094 | 0,988512389 | chr13 | + |
| Nr2f1 | 5,34 | -0,1322 | 0,88569455 | chr13 | - |
| Nsa2 | 2376,16 | 0,0549 | 0,895961198 | chr13 | - |
| Nsd1 | 863,10 | -0,0190 | 0,963078727 | chr13 | + |
| Nsun2 | 1508,11 | -0,3702 | 0,138292882 | chr13 | + |
| Nup153 | 358,94 | 0,1826 | 0,566425551 | chr13 | - |
| Ogn | 83,72 | 0,5359 | 0,239641183 | chr13 | + |
| Omd | 70,03 | 0,3678 | 0,468682808 | chr13 | + |
| Paip1 | 1183,29 | -0,2691 | 0,411552342 | chr13 | + |
| Pak1ip1 | 590,15 | -0,3396 | 0,193295503 | chr13 | + |
| Papd4 | 437,07 | 0,0560 | 0,905147397 | chr13 | - |
| Papd7 | 267,89 | -0,3592 | 0,244197385 | chr13 | - |
| Parp8 | 23,65 | 0,5969 | 0,313882736 | chr13 | - |
| Pcbd2 | 182,66 | -0,1638 | 0,679434494 | chr13 | + |
| Pdcd6 | 1400,13 | -0,2075 | 0,442165263 | chr13 | - |
| Pde4d | 95,15 | 0,8139 | 0,062942876 | chr13 | + |
| Pde8b | 10,64 | -0,1290 | 0,884370894 | chr13 | - |
| Pdlim7 | 69,10 | 0,1079 | 0,88569455 | chr13 | - |
| Pfkp | 38,22 | 0,4829 | 0,349391987 | chr13 | - |
| Pgbd1 | 15,72 | 0,8032 | 0,170348649 | chr13 | - |
| Phactr1 | 9,44 | 0,1026 | 0,911210861 | chr13 | + |
| Phf2 | 149,30 | 0,2922 | 0,546076999 | chr13 | - |
| Pik3r1 | 1049,30 | -0,0241 | 0,970125173 | chr13 | - |
| Pitrm1 | 487,20 | -0,0138 | 0,977120448 | chr13 | + |
| Plk2 | 153,31 | -0,2484 | 0,545220581 | chr13 | + |
| Poc5 | 63,79 | 0,2387 | 0,618749555 | chr13 | + |
| Polk | 94,54 | 0,4015 | 0,367138036 | chr13 | - |
| Polr3g | 176,35 | -0,5109 | 0,140027516 | chr13 | - |
| Plpp1 | 363,58 | 0,1857 | 0,629147953 | chr13 | + |
| Ppp1r3g | 165,15 | -0,1334 | 0,87995617 | chr13 | + |
| Ppwd1 | 205,35 | 0,3600 | 0,277831408 | chr13 | - |
| Prelid1 | 1895,03 | -0,2746 | 0,470730504 | chr13 | + |
| Prpf4b | 492,05 | 0,0319 | 0,944925159 | chr13 | + |
| Psma2 | 2887,93 | -0,1246 | 0,733803092 | chr13 | + |
| Psmg4 | 83,42 | -0,2153 | 0,751230284 | chr13 | + |
| Ptcd2 | 170,70 | -0,0507 | 0,911462802 | chr13 | - |
| Ptpdc1 | 19,83 | 0,6380 | 0,239157449 | chr13 | - |

| | | | | | |
|---|---|---|---|---|---|
| Pxdc1 | 705,86 | -0,4184 | 0,12746076 | chr13 | - |
| Rab24 | 397,71 | -0,5024 | 0,155681449 | chr13 | - |
| Rad17 | 240,43 | 0,0082 | 0,988171576 | chr13 | - |
| Rala | 578,53 | -0,2544 | 0,453845946 | chr13 | - |
| Ranbp9 | 638,55 | -0,1424 | 0,680997554 | chr13 | - |
| Rasa1 | 417,38 | 0,0339 | 0,943305207 | chr13 | - |
| Rasgrf2 | 4,89 | -0,3741 | 0,617869186 | chr13 | - |
| Rfesd | 195,70 | 0,3383 | 0,487862806 | chr13 | - |
| Rgs14 | 4,27 | 0,6136 | 0,343698954 | chr13 | + |
| Rgs7bp | 14,33 | -0,2797 | 0,693242675 | chr13 | - |
| Riok1 | 106,21 | 0,3863 | 0,279404793 | chr13 | + |
| Rmi1 | 325,10 | 0,0883 | 0,828012328 | chr13 | + |
| Rnf144b | 732,63 | 0,3525 | 0,287543614 | chr13 | + |
| Rnf180 | 2,98 | 0,1678 | 0,835531954 | chr13 | - |
| Rnf44 | 410,71 | 0,0451 | 0,938364545 | chr13 | - |
| Rreb1 | 231,35 | 0,0530 | 0,935055331 | chr13 | + |
| S1pr3 | 25,56 | 0,8919 | 0,065791837 | chr13 | + |
| Scamp1 | 1332,20 | 0,1432 | 0,714302449 | chr13 | - |
| Sdha | 8894,54 | -0,2812 | 0,219049665 | chr13 | - |
| Sema4d | 11,18 | 0,6028 | 0,335636465 | chr13 | - |
| Serf1 | 83,97 | -0,6585 | 0,220525617 | chr13 | + |
| Serinc5 | 584,29 | -0,1796 | 0,696074872 | chr13 | + |
| Serpinb6a | 463,70 | 0,3388 | 0,212649167 | chr13 | - |
| Serpinb6b | 71,47 | 0,0754 | 0,890498303 | chr13 | + |
| Serpinb6d | 2,09 | 0,5713 | NA | chr13 | + |
| Serpinb9 | 235,22 | 0,6943 | 0,063957057 | chr13 | + |
| Sfxn1 | 1752,83 | 0,0431 | 0,932528423 | chr13 | + |
| Sgtb | 35,19 | 0,0023 | 0,997464504 | chr13 | + |
| Sirt5 | 201,42 | 0,6124 | 0,093189314 | chr13 | + |
| Skiv2l2 | 560,98 | 0,2952 | 0,428072237 | chr13 | - |
| Slc12a7 | 207,88 | 0,1290 | 0,771003172 | chr13 | + |
| Slc17a1 | 357,43 | 0,4743 | 0,168419405 | chr13 | + |
| Slc17a2 | 5241,42 | -0,2454 | 0,513640882 | chr13 | + |
| Slc17a3 | 1628,08 | -0,1995 | 0,620509787 | chr13 | + |
| Slc17a4 | 969,92 | 0,6600 | 0,104029003 | chr13 | - |
| Slc22a23 | 939,77 | -0,3642 | 0,37806194 | chr13 | - |
| Slc25a48 | 35,40 | -0,3320 | 0,546407458 | chr13 | + |
| Slc28a3 | 3,00 | -0,3754 | 0,599116638 | chr13 | - |
| Slc30a5 | 951,94 | 0,2996 | 0,392789224 | chr13 | - |
| Slc35b3 | 217,45 | -0,0175 | 0,975879219 | chr13 | - |
| Slc35d2 | 803,07 | -0,0097 | 0,98507181 | chr13 | - |
| Slc38a9 | 223,91 | -0,1004 | 0,81957953 | chr13 | + |
| Slc6a18 | 16,27 | -0,1339 | 0,869631925 | chr13 | - |
| Slc6a19 | 4,36 | 0,8614 | NA | chr13 | - |
| Smad5 | 432,04 | -0,1337 | 0,677398032 | chr13 | + |
| Smim15 | 474,95 | 0,3811 | 0,22361789 | chr13 | + |
| Smn1 | 86,70 | -0,1716 | 0,730715292 | chr13 | + |
| Snrnp48 | 352,14 | -0,5695 | 0,202636202 | chr13 | + |
| Snx18 | 332,79 | 0,1619 | 0,618749555 | chr13 | - |
| Sox4 | 19,62 | 0,9631 | 0,073551485 | chr13 | - |

| | | | | | |
|---|---|---|---|---|---|
| Spin1 | 1414,15 | 0,1468 | 0,671272065 | chr13 | + |
| Sptlc1 | 582,64 | 0,0634 | 0,88569455 | chr13 | - |
| Srd5a1 | 1634,34 | -0,4855 | 0,123405092 | chr13 | - |
| Srek1 | 308,29 | 0,1066 | 0,817637582 | chr13 | - |
| Srek1ip1 | 329,61 | 0,2506 | 0,527966188 | chr13 | + |
| Ssbp2 | 23,28 | -0,4769 | 0,444071101 | chr13 | + |
| Ssr1 | 2290,92 | 0,0238 | 0,955504626 | chr13 | - |
| Stard3nl | 159,37 | 0,3219 | 0,424863329 | chr13 | - |
| Sugct | 748,22 | -0,2157 | 0,607061881 | chr13 | - |
| Susd3 | 3,32 | -0,1927 | 0,811801336 | chr13 | - |
| Syk | 74,94 | 0,0842 | 0,88569455 | chr13 | + |
| Taf9 | 771,90 | 0,0524 | 0,925508783 | chr13 | + |
| Tbc1d7 | 132,55 | -0,3277 | 0,357345082 | chr13 | - |
| Tbca | 2120,39 | -0,1106 | 0,781786246 | chr13 | + |
| Tbce | 365,44 | -0,3090 | 0,402148509 | chr13 | - |
| Tdp2 | 399,56 | -0,3491 | 0,299089342 | chr13 | + |
| Tert | 48,63 | 0,6740 | 0,191230937 | chr13 | + |
| Tfap2a | 3,92 | 0,1964 | 0,816082306 | chr13 | - |
| Tgfbi | 553,79 | 0,5046 | 0,087515981 | chr13 | + |
| Thoc3 | 352,43 | -0,2019 | 0,601597387 | chr13 | - |
| Tifab | 52,05 | 0,4070 | 0,361776578 | chr13 | - |
| Tmed9 | 1602,87 | -0,0044 | 0,993487304 | chr13 | + |
| Tmem161b | 207,24 | 0,0243 | 0,970399403 | chr13 | + |
| Tmem167 | 1406,82 | -0,1685 | 0,679975413 | chr13 | + |
| Tmem170b | 411,09 | -0,2017 | 0,548818911 | chr13 | + |
| Tmem171 | 2,71 | 0,1269 | 0,88306163 | chr13 | - |
| Tnpo1 | 750,92 | 0,5461 | 0,075427239 | chr13 | - |
| Tppp | 341,93 | 0,5375 | 0,250303145 | chr13 | + |
| Trappc13 | 609,81 | 0,1290 | 0,744313101 | chr13 | - |
| Trim23 | 177,37 | 0,4437 | 0,343973842 | chr13 | + |
| Trim27 | 267,49 | -0,4625 | 0,191022172 | chr13 | + |
| Trip13 | 16,49 | -0,4838 | 0,423089018 | chr13 | - |
| Tspan17 | 36,82 | -0,0138 | 0,986889999 | chr13 | + |
| Ttc37 | 257,21 | 0,3185 | 0,444852563 | chr13 | + |
| Tubb2a | 241,56 | 0,0515 | 0,931356238 | chr13 | - |
| Tubb2b | 2,50 | 0,4702 | NA | chr13 | - |
| Txndc15 | 1352,64 | -0,5664 | 0,055150337 | chr13 | + |
| Txndc5 | 2547,58 | -0,2510 | 0,526620417 | chr13 | - |
| Ubqln1 | 1807,53 | -0,2278 | 0,369767166 | chr13 | - |
| Uimc1 | 366,12 | -0,0807 | 0,862616891 | chr13 | - |
| Unc5a | 6,56 | -0,0614 | 0,951806422 | chr13 | + |
| Uqcrb | 7755,10 | -0,1568 | 0,684535373 | chr13 | - |
| Uqcrfs1 | 3673,60 | -0,3550 | 0,179535584 | chr13 | - |
| Utp15 | 160,71 | 0,0297 | 0,952188756 | chr13 | - |
| Vcan | 7,19 | 0,8580 | 0,155520806 | chr13 | - |
| Vps41 | 1043,00 | -0,1534 | 0,716183804 | chr13 | + |
| Wdr37 | 235,04 | 0,0841 | 0,87623846 | chr13 | - |
| Wdr41 | 189,80 | -0,1613 | 0,692394791 | chr13 | + |
| Wnk2 | 2,88 | 0,7108 | 0,23305528 | chr13 | - |
| Wrnip1 | 626,89 | 0,1360 | 0,799261258 | chr13 | + |

| | | | | | |
|---|---|---|---|---|---|
| Yae1d1 | 509,68 | 0,1081 | 0,82704652 | chr13 | - |
| Zbed3 | 31,02 | 0,0108 | 0,990189789 | chr13 | + |
| Zcchc6 | 1453,53 | 0,2611 | 0,440094295 | chr13 | - |
| Zcchc9 | 201,67 | -0,2830 | 0,394132098 | chr13 | - |
| Zfp131 | 202,70 | -0,3546 | 0,341412029 | chr13 | - |
| Zfp169 | 18,17 | 0,3679 | 0,581893426 | chr13 | - |
| Zfp184 | 4,00 | 0,1397 | 0,877822516 | chr13 | + |
| Zfp273 | 41,17 | 0,4955 | 0,309952056 | chr13 | + |
| Zfp322a | 218,80 | 0,3433 | 0,31835964 | chr13 | - |
| Zfp346 | 69,20 | -0,0659 | 0,909774279 | chr13 | + |
| Zfp366 | 5,43 | 0,8556 | 0,158793576 | chr13 | + |
| Zfp367 | 335,71 | -0,0234 | 0,969512928 | chr13 | - |
| Zfp429 | 32,96 | 0,0921 | 0,903494966 | chr13 | - |
| Zfp455 | 39,09 | 0,6693 | 0,178168438 | chr13 | + |
| Zfp456 | 41,00 | 0,2545 | 0,630381755 | chr13 | - |
| Zfp458 | 13,72 | 1,0139 | 0,063376737 | chr13 | - |
| Zfp459 | 8,61 | 0,2141 | 0,7950804 | chr13 | - |
| Zfp493 | 30,07 | 0,6614 | 0,173352491 | chr13 | + |
| Zfp58 | 112,20 | 0,2935 | 0,444265933 | chr13 | - |
| Zfp595 | 88,97 | 0,2595 | 0,549296977 | chr13 | - |
| Zfp65 | 247,45 | 0,4074 | 0,196659616 | chr13 | - |
| Zfp708 | 103,85 | 0,4534 | 0,215046281 | chr13 | - |
| Zfp712 | 15,87 | 0,9062 | 0,100179069 | chr13 | - |
| Zfp748 | 109,25 | 0,5455 | 0,099893252 | chr13 | - |
| Zfp825 | 652,94 | 0,1301 | 0,734795364 | chr13 | - |
| Zfp85 | 50,73 | 0,3696 | 0,501682426 | chr13 | - |
| Zfp85os | 10,26 | 0,5223 | 0,430730682 | chr13 | + |
| Zfp87 | 120,53 | 0,0517 | 0,923331663 | chr13 | - |
| Zfp874a | 566,52 | 0,0464 | 0,929524781 | chr13 | - |
| Zfp874b | 222,42 | 0,1462 | 0,716169774 | chr13 | - |
| Zfp953 | 112,72 | 0,4298 | 0,339404199 | chr13 | - |
| Zfyve16 | 122,17 | 0,2411 | 0,612582336 | chr13 | - |
| Zmynd11 | 461,80 | -0,1477 | 0,653474875 | chr13 | - |
| Zscan12 | 123,61 | 0,2454 | 0,55128576 | chr13 | + |
| Zscan26 | 731,98 | 0,1324 | 0,795226131 | chr13 | - |
| Zswim6 | 59,88 | -0,0455 | 0,947294154 | chr13 | - |
| 1700011H14Rik | 26,09 | 0,3051 | 0,658008155 | chr14 | - |
| 1700092C10Rik | 5,13 | -0,0803 | 0,93538092 | chr14 | - |
| Lrmda | 11,80 | -0,2649 | 0,740269415 | chr14 | + |
| 1700123O20Rik | 764,19 | -0,1325 | 0,753582185 | chr14 | + |
| 1810011H11Rik | 25,39 | 0,3902 | 0,535573923 | chr14 | + |
| Rtraf | 1983,07 | -0,4950 | 0,092349447 | chr14 | - |
| 3830406C13Rik | 469,34 | 0,4326 | 0,23655468 | chr14 | + |
| 4930452B06Rik | 28,17 | 0,1527 | 0,840322097 | chr14 | - |
| 4931414P19Rik | 42,81 | 0,7592 | 0,065893559 | chr14 | - |
| Rubcnl | 10,10 | -0,7143 | 0,237903839 | chr14 | + |
| 6230400D17Rik | 2,31 | -0,4773 | NA | chr14 | - |
| 9930012K11Rik | 6,90 | 0,0671 | 0,947182472 | chr14 | - |
| A330035P11Rik | 20,52 | -0,4743 | 0,407597046 | chr14 | - |
| Abcc4 | 40,35 | 0,2858 | 0,619039366 | chr14 | - |

| | | | | | |
|---|---|---|---|---|---|
| Abhd4 | 508,67 | 0,0380 | 0,943064069 | chr14 | + |
| Abhd6 | 946,84 | -0,4112 | 0,309392522 | chr14 | + |
| Acin1 | 522,06 | 0,1041 | 0,826107735 | chr14 | - |
| Acox2 | 2844,97 | -0,0429 | 0,942982044 | chr14 | - |
| Actr8 | 333,25 | -0,2739 | 0,327939573 | chr14 | + |
| Adam2 | 2,83 | -0,2236 | 0,771044887 | chr14 | - |
| Adam28 | 3,57 | -0,1760 | 0,834244884 | chr14 | - |
| Adamdec1 | 53,13 | 0,0907 | 0,905147397 | chr14 | - |
| Adcy4 | 52,04 | -0,2817 | 0,658008155 | chr14 | - |
| Adk | 14666,49 | 0,0938 | 0,808864467 | chr14 | + |
| Adra1a | 38,65 | -0,7355 | 0,124458443 | chr14 | + |
| Akap11 | 608,89 | 0,2734 | 0,449506818 | chr14 | - |
| Amer2 | 3,90 | -0,7130 | 0,242678725 | chr14 | + |
| Ang | 8071,10 | -0,6514 | 0,098348401 | chr14 | + |
| Ankrd28 | 397,78 | 0,0264 | 0,967536738 | chr14 | - |
| Anxa11 | 372,76 | -0,1866 | 0,681917507 | chr14 | + |
| Anxa7 | 1836,56 | -0,4060 | 0,110056031 | chr14 | - |
| Ap1g2 | 98,29 | 0,3721 | 0,440917912 | chr14 | - |
| Ap3m1 | 3300,06 | 0,4736 | 0,054219322 | chr14 | - |
| Ap5m1 | 216,96 | 0,0180 | 0,979212524 | chr14 | + |
| Apex1 | 653,90 | -0,2411 | 0,491510849 | chr14 | + |
| Appl1 | 305,45 | 0,5427 | 0,081916308 | chr14 | - |
| Arf4 | 1959,68 | -0,0674 | 0,875636241 | chr14 | + |
| Arhgap22 | 4,22 | -0,0414 | 0,969176357 | chr14 | + |
| Arhgef3 | 557,02 | 0,2251 | 0,505015502 | chr14 | + |
| Arhgef40 | 61,79 | -0,4741 | 0,268690208 | chr14 | + |
| Arl11 | 8,83 | 0,2323 | 0,775458257 | chr14 | + |
| Atg14 | 374,88 | -0,1438 | 0,701666523 | chr14 | - |
| Atp8a2 | 2,61 | -0,2871 | 0,696074872 | chr14 | - |
| Atxn7 | 298,57 | -0,2931 | 0,387366838 | chr14 | + |
| Bap1 | 374,74 | -0,2693 | 0,413323375 | chr14 | + |
| Bcl2l2 | 432,90 | 0,0406 | 0,932012942 | chr14 | + |
| Bin3 | 237,75 | -0,0288 | 0,957837964 | chr14 | + |
| Blk | 2,35 | 0,8596 | NA | chr14 | - |
| Bmp1 | 1070,11 | -0,2170 | 0,606429785 | chr14 | - |
| Bmpr1a | 695,55 | 0,3882 | 0,28179192 | chr14 | - |
| Bnip3l | 2444,77 | -0,2154 | 0,514309898 | chr14 | - |
| Bora | 5,83 | -0,2762 | 0,733541536 | chr14 | + |
| Btd | 925,07 | -0,2950 | 0,36985369 | chr14 | + |
| C1qtnf9 | 8,55 | 0,1029 | 0,912010517 | chr14 | + |
| Cab39l | 1304,20 | 0,1236 | 0,716169774 | chr14 | + |
| Cacna1d | 6,32 | -0,5745 | 0,391792082 | chr14 | - |
| Camk2g | 108,06 | -0,4257 | 0,269107101 | chr14 | - |
| Capn7 | 684,87 | -0,1274 | 0,721406886 | chr14 | + |
| Cbln3 | 13,31 | 0,6402 | 0,310149361 | chr14 | - |
| Ccar2 | 206,30 | 0,4219 | 0,225613833 | chr14 | - |
| Ccdc25 | 1086,53 | -0,1298 | 0,783937395 | chr14 | + |
| Ccdc66 | 54,85 | 0,7834 | 0,12746076 | chr14 | - |
| Ccser2 | 587,93 | -0,1803 | 0,637300541 | chr14 | - |
| Cdadc1 | 1221,74 | -0,2067 | 0,526749174 | chr14 | - |

| | | | | | |
|---|---|---|---|---|---|
| Cdca2 | 8,03 | 0,3001 | 0,703944394 | chr14 | - |
| Cdh24 | 3,37 | 0,0963 | 0,918104715 | chr14 | - |
| Cdkn3 | 11,05 | 0,1062 | 0,907984163 | chr14 | + |
| Cebpe | 58,45 | -0,1717 | 0,776467957 | chr14 | - |
| Cenpj | 75,60 | 0,4906 | 0,408720522 | chr14 | - |
| Chchd1 | 1070,32 | -0,3215 | 0,280299373 | chr14 | + |
| Chd8 | 362,02 | 0,0610 | 0,88099064 | chr14 | - |
| Chdh | 2771,42 | 0,1611 | 0,627992485 | chr14 | + |
| Chmp7 | 615,62 | -0,2301 | 0,433302067 | chr14 | - |
| Chrna2 | 119,92 | -0,0955 | 0,906066614 | chr14 | + |
| Cideb | 4004,08 | -0,0220 | 0,955234619 | chr14 | - |
| Cln5 | 693,52 | 0,0764 | 0,867463624 | chr14 | + |
| Clu | 20807,52 | -0,1913 | 0,603818311 | chr14 | + |
| Clybl | 790,65 | -0,2670 | 0,287640298 | chr14 | + |
| Cog3 | 304,69 | 0,0596 | 0,89040553 | chr14 | - |
| Commd6 | 994,53 | -0,0163 | 0,975445719 | chr14 | - |
| Comtd1 | 37,84 | -0,6893 | 0,23219679 | chr14 | - |
| Cpb2 | 13986,30 | -0,0542 | 0,895594598 | chr14 | + |
| Ctsb | 12526,12 | 0,1221 | 0,661068458 | chr14 | + |
| Cysltr2 | 3,02 | 0,5825 | 0,31302658 | chr14 | - |
| D830030K20Rik | 11,53 | -0,4019 | 0,572534156 | chr14 | + |
| Dach1 | 5,87 | 0,7475 | 0,233364856 | chr14 | - |
| Dad1 | 2371,33 | -0,2993 | 0,280102462 | chr14 | - |
| Dcaf11 | 5816,46 | -0,2338 | 0,559525694 | chr14 | + |
| Dcp1a | 83,01 | -0,0830 | 0,878605672 | chr14 | + |
| Ddhd1 | 239,49 | -0,2512 | 0,563218321 | chr14 | - |
| Dennd6a | 389,90 | -0,0608 | 0,895594598 | chr14 | + |
| Dgkh | 5,57 | -0,5560 | 0,409632036 | chr14 | - |
| Dhrs1 | 2859,39 | -0,0328 | 0,948314012 | chr14 | - |
| Dhrs4 | 2603,31 | -0,3384 | 0,443287405 | chr14 | + |
| Dis3 | 112,07 | -0,0670 | 0,898098997 | chr14 | - |
| Dleu2 | 103,53 | -0,2596 | 0,610703272 | chr14 | - |
| Dlg5 | 38,30 | 0,1000 | 0,887518458 | chr14 | - |
| Dlgap5 | 5,42 | 0,0757 | 0,940390384 | chr14 | - |
| Dmtn | 4,70 | 0,0681 | 0,946011436 | chr14 | - |
| Dnah1 | 3,00 | 0,9068 | 0,08440425 | chr14 | - |
| Dnajc15 | 663,80 | -0,2928 | 0,342277341 | chr14 | - |
| Dnajc3 | 5584,62 | 0,0061 | 0,991447068 | chr14 | + |
| Dnajc9 | 108,69 | 0,2704 | 0,499958086 | chr14 | - |
| Dnase1l3 | 906,21 | -0,0827 | 0,855769633 | chr14 | - |
| Dock5 | 292,71 | -0,0851 | 0,834675238 | chr14 | - |
| Dock9 | 188,14 | 0,3309 | 0,460127107 | chr14 | - |
| Dok2 | 33,63 | 0,0893 | 0,914454715 | chr14 | + |
| Dph3 | 515,85 | -0,2796 | 0,413537476 | chr14 | - |
| Dpysl2 | 120,61 | 0,1107 | 0,83796616 | chr14 | - |
| Dzip1 | 12,58 | 0,8235 | 0,147364898 | chr14 | - |
| E330034G19Rik | 2,37 | 0,2684 | NA | chr14 | + |
| Eaf1 | 551,34 | -0,1802 | 0,603196033 | chr14 | + |
| Ear2 | 18,27 | -0,1664 | 0,822888019 | chr14 | + |
| Ebpl | 1653,31 | -0,0394 | 0,943108135 | chr14 | - |

| | | | | | |
|---|---|---|---|---|---|
| Ecd | 391,56 | -0,0255 | 0,958614145 | chr14 | - |
| Ednrb | 309,91 | 0,4109 | 0,209104214 | chr14 | - |
| Elf1 | 298,72 | 0,1407 | 0,69353394 | chr14 | + |
| Elp3 | 411,16 | 0,3329 | 0,318765186 | chr14 | - |
| Emc9 | 325,72 | -0,3258 | 0,514682351 | chr14 | - |
| Ephx2 | 15821,76 | -0,1774 | 0,612061669 | chr14 | - |
| Epsti1 | 61,54 | 0,3576 | 0,43260167 | chr14 | + |
| Erc2 | 73,54 | -0,5784 | 0,139781433 | chr14 | + |
| Ercc6 | 159,78 | -0,0362 | 0,944925159 | chr14 | + |
| Ero1l | 500,41 | -0,0298 | 0,947045073 | chr14 | - |
| Esco2 | 30,46 | -0,1264 | 0,87618436 | chr14 | - |
| Esd | 8206,66 | -0,1901 | 0,535475471 | chr14 | + |
| Exoc5 | 421,84 | -0,2528 | 0,430730682 | chr14 | - |
| Extl3 | 314,69 | 0,0906 | 0,837132801 | chr14 | - |
| Fam107a | 11,27 | -0,0532 | 0,957318111 | chr14 | - |
| Fam124a | 181,15 | -0,1788 | 0,651662099 | chr14 | + |
| Fam149b | 273,59 | -0,2185 | 0,500796855 | chr14 | + |
| Fam160b2 | 107,64 | -0,1387 | 0,799751657 | chr14 | - |
| Fam167a | 26,01 | -0,1579 | 0,820554805 | chr14 | + |
| Fam208a | 630,55 | 0,5272 | 0,104667853 | chr14 | + |
| Fam213a | 1773,77 | 0,5504 | 0,052523656 | chr14 | - |
| Fam25c | 1132,00 | -0,3774 | 0,433006548 | chr14 | - |
| Fam35a | 1004,91 | 0,3252 | 0,209104214 | chr14 | - |
| Farp1 | 193,83 | -0,2909 | 0,502046579 | chr14 | + |
| Fbxl3 | 733,02 | 0,2553 | 0,388402325 | chr14 | - |
| Fbxo34 | 326,00 | -0,1752 | 0,636381742 | chr14 | + |
| Fdft1 | 1983,89 | -0,3966 | 0,322192824 | chr14 | - |
| Fermt2 | 2360,69 | 0,4457 | 0,157077211 | chr14 | - |
| Fgf9 | 3,26 | 0,0174 | 0,986818215 | chr14 | + |
| Fhit | 150,20 | 0,6554 | 0,065752252 | chr14 | - |
| Flnb | 1042,01 | 0,1884 | 0,540805163 | chr14 | + |
| Fndc3a | 1097,84 | 0,2017 | 0,640497332 | chr14 | - |
| Fut11 | 64,16 | 0,4056 | 0,374019679 | chr14 | + |
| Fzd3 | 8,69 | -0,3794 | 0,608287564 | chr14 | - |
| Galnt15 | 7,05 | 0,3986 | 0,589786442 | chr14 | + |
| Gata4 | 143,01 | 0,2649 | 0,517431746 | chr14 | - |
| Gch1 | 2487,34 | -0,4918 | 0,054144207 | chr14 | - |
| Gdf10 | 82,35 | 0,4720 | 0,342081078 | chr14 | + |
| Gdf2 | 401,04 | 0,4583 | 0,066915827 | chr14 | + |
| Gfra2 | 13,82 | 0,3460 | 0,623104286 | chr14 | + |
| Ggact | 1269,39 | -0,2905 | 0,303046806 | chr14 | - |
| Ghitm | 9293,87 | 0,2174 | 0,439437431 | chr14 | - |
| Gjb2 | 3593,49 | -0,1960 | 0,574684302 | chr14 | - |
| Glt8d1 | 161,91 | 0,1588 | 0,744313101 | chr14 | + |
| Glud1 | 15794,67 | -0,3201 | 0,227835508 | chr14 | + |
| Gm10364 | 3,49 | 0,0096 | 0,992733608 | chr14 | + |
| Gm10845 | 293,41 | -0,1140 | 0,793407482 | chr14 | - |
| Gm16973 | 53,18 | -0,3773 | 0,422960414 | chr14 | - |
| Gm17066 | 183,52 | -0,1837 | 0,690021525 | chr14 | - |
| Gm1821 | 345,24 | -0,2395 | 0,627875496 | chr14 | + |

| | | | | | |
|---|---|---|---|---|---|
| Gm3002 | 2,20 | 0,0737 | NA | chr14 | + |
| Gm3219 | 148,03 | -0,6096 | 0,050835053 | chr14 | + |
| Gm4285 | 34,93 | -0,1960 | 0,760028388 | chr14 | - |
| Gm5622 | 3,54 | 0,3489 | 0,640132917 | chr14 | + |
| Gm6498 | 4,59 | -0,6253 | 0,339404199 | chr14 | + |
| Gmfb | 959,16 | 0,0586 | 0,90870562 | chr14 | - |
| Gmpr2 | 351,39 | 0,0026 | 0,996644671 | chr14 | + |
| Gng2 | 59,15 | 0,4161 | 0,318113166 | chr14 | - |
| Gnl3 | 575,67 | -0,4815 | 0,163729985 | chr14 | - |
| Gnpnat1 | 2080,13 | -0,1231 | 0,813415373 | chr14 | - |
| Gnrh1 | 5,92 | -0,4228 | 0,565013157 | chr14 | + |
| Gpalpp1 | 321,43 | 0,1888 | 0,618381442 | chr14 | - |
| Gpc6 | 53,97 | 0,1339 | 0,849477252 | chr14 | + |
| Gpr18 | 4,20 | 0,2366 | 0,770645004 | chr14 | - |
| Gpr180 | 373,14 | -0,4114 | 0,162272166 | chr14 | + |
| Gpr183 | 6,96 | 0,2689 | 0,740716918 | chr14 | - |
| Gtf2f2 | 179,37 | -0,2388 | 0,549694981 | chr14 | - |
| Gzmb | 11,42 | 0,3920 | 0,582500211 | chr14 | - |
| Hacl1 | 5033,64 | -0,2302 | 0,483930201 | chr14 | - |
| Haus4 | 110,42 | -0,3759 | 0,263829098 | chr14 | - |
| Hmbox1 | 43,32 | 0,1064 | 0,874167297 | chr14 | - |
| Hnrnpc | 2467,28 | 0,2161 | 0,513570549 | chr14 | - |
| Homez | 68,23 | 0,0953 | 0,874167297 | chr14 | - |
| Hr | 141689,40 | -0,1538 | 0,739141404 | chr14 | + |
| Ift88 | 102,83 | 0,2173 | 0,615783353 | chr14 | + |
| Il17rd | 9,90 | 0,7830 | 0,194197016 | chr14 | + |
| Ints9 | 122,49 | 0,0175 | 0,980767643 | chr14 | + |
| Ipo4 | 241,51 | -0,4455 | 0,208540118 | chr14 | - |
| Ipo5 | 1017,55 | -0,3766 | 0,186556074 | chr14 | + |
| Irf9 | 639,24 | 0,0313 | 0,94647679 | chr14 | + |
| Acod1 | 2,21 | 0,3258 | NA | chr14 | + |
| Itgbl1 | 16,51 | 0,7642 | 0,198455103 | chr14 | + |
| Itih1 | 11463,21 | -0,1499 | 0,576265028 | chr14 | - |
| Itih3 | 14760,99 | 0,0934 | 0,806012703 | chr14 | - |
| Itih4 | 32690,32 | 0,0114 | 0,985849808 | chr14 | + |
| Itm2b | 16600,71 | -0,3527 | 0,23305528 | chr14 | - |
| Kat6b | 119,24 | 0,2208 | 0,66477796 | chr14 | + |
| Kbtbd7 | 215,43 | -0,2114 | 0,603013282 | chr14 | + |
| Kcnk5 | 755,00 | -0,1438 | 0,821346657 | chr14 | - |
| Kctd6 | 112,28 | -0,3540 | 0,413323375 | chr14 | + |
| Khnyn | 245,96 | 0,5826 | 0,23347764 | chr14 | + |
| Kif13b | 157,13 | 0,6314 | 0,124945987 | chr14 | + |
| Klf12 | 153,95 | 0,2932 | 0,570801749 | chr14 | - |
| Kpna3 | 763,82 | -0,0429 | 0,925898178 | chr14 | - |
| Ktn1 | 345,42 | 0,1363 | 0,769969083 | chr14 | + |
| Lacc1 | 126,78 | 0,4484 | 0,250727316 | chr14 | - |
| Lats2 | 189,46 | -0,3416 | 0,330697335 | chr14 | - |
| Lcp1 | 189,40 | -0,0254 | 0,972380647 | chr14 | + |
| Ldb3 | 4,29 | 0,5075 | 0,448865116 | chr14 | - |
| Lgals3 | 1135,50 | 0,1251 | 0,780902054 | chr14 | + |

| | | | | | |
|---|---|---|---|---|---|
| Lgi3 | 3,43 | 0,5057 | 0,424863329 | chr14 | + |
| Loxl2 | 125,42 | 0,4692 | 0,170992427 | chr14 | + |
| Lrch1 | 107,84 | 0,3333 | 0,474644284 | chr14 | - |
| Lrit1 | 202,24 | -0,4625 | 0,236106121 | chr14 | + |
| Lrp10 | 400,08 | -0,3332 | 0,293711445 | chr14 | + |
| Lrrc18 | 7,80 | 0,6692 | 0,298116176 | chr14 | + |
| Lrtm1 | 58,83 | -0,2801 | 0,704956933 | chr14 | + |
| Ltb4r1 | 9,82 | -0,5886 | 0,370803111 | chr14 | + |
| Ltb4r2 | 18,50 | 0,4409 | 0,517321822 | chr14 | + |
| Mapk1ip1l | 1017,05 | -0,1288 | 0,726240854 | chr14 | + |
| Mapk8 | 452,35 | -0,1761 | 0,651267254 | chr14 | - |
| Mat1a | 66856,71 | 0,4667 | 0,13621142 | chr14 | + |
| Mbl1 | 5948,03 | -0,2916 | 0,399992251 | chr14 | + |
| Mbnl2 | 534,34 | -0,5164 | 0,056273421 | chr14 | + |
| Med4 | 245,37 | -0,0232 | 0,966714122 | chr14 | + |
| Mettl17 | 123,45 | -0,1346 | 0,809461729 | chr14 | + |
| Mettl3 | 140,95 | 0,3198 | 0,452544998 | chr14 | - |
| Mettl6 | 254,80 | 0,0877 | 0,862421004 | chr14 | - |
| Mipep | 713,83 | 0,1341 | 0,6951078 | chr14 | + |
| Mir17hg | 55,42 | 0,1065 | 0,863490375 | chr14 | + |
| Mmp14 | 268,65 | 0,4745 | 0,194085468 | chr14 | + |
| Mmrn2 | 200,12 | -0,0853 | 0,838868977 | chr14 | + |
| Mphosph8 | 304,45 | 0,1733 | 0,723157997 | chr14 | + |
| Mrpl52 | 499,26 | -0,3731 | 0,619691963 | chr14 | + |
| Mrpl57 | 299,34 | -0,4016 | 0,588219472 | chr14 | + |
| Mrps16 | 950,70 | -0,0080 | 0,985269339 | chr14 | - |
| Msra | 3414,77 | -0,3031 | 0,199849806 | chr14 | - |
| Mss51 | 10,29 | -0,7724 | 0,20624753 | chr14 | - |
| Mtmr6 | 510,84 | 0,0688 | 0,884003681 | chr14 | + |
| Mtmr9 | 149,20 | 0,5377 | 0,183723189 | chr14 | - |
| Mtrf1 | 247,76 | 0,1093 | 0,812543852 | chr14 | + |
| Mustn1 | 62,05 | 0,5852 | 0,192379403 | chr14 | + |
| Mycbp2 | 397,96 | 0,3821 | 0,306398925 | chr14 | - |
| Mzt1 | 298,77 | 0,0247 | 0,966776406 | chr14 | - |
| Eef1akmt1 | 186,50 | -0,2766 | 0,57931481 | chr14 | - |
| Naa30 | 468,69 | -0,2565 | 0,469728046 | chr14 | + |
| Ncoa4 | 4414,12 | 0,1129 | 0,777753633 | chr14 | + |
| Ndfip2 | 642,90 | -0,0994 | 0,771142222 | chr14 | + |
| Ndrg2 | 19108,84 | -0,2090 | 0,366528137 | chr14 | - |
| Ndst2 | 211,44 | -0,1102 | 0,790166895 | chr14 | - |
| Nedd8 | 1935,56 | -0,1722 | 0,617467883 | chr14 | - |
| Neil2 | 11,55 | -0,2172 | 0,783422709 | chr14 | - |
| Nek4 | 84,95 | 0,1900 | 0,720311335 | chr14 | + |
| Nfatc4 | 5,00 | -0,0704 | 0,943346405 | chr14 | + |
| Ngdn | 468,83 | -0,1806 | 0,565200801 | chr14 | + |
| Ngly1 | 867,56 | -0,0707 | 0,862616891 | chr14 | + |
| Nid2 | 78,56 | 0,1015 | 0,862616891 | chr14 | + |
| Nisch | 940,31 | -0,1015 | 0,825518458 | chr14 | - |
| Nkiras1 | 363,36 | -0,2693 | 0,33879753 | chr14 | + |
| Nop9 | 262,33 | -0,2894 | 0,442222987 | chr14 | + |

| | | | | | |
|---|---|---|---|---|---|
| Nr1d2 | 2063,32 | -0,1131 | 0,813323745 | chr14 | - |
| Nt5dc2 | 58,01 | -0,7513 | 0,073845576 | chr14 | + |
| Nudt13 | 209,14 | 0,2849 | 0,490113662 | chr14 | + |
| Nudt15 | 21,82 | -0,1337 | 0,857773796 | chr14 | - |
| Nudt18 | 174,65 | 0,1340 | 0,77634365 | chr14 | + |
| Nufip1 | 151,40 | -0,1004 | 0,847796138 | chr14 | + |
| Nupl1 | 115,66 | 0,0949 | 0,863983232 | chr14 | - |
| Nynrin | 18,17 | -0,4243 | 0,494456563 | chr14 | + |
| Ogdhl | 1,95 | 0,5638 | NA | chr14 | + |
| Osgep | 244,80 | -0,2267 | 0,615304988 | chr14 | - |
| Oxa1l | 510,69 | -0,2668 | 0,408372125 | chr14 | + |
| Oxnad1 | 506,29 | 0,1595 | 0,695575308 | chr14 | + |
| Oxsm | 751,69 | 0,2200 | 0,626333589 | chr14 | - |
| Parg | 266,63 | 0,0493 | 0,906998011 | chr14 | + |
| Parp4 | 130,74 | -0,1457 | 0,744486882 | chr14 | + |
| Pbk | 10,92 | 0,4522 | 0,512939428 | chr14 | + |
| Pbrm1 | 487,42 | 0,0820 | 0,875717638 | chr14 | + |
| Pcca | 1996,38 | 0,2234 | 0,478251242 | chr14 | + |
| Pcdh17 | 20,03 | 0,3474 | 0,595228746 | chr14 | + |
| Pde12 | 162,88 | -0,0425 | 0,945352364 | chr14 | - |
| Pdhb | 11603,91 | -0,0814 | 0,874763133 | chr14 | - |
| Pdlim2 | 31,08 | 0,1494 | 0,823831306 | chr14 | - |
| Peli2 | 279,80 | -0,2572 | 0,424735065 | chr14 | + |
| Phf11a | 36,60 | 0,3626 | 0,540815736 | chr14 | - |
| Phf11b | 31,66 | 0,6612 | 0,197447777 | chr14 | - |
| Phf11c | 105,92 | 0,2496 | 0,736631536 | chr14 | - |
| Phf11d | 78,32 | 0,7729 | 0,093065906 | chr14 | - |
| Phf7 | 152,22 | -0,5083 | 0,120247725 | chr14 | - |
| Pibf1 | 73,36 | 0,3673 | 0,402148509 | chr14 | + |
| Pinx1 | 48,74 | 0,3399 | 0,512609895 | chr14 | + |
| Plau | 34,81 | 0,3379 | 0,587879802 | chr14 | + |
| Pnp | 2125,55 | 0,3207 | 0,357703576 | chr14 | + |
| Pnp2 | 9,00 | -0,2536 | 0,749603956 | chr14 | + |
| Polr3a | 184,22 | -0,1883 | 0,638379056 | chr14 | - |
| Polr3d | 78,94 | 0,0956 | 0,866762614 | chr14 | - |
| Ppif | 527,52 | 0,2339 | 0,581157554 | chr14 | + |
| Ppp1r3e | 18,28 | -0,2317 | 0,754953629 | chr14 | - |
| Ppp3cb | 369,64 | -0,0681 | 0,868479494 | chr14 | - |
| Prkcd | 163,42 | -0,0078 | 0,990189789 | chr14 | - |
| Prmt5 | 201,54 | -0,1825 | 0,692167613 | chr14 | - |
| Psmb11 | 10,36 | 0,8961 | 0,124483292 | chr14 | + |
| Psmb5 | 1362,57 | -0,3579 | 0,286763659 | chr14 | - |
| Psmc6 | 2434,90 | 0,0632 | 0,884454567 | chr14 | + |
| Psmd6 | 1708,97 | -0,2243 | 0,432420089 | chr14 | - |
| Psme1 | 1564,48 | -0,1901 | 0,682099278 | chr14 | + |
| Psme2 | 1723,07 | 0,0172 | 0,972177521 | chr14 | - |
| Pspc1 | 63,06 | 0,4485 | 0,420351643 | chr14 | - |
| Ptger2 | 25,54 | 0,1304 | 0,857773796 | chr14 | + |
| Ptk2b | 307,78 | -0,2266 | 0,631758049 | chr14 | - |
| Ptprg | 449,90 | -0,0455 | 0,917042614 | chr14 | + |

| | | | | | |
|---|---|---|---|---|---|
| Pxk | 310,85 | -0,0501 | 0,931322892 | chr14 | + |
| R3hcc1 | 159,04 | 0,0866 | 0,865418807 | chr14 | - |
| Rab2b | 57,13 | -0,0599 | 0,932012942 | chr14 | - |
| Rabggta | 375,61 | -0,1212 | 0,77350788 | chr14 | - |
| Rarb | 83,43 | 0,2605 | 0,719263094 | chr14 | - |
| Rb1 | 125,85 | 0,1294 | 0,784505535 | chr14 | - |
| Rbm26 | 147,14 | 0,4944 | 0,140539066 | chr14 | - |
| Rcbtb1 | 1005,82 | -0,3446 | 0,172087384 | chr14 | + |
| Rcbtb2 | 813,41 | 0,0148 | 0,9774226 | chr14 | + |
| Rec8 | 3,86 | -0,7057 | 0,23475207 | chr14 | + |
| Reep4 | 75,98 | -0,0024 | 0,996987906 | chr14 | + |
| Rem2 | 2,15 | 0,1702 | NA | chr14 | + |
| Rft1 | 332,47 | 0,0423 | 0,937247252 | chr14 | + |
| Rhobtb2 | 95,99 | -0,4086 | 0,296539208 | chr14 | - |
| Ripk3 | 25,38 | 0,0909 | 0,906066614 | chr14 | - |
| Rnase10 | 2,48 | 0,7187 | NA | chr14 | + |
| Rnase13 | 3,88 | 1,0320 | 0,058961472 | chr14 | - |
| Rnase4 | 29721,34 | -0,5346 | 0,156728357 | chr14 | + |
| Rnase6 | 23,19 | 0,6415 | 0,210781641 | chr14 | + |
| Rnaseh2b | 204,91 | -0,3076 | 0,403302288 | chr14 | + |
| Rnf219 | 26,04 | 0,6405 | 0,226526086 | chr14 | - |
| Rnf31 | 138,42 | -0,1050 | 0,840150496 | chr14 | + |
| Rpl15 | 6691,43 | -0,3990 | 0,196431964 | chr14 | - |
| Rpp14 | 356,33 | -0,0787 | 0,866208949 | chr14 | + |
| Rpph1 | 61,78 | -0,7408 | 0,174017104 | chr14 | - |
| Rps24 | 10776,13 | -0,4868 | 0,055404773 | chr14 | + |
| Samd4 | 37,92 | 0,4364 | 0,415700483 | chr14 | + |
| Samd8 | 880,13 | -0,0191 | 0,974449819 | chr14 | + |
| Sap18 | 343,60 | -0,2131 | 0,582500211 | chr14 | + |
| Saysd1 | 205,25 | -0,4398 | 0,37980999 | chr14 | - |
| Scara3 | 56,43 | 0,6922 | 0,177510828 | chr14 | - |
| Scara5 | 249,38 | 0,5558 | 0,409632036 | chr14 | + |
| Sdr39u1 | 157,76 | 0,4245 | 0,193696579 | chr14 | - |
| Sec24c | 1101,62 | -0,1346 | 0,692120295 | chr14 | + |
| Selenok | 2520,14 | 0,0647 | 0,883099159 | chr14 | + |
| Sema3g | 2,59 | -0,2609 | 0,726155531 | chr14 | + |
| Setdb2 | 237,55 | 0,4426 | 0,433006548 | chr14 | - |
| Sfmbt1 | 577,58 | -0,3403 | 0,231370727 | chr14 | + |
| Sh3bp5 | 742,58 | 0,5271 | 0,066396922 | chr14 | - |
| Shisa2 | 5,57 | 0,2721 | 0,738282454 | chr14 | + |
| Ska3 | 7,24 | 0,0357 | 0,973516404 | chr14 | - |
| Slain1 | 4,76 | 0,8153 | 0,167449045 | chr14 | + |
| Slc15a1 | 5,30 | 0,2393 | 0,773303025 | chr14 | - |
| Slc22a17 | 9,03 | -0,3514 | 0,634446856 | chr14 | - |
| Slc25a30 | 278,85 | 0,3725 | 0,562977316 | chr14 | - |
| Slc39a14 | 2519,95 | -0,0435 | 0,926388088 | chr14 | - |
| Slc39a2 | 18,10 | -0,5830 | 0,361307073 | chr14 | + |
| Slc4a7 | 134,45 | -0,2139 | 0,618224327 | chr14 | + |
| Slc7a7 | 82,31 | 0,2567 | 0,630381755 | chr14 | - |
| Slc7a8 | 46,23 | 0,2610 | 0,674998094 | chr14 | - |

| | | | | | |
|---|---|---|---|---|---|
| Slmap | 973,44 | -0,0899 | 0,816359983 | chr14 | - |
| Smim4 | 302,60 | -0,5943 | 0,220997862 | chr14 | - |
| Sncg | 3,95 | -0,0821 | 0,933919983 | chr14 | - |
| Snora31 | 9,52 | -0,1890 | 0,822888019 | chr14 | + |
| Snord69 | 3,20 | 0,0397 | 0,969159464 | chr14 | - |
| Socs4 | 57,35 | 0,3959 | 0,405665135 | chr14 | + |
| Sorbs3 | 474,98 | 0,5677 | 0,227218378 | chr14 | - |
| Sox7 | 50,98 | 0,8177 | 0,113991396 | chr14 | + |
| Spata13 | 335,27 | 0,3951 | 0,30944949 | chr14 | + |
| Spcs1 | 2709,12 | -0,1821 | 0,599696232 | chr14 | - |
| Spry2 | 16,11 | 0,0309 | 0,976127317 | chr14 | - |
| Spryd7 | 421,40 | -0,0366 | 0,935299476 | chr14 | - |
| Stab1 | 455,46 | -0,3168 | 0,390554711 | chr14 | - |
| Stc1 | 2,11 | 0,6137 | NA | chr14 | + |
| Stk24 | 209,16 | -0,0022 | 0,996826205 | chr14 | - |
| Styx | 146,60 | 0,2911 | 0,470739771 | chr14 | + |
| Sucla2 | 3565,74 | -0,2762 | 0,362293834 | chr14 | + |
| Sugt1 | 759,46 | -0,1548 | 0,67817958 | chr14 | + |
| Supt16 | 682,50 | 0,0045 | 0,992196557 | chr14 | - |
| Syt15 | 9,52 | 0,0288 | 0,979155888 | chr14 | + |
| Tbc1d4 | 49,38 | -0,2429 | 0,638713033 | chr14 | - |
| Tdrd3 | 169,48 | -0,2315 | 0,523118038 | chr14 | + |
| Tep1 | 191,48 | -0,0509 | 0,925898178 | chr14 | - |
| Tgds | 764,34 | -0,1359 | 0,744486882 | chr14 | - |
| Thoc7 | 976,00 | 0,0243 | 0,957017397 | chr14 | - |
| Thrb | 697,26 | 0,1290 | 0,796541442 | chr14 | + |
| Thtpa | 388,89 | -0,0836 | 0,874932716 | chr14 | + |
| Timm23 | 2033,70 | -0,3901 | 0,187852299 | chr14 | - |
| Tinf2 | 157,58 | -0,2397 | 0,566815451 | chr14 | - |
| Tkt | 1829,97 | -0,2589 | 0,517431746 | chr14 | + |
| Tlr11 | 7,75 | 0,7112 | 0,263764485 | chr14 | + |
| Tm9sf1 | 735,71 | -0,2040 | 0,516624153 | chr14 | - |
| Tm9sf2 | 3337,26 | -0,1658 | 0,630381755 | chr14 | + |
| Tmem110 | 130,23 | -0,0861 | 0,888917253 | chr14 | + |
| Tmem260 | 333,01 | 0,0793 | 0,855062146 | chr14 | + |
| Tmem55b | 309,68 | -0,2782 | 0,397488617 | chr14 | - |
| Tmtc4 | 81,97 | 0,3146 | 0,437620234 | chr14 | - |
| Tnfrsf10b | 13,11 | 0,2108 | 0,792197952 | chr14 | + |
| Tnfrsf19 | 70,16 | -0,6517 | 0,208536139 | chr14 | - |
| Tnnc1 | 22,52 | 0,5819 | 0,357345082 | chr14 | + |
| Top2b | 874,77 | -0,0397 | 0,938661865 | chr14 | + |
| Tox4 | 393,02 | 0,0492 | 0,915435 | chr14 | + |
| Tpt1 | 23100,12 | -0,5946 | 0,050838368 | chr14 | + |
| Trim13 | 37,16 | 0,3240 | 0,61795369 | chr14 | + |
| Trim35 | 78,89 | 0,2177 | 0,636447281 | chr14 | + |
| Tsc22d1 | 630,88 | -0,8014 | 0,055386722 | chr14 | + |
| Tspan14 | 396,90 | 0,3021 | 0,303750468 | chr14 | - |
| Ttc5 | 184,27 | 0,4725 | 0,255306897 | chr14 | - |
| Txndc16 | 106,89 | 0,2160 | 0,679975413 | chr14 | - |
| Ubac2 | 365,96 | -0,2391 | 0,556171512 | chr14 | + |

| | | | | | |
|---|---|---|---|---|---|
| Ube2e1 | 894,17 | -0,1285 | 0,759012157 | chr14 | - |
| Ube2e2 | 177,89 | -0,3685 | 0,317958264 | chr14 | - |
| Uchl3 | 381,27 | 0,0350 | 0,953911274 | chr14 | + |
| Uggt2 | 36,71 | 0,8182 | 0,101796122 | chr14 | - |
| Usp54 | 61,89 | 0,2204 | 0,717105204 | chr14 | - |
| Vcl | 529,12 | 0,1351 | 0,807278226 | chr14 | + |
| Vdac2 | 1522,13 | -0,3188 | 0,208577471 | chr14 | + |
| Vstm4 | 29,88 | 0,2683 | 0,646093772 | chr14 | + |
| Vwa8 | 2001,60 | -0,2730 | 0,457587064 | chr14 | + |
| Wapl | 902,95 | -0,0938 | 0,815149624 | chr14 | + |
| Wbp4 | 308,59 | -0,0291 | 0,952710435 | chr14 | - |
| Wdfy2 | 28,77 | 0,3856 | 0,483882557 | chr14 | + |
| Wdfy4 | 40,46 | 0,5288 | 0,228941765 | chr14 | - |
| Wdhd1 | 30,81 | -0,1521 | 0,832087093 | chr14 | - |
| Wnt5a | 6,85 | -0,1503 | 0,869020534 | chr14 | + |
| Xpo4 | 43,67 | 0,1768 | 0,76277049 | chr14 | - |
| Xpo7 | 718,61 | -0,1193 | 0,746827664 | chr14 | - |
| Zc3h13 | 282,45 | 0,2579 | 0,482009343 | chr14 | + |
| Zcchc24 | 1228,49 | 0,0586 | 0,89879469 | chr14 | - |
| Zdhhc20 | 358,65 | -0,0754 | 0,877283616 | chr14 | - |
| Zfhx2 | 62,43 | 0,4651 | 0,383114143 | chr14 | - |
| Zfp219 | 116,82 | -0,2191 | 0,754953629 | chr14 | - |
| Zfp395 | 357,36 | 0,1064 | 0,857478182 | chr14 | + |
| Zfp488 | 98,35 | -0,3476 | 0,416860553 | chr14 | - |
| Zfp503 | 21,31 | -0,0052 | 0,995836778 | chr14 | - |
| Zmiz1 | 268,13 | -0,2309 | 0,509917962 | chr14 | + |
| Zmym2 | 489,59 | -0,0825 | 0,862616891 | chr14 | + |
| Zmym5 | 615,48 | -0,1298 | 0,738098909 | chr14 | - |
| Zswim8 | 312,32 | -0,4304 | 0,344149271 | chr14 | + |
| 1110038F14Rik | 71,67 | 0,0605 | 0,937252102 | chr15 | + |
| 1700001L05Rik | 166,78 | 0,0396 | 0,942935841 | chr15 | - |
| 1700025B11Rik | 4,88 | -0,0651 | 0,948314012 | chr15 | - |
| 1700088E04Rik | 4,99 | 0,3625 | 0,631867311 | chr15 | - |
| 1810021B22Rik | 19,56 | -0,4977 | 0,403758222 | chr15 | - |
| 2010109I03Rik | 9,00 | -0,3358 | 0,663873104 | chr15 | - |
| 2410089E03Rik | 77,65 | 0,6562 | 0,132327977 | chr15 | + |
| 4921515E04Rik | 3,91 | -0,0339 | 0,974449819 | chr15 | - |
| 4930556M19Rik | 56,26 | 0,2197 | 0,657774838 | chr15 | + |
| 5031439G07Rik | 269,12 | 0,0502 | 0,919961951 | chr15 | - |
| 6030458C11Rik | 256,53 | -0,0248 | 0,964620526 | chr15 | - |
| 8430426J06Rik | 4,30 | 0,1836 | 0,83097362 | chr15 | - |
| 9130401M01Rik | 169,56 | -0,4524 | 0,281016665 | chr15 | - |
| 9330020H09Rik | 21,17 | -0,1779 | 0,800238939 | chr15 | + |
| 9930014A18Rik | 2,72 | 0,1393 | 0,871084117 | chr15 | + |
| A1bg | 9,63 | 1,0451 | 0,068097992 | chr15 | - |
| A330009N23Rik | 14,04 | 0,3005 | 0,671272065 | chr15 | - |
| A4galt | 5,26 | 0,3680 | 0,618723125 | chr15 | - |
| Aaas | 128,24 | -0,2773 | 0,522768066 | chr15 | - |
| Aard | 22,11 | 0,0987 | 0,903337956 | chr15 | + |
| Aco2 | 2702,89 | -0,1706 | 0,512939428 | chr15 | + |

| | | | | | |
|---|---|---|---|---|---|
| Acvr1b | 89,19 | 0,3198 | 0,449440511 | chr15 | + |
| Acvrl1 | 39,86 | 0,4083 | 0,382591532 | chr15 | + |
| Adamts12 | 4,02 | 0,3065 | 0,68361653 | chr15 | + |
| Adcy6 | 416,52 | -0,0986 | 0,859752964 | chr15 | - |
| Adsl | 207,90 | -0,0579 | 0,907270824 | chr15 | + |
| Ago2 | 58,88 | 0,3279 | 0,552949315 | chr15 | - |
| Agxt2 | 3241,87 | 0,3812 | 0,247584138 | chr15 | + |
| Alg12 | 75,04 | 0,1220 | 0,834507118 | chr15 | - |
| Amacr | 2228,66 | -0,3077 | 0,279128885 | chr15 | + |
| Amhr2 | 3,12 | 0,5897 | 0,339060647 | chr15 | + |
| Ank | 383,99 | 0,3659 | 0,291140498 | chr15 | + |
| Ankrd33b | 488,82 | -0,0358 | 0,950795699 | chr15 | - |
| Ankrd46 | 1178,18 | 0,2298 | 0,649854282 | chr15 | - |
| Ankrd54 | 71,84 | -0,2810 | 0,542478166 | chr15 | - |
| Ano6 | 128,81 | 0,1035 | 0,840150496 | chr15 | + |
| Apobec3 | 28,26 | 0,6285 | 0,220997862 | chr15 | + |
| Apol10a | 6,53 | 0,9098 | 0,110056031 | chr15 | + |
| Apol10b | 10,40 | 0,5685 | 0,391247721 | chr15 | - |
| Apol7a | 2307,73 | 0,0753 | 0,913293028 | chr15 | - |
| Apol7c | 5,52 | 0,6415 | 0,323355495 | chr15 | - |
| Apol7e | 2,39 | 0,0931 | NA | chr15 | + |
| Apol9a | 664,02 | 0,4982 | 0,287543614 | chr15 | - |
| Apol9b | 1043,07 | -0,4890 | 0,212297691 | chr15 | + |
| Arf3 | 956,79 | 0,1981 | 0,529603347 | chr15 | - |
| Arfgap3 | 111,80 | 0,0606 | 0,918946301 | chr15 | - |
| Arhgap39 | 29,13 | 0,2477 | 0,687070622 | chr15 | - |
| Arid2 | 265,67 | -0,0467 | 0,913017292 | chr15 | + |
| Asap1 | 147,32 | -0,1722 | 0,679380528 | chr15 | - |
| Asb8 | 360,79 | -0,1983 | 0,609093828 | chr15 | - |
| Atad2 | 92,92 | 0,3650 | 0,390859159 | chr15 | - |
| Atf1 | 385,77 | -0,0511 | 0,918104715 | chr15 | + |
| Atf7 | 22,28 | 0,9140 | 0,069527621 | chr15 | - |
| Atg101 | 625,35 | -0,0032 | 0,995836778 | chr15 | + |
| Atp5g2 | 3120,83 | -0,2360 | 0,456310852 | chr15 | - |
| Atp6v1c1 | 733,95 | -0,1935 | 0,505514525 | chr15 | + |
| Atxn10 | 1307,68 | 0,0477 | 0,907366343 | chr15 | + |
| AW549877 | 719,30 | 0,3730 | 0,274221528 | chr15 | - |
| Azin1 | 2758,59 | 0,0917 | 0,86802587 | chr15 | - |
| Basp1 | 17,64 | 0,6292 | 0,25536359 | chr15 | - |
| BC024139 | 78,14 | -0,6139 | 0,166418695 | chr15 | - |
| Bcdin3d | 42,24 | 0,3763 | 0,462351393 | chr15 | - |
| Bik | 132,11 | -0,0433 | 0,953911274 | chr15 | + |
| Bin2 | 47,99 | 0,4522 | 0,433006548 | chr15 | - |
| Bop1 | 267,81 | -0,3249 | 0,449196578 | chr15 | - |
| Brd1 | 299,33 | -0,3470 | 0,320685909 | chr15 | - |
| Brix1 | 490,36 | -0,0883 | 0,843029232 | chr15 | - |
| C030006K11Rik | 255,60 | -0,2418 | 0,50541673 | chr15 | - |
| C1ql4 | 2,15 | 0,7754 | NA | chr15 | - |
| C1qtnf3 | 7,91 | 0,0009 | 0,998994678 | chr15 | + |
| C1qtnf6 | 28,31 | 0,5554 | 0,312993461 | chr15 | - |

| | | | | | |
|---|---|---|---|---|---|
| C230037L18Rik | 2,48 | 0,1609 | NA | chr15 | + |
| C330013E15Rik | 10,88 | 0,5414 | 0,418345592 | chr15 | - |
| C6 | 3881,22 | -0,5700 | 0,235271103 | chr15 | + |
| C7 | 15,09 | 0,1589 | 0,854690386 | chr15 | - |
| Cacnb3 | 7,63 | 0,9299 | 0,118025078 | chr15 | + |
| Calcoco1 | 407,61 | -0,2681 | 0,502477511 | chr15 | - |
| Card10 | 146,44 | 0,3077 | 0,564849438 | chr15 | - |
| Card6 | 216,58 | 0,4701 | 0,137147675 | chr15 | - |
| Cbx5 | 531,73 | -0,4355 | 0,149536902 | chr15 | - |
| Cbx6 | 80,03 | 0,0663 | 0,906081073 | chr15 | - |
| Cbx7 | 76,19 | 0,0034 | 0,995750189 | chr15 | - |
| Cby1 | 120,91 | -0,1321 | 0,826227466 | chr15 | + |
| Ccdc134 | 50,83 | -0,3094 | 0,571350093 | chr15 | + |
| Ccdc166 | 14,93 | -0,2367 | 0,766152518 | chr15 | - |
| Ccnt1 | 49,48 | 0,7693 | 0,080029766 | chr15 | - |
| Cct5 | 3653,30 | -0,1247 | 0,699742099 | chr15 | - |
| Cdc42ep1 | 283,97 | 0,0628 | 0,912853091 | chr15 | + |
| Cdh18 | 4,41 | 0,5169 | 0,437620234 | chr15 | + |
| Cdpf1 | 80,67 | 0,2931 | 0,514682351 | chr15 | - |
| Cela1 | 673,92 | 0,1201 | 0,810346289 | chr15 | - |
| Celsr1 | 124,93 | 0,4460 | 0,408496057 | chr15 | - |
| Cenpm | 20,82 | 0,9840 | 0,069527621 | chr15 | - |
| Cerk | 56,18 | 0,3917 | 0,416860553 | chr15 | - |
| Cers5 | 45,42 | 0,2636 | 0,611914561 | chr15 | - |
| Chadl | 3,22 | -0,0382 | 0,969195493 | chr15 | - |
| Chkb | 20,41 | -0,9610 | 0,050277949 | chr15 | - |
| Chrac1 | 336,34 | -0,2231 | 0,588894341 | chr15 | + |
| Cmbl | 3843,10 | 0,2428 | 0,464245504 | chr15 | + |
| Commd5 | 243,04 | 0,1382 | 0,721569914 | chr15 | + |
| Copz1 | 3094,08 | -0,0909 | 0,843102154 | chr15 | + |
| Cox14 | 405,72 | -0,2404 | 0,71673612 | chr15 | + |
| Cox6c | 10105,75 | -0,4903 | 0,068746026 | chr15 | - |
| Cpq | 2013,43 | 0,1942 | 0,600688917 | chr15 | + |
| Cpsf1 | 354,03 | -0,3587 | 0,333886933 | chr15 | - |
| Creld2 | 832,39 | 0,3036 | 0,626412404 | chr15 | + |
| Csad | 5610,18 | 0,0679 | 0,904806119 | chr15 | - |
| Csdc2 | 4,80 | 0,3021 | 0,700925755 | chr15 | + |
| Csf2rb2 | 40,58 | 0,2121 | 0,761934432 | chr15 | - |
| Csnk1e | 63,70 | -0,2100 | 0,672907874 | chr15 | - |
| Csrnp2 | 29,89 | -0,4791 | 0,341412029 | chr15 | - |
| Cyb5r3 | 7005,09 | -0,0181 | 0,970419105 | chr15 | - |
| Cyp2d10 | 9548,08 | -0,3879 | 0,080350582 | chr15 | - |
| Cyp2d11 | 10,75 | -0,6938 | 0,273018284 | chr15 | - |
| Cyp2d22 | 3129,23 | -0,1235 | 0,728619917 | chr15 | - |
| Cyp2d34 | 2,82 | -0,3123 | 0,672918705 | chr15 | - |
| Cyp2d9 | 20379,17 | -0,0623 | 0,866934682 | chr15 | + |
| Cyth4 | 101,07 | 0,5467 | 0,100588364 | chr15 | + |
| Fam91a1 | 1473,40 | -0,0067 | 0,98947313 | chr15 | + |
| D730005E14Rik | 3,18 | 0,0335 | 0,975445719 | chr15 | - |
| Dab2 | 238,55 | 0,0642 | 0,898098997 | chr15 | + |

| | | | | | |
|---|---|---|---|---|---|
| Dap | 2061,58 | -0,4094 | 0,286640164 | chr15 | + |
| Dazap2 | 2005,21 | 0,0199 | 0,963400641 | chr15 | + |
| Dcaf13 | 693,97 | 0,0955 | 0,804754692 | chr15 | + |
| Ddx17 | 2111,95 | -0,1815 | 0,660992324 | chr15 | - |
| Ddx23 | 496,88 | -0,1836 | 0,565470336 | chr15 | - |
| Dennd3 | 57,47 | 0,3260 | 0,468677164 | chr15 | + |
| Dennd6b | 60,86 | -0,7463 | 0,095599478 | chr15 | - |
| Deptor | 3592,27 | -0,3611 | 0,233066911 | chr15 | + |
| Derl1 | 2763,65 | -0,2012 | 0,507474908 | chr15 | - |
| Desi1 | 626,81 | -0,3615 | 0,196118975 | chr15 | - |
| Dhh | 3,03 | 0,2908 | 0,695978671 | chr15 | - |
| Dip2b | 102,50 | 0,2791 | 0,526620417 | chr15 | + |
| Dnajc22 | 1979,66 | 0,0954 | 0,811137679 | chr15 | + |
| Dnal4 | 60,78 | 0,0418 | 0,950819236 | chr15 | - |
| Dpys | 3110,37 | -0,1912 | 0,511697151 | chr15 | - |
| Drosha | 116,04 | -0,3028 | 0,500274031 | chr15 | + |
| E330033B04Rik | 15,90 | -0,4744 | 0,481768228 | chr15 | - |
| Washc5 | 425,35 | 0,1800 | 0,67176872 | chr15 | - |
| Ebag9 | 407,88 | 0,3245 | 0,318113166 | chr15 | + |
| Eef1d | 1664,51 | -0,2597 | 0,354338694 | chr15 | - |
| Efr3a | 808,37 | -0,2312 | 0,394573343 | chr15 | + |
| Egflam | 8,29 | 0,1710 | 0,845508888 | chr15 | - |
| Eif3d | 790,04 | -0,3051 | 0,341871878 | chr15 | - |
| Eif3e | 2336,45 | -0,2260 | 0,555044685 | chr15 | - |
| Eif3h | 1929,76 | -0,1906 | 0,622736166 | chr15 | - |
| Eif3l | 1219,82 | -0,2968 | 0,279939661 | chr15 | + |
| Eif4b | 4479,03 | -0,2441 | 0,462797865 | chr15 | + |
| Elfn2 | 1,67 | 0,4838 | NA | chr15 | - |
| Emc2 | 1018,80 | -0,1778 | 0,661449625 | chr15 | + |
| Endou | 16,06 | 0,1402 | 0,869687361 | chr15 | - |
| Enpp2 | 1566,99 | -0,5449 | 0,062791791 | chr15 | - |
| Eny2 | 1818,06 | -0,1738 | 0,637616133 | chr15 | + |
| Ep300 | 230,77 | 0,0584 | 0,900327088 | chr15 | + |
| Eppk1 | 183,70 | 0,4794 | 0,217835326 | chr15 | - |
| Erich5 | 2,23 | -0,2576 | NA | chr15 | + |
| Espl1 | 3,57 | 0,2442 | 0,764363417 | chr15 | + |
| Exosc4 | 167,89 | -0,5577 | 0,25884542 | chr15 | + |
| Ext1 | 609,40 | -0,2470 | 0,442074728 | chr15 | - |
| Fam105a | 45,06 | 0,2614 | 0,658008155 | chr15 | - |
| Fam109b | 3,57 | -0,0920 | 0,922855195 | chr15 | + |
| Fam118a | 56,53 | 0,4397 | 0,402726862 | chr15 | + |
| Retreg1 | 2039,76 | 0,6817 | 0,111745857 | chr15 | + |
| Fam173b | 161,15 | -0,2128 | 0,567936438 | chr15 | + |
| Fam19a5 | 16,74 | 0,3034 | 0,675658116 | chr15 | + |
| Hgh1 | 65,76 | -0,2829 | 0,575906025 | chr15 | + |
| Fam227a | 13,48 | -0,3873 | 0,565929347 | chr15 | - |
| Fam49b | 172,98 | 0,0859 | 0,862616891 | chr15 | - |
| Fam83f | 22,58 | 0,5789 | 0,388402325 | chr15 | + |
| Fam83h | 90,34 | 0,1533 | 0,760825768 | chr15 | - |
| Fam84b | 104,45 | -0,0044 | 0,995142284 | chr15 | - |

| | | | | | |
|---|---|---|---|---|---|
| Fbln1 | 15,07 | 0,0888 | 0,923778914 | chr15 | + |
| Fbxl6 | 121,89 | -0,4208 | 0,279128885 | chr15 | - |
| Fbxl7 | 16,27 | -0,2033 | 0,793608383 | chr15 | - |
| Fbxo32 | 87,45 | 0,0475 | 0,944269942 | chr15 | - |
| Fbxo4 | 221,84 | -0,3980 | 0,186588546 | chr15 | - |
| Fbxo43 | 2,30 | 0,1059 | NA | chr15 | - |
| Fignl2 | 123,27 | 0,3111 | 0,394599707 | chr15 | - |
| Fkbp11 | 686,78 | 0,1541 | 0,738007269 | chr15 | - |
| Fmnl3 | 45,48 | 0,2181 | 0,705135797 | chr15 | - |
| Foxred2 | 4,79 | 0,4508 | 0,517431746 | chr15 | - |
| Fyb | 119,97 | 0,2916 | 0,438110291 | chr15 | + |
| Fzd6 | 58,58 | 0,1416 | 0,802314886 | chr15 | + |
| Galnt6 | 17,19 | 0,0128 | 0,989980239 | chr15 | - |
| Gcat | 1792,39 | 0,4721 | 0,25568151 | chr15 | + |
| Gga1 | 208,17 | -0,1858 | 0,709119038 | chr15 | + |
| Ghr | 9670,21 | -0,1669 | 0,705982644 | chr15 | - |
| Gm10865 | 2,87 | -0,0471 | 0,961556353 | chr15 | - |
| Mirt2 | 34,24 | -0,2736 | 0,705870256 | chr15 | - |
| Gm16576 | 4,53 | 0,2455 | 0,765061508 | chr15 | + |
| Gm19782 | 2,35 | 0,5561 | NA | chr15 | + |
| Gm20324 | 10,99 | -0,5184 | 0,435489536 | chr15 | + |
| Hnrnpa1l2-ps2 | 2,43 | -0,0783 | NA | chr15 | - |
| Gm8221 | 5,02 | 0,4236 | 0,562291762 | chr15 | + |
| Gm9920 | 4,09 | 0,1406 | 0,876634156 | chr15 | - |
| Golph3 | 1192,92 | -0,1667 | 0,651859853 | chr15 | + |
| Gpaa1 | 187,24 | -0,5776 | 0,163594932 | chr15 | + |
| Gpd1 | 5127,81 | -0,2118 | 0,666980604 | chr15 | + |
| Gpihbp1 | 509,22 | -0,0960 | 0,794944296 | chr15 | + |
| Gpt | 3584,50 | -0,0232 | 0,956727596 | chr15 | + |
| Gramd4 | 54,37 | 0,6008 | 0,162272166 | chr15 | + |
| Grap2 | 9,76 | 0,4907 | 0,483081622 | chr15 | + |
| Grasp | 16,99 | -0,2149 | 0,773299581 | chr15 | + |
| Grhl2 | 2,23 | 0,2120 | NA | chr15 | + |
| Grina | 1399,77 | -0,5747 | 0,131898706 | chr15 | + |
| Gsdmd | 562,63 | 0,4618 | 0,201155574 | chr15 | + |
| Gtpbp1 | 239,54 | -0,2898 | 0,39766425 | chr15 | + |
| Gtse1 | 9,94 | -0,5789 | 0,358287811 | chr15 | + |
| Gxylt1 | 288,95 | -0,3325 | 0,261264389 | chr15 | - |
| H1f0 | 1990,38 | 0,0831 | 0,839991734 | chr15 | + |
| Hdac10 | 133,25 | -0,0111 | 0,986661276 | chr15 | - |
| Hdac7 | 52,21 | -0,0889 | 0,89130564 | chr15 | - |
| Hnrnpa1 | 101,55 | -0,0618 | 0,916285969 | chr15 | + |
| Rida | 25885,40 | -0,0812 | 0,863101638 | chr15 | - |
| Hsf1 | 119,15 | -0,5334 | 0,185084413 | chr15 | + |
| Ift27 | 144,03 | -0,3226 | 0,370073748 | chr15 | - |
| Igfbp6 | 10,12 | -0,3437 | 0,64893397 | chr15 | + |
| Il2rb | 24,22 | 0,3974 | 0,483930201 | chr15 | - |
| Il7r | 15,07 | 0,6448 | 0,280299373 | chr15 | - |
| Irak4 | 151,61 | 0,2397 | 0,54401105 | chr15 | + |
| Itga5 | 203,77 | -0,0444 | 0,943696477 | chr15 | - |

| | | | | | |
|---|---|---|---|---|---|
| Itgb7 | 17,39 | 0,4605 | 0,468682808 | chr15 | - |
| Josd1 | 670,46 | -0,3671 | 0,192260732 | chr15 | - |
| Jrk | 12,67 | 0,6263 | 0,321255221 | chr15 | - |
| Kansl2 | 468,38 | -0,4916 | 0,088351774 | chr15 | - |
| Kctd17 | 53,50 | -0,0813 | 0,925898178 | chr15 | + |
| Kdelr3 | 11,60 | 0,9281 | 0,104816152 | chr15 | + |
| Khdrbs3 | 189,55 | 0,2179 | 0,616263238 | chr15 | + |
| Kif21a | 723,88 | -0,0234 | 0,965271905 | chr15 | - |
| Klf10 | 571,35 | 0,6319 | 0,240304485 | chr15 | - |
| Kmt2d | 320,17 | -0,0078 | 0,99137621 | chr15 | - |
| Krt18 | 3080,36 | 0,4040 | 0,183965146 | chr15 | + |
| Krt79 | 3,10 | 0,2697 | 0,688545912 | chr15 | - |
| Krt8 | 3595,18 | 0,0871 | 0,865391338 | chr15 | - |
| Krt80 | 2,31 | 0,4068 | NA | chr15 | - |
| L3mbtl2 | 198,25 | -0,2962 | 0,413323375 | chr15 | + |
| Laptm4b | 405,32 | -0,1610 | 0,641854125 | chr15 | + |
| Larp4 | 1288,17 | 0,0867 | 0,85496895 | chr15 | + |
| Rtl6 | 3,47 | 0,7003 | 0,254141333 | chr15 | - |
| Letmd1 | 35,42 | 0,0260 | 0,976528475 | chr15 | + |
| Lgals1 | 3,46 | -0,0364 | 0,971699327 | chr15 | + |
| Lgals2 | 123,04 | 0,1438 | 0,767097454 | chr15 | - |
| Lifr | 6393,33 | -0,3212 | 0,381025284 | chr15 | + |
| Lmbr1l | 226,75 | -0,3197 | 0,570187994 | chr15 | - |
| Lmf2 | 189,33 | -0,0820 | 0,895663283 | chr15 | - |
| Lrp12 | 60,94 | 0,0764 | 0,907233333 | chr15 | - |
| Lrrc14 | 93,19 | 0,0167 | 0,979212524 | chr15 | + |
| Lrrk2 | 32,95 | 0,5280 | 0,314431944 | chr15 | + |
| Ly6c1 | 15,84 | 0,8130 | 0,161930523 | chr15 | - |
| Ly6c2 | 54,67 | 0,0430 | 0,953751362 | chr15 | - |
| Ly6f | 12,50 | 0,5096 | 0,439006534 | chr15 | + |
| Ly6i | 5,83 | 0,3911 | 0,6016222 | chr15 | - |
| Lynx1 | 24,94 | 0,7642 | 0,143085377 | chr15 | - |
| Maf1 | 439,96 | -0,0126 | 0,981991128 | chr15 | + |
| Maff | 953,00 | -0,4955 | 0,340429934 | chr15 | + |
| Mal2 | 976,18 | -0,0521 | 0,926353842 | chr15 | + |
| Map3k12 | 10,60 | 0,0863 | 0,925382443 | chr15 | - |
| Mapk11 | 13,57 | -0,0231 | 0,981991128 | chr15 | - |
| Mapk12 | 50,84 | -0,3912 | 0,390780993 | chr15 | - |
| March6 | 2039,50 | -0,3902 | 0,099757726 | chr15 | - |
| Matn2 | 46,68 | 0,3729 | 0,533433508 | chr15 | + |
| Mcat | 291,73 | -0,2413 | 0,487461918 | chr15 | - |
| Mcrs1 | 93,23 | -0,0956 | 0,876857496 | chr15 | - |
| Med30 | 74,21 | 0,0198 | 0,978910193 | chr15 | + |
| Mettl7a1 | 4847,03 | -0,1965 | 0,525233119 | chr15 | + |
| Mettl7a2 | 6,15 | 0,7284 | 0,247921371 | chr15 | + |
| Methig1 | 2,84 | 0,6452 | 0,277831408 | chr15 | + |
| Mfng | 23,91 | 0,2067 | 0,755246573 | chr15 | - |
| Mfsd3 | 175,50 | 0,0245 | 0,969512928 | chr15 | + |
| Mfsd5 | 229,52 | -0,2784 | 0,523689387 | chr15 | + |
| Mgat3 | 4,19 | 0,2274 | 0,781051459 | chr15 | + |

| | | | | | |
|---|---|---|---|---|---|
| Micall1 | 157,43 | -0,1521 | 0,716811839 | chr15 | + |
| Mief1 | 460,85 | -0,3631 | 0,162648376 | chr15 | + |
| Lncppara | 7,67 | -0,0421 | 0,969130697 | chr15 | + |
| Mkl1 | 52,97 | -0,3547 | 0,521462295 | chr15 | - |
| Mpst | 830,95 | -0,5159 | 0,170992427 | chr15 | + |
| Mroh1 | 228,09 | -0,4505 | 0,248669578 | chr15 | + |
| Mroh6 | 13,88 | 0,3071 | 0,690021525 | chr15 | - |
| Mrpl13 | 1075,68 | -0,2763 | 0,376984775 | chr15 | - |
| Mtbp | 27,32 | 0,1763 | 0,811137679 | chr15 | + |
| Mtdh | 790,26 | 0,2290 | 0,483930201 | chr15 | + |
| Mtmr12 | 90,03 | -0,0286 | 0,960688549 | chr15 | + |
| Mtss1 | 2896,81 | -0,0179 | 0,976304651 | chr15 | - |
| Myc | 37,08 | -0,1546 | 0,848389544 | chr15 | + |
| Myg1 | 230,73 | -0,1804 | 0,633486481 | chr15 | + |
| Myh9 | 1204,88 | 0,2564 | 0,501000077 | chr15 | - |
| Myo10 | 441,17 | 0,1444 | 0,717851252 | chr15 | + |
| Nadk2 | 3203,24 | 0,1726 | 0,671790933 | chr15 | + |
| Naga | 384,73 | -0,1383 | 0,721421427 | chr15 | - |
| Naprt | 1188,91 | -0,2657 | 0,533903589 | chr15 | - |
| Ncaph2 | 887,49 | -0,1048 | 0,765219833 | chr15 | + |
| Ncf4 | 22,85 | -0,0031 | 0,997338104 | chr15 | + |
| Nckap1l | 156,82 | 0,4049 | 0,278686919 | chr15 | + |
| Nckap5l | 3,37 | 0,0811 | 0,93056206 | chr15 | - |
| Ndrg1 | 154,33 | -0,5606 | 0,066915827 | chr15 | - |
| Ndufa6 | 3957,87 | -0,3304 | 0,595460103 | chr15 | - |
| Ndufb9 | 4546,73 | -0,5130 | 0,159562415 | chr15 | + |
| Nfam1 | 52,10 | 0,3770 | 0,435695618 | chr15 | - |
| Nfe2 | 4,44 | 0,2581 | 0,747375272 | chr15 | - |
| Snu13 | 919,24 | -0,0540 | 0,896013441 | chr15 | - |
| Nipal2 | 8,95 | 0,5757 | 0,380302201 | chr15 | - |
| Nipbl | 635,74 | 0,1468 | 0,676716141 | chr15 | - |
| Nol12 | 144,34 | -0,1289 | 0,814156035 | chr15 | + |
| Nov | 3,80 | -0,4204 | 0,552537331 | chr15 | + |
| Npff | 19,56 | -0,8350 | 0,12803328 | chr15 | - |
| Npr3 | 49,95 | 0,3560 | 0,581990132 | chr15 | - |
| Nr4a1 | 69,35 | 0,5048 | 0,452286381 | chr15 | + |
| Nsmce2 | 236,48 | 0,5242 | 0,094281875 | chr15 | + |
| Nudcd1 | 383,33 | 0,1136 | 0,828422381 | chr15 | - |
| Nup155 | 407,82 | 0,1826 | 0,610189036 | chr15 | + |
| Nup50 | 598,06 | -0,3033 | 0,310273443 | chr15 | + |
| Odf3b | 51,22 | 0,2895 | 0,63580052 | chr15 | - |
| Oplah | 1608,54 | -0,2257 | 0,646093772 | chr15 | - |
| Osmr | 83,18 | 0,2689 | 0,587879802 | chr15 | - |
| Otulin | 648,54 | -0,4989 | 0,169324582 | chr15 | - |
| Oxr1 | 1165,40 | 0,2122 | 0,543779254 | chr15 | + |
| Pabpc1 | 3046,52 | -0,3434 | 0,135247543 | chr15 | - |
| Pacsin2 | 693,72 | -0,1138 | 0,77333268 | chr15 | - |
| Parp10 | 265,72 | 0,3017 | 0,452062186 | chr15 | - |
| Parvb | 36,30 | 0,2601 | 0,67311371 | chr15 | + |
| Parvg | 30,75 | 0,8423 | 0,057877889 | chr15 | + |

| | | | | | |
|---|---|---|---|---|---|
| Pcbp2 | 2676,52 | -0,5140 | 0,05129496 | chr15 | + |
| Pced1b | 19,15 | 0,1100 | 0,890761998 | chr15 | + |
| Pde1b | 16,06 | 0,6642 | 0,235271103 | chr15 | + |
| Pdgfb | 13,27 | 0,1506 | 0,859809354 | chr15 | - |
| Pdxp | 19,53 | 1,0835 | 0,052169847 | chr15 | + |
| Pdzd2 | 13,76 | 0,5180 | 0,411295797 | chr15 | - |
| Peg13 | 60,79 | 0,4329 | 0,376013337 | chr15 | - |
| Pfdn5 | 2423,28 | -0,1504 | 0,644046286 | chr15 | + |
| Pfkm | 269,41 | -0,1967 | 0,571110938 | chr15 | + |
| Phf20l1 | 435,89 | -0,2315 | 0,545123886 | chr15 | + |
| Phf21b | 3,87 | -0,2858 | 0,715754232 | chr15 | - |
| Phf5a | 600,03 | -0,3118 | 0,379417606 | chr15 | - |
| Pick1 | 152,68 | -0,2199 | 0,631914678 | chr15 | + |
| Pkhd1l1 | 4,60 | 0,4484 | 0,526620417 | chr15 | + |
| Pla2g6 | 389,48 | -0,3797 | 0,416450614 | chr15 | - |
| Plec | 400,53 | -0,4240 | 0,352333619 | chr15 | - |
| Plxnb2 | 1632,63 | -0,1238 | 0,744313101 | chr15 | - |
| Pnpla3 | 29,42 | 0,8293 | NA | chr15 | + |
| Pnpla5 | 15,08 | 1,7676 | NA | chr15 | - |
| Poldip3 | 473,21 | 0,1202 | 0,761282414 | chr15 | - |
| Polr2f | 441,87 | 0,0313 | 0,965429748 | chr15 | + |
| Polr2k | 753,31 | 0,1811 | 0,666391696 | chr15 | + |
| Polr3h | 136,31 | 0,1223 | 0,811137679 | chr15 | - |
| Pop1 | 85,55 | 0,0635 | 0,919467514 | chr15 | + |
| Pou6f1 | 63,89 | -0,3473 | 0,424001874 | chr15 | - |
| Ppara | 4737,73 | -0,1070 | 0,813415373 | chr15 | + |
| Pphln1 | 275,02 | 0,0435 | 0,931322892 | chr15 | + |
| Ppp1r16a | 237,63 | -0,4665 | 0,290028383 | chr15 | + |
| Ppp6r2 | 182,44 | 0,0110 | 0,98507181 | chr15 | + |
| Prickle1 | 55,13 | -0,0523 | 0,944541717 | chr15 | - |
| Prkaa1 | 243,52 | 0,0846 | 0,857400768 | chr15 | + |
| Prkag1 | 609,58 | -0,1564 | 0,671461631 | chr15 | - |
| Prpf40b | 98,92 | -0,3233 | 0,537935014 | chr15 | + |
| Prph | 2,47 | 0,4499 | NA | chr15 | + |
| Prr13 | 677,38 | -0,2239 | 0,565034228 | chr15 | + |
| Prr5 | 31,98 | -0,4159 | 0,440094295 | chr15 | + |
| Ptger4 | 8,66 | 0,2538 | 0,755670612 | chr15 | - |
| Ptk2 | 260,01 | 0,0216 | 0,964760801 | chr15 | - |
| Ptp4a3 | 89,53 | 0,7667 | 0,10463342 | chr15 | + |
| Puf60 | 529,05 | -0,1656 | 0,69672367 | chr15 | - |
| Pus7l | 26,39 | 0,2588 | 0,685921923 | chr15 | - |
| Pvt1 | 32,05 | -0,8120 | 0,128835828 | chr15 | + |
| Pycrl | 440,10 | -0,5496 | 0,082128644 | chr15 | - |
| Rac2 | 124,46 | -0,0240 | 0,969414042 | chr15 | - |
| Rad1 | 115,75 | -0,3078 | 0,479400504 | chr15 | + |
| Racgap1 | 16,47 | -0,3420 | 0,62106443 | chr15 | - |
| Rad21 | 857,42 | 0,1917 | 0,544004738 | chr15 | - |
| Rai14 | 311,85 | -0,0629 | 0,90048192 | chr15 | - |
| Ranbp3l | 19,38 | -0,8012 | 0,157077211 | chr15 | + |
| Rangap1 | 739,70 | -0,4096 | 0,325000508 | chr15 | - |

| | | | | | |
|---|---|---|---|---|---|
| Rapgef3 | 24,06 | -0,0506 | 0,953764743 | chr15 | - |
| Rarg | 51,19 | 0,4044 | 0,445734083 | chr15 | - |
| Rbfox2 | 274,86 | 0,3602 | 0,219278353 | chr15 | - |
| Rbx1 | 1225,27 | -0,2619 | 0,38215614 | chr15 | + |
| Recql4 | 17,63 | 0,5235 | 0,434164637 | chr15 | - |
| Rhebl1 | 13,06 | -0,0275 | 0,979051488 | chr15 | - |
| Rictor | 276,35 | -0,0184 | 0,972380647 | chr15 | + |
| Rims2 | 3,80 | -0,4259 | 0,553833033 | chr15 | + |
| Rnd1 | 176,52 | -0,0470 | 0,953751362 | chr15 | - |
| Rnf139 | 1174,30 | 0,1114 | 0,784087358 | chr15 | + |
| Rnf19a | 527,45 | -0,1489 | 0,666980604 | chr15 | - |
| Ropn1l | 158,55 | 0,1745 | 0,737094181 | chr15 | - |
| Rpap3 | 175,00 | 0,6613 | 0,077179074 | chr15 | - |
| Rpl30 | 5,69 | -0,0767 | 0,939403124 | chr15 | - |
| Rpl8 | 4126,59 | -0,6945 | 0,07229395 | chr15 | + |
| Rps19bp1 | 140,70 | -0,6223 | 0,093632322 | chr15 | - |
| Rrm2b | 76,66 | 0,3843 | 0,449823999 | chr15 | - |
| Rrp7a | 228,64 | -0,1968 | 0,682099278 | chr15 | - |
| Samm50 | 1598,31 | -0,1521 | 0,752586238 | chr15 | + |
| Sbf1 | 273,22 | 0,1606 | 0,737094181 | chr15 | - |
| Scaf11 | 659,24 | -0,1345 | 0,682080939 | chr15 | - |
| Scn8a | 6,47 | 0,2019 | 0,814156035 | chr15 | + |
| Sco2 | 190,38 | -0,3898 | 0,317387674 | chr15 | - |
| Scrib | 166,03 | -0,4651 | 0,387186245 | chr15 | - |
| Scube1 | 4,43 | 0,8178 | 0,169529667 | chr15 | - |
| Sdc2 | 4097,67 | 0,2431 | 0,347052012 | chr15 | + |
| Selenoo | 1143,87 | -0,3225 | 0,319080738 | chr15 | + |
| Sema5a | 19,53 | -0,0834 | 0,922904986 | chr15 | + |
| Senp1 | 106,29 | -0,2623 | 0,524801822 | chr15 | - |
| Selenop | 111243,11 | -0,1360 | 0,617490375 | chr15 | + |
| Serhl | 505,99 | -0,0055 | 0,991295673 | chr15 | + |
| Sgsm3 | 84,60 | -0,3794 | 0,4345327 | chr15 | + |
| Sh3bp1 | 10,89 | 0,6106 | 0,338543801 | chr15 | + |
| Shank3 | 38,76 | 0,1290 | 0,86309397 | chr15 | + |
| Sharpin | 264,00 | -0,3762 | 0,319080738 | chr15 | - |
| Skp2 | 38,16 | 0,6229 | 0,145168891 | chr15 | - |
| Sla | 84,09 | 0,3148 | 0,607677923 | chr15 | - |
| Slc11a2 | 406,14 | -0,3246 | 0,377944963 | chr15 | - |
| Slc1a3 | 3,92 | -0,1624 | 0,847796138 | chr15 | - |
| Slc25a17 | 1679,44 | -0,3397 | 0,425370435 | chr15 | - |
| Slc25a32 | 321,43 | 0,4437 | 0,301101755 | chr15 | - |
| Slc2a13 | 8,14 | 0,1462 | 0,871713688 | chr15 | - |
| Slc30a8 | 4,00 | 0,9551 | 0,093284325 | chr15 | + |
| Slc38a1 | 24,29 | 0,3356 | 0,567046015 | chr15 | - |
| Slc38a4 | 27061,20 | -0,2303 | 0,554660735 | chr15 | - |
| Slc39a4 | 1310,69 | 0,0009 | 0,998482911 | chr15 | - |
| Slc45a2 | 3,06 | 0,1046 | 0,903337956 | chr15 | + |
| Slc45a4 | 72,31 | 0,1164 | 0,845910956 | chr15 | - |
| Slc48a1 | 703,61 | -0,2008 | 0,557547051 | chr15 | + |
| Slc4a8 | 10,03 | -0,4823 | 0,475773557 | chr15 | + |

| | | | | | |
|---|---|---|---|---|---|
| Smagp | 196,31 | 0,0334 | 0,951806422 | chr15 | - |
| Smarcd1 | 39,73 | -0,2419 | 0,681592614 | chr15 | + |
| Smc1b | 14,82 | -0,3923 | 0,583144822 | chr15 | - |
| Smdt1 | 895,98 | -0,3736 | 0,477326847 | chr15 | + |
| Smug1 | 194,95 | 0,0730 | 0,864697132 | chr15 | - |
| Snhg18 | 47,29 | -0,0415 | 0,955234619 | chr15 | - |
| Snord72 | 4,36 | -0,5144 | 0,451533216 | chr15 | + |
| Snord83b | 2,75 | 0,2590 | 0,738918887 | chr15 | - |
| Soat2 | 743,60 | 0,1447 | 0,76277049 | chr15 | + |
| Sp1 | 118,87 | 0,1760 | 0,700754601 | chr15 | + |
| Spag1 | 13,50 | 0,0190 | 0,985488456 | chr15 | + |
| Spats2 | 38,20 | 0,2186 | 0,7009274 | chr15 | + |
| Spryd3 | 309,54 | 0,4631 | 0,186556074 | chr15 | - |
| Sqle | 984,83 | -0,3979 | 0,408202808 | chr15 | + |
| Srebf2 | 350,04 | 0,2715 | 0,527609356 | chr15 | + |
| Stk3 | 231,47 | 0,3410 | 0,423164161 | chr15 | - |
| Sub1 | 1966,54 | 0,0711 | 0,874932716 | chr15 | - |
| Sun2 | 1123,71 | -0,0868 | 0,861767698 | chr15 | - |
| Syngr1 | 7,74 | 0,6776 | 0,293001163 | chr15 | + |
| Tab1 | 47,41 | 0,5488 | 0,195952045 | chr15 | + |
| Taf2 | 247,43 | -0,2916 | 0,413323375 | chr15 | - |
| Tarbp2 | 175,16 | -0,4755 | 0,115975046 | chr15 | + |
| Tars | 1770,80 | -0,0483 | 0,921657556 | chr15 | - |
| Tatdn1 | 258,38 | 0,0275 | 0,960601359 | chr15 | - |
| Tbc1d22a | 188,25 | -0,1519 | 0,709119038 | chr15 | + |
| Tbc1d31 | 145,99 | 0,6878 | 0,092923763 | chr15 | + |
| Tcf20 | 320,92 | 0,1283 | 0,763240039 | chr15 | - |
| Tef | 2552,84 | -0,2970 | 0,239792389 | chr15 | + |
| Tns2 | 592,55 | -0,6025 | 0,227017349 | chr15 | + |
| Tfcp2 | 67,68 | 0,0111 | 0,988414083 | chr15 | - |
| Them6 | 18,09 | -0,0720 | 0,939351122 | chr15 | + |
| Tigd5 | 24,51 | -0,2554 | 0,689465734 | chr15 | + |
| Tmbim6 | 15771,13 | -0,1440 | 0,748732136 | chr15 | + |
| Tmem106c | 255,11 | 0,1994 | 0,617155778 | chr15 | + |
| Tmem117 | 2,40 | 0,1027 | NA | chr15 | + |
| Tmem184b | 238,09 | -0,1401 | 0,800238939 | chr15 | - |
| Tmem65 | 132,01 | -0,1579 | 0,695967514 | chr15 | - |
| Tmem71 | 30,88 | 0,8968 | 0,096593506 | chr15 | - |
| Tmprss6 | 6213,30 | -0,1178 | 0,783264136 | chr15 | - |
| Tnrc6b | 446,76 | 0,0594 | 0,895663283 | chr15 | + |
| Tob2 | 432,22 | -0,0124 | 0,986692637 | chr15 | - |
| Tomm22 | 1068,50 | -0,2013 | 0,437620234 | chr15 | + |
| Tonsl | 7,19 | -0,3251 | 0,671272065 | chr15 | - |
| Top1mt | 219,08 | -0,0805 | 0,848486057 | chr15 | - |
| Trabd | 1004,01 | -0,4100 | 0,381800527 | chr15 | + |
| Trappc9 | 123,28 | -0,3398 | 0,472408228 | chr15 | - |
| Trib1 | 557,03 | -0,1224 | 0,811801336 | chr15 | + |
| Trio | 127,78 | 0,2227 | 0,607677923 | chr15 | - |
| Triobp | 130,66 | -0,2611 | 0,6951078 | chr15 | + |
| Trmu | 93,29 | -0,2587 | 0,52307695 | chr15 | + |

| | | | | | |
|---|---|---|---|---|---|
| Trps1 | 16,64 | 0,5353 | 0,403302288 | chr15 | - |
| Tspo | 1092,76 | -0,1418 | 0,820051729 | chr15 | + |
| Tspyl5 | 3,98 | -0,8599 | 0,153470778 | chr15 | - |
| Tst | 3460,57 | -0,1673 | 0,751097875 | chr15 | - |
| Tsta3 | 256,54 | 0,0098 | 0,984995389 | chr15 | - |
| Ttc38 | 1638,12 | 0,2770 | 0,360045563 | chr15 | + |
| Ttll1 | 44,49 | -0,2818 | 0,580137509 | chr15 | - |
| Ttll12 | 102,86 | -0,0841 | 0,894535754 | chr15 | - |
| Ttll8 | 5,99 | -0,3408 | 0,653708291 | chr15 | - |
| Tuba1a | 62,75 | 0,5706 | 0,215519599 | chr15 | - |
| Tuba1b | 295,56 | 0,1220 | 0,783422709 | chr15 | - |
| Tuba1c | 289,57 | -0,1888 | 0,745081091 | chr15 | + |
| Tubgcp6 | 113,88 | -0,1612 | 0,732937334 | chr15 | - |
| Twf1 | 1173,83 | -0,0074 | 0,988512389 | chr15 | - |
| Txn2 | 3222,93 | -0,3131 | 0,271262155 | chr15 | - |
| Tymp | 301,22 | 0,0992 | 0,869067375 | chr15 | - |
| Ubr5 | 1266,83 | -0,3175 | 0,271880887 | chr15 | - |
| Ugt3a1 | 2374,55 | -0,1746 | 0,66585613 | chr15 | + |
| Utp23 | 134,63 | 0,2780 | 0,428072237 | chr15 | + |
| Vps13b | 646,73 | -0,3133 | 0,356901218 | chr15 | + |
| Vps28 | 1093,08 | -0,0572 | 0,890887823 | chr15 | - |
| Wdr70 | 56,86 | -0,3002 | 0,604491975 | chr15 | - |
| Wdyhv1 | 124,28 | 0,0411 | 0,937947106 | chr15 | + |
| Wisp1 | 27,94 | 0,7560 | 0,146101708 | chr15 | + |
| Xpnpep3 | 131,56 | 0,1825 | 0,738937653 | chr15 | + |
| Xrcc6 | 425,31 | 0,1201 | 0,729244932 | chr15 | + |
| Yaf2 | 219,13 | -0,3673 | 0,430392697 | chr15 | - |
| Ywhaz | 2043,25 | -0,1634 | 0,599987908 | chr15 | - |
| Zbed4 | 86,97 | 0,0139 | 0,98507181 | chr15 | + |
| Zc3h3 | 52,51 | -0,0038 | 0,99636982 | chr15 | - |
| Zc3h7b | 392,06 | 0,4348 | 0,072528427 | chr15 | + |
| Zfat | 22,45 | 0,2036 | 0,763597696 | chr15 | - |
| Zfp251 | 30,82 | 0,5074 | 0,340674125 | chr15 | - |
| Zfp385a | 119,66 | 0,0137 | 0,986818215 | chr15 | - |
| Zfp41 | 27,07 | 0,8690 | 0,114525976 | chr15 | + |
| Zfp622 | 370,86 | -0,1386 | 0,715470876 | chr15 | + |
| Zfp623 | 205,58 | -0,3311 | 0,347104845 | chr15 | + |
| Zfp641 | 3,58 | -0,6256 | 0,317417662 | chr15 | - |
| Zfp647 | 11,78 | -0,0674 | 0,945208438 | chr15 | - |
| Zfp7 | 36,64 | 0,0586 | 0,943064069 | chr15 | + |
| Zfp706 | 1493,79 | -0,4023 | 0,110800147 | chr15 | - |
| Zfp707 | 262,19 | -0,4548 | 0,447848007 | chr15 | + |
| Zfp740 | 350,57 | 0,0493 | 0,912418045 | chr15 | + |
| Zfpm2 | 7,98 | 0,0770 | 0,93871506 | chr15 | + |
| Zfr | 717,22 | 0,0024 | 0,99636982 | chr15 | + |
| Zhx1 | 800,40 | -0,0926 | 0,828675276 | chr15 | - |
| Zhx2 | 170,64 | -0,1549 | 0,754953629 | chr15 | + |
| 0610012G03Rik | 192,50 | -0,5829 | 0,1100164 | chr16 | - |
| 1110004E09Rik | 212,30 | -0,4480 | 0,242678725 | chr16 | - |
| Lppos | 108,91 | -0,4861 | 0,194460708 | chr16 | - |

| | | | | | |
|---|---|---|---|---|---|
| 1300002E11Rik | 189,96 | -0,1028 | 0,820554805 | chr16 | + |
| 1700007L15Rik | 34,52 | -0,4850 | 0,302314196 | chr16 | - |
| Rubcn | 229,15 | 0,0974 | 0,816914587 | chr16 | - |
| 1700037C18Rik | 7,69 | 0,9557 | 0,103014216 | chr16 | - |
| 1700119H24Rik | 15,54 | 0,6122 | 0,330791426 | chr16 | - |
| 1810013L24Rik | 841,80 | -0,4818 | 0,073280923 | chr16 | + |
| 1810053B23Rik | 27,01 | 0,4267 | 0,546882594 | chr16 | - |
| Zbtb11os1 | 25,37 | -0,3241 | 0,617869186 | chr16 | - |
| 2510002D24Rik | 181,46 | -0,5859 | 0,163729985 | chr16 | + |
| 2510009E07Rik | 140,62 | 0,3744 | 0,253291899 | chr16 | - |
| Ccdc191 | 73,84 | 0,0635 | 0,917616895 | chr16 | + |
| Mir99ahg | 8,41 | 0,0170 | 0,987278221 | chr16 | + |
| 2900011O08Rik | 2,78 | 0,2672 | 0,696074872 | chr16 | + |
| 3110001I22Rik | 46,09 | -0,1892 | 0,75912084 | chr16 | + |
| 4632428C04Rik | 15,16 | -0,0635 | 0,944541717 | chr16 | + |
| 4930451G09Rik | 5,55 | 0,0579 | 0,953314701 | chr16 | + |
| 4930453N24Rik | 349,55 | 0,0002 | 0,999351454 | chr16 | - |
| Dubr | 4,49 | 0,5722 | 0,392217524 | chr16 | - |
| A630089N07Rik | 147,57 | 0,4651 | 0,131674744 | chr16 | - |
| Morf4l1-ps1 | 18,62 | -0,4419 | 0,513222464 | chr16 | + |
| A930006K02Rik | 4,23 | -0,1998 | 0,812543852 | chr16 | + |
| Abat | 3917,65 | -0,1333 | 0,756319626 | chr16 | + |
| Abcc1 | 26,83 | -0,0591 | 0,941757244 | chr16 | + |
| Abcc5 | 49,40 | 0,5321 | 0,25884542 | chr16 | - |
| Abcf3 | 784,20 | -0,1858 | 0,62632688 | chr16 | + |
| Abhd10 | 118,99 | -0,1619 | 0,713100089 | chr16 | - |
| Abi3bp | 70,18 | 0,2467 | 0,587971243 | chr16 | + |
| Acap2 | 279,74 | 0,2633 | 0,357345082 | chr16 | - |
| Adamts1 | 137,34 | 0,4399 | 0,208689106 | chr16 | - |
| Adamts5 | 37,90 | -0,0606 | 0,93807111 | chr16 | - |
| Adcy5 | 9,02 | 0,8157 | 0,183723189 | chr16 | + |
| Adcy9 | 349,73 | 0,0025 | 0,996644671 | chr16 | - |
| Adprh | 82,93 | 0,0059 | 0,993445259 | chr16 | - |
| Ahsg | 245204,92 | -0,0147 | 0,974008692 | chr16 | + |
| Aifm3 | 52,19 | -0,5405 | 0,217465538 | chr16 | + |
| Alg1 | 245,50 | -0,3342 | 0,263205799 | chr16 | + |
| Alg3 | 142,56 | -0,3311 | 0,45432671 | chr16 | - |
| Anks3 | 126,90 | -0,4328 | 0,335637765 | chr16 | - |
| Ap2m1 | 3616,59 | -0,2058 | 0,535475471 | chr16 | + |
| App | 767,54 | 0,0510 | 0,899148878 | chr16 | - |
| Arhgap31 | 120,49 | 0,1966 | 0,665144558 | chr16 | - |
| Arl13b | 120,22 | 0,1454 | 0,766299799 | chr16 | - |
| Arl6 | 97,72 | 0,6823 | 0,05129496 | chr16 | - |
| Arvcf | 125,76 | 0,1263 | 0,834675238 | chr16 | + |
| Atg3 | 2168,39 | -0,2260 | 0,535573923 | chr16 | + |
| Atp13a3 | 2398,36 | 0,0801 | 0,854690386 | chr16 | - |
| Atp5j | 8104,99 | -0,3328 | 0,227688754 | chr16 | - |
| Atp5o | 5293,16 | -0,2826 | 0,217231457 | chr16 | - |
| Atp6v1a | 1152,11 | 0,1906 | 0,526620417 | chr16 | - |
| AU021092 | 34,61 | 0,5121 | 0,303046806 | chr16 | - |

| | | | | | |
|---|---|---|---|---|---|
| B130034C11Rik | 9,05 | -0,1381 | 0,877942095 | chr16 | + |
| B3gnt5 | 3,63 | 0,0147 | 0,988512389 | chr16 | + |
| B4galt4 | 37,00 | 0,4357 | 0,370069913 | chr16 | + |
| Bace2 | 118,87 | 0,4897 | 0,15168644 | chr16 | + |
| Bach1 | 680,79 | -0,4124 | 0,212149875 | chr16 | + |
| Bbx | 202,58 | 0,1697 | 0,753582185 | chr16 | - |
| BC002163 | 11,63 | -0,1432 | 0,872326129 | chr16 | + |
| Nepro | 79,45 | -0,3570 | 0,342277341 | chr16 | + |
| BC031361 | 6,08 | -0,1160 | 0,901445192 | chr16 | - |
| Bcl6 | 373,66 | -0,0530 | 0,957318111 | chr16 | - |
| Bdh1 | 4456,97 | -0,0769 | 0,869442758 | chr16 | + |
| Bfar | 1219,67 | -0,3026 | 0,272751761 | chr16 | + |
| Boc | 4,13 | -0,1757 | 0,840418613 | chr16 | - |
| Brwd1 | 426,97 | -0,1346 | 0,695935333 | chr16 | - |
| Btla | 18,32 | -0,0962 | 0,906240001 | chr16 | + |
| C2cd2 | 1761,25 | -0,5250 | 0,0545287 | chr16 | - |
| C330027C09Rik | 33,28 | 0,6350 | 0,193343482 | chr16 | + |
| Camk2n2 | 12,25 | -0,4454 | 0,512609895 | chr16 | - |
| Car15 | 7,16 | -0,4872 | 0,483930201 | chr16 | - |
| Carhsp1 | 2062,67 | 0,1968 | 0,45607208 | chr16 | - |
| Cblb | 115,00 | -0,3900 | 0,452418751 | chr16 | + |
| Cbr1 | 987,01 | -0,3914 | 0,413156981 | chr16 | + |
| Cbr3 | 10,85 | -0,0714 | 0,942982044 | chr16 | + |
| Ccdc14 | 18,37 | 0,6931 | 0,208540118 | chr16 | + |
| Ccdc50 | 1101,71 | -0,0609 | 0,869442758 | chr16 | + |
| Ccdc58 | 1113,80 | -0,3178 | 0,417835365 | chr16 | + |
| Ccdc80 | 244,88 | 0,3163 | 0,462797865 | chr16 | + |
| Cct8 | 3511,56 | 0,0495 | 0,893706091 | chr16 | - |
| Cd200 | 23,16 | 0,3105 | 0,617219405 | chr16 | - |
| Cd200r1 | 16,75 | 0,7999 | 0,132461478 | chr16 | + |
| Cd200r4 | 3,62 | 0,3764 | 0,587065518 | chr16 | + |
| Cd47 | 672,27 | -0,4895 | 0,136585635 | chr16 | + |
| Cd80 | 9,77 | 0,4592 | 0,517431746 | chr16 | + |
| Cd86 | 149,75 | 0,3560 | 0,346467419 | chr16 | - |
| Cd96 | 4,80 | -0,3364 | 0,655923828 | chr16 | - |
| Cdc45 | 25,56 | -0,4493 | 0,500881372 | chr16 | - |
| Cdip1 | 1406,97 | 0,2221 | 0,582630294 | chr16 | - |
| Cebpd | 33,03 | -0,3162 | 0,637721381 | chr16 | + |
| Cep19 | 112,79 | -0,1040 | 0,829255454 | chr16 | + |
| Cep97 | 72,52 | -0,3510 | 0,43417568 | chr16 | - |
| Cggbp1 | 899,51 | -0,1880 | 0,523048388 | chr16 | + |
| Chaf1b | 30,40 | 0,0427 | 0,955803452 | chr16 | + |
| Chmp2b | 431,54 | 0,1264 | 0,740663168 | chr16 | - |
| Chrd | 50,09 | -0,4252 | 0,368428658 | chr16 | + |
| Ciita | 16,47 | 0,9592 | 0,06491165 | chr16 | + |
| Clcn2 | 463,94 | -0,3010 | 0,290238703 | chr16 | - |
| Cldn14 | 104,97 | 0,5911 | 0,062190624 | chr16 | - |
| Cldnd1 | 389,45 | 0,1194 | 0,781562509 | chr16 | + |
| Cldn5 | 81,14 | -0,2246 | 0,713035456 | chr16 | + |
| Cldn8 | 6,21 | -0,1059 | 0,910916841 | chr16 | - |

| | | | | | |
|---|---|---|---|---|---|
| Clec16a | 262,44 | -0,4751 | 0,080075953 | chr16 | + |
| Clic6 | 5,37 | 0,2789 | 0,712967109 | chr16 | + |
| Cluap1 | 138,60 | -0,2255 | 0,634394209 | chr16 | + |
| Cmss1 | 100,51 | 0,4605 | 0,266466163 | chr16 | - |
| Col8a1 | 17,00 | 0,1827 | 0,81349332 | chr16 | - |
| Comt | 15703,19 | 0,0912 | 0,8763719 | chr16 | - |
| Coro7 | 129,73 | 0,2028 | 0,713614788 | chr16 | - |
| Cox17 | 860,48 | -0,0790 | 0,913936879 | chr16 | + |
| Cpn2 | 5722,98 | 0,1751 | 0,644714491 | chr16 | - |
| Cpox | 1465,76 | 0,3309 | 0,437233246 | chr16 | + |
| Cpped1 | 894,90 | 0,0706 | 0,876857496 | chr16 | - |
| Crebbp | 71,26 | -0,3075 | 0,515706574 | chr16 | - |
| Crkl | 524,94 | 0,0035 | 0,993487304 | chr16 | + |
| Cryzl1 | 248,27 | -0,2570 | 0,47939388 | chr16 | - |
| Cyyr1 | 11,38 | 0,6466 | 0,299245205 | chr16 | - |
| Dcbld2 | 46,30 | -0,1575 | 0,798276318 | chr16 | + |
| Dgcr14 | 56,16 | -0,1516 | 0,802338165 | chr16 | - |
| Dgcr2 | 868,58 | -0,0994 | 0,784087358 | chr16 | - |
| Dgcr6 | 452,41 | -0,2639 | 0,648735143 | chr16 | + |
| Dgcr8 | 170,00 | -0,2694 | 0,497745835 | chr16 | - |
| Dlg1 | 813,80 | 0,2299 | 0,414291323 | chr16 | + |
| Dnaja3 | 1723,25 | -0,0871 | 0,83176435 | chr16 | + |
| Dnajb11 | 1744,20 | -0,2615 | 0,564724076 | chr16 | + |
| Dnajc28 | 258,52 | 0,2062 | 0,683817601 | chr16 | - |
| Dnase1 | 3,53 | 0,5456 | 0,402107622 | chr16 | + |
| Dnm1l | 760,18 | 0,0675 | 0,869544763 | chr16 | - |
| Donson | 29,90 | 0,4438 | 0,413537476 | chr16 | - |
| Dopey2 | 251,97 | -0,0690 | 0,893432234 | chr16 | + |
| Dscr3 | 521,61 | 0,1783 | 0,614209897 | chr16 | - |
| Dtx3l | 767,40 | 0,1858 | 0,651407171 | chr16 | - |
| Dvl3 | 65,87 | -0,1167 | 0,855976995 | chr16 | + |
| Dyrk1a | 348,91 | -0,2311 | 0,435611308 | chr16 | + |
| Dzip3 | 47,19 | 0,3229 | 0,540877577 | chr16 | - |
| E330011O21Rik | 91,14 | 0,5689 | 0,323328805 | chr16 | - |
| Eaf2 | 90,23 | 0,5935 | 0,086898554 | chr16 | - |
| Ece2 | 186,69 | -0,1488 | 0,717728644 | chr16 | + |
| Ehhadh | 6169,40 | 0,4095 | 0,437265048 | chr16 | - |
| Eif2b5 | 607,50 | -0,3062 | 0,411295797 | chr16 | + |
| Eif4a2 | 6995,32 | -0,1020 | 0,840322097 | chr16 | + |
| Eif4g1 | 4409,24 | -0,3380 | 0,223917823 | chr16 | + |
| Epha3 | 9,85 | 0,9873 | 0,089263853 | chr16 | - |
| Ephb3 | 4,12 | 0,2434 | 0,761732176 | chr16 | + |
| Ercc4 | 79,06 | 0,1773 | 0,701666523 | chr16 | + |
| Erg | 71,84 | 0,4780 | 0,220777692 | chr16 | - |
| Ets2 | 434,98 | 0,1604 | 0,690639808 | chr16 | + |
| Etv5 | 67,15 | 0,0944 | 0,8763719 | chr16 | - |
| Fam131a | 6,50 | 0,1302 | 0,886483612 | chr16 | + |
| Fam162a | 914,56 | -0,5110 | 0,149221118 | chr16 | - |
| Fam43a | 61,62 | -0,0124 | 0,988512389 | chr16 | + |
| Eef2kmt | 136,24 | 0,0740 | 0,877949305 | chr16 | - |

| | | | | | |
|---|---|---|---|---|---|
| Fbxo45 | 212,35 | -0,0717 | 0,872053924 | chr16 | - |
| Fgd4 | 62,20 | 0,4259 | 0,309392522 | chr16 | - |
| Fgf12 | 3,65 | -0,2156 | 0,793608383 | chr16 | - |
| Filip1l | 109,68 | 0,5728 | 0,104658336 | chr16 | + |
| Fopnl | 475,61 | 0,1027 | 0,840953217 | chr16 | - |
| Fyttd1 | 705,77 | 0,0097 | 0,98507181 | chr16 | + |
| G730013B05Rik | 12,40 | 0,4807 | 0,479400504 | chr16 | + |
| Gabpa | 661,88 | 0,2284 | 0,606429785 | chr16 | + |
| Gart | 531,21 | -0,0787 | 0,860606321 | chr16 | - |
| Gbe1 | 1932,32 | 0,1671 | 0,701666523 | chr16 | + |
| Gcsam | 6,31 | -0,2210 | 0,794671748 | chr16 | + |
| Glis2 | 21,70 | 0,6153 | 0,260501558 | chr16 | + |
| Glyr1 | 1795,81 | 0,3222 | 0,307080143 | chr16 | - |
| Gm10785 | 4,80 | 0,0139 | 0,989908794 | chr16 | + |
| Gm10789 | 2,25 | 0,2546 | NA | chr16 | + |
| Gm10791 | 15,69 | 0,0841 | 0,920510254 | chr16 | + |
| Gm15760 | 2,75 | -0,3414 | 0,62964307 | chr16 | - |
| Gm16861 | 3,52 | 0,1134 | 0,899497201 | chr16 | - |
| Gm19522 | 63,79 | -0,2905 | 0,683522146 | chr16 | - |
| Gm20319 | 1038,04 | 0,1653 | 0,757395951 | chr16 | - |
| Gm4262 | 5,17 | 0,4795 | 0,495019454 | chr16 | + |
| Usf3 | 548,42 | -0,3581 | 0,23807108 | chr16 | + |
| Gnb1l | 17,90 | -0,4630 | 0,482787957 | chr16 | + |
| Golgb1 | 869,99 | 0,0283 | 0,95728901 | chr16 | + |
| Gramd1c | 595,35 | 0,0955 | 0,834675238 | chr16 | - |
| Gsk3b | 428,87 | -0,0705 | 0,847419446 | chr16 | + |
| Gspt1 | 2114,21 | -0,0797 | 0,850915997 | chr16 | - |
| Gtf2e1 | 293,41 | -0,3695 | 0,343440412 | chr16 | - |
| Gtpbp8 | 130,44 | 0,3348 | 0,486005862 | chr16 | - |
| Hcls1 | 124,92 | 0,3749 | 0,352372205 | chr16 | + |
| Heg1 | 46,78 | 0,5298 | 0,219049665 | chr16 | + |
| Hes1 | 72,62 | 0,3992 | 0,5252592 | chr16 | + |
| Hic2 | 45,87 | 0,1450 | 0,860107955 | chr16 | + |
| Hira | 156,17 | -0,2511 | 0,566893799 | chr16 | + |
| Hlcs | 176,48 | 0,5277 | 0,209412299 | chr16 | - |
| Hmgn1 | 943,19 | -0,2489 | 0,442165263 | chr16 | - |
| Hmox2 | 1520,12 | 0,0794 | 0,81323711 | chr16 | + |
| Hrg | 11300,64 | -0,3709 | 0,15139513 | chr16 | + |
| Hspa13 | 806,19 | 0,2301 | 0,639465027 | chr16 | - |
| Hspbap1 | 36,56 | 0,6017 | 0,199292942 | chr16 | + |
| Hunk | 48,89 | 0,6385 | 0,220418266 | chr16 | + |
| Ifnar1 | 520,93 | 0,0780 | 0,823949825 | chr16 | + |
| Ifngr2 | 333,32 | 0,3945 | 0,282189277 | chr16 | + |
| Ift57 | 14,59 | 0,5555 | 0,366318165 | chr16 | + |
| Igf2bp2 | 21,50 | 0,9049 | 0,082772365 | chr16 | - |
| Igsf11 | 252,02 | 0,0190 | 0,969512928 | chr16 | + |
| Igsf5 | 836,44 | -0,3927 | 0,231622612 | chr16 | + |
| Il10rb | 150,11 | 0,1795 | 0,665411671 | chr16 | + |
| Ildr1 | 3,16 | 0,4768 | 0,470905336 | chr16 | + |
| Impg2 | 566,41 | -0,3985 | 0,165530407 | chr16 | + |

| | | | | | |
|---|---|---|---|---|---|
| Iqcb1 | 95,45 | -0,5599 | 0,152078239 | chr16 | + |
| Itgb5 | 764,67 | 0,0391 | 0,923778914 | chr16 | + |
| Itsn1 | 231,06 | 0,1686 | 0,671813639 | chr16 | + |
| Jam2 | 272,36 | 0,2470 | 0,487880306 | chr16 | + |
| Kcnj15 | 10,61 | 0,0169 | 0,986889999 | chr16 | + |
| Klhl22 | 206,18 | 0,0659 | 0,895594598 | chr16 | + |
| Klhl24 | 1261,58 | 0,3297 | 0,433006548 | chr16 | + |
| Klhl6 | 73,81 | 0,4855 | 0,263245957 | chr16 | - |
| Kng1 | 50936,43 | -0,2654 | 0,386806294 | chr16 | + |
| Kng2 | 5805,22 | -0,2646 | 0,433006548 | chr16 | - |
| Kpna1 | 736,38 | -0,1190 | 0,793250646 | chr16 | + |
| P3h2 | 10,41 | -0,0244 | 0,981991128 | chr16 | - |
| Liph | 2,42 | -0,2567 | NA | chr16 | - |
| Litaf | 1096,30 | -0,3550 | 0,411460129 | chr16 | - |
| Lmln | 11,56 | 0,5903 | 0,370803111 | chr16 | + |
| Lpp | 798,97 | -0,1452 | 0,724652712 | chr16 | + |
| Lrch3 | 125,63 | 0,1022 | 0,854019408 | chr16 | + |
| Lrrc58 | 1239,65 | -0,5127 | 0,072865004 | chr16 | + |
| Lsg1 | 284,78 | -0,3479 | 0,24947031 | chr16 | - |
| Ltn1 | 568,87 | 0,0880 | 0,862616891 | chr16 | - |
| Lztr1 | 436,36 | -0,0518 | 0,901724181 | chr16 | + |
| Map3k13 | 199,92 | 0,6601 | 0,15392197 | chr16 | + |
| Mapk1 | 1625,40 | -0,0309 | 0,941757244 | chr16 | + |
| Marf1 | 1294,73 | -0,0723 | 0,838868977 | chr16 | - |
| Masp1 | 758,30 | 0,4639 | 0,264462538 | chr16 | - |
| Mb21d2 | 328,73 | -0,1288 | 0,792197952 | chr16 | - |
| Mcm4 | 70,20 | 0,2586 | 0,655452587 | chr16 | - |
| Med15 | 223,65 | 0,0035 | 0,994765486 | chr16 | - |
| Mefv | 8,66 | 0,2783 | 0,724955742 | chr16 | - |
| Mettl22 | 165,68 | -0,2951 | 0,463565059 | chr16 | + |
| Meltf | 2,24 | 0,8056 | NA | chr16 | + |
| Mgrn1 | 610,92 | -0,3272 | 0,329621696 | chr16 | + |
| Riox2 | 253,09 | 0,3816 | 0,3025413 | chr16 | + |
| Mir568 | 11,00 | 0,4035 | 0,576265028 | chr16 | + |
| Mis18a | 47,48 | 0,0023 | 0,997458998 | chr16 | - |
| Mkl2 | 5,18 | 0,0217 | 0,984995389 | chr16 | + |
| Morc3 | 364,61 | 0,3237 | 0,465196538 | chr16 | + |
| Mpv17l | 661,72 | -0,1237 | 0,842502441 | chr16 | + |
| Mrap | 753,53 | -0,1510 | 0,695978671 | chr16 | + |
| Mrpl39 | 905,29 | -0,1653 | 0,657237768 | chr16 | - |
| Mrpl40 | 496,25 | -0,1904 | 0,587007634 | chr16 | - |
| Mrps6 | 139,78 | -0,2123 | 0,66477796 | chr16 | + |
| Mx1 | 35,20 | 0,5067 | 0,394162656 | chr16 | - |
| Mx2 | 115,26 | 0,4544 | 0,38893116 | chr16 | + |
| Myh11 | 36,36 | -0,0154 | 0,985849808 | chr16 | - |
| Mylk | 625,79 | 0,0536 | 0,915621354 | chr16 | + |
| Mzt2 | 270,90 | 0,1203 | 0,767917363 | chr16 | - |
| N6amt1 | 141,63 | 0,0606 | 0,905147397 | chr16 | + |
| Naa50 | 1887,43 | -0,0307 | 0,952576171 | chr16 | + |
| Nagpa | 116,65 | -0,0517 | 0,93551479 | chr16 | - |

| | | | | | |
|---|---|---|---|---|---|
| Ncbp2 | 466,00 | -0,2663 | 0,47680357 | chr16 | + |
| Nde1 | 220,09 | -0,0610 | 0,909834629 | chr16 | + |
| Ndufb4 | 2356,68 | -0,4094 | 0,239392998 | chr16 | - |
| Nfkbiz | 135,02 | -0,0622 | 0,939351122 | chr16 | - |
| Nit2 | 5150,61 | 0,1438 | 0,736984547 | chr16 | - |
| Nr1i2 | 253,42 | -0,0784 | 0,871983898 | chr16 | - |
| Nrip1 | 35,77 | 0,0981 | 0,895594598 | chr16 | - |
| Nrros | 75,79 | -0,0727 | 0,893432234 | chr16 | - |
| Nsun3 | 68,65 | 0,4514 | 0,413323375 | chr16 | - |
| Ntan1 | 296,43 | -0,0667 | 0,89879469 | chr16 | + |
| Nudt16l1 | 244,95 | 0,2960 | 0,437710831 | chr16 | + |
| Nxpe3 | 18,09 | 0,1933 | 0,801547461 | chr16 | - |
| Olig1 | 22,34 | -0,7907 | 0,145466971 | chr16 | + |
| Opa1 | 1321,34 | -0,2666 | 0,268487435 | chr16 | + |
| Osbpl11 | 222,14 | -0,2059 | 0,570801749 | chr16 | + |
| Pak2 | 489,20 | 0,0471 | 0,913749296 | chr16 | - |
| Pam16 | 423,96 | -0,4321 | 0,421362686 | chr16 | - |
| Parl | 460,92 | -0,1753 | 0,672801255 | chr16 | - |
| Parn | 320,80 | 0,2212 | 0,616263238 | chr16 | - |
| Parp9 | 1443,28 | 0,2949 | 0,54877155 | chr16 | + |
| Paxbp1 | 278,79 | -0,1311 | 0,754889632 | chr16 | - |
| Pcnp | 1717,76 | 0,0183 | 0,969414042 | chr16 | - |
| Pcyt1a | 4483,61 | -0,6863 | 0,050148252 | chr16 | + |
| Pdia5 | 1365,56 | 0,1247 | 0,789064438 | chr16 | - |
| Pdxdc1 | 1652,78 | -0,3684 | 0,165182527 | chr16 | - |
| Phldb2 | 461,82 | 0,3180 | 0,52377705 | chr16 | - |
| Pi4ka | 411,08 | -0,4111 | 0,176249471 | chr16 | - |
| Pigp | 433,41 | 0,3268 | 0,468217847 | chr16 | - |
| Pigx | 259,65 | -0,4927 | 0,191276087 | chr16 | - |
| Pigz | 3,74 | -0,3579 | 0,633667772 | chr16 | + |
| Pkp2 | 463,82 | -0,0199 | 0,969618147 | chr16 | + |
| Pla1a | 1220,83 | -0,0835 | 0,802594059 | chr16 | - |
| Plcxd2 | 1941,50 | 0,1443 | 0,709237475 | chr16 | - |
| Pmm2 | 728,29 | -0,1147 | 0,809065986 | chr16 | + |
| Polq | 2,71 | -0,1321 | 0,879759297 | chr16 | + |
| Polr2h | 270,17 | -0,1309 | 0,764562081 | chr16 | + |
| Ppil2 | 704,54 | 0,0066 | 0,988512389 | chr16 | - |
| Ppm1f | 480,18 | 0,2593 | 0,413323375 | chr16 | + |
| Ppp1r2 | 623,61 | -0,4054 | 0,175240442 | chr16 | - |
| Prdm15 | 62,46 | -0,2046 | 0,6841486 | chr16 | - |
| Prkdc | 124,59 | 0,0935 | 0,888648505 | chr16 | + |
| Prm1 | 3,30 | 0,1197 | NA | chr16 | - |
| Prm2 | 3,77 | 0,1202 | NA | chr16 | - |
| Prodh | 2583,43 | 0,1409 | 0,792709808 | chr16 | - |
| Pros1 | 1399,60 | -0,2421 | 0,335264505 | chr16 | + |
| Psmd2 | 3269,50 | -0,2204 | 0,501951423 | chr16 | + |
| Psmg1 | 448,54 | -0,2044 | 0,581157554 | chr16 | - |
| Hacd2 | 1455,44 | 0,0450 | 0,932968636 | chr16 | + |
| Qtrt2 | 84,24 | -0,1337 | 0,815253842 | chr16 | - |
| Rabl3 | 213,78 | 0,1324 | 0,813323745 | chr16 | + |

| | | | | | |
|---|---|---|---|---|---|
| Ranbp1 | 376,54 | -0,1209 | 0,831759716 | chr16 | - |
| Rcan1 | 390,33 | 0,2648 | 0,574684302 | chr16 | - |
| Rfc4 | 96,60 | 0,1784 | 0,755670612 | chr16 | - |
| Ripk4 | 118,18 | -0,0116 | 0,985628536 | chr16 | - |
| Ripply3 | 44,27 | 0,0730 | 0,917042614 | chr16 | + |
| Rmi2 | 45,89 | 0,0378 | 0,95728901 | chr16 | + |
| Rnf168 | 280,29 | 0,2129 | 0,54795017 | chr16 | + |
| Robo2 | 29,94 | 0,4940 | 0,356860747 | chr16 | - |
| Rogdi | 216,71 | 0,3093 | 0,564849438 | chr16 | - |
| Rpl31-ps12 | 2,74 | -0,4266 | 0,516581635 | chr16 | - |
| Rpl35a | 7999,94 | -0,4919 | 0,10175355 | chr16 | + |
| Rrn3 | 282,78 | -0,2521 | 0,464991549 | chr16 | + |
| Rsl1d1 | 905,71 | -0,0773 | 0,869695212 | chr16 | - |
| Rtp4 | 334,46 | 0,5401 | 0,151610372 | chr16 | + |
| Runx1 | 8,24 | 0,7584 | 0,225575035 | chr16 | - |
| Rwdd2b | 183,46 | 0,0194 | 0,972909166 | chr16 | - |
| Samsn1 | 16,79 | 0,2693 | 0,697553093 | chr16 | - |
| Scaf4 | 134,49 | -0,3662 | 0,303938245 | chr16 | - |
| Scarf2 | 21,31 | -0,2091 | 0,793415854 | chr16 | + |
| Sdf2l1 | 418,06 | 0,0825 | 0,913452466 | chr16 | - |
| Sec22a | 410,13 | 0,1818 | 0,712581071 | chr16 | - |
| Sema5b | 14,64 | 0,8343 | 0,137618722 | chr16 | + |
| Senp2 | 397,41 | -0,4967 | 0,077430712 | chr16 | + |
| Senp5 | 493,07 | 0,0087 | 0,988512389 | chr16 | - |
| Serpind1 | 17281,94 | 0,1504 | 0,620918751 | chr16 | + |
| Sh3bgr | 2,49 | 0,3406 | NA | chr16 | + |
| Shisa9 | 3,78 | -0,2182 | 0,792536288 | chr16 | + |
| Slc12a8 | 8,99 | 0,2114 | 0,804380135 | chr16 | + |
| Slc15a2 | 90,17 | -0,7797 | 0,128835828 | chr16 | - |
| Slc25a1 | 1616,96 | -0,4828 | 0,130847434 | chr16 | - |
| Slc35a5 | 423,78 | 0,3383 | 0,36035337 | chr16 | - |
| Slc5a3 | 169,84 | -0,2250 | 0,724652712 | chr16 | + |
| Slx4 | 28,25 | -0,1758 | 0,783422709 | chr16 | - |
| Smco1 | 2,71 | 0,1938 | 0,792053919 | chr16 | + |
| Smim11 | 386,79 | -0,3454 | 0,317123683 | chr16 | + |
| Smim22 | 153,36 | 0,5833 | 0,190066764 | chr16 | + |
| Smpd4 | 144,54 | -0,1674 | 0,712581071 | chr16 | + |
| Snai2 | 59,01 | -0,1046 | 0,868115228 | chr16 | + |
| Snap29 | 851,49 | 0,0762 | 0,847964947 | chr16 | + |
| Snn | 26,30 | 0,1216 | 0,87206597 | chr16 | + |
| Snora81 | 5,40 | -0,2073 | 0,809065986 | chr16 | + |
| Snord2 | 4,35 | 0,3935 | 0,571043431 | chr16 | + |
| Snx29 | 59,90 | 0,2100 | 0,736631536 | chr16 | + |
| Snx4 | 614,26 | 0,1484 | 0,666980604 | chr16 | + |
| Sod1 | 12606,50 | -0,5514 | 0,157020992 | chr16 | + |
| Son | 1515,62 | -0,0503 | 0,899463828 | chr16 | + |
| Spice1 | 43,01 | 0,4520 | 0,416983017 | chr16 | + |
| Srl | 15,64 | -0,1698 | 0,827617374 | chr16 | - |
| St3gal6 | 60,13 | 0,6113 | 0,169293687 | chr16 | - |
| St6gal1 | 1604,81 | 0,1255 | 0,783418712 | chr16 | + |

| | | | | | |
|---|---|---|---|---|---|
| Stxbp5l | 3,98 | -0,7933 | 0,187706614 | chr16 | - |
| Synj1 | 147,89 | -0,0371 | 0,946138414 | chr16 | - |
| Tagln3 | 4,54 | -0,7451 | 0,220418266 | chr16 | - |
| Tango2 | 803,08 | 0,1274 | 0,753582185 | chr16 | - |
| Tbc1d23 | 195,66 | 0,0776 | 0,871700795 | chr16 | - |
| Tbccd1 | 364,00 | -0,1777 | 0,62632688 | chr16 | - |
| Tctex1d2 | 55,39 | 0,7328 | 0,086823025 | chr16 | + |
| Tfap4 | 40,70 | 0,3551 | 0,527820373 | chr16 | - |
| Tfg | 628,54 | -0,2303 | 0,425493131 | chr16 | - |
| Tfrc | 586,60 | 0,6570 | 0,179535584 | chr16 | + |
| Thap7 | 104,61 | -0,3781 | 0,581489212 | chr16 | - |
| Thpo | 163,82 | 0,1073 | 0,859981869 | chr16 | - |
| Tiam1 | 14,30 | -0,0899 | 0,920510254 | chr16 | - |
| Tigit | 8,61 | -0,0164 | 0,987867023 | chr16 | - |
| Timmdc1 | 593,80 | -0,3972 | 0,10162006 | chr16 | - |
| Tmem186 | 197,65 | 0,2201 | 0,539135765 | chr16 | - |
| Tmem191c | 5,02 | -0,1812 | 0,835640396 | chr16 | + |
| Tmem39a | 163,30 | 0,2282 | 0,610005889 | chr16 | + |
| Tmem44 | 19,20 | 0,0375 | 0,970125173 | chr16 | - |
| Tmem50b | 122,78 | 0,0150 | 0,981566938 | chr16 | - |
| Tnk2 | 143,50 | -0,4798 | 0,340674125 | chr16 | + |
| Tnp2 | 3,06 | 0,1194 | NA | chr16 | - |
| Tomm70a | 2573,38 | -0,3044 | 0,37347595 | chr16 | + |
| Top3b | 200,13 | 0,1652 | 0,680745326 | chr16 | + |
| Tra2b | 1025,03 | 0,4459 | 0,191230937 | chr16 | - |
| Trap1 | 2938,34 | -0,1389 | 0,630381755 | chr16 | - |
| Trmt10c | 408,61 | -0,0102 | 0,986642823 | chr16 | - |
| Trmt2a | 161,43 | -0,0678 | 0,884450766 | chr16 | + |
| Tssk1 | 2,46 | 0,1640 | NA | chr16 | + |
| Txndc11 | 252,72 | -0,4149 | 0,33288309 | chr16 | - |
| Txnrd2 | 505,06 | -0,0237 | 0,95463264 | chr16 | + |
| Ubald1 | 111,44 | -0,2145 | 0,751230284 | chr16 | - |
| Ube2l3 | 2072,69 | -0,0299 | 0,937853416 | chr16 | - |
| Ube2v2 | 929,19 | 0,3888 | 0,250303145 | chr16 | - |
| Ubn1 | 461,39 | -0,0707 | 0,869825447 | chr16 | + |
| Ubxn7 | 621,01 | 0,0405 | 0,926388088 | chr16 | + |
| Ufd1 | 714,34 | -0,2780 | 0,263205799 | chr16 | + |
| Umps | 619,99 | 0,5008 | 0,096308851 | chr16 | - |
| Upk1b | 8,37 | 0,2366 | 0,775458257 | chr16 | - |
| Urb1 | 46,38 | 0,0878 | 0,88569455 | chr16 | - |
| Usp16 | 666,24 | -0,0649 | 0,881809902 | chr16 | + |
| Usp25 | 399,50 | -0,2657 | 0,509550737 | chr16 | + |
| Usp7 | 850,00 | -0,3077 | 0,217231457 | chr16 | - |
| Vasn | 67,34 | 0,2230 | 0,65309424 | chr16 | + |
| Vgll3 | 2,54 | 0,3157 | NA | chr16 | + |
| Vps8 | 96,81 | 0,1735 | 0,716533507 | chr16 | + |
| Wdr53 | 230,68 | -0,0686 | 0,88286244 | chr16 | + |
| Wdr5b | 33,28 | 0,5394 | 0,3696571 | chr16 | + |
| Wrb | 60,86 | 0,2707 | 0,562477407 | chr16 | + |
| Xxylt1 | 23,85 | 0,4773 | 0,426922409 | chr16 | - |

| | | | | | |
|---|---|---|---|---|---|
| Yars2 | 285,78 | -0,3433 | 0,266466163 | chr16 | + |
| Yeats2 | 106,33 | -0,0214 | 0,972672419 | chr16 | + |
| Ypel1 | 2,19 | 0,5213 | NA | chr16 | + |
| Zbtb11 | 223,48 | -0,0782 | 0,882756477 | chr16 | + |
| Zbtb20 | 54,53 | -0,8187 | 0,077385039 | chr16 | + |
| Zbtb21 | 76,77 | 0,3601 | 0,412935998 | chr16 | - |
| Zc3h7a | 501,85 | -0,2257 | 0,517321822 | chr16 | - |
| Zdhhc8 | 53,90 | 0,0230 | 0,979943138 | chr16 | - |
| Zfp148 | 684,54 | -0,0210 | 0,965770119 | chr16 | + |
| Zfp174 | 33,99 | -0,2791 | 0,684297941 | chr16 | + |
| Zfp263 | 192,86 | 0,0166 | 0,974449819 | chr16 | + |
| Zfp654 | 261,24 | -0,3396 | 0,289820918 | chr16 | - |
| Mettl26 | 1597,19 | -0,6109 | 0,173210622 | chr17 | + |
| 1110020A21Rik | 43,18 | -0,7004 | 0,096695555 | chr17 | - |
| 1110038B12Rik | 208,82 | -0,7834 | 0,078316133 | chr17 | - |
| 1600012H06Rik | 602,18 | 0,0708 | 0,88286244 | chr17 | + |
| 1700001C19Rik | 135,04 | -0,1222 | 0,821346657 | chr17 | - |
| 1700010I14Rik | 7,08 | 0,8617 | 0,155946567 | chr17 | + |
| 1700102H20Rik | 2,63 | -0,3976 | 0,564849438 | chr17 | + |
| 2310039H08Rik | 435,52 | -0,1365 | 0,881097925 | chr17 | + |
| 2410015M20Rik | 2781,95 | -0,2915 | 0,308448866 | chr17 | - |
| 2700054A10Rik | 9,99 | 0,2268 | 0,788000186 | chr17 | - |
| 2700099C18Rik | 6,74 | -0,0341 | 0,975445719 | chr17 | + |
| Zfp983 | 127,86 | 0,4962 | 0,148243392 | chr17 | + |
| 4732491K20Rik | 24,56 | 0,2003 | 0,762940356 | chr17 | + |
| 4833418N02Rik | 21,25 | -0,1266 | 0,876803328 | chr17 | - |
| 4930470H14Rik | 86,50 | -0,3389 | 0,384986291 | chr17 | - |
| 4930506C21Rik | 8,30 | 0,0740 | 0,941250497 | chr17 | - |
| 4930539E08Rik | 16,23 | 0,7220 | 0,202636202 | chr17 | - |
| 4933401D09Rik | 3,35 | 0,9452 | 0,072134655 | chr17 | + |
| 9030025P20Rik | 21,79 | -0,2736 | 0,670191619 | chr17 | + |
| 9130008F23Rik | 4,37 | 0,6177 | 0,332014851 | chr17 | - |
| 9530082P21Rik | 39,05 | 0,4846 | 0,324580766 | chr17 | + |
| A530064D06Rik | 6,08 | 0,2094 | 0,806012703 | chr17 | - |
| A930015D03Rik | 2,66 | 0,3110 | 0,658008155 | chr17 | + |
| AA388235 | 14,25 | 0,4509 | 0,493980363 | chr17 | + |
| Aars2 | 179,55 | -0,1445 | 0,813323745 | chr17 | + |
| Abca3 | 874,59 | 0,1967 | 0,583518171 | chr17 | + |
| Abcc10 | 69,31 | -0,6786 | 0,171869196 | chr17 | - |
| Abcf1 | 789,50 | -0,4750 | 0,117246861 | chr17 | - |
| Abcg1 | 106,60 | 0,0138 | 0,98429278 | chr17 | + |
| Abcg5 | 1497,47 | -0,6581 | 0,052711367 | chr17 | - |
| Abhd16a | 416,09 | -0,0785 | 0,89879469 | chr17 | + |
| Acat2 | 887,90 | -0,0171 | 0,972909166 | chr17 | - |
| Acat3 | 2276,14 | 0,1134 | 0,767131243 | chr17 | - |
| Adamts10 | 58,79 | -0,1824 | 0,795812563 | chr17 | + |
| Ager | 4,71 | -0,0178 | 0,986691613 | chr17 | + |
| Agpat1 | 94,56 | -0,4234 | 0,295666715 | chr17 | + |
| Agpat4 | 18,12 | 0,0913 | 0,914198498 | chr17 | + |
| AI413582 | 34,93 | 0,3497 | 0,537660048 | chr17 | - |

| | | | | | |
|---|---|---|---|---|---|
| AI661453 | 264,85 | 0,0236 | 0,972909166 | chr17 | + |
| Aif1 | 227,34 | 0,1268 | 0,807278226 | chr17 | - |
| Airn | 6,35 | -0,3566 | 0,637152434 | chr17 | + |
| Akap8 | 652,47 | -0,1791 | 0,546240378 | chr17 | - |
| Akap8l | 310,52 | -0,6288 | 0,110128525 | chr17 | - |
| Alkbh7 | 94,32 | -0,2915 | 0,637139802 | chr17 | + |
| Amdhd2 | 194,59 | -0,2441 | 0,669280735 | chr17 | - |
| Angptl4 | 3686,41 | -0,4404 | 0,310756756 | chr17 | - |
| Ankrd12 | 127,77 | 0,4463 | 0,246159206 | chr17 | - |
| Anks1 | 194,74 | 0,3703 | 0,419539452 | chr17 | + |
| Apom | 7728,07 | -0,0870 | 0,814733578 | chr17 | - |
| Arhgap28 | 7,84 | 0,1070 | 0,909834629 | chr17 | - |
| Arid1b | 256,30 | -0,0837 | 0,838868977 | chr17 | + |
| Atf6b | 194,28 | 0,0752 | 0,895594598 | chr17 | + |
| Atl2 | 2515,36 | -0,0709 | 0,869825447 | chr17 | - |
| Atp6v0e | 961,15 | -0,3216 | 0,336671811 | chr17 | + |
| Atp6v1g2 | 13,63 | -0,5766 | 0,33244762 | chr17 | + |
| Axin1 | 262,57 | -0,3340 | 0,406993487 | chr17 | + |
| B3galt4 | 4,95 | -0,0931 | 0,922935681 | chr17 | - |
| B430306N03Rik | 39,78 | 0,4078 | 0,430392697 | chr17 | + |
| Cahm | 3,11 | 0,1224 | 0,890887823 | chr17 | + |
| Bag6 | 572,38 | -0,2689 | 0,535385445 | chr17 | + |
| Bak1 | 163,90 | 0,4905 | 0,321599847 | chr17 | - |
| BC003965 | 382,82 | -0,3055 | 0,394187455 | chr17 | + |
| BC004004 | 1762,50 | -0,2442 | 0,349661709 | chr17 | + |
| Platr17 | 6,56 | -0,0226 | 0,98429278 | chr17 | + |
| BC051226 | 32,31 | -0,4252 | 0,405233176 | chr17 | - |
| Birc6 | 919,23 | 0,1217 | 0,671975571 | chr17 | + |
| Bnip1 | 207,37 | 0,0675 | 0,885244858 | chr17 | + |
| Brd2 | 1146,99 | -0,2521 | 0,395040761 | chr17 | - |
| Brd4 | 293,30 | -0,3208 | 0,365662949 | chr17 | - |
| Brpf3 | 263,05 | -0,5077 | 0,231675652 | chr17 | + |
| Btbd9 | 226,37 | 0,0527 | 0,913017292 | chr17 | - |
| Bysl | 117,71 | -0,5288 | 0,212297691 | chr17 | - |
| C030034I22Rik | 15,36 | 0,1470 | 0,859139694 | chr17 | + |
| C2 | 3436,43 | 0,2083 | 0,522469399 | chr17 | - |
| C3 | 83400,39 | -0,1969 | 0,583782269 | chr17 | - |
| C4b | 11012,72 | -0,3318 | 0,289430713 | chr17 | - |
| Calm2 | 3894,04 | 0,2935 | 0,327028954 | chr17 | - |
| Camkmt | 10,68 | 0,3053 | 0,693158302 | chr17 | + |
| Capn11 | 2,13 | 0,1741 | NA | chr17 | - |
| Capn15 | 53,48 | -0,2513 | 0,734655193 | chr17 | - |
| Catsperd | 29,20 | 0,8544 | 0,062031449 | chr17 | + |
| Cbs | 4829,42 | -0,1157 | 0,787418695 | chr17 | - |
| Ccdc167 | 253,08 | -0,2358 | 0,559977981 | chr17 | - |
| Ccdc78 | 2,56 | -0,4793 | NA | chr17 | + |
| Ccdc94 | 50,49 | -0,1391 | 0,804754692 | chr17 | + |
| Cchcr1 | 25,74 | 0,3889 | 0,533686433 | chr17 | + |
| Ccnd3 | 343,47 | -0,1540 | 0,75436056 | chr17 | + |
| Ccnf | 305,44 | -0,0149 | 0,984995389 | chr17 | - |

| | | | | | |
|---|---|---|---|---|---|
| Cd2ap | 656,05 | 0,0584 | 0,905147397 | chr17 | - |
| Cd320 | 104,60 | -0,2430 | 0,612061669 | chr17 | + |
| Cdc42ep3 | 16,60 | -0,1382 | 0,862867001 | chr17 | - |
| Cdc5l | 436,86 | -0,0398 | 0,92446794 | chr17 | - |
| Cdkl4 | 5,84 | -0,2613 | 0,749992169 | chr17 | - |
| Cebpz | 789,62 | -0,1365 | 0,740352128 | chr17 | - |
| Cebpzos | 372,53 | -0,0436 | 0,925898178 | chr17 | + |
| Cenpq | 70,39 | 0,3310 | 0,458926944 | chr17 | - |
| Cfb | 28125,38 | 0,0455 | 0,912811384 | chr17 | - |
| Chaf1a | 22,58 | 0,0526 | 0,952989802 | chr17 | + |
| Chd1 | 465,14 | -0,0510 | 0,923778914 | chr17 | + |
| Clcn7 | 548,28 | 0,0732 | 0,847964947 | chr17 | + |
| Clic1 | 435,14 | 0,2789 | 0,356901218 | chr17 | + |
| Clpp | 570,62 | -0,4265 | 0,319311067 | chr17 | + |
| Cmtr1 | 666,36 | 0,1358 | 0,74829088 | chr17 | + |
| Cnpy3 | 370,81 | -0,2500 | 0,576267127 | chr17 | - |
| Col11a2 | 5,33 | -0,4672 | 0,513640882 | chr17 | + |
| Cox7a2l | 2802,94 | -0,4011 | 0,120261039 | chr17 | - |
| Cramp1l | 127,46 | 0,0035 | 0,995735297 | chr17 | - |
| Crb3 | 121,08 | -0,0911 | 0,906998011 | chr17 | + |
| Crebrf | 792,69 | -0,0181 | 0,977170685 | chr17 | + |
| Crim1 | 210,80 | -0,1237 | 0,79002616 | chr17 | + |
| Crip3 | 7,31 | -0,7932 | 0,198587755 | chr17 | + |
| Cript | 1016,65 | -0,2724 | 0,409970248 | chr17 | + |
| Csnk2b | 1082,99 | -0,5058 | 0,075466772 | chr17 | - |
| Cul7 | 87,10 | -0,2528 | 0,682099278 | chr17 | + |
| Cul9 | 52,78 | 0,7071 | 0,125541699 | chr17 | - |
| Cuta | 605,56 | -0,2957 | 0,524808191 | chr17 | - |
| Cyp1b1 | 16,49 | 0,7138 | 0,228941765 | chr17 | - |
| Cyp21a1 | 47,94 | -0,1568 | 0,857253251 | chr17 | - |
| Cyp4f14 | 3930,56 | 0,1198 | 0,697643792 | chr17 | - |
| Cyp4f15 | 1866,25 | 0,0833 | 0,805820649 | chr17 | + |
| Cyp4f17 | 67,53 | -0,1799 | 0,784877345 | chr17 | + |
| Cyp4f39 | 2,35 | -0,0044 | NA | chr17 | + |
| Cyp4f41-ps | 2,62 | 0,2829 | 0,682099278 | chr17 | - |
| D17Ertd648e | 3,21 | -0,1859 | 0,814990243 | chr17 | - |
| D17H6S53E | 195,35 | -0,0236 | 0,966776406 | chr17 | + |
| Mydgf | 1125,88 | -0,2722 | 0,64665416 | chr17 | - |
| D17Wsu92e | 3135,51 | -0,2948 | 0,309412451 | chr17 | - |
| D330041H03Rik | 4,62 | -0,4266 | 0,561773364 | chr17 | - |
| Daam2 | 28,56 | 0,7325 | 0,116833055 | chr17 | - |
| Dact2 | 98,75 | -0,1660 | 0,82942592 | chr17 | - |
| Daxx | 148,63 | 0,1622 | 0,695978671 | chr17 | + |
| Ddah2 | 40,99 | -0,1602 | 0,821346657 | chr17 | + |
| Ddr1 | 27,59 | 0,6727 | 0,191230937 | chr17 | - |
| Ddx11 | 58,22 | 0,0278 | 0,969512928 | chr17 | + |
| Ddx39b | 517,78 | -0,1162 | 0,807278226 | chr17 | + |
| Decr2 | 3429,05 | -0,0726 | 0,88569455 | chr17 | - |
| Def6 | 3,46 | 0,4270 | 0,539569536 | chr17 | + |
| Dennd1c | 11,09 | 0,1664 | 0,842940743 | chr17 | - |

| | | | | | |
|---|---|---|---|---|---|
| Dhx16 | 364,09 | -0,4966 | 0,104101739 | chr17 | + |
| Dhx57 | 140,83 | 0,3970 | 0,242830621 | chr17 | - |
| Dll1 | 41,87 | 0,5038 | 0,312022245 | chr17 | - |
| Dnah8 | 5,27 | -0,0697 | 0,943108135 | chr17 | + |
| Dnase1l2 | 13,91 | -0,1540 | 0,859745168 | chr17 | - |
| Dpp9 | 319,55 | -0,3301 | 0,4132425 | chr17 | - |
| Dpy30 | 526,98 | -0,0799 | 0,846704266 | chr17 | - |
| Dreh | 71,20 | -0,0571 | 0,945521759 | chr17 | + |
| Dus3l | 97,82 | -0,5055 | 0,301130682 | chr17 | + |
| Dusp1 | 343,16 | -0,4933 | 0,294750184 | chr17 | - |
| Dxo | 184,54 | -0,0632 | 0,920157635 | chr17 | + |
| Dync2li1 | 25,28 | 0,8831 | 0,066283273 | chr17 | + |
| Dynlt1a | 37,40 | 0,1562 | 0,825918365 | chr17 | - |
| E130008D07Rik | 16,49 | -0,4434 | 0,486009467 | chr17 | - |
| E4f1 | 77,98 | -0,5903 | 0,217835326 | chr17 | - |
| Ebi3 | 9,45 | 0,4055 | 0,577257466 | chr17 | + |
| Eci1 | 1793,36 | -0,2656 | 0,592333512 | chr17 | + |
| Efna5 | 94,44 | 0,6892 | 0,103715866 | chr17 | - |
| Egfl8 | 7,84 | -0,2099 | 0,801768605 | chr17 | - |
| Ehd3 | 933,81 | 0,2282 | 0,451182001 | chr17 | + |
| Ehmt2 | 505,45 | -0,3482 | 0,394162656 | chr17 | + |
| Eif2ak2 | 458,14 | 0,2775 | 0,588219472 | chr17 | - |
| Eme2 | 89,57 | 0,0967 | 0,869515614 | chr17 | - |
| Emilin2 | 27,94 | -0,2617 | 0,693242675 | chr17 | - |
| Eml4 | 225,65 | -0,1850 | 0,649565939 | chr17 | + |
| Adgre1 | 492,91 | 0,4509 | 0,151155593 | chr17 | + |
| Adgre4 | 183,62 | 0,5280 | 0,095599478 | chr17 | + |
| Enpp4 | 346,15 | 0,5475 | 0,083510154 | chr17 | - |
| Enpp5 | 761,61 | 0,4047 | 0,29233966 | chr17 | + |
| Epb41l3 | 42,20 | 0,5379 | 0,241800076 | chr17 | + |
| Epcam | 33,48 | 0,3868 | 0,500489603 | chr17 | + |
| Ergic1 | 881,10 | 0,0174 | 0,977141653 | chr17 | + |
| Ermard | 110,50 | 0,0032 | 0,996644671 | chr17 | + |
| Ezr | 74,89 | 0,1111 | 0,849477252 | chr17 | - |
| Fahd1 | 943,93 | -0,0220 | 0,970227029 | chr17 | - |
| Fam120b | 357,79 | 0,0012 | 0,997464504 | chr17 | + |
| Fam173a | 614,30 | -0,1639 | 0,793608383 | chr17 | - |
| Mcrip2 | 343,06 | -0,7146 | 0,101647827 | chr17 | - |
| Fam98a | 746,99 | -0,2056 | 0,607310966 | chr17 | - |
| Fance | 74,94 | -0,3791 | 0,4207546 | chr17 | + |
| Fbxl16 | 2,41 | -0,2164 | NA | chr17 | + |
| Fbxl17 | 413,95 | -0,0922 | 0,822067344 | chr17 | - |
| Fbxo11 | 419,94 | -0,0110 | 0,981991128 | chr17 | - |
| Fem1a | 271,96 | 0,1253 | 0,767446843 | chr17 | + |
| Fer | 126,24 | 0,1744 | 0,751956685 | chr17 | + |
| Fez2 | 465,92 | -0,1561 | 0,721834843 | chr17 | - |
| Fgd2 | 29,90 | -0,2140 | 0,747522909 | chr17 | + |
| Fgfr1op | 320,54 | -0,3662 | 0,211137484 | chr17 | + |
| Fkbp5 | 1356,87 | 0,3617 | 0,431444548 | chr17 | - |
| Fkbpl | 67,06 | -0,2606 | 0,590228824 | chr17 | + |

| | | | | | |
|---|---|---|---|---|---|
| Flot1 | 217,39 | -0,1588 | 0,769963557 | chr17 | + |
| Flywch1 | 64,78 | -0,0334 | 0,969159464 | chr17 | - |
| Flywch2 | 16,42 | -0,1281 | 0,886446111 | chr17 | - |
| Fndc1 | 3,28 | 0,1872 | 0,809582276 | chr17 | - |
| Foxn2 | 242,40 | -0,0740 | 0,875717638 | chr17 | + |
| Foxp4 | 117,43 | -0,2756 | 0,626400494 | chr17 | - |
| Fpr1 | 121,15 | 0,4405 | 0,29645146 | chr17 | - |
| Fpr2 | 50,77 | 0,0812 | 0,899463828 | chr17 | + |
| Frs3 | 12,77 | -0,2172 | 0,790595509 | chr17 | + |
| Gabbr1 | 25,48 | -0,0707 | 0,932012942 | chr17 | + |
| Galm | 3362,42 | 0,1915 | 0,549838805 | chr17 | + |
| Gemin6 | 122,95 | -0,0723 | 0,904179121 | chr17 | + |
| Gfer | 332,15 | -0,5114 | 0,069085297 | chr17 | - |
| Ggnbp1 | 85,52 | -0,6413 | 0,126720006 | chr17 | + |
| Bicral | 109,09 | 0,0189 | 0,974449819 | chr17 | - |
| Gm10509 | 32,96 | 0,3216 | 0,572269858 | chr17 | + |
| Gm11127 | 26,07 | -0,7100 | 0,252815255 | chr17 | - |
| Gm16386 | 10,81 | 0,8061 | 0,176249471 | chr17 | - |
| Gm16702 | 18,20 | -0,2471 | 0,722152554 | chr17 | - |
| Gm16712 | 2,53 | 0,5451 | NA | chr17 | - |
| Gm3435 | 6,57 | -0,6442 | 0,322327153 | chr17 | + |
| Gm6034 | 4,35 | 0,4619 | 0,500489603 | chr17 | + |
| Gm6402 | 78,16 | -0,5590 | 0,271262155 | chr17 | - |
| Gm6548 | 655,71 | 0,3260 | 0,568858283 | chr17 | - |
| Gm8909 | 3,95 | 0,3178 | 0,673822592 | chr17 | - |
| Gnl1 | 230,85 | -0,4407 | 0,213281186 | chr17 | + |
| Gnmt | 19700,26 | -0,2560 | 0,653708291 | chr17 | - |
| Gnptg | 386,52 | -0,0718 | 0,893432234 | chr17 | - |
| Gpank1 | 107,65 | -0,4632 | 0,191230937 | chr17 | + |
| Gpatch11 | 208,09 | -0,1348 | 0,782304667 | chr17 | + |
| Gpr108 | 450,71 | -0,4289 | 0,183821359 | chr17 | - |
| Adgrf1 | 253,56 | -0,2147 | 0,717728644 | chr17 | + |
| Adgrf5 | 549,99 | 0,0881 | 0,847964947 | chr17 | + |
| Gpsm3 | 34,79 | -0,0049 | 0,995142284 | chr17 | + |
| Gtf2f1 | 322,67 | -0,3687 | 0,237903839 | chr17 | - |
| Gtf2h4 | 206,86 | -0,2279 | 0,583782269 | chr17 | - |
| Gtf2h5 | 1015,84 | -0,1769 | 0,647184856 | chr17 | + |
| Gtpbp2 | 336,64 | -0,2538 | 0,60828654 | chr17 | + |
| Guca1a | 50,81 | -0,5639 | 0,161576375 | chr17 | - |
| Guca1b | 3,27 | 0,1684 | 0,842668735 | chr17 | + |
| H2-Aa | 1520,75 | 0,4557 | 0,357345082 | chr17 | - |
| H2-Ab1 | 526,29 | 0,4744 | 0,217175999 | chr17 | + |
| H2-D1 | 4009,21 | 0,3027 | 0,425066328 | chr17 | + |
| H2-DMa | 67,90 | 0,0953 | 0,881086817 | chr17 | + |
| H2-DMb1 | 63,88 | 0,5986 | 0,177510828 | chr17 | + |
| H2-DMb2 | 3,11 | 0,0917 | 0,92066291 | chr17 | + |
| H2-Eb1 | 443,89 | 0,4647 | 0,232472267 | chr17 | + |
| H2-Eb2 | 4,19 | 0,1960 | 0,81909949 | chr17 | + |
| H2-K1 | 20583,47 | -0,0415 | 0,943108135 | chr17 | - |
| H2-K2 | 115,90 | 0,1902 | 0,826107735 | chr17 | - |

| | | | | | |
|---|---|---|---|---|---|
| Pfdn6 | 629,48 | -0,4527 | 0,145765078 | chr17 | - |
| H2-M2 | 8,82 | 0,1536 | 0,864175561 | chr17 | - |
| H2-M3 | 67,30 | 0,6127 | 0,125513994 | chr17 | + |
| H2-Oa | 7,43 | 0,1547 | 0,862421004 | chr17 | + |
| H2-Ob | 18,08 | -0,1662 | 0,834675238 | chr17 | + |
| H2-Q5 | 133,34 | -0,2555 | 0,656102596 | chr17 | + |
| H2-T22 | 40,96 | 0,8725 | 0,060467148 | chr17 | - |
| H2-T23 | 2992,98 | 0,3950 | 0,357345082 | chr17 | - |
| H2-T24 | 84,95 | 0,6515 | 0,092096287 | chr17 | - |
| Haao | 10164,41 | -0,0916 | 0,821346657 | chr17 | - |
| Hagh | 5035,45 | -0,5263 | 0,050835053 | chr17 | + |
| Haghl | 71,27 | 0,1466 | 0,802632263 | chr17 | - |
| Hcfc1r1 | 495,67 | -0,4011 | 0,214347061 | chr17 | + |
| Hdgfl2 | 231,47 | -0,2670 | 0,569876243 | chr17 | + |
| Heatr5b | 180,28 | 0,0935 | 0,849477252 | chr17 | - |
| Hmga1 | 3,05 | 0,1343 | 0,86112413 | chr17 | + |
| Hnrnpll | 419,38 | -0,2456 | 0,415700483 | chr17 | - |
| Hnrnpm | 922,45 | -0,1867 | 0,614153379 | chr17 | - |
| Hs3st6 | 10,63 | -0,8776 | 0,147367695 | chr17 | + |
| Hsp90ab1 | 10524,33 | -0,1446 | 0,740352128 | chr17 | - |
| Hspa1a | 10,11 | -1,0586 | 0,063376737 | chr17 | - |
| Ier3 | 13,95 | 0,0956 | 0,916757648 | chr17 | + |
| Ift140 | 93,94 | 0,0314 | 0,966776406 | chr17 | + |
| Igf2r | 577,18 | -0,1524 | 0,619643473 | chr17 | - |
| Igfals | 1202,73 | -0,5028 | 0,110056031 | chr17 | + |
| Fam234a | 539,93 | -0,4932 | 0,119325906 | chr17 | - |
| Itpr3 | 19,93 | 0,4354 | 0,491188481 | chr17 | + |
| Jmjd8 | 733,77 | 0,0004 | 0,998994678 | chr17 | + |
| Kank3 | 77,20 | -0,2492 | 0,611365326 | chr17 | + |
| Kat2b | 1473,64 | -0,1140 | 0,795812563 | chr17 | + |
| Kctd20 | 301,27 | 0,0897 | 0,869687361 | chr17 | + |
| Kctd5 | 67,60 | -0,3289 | 0,491188481 | chr17 | - |
| Kdm4b | 126,04 | -0,6191 | 0,140539066 | chr17 | + |
| Khsrp | 270,09 | -0,2288 | 0,682348071 | chr17 | - |
| Kifc5b | 3,56 | -0,1805 | 0,83293655 | chr17 | + |
| Klc4 | 780,33 | -0,3030 | 0,409839891 | chr17 | - |
| Klhdc3 | 587,09 | -0,5161 | 0,230636762 | chr17 | - |
| Kremen2 | 2,68 | -0,0234 | 0,98142867 | chr17 | - |
| Lama1 | 57,63 | 0,3416 | 0,461603281 | chr17 | + |
| Lclat1 | 653,68 | 0,0602 | 0,875636241 | chr17 | + |
| Lemd2 | 266,52 | -0,2498 | 0,564849438 | chr17 | - |
| Lix1 | 8,79 | 0,1875 | 0,822888019 | chr17 | + |
| Lmf1 | 540,85 | -0,1697 | 0,647184856 | chr17 | + |
| Lonp1 | 1576,81 | -0,2781 | 0,390108497 | chr17 | - |
| Lrpprc | 2322,83 | -0,2310 | 0,462174993 | chr17 | - |
| Lsm2 | 119,23 | -0,1291 | 0,764664236 | chr17 | + |
| Lst1 | 91,99 | 0,5319 | 0,239641183 | chr17 | - |
| Ltb | 15,33 | 0,9199 | 0,088679016 | chr17 | + |
| Ltbp1 | 41,66 | 0,4419 | 0,379511614 | chr17 | + |
| Luc7l | 460,85 | -0,2104 | 0,621011348 | chr17 | + |

| | | | | | |
|---|---|---|---|---|---|
| Ly6g5b | 4,11 | -0,4033 | 0,572115465 | chr17 | - |
| Mad2l1bp | 78,86 | 0,0573 | 0,932528423 | chr17 | - |
| Man2a1 | 4513,35 | 0,0015 | 0,997302965 | chr17 | + |
| Map3k4 | 185,05 | -0,0446 | 0,941754349 | chr17 | - |
| Map4k3 | 439,99 | 0,0359 | 0,943305207 | chr17 | - |
| Mapk13 | 3,26 | 0,0596 | 0,950819236 | chr17 | + |
| Mapk14 | 1194,22 | -0,1853 | 0,615372313 | chr17 | + |
| Mapk8ip3 | 163,56 | -0,0847 | 0,882318627 | chr17 | - |
| Mcfd2 | 3913,18 | -0,0445 | 0,92369436 | chr17 | - |
| Mdc1 | 126,39 | 0,4323 | 0,19521384 | chr17 | + |
| Mdfi | 3,67 | 0,2822 | 0,717576816 | chr17 | - |
| Mea1 | 1027,73 | -0,2854 | 0,264965714 | chr17 | + |
| Med20 | 248,21 | -0,3046 | 0,279939661 | chr17 | + |
| Memo1 | 646,57 | -0,2711 | 0,390708488 | chr17 | - |
| Metrn | 67,59 | -0,0491 | 0,95728901 | chr17 | - |
| Mllt1 | 140,20 | -0,3781 | 0,421823262 | chr17 | - |
| Afdn | 590,17 | 0,1819 | 0,610359218 | chr17 | + |
| Mlst8 | 119,44 | 0,1068 | 0,834507118 | chr17 | - |
| Morn2 | 119,72 | 0,2254 | 0,594140852 | chr17 | + |
| Mpc1 | 4744,36 | 0,0581 | 0,930684299 | chr17 | + |
| Mpnd | 684,57 | -0,5460 | 0,068521927 | chr17 | + |
| Mrpl14 | 804,91 | -0,2902 | 0,561310046 | chr17 | + |
| Mrpl2 | 227,00 | 0,0186 | 0,969892522 | chr17 | + |
| Mrpl18 | 514,23 | -0,5187 | 0,274683426 | chr17 | - |
| Mrpl28 | 469,30 | -0,4170 | 0,301759036 | chr17 | + |
| Mrps10 | 561,75 | -0,2958 | 0,445734083 | chr17 | + |
| Mrps18a | 452,15 | -0,4229 | 0,194733334 | chr17 | + |
| Mrps18b | 716,22 | -0,3486 | 0,342638186 | chr17 | - |
| Mrps34 | 319,21 | -0,5370 | 0,127550707 | chr17 | + |
| Msh2 | 187,71 | 0,1198 | 0,784505535 | chr17 | + |
| Msh5 | 11,49 | 0,0703 | 0,94366064 | chr17 | - |
| Msh6 | 238,62 | -0,0815 | 0,864697132 | chr17 | + |
| Msln | 20,32 | 0,4344 | 0,538042836 | chr17 | - |
| Msrb1 | 7285,41 | -0,5277 | 0,106587153 | chr17 | + |
| Mta3 | 120,06 | -0,0349 | 0,953900048 | chr17 | + |
| Mtch1 | 734,30 | -0,3179 | 0,222351922 | chr17 | - |
| Mtcl1 | 5,35 | 0,7071 | 0,248157277 | chr17 | - |
| Mut | 3737,35 | 0,1961 | 0,622736166 | chr17 | + |
| Myl12a | 897,88 | 0,2104 | 0,540908104 | chr17 | - |
| Myl12b | 843,57 | 0,3093 | 0,394187455 | chr17 | - |
| Myo1f | 33,21 | 0,2163 | 0,725790752 | chr17 | + |
| Myom1 | 28,29 | -0,3168 | 0,630381755 | chr17 | + |
| Narfl | 325,66 | -0,0523 | 0,910351158 | chr17 | + |
| Ndc80 | 8,25 | 0,2816 | 0,726111471 | chr17 | - |
| Ndufa7 | 2162,62 | -0,2153 | 0,688902549 | chr17 | + |
| Ndufaf7 | 300,98 | 0,1135 | 0,812490734 | chr17 | + |
| Ndufb10 | 1822,23 | -0,4858 | 0,229100935 | chr17 | - |
| Ndufv2 | 2998,59 | -0,3252 | 0,377249969 | chr17 | - |
| Nelfe | 224,94 | -0,0813 | 0,879335212 | chr17 | + |
| Neu1 | 491,31 | -0,5095 | 0,12254751 | chr17 | + |

| | | | | | |
|---|---|---|---|---|---|
| Neurl1b | 12,38 | 0,4154 | 0,564702983 | chr17 | + |
| Nfkbie | 33,36 | 0,6312 | 0,207419757 | chr17 | + |
| Nfkbil1 | 53,90 | -0,3456 | 0,612061669 | chr17 | - |
| Nfya | 119,00 | 0,5788 | 0,078539732 | chr17 | - |
| Nlrc4 | 7,10 | 0,8472 | 0,165611157 | chr17 | - |
| Nme3 | 242,30 | -0,2995 | 0,632226809 | chr17 | + |
| Nme4 | 19,29 | -0,3804 | 0,557219065 | chr17 | - |
| Notch3 | 13,96 | -0,5357 | 0,391036013 | chr17 | - |
| Notch4 | 29,59 | 0,1910 | 0,782080553 | chr17 | + |
| Noxo1 | 6,86 | -0,1809 | 0,834675238 | chr17 | + |
| Nrm | 12,64 | -0,1019 | 0,90870562 | chr17 | + |
| Nrxn1 | 65,60 | 0,3296 | 0,537078872 | chr17 | - |
| Nthl1 | 72,32 | -0,1485 | 0,802706588 | chr17 | + |
| Ntn3 | 33,81 | -0,3828 | 0,480021428 | chr17 | - |
| Nubp2 | 455,39 | -0,1434 | 0,74259737 | chr17 | - |
| Nudt12 | 649,11 | 0,4570 | 0,271363618 | chr17 | - |
| Nudt3 | 460,02 | -0,2435 | 0,486225833 | chr17 | - |
| Oard1 | 682,67 | -0,1384 | 0,690021525 | chr17 | + |
| Paqr4 | 21,51 | 0,3435 | 0,570235502 | chr17 | - |
| Park2 | 24,88 | 0,5341 | 0,376702038 | chr17 | + |
| Pbx2 | 220,86 | -0,1450 | 0,764363417 | chr17 | + |
| Pdcd2 | 244,95 | -0,3711 | 0,266721655 | chr17 | - |
| Pde10a | 6,61 | -0,1225 | 0,89565156 | chr17 | + |
| Pde9a | 227,87 | 0,1130 | 0,834507118 | chr17 | + |
| Pdpk1 | 609,00 | 0,1780 | 0,636260727 | chr17 | - |
| Pex6 | 914,35 | -0,4452 | 0,340674125 | chr17 | + |
| Pglyrp2 | 155,02 | 0,2974 | 0,492901267 | chr17 | - |
| Phf1 | 136,83 | -0,5952 | 0,198075724 | chr17 | + |
| Pi16 | 2,14 | 0,0791 | NA | chr17 | + |
| Pigq | 579,25 | -0,0673 | 0,88569455 | chr17 | - |
| Pim1 | 114,08 | -0,2780 | 0,66442961 | chr17 | + |
| Pisd-ps2 | 61,08 | 0,2377 | 0,682596002 | chr17 | - |
| Pja2 | 628,98 | 0,2011 | 0,615783353 | chr17 | - |
| Pkd1 | 205,03 | -0,0923 | 0,842502441 | chr17 | + |
| Pkdcc | 211,27 | -0,2545 | 0,583782269 | chr17 | + |
| Pkmyt1 | 7,76 | -0,6444 | 0,323355495 | chr17 | + |
| Pknox1 | 188,84 | 0,1192 | 0,778952855 | chr17 | + |
| Pla2g7 | 130,02 | 0,1501 | 0,826227466 | chr17 | + |
| Plcl2 | 200,23 | -0,5142 | 0,105724638 | chr17 | + |
| Plekhh2 | 20,95 | 0,0504 | 0,952188756 | chr17 | + |
| Plg | 88,16 | 0,0385 | 0,953751362 | chr17 | + |
| Plin3 | 595,19 | 0,4455 | 0,236955722 | chr17 | - |
| Plin5 | 718,30 | -0,2058 | 0,740663168 | chr17 | - |
| Pnldc1 | 37,20 | -0,2600 | 0,691615955 | chr17 | - |
| Polh | 61,47 | 0,2856 | 0,559634089 | chr17 | - |
| Polr1c | 523,97 | -0,0891 | 0,817637582 | chr17 | - |
| Pot1b | 105,31 | 0,3317 | 0,43131925 | chr17 | - |
| Pp2d1 | 1,94 | 0,4753 | NA | chr17 | - |
| Ppard | 45,56 | -0,0971 | 0,893432234 | chr17 | + |
| Ppil1 | 137,37 | -0,3085 | 0,41633338 | chr17 | - |

| | | | | | |
|---|---|---|---|---|---|
| Ppm1b | 1450,64 | -0,1625 | 0,627637264 | chr17 | + |
| Ppp1r10 | 260,61 | -0,4869 | 0,335636465 | chr17 | + |
| Ppp1r11 | 614,06 | -0,0299 | 0,957017397 | chr17 | - |
| Ppp1r18 | 56,66 | 0,3819 | 0,376481973 | chr17 | + |
| Ppp1r21 | 253,90 | 0,0556 | 0,913452466 | chr17 | + |
| Ppp2r1a | 712,80 | -0,0768 | 0,875229732 | chr17 | + |
| Ppp2r5d | 351,64 | -0,5260 | 0,073523126 | chr17 | - |
| Ppp4r1 | 254,62 | -0,1472 | 0,70131463 | chr17 | + |
| Ppt2 | 331,01 | -0,1966 | 0,644399192 | chr17 | - |
| Pram1 | 10,39 | -0,2707 | 0,721569914 | chr17 | + |
| Prdm9 | 67,62 | 0,0124 | 0,986637152 | chr17 | - |
| Prepl | 376,28 | 0,0605 | 0,89879469 | chr17 | - |
| Prickle4 | 15,63 | -0,4947 | 0,428919534 | chr17 | - |
| Prkce | 180,33 | -0,1900 | 0,673340429 | chr17 | + |
| Prkd3 | 3033,57 | 0,0528 | 0,910948764 | chr17 | - |
| Prr22 | 2,23 | -0,1811 | NA | chr17 | + |
| Prr3 | 94,49 | -0,5032 | 0,331535469 | chr17 | - |
| Prrc2a | 474,19 | -0,1979 | 0,732140841 | chr17 | - |
| Prss30 | 2,36 | -0,1098 | NA | chr17 | - |
| Psmb1 | 3905,03 | -0,3392 | 0,323328805 | chr17 | - |
| Psmb8 | 667,78 | 0,4785 | 0,121424485 | chr17 | + |
| Ptcra | 12,23 | 0,5159 | 0,443670281 | chr17 | - |
| Ptk7 | 14,95 | 0,7234 | 0,235271103 | chr17 | - |
| Ptprm | 35,39 | 0,2147 | 0,7023898 | chr17 | - |
| Ptprs | 35,98 | 0,5112 | 0,347601558 | chr17 | - |
| Pxt1 | 3,23 | 0,7059 | 0,225814422 | chr17 | - |
| Qk | 908,85 | 0,1788 | 0,61720787 | chr17 | - |
| Rab11b | 812,02 | -0,1643 | 0,651839661 | chr17 | - |
| Rab11fip3 | 125,16 | 0,1501 | 0,761934432 | chr17 | - |
| Rab12 | 352,04 | 0,0841 | 0,869442758 | chr17 | - |
| Rab26os | 62,92 | -0,4631 | 0,262757002 | chr17 | + |
| Rab31 | 146,24 | 0,1519 | 0,727354216 | chr17 | - |
| Rab40c | 189,27 | -0,2970 | 0,471654089 | chr17 | - |
| Rab5a | 789,38 | -0,2312 | 0,516624153 | chr17 | + |
| Ralbp1 | 297,54 | -0,2174 | 0,595460103 | chr17 | - |
| Ranbp3 | 274,15 | -0,4185 | 0,298476336 | chr17 | + |
| Rasal3 | 9,74 | -0,1090 | 0,905147397 | chr17 | - |
| Rasgrp3 | 137,17 | 0,5399 | 0,08440425 | chr17 | + |
| Rcan2 | 77,83 | 0,0878 | 0,884882423 | chr17 | + |
| Rftn1 | 19,10 | -0,0368 | 0,969195493 | chr17 | - |
| Rfx2 | 9,21 | -0,1205 | 0,89468939 | chr17 | - |
| Rgl2 | 120,40 | -0,1826 | 0,777753633 | chr17 | + |
| Rgmb | 17,52 | 0,2427 | 0,749603956 | chr17 | - |
| Rhbdl1 | 30,03 | -0,8564 | 0,076193999 | chr17 | - |
| Rhoq | 151,42 | 0,0222 | 0,972672419 | chr17 | + |
| Rhot2 | 408,41 | -0,2742 | 0,472650412 | chr17 | - |
| Ring1 | 160,77 | -0,0741 | 0,912010517 | chr17 | - |
| Riok2 | 701,07 | 0,1339 | 0,771616285 | chr17 | + |
| Rmdn2 | 769,72 | -0,2573 | 0,506780377 | chr17 | + |
| Rnaset2b | 34,84 | -0,3959 | 0,506062256 | chr17 | + |

| | | | | | |
|---|---|---|---|---|---|
| Rnf39 | 40,87 | 0,4835 | 0,396411176 | chr17 | + |
| Rnf5 | 1363,88 | -0,3271 | 0,315238703 | chr17 | - |
| Rnf8 | 172,24 | -0,2939 | 0,41739988 | chr17 | + |
| Rnps1 | 297,57 | -0,1673 | 0,738645439 | chr17 | + |
| Rpl10a | 5180,01 | -0,3406 | 0,496793569 | chr17 | + |
| Rpl36 | 2520,17 | -0,4144 | 0,574604495 | chr17 | + |
| Rpl7l1 | 1436,67 | 0,0303 | 0,952989802 | chr17 | - |
| Rpp21 | 219,92 | -0,4943 | 0,299659421 | chr17 | - |
| Rps28 | 2747,03 | -0,4119 | 0,191230937 | chr17 | - |
| Rps6ka2 | 30,76 | -0,0210 | 0,981067951 | chr17 | + |
| Rpusd1 | 26,09 | 0,3269 | 0,570017362 | chr17 | + |
| Rrp1b | 42,53 | 0,4627 | 0,362428442 | chr17 | + |
| Rrp36 | 153,36 | -0,0203 | 0,971699327 | chr17 | - |
| Rsph1 | 3,76 | 0,0542 | 0,951934337 | chr17 | - |
| Rsph3a | 37,80 | -0,1873 | 0,793239112 | chr17 | + |
| Rsph3b | 23,77 | -0,2128 | 0,744313101 | chr17 | - |
| Rsph9 | 17,67 | -0,3332 | 0,624293109 | chr17 | - |
| Runx2 | 7,87 | 0,4024 | 0,579028345 | chr17 | - |
| Rxrb | 369,13 | -0,2665 | 0,545758572 | chr17 | + |
| Safb | 344,92 | -0,3838 | 0,377337243 | chr17 | + |
| Safb2 | 324,17 | -0,3049 | 0,560449199 | chr17 | - |
| Satb1 | 5,13 | -0,3598 | 0,634896352 | chr17 | - |
| Scaf8 | 250,76 | 0,1006 | 0,823734515 | chr17 | + |
| Sema6b | 42,74 | -0,1953 | 0,762922832 | chr17 | - |
| Serac1 | 66,91 | -0,1955 | 0,690415209 | chr17 | - |
| Sft2d1 | 344,93 | -0,2515 | 0,672205411 | chr17 | + |
| Sgo1 | 8,25 | -0,2970 | 0,707094339 | chr17 | - |
| Sh3gl1 | 219,34 | -0,0193 | 0,976528475 | chr17 | - |
| Sik1 | 294,28 | -0,3105 | 0,600401495 | chr17 | - |
| Skiv2l | 589,44 | -0,2917 | 0,471089317 | chr17 | - |
| Slc22a1 | 4772,95 | -0,4412 | 0,106945732 | chr17 | - |
| Slc22a3 | 66,94 | 0,1796 | 0,772525785 | chr17 | - |
| Slc25a27 | 13,71 | -0,0697 | 0,944005302 | chr17 | - |
| Slc29a1 | 3023,06 | 0,1082 | 0,794507134 | chr17 | - |
| Slc30a6 | 252,79 | 0,0904 | 0,826952398 | chr17 | + |
| Slc35b2 | 628,40 | 0,0648 | 0,869560578 | chr17 | + |
| Slc37a1 | 40,46 | 0,5622 | 0,358449334 | chr17 | + |
| Slc39a7 | 1389,59 | -0,3996 | 0,210391102 | chr17 | - |
| Slc44a4 | 9,05 | 0,1539 | 0,862616891 | chr17 | + |
| Slc9a3r2 | 176,30 | -0,4279 | 0,340033388 | chr17 | - |
| Smchd1 | 266,85 | 0,2446 | 0,575705463 | chr17 | - |
| Smoc2 | 135,13 | 0,4574 | 0,354139064 | chr17 | + |
| Snhg9 | 5,37 | -0,1793 | 0,837503804 | chr17 | - |
| Snora64 | 7,78 | 0,2269 | 0,785343688 | chr17 | + |
| Snora78 | 2,23 | -0,3573 | NA | chr17 | - |
| Snrpc | 277,94 | 0,1175 | 0,811619309 | chr17 | + |
| Snx9 | 327,30 | 0,0324 | 0,947182472 | chr17 | + |
| Socs5 | 102,01 | 0,1194 | 0,828658796 | chr17 | + |
| Sod2 | 3205,82 | -0,3799 | 0,119325906 | chr17 | + |
| Sos1 | 538,32 | -0,0874 | 0,817637582 | chr17 | - |

| | | | | | |
|---|---|---|---|---|---|
| Spaca6 | 13,05 | -0,8118 | 0,180849039 | chr17 | + |
| Spast | 539,82 | -0,3066 | 0,34812896 | chr17 | + |
| Spsb3 | 143,96 | -0,2220 | 0,630669284 | chr17 | + |
| Srbd1 | 313,83 | 0,3281 | 0,490650959 | chr17 | - |
| Srf | 108,33 | -0,0388 | 0,953751362 | chr17 | - |
| Srpk1 | 262,53 | -0,2975 | 0,483930201 | chr17 | - |
| Srrm2 | 1320,77 | -0,3141 | 0,430392697 | chr17 | + |
| Srsf3 | 1438,39 | 0,3408 | 0,493579462 | chr17 | + |
| Srsf7 | 678,02 | 0,3439 | 0,339404199 | chr17 | - |
| Stap2 | 601,62 | -0,3228 | 0,408496057 | chr17 | - |
| Stk19 | 93,36 | -0,4757 | 0,372536142 | chr17 | - |
| Stk38 | 715,10 | -0,3539 | 0,251516758 | chr17 | - |
| Ston1 | 39,15 | 0,8135 | 0,08440425 | chr17 | + |
| Strn | 58,17 | 0,1338 | 0,813415373 | chr17 | - |
| Stub1 | 459,34 | -0,5241 | 0,144343163 | chr17 | - |
| Sult1c2 | 141,38 | 0,2807 | 0,637638452 | chr17 | - |
| Supt3 | 61,19 | -0,1635 | 0,754989978 | chr17 | + |
| Syngap1 | 5,15 | 0,3921 | 0,600032204 | chr17 | + |
| Synj2 | 92,10 | 0,2822 | 0,557396844 | chr17 | + |
| Taf11 | 249,98 | -0,1477 | 0,675478019 | chr17 | - |
| Taf8 | 125,46 | -0,2411 | 0,571942598 | chr17 | - |
| Tagap | 4,65 | -0,0419 | 0,969130697 | chr17 | + |
| Tagap1 | 56,02 | -0,1237 | 0,834675238 | chr17 | - |
| Tap2 | 243,50 | 0,3229 | 0,47669554 | chr17 | + |
| Tapbp | 1096,24 | -0,1915 | 0,582441358 | chr17 | + |
| Tbc1d22b | 35,20 | -0,3604 | 0,527964623 | chr17 | + |
| Tbc1d24 | 294,87 | 0,2014 | 0,607310966 | chr17 | - |
| Tbc1d5 | 163,48 | 0,0795 | 0,866208949 | chr17 | - |
| Tbcc | 52,83 | 0,3279 | 0,478580716 | chr17 | + |
| Tbl3 | 206,07 | -0,3691 | 0,293404423 | chr17 | - |
| Tbp | 159,99 | 0,1037 | 0,813323745 | chr17 | + |
| Elob | 1833,06 | -0,3568 | 0,525098274 | chr17 | - |
| Tcf19 | 37,09 | -0,2807 | 0,604507219 | chr17 | - |
| Tcp1 | 2690,77 | -0,2925 | 0,298116176 | chr17 | + |
| Tcte2 | 5,39 | -0,5247 | 0,443292206 | chr17 | - |
| Tead3 | 60,88 | 0,0822 | 0,906240001 | chr17 | - |
| Telo2 | 66,92 | -0,0233 | 0,971699327 | chr17 | - |
| Tfb1m | 165,38 | 0,3597 | 0,41682625 | chr17 | - |
| Tfeb | 57,67 | -0,7931 | 0,131014868 | chr17 | + |
| Tgif1 | 251,03 | 0,3812 | 0,38301652 | chr17 | - |
| Thada | 89,87 | 0,0871 | 0,900327088 | chr17 | - |
| Thbs2 | 47,61 | 0,5209 | 0,306891735 | chr17 | - |
| Thoc6 | 102,04 | -0,1602 | 0,759339376 | chr17 | - |
| Thumpd2 | 37,11 | -0,0147 | 0,986818215 | chr17 | - |
| Ticam1 | 162,27 | 0,1471 | 0,77154104 | chr17 | - |
| Tjap1 | 53,66 | -0,4170 | 0,386872469 | chr17 | - |
| Tmem178 | 10,89 | 0,6423 | 0,294304263 | chr17 | + |
| Tmem181a | 42,15 | 0,1927 | 0,739141404 | chr17 | + |
| Tmem181b-ps | 28,98 | 0,4178 | 0,492095471 | chr17 | - |
| Tmem204 | 292,87 | 0,4512 | 0,201443085 | chr17 | - |

| | | | | | |
|---|---|---|---|---|---|
| Tmem242 | 920,02 | -0,0866 | 0,804380135 | chr17 | - |
| Tmem63b | 642,90 | -0,3913 | 0,263205799 | chr17 | - |
| Tmem8 | 395,61 | 0,2133 | 0,618028988 | chr17 | + |
| Tnf | 3,89 | 0,1116 | 0,905147397 | chr17 | - |
| Tnfaip8l1 | 407,79 | -0,4378 | 0,340012332 | chr17 | + |
| Tnfrsf12a | 150,65 | -0,3370 | 0,618749555 | chr17 | - |
| Tnfrsf21 | 79,24 | 0,2929 | 0,540877577 | chr17 | + |
| Tnfsf14 | 4,49 | 0,2693 | 0,740024898 | chr17 | - |
| Tnxb | 101,71 | -0,6372 | 0,110056031 | chr17 | + |
| Tomm6 | 1237,43 | -0,4583 | 0,370069913 | chr17 | - |
| Tomm6os | 4,29 | -0,8083 | 0,186048813 | chr17 | + |
| Traf7 | 136,98 | -0,2527 | 0,559032648 | chr17 | - |
| Trem2 | 14,43 | 0,6628 | 0,28821012 | chr17 | + |
| Trem3 | 2,14 | 0,1919 | NA | chr17 | + |
| Treml2 | 7,89 | 0,3166 | 0,682099278 | chr17 | + |
| Treml4 | 87,62 | 0,1815 | 0,736302391 | chr17 | + |
| Trerf1 | 2,59 | 0,2375 | 0,761282414 | chr17 | + |
| Trim26 | 478,60 | 0,3691 | 0,240623485 | chr17 | + |
| Trim39 | 158,72 | -0,4799 | 0,227688754 | chr17 | - |
| Trip10 | 1985,78 | -0,0272 | 0,950625589 | chr17 | + |
| Trmt61b | 61,25 | 0,2900 | 0,576013884 | chr17 | - |
| Trp53cor1 | 2,02 | 0,5820 | NA | chr17 | - |
| Tsc2 | 401,84 | -0,4867 | 0,13621142 | chr17 | - |
| Tsr3 | 171,67 | -0,2772 | 0,480652365 | chr17 | + |
| Ttbk1 | 8,63 | 0,6891 | 0,271016733 | chr17 | - |
| Ttc27 | 12,92 | 0,3965 | 0,562291762 | chr17 | + |
| Ttc7 | 436,90 | 0,0295 | 0,963868185 | chr17 | + |
| Tubb4a | 2,86 | 0,0839 | 0,925700434 | chr17 | - |
| Tubb5 | 752,27 | -0,0861 | 0,872774688 | chr17 | - |
| Tulp4 | 388,39 | -0,1691 | 0,619667783 | chr17 | + |
| Twsg1 | 812,20 | -0,0875 | 0,813510321 | chr17 | - |
| U2af1 | 947,86 | -0,1011 | 0,815690724 | chr17 | - |
| Ube2i | 855,24 | -0,0075 | 0,986889999 | chr17 | - |
| Ubr2 | 929,68 | -0,2833 | 0,255281519 | chr17 | - |
| Ubxn6 | 544,39 | -0,4910 | 0,246313798 | chr17 | - |
| Uhrf1 | 10,63 | -0,2862 | 0,710225448 | chr17 | + |
| Uhrf1bp1 | 301,79 | -0,0484 | 0,916852736 | chr17 | + |
| Unc93a | 31,24 | 0,5633 | 0,291574528 | chr17 | - |
| Unkl | 228,85 | 0,1318 | 0,765009643 | chr17 | + |
| Uqcc2 | 588,50 | -0,4474 | 0,357290285 | chr17 | - |
| Usp49 | 37,27 | 0,1317 | 0,849477252 | chr17 | + |
| Vapa | 1258,73 | -0,3444 | 0,191889357 | chr17 | - |
| Vars | 208,86 | -0,6753 | 0,098825827 | chr17 | + |
| Vars2 | 177,34 | -0,4196 | 0,270497584 | chr17 | - |
| Vav1 | 55,19 | 0,1016 | 0,881274343 | chr17 | + |
| Vegfa | 1155,29 | -0,1691 | 0,738282454 | chr17 | - |
| Vmac | 143,63 | -0,1769 | 0,691018693 | chr17 | - |
| Vps52 | 163,12 | -0,1188 | 0,811768517 | chr17 | + |
| Vwa7 | 4,07 | -0,1234 | 0,89147692 | chr17 | + |
| Washc1 | 190,69 | -0,2203 | 0,611914561 | chr17 | + |

| | | | | | |
|---|---|---|---|---|---|
| Wdr24 | 171,49 | -0,1081 | 0,809842452 | chr17 | + |
| Wdr4 | 141,34 | -0,5209 | 0,138292882 | chr17 | - |
| Wdr43 | 431,76 | -0,0138 | 0,981067951 | chr17 | + |
| Wdr46 | 195,36 | -0,3310 | 0,305042283 | chr17 | + |
| Wdr90 | 43,61 | 0,1289 | 0,850915997 | chr17 | - |
| Wfikkn1 | 3,41 | -0,4344 | 0,535573923 | chr17 | - |
| Wiz | 89,40 | -0,3683 | 0,397488617 | chr17 | - |
| Wtap | 361,11 | 0,1682 | 0,739722032 | chr17 | - |
| Xdh | 1636,23 | -0,0427 | 0,939521529 | chr17 | - |
| Xpo5 | 202,62 | 0,1011 | 0,830330662 | chr17 | + |
| Yipf3 | 578,28 | -0,4180 | 0,194733334 | chr17 | + |
| Yipf4 | 626,41 | 0,0966 | 0,831614489 | chr17 | + |
| Ypel5 | 1481,01 | -0,3321 | 0,235102108 | chr17 | + |
| Zbtb12 | 54,75 | -0,1673 | 0,82704652 | chr17 | + |
| Zbtb14 | 158,80 | 0,1637 | 0,682099278 | chr17 | + |
| Zbtb22 | 146,19 | -0,0219 | 0,975445719 | chr17 | + |
| Zbtb9 | 110,78 | 0,2901 | 0,444265933 | chr17 | + |
| Zfand3 | 939,51 | -0,3850 | 0,319311067 | chr17 | + |
| Zfp101 | 112,17 | 0,3239 | 0,398141536 | chr17 | - |
| Zfp119a | 96,73 | 0,4918 | 0,233544599 | chr17 | - |
| Zfp119b | 107,32 | 0,5016 | 0,168720631 | chr17 | - |
| Zfp160 | 167,39 | 0,2143 | 0,628393534 | chr17 | + |
| Zfp213 | 83,12 | 0,4609 | 0,310273443 | chr17 | - |
| Zfp318 | 338,73 | -0,4588 | 0,103473805 | chr17 | + |
| Zfp36l2 | 717,64 | -0,4688 | 0,332014851 | chr17 | - |
| Zfp40 | 48,80 | 0,1284 | 0,838868977 | chr17 | - |
| Zfp414 | 40,52 | -0,2016 | 0,747526723 | chr17 | + |
| Zfp472 | 93,79 | 0,2846 | 0,516856098 | chr17 | + |
| Zfp51 | 113,25 | 0,5008 | 0,161161606 | chr17 | + |
| Zfp52 | 29,68 | 0,6588 | 0,224150405 | chr17 | + |
| Zfp523 | 143,28 | -0,4408 | 0,378908981 | chr17 | + |
| Zfp54 | 92,87 | 0,6109 | 0,110800147 | chr17 | + |
| Zfp57 | 7,98 | -0,0602 | 0,951806422 | chr17 | + |
| Zfp758 | 151,18 | 0,5519 | 0,156728357 | chr17 | + |
| Zfp760 | 103,14 | 0,4022 | 0,352462246 | chr17 | + |
| Zfp81 | 29,92 | 0,5230 | 0,302314196 | chr17 | - |
| Zfp811 | 3,80 | -0,1093 | 0,901724181 | chr17 | - |
| Zfp820 | 63,43 | 0,5888 | 0,154188248 | chr17 | - |
| Zfp871 | 1430,77 | -0,1657 | 0,69567579 | chr17 | - |
| Zfp942 | 303,67 | 0,5030 | 0,152776166 | chr17 | - |
| Zfp943 | 230,00 | 0,0716 | 0,886483612 | chr17 | + |
| Zfp944 | 241,79 | 0,5039 | 0,124135408 | chr17 | - |
| Zfp945 | 265,20 | 0,2731 | 0,5751772 | chr17 | - |
| Zfp946 | 379,88 | 0,1702 | 0,707267324 | chr17 | + |
| Zfp948 | 96,38 | 0,3465 | 0,464245504 | chr17 | + |
| Zfp959 | 146,11 | 0,5212 | 0,065752252 | chr17 | + |
| Zfp960 | 45,54 | 0,6203 | 0,203863873 | chr17 | + |
| Zfp97 | 44,56 | 0,6464 | 0,134008263 | chr17 | + |
| Znrd1 | 345,16 | 0,0585 | 0,906240001 | chr17 | - |
| Znrd1as | 286,08 | 0,0338 | 0,953751362 | chr17 | + |

| | | | | | |
|---|---|---|---|---|---|
| 0610009O20Rik | 280,01 | 0,1113 | 0,796874179 | chr18 | + |
| 1010001N08Rik | 27,65 | -0,2876 | 0,64675918 | chr18 | - |
| 1500015A07Rik | 21,92 | -0,1450 | 0,850396791 | chr18 | + |
| 1700066B19Rik | 2,50 | 0,3618 | NA | chr18 | + |
| 1700086O06Rik | 22,77 | -0,0863 | 0,912811384 | chr18 | - |
| Epb41l4aos | 94,51 | -0,3297 | 0,668382348 | chr18 | + |
| 2700062C07Rik | 127,30 | 0,1459 | 0,717188441 | chr18 | + |
| 3110002H16Rik | 387,76 | -0,2390 | 0,392025719 | chr18 | + |
| 4833419F23Rik | 7,11 | 0,3294 | 0,669280735 | chr18 | + |
| 4930503L19Rik | 28,58 | 0,0675 | 0,940578009 | chr18 | - |
| 4930511M06Rik | 18,27 | -0,4401 | 0,464848266 | chr18 | - |
| 4933408B17Rik | 3,57 | 0,0319 | 0,976788115 | chr18 | - |
| 8030462N17Rik | 88,63 | 0,0387 | 0,952015353 | chr18 | - |
| 9030625G05Rik | 3,82 | 0,4743 | 0,440094295 | chr18 | - |
| A830052D11Rik | 26,49 | -0,3857 | 0,498040757 | chr18 | - |
| A930012L18Rik | 4,54 | 0,6063 | 0,346659901 | chr18 | + |
| Abhd3 | 976,68 | -0,2721 | 0,298774903 | chr18 | - |
| Ablim3 | 260,91 | -0,4005 | 0,374392563 | chr18 | - |
| Acaa2 | 19983,55 | -0,2867 | 0,31970856 | chr18 | + |
| Adrb2 | 19,83 | -0,0440 | 0,958789475 | chr18 | - |
| Afap1l1 | 81,71 | 0,3045 | 0,498546742 | chr18 | - |
| Afg3l2 | 1069,54 | -0,2065 | 0,501000077 | chr18 | - |
| Aldh7a1 | 6662,10 | -0,1729 | 0,525033727 | chr18 | - |
| Ammecr1l | 1107,03 | 0,1260 | 0,740352128 | chr18 | + |
| Ankhd1 | 474,65 | 0,3252 | 0,245406077 | chr18 | + |
| Ankrd29 | 8,93 | 0,1390 | 0,875621033 | chr18 | - |
| Ap3s1 | 589,99 | -0,3577 | 0,3845875 | chr18 | + |
| Apbb3 | 159,96 | 0,2818 | 0,605371398 | chr18 | - |
| Apc | 398,18 | -0,0823 | 0,826029548 | chr18 | + |
| Aqp4 | 87,37 | -0,1605 | 0,795600492 | chr18 | - |
| Arap3 | 118,25 | -0,1751 | 0,717725572 | chr18 | - |
| Arhgap12 | 515,21 | 0,0498 | 0,909889201 | chr18 | - |
| Arhgef37 | 85,93 | 0,4763 | 0,385522252 | chr18 | - |
| Atg12 | 608,36 | -0,0867 | 0,841883711 | chr18 | - |
| Atp5a1 | 15573,32 | -0,2923 | 0,197774355 | chr18 | + |
| Atp8b1 | 663,24 | 0,3051 | 0,319805888 | chr18 | - |
| Atp9b | 428,96 | 0,0266 | 0,95463264 | chr18 | - |
| AW554918 | 38,87 | 0,2865 | 0,608287564 | chr18 | + |
| B430212C06Rik | 2,73 | 0,8087 | 0,124945987 | chr18 | - |
| B4galt6 | 41,40 | 0,3471 | 0,513640882 | chr18 | - |
| Bambi | 69,72 | -0,3776 | 0,433687525 | chr18 | + |
| BC031181 | 1323,79 | -0,0073 | 0,990189789 | chr18 | + |
| Brd8 | 213,42 | 0,1611 | 0,740663168 | chr18 | - |
| C330018D20Rik | 302,32 | 0,7150 | 0,051507926 | chr18 | - |
| Camk4 | 4,28 | 0,0691 | 0,944852187 | chr18 | + |
| Ccbe1 | 90,01 | 0,2742 | 0,585010146 | chr18 | - |
| Ccdc112 | 3,68 | -0,2017 | 0,8112244 | chr18 | - |
| Ccdc68 | 38,03 | 0,3914 | 0,449823999 | chr18 | + |
| Ccny | 1239,86 | 0,0304 | 0,947294154 | chr18 | - |
| Cd14 | 88,63 | 0,3204 | 0,581489212 | chr18 | - |

| | | | | | |
|---|---|---|---|---|---|
| Cd226 | 7,67 | 0,8260 | 0,177510828 | chr18 | + |
| Cd74 | 1317,55 | 0,4371 | 0,250762967 | chr18 | + |
| Cdc23 | 150,57 | 0,2165 | 0,604679897 | chr18 | - |
| Cdh2 | 1473,16 | -0,0510 | 0,907824791 | chr18 | - |
| Cdo1 | 20689,39 | -0,3718 | 0,440094295 | chr18 | - |
| Celf4 | 4,18 | -0,5534 | 0,406538603 | chr18 | - |
| Cep120 | 215,64 | 0,2110 | 0,594362475 | chr18 | - |
| Cep192 | 45,93 | 0,4315 | 0,402148509 | chr18 | + |
| Cep76 | 72,21 | -0,2605 | 0,58928611 | chr18 | - |
| Chmp1b | 602,38 | 0,2758 | 0,413615196 | chr18 | + |
| Chsy3 | 54,50 | -0,6907 | 0,078316133 | chr18 | + |
| Cndp2 | 730,02 | 0,2162 | 0,517086052 | chr18 | - |
| Commd10 | 120,16 | 0,2585 | 0,55880551 | chr18 | + |
| Crem | 678,15 | -0,3345 | 0,466983871 | chr18 | - |
| Csf1r | 789,12 | 0,0282 | 0,950625589 | chr18 | + |
| Csnk1a1 | 802,11 | 0,0043 | 0,992196557 | chr18 | + |
| Csnk1g3 | 720,95 | -0,1235 | 0,791544358 | chr18 | + |
| Ctdp1 | 77,09 | -0,5439 | 0,235194942 | chr18 | - |
| Ctif | 298,06 | -0,1070 | 0,862599499 | chr18 | - |
| Ctnna1 | 1570,26 | -0,0310 | 0,947294154 | chr18 | + |
| Cxxc1 | 344,76 | -0,3475 | 0,300779985 | chr18 | + |
| Cxxc5 | 443,30 | -0,3498 | 0,504721936 | chr18 | + |
| Cyb5a | 28127,98 | -0,2890 | 0,466812905 | chr18 | + |
| Cystm1 | 110,69 | -0,0397 | 0,957444102 | chr18 | + |
| Dcc | 3,76 | -0,3334 | 0,656913078 | chr18 | - |
| Dcp2 | 295,17 | 0,3320 | 0,338536613 | chr18 | + |
| Dctn4 | 456,80 | 0,0559 | 0,900980922 | chr18 | + |
| Diaph1 | 346,76 | 0,2395 | 0,463352295 | chr18 | - |
| Dmxl1 | 382,97 | 0,1305 | 0,799103023 | chr18 | + |
| Dnd1 | 7,27 | -0,3262 | 0,672801255 | chr18 | - |
| Dpysl3 | 40,12 | 0,1907 | 0,746827664 | chr18 | - |
| Dsc2 | 991,66 | 0,1742 | 0,757883527 | chr18 | - |
| Dsg1a | 3,96 | -0,6661 | 0,276877002 | chr18 | + |
| Dsg1c | 46,69 | -0,4847 | 0,464752915 | chr18 | + |
| Dsg2 | 2086,80 | 0,4523 | 0,129703094 | chr18 | + |
| Dtna | 4,11 | 0,1555 | 0,86019237 | chr18 | + |
| Dtwd2 | 81,11 | 0,2477 | 0,568858283 | chr18 | - |
| Egr1 | 295,03 | 0,8075 | 0,175181057 | chr18 | + |
| Eif1a | 1388,78 | 0,4753 | 0,132069532 | chr18 | + |
| Eif4ebp3 | 941,10 | -0,3502 | 0,473786638 | chr18 | + |
| Elac1 | 110,46 | 0,3250 | 0,522469399 | chr18 | - |
| Elp2 | 675,17 | 0,2009 | 0,672205411 | chr18 | + |
| Eno1b | 10,99 | 0,0298 | 0,978280656 | chr18 | + |
| Epb41l4a | 3,19 | -0,3079 | 0,674998094 | chr18 | - |
| Epc1 | 222,29 | -0,2631 | 0,39766425 | chr18 | - |
| Epg5 | 289,16 | -0,0298 | 0,953487537 | chr18 | + |
| Ercc3 | 305,24 | -0,2471 | 0,559252792 | chr18 | + |
| Esco1 | 294,01 | 0,3016 | 0,366528137 | chr18 | - |
| Etf1 | 2023,16 | -0,1890 | 0,549054641 | chr18 | - |
| F830016B08Rik | 153,05 | 0,1296 | 0,827995984 | chr18 | + |

| | | | | | |
|---|---|---|---|---|---|
| Fam13b | 238,53 | 0,0082 | 0,986889999 | chr18 | - |
| Fam53c | 247,65 | -0,2259 | 0,509917962 | chr18 | + |
| Fbxo38 | 441,14 | 0,0249 | 0,952541585 | chr18 | - |
| Fchsd1 | 15,72 | 0,6370 | 0,288409641 | chr18 | - |
| Fech | 2372,08 | -0,2784 | 0,29233966 | chr18 | - |
| Fem1c | 113,12 | 0,4282 | 0,285499423 | chr18 | - |
| Fgf1 | 1609,92 | -0,1742 | 0,724652712 | chr18 | - |
| Fhod3 | 14,39 | -0,3925 | 0,588008527 | chr18 | + |
| G630055G22Rik | 3,21 | 0,3554 | 0,620367088 | chr18 | + |
| G630071F17Rik | 5,33 | -0,0947 | 0,921261454 | chr18 | + |
| Galnt1 | 1064,11 | -0,1276 | 0,7595822 | chr18 | + |
| Garem1 | 178,99 | 0,1978 | 0,660583784 | chr18 | - |
| Gata6 | 53,09 | 0,0809 | 0,907233333 | chr18 | + |
| Gm10556 | 5,56 | 0,2046 | 0,8112244 | chr18 | + |
| Bvht | 20,45 | 0,1624 | 0,823734515 | chr18 | + |
| Gm4013 | 103,67 | -0,6487 | 0,053014128 | chr18 | - |
| Gm4841 | 80,29 | 0,5460 | 0,24579162 | chr18 | - |
| Gm4951 | 2294,57 | 0,4302 | 0,323960783 | chr18 | + |
| Gm6277 | 2,43 | 0,5532 | NA | chr18 | - |
| Gnal | 12,92 | -0,7157 | 0,23305528 | chr18 | + |
| Gnpda1 | 224,80 | 0,4304 | 0,27946684 | chr18 | - |
| Gramd3 | 860,49 | -0,3817 | 0,168336507 | chr18 | + |
| Grpel2 | 1137,51 | -0,1861 | 0,750954651 | chr18 | - |
| Gypc | 109,67 | 0,5631 | 0,244000233 | chr18 | - |
| Hars | 697,34 | -0,0580 | 0,88569455 | chr18 | - |
| Hars2 | 402,81 | -0,1765 | 0,592873282 | chr18 | + |
| Haus1 | 130,16 | -0,0562 | 0,925898178 | chr18 | - |
| Hbegf | 17,46 | 0,6203 | 0,29105938 | chr18 | - |
| Hdac3 | 520,94 | -0,1273 | 0,765231448 | chr18 | - |
| Hdhd2 | 507,78 | -0,0570 | 0,88127014 | chr18 | + |
| Hmgxb3 | 138,44 | -0,2482 | 0,501951423 | chr18 | - |
| Hsbp1l1 | 77,97 | 0,5279 | 0,201179976 | chr18 | - |
| Hsd17b4 | 6568,11 | -0,4382 | 0,145007441 | chr18 | + |
| Hspa9 | 10869,44 | -0,2029 | 0,423813652 | chr18 | - |
| Ier3ip1 | 710,89 | -0,4902 | 0,239971296 | chr18 | + |
| Igip | 226,77 | -0,4083 | 0,214659181 | chr18 | + |
| Iigp1 | 10355,33 | 0,3051 | 0,473368945 | chr18 | + |
| Ik | 1309,33 | 0,0822 | 0,842925053 | chr18 | + |
| Impa2 | 24,48 | -0,2317 | 0,757565313 | chr18 | + |
| Impact | 340,16 | 0,6066 | 0,114525976 | chr18 | + |
| Ino80c | 257,86 | 0,1670 | 0,678676127 | chr18 | - |
| Isoc1 | 982,05 | -0,1399 | 0,695967514 | chr18 | + |
| Iws1 | 376,78 | 0,0644 | 0,876803328 | chr18 | + |
| Kcnn2 | 347,80 | 0,0464 | 0,937853416 | chr18 | + |
| Kctd1 | 6,54 | -0,0780 | 0,938364545 | chr18 | - |
| Kdm3b | 270,49 | 0,3128 | 0,384921569 | chr18 | + |
| Kif20a | 22,73 | 0,4961 | 0,416450614 | chr18 | + |
| Kif5b | 703,23 | 0,0273 | 0,958859839 | chr18 | - |
| Lama3 | 144,16 | 0,6384 | 0,068488954 | chr18 | + |
| Lars | 678,55 | 0,0276 | 0,96674257 | chr18 | - |

| | | | | | |
|---|---|---|---|---|---|
| Ldlrad4 | 45,26 | -0,3258 | 0,524801822 | chr18 | + |
| Lims2 | 649,28 | -0,3899 | 0,325839875 | chr18 | + |
| Lipg | 341,82 | 0,1540 | 0,753233421 | chr18 | - |
| Lman1 | 4128,07 | -0,0228 | 0,952576171 | chr18 | - |
| Lmnb1 | 69,82 | 0,5390 | 0,173216656 | chr18 | + |
| Lox | 9,71 | 0,7039 | 0,260028246 | chr18 | - |
| Malt1 | 117,45 | -0,0285 | 0,960969671 | chr18 | + |
| Map3k2 | 484,74 | -0,1204 | 0,772705111 | chr18 | + |
| Map3k8 | 44,97 | 0,2923 | 0,56353816 | chr18 | - |
| Mapre2 | 241,04 | -0,1767 | 0,692167613 | chr18 | + |
| March3 | 5,86 | 0,2548 | 0,757395951 | chr18 | - |
| Matr3 | 1720,80 | 0,2012 | 0,598404146 | chr18 | + |
| Mbd2 | 619,32 | -0,2908 | 0,274206913 | chr18 | + |
| Mbp | 14,99 | 0,3828 | 0,587879802 | chr18 | + |
| Mcc | 403,31 | -0,0408 | 0,943305207 | chr18 | - |
| Me2 | 61,64 | 0,3550 | 0,412895523 | chr18 | - |
| Mep1b | 4,09 | 0,2800 | 0,725248714 | chr18 | + |
| Mex3c | 323,00 | -0,0869 | 0,844618872 | chr18 | + |
| Mib1 | 169,30 | 0,2916 | 0,403302288 | chr18 | + |
| Mir122 | 5,70 | -0,8211 | 0,182954297 | chr18 | + |
| Carmn | 6,67 | -0,4087 | 0,579355496 | chr18 | - |
| Mkx | 32,09 | -0,6043 | 0,342277341 | chr18 | - |
| Mocos | 809,77 | -0,0192 | 0,970667732 | chr18 | + |
| Mpp7 | 124,95 | -0,0314 | 0,968763097 | chr18 | - |
| Mppe1 | 114,10 | 0,4239 | 0,391907358 | chr18 | - |
| Mtpap | 260,19 | -0,0552 | 0,912811384 | chr18 | + |
| Myo5b | 283,49 | 0,3003 | 0,476319975 | chr18 | + |
| Myoz3 | 7,73 | -0,4379 | 0,543578711 | chr18 | - |
| Mzb1 | 4,70 | 0,1437 | 0,874467051 | chr18 | - |
| Napg | 271,92 | -0,0662 | 0,88569455 | chr18 | + |
| Nars | 2177,17 | -0,1138 | 0,760736495 | chr18 | - |
| Ndfip1 | 1901,09 | -0,1920 | 0,495713543 | chr18 | + |
| Ndst1 | 1015,45 | 0,4777 | 0,131674744 | chr18 | - |
| Ndufa2 | 2561,15 | -0,3900 | 0,413323375 | chr18 | - |
| Nedd4l | 485,19 | 0,6688 | 0,077224204 | chr18 | + |
| Nfatc1 | 87,34 | -0,2804 | 0,48546604 | chr18 | - |
| Nme5 | 5,25 | 1,0036 | 0,078316133 | chr18 | - |
| Npc1 | 2103,78 | 0,0164 | 0,97437659 | chr18 | - |
| Nr3c1 | 1372,16 | 0,0799 | 0,844155615 | chr18 | - |
| Nrep | 963,55 | 0,3718 | 0,621775664 | chr18 | - |
| Onecut2 | 1264,67 | -0,3282 | 0,298814514 | chr18 | + |
| Osbpl1a | 2178,67 | 0,0173 | 0,966776406 | chr18 | - |
| Paip2 | 2364,07 | -0,0964 | 0,815188933 | chr18 | + |
| Pard6g | 6,57 | 0,1424 | 0,875717638 | chr18 | + |
| Pcdh1 | 2486,57 | -0,2072 | 0,557563183 | chr18 | - |
| Pcdh12 | 355,01 | -0,0543 | 0,932531696 | chr18 | - |
| Pcdhb12 | 2,59 | -0,0411 | 0,966637266 | chr18 | + |
| Pcdhb14 | 8,94 | 0,6170 | 0,339967066 | chr18 | + |
| Pcdhb16 | 6,36 | 0,3328 | 0,666980604 | chr18 | + |
| Pcdhb17 | 7,23 | 0,0605 | 0,952076682 | chr18 | + |

| | | | | | |
|---|---|---|---|---|---|
| Pcdhb18 | 2,96 | 0,5342 | 0,39766425 | chr18 | + |
| Pcdhb20 | 2,60 | 0,2351 | 0,764664236 | chr18 | + |
| Pcdhb22 | 2,48 | 0,1825 | NA | chr18 | + |
| Pcdhb4 | 2,13 | 0,0281 | NA | chr18 | + |
| Pcdhga1 | 1,80 | 0,5041 | NA | chr18 | + |
| Pcdhga12 | 2,61 | 0,0679 | 0,939403124 | chr18 | + |
| Pcdhgc3 | 2,34 | 0,2500 | NA | chr18 | + |
| Pcdhga2 | 1,90 | 0,5615 | NA | chr18 | + |
| Pcdhga4 | 4,85 | 0,5461 | 0,417486404 | chr18 | + |
| Pcdhgb2 | 176,92 | -0,0874 | 0,855566249 | chr18 | + |
| Pcyox1l | 11,78 | -0,1076 | 0,905147397 | chr18 | - |
| Pdgfrb | 92,37 | 0,2149 | 0,694914738 | chr18 | + |
| Pfdn1 | 1002,61 | 0,0679 | 0,871084117 | chr18 | - |
| Pggt1b | 9267,09 | -0,2659 | 0,353377472 | chr18 | - |
| Phax | 363,93 | -0,1804 | 0,640948606 | chr18 | + |
| Pias2 | 514,55 | -0,4464 | 0,101796122 | chr18 | + |
| Piezo2 | 10,10 | -0,0332 | 0,975445719 | chr18 | - |
| Pik3c3 | 73,46 | 0,9171 | 0,10210139 | chr18 | + |
| Pkd2l2 | 4,17 | 0,0587 | 0,952989802 | chr18 | + |
| Pmaip1 | 23,43 | 0,5164 | 0,358990473 | chr18 | + |
| Poli | 59,73 | 0,0960 | 0,87206597 | chr18 | - |
| Polr2d | 142,46 | 0,0837 | 0,880810135 | chr18 | + |
| Ppargc1b | 49,42 | 0,0184 | 0,985702623 | chr18 | - |
| Ppp2r2b | 2,91 | 0,1588 | 0,851988359 | chr18 | - |
| Pqlc1 | 1214,10 | 0,0551 | 0,915227882 | chr18 | + |
| Proc | 9219,71 | -0,3857 | 0,204328779 | chr18 | - |
| Prr16 | 410,63 | -0,3367 | 0,357290285 | chr18 | + |
| Prrc1 | 317,52 | -0,1520 | 0,696532876 | chr18 | + |
| Psma8 | 24,90 | 0,4463 | 0,489321662 | chr18 | + |
| Psmg2 | 337,08 | 0,1126 | 0,785857392 | chr18 | + |
| Ptpn2 | 262,50 | -0,0590 | 0,893432234 | chr18 | - |
| Pura | 25,09 | -0,3510 | 0,578442407 | chr18 | + |
| Rab18 | 2329,93 | 0,2064 | 0,561773364 | chr18 | + |
| Rab27b | 24,72 | 0,2981 | 0,645204605 | chr18 | - |
| Rbbp8 | 170,55 | 0,0359 | 0,949529403 | chr18 | + |
| Rbfa | 272,24 | -0,5349 | 0,06737212 | chr18 | - |
| Rbm22 | 181,00 | 0,2076 | 0,583654143 | chr18 | + |
| Rbm27 | 301,35 | -0,1021 | 0,798995815 | chr18 | + |
| Reep5 | 428,41 | 0,0297 | 0,952188756 | chr18 | - |
| Rell2 | 2,43 | -0,3864 | NA | chr18 | + |
| Riok3 | 1524,71 | -0,3990 | 0,067820892 | chr18 | + |
| Rnf125 | 459,19 | 0,4959 | 0,242907451 | chr18 | + |
| Rnf138 | 285,92 | -0,1531 | 0,703924654 | chr18 | + |
| Rnf14 | 1237,72 | -0,0655 | 0,879476378 | chr18 | + |
| Rnf165 | 9,46 | 0,3569 | 0,626098415 | chr18 | - |
| Rnmt | 278,65 | -0,0957 | 0,83176435 | chr18 | + |
| Rock1 | 672,89 | 0,1294 | 0,762940356 | chr18 | - |
| Rpl17 | 9656,87 | -0,1449 | 0,692167613 | chr18 | + |
| Rttn | 25,55 | 0,6469 | 0,313447648 | chr18 | + |
| Sap130 | 233,57 | -0,1170 | 0,831240409 | chr18 | + |

| | | | | | |
|---|---|---|---|---|---|
| Scarna17 | 3,67 | -0,5049 | 0,458688593 | chr18 | - |
| Sec11c | 613,45 | -0,5956 | 0,112275385 | chr18 | + |
| Seh1l | 759,35 | -0,0509 | 0,906171442 | chr18 | + |
| Sema6a | 77,71 | 0,0360 | 0,950819236 | chr18 | - |
| Setbp1 | 7,21 | 0,3645 | 0,630199445 | chr18 | - |
| Sft2d3 | 51,36 | -0,1061 | 0,857665028 | chr18 | - |
| Sh3tc2 | 14,94 | 0,4009 | 0,542534449 | chr18 | + |
| Sil1 | 730,11 | -0,3103 | 0,376702038 | chr18 | - |
| Ska1 | 2,29 | 0,0998 | NA | chr18 | - |
| Slc12a2 | 134,10 | -0,4745 | 0,178097604 | chr18 | + |
| Slc14a1 | 2,65 | 0,0666 | 0,943108135 | chr18 | - |
| Slc23a1 | 1103,02 | 0,3789 | 0,358968582 | chr18 | - |
| Slc25a2 | 3,79 | -0,6063 | 0,317123683 | chr18 | - |
| Slc25a46 | 980,22 | -0,1944 | 0,618509515 | chr18 | - |
| Slc26a2 | 30,68 | 0,6309 | 0,186783528 | chr18 | - |
| Slc27a6 | 3,30 | -0,0743 | 0,932528423 | chr18 | + |
| Slc35a4 | 173,62 | -0,0471 | 0,918997005 | chr18 | + |
| Slc39a6 | 50,37 | -0,2287 | 0,731048653 | chr18 | - |
| Slc6a7 | 2,58 | -0,1173 | 0,88569455 | chr18 | - |
| Smad2 | 349,16 | -0,1173 | 0,760844082 | chr18 | + |
| Smad4 | 309,67 | -0,3800 | 0,253908866 | chr18 | - |
| Smad7 | 75,34 | 0,2068 | 0,680276822 | chr18 | + |
| Smim3 | 12,69 | 0,0084 | 0,993445259 | chr18 | - |
| Sncaip | 7,37 | 0,0718 | 0,943108135 | chr18 | + |
| Snrpd1 | 427,84 | 0,1788 | 0,709191439 | chr18 | + |
| Snx2 | 1224,47 | -0,0180 | 0,967498977 | chr18 | + |
| Snx24 | 209,13 | 0,2569 | 0,489636271 | chr18 | + |
| Socs6 | 150,81 | -0,1239 | 0,786009002 | chr18 | - |
| Spata24 | 35,95 | -0,1394 | 0,841579427 | chr18 | - |
| Spink1 | 53,49 | 1,7823 | NA | chr18 | - |
| Spire1 | 8,37 | 0,0014 | 0,998482911 | chr18 | - |
| Spry4 | 109,25 | -0,1308 | 0,8763719 | chr18 | - |
| Sra1 | 1357,26 | -0,4248 | 0,092096287 | chr18 | - |
| Srfbp1 | 94,47 | 0,2867 | 0,489795964 | chr18 | + |
| Srp19 | 446,16 | -0,1554 | 0,764714786 | chr18 | + |
| Ss18 | 521,11 | -0,0567 | 0,905274865 | chr18 | - |
| St8sia3 | 8,30 | 0,1092 | 0,907824791 | chr18 | + |
| Stard4 | 1056,03 | 0,0155 | 0,976958093 | chr18 | - |
| Stard6 | 7,08 | 0,5051 | 0,468011283 | chr18 | + |
| Svil | 226,58 | 0,0785 | 0,910916841 | chr18 | + |
| Synpo | 81,77 | 0,3883 | 0,332496065 | chr18 | - |
| Taf4b | 38,42 | -0,4992 | 0,2894527 | chr18 | + |
| Tcerg1 | 187,01 | -0,2987 | 0,43666139 | chr18 | + |
| Tcf4 | 254,61 | 0,0619 | 0,899463828 | chr18 | + |
| Tcof1 | 54,47 | 0,0719 | 0,929418223 | chr18 | - |
| Thoc1 | 445,33 | 0,0306 | 0,960969671 | chr18 | + |
| Ticam2 | 9,12 | 0,4716 | 0,492254415 | chr18 | - |
| Timm21 | 536,31 | 0,2028 | 0,581990132 | chr18 | - |
| Tmco6 | 80,42 | 0,1188 | 0,844249702 | chr18 | + |
| Tmed7 | 2281,04 | 0,2142 | 0,581394499 | chr18 | - |

| | | | | | |
|---|---|---|---|---|---|
| Tmem173 | 62,43 | 0,2674 | 0,565029336 | chr18 | - |
| Tmem241 | 102,71 | 0,6585 | 0,075380094 | chr18 | - |
| Tnfaip8 | 64,19 | 0,2418 | 0,611914561 | chr18 | + |
| Tpgs2 | 89,58 | -0,4130 | 0,285271567 | chr18 | - |
| Trappc8 | 683,97 | -0,2066 | 0,54375285 | chr18 | - |
| Trim36 | 13,26 | -0,2997 | 0,690557571 | chr18 | - |
| Tshz1 | 213,30 | -0,0576 | 0,912811384 | chr18 | - |
| Tslp | 8,60 | 0,2449 | 0,764363417 | chr18 | + |
| Ttc39c | 4304,26 | -0,1487 | 0,790692717 | chr18 | + |
| Tubb6 | 79,00 | 0,2466 | 0,680997554 | chr18 | + |
| Txnl1 | 2026,46 | -0,3878 | 0,209412299 | chr18 | - |
| Txnl4a | 980,48 | -0,1438 | 0,725810792 | chr18 | + |
| Ube2d2a | 1232,74 | -0,3525 | 0,213663916 | chr18 | + |
| Usp14 | 974,46 | -0,1638 | 0,683417125 | chr18 | - |
| Vaultrc5 | 2,36 | -0,6356 | NA | chr18 | + |
| Wdr33 | 529,60 | 0,0583 | 0,88569455 | chr18 | + |
| Wdr36 | 494,50 | 0,0434 | 0,943108135 | chr18 | + |
| Wdr55 | 286,32 | -0,1563 | 0,677398032 | chr18 | + |
| Wdr7 | 353,58 | -0,3372 | 0,234220487 | chr18 | + |
| Yipf5 | 821,51 | -0,0563 | 0,894931973 | chr18 | - |
| Ythdc2 | 124,82 | 0,0704 | 0,912010517 | chr18 | + |
| Zadh2 | 2058,40 | 0,1745 | 0,632136487 | chr18 | + |
| Zbtb7c | 718,54 | 0,1279 | 0,859481826 | chr18 | + |
| Zeb1 | 235,10 | 0,2067 | 0,636447281 | chr18 | + |
| Zfp24 | 441,07 | 0,1288 | 0,720379739 | chr18 | - |
| Zfp236 | 186,94 | 0,0854 | 0,882124635 | chr18 | - |
| Zfp35 | 310,66 | 0,1755 | 0,656102596 | chr18 | + |
| Zfp397 | 260,76 | 0,1019 | 0,83669059 | chr18 | + |
| Zfp407 | 75,51 | 0,5609 | 0,217231457 | chr18 | - |
| Zfp438 | 16,78 | 0,7406 | 0,196626688 | chr18 | - |
| Zfp516 | 69,04 | 0,6921 | 0,070410876 | chr18 | + |
| Zfp521 | 8,23 | 0,2103 | 0,804380135 | chr18 | - |
| Zfp532 | 36,23 | 0,2221 | 0,694473101 | chr18 | + |
| Zmat2 | 1314,65 | -0,0212 | 0,971430847 | chr18 | + |
| 1110059E24Rik | 579,45 | -0,2619 | 0,581990132 | chr19 | - |
| Hectd2os | 1193,39 | -0,1520 | 0,81937394 | chr19 | - |
| 1700018L02Rik | 40,15 | 0,3262 | 0,633994747 | chr19 | + |
| 1810009A15Rik | 4,79 | 0,0311 | 0,9774226 | chr19 | + |
| 1810055G02Rik | 440,15 | -0,6872 | 0,081851577 | chr19 | + |
| Ifit1bl2 | 12,14 | 0,5228 | 0,432778144 | chr19 | - |
| 2010003K11Rik | 349,16 | 0,6281 | 0,156280728 | chr19 | - |
| Borcs7 | 150,91 | -0,2373 | 0,564849438 | chr19 | + |
| Carnmt1 | 315,29 | -0,1749 | 0,66477796 | chr19 | + |
| 2700046G09Rik | 13,36 | 0,7470 | 0,196118975 | chr19 | + |
| 2700081O15Rik | 39,32 | -0,1419 | 0,815176408 | chr19 | + |
| 4430402I18Rik | 14,54 | 0,8092 | 0,170818911 | chr19 | - |
| 4930481A15Rik | 117,82 | 0,0023 | 0,996945533 | chr19 | + |
| Shtn1 | 556,30 | 0,1431 | 0,782398264 | chr19 | - |
| 4930526L06Rik | 4,76 | -0,0992 | 0,916852736 | chr19 | + |
| 4930552P12Rik | 5,14 | 0,8780 | 0,142097482 | chr19 | - |

| | | | | | |
|---|---|---|---|---|---|
| 4931403E22Rik | 2,24 | -0,2475 | NA | chr19 | - |
| 4931408D14Rik | 351,48 | 0,6482 | 0,177510828 | chr19 | - |
| 4933411K16Rik | 15,65 | 0,2329 | 0,771786001 | chr19 | + |
| 5730408K05Rik | 10,77 | -0,7071 | 0,249215983 | chr19 | - |
| Mirt1 | 2,46 | -0,2380 | NA | chr19 | - |
| 9130011E15Rik | 112,37 | 0,2813 | 0,4891753 | chr19 | - |
| 9930021J03Rik | 340,57 | 0,3058 | 0,413323375 | chr19 | - |
| A1cf | 1427,37 | 0,4205 | 0,248800492 | chr19 | + |
| A330032B11Rik | 10,26 | -0,1104 | 0,901472073 | chr19 | + |
| A330040F15Rik | 4,76 | -0,1315 | 0,88569455 | chr19 | - |
| Ccdc186 | 46,02 | 0,7299 | 0,154269521 | chr19 | - |
| A930001C03Rik | 17,25 | -0,1031 | 0,91071941 | chr19 | + |
| A930007I19Rik | 3,57 | -0,6367 | 0,286640164 | chr19 | - |
| Abcc2 | 3627,41 | -0,0176 | 0,975528401 | chr19 | + |
| Abhd17b | 364,36 | -0,5005 | 0,172473519 | chr19 | + |
| Ablim1 | 624,45 | 0,7860 | 0,064499727 | chr19 | - |
| Acsl5 | 9887,32 | 0,3975 | 0,287543614 | chr19 | + |
| Acta2 | 61,31 | 0,8020 | 0,085593171 | chr19 | - |
| Actr1a | 709,64 | -0,1210 | 0,79351129 | chr19 | - |
| Acy3 | 1491,35 | 0,3331 | 0,357703576 | chr19 | + |
| Add3 | 198,37 | -0,1447 | 0,7224367 | chr19 | + |
| Adrb1 | 4,30 | 0,1324 | 0,881159817 | chr19 | + |
| Grk2 | 331,52 | -0,3408 | 0,271418885 | chr19 | - |
| Afap1l2 | 9,28 | 0,4699 | 0,494656683 | chr19 | - |
| Ahnak | 547,88 | 0,4595 | 0,332668588 | chr19 | + |
| Uqcc3 | 455,57 | -0,3540 | 0,296372739 | chr19 | - |
| AI837181 | 149,22 | -0,2000 | 0,704548549 | chr19 | + |
| AI846148 | 30,08 | -0,6718 | 0,223523653 | chr19 | - |
| Aip | 312,76 | -0,2078 | 0,620367088 | chr19 | - |
| Ak3 | 7718,31 | -0,2078 | 0,517480468 | chr19 | - |
| Aldh18a1 | 18,53 | 0,0589 | 0,947294154 | chr19 | - |
| Aldh1a1 | 24469,22 | 0,0169 | 0,981442074 | chr19 | + |
| Aldh1a7 | 4796,62 | -0,2442 | 0,472408228 | chr19 | - |
| Aldh3b1 | 55,11 | 0,1293 | 0,838868977 | chr19 | - |
| Ankrd13d | 2,34 | -0,1040 | NA | chr19 | - |
| Ankrd2 | 3,49 | -0,1819 | 0,818405803 | chr19 | + |
| Anxa1 | 153,62 | 0,4772 | 0,258443121 | chr19 | - |
| Ap5b1 | 12,77 | 0,0707 | 0,943717557 | chr19 | + |
| Apba1 | 2,01 | 0,3739 | NA | chr19 | + |
| Arhgap19 | 69,39 | 0,3822 | 0,444265933 | chr19 | - |
| Arl2 | 57,17 | -0,4227 | 0,340229346 | chr19 | - |
| Arl3 | 246,40 | -0,2265 | 0,629969991 | chr19 | - |
| As3mt | 784,04 | 0,0018 | 0,997464504 | chr19 | + |
| Asah2 | 400,34 | 0,6544 | 0,054141872 | chr19 | - |
| Atad1 | 1744,20 | 0,1812 | 0,677019009 | chr19 | - |
| Atg2a | 357,89 | -0,6162 | 0,098598649 | chr19 | + |
| Atl3 | 2383,48 | 0,5352 | 0,054219322 | chr19 | + |
| Atrnl1 | 442,16 | -0,4755 | 0,110973039 | chr19 | + |
| Avpi1 | 380,47 | -0,2103 | 0,754913678 | chr19 | - |
| AW112010 | 4126,68 | 0,1814 | 0,660583784 | chr19 | - |

| | | | | | |
|---|---|---|---|---|---|
| B230217O12Rik | 16,70 | -0,0288 | 0,9774226 | chr19 | - |
| B3gat3 | 372,43 | -0,4210 | 0,291231135 | chr19 | + |
| Bad | 269,93 | -0,3482 | 0,466451498 | chr19 | + |
| Banf1 | 673,11 | -0,2404 | 0,490047155 | chr19 | - |
| Batf2 | 7,27 | -0,3100 | 0,689526649 | chr19 | + |
| Bbip1 | 438,37 | 0,0669 | 0,887769034 | chr19 | - |
| Bbs1 | 19,20 | 0,6019 | 0,302314196 | chr19 | - |
| BC037704 | 4,52 | -0,1653 | 0,850915997 | chr19 | + |
| Bloc1s2 | 364,42 | -0,1887 | 0,655452587 | chr19 | - |
| Brms1 | 254,89 | -0,3578 | 0,322073711 | chr19 | + |
| Bscl2 | 529,18 | -0,4278 | 0,252617603 | chr19 | + |
| Btaf1 | 407,07 | -0,1043 | 0,797253751 | chr19 | + |
| Btrc | 167,64 | -0,1586 | 0,724641234 | chr19 | + |
| C030016D13Rik | 3,80 | -0,3049 | 0,692167613 | chr19 | + |
| Ric1 | 311,03 | 0,1747 | 0,672218442 | chr19 | + |
| C730002L08Rik | 20,71 | 1,0647 | 0,05009592 | chr19 | - |
| Cabp2 | 5,98 | 0,7214 | 0,253832835 | chr19 | + |
| Cabp4 | 4,15 | 0,2848 | 0,71963262 | chr19 | - |
| Cacul1 | 1104,40 | -0,0780 | 0,837417417 | chr19 | - |
| Calhm2 | 27,41 | -0,1696 | 0,7950804 | chr19 | - |
| Capn1 | 207,72 | -0,0597 | 0,907366343 | chr19 | - |
| Carns1 | 14,81 | -0,6515 | 0,249413774 | chr19 | - |
| Casp7 | 443,08 | -0,0246 | 0,968625947 | chr19 | + |
| Cbwd1 | 175,63 | -0,0013 | 0,997549615 | chr19 | - |
| Ccdc85b | 62,15 | -0,5943 | 0,335528734 | chr19 | - |
| Ccdc86 | 90,27 | -0,3227 | 0,517783511 | chr19 | - |
| Ccdc88b | 17,40 | 0,4237 | 0,515706574 | chr19 | - |
| Ccnj | 49,98 | 0,0656 | 0,925898178 | chr19 | + |
| Ccs | 3809,15 | 0,1611 | 0,618512306 | chr19 | - |
| Cd248 | 8,39 | -0,0277 | 0,979212524 | chr19 | + |
| Cd274 | 44,12 | 0,5208 | 0,291574528 | chr19 | + |
| Cd5 | 1,87 | 0,0815 | NA | chr19 | - |
| Cd6 | 2,19 | 0,1892 | NA | chr19 | - |
| Cdc37l1 | 410,50 | -0,3042 | 0,308448866 | chr19 | + |
| Cdc42bpg | 148,53 | -0,1268 | 0,84175507 | chr19 | + |
| Cdc42ep2 | 51,23 | -0,1011 | 0,867206921 | chr19 | - |
| Cdca5 | 5,36 | -0,4539 | 0,527966188 | chr19 | + |
| Cdk2ap2 | 478,51 | -0,2143 | 0,744313101 | chr19 | + |
| Cep55 | 9,65 | -0,0637 | 0,949529403 | chr19 | + |
| Cep78 | 27,60 | 0,0985 | 0,909774279 | chr19 | - |
| Cfl1 | 3514,60 | -0,1374 | 0,707108599 | chr19 | + |
| Ch25h | 2,38 | 0,3586 | NA | chr19 | - |
| Chka | 250,65 | -0,0935 | 0,87093463 | chr19 | + |
| Chrm1 | 3,74 | -0,0616 | 0,950625589 | chr19 | + |
| Chuk | 2444,32 | -0,3424 | 0,384509329 | chr19 | - |
| Clcf1 | 4,40 | 0,3573 | 0,607182487 | chr19 | + |
| Cnnm1 | 2,18 | -0,3364 | NA | chr19 | + |
| Cnnm2 | 54,39 | -0,3810 | 0,41682625 | chr19 | + |
| Cntf | 8,69 | -0,9337 | 0,115583894 | chr19 | - |
| Col17a1 | 3,78 | 0,6236 | 0,272891337 | chr19 | - |

| | | | | | |
|---|---|---|---|---|---|
| Coro1b | 889,23 | -0,2167 | 0,50822045 | chr19 | + |
| Cox15 | 363,39 | 0,0266 | 0,961928495 | chr19 | - |
| Cox8a | 2664,01 | -0,2356 | 0,776119796 | chr19 | - |
| Cpeb3 | 107,42 | -0,2065 | 0,660639092 | chr19 | - |
| Cpn1 | 3382,07 | -0,0869 | 0,859745168 | chr19 | - |
| Cpsf7 | 353,43 | -0,3565 | 0,283971503 | chr19 | + |
| Cpt1a | 5667,44 | -0,1470 | 0,708319716 | chr19 | + |
| Csf2ra | 37,74 | 0,7209 | 0,120468874 | chr19 | - |
| Cstf2t | 506,92 | 0,5053 | 0,058138256 | chr19 | + |
| Ctsf | 452,95 | -0,4213 | 0,160683257 | chr19 | + |
| Ctsw | 9,26 | 0,6568 | 0,306891735 | chr19 | - |
| Cuedc2 | 211,64 | 0,0133 | 0,985269339 | chr19 | - |
| Cutc | 317,11 | -0,2005 | 0,655452587 | chr19 | + |
| Cwf19l1 | 300,31 | -0,0469 | 0,935309554 | chr19 | - |
| Cyb561a3 | 325,45 | 0,2963 | 0,347246285 | chr19 | + |
| Cyp17a1 | 234,81 | -0,6788 | 0,198314967 | chr19 | - |
| Cyp26a1 | 155,00 | -0,4292 | 0,557507087 | chr19 | + |
| Cyp2c37 | 1801,86 | -0,1913 | 0,822666277 | chr19 | + |
| Cyp2c38 | 747,88 | 0,0228 | 0,964556444 | chr19 | - |
| Cyp2c39 | 1718,77 | 0,3989 | 0,564724076 | chr19 | + |
| Cyp2c40 | 258,80 | -0,3097 | 0,611268902 | chr19 | - |
| Cyp2c50 | 7819,18 | -0,5623 | 0,394890234 | chr19 | + |
| Cyp2c54 | 2995,81 | -0,7146 | 0,152776166 | chr19 | - |
| Cyp2c55 | 242,02 | 0,0934 | 0,919467514 | chr19 | + |
| Cyp2c67 | 8589,36 | 0,4634 | 0,246159206 | chr19 | - |
| Cyp2c68 | 4087,38 | -0,2485 | 0,619691963 | chr19 | - |
| Cyp2c69 | 256,18 | -0,5399 | 0,23305528 | chr19 | - |
| Cyp2c70 | 18062,00 | -0,3841 | 0,128604828 | chr19 | - |
| D030056L22Rik | 134,00 | 0,0716 | 0,890761998 | chr19 | + |
| Pum3 | 390,85 | 0,0477 | 0,93056206 | chr19 | - |
| D330050I16Rik | 9,21 | -0,2391 | 0,764714786 | chr19 | + |
| Dagla | 6,59 | -0,0656 | 0,947819654 | chr19 | - |
| Tkfc | 6206,50 | 0,2886 | 0,317123683 | chr19 | - |
| Dclre1a | 363,53 | -0,1575 | 0,832450834 | chr19 | - |
| Ddb1 | 3641,82 | -0,0395 | 0,925898178 | chr19 | + |
| Dmrt2 | 2,75 | -0,1799 | 0,822888019 | chr19 | + |
| Dnajc4 | 134,88 | -0,1115 | 0,857773796 | chr19 | - |
| Dnmbp | 335,15 | -0,2191 | 0,670594461 | chr19 | - |
| Doc2g | 21,23 | -0,4417 | 0,517431746 | chr19 | + |
| Dock8 | 417,56 | -0,2540 | 0,433006548 | chr19 | + |
| Dpcd | 237,78 | 0,0191 | 0,970419105 | chr19 | + |
| Dpf2 | 310,60 | -0,1099 | 0,784693857 | chr19 | - |
| Dpp3 | 776,33 | 0,0636 | 0,892990026 | chr19 | - |
| Drap1 | 543,30 | -0,2500 | 0,517431746 | chr19 | - |
| Dusp5 | 3,25 | -0,0668 | 0,945274006 | chr19 | + |
| Efemp2 | 45,26 | 0,1763 | 0,781581659 | chr19 | + |
| Ehbp1l1 | 86,54 | 0,3289 | 0,469816518 | chr19 | - |
| Eif1ad | 313,08 | -0,3558 | 0,263617984 | chr19 | + |
| Eif3a | 2115,58 | -0,1916 | 0,526946981 | chr19 | - |
| Eml3 | 231,10 | -0,3190 | 0,357779158 | chr19 | + |

| | | | | | |
|---|---|---|---|---|---|
| Entpd1 | 138,39 | 0,4540 | 0,156728357 | chr19 | + |
| Entpd7 | 36,46 | -0,0120 | 0,988512389 | chr19 | + |
| Erlin1 | 1153,77 | 0,1970 | 0,602175054 | chr19 | - |
| Ermp1 | 876,07 | 0,2688 | 0,477459149 | chr19 | - |
| Esrra | 256,11 | -0,3745 | 0,47680357 | chr19 | - |
| Exoc6 | 362,22 | 0,1698 | 0,633486481 | chr19 | + |
| Exosc1 | 136,24 | -0,3592 | 0,258561012 | chr19 | - |
| Fads2 | 4825,77 | -0,2922 | 0,519599979 | chr19 | - |
| Fam122a | 52,93 | 0,0633 | 0,925382443 | chr19 | - |
| Fam160b1 | 557,97 | -0,1581 | 0,703924654 | chr19 | + |
| Slf2 | 271,37 | -0,0241 | 0,955803452 | chr19 | + |
| Fam189a2 | 14,30 | -0,1690 | 0,844029674 | chr19 | - |
| Fam204a | 98,70 | 0,0081 | 0,992196557 | chr19 | - |
| Fam45a | 460,85 | 0,1189 | 0,749483335 | chr19 | + |
| Fam89b | 112,89 | -0,5257 | 0,146305568 | chr19 | - |
| Fas | 424,63 | 0,2663 | 0,402148509 | chr19 | + |
| Fau | 4127,01 | -0,5127 | 0,302314196 | chr19 | + |
| Fbxl15 | 60,99 | -0,5336 | 0,381800527 | chr19 | + |
| Fbxw4 | 129,50 | -0,3360 | 0,320305617 | chr19 | - |
| Fen1 | 69,65 | 0,2008 | 0,708011891 | chr19 | - |
| Fermt3 | 43,53 | 0,0830 | 0,909834629 | chr19 | - |
| Ffar4 | 5,18 | 1,0244 | 0,072134655 | chr19 | + |
| Fgfbp3 | 3,84 | -0,0168 | 0,987384829 | chr19 | - |
| Fibp | 242,22 | -0,4241 | 0,177510828 | chr19 | + |
| Fkbp2 | 1215,52 | -0,4184 | 0,048483358 | chr19 | - |
| Flrt1 | 6,42 | 0,0755 | 0,940538622 | chr19 | - |
| Fra10ac1 | 239,63 | 0,0611 | 0,920649716 | chr19 | - |
| Frat1 | 40,16 | -0,1415 | 0,842055683 | chr19 | + |
| Frat2 | 39,58 | -0,4640 | 0,421362686 | chr19 | - |
| Frmd8 | 225,89 | -0,1307 | 0,829026521 | chr19 | - |
| Fth1 | 22832,91 | -0,3279 | 0,423995253 | chr19 | + |
| Gal3st3 | 12,43 | -0,0670 | 0,943593159 | chr19 | + |
| Ganab | 3535,51 | -0,1203 | 0,757695155 | chr19 | + |
| Gbf1 | 462,64 | -0,2900 | 0,480124139 | chr19 | + |
| Gcnt1 | 13,05 | 0,1706 | 0,831614489 | chr19 | - |
| Gda | 714,39 | 0,2560 | 0,572666827 | chr19 | - |
| Gfra1 | 2954,44 | -0,3225 | 0,404511819 | chr19 | - |
| Gldc | 1560,80 | 0,1820 | 0,624824844 | chr19 | - |
| Glis3 | 2,67 | 0,7851 | 0,166993075 | chr19 | - |
| Glyat | 8632,22 | -0,3704 | 0,290819569 | chr19 | + |
| Ifit1bl1 | 11,28 | 0,7818 | 0,198288927 | chr19 | - |
| Gm7102 | 35,84 | 0,6713 | 0,131674744 | chr19 | - |
| Gm960 | 3,46 | -0,4084 | 0,565013157 | chr19 | - |
| Gm9895 | 23,97 | -0,0014 | 0,998302437 | chr19 | + |
| Gna14 | 62,56 | -0,4322 | 0,491188481 | chr19 | + |
| Gnaq | 442,46 | 0,1437 | 0,691615955 | chr19 | + |
| Got1 | 5232,95 | 0,2057 | 0,749603956 | chr19 | - |
| Gpam | 1207,03 | 0,4966 | 0,197844837 | chr19 | - |
| Gpr137 | 242,47 | -0,3808 | 0,350199328 | chr19 | - |
| Gsto1 | 1574,09 | -0,4325 | 0,248800492 | chr19 | + |

| | | | | | |
|---|---|---|---|---|---|
| Habp2 | 3239,58 | -0,0662 | 0,852923957 | chr19 | + |
| Hells | 18,12 | -0,0529 | 0,953638766 | chr19 | + |
| Hhex | 866,35 | -0,3514 | 0,451750537 | chr19 | + |
| Hif1an | 490,13 | -0,0191 | 0,966361108 | chr19 | + |
| Hnrnpul2 | 1565,00 | 0,0356 | 0,926353842 | chr19 | + |
| Hoga1 | 2803,75 | 0,0866 | 0,820051729 | chr19 | + |
| Hps1 | 93,34 | 0,3891 | 0,339404075 | chr19 | - |
| Hps6 | 39,13 | -0,1223 | 0,843171378 | chr19 | + |
| Hspa12a | 29,48 | 0,2021 | 0,75616611 | chr19 | - |
| Ide | 1267,80 | 0,3663 | 0,167820864 | chr19 | - |
| Ifit1 | 444,22 | 0,8060 | 0,05010744 | chr19 | + |
| Ifit3 | 398,39 | 0,6708 | 0,083560273 | chr19 | + |
| Ighmbp2 | 43,93 | -0,1547 | 0,813323745 | chr19 | - |
| Incenp | 22,11 | 0,1403 | 0,848381384 | chr19 | - |
| Insl6 | 4,18 | 0,4647 | 0,507785804 | chr19 | - |
| Ints5 | 155,72 | -0,1597 | 0,754913678 | chr19 | + |
| Itprip | 16,42 | -0,5689 | 0,32411521 | chr19 | - |
| Jak2 | 363,15 | -0,0427 | 0,93056206 | chr19 | + |
| Kank1 | 495,35 | 0,0639 | 0,883669918 | chr19 | + |
| Kat5 | 173,59 | 0,1057 | 0,795889374 | chr19 | - |
| Kazald1 | 11,54 | -0,7966 | 0,165257806 | chr19 | + |
| Kdm2a | 643,33 | -0,0243 | 0,952710435 | chr19 | - |
| Keg1 | 6390,57 | -0,4811 | 0,320769368 | chr19 | + |
| Kif11 | 26,42 | -0,0582 | 0,942935841 | chr19 | + |
| Kif20b | 17,86 | 0,6292 | 0,266466163 | chr19 | + |
| Klc2 | 56,88 | -0,3443 | 0,572283367 | chr19 | - |
| Klf9 | 1191,52 | 0,3065 | 0,388800969 | chr19 | + |
| Lcor | 15,73 | 0,6466 | 0,29259208 | chr19 | + |
| Ldb1 | 365,15 | 0,1483 | 0,747850382 | chr19 | - |
| Lipa | 3830,89 | 0,2057 | 0,3995596 | chr19 | - |
| Loxl4 | 28,00 | 0,8160 | 0,099247572 | chr19 | - |
| Lpxn | 49,77 | 0,1756 | 0,781281681 | chr19 | + |
| Lrp5 | 334,49 | 0,2837 | 0,535233812 | chr19 | - |
| Lrrn4cl | 2,53 | 0,0575 | NA | chr19 | + |
| Ltbp3 | 21,08 | -0,2121 | 0,762922832 | chr19 | + |
| Lzts2 | 66,75 | 0,0015 | 0,998203916 | chr19 | + |
| Macrod1 | 792,02 | 0,1241 | 0,749224764 | chr19 | + |
| Malat1 | 3575,12 | -0,5993 | 0,188391337 | chr19 | - |
| Mamdc2 | 48,08 | 0,2285 | 0,687523396 | chr19 | - |
| Map3k11 | 274,56 | -0,4166 | 0,432778144 | chr19 | + |
| Map4k2 | 135,32 | -0,2748 | 0,479578055 | chr19 | + |
| Mark2 | 148,20 | 0,3476 | 0,449778593 | chr19 | - |
| Mbl2 | 11769,67 | -0,2392 | 0,433006548 | chr19 | + |
| Men1 | 198,48 | -0,3039 | 0,432704831 | chr19 | + |
| Mgea5 | 1106,27 | -0,3180 | 0,228941765 | chr19 | - |
| Minpp1 | 536,82 | -0,1553 | 0,628566169 | chr19 | + |
| Mms19 | 328,78 | -0,1917 | 0,621449493 | chr19 | - |
| Morn4 | 10,43 | 0,7044 | 0,249470803 | chr19 | - |
| Mrpl11 | 643,20 | -0,3777 | 0,286306833 | chr19 | + |
| Mrpl16 | 695,84 | -0,0818 | 0,826164292 | chr19 | + |

| | | | | | |
|---|---|---|---|---|---|
| Mrpl21 | 382,98 | -0,3092 | 0,345819394 | chr19 | + |
| Mrpl43 | 336,95 | -0,4682 | 0,283971503 | chr19 | - |
| Mrpl49 | 533,31 | -0,3459 | 0,220525617 | chr19 | - |
| Ms4a1 | 12,02 | 0,0390 | 0,969275786 | chr19 | - |
| Ms4a4b | 24,34 | 0,3080 | 0,623572684 | chr19 | + |
| Ms4a4c | 24,63 | 0,4582 | 0,462469544 | chr19 | + |
| Ms4a4d | 149,94 | 0,5689 | 0,178616571 | chr19 | + |
| Ms4a6b | 54,22 | 0,5205 | 0,358895936 | chr19 | + |
| Ms4a6c | 171,11 | 0,3998 | 0,257994637 | chr19 | + |
| Ms4a6d | 100,01 | 0,3552 | 0,437584445 | chr19 | - |
| Ms4a7 | 32,33 | 0,5848 | 0,304781747 | chr19 | - |
| Ms4a8a | 34,33 | 0,2970 | 0,631758049 | chr19 | - |
| Mta2 | 753,86 | 0,0617 | 0,881086817 | chr19 | + |
| Mus81 | 82,16 | -0,3230 | 0,437402307 | chr19 | - |
| Mxi1 | 359,60 | -0,2111 | 0,571350093 | chr19 | + |
| Myof | 50,11 | 0,7424 | 0,087710179 | chr19 | - |
| Myrf | 271,89 | 0,2441 | 0,52377705 | chr19 | - |
| Naa40 | 121,55 | 0,1024 | 0,844511464 | chr19 | - |
| Ndufb8 | 2425,51 | -0,2640 | 0,614808234 | chr19 | - |
| Ndufs8 | 2249,87 | -0,4061 | 0,178452664 | chr19 | - |
| Ndufv1 | 2656,39 | -0,3832 | 0,205871072 | chr19 | - |
| Neat1 | 2419,75 | 0,3124 | 0,618749555 | chr19 | - |
| Neurl1a | 4,26 | 0,4143 | 0,570187994 | chr19 | + |
| Nfkb2 | 71,10 | 0,2732 | 0,615783353 | chr19 | + |
| Nhlrc2 | 201,01 | 0,0530 | 0,93411896 | chr19 | + |
| Nmrk1 | 712,85 | 0,1424 | 0,829999497 | chr19 | + |
| Noc3l | 116,34 | 0,5005 | 0,135521325 | chr19 | - |
| Nolc1 | 323,83 | -0,2704 | 0,508268404 | chr19 | + |
| Npm3 | 239,51 | -0,0373 | 0,947612608 | chr19 | - |
| Nrxn2 | 6,78 | 0,2442 | 0,767446843 | chr19 | + |
| Nt5c2 | 264,40 | -0,1010 | 0,831614489 | chr19 | - |
| Nudt22 | 134,66 | -0,0934 | 0,872644982 | chr19 | - |
| Nxf1 | 737,66 | -0,1671 | 0,65036605 | chr19 | + |
| Stn1 | 63,13 | 0,3153 | 0,536042479 | chr19 | - |
| Osbp | 3469,52 | -0,1386 | 0,708319716 | chr19 | + |
| Ostf1 | 549,36 | -0,0606 | 0,89879469 | chr19 | - |
| Otub1 | 548,84 | -0,3748 | 0,211542715 | chr19 | - |
| Pacs1 | 35,97 | 0,1590 | 0,794116376 | chr19 | - |
| Pank1 | 459,60 | -0,0284 | 0,951806422 | chr19 | - |
| Patl1 | 190,75 | -0,1009 | 0,816750599 | chr19 | + |
| Pcgf5 | 96,20 | -0,3536 | 0,374589794 | chr19 | + |
| Pcgf6 | 212,30 | -0,4173 | 0,214673153 | chr19 | - |
| Pcnx3 | 211,33 | -0,2013 | 0,734981618 | chr19 | - |
| Pcsk5 | 96,14 | -0,0236 | 0,978158168 | chr19 | - |
| Pcx | 5457,66 | -0,3253 | 0,27946684 | chr19 | + |
| Pdcd11 | 262,71 | 0,1617 | 0,712500177 | chr19 | + |
| Pdcd4 | 1904,97 | -0,0688 | 0,836045144 | chr19 | + |
| Pde6c | 15,56 | 0,2944 | 0,710796708 | chr19 | + |
| Pdlim1 | 440,09 | 0,3640 | 0,310109812 | chr19 | - |
| Pdzd8 | 535,11 | -0,3634 | 0,230748575 | chr19 | - |

| | | | | | |
|---|---|---|---|---|---|
| Peli3 | 4,92 | -0,2316 | 0,782080553 | chr19 | - |
| Pgam1 | 2846,82 | -0,1684 | 0,594322944 | chr19 | + |
| Pgm5 | 28,98 | -0,3841 | 0,515706574 | chr19 | - |
| Pi4k2a | 534,83 | -0,1120 | 0,770796088 | chr19 | + |
| Pik3ap1 | 322,29 | 0,1582 | 0,794944296 | chr19 | - |
| Pitpnm1 | 40,82 | -0,1401 | 0,860107955 | chr19 | + |
| Pla2g16 | 319,61 | 0,3684 | 0,228828011 | chr19 | + |
| Plcb3 | 161,21 | 0,0258 | 0,963027434 | chr19 | - |
| Plce1 | 160,77 | 0,0835 | 0,857478182 | chr19 | + |
| Plekhs1 | 2,94 | -0,0569 | 0,951934337 | chr19 | + |
| Pola2 | 94,95 | 0,0068 | 0,991447068 | chr19 | - |
| Pold4 | 372,94 | 0,3952 | 0,237903839 | chr19 | + |
| Poll | 84,30 | 0,0633 | 0,925508783 | chr19 | - |
| Polr2g | 477,47 | 0,0180 | 0,979713225 | chr19 | - |
| Plpp6 | 167,68 | 0,4040 | 0,37274915 | chr19 | + |
| Ppp1ca | 1845,70 | -0,0401 | 0,941757244 | chr19 | + |
| Ppp1r14b | 156,94 | -0,5110 | 0,209630049 | chr19 | + |
| Ppp6r3 | 942,21 | 0,1066 | 0,736302391 | chr19 | - |
| Pprc1 | 111,14 | -0,5423 | 0,220022322 | chr19 | + |
| Prdx3 | 3435,53 | -0,0439 | 0,918077233 | chr19 | - |
| Prdx5 | 4536,52 | -0,2163 | 0,670139713 | chr19 | - |
| Prkg1 | 28,08 | 0,3603 | 0,539569536 | chr19 | - |
| Prpf19 | 1813,83 | -0,1649 | 0,563763821 | chr19 | + |
| Prune2 | 9,76 | 0,2476 | 0,755670612 | chr19 | + |
| Ptar1 | 20,03 | -0,0918 | 0,912399044 | chr19 | + |
| Pten | 2868,46 | -0,4111 | 0,118025078 | chr19 | + |
| Ptprcap | 7,61 | -0,3604 | 0,632136487 | chr19 | + |
| Pygm | 11,70 | 0,5536 | 0,380049722 | chr19 | + |
| Pyroxd2 | 404,75 | 0,0094 | 0,988512389 | chr19 | - |
| R3hcc1l | 283,19 | -0,0986 | 0,81045863 | chr19 | + |
| Rab11fip2 | 121,64 | 0,4616 | 0,277366051 | chr19 | - |
| Rab1b | 1950,86 | -0,0495 | 0,9163988 | chr19 | - |
| Rab3il1 | 73,64 | 0,3217 | 0,428625654 | chr19 | + |
| Rad9a | 36,11 | 0,2182 | 0,732933983 | chr19 | - |
| Ranbp6 | 410,97 | 0,3598 | 0,28160417 | chr19 | - |
| Rasgrp2 | 151,82 | 0,0482 | 0,931322892 | chr19 | + |
| Rbm14 | 127,74 | -0,1611 | 0,790473751 | chr19 | - |
| Rbm20 | 52,94 | 0,6366 | 0,123183417 | chr19 | + |
| Rbm4 | 8,79 | -0,4296 | 0,54877155 | chr19 | - |
| Rbm4b | 138,59 | 0,1549 | 0,733541536 | chr19 | + |
| Rce1 | 92,38 | -0,1370 | 0,803874753 | chr19 | - |
| Rela | 394,75 | -0,3459 | 0,351934331 | chr19 | + |
| Rfk | 896,60 | -0,0092 | 0,984470281 | chr19 | + |
| Rfx3 | 62,20 | 0,8218 | 0,050606293 | chr19 | - |
| Rhod | 396,35 | -0,5504 | 0,277366051 | chr19 | - |
| Rin1 | 3,57 | 0,3705 | 0,61720787 | chr19 | + |
| Rnaseh2c | 340,19 | -0,6367 | 0,327028954 | chr19 | + |
| Rnls | 17,85 | 0,9121 | 0,081609555 | chr19 | - |
| Rom1 | 35,57 | -0,4394 | 0,394187455 | chr19 | - |
| Rorb | 4,80 | 0,1047 | 0,91071941 | chr19 | - |

| | | | | | |
|---|---|---|---|---|---|
| Rpp30 | 162,08 | 0,0840 | 0,873853413 | chr19 | + |
| Rps6ka4 | 79,64 | 0,0794 | 0,89879469 | chr19 | - |
| Rps6kb2 | 135,61 | -0,5382 | 0,170839779 | chr19 | - |
| Rrp12 | 69,22 | -0,3932 | 0,397488617 | chr19 | - |
| Rtn3 | 1881,69 | -0,1109 | 0,724652712 | chr19 | - |
| Sac3d1 | 52,01 | -0,4179 | 0,462033601 | chr19 | - |
| Sart1 | 276,91 | -0,4111 | 0,298176352 | chr19 | - |
| Scd1 | 133661,47 | -0,0036 | 0,996644671 | chr19 | - |
| Scd2 | 90,59 | 0,2990 | 0,626400494 | chr19 | + |
| Scd3 | 11,72 | 0,4357 | 0,513917577 | chr19 | + |
| Scd4 | 3,35 | 0,2974 | 0,695978671 | chr19 | + |
| Scyl1 | 435,39 | -0,5369 | 0,091510375 | chr19 | - |
| Sdhaf2 | 676,71 | -0,1526 | 0,649305665 | chr19 | - |
| Sec31b | 55,61 | 0,0191 | 0,981931319 | chr19 | - |
| Sema4g | 3067,82 | -0,2597 | 0,501991823 | chr19 | + |
| Sf1 | 715,58 | -0,2565 | 0,536622488 | chr19 | + |
| Sf3b2 | 1183,26 | 0,0735 | 0,860078935 | chr19 | - |
| Sfr1 | 745,17 | -0,3341 | 0,237935228 | chr19 | + |
| Sfxn2 | 626,60 | -0,0289 | 0,952576171 | chr19 | + |
| Sfxn3 | 37,27 | 0,1898 | 0,744618461 | chr19 | + |
| Sfxn4 | 22,63 | 0,0361 | 0,968726083 | chr19 | - |
| Sgms1 | 156,36 | -0,0939 | 0,873682657 | chr19 | - |
| Sh3pxd2a | 263,82 | -0,0036 | 0,995735297 | chr19 | - |
| Shoc2 | 534,69 | 0,1117 | 0,782080553 | chr19 | + |
| Sipa1 | 89,76 | -0,2417 | 0,647914259 | chr19 | - |
| Slc16a12 | 1431,29 | -0,1259 | 0,820554805 | chr19 | - |
| Slc18a2 | 6,77 | 0,0987 | 0,917470917 | chr19 | + |
| Slc1a1 | 4,40 | 0,9464 | 0,080271426 | chr19 | + |
| Slc22a29 | 4,47 | -0,1399 | 0,87623846 | chr19 | - |
| Slc25a45 | 471,84 | -0,5386 | 0,241312279 | chr19 | + |
| Slc29a2 | 9,82 | -0,6414 | 0,319674179 | chr19 | + |
| Slc3a2 | 518,92 | -0,2329 | 0,537935014 | chr19 | - |
| Slk | 714,26 | 0,1309 | 0,711248152 | chr19 | + |
| Smarca2 | 1513,27 | -0,1740 | 0,589677579 | chr19 | + |
| Smc5 | 343,96 | 0,1970 | 0,564724076 | chr19 | - |
| Smndc1 | 274,05 | -0,0580 | 0,907233333 | chr19 | - |
| Snhg1 | 334,50 | -0,5065 | 0,255722726 | chr19 | + |
| Snord22 | 17,18 | -0,6497 | 0,267137829 | chr19 | + |
| Snx15 | 127,55 | -0,3516 | 0,396411176 | chr19 | - |
| Snx32 | 33,32 | -0,6970 | 0,137645772 | chr19 | - |
| Sorbs1 | 702,45 | 0,2020 | 0,60845063 | chr19 | - |
| Sptbn2 | 1061,74 | 0,1341 | 0,769969083 | chr19 | + |
| Ssh3 | 96,81 | -0,2452 | 0,6841486 | chr19 | - |
| Sssca1 | 91,85 | -0,1881 | 0,782474411 | chr19 | - |
| Stip1 | 1117,16 | 0,0805 | 0,871802627 | chr19 | - |
| Stx3 | 27,98 | 0,0129 | 0,988512389 | chr19 | - |
| Stxbp3-ps | 3,29 | -0,2007 | 0,806401686 | chr19 | - |
| Sufu | 45,45 | -0,0694 | 0,915071511 | chr19 | + |
| Kmt5b | 324,58 | -0,1353 | 0,691169933 | chr19 | + |
| Syt12 | 21,58 | 0,6388 | 0,29233966 | chr19 | - |

| | | | | | |
|---|---|---|---|---|---|
| Syt7 | 11,25 | 0,9158 | 0,117722866 | chr19 | + |
| Syvn1 | 613,51 | -0,1621 | 0,856076562 | chr19 | + |
| Taf5 | 70,39 | -0,0759 | 0,89879469 | chr19 | + |
| Taf6l | 32,24 | 0,0122 | 0,988414083 | chr19 | - |
| Tbc1d10c | 6,41 | 0,5439 | 0,423813652 | chr19 | - |
| Tbc1d12 | 137,06 | -0,0919 | 0,876803328 | chr19 | + |
| Tcf7l2 | 227,73 | -0,5171 | 0,06162809 | chr19 | + |
| Tcirg1 | 512,15 | -0,3722 | 0,319080738 | chr19 | - |
| Tctn3 | 18,07 | 0,3220 | 0,653286041 | chr19 | - |
| Tigd3 | 5,27 | -0,3600 | 0,634623679 | chr19 | - |
| Tjp2 | 160,74 | 0,2228 | 0,598404146 | chr19 | - |
| Tle4 | 185,21 | -0,0789 | 0,885099576 | chr19 | - |
| Tm9sf3 | 3803,27 | -0,0905 | 0,830330662 | chr19 | - |
| Tmem109 | 282,35 | -0,2781 | 0,433006548 | chr19 | - |
| Tmem134 | 474,54 | -0,4556 | 0,217231457 | chr19 | + |
| Tmem138 | 22,54 | -0,0372 | 0,967498977 | chr19 | - |
| Tmem179b | 276,54 | 0,0044 | 0,993991445 | chr19 | - |
| Mfsd13a | 33,47 | 0,1646 | 0,830987741 | chr19 | + |
| Tmem2 | 222,10 | 0,1464 | 0,691018693 | chr19 | + |
| Tmem216 | 91,10 | -0,2625 | 0,598404146 | chr19 | - |
| Tmem223 | 395,31 | -0,6733 | 0,074317353 | chr19 | + |
| Tmem258 | 29,79 | 0,0782 | 0,92028484 | chr19 | + |
| Tnks2 | 1034,83 | 0,0196 | 0,968565076 | chr19 | + |
| Trim8 | 251,97 | -0,3967 | 0,328388243 | chr19 | + |
| Trmt112 | 438,38 | -0,3809 | 0,420444431 | chr19 | + |
| Trpm6 | 6,09 | 0,3986 | 0,585752237 | chr19 | + |
| Trpt1 | 28,07 | -0,0488 | 0,952541585 | chr19 | + |
| Trub1 | 194,72 | -0,1067 | 0,802632263 | chr19 | + |
| Ttc9c | 463,63 | -0,0441 | 0,930202709 | chr19 | - |
| Tut1 | 115,65 | -0,2146 | 0,624676895 | chr19 | + |
| Ubtd1 | 77,25 | -0,6448 | 0,21455748 | chr19 | + |
| Ubxn1 | 1142,18 | -0,5910 | 0,106945732 | chr19 | + |
| Uhrf2 | 494,24 | -0,0559 | 0,884858966 | chr19 | + |
| Unc93b1 | 436,39 | -0,0226 | 0,972880126 | chr19 | + |
| Usmg5 | 4971,16 | -0,4188 | 0,173590837 | chr19 | - |
| Vegfb | 93,12 | -0,7520 | 0,078201192 | chr19 | - |
| Vldlr | 31,73 | -0,2253 | 0,761732176 | chr19 | + |
| Vps13a | 332,29 | -0,2023 | 0,566893799 | chr19 | - |
| Vps37c | 111,12 | 0,0119 | 0,986691613 | chr19 | + |
| Vps51 | 124,73 | -0,1920 | 0,743472775 | chr19 | - |
| Vti1a | 142,38 | 0,2544 | 0,564724076 | chr19 | + |
| Vwce | 336,83 | -0,0167 | 0,980015446 | chr19 | + |
| Wbp1l | 2754,05 | -0,4087 | 0,253278995 | chr19 | + |
| Wdr74 | 304,88 | -0,4227 | 0,138749559 | chr19 | + |
| Cfap43 | 33,06 | 0,3468 | 0,442222987 | chr19 | - |
| Xpnpep1 | 433,06 | -0,1404 | 0,680920887 | chr19 | - |
| Yif1a | 715,36 | 0,0959 | 0,827995984 | chr19 | + |
| Zbtb3 | 5,41 | -0,1826 | 0,834675238 | chr19 | + |
| Zdhhc16 | 235,49 | -0,4591 | 0,09512379 | chr19 | + |
| Zdhhc24 | 39,89 | -0,4160 | 0,384370645 | chr19 | + |

| | | | | | |
|---|---|---|---|---|---|
| Zdhhc6 | 1087,70 | -0,4547 | 0,058961472 | chr19 | - |
| Zfand5 | 740,17 | -0,2821 | 0,345583636 | chr19 | + |
| Zfp91 | 1717,10 | -0,2928 | 0,394399177 | chr19 | - |
| Zfpl1 | 111,02 | -0,1787 | 0,738282454 | chr19 | - |
| Zfyve27 | 239,82 | -0,2090 | 0,574727653 | chr19 | + |
| Znhit2 | 83,33 | -0,5881 | 0,287543614 | chr19 | + |
| 1110012L19Rik | 423,84 | -0,0398 | 0,943108135 | chrX | + |
| 1810030O07Rik | 127,40 | 0,0006 | 0,998789109 | chrX | - |
| 2010204K13Rik | 9,90 | 1,0420 | 0,057660447 | chrX | - |
| 2010308F09Rik | 5,15 | 0,6241 | 0,342654396 | chrX | - |
| Nbdy | 49,76 | 0,3770 | 0,407597046 | chrX | + |
| 2610002M06Rik | 439,75 | 0,3067 | 0,345598364 | chrX | - |
| Stk26 | 3,87 | -0,0072 | 0,994483849 | chrX | + |
| 2810403D21Rik | 26,53 | 0,2829 | 0,701104843 | chrX | + |
| Bcor | 16,35 | 0,1376 | 0,875746336 | chrX | - |
| Kantr | 41,14 | 0,3650 | 0,474644284 | chrX | - |
| Cldn34c1 | 8,56 | -0,0130 | 0,990189789 | chrX | + |
| 3830403N18Rik | 4,53 | 0,0178 | 0,986661276 | chrX | + |
| 4930513O06Rik | 4,02 | -0,2722 | 0,724652712 | chrX | + |
| 4933407K13Rik | 11,29 | 0,4724 | 0,48486694 | chrX | - |
| 5430427O19Rik | 11,68 | -0,1479 | 0,869067375 | chrX | + |
| 5530601H04Rik | 75,42 | 0,5433 | 0,194733334 | chrX | - |
| 5730405O15Rik | 5,39 | 0,3650 | 0,629719515 | chrX | + |
| 6030498E09Rik | 3,46 | 0,3976 | 0,57626953 | chrX | + |
| Smim10l2a | 2,31 | -0,1638 | NA | chrX | + |
| A230072C01Rik | 99,11 | -0,5222 | 0,10473347 | chrX | + |
| A230072E10Rik | 2,63 | 0,1924 | 0,809792004 | chrX | - |
| P2ry10b | 13,26 | 0,6982 | 0,235271103 | chrX | + |
| Bclaf3 | 41,38 | 0,4306 | 0,41633338 | chrX | + |
| AA414768 | 3,34 | 0,4071 | 0,565013157 | chrX | + |
| Abcb7 | 1077,04 | 0,3623 | 0,250533705 | chrX | - |
| Abcd1 | 190,99 | -0,0392 | 0,944925159 | chrX | + |
| Ace2 | 18,92 | -0,1748 | 0,811058993 | chrX | + |
| Acot9 | 111,92 | -0,0085 | 0,988512389 | chrX | + |
| Acsl4 | 1518,67 | -0,2638 | 0,522469399 | chrX | - |
| Aifm1 | 2518,29 | -0,0931 | 0,786009002 | chrX | - |
| Akap17b | 91,95 | 0,0598 | 0,920978973 | chrX | - |
| Alg13 | 114,23 | 0,0380 | 0,948314012 | chrX | + |
| Amer1 | 106,73 | 0,0490 | 0,928423971 | chrX | - |
| Ammecr1 | 9,45 | 0,4450 | 0,52377705 | chrX | - |
| Amot | 220,09 | 0,6017 | 0,058961472 | chrX | - |
| Ap1s2 | 68,27 | 0,0507 | 0,937853416 | chrX | + |
| Apex2 | 94,91 | -0,2596 | 0,581394499 | chrX | - |
| Apoo | 257,27 | -0,4000 | 0,277366051 | chrX | + |
| Apool | 866,96 | -0,0137 | 0,979212524 | chrX | + |
| Araf | 779,03 | -0,0128 | 0,976304651 | chrX | + |
| Arhgap4 | 70,07 | 0,1215 | 0,854809325 | chrX | - |
| Arhgap6 | 75,79 | 0,0739 | 0,915621354 | chrX | + |
| Arhgef6 | 65,26 | 0,1737 | 0,770543183 | chrX | - |
| Armcx1 | 54,30 | 0,3766 | 0,434177785 | chrX | + |

| | | | | | |
|---|---|---|---|---|---|
| Armcx2 | 48,24 | 0,7779 | 0,084139273 | chrX | - |
| Armcx3 | 281,32 | 0,2743 | 0,452803479 | chrX | + |
| Armcx4 | 61,06 | 0,5252 | 0,235228316 | chrX | + |
| Armcx5 | 24,42 | 0,4772 | 0,404511819 | chrX | + |
| Armcx6 | 19,66 | 0,0022 | 0,997549615 | chrX | - |
| Atg4a | 352,81 | -0,2192 | 0,572115465 | chrX | + |
| Atp11c | 4129,34 | -0,0454 | 0,938006162 | chrX | - |
| Atp6ap1 | 1462,53 | -0,1564 | 0,638379056 | chrX | + |
| Atp6ap2 | 988,96 | -0,2222 | 0,469528266 | chrX | + |
| Atp7a | 106,72 | 0,2719 | 0,59810474 | chrX | + |
| Atrx | 768,23 | 0,4436 | 0,186556074 | chrX | - |
| Awat2 | 44,26 | 0,7843 | 0,089779423 | chrX | - |
| B630019K06Rik | 11,22 | -0,8361 | 0,154435937 | chrX | + |
| Tmem185a | 264,13 | -0,0145 | 0,980154061 | chrX | - |
| BC065397 | 16,32 | 0,0673 | 0,940263679 | chrX | + |
| Bcap31 | 3417,67 | -0,1500 | 0,618700706 | chrX | - |
| Bcor | 90,42 | 0,0818 | 0,895594598 | chrX | - |
| Bcorl1 | 58,35 | -0,0920 | 0,892875227 | chrX | + |
| Bex2 | 3,56 | 0,4881 | 0,472359592 | chrX | - |
| Bex4 | 13,29 | 0,7615 | 0,207617402 | chrX | + |
| Bgn | 1122,44 | 0,2365 | 0,503596333 | chrX | + |
| Bhlhb9 | 108,04 | 0,6990 | 0,110056031 | chrX | + |
| Bmx | 3,72 | -0,2783 | 0,720531276 | chrX | - |
| Brcc3 | 775,34 | -0,0459 | 0,918104715 | chrX | + |
| Btk | 30,79 | 0,2735 | 0,645264604 | chrX | - |
| C1galt1c1 | 635,04 | -0,2269 | 0,552507143 | chrX | - |
| C330007P06Rik | 721,86 | 0,2067 | 0,60173351 | chrX | - |
| C430049B03Rik | 5,69 | -0,4821 | 0,495988702 | chrX | - |
| Cacna1f | 4,27 | -0,4058 | 0,570278616 | chrX | + |
| Car5b | 64,79 | -0,5756 | 0,157433776 | chrX | - |
| Cask | 248,85 | -0,3900 | 0,217231457 | chrX | - |
| Ccdc160 | 5,51 | 0,0952 | 0,920978973 | chrX | + |
| Ccdc22 | 214,87 | -0,4868 | 0,087710179 | chrX | - |
| Cd99l2 | 556,13 | -0,1519 | 0,617391963 | chrX | - |
| Cdk16 | 243,69 | -0,3881 | 0,267344282 | chrX | + |
| Cdkl5 | 17,39 | 0,5989 | 0,336398046 | chrX | - |
| Cdx4 | 2,10 | 0,7849 | NA | chrX | + |
| Cenpi | 5,54 | 0,5509 | 0,41633338 | chrX | + |
| Cetn2 | 409,63 | 0,0296 | 0,950795699 | chrX | - |
| Cfp | 476,36 | 0,0669 | 0,878081757 | chrX | - |
| Chic1 | 183,14 | 0,7397 | 0,088510192 | chrX | + |
| Chm | 271,91 | 0,4582 | 0,22361789 | chrX | - |
| Chrdl1 | 8,56 | 0,6333 | 0,327202757 | chrX | - |
| Chst7 | 10,22 | 0,1990 | 0,807278226 | chrX | + |
| Clcn5 | 14,29 | 0,3679 | 0,587223505 | chrX | - |
| Cldn2 | 1710,80 | 0,5140 | 0,22384344 | chrX | + |
| Col4a5 | 38,01 | 0,3148 | 0,627875496 | chrX | + |
| Col4a6 | 2,51 | -0,1930 | NA | chrX | - |
| Cox7b | 8626,47 | -0,3508 | 0,321487973 | chrX | + |
| Cstf2 | 221,85 | 0,1867 | 0,621693318 | chrX | + |

| | | | | | |
|---|---|---|---|---|---|
| Cul4b | 758,69 | -0,1362 | 0,743138956 | chrX | - |
| Cxcr3 | 6,41 | 0,8552 | 0,158793576 | chrX | - |
| Rtl8a | 5,68 | 0,3297 | 0,670684766 | chrX | - |
| Rtl8b | 14,23 | -0,0217 | 0,98429278 | chrX | - |
| Rtl8c | 20,49 | -0,3073 | 0,627992485 | chrX | + |
| Cybb | 337,82 | 0,5546 | 0,112715106 | chrX | - |
| Cysltr1 | 21,31 | 0,7661 | 0,161077273 | chrX | - |
| Dcx | 9,40 | 0,6185 | 0,340033388 | chrX | - |
| Ints6l | 60,11 | 0,1190 | 0,865720762 | chrX | + |
| Ddx3x | 4586,48 | 0,2160 | 0,602938667 | chrX | + |
| Diaph2 | 287,73 | -0,0348 | 0,948200011 | chrX | + |
| Dkc1 | 373,25 | -0,0128 | 0,983780811 | chrX | + |
| Dlg3 | 233,98 | -0,1469 | 0,723157997 | chrX | + |
| Dnase1l1 | 115,83 | -0,0283 | 0,95990578 | chrX | - |
| Dock11 | 30,63 | 0,4464 | 0,404511819 | chrX | + |
| Dusp9 | 3,12 | 0,7838 | 0,186292552 | chrX | + |
| Dynlt3 | 1778,88 | 0,0247 | 0,964556444 | chrX | - |
| Eda | 15,38 | -0,2565 | 0,713796287 | chrX | + |
| Eda2r | 10,63 | 0,7531 | 0,215958978 | chrX | - |
| Efnb1 | 103,88 | 0,1307 | 0,820577263 | chrX | + |
| Eif1ax | 532,04 | -0,1058 | 0,847419446 | chrX | + |
| Eif2s3x | 2353,50 | -0,0093 | 0,986818215 | chrX | - |
| Elf4 | 33,84 | 0,1279 | 0,856738348 | chrX | - |
| Elk1 | 91,93 | 0,2136 | 0,691108142 | chrX | - |
| Emd | 242,08 | -0,1751 | 0,631914678 | chrX | + |
| Enox2 | 258,85 | 0,2239 | 0,6215219 | chrX | - |
| Ercc6l | 9,32 | 0,5420 | 0,411104454 | chrX | - |
| F630028O10Rik | 31,35 | 0,5628 | 0,252093318 | chrX | + |
| F8 | 574,54 | 0,3842 | 0,233364856 | chrX | - |
| F8a | 60,34 | -0,0070 | 0,992812238 | chrX | + |
| F9 | 6895,99 | 0,2578 | 0,455467885 | chrX | + |
| Fam120c | 111,02 | 0,3319 | 0,482925724 | chrX | + |
| Fam122b | 37,39 | 0,3638 | 0,522513339 | chrX | - |
| Fam199x | 165,27 | -0,0128 | 0,984995389 | chrX | + |
| Fam3a | 200,48 | -0,1989 | 0,63353315 | chrX | - |
| Fam50a | 291,32 | -0,4259 | 0,215046281 | chrX | + |
| Fancb | 14,53 | -0,4072 | 0,53237472 | chrX | + |
| Fgd1 | 14,51 | 0,2202 | 0,765858489 | chrX | + |
| Fhl1 | 115,01 | 0,4031 | 0,273713146 | chrX | + |
| Vegfd | 17,09 | -0,0756 | 0,932525817 | chrX | + |
| Firre | 276,23 | -0,1461 | 0,823578308 | chrX | - |
| Flna | 360,61 | 0,4054 | 0,169113943 | chrX | - |
| Fmr1 | 387,00 | 0,3866 | 0,217231457 | chrX | + |
| Foxo4 | 552,72 | 0,0709 | 0,871700795 | chrX | + |
| Ftsj1 | 410,00 | -0,2503 | 0,537289795 | chrX | - |
| Ftx | 32,74 | 0,2926 | 0,593645254 | chrX | - |
| Fundc1 | 226,17 | 0,3612 | 0,318767824 | chrX | - |
| Fundc2 | 255,48 | 0,4170 | 0,339404199 | chrX | + |
| G530011O06Rik | 139,45 | -0,0750 | 0,907233333 | chrX | - |
| Gab3 | 7,07 | 0,2500 | 0,760248244 | chrX | - |

| | | | | | |
|---|---|---|---|---|---|
| Gabra3 | 38,66 | 0,3047 | 0,631200781 | chrX | - |
| Gdi1 | 965,47 | 0,0009 | 0,998195551 | chrX | + |
| Gemin8 | 28,78 | 0,3133 | 0,581157554 | chrX | + |
| Gjb1 | 9069,61 | -0,0706 | 0,853450823 | chrX | + |
| Gla | 155,34 | 0,2962 | 0,422668474 | chrX | - |
| Glod5 | 3,87 | 0,8566 | 0,140278812 | chrX | - |
| Gm6377 | 29,91 | 0,8734 | 0,077224204 | chrX | + |
| Gm6787 | 6,11 | 0,3951 | 0,591108504 | chrX | + |
| Gm9159 | 14,79 | -0,0311 | 0,974809304 | chrX | + |
| Gnl3l | 258,21 | 0,2408 | 0,458689654 | chrX | - |
| Gpc3 | 22,59 | 0,2337 | 0,750672503 | chrX | - |
| Gpc4 | 632,49 | -0,0388 | 0,941757244 | chrX | - |
| Gpkow | 350,37 | 0,1650 | 0,699742099 | chrX | + |
| Gpm6b | 28,81 | 0,4991 | 0,430392697 | chrX | + |
| Gpr174 | 4,40 | 0,2356 | 0,77551946 | chrX | + |
| Gpr34 | 4,24 | 0,7021 | 0,258561012 | chrX | + |
| Gprasp1 | 91,24 | 0,5548 | 0,208689106 | chrX | + |
| Gprasp2 | 3,76 | 0,1729 | 0,840953217 | chrX | + |
| Gria3 | 18,00 | -1,0004 | 0,054188122 | chrX | + |
| Gripap1 | 281,63 | -0,0139 | 0,979212524 | chrX | + |
| Gspt2 | 7,54 | 0,8604 | 0,15398845 | chrX | + |
| Gk | 2014,72 | 0,2218 | 0,491229985 | chrX | - |
| Haus7 | 250,54 | 0,1058 | 0,830330662 | chrX | - |
| Hccs | 217,70 | -0,2059 | 0,693242675 | chrX | - |
| Hcfc1 | 328,70 | -0,0101 | 0,98507181 | chrX | - |
| Hdac6 | 174,94 | -0,2770 | 0,451688988 | chrX | - |
| Hdac8 | 103,45 | 0,5813 | 0,083804189 | chrX | - |
| Heph | 43,65 | 0,3302 | 0,525233119 | chrX | + |
| Hmgb3 | 326,52 | -0,0635 | 0,920482551 | chrX | + |
| Hmgn5 | 1283,01 | 0,0818 | 0,842502441 | chrX | - |
| Hnrnph2 | 1386,87 | 0,2294 | 0,528512451 | chrX | + |
| Hprt | 1427,80 | -0,2995 | 0,312960746 | chrX | + |
| Hsd17b10 | 3007,74 | -0,2471 | 0,380021816 | chrX | + |
| Htatsf1 | 486,99 | -0,0834 | 0,852405207 | chrX | + |
| Huwe1 | 1626,87 | -0,2318 | 0,462039583 | chrX | + |
| Idh3g | 1159,14 | -0,2014 | 0,471291736 | chrX | - |
| Ids | 141,11 | 0,2636 | 0,549694981 | chrX | - |
| Igbp1 | 1375,35 | -0,2408 | 0,47051491 | chrX | + |
| Igsf1 | 3,09 | -0,8624 | 0,136585635 | chrX | - |
| Ikbkg | 1279,52 | -0,1416 | 0,691615955 | chrX | + |
| Il13ra1 | 1084,92 | 0,1452 | 0,7950804 | chrX | + |
| Il13ra2 | 17,48 | 0,4280 | 0,53490582 | chrX | - |
| Il2rg | 57,69 | 0,2408 | 0,682099278 | chrX | - |
| Iqsec2 | 67,76 | -0,0081 | 0,991035244 | chrX | + |
| Itm2a | 31,05 | -0,3878 | 0,515706574 | chrX | - |
| Jade3 | 77,84 | -0,0220 | 0,975879219 | chrX | + |
| Jpx | 41,42 | 0,2388 | 0,662842062 | chrX | + |
| Kctd12b | 91,44 | 0,5542 | 0,215046281 | chrX | - |
| Kdm5c | 667,42 | -0,0812 | 0,849424132 | chrX | + |
| Kdm6a | 242,73 | 0,2255 | 0,577347444 | chrX | + |

| | | | | | |
|---|---|---|---|---|---|
| Kif4 | 11,87 | 0,2055 | 0,807278226 | chrX | + |
| Klhl15 | 16,47 | -0,3276 | 0,63367203 | chrX | + |
| Klhl4 | 20,15 | 0,4600 | 0,475341278 | chrX | + |
| Lage3 | 470,25 | -0,0536 | 0,905147397 | chrX | - |
| Lamp2 | 16930,28 | 0,0275 | 0,955234619 | chrX | - |
| Las1l | 817,21 | -0,0013 | 0,997464504 | chrX | - |
| Lonrf3 | 127,75 | -0,2572 | 0,585791602 | chrX | + |
| Mageb16-ps1 | 11,43 | -0,1945 | 0,811093035 | chrX | - |
| Maged1 | 1312,60 | -0,0424 | 0,935880676 | chrX | - |
| Maged2 | 122,33 | 0,2383 | 0,544991268 | chrX | - |
| Magee1 | 35,30 | -0,0241 | 0,976528475 | chrX | + |
| Mageh1 | 65,54 | 0,6740 | 0,075163457 | chrX | - |
| Magix | 164,71 | 0,1210 | 0,820051729 | chrX | - |
| Magt1 | 1792,70 | 0,1543 | 0,711258571 | chrX | - |
| Mamld1 | 3,98 | 0,2951 | 0,69945599 | chrX | + |
| Maoa | 2,96 | 0,2669 | 0,717188441 | chrX | + |
| Maob | 4341,13 | 0,3028 | 0,317453963 | chrX | - |
| Map3k15 | 8,99 | -0,5676 | 0,39354827 | chrX | + |
| Mbnl3 | 26,24 | 0,2127 | 0,762940356 | chrX | - |
| Mbtps2 | 171,21 | 0,3323 | 0,456560888 | chrX | - |
| Mcts1 | 880,61 | -0,1442 | 0,765219833 | chrX | + |
| Mecp2 | 256,63 | 0,0130 | 0,980186108 | chrX | - |
| Med14 | 229,39 | -0,1468 | 0,713100089 | chrX | - |
| Mid1 | 39,79 | 0,0159 | 0,985212135 | chrX | + |
| Mid2 | 113,66 | 0,2843 | 0,592000353 | chrX | + |
| Mmgt1 | 593,36 | -0,2407 | 0,523438189 | chrX | - |
| Morc4 | 52,44 | 0,5815 | 0,217208792 | chrX | - |
| Morf4l2 | 2903,75 | 0,4260 | 0,216874749 | chrX | - |
| Mospd1 | 340,16 | 0,0933 | 0,856916818 | chrX | - |
| Mospd2 | 650,69 | -0,0943 | 0,834674428 | chrX | - |
| Msl3 | 554,51 | 0,0403 | 0,942982044 | chrX | - |
| Msn | 393,11 | 0,3414 | 0,23347764 | chrX | + |
| Map7d3 | 2,39 | 0,5522 | NA | chrX | - |
| Mtcp1 | 57,08 | -0,5393 | 0,186118652 | chrX | - |
| Mtm1 | 241,82 | 0,0014 | 0,997729735 | chrX | + |
| Mtmr1 | 478,72 | -0,2749 | 0,39810264 | chrX | + |
| Naa10 | 335,67 | -0,6255 | 0,073076939 | chrX | - |
| Nap1l3 | 13,21 | 0,3400 | 0,63960646 | chrX | - |
| Ndufa1 | 3232,13 | -0,3113 | 0,286525456 | chrX | - |
| Ndufb11 | 3557,23 | -0,5031 | 0,264109601 | chrX | - |
| Bex3 | 22,44 | 0,1036 | 0,901481982 | chrX | + |
| Nhsl2 | 18,12 | 0,0551 | 0,949273123 | chrX | + |
| Nkap | 167,90 | 0,1579 | 0,695967514 | chrX | + |
| Nkrf | 67,65 | -0,3277 | 0,522768066 | chrX | - |
| Nono | 2137,90 | 0,0747 | 0,844155615 | chrX | + |
| Nsdhl | 1135,90 | -0,5569 | 0,133332328 | chrX | + |
| Nudt11 | 2,26 | -0,4952 | NA | chrX | + |
| Nxt2 | 663,56 | 0,4314 | 0,277831408 | chrX | + |
| Ocrl | 95,15 | 0,2822 | 0,549838805 | chrX | + |
| Ofd1 | 53,68 | 0,4989 | 0,305620059 | chrX | - |

| | | | | | |
|---|---|---|---|---|---|
| Ogt | 1044,95 | -0,1299 | 0,747522909 | chrX | + |
| Ophn1 | 20,66 | 0,6330 | 0,228930646 | chrX | - |
| Otc | 17716,89 | -0,2154 | 0,549694981 | chrX | + |
| Otud5 | 501,84 | -0,1825 | 0,636260727 | chrX | + |
| P2ry10 | 8,55 | 0,4502 | 0,528306301 | chrX | + |
| Pak3 | 2,41 | 0,5262 | NA | chrX | + |
| Pcsk1n | 4610,39 | -0,1190 | 0,764363417 | chrX | + |
| Pbdc1 | 286,07 | -0,1057 | 0,807278226 | chrX | + |
| Pcyt1b | 1407,53 | -0,0768 | 0,844511464 | chrX | + |
| Pdha1 | 2432,16 | -0,2937 | 0,292182106 | chrX | - |
| Pdk3 | 10,64 | 0,7307 | 0,228182471 | chrX | - |
| Pdzd11 | 559,10 | 0,0948 | 0,850572005 | chrX | - |
| Pdzd4 | 5,56 | -0,0101 | 0,992286283 | chrX | - |
| Pfkfb1 | 827,73 | -0,0733 | 0,88880917 | chrX | + |
| Pgk1 | 294,26 | -0,6339 | 0,149536902 | chrX | + |
| Pgrmc1 | 22432,35 | -0,4887 | 0,05081654 | chrX | + |
| Phex | 4,32 | 0,3064 | 0,693242675 | chrX | - |
| Phf6 | 143,11 | 0,4093 | 0,33637288 | chrX | + |
| Phf8 | 148,82 | -0,2013 | 0,651069698 | chrX | + |
| Phka1 | 112,78 | -0,2445 | 0,606429785 | chrX | - |
| Phka2 | 286,01 | 0,0016 | 0,997464504 | chrX | + |
| Piga | 118,22 | 0,2519 | 0,587879802 | chrX | + |
| Pim2 | 43,85 | -0,4193 | 0,416983017 | chrX | + |
| Pin4 | 768,24 | -0,4934 | 0,077224204 | chrX | + |
| Pir | 539,06 | 0,1156 | 0,811137679 | chrX | + |
| Pja1 | 965,25 | -0,1850 | 0,537952866 | chrX | - |
| Plp1 | 26,73 | 0,1673 | 0,802683933 | chrX | + |
| Plp2 | 569,45 | -0,0064 | 0,990189789 | chrX | - |
| Pls3 | 343,63 | 0,3568 | 0,381192505 | chrX | - |
| Plxna3 | 5,99 | 0,3818 | 0,611887546 | chrX | + |
| Plxnb3 | 3,59 | -0,2348 | 0,773303025 | chrX | + |
| Porcn | 30,03 | -0,3891 | 0,490047155 | chrX | - |
| Ppp1r3f | 1,87 | 0,5509 | NA | chrX | - |
| Pqbp1 | 324,55 | -0,2191 | 0,61779384 | chrX | - |
| Praf2 | 46,67 | -0,6070 | 0,229821529 | chrX | + |
| Prdx4 | 2988,19 | 0,0627 | 0,913749296 | chrX | - |
| Prickle3 | 27,89 | -0,3507 | 0,608287564 | chrX | + |
| Prkx | 49,52 | 0,1310 | 0,824000947 | chrX | - |
| Prps1 | 765,12 | 0,1620 | 0,637633057 | chrX | + |
| Prps2 | 305,14 | 0,5614 | 0,06084058 | chrX | - |
| Prrg3 | 12,46 | -0,2514 | 0,750261829 | chrX | + |
| Psmd10 | 354,05 | 0,0532 | 0,917614646 | chrX | - |
| Rab9 | 1171,96 | -0,3002 | 0,41552947 | chrX | - |
| Rap2c | 636,24 | 0,2526 | 0,439315481 | chrX | - |
| Rbbp7 | 1753,65 | 0,2109 | 0,471654089 | chrX | + |
| Rbm10 | 270,98 | -0,1668 | 0,720648137 | chrX | + |
| Rbm3 | 1007,80 | 0,2603 | 0,603013282 | chrX | - |
| Rbm41 | 123,28 | 0,6044 | 0,109852731 | chrX | - |
| Rbmx | 203,93 | 0,0033 | 0,995194784 | chrX | - |
| Rbmx2 | 28,84 | 0,3333 | 0,568858283 | chrX | + |

| | | | | | |
|---|---|---|---|---|---|
| Renbp | 77,46 | 0,0682 | 0,913749296 | chrX | - |
| Rtl5 | 10,01 | 0,2337 | 0,767097454 | chrX | - |
| Rgn | 34632,94 | -0,1728 | 0,730057183 | chrX | + |
| Rhox8 | 13,39 | -0,3162 | 0,67176872 | chrX | - |
| Ribc1 | 26,78 | 0,0493 | 0,952576171 | chrX | - |
| Ripply1 | 9,40 | 0,7454 | 0,227528657 | chrX | - |
| Rlim | 845,73 | 0,0350 | 0,944269942 | chrX | - |
| Rnf113a1 | 17,54 | 0,0745 | 0,938131508 | chrX | + |
| Rnf128 | 1794,24 | -0,1161 | 0,789520165 | chrX | + |
| Rp2 | 202,24 | 0,4654 | 0,204851518 | chrX | + |
| Rpgr | 35,50 | 0,2863 | 0,614153379 | chrX | - |
| Rpl36a | 4833,29 | -0,3814 | 0,23700863 | chrX | + |
| Rpl39 | 4758,18 | -0,4803 | 0,08440425 | chrX | - |
| Rps4x | 10751,96 | -0,3470 | 0,202636202 | chrX | - |
| Rragb | 3,58 | 0,9080 | 0,101361743 | chrX | + |
| Sash3 | 27,60 | 0,6093 | 0,210808905 | chrX | + |
| Sat1 | 1877,14 | -0,2406 | 0,494351553 | chrX | - |
| Serpina7 | 615,94 | -0,6551 | 0,19521384 | chrX | - |
| Sh2d1a | 3,41 | -0,0681 | 0,942425773 | chrX | + |
| Sh3bgrl | 2860,07 | 0,2194 | 0,588008527 | chrX | + |
| Sh3kbp1 | 42,62 | 0,2730 | 0,629969991 | chrX | + |
| Shroom2 | 266,45 | -0,0775 | 0,85496895 | chrX | - |
| Shroom4 | 2,50 | 0,3092 | NA | chrX | + |
| Siah1b | 8,98 | -0,2103 | 0,803738851 | chrX | - |
| Slc10a3 | 72,89 | 0,0842 | 0,88127014 | chrX | - |
| Slc16a2 | 2092,64 | 0,0049 | 0,990382071 | chrX | - |
| Slc25a14 | 66,35 | -0,0334 | 0,958903346 | chrX | + |
| Slc25a5 | 10043,64 | -0,1476 | 0,612344152 | chrX | + |
| Slc25a53 | 11,97 | 0,4467 | 0,517431746 | chrX | - |
| Slc35a2 | 170,07 | -0,2940 | 0,486771962 | chrX | + |
| Slc6a8 | 115,24 | 0,6350 | 0,093542539 | chrX | + |
| Slc9a6 | 252,68 | -0,3458 | 0,287408362 | chrX | + |
| Slc9a7 | 4,87 | -0,0244 | 0,981991128 | chrX | - |
| Smarca1 | 3,55 | 0,7040 | 0,24947031 | chrX | - |
| Smc1a | 1074,48 | 0,1017 | 0,83796616 | chrX | + |
| Smim9 | 3,17 | -0,2250 | 0,772961171 | chrX | - |
| Sms | 135,18 | -0,1497 | 0,716547565 | chrX | - |
| Snora70 | 13,73 | -0,7164 | 0,218722998 | chrX | + |
| Snx12 | 220,79 | 0,3255 | 0,323749114 | chrX | - |
| Spin2c | 13,04 | 0,5622 | 0,376702038 | chrX | + |
| Spin4 | 6,31 | 0,9368 | 0,114599337 | chrX | - |
| Srpx | 10,51 | 0,6433 | 0,321708301 | chrX | - |
| Srpx2 | 37,41 | -0,1224 | 0,858880859 | chrX | + |
| Stag2 | 1206,32 | 0,1711 | 0,70880025 | chrX | + |
| Stard8 | 150,24 | 0,1710 | 0,682099278 | chrX | + |
| Suv39h1 | 73,12 | -0,1098 | 0,848745988 | chrX | - |
| Syap1 | 960,49 | 0,0592 | 0,905147397 | chrX | - |
| Syn1 | 3,72 | -0,0163 | 0,986997724 | chrX | - |
| Syp | 13,13 | 0,4007 | 0,564724076 | chrX | + |
| Sytl4 | 54,33 | 0,8232 | 0,051447413 | chrX | - |

| | | | | | |
|---|---|---|---|---|---|
| Tab3 | 164,71 | -0,1503 | 0,740269415 | chrX | + |
| Taf1 | 225,65 | 0,0364 | 0,951806422 | chrX | + |
| Taf9b | 32,20 | 0,2125 | 0,719668754 | chrX | - |
| Taz | 377,43 | -0,1946 | 0,600688917 | chrX | + |
| Tbc1d25 | 28,87 | -0,3019 | 0,604491975 | chrX | - |
| Tbc1d8b | 507,92 | 0,2987 | 0,451967425 | chrX | + |
| Tbl1x | 147,65 | 0,1479 | 0,757695155 | chrX | + |
| Tceal1 | 25,87 | 0,4925 | 0,398132968 | chrX | + |
| Tceal5 | 2,31 | 0,4444 | NA | chrX | - |
| Tceanc | 34,12 | 0,2193 | 0,717576273 | chrX | - |
| Tfe3 | 257,04 | -0,2007 | 0,66477796 | chrX | + |
| Thoc2 | 409,97 | 0,5057 | 0,113136277 | chrX | - |
| Timm17b | 521,81 | -0,2237 | 0,614056134 | chrX | + |
| Timm8a1 | 1386,37 | -0,1474 | 0,740663168 | chrX | - |
| Tlr13 | 61,02 | 0,4632 | 0,248800492 | chrX | + |
| Tmem164 | 173,31 | 0,3625 | 0,28821012 | chrX | + |
| Tmem29 | 268,27 | 0,3990 | 0,251005984 | chrX | - |
| Tmem35a | 3,81 | 0,6410 | 0,306414953 | chrX | + |
| Tmem47 | 102,45 | 0,4847 | 0,238794764 | chrX | + |
| Tmsb15b1 | 21,30 | 0,5567 | 0,317646908 | chrX | - |
| Tmsb4x | 3171,27 | 0,2125 | 0,558489002 | chrX | - |
| Trappc2 | 123,27 | 0,2856 | 0,534939703 | chrX | + |
| Trmt2b | 622,31 | 0,1130 | 0,794116376 | chrX | - |
| Tro | 2,30 | 0,1163 | NA | chrX | - |
| Tsc22d3 | 2124,88 | -0,5987 | 0,191230937 | chrX | - |
| Tspan6 | 16,24 | 0,8355 | 0,151287224 | chrX | - |
| Tspan7 | 409,03 | 0,0590 | 0,880810135 | chrX | + |
| Tspyl2 | 97,27 | -0,1059 | 0,848993225 | chrX | - |
| Tsr2 | 170,17 | -0,0335 | 0,948647685 | chrX | - |
| Txlng | 510,89 | 0,0846 | 0,848745988 | chrX | - |
| Uba1 | 1978,31 | -0,1382 | 0,7588734 | chrX | + |
| Ube2a | 519,53 | -0,3443 | 0,379104437 | chrX | + |
| Ubl4a | 288,90 | -0,2783 | 0,391329882 | chrX | - |
| Ubqln2 | 175,05 | -0,4661 | 0,114805312 | chrX | + |
| Upf3b | 270,97 | -0,3604 | 0,248510194 | chrX | - |
| Uprt | 5,08 | 0,8485 | 0,156728357 | chrX | + |
| Usp11 | 9,65 | 0,5180 | 0,437620234 | chrX | + |
| Usp27x | 13,33 | -0,2038 | 0,803874753 | chrX | - |
| Usp9x | 2346,23 | -0,1844 | 0,565966999 | chrX | + |
| Utp14a | 340,51 | -0,2791 | 0,401382558 | chrX | + |
| Vbp1 | 359,08 | 0,5226 | 0,080029766 | chrX | + |
| Vma21 | 600,66 | -0,0019 | 0,996826205 | chrX | + |
| Vsig4 | 1077,90 | 0,1588 | 0,684228218 | chrX | - |
| Was | 26,79 | 0,4664 | 0,38519619 | chrX | - |
| Tceal9 | 553,32 | 0,3423 | 0,169907896 | chrX | + |
| Wdr13 | 5,99 | 0,0264 | 0,981025197 | chrX | - |
| Wdr44 | 66,73 | 0,5519 | 0,166993075 | chrX | + |
| Wdr45 | 895,30 | -0,4127 | 0,112275385 | chrX | + |
| Xiap | 2050,33 | 0,2172 | 0,526375055 | chrX | + |
| Xist | 2,42 | 0,2172 | NA | chrX | - |

| | | | | | |
|---|---|---|---|---|---|
| Xk | 82,62 | -0,2532 | 0,632136487 | chrX | + |
| Xkrx | 2,12 | -0,0878 | NA | chrX | - |
| Xlr | 29,90 | 0,3167 | 0,626098415 | chrX | - |
| Xlr3a | 103,04 | 0,1701 | 0,845988264 | chrX | - |
| Xlr4b | 7,16 | 0,2987 | 0,558489002 | chrX | + |
| Yipf6 | 697,45 | 0,0215 | 0,969130697 | chrX | + |
| Yy2 | 11,55 | 0,6099 | 0,345491882 | chrX | - |
| Zbtb33 | 529,38 | 0,4810 | 0,14723273 | chrX | + |
| Zc4h2 | 36,14 | 0,5637 | 0,303750468 | chrX | - |
| Zdhhc15 | 4,74 | 0,4077 | 0,569876243 | chrX | - |
| Zdhhc9 | 1200,29 | -0,0106 | 0,98398644 | chrX | - |
| Zfp182 | 126,25 | 0,6373 | 0,100767966 | chrX | - |
| Zfp185 | 4,37 | 0,6798 | 0,287543614 | chrX | + |
| Zfp275 | 239,96 | 0,3900 | 0,205441527 | chrX | + |
| Zfp280c | 82,21 | 0,6205 | 0,117246861 | chrX | - |
| Zfp300 | 20,41 | -0,1985 | 0,783422709 | chrX | - |
| Zfx | 270,20 | 0,2100 | 0,642318814 | chrX | - |
| Zmym3 | 145,47 | -0,2577 | 0,514948445 | chrX | - |
| Zrsr2 | 114,81 | -0,0118 | 0,984995389 | chrX | - |
| Zxda | 35,08 | -0,5451 | 0,291313478 | chrX | - |
| Zxdb | 28,37 | -0,0894 | 0,899463828 | chrX | + |
| Ddx3y | 1159,09 | 0,5028 | 0,18993243 | chrY | - |
| Eif2s3y | 1520,63 | 0,0131 | 0,980379461 | chrY | + |
| Kdm5d | 237,55 | 0,6347 | 0,07450906 | chrY | + |
| Uba1y | 8,06 | -0,4480 | 0,528306301 | chrY | + |
| Uty | 280,96 | 0,5931 | 0,10394222 | chrY | - |
| Adora1 | 558,0258406 | 1,068714996 | 3,52E-06 | Chr1 | - |
| Btg2 | 289,9862211 | 1,000148207 | 0,000338147 | Chr1 | - |
| Ccdc181 | 89,2614866 | 0,787724763 | 0,01711988 | Chr1 | + |
| Cd244 | 13,80542759 | 1,312951317 | 0,009000221 | Chr1 | + |
| Cd34 | 62,85920594 | 0,844399641 | 0,042803116 | Chr1 | + |
| Cd46 | 13,78886225 | 1,522716332 | 0,001100299 | Chr1 | - |
| Chil1 | 7,052046148 | 1,525160624 | 0,002162884 | Chr1 | + |
| Cnga3 | 8,27395137 | 1,907824504 | 4,24E-05 | Chr1 | + |
| Col4a3 | 15,56786367 | 1,570195738 | 0,001415734 | Chr1 | + |
| Col5a2 | 416,1712747 | 2,617492523 | 1,10E-19 | Chr1 | - |
| Cr1l | 1526,147417 | 0,528718657 | 0,022395757 | Chr1 | - |
| Cr2 | 6,552390849 | 1,194500334 | 0,028862631 | Chr1 | - |
| Csrp1 | 246,306641 | 0,774629621 | 0,000747245 | Chr1 | + |
| Efcab2 | 102,3484609 | 1,549215725 | 3,30E-07 | Chr1 | + |
| Fam129a | 41,79123244 | 0,838298812 | 0,047921745 | Chr1 | + |
| Fmo2 | 298,4862998 | 0,788775904 | 0,043054595 | Chr1 | - |
| Fmo3 | 93,2763624 | 1,176646157 | 0,022395757 | Chr1 | - |
| Gls | 127,4854708 | 0,633939322 | 0,037660849 | Chr1 | - |
| Gm16897 | 8,11488606 | 1,905725166 | 4,31E-05 | Chr1 | - |
| Gm19705 | 38,23555259 | 1,042347445 | 0,019113605 | Chr1 | + |
| Gpr35 | 19,61715249 | 1,026264547 | 0,033493066 | Chr1 | + |
| Hhipl2 | 9,472994684 | 1,520143627 | 0,00208761 | Chr1 | + |
| Ier5 | 78,9199725 | 1,11451532 | 0,00018056 | Chr1 | - |
| Ifi205 | 26,7388254 | 0,93243436 | 0,035186081 | Chr1 | - |

| | | | | | |
|---|---|---|---|---|---|
| Kcnt2 | 51,59229875 | 1,748179568 | 1,49E-05 | Chr1 | + |
| Ly9 | 50,28074732 | 0,762314602 | 0,043954241 | Chr1 | - |
| Nbeal1 | 144,1529039 | 0,923067174 | 0,014488154 | Chr1 | + |
| Pkhd1 | 177,2545668 | 0,847941693 | 0,022642478 | Chr1 | - |
| Pms1 | 161,8686132 | 0,86456017 | 0,009614825 | Chr1 | - |
| Prelp | 607,3063142 | 0,511245744 | 0,033836468 | Chr1 | - |
| Ptprc | 232,5475162 | 0,812383391 | 0,0063705 | Chr1 | - |
| Rgs5 | 236,8436256 | 1,090890778 | 5,89E-05 | Chr1 | + |
| Rps6kc1 | 217,1823045 | 1,114159479 | 0,000132242 | Chr1 | - |
| Selp | 25,58739175 | 1,048872731 | 0,026733412 | Chr1 | + |
| Serpinb8 | 55,91195578 | 1,030515984 | 0,011248868 | Chr1 | + |
| Sp110 | 69,41085369 | 0,97975157 | 0,01157497 | Chr1 | - |
| Spats2l | 56,38811727 | 1,378738156 | 0,001492349 | Chr1 | + |
| Spp2 | 5667,511934 | 0,570410458 | 0,026196265 | Chr1 | + |
| Tmem237 | 81,55329256 | 0,96757102 | 0,00552253 | Chr1 | - |
| Trip12 | 129,6001773 | 0,656731653 | 0,045315053 | Chr1 | - |
| Vil1 | 6,468875243 | 1,444840875 | 0,004663891 | Chr1 | + |
| Platr14 | 17,81948986 | 2,612426868 | 9,66E-10 | chr2 | + |
| B3galt1 | 1039,18083 | 0,870143269 | 0,008539289 | chr2 | + |
| Bmyc | 111,1604805 | 1,293708441 | 0,013752458 | chr2 | + |
| Calcrl | 367,5152209 | 0,746340141 | 0,04497473 | chr2 | - |
| Ccdc141 | 203,2845893 | 0,910587171 | 0,000368685 | chr2 | - |
| Cd93 | 159,7988717 | 0,607907336 | 0,0348796 | chr2 | - |
| Dab2ip | 130,4956406 | 0,731192281 | 0,03992849 | chr2 | + |
| Dbndd2 | 122,3447512 | 0,646801721 | 0,03261247 | chr2 | + |
| Dhrs9 | 24,91980786 | 1,311423079 | 0,005437641 | chr2 | + |
| Dnajc10 | 782,1947691 | 0,556457673 | 0,03661192 | chr2 | + |
| Eid1 | 214,5607324 | 0,830757165 | 0,000961755 | chr2 | + |
| Fam129b | 239,1726222 | 1,830333363 | 2,71E-09 | chr2 | + |
| Fermt1 | 16,55759387 | 1,653690446 | 0,000304342 | chr2 | - |
| Fmn1 | 215,5817073 | 1,045352948 | 0,002587428 | chr2 | + |
| Gm14124 | 5,785206186 | 1,298600473 | 0,012626346 | chr2 | + |
| Gm14139 | 5,959224028 | 1,518548301 | 0,002073441 | chr2 | + |
| Gtdc1 | 160,831173 | 1,283144346 | 2,46E-07 | chr2 | - |
| Il1rn | 108,0442411 | 1,082823898 | 0,002387116 | chr2 | + |
| Itih5 | 827,1843833 | 1,449364548 | 1,41E-11 | chr2 | + |
| Lrp2 | 29,72486355 | 1,647526256 | 0,000328253 | chr2 | - |
| Mettl15 | 102,0364141 | 0,999380893 | 0,003158251 | chr2 | - |
| Mga | 505,2555073 | 0,786929169 | 0,003798776 | chr2 | + |
| Myef2 | 166,9839342 | 0,687058892 | 0,023381506 | chr2 | - |
| Napb | 395,8284803 | 3,640677203 | 3,45E-34 | chr2 | - |
| Nmi | 739,283815 | 0,856766646 | 0,014703567 | chr2 | - |
| Nphp1 | 120,4270262 | 0,759230908 | 0,033836468 | chr2 | - |
| Obp2a | 8615,5635 | 7,153773635 | 4,26E-113 | chr2 | + |
| Olfr1033 | 206,1669164 | 0,626126579 | 0,042951355 | chr2 | + |
| Pak6 | 12,92346851 | 1,385647088 | 0,007036141 | chr2 | + |
| Pygb | 140,6467829 | 0,763838787 | 0,017102722 | chr2 | + |
| Rbm43 | 123,3744467 | 1,006668804 | 0,011778732 | chr2 | - |
| Samd10 | 34,16002157 | 1,119969978 | 0,004837875 | chr2 | - |
| Skida1 | 12,76430637 | 1,124583905 | 0,034704076 | chr2 | - |

| | | | | | |
|---|---|---|---|---|---|
| Slc13a3 | 190,3262805 | 0,798103196 | 0,020664082 | chr2 | - |
| Slpi | 113,9034059 | 1,688474746 | 8,72E-07 | chr2 | - |
| Snhg11 | 236,3029118 | 1,750697255 | 2,17E-06 | chr2 | + |
| Src | 47,86477863 | 1,28680343 | 0,001466109 | chr2 | + |
| Tcfl5 | 39,63208553 | 3,680573935 | 9,39E-22 | chr2 | - |
| Trdmt1 | 61,16654816 | 1,30871656 | 0,001857909 | chr2 | - |
| Ttc30b | 7,008194404 | 1,149775972 | 0,037660849 | chr2 | - |
| Uap1l1 | 144,4452808 | 1,106503647 | 1,01E-05 | chr2 | - |
| Ube2l6 | 2492,65762 | 0,906485084 | 0,000197378 | chr2 | + |
| Wfdc12 | 355,2179321 | 5,172619603 | 8,67E-53 | chr2 | - |
| Wfdc15b | 25,91980068 | 1,82152364 | 5,89E-05 | chr2 | - |
| Wfdc2 | 350,7531028 | 2,495420497 | 7,95E-23 | chr2 | + |
| Zbp1 | 485,7619492 | 1,560261491 | 1,91E-06 | chr2 | - |
| Zfp120 | 253,1621832 | 0,878685222 | 0,001289374 | chr2 | - |
| Zfp345 | 11,77513163 | 2,073777505 | 3,82E-06 | chr2 | - |
| Zfp770 | 149,0849614 | 0,929173761 | 0,001448591 | chr2 | - |
| Acad9 | 816,4419474 | 0,571404464 | 0,017210327 | chr3 | + |
| Amy1 | 12509,06312 | 0,714472956 | 0,031250949 | chr3 | - |
| Ank2 | 38,4438595 | 0,987983323 | 0,010993925 | chr3 | - |
| Anxa5 | 1720,139071 | 0,785709947 | 0,006224175 | chr3 | - |
| Bche | 3630,878878 | 0,747814942 | 0,043328016 | chr3 | - |
| Bglap | 3,582044473 | 1,140459141 | 0,028526877 | chr3 | - |
| Bglap3 | 1137,292917 | 7,265902439 | 1,57E-148 | chr3 | - |
| Car14 | 1401,529208 | 0,553471049 | 0,042602547 | chr3 | - |
| Cgn | 267,434322 | 0,709555657 | 0,014703567 | chr3 | - |
| Clca1 | 78,78875689 | 0,99260168 | 0,004704331 | chr3 | - |
| Col25a1 | 37,86445346 | 1,776908396 | 0,000136133 | chr3 | + |
| E130311K13Rik | 118,7932323 | 0,877911461 | 0,016854045 | chr3 | - |
| Ect2 | 38,36749319 | 1,130730497 | 0,01865562 | chr3 | - |
| Fabp12 | 12,37599746 | 1,541557894 | 0,001760215 | chr3 | - |
| Gbp2 | 181,9728754 | 0,823745857 | 0,001540278 | chr3 | + |
| Gbp3 | 260,5648915 | 0,954056659 | 0,000557541 | chr3 | + |
| Gbp7 | 345,4364776 | 1,101660175 | 0,000852014 | chr3 | + |
| Gipc2 | 14,61229458 | 1,768516797 | 0,000208779 | chr3 | - |
| Gm20752 | 20,01448974 | 1,222976662 | 0,018778033 | chr3 | + |
| Gimd1 | 96,17532284 | 1,334673622 | 0,000682221 | chr3 | + |
| Hao2 | 31,27098371 | 1,980919168 | 7,57E-06 | chr3 | - |
| Ifi44 | 244,9893236 | 1,75007843 | 3,53E-12 | chr3 | - |
| Il7 | 10,58188101 | 1,122545063 | 0,0400769 | chr3 | - |
| Ints12 | 176,4550587 | 0,84625329 | 0,027149801 | chr3 | + |
| Lrif1 | 235,2149497 | 0,813851415 | 0,007909756 | chr3 | + |
| Mgst2 | 6,076815482 | 1,199772565 | 0,027483386 | chr3 | + |
| Mtmr11 | 43,94414334 | 1,160442412 | 0,003531077 | chr3 | + |
| P2ry13 | 107,1602959 | 0,630655299 | 0,038944122 | chr3 | - |
| Palmd | 626,4092294 | 0,728067121 | 0,016497036 | chr3 | - |
| Pbxip1 | 251,2137521 | 0,622499391 | 0,014620211 | chr3 | + |
| Plrg1 | 135,8799601 | 2,251156473 | 1,13E-08 | chr3 | + |
| Rfx5 | 127,5676217 | 0,988278672 | 0,001216836 | chr3 | + |
| Rhoc | 303,1930858 | 0,858395987 | 0,000725675 | chr3 | + |
| S100a10 | 2848,413506 | 0,666314148 | 0,045665077 | chr3 | + |

| | | | | | |
|---|---|---|---|---|---|
| S100a11 | 361,0074868 | 1,112515288 | 0,00035788 | chr3 | + |
| S100a4 | 38,86804583 | 0,970764326 | 0,03196404 | chr3 | + |
| Sema4a | 903,5008227 | 0,63692349 | 0,040153429 | chr3 | - |
| Slc10a5 | 640,1403158 | 0,755904615 | 0,048623583 | chr3 | - |
| Sprr1a | 8,80338552 | 1,205302162 | 0,022027814 | chr3 | - |
| Sucnr1 | 1128,769277 | 0,970742056 | 0,010220245 | chr3 | + |
| Supt20 | 99,07745445 | 0,797135332 | 0,046074596 | chr3 | + |
| Tnfsf10 | 240,2832743 | 1,003202427 | 0,005437701 | chr3 | + |
| Tuft1 | 137,0931777 | 0,69919673 | 0,03196404 | chr3 | - |
| Vcam1 | 560,6302691 | 0,87259343 | 0,009528797 | chr3 | - |
| Veph1 | 18,88303747 | 1,545098549 | 0,001244855 | chr3 | - |
| Vtcn1 | 11,30599784 | 1,267403281 | 0,012917631 | chr3 | + |
| Zfp979 | 38,68427515 | 2,75465916 | 5,84E-11 | chr4 | - |
| 4930480G23Rik | 137,5900418 | 0,703126207 | 0,041373145 | chr4 | - |
| 4933438K21Rik | 9,703182946 | 1,434122476 | 0,004317857 | chr4 | - |
| Acnat2 | 347,1599271 | 1,172280952 | 0,000429211 | chr4 | - |
| Aldh1b1 | 2583,763818 | 1,690262737 | 1,19E-19 | chr4 | + |
| Aqp7 | 26,0568193 | 1,739119 | 8,42E-05 | chr4 | - |
| Arhgef16 | 60,68199741 | 0,882718767 | 0,022925104 | chr4 | - |
| Atp6v0d2 | 237,2068961 | 1,701872485 | 0,000412839 | chr4 | - |
| Atp8b5 | 16,67260017 | 1,501835454 | 0,001211545 | chr4 | + |
| BC039966 | 10,61138062 | 2,093018389 | 4,03E-06 | chr4 | - |
| Bmp8b | 11,42050792 | 2,149483342 | 1,63E-06 | chr4 | + |
| Ccdc171 | 115,2486657 | 1,305185646 | 2,48E-05 | chr4 | + |
| Cdkn2c | 150,0492963 | 0,655178766 | 0,044481388 | chr4 | - |
| Dnaic1 | 72,44639518 | 1,024251931 | 0,03196404 | chr4 | + |
| Exo5 | 69,4886132 | 0,847939053 | 0,010594723 | chr4 | - |
| Gabbr2 | 132,6645152 | 0,879542777 | 0,016718239 | chr4 | - |
| Gm12718 | 120,4680762 | 3,548745048 | 1,27E-26 | chr4 | + |
| Zfp987 | 18,21087767 | 1,289637995 | 0,008769355 | chr4 | + |
| Zfp982 | 126,2829872 | 1,347920311 | 0,000155769 | chr4 | + |
| Zfp984 | 127,3238758 | 2,303525419 | 2,78E-17 | chr4 | - |
| Zfp981 | 8,897912649 | 2,012554368 | 1,14E-05 | chr4 | + |
| Zfp992 | 22,89414619 | 1,5570614 | 0,0005034 | chr4 | + |
| Impad1 | 621,0496551 | 0,78099927 | 0,014352044 | chr4 | - |
| Patj | 234,1203981 | 2,534908336 | 3,05E-24 | chr4 | + |
| Klhdc7a | 471,1378957 | 0,901090447 | 0,034522477 | chr4 | - |
| Lepr | 238,8374278 | 1,292513909 | 0,004306468 | chr4 | + |
| Megf6 | 119,389932 | 1,024834661 | 0,008728311 | chr4 | + |
| Msantd3 | 6,35146565 | 1,106223606 | 0,048435774 | chr4 | + |
| Mup21 | 20209,26927 | 3,877353202 | 2,25E-50 | chr4 | - |
| Mycl | 204,5923726 | 0,953908658 | 0,016705702 | chr4 | + |
| Orm3 | 139,7676487 | 2,097657377 | 2,62E-10 | chr4 | + |
| Pars2 | 56,29963631 | 0,888486162 | 0,017119669 | chr4 | + |
| Pgd | 811,0183979 | 0,718783473 | 0,035763561 | chr4 | - |
| Plag1 | 6,852460297 | 1,270347821 | 0,017286693 | chr4 | - |
| Ptgr1 | 193,3942618 | 1,221665185 | 9,36E-06 | chr4 | - |
| Skint9 | 4,543585524 | 1,362418211 | 0,007218232 | chr4 | - |
| Slc5a9 | 4,263001239 | 1,17635642 | 0,02272099 | chr4 | - |
| Smarca5-ps | 8,071984124 | 1,169955124 | 0,032954496 | chr4 | + |

| | | | | | |
|---|---|---|---|---|---|
| Spaca1 | 8,634011146 | 1,660687934 | 0,00064465 | chr4 | - |
| Themis2 | 112,1325453 | 0,773458234 | 0,03905306 | chr4 | - |
| Dmac1 | 791,3457784 | 1,118862423 | 1,92E-05 | chr4 | - |
| Tstd2 | 1027,153629 | 1,464987924 | 4,78E-08 | chr4 | - |
| Tstd3 | 771,7094154 | 0,683296826 | 0,027907786 | chr4 | - |
| Ttc39a | 46,64664503 | 1,746527967 | 7,81E-06 | chr4 | + |
| Ubxn2b | 73,57250484 | 0,902672785 | 0,006796468 | chr4 | + |
| Zfp933 | 129,404934 | 0,85459798 | 0,005257528 | chr4 | - |
| Zmym6 | 174,0921726 | 1,167763813 | 4,07E-05 | chr4 | + |
| Znf41-ps | 48,43954732 | 2,73567188 | 7,68E-16 | chr4 | + |
| 2310034O05Rik | 24,64713388 | 2,164332162 | 1,12E-07 | chr5 | + |
| A830010M20Rik | 151,293423 | 2,344314174 | 1,72E-19 | chr5 | + |
| Abcb1a | 68,36568484 | 1,059082668 | 0,006385096 | chr5 | + |
| Acad12 | 87,75934778 | 0,791627783 | 0,049774092 | chr5 | - |
| Afp | 65,33735904 | 1,912855277 | 1,99E-09 | chr5 | + |
| Gpat3 | 186,1778329 | 1,945900476 | 9,38E-09 | chr5 | + |
| Btbd8 | 12,19280096 | 1,919037504 | 2,98E-05 | chr5 | + |
| Cldn4 | 4,240267624 | 1,133414259 | 0,034599171 | chr5 | - |
| Cxcl10 | 99,92809461 | 1,009791639 | 0,002215245 | chr5 | - |
| D130017N08Rik | 11,75966759 | 1,256852679 | 0,017503197 | chr5 | + |
| Drc1 | 39,57567389 | 1,19585288 | 0,010521428 | chr5 | + |
| Eln | 51,24869767 | 0,934298348 | 0,03196404 | chr5 | - |
| Fam126a | 159,6065756 | 0,764542554 | 0,00285195 | chr5 | - |
| Fam184b | 11,80893952 | 1,323411715 | 0,010105755 | chr5 | - |
| Gbp10 | 206,5897235 | 0,910810926 | 0,028940327 | chr5 | - |
| Gbp6 | 295,3771399 | 0,88348215 | 0,019328711 | chr5 | - |
| Gbp8 | 116,6315359 | 1,509967048 | 3,89E-06 | chr5 | - |
| Gbp9 | 103,8035756 | 1,264234589 | 5,16E-05 | chr5 | - |
| Gm4961 | 13,23415046 | 1,98801191 | 1,40E-05 | chr5 | + |
| Gpc2 | 11,86833604 | 1,46606097 | 0,002828071 | chr5 | - |
| Hgf | 140,8569251 | 1,425692691 | 3,07E-06 | chr5 | + |
| Hpse | 12,34958216 | 1,118612552 | 0,041002418 | chr5 | - |
| Lfng | 1126,185169 | 1,070647461 | 1,51E-05 | chr5 | + |
| Mmd2 | 367,2206572 | 1,150450227 | 0,001465109 | chr5 | - |
| Nipal1 | 274,1008174 | 2,368843286 | 1,10E-12 | chr5 | + |
| Oas1a | 86,42964964 | 0,90043983 | 0,027149801 | chr5 | - |
| Oasl1 | 301,2067524 | 1,433332463 | 0,000181364 | chr5 | + |
| Pgm1 | 107,2806119 | 0,729895022 | 0,022642478 | chr5 | + |
| Rad9b | 14,22900816 | 1,193337608 | 0,01644885 | chr5 | - |
| Rgs12 | 88,79585829 | 1,031655476 | 0,004194139 | chr5 | + |
| Sema3d | 46,32527779 | 0,957866622 | 0,049900203 | chr5 | + |
| Serpine1 | 39,17917453 | 0,890583003 | 0,048623583 | chr5 | - |
| Sgcb | 52,31016111 | 0,761452954 | 0,046472053 | chr5 | - |
| Sgsm1 | 128,0067419 | 1,681442894 | 6,44E-07 | chr5 | - |
| Spon2 | 241,6599665 | 1,697644808 | 4,06E-09 | chr5 | - |
| Spp1 | 2347,226458 | 1,154337093 | 0,00241197 | chr5 | + |
| Stag3 | 15,38451344 | 1,538603707 | 0,000908604 | chr5 | + |
| Stap1 | 100,0038663 | 2,784580051 | 3,16E-17 | chr5 | + |
| Steap4 | 3646,264517 | 0,988017473 | 0,022287028 | chr5 | + |
| Sult1b1 | 1445,751387 | 0,891320293 | 0,01896692 | chr5 | - |

| | | | | | |
|---|---|---|---|---|---|
| Tbc1d19 | 82,23906899 | 0,921124686 | 0,013849558 | chr5 | + |
| Trim56 | 66,74985531 | 1,176127555 | 0,008885032 | chr5 | - |
| Ttc28 | 6,591901532 | 1,656959048 | 0,000619592 | chr5 | + |
| Uba6 | 133,3376269 | 0,872628559 | 0,008098959 | chr5 | - |
| Vsig10 | 60,56093651 | 0,799746629 | 0,043876034 | chr5 | + |
| Zfp157 | 117,0009221 | 0,764564877 | 0,010705465 | chr5 | + |
| Zfp605 | 63,64056225 | 1,395626802 | 9,90E-05 | chr5 | + |
| Zfp951 | 58,27348492 | 1,98198027 | 1,46E-10 | chr5 | - |
| Adamts9 | 30,93085626 | 1,216948553 | 0,007211123 | chr6 | - |
| Agk | 164,6292533 | 0,813747229 | 0,025271405 | chr6 | + |
| Atp2b2 | 22,73037647 | 1,690476636 | 0,000305393 | chr6 | - |
| Atp6v0e2 | 26,16196458 | 1,483486021 | 0,000326485 | chr6 | + |
| B4galnt3 | 27,58344443 | 1,055682031 | 0,033957292 | chr6 | - |
| BC048671 | 8,725077561 | 1,64861898 | 0,000700981 | chr6 | + |
| Bcl2l14 | 35,94823132 | 1,197180066 | 0,024709274 | chr6 | + |
| Bmp10 | 6,781032893 | 1,135089185 | 0,04104012 | chr6 | + |
| C1s1 | 13061,85666 | 0,588915987 | 0,009803754 | chr6 | - |
| Ccdc91 | 598,5374697 | 0,517459666 | 0,017401276 | chr6 | + |
| Clec7a | 175,3203145 | 0,853785843 | 0,011537428 | chr6 | - |
| Cntn6 | 3,152534435 | 1,06707077 | 0,042234184 | chr6 | + |
| D630045J12Rik | 46,88459938 | 0,762466294 | 0,049463927 | chr6 | - |
| Dqx1 | 150,7246053 | 1,076583098 | 0,001274463 | chr6 | + |
| Emp1 | 81,93755469 | 1,289892864 | 0,001908703 | chr6 | + |
| Foxp2 | 83,87278092 | 0,833148628 | 0,014352044 | chr6 | + |
| Ggct | 180,1475678 | 0,772627254 | 0,009803754 | chr6 | - |
| Nat8f7 | 17,23459728 | 2,253332947 | 3,20E-07 | chr6 | - |
| H1fx | 5,708239462 | 1,355836802 | 0,007211123 | chr6 | - |
| Herc6 | 253,9325714 | 0,672160832 | 0,022759599 | chr6 | + |
| Klrb1b | 5,37324845 | 1,121219318 | 0,043954241 | chr6 | - |
| Mansc4 | 19,89999572 | 1,046701291 | 0,037864464 | chr6 | - |
| Mob1a | 179,902789 | 0,690008766 | 0,026151101 | chr6 | + |
| Mturn | 35,54709529 | 1,639894585 | 0,000669752 | chr6 | + |
| Nat8 | 677,3868532 | 2,007895385 | 8,85E-10 | chr6 | - |
| Nup210 | 267,5636784 | 0,639892837 | 0,049606348 | chr6 | - |
| Osbpl3 | 25,02296474 | 1,211904538 | 0,006396217 | chr6 | - |
| Parp11 | 105,4119968 | 0,973613175 | 0,003074994 | chr6 | + |
| Pot1a | 102,5910669 | 0,714586379 | 0,037343379 | chr6 | - |
| Ppp1r9a | 269,8900026 | 1,788613777 | 5,16E-09 | chr6 | + |
| Samd9l | 681,5085203 | 1,363114041 | 7,30E-06 | chr6 | - |
| Smo | 115,0083055 | 0,657212449 | 0,037185968 | chr6 | + |
| Tmem209 | 200,4858866 | 0,764089914 | 0,004864608 | chr6 | - |
| Tra2a | 672,1428937 | 0,721482141 | 0,02522906 | chr6 | - |
| Trim24 | 807,7044658 | 1,783448984 | 1,76E-14 | chr6 | + |
| Tspan9 | 593,9850229 | 0,604215506 | 0,017856953 | chr6 | - |
| Vamp5 | 289,484513 | 0,885415195 | 0,002280106 | chr6 | - |
| 2810047C21Rik1 | 29,55303649 | 2,631849536 | 4,49E-11 | chr7 | - |
| Abhd2 | 260,2455392 | 1,63604243 | 5,76E-09 | chr7 | + |
| Aqp8 | 1984,582077 | 0,904661015 | 0,046059699 | chr7 | + |
| Catsperg2 | 184,1866692 | 4,664978934 | 1,45E-36 | chr7 | - |
| Chp2 | 59,98766685 | 0,864103664 | 0,043636165 | chr7 | + |

| | | | | | |
|---|---|---|---|---|---|
| Crym | 28,90682194 | 1,464950422 | 0,000404796 | chr7 | - |
| Csrp3 | 902,6190197 | 1,022920452 | 6,84E-06 | chr7 | - |
| Cyp2a22 | 637,451666 | 1,061227951 | 0,00055932 | chr7 | - |
| Cyp2a4 | 4969,118811 | 1,088738496 | 0,034199883 | chr7 | + |
| Cyp2b10 | 524,8666731 | 4,060631815 | 1,41E-38 | chr7 | + |
| Cyp2b13 | 91,58147639 | 1,801613326 | 0,000119831 | chr7 | + |
| Cyp2b9 | 747,3317114 | 2,678014678 | 4,51E-10 | chr7 | + |
| Det1 | 117,5464051 | 1,477759713 | 6,09E-07 | chr7 | - |
| Dpy19l3 | 456,4670633 | 3,249533581 | 3,35E-41 | chr7 | - |
| Fam53b | 110,046893 | 0,851502539 | 0,025287128 | chr7 | - |
| Far1 | 72,17247661 | 0,840675747 | 0,037414363 | chr7 | + |
| Fxyd3 | 18,55136718 | 1,04436363 | 0,049007441 | chr7 | - |
| Fxyd5 | 211,3470938 | 1,169697197 | 8,56E-07 | chr7 | - |
| Gapdhs | 18,84187907 | 1,858632442 | 8,55E-06 | chr7 | - |
| H19 | 213,4076374 | 2,124768952 | 2,67E-06 | chr7 | - |
| Hddc3 | 45,08379681 | 0,850927104 | 0,038478567 | chr7 | + |
| Il18bp | 432,3290563 | 0,596153147 | 0,013040519 | chr7 | - |
| Kcne3 | 49,18039452 | 0,913887784 | 0,043490691 | chr7 | + |
| Klk1b4 | 161,4782993 | 0,866840265 | 0,000962901 | chr7 | + |
| Lgi4 | 17,31281404 | 1,718048826 | 0,000257605 | chr7 | + |
| Lipt2 | 60,49747044 | 0,902017955 | 0,009224107 | chr7 | + |
| Megf8 | 33,889825 | 1,117992578 | 0,030188319 | chr7 | + |
| Mfge8 | 301,1619743 | 0,823745236 | 0,016244769 | chr7 | - |
| Mogat2 | 9,731383203 | 1,385644125 | 0,00761647 | chr7 | - |
| Mrgprb2 | 5,327443134 | 1,378359665 | 0,007490652 | chr7 | - |
| Myadm | 446,4066896 | 0,546320712 | 0,023606121 | chr7 | + |
| Neu3 | 46,58023375 | 1,303868838 | 0,000358943 | chr7 | - |
| Pak1 | 103,2840867 | 1,301791285 | 6,53E-06 | chr7 | + |
| Plekha1 | 183,5668923 | 0,551729224 | 0,043954241 | chr7 | + |
| Ric3 | 30,06401847 | 2,010448957 | 1,43E-07 | chr7 | - |
| Rnf141 | 264,0015921 | 1,040464144 | 0,000192616 | chr7 | - |
| Serpinh1 | 464,937713 | 0,674869876 | 0,010046916 | chr7 | - |
| Slc7a9 | 28,86390437 | 2,754850133 | 1,85E-11 | chr7 | + |
| Tarm1 | 3,4128013 | 1,03883376 | 0,047749515 | chr7 | - |
| Tnfrsf22 | 9,863767095 | 1,267505273 | 0,017119669 | chr7 | - |
| Trim21 | 179,6244937 | 0,745334544 | 0,023367724 | chr7 | - |
| Trim30a | 292,7444386 | 1,205670272 | 1,93E-06 | chr7 | - |
| Trim30d | 227,270524 | 1,121492848 | 0,001857909 | chr7 | - |
| Trim68 | 52,24493908 | 1,485750799 | 1,11E-06 | chr7 | - |
| Trpm4 | 24,58571908 | 0,993030371 | 0,047749515 | chr7 | - |
| Ucp2 | 493,4324068 | 0,870850769 | 0,030011851 | chr7 | + |
| Zfp667 | 235,6204116 | 2,418820971 | 2,89E-18 | chr7 | + |
| Zfp719 | 108,0442944 | 0,888657991 | 0,005929858 | chr7 | + |
| Zfp773 | 144,5026741 | 2,29394421 | 1,56E-14 | chr7 | - |
| Zfp78 | 35,14112604 | 1,218722531 | 0,003263682 | chr7 | + |
| Zfp84 | 216,7620301 | 0,865694271 | 0,00015014 | chr7 | + |
| Zfp850 | 47,11409235 | 2,089142379 | 2,84E-09 | chr7 | - |
| Zfp941 | 20,01438055 | 1,358707074 | 0,003442884 | chr7 | - |
| Zik1 | 16,09672215 | 1,050771988 | 0,040364755 | chr7 | - |
| Zscan18 | 41,73221037 | 2,926448244 | 1,09E-14 | chr7 | - |

| | | | | | |
|---|---|---|---|---|---|
| 1700029J07Rik | 14,84314711 | 1,17710994 | 0,021797201 | chr8 | - |
| 4933408N05Rik | 12,1332958 | 1,133297409 | 0,03490297 | chr8 | - |
| Abcc12 | 51,75468922 | 2,644409472 | 1,40E-11 | chr8 | - |
| Atp7b | 141,3061107 | 0,958117312 | 0,001268282 | chr8 | - |
| Atxn1l | 331,7348564 | 0,741148746 | 0,040941746 | chr8 | - |
| Cd209c | 50,21202893 | 1,668581206 | 9,10E-06 | chr8 | - |
| Cdh1 | 1095,461392 | 1,083425756 | 0,000215352 | chr8 | + |
| Champ1 | 113,9934811 | 0,823983967 | 0,006396217 | chr8 | + |
| Cib3 | 6353,340862 | 4,186763309 | 5,95E-40 | chr8 | - |
| Defb1 | 63,34046734 | 1,677164206 | 9,89E-05 | chr8 | + |
| Glg1 | 191,2243168 | 0,658127956 | 0,013235512 | chr8 | - |
| Tarbp1 | 27,60749616 | 0,988486381 | 0,035842812 | chr8 | - |
| Got1l1 | 13,84342624 | 2,036038865 | 8,29E-06 | chr8 | - |
| Hapln4 | 57,8448265 | 1,580305773 | 1,56E-05 | chr8 | + |
| Map10 | 37,63844099 | 2,543876301 | 3,76E-13 | chr8 | + |
| March1 | 56,87226943 | 1,139896043 | 0,000682221 | chr8 | + |
| Mboat4 | 48,0464829 | 1,707887687 | 4,42E-07 | chr8 | + |
| Mlkl | 97,82403905 | 1,29501832 | 0,000209363 | chr8 | - |
| Mmp15 | 689,5166805 | 1,11624893 | 8,01E-05 | chr8 | + |
| Pomk | 128,9599738 | 0,914452863 | 0,021120002 | chr8 | - |
| Slc10a2 | 1058,176483 | 2,833672521 | 4,58E-40 | chr8 | - |
| Slc25a4 | 146,5307814 | 0,784474593 | 0,039523356 | chr8 | - |
| Slc35f3 | 14,93144294 | 2,592444869 | 1,21E-09 | chr8 | + |
| Tlr3 | 112,0310708 | 0,756576396 | 0,024715096 | chr8 | - |
| Tmem184c | 346,257715 | 0,929609553 | 0,000151943 | chr8 | - |
| Tpm4 | 610,5261724 | 0,58229124 | 0,033058809 | chr8 | + |
| Zfp709 | 69,48362564 | 1,027942595 | 0,001196968 | chr8 | + |
| Zfp791 | 81,6620467 | 1,11806017 | 0,004582939 | chr8 | - |
| Zfp882 | 85,02108676 | 1,427477321 | 5,43E-08 | chr8 | + |
| 1700048O20Rik | 116,3722494 | 2,210238649 | 3,55E-10 | chr9 | + |
| 1700057G04Rik | 40,20488909 | 3,045560097 | 1,85E-16 | chr9 | + |
| 1810064F22Rik | 46,57251684 | 1,795683366 | 2,68E-08 | chr9 | - |
| 2010315B03Rik | 373,4848134 | 1,391315604 | 1,29E-07 | chr9 | - |
| 9430037G07Rik | 49,90762201 | 0,852763062 | 0,044362742 | chr9 | - |
| Acrv1 | 3,585446192 | 1,120675056 | 0,031776207 | chr9 | + |
| Anxa2 | 534,7372875 | 0,953010388 | 5,09E-05 | chr9 | + |
| Aph1b | 55,78731516 | 1,150018113 | 0,001769284 | chr9 | - |
| Bbs4 | 115,9919989 | 1,373076191 | 3,85E-05 | chr9 | - |
| Bmp5 | 129,807753 | 0,804491106 | 0,016705702 | chr9 | + |
| Calml4 | 209,4351699 | 2,796234274 | 1,24E-20 | chr9 | + |
| Casp12 | 62,72734218 | 0,805081652 | 0,017119669 | chr9 | + |
| Ccr2 | 81,3207337 | 1,150833024 | 0,002588169 | chr9 | + |
| Ccr5 | 153,6312805 | 0,783753127 | 0,020312936 | chr9 | + |
| Cd276 | 109,6737037 | 0,776509096 | 0,048412358 | chr9 | - |
| Clmp | 15,98172405 | 1,132587221 | 0,028835369 | chr9 | + |
| Cryab | 58,09819191 | 0,811359882 | 0,021120002 | chr9 | + |
| Entpd3 | 11,70361336 | 1,54405538 | 0,001705438 | chr9 | + |
| Fam198a | 311,7733257 | 2,472164996 | 1,28E-16 | chr9 | + |
| Fam81a | 161,9560005 | 3,409032224 | 8,38E-29 | chr9 | - |
| Fbxw16 | 36,85683758 | 3,495434325 | 5,04E-19 | chr9 | - |

| | | | | | |
|---|---|---|---|---|---|
| Fbxw19 | 52,79003255 | 4,001892332 | 9,03E-27 | chr9 | - |
| Ky | 16,28568478 | 2,645382067 | 4,77E-10 | chr9 | + |
| Mmp12 | 30,19562744 | 1,138803941 | 0,03196404 | chr9 | + |
| Nt5e | 148,5644507 | 0,780414655 | 0,016578697 | chr9 | + |
| Parp16 | 1210,226471 | 1,684105675 | 4,85E-08 | chr9 | + |
| Plxnb1 | 763,266751 | 0,747318565 | 0,002587428 | chr9 | + |
| Rab27a | 156,5307513 | 2,122539672 | 3,69E-11 | chr9 | + |
| Rbp1 | 786,1239031 | 1,513788562 | 1,79E-12 | chr9 | + |
| Scn4b | 13,95050652 | 1,795019501 | 0,000154433 | chr9 | + |
| Slc35f2 | 70,45662132 | 3,208698376 | 7,60E-20 | chr9 | + |
| Sorl1 | 17,16302158 | 1,084734868 | 0,043954241 | chr9 | - |
| Stag1 | 508,8382189 | 0,651395012 | 0,005560758 | chr9 | + |
| Tmem45b | 11,34670269 | 1,31598624 | 0,010122278 | chr9 | - |
| Tmie | 224,9792589 | 1,198325155 | 1,78E-06 | chr9 | - |
| Tmprss4 | 15,94388509 | 1,391747581 | 0,00425478 | chr9 | - |
| Uba7 | 218,4773037 | 0,869028269 | 0,004879833 | chr9 | + |
| Zfp105 | 22,00472669 | 1,619982295 | 0,000215477 | chr9 | + |
| Zfp26 | 157,1484861 | 0,753120993 | 0,004030047 | chr9 | - |
| Zfp266 | 409,6475519 | 0,770406129 | 0,016910829 | chr9 | - |
| Zfp426 | 233,4773372 | 0,626362922 | 0,013266719 | chr9 | - |
| Zfp445 | 2463,420299 | 1,748450031 | 1,08E-15 | chr9 | - |
| Zfp558 | 24,85464853 | 1,176144549 | 0,01054773 | chr9 | - |
| Zfp809 | 329,8213172 | 0,660893907 | 0,048412358 | chr9 | + |
| Zfp872 | 21,41180196 | 1,964710354 | 1,18E-05 | chr9 | + |
| Zkscan7 | 152,7933305 | 2,170259175 | 1,09E-09 | chr9 | + |
| 1700025F24Rik | 13,87278831 | 2,116628025 | 3,00E-06 | chr10 | + |
| 1810014B01Rik | 59,27530903 | 0,892813341 | 0,009987469 | chr10 | + |
| Fam241b | 37,79737478 | 0,958086234 | 0,033758254 | chr10 | - |
| 4930459C07Rik | 22,52032298 | 2,02406727 | 9,48E-06 | chr10 | + |
| Acss3 | 198,4577799 | 1,95248095 | 2,46E-12 | chr10 | - |
| Crybg1 | 376,8930577 | 0,845102834 | 0,025898283 | chr10 | - |
| AU041133 | 51,89291148 | 1,344550886 | 0,002292524 | chr10 | + |
| Rdh16f2 | 3042,766015 | 1,071894598 | 0,000269384 | chr10 | + |
| Bicc1 | 17,39072995 | 1,199765956 | 0,015699042 | chr10 | - |
| Cd63 | 266,703791 | 1,329363492 | 3,10E-05 | chr10 | + |
| Cep83 | 196,6318569 | 0,655705586 | 0,038478567 | chr10 | + |
| Cep83os | 48,92333415 | 0,947130276 | 0,025869348 | chr10 | - |
| Dcn | 4885,246593 | 0,705943046 | 0,030059503 | chr10 | + |
| Esr1 | 230,0358323 | 1,109526716 | 0,005923756 | chr10 | + |
| Fam19a2 | 20,09248664 | 1,186631844 | 0,010987722 | chr10 | + |
| Fuca2 | 331,253269 | 1,382214203 | 5,32E-08 | chr10 | + |
| Grip1 | 12,43016164 | 2,27112944 | 3,20E-07 | chr10 | + |
| Hebp2 | 27,57416843 | 1,331422442 | 0,000956991 | chr10 | - |
| Hkdc1 | 4,788474316 | 1,094187147 | 0,048245026 | chr10 | - |
| Ipcef1 | 32,81699463 | 2,932912883 | 3,00E-14 | chr10 | - |
| Lrp11 | 39,13305101 | 0,990994921 | 0,04104012 | chr10 | + |
| Lum | 472,346867 | 0,883693732 | 0,011232874 | chr10 | + |
| Nab2 | 240,9393796 | 0,9227626 | 0,017229721 | chr10 | - |
| Nhsl1 | 170,1896837 | 0,878566385 | 0,007815602 | chr10 | + |
| Rdh5 | 272,6690332 | 0,886325666 | 0,007390385 | chr10 | - |

| | | | | | |
|---|---|---|---|---|---|
| Slc17a8 | 966,6416504 | 1,668039027 | 4,45E-07 | chr10 | - |
| Smim24 | 20,56493692 | 1,480764279 | 0,000959367 | chr10 | + |
| Syt1 | 321,9836776 | 0,627564986 | 0,01279625 | chr10 | - |
| Themis | 8,254344349 | 1,23521744 | 0,019188165 | chr10 | + |
| Unc5b | 82,29441621 | 1,130505003 | 0,000317923 | chr10 | - |
| Zc3h12d | 45,99661543 | 1,182475131 | 0,002156111 | chr10 | + |
| Zfp781 | 5,482333965 | 1,194114708 | 0,026819656 | chr10 | - |
| Zfp873 | 68,65767628 | 1,381209063 | 0,000307558 | chr10 | + |
| Zfp938 | 150,2054405 | 0,991606619 | 0,001100299 | chr10 | - |
| 9930111J21Rik1 | 11,73875209 | 1,182445233 | 0,024491718 | chr11 | - |
| Acaca | 2350,999906 | 0,769546175 | 0,027182749 | chr11 | + |
| Ace | 16,88985686 | 1,146168332 | 0,020449998 | chr11 | + |
| Acly | 4558,523425 | 0,816529499 | 0,047749515 | chr11 | - |
| Actg1 | 2211,775069 | 0,911185323 | 0,001857909 | chr11 | - |
| Arsg | 501,1267152 | 1,04633315 | 0,003747365 | chr11 | + |
| Ccl2 | 22,64169195 | 1,121871351 | 0,016788778 | chr11 | + |
| Cd300lb | 35,12420328 | 0,894574011 | 0,031214237 | chr11 | - |
| Cldn7 | 13,87815581 | 1,200861054 | 0,024615659 | chr11 | + |
| Clint1 | 1079,321702 | 1,345789753 | 1,23E-10 | chr11 | + |
| Cnp | 332,5735153 | 1,039586487 | 0,000194799 | chr11 | + |
| Fancl | 257,6305079 | 0,823494521 | 0,020795773 | chr11 | + |
| Fbxo47 | 12,65678049 | 1,868846652 | 3,58E-05 | chr11 | - |
| Flt4 | 534,9942889 | 1,377584154 | 5,63E-09 | chr11 | + |
| Gal3st1 | 21,54957098 | 1,290431074 | 0,004261355 | chr11 | + |
| Gm11437 | 158,8542403 | 1,686922601 | 1,08E-07 | chr11 | - |
| Gm2a | 4561,636579 | 1,090967261 | 2,85E-08 | chr11 | + |
| Krt23 | 63,93884773 | 1,277058806 | 0,004580356 | chr11 | - |
| Limd2 | 302,797653 | 1,58025749 | 3,63E-13 | chr11 | - |
| Meox1 | 10,48550641 | 1,243457334 | 0,019726376 | chr11 | - |
| Myh10 | 230,0954519 | 0,683543769 | 0,042901157 | chr11 | + |
| Nptx1 | 4,600910495 | 1,31052455 | 0,010137328 | chr11 | - |
| P2rx5 | 4,102151414 | 1,070375827 | 0,048435774 | chr11 | + |
| Plek | 116,3546775 | 1,219212113 | 2,67E-05 | chr11 | - |
| Pnpo | 3423,183362 | 0,571050949 | 0,011535392 | chr11 | - |
| Prkca | 163,5670559 | 0,965128121 | 0,001316719 | chr11 | - |
| Rasl10b | 5,252485746 | 1,147409854 | 0,035374384 | chr11 | + |
| Rpa1 | 746,5814741 | 0,937427361 | 0,000271108 | chr11 | - |
| Rtn4 | 3537,463915 | 1,286593396 | 2,79E-05 | chr11 | + |
| Slc13a5 | 237,8092019 | 2,595002051 | 2,78E-12 | chr11 | - |
| Slc16a5 | 311,3462 | 1,977546208 | 1,53E-15 | chr11 | + |
| Slc43a2 | 216,1536569 | 0,731957385 | 0,027633464 | chr11 | + |
| Slc47a2 | 7,188967537 | 1,713843309 | 0,000342686 | chr11 | - |
| Smtnl2 | 48,3702091 | 1,825825104 | 1,70E-05 | chr11 | - |
| Smyd4 | 53,4542617 | 1,027698318 | 0,004524574 | chr11 | + |
| Spns3 | 6,949640715 | 1,379433045 | 0,007975041 | chr11 | - |
| Sstr2 | 20,43470026 | 1,418473201 | 0,002439392 | chr11 | + |
| Stk10 | 70,04458934 | 0,900381433 | 0,017827372 | chr11 | + |
| Tex19.1 | 5,804820855 | 1,252502999 | 0,016705702 | chr11 | + |
| Tgtp1 | 36,70535605 | 1,181266166 | 0,006928647 | chr11 | - |
| Thra | 77,97389244 | 0,894269568 | 0,01568138 | chr11 | + |

| | | | | | |
|---|---|---|---|---|---|
| 2410018L13Rik | 29,21909471 | 1,227655095 | 0,004864608 | chr12 | - |
| 5730507C01Rik | 440,7917877 | 4,038948551 | 1,25E-58 | chr12 | + |
| 9030624G23Rik | 107,4464786 | 3,673756885 | 1,40E-36 | chr12 | - |
| Wdcp | 53,65716019 | 0,965854568 | 0,005020028 | chr12 | + |
| D430020J02Rik | 80,30933043 | 1,673047701 | 2,86E-06 | chr12 | - |
| Dcaf4 | 41,84922736 | 1,130790624 | 0,013067104 | chr12 | + |
| Dpf3 | 11,69202778 | 1,163014501 | 0,032427477 | chr12 | - |
| G2e3 | 92,69591224 | 0,680900675 | 0,049206241 | chr12 | + |
| Id2 | 2885,869364 | 0,642827051 | 0,044330288 | chr12 | - |
| Ifi27l2b | 155,3074178 | 1,286827884 | 0,001266354 | chr12 | - |
| Flvcr2 | 61,01487651 | 1,336036971 | 3,85E-05 | chr12 | + |
| Rad51b | 125,2404141 | 2,096948915 | 1,20E-08 | chr12 | + |
| Rapgef5 | 70,8179424 | 0,878813364 | 0,009482451 | chr12 | + |
| Rian | 38,74299296 | 2,345569104 | 7,90E-10 | chr12 | + |
| Sdc1 | 3593,691905 | 0,500999464 | 0,048479068 | chr12 | + |
| Serpina3i | 27,76849158 | 1,08091166 | 0,048691753 | chr12 | + |
| Sptb | 23,06200617 | 1,501196876 | 0,000354526 | chr12 | - |
| Sptlc2 | 419,9228603 | 0,547192803 | 0,044732355 | chr12 | - |
| Ston2 | 3,971964188 | 1,207914939 | 0,019277154 | chr12 | - |
| Taf1b | 208,7969982 | 1,015885775 | 0,00168072 | chr12 | + |
| Tdrd9 | 3,213035549 | 1,036471197 | 0,048483358 | chr12 | + |
| Tpo | 75,55565622 | 2,264625916 | 3,16E-10 | chr12 | - |
| 2410141K09Rik | 17,15641487 | 2,46668245 | 5,76E-09 | chr13 | - |
| 4933416M06Rik | 3,396083879 | 1,039333681 | 0,0474782 | chr13 | + |
| 6720489N17Rik | 17,07460903 | 1,494357103 | 0,001034252 | chr13 | - |
| Zfp729b | 182,7921585 | 0,730607715 | 0,014620211 | chr13 | - |
| Adamts6 | 9,044262703 | 1,458881464 | 0,004086561 | chr13 | + |
| BC052688 | 178,406793 | 5,157160868 | 5,25E-53 | chr13 | + |
| Bhmt | 57162,32616 | 0,908896108 | 0,024715096 | chr13 | - |
| Cxcl14 | 27,20322927 | 1,216137729 | 0,009731576 | chr13 | - |
| Eprn | 8,535998213 | 1,180308682 | 0,031250949 | chr13 | - |
| Fam208b | 257,713354 | 0,702646981 | 0,00320933 | chr13 | - |
| Foxq1 | 167,5180594 | 0,94198451 | 0,044732355 | chr13 | + |
| Gm10324 | 11,39904331 | 2,134597966 | 1,98E-06 | chr13 | + |
| Lncenc1 | 426,6475758 | 6,477915735 | 4,22E-93 | chr13 | + |
| Gm3604 | 18,90261422 | 2,029596987 | 4,90E-06 | chr13 | - |
| Golm1 | 262,1763217 | 1,658275879 | 2,74E-12 | chr13 | - |
| Adgrv1 | 1936,968446 | 3,953488661 | 1,58E-55 | chr13 | - |
| Hexb | 436,2350234 | 1,436451363 | 6,52E-11 | chr13 | - |
| Inhba | 40,88961014 | 1,556423544 | 5,52E-05 | chr13 | + |
| Klf6 | 220,7125639 | 0,671018326 | 0,014243688 | chr13 | + |
| Mtr | 77,17203399 | 0,933960833 | 0,003981116 | chr13 | - |
| Naip2 | 179,1540499 | 1,492791419 | 6,97E-06 | chr13 | - |
| Nhlrc1 | 18,29753374 | 1,248160999 | 0,0063705 | chr13 | - |
| Nid1 | 364,2226965 | 0,759702821 | 0,017102722 | chr13 | + |
| Ocln | 740,3389191 | 1,332693981 | 4,57E-05 | chr13 | - |
| Ptch1 | 67,01973686 | 1,062855042 | 0,012896621 | chr13 | - |
| Rpp40 | 1776,896822 | 3,475656289 | 8,90E-41 | chr13 | - |
| Rsl1 | 51,83738136 | 0,947099778 | 0,005354992 | chr13 | + |
| Rslcan18 | 54,2397273 | 0,818102929 | 0,034199883 | chr13 | - |

| | | | | | |
|---|---|---|---|---|---|
| Serpinb1a | 53,98306327 | 1,009379819 | 0,018716187 | chr13 | - |
| Tpmt | 2547,064915 | 0,958502386 | 0,001919571 | chr13 | - |
| Ube2ql1 | 5,669362357 | 1,309881262 | 0,013065175 | chr13 | - |
| Xrcc4 | 219,3986791 | 0,843963282 | 0,011768776 | chr13 | - |
| Zfp369 | 22,2783221 | 1,401899641 | 0,001857909 | chr13 | + |
| Zfp72 | 32,47368126 | 1,038706932 | 0,011232874 | chr13 | - |
| Zfp759 | 73,27401332 | 0,880479914 | 0,012974293 | chr13 | + |
| Zfp808 | 74,09740682 | 2,923741867 | 9,85E-18 | chr13 | + |
| Zfp934 | 19,98414162 | 2,405897132 | 4,06E-09 | chr13 | - |
| Zfp935 | 111,6025942 | 1,220585745 | 0,00015014 | chr13 | - |
| Zkscan3 | 598,8586215 | 0,677002701 | 0,018208572 | chr13 | - |
| Zkscan4 | 16,13374224 | 1,07822733 | 0,035665157 | chr13 | + |
| Zkscan8 | 123,4194956 | 0,750722051 | 0,014463104 | chr13 | - |
| Ajuba | 76,33752105 | 0,837598755 | 0,031698865 | chr14 | - |
| Anxa8 | 12,52186465 | 2,112647231 | 2,86E-06 | chr14 | + |
| Ccdc122 | 70,21429803 | 0,761311531 | 0,030855688 | chr14 | + |
| Diaph3 | 266,0748217 | 5,675753821 | 3,17E-71 | chr14 | - |
| Grid1 | 101,8713019 | 4,083147192 | 7,55E-39 | chr14 | + |
| Il17rb | 238,7251487 | 0,99107126 | 0,000155071 | chr14 | - |
| Il3ra | 21,68536119 | 1,282129448 | 0,012926258 | chr14 | + |
| Kctd12 | 287,2650591 | 0,675493884 | 0,026467698 | chr14 | - |
| Micu2 | 1502,540326 | 1,116148711 | 3,94E-07 | chr14 | - |
| Nuggc | 15,51216038 | 2,510277914 | 3,69E-09 | chr14 | + |
| Parp2 | 134,0499373 | 0,772393951 | 0,03437957 | chr14 | + |
| Pck2 | 66,11264918 | 0,884933319 | 0,006351835 | chr14 | + |
| Ppp3cc | 52,30532095 | 0,979493107 | 0,02272099 | chr14 | - |
| Rnf17 | 32,39500665 | 3,253762025 | 3,97E-17 | chr14 | + |
| Sftpd | 42,96039448 | 2,229490203 | 9,46E-10 | chr14 | - |
| Tgm1 | 339,1681656 | 0,963697624 | 0,006172818 | chr14 | - |
| 2610037D02Rik | 25,02064112 | 2,277933068 | 6,09E-09 | chr15 | - |
| 9030619P08Rik | 628,404009 | 1,428487683 | 2,48E-06 | chr15 | - |
| Abcd2 | 679,8471497 | 1,665130238 | 6,02E-07 | chr15 | - |
| Alg10b | 287,2485807 | 0,781638775 | 0,028367787 | chr15 | + |
| Amigo2 | 182,2350313 | 0,627222745 | 0,033496932 | chr15 | - |
| Angpt1 | 19,18701537 | 1,031378641 | 0,046603699 | chr15 | - |
| Anxa13 | 46,48364524 | 1,128270839 | 0,007909756 | chr15 | - |
| Capsl | 15,30062353 | 1,031449982 | 0,046513257 | chr15 | + |
| Col14a1 | 797,3023989 | 0,602075737 | 0,047921745 | chr15 | + |
| Colec10 | 794,3796003 | 0,699177636 | 0,030454862 | chr15 | + |
| Csf2rb | 313,6066278 | 1,773400693 | 2,34E-08 | chr15 | + |
| Cyp2d12 | 42,38550344 | 1,37837723 | 0,000468196 | chr15 | + |
| Dnah5 | 28,10647318 | 2,508550705 | 4,06E-09 | chr15 | + |
| Fam83a | 23,19855076 | 1,091242139 | 0,022611041 | chr15 | + |
| Krt7 | 17,06466418 | 1,028028631 | 0,042487153 | chr15 | + |
| Lmbrd2 | 192,8673006 | 1,071813918 | 0,003922197 | chr15 | + |
| Ly6a | 1178,147825 | 2,019317638 | 6,60E-10 | chr15 | - |
| Ly6d | 48,83933077 | 1,469938039 | 0,000242589 | chr15 | - |
| Ly6e | 6314,784266 | 1,099509802 | 5,79E-05 | chr15 | + |
| Mettl7a3 | 52,48353913 | 3,149792268 | 6,24E-18 | chr15 | + |
| Miox | 6,781765377 | 1,594493384 | 0,001202226 | chr15 | + |

| | | | | | |
|---|---|---|---|---|---|
| Ncald | 205,1596863 | 0,764542235 | 0,034199883 | chr15 | - |
| Oxct1 | 159,0400031 | 0,878767505 | 0,002073958 | chr15 | + |
| Pmm1 | 154,9264984 | 1,929324791 | 2,81E-10 | chr15 | - |
| Prlr | 2281,983076 | 0,906275794 | 0,005937526 | chr15 | + |
| Rabl2 | 54,42780903 | 1,009220239 | 0,003950503 | chr15 | - |
| Sntb1 | 271,2881776 | 1,053439853 | 0,001487543 | chr15 | - |
| Sult4a1 | 4,845560391 | 1,230014201 | 0,019665653 | chr15 | - |
| Sybu | 18,64265449 | 2,616412774 | 3,08E-10 | chr15 | - |
| Tnfrsf11b | 100,9832812 | 1,251320127 | 8,04E-06 | chr15 | - |
| Trmt12 | 68,83533698 | 0,961223839 | 0,03196404 | chr15 | + |
| 2010005H15Rik | 221,0854256 | 4,182953112 | 1,92E-52 | chr16 | + |
| Alcam | 1366,782222 | 0,663603316 | 0,0073558 | chr16 | - |
| Cldn1 | 1487,751069 | 1,08906947 | 4,06E-09 | chr16 | - |
| Cxadr | 2494,871474 | 0,782214141 | 0,000153705 | chr16 | + |
| Emp2 | 343,4089943 | 1,045596961 | 9,04E-06 | chr16 | - |
| Fstl1 | 201,6476071 | 0,685041138 | 0,030552507 | chr16 | + |
| Gm609 | 5,109703402 | 1,240361236 | 0,018436179 | chr16 | - |
| Hrasls | 22,09811466 | 2,92881465 | 1,33E-12 | chr16 | + |
| Iqcg | 49,72034151 | 0,930647042 | 0,024390594 | chr16 | - |
| Ncam2 | 39,90303338 | 2,29606616 | 2,23E-09 | chr16 | + |
| Nmral1 | 249,7951661 | 0,929060364 | 0,001007069 | chr16 | - |
| Poglut1 | 342,1552461 | 0,669242175 | 0,014941851 | chr16 | - |
| Ppl | 249,9728295 | 1,225225807 | 8,95E-07 | chr16 | - |
| Retnlg | 12,14298122 | 1,347769652 | 0,009894756 | chr16 | + |
| Robo1 | 136,6162096 | 1,771899444 | 6,93E-09 | chr16 | + |
| Senp7 | 145,7573018 | 1,453738225 | 8,93E-09 | chr16 | + |
| Slc7a4 | 25,27342181 | 1,476461249 | 0,000693733 | chr16 | - |
| Spidr | 118,639591 | 0,79267102 | 0,049461601 | chr16 | - |
| Tmem45a | 24,77104325 | 0,95675765 | 0,042803116 | chr16 | - |
| Tmprss2 | 99,03451139 | 1,066424402 | 0,000638626 | chr16 | - |
| Ttc3 | 97,34930584 | 0,939958699 | 0,001857909 | chr16 | + |
| Zfp995 | 53,86094861 | 1,160868032 | 0,003386635 | chr17 | - |
| Btnl4 | 5,972755189 | 1,144104364 | 0,034522477 | chr17 | - |
| Btnl5-ps | 7,573243256 | 1,821167309 | 0,000116264 | chr17 | - |
| Btnl6 | 6,499454514 | 1,628843206 | 0,000795611 | chr17 | - |
| C4a | 427,7813857 | 1,404880496 | 0,000251317 | chr17 | - |
| H2-T-ps | 60,48252261 | 2,895851155 | 4,04E-20 | chr17 | - |
| Cdkn1a | 49,58535127 | 1,118069387 | 0,007815602 | chr17 | + |
| Cldn6 | 12,8061472 | 1,16082866 | 0,026819656 | chr17 | + |
| Clic5 | 74,86918211 | 1,297220361 | 0,008131231 | chr17 | + |
| Clip4 | 23,85231017 | 1,65382036 | 0,00011082 | chr17 | + |
| Crisp2 | 4,696731943 | 1,367642028 | 0,006907987 | chr17 | - |
| Cyp4f16 | 101,6811991 | 1,627656329 | 1,36E-09 | chr17 | + |
| Dnph1 | 167,6510353 | 0,567231929 | 0,045665077 | chr17 | + |
| Gm1976 | 31,99998719 | 1,741085953 | 2,45E-05 | chr17 | - |
| Gm20939 | 130,9349752 | 1,207602537 | 0,00046773 | chr17 | + |
| Zfp994 | 299,641859 | 0,87080886 | 0,00062456 | chr17 | - |
| Gm9992 | 13,46311385 | 1,575037819 | 0,000961755 | chr17 | - |
| H2-Bl | 42,08661874 | 2,154228926 | 4,57E-09 | chr17 | - |
| H2-Q1 | 546,9206191 | 2,263108607 | 7,07E-13 | chr17 | + |

| | | | | | |
|---|---|---|---|---|---|
| H2-Q2 | 24,37739861 | 1,257986136 | 0,005845973 | chr17 | + |
| H2-Q4 | 482,5396911 | 0,609934435 | 0,013576999 | chr17 | + |
| H2-T10 | 40,02452281 | 1,016829124 | 0,008977697 | chr17 | - |
| Jpt2 | 618,7111943 | 0,871331376 | 0,000725735 | chr17 | - |
| Hsf2bp | 34,66485841 | 2,916138887 | 8,83E-14 | chr17 | - |
| Lbh | 140,8280637 | 0,898604818 | 0,004109461 | chr17 | + |
| Lnpep | 81,26998919 | 1,030308218 | 0,004030047 | chr17 | - |
| Mas1 | 38,51860408 | 2,935083605 | 1,54E-12 | chr17 | - |
| Meiob | 294,6836169 | 1,272370387 | 0,000795611 | chr17 | + |
| Mettl4 | 100,9264693 | 0,964647663 | 0,004372535 | chr17 | - |
| Prss41 | 4,577455238 | 1,245334227 | 0,015256665 | chr17 | - |
| Psmb9 | 638,569947 | 1,243984824 | 3,37E-06 | chr17 | - |
| Qpct | 521,8114326 | 1,002634259 | 0,001492349 | chr17 | + |
| Slc22a7 | 961,9869562 | 0,719580837 | 0,030188319 | chr17 | - |
| Slc8a1 | 36,5456645 | 1,084627036 | 0,005020028 | chr17 | - |
| Srd5a2 | 19,53006899 | 1,005744704 | 0,038313544 | chr17 | - |
| Tap1 | 300,6208513 | 0,807057249 | 0,003255089 | chr17 | + |
| Tff2 | 10,14429785 | 1,208235384 | 0,025287128 | chr17 | - |
| Tff3 | 473,5279157 | 1,864704849 | 2,53E-11 | chr17 | - |
| Ubd | 46,21449363 | 1,913283071 | 7,73E-07 | chr17 | + |
| Vmn2r-ps129 | 26,7302955 | 1,070046366 | 0,033460797 | chr17 | + |
| Zfp13 | 29,96048384 | 2,548051706 | 3,44E-11 | chr17 | - |
| Zfp229 | 100,7884641 | 0,931390024 | 0,003651689 | chr17 | + |
| Zfp53 | 126,2439286 | 1,31238897 | 2,62E-07 | chr17 | + |
| Zfp563 | 121,1330858 | 0,820699988 | 0,003454129 | chr17 | + |
| Zfp677 | 44,33257068 | 2,602479382 | 3,74E-13 | chr17 | + |
| Zfp763 | 104,0419361 | 2,085710822 | 6,50E-14 | chr17 | - |
| Zfp799 | 63,17174472 | 0,905898179 | 0,007053313 | chr17 | - |
| Zfp870 | 122,9126837 | 0,80892586 | 0,038283691 | chr17 | - |
| Zfp947 | 78,62483164 | 1,041191377 | 0,00071662 | chr17 | - |
| Zfp952 | 122,4847631 | 0,908892317 | 0,001927729 | chr17 | + |
| Zfp955a | 154,93934 | 1,060114454 | 6,29E-05 | chr17 | - |
| Zfp955b | 99,91178922 | 1,091009581 | 0,001539492 | chr17 | + |
| Jcad | 146,9444131 | 1,296190961 | 1,28E-08 | chr18 | + |
| Adnp2 | 130,8166864 | 1,097166957 | 6,96E-06 | chr18 | - |
| Cables1 | 53,45775535 | 1,42366376 | 0,000442223 | chr18 | + |
| Cfap53 | 120,4646663 | 4,044576725 | 8,84E-30 | chr18 | + |
| Ecscr | 113,5567772 | 0,663690015 | 0,02742883 | chr18 | - |
| Greb1l | 18,88138178 | 1,381039316 | 0,003393958 | chr18 | + |
| Mbd1 | 2717,088182 | 2,156485076 | 1,09E-14 | chr18 | + |
| Myo7b | 3,743352054 | 1,111085829 | 0,032846126 | chr18 | - |
| Pcdhga3 | 6,383314916 | 1,193434886 | 0,027954357 | chr18 | + |
| Ppic | 129,5272659 | 0,955904938 | 0,00247095 | chr18 | - |
| Prelid2 | 49,0316917 | 2,673128608 | 5,44E-13 | chr18 | - |
| Pstpip2 | 187,0908066 | 0,813606167 | 0,00393493 | chr18 | + |
| Rprd1a | 605,4651026 | 1,112307617 | 2,48E-06 | chr18 | - |
| Slc4a9 | 41,23049218 | 1,517689512 | 0,000155769 | chr18 | + |
| Spink5 | 65,34908972 | 2,842090151 | 1,65E-16 | chr18 | + |
| Taf7 | 134,0589293 | 0,789741067 | 0,03196404 | chr18 | - |
| Tmx3 | 435,8638823 | 0,618255603 | 0,029831049 | chr18 | + |

| | | | | | |
|---|---|---|---|---|---|
| Zfp608 | 26,32119621 | 1,013721126 | 0,024221418 | chr18 | - |
| Aldh3b3 | 9,383929772 | 1,931104015 | 3,08E-05 | chr19 | + |
| Ankrd1 | 16,66256093 | 1,270349374 | 0,01054773 | chr19 | - |
| Asrgl1 | 209,5524953 | 0,769082776 | 0,0348796 | chr19 | - |
| Gstp3 | 1162,935809 | 1,91063053 | 5,31E-10 | chr19 | - |
| Blnk | 88,00978817 | 1,161637641 | 0,01040626 | chr19 | - |
| Dntt | 92,61162153 | 1,923003872 | 1,10E-10 | chr19 | + |
| Dtx4 | 163,9042905 | 0,807298531 | 0,018707375 | chr19 | - |
| E030003E18Rik | 25,03156092 | 1,014663689 | 0,034963938 | chr19 | + |
| Ehd1 | 466,214831 | 0,550710857 | 0,01048884 | chr19 | + |
| Fads3 | 58,27685632 | 1,245314843 | 0,000561937 | chr19 | + |
| Fam111a | 101,3478187 | 0,742913781 | 0,014620211 | chr19 | + |
| Gm4952 | 3110,446904 | 0,763908313 | 0,011248868 | chr19 | + |
| Grk5 | 44,80133911 | 0,910026226 | 0,024054827 | chr19 | + |
| Gsto2 | 8,878049103 | 1,248793252 | 0,019739765 | chr19 | + |
| Hectd2 | 96,22962713 | 2,089959641 | 1,21E-14 | chr19 | + |
| Ifit3b | 114,5101013 | 0,921621183 | 0,010780375 | chr19 | + |
| Ifit2 | 209,085211 | 1,089465551 | 3,82E-05 | chr19 | + |
| Il33 | 71,35334495 | 1,030623944 | 0,015743368 | chr19 | + |
| Lipo1 | 132,5895001 | 2,188185809 | 2,56E-17 | chr19 | - |
| Mpeg1 | 2232,256083 | 0,992179585 | 2,30E-06 | chr19 | + |
| Papss2 | 3622,227476 | 0,708908414 | 0,034522477 | chr19 | + |
| Pnliprp1 | 10,78775594 | 1,724825204 | 0,000314559 | chr19 | + |
| Slc15a3 | 167,2537542 | 0,662769289 | 0,0386619 | chr19 | + |
| Slc22a26 | 64,26272195 | 1,73310502 | 8,08E-05 | chr19 | - |
| Slc22a27 | 175,5199579 | 1,875774027 | 5,72E-05 | chr19 | - |
| Smc3 | 74,28126247 | 0,8250081 | 0,008072797 | chr19 | + |
| Stambpl1 | 22,63577538 | 1,554350462 | 0,000298384 | chr19 | + |
| Zfp518a | 357,4051255 | 1,98330401 | 2,72E-17 | chr19 | + |
| Arhgef9 | 81,36274789 | 1,138527286 | 0,000596282 | chrX | - |
| Bex1 | 11,20042115 | 1,60314948 | 0,001083279 | chrX | - |
| Brwd3 | 73,16368135 | 0,915626045 | 0,00605445 | chrX | - |
| Capn6 | 17,38576737 | 1,335594219 | 0,005540839 | chrX | - |
| Ccdc120 | 20,86080882 | 1,113033453 | 0,040095945 | chrX | - |
| Ctps2 | 372,8689604 | 1,124004796 | 0,000119831 | chrX | + |
| D330045A20Rik | 36,75499426 | 3,374928154 | 1,31E-18 | chrX | + |
| Dcaf12l1 | 85,49005985 | 0,641443347 | 0,044683691 | chrX | - |
| Dmd | 553,6401887 | 0,788679328 | 0,01093117 | chrX | + |
| Fmr1nb | 7,008965193 | 1,394779559 | 0,006233369 | chrX | + |
| G6pdx | 142,267533 | 0,988995684 | 0,001032732 | chrX | - |
| Klhl13 | 222,8809467 | 1,180774598 | 6,96E-06 | chrX | - |
| Mid1ip1 | 507,2460872 | 0,659594272 | 0,042188881 | chrX | + |
| P2ry4 | 69,98349491 | 0,86661396 | 0,046650967 | chrX | - |
| Pola1 | 129,2099582 | 0,830649451 | 0,011585744 | chrX | - |
| Prrg1 | 27,52373467 | 1,319765873 | 0,002622865 | chrX | - |
| Rhox5 | 18,97401297 | 1,925054754 | 1,22E-05 | chrX | + |
| Rps6ka3 | 268,7824819 | 0,839866427 | 0,001342019 | chrX | + |
| S100g | 10,78227043 | 1,22538966 | 0,020728549 | chrX | - |
| Sytl5 | 15,69025604 | 1,52164249 | 0,002145277 | chrX | + |
| Tceal8 | 746,822696 | 0,864837802 | 0,002069342 | chrX | - |

| | | | | | |
|---|---|---|---|---|---|
| Timp1 | 19,70184499 | 1,368027995 | 0,007909756 | chrX | + |
| Tlr7 | 86,4919995 | 0,667461657 | 0,044481388 | chrX | - |
| Tlr8 | 34,4785309 | 1,040212664 | 0,025808311 | chrX | - |
| Xlr3b | 8,283125138 | 1,544439717 | 0,001927729 | chrX | + |
| Zfp449 | 14,95548955 | 1,329241891 | 0,006894028 | chrX | + |
| Zmat1 | 106,0103339 | 0,738494248 | 0,019277154 | chrX | - |
| 1700066M21Rik | 538,4111201 | -0,644110009 | 0,005533151 | Chr1 | + |
| Abcb6 | 896,1789315 | -0,91246533 | 1,12E-05 | Chr1 | - |
| Aox1 | 1356,637989 | -0,598215697 | 0,022611041 | Chr1 | + |
| C730036E19Rik | 520,277397 | -1,16398717 | 0,004151737 | Chr1 | + |
| Coa5 | 1441,660602 | -0,454871033 | 0,047966165 | Chr1 | - |
| Creg1 | 4359,352559 | -0,915064517 | 0,001915773 | Chr1 | + |
| Cyp27a1 | 5717,527997 | -0,829696885 | 1,30E-05 | Chr1 | + |
| Ephx1 | 6008,228483 | -0,523571558 | 0,023226367 | Chr1 | - |
| Gm4956 | 178,493242 | -2,838447749 | 8,63E-21 | Chr1 | - |
| Grem2 | 256,0683883 | -1,102587291 | 0,001568497 | Chr1 | - |
| Hes6 | 1122,00645 | -0,948195129 | 0,000384527 | Chr1 | - |
| Hjurp | 149,1690336 | -0,96758871 | 0,001198245 | Chr1 | - |
| Ildr2 | 384,5264691 | -0,941373725 | 0,002145277 | Chr1 | + |
| Ilkap | 445,4364392 | -0,576990191 | 0,016705702 | Chr1 | - |
| Ivns1abp | 2703,432373 | -0,637930894 | 0,021610664 | Chr1 | + |
| Marc1 | 5892,110815 | -0,531560298 | 0,016732031 | Chr1 | - |
| Mpp4 | 1298,441873 | -0,540448456 | 0,04056125 | Chr1 | - |
| Ndufb3 | 2203,48064 | -0,660961334 | 0,026272185 | Chr1 | + |
| Ogfrl1 | 215,1450928 | -0,776667801 | 0,009803754 | Chr1 | - |
| Pcp4l1 | 102,604881 | -1,701834365 | 3,98E-09 | Chr1 | - |
| Ptp4a1 | 760,8921396 | -1,028640971 | 0,000269384 | Chr1 | - |
| Raph1 | 861,2511833 | -0,607697169 | 0,044732355 | Chr1 | - |
| Rnf152 | 360,3569034 | -0,98954671 | 0,00068116 | Chr1 | - |
| Serpine2 | 33,41144841 | -0,967417625 | 0,039399445 | Chr1 | - |
| Susd4 | 57,72751939 | -4,185333375 | 4,58E-33 | Chr1 | + |
| Tcf24 | 119,7530828 | -0,983143741 | 0,030939931 | Chr1 | - |
| Ube2f | 616,006146 | -0,472469335 | 0,046298439 | Chr1 | + |
| Ugt1a1 | 1871,552677 | -0,69834176 | 0,030454862 | Chr1 | + |
| Ugt1a9 | 413,6948222 | -1,608972429 | 1,69E-07 | Chr1 | + |
| Zfand2b | 324,6414625 | -0,819043249 | 0,003163564 | Chr1 | + |
| 1110008F13Rik | 1583,950577 | -0,787642872 | 0,000908604 | chr2 | + |
| 6430550D23Rik | 14,9216654 | -1,055247466 | 0,03196404 | chr2 | - |
| Ahcy | 37510,37122 | -0,510232408 | 0,013670896 | chr2 | - |
| Bdnf | 19,36122137 | -1,273228127 | 0,005018061 | chr2 | + |
| Bmp7 | 30,06112954 | -0,88683366 | 0,042234184 | chr2 | - |
| C8g | 11292,22827 | -0,58682036 | 0,019663776 | chr2 | - |
| Cebpb | 206,1531011 | -0,872444664 | 0,000298384 | chr2 | + |
| Celf2 | 118,9605397 | -0,930235645 | 0,001115822 | chr2 | - |
| Cobll1 | 496,7415725 | -0,639989719 | 0,024454584 | chr2 | - |
| Ctcflos | 52,7739932 | -1,347323382 | 8,08E-05 | chr2 | + |
| Mindy3 | 516,6314897 | -0,570587131 | 0,014565073 | chr2 | - |
| Gm14057 | 22,27032818 | -1,165217926 | 0,025822683 | chr2 | - |
| Gnas | 4263,339476 | -0,81260843 | 0,000124884 | chr2 | + |
| H13 | 1618,349838 | -0,620770455 | 0,011960997 | chr2 | + |

| | | | | | |
|---|---|---|---|---|---|
| Mcm10 | 1605,6551 | -1,226243171 | 6,77E-07 | chr2 | - |
| Meig1 | 8,236144787 | -1,097798973 | 0,049992049 | chr2 | - |
| Nop56 | 403,4524328 | -0,624755143 | 0,003922197 | chr2 | + |
| Ola1 | 716,2631354 | -0,532618867 | 0,011041098 | chr2 | - |
| Phyh | 30497,55648 | -0,727334462 | 1,30E-05 | chr2 | + |
| Phyhd1 | 1199,116898 | -0,532450802 | 0,008098959 | chr2 | + |
| Rab22a | 260,0719441 | -0,546646049 | 0,0348796 | chr2 | + |
| Rpl35 | 5579,612292 | -0,550547872 | 0,034963938 | chr2 | - |
| Sdccag3 | 418,5814419 | -0,595578198 | 0,038403403 | chr2 | - |
| Slc27a2 | 20045,56199 | -0,684413865 | 0,017105098 | chr2 | + |
| Sp5 | 4,851568089 | -1,513694978 | 0,001915773 | chr2 | + |
| Spc25 | 170,5006772 | -0,78985868 | 0,047966165 | chr2 | - |
| Srsf6 | 1102,418921 | -0,512931997 | 0,025808311 | chr2 | + |
| Strbp | 643,6659897 | -0,498892401 | 0,026124007 | chr2 | - |
| Tmx4 | 471,2790651 | -0,892045611 | 0,001988185 | chr2 | - |
| Ubac1 | 274,5633413 | -0,543425518 | 0,04378035 | chr2 | - |
| Vapb | 1031,587829 | -0,585229695 | 0,019869728 | chr2 | + |
| Zc3h8 | 83,36336563 | -0,729924976 | 0,030454862 | chr2 | - |
| Atp1a1 | 4832,202634 | -0,863851415 | 2,66E-06 | chr3 | - |
| Etnppl | 2044,480906 | -1,207206791 | 0,00010504 | chr3 | + |
| Gstm1 | 25979,90237 | -0,571109227 | 0,01070874 | chr3 | - |
| Hsd3b5 | 3765,837244 | -1,385859701 | 0,005687763 | chr3 | - |
| Larp1b | 1271,667748 | -0,453790392 | 0,032846126 | chr3 | + |
| Mme | 332,0678451 | -1,005438687 | 0,002500373 | chr3 | + |
| P2ry1 | 125,0450409 | -0,781594698 | 0,008957506 | chr3 | + |
| Pla2g12a | 176,0599251 | -0,978783111 | 0,000302628 | chr3 | + |
| Rarres1 | 586,5012731 | -2,588203394 | 1,25E-14 | chr3 | - |
| Sh3d19 | 975,8934199 | -0,491853861 | 0,032791549 | chr3 | + |
| Syde2 | 85,22154042 | -0,729903244 | 0,025808311 | chr3 | + |
| Tsacc | 78,06476847 | -0,723272343 | 0,037533773 | chr3 | - |
| Tsc22d2 | 423,6944948 | -0,622399621 | 0,037533773 | chr3 | + |
| Uox | 59043,10444 | -0,591011395 | 0,017102722 | chr3 | + |
| 1110065P20Rik | 147,8682338 | -0,831333072 | 0,026733412 | chr4 | - |
| AI314180 | 1439,860548 | -0,480534477 | 0,037117133 | chr4 | - |
| Akr1a1 | 10719,87188 | -0,45462316 | 0,031951285 | chr4 | - |
| Baat | 4701,103712 | -0,707896278 | 0,023606121 | chr4 | - |
| Bag1 | 1152,788264 | -0,631003807 | 0,019663776 | chr4 | - |
| Car8 | 1332,834167 | -0,660437436 | 0,008223807 | chr4 | - |
| Col27a1 | 308,754073 | -0,746271636 | 0,035724719 | chr4 | + |
| Echdc2 | 1876,403409 | -1,138383023 | 0,00139651 | chr4 | + |
| Errfi1 | 12744,77272 | -0,765612305 | 0,046603699 | chr4 | + |
| Gba2 | 117,2823393 | -0,848199673 | 0,013565256 | chr4 | - |
| Gm11837 | 29,9973304 | -1,109308431 | 0,009685588 | chr4 | + |
| Mrpl20 | 1183,324984 | -0,65604746 | 0,017949633 | chr4 | + |
| Mup1 | 3029,816996 | -2,226987385 | 6,44E-07 | chr4 | - |
| Mup12 | 633,5208724 | -1,162270226 | 0,00828198 | chr4 | - |
| Mup13 | 155,9968217 | -2,112522679 | 3,00E-14 | chr4 | - |
| Mup14 | 5236,828132 | -1,26560005 | 0,017500348 | chr4 | - |
| Mup3 | 226720,6369 | -1,209577798 | 8,06E-06 | chr4 | - |
| Mup5 | 317,2788623 | -1,217691693 | 0,000192616 | chr4 | - |

| | | | | | |
|---|---|---|---|---|---|
| Mup6 | 317,2949593 | -1,382562994 | 0,00011349 | chr4 | + |
| Mup7 | 66784,96087 | -2,677886781 | 5,36E-10 | chr4 | - |
| Mup8 | 138,7101537 | -1,199459609 | 0,008370287 | chr4 | - |
| Mup9 | 697,1522412 | -1,772824569 | 0,000194995 | chr4 | - |
| Paqr7 | 926,9943208 | -0,868524364 | 0,002455251 | chr4 | + |
| Pithd1 | 272,5765026 | -0,633379588 | 0,017119669 | chr4 | - |
| Rcc2 | 282,1605626 | -0,700056004 | 0,045207748 | chr4 | + |
| Rpl11 | 7046,793009 | -0,595773818 | 0,012626346 | chr4 | - |
| Rps6 | 9695,433416 | -0,545415035 | 0,006385096 | chr4 | - |
| Rps8 | 6463,737181 | -0,554863766 | 0,005314525 | chr4 | - |
| Slc25a51 | 1321,971094 | -0,551332303 | 0,034199883 | chr4 | - |
| Tmem57 | 778,4759779 | -0,606779542 | 0,031214237 | chr4 | - |
| Ubap1 | 596,5111698 | -0,707262063 | 0,005928082 | chr4 | + |
| Vcp | 6147,345213 | -0,519024539 | 0,030454862 | chr4 | - |
| 9130230L23Rik | 3,279959445 | -1,139042563 | 0,026893975 | chr5 | - |
| Atp5j2 | 1073,898469 | -0,644279935 | 0,013620546 | chr5 | - |
| Brap | 1646,915599 | -0,690822954 | 0,036050684 | chr5 | + |
| Crygn | 16,55952544 | -1,924058774 | 5,65E-06 | chr5 | - |
| Cyp3a25 | 12340,02059 | -0,849626141 | 0,002215245 | chr5 | - |
| Cyp3a59 | 1006,03015 | -2,154456521 | 3,62E-17 | chr5 | + |
| Dancr | 34,71748176 | -1,122206734 | 0,015737606 | chr5 | + |
| Fras1 | 40,10602527 | -1,271903359 | 0,003400929 | chr5 | + |
| Gm16063 | 4,245237181 | -1,427468271 | 0,003255089 | chr5 | + |
| Gtf2ird2 | 60,86516459 | -1,001808939 | 0,008956983 | chr5 | + |
| Gtf3a | 117,9218407 | -0,637744656 | 0,030459905 | chr5 | + |
| Hopx | 1099,143009 | -0,723437858 | 0,006218311 | chr5 | - |
| Hpd | 25785,81549 | -0,462573348 | 0,044747609 | chr5 | - |
| Hspb1 | 87,62420705 | -1,550258397 | 0,00032529 | chr5 | + |
| Iscu | 955,7956002 | -0,443041172 | 0,042487153 | chr5 | + |
| Mapkapk5 | 431,1158311 | -0,50496884 | 0,034199883 | chr5 | - |
| Mn1 | 85,44507389 | -2,598158798 | 1,11E-16 | chr5 | + |
| Mospd3 | 181,9457137 | -0,735655888 | 0,043912298 | chr5 | - |
| Noc4l | 102,730026 | -0,651264604 | 0,040670349 | chr5 | - |
| Polr1d | 1091,851096 | -0,538068087 | 0,021296181 | chr5 | + |
| Pop5 | 334,3852836 | -0,681550505 | 0,014755854 | chr5 | + |
| Psmd9 | 577,5444168 | -0,903107973 | 0,002206469 | chr5 | + |
| Rplp0 | 7508,335091 | -0,661861129 | 0,006862198 | chr5 | + |
| Slc46a3 | 671,755421 | -0,758790095 | 0,02233571 | chr5 | - |
| Suds3 | 1138,714637 | -0,706168333 | 0,009731576 | chr5 | - |
| Ugdh | 2597,984842 | -0,660431537 | 0,002015907 | chr5 | - |
| Ugt2b1 | 16820,65322 | -1,01000218 | 7,57E-05 | chr5 | - |
| Ugt2b35 | 4641,067886 | -0,777197996 | 0,026819656 | chr5 | + |
| Ugt2b5 | 15617,53631 | -1,204264075 | 7,55E-06 | chr5 | - |
| Ung | 114,7905694 | -0,908719773 | 0,031334451 | chr5 | + |
| Aass | 4694,947856 | -0,46642035 | 0,035622397 | chr6 | - |
| Abcg2 | 2103,458892 | -0,753722604 | 0,017119669 | chr6 | + |
| Asns | 78,4254407 | -1,057151955 | 0,003109254 | chr6 | - |
| A2ml1 | 891,0928346 | -0,747157766 | 0,007884642 | chr6 | - |
| Cbx3 | 1091,284842 | -0,677564773 | 0,005540839 | chr6 | + |
| Cmas | 660,5969675 | -0,525201998 | 0,021120002 | chr6 | + |

| | | | | | |
|---|---|---|---|---|---|
| Fabp1 | 185249,4478 | -0,92549447 | 0,018952788 | chr6 | + |
| Gm10319 | 877,5886013 | -0,719612489 | 0,031555845 | chr6 | + |
| Gm20594 | 1767,508153 | -0,894721328 | 0,04255442 | chr6 | + |
| Grm8 | 8,903557837 | -1,495605142 | 0,002744629 | chr6 | - |
| Itpr1 | 640,7116591 | -0,55772336 | 0,0065641 | chr6 | + |
| Itpr2 | 457,5741279 | -0,735957199 | 0,000209903 | chr6 | - |
| Kcmf1 | 722,3071365 | -0,788013078 | 0,000332602 | chr6 | - |
| Mansc1 | 268,8337326 | -1,016357902 | 0,000757552 | chr6 | - |
| Mrpl19 | 1003,586429 | -0,590812315 | 0,002292524 | chr6 | - |
| Mthfd2 | 24,27416429 | -1,2801751 | 0,001857909 | chr6 | - |
| Mug1 | 50505,14941 | -0,538932189 | 0,024860131 | chr6 | + |
| Ndufa4 | 15014,61604 | -0,611798181 | 0,025808311 | chr6 | - |
| Pparg | 94,23140899 | -1,211751148 | 0,000196359 | chr6 | + |
| Sem1 | 1805,554042 | -0,587728909 | 0,0392417 | chr6 | - |
| Slc15a5 | 66,88372432 | -1,285667503 | 0,001762813 | chr6 | - |
| Slco1a1 | 9915,09478 | -1,085615765 | 0,000304342 | chr6 | - |
| Tatdn2 | 425,8017944 | -0,547972133 | 0,042602547 | chr6 | + |
| Tmcc1 | 607,6821215 | -0,647478162 | 0,031858679 | chr6 | - |
| Tspan33 | 178,0614496 | -1,162769251 | 0,00320933 | chr6 | + |
| Txnrd3 | 106,3262306 | -0,773632883 | 0,010993925 | chr6 | + |
| 0610005C13Rik | 1832,578989 | -0,769159349 | 0,002588169 | chr7 | - |
| Acsm2 | 51,19588206 | -2,04591194 | 9,66E-10 | chr7 | + |
| Bag3 | 279,2688218 | -0,530745538 | 0,03238963 | chr7 | + |
| BC024386 | 3914,888947 | -0,810974427 | 0,007481032 | chr7 | + |
| Cadm4 | 120,4414432 | -1,383893193 | 0,000397498 | chr7 | + |
| Clns1a | 459,4197094 | -0,505708168 | 0,0348796 | chr7 | + |
| Clpb | 557,3090906 | -0,609026069 | 0,018278979 | chr7 | + |
| Cpeb1 | 25,48526043 | -1,493574463 | 0,000261732 | chr7 | - |
| Ctsc | 1600,360607 | -0,524486577 | 0,012392986 | chr7 | + |
| Cyp2a5 | 7713,567049 | -0,97011873 | 0,001289374 | chr7 | + |
| Cyp2e1 | 96588,97949 | -0,93971779 | 7,81E-05 | chr7 | + |
| Cyp2f2 | 23331,23482 | -0,651684872 | 0,003178125 | chr7 | + |
| Cyp2r1 | 320,4577794 | -0,717290451 | 0,017309231 | chr7 | - |
| Fam98c | 60,36051215 | -0,986933643 | 0,02758864 | chr7 | - |
| Fgf21 | 20,52856378 | -1,518622851 | 0,002091619 | chr7 | - |
| Ftl1 | 50529,05053 | -0,638243712 | 0,046298439 | chr7 | - |
| Gde1 | 899,3120328 | -1,16737678 | 6,11E-07 | chr7 | - |
| Hras | 494,9467359 | -0,73729016 | 0,013501349 | chr7 | - |
| Hspb6 | 125,5793587 | -0,947532238 | 0,003614138 | chr7 | + |
| Kcnc3 | 32,00650061 | -0,981588725 | 0,022027814 | chr7 | + |
| Leng8 | 339,363623 | -0,867905169 | 0,018849928 | chr7 | + |
| Mical2 | 88,2324174 | -0,659947803 | 0,049463927 | chr7 | + |
| Nox4 | 474,8495996 | -1,721020852 | 2,14E-07 | chr7 | + |
| Pcsk6 | 908,4042866 | -0,914678166 | 2,45E-05 | chr7 | + |
| Pde3b | 995,5614852 | -0,607780625 | 0,017119669 | chr7 | + |
| Pdilt | 246,8633311 | -0,855183715 | 0,009803754 | chr7 | - |
| Phkg2 | 412,4092324 | -0,618481402 | 0,010054922 | chr7 | + |
| Plekhb1 | 240,3775018 | -0,924240557 | 0,027397756 | chr7 | - |
| Plekhf1 | 159,9078327 | -0,846507487 | 0,03704255 | chr7 | - |
| Ptov1 | 301,1988899 | -0,900221964 | 0,010383158 | chr7 | - |

| | | | | | |
|---|---|---|---|---|---|
| Rab38 | 7,322750129 | -1,312049125 | 0,013067104 | chr7 | + |
| Rpl28 | 3981,915709 | -0,586656586 | 0,036872088 | chr7 | + |
| Sephs2 | 13436,53639 | -0,557984171 | 0,008539289 | chr7 | - |
| Sirt2 | 550,1761773 | -0,683271751 | 0,008205742 | chr7 | + |
| Smpd1 | 632,2932347 | -0,658436163 | 0,020741262 | chr7 | + |
| Stard10 | 7585,434527 | -0,717659058 | 0,016559824 | chr7 | + |
| Syt3 | 16,61578002 | -1,502129208 | 0,000757552 | chr7 | + |
| Tmem147 | 1486,052332 | -0,579382732 | 0,015571226 | chr7 | - |
| Tmem160 | 97,1578225 | -1,059902497 | 0,012896621 | chr7 | + |
| Tmem219 | 596,3526325 | -0,643940993 | 0,0386619 | chr7 | - |
| Tsku | 1423,936757 | -1,011397236 | 0,020449998 | chr7 | - |
| Ube2s | 202,2459266 | -0,770903617 | 0,027857654 | chr7 | - |
| Uri1 | 513,2519891 | -0,604309303 | 0,008464 | chr7 | - |
| Zfp511 | 67,43061757 | -0,964337514 | 0,012476018 | chr7 | + |
| Zfp592 | 324,5691571 | -0,534590325 | 0,045343803 | chr7 | + |
| 1190005I06Rik | 35,82764207 | -1,316486946 | 0,008539289 | chr8 | - |
| 1810019D21Rik | 338,7674472 | -0,91732997 | 0,023294177 | chr8 | + |
| 4833411C07Rik | 124,841746 | -0,880118177 | 0,028728461 | chr8 | + |
| Abcb10 | 571,8890526 | -0,72565963 | 0,005140595 | chr8 | - |
| Gpat4 | 2341,247034 | -0,913112851 | 1,01E-07 | chr8 | - |
| B930025P03Rik | 26,28050786 | -1,314421474 | 0,008960501 | chr8 | - |
| Ces1e | 3543,819918 | -0,98772827 | 0,001073079 | chr8 | - |
| Ces2a | 10505,76775 | -0,609939656 | 0,019665653 | chr8 | + |
| Ces3a | 33618,89322 | -0,626931722 | 0,012305883 | chr8 | + |
| Cmc2 | 295,0967043 | -0,523794086 | 0,049614731 | chr8 | - |
| Cox4i1 | 6670,948443 | -0,608285071 | 0,033496932 | chr8 | + |
| Cyb5b | 6963,951998 | -0,732901661 | 0,000162123 | chr8 | + |
| F10 | 7726,107873 | -0,748118225 | 0,000111314 | chr8 | + |
| Fbxo31 | 464,7129207 | -0,771835035 | 0,020741262 | chr8 | - |
| Hp | 73195,35223 | -0,932674272 | 0,016705702 | chr8 | - |
| Hsd17b2 | 2746,980391 | -0,616910636 | 0,003968497 | chr8 | + |
| Irf2bp2 | 663,1048968 | -0,548054485 | 0,036991353 | chr8 | - |
| Isyna1 | 72,53910018 | -1,013318304 | 0,034963938 | chr8 | + |
| Map1lc3b | 738,121321 | -0,862521541 | 0,03992849 | chr8 | + |
| Mcoln1 | 304,3251497 | -0,577619925 | 0,018443497 | chr8 | + |
| Mt1 | 1918,248006 | -0,868118811 | 0,028145276 | chr8 | + |
| Osgin1 | 2309,300593 | -0,981456155 | 0,03345025 | chr8 | + |
| Pdgfrl | 22,53317639 | -1,385597404 | 0,004879833 | chr8 | + |
| Slc7a6os | 210,7515893 | -0,541544125 | 0,043162954 | chr8 | - |
| Tdrp | 166,021866 | -0,711293908 | 0,018319769 | chr8 | - |
| Zfp703 | 57,23916041 | -1,194855707 | 0,003747365 | chr8 | + |
| 1190002N15Rik | 604,671199 | -0,823502521 | 0,004467131 | chr9 | - |
| Acaa1b | 5790,000759 | -0,969590682 | 0,033600431 | chr9 | - |
| Acp5 | 1590,528387 | -0,858228559 | 0,002413927 | chr9 | - |
| Arih2 | 576,3734295 | -0,615167035 | 0,008898973 | chr9 | - |
| Atg4d | 274,632413 | -0,756247551 | 0,017852348 | chr9 | + |
| Bud13 | 75,97780492 | -1,264084042 | 0,000194337 | chr9 | + |
| C2cd2l | 392,2698335 | -0,665619188 | 0,043054595 | chr9 | - |
| Ccdc84 | 118,6624153 | -0,936304786 | 0,018960647 | chr9 | - |
| Cox5a | 2439,981447 | -0,626636587 | 0,033058809 | chr9 | + |

| | | | | | |
|---|---|---|---|---|---|
| Cox7a2 | 6880,151559 | -0,515711831 | 0,036809024 | chr9 | - |
| Cspg5 | 17,69854646 | -2,118479443 | 1,78E-06 | chr9 | + |
| Cyp1a2 | 6528,198282 | -1,252432486 | 3,65E-12 | chr9 | - |
| Ss18l2 | 1029,503423 | -0,628433853 | 0,014891077 | chr9 | + |
| Endod1 | 59,63958073 | -1,17842219 | 0,000317571 | chr9 | - |
| Gm10658 | 192,7452731 | -0,649041189 | 0,023062773 | chr9 | - |
| Gm16551 | 74,87856489 | -1,835795327 | 1,09E-06 | chr9 | + |
| Gpx1 | 23078,11555 | -0,91419591 | 0,032626917 | chr9 | + |
| Gsta4 | 849,5670499 | -1,084844743 | 3,57E-05 | chr9 | + |
| Lactb | 757,8313305 | -0,660423338 | 0,006253984 | chr9 | - |
| Mto1 | 200,3394468 | -0,540458127 | 0,046333585 | chr9 | + |
| Plscr4 | 445,8707509 | -0,773304972 | 0,02937175 | chr9 | + |
| Rp9 | 331,1343985 | -0,6386828 | 0,026541471 | chr9 | - |
| Rpl14 | 1992,262977 | -0,523275412 | 0,01644885 | chr9 | + |
| Rpsa | 13835,96581 | -0,600336374 | 0,001059132 | chr9 | + |
| Slc25a38 | 197,6076218 | -0,744670602 | 0,029376596 | chr9 | + |
| Wdr72 | 105,7410374 | -1,436462953 | 4,25E-05 | chr9 | + |
| Xylb | 1509,769666 | -0,54222652 | 0,048412358 | chr9 | + |
| Zpr1 | 962,9520765 | -1,18708708 | 5,68E-10 | chr9 | + |
| 5730435O14Rik | 14,60787249 | -1,199070292 | 0,020156171 | chr10 | + |
| Avpr1a | 134,9273051 | -0,710895917 | 0,017119669 | chr10 | + |
| B4galnt1 | 512,4189125 | -0,726611872 | 0,034199883 | chr10 | + |
| C730027H18Rik | 314,3969552 | -0,914975635 | 0,000574674 | chr10 | - |
| Cdc34 | 620,2903492 | -0,798462396 | 0,025898283 | chr10 | + |
| Chpt1 | 2194,716263 | -0,909540347 | 0,000121482 | chr10 | - |
| Coq10a | 371,8756189 | -0,839894556 | 0,000930482 | chr10 | - |
| Cpm | 30,84615509 | -1,004003284 | 0,038478567 | chr10 | + |
| Csrp2 | 643,3087034 | -1,088813454 | 0,00094768 | chr10 | + |
| Fam184a | 3,643623303 | -1,318168282 | 0,008847001 | chr10 | - |
| Klf16 | 64,38674723 | -0,884463435 | 0,008951259 | chr10 | - |
| Mars | 738,8366182 | -0,680546769 | 0,007909756 | chr10 | - |
| Nr2c1 | 1493,705581 | -1,135961504 | 9,53E-12 | chr10 | + |
| Nt5dc3 | 166,919883 | -0,624464727 | 0,043437789 | chr10 | + |
| Pah | 24058,84111 | -0,662581935 | 0,027397756 | chr10 | + |
| Pbld2 | 2473,855154 | -0,98564815 | 8,05E-06 | chr10 | + |
| Pcsk4 | 151,5964638 | -1,393040209 | 0,00011603 | chr10 | - |
| Rdh9 | 1098,233523 | -0,848744952 | 0,031498565 | chr10 | + |
| Reep6 | 3419,256485 | -0,975954233 | 0,00086015 | chr10 | + |
| Rpl41 | 9518,483844 | -0,580243458 | 0,019328711 | chr10 | - |
| Rps15 | 7049,478235 | -0,573529632 | 0,017102722 | chr10 | + |
| Slc19a1 | 101,7863554 | -0,882924966 | 0,005235114 | chr10 | + |
| Slc25a3 | 3931,125728 | -0,471526456 | 0,039207193 | chr10 | - |
| Slc35e3 | 350,2123026 | -0,602230538 | 0,016819129 | chr10 | - |
| Tle2 | 67,86470317 | -1,020460691 | 0,040651668 | chr10 | + |
| Tmem5 | 312,1983274 | -0,653751491 | 0,037519451 | chr10 | - |
| Zfc3h1 | 413,2961496 | -0,721802597 | 0,000848471 | chr10 | + |
| Aatk | 27,85027026 | -1,064744841 | 0,011248868 | chr11 | - |
| Adcy1 | 18,95404848 | -1,126926707 | 0,014703567 | chr11 | + |
| Cbx1 | 154,1745469 | -2,413763144 | 1,74E-24 | chr11 | + |
| Ccdc157 | 193,4160498 | -0,944909284 | 0,000120613 | chr11 | - |

| | | | | | |
|---|---|---|---|---|---|
| Ccl9 | 2544,583533 | -0,939411908 | 0,004117382 | chr11 | - |
| Coasy | 1902,529745 | -0,477043082 | 0,019608972 | chr11 | + |
| D630024D03Rik | 27,44762605 | -1,036412786 | 0,014752457 | chr11 | - |
| Dhrs13 | 34,77150824 | -1,279589912 | 0,000258327 | chr11 | + |
| Dus1l | 827,5388555 | -0,827370416 | 0,001332784 | chr11 | - |
| Epop | 30,67052722 | -1,156248947 | 0,007398297 | chr11 | - |
| Egfr | 5863,448674 | -0,811299881 | 0,000837089 | chr11 | + |
| Emc6 | 976,6712176 | -0,692317174 | 0,036991353 | chr11 | - |
| Gcgr | 2500,107348 | -0,694824911 | 0,02758864 | chr11 | + |
| Gm16062 | 50,51785427 | -1,05382521 | 0,024221418 | chr11 | + |
| Mafg | 146,6980623 | -1,158728721 | 0,014243688 | chr11 | - |
| Mien1 | 1264,874824 | -0,548669885 | 0,014703567 | chr11 | - |
| Mir22hg | 696,5055869 | -0,688011408 | 0,016688812 | chr11 | + |
| Mrps24 | 385,2728373 | -0,647050188 | 0,024779229 | chr11 | - |
| Nme1 | 1798,527939 | -0,470630563 | 0,038150816 | chr11 | - |
| Nme2 | 6164,184584 | -0,659750697 | 0,010642415 | chr11 | - |
| Ppia | 15966,92849 | -0,592170754 | 0,018669488 | chr11 | + |
| Pwwp2a | 161,0000854 | -0,736164853 | 0,010156653 | chr11 | + |
| Rpl38 | 3859,258122 | -0,600926035 | 0,019277154 | chr11 | + |
| Traf4 | 426,8732646 | -0,652761553 | 0,048483358 | chr11 | - |
| Tubg1 | 229,4193248 | -0,690877487 | 0,047493774 | chr11 | + |
| Zfp354a | 58,95932929 | -0,960408629 | 0,006200512 | chr11 | + |
| Cyp46a1 | 11,18702077 | -1,641020231 | 0,000628663 | chr12 | + |
| Dio3os | 4,667044602 | -1,373940509 | 0,005485821 | chr12 | - |
| Dtnb | 238,6548768 | -0,679535789 | 0,042736499 | chr12 | + |
| Eif5 | 4824,59089 | -0,494791543 | 0,038143836 | chr12 | + |
| Gm20337 | 19,75553193 | -1,009722439 | 0,02696191 | chr12 | - |
| Hspa2 | 31,79447532 | -1,071664111 | 0,017829148 | chr12 | + |
| Mycn | 8,689688425 | -1,258711877 | 0,017323407 | chr12 | - |
| Nampt | 2059,347181 | -0,621098879 | 0,014683094 | chr12 | + |
| Pomt2 | 295,8080509 | -0,625738996 | 0,011009293 | chr12 | - |
| Serpina12 | 3151,532492 | -1,340907484 | 1,81E-06 | chr12 | - |
| Serpina1c | 50060,81326 | -0,911836967 | 1,91E-06 | chr12 | - |
| Serpina3k | 301428,0127 | -1,213049469 | 1,12E-06 | chr12 | + |
| Serpina6 | 3116,735822 | -1,610950643 | 4,34E-11 | chr12 | - |
| Tc2n | 46,9013137 | -1,033766304 | 0,019869728 | chr12 | - |
| Timm9 | 625,0945781 | -1,015689281 | 5,43E-08 | chr12 | - |
| Ttc7b | 186,4605596 | -0,771787399 | 0,006385226 | chr12 | - |
| Ak6 | 328,4149114 | -0,652186714 | 0,03196404 | chr13 | + |
| Akr1c12 | 1923,180662 | -0,831816114 | 0,001083279 | chr13 | - |
| Akr1c13 | 2353,105359 | -0,812848076 | 0,00138517 | chr13 | + |
| Akr1c19 | 2139,348507 | -1,502707553 | 5,08E-07 | chr13 | + |
| Cox7c | 1958,473203 | -0,524451033 | 0,02272099 | chr13 | - |
| Dtnbp1 | 450,3938188 | -0,600037676 | 0,026151101 | chr13 | - |
| Fars2 | 601,9241558 | -0,644930001 | 0,007355049 | chr13 | + |
| Gadd45g | 113,8596962 | -1,656399133 | 0,000122644 | chr13 | + |
| Gcnt4 | 111,2146508 | -0,756069945 | 0,039560456 | chr13 | + |
| Glrx | 1009,294578 | -1,067071389 | 1,11E-06 | chr13 | + |
| Gmnn | 216,8788221 | -0,685560478 | 0,008893706 | chr13 | - |
| Hist1h2bc | 1222,523623 | -0,939436569 | 0,034522477 | chr13 | + |

| | | | | | |
|---|---|---|---|---|---|
| Lyrm4 | 229,8272558 | -0,652148676 | 0,022611041 | chr13 | - |
| Map1b | 298,1743944 | -0,94841449 | 0,000948912 | chr13 | - |
| Marcksl1-ps4 | 175,272919 | -1,34709911 | 5,93E-06 | chr13 | + |
| Nrn1 | 2855,35176 | -1,933933515 | 1,46E-18 | chr13 | - |
| Ntrk2 | 17,23836454 | -1,261876998 | 0,017729939 | chr13 | + |
| Pelo | 166,8603705 | -0,794544272 | 0,022982029 | chr13 | - |
| Ptdss1 | 637,5256584 | -0,6422181 | 0,003747365 | chr13 | + |
| Ripk1 | 618,3703734 | -0,459543485 | 0,047966165 | chr13 | + |
| Rps23 | 3290,728746 | -0,486690795 | 0,034767571 | chr13 | + |
| Simc1 | 97,88599324 | -0,921245056 | 0,00094424 | chr13 | + |
| Smim13 | 588,1232088 | -0,521217664 | 0,044362742 | chr13 | + |
| Tmem14c | 2428,362218 | -0,475245907 | 0,026124007 | chr13 | + |
| Bmp4 | 40,09312736 | -1,181661925 | 0,001926128 | chr14 | - |
| Cgrrf1 | 790,2026352 | -0,568576463 | 0,044937637 | chr14 | + |
| Cnih1 | 1011,750131 | -0,488391647 | 0,025882765 | chr14 | - |
| Cryl1 | 291,0043372 | -0,832560979 | 0,000286779 | chr14 | - |
| Dct | 59,40851357 | -3,092593292 | 1,18E-17 | chr14 | - |
| Fitm1 | 201,2472608 | -2,023446666 | 8,26E-12 | chr14 | + |
| Gulo | 6342,620839 | -0,990465564 | 3,00E-08 | chr14 | - |
| Ints6 | 253,8719318 | -0,592489395 | 0,017896685 | chr14 | - |
| Cnmd | 101,2686916 | -1,461698044 | 0,001031759 | chr14 | - |
| Lmo7 | 192,4108423 | -0,702629537 | 0,034416582 | chr14 | + |
| Lrit2 | 94,6279542 | -0,759483973 | 0,044355016 | chr14 | + |
| Pabpn1 | 986,6891034 | -0,608516375 | 0,03945775 | chr14 | + |
| Ppp2r2a | 306,4867432 | -0,554617499 | 0,023381506 | chr14 | - |
| Rap2a | 367,7974272 | -0,72056876 | 0,028145276 | chr14 | + |
| Sall2 | 33,79392629 | -1,036812489 | 0,026884253 | chr14 | - |
| Sftpa1 | 32,60530444 | -1,084088026 | 0,041208422 | chr14 | + |
| Slc25a37 | 543,4421577 | -0,641976362 | 0,00444528 | chr14 | - |
| Adck5 | 260,0829885 | -0,723402667 | 0,02742883 | chr15 | + |
| Arsa | 612,070542 | -0,668972201 | 0,022611041 | chr15 | - |
| Atf4 | 1202,816507 | -0,511716575 | 0,048412358 | chr15 | + |
| C9 | 15178,64109 | -1,359728005 | 1,29E-07 | chr15 | + |
| Cpne8 | 90,21688904 | -0,932451654 | 0,001171489 | chr15 | - |
| Cyc1 | 3204,267193 | -0,523288901 | 0,033600431 | chr15 | + |
| Cyhr1 | 883,3730779 | -0,530512752 | 0,016705702 | chr15 | - |
| Cyp2d13 | 1367,587918 | -1,000595749 | 0,00444528 | chr15 | - |
| Cyp2d26 | 9004,81683 | -0,950254211 | 2,57E-05 | chr15 | - |
| Cyp2d37-ps | 317,3572454 | -0,828832532 | 0,031858679 | chr15 | - |
| Cyp2d40 | 767,2762009 | -1,421067031 | 6,83E-06 | chr15 | - |
| Dgat1 | 293,6858981 | -0,696838074 | 0,006796468 | chr15 | - |
| Dnajc21 | 360,4442934 | -0,588900973 | 0,047116916 | chr15 | - |
| Kifc2 | 46,93726944 | -0,922561804 | 0,03196404 | chr15 | + |
| Mapk15 | 50,67897872 | -2,004008299 | 1,22E-07 | chr15 | + |
| Nrbp2 | 384,0812827 | -1,003924597 | 0,003263682 | chr15 | - |
| Pim3 | 735,3126895 | -0,741054292 | 0,010521428 | chr15 | + |
| Rpl37 | 3029,136015 | -0,713502993 | 0,000332602 | chr15 | + |
| Slc38a2 | 3260,065481 | -0,831727158 | 0,01070874 | chr15 | - |
| Slc52a2 | 87,73962808 | -0,741247593 | 0,04104012 | chr15 | + |
| Ugt3a2 | 8340,983305 | -0,617140951 | 0,022503908 | chr15 | + |

| | | | | | |
|---|---|---|---|---|---|
| Zcrb1 | 333,4242399 | -0,73327156 | 0,016578697 | chr15 | - |
| Crybg3 | 114,223222 | -0,709717317 | 0,042365922 | chr16 | - |
| D16Ertd472e | 187,841275 | -0,64276439 | 0,03417463 | chr16 | - |
| Dexi | 146,4860812 | -0,664146742 | 0,034968569 | chr16 | - |
| Dirc2 | 468,8205309 | -0,531777778 | 0,038989195 | chr16 | - |
| Fetub | 7196,39517 | -0,655889616 | 0,005350521 | chr16 | + |
| Hgd | 13285,10159 | -0,552052108 | 0,030459905 | chr16 | + |
| Ifnar2 | 954,1631176 | -0,744063349 | 0,000961755 | chr16 | + |
| Kalrn | 160,0811512 | -0,73638984 | 0,021755974 | chr16 | - |
| Naa60 | 1325,86877 | -0,778211375 | 0,000695763 | chr16 | + |
| Nubp1 | 221,5368379 | -0,668604294 | 0,021655699 | chr16 | + |
| Nectin3 | 766,5176088 | -0,681954611 | 0,019387746 | chr16 | - |
| Rpl24 | 5672,037317 | -0,543513071 | 0,007994307 | chr16 | + |
| Setd4 | 75,18121423 | -0,906651251 | 0,016191769 | chr16 | - |
| Tmem41a | 117,2002681 | -1,031455921 | 0,002904517 | chr16 | - |
| Tedc2 | 1058,369584 | -0,564024021 | 0,03992849 | chr17 | - |
| 2310061I04Rik | 248,9843069 | -0,6784944 | 0,019682666 | chr17 | - |
| Abcg8 | 1041,401623 | -1,190912235 | 8,32E-11 | chr17 | + |
| Atat1 | 132,1343441 | -0,836668455 | 0,030454862 | chr17 | - |
| Cyp39a1 | 880,007474 | -0,881225768 | 0,00164095 | chr17 | + |
| Cyp4f13 | 813,5021439 | -0,667226138 | 0,015058402 | chr17 | - |
| Dlgap1 | 6,877808516 | -1,195356078 | 0,028728461 | chr17 | + |
| Epas1 | 1589,748272 | -0,542338264 | 0,028940327 | chr17 | + |
| Glo1 | 9075,137634 | -0,983341149 | 2,30E-09 | chr17 | - |
| H2-Ke6 | 735,5175747 | -0,670671573 | 0,002451531 | chr17 | - |
| H2-Q10 | 24174,13561 | -0,798357419 | 0,006870993 | chr17 | + |
| Hspa1b | 57,09307647 | -1,145082097 | 0,017708051 | chr17 | - |
| March2 | 753,2066879 | -0,698062614 | 0,000315726 | chr17 | - |
| Mocs1 | 1567,252164 | -0,563411671 | 0,031151848 | chr17 | + |
| Ndufa11 | 1805,792721 | -0,494074799 | 0,034083155 | chr17 | + |
| Ndufv3 | 2235,000886 | -0,458440251 | 0,020156171 | chr17 | + |
| Pgp | 74,02513898 | -1,058499734 | 0,02578402 | chr17 | + |
| Phf10 | 324,4234793 | -0,629945038 | 0,010050828 | chr17 | - |
| Pnpla1 | 202,0736682 | -1,150777426 | 0,00177917 | chr17 | + |
| Rps10 | 3619,553184 | -0,57289279 | 0,025873383 | chr17 | - |
| Rps18 | 11860,74473 | -0,515791908 | 0,028940327 | chr17 | - |
| Rps2 | 5961,419848 | -0,665046108 | 0,029376596 | chr17 | + |
| Slc25a23 | 2096,064325 | -0,631831091 | 0,004657848 | chr17 | - |
| Slc3a1 | 795,9799556 | -1,139791771 | 0,000623221 | chr17 | + |
| Tiam2 | 129,631331 | -1,146905299 | 1,34E-05 | chr17 | + |
| Zdhhc14 | 41,38884893 | -1,275762128 | 0,000244283 | chr17 | + |
| Zfp598 | 372,811528 | -0,73359399 | 0,046298439 | chr17 | + |
| Arhgap26 | 220,8291958 | -1,5123532 | 6,77E-08 | chr18 | + |
| Bin1 | 159,0095572 | -0,84662723 | 0,007878532 | chr18 | + |
| Cabyr | 77,56489549 | -2,119597388 | 3,53E-07 | chr18 | + |
| Colec12 | 545,1987743 | -2,238756689 | 1,12E-18 | chr18 | + |
| Cul2 | 771,8930521 | -0,734893036 | 0,001071697 | chr18 | + |
| Dnajc18 | 167,4114024 | -0,640147165 | 0,027633464 | chr18 | - |
| Dym | 396,161322 | -0,638765457 | 0,017016792 | chr18 | + |
| Fam210a | 1355,227568 | -0,58108596 | 0,017729939 | chr18 | - |

| | | | | | |
|---|---|---|---|---|---|
| Fzd8 | 62,31998752 | -1,350259553 | 4,31E-05 | chr18 | + |
| Mapk4 | 3,113999258 | -1,260458962 | 0,010854611 | chr18 | - |
| Prob1 | 58,51941249 | -1,566904586 | 3,76E-06 | chr18 | - |
| Rps14 | 8307,239673 | -0,509429 | 0,0392417 | chr18 | + |
| Snhg4 | 37,2501845 | -1,212922587 | 0,001539492 | chr18 | + |
| Ttr | 182673,4231 | -0,58372063 | 0,009227081 | chr18 | + |
| Wac | 933,2076512 | -0,477180751 | 0,020344623 | chr18 | + |
| Cyp2c29 | 16036,68672 | -2,600931177 | 7,65E-25 | chr19 | + |
| Cyp2c23 | 4260,324875 | -2,203233987 | 2,18E-20 | chr19 | - |
| Cyp2c53-ps | 4,624695965 | -1,311398195 | 0,011778732 | chr19 | + |
| Eef1g | 2553,891764 | -0,667087489 | 0,000937112 | chr19 | + |
| Elovl3 | 1058,631048 | -1,802973979 | 5,39E-15 | chr19 | + |
| Fads1 | 4990,291872 | -0,701850469 | 0,000208779 | chr19 | + |
| Fxn | 166,740376 | -0,835153934 | 0,036091001 | chr19 | - |
| Gm10768 | 2270,568548 | -0,634808051 | 0,012392986 | chr19 | + |
| Gstp1 | 8787,825262 | -1,105618918 | 7,57E-05 | chr19 | - |
| Gstp2 | 67,82359902 | -1,801834677 | 3,84E-09 | chr19 | - |
| March5 | 1809,016424 | -0,54549909 | 0,034199883 | chr19 | + |
| Marveld1 | 368,6298225 | -0,750045224 | 0,007681366 | chr19 | + |
| Tesmin | 20,63907713 | -0,967018889 | 0,044651845 | chr19 | + |
| Nudt8 | 183,3906987 | -0,894085691 | 0,003604079 | chr19 | + |
| Plgrkt | 69,77325539 | -0,939070235 | 0,013794098 | chr19 | - |
| Ppp1r3c | 440,8905019 | -0,750832723 | 0,00320933 | chr19 | - |
| Ppp2r5b | 68,05770887 | -0,917446215 | 0,029506282 | chr19 | - |
| Psat1 | 24,89152526 | -1,079534948 | 0,010697902 | chr19 | - |
| Rbp4 | 40817,19012 | -1,304125645 | 1,91E-07 | chr19 | - |
| Rcl1 | 3552,99437 | -0,476571246 | 0,020553613 | chr19 | + |
| Slc22a28 | 244,5963716 | -1,363232248 | 0,000372363 | chr19 | - |
| Slc22a30 | 1806,794155 | -0,917439801 | 5,78E-06 | chr19 | - |
| Slc25a28 | 137,3734557 | -0,732955419 | 0,026541471 | chr19 | - |
| Slc35g1 | 431,6298184 | -0,887139965 | 0,017300648 | chr19 | + |
| Tm7sf2 | 1861,253733 | -0,591651459 | 0,039766485 | chr19 | - |
| Alas2 | 2078,499095 | -0,805951322 | 0,016167446 | chrX | + |
| Ebp | 3961,507667 | -0,441795817 | 0,046074596 | chrX | - |
| Adgrg2 | 8,543428139 | -1,532339374 | 0,00213548 | chrX | + |
| Irak1 | 646,8207928 | -0,509970639 | 0,02742883 | chrX | - |
| Nup62cl | 14,54162345 | -1,695475556 | 0,000304499 | chrX | - |
| Rai2 | 18,23277787 | -1,02263942 | 0,044937637 | chrX | + |
| Rpl10 | 17282,89843 | -0,517478478 | 0,012536589 | chrX | + |
| Ssr4 | 3109,782654 | -0,713263281 | 0,00444528 | chrX | + |
| Uxt | 530,2509432 | -0,52845814 | 0,037864464 | chrX | - |
