## Supplementary material for "The mouse HP1 proteins are essential for preventing liver tumorigenesis": suppl table2

**Supplementary Table 2** : functional clustering of genes up-regulated upon loss of HP1 witin hepatocytes (https://david.ncifcrf.gov/)

| Annotation | Enrichment Score: 12.291891875812565 | | | | | | | | | | | |
|---|---|---|---|---|---|---|---|---|---|---|---|---|
| Category | Term | Count | % | PValue | Genes | List Total | Pop Hits | Pop Total | Fold Enrichment | Bonferroni | Benjamini | FDR |
| INTERPRO | IPR001909:Krueppel-associated box | 61 | 8,6 | 5,80E-26 | GM14124, ZFP13, 5730507C01RIK, ZKSCAN8, ZFP872, ZFP84, ZKSCAN3, ZFP870, ZFP935, ZFP934, ZFP933, ZFP781, ZFP882, ZFP938, GM14139, ZFP426, RSLCAN18, ZFP78, ZFP26, ZFP799, ZFP120, ZFP791, AU041133, ZFP558, GM3604, ZFP229, ZFP369, ZFP266, ZFP719, ZFP850, ZFP763, ZFP759, ZFP563, 2010315B03RIK, ZFP952, ZFP951, ZFP667, ZFP157, 2410141K09RIK, RSL1, 1700048O20RIK, ZFP72, ZFP53, ZFP345, ZFP605, ZFP955B, ZFP955A, ZFP445, GM20939, 6720489N17RIK, ZFP709, 2410018L13RIK, ZFP809, ZFP808, ZFP941, ZFP677, 9030624G23RIK, ZFP773, ZFP947, GM10324, ZIK1 | 645 | 373 | 20594 | 5,22 | 6,25E-23 | 6,25E-23 | 9,25E-23 |
| SMART | SM00349:KRAB | 60 | 8,46 | 6,13E-23 | GM14124, ZFP13, 5730507C01RIK, ZKSCAN8, ZFP872, ZFP84, ZKSCAN3, ZFP870, ZFP935, ZFP934, ZFP933, ZFP781, ZFP882, ZFP938, GM14139, ZFP426, RSLCAN18, ZFP78, ZFP26, ZFP799, ZFP120, ZFP791, AU041133, ZFP558, GM3604, ZFP229, ZFP369, ZFP266, ZFP719, ZFP850, ZFP763, ZFP759, ZFP563, 2010315B03RIK, ZFP952, ZFP951, ZFP667, 2410141K09RIK, RSL1, ZFP157, 1700048O20RIK, ZFP53, ZFP345, ZFP605, ZFP955B, ZFP955A, ZFP445, GM20939, 6720489N17RIK, ZFP709, 2410018L13RIK, ZFP809, ZFP808, ZFP941, ZFP677, 9030624G23RIK, ZFP773, ZFP947, GM10324, ZIK1 | 379 | 367 | 10425 | 4,50 | 1,38E-20 | 1,38E-20 | 7,83E-20 |
| INTERPRO | IPR015880:Zinc finger, C2H2-like | 70 | 9,87 | 1,02E-17 | ZKSCAN7, ZMAT1, GM14124, ZFP13, 5730507C01RIK, ZKSCAN8, ZFP872, ZFP84, ZKSCAN3, ZFP870, ZKSCAN4, ZFP935, ZFP934, ZFP873, ZFP933, ZFP882, ZFP938, ZFP518A, ZFP426, GM14139, RSLCAN18, ZFP78, CHAMP1, ZFP26, ZFP799, ZFP120, ZFP791, AU041133, ZFP558, ZSCAN18, GM3604, ZFP229, ZFP369, ZFP266, DPF3, ZFP719, ZFP850, ZFP763, ZFP759, ZFP563, 2010315B03RIK, ZFP952, ZFP951, ZFP667, ZFP157, 2410141K09RIK, RSL1, ZFP72, PLAG1, ZFP53, ZFP345, KLF6, ZFP605, ZFP955B, ZFP955A, ZFP449, ZFP445, GM20939, ZFP709, ZFP809, ZFP808, ADNP2, ZFP941, ZFP105, ZFP770, ZFP677, ZFP773, ZFP947, GM10324, ZIK1 | 645 | 693 | 20594 | 3,23 | 1,10E-14 | 5,49E-15 | 1,62E-14 |

| | | | | | | | | | | | | |
|---|---|---|---|---|---|---|---|---|---|---|---|---|
| INTERPRO | IPR007087:Zinc finger, C2H2 | 71 | 10 | 4,52E-17 | ZKSCAN7, ZMAT1, GM14124, ZFP13, 5730507C01RIK, ZKSCAN8, ZFP872, ZFP84, ZKSCAN3, ZFP870, ZKSCAN4, ZFP935, ZFP934, ZFP873, ZFP933, ZFP882, ZFP938, ZFP518A, ZFP426, GM14139, RSLCAN18, ZFP78, CHAMP1, ZFP26, ZFP799, ZFP120, ZFP791, AU041133, ZFP558, ZSCAN18, GM3604, ZFP229, ZFP369, ZFP266, DPF3, ZFP719, ZFP850, ZFP763, ZFP759, ZFP563, 2010315B03RIK, ZFP952, ZFP951, ZFP667, ZFP157, 2410141K09RIK, RSL1, ZFP72, PLAG1, ZFP53, ZFP345, KLF6, ZFP605, ZFP955B, ZFP955A, ZFP449, ZFP445, GM20939, ZFP709, ZFP608, ZFP809, ZFP808, ADNP2, ZFP941, ZFP105, ZFP770, ZFP677, ZFP773, ZFP947, ZIK1, GM10324 | 645 | 731 | 20594 | 3,10 | 4,87E-14 | 1,62E-14 | 7,21E-14 |
| INTERPRO | IPR013087:Zinc finger C2H2-type/integrase DNA-binding domain | 66 | 9,31 | 8,94E-17 | ZKSCAN7, GM14124, ZFP13, 5730507C01RIK, ZKSCAN8, ZFP872, ZFP84, ZKSCAN3, ZFP870, ZKSCAN4, ZFP935, ZFP934, ZFP873, ZFP933, ZFP882, ZFP938, ZFP426, GM14139, RSLCAN18, ZFP78, CHAMP1, ZFP26, ZFP799, ZFP120, ZFP791, AU041133, ZFP558, GM3604, ZSCAN18, ZFP229, ZFP369, ZFP266, ZFP719, ZFP850, ZFP763, ZFP759, ZFP563, 2010315B03RIK, ZFP952, ZFP951, ZFP667, ZFP157, 2410141K09RIK, RSL1, ZFP72, PLAG1, ZFP53, ZFP345, KLF6, ZFP605, ZFP955B, ZFP955A, ZFP449, ZFP445, GM20939, ZFP709, ZFP809, ZFP808, ZFP941, ZFP105, ZFP770, ZFP677, ZFP773, ZFP947, GM10324, ZIK1 | 645 | 652 | 20594 | 3,23 | 1,20E-13 | 3,00E-14 | 1,78E-13 |
| SMART | SM00355:ZnF_C2H2 | 70 | 9,87 | 5,72E-15 | ZKSCAN7, ZMAT1, GM14124, ZFP13, 5730507C01RIK, ZKSCAN8, ZFP872, ZFP84, ZKSCAN3, ZFP870, ZKSCAN4, ZFP935, ZFP934, ZFP873, ZFP933, ZFP882, ZFP938, ZFP518A, ZFP426, GM14139, RSLCAN18, ZFP78, CHAMP1, ZFP26, ZFP799, ZFP120, ZFP791, AU041133, ZFP558, ZSCAN18, GM3604, ZFP229, ZFP369, ZFP266, DPF3, ZFP719, ZFP850, ZFP763, ZFP759, ZFP563, 2010315B03RIK, ZFP952, ZFP951, ZFP667, ZFP157, 2410141K09RIK, RSL1, ZFP72, PLAG1, ZFP53, ZFP345, KLF6, ZFP605, ZFP955B, ZFP955A, ZFP449, ZFP445, GM20939, ZFP709, ZFP809, ZFP808, ADNP2, ZFP941, ZFP105, ZFP770, ZFP677, ZFP773, ZFP947, GM10324, ZIK1 | 379 | 693 | 10425 | 2,78 | 1,30E-12 | 6,52E-13 | 7,38E-12 |
| UP_KEYWOI | Metal-binding | 171 | 24,1 | 1,85E-12 | ZFP935, ZFP934, ATP2B2, ZFP933, G2E3, RNF141, DNTT, MIOX, PLS1, ZFP882, SPON2, CIB3, ZFP938, AMY1, CLCA1, CRYAB, NCALD, CHP2, CYP2B13, ZFP78, CYP2B10, ZFP799, SP140, ZFP120, BHMT, ZFP791, PGM1, GM3604, OAS1A, ZFP266, RNF17, CYP2B9, ZFP763, ZFP759, ZFP563, TIMP1, ZFP952, ZFP951, ACE, TRIM68, RSL1, ZFP157, DMD, STAMBPL1, PLAG1, ZFP345, KLF6, ZFP605, ACACA, S100A11, ZFP449, SMYD4, HDDC3, CSRP1, GM20939, ZFP445, CSRP3, ZFP608, FOXP2, ZFP809, TRIM56, ZFP808, CDKN1A, ZFP941, ZFP770, MTR, ZFP773, SYTL5, S100G, ZFP947, ZIK1, GM10324, CLEC7A, ATP7B, TAF1B, ZKSCAN7, ZMAT1, STEAP4, GM14124, ZFP13, ZKSCAN8, ARSG, HR, ZKSCAN3, ZFP872, ZFP870, ZKSCAN4, FANCL, LNPEP, TPO, ATP8B5, NT5E, PRKCA, ZFP518A, GM14139, ZFP426, CAR14, CAPSL, VIL1, ESR1, RSLCAN18, MMP15, MBD1, CHAMP1, ZFP26, PMM1, MMP12, AU041133, ZFP558, ZSCAN18, ZFP369, ZFP229, CYP2A4, DPF3, MOB1A, ZFP719, ZFP850, POLA1, CDH1, NHLRC1, 2010315B03RIK, TPM4, RPA1, TRIM30A, NPTX1, ANXA8, NAIP2, ZFP667, TRIM30D, 2410141K09RIK, TGM1, PPP3CC, LIMD2, EHD1, ZC3H12D, ZFP72, BGLAP3, DTX4, ZFP53, BGLAP, SLC8A1, PNLIPRP1, ZFP955B, ZFP955A, ACLY, TRIM24, EXO5, PCK2, ZFP709, COL5A2, TRIM21, AJUBA, MARCH1, QPCT, CYP2A22, AFP, ADNP2, PRLR, CYP4F16, ZFP677, C1S1, LRP2 | 678 | 3395 | 22680 | 1,68 | 6,17E-10 | 6,17E-10 | 2,52E-09 |

| | | | | | | | | | | | | |
|---|---|---|---|---|---|---|---|---|---|---|---|---|
| UP_KEYWO | Zinc | 119 | 16,8 | 1,00E-11 | 5730507C01RIK, THRA, KLK1B4, ZFP84, ZFP935, ZFP934, G2E3, ZFP933, RNF141, ZFP882, ZFP938, CLCA1, CRYAB, ZFP78, ZFP799, SP140, ZFP120, BHMT, ZFP791, GM3604, ZFP266, RNF17, ZFP763, ZFP563, ZFP759, TIMP1, ZFP952, ACE, ZFP951, TRIM68, ZFP157, DMD, RSL1, STAMBPL1, PLAG1, KLF6, ZFP345, ZFP605, ZFP449, SMYD4, CSRP1, GM20939, ZFP445, CSRP3, ZFP608, FOXP2, ZFP809, TRIM56, ZFP808, CDKN1A, ZFP941, ZFP770, MTR, ZFP773, ZFP947, SYTL5, ZIK1, GM10324, TAF1B, ZKSCAN7, ZMAT1, ZFP13, GM14124, ZKSCAN8, HR, ZFP872, ZKSCAN3, ZFP870, ZKSCAN4, LNPEP, FANCL, NT5E, PRKCA, ZFP518A, GM14139, ZFP426, CAR14, ESR1, RSLCAN18, MMP15, CHAMP1, MBD1, MMP12, ZFP26, AU041133, ZFP558, ZSCAN18, ZFP369, ZFP229, DPF3, MOB1A, POLA1, ZFP850, ZFP719, NHLRC1, 2010315B03RIK, RPA1, TRIM30A, NAIP2, ZFP667, 2410141K09RIK, TRIM30D, LIMD2, PPP3CC, ZC3H12D, ZFP72, DTX4, ZFP53, ZFP955B, ZFP955A, TRIM24, ZFP709, TRIM21, AJUBA, MARCH1, QPCT, ADNP2, PRLR, ZFP677 | 678 | 2099 | 22680 | 1,90 | 3,34E-09 | 1,67E-09 | 1,36E-08 |
| UP_KEYWO | Zinc-finger | 97 | 13,7 | 1,08E-11 | THRA, 5730507C01RIK, ZFP84, ZFP935, ZFP934, ZFP933, G2E3, RNF141, ZFP882, ZFP938, ZFP78, ZFP799, SP140, ZFP120, ZFP791, GM3604, ZFP266, RNF17, ZFP763, ZFP563, ZFP759, ZFP952, ZFP951, TRIM68, RSL1, ZFP157, DMD, PLAG1, ZFP345, KLF6, ZFP605, ZFP449, SMYD4, ZFP445, GM20939, ZFP608, FOXP2, ZFP809, TRIM56, ZFP808, CDKN1A, ZFP941, ZFP770, ZFP773, SYTL5, ZFP947, GM10324, ZIK1, ZMAT1, ZKSCAN7, TAF1B, GM14124, ZFP13, ZKSCAN8, HR, ZKSCAN3, ZFP872, ZFP870, ZKSCAN4, FANCL, PRKCA, ZFP518A, GM14139, ZFP426, ESR1, RSLCAN18, CHAMP1, MBD1, ZFP26, AU041133, ZFP558, ZSCAN18, ZFP369, ZFP229, DPF3, ZFP850, ZFP719, POLA1, NHLRC1, 2010315B03RIK, RPA1, TRIM30A, ZFP667, TRIM30D, 2410141K09RIK, ZC3H12D, ZFP72, DTX4, ZFP53, ZFP955B, ZFP955A, TRIM24, ZFP709, TRIM21, MARCH1, ADNP2, ZFP677 | 678 | 1565 | 22680 | 2,07 | 3,59E-09 | 1,20E-09 | 1,46E-08 |
| GOTERM_C | GO:0005622~intracellular | 94 | 13,3 | 1,38E-08 | RTN4, ZMAT1, ZFP13, GM14124, 5730507C01RIK, ZKSCAN8, TUFT1, DNAJC10, TLR3, ZFP872, ZKSCAN3, ZFP84, TLR8, SENP7, ZFP870, LNPEP, ZFP935, ZFP934, ZFP781, ZFP933, ANK2, RASL10B, RHOC, ZFP882, ZFP938, RAB27A, RABL2, ZFP426, GM14139, NCALD, RSLCAN18, ZFP26, ZFP799, DCAF4, SGSM1, ZFP120, AU041133, ZFP558, GM3604, ZFP229, ZFP369, ZFP266, MOB1A, ZFP850, NAPB, ZFP763, ZFP759, ZFP563, 2010315B03RIK, LY6A, TRIM30A, ZFP952, ZFP951, ZFP667, TRIM68, ZFP157, 2410141K09RIK, RSL1, TRIM30D, TFF2, MAS1, 1700048O20RIK, DEFB1, CAPN6, ZFP53, ZFP345, ZFP605, PLEK, LGALS1, ZFP955B, ZFP955A, TRIM24, ANXA5, GM20939, ZFP445, 6720489N17RIK, ZFP709, TRIM21, ZFP809, 2410018L13RIK, ZFP808, TRIM56, RPS6KA3, ZFP941, PRLR, ZFP677, MTR, 9030624G23RIK, ZFP773, ZFP947, SYTL5, ZIK1, TGTP1, GM10324 | 633 | 1599 | 19662 | 1,83 | 5,48E-06 | 2,74E-06 | 1,92E-05 |

| | | | | | | | | | | | | |
|---|---|---|---|---|---|---|---|---|---|---|---|---|
| GOTERM_M | GO:0003676~nucleic acid binding | 79 | 11,1 | 1,91E-08 | ZKSCAN7, ZMAT1, ZFP13, 5730507C01RIK, ZKSCAN8, ZFP872, ZKSCAN3, ZFP84, ZFP870, ZKSCAN4, ZFP935, ZFP934, ZFP873, ZFP781, ZFP933, ZFP882, ZFP938, ZFP426, GM14139, METTL4, RBM43, RSLCAN18, ZFP78, CHAMP1, ZFP26, ZFP799, ZFP120, ZFP791, AU041133, ZFP558, ZSCAN18, GM3604, ZFP229, ZFP369, ZFP266, TNFRSF22, DPF3, TRA2A, TDRD9, POLA1, ZFP719, ZFP850, MYEF2, ZFP763, BICC1, ZFP759, ZFP563, 2010315B03RIK, RPA1, ZFP952, ZFP951, ZFP667, ZFP157, 2410141K09RIK, 1700048O20RIK, ZFP72, PLAG1, KLF6, ZFP53, ZFP345, ZFP605, ZFP955B, ZMYM6, ZFP955A, ZFP445, GM20939, 6720489N17RIK, ZFP709, ZFP809, 2410018L13RIK, ZFP808, ZFP941, ZFP770, ZFP677, 9030624G23RIK, ZFP773, ZFP947, ZIK1, GM10324 | 575 | 1237 | 17446 | 1,94 | 1,42E-05 | 1,42E-05 | 2,90E-05 |
| GOTERM_M | GO:0046872~metal ion binding | 165 | 23,3 | 2,50E-08 | S100A4, SYT1, IMPAD1, 5730507C01RIK, THRA, KLK1B4, ZFP84, CD209C, ZFP935, ZFP934, ATP2B2, ZFP933, G2E3, RNF141, DNTT, MIOX, PLS1, ZFP882, SPON2, CIB3, ZFP938, AMY1, CLCA1, CRYAB, NCALD, CHP2, ZFP78, CYP2B10, ZFP799, ZFP120, ZFP791, PGM1, GM3604, OAS1A, ZFP266, RNF17, CYP2B9, ZFP763, ZFP759, ZFP563, TIMP1, ZFP952, ZFP951, ACE, TRIM68, ZFP157, DMD, RSL1, STAMBPL1, PLAG1, ZFP345, KLF6, ZFP605, ACACA, S100A11, ZFP449, SMYD4, HDDC3, CSRP1, GM20939, ZFP445, CSRP3, ZFP608, FOXP2, ZFP809, TRIM56, ZFP808, CDKN1A, ZFP941, ZFP770, MTR, ZFP773, SYTL5, S100G, ZFP947, ZIK1, GM10324, CLEC7A, ATP7B, TAF1B, ZKSCAN7, ZMAT1, STEAP4, GM14124, ZFP13, ZKSCAN8, ARSG, HR, ZKSCAN3, ZFP872, ZFP870, ZKSCAN4, FANCL, LNPEP, ZFP873, TPO, ATP8B5, NT5E, PRKCA, ZFP518A, GM14139, ZFP426, CAR14, CAPSL, VIL1, ESR1, RSLCAN18, MMP15, MBD1, CHAMP1, ZFP26, PMM1, MMP12, AU041133, ZFP558, ZSCAN18, ZFP369, ZFP229, DPF3, MOB1A, ZFP719, ZFP850, POLA1, CDH1, NHLRC1, 2010315B03RIK, TPM4, RPA1, TRIM30A, NPTX1, ANXA8, NAIP2, ZFP667, 2410141K09RIK, TGM1, PPP3CC, LIMD2, EHD1, ZC3H12D, ZFP72, BGLAP3, DTX4, ZFP53, BGLAP, SLC8A1, PNLIPRP1, ZFP955B, ZFP955A, ACLY, TRIM24, EXO5, PCK2, ZFP709, COL5A2, TRIM21, AJUBA, MARCH1, QPCT, AFP, ADNP2, ZFP105, PRLR, ZFP677, C1S1, LRP2 | 575 | 3355 | 17446 | 1,49 | 1,86E-05 | 9,29E-06 | 3,80E-05 |
| GOTERM_M | GO:0003700~transcription factor activity, sequence-specific DNA binding | 44 | 6,21 | 6,74E-03 | ZKSCAN7, GM14124, ZFP13, THRA, ZKSCAN8, ZFP850, ZKSCAN3, ZKSCAN4, ZFP935, ZFP934, ZFP873, FOXQ1, 2410141K09RIK, ZFP72, ZFP518A, PLAG1, ZFP345, KLF6, ZFP53, GM14139, RFX5, TAF7, ESR1, ZFP449, RSLCAN18, BMYC, ZFP445, MYCL, ZFP26, FOXP2, BTBD8, ZFP808, ZFP105, ZFP941, TCFL5, ZFP770, MEOX1, ZFP677, MGA, GM10324, ZIK1, GM3604, ZFP229, ZFP369 | 575 | 883 | 17446 | 1,51 | 9,93E-01 | 5,67E-01 | 9,77E+00 |

| | | | | | | | | | | | | |
|---|---|---|---|---|---|---|---|---|---|---|---|---|
| GOTERM_BI | GO:0006355~regulation of transcription, DNA-templated | 94 | 13,3 | 1,10E-02 | TAF1B, ZFP13, GM14124, 5730507C01RIK, THRA, HR, TCEAL8, ZFP872, ZKSCAN3, ZFP84, ZFP870, ZFP935, ZFP934, ZFP873, ZFP781, ZFP933, RNF141, LBH, ZFP882, ZFP938, ZFP518A, ZFP426, GM14139, LRIF1, ESR1, RSLCAN18, ZFP78, MBD1, MYCL, ZFP26, ZFP799, TCFL5, BTG2, ZFP120, NAB2, ZFP791, AU041133, MGA, ZFP558, GM3604, ATXN1L, ZFP229, ZFP369, ZFP266, DPF3, EID1, ZFP719, ZFP850, ZFP763, ZFP759, ZFP563, 2010315B03RIK, TRIM30A, ZFP952, FOXQ1, ZFP951, ZFP667, DMD, ZFP157, 2410141K09RIK, 1700048O20RIK, ZFP72, PLAG1, KLF6, ZFP345, ZFP53, ZFP605, RFX5, ZFP955B, ZFP955A, TAF7, BMYC, TRIM24, GM20939, ZFP445, 6720489N17RIK, ZFP709, FOXP2, AJUBA, ZFP809, 2410018L13RIK, ZFP808, ADNP2, ZFP941, ID2, ZFP770, MEOX1, ZFP677, 9030624G23RIK, ZFP773, ZFP947, ZIK1, GM10324, IFI205 | 588 | 2279 | 18082 | 1,27 | 1,00E+00 | 7,63E-01 | 1,77E+01 |
| Annotation | Enrichment Score: 5.777288767228272 | | | | | | | | | | | |
| Category | Term | Count | % | PValue | Genes | List To | Pop Hit | Pop Tota | Fold Enr | Bonferroni | Benjamini | FDR |
| UP_SEQ_FE/ | signal peptide | 145 | 20,5 | 3,72E-08 | BMP10, WFDC15B, KLK1B4, VTCN1, OBP2A, TLR3, FSTL1, RETNLG, TLR7, TLR8, CXCL10, IL17RB, UNC5B, ROBO1, CD46, SERPINE1, LGI4, SPON2, GRID1, HHIPL2, AMY1, CLCA1, C4A, PLXNB1, CRISP2, TMIE, COLEC10, ACRV1, MFGE8, PRELP, NCAM2, WFDC12, CD34, LRP11, CD300LB, WFDC2, SMIM24, GLG1, FXYD3, CCL2, TMX3, ELN, FAM19A2, RIC3, MANSC1, FXYD5, SERPINH1, TIMP1, ALCAM, ACE, CHIL1, BCHE, IMPG2, TFF2, SFTPD, TFF3, DEFB1, SPP2, SPP1, H2-Q2, IL1RN, H2-Q1, CD276, HGF, COL14A1, CXCL14, FAM198A, ECSCR, IL3RA, BMP8B, BMP5, SKINT9, GM2A, ARSG, MPEG1, LEPR, HEXB, PRSS41, SORL1, DNAJC10, LY9, GABBR2, CXADR, SPINK5, GPC2, TNFRSF11B, CD93, SEMA3D, CSF2RB, TPO, ITIH5, ANGPT1, CALCRL, NT5E, CLMP, CNTN6, CAR14, DHRS9, MMP15, POGLUT1, MMP12, INHBA, SMO, AMIGO2, SDC1, IL18BP, H2-BL, SLPI, SPACA1, CR1L, SEMA4A, CD244, MEGF9, LUM, CDH1, DCN, VCAM1, LY6A, NPTX1, LY6D, LY6E, NUP210, CATSPERG2, BGLAP3, SELP, COL4A3, PTPRC, BGLAP, PNLIPRP1, SLC8A1, HAPLN4, CR2, IL7, FLT4, NID1, SLC10A5, FUCA2, COL5A2, QPCT, AFP, PRLR, SCN4B, C1S1, LRP2, ORM3 | 544 | 3124 | 18012 | 1,54 | 5,78E-05 | 5,78E-05 | 6,21E-05 |

| | | | | | | | | | | | | |
|---|---|---|---|---|---|---|---|---|---|---|---|---|
| UP_KEYWO | Glycoprotein | 166 | 23,4 | 2,34E-07 | BMP10, SYT1, IMPAD1, SLC13A5, AQP8, VTCN1, OBP2A, TLR3, FSTL1, TSPAN9, SLC7A4, CD209C, ADORA1, TLR7, KLHDC7A, TLR8, IL17RB, UNC5B, ROBO1, CD46, SERPINE1, LGI4, SPON2, GRID1, MRGPRB2, HHIPL2, CLCA1, SLC22A27, PLXNB1, COLEC10, MFGE8, ABCC12, TMEM209, PRELP, NCAM2, SSTR2, ABCB1A, CCR5, CD34, LRP11, EMP2, CD300LB, EMP1, GLG1, CCL2, POMK, TMX3, TARM1, MANSC1, FXYD5, SERPINH1, TIMP1, ALCAM, ACE, P2RY4, CHIL1, BCHE, IMPG2, SFTPD, GAL3ST1, SPP1, IL1RN, CD276, HGF, CD63, TNFSF10, GPR35, COL14A1, ECSCR, SLC13A3, CLEC7A, IL3RA, BMP5, BMP8B, SKINT9, GM2A, ARSG, MPEG1, LEPR, HEXB, PRSS41, SORL1, DNAJC10, LY9, GABBR2, CXADR, LNPEP, GPC2, TNFRSF11B, CD93, SEMA3D, CSF2RB, TPO, ITIH5, ANGPT1, NIPAL1, CALCRL, NT5E, GOLM1, SLC43A2, VSIG10, CLMP, CNTN6, CAR14, ESR1, COL25A1, MMP15, LMBRD2, POGLUT1, MMP12, RDH5, AMIGO2, SMO, INHBA, SDC1, IL18BP, SPACA1, CR1L, SEMA4A, SGCB, TNFRSF22, MEGF9, CD244, LUM, ABHD2, CDH1, DCN, VCAM1, LY6A, NPTX1, LY6D, LY6E, NUP210, ABCD2, MAS1, SUCNR1, CATSPERG2, KCNE3, TMPRSS2, SELP, COL4A3, PTPRC, HAPLN4, SLC8A1, CR2, IL7, B3GALT1, FLT4, NID1, SLC10A5, SLC10A2, FUCA2, COL5A2, TMPRSS4, QPCT, SLC17A8, AFP, P2RY13, PRLR, SLC5A9, SCN4B, PTCH1, C1S1, LRP2, SLC15A3, ORM3 | 678 | 3815 | 22680 | 1,46 | 7,80E-05 | 1,95E-05 | 3,18E-04 |
| UP_SEQ_FE | glycosylation site:N-linked (GlcNAc...) | 153 | 21,6 | 1,80E-06 | S100A4, BMP10, SYT1, SLC13A5, AQP8, VTCN1, OBP2A, TLR3, FSTL1, TSPAN9, SLC7A4, CD209C, ADORA1, TLR7, KLHDC7A, TLR8, IL17RB, UNC5B, ROBO1, CD46, SERPINE1, LGI4, GRID1, MRGPRB2, HHIPL2, CLCA1, C4A, PLXNB1, COLEC10, MFGE8, ABCC12, TMEM209, PRELP, NCAM2, SSTR2, ABCB1A, CD34, LRP11, CD300LB, EMP2, EMP1, GLG1, CCL2, TMX3, MANSC1, SERPINH1, TIMP1, ALCAM, ACE, P2RY4, CHIL1, BCHE, IMPG2, SFTPD, GAL3ST1, SPP1, H2-Q2, IL1RN, H2-Q1, CD276, HGF, CD63, TNFSF10, GPR35, COL14A1, SLC13A3, CLEC7A, IL3RA, BMP5, BMP8B, SKINT9, GM2A, ARSG, MPEG1, LEPR, HEXB, PRSS41, SORL1, DNAJC10, LY9, GABBR2, CXADR, GM9992, LNPEP, TNFRSF11B, CD93, SEMA3D, CSF2RB, TPO, ITIH5, ANGPT1, NIPAL1, CALCRL, NT5E, GOLM1, SLC43A2, CLMP, CNTN6, CAR14, MMP15, LMBRD2, MMP12, AMIGO2, INHBA, SMO, SDC1, IL18BP, SPACA1, CR1L, SEMA4A, SGCB, TNFRSF22, MEGF9, CD244, CDH1, DCN, VCAM1, NPTX1, LY6E, NUP210, ABCD2, MAS1, SUCNR1, CATSPERG2, KCNE3, TMPRSS2, SELP, COL4A3, PTPRC, HAPLN4, SLC8A1, CR2, IL7, B3GALT1, FLT4, NID1, SLC10A5, SLC10A2, FUCA2, COL5A2, TMPRSS4, QPCT, SLC17A8, AFP, P2RY13, PRLR, SLC5A9, SCN4B, PTCH1, C1S1, LRP2, SLC15A3, ORM3 | 544 | 3563 | 18012 | 1,42 | 2,80E-03 | 1,40E-03 | 3,01E-03 |

| | | | | | | | | | | | | |
|---|---|---|---|---|---|---|---|---|---|---|---|---|
| UP_KEYWO | Disulfide bond | 137 | 19,3 | 2,69E-06 | BMP10, WFDC15B, KLK1B4, VTCN1, OBP2A, TLR3, SLC7A9, FSTL1, ADORA1, CD209C, TLR8, CXCL10, UNC5B, ROBO1, CD46, SPON2, RAB27A, HHIPL2, AMY1, C4A, PLXNB1, CRISP2, COLEC10, MFGE8, PRELP, NCAM2, WFDC12, ADAMTS6, SSTR2, ADAMTS9, CCR5, LRP11, CCR2, CD300LB, WFDC2, CCL2, TMX3, ELN, TARM1, TIMP1, ALCAM, ACE, P2RY4, CHIL1, BCHE, IMPG2, TFF2, SFTPD, TFF3, H2-T10, DEFB1, SPP2, H2-Q2, H2-Q4, MICU2, IL1RN, H2-Q1, CD276, S100A11, HGF, GPR35, COL14A1, CXCL14, CLEC7A, KLRB1B, BMP8B, BMP5, SKINT9, GM2A, LEPR, PRSS41, SORL1, HEXB, DNAJC10, LY9, GABBR2, CXADR, TNFRSF11B, CD93, SEMA3D, CSF2RB, TPO, ANGPT1, CALCRL, NT5E, VSIG10, CLMP, CNTN6, CAR14, MMP15, MMP12, BTNL4, INHBA, SMO, AMIGO2, MUP21, IL18BP, H2-BL, CLDN1, SLPI, CR1L, SEMA4A, SGCB, TNFRSF22, CD244, MEGF9, LUM, CDH1, DCN, VCAM1, LY6A, NPTX1, LY6D, LY6E, PRRG1, SUCNR1, BGLAP3, SELP, COL4A3, TMPRSS2, BGLAP, PNLIPRP1, HAPLN4, CR2, IL7, FLT4, NID1, COL5A2, TMPRSS4, QPCT, P2RY13, AFP, PRLR, SCN4B, C1S1, LRP2, ORM3 | 678 | 3124 | 22680 | 1,47 | 8,95E-04 | 1,49E-04 | 3,65E-03 |
| UP_SEQ_FE | disulfide bond | 113 | 15,9 | 9,23E-06 | BMP10, WFDC15B, KLK1B4, VTCN1, OBP2A, TLR3, FSTL1, ADORA1, CD209C, CXCL10, UNC5B, ROBO1, CD46, SPON2, RAB27A, AMY1, C4A, CRISP2, COLEC10, MFGE8, PRELP, WFDC12, NCAM2, SSTR2, CCR5, LRP11, CCR2, CD300LB, WFDC2, CCL2, TMX3, ELN, TIMP1, ALCAM, ACE, P2RY4, CHIL1, BCHE, NMRAL1, IMPG2, TFF2, SFTPD, TFF3, DEFB1, SPP2, H2-Q2, H2-Q1, IL1RN, CD276, S100A11, HGF, GPR35, CXCL14, CLEC7A, KLRB1B, BMP8B, BMP5, SKINT9, GM2A, LEPR, PRSS41, SORL1, HEXB, LY9, CXADR, TNFRSF11B, CD93, SEMA3D, CSF2RB, TPO, ANGPT1, CLMP, CNTN6, CAR14, MMP15, MMP12, INHBA, SMO, AMIGO2, IL18BP, SLPI, CR1L, SEMA4A, SGCB, TNFRSF22, CD244, MEGF9, LUM, DCN, VCAM1, LY6A, NPTX1, LY6D, LY6E, SUCNR1, BGLAP3, COL4A3, SELP, TMPRSS2, BGLAP, PNLIPRP1, HAPLN4, CR2, FLT4, NID1, TMPRSS4, P2RY13, AFP, PRLR, SCN4B, LRP2, C1S1, ORM3 | 544 | 2510 | 18012 | 1,49 | 1,42E-02 | 4,77E-03 | 1,54E-02 |
| UP_KEYWO | Signal | 178 | 25,1 | 5,55E-05 | CXCL10, IL17RB, UNC5B, ROBO1, CD46, SERPINE1, BC089597, RAPGEF5, LGI4, SPON2, GRID1, HHIPL2, AMY1, CLCA1, C4A, PLXNB1, CRISP2, TMIE, COLEC10, CYP2B13, ACRV1, MFGE8, CYP2B10, PRELP, NCAM2, WFDC12, ADAMTS6, ADAMTS9, ZFP120, CD34, LRP11, CD300LB, WFDC2, SMIM24, GLG1, GM1976, FXYD3, CCL2, PKHD1, ELN, TMX3, FAM19A2, TARM1, RIC3, MANSC1, FXYD5, SERPINH1, TIMP1, ALCAM, ACE, CHIL1, BCHE, IMPG2, TFF2, SFTPD, STAMBPL1, TFF3, H2-T10, DEFB1, SPP2, SPP1, H2-Q2, H2-Q4, IL1RN, H2-Q1, ACACA, CD276, HGF, 9030619P08RIK, GPR35, COL14A1, CXCL14, FAM198A, IFI27L2B, PPIC, ECSCR, IL3RA, BMP8B, BMP5, SKINT9, GM2A, MPEG1, ARSG, LEPR, CLDN6, HEXB, PRSS41, SORL1, DNAJC10, GABBR2, LY9, CXADR, SPINK5, PCDHGA2, GPC2, TNFRSF11B, CD93, SEMA3D, CSF2RB, TPO, ITIH5, ANGPT1, CALCRL, NT5E, VSIG10, CLMP, CNTN6, CAR14, DHRS9, MMP15, BTNL6, POGLUT1, BTNL4, MMP12, RDH5, AMIGO2, SMO, INHBA, SDC1, MUP21, LIPO1, IL18BP, ALDH1B1, H2-BL, SLPI, SPACA1, CR1L, SEMA4A, MEGF9, CD244, MOB1A, LUM, CDH1, DCN, VCAM1, LY6A, NPTX1, LY6D, LY6E, NUP210, GM609, B4GALNT3, CATSPERG2, BGLAP3, SELP, COL4A3, PTPRC, BGLAP, HAPLN4, SLC8A1, PNLIPRP1, CR2, IL7, FLT4, NID1, SLC10A5, FUCA2, COL5A2, QPCT, CYP2A22, AFP, PRLR, SCN4B, PTCH1, C1S1, LRP2, ORM3 | 678 | 4543 | 22680 | 1,31 | 1,83E-02 | 2,31E-03 | 7,54E-02 |

| | | | | | | | | | | | | |
|---|---|---|---|---|---|---|---|---|---|---|---|---|
| Annotation | Enrichment Score: 4.2034750242852095 | | | | | | | | | | | |
| Category | Term | Count | % | PValue | Genes | List To | Pop Hit | Pop Tot | Fold Enr | Bonferroni | Benjamini | FDR |
| UP_SEQ_FE | signal peptide | 145 | 20,5 | 3,72E-08 | BMP10, WFDC15B, KLK1B4, VTCN1, OBP2A, TLR3, FSTL1, RETNLG, TLR7, TLR8, CXCL10, IL17RB, UNC5B, ROBO1, CD46, SERPINE1, LGI4, SPON2, GRID1, HHIPL2, AMY1, CLCA1, C4A, PLXNB1, CRISP2, TMIE, COLEC10, ACRV1, MFGE8, PRELP, NCAM2, WFDC12, CD34, LRP11, CD300LB, WFDC2, SMIM24, GLG1, FXYD3, CCL2, TMX3, ELN, FAM19A2, RIC3, MANSC1, FXYD5, SERPINH1, TIMP1, ALCAM, ACE, CHIL1, BCHE, IMPG2, TFF2, SFTPD, TFF3, DEFB1, SPP2, SPP1, H2-Q2, IL1RN, H2-Q1, CD276, HGF, COL14A1, CXCL14, FAM198A, ECSCR, IL3RA, BMP8B, BMP5, SKINT9, GM2A, ARSG, MPEG1, LEPR, HEXB, PRSS41, SORL1, DNAJC10, LY9, GABBR2, CXADR, SPINK5, GPC2, TNFRSF11B, CD93, SEMA3D, CSF2RB, TPO, ITIH5, ANGPT1, CALCRL, NT5E, CLMP, CNTN6, CAR14, DHRS9, MMP15, POGLUT1, MMP12, INHBA, SMO, AMIGO2, SDC1, IL18BP, H2-BL, SLPI, SPACA1, CR1L, SEMA4A, CD244, MEGF9, LUM, CDH1, DCN, VCAM1, LY6A, NPTX1, LY6D, LY6E, NUP210, CATSPERG2, BGLAP3, SELP, COL4A3, PTPRC, BGLAP, PNLIPRP1, SLC8A1, HAPLN4, CR2, IL7, FLT4, NID1, SLC10A5, FUCA2, COL5A2, QPCT, AFP, PRLR, SCN4B, C1S1, LRP2, ORM3 | 544 | 3124 | 18012 | 1,54 | 5,78E-05 | 5,78E-05 | 6,21E-05 |
| GOTERM_C | GO:0005615~extracellular space | 75 | 10,6 | 1,56E-04 | S100A4, BMP10, KLK1B4, ARSG, LEPR, 2010005H15RIK, SORL1, HEXB, FSTL1, CNP, RETNLG, CXCL10, ACTG1, TNFRSF11B, GPC2, SERPINE1, SEMA3D, LGI4, TPO, STAG3, ANGPT1, SPON2, GOLM1, AMY1, CLCA1, C4A, COL25A1, MFGE8, PRELP, INHBA, ADAMTS9, IL18BP, SERPINB8, SERPINA3I, SLPI, SEMA4A, CCL2, LUM, SERPINB1A, IL33, DCN, SERPINH1, TIMP1, VCAM1, ACE, CHIL1, BCHE, TFF2, SFTPD, TFF3, DEFB1, BGLAP3, SPP1, SELP, BGLAP, PNLIPRP1, E130311K13RIK, IL7, LGALS1, IL1RN, S100A11, HGF, FUCA2, ANXA5, CD63, ANXA2, AFP, TNFSF10, COL14A1, CXCL14, ANXA13, LRP2, BMP5, BMP8B, ORM3 | 633 | 1504 | 19662 | 1,55 | 6,00E-02 | 1,54E-02 | 2,17E-01 |
| UP_KEYWO | Secreted | 73 | 10,3 | 1,33E-03 | BMP10, WFDC15B, TUFT1, OBP2A, LEPR, SORL1, FSTL1, CXADR, IL17RB, CXCL10, TNFRSF11B, GPC2, CD46, SERPINE1, SEMA3D, ITIH5, LGI4, ANGPT1, SPON2, HHIPL2, AMY1, CLCA1, C4A, CRISP2, COLEC10, MFGE8, MMP12, PRELP, WFDC12, INHBA, MUP21, SDC1, IL18BP, SLPI, WFDC2, TNFRSF22, CCL2, LUM, ELN, IL33, DCN, TIMP1, ACE, CHIL1, BCHE, TFF2, SFTPD, TFF3, DEFB1, SPP2, BGLAP3, SPP1, COL4A3, TMPRSS2, BGLAP, PNLIPRP1, HAPLN4, IL7, LGALS1, IL1RN, NID1, FUCA2, CD63, COL5A2, ANXA2, QPCT, AFP, COL14A1, CXCL14, FAM198A, BMP5, BMP8B, ORM3 | 678 | 1685 | 22680 | 1,45 | 3,58E-01 | 3,95E-02 | 1,79E+00 |

| | | | | | | | | | | | | |
|---|---|---|---|---|---|---|---|---|---|---|---|---|
| GOTERM_C | GO:0005576~extracellular region | 79 | 11,1 | 1,99E-03 | BMP10, WFDC15B, TUFT1, OBP2A, LEPR, SORL1, FSTL1, CXADR, SPINK5, IL17RB, CXCL10, TNFRSF11B, GPC2, CD46, SERPINE1, SEMA3D, ITIH5, LGI4, ANGPT1, SPON2, HHIPL2, AMY1, CLCA1, C4A, CRISP2, COLEC10, COL25A1, MFGE8, MMP12, PRELP, WFDC12, INHBA, MUP21, SDC1, IL18BP, CD34, SLPI, WFDC2, TNFRSF22, CCL2, LUM, ELN, IL33, DCN, TIMP1, ACE, PRRG1, CHIL1, BCHE, TFF2, SFTPD, TFF3, DEFB1, SPP2, BGLAP3, SPP1, COL4A3, TMPRSS2, BGLAP, HAPLN4, PNLIPRP1, IL7, LGALS1, IL1RN, NID1, HGF, FUCA2, CD63, COL5A2, ANXA2, QPCT, AFP, COL14A1, CXCL14, FAM198A, PTCH1, C1S1, BMP5, BMP8B | 633 | 1753 | 19662 | 1,40 | 5,48E-01 | 1,24E-01 | 2,74E+00 |
| Annotation | Enrichment Score: 4.184529219386322 | | | | | | | | | | | |
| Category | Term | Count | % | PValue | Genes | List To | Pop Hit | Pop Tota | Fold Enr | Bonferroni | Benjamini | FDR |
| UP_KEYWO | Immunity | 33 | 4,65 | 5,17E-07 | CD244, VTCN1, TLR3, LY9, TLR7, TLR8, SRC, ALCAM, NAIP2, TAP1, OASL1, SFTPD, SPON2, ZBP1, H2-Q2, TRPM4, DAB2IP, CR2, THEMIS, HERC6, H2-Q1, PSMB9, MARCH1, IFIT2, TRIM56, SLPI, TGTP1, OAS1A, CLEC7A, C1S1, CD300LB, SEMA4A, CR1L | 678 | 401 | 22680 | 2,75 | 1,72E-04 | 3,44E-05 | 7,02E-04 |
| UP_KEYWO | Innate immunity | 21 | 2,96 | 4,01E-05 | CD244, DAB2IP, CR2, HERC6, TLR3, LY9, TLR7, TLR8, TRIM56, IFIT2, NAIP2, OASL1, SFTPD, SLPI, TGTP1, CLEC7A, OAS1A, C1S1, SPON2, CR1L, ZBP1 | 678 | 241 | 22680 | 2,91 | 1,33E-02 | 1,91E-03 | 5,45E-02 |
| GOTERM_B | GO:0002376~immune system process | 28 | 3,95 | 1,46E-04 | CD244, VTCN1, TLR3, LY9, TLR7, TLR8, SRC, ALCAM, NAIP2, TAP1, OASL1, SFTPD, SPON2, ZBP1, TRPM4, DAB2IP, CR2, THEMIS, HERC6, PSMB9, MARCH1, IFIT2, TRIM56, SLPI, C1S1, CD300LB, SEMA4A, CR1L | 588 | 383 | 18082 | 2,25 | 2,89E-01 | 1,57E-01 | 2,55E-01 |
| GOTERM_B | GO:0045087~innate immune response | 24 | 3,39 | 6,03E-03 | CD244, DAB2IP, CR2, HERC6, TLR3, LY9, TLR7, SRC, TRIM21, TLR8, ZFP809, TRIM56, IFIT2, NAIP2, OASL1, SFTPD, SLPI, CLEC7A, OAS1A, C1S1, SPON2, DEFB1, CR1L, ZBP1 | 588 | 400 | 18082 | 1,85 | 1,00E+00 | 6,91E-01 | 1,00E+01 |
| Annotation | Enrichment Score: 3.3833934412272066 | | | | | | | | | | | |
| Category | Term | Count | % | PValue | Genes | List To | Pop Hit | Pop Tota | Fold Enr | Bonferroni | Benjamini | FDR |
| INTERPRO | IPR003309:Transcription regulator SCAN | 8 | 1,13 | 2,95E-04 | ZKSCAN7, ZKSCAN8, ZFP449, ZSCAN18, ZFP369, ZKSCAN3, ZFP445, ZKSCAN4 | 645 | 42 | 20594 | 6,08 | 2,72E-01 | 4,43E-02 | 4,69E-01 |
| INTERPRO | IPR008916:Retrovirus capsid, C-terminal | 8 | 1,13 | 3,42E-04 | ZKSCAN7, ZKSCAN8, ZFP449, ZSCAN18, ZFP369, ZKSCAN3, ZFP445, ZKSCAN4 | 645 | 43 | 20594 | 5,94 | 3,09E-01 | 4,51E-02 | 5,45E-01 |
| SMART | SM00431:SCAN | 8 | 1,13 | 7,02E-04 | ZKSCAN7, ZKSCAN8, ZFP449, ZSCAN18, ZFP369, ZKSCAN3, ZFP445, ZKSCAN4 | 379 | 42 | 10425 | 5,24 | 1,47E-01 | 5,15E-02 | 8,93E-01 |
| Annotation | Enrichment Score: 2.610010363501475 | | | | | | | | | | | |
| Category | Term | Count | % | PValue | Genes | List To | Pop Hit | Pop Tota | Fold Enr | Bonferroni | Benjamini | FDR |
| INTERPRO | IPR003191:Guanylate-binding protein, C-terminal | 7 | 0,99 | 2,21E-06 | GBP8, GBP7, GBP6, GBP9, GBP10, GBP3, GBP2 | 645 | 14 | 20594 | 15,96 | 2,38E-03 | 4,77E-04 | 3,53E-03 |

| | | | | | | | | | | | | |
|---|---|---|---|---|---|---|---|---|---|---|---|---|
| INTERPRO | IPR015894:Guanylate-binding protein, N-terminal | 7 | 0,99 | 3,59E-06 | GBP8, GBP7, GBP6, GBP9, GBP10, GBP3, GBP2 | 645 | 15 | 20594 | 14,90 | 3,87E-03 | 6,45E-04 | 5,73E-03 |
| GOTERM_C | GO:0020005~symbiont-containing vacuole membrane | 5 | 0,71 | 6,68E-05 | GBP7, GBP6, GBP9, GBP3, GBP2 | 633 | 8 | 19662 | 19,41 | 2,62E-02 | 8,82E-03 | 9,30E-02 |
| GOTERM_B | GO:0044406~adhesion of symbiont to host | 5 | 0,71 | 1,22E-04 | GBP7, GBP6, GBP9, GBP3, GBP2 | 588 | 9 | 18082 | 17,08 | 2,47E-01 | 2,47E-01 | 2,13E-01 |
| GOTERM_B | GO:0071346~cellular response to interferon-gamma | 9 | 1,27 | 1,57E-03 | GBP8, GBP7, GBP6, CCL2, GBP9, GBP10, TLR3, GBP3, GBP2 | 588 | 68 | 18082 | 4,07 | 9,74E-01 | 4,56E-01 | 2,71E+00 |
| GOTERM_B | GO:0035458~cellular response to interferon-beta | 7 | 0,99 | 1,95E-03 | 9930111J21RIK1, GBP6, TLR3, TGTP1, GBP3, GBP2, IFI205 | 588 | 41 | 18082 | 5,25 | 9,89E-01 | 4,78E-01 | 3,36E+00 |
| GOTERM_B | GO:0042832~defense response to protozoan | 6 | 0,85 | 2,58E-03 | GBP7, GBP6, GBP9, GBP10, GBP3, GBP2 | 588 | 30 | 18082 | 6,15 | 9,98E-01 | 4,88E-01 | 4,42E+00 |
| GOTERM_B | GO:0050830~defense response to Gram-positive bacterium | 7 | 0,99 | 8,20E-02 | GBP7, GBP6, GBP9, GBP10, DEFB1, GBP3, GBP2 | 588 | 93 | 18082 | 2,31 | 1,00E+00 | 9,70E-01 | 7,76E+01 |
| INTERPRO | IPR027417:P-loop containing nucleoside triphosphate hydrolase | 36 | 5,08 | 1,19E-01 | 9930111J21RIK1, RAD51B, MYO7B, TDRD9, DQX1, CTPS2, CNP, SAMD9L, DNAH5, SULT4A1, NAIP2, TAP1, RASL10B, ABCD2, GBP10, RHOC, EHD1, PAPSS2, GAL3ST1, RAB27A, GBP8, RABL2, GBP7, GBP6, GBP9, IFI44, ABCC12, SMC2, NUGGC, ABCB1A, SULT1B1, TGTP1, THNSL1, GBP3, GBP2, MYH10 | 645 | 909 | 20594 | 1,26 | 1,00E+00 | 9,61E-01 | 8,67E+01 |
| UP_KEYWO | GTP-binding | 14 | 1,97 | 1,94E-01 | RABL2, GBP8, GBP7, GBP6, GBP9, PCK2, NUGGC, RASL10B, GBP10, TGTP1, RHOC, GBP3, GBP2, RAB27A | 678 | 332 | 22680 | 1,41 | 1,00E+00 | 7,16E-01 | 9,46E+01 |
| GOTERM_M | GO:0005525~GTP binding | 17 | 2,4 | 1,97E-01 | 9930111J21RIK1, GBP8, RABL2, GBP7, GBP6, GBP9, PCK2, NUGGC, P2RX5, RASL10B, GBP10, TGTP1, RHOC, EHD1, GBP3, GBP2, RAB27A | 575 | 383 | 17446 | 1,35 | 1,00E+00 | 9,71E-01 | 9,65E+01 |
| GOTERM_M | GO:0003924~GTPase activity | 10 | 1,41 | 2,53E-01 | 9930111J21RIK1, GBP8, GBP7, GBP6, GBP9, GBP10, TGTP1, GBP3, GBP2, RAB27A | 575 | 209 | 17446 | 1,45 | 1,00E+00 | 9,78E-01 | 9,88E+01 |
| Annotation | Enrichment Score: 2.536060067955947 | | | | | | | | | | | |
| Category | Term | Count | % | PValue | Genes | List To | Pop Hit | Pop Tota | Fold Enr | Bonferroni | Benjamini | FDR |

| | | | | | | | | | | | | |
|---|---|---|---|---|---|---|---|---|---|---|---|---|
| UP_KEYWO | Glycoprotein | 166 | 23,4 | 2,34E-07 | BMP10, SYT1, IMPAD1, SLC13A5, AQP8, VTCN1, OBP2A, TLR3, FSTL1, TSPAN9, SLC7A4, CD209C, ADORA1, TLR7, KLHDC7A, TLR8, IL17RB, UNC5B, ROBO1, CD46, SERPINE1, LGI4, SPON2, GRID1, MRGPRB2, HHIPL2, CLCA1, SLC22A27, PLXNB1, COLEC10, MFGE8, ABCC12, TMEM209, PRELP, NCAM2, SSTR2, ABCB1A, CCR5, CD34, LRP11, EMP2, CD300LB, EMP1, GLG1, CCL2, POMK, TMX3, TARM1, MANSC1, FXYD5, SERPINH1, TIMP1, ALCAM, ACE, P2RY4, CHIL1, BCHE, IMPG2, SFTPD, GAL3ST1, SPP1, IL1RN, CD276, HGF, CD63, TNFSF10, GPR35, COL14A1, ECSCR, SLC13A3, CLEC7A, IL3RA, BMP5, BMP8B, SKINT9, GM2A, ARSG, MPEG1, LEPR, HEXB, PRSS41, SORL1, DNAJC10, LY9, GABBR2, CXADR, LNPEP, GPC2, TNFRSF11B, CD93, SEMA3D, CSF2RB, TPO, ITIH5, ANGPT1, NIPAL1, CALCRL, NT5E, GOLM1, SLC43A2, VSIG10, CLMP, CNTN6, CAR14, ESR1, COL25A1, MMP15, LMBRD2, POGLUT1, MMP12, RDH5, AMIGO2, SMO, INHBA, SDC1, IL18BP, SPACA1, CR1L, SEMA4A, SGCB, TNFRSF22, MEGF9, CD244, LUM, ABHD2, CDH1, DCN, VCAM1, LY6A, NPTX1, LY6D, LY6E, NUP210, ABCD2, MAS1, SUCNR1, CATSPERG2, KCNE3, TMPRSS2, SELP, COL4A3, PTPRC, HAPLN4, SLC8A1, CR2, IL7, B3GALT1, FLT4, NID1, SLC10A5, SLC10A2, FUCA2, COL5A2, TMPRSS4, QPCT, SLC17A8, AFP, P2RY13, PRLR, SLC5A9, SCN4B, PTCH1, C1S1, LRP2, SLC15A3, ORM3 | 678 | 3815 | 22680 | 1,46 | 7,80E-05 | 1,95E-05 | 3,18E-04 |
| UP_SEQ_FE | glycosylation site:N-linked (GlcNAc...) | 153 | 21,6 | 1,80E-06 | S100A4, BMP10, SYT1, SLC13A5, AQP8, VTCN1, OBP2A, TLR3, FSTL1, TSPAN9, SLC7A4, CD209C, ADORA1, TLR7, KLHDC7A, TLR8, IL17RB, UNC5B, ROBO1, CD46, SERPINE1, LGI4, GRID1, MRGPRB2, HHIPL2, CLCA1, C4A, PLXNB1, COLEC10, MFGE8, ABCC12, TMEM209, PRELP, NCAM2, SSTR2, ABCB1A, CD34, LRP11, CD300LB, EMP2, EMP1, GLG1, CCL2, TMX3, MANSC1, SERPINH1, TIMP1, ALCAM, ACE, P2RY4, CHIL1, BCHE, IMPG2, SFTPD, GAL3ST1, SPP1, H2-Q2, IL1RN, H2-Q1, CD276, HGF, CD63, TNFSF10, GPR35, COL14A1, SLC13A3, CLEC7A, IL3RA, BMP5, BMP8B, SKINT9, GM2A, ARSG, MPEG1, LEPR, HEXB, PRSS41, SORL1, DNAJC10, LY9, GABBR2, CXADR, GM9992, LNPEP, TNFRSF11B, CD93, SEMA3D, CSF2RB, TPO, ITIH5, ANGPT1, NIPAL1, CALCRL, NT5E, GOLM1, SLC43A2, CLMP, CNTN6, CAR14, MMP15, LMBRD2, MMP12, AMIGO2, INHBA, SMO, SDC1, IL18BP, SPACA1, CR1L, SEMA4A, SGCB, TNFRSF22, MEGF9, CD244, CDH1, DCN, VCAM1, NPTX1, LY6E, NUP210, ABCD2, MAS1, SUCNR1, CATSPERG2, KCNE3, TMPRSS2, SELP, COL4A3, PTPRC, HAPLN4, SLC8A1, CR2, IL7, B3GALT1, FLT4, NID1, SLC10A5, SLC10A2, FUCA2, COL5A2, TMPRSS4, QPCT, SLC17A8, AFP, P2RY13, PRLR, SLC5A9, SCN4B, PTCH1, C1S1, LRP2, SLC15A3, ORM3 | 544 | 3563 | 18012 | 1,42 | 2,80E-03 | 1,40E-03 | 3,01E-03 |

| | | | | | | | | | | | | |
|---|---|---|---|---|---|---|---|---|---|---|---|---|
| UP_SEQ_FE/ | topological domain:Extracellular | 103 | 14,5 | 1,48E-05 | S100A4, OCLN, AQP8, SLC7A9, AQP7, TSPAN9, TLR7, ADORA1, TLR8, IL17RB, ATP2B2, UNC5B, ROBO1, CD46, SLC51B, GRID1, MRGPRB2, PLXNB1, TMIE, NCAM2, SSTR2, ABCB1A, CCR5, CD34, LRP11, CCR2, CD300LB, GLG1, FXYD3, MANSC1, FXYD5, ALCAM, ACE, P2RY4, IMPG2, H2-Q2, H2-Q1, CD276, CD63, GPR35, TNFSF10, ECSCR, CLEC7A, KLRB1B, IL3RA, ATP7B, CLDN7, SKINT9, CLDN4, MPEG1, LEPR, CLDN6, SORL1, LY9, GABBR2, CXADR, LNPEP, CD93, CSF2RB, TPO, MMD2, NIPAL1, CALCRL, TRPM4, CLMP, CAR14, FMR1NB, COL25A1, MMP15, LMBRD2, SMO, AMIGO2, SDC1, ALG10B, CLDN1, SPACA1, SEMA4A, CR1L, SGCB, TNFRSF22, CD244, MEGF9, CDH1, SLC47A2, VCAM1, MAS1, CATSPERG2, SUCNR1, PTPRC, SELP, TMPRSS2, SLC8A1, CR2, FLT4, SLC10A5, SLC10A2, TMPRSS4, P2RY13, PRLR, SCN4B, SLC5A9, PTCH1, LRP2 | 544 | 2256 | 18012 | 1,51 | 2,27E-02 | 5,73E-03 | 2,47E-02 |
| UP_SEQ_FE/ | transmembrane region | 168 | 23,7 | 1,38E-04 | SLC7A4, ADORA1, TLR7, KLHDC7A, TLR8, IL17RB, ATP2B2, UNC5B, ROBO1, CD46, SLC51B, GRID1, MRGPRB2, SPTLC2, PLXNB1, TMIE, ABCC12, TMEM209, CNGA3, PARP16, NCAM2, SSTR2, ABCB1A, CCR5, CD34, CCR2, LRP11, VAMP5, TMEM184C, CD300LB, EMP2, EMP1, SMIM24, GLG1, FXYD3, POMK, TMX3, RIC3, MANSC1, FXYD5, ALCAM, ACE, P2RY4, IMPG2, SYBU, SLC35F2, SLC35F3, GAL3ST1, H2-Q2, 2010107G23RIK, SPNS3, H2-Q1, CD276, CD63, TNFSF10, GPR35, ATP6V0E2, UCP2, ECSCR, SLC13A3, CLEC7A, KLRB1B, IL3RA, D630045J12RIK, ATP7B, RTN4, CLDN7, STEAP4, SKINT9, CLDN4, MPEG1, LEPR, CLDN6, SORL1, TMEM237, LY9, DPY19L3, GABBR2, CXADR, GM9992, FAR1, LNPEP, CD93, AGPAT9, CSF2RB, TPO, NIPAL1, MMD2, SRD5A2, CALCRL, ATP8B5, GOLM1, SLC43A2, TRPM4, CLMP, SLC25A4, FMR1NB, CAR14, SLC22A7, COL25A1, MMP15, LMBRD2, MYADM, AMIGO2, SMO, SDC1, ALG10B, GM11437, CLIC5, CLDN1, SPACA1, CR1L, SEMA4A, SGCB, TNFRSF22, MEGF9, CD244, APH1B, ABHD2, CDH1, SLC47A2, VCAM1, MFSD7C, NUP210, TAP1, ABCD2, MAS1, B4GALNT3, SUCNR1, CATSPERG2, KCNE3, TMEM45A, TMPRSS2, NAT8, SELP, MOGAT2, PTPRC, SLC8A1, E130311K13RIK, CR2, TMEM45B, B3GALT1, FLT4, FADS3, SLC10A5, SLC10A2, TMPRSS4, MARCH1, SLC17A8, P2RY13, PRLR, TMEM261, SLC5A9, SCN4B, PTCH1, MBOAT4, LRP2, SLC15A3 | 544 | 4312 | 18012 | 1,29 | 1,93E-01 | 4,20E-02 | 2,30E-01 |
| UP_SEQ_FE/ | topological domain:Cytoplasmic | 116 | 16,4 | 7,29E-04 | S100A4, SYT1, OCLN, AQP8, TLR3, SLC7A9, AQP7, TSPAN9, TLR7, ADORA1, TLR8, IL17RB, ATP2B2, UNC5B, ROBO1, CD46, SLC51B, GRID1, MRGPRB2, PLXNB1, TMIE, NCAM2, SSTR2, ABCB1A, CCR5, CD34, LRP11, CCR2, VAMP5, CD300LB, GLG1, FXYD3, TMX3, RIC3, MANSC1, FXYD5, ALCAM, ACE, P2RY4, IMPG2, GAL3ST1, CD276, CD63, TNFSF10, GPR35, ECSCR, CLEC7A, KLRB1B, IL3RA, ATP7B, RTN4, CLDN7, SKINT9, CLDN4, MPEG1, LEPR, CLDN6, SORL1, LY9, GABBR2, CXADR, LNPEP, CD93, CSF2RB, TPO, MMD2, NIPAL1, CALCRL, GOLM1, TRPM4, CLMP, FMR1NB, CAR14, COL25A1, MMP15, LMBRD2, SMO, AMIGO2, SDC1, ALG10B, CLDN1, SPACA1, CR1L, SEMA4A, SGCB, TNFRSF22, CD244, MEGF9, CDH1, SLC47A2, VCAM1, NUP210, TAP1, MAS1, B4GALNT3, CATSPERG2, SUCNR1, KCNE3, PTPRC, SELP, TMPRSS2, SLC8A1, CR2, B3GALT1, FLT4, FADS3, SLC10A5, SLC10A2, TMPRSS4, P2RY13, SLC17A8, PRLR, SCN4B, SLC5A9, PTCH1, LRP2 | 544 | 2880 | 18012 | 1,33 | 6,78E-01 | 1,72E-01 | 1,21E+00 |

| | | | | | | | | | | | | |
|---|---|---|---|---|---|---|---|---|---|---|---|---|
| GOTERM_C( | GO:0016020~membrane | 257 | 36,2 | 5,21E-03 | RAB27A, MRGPRB2, GBP6, DAB2IP, TMIE, ABCC12, PARP16, SSTR2, CCR5, ABCB1A, CD34, CCR2, TMEM184C, GLG1, POMK, CYP2B9, TMX3, CCDC91, MANSC1, RIC3, ALCAM, ACE, DMD, IMPG2, SYBU, BLNK, SPNS3, OSBPL3, HGF, CD63, ATP6V0E2, RGS5, SYTL5, GRK5, KLRB1B, D630045J12RIK, RTN4, STEAP4, MPEG1, GRIP1, PRSS41, FERMT1, DNAJC10, LY9, LNPEP, FAR1, VEPH1, AGPAT9, MMD2, FAM129B, FAM129A, ATP6V0D2, NT5E, GOLM1, CLMP, SLC25A4, CAR14, FMR1NB, SLC22A7, COL25A1, MMP15, MYADM, RDH5, FMN1, AMIGO2, ALG10B, CLIC5, SPACA1, CR1L, SEMA4A, SGCB, MEGF9, SERPINB1A, ABHD2, LY6A, LY6D, LY6E, MFSD7C, NUP210, FMO2, FMO3, MAS1, ABCD2, IPCEF1, AGK, KCNE3, NAT8, TMPRSS2, MOGAT2, SELP, SLC8A1, E130311K13RIK, PLEK, FADS3, ACLY, TMPRSS4, MARCH1, P2RY13, SLC16A5, PTCH1, MBOAT4, LRP2, SLC15A3, GREB1L, FAM126A, PYGB, IMPAD1, OCLN, SLC13A5, SLC7A9, TLR3, CNP, TSPAN9, SLC7A4, TLR7, TLR8, IL17RB, RNF141, ANK2, UNC5B, ROBO1, MLKL, SLC51B, SLC22A27, SPTLC2, PLXNB1, CHP2, RPS6KC1, MFGE8, CNGA3, CYP2B10, TMEM209, NCAM2, SGSM1, HRASLS, LRP11, VAMP5, NEU3, EMP2, GBP3, CD300LB, GBP2, EMP1, SPTB, SMIM24, BBS4, FXYD3, STK10, NAPB, TARM1, FXYD5, SRC, P2RY4, BCHE, PPL, STAMBPL1, SLC35F2, SLC35F3, GAL3ST1, H2-Q2, 2010107G23RIK, H2-Q4, H2-Q1, CD276, TNFSF10, GPR35, UCP2, ECSCR, IFI27L2B, | 633 | 6998 | 19662 | 1,14 | 8,75E-01 | 2,06E-01 | 7,02E+00 |
| UP_KEYWO | Membrane | 268 | 37,8 | 2,78E-01 | GRID1, RAB27A, MRGPRB2, GBP6, DAB2IP, TMIE, GBP9, ABCC12, PARP16, SSTR2, CCR5, ABCB1A, CD34, CCR2, TMEM184C, GLG1, GM1976, POMK, CYP2B9, TMX3, CCDC91, MANSC1, RIC3, ALCAM, ACE, DMD, IMPG2, SYBU, OLFR1033, BLNK, SPNS3, OSBPL3, S100A10, CD63, ATP6V0E2, RGS5, SYTL5, GRK5, CLEC7A, KLRB1B, D630045J12RIK, RTN4, STEAP4, MPEG1, GRIP1, PRSS41, FERMT1, DNAJC10, LY9, LNPEP, FAR1, VEPH1, PBXIP1, AGPAT9, SEMA3D, ANGPT1, MMD2, FAM129B, FAM129A, NT5E, GOLM1, CLMP, SLC25A4, CAR14, FMR1NB, SLC22A7, COL25A1, MMP15, MYADM, RDH5, FMN1, AMIGO2, ALG10B, H2-BL, CLIC5, SPACA1, CR1L, SEMA4A, SGCB, MEGF9, 1810064F22RIK, ABHD2, LY6A, LY6D, LY6E, MFSD7C, NUP210, FMO2, FMO3, MAS1, ABCD2, IPCEF1, METTL7A3, AGK, KCNE3, NAT8, TMPRSS2, MOGAT2, SELP, SLC8A1, E130311K13RIK, FADS3, TMPRSS4, MARCH1, P2RY13, SLC16A5, CYP4F16, PTCH1, MBOAT4, LRP2, SLC15A3, GREB1L, FAM126A, IMPAD1, OCLN, SLC13A5, SLC7A9, TLR3, CNP, TSPAN9, SLC7A4, TLR7, TLR8, IL17RB, RNF141, ANK2, UNC5B, ROBO1, MLKL, SLC51B, SLC22A27, SPTLC2, SLC22A26, PLXNB1, CHP2, RPS6KC1, MFGE8, CNGA3, CYP2B10, TMEM209, NCAM2, SGSM1, KCNT2, HRASLS, LRP11, VAMP5, NEU3, EMP2, GBP3, CD300LB, GBP2, EMP1, SMIM24, BBS4, FXYD3, PKHD1, STK10, BICC1, NAPB, TARM1, FXYD5, SRC, P2RY4, BCHE, PPL, | 678 | 8683 | 22680 | 1,03 | 1,00E+00 | 7,97E-01 | 9,88E+01 |

| | | | | | | | | | | | | |
|---|---|---|---|---|---|---|---|---|---|---|---|---|
| UP_KEYWO | Transmembrane | 209 | 29,5 | 4,98E-01 | MRGPRB2, TMIE, GBP9, ABCC12, PARP16, SSTR2, ABCB1A, CCR5, CD34, CCR2, TMEM184C, GLG1, GM1976, POMK, TMX3, RIC3, MANSC1, ALCAM, ACE, IMPG2, SYBU, OLFR1033, SPNS3, S100A10, CD63, ATP6V0E2, CLEC7A, KLRB1B, D630045J12RIK, RTN4, STEAP4, MPEG1, DNAJC10, LY9, LNPEP, FAR1, PBXIP1, AGPAT9, SEMA3D, ANGPT1, MMD2, NT5E, GOLM1, CLMP, SLC25A4, CAR14, FMR1NB, SLC22A7, COL25A1, MMP15, MYADM, RDH5, AMIGO2, ALG10B, CLIC5, H2-BL, SPACA1, CR1L, SEMA4A, SGCB, MEGF9, 1810064F22RIK, ABHD2, LY6E, MFSD7C, NUP210, FMO2, FMO3, ABCD2, MAS1, METTL7A3, KCNE3, TMPRSS2, MOGAT2, SELP, NAT8, SLC8A1, E130311K13RIK, FADS3, TMPRSS4, MARCH1, P2RY13, SLC16A5, CYP4F16, PTCH1, MBOAT4, LRP2, SLC15A3, GREB1L, IMPAD1, SLC13A5, OCLN, TLR3, SLC7A9, TSPAN9, SLC7A4, TLR7, TLR8, IL17RB, UNC5B, ROBO1, SLC51B, SLC22A27, SLC22A26, SPTLC2, PLXNB1, CNGA3, TMEM209, NCAM2, KCNT2, HRASLS, LRP11, VAMP5, EMP2, CD300LB, EMP1, SMIM24, FXYD3, PKHD1, BICC1, TARM1, FXYD5, P2RY4, BCHE, SLC35F2, SLC35F3, H2-T10, ENTPD3, GAL3ST1, H2-Q2, 2010107G23RIK, H2-Q4, H2-Q1, CD276, P2RX5, TNFSF10, GPR35, UCP2, IFI27L2B, ECSCR, SLC13A3, IL3RA, ATP7B, CLDN7, SKINT9, CLDN4, CLDN6, LEPR, SORL1, TMEM237, GABBR2, DPY19L3, CXADR, GM9992, PCDHGA2, CD93, CSF2RB, TPO, NIPAL1, SRD5A2, CALCRL, | 678 | 6955 | 22680 | 1,01 | 1,00E+00 | 9,06E-01 | 1,00E+02 |
| UP_KEYWO | Transmembrane helix | 207 | 29,2 | 5,52E-01 | MRGPRB2, TMIE, GBP9, ABCC12, PARP16, SSTR2, ABCB1A, CCR5, CD34, CCR2, TMEM184C, GLG1, GM1976, POMK, TMX3, RIC3, MANSC1, ALCAM, ACE, IMPG2, SYBU, OLFR1033, SPNS3, S100A10, CD63, ATP6V0E2, CLEC7A, KLRB1B, D630045J12RIK, RTN4, STEAP4, MPEG1, DNAJC10, LY9, LNPEP, FAR1, PBXIP1, AGPAT9, SEMA3D, ANGPT1, MMD2, NT5E, GOLM1, CLMP, SLC25A4, CAR14, FMR1NB, SLC22A7, COL25A1, MMP15, MYADM, RDH5, AMIGO2, ALG10B, CLIC5, H2-BL, SPACA1, CR1L, SEMA4A, SGCB, MEGF9, 1810064F22RIK, ABHD2, LY6E, MFSD7C, NUP210, ABCD2, MAS1, METTL7A3, KCNE3, TMPRSS2, MOGAT2, SELP, NAT8, SLC8A1, E130311K13RIK, FADS3, TMPRSS4, MARCH1, P2RY13, SLC16A5, CYP4F16, PTCH1, MBOAT4, LRP2, SLC15A3, GREB1L, IMPAD1, SLC13A5, OCLN, TLR3, SLC7A9, TSPAN9, SLC7A4, TLR7, TLR8, IL17RB, UNC5B, ROBO1, SLC51B, SLC22A27, SLC22A26, SPTLC2, PLXNB1, CNGA3, TMEM209, NCAM2, KCNT2, HRASLS, LRP11, VAMP5, EMP2, CD300LB, EMP1, SMIM24, FXYD3, PKHD1, BICC1, TARM1, FXYD5, P2RY4, BCHE, SLC35F2, SLC35F3, H2-T10, ENTPD3, GAL3ST1, H2-Q2, 2010107G23RIK, H2-Q4, H2-Q1, CD276, P2RX5, TNFSF10, GPR35, UCP2, IFI27L2B, ECSCR, SLC13A3, IL3RA, ATP7B, CLDN7, SKINT9, CLDN4, CLDN6, LEPR, SORL1, TMEM237, GABBR2, DPY19L3, CXADR, GM9992, PCDHGA2, CD93, CSF2RB, TPO, NIPAL1, SRD5A2, CALCRL, ATP8B5, SLC43A2, | 678 | 6938 | 22680 | 1,00 | 1,00E+00 | 9,20E-01 | 1,00E+02 |

| | | | | | | | | | | | | |
|---|---|---|---|---|---|---|---|---|---|---|---|---|
| GOTERM_C( | GO:0005886~plasma membrane | 156 | 22 | 5,78E-01 | SYT1, OCLN, SLC13A5, AQP8, VTCN1, SLC7A9, CNP, AQP7, TSPAN9, ADORA1, TLR7, IL17RB, ACTG1, ATP2B2, ANK2, UNC5B, ROBO1, CD46, MLKL, SLC51B, GRID1, MRGPRB2, DAB2IP, SLC22A27, CRYAB, PLXNB1, CHP2, CNGA3, NCAM2, SSTR2, ABCB1A, KCNT2, CCR5, CD34, CCR2, VAMP5, NEU3, CD300LB, EMP2, EMP1, SPTB, GLG1, BBS4, STK10, SRC, ALCAM, ACE, P2RY4, PPL, DMD, OLFR1033, ENTPD3, H2-T10, BLNK, 1700057G04RIK, H2-Q2, H2-Q4, OSBPL3, H2-Q1, CD63, HN1L, P2RX5, GPR35, RGS5, ECSCR, SYTL5, SLC13A3, CLEC7A, GRK5, KLRB1B, IL3RA, MYH10, PLEKHA1, KCTD12, ATP7B, RTN4, CLDN7, STEAP4, CLDN4, GRIP1, LEPR, CLDN6, PRSS41, FERMT1, LY9, GABBR2, CXADR, PCDHGA2, LNPEP, VEPH1, GPC2, CD93, RASL10B, SNTB1, TPO, RHOC, ANGPT1, FAM129B, CALCRL, PAK1, ATP8B5, FAM129A, NT5E, SLC43A2, TRPM4, PRKCA, CLMP, CNTN6, SLC22A7, VIL1, ESR1, ARHGEF9, MYADM, FMN1, SMO, AMIGO2, ALG10B, H2-BL, CLDN1, SEMA4A, SGCB, TNFRSF22, CD244, APH1B, ABHD2, CDH1, VCAM1, LY6A, NPTX1, ANXA8, LY6D, RGS12, LY6E, MFSD7C, MAS1, IPCEF1, SUCNR1, EHD1, KCNE3, TMPRSS2, PTPRC, SLC8A1, FLT4, ACLY, ANXA5, ZFP709, ANXA2, AJUBA, MARCH1, P2RY13, SLC16A5, ANXA13, SCN4B, PTCH1, MGST2, FAM126A | 633 | 4874 | 19662 | 0,99 | 1,00E+00 | 9,74E-01 | 1,00E+02 |
| GOTERM_C( | GO:0016021~integral component of membrane | 209 | 29,5 | 8,76E-01 | MRGPRB2, CLCA1, TMIE, GBP9, ABCC12, PARP16, SSTR2, CCR5, ABCB1A, CD34, CCR2, TMEM184C, GLG1, GM1976, POMK, TMX3, RIC3, MANSC1, ALCAM, ACE, IMPG2, SYBU, OLFR1033, SPNS3, S100A10, CD63, ATP6V0E2, CLEC7A, KLRB1B, D630045J12RIK, RTN4, STEAP4, MPEG1, DNAJC10, LY9, LNPEP, FAR1, PBXIP1, AGPAT9, SEMA3D, ANGPT1, MMD2, NT5E, GOLM1, CLMP, SLC25A4, CAR14, FMR1NB, SLC22A7, COL25A1, MMP15, MYADM, RDH5, AMIGO2, ALG10B, CLIC5, H2-BL, SPACA1, CR1L, SEMA4A, SGCB, MEGF9, 1810064F22RIK, ABHD2, LY6E, MFSD7C, NUP210, FMO2, FMO3, ABCD2, MAS1, METTL7A3, KCNE3, TMPRSS2, MOGAT2, SELP, NAT8, SLC8A1, E130311K13RIK, FADS3, TMPRSS4, MARCH1, P2RY13, SLC16A5, CYP4F16, PTCH1, MBOAT4, LRP2, SLC15A3, GREB1L, IMPAD1, SLC13A5, OCLN, TLR3, SLC7A9, TSPAN9, SLC7A4, TLR7, TLR8, IL17RB, UNC5B, ROBO1, SLC51B, SLC22A27, SLC22A26, SPTLC2, PLXNB1, CNGA3, TMEM209, NCAM2, KCNT2, HRASLS, LRP11, VAMP5, EMP2, CD300LB, EMP1, SMIM24, FXYD3, PKHD1, BICC1, TARM1, FXYD5, P2RY4, BCHE, SLC35F2, SLC35F3, H2-T10, ENTPD3, GAL3ST1, H2-Q2, 2010107G23RIK, H2-Q4, H2-Q1, CD276, TNFSF10, GPR35, UCP2, IFI27L2B, ECSCR, SLC13A3, IL3RA, ATP7B, CLDN7, SKINT9, CLDN4, CLDN6, LEPR, SORL1, TMEM237, GABBR2, DPY19L3, CXADR, GM9992, PCDHGA2, CD93, CSF2RB, TPO, NIPAL1, SRD5A2, CALCRL, ATP8B5, | 633 | 6878 | 19662 | 0,94 | 1,00E+00 | 9,99E-01 | 1,00E+02 |
| Annotation | Enrichment Score: 2.1455953567013775 | | | | | | | | | | | |
| Category | Term | Count | % | PValue | Genes | List To | Pop Hit | Pop Tota | Fold Enr | Bonferroni | Benjamini | FDR |
| UP_KEYWO | Annexin | 4 | 0,56 | 4,76E-03 | ANXA8, ANXA13, ANXA5, ANXA2 | 678 | 12 | 22680 | 11,15 | 7,96E-01 | 1,15E-01 | 6,27E+00 |
| INTERPRO | IPR001464:Annexin | 4 | 0,56 | 5,42E-03 | ANXA8, ANXA13, ANXA5, ANXA2 | 645 | 12 | 20594 | 10,64 | 9,97E-01 | 4,43E-01 | 8,31E+00 |
| INTERPRO | IPR018252:Annexin repeat, conserved site | 4 | 0,56 | 5,42E-03 | ANXA8, ANXA13, ANXA5, ANXA2 | 645 | 12 | 20594 | 10,64 | 9,97E-01 | 4,43E-01 | 8,31E+00 |

| | | | | | | | | | | | | |
|---|---|---|---|---|---|---|---|---|---|---|---|---|
| INTERPRO | IPR018502:Annexin repeat | 4 | 0,56 | 5,42E-03 | ANXA8, ANXA13, ANXA5, ANXA2 | 645 | 12 | 20594 | 10,64 | 9,97E-01 | 4,43E-01 | 8,31E+00 |
| UP_KEYWO | Calcium/phospholipid-binding | 4 | 0,56 | 6,05E-03 | ANXA8, ANXA13, ANXA5, ANXA2 | 678 | 13 | 22680 | 10,29 | 8,68E-01 | 1,26E-01 | 7,91E+00 |
| UP_SEQ_FE | repeat:Annexin 4 | 4 | 0,56 | 7,74E-03 | ANXA8, ANXA13, ANXA5, ANXA2 | 544 | 14 | 18012 | 9,46 | 1,00E+00 | 7,79E-01 | 1,22E+01 |
| UP_SEQ_FE | repeat:Annexin 2 | 4 | 0,56 | 7,74E-03 | ANXA8, ANXA13, ANXA5, ANXA2 | 544 | 14 | 18012 | 9,46 | 1,00E+00 | 7,79E-01 | 1,22E+01 |
| UP_SEQ_FE | repeat:Annexin 3 | 4 | 0,56 | 7,74E-03 | ANXA8, ANXA13, ANXA5, ANXA2 | 544 | 14 | 18012 | 9,46 | 1,00E+00 | 7,79E-01 | 1,22E+01 |
| UP_SEQ_FE | repeat:Annexin 1 | 4 | 0,56 | 7,74E-03 | ANXA8, ANXA13, ANXA5, ANXA2 | 544 | 14 | 18012 | 9,46 | 1,00E+00 | 7,79E-01 | 1,22E+01 |
| SMART | SM00335:ANX | 4 | 0,56 | 8,15E-03 | ANXA8, ANXA13, ANXA5, ANXA2 | 379 | 12 | 10425 | 9,17 | 8,43E-01 | 3,70E-01 | 9,93E+00 |
| GOTERM_M | GO:0005544~calcium-dependent phospholipid binding | 6 | 0,85 | 1,86E-02 | SYT1, ANXA8, SYTL5, ANXA13, ANXA5, ANXA2 | 575 | 47 | 17446 | 3,87 | 1,00E+00 | 8,26E-01 | 2,49E+01 |
| Annotation | Enrichment Score: 1.5562002715526824 | | | | | | | | | | | |
| Category | Term | Count | % | PValue | Genes | List To | Pop Hit | Pop Tota | Fold Enr | Bonferroni | Benjamini | FDR |
| GOTERM_M | GO:0050661~NADP binding | 7 | 0,99 | 1,60E-03 | GAPDHS, FMO2, PGD, G6PDX, MIOX, FMO3, CRYM | 575 | 39 | 17446 | 5,45 | 6,96E-01 | 3,27E-01 | 2,40E+00 |
| UP_SEQ_FE | binding site:NADP | 4 | 0,56 | 3,83E-02 | PTGR1, NMRAL1, G6PDX, CRYM | 544 | 25 | 18012 | 5,30 | 1,00E+00 | 9,52E-01 | 4,79E+01 |
| UP_KEYWO | NADP | 10 | 1,41 | 7,98E-02 | FAR1, PTGR1, NMRAL1, FMO2, PGD, G6PDX, FMO3, DHRS9, SRD5A2, CRYM | 678 | 175 | 22680 | 1,91 | 1,00E+00 | 5,57E-01 | 6,77E+01 |
| UP_SEQ_FE | nucleotide phosphate-binding region:NADP | 5 | 0,71 | 1,22E-01 | PTGR1, NMRAL1, FMO2, FMO3, CRYM | 544 | 63 | 18012 | 2,63 | 1,00E+00 | 9,84E-01 | 8,86E+01 |
| Annotation | Enrichment Score: 1.531595436094844 | | | | | | | | | | | |
| Category | Term | Count | % | PValue | Genes | List To | Pop Hit | Pop Tota | Fold Enr | Bonferroni | Benjamini | FDR |
| UP_KEYWO | Tight junction | 10 | 1,41 | 8,19E-04 | CLDN7, OCLN, CLMP, CLDN4, CGN, CLDN6, CLDN1, CXADR, ECT2, NPHP1 | 678 | 83 | 22680 | 4,03 | 2,39E-01 | 2,69E-02 | 1,11E+00 |
| GOTERM_C | GO:0016327~apicolateral plasma membrane | 5 | 0,71 | 4,09E-03 | CLDN7, OCLN, CLDN4, CLDN6, CXADR | 633 | 21 | 19662 | 7,40 | 8,04E-01 | 2,08E-01 | 5,56E+00 |
| GOTERM_C | GO:0005923~bicellular tight junction | 11 | 1,55 | 9,77E-03 | CLDN7, OCLN, CLMP, CLDN4, CGN, CLDN6, CLDN1, CXADR, ECT2, ATP7B, NPHP1 | 633 | 131 | 19662 | 2,61 | 9,80E-01 | 2,60E-01 | 1,28E+01 |
| GOTERM_B | GO:0070830~bicellular tight junction assembly | 5 | 0,71 | 1,07E-02 | OCLN, CGN, CLDN1, CDH1, ECT2 | 588 | 27 | 18082 | 5,69 | 1,00E+00 | 7,70E-01 | 1,71E+01 |
| UP_SEQ_FE | region of interest:Interactions with TJP1, TJP2 and TJP3 | 3 | 0,42 | 1,72E-02 | CLDN7, CLDN4, CLDN6 | 544 | 7 | 18012 | 14,19 | 1,00E+00 | 9,33E-01 | 2,52E+01 |
| INTERPRO | IPR004031:PMP-22/EMP/MP20/Claudin | 6 | 0,85 | 2,28E-02 | CLDN7, CLDN4, CLDN6, CLDN1, EMP2, EMP1 | 645 | 52 | 20594 | 3,68 | 1,00E+00 | 8,31E-01 | 3,08E+01 |

| | | | | | | | | | | | | |
|---|---|---|---|---|---|---|---|---|---|---|---|---|
| KEGG_PATH | mmu04530:Tight junction | 10 | 1,41 | 2,39E-02 | ACTG1, PRKCA, CLDN7, OCLN, CLDN4, CGN, CLDN6, CLDN1, SRC, MYH10 | 233 | 139 | 7720 | 2,38 | 9,97E-01 | 6,93E-01 | 2,69E+01 |
| INTERPRO | IPR017974:Claudin, conserved site | 4 | 0,56 | 3,39E-02 | CLDN7, CLDN4, CLDN6, CLDN1 | 645 | 23 | 20594 | 5,55 | 1,00E+00 | 8,58E-01 | 4,23E+01 |
| GOTERM_BI | GO:0016338~calcium-independent cell-cell adhesion via plasma membrane cell-adhesion molecules | 4 | 0,56 | 4,15E-02 | CLDN7, CLDN4, CLDN6, CLDN1 | 588 | 24 | 18082 | 5,13 | 1,00E+00 | 9,21E-01 | 5,24E+01 |
| GOTERM_BI | GO:0045216~cell-cell junction organization | 4 | 0,56 | 4,15E-02 | OCLN, CLDN6, CLDN1, CXADR | 588 | 24 | 18082 | 5,13 | 1,00E+00 | 9,21E-01 | 5,24E+01 |
| KEGG_PATH | mmu04670:Leukocyte transendothelial migration | 8 | 1,13 | 7,17E-02 | ACTG1, PRKCA, VCAM1, CLDN7, OCLN, CLDN4, CLDN6, CLDN1 | 233 | 121 | 7720 | 2,19 | 1,00E+00 | 7,80E-01 | 6,18E+01 |
| GOTERM_C( | GO:0016328~lateral plasma membrane | 5 | 0,71 | 9,51E-02 | CLDN7, OCLN, CLDN4, CLDN1, CDH1 | 633 | 54 | 19662 | 2,88 | 1,00E+00 | 6,39E-01 | 7,52E+01 |
| KEGG_PATH | mmu05160:Hepatitis C | 8 | 1,13 | 1,15E-01 | CLDN7, CDKN1A, OCLN, CLDN4, CLDN6, CLDN1, TLR3, OAS1A | 233 | 136 | 7720 | 1,95 | 1,00E+00 | 7,74E-01 | 7,94E+01 |
| INTERPRO | IPR006187:Claudin | 4 | 0,56 | 1,36E-01 | CLDN7, CLDN4, CLDN6, CLDN1 | 645 | 41 | 20594 | 3,11 | 1,00E+00 | 9,72E-01 | 9,03E+01 |
| GOTERM_M | GO:0005198~structural molecule activity | 9 | 1,27 | 5,19E-01 | CLDN7, CLDN4, SPRR1A, CLDN6, KRT7, 2010005H15RIK, CLDN1, SNTB1, KRT23 | 575 | 236 | 17446 | 1,16 | 1,00E+00 | 9,98E-01 | 1,00E+02 |
| Annotation | Enrichment Score: 1.4340358174913816 | | | | | | | | | | | |
| Category | Term | Count | % | PValue | Genes | List To | Pop Hit | Pop Tota | Fold Enr | Bonferroni | Benjamini | FDR |
| INTERPRO | IPR016040:NAD(P)-binding domain | 17 | 2,4 | 5,45E-04 | STEAP4, PTGR1, G6PDX, PGD, UBA7, UBA6, DHRS9, ACLY, RDH5, GAPDHS, FAR1, KCNT2, NMRAL1, FMO2, FMO3, BC089597, CRYM | 645 | 199 | 20594 | 2,73 | 4,44E-01 | 6,32E-02 | 8,66E-01 |
| UP_KEYWO | Oxidoreductase | 30 | 4,23 | 1,77E-02 | STEAP4, CYP2B9, CYP2D12, G6PDX, PGD, DNAJC10, HR, FAR1, FMO2, FMO3, MIOX, PNPO, TPO, GSTO2, SRD5A2, ACAD9, PTGR1, FADS3, DHRS9, CYP2B13, CYP2B10, RDH5, GAPDHS, CYP2A22, ALDH1B1, CYP4F16, HAO2, CRYM, CYP2A4, ACAD12 | 678 | 639 | 22680 | 1,57 | 9,97E-01 | 2,81E-01 | 2,15E+01 |
| UP_KEYWO | NADP | 10 | 1,41 | 7,98E-02 | FAR1, PTGR1, NMRAL1, FMO2, PGD, G6PDX, FMO3, DHRS9, SRD5A2, CRYM | 678 | 175 | 22680 | 1,91 | 1,00E+00 | 5,57E-01 | 6,77E+01 |
| GOTERM_M | GO:0016491~oxidoreductase activity | 25 | 3,53 | 1,97E-01 | STEAP4, CYP2B9, PGD, G6PDX, DNAJC10, HR, FAR1, FMO2, FMO3, MIOX, BC089597, TPO, PNPO, GSTO2, SRD5A2, ACAD9, PTGR1, FADS3, DHRS9, CYP2B10, RDH5, GAPDHS, ALDH1B1, HAO2, CRYM | 575 | 604 | 17446 | 1,26 | 1,00E+00 | 9,73E-01 | 9,64E+01 |
| GOTERM_BI | GO:0055114~oxidation-reduction process | 24 | 3,39 | 4,47E-01 | STEAP4, PTGR1, CYP2B9, G6PDX, PGD, FADS3, HR, DNAJC10, DHRS9, CYP2B10, RDH5, GAPDHS, FAR1, ALDH1B1, FMO2, MIOX, HAO2, FMO3, PNPO, TPO, GSTO2, SRD5A2, ACAD9, CRYM | 588 | 676 | 18082 | 1,09 | 1,00E+00 | 1,00E+00 | 1,00E+02 |

| Annotation | Enrichment Score: 1.3677231318128533 | | | | | | | | | | | |
|---|---|---|---|---|---|---|---|---|---|---|---|---|
| Category | Term | Count | % | PValue | Genes | List To | Pop Hit | Pop Tota | Fold Enr | Bonferroni | Benjamini | FDR |
| GOTERM_B | GO:0009615~response to virus | 8 | 1,13 | 1,93E-02 | IFIT2, OASL1, TLR3, TGTP1, OAS1A, SRC, TLR8, CXCL10 | 588 | 84 | 18082 | 2,93 | 1,00E+00 | 8,50E-01 | 2,89E+01 |
| GOTERM_B | GO:0051607~defense response to virus | 12 | 1,69 | 2,06E-02 | IFIT2, PTPRC, TRIM56, OASL1, TLR3, OAS1A, IL33, SPON2, TLR7, TLR8, ZBP1, CXCL10 | 588 | 167 | 18082 | 2,21 | 1,00E+00 | 8,45E-01 | 3,05E+01 |
| GOTERM_M | GO:0003725~double-stranded RNA binding | 5 | 0,71 | 1,99E-01 | OASL1, TLR3, OAS1A, TLR7, TLR8 | 575 | 70 | 17446 | 2,17 | 1,00E+00 | 9,70E-01 | 9,65E+01 |
| Annotation | Enrichment Score: 1.298766410539755 | | | | | | | | | | | |
| Category | Term | Count | % | PValue | Genes | List To | Pop Hit | Pop Tota | Fold Enr | Bonferroni | Benjamini | FDR |
| INTERPRO | IPR001751:S100/Calbindin-D9k, conserved site | 4 | 0,56 | 1,75E-02 | S100A4, S100A11, S100G, S100A10 | 645 | 18 | 20594 | 7,10 | 1,00E+00 | 7,69E-01 | 2,45E+01 |
| INTERPRO | IPR013787:S100/CaBP-9k-type, calcium binding, subdomain | 4 | 0,56 | 5,11E-02 | S100A4, S100A11, S100G, S100A10 | 645 | 27 | 20594 | 4,73 | 1,00E+00 | 9,05E-01 | 5,66E+01 |
| SMART | SM01394:SM01394 | 4 | 0,56 | 6,64E-02 | S100A4, S100A11, S100G, S100A10 | 379 | 26 | 10425 | 4,23 | 1,00E+00 | 9,25E-01 | 5,85E+01 |
| UP_SEQ_FE | calcium-binding region:2; high affinity | 3 | 0,42 | 7,35E-02 | S100A4, S100A11, S100G | 544 | 15 | 18012 | 6,62 | 1,00E+00 | 9,69E-01 | 7,20E+01 |
| UP_SEQ_FE | calcium-binding region:1; low affinity | 3 | 0,42 | 7,35E-02 | S100A4, S100A11, S100G | 544 | 15 | 18012 | 6,62 | 1,00E+00 | 9,69E-01 | 7,20E+01 |
| Annotation | Enrichment Score: 1.2493959155905938 | | | | | | | | | | | |
| Category | Term | Count | % | PValue | Genes | List To | Pop Hit | Pop Tota | Fold Enr | Bonferroni | Benjamini | FDR |
| UP_KEYWO | Sushi | 6 | 0,85 | 1,27E-02 | SELP, CR2, CD46, TPO, C1S1, CR1L | 678 | 47 | 22680 | 4,27 | 9,86E-01 | 2,22E-01 | 1,60E+01 |
| UP_SEQ_FE | domain:Sushi 2 | 5 | 0,71 | 2,04E-02 | SELP, CR2, CD46, C1S1, CR1L | 544 | 35 | 18012 | 4,73 | 1,00E+00 | 9,46E-01 | 2,91E+01 |
| UP_SEQ_FE | domain:Sushi 1 | 5 | 0,71 | 2,04E-02 | SELP, CR2, CD46, C1S1, CR1L | 544 | 35 | 18012 | 4,73 | 1,00E+00 | 9,46E-01 | 2,91E+01 |
| UP_SEQ_FE | domain:Sushi 4 | 4 | 0,56 | 2,12E-02 | SELP, CR2, CD46, CR1L | 544 | 20 | 18012 | 6,62 | 1,00E+00 | 9,23E-01 | 3,01E+01 |
| INTERPRO | IPR000436:Sushi/SCR/CCP | 6 | 0,85 | 3,04E-02 | SELP, CR2, CD46, TPO, C1S1, CR1L | 645 | 56 | 20594 | 3,42 | 1,00E+00 | 8,59E-01 | 3,89E+01 |
| UP_SEQ_FE | domain:Sushi 3 | 4 | 0,56 | 3,83E-02 | SELP, CR2, CD46, CR1L | 544 | 25 | 18012 | 5,30 | 1,00E+00 | 9,52E-01 | 4,79E+01 |
| SMART | SM00032:CCP | 6 | 0,85 | 4,52E-02 | SELP, CR2, CD46, TPO, C1S1, CR1L | 379 | 54 | 10425 | 3,06 | 1,00E+00 | 8,77E-01 | 4,47E+01 |
| UP_SEQ_FE | domain:Sushi 5 | 3 | 0,42 | 7,35E-02 | SELP, CR2, CR1L | 544 | 15 | 18012 | 6,62 | 1,00E+00 | 9,69E-01 | 7,20E+01 |
| UP_KEYWO | Complement pathway | 3 | 0,42 | 1,92E-01 | CR2, C1S1, CR1L | 678 | 27 | 22680 | 3,72 | 1,00E+00 | 7,25E-01 | 9,45E+01 |

| | | | | | | | | | | | | |
|---|---|---|---|---|---|---|---|---|---|---|---|---|
| KEGG_PATH | mmu04610:Complement and coagulation cascades | 4 | 0,56 | 4,01E-01 | CR2, CD46, SERPINE1, C1S1 | 233 | 76 | 7720 | 1,74 | 1,00E+00 | 9,10E-01 | 9,99E+01 |
| GOTERM_BI | GO:0006958~complement activation, classical pathway | 3 | 0,42 | 5,37E-01 | CR2, C1S1, CR1L | 588 | 55 | 18082 | 1,68 | 1,00E+00 | 1,00E+00 | 1,00E+02 |
| Annotation | Enrichment Score: 1.2046142592410185 | | | | | | | | | | | |
| Category | Term | Count | % | PValue | Genes | List To | Pop Hit | Pop Tota | Fold Enr | Bonferroni | Benjamini | FDR |
| UP_KEYWO | GPI-anchor | 10 | 1,41 | 2,48E-02 | LY6A, VCAM1, NCAM2, GPC2, LY6D, LY6E, VTCN1, CNTN6, PRSS41, NT5E | 678 | 140 | 22680 | 2,39 | 1,00E+00 | 3,41E-01 | 2,89E+01 |
| GOTERM_C | GO:0031225~anchored component of membrane | 10 | 1,41 | 3,85E-02 | LY6A, VCAM1, NCAM2, GPC2, LY6D, LY6E, VTCN1, CNTN6, PRSS41, NT5E | 633 | 141 | 19662 | 2,20 | 1,00E+00 | 5,25E-01 | 4,22E+01 |
| UP_SEQ_FE | propeptide:Removed in mature form | 13 | 1,83 | 5,70E-02 | LY6A, GPC2, LY6D, LY6E, VTCN1, CNTN6, SORL1, PRSS41, RASL10B, RHOC, NT5E, GBP2, PSMB9 | 544 | 238 | 18012 | 1,81 | 1,00E+00 | 9,52E-01 | 6,25E+01 |
| UP_SEQ_FE | lipid moiety-binding region:GPI-anchor amidated serine | 4 | 0,56 | 2,79E-01 | GPC2, LY6D, CNTN6, NT5E | 544 | 61 | 18012 | 2,17 | 1,00E+00 | 9,98E-01 | 9,96E+01 |
| Annotation | Enrichment Score: 1.170720928954318 | | | | | | | | | | | |
| Category | Term | Count | % | PValue | Genes | List To | Pop Hit | Pop Tota | Fold Enr | Bonferroni | Benjamini | FDR |
| INTERPRO | IPR011042:Six-bladed beta-propeller, TolB-like | 6 | 0,85 | 8,71E-03 | HHIPL2, TRIM56, SORL1, NID1, NHLRC1, LRP2 | 645 | 41 | 20594 | 4,67 | 1,00E+00 | 5,75E-01 | 1,30E+01 |
| UP_SEQ_FE | repeat:LDL-receptor class B 4 | 3 | 0,42 | 6,50E-02 | SORL1, NID1, LRP2 | 544 | 14 | 18012 | 7,10 | 1,00E+00 | 9,65E-01 | 6,74E+01 |
| UP_SEQ_FE | repeat:LDL-receptor class B 2 | 3 | 0,42 | 6,50E-02 | SORL1, NID1, LRP2 | 544 | 14 | 18012 | 7,10 | 1,00E+00 | 9,65E-01 | 6,74E+01 |
| UP_SEQ_FE | repeat:LDL-receptor class B 3 | 3 | 0,42 | 6,50E-02 | SORL1, NID1, LRP2 | 544 | 14 | 18012 | 7,10 | 1,00E+00 | 9,65E-01 | 6,74E+01 |
| UP_SEQ_FE | repeat:LDL-receptor class B 1 | 3 | 0,42 | 6,50E-02 | SORL1, NID1, LRP2 | 544 | 14 | 18012 | 7,10 | 1,00E+00 | 9,65E-01 | 6,74E+01 |
| INTERPRO | IPR000033:LDLR class B repeat | 3 | 0,42 | 7,84E-02 | SORL1, NID1, LRP2 | 645 | 15 | 20594 | 6,39 | 1,00E+00 | 9,41E-01 | 7,28E+01 |
| SMART | SM00135:LY | 3 | 0,42 | 1,01E-01 | SORL1, NID1, LRP2 | 379 | 15 | 10425 | 5,50 | 1,00E+00 | 9,10E-01 | 7,43E+01 |
| UP_KEYWO | EGF-like domain | 9 | 1,27 | 3,51E-01 | SELP, CD93, IMPG2, SORL1, TPO, NID1, MFGE8, LRP2, C1S1 | 678 | 224 | 22680 | 1,34 | 1,00E+00 | 8,31E-01 | 9,97E+01 |
| Annotation | Enrichment Score: 1.1616452279237515 | | | | | | | | | | | |

| Category | Term | Count | % | PValue | Genes | List To | Pop Hit | Pop Tot | Fold Enr | Bonferroni | Benjamini | FDR |
|---|---|---|---|---|---|---|---|---|---|---|---|---|
| GOTERM_BI | GO:0006814~sodium ion transport | 9 | 1,27 | 4,91E-02 | SLC17A8, SLC8A1, SLC13A5, CHP2, SLC5A9, SCN4B, SLC13A3, SLC10A5, SLC10A2 | 588 | 124 | 18082 | 2,23 | 1,00E+00 | 9,31E-01 | 5,86E+01 |
| UP_KEYWO | Sodium transport | 8 | 1,13 | 5,27E-02 | SLC17A8, SLC8A1, SLC13A5, SLC5A9, SCN4B, SLC13A3, SLC10A5, SLC10A2 | 678 | 113 | 22680 | 2,37 | 1,00E+00 | 4,87E-01 | 5,21E+01 |
| UP_KEYWO | Sodium | 8 | 1,13 | 6,83E-02 | SLC17A8, SLC8A1, SLC13A5, SLC5A9, SCN4B, SLC13A3, SLC10A5, SLC10A2 | 678 | 120 | 22680 | 2,23 | 1,00E+00 | 5,44E-01 | 6,17E+01 |
| GOTERM_M | GO:0015293~symporter activity | 8 | 1,13 | 7,65E-02 | SLC17A8, SLC16A5, SLC13A5, SLC5A9, SLC13A3, SLC10A5, SLC10A2, SLC15A3 | 575 | 112 | 17446 | 2,17 | 1,00E+00 | 9,06E-01 | 7,02E+01 |
| UP_KEYWO | Symport | 7 | 0,99 | 1,15E-01 | SLC17A8, SLC16A5, SLC13A5, SLC13A3, SLC10A5, SLC10A2, SLC15A3 | 678 | 111 | 22680 | 2,11 | 1,00E+00 | 6,21E-01 | 8,10E+01 |
| Annotation | Enrichment Score: 1.1245328203023952 | | | | | | | | | | | |
| Category | Term | Count | % | PValue | Genes | List To | Pop Hit | Pop Tot | Fold Enr | Bonferroni | Benjamini | FDR |
| GOTERM_BI | GO:0010881~regulation of cardiac muscle contraction by regulation of the release of sequestered calcium ion | 3 | 0,42 | 2,59E-02 | SLC8A1, ANK2, DMD | 588 | 8 | 18082 | 11,53 | 1,00E+00 | 8,70E-01 | 3,68E+01 |
| GOTERM_C | GO:0030018~Z disc | 9 | 1,27 | 4,69E-02 | BMP10, SLC8A1, ANK2, CRYAB, DMD, KY, PAK1, ANXA5, CSRP3 | 633 | 124 | 19662 | 2,25 | 1,00E+00 | 5,34E-01 | 4,88E+01 |
| GOTERM_BI | GO:0060048~cardiac muscle contraction | 5 | 0,71 | 6,97E-02 | SLC8A1, ANK2, DMD, SCN4B, CSRP3 | 588 | 48 | 18082 | 3,20 | 1,00E+00 | 9,56E-01 | 7,18E+01 |
| GOTERM_C | GO:0042383~sarcolemma | 8 | 1,13 | 8,62E-02 | VCAM1, SLC8A1, ANK2, DMD, SNTB1, ANXA5, SGCB, ANXA2 | 633 | 118 | 19662 | 2,11 | 1,00E+00 | 6,31E-01 | 7,16E+01 |
| GOTERM_BI | GO:0002027~regulation of heart rate | 3 | 0,42 | 3,27E-01 | SLC8A1, ANK2, DMD | 588 | 36 | 18082 | 2,56 | 1,00E+00 | 9,99E-01 | 9,99E+01 |
| Annotation | Enrichment Score: 1.1164206942570611 | | | | | | | | | | | |
| Category | Term | Count | % | PValue | Genes | List To | Pop Hit | Pop Tot | Fold Enr | Bonferroni | Benjamini | FDR |
| GOTERM_M | GO:0004867~serine-type endopeptidase inhibitor activity | 10 | 1,41 | 2,04E-02 | WFDC12, SERPINB8, SERPINA3I, SERPINE1, SERPINB1A, ITIH5, SLPI, SPINK5, SERPINH1, WFDC2 | 575 | 123 | 17446 | 2,47 | 1,00E+00 | 7,83E-01 | 2,69E+01 |
| GOTERM_M | GO:0030414~peptidase inhibitor activity | 10 | 1,41 | 2,04E-02 | WFDC12, WFDC15B, SERPINB8, SERPINE1, SERPINB1A, ITIH5, SLPI, SPINK5, WFDC2, TIMP1 | 575 | 123 | 17446 | 2,47 | 1,00E+00 | 7,83E-01 | 2,69E+01 |
| GOTERM_BI | GO:0010951~negative regulation of endopeptidase activity | 4 | 0,56 | 2,24E-02 | SERPINB8, SERPINE1, SLPI, TIMP1 | 588 | 19 | 18082 | 6,47 | 1,00E+00 | 8,59E-01 | 3,27E+01 |
| INTERPRO | IPR008197:Whey acidic protein-type 4-disulphide core | 4 | 0,56 | 3,01E-02 | WFDC12, WFDC15B, SLPI, WFDC2 | 645 | 22 | 20594 | 5,81 | 1,00E+00 | 8,73E-01 | 3,86E+01 |

| | | | | | | | | | | | | |
|---|---|---|---|---|---|---|---|---|---|---|---|---|
| UP_KEYWO | Serine protease inhibitor | 7 | 0,99 | 3,97E-02 | WFDC12, SERPINB8, SERPINE1, SERPINB1A, ITIH5, SLPI, WFDC2 | 678 | 84 | 22680 | 2,79 | 1,00E+00 | 4,44E-01 | 4,23E+01 |
| UP_KEYWO | Protease inhibitor | 8 | 1,13 | 7,07E-02 | WFDC12, SERPINB8, SERPINE1, SERPINB1A, ITIH5, SLPI, WFDC2, TIMP1 | 678 | 121 | 22680 | 2,21 | 1,00E+00 | 5,45E-01 | 6,31E+01 |
| GOTERM_B | GO:0010466~negative regulation of peptidase activity | 8 | 1,13 | 8,66E-02 | WFDC12, SERPINB8, SERPINE1, SERPINB1A, ITIH5, SLPI, WFDC2, TIMP1 | 588 | 117 | 18082 | 2,10 | 1,00E+00 | 9,74E-01 | 7,95E+01 |
| INTERPRO | IPR000215:Serpin family | 5 | 0,71 | 1,57E-01 | SERPINB8, SERPINA3I, SERPINE1, SERPINB1A, SERPINH1 | 645 | 67 | 20594 | 2,38 | 1,00E+00 | 9,71E-01 | 9,35E+01 |
| INTERPRO | IPR023796:Serpin domain | 5 | 0,71 | 1,57E-01 | SERPINB8, SERPINA3I, SERPINE1, SERPINB1A, SERPINH1 | 645 | 67 | 20594 | 2,38 | 1,00E+00 | 9,71E-01 | 9,35E+01 |
| INTERPRO | IPR023795:Protease inhibitor I4, serpin, conserved site | 4 | 0,56 | 2,22E-01 | SERPINB8, SERPINE1, SERPINB1A, SERPINH1 | 645 | 52 | 20594 | 2,46 | 1,00E+00 | 9,84E-01 | 9,82E+01 |
| SMART | SM00093:SERPIN | 5 | 0,71 | 2,25E-01 | SERPINB8, SERPINA3I, SERPINE1, SERPINB1A, SERPINH1 | 379 | 67 | 10425 | 2,05 | 1,00E+00 | 9,66E-01 | 9,61E+01 |
| UP_SEQ_FE | site:Reactive bond | 3 | 0,42 | 4,78E-01 | SERPINB8, SERPINE1, SERPINB1A | 544 | 53 | 18012 | 1,87 | 1,00E+00 | 1,00E+00 | 1,00E+02 |
| Annotation | Enrichment Score: 1.0856289726512776 | | | | | | | | | | | |
| Category | Term | Count | % | PValue | Genes | List To | Pop Hit | Pop Tot | Fold Enr | Bonferroni | Benjamini | FDR |
| GOTERM_M | GO:0005509~calcium ion binding | 38 | 5,36 | 3,08E-03 | S100A4, SYT1, FSTL1, CDH1, PCDHGA2, ATP2B2, ANXA8, CALML4, CD93, PRRG1, PLS1, EFCAB2, TPO, EHD1, CIB3, BGLAP3, SELP, AMY1, PNLIPRP1, BGLAP, SLC8A1, NCALD, CAPSL, MICU2, VIL1, S100A11, CHP2, S100A10, NID1, MMP15, ANXA5, MMP12, ANXA2, S100G, SYTL5, ANXA13, C1S1, LRP2 | 575 | 699 | 17446 | 1,65 | 8,99E-01 | 4,36E-01 | 4,58E+00 |
| INTERPRO | IPR002048:EF-hand domain | 14 | 1,97 | 2,39E-02 | S100A4, NCALD, CAPSL, MICU2, CHP2, S100A11, S100A10, FSTL1, CALML4, PLS1, EFCAB2, S100G, EHD1, CIB3 | 645 | 223 | 20594 | 2,00 | 1,00E+00 | 8,24E-01 | 3,20E+01 |
| UP_SEQ_FE | domain:EF-hand 2 | 11 | 1,55 | 3,71E-02 | S100A4, CALML4, NCALD, MICU2, CAPSL, PLS1, CHP2, S100A11, EFCAB2, S100G, FSTL1 | 544 | 173 | 18012 | 2,11 | 1,00E+00 | 9,55E-01 | 4,68E+01 |
| UP_SEQ_FE | domain:EF-hand 1 | 11 | 1,55 | 3,84E-02 | S100A4, CALML4, NCALD, MICU2, CAPSL, PLS1, CHP2, S100A11, EFCAB2, S100G, FSTL1 | 544 | 174 | 18012 | 2,09 | 1,00E+00 | 9,45E-01 | 4,79E+01 |
| INTERPRO | IPR011992:EF-hand-like domain | 15 | 2,12 | 4,95E-02 | S100A4, NCALD, CAPSL, MICU2, CHP2, S100A11, S100A10, FSTL1, CALML4, DMD, PLS1, EFCAB2, S100G, EHD1, CIB3 | 645 | 273 | 20594 | 1,75 | 1,00E+00 | 9,07E-01 | 5,55E+01 |
| SMART | SM00054:EFh | 10 | 1,41 | 8,09E-02 | S100A4, CALML4, NCALD, MICU2, CAPSL, PLS1, CHP2, S100A11, EFCAB2, S100G | 379 | 145 | 10425 | 1,90 | 1,00E+00 | 9,08E-01 | 6,60E+01 |
| UP_SEQ_FE | domain:EF-hand 4 | 5 | 0,71 | 1,02E-01 | CALML4, NCALD, MICU2, CAPSL, CHP2 | 544 | 59 | 18012 | 2,81 | 1,00E+00 | 9,76E-01 | 8,34E+01 |
| INTERPRO | IPR018247:EF-Hand 1, calcium-binding site | 9 | 1,27 | 1,83E-01 | S100A4, NCALD, CAPSL, PLS1, CHP2, S100A11, S100G, EHD1, CIB3 | 645 | 175 | 20594 | 1,64 | 1,00E+00 | 9,81E-01 | 9,60E+01 |
| UP_SEQ_FE | domain:EF-hand 3 | 5 | 0,71 | 2,94E-01 | CALML4, NCALD, MICU2, CAPSL, CHP2 | 544 | 91 | 18012 | 1,82 | 1,00E+00 | 9,99E-01 | 9,97E+01 |
| UP_SEQ_FE | calcium-binding region:2 | 4 | 0,56 | 6,72E-01 | NCALD, CAPSL, PLS1, CHP2 | 544 | 114 | 18012 | 1,16 | 1,00E+00 | 1,00E+00 | 1,00E+02 |
| UP_SEQ_FE | calcium-binding region:1 | 4 | 0,56 | 7,36E-01 | NCALD, CAPSL, PLS1, CHP2 | 544 | 126 | 18012 | 1,05 | 1,00E+00 | 1,00E+00 | 1,00E+02 |

| Annotation | Enrichment Score: 1.0353382993539342 | | | | | | | | | | | |
|---|---|---|---|---|---|---|---|---|---|---|---|---|
| Category | Term | Count | % | PValue | Genes | List To | Pop Hit | Pop Tota | Fold Enr | Bonferroni | Benjamini | FDR |
| UP_KEYWO | Adaptive immunity | 8 | 1,13 | 2,74E-02 | ALCAM, TRPM4, CD244, THEMIS, VTCN1, TAP1, LY9, SEMA4A | 678 | 98 | 22680 | 2,73 | 1,00E+00 | 3,57E-01 | 3,15E+01 |
| UP_SEQ_FE | domain:Ig-like V-type 2 | 3 | 0,42 | 1,21E-01 | ALCAM, VTCN1, LY9 | 544 | 20 | 18012 | 4,97 | 1,00E+00 | 9,84E-01 | 8,83E+01 |
| UP_SEQ_FE | domain:Ig-like V-type 1 | 3 | 0,42 | 1,31E-01 | ALCAM, VTCN1, LY9 | 544 | 21 | 18012 | 4,73 | 1,00E+00 | 9,84E-01 | 9,04E+01 |
| GOTERM_BI | GO:0002250~adaptive immune response | 8 | 1,13 | 1,67E-01 | ALCAM, TRPM4, CD244, THEMIS, VTCN1, TAP1, LY9, SEMA4A | 588 | 139 | 18082 | 1,77 | 1,00E+00 | 9,92E-01 | 9,59E+01 |
| Annotation | Enrichment Score: 0.9941657739993369 | | | | | | | | | | | |
| Category | Term | Count | % | PValue | Genes | List To | Pop Hit | Pop Tota | Fold Enr | Bonferroni | Benjamini | FDR |
| INTERPRO | IPR018159:Spectrin/alpha-actinin | 4 | 0,56 | 6,09E-02 | CCDC141, PPL, DMD, SPTB | 645 | 29 | 20594 | 4,40 | 1,00E+00 | 9,26E-01 | 6,33E+01 |
| SMART | SM00150:SPEC | 4 | 0,56 | 8,63E-02 | CCDC141, PPL, DMD, SPTB | 379 | 29 | 10425 | 3,79 | 1,00E+00 | 8,96E-01 | 6,85E+01 |
| UP_SEQ_FE | repeat:Spectrin 3 | 3 | 0,42 | 9,16E-02 | PPL, DMD, SPTB | 544 | 17 | 18012 | 5,84 | 1,00E+00 | 9,71E-01 | 7,99E+01 |
| UP_SEQ_FE | repeat:Spectrin 2 | 3 | 0,42 | 1,11E-01 | PPL, DMD, SPTB | 544 | 19 | 18012 | 5,23 | 1,00E+00 | 9,81E-01 | 8,59E+01 |
| UP_SEQ_FE | repeat:Spectrin 1 | 3 | 0,42 | 1,11E-01 | PPL, DMD, SPTB | 544 | 19 | 18012 | 5,23 | 1,00E+00 | 9,81E-01 | 8,59E+01 |
| INTERPRO | IPR002017:Spectrin repeat | 3 | 0,42 | 1,83E-01 | CCDC141, DMD, SPTB | 645 | 25 | 20594 | 3,83 | 1,00E+00 | 9,80E-01 | 9,60E+01 |
| Annotation | Enrichment Score: 0.9763778018895578 | | | | | | | | | | | |
| Category | Term | Count | % | PValue | Genes | List To | Pop Hit | Pop Tota | Fold Enr | Bonferroni | Benjamini | FDR |
| GOTERM_BI | GO:0090026~positive regulation of monocyte chemotaxis | 4 | 0,56 | 9,48E-03 | CCL2, CCR2, SERPINE1, CXCL10 | 588 | 14 | 18082 | 8,79 | 1,00E+00 | 7,51E-01 | 1,54E+01 |
| GOTERM_BI | GO:0070098~chemokine-mediated signaling pathway | 5 | 0,71 | 1,03E-01 | GPR35, CCL2, CCR2, TFF2, CXCL10 | 588 | 55 | 18082 | 2,80 | 1,00E+00 | 9,82E-01 | 8,50E+01 |
| GOTERM_BI | GO:0006935~chemotaxis | 6 | 0,85 | 3,37E-01 | CCL2, CCR5, ROBO1, ECSCR, CCR2, CXCL10 | 588 | 118 | 18082 | 1,56 | 1,00E+00 | 9,99E-01 | 9,99E+01 |
| KEGG_PATH | mmu04062:Chemokine signaling pathway | 8 | 1,13 | 3,78E-01 | CCL2, CXCL14, CCR5, CCR2, GRK5, PAK1, SRC, CXCL10 | 233 | 196 | 7720 | 1,35 | 1,00E+00 | 9,06E-01 | 9,98E+01 |
| Annotation | Enrichment Score: 0.956280717223684 | | | | | | | | | | | |
| Category | Term | Count | % | PValue | Genes | List To | Pop Hit | Pop Tota | Fold Enr | Bonferroni | Benjamini | FDR |

| | | | | | | | | | | | | |
|---|---|---|---|---|---|---|---|---|---|---|---|---|
| UP_SEQ_FE | domain:KRAB | 7 | 0,99 | 3,40E-03 | ZFP13, ZFP426, ZFP120, ZFP667, ZIK1, ZFP445, ZFP26 | 544 | 49 | 18012 | 4,73 | 9,95E-01 | 5,30E-01 | 5,52E+00 |
| UP_SEQ_FE | zinc finger region:C2H2-type 8 | 8 | 1,13 | 6,86E-02 | ZFP13, ZFP770, ZFP426, ZFP120, ZFP667, ZIK1, ZFP445, ZFP26 | 544 | 119 | 18012 | 2,23 | 1,00E+00 | 9,65E-01 | 6,94E+01 |
| UP_SEQ_FE | zinc finger region:C2H2-type 9 | 7 | 0,99 | 8,48E-02 | ZFP770, ZFP426, ZFP120, ZFP667, ZIK1, ZFP445, ZFP26 | 544 | 101 | 18012 | 2,29 | 1,00E+00 | 9,73E-01 | 7,72E+01 |
| UP_SEQ_FE | zinc finger region:C2H2-type 6 | 9 | 1,27 | 8,94E-02 | PLAG1, ZFP13, ZFP770, ZFP426, ZFP120, ZFP667, ZIK1, ZFP445, ZFP26 | 544 | 152 | 18012 | 1,96 | 1,00E+00 | 9,74E-01 | 7,91E+01 |
| UP_SEQ_FE | zinc finger region:C2H2-type 7 | 8 | 1,13 | 9,55E-02 | PLAG1, ZFP13, ZFP426, ZFP120, ZFP667, ZIK1, ZFP445, ZFP26 | 544 | 129 | 18012 | 2,05 | 1,00E+00 | 9,73E-01 | 8,13E+01 |
| UP_SEQ_FE | zinc finger region:C2H2-type 4 | 11 | 1,55 | 1,16E-01 | ZFP518A, PLAG1, ZFP13, ADNP2, ZFP770, ZFP426, ZFP120, ZFP667, ZIK1, ZFP445, ZFP26 | 544 | 215 | 18012 | 1,69 | 1,00E+00 | 9,83E-01 | 8,71E+01 |
| UP_SEQ_FE | zinc finger region:C2H2-type 10 | 6 | 0,85 | 1,36E-01 | ZFP770, ZFP426, ZFP120, ZFP667, ZFP445, ZFP26 | 544 | 90 | 18012 | 2,21 | 1,00E+00 | 9,85E-01 | 9,12E+01 |
| UP_SEQ_FE | zinc finger region:C2H2-type 3 | 12 | 1,69 | 1,65E-01 | ZFP518A, PLAG1, KLF6, ZFP13, ADNP2, ZFP770, ZFP426, ZFP120, ZFP667, ZIK1, ZFP445, ZFP26 | 544 | 261 | 18012 | 1,52 | 1,00E+00 | 9,93E-01 | 9,51E+01 |
| UP_SEQ_FE | zinc finger region:C2H2-type 1 | 12 | 1,69 | 1,69E-01 | ZFP518A, PLAG1, KLF6, ZFP13, ADNP2, ZFP770, ZFP426, ZFP120, ZFP667, ZIK1, ZFP445, ZFP26 | 544 | 263 | 18012 | 1,51 | 1,00E+00 | 9,93E-01 | 9,54E+01 |
| UP_SEQ_FE | zinc finger region:C2H2-type 2 | 12 | 1,69 | 1,86E-01 | ZFP518A, PLAG1, KLF6, ZFP13, ADNP2, ZFP770, ZFP426, ZFP120, ZFP667, ZIK1, ZFP445, ZFP26 | 544 | 268 | 18012 | 1,48 | 1,00E+00 | 9,94E-01 | 9,68E+01 |
| UP_SEQ_FE | zinc finger region:C2H2-type 11 | 5 | 0,71 | 1,90E-01 | ZFP770, ZFP426, ZFP667, ZFP445, ZFP26 | 544 | 75 | 18012 | 2,21 | 1,00E+00 | 9,94E-01 | 9,70E+01 |
| UP_SEQ_FE | zinc finger region:C2H2-type 5 | 9 | 1,27 | 2,32E-01 | ZFP518A, PLAG1, ZFP13, ZFP770, ZFP426, ZFP667, ZIK1, ZFP445, ZFP26 | 544 | 194 | 18012 | 1,54 | 1,00E+00 | 9,97E-01 | 9,88E+01 |
| UP_SEQ_FE | zinc finger region:C2H2-type 12 | 3 | 0,42 | 6,12E-01 | ZFP667, ZFP445, ZFP26 | 544 | 68 | 18012 | 1,46 | 1,00E+00 | 1,00E+00 | 1,00E+02 |
| Annotation | Enrichment Score: 0.9515493889871464 | | | | | | | | | | | |
| Category | Term | Count | % | PValue | Genes | List To | Pop Hit | Pop Tot | Fold Enr | Bonferroni | Benjamini | FDR |
| INTERPRO | IPR008160:Collagen triple helix repeat | 6 | 0,85 | 8,94E-02 | COL4A3, COL14A1, SFTPD, COLEC10, COL25A1, COL5A2 | 645 | 76 | 20594 | 2,52 | 1,00E+00 | 9,44E-01 | 7,75E+01 |
| UP_KEYWO | Collagen | 6 | 0,85 | 1,10E-01 | COL4A3, COL14A1, SFTPD, COLEC10, COL25A1, COL5A2 | 678 | 85 | 22680 | 2,36 | 1,00E+00 | 6,13E-01 | 7,96E+01 |
| UP_KEYWO | Hydroxylation | 6 | 0,85 | 1,23E-01 | COL14A1, ELN, SFTPD, COL25A1, C1S1, COL5A2 | 678 | 88 | 22680 | 2,28 | 1,00E+00 | 6,21E-01 | 8,32E+01 |
| GOTERM_C | GO:0005581~collagen trimer | 6 | 0,85 | 1,29E-01 | COL4A3, COL14A1, SFTPD, COLEC10, COL25A1, COL5A2 | 633 | 83 | 19662 | 2,25 | 1,00E+00 | 6,96E-01 | 8,53E+01 |
| Annotation | Enrichment Score: 0.9326072354612176 | | | | | | | | | | | |
| Category | Term | Count | % | PValue | Genes | List To | Pop Hit | Pop Tot | Fold Enr | Bonferroni | Benjamini | FDR |
| GOTERM_M | GO:0030246~carbohydrate binding | 15 | 2,12 | 2,01E-02 | GLG1, SELP, C4A, LGALS1, G6PDX, PGD, HEXB, COLEC10, CD209C, CD93, CD34, SFTPD, CLEC7A, KLRB1B, PYGB | 575 | 229 | 17446 | 1,99 | 1,00E+00 | 8,13E-01 | 2,66E+01 |
| UP_SEQ_FE | domain:C-type lectin | 7 | 0,99 | 8,17E-02 | SELP, CD93, SFTPD, COLEC10, CLEC7A, KLRB1B, CD209C | 544 | 100 | 18012 | 2,32 | 1,00E+00 | 9,75E-01 | 7,59E+01 |

| | | | | | | | | | | | | |
|---|---|---|---|---|---|---|---|---|---|---|---|---|
| INTERPRO | IPR016187:C-type lectin fold | 9 | 1,27 | 8,54E-02 | COL4A3, SELP, HAPLN4, CD93, SFTPD, COLEC10, CLEC7A, KLRB1B, CD209C | 645 | 145 | 20594 | 1,98 | 1,00E+00 | 9,46E-01 | 7,59E+01 |
| INTERPRO | IPR016186:C-type lectin-like | 8 | 1,13 | 1,38E-01 | SELP, HAPLN4, CD93, SFTPD, COLEC10, CLEC7A, KLRB1B, CD209C | 645 | 137 | 20594 | 1,86 | 1,00E+00 | 9,70E-01 | 9,07E+01 |
| UP_KEYWO | Lectin | 9 | 1,27 | 1,47E-01 | GLG1, SELP, CD93, LGALS1, SFTPD, COLEC10, CLEC7A, KLRB1B, CD209C | 678 | 173 | 22680 | 1,74 | 1,00E+00 | 6,76E-01 | 8,85E+01 |
| INTERPRO | IPR001304:C-type lectin | 7 | 0,99 | 1,93E-01 | SELP, CD93, SFTPD, COLEC10, CLEC7A, KLRB1B, CD209C | 645 | 124 | 20594 | 1,80 | 1,00E+00 | 9,80E-01 | 9,67E+01 |
| INTERPRO | IPR018378:C-type lectin, conserved site | 4 | 0,56 | 2,13E-01 | SELP, SFTPD, COLEC10, CD209C | 645 | 51 | 20594 | 2,50 | 1,00E+00 | 9,84E-01 | 9,78E+01 |
| SMART | SM00034:CLECT | 7 | 0,99 | 2,94E-01 | SELP, CD93, SFTPD, COLEC10, CLEC7A, KLRB1B, CD209C | 379 | 124 | 10425 | 1,55 | 1,00E+00 | 9,81E-01 | 9,88E+01 |
| Annotation | Enrichment Score: 0.9321754415472475 | | | | | | | | | | | |
| Category | Term | Count | % | PValue | Genes | List To | Pop Hit | Pop Tota | Fold Enr | Bonferroni | Benjamini | FDR |
| UP_SEQ_FE | domain:Fibronectin type-III 2 | 9 | 1,27 | 3,41E-02 | NCAM2, PTPRC, COL14A1, PRLR, ROBO1, CNTN6, LEPR, SORL1, CSF2RB | 544 | 124 | 18012 | 2,40 | 1,00E+00 | 9,73E-01 | 4,40E+01 |
| UP_SEQ_FE | domain:Fibronectin type-III 1 | 9 | 1,27 | 3,56E-02 | NCAM2, PTPRC, COL14A1, PRLR, ROBO1, CNTN6, LEPR, SORL1, CSF2RB | 544 | 125 | 18012 | 2,38 | 1,00E+00 | 9,63E-01 | 4,53E+01 |
| SMART | SM00060:FN3 | 9 | 1,27 | 1,92E-01 | NCAM2, PTPRC, COL14A1, PRLR, ROBO1, CNTN6, LEPR, SORL1, CSF2RB | 379 | 153 | 10425 | 1,62 | 1,00E+00 | 9,59E-01 | 9,34E+01 |
| UP_SEQ_FE | domain:Fibronectin type-III 3 | 5 | 0,71 | 1,96E-01 | COL14A1, ROBO1, CNTN6, LEPR, SORL1 | 544 | 76 | 18012 | 2,18 | 1,00E+00 | 9,95E-01 | 9,74E+01 |
| INTERPRO | IPR003961:Fibronectin, type III | 10 | 1,41 | 2,20E-01 | NCAM2, PTPRC, COL14A1, PRLR, ROBO1, CNTN6, LEPR, SORL1, CSF2RB, IL3RA | 645 | 211 | 20594 | 1,51 | 1,00E+00 | 9,85E-01 | 9,81E+01 |
| UP_SEQ_FE | domain:Fibronectin type-III 4 | 4 | 0,56 | 2,54E-01 | COL14A1, CNTN6, LEPR, SORL1 | 544 | 58 | 18012 | 2,28 | 1,00E+00 | 9,98E-01 | 9,93E+01 |
| Annotation | Enrichment Score: 0.9241483091458282 | | | | | | | | | | | |
| Category | Term | Count | % | PValue | Genes | List To | Pop Hit | Pop Tota | Fold Enr | Bonferroni | Benjamini | FDR |
| GOTERM_M | GO:0042605~peptide antigen binding | 7 | 0,99 | 3,38E-03 | H2-Q2, H2-Q4, H2-BL, TAP1, H2-Q1, SLC7A9, H2-T10 | 575 | 45 | 17446 | 4,72 | 9,19E-01 | 3,95E-01 | 5,02E+00 |
| KEGG_PATH | mmu05416:Viral myocarditis | 8 | 1,13 | 9,35E-03 | ACTG1, H2-Q2, H2-BL, DMD, H2-Q1, H2-T10, CXADR, SGCB | 233 | 79 | 7720 | 3,36 | 8,99E-01 | 5,34E-01 | 1,14E+01 |
| GOTERM_C | GO:0005797~Golgi medial cisterna | 4 | 0,56 | 1,61E-02 | H2-Q2, GLG1, H2-Q4, H2-Q1 | 633 | 17 | 19662 | 7,31 | 9,98E-01 | 3,69E-01 | 2,02E+01 |
| GOTERM_M | GO:0046977~TAP binding | 3 | 0,42 | 2,65E-02 | H2-Q2, H2-Q4, H2-Q1 | 575 | 8 | 17446 | 11,38 | 1,00E+00 | 8,37E-01 | 3,35E+01 |

| | | | | | | | | | | | | |
|---|---|---|---|---|---|---|---|---|---|---|---|---|
| GOTERM_BI | GO:0002474~antigen processing and presentation of peptide antigen via MHC class I | 5 | 0,71 | 2,85E-02 | H2-Q2, H2-Q4, H2-BL, H2-Q1, H2-T10 | 588 | 36 | 18082 | 4,27 | 1,00E+00 | 8,86E-01 | 3,97E+01 |
| KEGG_PATH | mmu04145:Phagosome | 11 | 1,55 | 3,67E-02 | ACTG1, H2-Q2, ATP6V0E2, H2-BL, TAP1, H2-Q1, SFTPD, H2-T10, CLEC7A, CD209C, ATP6V0D2 | 233 | 174 | 7720 | 2,09 | 1,00E+00 | 7,28E-01 | 3,83E+01 |
| INTERPRO | IPR001039:MHC class I, alpha chain, alpha1/alpha2 | 5 | 0,71 | 5,15E-02 | H2-Q2, H2-Q4, H2-BL, H2-Q1, H2-T10 | 645 | 45 | 20594 | 3,55 | 1,00E+00 | 8,98E-01 | 5,69E+01 |
| GOTERM_M | GO:0030881~beta-2-microglobulin binding | 3 | 0,42 | 6,63E-02 | H2-Q2, H2-Q4, H2-Q1 | 575 | 13 | 17446 | 7,00 | 1,00E+00 | 9,02E-01 | 6,48E+01 |
| GOTERM_M | GO:0042608~T cell receptor binding | 3 | 0,42 | 7,57E-02 | H2-Q2, H2-Q4, H2-Q1 | 575 | 14 | 17446 | 6,50 | 1,00E+00 | 9,13E-01 | 6,98E+01 |
| GOTERM_C | GO:0042612~MHC class I protein complex | 3 | 0,42 | 9,20E-02 | H2-Q2, H2-Q4, H2-Q1 | 633 | 16 | 19662 | 5,82 | 1,00E+00 | 6,36E-01 | 7,39E+01 |
| KEGG_PATH | mmu04612:Antigen processing and presentation | 6 | 0,85 | 9,99E-02 | H2-Q2, RFX5, H2-BL, TAP1, H2-Q1, H2-T10 | 233 | 82 | 7720 | 2,42 | 1,00E+00 | 7,60E-01 | 7,44E+01 |
| INTERPRO | IPR011161:MHC class I-like antigen recognition | 5 | 0,71 | 1,08E-01 | H2-Q2, H2-Q4, H2-BL, H2-Q1, H2-T10 | 645 | 58 | 20594 | 2,75 | 1,00E+00 | 9,60E-01 | 8,37E+01 |
| GOTERM_C | GO:0070971~endoplasmic reticulum exit site | 3 | 0,42 | 1,23E-01 | H2-Q2, H2-Q4, H2-Q1 | 633 | 19 | 19662 | 4,90 | 1,00E+00 | 6,96E-01 | 8,40E+01 |
| KEGG_PATH | mmu05320:Autoimmune thyroid disease | 5 | 0,71 | 1,64E-01 | H2-Q2, H2-BL, H2-Q1, TPO, H2-T10 | 233 | 71 | 7720 | 2,33 | 1,00E+00 | 8,03E-01 | 9,02E+01 |
| INTERPRO | IPR011162:MHC classes I/II-like antigen recognition protein | 5 | 0,71 | 1,69E-01 | H2-Q2, H2-Q4, H2-BL, H2-Q1, H2-T10 | 645 | 69 | 20594 | 2,31 | 1,00E+00 | 9,75E-01 | 9,48E+01 |
| GOTERM_M | GO:0042277~peptide binding | 6 | 0,85 | 1,92E-01 | LNPEP, H2-Q2, H2-Q4, PPIC, H2-Q1, MAS1 | 575 | 93 | 17446 | 1,96 | 1,00E+00 | 9,77E-01 | 9,61E+01 |
| KEGG_PATH | mmu05332:Graft-versus-host disease | 4 | 0,56 | 2,05E-01 | H2-Q2, H2-BL, H2-Q1, H2-T10 | 233 | 52 | 7720 | 2,55 | 1,00E+00 | 8,35E-01 | 9,49E+01 |
| KEGG_PATH | mmu05330:Allograft rejection | 4 | 0,56 | 2,37E-01 | H2-Q2, H2-BL, H2-Q1, H2-T10 | 233 | 56 | 7720 | 2,37 | 1,00E+00 | 8,56E-01 | 9,70E+01 |

| | | | | | | | | | | | | |
|---|---|---|---|---|---|---|---|---|---|---|---|---|
| INTERPRO | IPR003006:Immunoglobulin/major histocompatibility complex, conserved site | 5 | 0,71 | 2,75E-01 | H2-Q2, H2-Q4, H2-BL, H2-Q1, H2-T10 | 645 | 85 | 20594 | 1,88 | 1,00E+00 | 9,92E-01 | 9,94E+01 |
| KEGG_PATH | mmu04940:Type I diabetes mellitus | 4 | 0,56 | 2,86E-01 | H2-Q2, H2-BL, H2-Q1, H2-T10 | 233 | 62 | 7720 | 2,14 | 1,00E+00 | 8,71E-01 | 9,87E+01 |
| INTERPRO | IPR003597:Immunoglobulin C1-set | 5 | 0,71 | 3,89E-01 | H2-Q2, H2-Q4, H2-BL, H2-Q1, H2-T10 | 645 | 101 | 20594 | 1,58 | 1,00E+00 | 9,98E-01 | 1,00E+02 |
| KEGG_PATH | mmu05168:Herpes simplex infection | 8 | 1,13 | 4,37E-01 | H2-Q2, CCL2, H2-BL, TAP1, H2-Q1, TLR3, H2-T10, OAS1A | 233 | 208 | 7720 | 1,27 | 1,00E+00 | 9,22E-01 | 9,99E+01 |
| SMART | SM00407:IGc1 | 5 | 0,71 | 4,78E-01 | H2-Q2, H2-Q4, H2-BL, H2-Q1, H2-T10 | 379 | 98 | 10425 | 1,40 | 1,00E+00 | 9,91E-01 | 1,00E+02 |
| KEGG_PATH | mmu05203:Viral carcinogenesis | 8 | 1,13 | 5,46E-01 | H2-Q2, CDKN1A, CCR5, H2-BL, H2-Q1, H2-T10, ATP6V0D2, SRC | 233 | 231 | 7720 | 1,15 | 1,00E+00 | 9,48E-01 | 1,00E+02 |
| KEGG_PATH | mmu05166:HTLV-I infection | 9 | 1,27 | 6,01E-01 | H2-Q2, VCAM1, CDKN1A, SLC25A4, CDKN2C, H2-BL, H2-Q1, PPP3CC, H2-T10 | 233 | 278 | 7720 | 1,07 | 1,00E+00 | 9,59E-01 | 1,00E+02 |
| KEGG_PATH | mmu05169:Epstein-Barr virus infection | 7 | 0,99 | 6,32E-01 | H2-Q2, CDKN1A, CR2, H2-BL, H2-Q1, H2-T10, ENTPD3 | 233 | 215 | 7720 | 1,08 | 1,00E+00 | 9,58E-01 | 1,00E+02 |
| KEGG_PATH | mmu04144:Endocytosis | 8 | 1,13 | 7,37E-01 | H2-Q2, CCR5, H2-BL, H2-Q1, H2-T10, GRK5, EHD1, SRC | 233 | 278 | 7720 | 0,95 | 1,00E+00 | 9,71E-01 | 1,00E+02 |
| Annotation | Enrichment Score: 0.9127080901453 | | | | | | | | | | | |
| Category | Term | Count | % | PValue | Genes | List To | Pop Hit | Pop Tot | Fold Enr | Bonferroni | Benjamini | FDR |
| INTERPRO | IPR008068:Cytochrome P450, E-class, group I, CYP2B-like | 3 | 0,42 | 1,35E-02 | CYP2B9, CYP2B13, CYP2B10 | 645 | 6 | 20594 | 15,96 | 1,00E+00 | 7,04E-01 | 1,95E+01 |
| GOTERM_BI | GO:0019373~epoxygenase P450 pathway | 5 | 0,71 | 1,54E-02 | CYP2A22, CYP2B9, CYP2B13, CYP2B10, CYP2A4 | 588 | 30 | 18082 | 5,13 | 1,00E+00 | 8,52E-01 | 2,38E+01 |
| GOTERM_M | GO:0008392~arachidonic acid epoxygenase activity | 6 | 0,85 | 1,86E-02 | CYP2A22, CYP2B9, CYP2D12, CYP2B13, CYP2B10, CYP2A4 | 575 | 47 | 17446 | 3,87 | 1,00E+00 | 8,26E-01 | 2,49E+01 |
| GOTERM_M | GO:0008395~steroid hydroxylase activity | 6 | 0,85 | 3,19E-02 | CYP2A22, CYP2B9, CYP2D12, CYP2B13, CYP2B10, CYP2A4 | 575 | 54 | 17446 | 3,37 | 1,00E+00 | 8,43E-01 | 3,89E+01 |
| INTERPRO | IPR002401:Cytochrome P450, E-class, group I | 7 | 0,99 | 4,56E-02 | CYP2A22, CYP4F16, CYP2B9, CYP2D12, CYP2B13, CYP2B10, CYP2A4 | 645 | 83 | 20594 | 2,69 | 1,00E+00 | 8,98E-01 | 5,25E+01 |
| UP_KEYWO | Monooxygenase | 9 | 1,27 | 4,62E-02 | CYP2A22, CYP4F16, FMO2, CYP2B9, CYP2D12, FMO3, CYP2B13, CYP2B10, CYP2A4 | 678 | 133 | 22680 | 2,26 | 1,00E+00 | 4,67E-01 | 4,74E+01 |

| | | | | | | | | | | | | |
|---|---|---|---|---|---|---|---|---|---|---|---|---|
| GOTERM_M | GO:0016712~oxidoreductase activity, acting on paired donors, with incorporation or reduction of molecular oxygen, reduced flavin or flavoprotein as one donor, and incorporation of one atom of oxygen | 5 | 0,71 | 4,85E-02 | CYP2A22, CYP2B9, CYP2D12, CYP2B13, CYP2B10 | 575 | 42 | 17446 | 3,61 | 1,00E+00 | 9,01E-01 | 5,30E+01 |
| INTERPRO | IPR017972:Cytochrome P450, conserved site | 7 | 0,99 | 8,04E-02 | CYP2A22, CYP4F16, CYP2B9, CYP2D12, CYP2B13, CYP2B10, CYP2A4 | 645 | 96 | 20594 | 2,33 | 1,00E+00 | 9,41E-01 | 7,38E+01 |
| GOTERM_C | GO:0031090~organelle membrane | 7 | 0,99 | 8,23E-02 | FMO2, CYP2B9, FMO3, DHRS9, SRD5A2, CYP2B10, CYP2A4 | 633 | 94 | 19662 | 2,31 | 1,00E+00 | 6,34E-01 | 6,98E+01 |
| UP_KEYWO | Microsome | 8 | 1,13 | 1,01E-01 | FMO2, CYP2B9, FMO3, DHRS9, OAS1A, SRD5A2, CYP2B10, CYP2A4 | 678 | 132 | 22680 | 2,03 | 1,00E+00 | 6,06E-01 | 7,63E+01 |
| INTERPRO | IPR001128:Cytochrome P450 | 7 | 0,99 | 1,08E-01 | CYP2A22, CYP4F16, CYP2B9, CYP2D12, CYP2B13, CYP2B10, CYP2A4 | 645 | 104 | 20594 | 2,15 | 1,00E+00 | 9,57E-01 | 8,37E+01 |
| GOTERM_B | GO:0017144~drug metabolic process | 3 | 0,42 | 1,14E-01 | FMO2, FMO3, CYP2B10 | 588 | 18 | 18082 | 5,13 | 1,00E+00 | 9,83E-01 | 8,81E+01 |
| GOTERM_M | GO:0020037~heme binding | 10 | 1,41 | 1,24E-01 | CYP2A22, MFSD7C, CYP4F16, CYP2B9, CYP2D12, TPO, CYP2B13, CYP2B10, SRC, CYP2A4 | 575 | 175 | 17446 | 1,73 | 1,00E+00 | 9,62E-01 | 8,67E+01 |
| KEGG_PATH | mmu00830:Retinol metabolism | 6 | 0,85 | 1,29E-01 | CYP2B9, DHRS9, CYP2B13, CYP2B10, CYP2A4, RDH5 | 233 | 89 | 7720 | 2,23 | 1,00E+00 | 8,00E-01 | 8,33E+01 |
| UP_KEYWO | Heme | 9 | 1,27 | 1,41E-01 | STEAP4, CYP2A22, CYP4F16, CYP2B9, CYP2D12, TPO, CYP2B13, CYP2B10, CYP2A4 | 678 | 171 | 22680 | 1,76 | 1,00E+00 | 6,66E-01 | 8,72E+01 |
| UP_SEQ_FE | metal ion-binding site:Iron (heme axial ligand) | 6 | 0,85 | 2,61E-01 | STEAP4, CYP2A22, CYP2B9, TPO, CYP2B10, CYP2A4 | 544 | 114 | 18012 | 1,74 | 1,00E+00 | 9,98E-01 | 9,94E+01 |
| KEGG_PATH | mmu00140:Steroid hormone biosynthesis | 5 | 0,71 | 2,65E-01 | CYP2B9, CYP2D12, CYP2B13, SRD5A2, CYP2B10 | 233 | 87 | 7720 | 1,90 | 1,00E+00 | 8,69E-01 | 9,82E+01 |
| GOTERM_M | GO:0004497~monooxygenase activity | 6 | 0,85 | 2,83E-01 | CYP4F16, FMO2, CYP2B9, CYP2D12, FMO3, CYP2B10 | 575 | 108 | 17446 | 1,69 | 1,00E+00 | 9,81E-01 | 9,94E+01 |
| KEGG_PATH | mmu05204:Chemical carcinogenesis | 5 | 0,71 | 2,99E-01 | CYP2B9, CYP2B13, GSTO2, CYP2B10, MGST2 | 233 | 92 | 7720 | 1,80 | 1,00E+00 | 8,79E-01 | 9,90E+01 |
| UP_KEYWO | Iron | 14 | 1,97 | 3,28E-01 | STEAP4, CYP2B9, CYP2D12, POLA1, HR, CYP2B13, EXO5, CYP2B10, CYP2A22, CYP4F16, MIOX, PPP3CC, TPO, CYP2A4 | 678 | 375 | 22680 | 1,25 | 1,00E+00 | 8,37E-01 | 9,95E+01 |
| GOTERM_M | GO:0070330~aromatase activity | 3 | 0,42 | 3,33E-01 | CYP2B9, CYP2B10, CYP2A4 | 575 | 36 | 17446 | 2,53 | 1,00E+00 | 9,89E-01 | 9,98E+01 |

| | | | | | | | | | | | | |
|---|---|---|---|---|---|---|---|---|---|---|---|---|
| GOTERM_M | GO:0016705~oxidoreductase activity, acting on paired donors, with incorporation or reduction of molecular oxygen | 5 | 0,71 | 4,11E-01 | CYP4F16, CYP2B9, CYP2D12, CYP2B13, CYP2B10 | 575 | 99 | 17446 | 1,53 | 1,00E+00 | 9,95E-01 | 1,00E+02 |
| GOTERM_M | GO:0005506~iron ion binding | 8 | 1,13 | 5,50E-01 | CYP2A22, CYP4F16, CYP2B9, CYP2D12, MIOX, CYP2B13, CYP2B10, CYP2A4 | 575 | 212 | 17446 | 1,14 | 1,00E+00 | 9,99E-01 | 1,00E+02 |
| COG_ONTO | Secondary metabolites biosynthesis, transport, and catabolism | 7 | 0,99 | 5,99E-01 | CYP2A22, CYP4F16, CYP2B9, CYP2D12, CYP2B13, CYP2B10, CYP2A4 | 114 | 117 | 2126 | 1,12 | 1,00E+00 | 9,63E-01 | 9,99E+01 |
| KEGG_PATH | mmu00590:Arachidonic acid metabolism | 3 | 0,42 | 7,53E-01 | CYP2B9, CYP2B13, CYP2B10 | 233 | 89 | 7720 | 1,12 | 1,00E+00 | 9,75E-01 | 1,00E+02 |
| Annotation | Enrichment Score: 0.9086540543564907 | | | | | | | | | | | |
| Category | Term | Count | % | PValue | Genes | List To | Pop Hit | Pop Tota | Fold Enr | Bonferroni | Benjamini | FDR |
| GOTERM_BI | GO:0040007~growth | 5 | 0,71 | 2,85E-02 | INHBA, BMP10, BMP5, BMP8B, FOXP2 | 588 | 36 | 18082 | 4,27 | 1,00E+00 | 8,86E-01 | 3,97E+01 |
| INTERPRO | IPR001111:Transforming growth factor-beta, N-terminal | 4 | 0,56 | 3,39E-02 | INHBA, BMP10, BMP5, BMP8B | 645 | 23 | 20594 | 5,55 | 1,00E+00 | 8,58E-01 | 4,23E+01 |
| GOTERM_BI | GO:0060395~SMAD protein signal transduction | 7 | 0,99 | 3,89E-02 | LNPEP, INHBA, AFP, BMP10, MEG3, BMP5, BMP8B | 588 | 77 | 18082 | 2,80 | 1,00E+00 | 9,24E-01 | 5,01E+01 |
| GOTERM_BI | GO:0043408~regulation of MAPK cascade | 6 | 0,85 | 4,22E-02 | INHBA, BMP10, E130311K13RIK, PAK1, BMP5, BMP8B | 588 | 59 | 18082 | 3,13 | 1,00E+00 | 9,19E-01 | 5,30E+01 |
| GOTERM_M | GO:0005125~cytokine activity | 13 | 1,83 | 5,07E-02 | BMP10, INHBA, TNFSF10, CCL2, CXCL14, IL7, IL1RN, IL33, BMP5, BMP8B, SPP1, TIMP1, CXCL10 | 575 | 214 | 17446 | 1,84 | 1,00E+00 | 8,83E-01 | 5,46E+01 |
| INTERPRO | IPR015615:Transforming growth factor-beta-related | 4 | 0,56 | 6,62E-02 | INHBA, BMP10, BMP5, BMP8B | 645 | 30 | 20594 | 4,26 | 1,00E+00 | 9,35E-01 | 6,65E+01 |
| INTERPRO | IPR017948:Transforming growth factor beta, conserved site | 4 | 0,56 | 7,73E-02 | INHBA, BMP10, BMP5, BMP8B | 645 | 32 | 20594 | 3,99 | 1,00E+00 | 9,45E-01 | 7,23E+01 |
| UP_KEYWO | Growth factor | 8 | 1,13 | 8,63E-02 | INHBA, BMP10, IL7, KLK1B4, HGF, BMP5, BMP8B, TIMP1 | 678 | 127 | 22680 | 2,11 | 1,00E+00 | 5,66E-01 | 7,07E+01 |
| INTERPRO | IPR001839:Transforming growth factor-beta, C-terminal | 4 | 0,56 | 1,08E-01 | INHBA, BMP10, BMP5, BMP8B | 645 | 37 | 20594 | 3,45 | 1,00E+00 | 9,54E-01 | 8,39E+01 |
| SMART | SM00204:TGFB | 4 | 0,56 | 1,32E-01 | INHBA, BMP10, BMP5, BMP8B | 379 | 35 | 10425 | 3,14 | 1,00E+00 | 9,15E-01 | 8,37E+01 |

| | | | | | | | | | | | | |
|---|---|---|---|---|---|---|---|---|---|---|---|---|
| GOTERM_M | GO:0005160~transforming growth factor beta receptor binding | 4 | 0,56 | 1,76E-01 | INHBA, BMP10, BMP5, BMP8B | 575 | 44 | 17446 | 2,76 | 1,00E+00 | 9,79E-01 | 9,47E+01 |
| GOTERM_BI | GO:0042981~regulation of apoptotic process | 10 | 1,41 | 2,00E-01 | INHBA, BMP10, TNFRSF11B, BCL2L14, ANK2, CASP12, ESR1, TRIM24, BMP5, BMP8B | 588 | 199 | 18082 | 1,55 | 1,00E+00 | 9,95E-01 | 9,80E+01 |
| GOTERM_M | GO:0008083~growth factor activity | 8 | 1,13 | 2,01E-01 | INHBA, BMP10, IL7, KLK1B4, HGF, BMP5, BMP8B, TIMP1 | 575 | 145 | 17446 | 1,67 | 1,00E+00 | 9,69E-01 | 9,67E+01 |
| GOTERM_BI | GO:0010862~positive regulation of pathway-restricted SMAD protein phosphorylation | 4 | 0,56 | 2,04E-01 | INHBA, BMP10, BMP5, BMP8B | 588 | 48 | 18082 | 2,56 | 1,00E+00 | 9,95E-01 | 9,81E+01 |
| KEGG_PATH | mmu04350:TGF-beta signaling pathway | 5 | 0,71 | 2,52E-01 | INHBA, ID2, DCN, BMP5, BMP8B | 233 | 85 | 7720 | 1,95 | 1,00E+00 | 8,60E-01 | 9,77E+01 |
| GOTERM_BI | GO:0051216~cartilage development | 4 | 0,56 | 5,00E-01 | LUM, BMP5, BMP8B, TIMP1 | 588 | 82 | 18082 | 1,50 | 1,00E+00 | 1,00E+00 | 1,00E+02 |
| UP_KEYWO | Cleavage on pair of basic residues | 8 | 1,13 | 6,12E-01 | INHBA, BMP10, BGLAP, SORL1, CDH1, MMP15, BMP5, BGLAP3 | 678 | 248 | 22680 | 1,08 | 1,00E+00 | 9,40E-01 | 1,00E+02 |
| GOTERM_BI | GO:0030509~BMP signaling pathway | 3 | 0,42 | 7,79E-01 | BMP10, BMP5, BMP8B | 588 | 87 | 18082 | 1,06 | 1,00E+00 | 1,00E+00 | 1,00E+02 |
| Annotation | Enrichment Score: 0.9070638360396251 | | | | | | | | | | | |
| Category | Term | Count | % | PValue | Genes | List To | Pop Hit | Pop Tot | Fold Enr | Bonferroni | Benjamini | FDR |
| UP_SEQ_FE | short sequence motif:Box 1 motif | 4 | 0,56 | 4,24E-02 | PRLR, LEPR, CSF2RB, IL3RA | 544 | 26 | 18012 | 5,09 | 1,00E+00 | 9,46E-01 | 5,14E+01 |
| GOTERM_M | GO:0004896~cytokine receptor activity | 5 | 0,71 | 5,98E-02 | PRLR, LEPR, CSF2RB, IL3RA, IL17RB | 575 | 45 | 17446 | 3,37 | 1,00E+00 | 8,99E-01 | 6,09E+01 |
| UP_SEQ_FE | short sequence motif:WSXWS motif | 4 | 0,56 | 8,19E-02 | PRLR, LEPR, CSF2RB, IL3RA | 544 | 34 | 18012 | 3,90 | 1,00E+00 | 9,72E-01 | 7,60E+01 |
| INTERPRO | IPR003961:Fibronectin, type III | 10 | 1,41 | 2,20E-01 | NCAM2, PTPRC, COL14A1, PRLR, ROBO1, CNTN6, LEPR, SORL1, CSF2RB, IL3RA | 645 | 211 | 20594 | 1,51 | 1,00E+00 | 9,85E-01 | 9,81E+01 |
| KEGG_PATH | mmu04630:Jak-STAT signaling pathway | 5 | 0,71 | 6,39E-01 | PRLR, IL7, LEPR, CSF2RB, IL3RA | 233 | 145 | 7720 | 1,14 | 1,00E+00 | 9,55E-01 | 1,00E+02 |
| Annotation | Enrichment Score: 0.9024238182280925 | | | | | | | | | | | |
| Category | Term | Count | % | PValue | Genes | List To | Pop Hit | Pop Tot | Fold Enr | Bonferroni | Benjamini | FDR |
| UP_SEQ_FE | metal ion-binding site:Calcium 1 | 5 | 0,71 | 5,26E-02 | PRKCA, SYT1, BGLAP, NPTX1, MMP12 | 544 | 47 | 18012 | 3,52 | 1,00E+00 | 9,50E-01 | 5,94E+01 |

| | | | | | | | | | | | | |
|---|---|---|---|---|---|---|---|---|---|---|---|---|
| UP_SEQ_FE/ | metal ion-binding site:Calcium 3 | 4 | 0,56 | 6,57E-02 | PRKCA, SYT1, BGLAP, MMP12 | 544 | 31 | 18012 | 4,27 | 1,00E+00 | 9,63E-01 | 6,78E+01 |
| UP_SEQ_FE/ | metal ion-binding site:Calcium 2 | 5 | 0,71 | 9,73E-02 | PRKCA, SYT1, BGLAP, NPTX1, MMP12 | 544 | 58 | 18012 | 2,85 | 1,00E+00 | 9,73E-01 | 8,19E+01 |
| UP_SEQ_FE/ | metal ion-binding site:Calcium 3; via carbonyl oxygen | 3 | 0,42 | 1,31E-01 | PRKCA, SYT1, MMP12 | 544 | 21 | 18012 | 4,73 | 1,00E+00 | 9,84E-01 | 9,04E+01 |
| UP_SEQ_FE/ | metal ion-binding site:Calcium 1; via carbonyl oxygen | 3 | 0,42 | 2,85E-01 | PRKCA, SYT1, NPTX1 | 544 | 35 | 18012 | 2,84 | 1,00E+00 | 9,98E-01 | 9,96E+01 |
| UP_SEQ_FE/ | metal ion-binding site:Calcium 2; via carbonyl oxygen | 3 | 0,42 | 3,07E-01 | PRKCA, SYT1, MMP12 | 544 | 37 | 18012 | 2,68 | 1,00E+00 | 9,99E-01 | 9,98E+01 |
| Annotation | Enrichment Score: 0.8816386977984355 | | | | | | | | | | | |
| Category | Term | Count | % | PValue | Genes | List To | Pop Hit | Pop Tota | Fold Enr | Bonferroni | Benjamini | FDR |
| UP_KEYWO | Immunoglobulin domain | 24 | 3,39 | 1,85E-02 | VSIG10, CD244, SKINT9, HAPLN4, CLMP, CNTN6, VTCN1, LEPR, FLT4, CD276, TARM1, LY9, CXADR, VCAM1, ALCAM, NCAM2, AMIGO2, IL18BP, UNC5B, ROBO1, SEMA3D, SCN4B, CD300LB, SEMA4A | 678 | 481 | 22680 | 1,67 | 9,98E-01 | 2,79E-01 | 2,24E+01 |
| INTERPRO | IPR003598:Immunoglobulin subtype 2 | 14 | 1,97 | 4,21E-02 | VSIG10, HAPLN4, CLMP, VTCN1, CNTN6, FLT4, CD276, CXADR, VCAM1, CCDC141, AMIGO2, NCAM2, UNC5B, ROBO1 | 645 | 242 | 20594 | 1,85 | 1,00E+00 | 8,90E-01 | 4,96E+01 |
| UP_SEQ_FE/ | domain:Ig-like C2-type 1 | 9 | 1,27 | 4,66E-02 | ALCAM, VCAM1, NCAM2, CLMP, ROBO1, CNTN6, FLT4, LY9, CXADR | 544 | 132 | 18012 | 2,26 | 1,00E+00 | 9,54E-01 | 5,49E+01 |
| UP_SEQ_FE/ | domain:Ig-like C2-type 2 | 9 | 1,27 | 4,83E-02 | ALCAM, VCAM1, NCAM2, CLMP, ROBO1, CNTN6, FLT4, LY9, CXADR | 544 | 133 | 18012 | 2,24 | 1,00E+00 | 9,54E-01 | 5,62E+01 |
| UP_SEQ_FE/ | domain:Ig-like C2-type 5 | 5 | 0,71 | 5,61E-02 | VCAM1, NCAM2, ROBO1, CNTN6, FLT4 | 544 | 48 | 18012 | 3,45 | 1,00E+00 | 9,55E-01 | 6,18E+01 |
| UP_SEQ_FE/ | domain:Ig-like C2-type 4 | 5 | 0,71 | 8,81E-02 | VCAM1, NCAM2, ROBO1, CNTN6, FLT4 | 544 | 56 | 18012 | 2,96 | 1,00E+00 | 9,75E-01 | 7,85E+01 |
| INTERPRO | IPR013098:Immunoglobulin I-set | 9 | 1,27 | 9,07E-02 | CCDC141, VCAM1, VSIG10, NCAM2, UNC5B, ROBO1, CNTN6, FLT4, CXADR | 645 | 147 | 20594 | 1,95 | 1,00E+00 | 9,42E-01 | 7,81E+01 |
| SMART | SM00408:IGc2 | 14 | 1,97 | 1,02E-01 | VSIG10, HAPLN4, CLMP, VTCN1, CNTN6, FLT4, CD276, CXADR, VCAM1, CCDC141, AMIGO2, NCAM2, UNC5B, ROBO1 | 379 | 242 | 10425 | 1,59 | 1,00E+00 | 8,90E-01 | 7,47E+01 |
| UP_SEQ_FE/ | domain:Ig-like C2-type 3 | 6 | 0,85 | 1,27E-01 | ALCAM, VCAM1, NCAM2, ROBO1, CNTN6, FLT4 | 544 | 88 | 18012 | 2,26 | 1,00E+00 | 9,84E-01 | 8,96E+01 |
| INTERPRO | IPR003599:Immunoglobulin subtype | 22 | 3,1 | 1,38E-01 | VSIG10, HAPLN4, CLMP, CNTN6, VTCN1, FLT4, CD276, TARM1, LY9, CXADR, BTNL6, BTNL4, CCDC141, VCAM1, ALCAM, NCAM2, AMIGO2, UNC5B, ROBO1, GM609, SCN4B, CD300LB | 645 | 518 | 20594 | 1,36 | 1,00E+00 | 9,71E-01 | 9,06E+01 |

| | | | | | | | | | | | | |
|---|---|---|---|---|---|---|---|---|---|---|---|---|
| INTERPRO | IPR013783:Immunoglobulin-like fold | 40 | 5,64 | 2,24E-01 | SKINT9, CD244, PKHD1, VTCN1, LEPR, SORL1, LY9, TARM1, CXADR, CCDC141, ALCAM, VCAM1, UNC5B, ROBO1, TGM1, SEMA3D, GM609, CSF2RB, H2-T10, H2-Q2, VSIG10, PTPRC, H2-Q4, HAPLN4, CLMP, PLXNB1, CNTN6, FLT4, H2-Q1, CD276, BTNL6, BTNL4, AMIGO2, NCAM2, COL14A1, PRLR, H2-BL, SCN4B, CD300LB, IL3RA | 645 | 1099 | 20594 | 1,16 | 1,00E+00 | 9,84E-01 | 9,82E+01 |
| UP_SEQ_FE | domain:Ig-like C2-type 6 | 3 | 0,42 | 2,40E-01 | VCAM1, CNTN6, FLT4 | 544 | 31 | 18012 | 3,20 | 1,00E+00 | 9,98E-01 | 9,90E+01 |
| SMART | SM00409:IG | 22 | 3,1 | 3,27E-01 | VSIG10, HAPLN4, CLMP, CNTN6, VTCN1, FLT4, CD276, TARM1, LY9, CXADR, BTNL6, BTNL4, CCDC141, VCAM1, ALCAM, NCAM2, AMIGO2, UNC5B, ROBO1, GM609, SCN4B, CD300LB | 379 | 518 | 10425 | 1,17 | 1,00E+00 | 9,79E-01 | 9,94E+01 |
| INTERPRO | IPR007110:Immunoglobulin-like domain | 32 | 4,51 | 3,48E-01 | SKINT9, CD244, VTCN1, LEPR, LY9, TARM1, CXADR, CCDC141, ALCAM, VCAM1, UNC5B, ROBO1, SEMA3D, GM609, H2-T10, H2-Q2, VSIG10, H2-Q4, HAPLN4, CLMP, CNTN6, FLT4, H2-Q1, CD276, BTNL6, BTNL4, AMIGO2, NCAM2, IL18BP, H2-BL, SCN4B, CD300LB | 645 | 920 | 20594 | 1,11 | 1,00E+00 | 9,97E-01 | 9,99E+01 |
| INTERPRO | IPR013106:Immunoglobulin V-set | 13 | 1,83 | 9,32E-01 | VSIG10, HAPLN4, CLMP, VTCN1, CD276, BTNL6, CXADR, BTNL4, NCAM2, ROBO1, GM609, SCN4B, CD300LB | 645 | 554 | 20594 | 0,75 | 1,00E+00 | 1,00E+00 | 1,00E+02 |
| SMART | SM00406:IGv | 9 | 1,27 | 9,88E-01 | VSIG10, NCAM2, HAPLN4, CLMP, ROBO1, GM609, CD276, CXADR, BTNL4 | 379 | 423 | 10425 | 0,59 | 1,00E+00 | 1,00E+00 | 1,00E+02 |
| Annotation | Enrichment Score: 0.8516848375338978 | | | | | | | | | | | |
| Category | Term | Count | % | PValue | Genes | List To | Pop Hit | Pop Tota | Fold Enr | Bonferroni | Benjamini | FDR |
| KEGG_PATH | mmu00500:Starch and sucrose metabolism | 4 | 0,56 | 7,02E-02 | AMY1, HKDC1, PGM1, PYGB | 233 | 32 | 7720 | 4,14 | 1,00E+00 | 8,01E-01 | 6,10E+01 |
| GOTERM_B | GO:0005975~carbohydrate metabolic process | 11 | 1,55 | 1,35E-01 | AMY1, HHIPL2, CHIL1, HKDC1, HEXB, PGD, G6PDX, PGM1, FUCA2, NEU3, PYGB | 588 | 206 | 18082 | 1,64 | 1,00E+00 | 9,87E-01 | 9,20E+01 |
| UP_KEYWO | Carbohydrate metabolism | 5 | 0,71 | 2,95E-01 | AMY1, G6PDX, PGM1, NEU3, PYGB | 678 | 92 | 22680 | 1,82 | 1,00E+00 | 8,11E-01 | 9,91E+01 |
| Annotation | Enrichment Score: 0.8094928443952004 | | | | | | | | | | | |
| Category | Term | Count | % | PValue | Genes | List To | Pop Hit | Pop Tota | Fold Enr | Bonferroni | Benjamini | FDR |
| KEGG_PATH | mmu05416:Viral myocarditis | 8 | 1,13 | 9,35E-03 | ACTG1, H2-Q2, H2-BL, DMD, H2-Q1, H2-T10, CXADR, SGCB | 233 | 79 | 7720 | 3,36 | 8,99E-01 | 5,34E-01 | 1,14E+01 |
| KEGG_PATH | mmu05410:Hypertrophic cardiomyopathy (HCM) | 5 | 0,71 | 2,13E-01 | ACTG1, ACE, DMD, TPM4, SGCB | 233 | 79 | 7720 | 2,10 | 1,00E+00 | 8,39E-01 | 9,55E+01 |
| KEGG_PATH | mmu05414:Dilated cardiomyopathy | 4 | 0,56 | 4,57E-01 | ACTG1, DMD, TPM4, SGCB | 233 | 83 | 7720 | 1,60 | 1,00E+00 | 9,24E-01 | 1,00E+02 |

| | | | | | | | | | | | | |
|---|---|---|---|---|---|---|---|---|---|---|---|---|
| KEGG_PATH | mmu05412:Arrhythmogenic right ventricular cardiomyopathy (ARVC) | 3 | 0,42 | 6,35E-01 | ACTG1, DMD, SGCB | 233 | 71 | 7720 | 1,40 | 1,00E+00 | 9,55E-01 | 1,00E+02 |
| Annotation | Enrichment Score: 0.762893028392063 | | | | | | | | | | | |
| Category | Term | Count | % | PValue | Genes | List To | Pop Hit | Pop Tot | Fold Enr | Bonferroni | Benjamini | FDR |
| UP_SEQ_FE | domain:PARP catalytic | 3 | 0,42 | 4,17E-02 | PARP16, PARP11, PARP2 | 544 | 11 | 18012 | 9,03 | 1,00E+00 | 9,50E-01 | 5,08E+01 |
| INTERPRO | IPR012317:Poly(ADP-ribose) polymerase, catalytic domain | 3 | 0,42 | 8,78E-02 | PARP16, PARP11, PARP2 | 645 | 16 | 20594 | 5,99 | 1,00E+00 | 9,46E-01 | 7,69E+01 |
| GOTERM_M | GO:0003950~NAD+ ADP-ribosyltransferase activity | 3 | 0,42 | 1,51E-01 | PARP16, PARP11, PARP2 | 575 | 21 | 17446 | 4,33 | 1,00E+00 | 9,72E-01 | 9,17E+01 |
| UP_KEYWO | NAD | 9 | 1,27 | 1,82E-01 | PARP16, GAPDHS, STEAP4, ALDH1B1, DHRS9, PARP11, PARP2, CRYM, RDH5 | 678 | 183 | 22680 | 1,65 | 1,00E+00 | 7,24E-01 | 9,35E+01 |
| UP_KEYWO | ADP-ribosylation | 3 | 0,42 | 3,36E-01 | PARP16, PARP11, PARP2 | 678 | 40 | 22680 | 2,51 | 1,00E+00 | 8,38E-01 | 9,96E+01 |
| UP_KEYWO | Glycosyltransferase | 9 | 1,27 | 3,57E-01 | PARP16, ALG10B, B3GALT1, PARP11, DPY19L3, GTDC1, PARP2, POGLUT1, PYGB | 678 | 225 | 22680 | 1,34 | 1,00E+00 | 8,33E-01 | 9,97E+01 |
| GOTERM_M | GO:0016757~transferase activity, transferring glycosyl groups | 9 | 1,27 | 3,79E-01 | PARP16, ALG10B, B3GALT1, PARP11, DPY19L3, GTDC1, PARP2, POGLUT1, PYGB | 575 | 208 | 17446 | 1,31 | 1,00E+00 | 9,95E-01 | 9,99E+01 |
| Annotation | Enrichment Score: 0.7576785381402362 | | | | | | | | | | | |
| Category | Term | Count | % | PValue | Genes | List To | Pop Hit | Pop Tot | Fold Enr | Bonferroni | Benjamini | FDR |
| INTERPRO | IPR000566:Lipocalin/cytosolic fatty-acid binding protein domain | 5 | 0,71 | 1,46E-01 | MUP21, RBP1, OBP2A, FABP12, ORM3 | 645 | 65 | 20594 | 2,46 | 1,00E+00 | 9,69E-01 | 9,19E+01 |
| INTERPRO | IPR012674:Calycin | 5 | 0,71 | 1,88E-01 | MUP21, RBP1, OBP2A, FABP12, ORM3 | 645 | 72 | 20594 | 2,22 | 1,00E+00 | 9,81E-01 | 9,64E+01 |
| INTERPRO | IPR011038:Calycin-like | 5 | 0,71 | 1,95E-01 | MUP21, RBP1, OBP2A, FABP12, ORM3 | 645 | 73 | 20594 | 2,19 | 1,00E+00 | 9,79E-01 | 9,68E+01 |
| Annotation | Enrichment Score: 0.7521655031565833 | | | | | | | | | | | |
| Category | Term | Count | % | PValue | Genes | List To | Pop Hit | Pop Tot | Fold Enr | Bonferroni | Benjamini | FDR |

| | | | | | | | | | | | | |
|---|---|---|---|---|---|---|---|---|---|---|---|---|
| KEGG_PATH | mmu00511:Other glycan degradation | 3 | 0,42 | 1,00E-01 | HEXB, FUCA2, NEU3 | 233 | 18 | 7720 | 5,52 | 1,00E+00 | 7,43E-01 | 7,46E+01 |
| GOTERM_BI | GO:0005975~carbohydrate metabolic process | 11 | 1,55 | 1,35E-01 | AMY1, HHIPL2, CHIL1, HKDC1, HEXB, PGD, G6PDX, PGM1, FUCA2, NEU3, PYGB | 588 | 206 | 18082 | 1,64 | 1,00E+00 | 9,87E-01 | 9,20E+01 |
| INTERPRO | IPR013781:Glycoside hydrolase, catalytic domain | 4 | 0,56 | 1,36E-01 | AMY1, CHIL1, HEXB, FUCA2 | 645 | 41 | 20594 | 3,11 | 1,00E+00 | 9,72E-01 | 9,03E+01 |
| UP_KEYWO | Glycosidase | 5 | 0,71 | 2,49E-01 | AMY1, DNPH1, HEXB, FUCA2, NEU3 | 678 | 85 | 22680 | 1,97 | 1,00E+00 | 7,69E-01 | 9,79E+01 |
| GOTERM_M | GO:0016798~hydrolase activity, acting on glycosyl bonds | 5 | 0,71 | 2,55E-01 | AMY1, DNPH1, HEXB, FUCA2, NEU3 | 575 | 78 | 17446 | 1,94 | 1,00E+00 | 9,77E-01 | 9,89E+01 |
| INTERPRO | IPR017853:Glycoside hydrolase, superfamily | 4 | 0,56 | 2,64E-01 | AMY1, CHIL1, HEXB, FUCA2 | 645 | 57 | 20594 | 2,24 | 1,00E+00 | 9,91E-01 | 9,92E+01 |
| Annotation | Enrichment Score: 0.7512220511220798 | | | | | | | | | | | |
| Category | Term | Count | % | PValue | Genes | List To | Pop Hit | Pop Tota | Fold Enr | Bonferroni | Benjamini | FDR |
| INTERPRO | IPR001849:Pleckstrin homology domain | 13 | 1,83 | 1,23E-01 | DAB2IP, OSBPL3, PLEK, STAP1, FERMT1, ARHGEF16, ARHGEF9, VEPH1, SNTB1, IPCEF1, FAM129B, PLEKHA1, SPTB | 645 | 262 | 20594 | 1,58 | 1,00E+00 | 9,63E-01 | 8,77E+01 |
| INTERPRO | IPR011993:Pleckstrin homology-like domain | 18 | 2,54 | 1,46E-01 | DAB2IP, OSBPL3, STAP1, PLEK, MYO7B, FERMT1, ARHGEF16, ARHGEF9, ECT2, VEPH1, MTMR11, RGS12, SNTB1, IPCEF1, FAM129B, NBEAL1, SPTB, PLEKHA1 | 645 | 409 | 20594 | 1,41 | 1,00E+00 | 9,67E-01 | 9,19E+01 |
| UP_SEQ_FE | domain:PH | 10 | 1,41 | 1,77E-01 | VEPH1, DAB2IP, OSBPL3, STAP1, FERMT1, ARHGEF16, IPCEF1, FAM129B, ARHGEF9, ECT2 | 544 | 208 | 18012 | 1,59 | 1,00E+00 | 9,93E-01 | 9,61E+01 |
| SMART | SM00233:PH | 12 | 1,69 | 3,12E-01 | VEPH1, DAB2IP, OSBPL3, STAP1, PLEK, FERMT1, ARHGEF16, SNTB1, IPCEF1, ARHGEF9, PLEKHA1, SPTB | 379 | 253 | 10425 | 1,30 | 1,00E+00 | 9,78E-01 | 9,92E+01 |
| Annotation | Enrichment Score: 0.7268061413735906 | | | | | | | | | | | |
| Category | Term | Count | % | PValue | Genes | List To | Pop Hit | Pop Tota | Fold Enr | Bonferroni | Benjamini | FDR |
| INTERPRO | IPR003879:Butyrophylin-like | 6 | 0,85 | 3,25E-02 | TRIM30A, TRIM68, TRIM30D, BTNL6, TRIM21, BTNL4 | 645 | 57 | 20594 | 3,36 | 1,00E+00 | 8,61E-01 | 4,09E+01 |
| SMART | SM00336:BBOX | 6 | 0,85 | 6,96E-02 | TRIM56, TRIM30A, TRIM68, TRIM30D, TRIM24, TRIM21 | 379 | 61 | 10425 | 2,71 | 1,00E+00 | 9,03E-01 | 6,02E+01 |
| INTERPRO | IPR000315:Zinc finger, B-box | 6 | 0,85 | 7,12E-02 | TRIM56, TRIM30A, TRIM68, TRIM30D, TRIM24, TRIM21 | 645 | 71 | 20594 | 2,70 | 1,00E+00 | 9,36E-01 | 6,92E+01 |
| INTERPRO | IPR003877:SPla/RYanodine receptor SPRY | 6 | 0,85 | 9,33E-02 | TRIM30A, TRIM68, TRIM30D, BTNL6, TRIM21, BTNL4 | 645 | 77 | 20594 | 2,49 | 1,00E+00 | 9,42E-01 | 7,90E+01 |
| INTERPRO | IPR001870:B30.2/SPRY domain | 6 | 0,85 | 1,18E-01 | TRIM30A, TRIM68, TRIM30D, BTNL6, TRIM21, BTNL4 | 645 | 83 | 20594 | 2,31 | 1,00E+00 | 9,64E-01 | 8,66E+01 |

| | | | | | | | | | | | | |
|---|---|---|---|---|---|---|---|---|---|---|---|---|
| SMART | SM00449:SPRY | 6 | 0,85 | 1,36E-01 | TRIM30A, TRIM68, TRIM30D, BTNL6, TRIM21, BTNL4 | 379 | 75 | 10425 | 2,20 | 1,00E+00 | 9,05E-01 | 8,45E+01 |
| INTERPRO | IPR017907:Zinc finger, RING-type, conserved site | 9 | 1,27 | 1,40E-01 | TRIM56, TRIM30A, RNF17, RNF141, TRIM68, TRIM30D, NHLRC1, TRIM24, TRIM21 | 645 | 163 | 20594 | 1,76 | 1,00E+00 | 9,66E-01 | 9,09E+01 |
| UP_SEQ_FE/ | zinc finger region:RING-type | 9 | 1,27 | 2,00E-01 | FANCL, TRIM56, TRIM30A, RNF17, RNF141, TRIM68, NHLRC1, TRIM24, TRIM21 | 544 | 186 | 18012 | 1,60 | 1,00E+00 | 9,94E-01 | 9,76E+01 |
| INTERPRO | IPR013320:Concanavalin A-like lectin/glucanase, subgroup | 10 | 1,41 | 2,20E-01 | TRIM30A, NPTX1, COL14A1, TRIM68, LGALS1, TRIM30D, BTNL6, TRIM21, BTNL4, NBEAL1 | 645 | 211 | 20594 | 1,51 | 1,00E+00 | 9,85E-01 | 9,81E+01 |
| GOTERM_BI | GO:0051865~protein autoubiquitination | 4 | 0,56 | 2,38E-01 | TRIM30A, RNF141, TRIM68, TRIM21 | 588 | 52 | 18082 | 2,37 | 1,00E+00 | 9,96E-01 | 9,91E+01 |
| UP_SEQ_FE/ | zinc finger region:B box-type | 3 | 0,42 | 3,63E-01 | TRIM30A, TRIM68, TRIM21 | 544 | 42 | 18012 | 2,37 | 1,00E+00 | 1,00E+00 | 9,99E+01 |
| UP_SEQ_FE/ | domain:B30.2/SPRY | 3 | 0,42 | 5,35E-01 | TRIM30A, TRIM68, TRIM21 | 544 | 59 | 18012 | 1,68 | 1,00E+00 | 1,00E+00 | 1,00E+02 |
| INTERPRO | IPR013083:Zinc finger, RING/FYVE/PHD-type | 15 | 2,12 | 5,40E-01 | DTX4, DPF3, TRIM24, NHLRC1, TRIM21, SP140, MARCH1, FANCL, TRIM56, TRIM30A, G2E3, RNF141, TRIM68, TRIM30D, SYTL5 | 645 | 445 | 20594 | 1,08 | 1,00E+00 | 1,00E+00 | 1,00E+02 |
| INTERPRO | IPR001841:Zinc finger, RING-type | 10 | 1,41 | 5,49E-01 | DTX4, TRIM56, TRIM30A, RNF17, RNF141, TRIM68, TRIM30D, NHLRC1, TRIM24, TRIM21 | 645 | 286 | 20594 | 1,12 | 1,00E+00 | 1,00E+00 | 1,00E+02 |
| SMART | SM00184:RING | 9 | 1,27 | 6,18E-01 | DTX4, TRIM56, TRIM30A, RNF141, TRIM68, TRIM30D, NHLRC1, TRIM24, TRIM21 | 379 | 234 | 10425 | 1,06 | 1,00E+00 | 9,97E-01 | 1,00E+02 |
| Annotation | Enrichment Score: 0.6762549059371291 | | | | | | | | | | | |
| Category | Term | Count | % | PValue | Genes | List To | Pop Hit | Pop Tot | Fold En | Bonferroni | Benjamini | FDR |
| GOTERM_BI | GO:0042384~cilium assembly | 9 | 1,27 | 5,93E-02 | RABL2, BBS4, CEP83, PKHD1, TTC30B, TTC30A2, TMEM237, EHD1, DNAH5 | 588 | 129 | 18082 | 2,15 | 1,00E+00 | 9,39E-01 | 6,57E+01 |
| GOTERM_C( | GO:0005929~cilium | 13 | 1,83 | 1,25E-01 | GAPDHS, BBS4, SMO, ATP2B2, TTC30B, TTC30A2, DRC1, TMEM237, PTCH1, DNAIC1, EHD1, DNAH5, NPHP1 | 633 | 257 | 19662 | 1,57 | 1,00E+00 | 6,94E-01 | 8,45E+01 |
| GOTERM_C( | GO:0030992~intraciliary transport particle B | 3 | 0,42 | 1,45E-01 | RABL2, TTC30B, TTC30A2 | 633 | 21 | 19662 | 4,44 | 1,00E+00 | 7,06E-01 | 8,88E+01 |
| GOTERM_BI | GO:0060271~cilium morphogenesis | 9 | 1,27 | 1,89E-01 | RABL2, BBS4, CEP83, PKHD1, TTC30B, TTC30A2, TMEM237, EHD1, DNAH5 | 588 | 170 | 18082 | 1,63 | 1,00E+00 | 9,94E-01 | 9,74E+01 |
| UP_SEQ_FE/ | repeat:TPR 7 | 4 | 0,56 | 1,99E-01 | IFIT2, BBS4, TTC30B, TTC30A2 | 544 | 51 | 18012 | 2,60 | 1,00E+00 | 9,95E-01 | 9,75E+01 |
| UP_KEYWO | Cilium | 9 | 1,27 | 2,13E-01 | BBS4, TTC30B, TTC30A2, DRC1, TMEM237, DNAIC1, EHD1, DNAH5, NPHP1 | 678 | 191 | 22680 | 1,58 | 1,00E+00 | 7,41E-01 | 9,61E+01 |
| GOTERM_BI | GO:0030030~cell projection organization | 8 | 1,13 | 2,20E-01 | BBS4, CEP83, PLEK, TTC30B, TTC30A2, TMEM237, EHD1, NPHP1 | 588 | 151 | 18082 | 1,63 | 1,00E+00 | 9,96E-01 | 9,87E+01 |
| UP_KEYWO | Cell projection | 25 | 3,53 | 2,21E-01 | BBS4, MYO7B, FERMT1, TMEM237, PALMD, DNAH5, ALCAM, RGS12, ROBO1, TTC30B, PAK1, EHD1, KCNE3, CABLES1, NPHP1, DAB2IP, OSBPL3, DRC1, VIL1, TTC30A2, DNAIC1, SMO, IQCG, ATXN1L, MYH10 | 678 | 678 | 22680 | 1,23 | 1,00E+00 | 7,50E-01 | 9,66E+01 |

| | | | | | | | | | | | | |
|---|---|---|---|---|---|---|---|---|---|---|---|---|
| GOTERM_C | GO:0035869~ciliary transition zone | 3 | 0,42 | 2,27E-01 | BBS4, TMEM237, NPHP1 | 633 | 28 | 19662 | 3,33 | 1,00E+00 | 8,45E-01 | 9,72E+01 |
| UP_KEYWO | Cilium biogenesis/degradation | 7 | 0,99 | 2,41E-01 | BBS4, CEP83, TTC30B, TTC30A2, TMEM237, EHD1, NPHP1 | 678 | 140 | 22680 | 1,67 | 1,00E+00 | 7,62E-01 | 9,76E+01 |
| UP_SEQ_FE | repeat:TPR 6 | 4 | 0,56 | 2,46E-01 | IFIT2, BBS4, TTC30B, TTC30A2 | 544 | 57 | 18012 | 2,32 | 1,00E+00 | 9,98E-01 | 9,91E+01 |
| UP_SEQ_FE | repeat:TPR 8 | 3 | 0,42 | 3,30E-01 | BBS4, TTC30B, TTC30A2 | 544 | 39 | 18012 | 2,55 | 1,00E+00 | 9,99E-01 | 9,99E+01 |
| GOTERM_C | GO:0036064~ciliary basal body | 3 | 0,42 | 8,69E-01 | BBS4, PKHD1, TTC30B | 633 | 109 | 19662 | 0,85 | 1,00E+00 | 9,99E-01 | 1,00E+02 |
| Annotation | Enrichment Score: 0.6592692957575353 | | | | | | | | | | | |
| Category | Term | Count | % | PValue | Genes | List To | Pop Hit | Pop Tot | Fold En | Bonferroni | Benjamini | FDR |
| UP_SEQ_FE | domain:CH 1 | 3 | 0,42 | 1,31E-01 | DMD, PLS1, SPTB | 544 | 21 | 18012 | 4,73 | 1,00E+00 | 9,84E-01 | 9,04E+01 |
| UP_SEQ_FE | domain:CH 2 | 3 | 0,42 | 1,31E-01 | DMD, PLS1, SPTB | 544 | 21 | 18012 | 4,73 | 1,00E+00 | 9,84E-01 | 9,04E+01 |
| INTERPRO | IPR001589:Actinin-type, actin-binding, conserved site | 3 | 0,42 | 1,61E-01 | DMD, PLS1, SPTB | 645 | 23 | 20594 | 4,16 | 1,00E+00 | 9,72E-01 | 9,39E+01 |
| INTERPRO | IPR001715:Calponin homology domain | 4 | 0,56 | 4,26E-01 | DMD, SMTNL2, PLS1, SPTB | 645 | 76 | 20594 | 1,68 | 1,00E+00 | 9,99E-01 | 1,00E+02 |
| SMART | SM00033:CH | 4 | 0,56 | 4,31E-01 | DMD, SMTNL2, PLS1, SPTB | 379 | 66 | 10425 | 1,67 | 1,00E+00 | 9,91E-01 | 9,99E+01 |
| Annotation | Enrichment Score: 0.6477476319264411 | | | | | | | | | | | |
| Category | Term | Count | % | PValue | Genes | List To | Pop Hit | Pop Tot | Fold En | Bonferroni | Benjamini | FDR |
| GOTERM_B | GO:0045356~positive regulation of interferon-alpha biosynthetic process | 3 | 0,42 | 6,04E-03 | TLR3, TLR7, TLR8 | 588 | 4 | 18082 | 23,06 | 1,00E+00 | 6,63E-01 | 1,01E+01 |
| GOTERM_B | GO:0045078~positive regulation of interferon-gamma biosynthetic process | 4 | 0,56 | 7,63E-03 | CD276, TLR3, TLR7, TLR8 | 588 | 13 | 18082 | 9,46 | 1,00E+00 | 7,21E-01 | 1,25E+01 |
| GOTERM_B | GO:0002224~toll-like receptor signaling pathway | 4 | 0,56 | 1,65E-02 | RPS6KA3, TLR3, TLR7, TLR8 | 588 | 17 | 18082 | 7,24 | 1,00E+00 | 8,15E-01 | 2,52E+01 |
| GOTERM_B | GO:0045359~positive regulation of interferon-beta biosynthetic process | 3 | 0,42 | 1,98E-02 | TLR3, TLR7, TLR8 | 588 | 7 | 18082 | 13,18 | 1,00E+00 | 8,46E-01 | 2,96E+01 |
| UP_SEQ_FE | repeat:LRR 22 | 3 | 0,42 | 3,48E-02 | TLR3, TLR7, TLR8 | 544 | 10 | 18012 | 9,93 | 1,00E+00 | 9,68E-01 | 4,46E+01 |
| UP_SEQ_FE | repeat:LRR 21 | 3 | 0,42 | 7,35E-02 | TLR3, TLR7, TLR8 | 544 | 15 | 18012 | 6,62 | 1,00E+00 | 9,69E-01 | 7,20E+01 |

| | | | | | | | | | | | | |
|---|---|---|---|---|---|---|---|---|---|---|---|---|
| UP_SEQ_FE/ | repeat:LRR 12 | 6 | 0,85 | 9,00E-02 | LUM, TLR3, DCN, TLR7, TLR8, PRELP | 544 | 79 | 18012 | 2,51 | 1,00E+00 | 9,72E-01 | 7,93E+01 |
| UP_KEYWOI | Inflammatory response | 9 | 1,27 | 1,10E-01 | DAB2IP, CCL2, CHIL1, NAIP2, TLR3, CLEC7A, TLR7, TLR8, CXCL10 | 678 | 161 | 22680 | 1,87 | 1,00E+00 | 6,21E-01 | 7,94E+01 |
| UP_SEQ_FE/ | repeat:LRR 20 | 3 | 0,42 | 1,11E-01 | TLR3, TLR7, TLR8 | 544 | 19 | 18012 | 5,23 | 1,00E+00 | 9,81E-01 | 8,59E+01 |
| GOTERM_BI | GO:0001774~microglial cell activation | 3 | 0,42 | 1,14E-01 | TLR3, TLR7, TLR8 | 588 | 18 | 18082 | 5,13 | 1,00E+00 | 9,83E-01 | 8,81E+01 |
| UP_SEQ_FE/ | repeat:LRR 11 | 6 | 0,85 | 1,27E-01 | LUM, TLR3, DCN, TLR7, TLR8, PRELP | 544 | 88 | 18012 | 2,26 | 1,00E+00 | 9,84E-01 | 8,96E+01 |
| UP_SEQ_FE/ | repeat:LRR 19 | 3 | 0,42 | 1,41E-01 | TLR3, TLR7, TLR8 | 544 | 22 | 18012 | 4,52 | 1,00E+00 | 9,86E-01 | 9,21E+01 |
| UP_SEQ_FE/ | domain:TIR | 3 | 0,42 | 1,73E-01 | TLR3, TLR7, TLR8 | 544 | 25 | 18012 | 3,97 | 1,00E+00 | 9,93E-01 | 9,58E+01 |
| UP_SEQ_FE/ | repeat:LRR 18 | 3 | 0,42 | 1,84E-01 | TLR3, TLR7, TLR8 | 544 | 26 | 18012 | 3,82 | 1,00E+00 | 9,94E-01 | 9,66E+01 |
| INTERPRO | IPR000157:Toll/interleukin-1 receptor homology (TIR) domain | 3 | 0,42 | 1,95E-01 | TLR3, TLR7, TLR8 | 645 | 26 | 20594 | 3,68 | 1,00E+00 | 9,78E-01 | 9,68E+01 |
| GOTERM_M | GO:0003725~double-stranded RNA binding | 5 | 0,71 | 1,99E-01 | OASL1, TLR3, OAS1A, TLR7, TLR8 | 575 | 70 | 17446 | 2,17 | 1,00E+00 | 9,70E-01 | 9,65E+01 |
| UP_SEQ_FE/ | repeat:LRR 10 | 6 | 0,85 | 2,00E-01 | LUM, TLR3, DCN, TLR7, TLR8, PRELP | 544 | 103 | 18012 | 1,93 | 1,00E+00 | 9,94E-01 | 9,76E+01 |
| SMART | SM00255:TIR | 3 | 0,42 | 2,02E-01 | TLR3, TLR7, TLR8 | 379 | 23 | 10425 | 3,59 | 1,00E+00 | 9,59E-01 | 9,44E+01 |
| UP_SEQ_FE/ | repeat:LRR 17 | 3 | 0,42 | 2,85E-01 | TLR3, TLR7, TLR8 | 544 | 35 | 18012 | 2,84 | 1,00E+00 | 9,98E-01 | 9,96E+01 |
| INTERPRO | IPR003591:Leucine-rich repeat, typical subtype | 8 | 1,13 | 3,08E-01 | AMIGO2, LUM, LGI4, TLR3, DCN, TLR7, TLR8, PRELP | 645 | 175 | 20594 | 1,46 | 1,00E+00 | 9,96E-01 | 9,97E+01 |
| UP_SEQ_FE/ | repeat:LRR 9 | 6 | 0,85 | 3,13E-01 | LUM, TLR3, DCN, TLR7, TLR8, PRELP | 544 | 123 | 18012 | 1,62 | 1,00E+00 | 9,99E-01 | 9,98E+01 |
| UP_SEQ_FE/ | repeat:LRR 16 | 3 | 0,42 | 3,52E-01 | TLR3, TLR7, TLR8 | 544 | 41 | 18012 | 2,42 | 1,00E+00 | 9,99E-01 | 9,99E+01 |
| KEGG_PATH | mmu04620:Toll-like receptor signaling pathway | 5 | 0,71 | 3,61E-01 | TLR3, TLR7, TLR8, SPP1, CXCL10 | 233 | 101 | 7720 | 1,64 | 1,00E+00 | 9,02E-01 | 9,97E+01 |
| UP_SEQ_FE/ | repeat:LRR 8 | 6 | 0,85 | 3,79E-01 | LUM, TLR3, DCN, TLR7, TLR8, PRELP | 544 | 134 | 18012 | 1,48 | 1,00E+00 | 1,00E+00 | 1,00E+02 |
| INTERPRO | IPR000483:Cysteine-rich flanking region, C-terminal | 4 | 0,56 | 4,26E-01 | AMIGO2, LGI4, TLR3, TLR7 | 645 | 76 | 20594 | 1,68 | 1,00E+00 | 9,99E-01 | 1,00E+02 |
| UP_SEQ_FE/ | repeat:LRR 15 | 3 | 0,42 | 4,27E-01 | TLR3, TLR7, TLR8 | 544 | 48 | 18012 | 2,07 | 1,00E+00 | 1,00E+00 | 1,00E+02 |
| SMART | SM00369:LRR_TYP | 8 | 1,13 | 4,50E-01 | AMIGO2, LUM, LGI4, TLR3, DCN, TLR7, TLR8, PRELP | 379 | 175 | 10425 | 1,26 | 1,00E+00 | 9,89E-01 | 1,00E+02 |
| GOTERM_BI | GO:0071260~cellular response to mechanical stimulus | 4 | 0,56 | 4,67E-01 | TLR3, ANKRD1, TLR7, TLR8 | 588 | 78 | 18082 | 1,58 | 1,00E+00 | 1,00E+00 | 1,00E+02 |
| INTERPRO | IPR000372:Leucine-rich repeat-containing N-terminal | 3 | 0,42 | 4,97E-01 | LUM, DCN, PRELP | 645 | 53 | 20594 | 1,81 | 1,00E+00 | 1,00E+00 | 1,00E+02 |
| UP_SEQ_FE/ | repeat:LRR 14 | 3 | 0,42 | 4,97E-01 | TLR3, TLR7, TLR8 | 544 | 55 | 18012 | 1,81 | 1,00E+00 | 1,00E+00 | 1,00E+02 |
| SMART | SM00082:LRRCT | 4 | 0,56 | 5,15E-01 | AMIGO2, LGI4, TLR3, TLR7 | 379 | 75 | 10425 | 1,47 | 1,00E+00 | 9,91E-01 | 1,00E+02 |
| UP_SEQ_FE/ | repeat:LRR 6 | 7 | 0,99 | 5,17E-01 | AMIGO2, LUM, TLR3, DCN, TLR7, TLR8, PRELP | 544 | 191 | 18012 | 1,21 | 1,00E+00 | 1,00E+00 | 1,00E+02 |

| | | | | | | | | | | | | |
|---|---|---|---|---|---|---|---|---|---|---|---|---|
| UP_SEQ_FE/ | repeat:LRR 7 | 6 | 0,85 | 5,20E-01 | LUM, TLR3, DCN, TLR7, TLR8, PRELP | 544 | 158 | 18012 | 1,26 | 1,00E+00 | 1,00E+00 | 1,00E+02 |
| SMART | SM00013:LRRNT | 3 | 0,42 | 5,78E-01 | LUM, DCN, PRELP | 379 | 53 | 10425 | 1,56 | 1,00E+00 | 9,96E-01 | 1,00E+02 |
| UP_SEQ_FE/ | repeat:LRR 13 | 3 | 0,42 | 5,87E-01 | TLR3, TLR7, TLR8 | 544 | 65 | 18012 | 1,53 | 1,00E+00 | 1,00E+00 | 1,00E+02 |
| UP_SEQ_FE/ | repeat:LRR 5 | 7 | 0,99 | 6,10E-01 | AMIGO2, LUM, TLR3, DCN, TLR7, TLR8, PRELP | 544 | 210 | 18012 | 1,10 | 1,00E+00 | 1,00E+00 | 1,00E+02 |
| UP_SEQ_FE/ | repeat:LRR 3 | 8 | 1,13 | 6,44E-01 | AMIGO2, LUM, LGI4, TLR3, DCN, TLR7, TLR8, PRELP | 544 | 253 | 18012 | 1,05 | 1,00E+00 | 1,00E+00 | 1,00E+02 |
| UP_SEQ_FE/ | repeat:LRR 4 | 7 | 0,99 | 6,88E-01 | AMIGO2, LUM, TLR3, DCN, TLR7, TLR8, PRELP | 544 | 228 | 18012 | 1,02 | 1,00E+00 | 1,00E+00 | 1,00E+02 |
| INTERPRO | IPR001611:Leucine-rich repeat | 8 | 1,13 | 7,04E-01 | AMIGO2, LUM, LGI4, TLR3, DCN, TLR7, TLR8, PRELP | 645 | 259 | 20594 | 0,99 | 1,00E+00 | 1,00E+00 | 1,00E+02 |
| UP_KEYWO | Leucine-rich repeat | 8 | 1,13 | 7,17E-01 | AMIGO2, LUM, LGI4, TLR3, DCN, TLR7, TLR8, PRELP | 678 | 275 | 22680 | 0,97 | 1,00E+00 | 9,66E-01 | 1,00E+02 |
| UP_SEQ_FE/ | repeat:LRR 2 | 8 | 1,13 | 7,23E-01 | AMIGO2, LUM, LGI4, TLR3, DCN, TLR7, TLR8, PRELP | 544 | 274 | 18012 | 0,97 | 1,00E+00 | 1,00E+00 | 1,00E+02 |
| UP_SEQ_FE/ | repeat:LRR 1 | 8 | 1,13 | 7,23E-01 | AMIGO2, LUM, LGI4, TLR3, DCN, TLR7, TLR8, PRELP | 544 | 274 | 18012 | 0,97 | 1,00E+00 | 1,00E+00 | 1,00E+02 |
| UP_SEQ_FE/ | compositionally biased region:Cys-rich | 4 | 0,56 | 7,78E-01 | SLC17A8, LUM, DCN, PRELP | 544 | 135 | 18012 | 0,98 | 1,00E+00 | 1,00E+00 | 1,00E+02 |
| Annotation | Enrichment Score: 0.6266150377990402 | | | | | | | | | | | |
| Category | Term | Count | % | PValue | Genes | List To | Pop Hit | Pop Tot | Fold Enr | Bonferroni | Benjamini | FDR |
| UP_SEQ_FE/ | domain:UPAR/Ly6 | 3 | 0,42 | 1,95E-01 | LY6A, LY6D, LY6E | 544 | 27 | 18012 | 3,68 | 1,00E+00 | 9,95E-01 | 9,73E+01 |
| SMART | SM00134:LU | 3 | 0,42 | 2,02E-01 | LY6A, LY6D, LY6E | 379 | 23 | 10425 | 3,59 | 1,00E+00 | 9,59E-01 | 9,44E+01 |
| INTERPRO | IPR016054:Ly-6 antigen / uPA receptor -like | 3 | 0,42 | 3,34E-01 | LY6A, LY6D, LY6E | 645 | 38 | 20594 | 2,52 | 1,00E+00 | 9,97E-01 | 9,98E+01 |
| Annotation | Enrichment Score: 0.6244678836590328 | | | | | | | | | | | |
| Category | Term | Count | % | PValue | Genes | List To | Pop Hit | Pop Tot | Fold Enr | Bonferroni | Benjamini | FDR |
| GOTERM_C | GO:0005903~brush border | 6 | 0,85 | 1,02E-01 | CGN, MYO7B, VIL1, PLS1, LRP2, MYH10 | 633 | 77 | 19662 | 2,42 | 1,00E+00 | 6,47E-01 | 7,76E+01 |
| GOTERM_M | GO:0003774~motor activity | 5 | 0,71 | 2,62E-01 | CGN, MYO7B, DNAIC1, DNAH5, MYH10 | 575 | 79 | 17446 | 1,92 | 1,00E+00 | 9,78E-01 | 9,90E+01 |
| GOTERM_C | GO:0016459~myosin complex | 3 | 0,42 | 5,02E-01 | CGN, MYO7B, MYH10 | 633 | 52 | 19662 | 1,79 | 1,00E+00 | 9,57E-01 | 1,00E+02 |
| Annotation | Enrichment Score: 0.6060748789381049 | | | | | | | | | | | |
| Category | Term | Count | % | PValue | Genes | List To | Pop Hit | Pop Tot | Fold Enr | Bonferroni | Benjamini | FDR |
| INTERPRO | IPR011527:ABC transporter, transmembrane domain, type 1 | 4 | 0,56 | 5,11E-02 | ABCB1A, TAP1, ABCD2, ABCC12 | 645 | 27 | 20594 | 4,73 | 1,00E+00 | 9,05E-01 | 5,66E+01 |

| | | | | | | | | | | | | |
|---|---|---|---|---|---|---|---|---|---|---|---|---|
| KEGG_PATH | mmu02010:ABC transporters | 4 | 0,56 | 1,59E-01 | ABCB1A, TAP1, ABCD2, ABCC12 | 233 | 46 | 7720 | 2,88 | 1,00E+00 | 8,17E-01 | 8,94E+01 |
| INTERPRO | IPR017871:ABC transporter, conserved site | 4 | 0,56 | 1,89E-01 | ABCB1A, TAP1, ABCD2, ABCC12 | 645 | 48 | 20594 | 2,66 | 1,00E+00 | 9,80E-01 | 9,65E+01 |
| GOTERM_M | GO:0042626~ATPase activity, coupled to transmembrane movement of substances | 4 | 0,56 | 2,18E-01 | ABCB1A, TAP1, ABCD2, ABCC12 | 575 | 49 | 17446 | 2,48 | 1,00E+00 | 9,70E-01 | 9,76E+01 |
| INTERPRO | IPR003439:ABC transporter-like | 4 | 0,56 | 2,30E-01 | ABCB1A, TAP1, ABCD2, ABCC12 | 645 | 53 | 20594 | 2,41 | 1,00E+00 | 9,84E-01 | 9,85E+01 |
| INTERPRO | IPR003593:AAA+ ATPase domain | 6 | 0,85 | 4,70E-01 | RAD51B, ABCB1A, TAP1, ABCD2, ABCC12, DNAH5 | 645 | 144 | 20594 | 1,33 | 1,00E+00 | 9,99E-01 | 1,00E+02 |
| SMART | SM00382:AAA | 6 | 0,85 | 6,03E-01 | RAD51B, ABCB1A, TAP1, ABCD2, ABCC12, DNAH5 | 379 | 144 | 10425 | 1,15 | 1,00E+00 | 9,96E-01 | 1,00E+02 |
| GOTERM_M | GO:0016887~ATPase activity | 7 | 0,99 | 6,48E-01 | ABCB1A, TAP1, DQX1, ABCD2, ABCC12, DNAH5, PMS1 | 575 | 200 | 17446 | 1,06 | 1,00E+00 | 9,99E-01 | 1,00E+02 |
| Annotation | Enrichment Score: 0.584517604722037 | | | | | | | | | | | |
| Category | Term | Count | % | PValue | Genes | List To | Pop Hit | Pop Tota | Fold Enr | Bonferroni | Benjamini | FDR |
| KEGG_PATH | mmu04360:Axon guidance | 8 | 1,13 | 9,33E-02 | PAK6, UNC5B, PLXNB1, ROBO1, SEMA3D, PPP3CC, PAK1, SEMA4A | 233 | 129 | 7720 | 2,05 | 1,00E+00 | 7,75E-01 | 7,18E+01 |
| UP_SEQ_FE | domain:Sema | 3 | 0,42 | 2,06E-01 | PLXNB1, SEMA3D, SEMA4A | 544 | 28 | 18012 | 3,55 | 1,00E+00 | 9,94E-01 | 9,79E+01 |
| INTERPRO | IPR001627:Semaphorin/CD100 antigen | 3 | 0,42 | 2,53E-01 | PLXNB1, SEMA3D, SEMA4A | 645 | 31 | 20594 | 3,09 | 1,00E+00 | 9,89E-01 | 9,90E+01 |
| GOTERM_BI | GO:0071526~semaphorin-plexin signaling pathway | 3 | 0,42 | 3,03E-01 | PLXNB1, SEMA3D, SEMA4A | 588 | 34 | 18082 | 2,71 | 1,00E+00 | 9,98E-01 | 9,98E+01 |
| SMART | SM00630:Sema | 3 | 0,42 | 3,11E-01 | PLXNB1, SEMA3D, SEMA4A | 379 | 31 | 10425 | 2,66 | 1,00E+00 | 9,82E-01 | 9,91E+01 |
| INTERPRO | IPR016201:Plexin-like fold | 3 | 0,42 | 4,13E-01 | PLXNB1, SEMA3D, SEMA4A | 645 | 45 | 20594 | 2,13 | 1,00E+00 | 9,99E-01 | 1,00E+02 |
| SMART | SM00423:PSI | 3 | 0,42 | 4,29E-01 | PLXNB1, SEMA3D, SEMA4A | 379 | 40 | 10425 | 2,06 | 1,00E+00 | 9,92E-01 | 9,99E+01 |
| Annotation | Enrichment Score: 0.5838174575724596 | | | | | | | | | | | |
| Category | Term | Count | % | PValue | Genes | List To | Pop Hit | Pop Tota | Fold Enr | Bonferroni | Benjamini | FDR |
| GOTERM_C | GO:0005913~cell-cell adherens junction | 15 | 2,12 | 1,45E-01 | RTN4, DAB2IP, ARHGEF16, CDH1, H1FX, SRC, ANXA2, PAK6, CGN, PPL, TGM1, GM609, FAM129B, EHD1, CLINT1 | 633 | 316 | 19662 | 1,47 | 1,00E+00 | 7,12E-01 | 8,87E+01 |
| GOTERM_M | GO:0098641~cadherin binding involved in cell-cell adhesion | 13 | 1,83 | 2,11E-01 | RTN4, PAK6, DAB2IP, CGN, PPL, ARHGEF16, CDH1, H1FX, FAM129B, EHD1, CLINT1, SRC, ANXA2 | 575 | 279 | 17446 | 1,41 | 1,00E+00 | 9,70E-01 | 9,73E+01 |

| | | | | | | | | | | | | |
|---|---|---|---|---|---|---|---|---|---|---|---|---|
| GOTERM_BI | GO:0098609~cell-cell adhesion | 7 | 0,99 | 5,80E-01 | PAK6, CGN, PPL, ARHGEF16, H1FX, FAM129B, CLINT1 | 588 | 189 | 18082 | 1,14 | 1,00E+00 | 1,00E+00 | 1,00E+02 |
| Annotation | Enrichment Score: 0.5651323351504497 | | | | | | | | | | | |
| Category | Term | Count | % | PValue | Genes | List To | Pop Hit | Pop Tota | Fold Enr | Bonferroni | Benjamini | FDR |
| INTERPRO | IPR002172:Low-density lipoprotein (LDL) receptor class A repeat | 5 | 0,71 | 7,06E-02 | TMPRSS2, LRP11, SORL1, LRP2, TMPRSS4 | 645 | 50 | 20594 | 3,19 | 1,00E+00 | 9,40E-01 | 6,89E+01 |
| INTERPRO | IPR023415:Low-density lipoprotein (LDL) receptor class A, conserved site | 4 | 0,56 | 1,51E-01 | TMPRSS2, LRP11, SORL1, LRP2 | 645 | 43 | 20594 | 2,97 | 1,00E+00 | 9,68E-01 | 9,26E+01 |
| SMART | SM00192:LDLa | 4 | 0,56 | 2,42E-01 | TMPRSS2, LRP11, SORL1, LRP2 | 379 | 47 | 10425 | 2,34 | 1,00E+00 | 9,69E-01 | 9,71E+01 |
| UP_KEYWO | Endocytosis | 4 | 0,56 | 6,88E-01 | LRP11, SORL1, LRP2, CLINT1 | 678 | 118 | 22680 | 1,13 | 1,00E+00 | 9,63E-01 | 1,00E+02 |
| GOTERM_BI | GO:0006897~endocytosis | 5 | 0,71 | 8,43E-01 | LRP11, SORL1, LRP2, EHD1, CLINT1 | 588 | 181 | 18082 | 0,85 | 1,00E+00 | 1,00E+00 | 1,00E+02 |
| Annotation | Enrichment Score: 0.5596298584344069 | | | | | | | | | | | |
| Category | Term | Count | % | PValue | Genes | List To | Pop Hit | Pop Tota | Fold Enr | Bonferroni | Benjamini | FDR |
| UP_KEYWO | Ion transport | 25 | 3,53 | 1,16E-01 | STEAP4, FXYD3, SLC13A5, AQP7, FXYD5, ATP2B2, NIPAL1, ATP6V0D2, KCNE3, GRID1, TRPM4, SLC8A1, CLCA1, SLC22A7, SLC10A5, CNGA3, SLC10A2, P2RX5, SLC17A8, ATP6V0E2, CLIC5, SCN4B, SLC5A9, SLC13A3, ATP7B | 678 | 619 | 22680 | 1,35 | 1,00E+00 | 6,16E-01 | 8,13E+01 |
| GOTERM_BI | GO:0006811~ion transport | 23 | 3,24 | 2,64E-01 | TRPM4, STEAP4, SLC8A1, FXYD3, SLC13A5, CLCA1, SLC22A7, SLC10A5, FXYD5, CNGA3, SLC10A2, ATP2B2, SLC17A8, ATP6V0E2, CLIC5, SCN4B, SLC5A9, NIPAL1, SLC13A3, ATP6V0D2, KCNE3, GRID1, ATP7B | 588 | 584 | 18082 | 1,21 | 1,00E+00 | 9,97E-01 | 9,95E+01 |
| UP_KEYWO | Transport | 60 | 8,46 | 3,92E-01 | STEAP4, SLC13A5, AQP8, OBP2A, SORL1, SLC7A9, AQP7, SLC7A4, ATP2B2, NIPAL1, SLC51B, ATP8B5, ATP6V0D2, GRID1, TRPM4, SLC22A27, CLCA1, SLC25A4, SLC22A7, CHP2, PARP11, ABCC12, CNGA3, ECT2, ABCB1A, CLIC5, BBS4, FXYD3, RBP1, CCDC91, NAPB, FXYD5, SLC47A2, MFSD7C, NUP210, TAP1, ABCD2, SLC35F2, SLC35F3, EHD1, KCNE3, SLC8A1, OSBPL3, SPNS3, FABP12, FADS3, SLC10A5, SLC10A2, CD63, P2RX5, SLC17A8, SLC16A5, ATP6V0E2, UCP2, SLC5A9, SCN4B, SLC13A3, SLC15A3, ATP7B, ORM3 | 678 | 1901 | 22680 | 1,06 | 1,00E+00 | 8,51E-01 | 9,99E+01 |

| | | | | | | | | | | | | |
|---|---|---|---|---|---|---|---|---|---|---|---|---|
| GOTERM_BI | GO:0006810~transport | 61 | 8,6 | 4,79E-01 | STEAP4, SLC13A5, AQP8, OBP2A, SORL1, SLC7A9, AQP7, SLC7A4, ATP2B2, NIPAL1, SLC51B, ATP8B5, ATP6V0D2, GRID1, TRPM4, SLC22A27, CLCA1, SLC25A4, SLC22A7, CHP2, PARP11, ABCC12, CNGA3, ECT2, ABCB1A, CLIC5, TMEM184C, BBS4, FXYD3, RBP1, CCDC91, NAPB, FXYD5, SLC47A2, MFSD7C, NUP210, TAP1, ABCD2, SLC35F2, SLC35F3, EHD1, KCNE3, SLC8A1, OSBPL3, SPNS3, FABP12, FADS3, SLC10A5, SLC10A2, CD63, AFP, SLC17A8, SLC16A5, ATP6V0E2, UCP2, SLC5A9, SCN4B, SLC13A3, SLC15A3, ATP7B, ORM3 | 588 | 1822 | 18082 | 1,03 | 1,00E+00 | 1,00E+00 | 1,00E+02 |
| Annotation | Enrichment Score: 0.557412564784264 | | | | | | | | | | | |
| Category | Term | Count | % | PValue | Genes | List To | Pop Hit | Pop Tota | Fold Enr | Bonferroni | Benjamini | FDR |
| INTERPRO | IPR008197:Whey acidic protein-type 4-disulphide core | 4 | 0,56 | 3,01E-02 | WFDC12, WFDC15B, SLPI, WFDC2 | 645 | 22 | 20594 | 5,81 | 1,00E+00 | 8,73E-01 | 3,86E+01 |
| UP_KEYWO | Antimicrobial | 5 | 0,71 | 4,77E-01 | WFDC12, CHIL1, WFDC15B, SLPI, DEFB1 | 678 | 119 | 22680 | 1,41 | 1,00E+00 | 8,94E-01 | 1,00E+02 |
| UP_KEYWO | Antibiotic | 4 | 0,56 | 5,50E-01 | WFDC12, WFDC15B, SLPI, DEFB1 | 678 | 96 | 22680 | 1,39 | 1,00E+00 | 9,20E-01 | 1,00E+02 |
| GOTERM_BI | GO:0042742~defense response to bacterium | 6 | 0,85 | 7,47E-01 | WFDC12, NAIP2, WFDC15B, SLPI, SPON2, DEFB1 | 588 | 191 | 18082 | 0,97 | 1,00E+00 | 1,00E+00 | 1,00E+02 |
| Annotation | Enrichment Score: 0.52085537003882 | | | | | | | | | | | |
| Category | Term | Count | % | PValue | Genes | List To | Pop Hit | Pop Tota | Fold Enr | Bonferroni | Benjamini | FDR |
| GOTERM_BI | GO:0030308~negative regulation of cell growth | 8 | 1,13 | 1,13E-01 | RTN4, INHBA, BMP10, CDKN1A, DAB2IP, CDKN2C, CRYAB, ZC3H12D | 588 | 125 | 18082 | 1,97 | 1,00E+00 | 9,83E-01 | 8,76E+01 |
| GOTERM_BI | GO:0000082~G1/S transition of mitotic cell cycle | 4 | 0,56 | 3,27E-01 | INHBA, CDKN1A, CDKN2C, CABLES1 | 588 | 62 | 18082 | 1,98 | 1,00E+00 | 9,99E-01 | 9,99E+01 |
| GOTERM_BI | GO:0007050~cell cycle arrest | 3 | 0,42 | 7,44E-01 | INHBA, CDKN1A, CDKN2C | 588 | 81 | 18082 | 1,14 | 1,00E+00 | 1,00E+00 | 1,00E+02 |
| Annotation | Enrichment Score: 0.5103653968411024 | | | | | | | | | | | |
| Category | Term | Count | % | PValue | Genes | List To | Pop Hit | Pop Tota | Fold Enr | Bonferroni | Benjamini | FDR |
| GOTERM_BI | GO:0060326~cell chemotaxis | 6 | 0,85 | 1,09E-01 | VCAM1, CCL2, CXCL14, ARHGEF16, HGF, CXCL10 | 588 | 78 | 18082 | 2,37 | 1,00E+00 | 9,83E-01 | 8,68E+01 |
| KEGG_PATH | mmu04062:Chemokine signaling pathway | 8 | 1,13 | 3,78E-01 | CCL2, CXCL14, CCR5, CCR2, GRK5, PAK1, SRC, CXCL10 | 233 | 196 | 7720 | 1,35 | 1,00E+00 | 9,06E-01 | 9,98E+01 |
| INTERPRO | IPR001811:Chemokine interleukin-8-like domain | 3 | 0,42 | 4,66E-01 | CCL2, CXCL14, CXCL10 | 645 | 50 | 20594 | 1,92 | 1,00E+00 | 9,99E-01 | 1,00E+02 |

| | | | | | | | | | | | | |
|---|---|---|---|---|---|---|---|---|---|---|---|---|
| GOTERM_M | GO:0008009~chemokine activity | 3 | 0,42 | 4,72E-01 | CCL2, CXCL14, CXCL10 | 575 | 48 | 17446 | 1,90 | 1,00E+00 | 9,97E-01 | 1,00E+02 |
| Annotation | Enrichment Score: 0.49964426952087077 | | | | | | | | | | | |
| Category | Term | Count | % | PValue | Genes | List To | Pop Hit | Pop Tota | Fold Enr | Bonferroni | Benjamini | FDR |
| UP_SEQ_FE | metal ion-binding site:Magnesium | 6 | 0,85 | 2,44E-01 | ATP2B2, DNTT, PGM1, ACLY, ATP8B5, ATP7B | 544 | 111 | 18012 | 1,79 | 1,00E+00 | 9,98E-01 | 9,91E+01 |
| UP_SEQ_FE | active site:4-aspartylphosphate intermediate | 3 | 0,42 | 2,62E-01 | ATP2B2, ATP8B5, ATP7B | 544 | 33 | 18012 | 3,01 | 1,00E+00 | 9,98E-01 | 9,94E+01 |
| INTERPRO | IPR023299:P-type ATPase, cytoplasmic domain N | 3 | 0,42 | 3,23E-01 | ATP2B2, ATP8B5, ATP7B | 645 | 37 | 20594 | 2,59 | 1,00E+00 | 9,97E-01 | 9,98E+01 |
| INTERPRO | IPR001757:Cation-transporting P-type ATPase | 3 | 0,42 | 3,23E-01 | ATP2B2, ATP8B5, ATP7B | 645 | 37 | 20594 | 2,59 | 1,00E+00 | 9,97E-01 | 9,98E+01 |
| INTERPRO | IPR008250:P-type ATPase, A domain | 3 | 0,42 | 3,23E-01 | ATP2B2, ATP8B5, ATP7B | 645 | 37 | 20594 | 2,59 | 1,00E+00 | 9,97E-01 | 9,98E+01 |
| INTERPRO | IPR018303:P-type ATPase, phosphorylation site | 3 | 0,42 | 3,23E-01 | ATP2B2, ATP8B5, ATP7B | 645 | 37 | 20594 | 2,59 | 1,00E+00 | 9,97E-01 | 9,98E+01 |
| INTERPRO | IPR023214:HAD-like domain | 4 | 0,56 | 4,59E-01 | ATP2B2, ATP8B5, PMM1, ATP7B | 645 | 80 | 20594 | 1,60 | 1,00E+00 | 9,99E-01 | 1,00E+02 |
| Annotation | Enrichment Score: 0.487557911276864 | | | | | | | | | | | |
| Category | Term | Count | % | PValue | Genes | List To | Pop Hit | Pop Tota | Fold Enr | Bonferroni | Benjamini | FDR |
| UP_SEQ_FE | repeat:TPR 5 | 5 | 0,71 | 1,27E-01 | IFIT2, BBS4, TTC30B, TTC30A2, TTC29 | 544 | 64 | 18012 | 2,59 | 1,00E+00 | 9,83E-01 | 8,97E+01 |
| UP_SEQ_FE | repeat:TPR 7 | 4 | 0,56 | 1,99E-01 | IFIT2, BBS4, TTC30B, TTC30A2 | 544 | 51 | 18012 | 2,60 | 1,00E+00 | 9,95E-01 | 9,75E+01 |
| UP_KEYWO | Cilium biogenesis/degradation | 7 | 0,99 | 2,41E-01 | BBS4, CEP83, TTC30B, TTC30A2, TMEM237, EHD1, NPHP1 | 678 | 140 | 22680 | 1,67 | 1,00E+00 | 7,62E-01 | 9,76E+01 |
| UP_SEQ_FE | repeat:TPR 6 | 4 | 0,56 | 2,46E-01 | IFIT2, BBS4, TTC30B, TTC30A2 | 544 | 57 | 18012 | 2,32 | 1,00E+00 | 9,98E-01 | 9,91E+01 |
| UP_SEQ_FE | repeat:TPR 4 | 5 | 0,71 | 2,61E-01 | IFIT2, BBS4, TTC30B, TTC30A2, TTC29 | 544 | 86 | 18012 | 1,93 | 1,00E+00 | 9,98E-01 | 9,94E+01 |
| UP_SEQ_FE | repeat:TPR 3 | 6 | 0,85 | 3,25E-01 | IFIT2, BBS4, TTC30B, TTC30A2, TTC29, TTC39A | 544 | 125 | 18012 | 1,59 | 1,00E+00 | 9,99E-01 | 9,99E+01 |
| UP_SEQ_FE | repeat:TPR 8 | 3 | 0,42 | 3,30E-01 | BBS4, TTC30B, TTC30A2 | 544 | 39 | 18012 | 2,55 | 1,00E+00 | 9,99E-01 | 9,99E+01 |
| INTERPRO | IPR019734:Tetratricopeptide repeat | 6 | 0,85 | 3,59E-01 | IFIT2, BBS4, TTC30B, TTC30A2, TTC29, TTC39A | 645 | 126 | 20594 | 1,52 | 1,00E+00 | 9,98E-01 | 9,99E+01 |
| UP_SEQ_FE | repeat:TPR 1 | 6 | 0,85 | 3,97E-01 | IFIT2, BBS4, TTC30B, TTC30A2, TTC29, TTC39A | 544 | 137 | 18012 | 1,45 | 1,00E+00 | 1,00E+00 | 1,00E+02 |
| UP_SEQ_FE | repeat:TPR 2 | 6 | 0,85 | 3,97E-01 | IFIT2, BBS4, TTC30B, TTC30A2, TTC29, TTC39A | 544 | 137 | 18012 | 1,45 | 1,00E+00 | 1,00E+00 | 1,00E+02 |
| UP_KEYWO | TPR repeat | 6 | 0,85 | 4,30E-01 | IFIT2, BBS4, TTC30B, TTC30A2, TTC29, TTC39A | 678 | 144 | 22680 | 1,39 | 1,00E+00 | 8,70E-01 | 1,00E+02 |
| SMART | SM00028:TPR | 6 | 0,85 | 4,34E-01 | IFIT2, BBS4, TTC30B, TTC30A2, TTC29, TTC39A | 379 | 119 | 10425 | 1,39 | 1,00E+00 | 9,90E-01 | 9,99E+01 |

| | | | | | | | | | | | | |
|---|---|---|---|---|---|---|---|---|---|---|---|---|
| INTERPRO | IPR011990:Tetratricopeptide-like helical | 8 | 1,13 | 4,63E-01 | IFIT2, BBS4, TTC30B, TTC30A2, TTC29, SMYD4, NAPB, TTC39A | 645 | 204 | 20594 | 1,25 | 1,00E+00 | 9,99E-01 | 1,00E+02 |
| INTERPRO | IPR013026:Tetratricopeptide repeat-containing domain | 4 | 0,56 | 7,29E-01 | IFIT2, BBS4, TTC30B, TTC29 | 645 | 120 | 20594 | 1,06 | 1,00E+00 | 1,00E+00 | 1,00E+02 |
| Annotation | Enrichment Score: 0.4771858186009551 | | | | | | | | | | | |
| Category | Term | Count | % | PValue | Genes | List To | Pop Hit | Pop Tota | Fold Enr | Bonferroni | Benjamini | FDR |
| UP_KEYWO | Ligase | 14 | 1,97 | 1,87E-01 | DTX4, ACACA, UBA6, CTPS2, TRIM24, NHLRC1, ACSS3, TRIM21, FANCL, MARCH1, TRIM56, TRIM68, PARS2, LIPT2 | 678 | 330 | 22680 | 1,42 | 1,00E+00 | 7,27E-01 | 9,40E+01 |
| GOTERM_M | GO:0004842~ubiquitin-protein transferase activity | 15 | 2,12 | 1,90E-01 | HERC6, UBA7, NHLRC1, TRIM24, KLHDC7A, TRIM21, FANCL, MARCH1, TRIM56, G2E3, RNF141, TRIM68, NAIP2, HECTD2, KLHL13 | 575 | 326 | 17446 | 1,40 | 1,00E+00 | 9,78E-01 | 9,59E+01 |
| GOTERM_M | GO:0016874~ligase activity | 16 | 2,26 | 2,17E-01 | DTX4, ACACA, UBA6, CTPS2, TRIM24, NHLRC1, ACSS3, TRIM21, MARCH1, FANCL, TRIM56, G2E3, TRIM68, PARS2, HECTD2, LIPT2 | 575 | 362 | 17446 | 1,34 | 1,00E+00 | 9,72E-01 | 9,76E+01 |
| UP_KEYWO | Ubl conjugation pathway | 20 | 2,82 | 5,14E-01 | DTX4, HERC6, UBA6, UBE2L6, TRIM24, NHLRC1, UBE2QL1, TRIM21, SENP7, MARCH1, FANCL, TRIM56, G2E3, TRIM68, KLHL13, STAMBPL1, HECTD2, DET1, UBD, FBXO47 | 678 | 631 | 22680 | 1,06 | 1,00E+00 | 9,07E-01 | 1,00E+02 |
| INTERPRO | IPR013083:Zinc finger, RING/FYVE/PHD-type | 15 | 2,12 | 5,40E-01 | DTX4, DPF3, TRIM24, NHLRC1, TRIM21, SP140, MARCH1, FANCL, TRIM56, TRIM30A, G2E3, RNF141, TRIM68, TRIM30D, SYTL5 | 645 | 445 | 20594 | 1,08 | 1,00E+00 | 1,00E+00 | 1,00E+02 |
| GOTERM_M | GO:0061630~ubiquitin protein ligase activity | 7 | 0,99 | 6,43E-01 | MARCH1, FANCL, TRIM56, G2E3, HERC6, NHLRC1, TRIM24 | 575 | 199 | 17446 | 1,07 | 1,00E+00 | 9,99E-01 | 1,00E+02 |
| Annotation | Enrichment Score: 0.47530507396914207 | | | | | | | | | | | |
| Category | Term | Count | % | PValue | Genes | List To | Pop Hit | Pop Tota | Fold Enr | Bonferroni | Benjamini | FDR |
| SMART | SM00005:DEATH | 3 | 0,42 | 2,56E-01 | TNFRSF11B, UNC5B, ANK2 | 379 | 27 | 10425 | 3,06 | 1,00E+00 | 9,70E-01 | 9,77E+01 |
| INTERPRO | IPR000488:Death domain | 3 | 0,42 | 2,76E-01 | TNFRSF11B, UNC5B, ANK2 | 645 | 33 | 20594 | 2,90 | 1,00E+00 | 9,92E-01 | 9,94E+01 |
| INTERPRO | IPR011029:Death-like domain | 4 | 0,56 | 5,30E-01 | TNFRSF11B, UNC5B, ANK2, CASP12 | 645 | 89 | 20594 | 1,43 | 1,00E+00 | 1,00E+00 | 1,00E+02 |
| Annotation | Enrichment Score: 0.4743124965760021 | | | | | | | | | | | |
| Category | Term | Count | % | PValue | Genes | List To | Pop Hit | Pop Tota | Fold Enr | Bonferroni | Benjamini | FDR |
| GOTERM_BI | GO:0098792~xenophagy | 6 | 0,85 | 2,43E-01 | FANCL, DPF3, CLDN7, CD93, PNPO, ANXA5 | 588 | 103 | 18082 | 1,79 | 1,00E+00 | 9,96E-01 | 9,92E+01 |

| | | | | | | | | | | | | |
|---|---|---|---|---|---|---|---|---|---|---|---|---|
| GOTERM_BI | GO:0002230~positive regulation of defense response to virus by host | 6 | 0,85 | 3,63E-01 | FANCL, DPF3, CLDN7, CD93, PNPO, ANXA5 | 588 | 122 | 18082 | 1,51 | 1,00E+00 | 9,99E-01 | 1,00E+02 |
| GOTERM_BI | GO:0098779~mitophagy in response to mitochondrial depolarization | 6 | 0,85 | 4,28E-01 | P2RX5, DPF3, CD93, PNPO, HSF2BP, ANXA5 | 588 | 132 | 18082 | 1,40 | 1,00E+00 | 1,00E+00 | 1,00E+02 |
| Annotation | Enrichment Score: 0.47134377508770914 | | | | | | | | | | | |
| Category | Term | Count | % | PValue | Genes | List To | Pop Hit | Pop Tota | Fold Enr | Bonferroni | Benjamini | FDR |
| UP_KEYWOI | Flavoprotein | 7 | 0,99 | 1,93E-01 | STEAP4, FMO2, HAO2, FMO3, PNPO, ACAD9, ACAD12 | 678 | 130 | 22680 | 1,80 | 1,00E+00 | 7,21E-01 | 9,46E+01 |
| GOTERM_M | GO:0050660~flavin adenine dinucleotide binding | 4 | 0,56 | 4,24E-01 | FMO2, FMO3, ACAD9, ACAD12 | 575 | 72 | 17446 | 1,69 | 1,00E+00 | 9,95E-01 | 1,00E+02 |
| UP_KEYWOI | FAD | 5 | 0,71 | 4,70E-01 | STEAP4, FMO2, FMO3, ACAD9, ACAD12 | 678 | 118 | 22680 | 1,42 | 1,00E+00 | 8,95E-01 | 1,00E+02 |
| Annotation | Enrichment Score: 0.46858181909226515 | | | | | | | | | | | |
| Category | Term | Count | % | PValue | Genes | List To | Pop Hit | Pop Tota | Fold Enr | Bonferroni | Benjamini | FDR |
| UP_SEQ_FE | short sequence motif:Cell attachment site | 7 | 0,99 | 3,19E-02 | COL4A3, COL14A1, NID1, MFGE8, LRP2, COL5A2, SPP1 | 544 | 79 | 18012 | 2,93 | 1,00E+00 | 9,73E-01 | 4,18E+01 |
| INTERPRO | IPR026823:Complement Clr-like EGF domain | 3 | 0,42 | 1,39E-01 | CD93, NID1, LRP2 | 645 | 21 | 20594 | 4,56 | 1,00E+00 | 9,68E-01 | 9,08E+01 |
| UP_SEQ_FE | domain:EGF-like 2 | 5 | 0,71 | 2,48E-01 | CD93, IMPG2, NID1, MFGE8, LRP2 | 544 | 84 | 18012 | 1,97 | 1,00E+00 | 9,97E-01 | 9,91E+01 |
| UP_KEYWOI | EGF-like domain | 9 | 1,27 | 3,51E-01 | SELP, CD93, IMPG2, SORL1, TPO, NID1, MFGE8, LRP2, C1S1 | 678 | 224 | 22680 | 1,34 | 1,00E+00 | 8,31E-01 | 9,97E+01 |
| INTERPRO | IPR001881:EGF-like calcium-binding | 6 | 0,85 | 3,59E-01 | SELP, CD93, TPO, NID1, LRP2, C1S1 | 645 | 126 | 20594 | 1,52 | 1,00E+00 | 9,98E-01 | 9,99E+01 |
| INTERPRO | IPR018097:EGF-like calcium-binding, conserved site | 5 | 0,71 | 3,60E-01 | CD93, TPO, NID1, LRP2, C1S1 | 645 | 97 | 20594 | 1,65 | 1,00E+00 | 9,98E-01 | 9,99E+01 |
| INTERPRO | IPR000152:EGF-type aspartate/asparagine hydroxylation site | 5 | 0,71 | 3,67E-01 | CD93, TPO, NID1, LRP2, C1S1 | 645 | 98 | 20594 | 1,63 | 1,00E+00 | 9,98E-01 | 9,99E+01 |
| UP_SEQ_FE | domain:EGF-like 1 | 5 | 0,71 | 4,31E-01 | CD93, IMPG2, NID1, MFGE8, LRP2 | 544 | 111 | 18012 | 1,49 | 1,00E+00 | 1,00E+00 | 1,00E+02 |
| SMART | SM00179:EGF_CA | 6 | 0,85 | 4,83E-01 | SELP, CD93, TPO, NID1, LRP2, C1S1 | 379 | 126 | 10425 | 1,31 | 1,00E+00 | 9,91E-01 | 1,00E+02 |
| INTERPRO | IPR013032:EGF-like, conserved site | 7 | 0,99 | 5,83E-01 | SELP, CD93, IMPG2, TPO, NID1, MFGE8, LRP2 | 645 | 197 | 20594 | 1,13 | 1,00E+00 | 1,00E+00 | 1,00E+02 |

| | | | | | | | | | | | | |
|---|---|---|---|---|---|---|---|---|---|---|---|---|
| INTERPRO | IPR000742:Epidermal growth factor-like domain | 8 | 1,13 | 6,14E-01 | SELP, MEGF9, CD93, IMPG2, TPO, NID1, MFGE8, LRP2 | 645 | 237 | 20594 | 1,08 | 1,00E+00 | 1,00E+00 | 1,00E+02 |
| SMART | SM00181:EGF | 7 | 0,99 | 6,46E-01 | SELP, MEGF9, CD93, TPO, NID1, MFGE8, LRP2 | 379 | 181 | 10425 | 1,06 | 1,00E+00 | 9,98E-01 | 1,00E+02 |
| INTERPRO | IPR009030:Insulin-like growth factor binding protein, N-terminal | 3 | 0,42 | 9,17E-01 | CD93, NID1, LRP2 | 645 | 130 | 20594 | 0,74 | 1,00E+00 | 1,00E+00 | 1,00E+02 |
| Annotation | Enrichment Score: 0.4644703456738068 | | | | | | | | | | | |
| Category | Term | Count | % | PValue | Genes | List To | Pop Hit | Pop Tota | Fold Enr | Bonferroni | Benjamini | FDR |
| UP_KEYWO | Calmodulin-binding | 7 | 0,99 | 2,36E-01 | TRPM4, ATP2B2, SLC8A1, SNTB1, IQCG, PPP3CC, MYH10 | 678 | 139 | 22680 | 1,68 | 1,00E+00 | 7,70E-01 | 9,74E+01 |
| GOTERM_M | GO:0005516~calmodulin binding | 8 | 1,13 | 3,92E-01 | TRPM4, SYT1, ATP2B2, SLC8A1, SNTB1, IQCG, PPP3CC, MYH10 | 575 | 182 | 17446 | 1,33 | 1,00E+00 | 9,94E-01 | 9,99E+01 |
| UP_SEQ_FE | region of interest:Calmodulin-binding | 3 | 0,42 | 4,37E-01 | TRPM4, SLC8A1, SNTB1 | 544 | 49 | 18012 | 2,03 | 1,00E+00 | 1,00E+00 | 1,00E+02 |
| Annotation | Enrichment Score: 0.3812479311611693 | | | | | | | | | | | |
| Category | Term | Count | % | PValue | Genes | List To | Pop Hit | Pop Tota | Fold Enr | Bonferroni | Benjamini | FDR |
| UP_KEYWO | Methyltransferase | 8 | 1,13 | 2,74E-01 | METTL4, TRDMT1, BHMT, MTR, SMYD4, TPMT, METTL15, METTL7A3 | 678 | 176 | 22680 | 1,52 | 1,00E+00 | 7,96E-01 | 9,87E+01 |
| UP_SEQ_FE | binding site:S-adenosyl-L-methionine | 3 | 0,42 | 3,95E-01 | MTR, TPMT, METTL15 | 544 | 45 | 18012 | 2,21 | 1,00E+00 | 1,00E+00 | 1,00E+02 |
| GOTERM_B | GO:0032259~methylation | 7 | 0,99 | 4,70E-01 | METTL4, TRDMT1, MTR, SMYD4, TPMT, METTL15, METTL7A3 | 588 | 169 | 18082 | 1,27 | 1,00E+00 | 1,00E+00 | 1,00E+02 |
| GOTERM_M | GO:0008168~methyltransferase activity | 7 | 0,99 | 4,78E-01 | METTL4, TRDMT1, BHMT, MTR, SMYD4, TPMT, METTL15 | 575 | 168 | 17446 | 1,26 | 1,00E+00 | 9,97E-01 | 1,00E+02 |
| UP_KEYWO | S-adenosyl-L-methionine | 6 | 0,85 | 5,11E-01 | TRDMT1, MTR, TRMT12, SMYD4, TPMT, METTL15 | 678 | 158 | 22680 | 1,27 | 1,00E+00 | 9,08E-01 | 1,00E+02 |
| Annotation | Enrichment Score: 0.3707566140307781 | | | | | | | | | | | |
| Category | Term | Count | % | PValue | Genes | List To | Pop Hit | Pop Tota | Fold Enr | Bonferroni | Benjamini | FDR |
| GOTERM_B | GO:0007129~synapsis | 3 | 0,42 | 2,54E-01 | TEX19.1, MEIOB, STAG3 | 588 | 30 | 18082 | 3,08 | 1,00E+00 | 9,97E-01 | 9,94E+01 |
| UP_KEYWO | Meiosis | 4 | 0,56 | 5,06E-01 | TEX19.1, TDRD9, MEIOB, STAG3 | 678 | 90 | 22680 | 1,49 | 1,00E+00 | 9,07E-01 | 1,00E+02 |
| GOTERM_B | GO:0051321~meiotic cell cycle | 4 | 0,56 | 6,00E-01 | TEX19.1, TDRD9, MEIOB, STAG3 | 588 | 95 | 18082 | 1,29 | 1,00E+00 | 1,00E+00 | 1,00E+02 |

| | | | | | | | | | | | | |
|---|---|---|---|---|---|---|---|---|---|---|---|---|
| Annotation | Enrichment Score: 0.31510761594477754 | | | | | | | | | | | |
| Category | Term | Count | % | PValue | Genes | List To | Pop Hit | Pop Tot | Fold Enr | Bonferroni | Benjamini | FDR |
| UP_KEYWO | LIM domain | 4 | 0,56 | 3,81E-01 | AJUBA, LIMD2, CSRP1, CSRP3 | 678 | 74 | 22680 | 1,81 | 1,00E+00 | 8,50E-01 | 9,99E+01 |
| INTERPRO | IPR001781:Zinc finger, LIM-type | 4 | 0,56 | 4,09E-01 | AJUBA, LIMD2, CSRP1, CSRP3 | 645 | 74 | 20594 | 1,73 | 1,00E+00 | 9,99E-01 | 1,00E+02 |
| UP_SEQ_FE | domain:LIM zinc-binding 2 | 3 | 0,42 | 4,47E-01 | AJUBA, CSRP1, CSRP3 | 544 | 50 | 18012 | 1,99 | 1,00E+00 | 1,00E+00 | 1,00E+02 |
| UP_SEQ_FE | domain:LIM zinc-binding 1 | 3 | 0,42 | 4,47E-01 | AJUBA, CSRP1, CSRP3 | 544 | 50 | 18012 | 1,99 | 1,00E+00 | 1,00E+00 | 1,00E+02 |
| SMART | SM00132:LIM | 4 | 0,56 | 5,06E-01 | AJUBA, LIMD2, CSRP1, CSRP3 | 379 | 74 | 10425 | 1,49 | 1,00E+00 | 9,91E-01 | 1,00E+02 |
| UP_SEQ_FE | short sequence motif:Nuclear localization signal | 9 | 1,27 | 8,16E-01 | AJUBA, PLAG1, TRIM30A, PBXIP1, CSRP1, TRIM24, CSRP3, MBD1, PARP2 | 544 | 343 | 18012 | 0,87 | 1,00E+00 | 1,00E+00 | 1,00E+02 |
| Annotation | Enrichment Score: 0.3063503760059623 | | | | | | | | | | | |
| Category | Term | Count | % | PValue | Genes | List To | Pop Hit | Pop Tot | Fold Enr | Bonferroni | Benjamini | FDR |
| KEGG_PATH | mmu04012:ErbB signaling pathway | 5 | 0,71 | 2,65E-01 | PRKCA, PAK6, CDKN1A, PAK1, SRC | 233 | 87 | 7720 | 1,90 | 1,00E+00 | 8,69E-01 | 9,82E+01 |
| KEGG_PATH | mmu05161:Hepatitis B | 5 | 0,71 | 6,44E-01 | PRKCA, CDKN1A, CASP12, TLR3, SRC | 233 | 146 | 7720 | 1,13 | 1,00E+00 | 9,52E-01 | 1,00E+02 |
| KEGG_PATH | mmu04921:Oxytocin signaling pathway | 5 | 0,71 | 7,05E-01 | ACTG1, PRKCA, CDKN1A, PPP3CC, SRC | 233 | 158 | 7720 | 1,05 | 1,00E+00 | 9,65E-01 | 1,00E+02 |
| Annotation | Enrichment Score: 0.2855452969144743 | | | | | | | | | | | |
| Category | Term | Count | % | PValue | Genes | List To | Pop Hit | Pop Tot | Fold Enr | Bonferroni | Benjamini | FDR |
| KEGG_PATH | mmu04015:Rap1 signaling pathway | 9 | 1,27 | 3,16E-01 | ACTG1, PRKCA, KLK1B4, FLT4, RAPGEF5, ANGPT1, CDH1, HGF, SRC | 233 | 214 | 7720 | 1,39 | 1,00E+00 | 8,90E-01 | 9,93E+01 |
| KEGG_PATH | mmu04520:Adherens junction | 3 | 0,42 | 6,42E-01 | ACTG1, CDH1, SRC | 233 | 72 | 7720 | 1,38 | 1,00E+00 | 9,53E-01 | 1,00E+02 |
| KEGG_PATH | mmu05100:Bacterial invasion of epithelial cells | 3 | 0,42 | 6,85E-01 | ACTG1, CDH1, SRC | 233 | 78 | 7720 | 1,27 | 1,00E+00 | 9,61E-01 | 1,00E+02 |
| Annotation | Enrichment Score: 0.27932797722261604 | | | | | | | | | | | |
| Category | Term | Count | % | PValue | Genes | List To | Pop Hit | Pop Tot | Fold Enr | Bonferroni | Benjamini | FDR |

| | | | | | | | | | | | | |
|---|---|---|---|---|---|---|---|---|---|---|---|---|
| UP_KEYWOI | Metalloprotease | 6 | 0,85 | 4,24E-01 | LNPEP, ACE, CLCA1, STAMBPL1, MMP15, MMP12 | 678 | 143 | 22680 | 1,40 | 1,00E+00 | 8,68E-01 | 9,99E+01 |
| GOTERM_M | GO:0008237~metallopeptidase activity | 7 | 0,99 | 4,31E-01 | LNPEP, ACE, ADAMTS9, CLCA1, STAMBPL1, MMP15, MMP12 | 575 | 160 | 17446 | 1,33 | 1,00E+00 | 9,95E-01 | 1,00E+02 |
| GOTERM_BI | GO:0006508~proteolysis | 20 | 2,82 | 5,30E-01 | CAPN6, TMPRSS2, CLCA1, KLK1B4, PRSS41, KY, HGF, MMP15, MMP12, SENP7, TMPRSS4, PSMB9, LNPEP, ADAMTS9, ACE, CD46, CASP12, STAMBPL1, ASRGL1, C1S1 | 588 | 582 | 18082 | 1,06 | 1,00E+00 | 1,00E+00 | 1,00E+02 |
| GOTERM_M | GO:0008233~peptidase activity | 17 | 2,4 | 6,32E-01 | TMPRSS2, CLCA1, PRSS41, KY, MMP15, MMP12, SENP7, TMPRSS4, PSMB9, LNPEP, ADAMTS6, ACE, CASP12, STAMBPL1, ASRGL1, SLPI, C1S1 | 575 | 516 | 17446 | 1,00 | 1,00E+00 | 9,99E-01 | 1,00E+02 |
| UP_KEYWOI | Protease | 16 | 2,26 | 6,54E-01 | TMPRSS2, CLCA1, PRSS41, KY, MMP15, SENP7, MMP12, TMPRSS4, PSMB9, LNPEP, ACE, CASP12, STAMBPL1, ASRGL1, SLPI, C1S1 | 678 | 542 | 22680 | 0,99 | 1,00E+00 | 9,51E-01 | 1,00E+02 |
| Annotation | Enrichment Score: 0.2770427198719615 | | | | | | | | | | | |
| Category | Term | Count | % | PValue | Genes | List To | Pop Hit | Pop Tota | Fold Enr | Bonferroni | Benjamini | FDR |
| UP_KEYWOI | Chloride | 4 | 0,56 | 3,23E-01 | AMY1, FXYD3, CLCA1, CLIC5 | 678 | 67 | 22680 | 2,00 | 1,00E+00 | 8,40E-01 | 9,95E+01 |
| GOTERM_M | GO:0005254~chloride channel activity | 3 | 0,42 | 6,19E-01 | FXYD3, CLCA1, CLIC5 | 575 | 63 | 17446 | 1,44 | 1,00E+00 | 9,99E-01 | 1,00E+02 |
| GOTERM_BI | GO:0006821~chloride transport | 3 | 0,42 | 7,38E-01 | FXYD3, CLCA1, CLIC5 | 588 | 80 | 18082 | 1,15 | 1,00E+00 | 1,00E+00 | 1,00E+02 |
| Annotation | Enrichment Score: 0.2371853575041954 | | | | | | | | | | | |
| Category | Term | Count | % | PValue | Genes | List To | Pop Hit | Pop Tota | Fold Enr | Bonferroni | Benjamini | FDR |
| UP_SEQ_FE | domain:Peptidase S1 | 6 | 0,85 | 2,61E-01 | TMPRSS2, KLK1B4, PRSS41, HGF, C1S1, TMPRSS4 | 544 | 114 | 18012 | 1,74 | 1,00E+00 | 9,98E-01 | 9,94E+01 |
| UP_SEQ_FE | active site:Charge relay system | 8 | 1,13 | 4,20E-01 | TMPRSS2, PNLIPRP1, BCHE, ACNAT2, PRSS41, ABHD2, C1S1, TMPRSS4 | 544 | 205 | 18012 | 1,29 | 1,00E+00 | 1,00E+00 | 1,00E+02 |
| INTERPRO | IPR009003:Trypsin-like cysteine/serine peptidase domain | 7 | 0,99 | 4,29E-01 | TMPRSS2, KLK1B4, FAM111A, PRSS41, HGF, C1S1, TMPRSS4 | 645 | 168 | 20594 | 1,33 | 1,00E+00 | 9,99E-01 | 1,00E+02 |
| INTERPRO | IPR001314:Peptidase S1A, chymotrypsin-type | 6 | 0,85 | 5,12E-01 | TMPRSS2, KLK1B4, PRSS41, HGF, C1S1, TMPRSS4 | 645 | 151 | 20594 | 1,27 | 1,00E+00 | 1,00E+00 | 1,00E+02 |
| INTERPRO | IPR001254:Peptidase S1 | 6 | 0,85 | 5,64E-01 | TMPRSS2, KLK1B4, PRSS41, HGF, C1S1, TMPRSS4 | 645 | 160 | 20594 | 1,20 | 1,00E+00 | 1,00E+00 | 1,00E+02 |
| SMART | SM00020:Tryp_SPc | 6 | 0,85 | 6,84E-01 | TMPRSS2, KLK1B4, PRSS41, HGF, C1S1, TMPRSS4 | 379 | 158 | 10425 | 1,04 | 1,00E+00 | 9,98E-01 | 1,00E+02 |
| GOTERM_M | GO:0004252~serine-type endopeptidase activity | 7 | 0,99 | 7,02E-01 | TMPRSS2, KLK1B4, PRSS41, HGF, C1S1, MMP12, TMPRSS4 | 575 | 212 | 17446 | 1,00 | 1,00E+00 | 1,00E+00 | 1,00E+02 |
| INTERPRO | IPR018114:Peptidase S1, trypsin family, active site | 4 | 0,56 | 8,06E-01 | TMPRSS2, KLK1B4, PRSS41, TMPRSS4 | 645 | 137 | 20594 | 0,93 | 1,00E+00 | 1,00E+00 | 1,00E+02 |

| | | | | | | | | | | | | |
|---|---|---|---|---|---|---|---|---|---|---|---|---|
| UP_KEYWO | Serine protease | 4 | 0,56 | 8,93E-01 | TMPRSS2, PRSS41, C1S1, TMPRSS4 | 678 | 173 | 22680 | 0,77 | 1,00E+00 | 9,94E-01 | 1,00E+02 |
| GOTERM_M | GO:0008236~serine-type peptidase activity | 4 | 0,56 | 9,08E-01 | TMPRSS2, PRSS41, C1S1, TMPRSS4 | 575 | 163 | 17446 | 0,74 | 1,00E+00 | 1,00E+00 | 1,00E+02 |
| Annotation | Enrichment Score: 0.23550633999485235 | | | | | | | | | | | |
| Category | Term | Count | % | PValue | Genes | List To | Pop Hit | Pop Tot | Fold Enr | Bonferroni | Benjamini | FDR |
| GOTERM_M | GO:0008237~metallopeptidase activity | 7 | 0,99 | 4,31E-01 | LNPEP, ACE, ADAMTS9, CLCA1, STAMBPL1, MMP15, MMP12 | 575 | 160 | 17446 | 1,33 | 1,00E+00 | 9,95E-01 | 1,00E+02 |
| INTERPRO | IPR024079:Metallopeptidase, catalytic domain | 4 | 0,56 | 5,88E-01 | ADAMTS6, ADAMTS9, MMP15, MMP12 | 645 | 97 | 20594 | 1,32 | 1,00E+00 | 1,00E+00 | 1,00E+02 |
| GOTERM_M | GO:0004222~metalloendopeptidase activity | 4 | 0,56 | 7,75E-01 | ADAMTS6, ADAMTS9, MMP15, MMP12 | 575 | 123 | 17446 | 0,99 | 1,00E+00 | 1,00E+00 | 1,00E+02 |
| Annotation | Enrichment Score: 0.2192520170304447 | | | | | | | | | | | |
| Category | Term | Count | % | PValue | Genes | List To | Pop Hit | Pop Tot | Fold Enr | Bonferroni | Benjamini | FDR |
| INTERPRO | IPR013761:Sterile alpha motif/pointed domain | 5 | 0,71 | 4,80E-01 | PPP1R9A, NAB2, SAMD10, BICC1, SAMD9L | 645 | 114 | 20594 | 1,40 | 1,00E+00 | 1,00E+00 | 1,00E+02 |
| INTERPRO | IPR001660:Sterile alpha motif domain | 4 | 0,56 | 5,45E-01 | PPP1R9A, SAMD10, BICC1, SAMD9L | 645 | 91 | 20594 | 1,40 | 1,00E+00 | 1,00E+00 | 1,00E+02 |
| UP_SEQ_FE | domain:SAM | 3 | 0,42 | 6,28E-01 | SAMD10, BICC1, SAMD9L | 544 | 70 | 18012 | 1,42 | 1,00E+00 | 1,00E+00 | 1,00E+02 |
| SMART | SM00454:SAM | 3 | 0,42 | 8,09E-01 | PPP1R9A, SAMD10, BICC1 | 379 | 83 | 10425 | 0,99 | 1,00E+00 | 1,00E+00 | 1,00E+02 |
| Annotation | Enrichment Score: 0.212805069102567 | | | | | | | | | | | |
| Category | Term | Count | % | PValue | Genes | List To | Pop Hit | Pop Tot | Fold Enr | Bonferroni | Benjamini | FDR |
| BIOCARTA | m_gpcrPathway:Signaling Pathway from G-Protein Families | 3 | 0,42 | 3,93E-01 | PRKCA, RPS6KA3, PPP3CC | 52 | 34 | 1289 | 2,19 | 1,00E+00 | 1,00E+00 | 9,97E+01 |
| KEGG_PATH | mmu04720:Long-term potentiation | 3 | 0,42 | 5,95E-01 | PRKCA, RPS6KA3, PPP3CC | 233 | 66 | 7720 | 1,51 | 1,00E+00 | 9,59E-01 | 1,00E+02 |
| KEGG_PATH | mmu04010:MAPK signaling pathway | 4 | 0,56 | 9,84E-01 | PRKCA, RPS6KA3, PPP3CC, PAK1 | 233 | 253 | 7720 | 0,52 | 1,00E+00 | 1,00E+00 | 1,00E+02 |
| Annotation | Enrichment Score: 0.20797391515180494 | | | | | | | | | | | |

| Category | Term | Count | % | PValue | Genes | List To | Pop Hit | Pop Tot | Fold Enr | Bonferroni | Benjamini | FDR |
|---|---|---|---|---|---|---|---|---|---|---|---|---|
| INTERPRO | IPR001965:Zinc finger, PHD-type | 4 | 0,56 | 5,45E-01 | DPF3, G2E3, TRIM24, SP140 | 645 | 91 | 20594 | 1,40 | 1,00E+00 | 1,00E+00 | 1,00E+02 |
| INTERPRO | IPR019786:Zinc finger, PHD-type, conserved site | 3 | 0,42 | 5,99E-01 | G2E3, TRIM24, SP140 | 645 | 64 | 20594 | 1,50 | 1,00E+00 | 1,00E+00 | 1,00E+02 |
| INTERPRO | IPR011011:Zinc finger, FYVE/PHD-type | 5 | 0,71 | 6,19E-01 | DPF3, G2E3, SYTL5, TRIM24, SP140 | 645 | 136 | 20594 | 1,17 | 1,00E+00 | 1,00E+00 | 1,00E+02 |
| SMART | SM00249:PHD | 4 | 0,56 | 6,46E-01 | DPF3, G2E3, TRIM24, SP140 | 379 | 91 | 10425 | 1,21 | 1,00E+00 | 9,97E-01 | 1,00E+02 |
| INTERPRO | IPR019787:Zinc finger, PHD-finger | 3 | 0,42 | 6,99E-01 | DPF3, TRIM24, SP140 | 645 | 77 | 20594 | 1,24 | 1,00E+00 | 1,00E+00 | 1,00E+02 |
| Annotation | Enrichment Score: 0.18776709072429665 | | | | | | | | | | | |
| Category | Term | Count | % | PValue | Genes | List To | Pop Hit | Pop Tot | Fold Enr | Bonferroni | Benjamini | FDR |
| GOTERM_C | GO:0005777~peroxisome | 5 | 0,71 | 6,11E-01 | FAR1, HRASLS, ACNAT2, HAO2, ABCD2 | 633 | 131 | 19662 | 1,19 | 1,00E+00 | 9,77E-01 | 1,00E+02 |
| UP_KEYWO | Peroxisome | 4 | 0,56 | 6,23E-01 | FAR1, ACNAT2, HAO2, ABCD2 | 678 | 107 | 22680 | 1,25 | 1,00E+00 | 9,42E-01 | 1,00E+02 |
| KEGG_PATH | mmu04146:Peroxisome | 3 | 0,42 | 7,18E-01 | FAR1, HAO2, ABCD2 | 233 | 83 | 7720 | 1,20 | 1,00E+00 | 9,68E-01 | 1,00E+02 |
| Annotation | Enrichment Score: 0.18352863503760677 | | | | | | | | | | | |
| Category | Term | Count | % | PValue | Genes | List To | Pop Hit | Pop Tot | Fold Enr | Bonferroni | Benjamini | FDR |
| UP_KEYWO | DNA recombination | 4 | 0,56 | 3,48E-01 | RPA1, XRCC4, RAD51B, SPIDR | 678 | 70 | 22680 | 1,91 | 1,00E+00 | 8,35E-01 | 9,97E+01 |
| GOTERM_BI | GO:0000724~double-strand break repair via homologous recombination | 4 | 0,56 | 3,80E-01 | RPA1, RAD51B, MEIOB, SPIDR | 588 | 68 | 18082 | 1,81 | 1,00E+00 | 9,99E-01 | 1,00E+02 |
| GOTERM_BI | GO:0006310~DNA recombination | 4 | 0,56 | 5,24E-01 | RPA1, XRCC4, RAD51B, SPIDR | 588 | 85 | 18082 | 1,45 | 1,00E+00 | 1,00E+00 | 1,00E+02 |
| GOTERM_BI | GO:0006281~DNA repair | 8 | 1,13 | 8,95E-01 | RPA1, FANCL, XRCC4, RAD51B, RAD9B, SPIDR, EXO5, PARP2 | 588 | 318 | 18082 | 0,77 | 1,00E+00 | 1,00E+00 | 1,00E+02 |
| UP_KEYWO | DNA repair | 6 | 0,85 | 9,22E-01 | RPA1, FANCL, XRCC4, RAD51B, SPIDR, EXO5 | 678 | 279 | 22680 | 0,72 | 1,00E+00 | 9,97E-01 | 1,00E+02 |
| GOTERM_BI | GO:0006974~cellular response to DNA damage stimulus | 10 | 1,41 | 9,31E-01 | RPA1, FANCL, XRCC4, CDKN1A, RAD51B, BTG2, ZFP791, SPIDR, EXO5, NUGGC | 588 | 420 | 18082 | 0,73 | 1,00E+00 | 1,00E+00 | 1,00E+02 |
| UP_KEYWO | DNA damage | 6 | 0,85 | 9,74E-01 | RPA1, FANCL, XRCC4, RAD51B, SPIDR, EXO5 | 678 | 336 | 22680 | 0,60 | 1,00E+00 | 1,00E+00 | 1,00E+02 |

| Annotation | Enrichment Score: 0.17729488759353984 | | | | | | | | | | | |
|---|---|---|---|---|---|---|---|---|---|---|---|---|
| Category | Term | Count | % | PValue | Genes | List To | Pop Hit | Pop Tot | Fold Enr | Bonferroni | Benjamini | FDR |
| UP_KEYWO | Nucleotide-binding | 57 | 8,04 | 3,25E-01 | RAD51B, MYO7B, DQX1, ACSS3, PAK6, ACTG1, ATP2B2, RASL10B, GBP10, RHOC, MLKL, PAK1, ATP8B5, NT5E, RAB27A, GBP8, TRPM4, PRKCA, RABL2, GBP7, GBP6, HKDC1, GBP9, RPS6KC1, ABCC12, CNGA3, ABCB1A, OAS1A, GBP3, GBP2, POMK, STK10, TDRD9, UBA6, CTPS2, UBE2QL1, SRC, DNAH5, NAIP2, PARS2, TAP1, ABCD2, EHD1, AGK, PAPSS2, FLT4, ACACA, UBE2L6, ACLY, PCK2, SMC2, NUGGC, RPS6KA3, TGTP1, GRK5, ATP7B, MYH10 | 678 | 1754 | 22680 | 1,09 | 1,00E+00 | 8,38E-01 | 9,95E+01 |
| UP_KEYWO | ATP-binding | 41 | 5,78 | 5,69E-01 | RAD51B, POMK, MYO7B, STK10, TDRD9, DQX1, UBA6, CTPS2, UBE2QL1, ACSS3, SRC, DNAH5, ACTG1, PAK6, ATP2B2, NAIP2, PARS2, TAP1, ABCD2, MLKL, PAK1, ATP8B5, EHD1, AGK, PAPSS2, PRKCA, TRPM4, HKDC1, FLT4, ACACA, UBE2L6, RPS6KC1, ACLY, ABCC12, SMC2, RPS6KA3, ABCB1A, OAS1A, GRK5, ATP7B, MYH10 | 678 | 1363 | 22680 | 1,01 | 1,00E+00 | 9,25E-01 | 1,00E+02 |
| GOTERM_M | GO:0005524~ATP binding | 45 | 6,35 | 8,20E-01 | RAD51B, POMK, MYO7B, STK10, TDRD9, DQX1, UBA6, CTPS2, UBE2QL1, ACSS3, SRC, DNAH5, ACTG1, PAK6, ATP2B2, NAIP2, P2RY4, PARS2, TAP1, OASL1, ABCD2, MLKL, PAK1, ATP8B5, EHD1, AGK, PAPSS2, PMS1, PRKCA, TRPM4, HKDC1, FLT4, ACACA, UBE2L6, RPS6KC1, ACLY, ABCC12, SMC2, P2RX5, RPS6KA3, ABCB1A, OAS1A, GRK5, ATP7B, MYH10 | 575 | 1507 | 17446 | 0,91 | 1,00E+00 | 1,00E+00 | 1,00E+02 |
| GOTERM_M | GO:0000166~nucleotide binding | 55 | 7,76 | 9,18E-01 | RAD51B, MYO7B, DQX1, ACSS3, PAK6, ACTG1, ATP2B2, RASL10B, RHOC, MLKL, PAK1, ATP8B5, NT5E, RAB27A, RABL2, PRKCA, TRPM4, RBM43, HKDC1, RPS6KC1, ABCC12, CNGA3, ABCB1A, GBP3, GBP2, NMI, POMK, STK10, TDRD9, TRA2A, POLA1, MYEF2, UBA6, CTPS2, UBE2QL1, DNAH5, SRC, NAIP2, PARS2, TAP1, ABCD2, EHD1, AGK, PAPSS2, FLT4, ACACA, UBE2L6, ACLY, PCK2, SMC2, NUGGC, RPS6KA3, GRK5, ATP7B, MYH10 | 575 | 1936 | 17446 | 0,86 | 1,00E+00 | 1,00E+00 | 1,00E+02 |
| UP_SEQ_FE | nucleotide phosphate-binding region:ATP | 23 | 3,24 | 9,33E-01 | PRKCA, RAD51B, STK10, FLT4, HKDC1, TDRD9, ACACA, DQX1, RPS6KC1, UBA6, ACLY, SMC2, SRC, DNAH5, PAK6, RPS6KA3, TAP1, ABCD2, GRK5, PAK1, EHD1, PAPSS2, MYH10 | 544 | 963 | 18012 | 0,79 | 1,00E+00 | 1,00E+00 | 1,00E+02 |
| Annotation | Enrichment Score: 0.17361958687772067 | | | | | | | | | | | |
| Category | Term | Count | % | PValue | Genes | List To | Pop Hit | Pop Tot | Fold Enr | Bonferroni | Benjamini | FDR |
| UP_KEYWO | Differentiation | 22 | 3,1 | 3,76E-01 | TEX19.1, EID1, SYT1, RNF17, MYO7B, TDRD9, BEX1, PARP11, ECT2, CSRP3, ROBO1, ECSCR, SEMA3D, VAMP5, IQCG, ANGPT1, CATSPERG2, SRD5A2, BMP8B, BMP5, SEMA4A, NPHP1 | 678 | 646 | 22680 | 1,14 | 1,00E+00 | 8,49E-01 | 9,98E+01 |
| GOTERM_BI | GO:0030154~cell differentiation | 24 | 3,39 | 7,13E-01 | TEX19.1, EID1, SYT1, RNF17, MYO7B, TDRD9, BEX1, PARP11, ECT2, CSRP3, PBXIP1, ROBO1, ECSCR, MEG3, SEMA3D, VAMP5, IQCG, ANGPT1, CATSPERG2, SRD5A2, SEMA4A, BMP8B, BMP5, NPHP1 | 588 | 780 | 18082 | 0,95 | 1,00E+00 | 1,00E+00 | 1,00E+02 |
| UP_KEYWO | Developmental protein | 26 | 3,67 | 8,12E-01 | BMP10, RNF17, SORL1, TDRD9, BEX1, BICC1, G2E3, LBH, UNC5B, ROBO1, SEMA3D, ANGPT1, CATSPERG2, DAB2IP, THEMIS, CSRP3, SMO, ID2, MEOX1, ECSCR, VAMP5, SUPT20, SEMA4A, CR1L, BMP8B, BMP5 | 678 | 976 | 22680 | 0,89 | 1,00E+00 | 9,83E-01 | 1,00E+02 |

| | | | | | | | | | | | | |
|---|---|---|---|---|---|---|---|---|---|---|---|---|
| GOTERM_BI | GO:0007275~multicellular organism development | 27 | 3,81 | 9,28E-01 | BMP10, RNF17, SORL1, TDRD9, BEX1, BICC1, G2E3, LBH, UNC5B, ROBO1, SEMA3D, ANGPT1, CATSPERG2, DAB2IP, THEMIS, ZMYM6, CSRP3, SMO, ID2, MEOX1, ECSCR, VAMP5, SUPT20, SEMA4A, CR1L, BMP8B, BMP5 | 588 | 1029 | 18082 | 0,81 | 1,00E+00 | 1,00E+00 | 1,00E+02 |
| Annotation | Enrichment Score: 0.17244649622250158 | | | | | | | | | | | |
| Category | Term | Count | % | PValue | Genes | List To | Pop Hit | Pop Tota | Fold Enr | Bonferroni | Benjamini | FDR |
| UP_SEQ_FE | domain:C2 | 3 | 0,42 | 6,28E-01 | PRKCA, CAPN6, DAB2IP | 544 | 70 | 18012 | 1,42 | 1,00E+00 | 1,00E+00 | 1,00E+02 |
| SMART | SM00239:C2 | 5 | 0,71 | 6,69E-01 | PRKCA, CAPN6, SYT1, DAB2IP, SYTL5 | 379 | 125 | 10425 | 1,10 | 1,00E+00 | 9,98E-01 | 1,00E+02 |
| INTERPRO | IPR000008:C2 calcium-dependent membrane targeting | 5 | 0,71 | 7,23E-01 | PRKCA, CAPN6, SYT1, DAB2IP, SYTL5 | 645 | 156 | 20594 | 1,02 | 1,00E+00 | 1,00E+00 | 1,00E+02 |
| Annotation | Enrichment Score: 0.16715657237918427 | | | | | | | | | | | |
| Category | Term | Count | % | PValue | Genes | List To | Pop Hit | Pop Tota | Fold Enr | Bonferroni | Benjamini | FDR |
| UP_KEYWO | Postsynaptic cell membrane | 6 | 0,85 | 6,07E-01 | ANK2, GRIP1, DMD, GABBR2, GRID1, KCTD12 | 678 | 176 | 22680 | 1,14 | 1,00E+00 | 9,39E-01 | 1,00E+02 |
| GOTERM_C | GO:0045202~synapse | 16 | 2,26 | 6,60E-01 | SYT1, CCL2, C4A, CRYAB, GRIP1, GABBR2, ADORA1, SLC17A8, ATP2B2, PPP1R9A, RGS12, ANK2, DMD, SNTB1, GRID1, KCTD12 | 633 | 505 | 19662 | 0,98 | 1,00E+00 | 9,82E-01 | 1,00E+02 |
| GOTERM_C | GO:0045211~postsynaptic membrane | 7 | 0,99 | 7,22E-01 | ANK2, GRIP1, DMD, GABBR2, ADORA1, GRID1, KCTD12 | 633 | 222 | 19662 | 0,98 | 1,00E+00 | 9,89E-01 | 1,00E+02 |
| UP_KEYWO | Synapse | 10 | 1,41 | 7,42E-01 | SYT1, SLC17A8, ATP2B2, RGS12, ANK2, GRIP1, DMD, GABBR2, GRID1, KCTD12 | 678 | 357 | 22680 | 0,94 | 1,00E+00 | 9,71E-01 | 1,00E+02 |
| Annotation | Enrichment Score: 0.15662109820302758 | | | | | | | | | | | |
| Category | Term | Count | % | PValue | Genes | List To | Pop Hit | Pop Tota | Fold Enr | Bonferroni | Benjamini | FDR |
| UP_SEQ_FE | domain:DH | 3 | 0,42 | 5,35E-01 | ARHGEF16, ARHGEF9, ECT2 | 544 | 59 | 18012 | 1,68 | 1,00E+00 | 1,00E+00 | 1,00E+02 |
| INTERPRO | IPR000219:Dbl homology (DH) domain | 3 | 0,42 | 6,32E-01 | ARHGEF16, ARHGEF9, ECT2 | 645 | 68 | 20594 | 1,41 | 1,00E+00 | 1,00E+00 | 1,00E+02 |
| SMART | SM00325:RhoGEF | 3 | 0,42 | 6,89E-01 | ARHGEF16, ARHGEF9, ECT2 | 379 | 65 | 10425 | 1,27 | 1,00E+00 | 9,98E-01 | 1,00E+02 |
| GOTERM_M | GO:0005089~Rho guanyl-nucleotide exchange factor activity | 3 | 0,42 | 7,05E-01 | ARHGEF16, ARHGEF9, ECT2 | 575 | 74 | 17446 | 1,23 | 1,00E+00 | 1,00E+00 | 1,00E+02 |
| GOTERM_BI | GO:0035023~regulation of Rho protein signal transduction | 3 | 0,42 | 7,18E-01 | ARHGEF16, ARHGEF9, ECT2 | 588 | 77 | 18082 | 1,20 | 1,00E+00 | 1,00E+00 | 1,00E+02 |

| | | | | | | | | | | | | |
|---|---|---|---|---|---|---|---|---|---|---|---|---|
| UP_KEYWOI | Guanine-nucleotide releasing factor | 4 | 0,56 | 7,63E-01 | ARHGEF16, RAPGEF5, ARHGEF9, ECT2 | 678 | 133 | 22680 | 1,01 | 1,00E+00 | 9,73E-01 | 1,00E+02 |
| GOTERM_M | GO:0005085~guanyl-nucleotide exchange factor activity | 4 | 0,56 | 8,91E-01 | ARHGEF16, RAPGEF5, ARHGEF9, ECT2 | 575 | 156 | 17446 | 0,78 | 1,00E+00 | 1,00E+00 | 1,00E+02 |
| Annotation | Enrichment Score: 0.1452717666218312 | | | | | | | | | | | |
| Category | Term | Count | % | PValue | Genes | List To | Pop Hit | Pop Tota | Fold Enr | Bonferroni | Benjamini | FDR |
| KEGG_PATH | mmu04510:Focal adhesion | 10 | 1,41 | 1,71E-01 | ACTG1, PRKCA, PAK6, COL4A3, FLT4, HGF, PAK1, COL5A2, SRC, SPP1 | 233 | 207 | 7720 | 1,60 | 1,00E+00 | 8,05E-01 | 9,12E+01 |
| KEGG_PATH | mmu04012:ErbB signaling pathway | 5 | 0,71 | 2,65E-01 | PRKCA, PAK6, CDKN1A, PAK1, SRC | 233 | 87 | 7720 | 1,90 | 1,00E+00 | 8,69E-01 | 9,82E+01 |
| BIOCARTA | m_pyk2Pathway:Links between Pyk2 and Map Kinases | 3 | 0,42 | 3,05E-01 | PRKCA, PAK1, SRC | 52 | 28 | 1289 | 2,66 | 1,00E+00 | 1,00E+00 | 9,85E+01 |
| BIOCARTA | m_at1rPathway:Angiotensin II mediated activation of JNK Pathway via Pyk2 dependent signaling | 3 | 0,42 | 3,79E-01 | PRKCA, PAK1, SRC | 52 | 33 | 1289 | 2,25 | 1,00E+00 | 1,00E+00 | 9,96E+01 |
| SMART | SM00133:S_TK_X | 3 | 0,42 | 4,89E-01 | PRKCA, RPS6KA3, GRK5 | 379 | 45 | 10425 | 1,83 | 1,00E+00 | 9,90E-01 | 1,00E+02 |
| INTERPRO | IPR000961:AGC-kinase, C-terminal | 3 | 0,42 | 4,97E-01 | PRKCA, RPS6KA3, GRK5 | 645 | 53 | 20594 | 1,81 | 1,00E+00 | 1,00E+00 | 1,00E+02 |
| UP_SEQ_FE | domain:AGC-kinase C-terminal | 3 | 0,42 | 4,97E-01 | PRKCA, RPS6KA3, GRK5 | 544 | 55 | 18012 | 1,81 | 1,00E+00 | 1,00E+00 | 1,00E+02 |
| GOTERM_BI | GO:0046777~protein autophosphorylation | 7 | 0,99 | 5,48E-01 | PRKCA, STK10, FLT4, GRK5, PAK1, TRIM24, SRC | 588 | 183 | 18082 | 1,18 | 1,00E+00 | 1,00E+00 | 1,00E+02 |
| GOTERM_M | GO:0004702~receptor signaling protein serine/threonine kinase activity | 3 | 0,42 | 5,64E-01 | PAK6, STK10, PAK1 | 575 | 57 | 17446 | 1,60 | 1,00E+00 | 9,99E-01 | 1,00E+02 |
| UP_SEQ_FE | active site:Proton acceptor | 19 | 2,68 | 8,07E-01 | PRKCA, STK10, FLT4, G6PDX, RPS6KC1, DHRS9, CNP, SRC, PAK6, QPCT, RPS6KA3, SULT1B1, ALDH1B1, HAO2, TPO, GRK5, PAK1, ACAD9, NEU3 | 544 | 710 | 18012 | 0,89 | 1,00E+00 | 1,00E+00 | 1,00E+02 |
| INTERPRO | IPR001245:Serine-threonine/tyrosine-protein kinase catalytic domain | 4 | 0,56 | 8,14E-01 | POMK, FLT4, MLKL, SRC | 645 | 139 | 20594 | 0,92 | 1,00E+00 | 1,00E+00 | 1,00E+02 |
| UP_SEQ_FE | nucleotide phosphate-binding region:ATP | 23 | 3,24 | 9,33E-01 | PRKCA, RAD51B, STK10, FLT4, HKDC1, TDRD9, ACACA, DQX1, RPS6KC1, UBA6, ACLY, SMC2, SRC, DNAH5, PAK6, RPS6KA3, TAP1, ABCD2, GRK5, PAK1, EHD1, PAPSS2, MYH10 | 544 | 963 | 18012 | 0,79 | 1,00E+00 | 1,00E+00 | 1,00E+02 |

| | | | | | | | | | | | | |
|---|---|---|---|---|---|---|---|---|---|---|---|---|
| GOTERM_M | GO:0004672~protein kinase activity | 13 | 1,83 | 9,37E-01 | PRKCA, POMK, FLT4, STK10, RPS6KC1, TRIM24, SRC, PAK6, RPS6KA3, MMD2, MLKL, PAK1, GRK5 | 575 | 531 | 17446 | 0,74 | 1,00E+00 | 1,00E+00 | 1,00E+02 |
| KEGG_PATH | mmu04810:Regulation of actin cytoskeleton | 4 | 0,56 | 9,59E-01 | ACTG1, PAK6, PAK1, SRC | 233 | 214 | 7720 | 0,62 | 1,00E+00 | 9,99E-01 | 1,00E+02 |
| INTERPRO | IPR000719:Protein kinase, catalytic domain | 11 | 1,55 | 9,63E-01 | PRKCA, PAK6, RPS6KA3, POMK, STK10, FLT4, RPS6KC1, MLKL, GRK5, PAK1, SRC | 645 | 515 | 20594 | 0,68 | 1,00E+00 | 1,00E+00 | 1,00E+02 |
| INTERPRO | IPR017441:Protein kinase, ATP binding site | 8 | 1,13 | 9,65E-01 | PRKCA, PAK6, RPS6KA3, STK10, FLT4, GRK5, PAK1, SRC | 645 | 394 | 20594 | 0,65 | 1,00E+00 | 1,00E+00 | 1,00E+02 |
| UP_KEYWO | Kinase | 14 | 1,97 | 9,80E-01 | PRKCA, POMK, FLT4, HKDC1, STK10, RPS6KC1, SRC, PAK6, RPS6KA3, CDKN1A, GRK5, PAK1, PAPSS2, AGK | 678 | 707 | 22680 | 0,66 | 1,00E+00 | 1,00E+00 | 1,00E+02 |
| GOTERM_B | GO:0016310~phosphorylation | 13 | 1,83 | 9,80E-01 | PRKCA, POMK, FLT4, HKDC1, STK10, RPS6KC1, SRC, PAK6, RPS6KA3, GRK5, PAK1, PAPSS2, AGK | 588 | 612 | 18082 | 0,65 | 1,00E+00 | 1,00E+00 | 1,00E+02 |
| INTERPRO | IPR011009:Protein kinase-like domain | 11 | 1,55 | 9,81E-01 | PRKCA, PAK6, RPS6KA3, POMK, STK10, FLT4, RPS6KC1, MLKL, GRK5, PAK1, SRC | 645 | 556 | 20594 | 0,63 | 1,00E+00 | 1,00E+00 | 1,00E+02 |
| GOTERM_B | GO:0006468~protein phosphorylation | 12 | 1,69 | 9,82E-01 | PRKCA, PAK6, RPS6KA3, POMK, STK10, FLT4, RPS6KC1, MLKL, GRK5, PAK1, TRIM24, SRC | 588 | 576 | 18082 | 0,64 | 1,00E+00 | 1,00E+00 | 1,00E+02 |
| UP_KEYWO | Serine/threonine-protein kinase | 7 | 0,99 | 9,83E-01 | PRKCA, PAK6, RPS6KA3, STK10, RPS6KC1, GRK5, PAK1 | 678 | 405 | 22680 | 0,58 | 1,00E+00 | 1,00E+00 | 1,00E+02 |
| UP_SEQ_FE | binding site:ATP | 11 | 1,55 | 9,83E-01 | PRKCA, PAK6, RPS6KA3, HKDC1, STK10, FLT4, RPS6KC1, GRK5, PAK1, EHD1, SRC | 544 | 583 | 18012 | 0,62 | 1,00E+00 | 1,00E+00 | 1,00E+02 |
| KEGG_PATH | mmu04010:MAPK signaling pathway | 4 | 0,56 | 9,84E-01 | PRKCA, RPS6KA3, PPP3CC, PAK1 | 233 | 253 | 7720 | 0,52 | 1,00E+00 | 1,00E+00 | 1,00E+02 |
| UP_SEQ_FE | domain:Protein kinase | 9 | 1,27 | 9,86E-01 | PRKCA, PAK6, POMK, STK10, FLT4, MLKL, GRK5, PAK1, SRC | 544 | 502 | 18012 | 0,59 | 1,00E+00 | 1,00E+00 | 1,00E+02 |
| GOTERM_M | GO:0016301~kinase activity | 13 | 1,83 | 9,94E-01 | PRKCA, POMK, FLT4, HKDC1, STK10, RPS6KC1, SRC, PAK6, RPS6KA3, GRK5, PAK1, PAPSS2, AGK | 575 | 674 | 17446 | 0,59 | 1,00E+00 | 1,00E+00 | 1,00E+02 |
| GOTERM_M | GO:0004674~protein serine/threonine kinase activity | 7 | 0,99 | 9,96E-01 | PRKCA, PAK6, RPS6KA3, STK10, RPS6KC1, GRK5, PAK1 | 575 | 428 | 17446 | 0,50 | 1,00E+00 | 1,00E+00 | 1,00E+02 |
| INTERPRO | IPR008271:Serine/threonine-protein kinase, active site | 4 | 0,56 | 9,98E-01 | PRKCA, RPS6KA3, STK10, PAK1 | 645 | 333 | 20594 | 0,38 | 1,00E+00 | 1,00E+00 | 1,00E+02 |
| SMART | SM00220:S_TKc | 6 | 0,85 | 9,98E-01 | PRKCA, RPS6KA3, STK10, RPS6KC1, GRK5, PAK1 | 379 | 380 | 10425 | 0,43 | 1,00E+00 | 1,00E+00 | 1,00E+02 |
| Annotation | Enrichment Score: 0.10154330132085404 | | | | | | | | | | | |
| Category | Term | Count | % | PValue | Genes | List To | Pop Hit | Pop Tot | Fold Enr | Bonferroni | Benjamini | FDR |

| | | | | | | | | | | | | |
|---|---|---|---|---|---|---|---|---|---|---|---|---|
| UP_SEQ_FE | lipid moiety-binding region:S-geranylgeranyl cysteine | 4 | 0,56 | 6,49E-01 | RASL10B, RHOC, GBP2, RAB27A | 544 | 110 | 18012 | 1,20 | 1,00E+00 | 1,00E+00 | 1,00E+02 |
| UP_KEYWO | Prenylation | 5 | 0,71 | 6,93E-01 | RASL10B, RHOC, CNP, GBP2, RAB27A | 678 | 157 | 22680 | 1,07 | 1,00E+00 | 9,63E-01 | 1,00E+02 |
| GOTERM_BI | GO:0007264~small GTPase mediated signal transduction | 7 | 0,99 | 7,82E-01 | RABL2, KLK1B4, RASL10B, RAPGEF5, RHOC, ARHGEF9, RAB27A | 588 | 236 | 18082 | 0,91 | 1,00E+00 | 1,00E+00 | 1,00E+02 |
| INTERPRO | IPR001806:Small GTPase superfamily | 4 | 0,56 | 7,94E-01 | RABL2, RASL10B, RHOC, RAB27A | 645 | 134 | 20594 | 0,95 | 1,00E+00 | 1,00E+00 | 1,00E+02 |
| UP_SEQ_FE | short sequence motif:Effector region | 3 | 0,42 | 8,08E-01 | RASL10B, RHOC, RAB27A | 544 | 100 | 18012 | 0,99 | 1,00E+00 | 1,00E+00 | 1,00E+02 |
| INTERPRO | IPR005225:Small GTP-binding protein domain | 4 | 0,56 | 8,93E-01 | RABL2, RASL10B, RHOC, RAB27A | 645 | 165 | 20594 | 0,77 | 1,00E+00 | 1,00E+00 | 1,00E+02 |
| UP_SEQ_FE | nucleotide phosphate-binding region:GTP | 6 | 0,85 | 9,66E-01 | RASL10B, RHOC, PCK2, GBP3, GBP2, RAB27A | 544 | 319 | 18012 | 0,62 | 1,00E+00 | 1,00E+00 | 1,00E+02 |
| Annotation | Enrichment Score: 0.10127193184608259 | | | | | | | | | | | |
| Category | Term | Count | % | PValue | Genes | List To | Pop Hit | Pop Tota | Fold Enr | Bonferroni | Benjamini | FDR |
| INTERPRO | IPR011598:Myc-type, basic helix-loop-helix (bHLH) domain | 4 | 0,56 | 6,91E-01 | TCFL5, ID2, MGA, MYCL | 645 | 113 | 20594 | 1,13 | 1,00E+00 | 1,00E+00 | 1,00E+02 |
| SMART | SM00353:HLH | 4 | 0,56 | 7,67E-01 | TCFL5, ID2, MGA, MYCL | 379 | 110 | 10425 | 1,00 | 1,00E+00 | 9,99E-01 | 1,00E+02 |
| UP_SEQ_FE | domain:Helix-loop-helix motif | 3 | 0,42 | 8,48E-01 | ID2, MGA, MYCL | 544 | 110 | 18012 | 0,90 | 1,00E+00 | 1,00E+00 | 1,00E+02 |
| GOTERM_M | GO:0046983~protein dimerization activity | 5 | 0,71 | 8,75E-01 | THRA, ID2, NUP210, MGA, MYCL | 575 | 190 | 17446 | 0,80 | 1,00E+00 | 1,00E+00 | 1,00E+02 |
| Annotation | Enrichment Score: 0.09504255470504386 | | | | | | | | | | | |
| Category | Term | Count | % | PValue | Genes | List To | Pop Hit | Pop Tota | Fold Enr | Bonferroni | Benjamini | FDR |
| KEGG_PATH | mmu04080:Neuroactive ligand-receptor interaction | 13 | 1,83 | 1,50E-01 | P2RX5, P2RY13, SSTR2, GPR35, THRA, PRLR, P2RY4, LEPR, MAS1, CALCRL, GABBR2, ADORA1, GRID1 | 233 | 285 | 7720 | 1,51 | 1,00E+00 | 8,07E-01 | 8,77E+01 |
| GOTERM_M | GO:0004871~signal transducer activity | 16 | 2,26 | 9,43E-01 | MRGPRB2, LGALS1, GABBR2, ECT2, ADORA1, P2RY13, SMO, SSTR2, GPR35, P2RY4, CCR5, CCR2, MAS1, SUCNR1, RHOC, CALCRL | 575 | 648 | 17446 | 0,75 | 1,00E+00 | 1,00E+00 | 1,00E+02 |

| | | | | | | | | | | | | |
|---|---|---|---|---|---|---|---|---|---|---|---|---|
| GOTERM_BI | GO:0007165~signal transduction | 29 | 4,09 | 9,89E-01 | LEPR, TLR3, GABBR2, TLR7, ADORA1, TLR8, TNFRSF11B, RGS12, ANK2, P2RY4, UNC5B, RASL10B, MAS1, SUCNR1, CALCRL, MRGPRB2, DAB2IP, PLXNB1, RPS6KC1, SMO, P2RY13, GPR35, SSTR2, CCR5, CD34, RGS5, CCR2, PTCH1, GRK5 | 588 | 1255 | 18082 | 0,71 | 1,00E+00 | 1,00E+00 | 1,00E+02 |
| GOTERM_BI | GO:0007186~G-protein coupled receptor signaling pathway | 17 | 2,4 | 1,00E+00 | MRGPRB2, CCL2, GABBR2, ADORA1, CXCL10, P2RY13, SMO, SSTR2, GPR35, CCR5, P2RY4, CCR2, RGS5, MAS1, OLFR1033, SUCNR1, CALCRL | 588 | 1706 | 18082 | 0,31 | 1,00E+00 | 1,00E+00 | 1,00E+02 |
| INTERPRO | IPR000276:G protein-coupled receptor, rhodopsin-like | 11 | 1,55 | 1,00E+00 | MRGPRB2, P2RY13, SSTR2, GPR35, CCR5, P2RY4, CCR2, MAS1, OLFR1033, SUCNR1, ADORA1 | 645 | 1458 | 20594 | 0,24 | 1,00E+00 | 1,00E+00 | 1,00E+02 |
| UP_KEYWO | G-protein coupled receptor | 14 | 1,97 | 1,00E+00 | MRGPRB2, GABBR2, ADORA1, P2RY13, SMO, SSTR2, GPR35, P2RY4, CCR5, CCR2, MAS1, OLFR1033, SUCNR1, CALCRL | 678 | 1741 | 22680 | 0,27 | 1,00E+00 | 1,00E+00 | 1,00E+02 |
| UP_KEYWO | Transducer | 14 | 1,97 | 1,00E+00 | MRGPRB2, GABBR2, ADORA1, P2RY13, SMO, SSTR2, GPR35, P2RY4, CCR5, CCR2, MAS1, OLFR1033, SUCNR1, CALCRL | 678 | 1801 | 22680 | 0,26 | 1,00E+00 | 1,00E+00 | 1,00E+02 |
| GOTERM_M | GO:0004930~G-protein coupled receptor activity | 14 | 1,97 | 1,00E+00 | MRGPRB2, GABBR2, ADORA1, P2RY13, SMO, SSTR2, GPR35, P2RY4, CCR5, CCR2, MAS1, OLFR1033, SUCNR1, CALCRL | 575 | 1749 | 17446 | 0,24 | 1,00E+00 | 1,00E+00 | 1,00E+02 |
| INTERPRO | IPR017452:GPCR, rhodopsin-like, 7TM | 11 | 1,55 | 1,00E+00 | MRGPRB2, P2RY13, SSTR2, GPR35, CCR5, P2RY4, CCR2, MAS1, OLFR1033, SUCNR1, ADORA1 | 645 | 1708 | 20594 | 0,21 | 1,00E+00 | 1,00E+00 | 1,00E+02 |
| Annotation | Enrichment Score: 0.07570602409214529 | | | | | | | | | | | |
| Category | Term | Count | % | PValue | Genes | List To | Pop Hit | Pop Tot | Fold Enr | Bonferroni | Benjamini | FDR |
| UP_SEQ_FE | domain:SH2 | 3 | 0,42 | 7,75E-01 | STAP1, SRC, BLNK | 544 | 93 | 18012 | 1,07 | 1,00E+00 | 1,00E+00 | 1,00E+02 |
| UP_KEYWO | SH2 domain | 3 | 0,42 | 8,25E-01 | STAP1, SRC, BLNK | 678 | 105 | 22680 | 0,96 | 1,00E+00 | 9,85E-01 | 1,00E+02 |
| INTERPRO | IPR000980:SH2 domain | 3 | 0,42 | 8,72E-01 | STAP1, SRC, BLNK | 645 | 113 | 20594 | 0,85 | 1,00E+00 | 1,00E+00 | 1,00E+02 |
| SMART | SM00252:SH2 | 3 | 0,42 | 8,93E-01 | STAP1, SRC, BLNK | 379 | 103 | 10425 | 0,80 | 1,00E+00 | 1,00E+00 | 1,00E+02 |
| Annotation | Enrichment Score: 0.07450638584228551 | | | | | | | | | | | |
| Category | Term | Count | % | PValue | Genes | List To | Pop Hit | Pop Tot | Fold Enr | Bonferroni | Benjamini | FDR |
| GOTERM_C | GO:0030496~midbody | 6 | 0,85 | 4,07E-01 | MAP10, KLHL13, PTCH1, ECT2, MYH10, ANXA2 | 633 | 130 | 19662 | 1,43 | 1,00E+00 | 9,35E-01 | 9,99E+01 |
| UP_KEYWO | Cell cycle | 16 | 2,26 | 8,41E-01 | EID1, DAB2IP, STK10, ECT2, SMC2, SRC, TRIM21, AJUBA, RPS6KA3, MAP10, CDKN1A, CDKN2C, KLHL13, STAG3, CABLES1, STAG1 | 678 | 626 | 22680 | 0,85 | 1,00E+00 | 9,88E-01 | 1,00E+02 |
| GOTERM_BI | GO:0007049~cell cycle | 16 | 2,26 | 9,00E-01 | EID1, DAB2IP, STK10, ECT2, SMC2, SRC, TRIM21, AJUBA, RPS6KA3, MAP10, CDKN1A, CDKN2C, KLHL13, STAG3, CABLES1, STAG1 | 588 | 614 | 18082 | 0,80 | 1,00E+00 | 1,00E+00 | 1,00E+02 |
| UP_KEYWO | Cell division | 6 | 0,85 | 9,87E-01 | MAP10, KLHL13, ECT2, SMC2, CABLES1, STAG1 | 678 | 372 | 22680 | 0,54 | 1,00E+00 | 1,00E+00 | 1,00E+02 |
| GOTERM_BI | GO:0051301~cell division | 6 | 0,85 | 9,94E-01 | MAP10, KLHL13, ECT2, SMC2, CABLES1, STAG1 | 588 | 374 | 18082 | 0,49 | 1,00E+00 | 1,00E+00 | 1,00E+02 |

| | | | | | | | | | | | | |
|---|---|---|---|---|---|---|---|---|---|---|---|---|
| UP_KEYWO | Mitosis | 3 | 0,42 | 9,97E-01 | KLHL13, SMC2, STAG1 | 678 | 259 | 22680 | 0,39 | 1,00E+00 | 1,00E+00 | 1,00E+02 |
| GOTERM_B | GO:0007067~mitotic nuclear division | 3 | 0,42 | 9,99E-01 | KLHL13, SMC2, STAG1 | 588 | 277 | 18082 | 0,33 | 1,00E+00 | 1,00E+00 | 1,00E+02 |
| Annotation | Enrichment Score: 0.06370585421243077 | | | | | | | | | | | |
| Category | Term | Count | % | PValue | Genes | List To | Pop Hit | Pop Tota | Fold Enr | Bonferroni | Benjamini | FDR |
| UP_KEYWO | Ion channel | 9 | 1,27 | 7,89E-01 | TRPM4, P2RX5, FXYD3, CLIC5, SCN4B, CNGA3, FXYD5, KCNE3, GRID1 | 678 | 336 | 22680 | 0,90 | 1,00E+00 | 9,78E-01 | 1,00E+02 |
| GOTERM_M | GO:0005244~voltage-gated ion channel activity | 4 | 0,56 | 8,22E-01 | TRPM4, CLIC5, SCN4B, KCNE3 | 575 | 134 | 17446 | 0,91 | 1,00E+00 | 1,00E+00 | 1,00E+02 |
| UP_KEYWO | Voltage-gated channel | 3 | 0,42 | 9,17E-01 | CLIC5, SCN4B, KCNE3 | 678 | 136 | 22680 | 0,74 | 1,00E+00 | 9,96E-01 | 1,00E+02 |
| GOTERM_B | GO:0034765~regulation of ion transmembrane transport | 3 | 0,42 | 9,36E-01 | CLIC5, SCN4B, KCNE3 | 588 | 135 | 18082 | 0,68 | 1,00E+00 | 1,00E+00 | 1,00E+02 |
| Annotation | Enrichment Score: 0.04775136478907559 | | | | | | | | | | | |
| Category | Term | Count | % | PValue | Genes | List To | Pop Hit | Pop Tota | Fold Enr | Bonferroni | Benjamini | FDR |
| UP_KEYWO | SH3 domain | 5 | 0,71 | 8,72E-01 | MYO7B, ARHGEF16, ARHGEF9, SRC, NPHP1 | 678 | 208 | 22680 | 0,80 | 1,00E+00 | 9,92E-01 | 1,00E+02 |
| UP_SEQ_FE | domain:SH3 | 4 | 0,56 | 8,81E-01 | ARHGEF16, ARHGEF9, SRC, NPHP1 | 544 | 166 | 18012 | 0,80 | 1,00E+00 | 1,00E+00 | 1,00E+02 |
| INTERPRO | IPR001452:Src homology-3 domain | 5 | 0,71 | 9,02E-01 | MYO7B, ARHGEF16, ARHGEF9, SRC, NPHP1 | 645 | 212 | 20594 | 0,75 | 1,00E+00 | 1,00E+00 | 1,00E+02 |
| SMART | SM00326:SH3 | 5 | 0,71 | 9,31E-01 | MYO7B, ARHGEF16, ARHGEF9, SRC, NPHP1 | 379 | 197 | 10425 | 0,70 | 1,00E+00 | 1,00E+00 | 1,00E+02 |
| Annotation | Enrichment Score: 0.038184078192659524 | | | | | | | | | | | |
| Category | Term | Count | % | PValue | Genes | List To | Pop Hit | Pop Tota | Fold Enr | Bonferroni | Benjamini | FDR |
| UP_SEQ_FE | repeat:ANK 5 | 3 | 0,42 | 8,41E-01 | ANK2, CDKN2C, ANKRD1 | 544 | 108 | 18012 | 0,92 | 1,00E+00 | 1,00E+00 | 1,00E+02 |
| UP_SEQ_FE | repeat:ANK 3 | 4 | 0,56 | 8,65E-01 | ANK2, CDKN2C, ANKRD1, CLIP4 | 544 | 160 | 18012 | 0,83 | 1,00E+00 | 1,00E+00 | 1,00E+02 |
| UP_SEQ_FE | repeat:ANK 4 | 3 | 0,42 | 8,99E-01 | ANK2, CDKN2C, ANKRD1 | 544 | 127 | 18012 | 0,78 | 1,00E+00 | 1,00E+00 | 1,00E+02 |
| UP_KEYWO | ANK repeat | 5 | 0,71 | 9,30E-01 | ANK2, CDKN2C, GLS, ANKRD1, CLIP4 | 678 | 240 | 22680 | 0,70 | 1,00E+00 | 9,97E-01 | 1,00E+02 |
| UP_SEQ_FE | repeat:ANK 1 | 4 | 0,56 | 9,33E-01 | ANK2, CDKN2C, ANKRD1, CLIP4 | 544 | 193 | 18012 | 0,69 | 1,00E+00 | 1,00E+00 | 1,00E+02 |
| UP_SEQ_FE | repeat:ANK 2 | 4 | 0,56 | 9,33E-01 | ANK2, CDKN2C, ANKRD1, CLIP4 | 544 | 193 | 18012 | 0,69 | 1,00E+00 | 1,00E+00 | 1,00E+02 |
| INTERPRO | IPR002110:Ankyrin repeat | 5 | 0,71 | 9,35E-01 | ANK2, CDKN2C, GLS, ANKRD1, CLIP4 | 645 | 232 | 20594 | 0,69 | 1,00E+00 | 1,00E+00 | 1,00E+02 |
| INTERPRO | IPR020683:Ankyrin repeat-containing domain | 5 | 0,71 | 9,47E-01 | ANK2, CDKN2C, GLS, ANKRD1, CLIP4 | 645 | 242 | 20594 | 0,66 | 1,00E+00 | 1,00E+00 | 1,00E+02 |

| | | | | | | | | | | | | |
|---|---|---|---|---|---|---|---|---|---|---|---|---|
| SMART | SM00248:ANK | 5 | 0,71 | 9,66E-01 | ANK2, CDKN2C, GLS, ANKRD1, CLIP4 | 379 | 225 | 10425 | 0,61 | 1,00E+00 | 1,00E+00 | 1,00E+02 |
| Annotation | Enrichment Score: 0.03002447164335894 | | | | | | | | | | | |
| Category | Term | Count | % | PValue | Genes | List To | Pop Hit | Pop Tota | Fold Enr | Bonferroni | Benjamini | FDR |
| INTERPRO | IPR015943:WD40/YVTN repeat-like-containing domain | 11 | 1,55 | 6,23E-01 | BRWD3, DCAF4, FBXW16, PLXNB1, FBXW19, SEMA3D, DCAF12L1, DNAIC1, NBEAL1, SEMA4A, BC068281 | 645 | 340 | 20594 | 1,03 | 1,00E+00 | 1,00E+00 | 1,00E+02 |
| INTERPRO | IPR017986:WD40-repeat-containing domain | 6 | 0,85 | 9,65E-01 | BRWD3, DCAF4, FBXW16, DCAF12L1, DNAIC1, NBEAL1 | 645 | 307 | 20594 | 0,62 | 1,00E+00 | 1,00E+00 | 1,00E+02 |
| INTERPRO | IPR001680:WD40 repeat | 5 | 0,71 | 9,66E-01 | BRWD3, DCAF4, DCAF12L1, DNAIC1, NBEAL1 | 645 | 263 | 20594 | 0,61 | 1,00E+00 | 1,00E+00 | 1,00E+02 |
| UP_SEQ_FE | repeat:WD 1 | 4 | 0,56 | 9,81E-01 | BRWD3, DCAF12L1, DNAIC1, BC068281 | 544 | 248 | 18012 | 0,53 | 1,00E+00 | 1,00E+00 | 1,00E+02 |
| UP_SEQ_FE | repeat:WD 2 | 4 | 0,56 | 9,81E-01 | BRWD3, DCAF12L1, DNAIC1, BC068281 | 544 | 248 | 18012 | 0,53 | 1,00E+00 | 1,00E+00 | 1,00E+02 |
| UP_KEYWO | WD repeat | 4 | 0,56 | 9,83E-01 | BRWD3, DCAF12L1, DNAIC1, BC068281 | 678 | 256 | 22680 | 0,52 | 1,00E+00 | 1,00E+00 | 1,00E+02 |
| SMART | SM00320:WD40 | 5 | 0,71 | 9,87E-01 | BRWD3, DCAF4, DCAF12L1, DNAIC1, NBEAL1 | 379 | 262 | 10425 | 0,52 | 1,00E+00 | 1,00E+00 | 1,00E+02 |
| UP_SEQ_FE | repeat:WD 4 | 3 | 0,42 | 9,93E-01 | BRWD3, DCAF12L1, DNAIC1 | 544 | 230 | 18012 | 0,43 | 1,00E+00 | 1,00E+00 | 1,00E+02 |
| UP_SEQ_FE | repeat:WD 3 | 3 | 0,42 | 9,95E-01 | BRWD3, DCAF12L1, DNAIC1 | 544 | 243 | 18012 | 0,41 | 1,00E+00 | 1,00E+00 | 1,00E+02 |
| Annotation | Enrichment Score: 0.019131714822143986 | | | | | | | | | | | |
| Category | Term | Count | % | PValue | Genes | List To | Pop Hit | Pop Tota | Fold Enr | Bonferroni | Benjamini | FDR |
| UP_KEYWO | DNA-binding | 41 | 5,78 | 9,02E-01 | TAF1B, ZMAT1, ZFP13, RAD51B, THRA, POT1A, POLA1, HR, MYEF2, H1FX, ZKSCAN3, RPA1, TRIM30A, FOXQ1, ZFP667, ZBP1, ZFP518A, PLAG1, KLF6, RHOX5, ZFP426, RFX5, ESR1, MEIOB, TRIM24, EXO5, ZFP445, MYCL, MBD1, TRIM21, ZFP26, FOXP2, ADNP2, ZFP120, ZFP770, MEOX1, MGA, ZIK1, ATXN1L, ZFP369, PARP2 | 678 | 1604 | 22680 | 0,86 | 1,00E+00 | 9,95E-01 | 1,00E+02 |
| GOTERM_M | GO:0003677~DNA binding | 51 | 7,19 | 9,43E-01 | ZMAT1, TAF1B, XRCC4, GM14124, RAD51B, ZFP13, THRA, POT1A, HR, H1FX, ANKRD1, ZKSCAN3, TLR8, RNF141, DNTT, OASL1, ZFP518A, ZFP426, ESR1, MEIOB, MYCL, MBD1, SP140, ZFP26, ZFP120, TCFL5, MGA, ATXN1L, ZFP369, POLA1, RPA1, TRIM30A, FOXQ1, ZFP667, ZBP1, PLAG1, KLF6, RHOX5, CR2, RFX5, ZMYM6, TRIM24, ZFP445, EXO5, TRIM21, FOXP2, ADNP2, ZFP770, MEOX1, ZIK1, PARP2 | 575 | 1847 | 17446 | 0,84 | 1,00E+00 | 1,00E+00 | 1,00E+02 |
| UP_KEYWO | Transcription | 44 | 6,21 | 9,71E-01 | TAF1B, ZKSCAN7, DPF3, EID1, THRA, ZKSCAN8, HR, MYEF2, TCEAL8, IL33, ZKSCAN3, ZKSCAN4, TRIM30A, FOXQ1, LBH, ZFP667, ZFP518A, PLAG1, KLF6, ZFP426, LRIF1, RFX5, TAF7, ESR1, ZFP449, TRIM24, ZFP445, MBD1, CSRP3, FOXP2, ZFP809, AJUBA, ADNP2, BTG2, ZFP120, ZFP770, ID2, MEOX1, NAB2, MGA, ZIK1, ATXN1L, ZFP369, IFI205 | 678 | 1859 | 22680 | 0,79 | 1,00E+00 | 9,99E-01 | 1,00E+02 |

| | | | | | | | | | | | | |
|---|---|---|---|---|---|---|---|---|---|---|---|---|
| UP_KEYWO | Transcription regulation | 42 | 5,92 | 9,75E-01 | TAF1B, ZKSCAN7, DPF3, EID1, THRA, ZKSCAN8, HR, TCEAL8, ZKSCAN3, ZKSCAN4, TRIM30A, FOXQ1, LBH, ZFP667, ZFP518A, PLAG1, KLF6, ZFP426, LRIF1, RFX5, TAF7, ESR1, ZFP449, TRIM24, ZFP445, MBD1, CSRP3, FOXP2, ZFP809, AJUBA, ADNP2, BTG2, ZFP120, ZFP770, ID2, MEOX1, NAB2, MGA, ZIK1, ATXN1L, ZFP369, IFI205 | 678 | 1799 | 22680 | 0,78 | 1,00E+00 | 1,00E+00 | 1,00E+02 |
| GOTERM_B | GO:0006351~transcription, DNA-templated | 44 | 6,21 | 9,96E-01 | TAF1B, ZKSCAN7, DPF3, EID1, THRA, ZKSCAN8, HR, MYEF2, TCEAL8, IL33, ZKSCAN3, ZKSCAN4, TRIM30A, FOXQ1, LBH, ZFP667, ZFP518A, PLAG1, KLF6, ZFP426, LRIF1, RFX5, TAF7, ESR1, ZFP449, TRIM24, ZFP445, MBD1, FOXP2, ZFP809, AJUBA, ADNP2, BTG2, ZFP120, ZFP770, ID2, MEOX1, NAB2, MGA, ZIK1, ATXN1L, ZSCAN18, ZFP369, IFI205 | 588 | 1885 | 18082 | 0,72 | 1,00E+00 | 1,00E+00 | 1,00E+02 |
| Annotation | Enrichment Score: 0.0071458693713676805 | | | | | | | | | | | |
| Category | Term | Count | % | PValue | Genes | List To | Pop Hit | Pop Tota | Fold Enr | Bonferroni | Benjamini | FDR |
| UP_KEYWO | Transit peptide | 10 | 1,41 | 9,70E-01 | PARS2, ALDH1B1, MICU2, OXCT1, GLS, LIPT2, ACAD9, PCK2, ACSS3, AGK | 678 | 510 | 22680 | 0,66 | 1,00E+00 | 9,99E-01 | 1,00E+02 |
| UP_SEQ_FE | transit peptide:Mitochondrion | 9 | 1,27 | 9,84E-01 | PRLR, PARS2, ALDH1B1, OXCT1, LIPT2, ACAD9, PCK2, ACSS3, AGK | 544 | 496 | 18012 | 0,60 | 1,00E+00 | 1,00E+00 | 1,00E+02 |
| UP_KEYWO | Mitochondrion | 19 | 2,68 | 9,97E-01 | PRKCA, SLC25A4, STAP1, MICU2, PCK2, ACSS3, SRC, GM4952, ALDH1B1, PARS2, UCP2, PPL, GLS, OXCT1, HEBP2, OAS1A, LIPT2, ACAD9, AGK | 678 | 1058 | 22680 | 0,60 | 1,00E+00 | 1,00E+00 | 1,00E+02 |
| Annotation | Enrichment Score: 0.002145056574423231 | | | | | | | | | | | |
| Category | Term | Count | % | PValue | Genes | List To | Pop Hit | Pop Tota | Fold Enr | Bonferroni | Benjamini | FDR |
| INTERPRO | IPR000210:BTB/POZ-like | 3 | 0,42 | 9,93E-01 | KLHL13, BTBD8, KCTD12 | 645 | 222 | 20594 | 0,43 | 1,00E+00 | 1,00E+00 | 1,00E+02 |
| INTERPRO | IPR011333:BTB/POZ fold | 3 | 0,42 | 9,95E-01 | KLHL13, BTBD8, KCTD12 | 645 | 232 | 20594 | 0,41 | 1,00E+00 | 1,00E+00 | 1,00E+02 |
| SMART | SM00225:BTB | 3 | 0,42 | 9,97E-01 | KLHL13, BTBD8, KCTD12 | 379 | 218 | 10425 | 0,38 | 1,00E+00 | 1,00E+00 | 1,00E+02 |
| Annotation | Enrichment Score: 4.708111564534914E-4 | | | | | | | | | | | |
| Category | Term | Count | % | PValue | Genes | List To | Pop Hit | Pop Tota | Fold Enr | Bonferroni | Benjamini | FDR |
| UP_KEYWO | Homeobox | 3 | 0,42 | 9,98E-01 | ADNP2, RHOX5, MEOX1 | 678 | 280 | 22680 | 0,36 | 1,00E+00 | 1,00E+00 | 1,00E+02 |
| INTERPRO | IPR001356:Homeodomain | 3 | 0,42 | 9,98E-01 | ADNP2, RHOX5, MEOX1 | 645 | 271 | 20594 | 0,35 | 1,00E+00 | 1,00E+00 | 1,00E+02 |
| SMART | SM00389:HOX | 3 | 0,42 | 9,99E-01 | ADNP2, RHOX5, MEOX1 | 379 | 266 | 10425 | 0,31 | 1,00E+00 | 1,00E+00 | 1,00E+02 |
| INTERPRO | IPR009057:Homeodomain-like | 3 | 0,42 | 1,00E+00 | ADNP2, RHOX5, MEOX1 | 645 | 345 | 20594 | 0,28 | 1,00E+00 | 1,00E+00 | 1,00E+02 |
