## Supplementary material for "The mouse HP1 proteins are essential for preventing liver tumorigenesis": suppl table3

**Supplementary Table 3** : functional clustering of genes down-regulated upon loss of HP1 witin hepatocytes (https://david.ncifcrf.g

| Annotation | Enrichment Score: 11.14831199606435 | | | | | | | | | | | |
|---|---|---|---|---|---|---|---|---|---|---|---|---|
| Category | Term | Count | % | PValue | Genes | List Total | Pop Hits | Pop Total | Enrichment | Bonferroni | Benjamini | FDR |
| GOTERM_BP_DIRECT | GO:0055114~oxidation-reduction process | 49 | 10,17 | 4,72E-13 | NDUFB3, TM7SF2, PHYHD1, CYP3A25, CYP2F2, HSD17B2, TMX4, CYC1, HSD3B5, UGDH, AASS, PAH, AKR1C13, GPX1, MTHFD2, CRYL1, CYP39A1, FXN, AKR1A1, CREG1, DUS1L, MTO1, GLRX, HPD, NOX4, NDUFA4, CYP46A1, DHRS13, MAOA, FADS1, MICAL2, CYP2C29, HGD, CYP2E1, CYB5B, CYP1A2, PHYH, NDUFA11, NDUFV3, MARC1, CYP27A1, UOX, AOX1, TXNRD3, CYP2D26, CYP3A59, H2-KE6, CYP2R1, GULO | 394 | 676 | 18082 | 3,326593 | 8,58E-10 | 8,58E-10 | 8,02E-10 |
| UP_KEYWORDS | Oxidoreductase | 43 | 8,921 | 1,76E-11 | TM7SF2, PHYHD1, CYP3A25, CYP2C44, HSD17B2, CYP2F2, HSD3B5, UGDH, AASS, PAH, AKR1C13, MTHFD2, GPX1, CRYL1, CYP39A1, FXN, AKR1A1, DUS1L, HPD, NOX4, CYP46A1, CYP2D40, DHRS13, MAOA, MICAL2, FADS1, CYP2C29, CYP4F13, HGD, CYP1A2, CYP2E1, PHYH, MARC1, CYP27A1, UOX, AOX1, CYP2A5, TXNRD3, CYP3A59, CYP2D26, H2-KE6, CYP2R1, GULO | 456 | 639 | 22680 | 3,346924 | 5,62E-09 | 5,62E-09 | 2,38E-08 |
| GOTERM_MF_DIRECT | GO:0016491~oxidoreductase activity | 43 | 8,921 | 4,31E-11 | TM7SF2, PHYHD1, CYP3A25, HSD17B2, CYP2F2, HSD3B5, UGDH, AASS, PAH, OSGIN1, AKR1C13, DCT, MTHFD2, GPX1, CRYL1, CYP39A1, FXN, AKR1A1, CREG1, DUS1L, MTO1, HPD, NOX4, CYP46A1, DHRS13, MAOA, MICAL2, FADS1, CYP2C29, HGD, CYP1A2, CYP2E1, PHYH, MARC1, CYP27A1, UOX, AKR1C19, AOX1, TXNRD3, CYP2D26, H2-KE6, CYP2R1, GULO | 385 | 604 | 17446 | 3,226017 | 2,72E-08 | 1,36E-08 | 6,42E-08 |
| Annotation | Enrichment Score: 9.452175437202708 | | | | | | | | | | | |
| Category | Term | Count | % | PValue | Genes | List Total | Pop Hits | Pop Total | Enrichment | Bonferroni | Benjamini | FDR |

| | | | | | | | | | | | | |
|---|---|---|---|---|---|---|---|---|---|---|---|---|
| GOTERM_CC_DIRECT | GO:0005783~endoplasmic reticulum | 68 | 14,11 | 4,37E-11 | TM7SF2, VAPB, PDE3B, HP, ILDR2, CSPG5, AI314180, TMEM147, POMT2, CYP39A1, H13, FITM1, ELOVL3, RPL10, CES1E, PTDSS1, HPD, EGFR, F10, CYP2C29, CYP2E1, CYP1A2, UGT1A1, ZDHHC14, DGAT1, ZFAND2B, EMC6, TXNRD3, CYP2D26, EEF1G, CTSC, GULO, SLC27A2, CYP3A25, CYP2F2, TMX4, HSD3B5, DTNBP1, TMCC1, UGT1A9, PLA2G12A, SERPINA1C, PDILT, AATK, NOX4, REEP6, EBP, H2-Q10, CYP46A1, MAP2K2, FADS1, CES3A, EPHX1, ATP1A1, MARCH5, ABCB6, ITPR1, ITPR2, MARCH2, GBA2, VCP, SLC35G1, PTP4A1, UGT2B5, CYP2R1, RAB38, CNIH1, SSR4 | 424 | 1323 | 19662 | 2,383477 | 1,56E-08 | 7,82E-09 | 6,00E-08 |
| UP_KEYWORDS | Endoplasmic reticulum | 52 | 10,79 | 9,53E-10 | TM7SF2, VAPB, ILDR2, CSPG5, AI314180, TMEM147, POMT2, CYP39A1, H13, FITM1, ELOVL3, CES1E, PTDSS1, EGFR, CYP2C29, CYP1A2, CYP2E1, UGT1A1, DGAT1, ZFAND2B, TXNRD3, CYP2A5, CYP2D26, GULO, SLC27A2, CYP3A25, CYP2F2, HSD3B5, DTNBP1, TMCC1, UGT1A9, PDILT, NOX4, REEP6, EBP, CYP46A1, FADS1, CES3A, EPHX1, MARCH5, ABCB6, ITPR1, ITPR2, MARCH2, GBA2, SLC35G1, VCP, PTP4A1, UGT2B5, CYP2R1, CNIH1, SSR4 | 456 | 997 | 22680 | 2,594098 | 3,04E-07 | 6,08E-08 | 1,28E-06 |
| GOTERM_CC_DIRECT | GO:0005789~endoplasmic reticulum membrane | 44 | 9,129 | 1,06E-09 | CYP3A25, CYP2F2, VAPB, HSD3B5, ILDR2, CSPG5, DTNBP1, TMCC1, POMT2, TMEM147, CYP39A1, H13, FITM1, ELOVL3, PTDSS1, HPD, NOX4, EGFR, REEP6, EBP, CYP46A1, FADS1, CYP2C29, EPHX1, CYP1A2, ABCB6, UGT1A1, MARCH5, ITPR1, ITPR2, MARCH2, GBA2, DGAT1, SLC35G1, VCP, CYP2A5, UGT2B5, CYP2D26, MOSPD3, CYP2R1, CNIH1, GULO, SSR4, SLC27A2 | 424 | 710 | 19662 | 2,873798 | 3,78E-07 | 1,26E-07 | 1,45E-06 |
| Annotation | Enrichment Score: 6.7754815908046035 | | | | | | | | | | | |
| Category | Term | Count | % | PValue | Genes | List Total | Pop Hits | Pop Total | Enrichment | Bonferroni | Benjamini | FDR |

| | | | | | | | | | | | | |
|---|---|---|---|---|---|---|---|---|---|---|---|---|
| GOTERM_MF_DIRECT | GO:0003735~structural constituent of ribosome | 28 | 5,809 | 3,48E-11 | RPL14, RPL35, RPL37, RPL38, RPS2, MRPL20, SLC25A23, RPLP0, SLC25A3, RPL10, MRPL18, SLC25A28, RPL11, RPS23, RPSA, MRPS24, RPL24, RPS6, RPS8, RPL28, RPS18, RPL41, RPS14, RPS15, SLC25A38, SLC25A37, RPS10, SLC25A51 | 385 | 264 | 17446 | 4,806061 | 2,19E-08 | 2,19E-08 | 5,17E-08 |
| UP_KEYWORDS | Ribosomal protein | 22 | 4,564 | 9,45E-10 | RPSA, RPL14, MRPS24, RPL35, RPL37, RPL24, RPL38, RPS6, RPS2, RPS8, RPL28, MRPL20, RPS18, RPL41, RPS14, RPLP0, RPS15, RPL10, MRPL18, RPL11, RPS10, RPS23 | 456 | 203 | 22680 | 5,3902 | 3,02E-07 | 7,54E-08 | 1,28E-06 |
| KEGG_PATHWAY | mmu03010:Ribosome | 21 | 4,357 | 1,05E-09 | RPSA, RPL14, RPL35, RPL37, RPL24, RPL38, RPS6, RPS2, RPS8, RPL28, MRPL20, RPS18, RPL41, RPS14, RPLP0, RPS15, RPL10, MRPL18, RPL11, RPS10, RPS23 | 205 | 145 | 7720 | 5,453995 | 2,41E-07 | 1,21E-07 | 1,35E-06 |
| GOTERM_BP_DIRECT | GO:0006412~translation | 32 | 6,639 | 1,07E-09 | RPL14, FARS2, EIF5, RPL35, RPL37, RPL38, RPS2, MRPL20, SLC25A23, SLC25A3, RPL10, MRPL18, SLC25A28, RPL11, RPS23, MARS, EGFR, RPSA, MRPS24, RPL24, RPS6, RPS8, RPL28, RPS18, RPL41, RPS14, RPS15, SLC25A38, SLC25A37, EEF1G, PELO, SLC25A51 | 394 | 401 | 18082 | 3,662316 | 1,95E-06 | 9,74E-07 | 1,82E-06 |
| GOTERM_CC_DIRECT | GO:0005840~ribosome | 20 | 4,149 | 2,60E-08 | RPSA, RPL14, MRPS24, RPL24, RPL38, RPS6, RPS2, RPS8, RPL28, MRPL20, RPS18, RPL41, RPS14, RPLP0, RPS15, RPL10, MRPL18, RPL11, RPS10, RPS23 | 424 | 188 | 19662 | 4,93326 | 9,31E-06 | 2,33E-06 | 3,57E-05 |
| UP_KEYWORDS | Ribonucleoprotein | 24 | 4,979 | 7,73E-08 | RPSA, RPL14, MRPS24, RPL35, RPL37, RPL24, RPL38, CPEB1, RPS6, RPS2, RPS8, RPL28, MRPL20, RPS18, RPL41, RPS14, RPLP0, RPS15, RPL10, MRPL18, RPL11, RPS10, NOP56, RPS23 | 456 | 308 | 22680 | 3,875598 | 2,46E-05 | 2,46E-06 | 1,04E-04 |
| GOTERM_CC_DIRECT | GO:0022627~cytosolic small ribosomal subunit | 9 | 1,867 | 1,63E-05 | RPSA, RPS18, RPS14, RPS15, RPS10, RPS6, RPS2, RPS23, RPS8 | 424 | 53 | 19662 | 7,8746 | 0,0058211 | 6,48E-04 | 0,0223758 |
| GOTERM_CC_DIRECT | GO:0030529~intracellular ribonucleoprotein complex | 21 | 4,357 | 2,22E-05 | PABPN1, RPSA, RPL14, MRPS24, RPL24, RPL38, CPEB1, RPS6, RPS2, RPS8, RPL28, MRPL20, RPS18, RPS14, RPLP0, RPS15, MRPL18, RPL11, RPS10, NOP56, RPS23 | 424 | 320 | 19662 | 3,043205 | 0,007904 | 7,21E-04 | 0,0304132 |
| GOTERM_CC_DIRECT | GO:0022625~cytosolic large ribosomal subunit | 9 | 1,867 | 7,69E-04 | RPL14, RPLP0, RPL35, RPL10, RPL11, RPL24, RPL37, RPL38, RPL28 | 424 | 91 | 19662 | 4,586305 | 0,2407108 | 0,0209597 | 1,0499972 |
| GOTERM_MF_DIRECT | GO:0044822~poly(A) RNA binding | 42 | 8,714 | 8,49E-04 | MRPL20, ZC3H8, ZFC3H1, BUD13, RPLP0, RPL10, RPL11, STRBP, MTO1, RPS23, PABPN1, CLNS1A, RPSA, H2-Q10, NOC4L, MRPS24, DNAJC21, RPL24, RPS6, RPL28, RPS8, NDUFV3, RPS18, RCC2, ZFP598, VCP, PPIA, SRSF6, NME1, RPS14, | 385 | 1113 | 17446 | 1,709973 | 0,4147419 | 0,0310206 | 1,2545762 |

| | | | | | | | | | | | | |
|---|---|---|---|---|---|---|---|---|---|---|---|---|
| Annotation | Enrichment Score: 6.006448397908353 | | | | | | | | | | | |
| Category | Term | Count | % | PValue | Genes | List Total | Pop Hits | Pop Total | Enrichment | Bonferroni | Benjamini | FDR |
| UP_KEYWORDS | Oxidoreductase | 43 | 8,921 | 1,76E-11 | TM7SF2, PHYHD1, CYP3A25, CYP2C44, HSD17B2, CYP2F2, HSD3B5, UGDH, AASS, PAH, AKR1C13, MTHFD2, GPX1, CRYL1, CYP39A1, FXN, AKR1A1, DUS1L, HPD, NOX4, CYP46A1, CYP2D40, DHRS13, MAOA, MICAL2, FADS1, CYP2C29, CYP4F13, HGD, CYP1A2, CYP2E1, PHYH, MARC1, CYP27A1, UOX, AOX1, CYP2A5, TXNRD3, CYP3A59, CYP2D26, H2-KE6, CYP2R1, GULO | 456 | 639 | 22680 | 3,346924 | 5,62E-09 | 5,62E-09 | 2,38E-08 |
| UP_KEYWORDS | Iron | 32 | 6,639 | 3,24E-11 | PHYHD1, CYP3A25, FTL1, CYP2C44, CYP2F2, CYC1, ACP5, PAH, COX5A, CYP39A1, FXN, NUBP1, SLC25A28, MOCS1, HPD, CYP46A1, CYP2D40, CYP2C29, CYP4F13, HGD, CYP2E1, CYB5B, CYP1A2, PHYH, ISCU, CYP27A1, AOX1, CYP2A5, CYP3A59, SLC25A37, CYP2D26, CYP2R1 | 456 | 375 | 22680 | 4,244211 | 1,03E-08 | 5,16E-09 | 4,36E-08 |
| INTERPRO | IPR001128:Cytochrome P450 | 16 | 3,32 | 5,02E-09 | CYP3A25, CYP2C44, CYP2F2, CYP46A1, CYP2D40, CYP2C29, CYP4F13, CYP2D37-PS, CYP1A2, CYP2E1, CYP39A1, CYP27A1, CYP2A5, CYP3A59, CYP2D26, CYP2R1 | 436 | 104 | 20594 | 7,266761 | 4,49E-06 | 1,12E-06 | 7,82E-06 |
| UP_KEYWORDS | Microsome | 17 | 3,527 | 9,91E-09 | TM7SF2, CYP3A25, CYP2F2, CYP46A1, CYP2C29, EPHX1, CYP1A2, CYP2E1, UGT1A1, UGT1A9, CYP39A1, UGT2B5, TXNRD3, CYP2A5, CYP2D26, CYP2R1, GULO | 456 | 132 | 22680 | 6,405502 | 3,16E-06 | 4,52E-07 | 1,34E-05 |
| UP_KEYWORDS | Monooxygenase | 17 | 3,527 | 1,11E-08 | CYP3A25, CYP2C44, CYP2F2, CYP46A1, CYP2D40, MICAL2, CYP2C29, CYP4F13, PAH, CYP1A2, CYP2E1, CYP39A1, CYP27A1, CYP2A5, CYP3A59, CYP2D26, CYP2R1 | 456 | 133 | 22680 | 6,357341 | 3,53E-06 | 4,41E-07 | 1,49E-05 |
| GOTERM_MF_DIRECT | GO:0016705~oxidoreductase activity, acting on paired donors, with incorporation or reduction of molecular oxygen | 15 | 3,112 | 3,45E-08 | CYP3A25, CYP2C44, CYP2F2, CYP46A1, CYP2D40, CYP2C29, CYP4F13, CYP2D37-PS, CYP1A2, CYP2E1, CYP39A1, CYP27A1, CYP2A5, CYP2D26, CYP2R1 | 385 | 99 | 17446 | 6,865801 | 2,18E-05 | 5,44E-06 | 5,13E-05 |
| GOTERM_MF_DIRECT | GO:0020037~heme binding | 19 | 3,942 | 6,33E-08 | CYP3A25, CYP2C44, CYP2F2, CYP46A1, CYP2D40, CYP2C29, CYC1, CYP4F13, CYP2D37-PS, CYB5B, CYP1A2, CYP2E1, ABCB6, CYP39A1, CYP27A1, CYP2A5, CYP3A59, CYP2D26, CYP2R1 | 385 | 175 | 17446 | 4,919837 | 4,00E-05 | 7,99E-06 | 9,42E-05 |

| | | | | | | | | | | | | |
|---|---|---|---|---|---|---|---|---|---|---|---|---|
| UP_KEYWORDS | Heme | 18 | 3,734 | 6,83E-08 | CYP3A25, CYP2C44, CYP2F2, CYP46A1, CYP2D40, CYP2C29, CYC1, CYP4F13, CYB5B, CYP1A2, CYP2E1, COX5A, CYP39A1, CYP27A1, CYP2A5, CYP3A59, CYP2D26, CYP2R1 | 456 | 171 | 22680 | 5,235457 | 2,18E-05 | 2,42E-06 | 9,21E-05 |
| GOTERM_MF_DIRECT | GO:0004497~monooxygenase activity | 15 | 3,112 | 1,06E-07 | CYP3A25, CYP2C44, CYP2F2, CYP46A1, CYP2D40, MICAL2, CYP2C29, CYP4F13, PAH, CYP1A2, CYP2E1, CYP39A1, CYP27A1, CYP2D26, CYP2R1 | 385 | 108 | 17446 | 6,293651 | 6,69E-05 | 1,11E-05 | 1,58E-04 |
| INTERPRO | IPR017972:Cytochrome P450, conserved site | 14 | 2,905 | 1,15E-07 | CYP3A25, CYP2C44, CYP2F2, CYP46A1, CYP2D40, CYP2C29, CYP4F13, CYP1A2, CYP2E1, CYP27A1, CYP2A5, CYP3A59, CYP2D26, CYP2R1 | 436 | 96 | 20594 | 6,888284 | 1,03E-04 | 1,28E-05 | 1,79E-04 |
| INTERPRO | IPR002401:Cytochrome P450, E-class, group I | 13 | 2,697 | 1,69E-07 | CYP2F2, CYP2C44, CYP46A1, CYP2D40, CYP2C29, CYP4F13, CYP1A2, CYP2E1, CYP39A1, CYP27A1, CYP2A5, CYP2D26, CYP2R1 | 436 | 83 | 20594 | 7,398088 | 1,52E-04 | 1,68E-05 | 2,64E-04 |
| GOTERM_MF_DIRECT | GO:0005506~iron ion binding | 20 | 4,149 | 2,45E-07 | CYP3A25, FTL1, CYP2C44, CYP2D40, CYP2F2, CYP46A1, CYP2C29, CYP4F13, PAH, CYP2D37-PS, CYP1A2, CYP2E1, ISCU, CYP39A1, CYP27A1, AOX1, CYP2A5, CYP3A59, CYP2D26, CYP2R1 | 385 | 212 | 17446 | 4,274933 | 1,55E-04 | 2,21E-05 | 3,64E-04 |
| GOTERM_CC_DIRECT | GO:0031090~organelle membrane | 13 | 2,697 | 8,18E-07 | TM7SF2, CYP3A25, CYP39A1, MAP1LC3A, CYP46A1, CYP2F2, CYP2C29, CYP2A5, CYP2D26, EPHX1, CYP2R1, CYP1A2, GULO | 424 | 94 | 19662 | 6,413238 | 2,93E-04 | 4,88E-05 | 0,0011221 |
| UP_SEQ_FEATURE | metal ion-binding site:Iron (heme axial ligand) | 14 | 2,905 | 1,82E-06 | CYP3A25, CYP2F2, CYP46A1, CYC1, CYP2C29, CYB5B, CYP1A2, CYP2E1, CYP39A1, CYP27A1, CYP2A5, CYP3A59, CYP2D26, CYP2R1 | 408 | 114 | 18012 | 5,421569 | 0,0019489 | 9,75E-04 | 0,0028959 |
| COG_ONTOLOGY | Secondary metabolites biosynthesis, transport, and catabolism | 15 | 3,112 | 2,71E-06 | CYP3A25, CYP2C44, CYP2F2, CYP46A1, CYP2D40, CYP2C29, CYP4F13, CYP1A2, CYP2E1, CYP39A1, CYP27A1, CYP2A5, CYP3A59, CYP2D26, CYP2R1 | 62 | 117 | 2126 | 4,396195 | 5,42E-05 | 5,42E-05 | 0,0020609 |
| GOTERM_MF_DIRECT | GO:0016712~oxidoreductase activity, acting on paired donors, with incorporation or reduction of molecular oxygen, reduced flavin or flavoprotein as one donor, and incorporation of one atom of oxygen | 9 | 1,867 | 3,14E-06 | CYP3A25, CYP2C44, CYP2F2, CYP2C29, CYP2A5, CYP2D26, CYP3A59, CYP2E1, CYP1A2 | 385 | 42 | 17446 | 9,710204 | 0,0019802 | 2,48E-04 | 0,0046712 |
| GOTERM_MF_DIRECT | GO:0008395~steroid hydroxylase activity | 8 | 1,66 | 1,72E-04 | CYP2D40, CYP2C44, CYP2F2, CYP2C29, CYP2A5, CYP2D26, CYP2R1, CYP2E1 | 385 | 54 | 17446 | 6,713228 | 0,1029552 | 0,0083228 | 0,2557312 |
| GOTERM_MF_DIRECT | GO:0008392~arachidonic acid epoxygenase activity | 7 | 1,452 | 5,48E-04 | CYP2D40, CYP2C44, CYP2F2, CYP2C29, CYP2A5, CYP2D26, CYP2E1 | 385 | 47 | 17446 | 6,748936 | 0,2922963 | 0,0227851 | 0,8114854 |
| GOTERM_BP_DIRECT | GO:0019373~epoxygenase P450 pathway | 5 | 1,037 | 0,00385111 | CYP2C44, CYP2F2, CYP2C29, CYP2A5, CYP2E1 | 394 | 30 | 18082 | 7,6489 | 0,9991016 | 0,2839749 | 6,3496124 |

| | | | | | | | | | | | | |
|---|---|---|---|---|---|---|---|---|---|---|---|---|
| GOTERM_MF_DIRECT | GO:0070330~aromatase activity | 5 | 1,037 | 0,00780616 | CYP3A25, CYP2C29, CYP2A5, CYP2D26, CYP1A2 | 385 | 36 | 17446 | 6,293651 | 0,9928812 | 0,1862012 | 11,000653 |
| KEGG_PATHWAY | mmu00591:Linoleic acid metabolism | 6 | 1,245 | 0,00990646 | CYP3A25, CYP2C44, PLA2G12A, CYP2C29, CYP2E1, CYP1A2 | 205 | 50 | 7720 | 4,519024 | 0,8977046 | 0,1189673 | 11,975006 |
| KEGG_PATHWAY | mmu00590:Arachidonic acid metabolism | 6 | 1,245 | 0,08606557 | GPX1, CYP2C44, PLA2G12A, CYP2C29, CYP4F13, CYP2E1 | 205 | 89 | 7720 | 2,538778 | 1 | 0,4270776 | 68,430837 |
| Annotation | Enrichment Score: 5.189817369973017 | | | | | | | | | | | |
| Category | Term | Count | % | PValue | Genes | List Total | Pop Hits | Pop Total | Enrichment | Bonferroni | Benjamini | FDR |
| UP_KEYWORDS | Mitochondrion | 53 | 11 | 2,49E-09 | ETNPPL, CMC2, CLPB, CYC1, ECHDC2, COX5A, FAM210A, MTHFD2, ALAS2, FXN, SLC25A23, TIMM9, SLC25A3, SLC25A28, ABCB10, TMEM14C, MTO1, LYRM4, COX4I1, CYB5B, NDUFA11, ISCU, ATG4D, CYP27A1, MARC1, SLC25A38, SLC25A37, H2-KE6, GSTP1, NDUFB3, COASY, UNG, FARS2, HSD3B5, COX7C, AASS, MRPL20, MRPL18, LACTB, NDUFA4, ATP5J2, COX7A2, FADS1, MAOA, MRPS24, MARCH5, ABCB6, ABCG2, NDUFV3, ACSM2, UOX, SLC25A51, URI1 | 456 | 1058 | 22680 | 2,491543 | 7,95E-07 | 1,33E-07 | 3,36E-06 |
| UP_KEYWORDS | Transit peptide | 25 | 5,187 | 1,12E-04 | COX4I1, AASS, ECHDC2, COX5A, PHYH, MRPL20, NDUFV3, MTHFD2, ISCU, ACSM2, ALAS2, FXN, CYP27A1, SLC25A3, MRPL18, ABCB10, ACAA1B, | 456 | 510 | 22680 | 2,43808 | 0,0351786 | 0,0023846 | 0,1513083 |
| UP_SEQ_FEATURE | transit peptide:Mitochondrion | 24 | 4,979 | 9,63E-04 | ECHDC2, AASS, COX5A, MRPL20, NDUFV3, MTHFD2, ISCU, ACSM2, ALAS2, MARC1, FXN, CYP27A1, NUDT8, SLC25A3, ABCB10, MRPL18, | 408 | 496 | 18012 | 2,136148 | 0,644736 | 0,2917546 | 1,5245512 |
| Annotation | Enrichment Score: 4.90866850472854 | | | | | | | | | | | |
| Category | Term | Count | % | PValue | Genes | List Total | Pop Hits | Pop Total | Enrichment | Bonferroni | Benjamini | FDR |
| INTERPRO | IPR002345:Lipocalin | 12 | 2,49 | 1,91E-11 | MUP5, RBP4, MUP7, MUP6, MUP1, MUP3, MUP8, MUP14, MUP13, MUP9, MUP12, C8G | 436 | 31 | 20594 | 18,28411 | 1,71E-08 | 1,71E-08 | 2,97E-08 |
| GOTERM_MF_DIRECT | GO:0036094~small molecule binding | 12 | 2,49 | 4,34E-11 | MUP5, RBP4, MUP7, MUP6, MUP1, MUP3, MUP8, MUP14, MUP13, MUP9, MUP12, C8G | 385 | 32 | 17446 | 16,99286 | 2,74E-08 | 9,13E-09 | 6,46E-08 |
| INTERPRO | IPR022272:Lipocalin conserved site | 12 | 2,49 | 8,71E-11 | MUP5, RBP4, MUP7, MUP6, MUP1, MUP3, MUP8, MUP14, MUP13, MUP9, MUP12, C8G | 436 | 35 | 20594 | 16,1945 | 7,79E-08 | 3,90E-08 | 1,36E-07 |
| INTERPRO | IPR002971:Major urinary protein | 10 | 2,075 | 1,75E-09 | MUP5, MUP7, MUP6, MUP1, MUP3, MUP8, MUP14, MUP13, MUP9, MUP12 | 436 | 26 | 20594 | 18,1669 | 1,57E-06 | 5,23E-07 | 2,73E-06 |

| | | | | | | | | | | | | |
|---|---|---|---|---|---|---|---|---|---|---|---|---|
| INTERPRO | IPR012674:Calycin | 13 | 2,697 | 3,31E-08 | MUP5, RBP4, MUP7, MUP6, MUP1, MUP3, FABP1, MUP8, MUP14, MUP9, MUP13, MUP12, C8G | 436 | 72 | 20594 | 8,528351 | 2,97E-05 | 5,93E-06 | 5,16E-05 |
| INTERPRO | IPR011038:Calycin-like | 13 | 2,697 | 3,89E-08 | MUP5, RBP4, MUP7, MUP6, MUP1, MUP3, FABP1, MUP8, MUP14, MUP9, MUP13, MUP12, C8G | 436 | 73 | 20594 | 8,411524 | 3,48E-05 | 5,81E-06 | 6,06E-05 |
| INTERPRO | IPR000566:Lipocalin/cytosolic fatty-acid binding protein domain | 12 | 2,49 | 1,06E-07 | MUP5, RBP4, MUP7, MUP6, MUP1, MUP3, MUP8, MUP14, MUP13, MUP9, MUP12, C8G | 436 | 65 | 20594 | 8,720113 | 9,46E-05 | 1,35E-05 | 1,65E-04 |
| UP_SEQ_FEATURE | chain:Major urinary protein 6 | 7 | 1,452 | 8,48E-07 | MUP7, MUP1, MUP8, MUP14, MUP13, MUP9, MUP12 | 408 | 16 | 18012 | 19,31434 | 9,10E-04 | 9,10E-04 | 0,0013521 |
| UP_SEQ_FEATURE | chain:Major urinary protein 2 | 7 | 1,452 | 8,48E-07 | MUP7, MUP1, MUP8, MUP14, MUP13, MUP9, MUP12 | 408 | 16 | 18012 | 19,31434 | 9,10E-04 | 9,10E-04 | 0,0013521 |
| UP_SEQ_FEATURE | chain:Major urinary protein 1 | 7 | 1,452 | 8,48E-07 | MUP7, MUP1, MUP8, MUP14, MUP13, MUP9, MUP12 | 408 | 16 | 18012 | 19,31434 | 9,10E-04 | 9,10E-04 | 0,0013521 |
| GOTERM_MF_DIRECT | GO:0005215~transporter activity | 18 | 3,734 | 6,62E-06 | MUP5, RBP4, MUP7, MUP6, MUP1, MUP3, STARD10, SLCO1A1, TIMM9, SLC25A38, ABCB10, FABP1, MUP8, MUP14, MUP13, MUP9, MUP12, SLC15A5 | 385 | 217 | 17446 | 3,758789 | 0,0041713 | 4,64E-04 | 0,0098508 |
| GOTERM_BP_DIRECT | GO:0045721~negative regulation of gluconeogenesis | 7 | 1,452 | 9,80E-06 | MUP5, PGP, MUP1, SERPINA12, MUP3, MUP9, MUP12 | 394 | 24 | 18082 | 13,38558 | 0,0176601 | 0,0059217 | 0,0166616 |
| GOTERM_BP_DIRECT | GO:0051055~negative regulation of lipid biosynthetic process | 6 | 1,245 | 1,69E-05 | MUP5, MUP1, SERPINA12, MUP3, MUP9, MUP12 | 394 | 16 | 18082 | 17,21003 | 0,0303439 | 0,0076738 | 0,0288124 |
| GOTERM_BP_DIRECT | GO:0006112~energy reserve metabolic process | 6 | 1,245 | 9,18E-05 | MUP5, MUP1, MUP3, GNAS, MUP9, MUP12 | 394 | 22 | 18082 | 12,51638 | 0,1536574 | 0,0206379 | 0,155896 |
| GOTERM_BP_DIRECT | GO:0061179~negative regulation of insulin secretion involved in cellular response to glucose stimulus | 6 | 1,245 | 9,18E-05 | MUP5, MUP1, MUP3, PIM3, MUP9, MUP12 | 394 | 22 | 18082 | 12,51638 | 0,1536574 | 0,0206379 | 0,155896 |
| GOTERM_MF_DIRECT | GO:0005009~insulin-activated receptor activity | 5 | 1,037 | 1,41E-04 | MUP5, MUP1, MUP3, MUP9, MUP12 | 385 | 13 | 17446 | 17,42857 | 0,0852161 | 0,0080643 | 0,2096887 |
| GOTERM_BP_DIRECT | GO:0035634~response to stilbenoid | 6 | 1,245 | 1,75E-04 | SLCO1A1, MUP1, MUP3, HSD3B5, CYP2A5, MUP12 | 394 | 25 | 18082 | 11,01442 | 0,2731362 | 0,0313981 | 0,297894 |
| GOTERM_BP_DIRECT | GO:0045834~positive regulation of lipid metabolic process | 5 | 1,037 | 2,48E-04 | MUP5, MUP1, MUP3, MUP9, MUP12 | 394 | 15 | 18082 | 15,2978 | 0,3630217 | 0,0401726 | 0,4208979 |
| GOTERM_BP_DIRECT | GO:0031649~heat generation | 5 | 1,037 | 2,48E-04 | MUP5, MUP1, MUP3, MUP9, MUP12 | 394 | 15 | 18082 | 15,2978 | 0,3630217 | 0,0401726 | 0,4208979 |
| GOTERM_BP_DIRECT | GO:0071396~cellular response to lipid | 5 | 1,037 | 2,48E-04 | MUP5, MUP1, MUP3, MUP9, MUP12 | 394 | 15 | 18082 | 15,2978 | 0,3630217 | 0,0401726 | 0,4208979 |
| GOTERM_BP_DIRECT | GO:0010888~negative regulation of lipid storage | 5 | 1,037 | 6,58E-04 | MUP5, MUP1, MUP3, MUP9, MUP12 | 394 | 19 | 18082 | 12,07721 | 0,6975206 | 0,0878768 | 1,112009 |
| GOTERM_MF_DIRECT | GO:0005550~pheromone binding | 10 | 2,075 | 7,25E-04 | MUP5, MUP7, MUP6, MUP1, MUP3, MUP8, MUP14, MUP13, MUP9, MUP12 | 385 | 110 | 17446 | 4,119481 | 0,3673132 | 0,0282058 | 1,07307 |
| GOTERM_BP_DIRECT | GO:0010907~positive regulation of glucose metabolic process | 5 | 1,037 | 9,81E-04 | MUP5, MUP1, MUP3, MUP9, MUP12 | 394 | 21 | 18082 | 10,927 | 0,8320884 | 0,1196637 | 1,6548188 |

| | | | | | | | | | | | | |
|---|---|---|---|---|---|---|---|---|---|---|---|---|
| GOTERM_BP_DIRECT | GO:0051897~positive regulation of protein kinase B signaling | 9 | 1,867 | 0,00124412 | NOX4, EGFR, MUP5, GPX1, F10, MUP1, MUP3, MUP9, MUP12 | 394 | 97 | 18082 | 4,258151 | 0,8959848 | 0,1400502 | 2,0942828 |
| GOTERM_BP_DIRECT | GO:0009060~aerobic respiration | 6 | 1,245 | 0,00148187 | MUP5, FXN, MUP1, MUP3, MUP9, MUP12 | 394 | 39 | 18082 | 7,060523 | 0,9325282 | 0,1550711 | 2,489777 |
| GOTERM_BP_DIRECT | GO:0045475~locomotor rhythm | 5 | 1,037 | 0,00224931 | MUP5, MUP1, MUP3, MUP9, MUP12 | 394 | 26 | 18082 | 8,825654 | 0,9833252 | 0,214013 | 3,7561441 |
| UP_KEYWORDS | Pheromone-binding | 4 | 0,83 | 0,00247627 | MUP5, MUP3, MUP9, MUP12 | 456 | 14 | 22680 | 14,21053 | 0,5465684 | 0,036962 | 3,2888652 |
| GOTERM_BP_DIRECT | GO:0042593~glucose homeostasis | 10 | 2,075 | 0,00251991 | MUP5, RBP4, MUP1, MUP3, PPARG, PDE3B, RPS6, GCGR, MUP9, MUP12 | 394 | 133 | 18082 | 3,450632 | 0,9898164 | 0,2249533 | 4,1989586 |
| GOTERM_BP_DIRECT | GO:0070584~mitochondrion morphogenesis | 5 | 1,037 | 0,0048841 | MUP5, MUP1, MUP3, MUP9, MUP12 | 394 | 32 | 18082 | 7,170844 | 0,9998638 | 0,3327518 | 7,9870769 |
| GOTERM_BP_DIRECT | GO:0010628~positive regulation of gene expression | 12 | 2,49 | 0,25264147 | BMP4, MUP5, HRAS, ATF4, MUP1, MUP3, NTRK2, ZPR1, MUP9, DTNBP1, MUP12, MYCN | 394 | 399 | 18082 | 1,380253 | 1 | 0,9954936 | 99,292402 |
| Annotation | Enrichment Score: 4.35123739329693 | | | | | | | | | | | |
| Category | Term | Count | % | PValue | Genes | List Total | Pop Hits | Pop Total | Enrichment | Bonferroni | Benjamini | FDR |
| KEGG_PATHWAY | mmu00140:Steroid hormone biosynthesis | 15 | 3,112 | 5,13E-08 | CYP3A25, HSD17B2, CYP2D40, CYP2C44, HSD3B5, CYP2C29, UGT2B1, CYP1A2, CYP2E1, UGT1A1, UGT1A9, UGT2B35, UGT2B5, CYP2D26, H2-KE6 | 205 | 87 | 7720 | 6,492851 | 1,17E-05 | 3,91E-06 | 6,57E-05 |
| KEGG_PATHWAY | mmu00980:Metabolism of xenobiotics by cytochrome P450 | 13 | 2,697 | 8,24E-08 | GSTA4, CYP2F2, UGT2B1, EPHX1, CYP1A2, CYP2E1, UGT1A1, GSTM1, UGT1A9, UGT2B35, UGT2B5, GSTP2, GSTP1 | 205 | 64 | 7720 | 7,64939 | 1,89E-05 | 4,72E-06 | 1,06E-04 |
| KEGG_PATHWAY | mmu05204:Chemical carcinogenesis | 15 | 3,112 | 1,06E-07 | GSTA4, CYP3A25, CYP2C44, CYP2C29, UGT2B1, EPHX1, CYP1A2, CYP2E1, UGT1A1, GSTM1, UGT1A9, UGT2B35, UGT2B5, GSTP2, GSTP1 | 205 | 92 | 7720 | 6,139979 | 2,43E-05 | 4,87E-06 | 1,36E-04 |
| KEGG_PATHWAY | mmu00982:Drug metabolism - cytochrome P450 | 13 | 2,697 | 1,18E-07 | GSTA4, MAOA, UGT2B1, CYP1A2, CYP2E1, UGT1A1, GSTM1, UGT1A9, UGT2B35, AOX1, UGT2B5, GSTP2, GSTP1 | 205 | 66 | 7720 | 7,417591 | 2,70E-05 | 4,50E-06 | 1,51E-04 |
| KEGG_PATHWAY | mmu00040:Pentose and glucuronate interconversions | 9 | 1,867 | 3,32E-06 | CRYL1, UGT1A9, UGT2B35, AKR1A1, UGT2B1, UGDH, UGT2B5, UGT1A1, XYLB | 205 | 36 | 7720 | 9,414634 | 7,60E-04 | 1,09E-04 | 0,0042523 |
| KEGG_PATHWAY | mmu00830:Retinol metabolism | 13 | 2,697 | 3,34E-06 | CYP3A25, CYP2C44, CYP2C29, UGT2B1, CYP1A2, UGT1A1, RDH9, UGT1A9, UGT2B35, DGAT1, AOX1, UGT2B5, CYP2A5 | 205 | 89 | 7720 | 5,500685 | 7,64E-04 | 9,55E-05 | 0,0042733 |
| KEGG_PATHWAY | mmu00053:Ascorbate and aldarate metabolism | 7 | 1,452 | 5,89E-05 | UGT1A9, UGT2B35, UGT2B1, UGDH, UGT2B5, GULO, UGT1A1 | 205 | 27 | 7720 | 9,763324 | 0,0133904 | 0,0014968 | 0,075391 |
| GOTERM_BP_DIRECT | GO:0009813~flavonoid biosynthetic process | 6 | 1,245 | 7,22E-05 | UGT1A9, UGT2B35, UGT2B1, UGT2B5, UGT1A1, UGT3A2 | 394 | 21 | 18082 | 13,1124 | 0,1229952 | 0,0216363 | 0,1226608 |
| GOTERM_BP_DIRECT | GO:0052696~flavonoid glucuronidation | 6 | 1,245 | 7,22E-05 | UGT1A9, UGT2B35, UGT2B1, UGT2B5, UGT1A1, UGT3A2 | 394 | 21 | 18082 | 13,1124 | 0,1229952 | 0,0216363 | 0,1226608 |

| | | | | | | | | | | | | |
|---|---|---|---|---|---|---|---|---|---|---|---|---|
| INTERPRO | IPR002213:UDP-glucuronosyl/UDP-glucosyltransferase | 6 | 1,245 | 8,04E-05 | UGT1A9, UGT2B35, UGT2B1, UGT2B5, UGT1A1, UGT3A2 | 436 | 22 | 20594 | 12,88198 | 0,0694429 | 0,0071713 | 0,1251787 |
| GOTERM_MF_DIRECT | GO:0015020~glucuronosyltransferase activity | 6 | 1,245 | 3,87E-04 | UGT1A9, UGT2B35, UGT2B1, UGT2B5, UGT1A1, UGT3A2 | 385 | 29 | 17446 | 9,375369 | 0,2165387 | 0,0172799 | 0,5734736 |
| KEGG_PATHWAY | mmu00860:Porphyrin and chlorophyll metabolism | 7 | 1,452 | 6,57E-04 | UGT1A9, ALAS2, UGT2B35, FXN, UGT2B1, UGT2B5, UGT1A1 | 205 | 41 | 7720 | 6,429506 | 0,139741 | 0,0149395 | 0,8385647 |
| KEGG_PATHWAY | mmu00983:Drug metabolism - other enzymes | 7 | 1,452 | 0,00211441 | UGT1A9, UGT2B35, UGT2B1, UGT2B5, CES2A, CES1E, UGT1A1 | 205 | 51 | 7720 | 5,168819 | 0,384126 | 0,0395878 | 2,6753082 |
| GOTERM_MF_DIRECT | GO:0016758~transferase activity, transferring hexosyl groups | 5 | 1,037 | 0,00454437 | UGT1A9, UGT2B5, UGT1A1, UGT3A2, B4GALNT1 | 385 | 31 | 17446 | 7,308756 | 0,9435293 | 0,1338539 | 6,5490551 |
| UP_KEYWORDS | Glycosyltransferase | 10 | 2,075 | 0,03844037 | POMT2, NAMPT, GCNT4, UGT1A9, UGT2B35, UGT2B1, UGT2B5, UGT1A1, UGT3A2, B4GALNT1 | 456 | 225 | 22680 | 2,210526 | 0,9999963 | 0,2867736 | 41,063875 |
| GOTERM_MF_DIRECT | GO:0016757~transferase activity, transferring glycosyl groups | 8 | 1,66 | 0,17662097 | POMT2, NAMPT, GCNT4, UGT1A9, UGT2B5, UGT1A1, UGT3A2, B4GALNT1 | 385 | 208 | 17446 | 1,742857 | 1 | 0,8616358 | 94,442481 |
| Annotation | Enrichment Score: 3.751042040151358 | | | | | | | | | | | |
| Category | Term | Count | % | PValue | Genes | List Total | Pop Hits | Pop Total | Enrichment | Bonferroni | Benjamini | FDR |
| GOTERM_CC_DIRECT | GO:0022627~cytosolic small ribosomal subunit | 9 | 1,867 | 1,63E-05 | RPSA, RPS18, RPS14, RPS15, RPS10, RPS6, RPS2, RPS23, RPS8 | 424 | 53 | 19662 | 7,8746 | 0,0058211 | 6,48E-04 | 0,0223758 |
| GOTERM_CC_DIRECT | GO:0015935~small ribosomal subunit | 6 | 1,245 | 3,49E-04 | RPSA, RPS18, RPS15, RPS6, RPS2, RPS23 | 424 | 29 | 19662 | 9,59434 | 0,1174713 | 0,0103596 | 0,4778622 |
| GOTERM_BP_DIRECT | GO:0000028~ribosomal small subunit assembly | 5 | 1,037 | 9,81E-04 | RPSA, RPS14, RPS15, RPS10, RPS2 | 394 | 21 | 18082 | 10,927 | 0,8320884 | 0,1196637 | 1,6548188 |
| Annotation | Enrichment Score: 2.524771980804137 | | | | | | | | | | | |
| Category | Term | Count | % | PValue | Genes | List Total | Pop Hits | Pop Total | Enrichment | Bonferroni | Benjamini | FDR |
| GOTERM_BP_DIRECT | GO:0006629~lipid metabolic process | 23 | 4,772 | 4,80E-04 | ECHDC2, CYP1A2, PNPLA1, CHPT1, GPX1, ACSM2, GBA2, ISYNA1, BAAT, CYP39A1, ELOVL3, PLA2G12A, H2-KE6, PTDSS1, ACAA1B, SLC27A2, | 394 | 459 | 18082 | 2,299669 | 0,5822024 | 0,070148 | 0,8128672 |
| UP_KEYWORDS | Lipid metabolism | 20 | 4,149 | 8,37E-04 | ECHDC2, CYP1A2, PNPLA1, CHPT1, GBA2, ACSM2, ISYNA1, BAAT, CYP39A1, ELOVL3, PLA2G12A, H2-KE6, PTDSS1, ACAA1B, SLC27A2 | 456 | 417 | 22680 | 2,38546 | 0,2343662 | 0,0147267 | 1,1228071 |
| UP_KEYWORDS | Fatty acid metabolism | 8 | 1,66 | 0,01263879 | ACSM2, BAAT, ELOVL3, FADS1, ECHDC2, H2-KE6, ACAA1B, SLC27A2 | 456 | 124 | 22680 | 3,208829 | 0,9827073 | 0,1444892 | 15,765005 |
| UP_KEYWORDS | Lipid biosynthesis | 9 | 1,867 | 0,01568153 | TM7SF2, EBP, ISYNA1, HSD17B2, ELOVL3, FADS1, H2-KE6, PTDSS1, CHPT1 | 456 | 160 | 22680 | 2,797697 | 0,9935395 | 0,1703426 | 19,199741 |
| Annotation | Enrichment Score: 2.4101356339356292 | | | | | | | | | | | |
| Category | Term | Count | % | PValue | Genes | List Total | Pop Hits | Pop Total | Enrichment | Bonferroni | Benjamini | FDR |
| GOTERM_BP_DIRECT | GO:0006749~glutathione metabolic process | 8 | 1,66 | 8,49E-05 | GSTM1, GPX1, GSTA4, TXNRD3, EEF1G, GLO1, GSTP2, GSTP1 | 394 | 49 | 18082 | 7,4928 | 0,1430692 | 0,0218154 | 0,1442864 |

| | | | | | | | | | | | | |
|---|---|---|---|---|---|---|---|---|---|---|---|---|
| INTERPRO | IPR004046:Glutathione S-transferase, C-terminal | 6 | 1,245 | 4,44E-04 | GSTM1, GSTA4, EEF1G, GSTP2, GSTP1, MARS | 436 | 31 | 20594 | 9,142054 | 0,3278098 | 0,0354662 | 0,6889123 |
| INTERPRO | IPR012336:Thioredoxin-like fold | 11 | 2,282 | 6,36E-04 | GSTM1, GPX1, GSTA4, TMX4, PDILT, TXNRD3, EEF1G, GSTP2, MIEN1, GSTP1, GLRX | 436 | 136 | 20594 | 3,820393 | 0,4342769 | 0,0463618 | 0,9865002 |
| UP_SEQ_FEATURE | domain:GST C-terminal | 6 | 1,245 | 0,00220545 | GSTM1, GSTA4, EEF1G, GSTP2, GSTP1, MARS | 408 | 41 | 18012 | 6,460545 | 0,9066374 | 0,4472315 | 3,458887 |
| INTERPRO | IPR010987:Glutathione S-transferase, C-terminal-like | 6 | 1,245 | 0,00276285 | GSTM1, GSTA4, EEF1G, GSTP2, GSTP1, MARS | 436 | 46 | 20594 | 6,160949 | 0,9159356 | 0,1734334 | 4,2179008 |
| UP_SEQ_FEATURE | domain:GST N-terminal | 5 | 1,037 | 0,00442178 | GSTM1, GSTA4, EEF1G, GSTP2, GSTP1 | 408 | 30 | 18012 | 7,357843 | 0,9914303 | 0,5476291 | 6,8216376 |
| INTERPRO | IPR004045:Glutathione S-transferase, N-terminal | 5 | 1,037 | 0,00494812 | GSTM1, GSTA4, EEF1G, GSTP2, GSTP1 | 436 | 33 | 20594 | 7,156658 | 0,9881989 | 0,2423037 | 7,4354972 |
| GOTERM_MF_DIRECT | GO:0004364~glutathione transferase activity | 5 | 1,037 | 0,00570978 | GSTM1, GSTA4, EEF1G, GSTP2, GSTP1 | 385 | 33 | 17446 | 6,865801 | 0,9730344 | 0,1514584 | 8,1628876 |
| GOTERM_MF_DIRECT | GO:0043295~glutathione binding | 3 | 0,622 | 0,04199903 | GSTA4, GSTP2, GSTP1 | 385 | 15 | 17446 | 9,062857 | 1 | 0,5286033 | 47,168589 |
| KEGG_PATHWAY | mmu00480:Glutathione metabolism | 5 | 1,037 | 0,05678277 | GSTM1, GPX1, GSTA4, GSTP2, GSTP1 | 205 | 55 | 7720 | 3,423503 | 0,9999985 | 0,3697394 | 52,713392 |
| GOTERM_MF_DIRECT | GO:0004602~glutathione peroxidase activity | 3 | 0,622 | 0,07075553 | GPX1, GSTP2, GSTP1 | 385 | 20 | 17446 | 6,797143 | 1 | 0,6767692 | 66,421631 |
| Annotation | Enrichment Score: 2.3915123601981705 | | | | | | | | | | | |
| Category | Term | Count | % | PValue | Genes | List Total | Pop Hits | Pop Total | Enrichment | Bonferroni | Benjamini | FDR |
| GOTERM_CC_DIRECT | GO:0005743~mitochondrial inner membrane | 27 | 5,602 | 3,37E-07 | NDUFB3, CYC1, HSD3B5, COX7C, COX5A, ALAS2, SLC25A23, TIMM9, SLC25A3, ABCB10, SLC25A28, TMEM14C, NDUFA4, ATP5J2, COX7A2, COX4I1, SLC3A1, CYB5B, ABCB6, NDUFA11, NDUFV3, CYP27A1, MARC1, SLC25A38, UGT2B5, SLC25A37, SLC25A51 | 424 | 387 | 19662 | 3,235301 | 1,21E-04 | 2,41E-05 | 4,63E-04 |
| UP_KEYWORDS | Mitochondrion inner membrane | 18 | 3,734 | 1,64E-05 | NDUFA4, NDUFB3, ATP5J2, COX7A2, CYC1, COX7C, COX4I1, COX5A, NDUFA11, NDUFV3, SLC25A23, TIMM9, SLC25A38, SLC25A3, ABCB10, SLC25A37, SLC25A28, SLC25A51 | 456 | 254 | 22680 | 3,524658 | 0,0052045 | 4,74E-04 | 0,0220609 |
| UP_KEYWORDS | Electron transport | 10 | 2,075 | 3,07E-04 | NDUFV3, NDUFA4, NDUFB3, FADS1, TMX4, CYC1, TXNRD3, CYB5B, GLRX, NDUFA11 | 456 | 107 | 22680 | 4,648303 | 0,0934021 | 0,0057514 | 0,4137476 |
| GOTERM_MF_DIRECT | GO:0004129~cytochrome-c oxidase activity | 5 | 1,037 | 0,00311428 | NDUFA4, COX7A2, COX7C, COX4I1, COX5A | 385 | 28 | 17446 | 8,091837 | 0,8602888 | 0,0984036 | 4,5325538 |
| GOTERM_CC_DIRECT | GO:0005751~mitochondrial respiratory chain complex IV | 4 | 0,83 | 0,00371058 | NDUFA4, COX7C, COX4I1, COX5A | 424 | 15 | 19662 | 12,36604 | 0,7357498 | 0,0849018 | 4,9734666 |
| KEGG_PATHWAY | mmu05010:Alzheimer's disease | 12 | 2,49 | 0,00714094 | COX7C, COX4I1, MME, COX5A, ITPR1, NDUFA11, ITPR2 | 205 | 177 | 7720 | 2,553121 | 0,8062412 | 0,1106144 | 8,7725478 |

| | | | | | | | | | | | | |
|---|---|---|---|---|---|---|---|---|---|---|---|---|
| KEGG_PATHWAY | mmu04932:Non-alcoholic fatty liver disease (NAFLD) | 11 | 2,282 | 0,00857244 | NDUFV3, NDUFA4, NDUFB3, ATF4, COX7A2, CYC1, COX7C, COX4I1, CYP2E1, COX5A, NDUFA11 | 205 | 157 | 7720 | 2,638496 | 0,8607593 | 0,1095013 | 10,443389 |
| KEGG_PATHWAY | mmu00190:Oxidative phosphorylation | 10 | 2,075 | 0,01119511 | NDUFV3, NDUFA4, NDUFB3, ATP5J2, COX7A2, CYC1, COX7C, COX4I1, COX5A, NDUFA11 | 205 | 139 | 7720 | 2,709247 | 0,9240848 | 0,1209442 | 13,431572 |
| KEGG_PATHWAY | mmu05016:Huntington's disease | 12 | 2,49 | 0,01569369 | PPARG, CYC1, COX7C, COX4I1, COX5A, ITPR1, NDUFA11 | 205 | 198 | 7720 | 2,282336 | 0,9732803 | 0,1584359 | 18,343936 |
| GOTERM_BP_DIRECT | GO:0006123~mitochondrial electron transport, cytochrome c to oxygen | 3 | 0,622 | 0,02692769 | COX7C, COX4I1, COX5A | 394 | 12 | 18082 | 11,47335 | 1 | 0,7879226 | 37,129401 |
| UP_KEYWORDS | Respiratory chain | 5 | 1,037 | 0,03583352 | NDUFV3, NDUFA4, NDUFB3, CYC1, NDUFA11 | 456 | 62 | 22680 | 4,011036 | 0,9999912 | 0,2829374 | 38,871887 |
| GOTERM_CC_DIRECT | GO:0070469~respiratory chain | 5 | 1,037 | 0,03605672 | NDUFV3, NDUFA4, NDUFB3, CYC1, NDUFA11 | 424 | 58 | 19662 | 3,997642 | 0,999998 | 0,4217692 | 39,58529 |
| KEGG_PATHWAY | mmu05012:Parkinson's disease | 9 | 1,867 | 0,04352164 | NDUFV3, NDUFA4, NDUFB3, COX7A2, CYC1, COX7C, COX4I1, COX5A, NDUFA11 | 205 | 149 | 7720 | 2,274677 | 0,9999625 | 0,3242397 | 43,451681 |
| KEGG_PATHWAY | mmu04260:Cardiac muscle contraction | 6 | 1,245 | 0,05247389 | COX7A2, CYC1, COX7C, COX4I1, ATP1A1, COX5A | 205 | 77 | 7720 | 2,934431 | 0,9999956 | 0,3564988 | 49,869964 |
| GOTERM_CC_DIRECT | GO:0005747~mitochondrial respiratory chain complex I | 4 | 0,83 | 0,08012069 | NDUFV3, NDUFA4, NDUFB3, NDUFA11 | 424 | 47 | 19662 | 3,946608 | 1 | 0,6433325 | 68,21018 |
| Annotation | Enrichment Score: 2.3286079095014918 | | | | | | | | | | | |
| Category | Term | Count | % | PValue | Genes | List Total | Pop Hits | Pop Total | Enrichment | Bonferroni | Benjamini | FDR |
| UP_KEYWORDS | Metal-binding | 101 | 20,95 | 3,66E-05 | ADCY1, ILKAP, LMO4, SYT3, RP9, NT5DC3, PDE3B, COX5A, MCM10, PGP, SLC25A23, TIMM9, CYHR1, CGRRF1, OLA1, COLEC12, CYP2E1, CYP1A2, CHPT1, BRAP, ZFP592, ISCU, NME2, ZFP598, NME1, CAR8, CYP2D26, PELO, ZFP511, KALRN, FRAS1, CYP3A25, CYP2C44, MME, ZCRB1, ACP5, PAH, ZC3H8, ARIH2, GLO1, TRAF4, MOCS1, CYP2D40, CYP46A1, GDE1, MAP2K2, KLF16, HGD, CSRP2, SALL2, NR1I3, RNF152, CYP2R1, GTF3A, PPARG, CYC1, CPEB1, CYP39A1, NUBP1, FXN, ZFP354A, MT1, ZPR1, HPD, MICAL2, CYP2C29, IRF2BP2, DNAJC21, CYB5B, CYP27A1, ZFAND2B, CYP2A5, CYP3A59, GNAS, PHYHD1, CPM, FTL1, CYP2F2, RPL37, SFTPA1, DCT, NUDT8, COL27A1, PLA2G12A, DTNB, ZFP703, CYP4F13, PHF10, ATP1A1, MARCH5, PHYH, SIRT2, CABYR, PLEKHF1, ACSM2, MARCH2, KCMF1, AOX1, ARSA, SP5, SMPD1 | 456 | 3395 | 22680 | 1,479653 | 0,0115926 | 8,97E-04 | 0,0492904 |

| | | | | | | | | | | | | |
|---|---|---|---|---|---|---|---|---|---|---|---|---|
| GOTERM_MF_DIRECT | GO:0046872~metal ion binding | 95 | 19,71 | 0,0060419 | RP9, NT5DC3, PDE3B, CPEB1, COX5A, MCM10, PGP, CYP39A1, FXN, NUBP1, SLC25A23, TIMM9, MT1, ZFP354A, ZPR1, CYHR1, HPD, CGRRF1, MICAL2, CYP2C29, OLA1, COLEC12, IRF2BP2, CYP2E1, CYB5B, CYP1A2, CHPT1, BRAP, ZFP592, ISCU, NME2, ZFP598, CYP27A1, NME1, ZFAND2B, CAR8, CYP2D26, PELO, GNAS, ZFP511, KALRN, FRAS1, PHYHD1, CPM, CYP3A25, CYP2F2, MME, ZCRB1, ACP5, RPL37, PAH, SFTPA1, ZC3H8, DCT, ARIH2, NUDT8, COL27A1, PLA2G12A, DTNB, GLO1, TRAF4, MOCS1, CYP46A1, GDE1, MAP2K2, | 385 | 3355 | 17446 | 1,283117 | 0,9781598 | 0,1531748 | 8,6180205 |
| UP_KEYWORDS | Zinc | 44 | 9,129 | 0,46783434 | RPL37, CPEB1, MCM10, ZC3H8, DCT, ARIH2, TIMM9, DTNB, ZFP354A, GLO1, ZPR1, CYHR1, TRAF4, CGRRF1, MICAL2, ZFP703, KLF16, PHF10, DNAJC21, IRF2BP2, CSRP2, MARCH5, SIRT2, BRAP, ZFP592, PLEKHF1, SALL2, MARCH2, NR1I3, | 456 | 2099 | 22680 | 1,042602 | 1 | 0,8824269 | 99,979821 |
| Annotation | Enrichment Score: 2.304911280767715 | | | | | | | | | | | |
| Category | Term | Count | % | PValue | Genes | List Total | Pop Hits | Pop Total | Enrichment | Bonferroni | Benjamini | FDR |
| UP_SEQ_FEATURE | repeat:Solcar 3 | 6 | 1,245 | 0,00404276 | SLC25A23, SLC25A38, SLC25A3, SLC25A28, SLC25A37, SLC25A51 | 408 | 47 | 18012 | 5,635795 | 0,9871026 | 0,5811095 | 6,2544723 |
| INTERPRO | IPR023395:Mitochondrial carrier domain | 6 | 1,245 | 0,00472567 | SLC25A23, SLC25A38, SLC25A3, SLC25A28, SLC25A37, SLC25A51 | 436 | 52 | 20594 | 5,450071 | 0,9855853 | 0,2462041 | 7,1126516 |
| INTERPRO | IPR018108:Mitochondrial substrate/solute carrier | 6 | 1,245 | 0,00472567 | SLC25A23, SLC25A38, SLC25A3, SLC25A28, SLC25A37, SLC25A51 | 436 | 52 | 20594 | 5,450071 | 0,9855853 | 0,2462041 | 7,1126516 |
| UP_SEQ_FEATURE | repeat:Solcar 2 | 6 | 1,245 | 0,00575332 | SLC25A23, SLC25A38, SLC25A3, SLC25A28, SLC25A37, SLC25A51 | 408 | 51 | 18012 | 5,193772 | 0,9979643 | 0,5391184 | 8,7888181 |
| UP_SEQ_FEATURE | repeat:Solcar 1 | 6 | 1,245 | 0,00575332 | SLC25A23, SLC25A38, SLC25A3, SLC25A28, SLC25A37, SLC25A51 | 408 | 51 | 18012 | 5,193772 | 0,9979643 | 0,5391184 | 8,7888181 |
| Annotation | Enrichment Score: 2.149455231081104 | | | | | | | | | | | |
| Category | Term | Count | % | PValue | Genes | List Total | Pop Hits | Pop Total | Enrichment | Bonferroni | Benjamini | FDR |
| UP_KEYWORDS | Phenylalanine catabolism | 3 | 0,622 | 0,00571032 | HGD, PAH, HPD | 456 | 6 | 22680 | 24,86842 | 0,8390745 | 0,0763543 | 7,4334325 |
| UP_SEQ_FEATURE | metal ion-binding site:Iron | 6 | 1,245 | 0,00677588 | PHYHD1, FTL1, HGD, PAH, PHYH, HPD | 408 | 53 | 18012 | 4,99778 | 0,9993259 | 0,5557377 | 10,273005 |
| GOTERM_BP_DIRECT | GO:0006559~L-phenylalanine catabolic process | 3 | 0,622 | 0,00920467 | HGD, PAH, HPD | 394 | 7 | 18082 | 19,6686 | 1 | 0,4761757 | 14,548313 |
| Annotation | Enrichment Score: 2.108987765453137 | | | | | | | | | | | |
| Category | Term | Count | % | PValue | Genes | List Total | Pop Hits | Pop Total | Enrichment | Bonferroni | Benjamini | FDR |

| | | | | | | | | | | | | |
|---|---|---|---|---|---|---|---|---|---|---|---|---|
| INTERPRO | IPR020471:Aldo/keto reductase subgroup | 4 | 0,83 | 0,00427087 | AKR1A1, AKR1C19, AKR1C12, AKR1C13 | 436 | 16 | 20594 | 11,80849 | 0,9783037 | 0,2393755 | 6,449295 |
| INTERPRO | IPR018170:Aldo/keto reductase, conserved site | 4 | 0,83 | 0,00510557 | AKR1A1, AKR1C19, AKR1C12, AKR1C13 | 436 | 17 | 20594 | 11,11387 | 0,9897573 | 0,2362248 | 7,6633741 |
| INTERPRO | IPR001395:Aldo/keto reductase | 4 | 0,83 | 0,00816781 | AKR1A1, AKR1C19, AKR1C12, AKR1C13 | 436 | 20 | 20594 | 9,446789 | 0,9993511 | 0,3348811 | 11,992335 |
| INTERPRO | IPR023210:NADP-dependent oxidoreductase domain | 4 | 0,83 | 0,00938187 | AKR1A1, AKR1C19, AKR1C12, AKR1C13 | 436 | 21 | 20594 | 8,996942 | 0,9997832 | 0,3585486 | 13,655423 |
| PIR_SUPERFAMILY | PIRSF000097:aldo-keto reductase | 4 | 0,83 | 0,01706495 | AKR1A1, AKR1C19, AKR1C12, AKR1C13 | 65 | 16 | 1807 | 6,95 | 0,6843811 | 0,6843811 | 16,182967 |
| Annotation | Enrichment Score: 1.9266522383324713 | | | | | | | | | | | |
| Category | Term | Count | % | PValue | Genes | List Total | Pop Hits | Pop Total | Enrichment | Bonferroni | Benjamini | FDR |
| UP_SEQ_FEATURE | domain:MIR 2 | 3 | 0,622 | 0,00992117 | POMT2, ITPR1, ITPR2 | 408 | 7 | 18012 | 18,92017 | 0,9999776 | 0,6572845 | 14,697594 |
| UP_SEQ_FEATURE | domain:MIR 1 | 3 | 0,622 | 0,00992117 | POMT2, ITPR1, ITPR2 | 408 | 7 | 18012 | 18,92017 | 0,9999776 | 0,6572845 | 14,697594 |
| UP_SEQ_FEATURE | domain:MIR 3 | 3 | 0,622 | 0,00992117 | POMT2, ITPR1, ITPR2 | 408 | 7 | 18012 | 18,92017 | 0,9999776 | 0,6572845 | 14,697594 |
| SMART | SM00472:MIR | 3 | 0,622 | 0,01103976 | POMT2, ITPR1, ITPR2 | 172 | 10 | 10425 | 18,18314 | 0,8344342 | 0,8344342 | 12,569805 |
| INTERPRO | IPR016093:MIR motif | 3 | 0,622 | 0,02158194 | POMT2, ITPR1, ITPR2 | 436 | 11 | 20594 | 12,88198 | 1 | 0,6053959 | 28,812012 |
| Annotation | Enrichment Score: 1.7054184369762884 | | | | | | | | | | | |
| Category | Term | Count | % | PValue | Genes | List Total | Pop Hits | Pop Total | Enrichment | Bonferroni | Benjamini | FDR |
| GOTERM_BP_DIRECT | GO:0006879~cellular iron ion homeostasis | 6 | 1,245 | 0,00282756 | ISCU, ALAS2, FTL1, FXN, NUBP1, ABCB6 | 394 | 45 | 18082 | 6,11912 | 0,9941877 | 0,2373346 | 4,7000774 |
| GOTERM_BP_DIRECT | GO:0016226~iron-sulfur cluster assembly | 3 | 0,622 | 0,05168352 | ISCU, FXN, NUBP1 | 394 | 17 | 18082 | 8,098835 | 1 | 0,8716092 | 59,433417 |
| GOTERM_MF_DIRECT | GO:0051536~iron-sulfur cluster binding | 5 | 1,037 | 0,05235777 | ISCU, FXN, NUBP1, AOX1, MOCS1 | 385 | 64 | 17446 | 3,540179 | 1 | 0,6003384 | 55,055359 |
| Annotation | Enrichment Score: 1.512880938987896 | | | | | | | | | | | |
| Category | Term | Count | % | PValue | Genes | List Total | Pop Hits | Pop Total | Enrichment | Bonferroni | Benjamini | FDR |
| UP_KEYWORDS | Lipid biosynthesis | 9 | 1,867 | 0,01568153 | TM7SF2, EBP, ISYNA1, HSD17B2, ELOVL3, FADS1, H2-KE6, PTDSS1, CHPT1 | 456 | 160 | 22680 | 2,797697 | 0,9935395 | 0,1703426 | 19,199741 |
| UP_KEYWORDS | Steroid biosynthesis | 4 | 0,83 | 0,04031506 | TM7SF2, EBP, HSD17B2, H2-KE6 | 456 | 38 | 22680 | 5,235457 | 0,999998 | 0,292095 | 42,594985 |
| GOTERM_BP_DIRECT | GO:0006694~steroid biosynthetic process | 5 | 1,037 | 0,04576162 | TM7SF2, EBP, HSD17B2, HSD3B5, H2-KE6 | 394 | 62 | 18082 | 3,701081 | 1 | 0,8746985 | 54,904446 |

| | | | | | | | | | | | | |
|---|---|---|---|---|---|---|---|---|---|---|---|---|
| Annotation | Enrichment Score: 1.4736670745056024 | | | | | | | | | | | |
| Category | Term | Count | % | PValue | Genes | List Total | Pop Hits | Pop Total | Enrichment | Bonferroni | Benjamini | FDR |
| INTERPRO | IPR012336:Thioredoxin-like fold | 11 | 2,282 | 6,36E-04 | GSTM1, GPX1, GSTA4, TMX4, PDILT, TXNRD3, EEF1G, GSTP2, MIEN1, GSTP1, GLRX | 436 | 136 | 20594 | 3,820393 | 0,4342769 | 0,0463618 | 0,9865002 |
| GOTERM_BP_DIRECT | GO:0045454~cell redox homeostasis | 5 | 1,037 | 0,05284656 | GPX1, TMX4, PDILT, TXNRD3, GLRX | 394 | 65 | 18082 | 3,530262 | 1 | 0,8720853 | 60,271032 |
| UP_KEYWORDS | Redox-active center | 4 | 0,83 | 0,06804431 | TMX4, TXNRD3, MIEN1, GLRX | 456 | 47 | 22680 | 4,232923 | 1 | 0,3931988 | 61,345479 |
| GOTERM_CC_DIRECT | GO:0005623~cell | 6 | 1,245 | 0,55702658 | FTL1, TMX4, NUP62CL, TXNRD3, FGF21, GLRX | 424 | 231 | 19662 | 1,204484 | 1 | 0,9862506 | 99,998596 |
| Annotation | Enrichment Score: 1.3637497471234148 | | | | | | | | | | | |
| Category | Term | Count | % | PValue | Genes | List Total | Pop Hits | Pop Total | Enrichment | Bonferroni | Benjamini | FDR |
| GOTERM_CC_DIRECT | GO:0005615~extracellular space | 52 | 10,79 | 8,21E-04 | SERPINE2, SERPINA6, CREG1, CES1E, TSKU, MUP12, TLE2, FGF21, C8G, IFNAR2, SERPINA3K, 1190002N15RIK, HSPB1, CTSC, GSTP2, GSTP1, RBP4, CPM, SERPINA12, CCL9, SFTPA1, NRN1, GREM2, ZFC3H1, AKR1A1, SERPINA1C, MRPL18, PCSK6, PLTP, PCSK4, BMP4, MUP5, H2-Q10, | 424 | 1504 | 19662 | 1,603309 | 0,2548563 | 0,0207935 | 1,1213001 |
| UP_KEYWORDS | Secreted | 47 | 9,751 | 0,02135111 | SFTPA1, GREM2, TTR, BDNF, LECT1, SERPINE2, SERPINA6, PLA2G12A, COL27A1, SERPINA1C, CREG1, MUP14, MUP13, TSKU, MUP12, PLTP, BMP4, MUP5, MUP7, MUP6, F10, MUP1, DHRS13, MUP3, FETUB, ENDOD1, FGF21, 1700066M21RIK, C8G, IFNAR2, SERPINA3K, | 456 | 1685 | 22680 | 1,387318 | 0,9989767 | 0,2113288 | 25,256265 |
| GOTERM_CC_DIRECT | GO:0005576~extracellular region | 50 | 10,37 | 0,03505939 | CCL9, HP, GREM2, GSTM1, TTR, BDNF, LECT1, SERPINE2, SERPINA6, COL27A1, PLA2G12A, SERPINA1C, CREG1, PCSK6, MUP14, TSKU, MUP13, MUP12, PLTP, MUP5, BMP4, FZD8, MUP7, MUP6, F10, MUP1, DHRS13, MUP3, FETUB, ENDOD1, FGF21, 1700066M21RIK, C8G, | 424 | 1753 | 19662 | 1,322665 | 0,9999972 | 0,4262163 | 38,721851 |

| | | | | | | | | | | | | |
|---|---|---|---|---|---|---|---|---|---|---|---|---|
| UP_KEYWORDS | Signal | 93 | 19,29 | 0,49453247 | ILDR2, CSPG5, RDH9, MTHFD2, TTR, BDNF, SERPINE2, FXN, SERPINA6, CREG1, CES1E, MUP14, TSKU, MUP13, MUP12, EGFR, F10, CYP2C29, FGF21, UGT1A1, C8G, IFNAR2, SERPINA3K, 1190002N15RIK, GRM8, PDGFRL, SUSD4, CTSC, GNAS, FRAS1, RBP4, CPM, C9, CYP2C44, SERPINA12, TMX4, MAPKAPK5, UGT2B1, FAM98C, CCL9, TMEM219, ACP5, SFTPA1, NRN1, GCGR, GREM2, TMCC1, UGT3A2, DCT, UGT1A9, COL27A1, PLA2G12A, SERPINA1C, PDILT, PCSK6, PCSK4, PLTP, MOCS1, AATK, | 456 | 4543 | 22680 | 1,018166 | 1 | 0,8890228 | 99,989922 |
| UP_SEQ_FEATURE | signal peptide | 72 | 14,94 | 0,49920914 | BDNF, SERPINE2, SERPINA6, CREG1, CES1E, MUP14, TSKU, MUP13, MUP12, EGFR, F10, FGF21, UGT1A1, C8G, IFNAR2, SERPINA3K, 1190002N15RIK, GRM8, PDGFRL, SUSD4, CTSC, FRAS1, RBP4, CPM, C9, SERPINA12, TMX4, CCL9, ACP5, SFTPA1, NRN1, GREM2, GCGR, UGT3A2, DCT, UGT1A9, COL27A1, PLA2G12A, SERPINA1C, PDILT, PCSK4, PLTP, MUP5, BMP4, FZD8, MUP7, | 408 | 3124 | 18012 | 1,017474 | 1 | 1 | 99,998373 |
| Annotation | Enrichment Score: 1.273203396946438 | | | | | | | | | | | |
| Category | Term | Count | % | PValue | Genes | List Total | Pop Hits | Pop Total | Enrichment | Bonferroni | Benjamini | FDR |
| INTERPRO | IPR023795:Protease inhibitor I4, serpin, conserved site | 5 | 1,037 | 0,02396279 | SERPINA3K, SERPINE2, SERPINA12, SERPINA6, SERPINA1C | 436 | 52 | 20594 | 4,541725 | 1 | 0,6272025 | 31,462902 |
| SMART | SM00093:SERPIN | 5 | 1,037 | 0,02407277 | SERPINA3K, SERPINE2, SERPINA12, SERPINA6, SERPINA1C | 172 | 67 | 10425 | 4,523169 | 0,980697 | 0,8610648 | 25,536117 |
| UP_KEYWORDS | Serine protease inhibitor | 6 | 1,245 | 0,02696915 | SERPINA3K, BC048546, MUG1, SERPINE2, SERPINA12, SERPINA1C | 456 | 84 | 22680 | 3,552632 | 0,9998369 | 0,2452225 | 30,840713 |
| UP_KEYWORDS | Protease inhibitor | 7 | 1,452 | 0,03532473 | SERPINA3K, BC048546, MUG1, SERPINE2, SERPINA12, FETUB, SERPINA1C | 456 | 121 | 22680 | 2,877338 | 0,9999896 | 0,2863941 | 38,43536 |
| GOTERM_BP_DIRECT | GO:0010466~negative regulation of peptidase activity | 7 | 1,452 | 0,04252869 | SERPINA3K, BC048546, MUG1, SERPINE2, SERPINA12, FETUB, SERPINA1C | 394 | 117 | 18082 | 2,745759 | 1 | 0,8749692 | 52,23528 |
| INTERPRO | IPR000215:Serpin family | 5 | 1,037 | 0,0532165 | SERPINA3K, SERPINE2, SERPINA12, SERPINA6, SERPINA1C | 436 | 67 | 20594 | 3,524921 | 1 | 0,8588196 | 57,334697 |
| INTERPRO | IPR023796:Serpin domain | 5 | 1,037 | 0,0532165 | SERPINA3K, SERPINE2, SERPINA12, SERPINA6, SERPINA1C | 436 | 67 | 20594 | 3,524921 | 1 | 0,8588196 | 57,334697 |
| GOTERM_MF_DIRECT | GO:0030414~peptidase inhibitor activity | 7 | 1,452 | 0,05461234 | SERPINA3K, BC048546, MUG1, SERPINE2, SERPINA12, FETUB, SERPINA1C | 385 | 123 | 17446 | 2,578862 | 1 | 0,5969294 | 56,619537 |
| GOTERM_MF_DIRECT | GO:0004867~serine-type endopeptidase inhibitor activity | 7 | 1,452 | 0,05461234 | SERPINA3K, BC048546, MUG1, SERPINE2, SERPINA12, SERPINA6, SERPINA1C | 385 | 123 | 17446 | 2,578862 | 1 | 0,5969294 | 56,619537 |
| GOTERM_MF_DIRECT | GO:0004866~endopeptidase inhibitor activity | 3 | 0,622 | 0,14833623 | BC048546, MUG1, SERPINA1C | 385 | 31 | 17446 | 4,385253 | 1 | 0,8521584 | 90,816359 |

| | | | | | | | | | | | | |
|---|---|---|---|---|---|---|---|---|---|---|---|---|
| UP_SEQ_FEATURE | site:Reactive bond | 3 | 0,622 | 0,33740357 | SERPINA3K, SERPINE2, SERPINA1C | 408 | 53 | 18012 | 2,49889 | 1 | 1 | 99,858715 |
| Annotation | Enrichment Score: 1.226082040740489 | | | | | | | | | | | |
| Category | Term | Count | % | PValue | Genes | List Total | Pop Hits | Pop Total | Enrichment | Bonferroni | Benjamini | FDR |
| KEGG_PATHWAY | mmu04915:Estrogen signaling pathway | 10 | 2,075 | 0,00108467 | EGFR, HRAS, ADCY1, ATF4, HSPA2, MAP2K2, GNAS, HSPA1B, ITPR1, ITPR2 | 205 | 98 | 7720 | 3,842708 | 0,2200488 | 0,0223398 | 1,3807558 |
| KEGG_PATHWAY | mmu04912:GnRH signaling pathway | 8 | 1,66 | 0,00845519 | EGFR, HRAS, ADCY1, ATF4, MAP2K2, GNAS, ITPR1, ITPR2 | 205 | 88 | 7720 | 3,423503 | 0,856937 | 0,114435 | 10,307602 |
| KEGG_PATHWAY | mmu04918:Thyroid hormone synthesis | 7 | 1,452 | 0,01018545 | GPX1, ADCY1, ATF4, GNAS, ATP1A1, ITPR1, ITPR2 | 205 | 70 | 7720 | 3,765854 | 0,9040978 | 0,1160819 | 12,29225 |
| BIOCARTA | arrestin-dependent Recruitment of Src Kinases in GPCR Signaling | 4 | 0,83 | 0,01249843 | HRAS, ADCY1, MAP2K2, GNAS | 32 | 21 | 1289 | 7,672619 | 0,7645782 | 0,5147972 | 13,347507 |
| KEGG_PATHWAY | mmu04540:Gap junction | 7 | 1,452 | 0,02587746 | EGFR, HRAS, ADCY1, MAP2K2, GNAS, ITPR1, ITPR2 | 205 | 86 | 7720 | 3,06523 | 0,997531 | 0,2213277 | 28,530119 |
| BIOCARTA | m_erkPathway:Erk1/Erk2 Mapk Signaling pathway | 4 | 0,83 | 0,04219162 | EGFR, HRAS, MAP2K2, GNAS | 32 | 33 | 1289 | 4,882576 | 0,9929686 | 0,7104253 | 38,800429 |
| BIOCARTA | :Role of ï¿½-arrestins in the activation and targeting of MAP kinases | 3 | 0,622 | 0,06084226 | ADCY1, MAP2K2, GNAS | 32 | 17 | 1289 | 7,108456 | 0,9992673 | 0,6997461 | 51,081875 |
| KEGG_PATHWAY | mmu04925:Aldosterone synthesis and secretion | 6 | 1,245 | 0,07680374 | ADCY1, ATF4, HSD3B5, GNAS, ITPR1, ITPR2 | 205 | 86 | 7720 | 2,62734 | 1 | 0,4256698 | 64,077661 |
| KEGG_PATHWAY | mmu04730:Long-term depression | 5 | 1,037 | 0,07708824 | HRAS, MAP2K2, GNAS, ITPR1, ITPR2 | 205 | 61 | 7720 | 3,086765 | 1 | 0,4174361 | 64,219229 |
| BIOCARTA | m_crebPathway:Transcription factor CREB and its extracellular signals | 3 | 0,622 | 0,11928644 | HRAS, ADCY1, GNAS | 32 | 25 | 1289 | 4,83375 | 0,9999995 | 0,8759169 | 76,469988 |
| KEGG_PATHWAY | mmu04270:Vascular smooth muscle contraction | 7 | 1,452 | 0,11949698 | ADCY1, MAP2K2, PLA2G12A, AVPR1A, GNAS, ITPR1, ITPR2 | 205 | 127 | 7720 | 2,075667 | 1 | 0,5174036 | 80,415191 |
| KEGG_PATHWAY | mmu04916:Melanogenesis | 6 | 1,245 | 0,12089612 | DCT, FZD8, HRAS, ADCY1, MAP2K2, GNAS | 205 | 99 | 7720 | 2,282336 | 1 | 0,5130952 | 80,810173 |
| BIOCARTA | m_mPRpathway:How Progesterone Initiates the Oocyte Maturation | 3 | 0,622 | 0,16080269 | HRAS, ADCY1, GNAS | 32 | 30 | 1289 | 4,028125 | 1 | 0,9195463 | 86,424683 |
| BIOCARTA | m_gpcrPathway:Signaling Pathway from G-Protein Families | 3 | 0,622 | 0,19571538 | HRAS, ADCY1, GNAS | 32 | 34 | 1289 | 3,554228 | 1 | 0,9183018 | 91,633569 |
| KEGG_PATHWAY | mmu04921:Oxytocin signaling pathway | 7 | 1,452 | 0,23953149 | EGFR, HRAS, ADCY1, MAP2K2, GNAS, ITPR1, ITPR2 | 205 | 158 | 7720 | 1,668416 | 1 | 0,6736323 | 97,004479 |
| KEGG_PATHWAY | mmu05200:Pathways in cancer | 12 | 2,49 | 0,48044923 | EGFR, BMP4, CUL2, FZD8, HRAS, ADCY1, EPAS1, MAP2K2, PPARG, GNAS, FGF21, TRAF4 | 205 | 397 | 7720 | 1,138293 | 1 | 0,8145182 | 99,977261 |
| KEGG_PATHWAY | mmu04015:Rap1 signaling pathway | 7 | 1,452 | 0,50042306 | EGFR, HRAS, ADCY1, MAP2K2, P2RY1, GNAS, FGF21 | 205 | 214 | 7720 | 1,231821 | 1 | 0,8256035 | 99,986239 |
| Annotation | Enrichment Score: 1.1634325547010769 | | | | | | | | | | | |
| Category | Term | Count | % | PValue | Genes | List Total | Pop Hits | Pop Total | Enrichment | Bonferroni | Benjamini | FDR |

| | | | | | | | | | | | | |
|---|---|---|---|---|---|---|---|---|---|---|---|---|
| GOTERM_MF_DIRECT | GO:0050660~flavin adenine dinucleotide binding | 6 | 1,245 | 0,0211584 | MAOA, AOX1, TXNRD3, GULO, DUS1L, MTO1 | 385 | 72 | 17446 | 3,77619 | 0,9999986 | 0,3720645 | 27,241459 |
| INTERPRO | oxidoreductase, FAD/NAD(P)-binding domain | 5 | 1,037 | 0,04410993 | MAOA, MICAL2, TXNRD3, OSGIN1, MTO1 | 436 | 63 | 20594 | 3,748726 | 1 | 0,8271721 | 50,474431 |
| UP_KEYWORDS | Flavoprotein | 7 | 1,452 | 0,04733517 | MAOA, MICAL2, AOX1, TXNRD3, GULO, DUS1L, MTO1 | 456 | 130 | 22680 | 2,678138 | 0,9999998 | 0,3142845 | 48,007295 |
| UP_KEYWORDS | FAD | 6 | 1,245 | 0,08924864 | MAOA, MICAL2, AOX1, TXNRD3, GULO, MTO1 | 456 | 118 | 22680 | 2,528992 | 1 | 0,462748 | 71,661241 |
| UP_SEQ_FEATURE | nucleotide phosphate-binding region:FAD | 3 | 0,622 | 0,38639597 | MICAL2, TXNRD3, MTO1 | 408 | 59 | 18012 | 2,244766 | 1 | 1 | 99,958485 |
| Annotation | Enrichment Score: 1.1302971409373632 | | | | | | | | | | | |
| Category | Term | Count | % | PValue | Genes | List Total | Pop Hits | Pop Total | Enrichment | Bonferroni | Benjamini | FDR |
| UP_SEQ_FEATURE | metal ion-binding site:Iron | 6 | 1,245 | 0,00677588 | PHYHD1, FTL1, HGD, PAH, PHYH, HPD | 408 | 53 | 18012 | 4,99778 | 0,9993259 | 0,5557377 | 10,273005 |
| UP_KEYWORDS | Dioxygenase | 4 | 0,83 | 0,23252216 | PHYHD1, HGD, PHYH, HPD | 456 | 83 | 22680 | 2,396956 | 1 | 0,7326224 | 97,183196 |
| GOTERM_MF_DIRECT | GO:0051213~dioxygenase activity | 4 | 0,83 | 0,25803499 | PHYHD1, HGD, PHYH, HPD | 385 | 80 | 17446 | 2,265714 | 1 | 0,9321423 | 98,818412 |
| Annotation | Enrichment Score: 1.1294683696003773 | | | | | | | | | | | |
| Category | Term | Count | % | PValue | Genes | List Total | Pop Hits | Pop Total | Enrichment | Bonferroni | Benjamini | FDR |
| UP_KEYWORDS | Differentiation | 23 | 4,772 | 0,01136152 | PAQR7, CSPG5, HES6, CSRP2, SIRT2, LECT1, SERPINE2, HSPA2, NME1, NTRK2, GADD45G, PDILT, TXNRD3, STRBP, ZPR1, BIN1, BMP7 | 456 | 646 | 22680 | 1,770816 | 0,9738804 | 0,1356719 | 14,283296 |
| GOTERM_BP_DIRECT | GO:0030154~cell differentiation | 24 | 4,979 | 0,08751409 | PAQR7, ILDR2, CSPG5, HES6, CSRP2, SIRT2, LECT1, SERPINE2, HSPA2, NME1, NTRK2, GADD45G, PDILT, TXNRD3, STRBP, ZPR1, BIN1, | 394 | 780 | 18082 | 1,412105 | 1 | 0,9433375 | 78,924404 |
| UP_KEYWORDS | Developmental protein | 26 | 5,394 | 0,1262261 | SERPINE2, PDILT, ZFP354A, STRBP, ZPR1, MN1, TRAF4, BMP4, EGFR, FZD8, EPAS1, MTL5, HES6, CSRP2, PTP4A1, NTRK2, GADD45G, TXNRD3, | 456 | 976 | 22680 | 1,324957 | 1 | 0,5700114 | 83,797397 |
| GOTERM_BP_DIRECT | GO:0007275~multicellular organism development | 27 | 5,602 | 0,24180376 | SERPINE2, PDILT, ZFP354A, STRBP, ZPR1, MN1, TRAF4, BMP4, EGFR, FZD8, EPAS1, GTF2IRD2, MTL5, HES6, CSRP2, PTP4A1, NTRK2, GADD45G, | 394 | 1029 | 18082 | 1,2042 | 1 | 0,994995 | 99,09616 |
| Annotation | Enrichment Score: 1.0339983382940998 | | | | | | | | | | | |
| Category | Term | Count | % | PValue | Genes | List Total | Pop Hits | Pop Total | Enrichment | Bonferroni | Benjamini | FDR |
| SMART | SM00261:FU | 3 | 0,622 | 0,0309546 | EGFR, FRAS1, PCSK6 | 172 | 17 | 10425 | 10,69596 | 0,9938659 | 0,8169446 | 31,647078 |
| INTERPRO | IPR006212:Furin-like repeat | 3 | 0,622 | 0,04911713 | EGFR, FRAS1, PCSK6 | 436 | 17 | 20594 | 8,335402 | 1 | 0,8471307 | 54,364674 |

| | | | | | | | | | | | | |
|---|---|---|---|---|---|---|---|---|---|---|---|---|
| INTERPRO | IPR009030:Insulin-like growth factor binding protein, N-terminal | 4 | 0,83 | 0,52005195 | EGFR, FRAS1, C9, PCSK6 | 436 | 130 | 20594 | 1,453352 | 1 | 0,9999985 | 99,998918 |
| Annotation | Enrichment Score: 1.0292516598556236 | | | | | | | | | | | |
| Category | Term | Count | % | PValue | Genes | List Total | Pop Hits | Pop Total | Enrichment | Bonferroni | Benjamini | FDR |
| UP_KEYWORDS | Fatty acid metabolism | 8 | 1,66 | 0,01263879 | ACSM2, BAAT, ELOVL3, FADS1, ECHDC2, H2-KE6, ACAA1B, SLC27A2 | 456 | 124 | 22680 | 3,208829 | 0,9827073 | 0,1444892 | 15,765005 |
| GOTERM_BP_DIRECT | GO:0006631~fatty acid metabolic process | 9 | 1,867 | 0,02088181 | ACSM2, CRYL1, BAAT, ELOVL3, FADS1, ECHDC2, H2-KE6, ACAA1B, SLC27A2 | 394 | 156 | 18082 | 2,647696 | 1 | 0,7216391 | 30,147482 |
| KEGG_PATHWAY | mmu01040:Biosynthesis of unsaturated fatty acids | 3 | 0,622 | 0,15892786 | BAAT, FADS1, ACAA1B | 205 | 27 | 7720 | 4,184282 | 1 | 0,5546407 | 89,110668 |
| GOTERM_BP_DIRECT | GO:0006633~fatty acid biosynthetic process | 4 | 0,83 | 0,21744384 | ACSM2, ELOVL3, FADS1, H2-KE6 | 394 | 74 | 18082 | 2,480724 | 1 | 0,9921336 | 98,452645 |
| GOTERM_BP_DIRECT | GO:0006635~fatty acid beta-oxidation | 3 | 0,622 | 0,24795107 | ECHDC2, ACAA1B, SLC27A2 | 394 | 44 | 18082 | 3,129096 | 1 | 0,9954667 | 99,212988 |
| UP_KEYWORDS | Fatty acid biosynthesis | 3 | 0,622 | 0,29519108 | ELOVL3, FADS1, H2-KE6 | 456 | 54 | 22680 | 2,763158 | 1 | 0,769697 | 99,107164 |
| Annotation | Enrichment Score: 1.0213030076100034 | | | | | | | | | | | |
| Category | Term | Count | % | PValue | Genes | List Total | Pop Hits | Pop Total | Enrichment | Bonferroni | Benjamini | FDR |
| GOTERM_CC_DIRECT | GO:0005913~cell-cell adherens junction | 12 | 2,49 | 0,08117696 | EGFR, COBLL1, CADM4, RPL14, VAPB, BAG3, EIF5, OLA1, EEF1G, RPL24, NOP56, RPS2 | 424 | 316 | 19662 | 1,760986 | 1 | 0,6358891 | 68,707461 |
| GOTERM_MF_DIRECT | GO:0098641~cadherin binding involved in cell-cell adhesion | 11 | 2,282 | 0,09030849 | EGFR, COBLL1, RPL14, VAPB, BAG3, EIF5, OLA1, EEF1G, RPL24, NOP56, RPS2 | 385 | 279 | 17446 | 1,786585 | 1 | 0,750658 | 75,525548 |
| GOTERM_BP_DIRECT | GO:0098609~cell-cell adhesion | 8 | 1,66 | 0,11774151 | COBLL1, RPL14, VAPB, BAG3, EIF5, OLA1, EEF1G, RPS2 | 394 | 189 | 18082 | 1,942578 | 1 | 0,9613592 | 88,113868 |
| Annotation | Enrichment Score: 1.0163250460552116 | | | | | | | | | | | |
| Category | Term | Count | % | PValue | Genes | List Total | Pop Hits | Pop Total | Enrichment | Bonferroni | Benjamini | FDR |
| UP_SEQ_FEATURE | domain:ABC transporter | 4 | 0,83 | 0,00515225 | ABCG8, ABCB10, ABCB6, ABCG2 | 408 | 16 | 18012 | 11,03676 | 0,996104 | 0,5473089 | 7,9056782 |
| GOTERM_MF_DIRECT | GO:0016887~ATPase activity | 10 | 2,075 | 0,03344449 | KIFC2, ABCG8, ATP5J2, VCP, CLPB, OLA1, ABCB10, ABCB6, AK6, ABCG2 | 385 | 200 | 17446 | 2,265714 | 1 | 0,4681039 | 39,701414 |
| SMART | SM00382:AAA | 6 | 1,245 | 0,08810521 | VCP, NUBP1, CLPB, ABCB10, ABCB6, ABCG2 | 172 | 144 | 10425 | 2,525436 | 0,9999997 | 0,976135 | 67,242458 |
| GOTERM_MF_DIRECT | GO:0042626~ATPase activity, coupled to transmembrane movement of substances | 4 | 0,83 | 0,09298796 | ABCG8, ABCB10, ABCB6, ABCG2 | 385 | 49 | 17446 | 3,699125 | 1 | 0,7533208 | 76,57596 |
| INTERPRO | IPR003439:ABC transporter-like | 4 | 0,83 | 0,10136302 | ABCG8, ABCB10, ABCB6, ABCG2 | 436 | 53 | 20594 | 3,564826 | 1 | 0,9710662 | 81,0758 |
| KEGG_PATHWAY | mmu02010:ABC transporters | 4 | 0,83 | 0,12103629 | ABCG8, ABCB10, ABCB6, ABCG2 | 205 | 46 | 7720 | 3,274655 | 1 | 0,5051105 | 80,849336 |

| | | | | | | | | | | | | |
|---|---|---|---|---|---|---|---|---|---|---|---|---|
| INTERPRO | IPR003593:AAA+ ATPase domain | 6 | 1,245 | 0,18943775 | VCP, NUBP1, CLPB, ABCB10, ABCB6, ABCG2 | 436 | 144 | 20594 | 1,968081 | 1 | 0,9971889 | 96,204803 |
| INTERPRO | IPR017871:ABC transporter, conserved site | 3 | 0,622 | 0,26938764 | ABCG8, ABCB10, ABCB6 | 436 | 48 | 20594 | 2,952122 | 1 | 0,9993842 | 99,247068 |
| INTERPRO | IPR027417:P-loop containing nucleoside triphosphate hydrolase | 17 | 3,527 | 0,80665119 | ABCB6, AK6, ABCG2, ABCG8, VCP, NUBP1, RAB22A, ABCB10, GNAS, RAB38 | 436 | 909 | 20594 | 0,883363 | 1 | 1 | 100 |
| Annotation | Enrichment Score: 0.9853303029895368 | | | | | | | | | | | |
| Category | Term | Count | % | PValue | Genes | List Total | Pop Hits | Pop Total | Enrichment | Bonferroni | Benjamini | FDR |
| GOTERM_CC_DIRECT | GO:0016020~membrane | 193 | 40,04 | 1,73E-05 | SLC52A2, FAM210A, AI314180, TMEM147, SERPINE2, RPLP0, RPL10, ABCB10, RPL11, TMEM14C, F10, TTC7B, OLA1, COLEC12, CYP1A2, CYP2E1, RPS18, RCC2, NME1, RPS14, RPS15, CYP2D26, RPS10, BIN1, GCNT4, CYP3A25, TMX4, MME, NRN1, TMCC1, RPS23, RPSA, NOC4L, CYP46A1, GDE1, MAP2K2, MAOA, RPS6, RPS8, ABCG2, ABCG8, NAA60, NTRK2, AVPR1A, CYP2R1, TMEM41A, CNIH1, SLC46A3, TM7SF2, CPEB1, DNAJC18, CYP39A1, MAP1LC3A, FITM1, SLC25A3, ERRFI1, ADCK5, RAP2A, CYP2C29, CYB5B, SLC3A1, MIEN1, NDUFA11, ZDHHC14, CYP27A1, GRM8, RIPK1, EEF1G, MOSPD3, GNAS, SUSD4, SLC27A2, PTOV1, C9, CYP2F2, HSD17B2, HSD3B5, RPL35, STARD10, SLC19A1, GCGR, LECT1, PLIN4, PCSK4, MARS, AATK, FZD8, DLGAP1, H2-Q10, COX7A2, FADS1, RPL24, ATP1A1, TMEM5, MARCH5, ITPR1, RPL28, ITPR2, MARCH2, GBA2, PTP4A1, RAB22A, DYM, NOP56, RAB38, SSR4, SLC15A5, KCNC3, PDE3B, ILDR2, CSPG5, COX5A, C2CD2L, DIRC2, ATAT1, TIAM2, H13, ELOVL3, SLC25A23, TIMM9, SLC25A28, MCOLN1, EGFR, CMAS, SMIM13, PNPLA1, CHPT1, OGFRL1, MARC1, SLC25A38, GULO, FRAS1, SLC38A2, COX7C, TMEM219, PAQR7, P2RY1, TRAF4, REEP6, ATP5J2, EPHX1, ENDOD1, ABCB6, AK6, NDUFV3, ATF4, SLC35G1, RNF152, PPIA, LRIT2, CYC1, RPS2, POMT2, PLEKHB1, TMEM57, PTDSS1, HPD, IRAK1, MPP6, COX4I1, UGT1A1, IFNAR2, DGAT1, SLC35E3, EMC6, H2-KE6, CTSC, NDUFB3, CPM, FTL1, DTNBP1, | 424 | 6998 | 19662 | 1,278925 | 0,0061748 | 6,19E-04 | 0,0237397 |

| | | | | | | | | | | | | |
|---|---|---|---|---|---|---|---|---|---|---|---|---|
| UP_SEQ_FEATURE | transmembrane region | 103 | 21,37 | 0,31377845 | CSPG5, SLC52A2, FAM210A, C2CD2L, TMEM147, DIRC2, H13, ELOVL3, SLC25A23, SLC25A28, ABCB10, MCOLN1, TMEM14C, EGFR, COLEC12, CHPT1, SLC25A38, SLC25A37, GULO, FRAS1, SLC38A2, TMX4, COX7C, MME, PAQR7, TMEM219, TMCC1, P2RY1, REEP6, CYP46A1, GDE1, NOC4L, MAOA, EPHX1, ABCB6, ABCG2, ABCG8, SLC35G1, NTRK2, AVPR1A, TMEM41A, CNIH1, SLC46A3, TM7SF2, LRIT2, CYC1, POMT2, DNAJC18, TMEM57, FITM1, SLC25A3, PTDSS1, ADCK5, CYB5B, UGT1A1, NDUFA11, IFNAR2, ZDHHC14, DGAT1, GRM8, SLC35E3, EMC6, MOSPD3, SUSD4, SLC27A2, NDUFB3, C9, | 408 | 4312 | 18012 | 1,054533 | 1 | 1 | 99,753011 |
| UP_KEYWORDS | Membrane | 176 | 36,51 | 0,48673105 | PDE3B, ILDR2, CSPG5, SLC52A2, COX5A, FAM210A, C2CD2L, TMEM147, DIRC2, ATAT1, H13, ELOVL3, SLC25A23, TIMM9, RPL10, SLC25A28, ABCB10, MCOLN1, TMEM14C, EGFR, SLC22A28, SMIM13, TTC7B, COLEC12, CYP1A2, CYP2E1, CHPT1, ZFP592, MARC1, SLC25A38, SLC25A37, CYP2D26, GULO, FRAS1, GCNT4, SLC38A2, CYP3A25, TMX4, COX7C, TMEM219, PAQR7, MME, NRN1, SLC22A30, TMCC1, P2RY1, STRBP, TRAF4, REEP6, RPSA, ATP5J2, CYP2D40, NOC4L, CYP46A1, GDE1, MAP2K2, MAOA, EPHX1, ENDOD1, ABCB6, RPS8, ABCG2, ABCG8, NDUFV3, ATF4, UGT2B35, SLC35G1, RNF152, NTRK2, NAA60, AVPR1A, FBXO31, CYP2R1, TMEM41A, CNIH1, SLC46A3, TM7SF2, LRIT2, CYC1, GM10658, CPEB1, DNAJC18, POMT2, PLEKHB1, CYP39A1, TMEM57, MAP1LC3A, 2310061I04RIK, FITM1, SLC25A3, PTDSS1, | 456 | 8683 | 22680 | 1,008141 | 1 | 0,8859374 | 99,987609 |

| | | | | | | | | | | | | |
|---|---|---|---|---|---|---|---|---|---|---|---|---|
| UP_KEYWORDS | Transmembrane helix | 135 | 28,01 | 0,71915555 | ILDR2, CSPG5, SLC52A2, FAM210A, C2CD2L, TMEM147, DIRC2, H13, ELOVL3, SLC25A23, RPL10, SLC25A28, ABCB10, MCOLN1, TMEM14C, EGFR, SLC22A28, SMIM13, COLEC12, CYP2E1, CHPT1, ZFP592, MARC1, SLC25A38, SLC25A37, CYP2D26, GULO, FRAS1, GCNT4, SLC38A2, CYP3A25, TMX4, COX7C, PAQR7, TMEM219, MME, NRN1, SLC22A30, TMCC1, P2RY1, STRBP, REEP6, ATP5J2, CYP2D40, CYP46A1, GDE1, NOC4L, MAOA, EPHX1, ENDOD1, ABCB6, ABCG2, ABCG8, UGT2B35, SLC35G1, RNF152, NTRK2, AVPR1A, FBXO31, CYP2R1, TMEM41A, CNIH1, SLC46A3, TM7SF2, LRIT2, CYC1, GM10658, DNAJC18, POMT2, TMEM57, 2310061I04RIK, FITM1, SLC25A3, PTDSS1, ADCK5, COX4I1, CYB5B, SLC3A1, UGT1A1, NDUFA11, IFNAR2, ZDHHC14, DGAT1, GRM8, SLC35E3, EMC6, | 456 | 6938 | 22680 | 0,967782 | 1 | 0,966771 | 99,999996 |
| UP_KEYWORDS | Transmembrane | 135 | 28,01 | 0,73080521 | ILDR2, CSPG5, SLC52A2, FAM210A, C2CD2L, TMEM147, DIRC2, H13, ELOVL3, SLC25A23, RPL10, SLC25A28, ABCB10, MCOLN1, TMEM14C, EGFR, SLC22A28, SMIM13, COLEC12, CYP2E1, CHPT1, ZFP592, MARC1, SLC25A38, SLC25A37, CYP2D26, GULO, FRAS1, GCNT4, SLC38A2, CYP3A25, TMX4, COX7C, PAQR7, TMEM219, MME, NRN1, SLC22A30, TMCC1, P2RY1, STRBP, REEP6, ATP5J2, CYP2D40, CYP46A1, GDE1, NOC4L, MAOA, EPHX1, ENDOD1, ABCB6, ABCG2, ABCG8, UGT2B35, SLC35G1, RNF152, NTRK2, AVPR1A, FBXO31, CYP2R1, TMEM41A, CNIH1, SLC46A3, TM7SF2, LRIT2, CYC1, GM10658, DNAJC18, POMT2, TMEM57, 2310061I04RIK, FITM1, SLC25A3, PTDSS1, ADCK5, COX4I1, CYB5B, SLC3A1, UGT1A1, NDUFA11, IFNAR2, ZDHHC14, DGAT1, GRM8, SLC35E3, EMC6, | 456 | 6955 | 22680 | 0,965417 | 1 | 0,9694559 | 99,999998 |

| | | | | | | | | | | | | |
|---|---|---|---|---|---|---|---|---|---|---|---|---|
| GOTERM_CC_DIRECT | GO:0016021~integral component of membrane | 138 | 28,63 | 0,8818599 | ILDR2, CSPG5, SLC52A2, FAM210A, C2CD2L, TMEM147, DIRC2, H13, ELOVL3, SLC25A23, SLC25A28, ABCB10, MCOLN1, TMEM14C, EGFR, SLC22A28, SMIM13, COLEC12, CYP2E1, CHPT1, ZFP592, MARC1, SLC25A38, SLC25A37, CYP2D26, GULO, FRAS1, GCNT4, SLC38A2, CYP3A25, TMX4, COX7C, PAQR7, TMEM219, MME, NRN1, SLC22A30, TMCC1, P2RY1, STRBP, REEP6, ATP5J2, CYP2D40, CYP46A1, GDE1, NOC4L, MAOA, EPHX1, ENDOD1, ABCB6, ABCG2, ABCG8, UGT2B35, SLC35G1, RNF152, NTRK2, AVPR1A, FBXO31, CYP2R1, TMEM41A, CNIH1, SLC46A3, TM7SF2, LRIT2, CYC1, GM10658, DNAJC18, POMT2, PLEKHB1, TMEM57, 2310061I04RIK, FITM1, SLC25A3, PTDSS1, ADCK5, COX4I1, MPP6, CYB5B, SLC3A1, UGT1A1, NDUFA11, IFNAR2, ZDHHC14, DGAT1, GRM8, SLC35E3, EMC6, | 424 | 6878 | 19662 | 0,930419 | 1 | 0,9997957 | 100 |
| Annotation | Enrichment Score: 0.968197538243669 | | | | | | | | | | | |
| Category | Term | Count | % | PValue | Genes | List Total | Pop Hits | Pop Total | Enrichment | Bonferroni | Benjamini | FDR |
| UP_KEYWORDS | Lipid biosynthesis | 9 | 1,867 | 0,01568153 | TM7SF2, EBP, ISYNA1, HSD17B2, ELOVL3, FADS1, H2-KE6, PTDSS1, CHPT1 | 456 | 160 | 22680 | 2,797697 | 0,9935395 | 0,1703426 | 19,199741 |
| GOTERM_BP_DIRECT | GO:0008654~phospholipid biosynthetic process | 4 | 0,83 | 0,13674018 | ISYNA1, FITM1, PTDSS1, CHPT1 | 394 | 59 | 18082 | 3,111417 | 1 | 0,9689346 | 91,790783 |
| UP_KEYWORDS | Phospholipid biosynthesis | 3 | 0,622 | 0,23554613 | ISYNA1, PTDSS1, CHPT1 | 456 | 46 | 22680 | 3,243707 | 1 | 0,7323767 | 97,329267 |
| UP_KEYWORDS | Phospholipid metabolism | 3 | 0,622 | 0,26536845 | ISYNA1, PTDSS1, CHPT1 | 456 | 50 | 22680 | 2,984211 | 1 | 0,7596664 | 98,438532 |
| Annotation | Enrichment Score: 0.9158984415937792 | | | | | | | | | | | |
| Category | Term | Count | % | PValue | Genes | List Total | Pop Hits | Pop Total | Enrichment | Bonferroni | Benjamini | FDR |
| GOTERM_BP_DIRECT | GO:0008202~steroid metabolic process | 6 | 1,245 | 0,0361474 | TM7SF2, EBP, CYP39A1, CYP46A1, CYP2E1, CYP1A2 | 394 | 84 | 18082 | 3,2781 | 1 | 0,8442111 | 46,524542 |
| UP_KEYWORDS | Steroid metabolism | 5 | 1,037 | 0,0615115 | TM7SF2, EBP, CYP39A1, CYP46A1, CYP1A2 | 456 | 74 | 22680 | 3,360597 | 1 | 0,3756023 | 57,526409 |
| UP_KEYWORDS | Sterol metabolism | 4 | 0,83 | 0,12818879 | TM7SF2, EBP, CYP46A1, CYP1A2 | 456 | 62 | 22680 | 3,208829 | 1 | 0,5689629 | 84,281467 |
| UP_KEYWORDS | Cholesterol metabolism | 3 | 0,622 | 0,30262499 | TM7SF2, EBP, CYP46A1 | 456 | 55 | 22680 | 2,712919 | 1 | 0,7753515 | 99,226146 |
| GOTERM_BP_DIRECT | GO:0008203~cholesterol metabolic process | 4 | 0,83 | 0,30529493 | TM7SF2, EBP, CYP46A1, CYP27A1 | 394 | 89 | 18082 | 2,062625 | 1 | 0,9972955 | 99,795662 |

| Annotation | Enrichment Score: 0.8696037853560504 | | | | | | | | | | | |
|---|---|---|---|---|---|---|---|---|---|---|---|---|
| Category | Term | Count | % | PValue | Genes | List Total | Pop Hits | Pop Total | Enrichment | Bonferroni | Benjamini | FDR |
| KEGG_PATHWAY | mmu04915:Estrogen signaling pathway | 10 | 2,075 | 0,00108467 | EGFR, HRAS, ADCY1, ATF4, HSPA2, MAP2K2, GNAS, HSPA1B, ITPR1, ITPR2 | 205 | 98 | 7720 | 3,842708 | 0,2200488 | 0,0223398 | 1,3807558 |
| KEGG_PATHWAY | mmu04912:GnRH signaling pathway | 8 | 1,66 | 0,00845519 | EGFR, HRAS, ADCY1, ATF4, MAP2K2, GNAS, ITPR1, ITPR2 | 205 | 88 | 7720 | 3,423503 | 0,856937 | 0,114435 | 10,307602 |
| KEGG_PATHWAY | mmu04918:Thyroid hormone synthesis | 7 | 1,452 | 0,01018545 | GPX1, ADCY1, ATF4, GNAS, ATP1A1, ITPR1, ITPR2 | 205 | 70 | 7720 | 3,765854 | 0,9040978 | 0,1160819 | 12,29225 |
| KEGG_PATHWAY | mmu04726:Serotonergic synapse | 9 | 1,867 | 0,02341623 | HRAS, CYP2D40, CYP2C44, MAOA, CYP2C29, CYP2D26, GNAS, ITPR1, ITPR2 | 205 | 132 | 7720 | 2,567627 | 0,9955998 | 0,210154 | 26,181828 |
| KEGG_PATHWAY | mmu04540:Gap junction | 7 | 1,452 | 0,02587746 | EGFR, HRAS, ADCY1, MAP2K2, GNAS, ITPR1, ITPR2 | 205 | 86 | 7720 | 3,06523 | 0,997531 | 0,2213277 | 28,530119 |
| KEGG_PATHWAY | mmu04720:Long-term potentiation | 6 | 1,245 | 0,02984439 | HRAS, ADCY1, ATF4, MAP2K2, ITPR1, ITPR2 | 205 | 66 | 7720 | 3,423503 | 0,9990302 | 0,2423524 | 32,1705 |
| KEGG_PATHWAY | mmu04922:Glucagon signaling pathway | 7 | 1,452 | 0,04866142 | ATF4, PHKG2, PDE3B, GNAS, GCGR, ITPR1, ITPR2 | 205 | 100 | 7720 | 2,636098 | 0,9999891 | 0,3449874 | 47,223536 |
| KEGG_PATHWAY | mmu04925:Aldosterone synthesis and secretion | 6 | 1,245 | 0,07680374 | ADCY1, ATF4, HSD3B5, GNAS, ITPR1, ITPR2 | 205 | 86 | 7720 | 2,62734 | 1 | 0,4256698 | 64,077661 |
| KEGG_PATHWAY | mmu04730:Long-term depression | 5 | 1,037 | 0,07708824 | HRAS, MAP2K2, GNAS, ITPR1, ITPR2 | 205 | 61 | 7720 | 3,086765 | 1 | 0,4174361 | 64,219229 |
| KEGG_PATHWAY | mmu04724:Glutamatergic synapse | 7 | 1,452 | 0,08363644 | ADCY1, DLGAP1, SLC38A2, GRM8, GNAS, ITPR1, ITPR2 | 205 | 115 | 7720 | 2,292259 | 1 | 0,4262679 | 67,338824 |
| KEGG_PATHWAY | mmu04270:Vascular smooth muscle contraction | 7 | 1,452 | 0,11949698 | ADCY1, MAP2K2, PLA2G12A, AVPR1A, GNAS, ITPR1, ITPR2 | 205 | 127 | 7720 | 2,075667 | 1 | 0,5174036 | 80,415191 |
| KEGG_PATHWAY | mmu04971:Gastric acid secretion | 5 | 1,037 | 0,12229767 | ADCY1, GNAS, ATP1A1, ITPR1, ITPR2 | 205 | 72 | 7720 | 2,615176 | 1 | 0,5007809 | 81,198462 |
| KEGG_PATHWAY | mmu04972:Pancreatic secretion | 6 | 1,245 | 0,12469795 | ADCY1, PLA2G12A, GNAS, ATP1A1, ITPR1, ITPR2 | 205 | 100 | 7720 | 2,259512 | 1 | 0,492252 | 81,846657 |
| KEGG_PATHWAY | mmu04728:Dopaminergic synapse | 7 | 1,452 | 0,14343964 | ATF4, PPP2R5B, MAOA, GNAS, ITPR1, PPP2R2A, ITPR2 | 205 | 134 | 7720 | 1,967237 | 1 | 0,5297011 | 86,242838 |
| KEGG_PATHWAY | mmu04970:Salivary secretion | 5 | 1,037 | 0,14587869 | ADCY1, GNAS, ATP1A1, ITPR1, ITPR2 | 205 | 77 | 7720 | 2,44536 | 1 | 0,5287067 | 86,736354 |
| KEGG_PATHWAY | mmu04921:Oxytocin signaling pathway | 7 | 1,452 | 0,23953149 | EGFR, HRAS, ADCY1, MAP2K2, GNAS, ITPR1, ITPR2 | 205 | 158 | 7720 | 1,668416 | 1 | 0,6736323 | 97,004479 |
| KEGG_PATHWAY | mmu04750:Inflammatory mediator regulation of TRP channels | 6 | 1,245 | 0,24006967 | ADCY1, CYP2C44, CYP2C29, GNAS, ITPR1, ITPR2 | 205 | 126 | 7720 | 1,793264 | 1 | 0,6681029 | 97,031525 |
| KEGG_PATHWAY | mmu04611:Platelet activation | 6 | 1,245 | 0,26503559 | ADCY1, COL27A1, P2RY1, GNAS, ITPR1, ITPR2 | 205 | 131 | 7720 | 1,724818 | 1 | 0,6973561 | 98,065045 |
| KEGG_PATHWAY | mmu04924:Renin secretion | 4 | 0,83 | 0,28919035 | PDE3B, GNAS, ITPR1, ITPR2 | 205 | 71 | 7720 | 2,121608 | 1 | 0,7223694 | 98,738934 |
| KEGG_PATHWAY | mmu04022:cGMP-PKG signaling pathway | 7 | 1,452 | 0,29813994 | ADCY1, ATF4, MAP2K2, PDE3B, ATP1A1, ITPR1, ITPR2 | 205 | 171 | 7720 | 1,541578 | 1 | 0,7238577 | 98,927891 |

| | | | | | | | | | | | | |
|---|---|---|---|---|---|---|---|---|---|---|---|---|
| KEGG_PATHWAY | mmu04114:Oocyte meiosis | 5 | 1,037 | 0,33138908 | ADCY1, PPP2R5B, CPEB1, ITPR1, ITPR2 | 205 | 110 | 7720 | 1,711752 | 1 | 0,752597 | 99,424282 |
| KEGG_PATHWAY | mmu04020:Calcium signaling pathway | 7 | 1,452 | 0,3405431 | EGFR, ADCY1, PHKG2, AVPR1A, GNAS, ITPR1, ITPR2 | 205 | 180 | 7720 | 1,464499 | 1 | 0,7539158 | 99,517489 |
| KEGG_PATHWAY | mmu04725:Cholinergic synapse | 5 | 1,037 | 0,34950637 | HRAS, ADCY1, ATF4, ITPR1, ITPR2 | 205 | 113 | 7720 | 1,666307 | 1 | 0,7550723 | 99,595085 |
| KEGG_PATHWAY | mmu04261:Adrenergic signaling in cardiomyocytes | 6 | 1,245 | 0,36384191 | ADCY1, ATF4, PPP2R5B, GNAS, ATP1A1, PPP2R2A | 205 | 150 | 7720 | 1,506341 | 1 | 0,7580162 | 99,69565 |
| KEGG_PATHWAY | mmu04911:Insulin secretion | 4 | 0,83 | 0,39800016 | ADCY1, ATF4, GNAS, ATP1A1 | 205 | 86 | 7720 | 1,75156 | 1 | 0,7876588 | 99,849928 |
| KEGG_PATHWAY | mmu05215:Prostate cancer | 4 | 0,83 | 0,41228573 | EGFR, HRAS, ATF4, MAP2K2 | 205 | 88 | 7720 | 1,711752 | 1 | 0,789963 | 99,889675 |
| GOTERM_CC_DIRECT | GO:0045121~membrane raft | 6 | 1,245 | 0,66767038 | EGFR, ADCY1, RIPK1, GNAS, ATP1A1, ITPR1 | 424 | 262 | 19662 | 1,061969 | 1 | 0,9947966 | 99,999973 |
| GOTERM_MF_DIRECT | GO:0008022~protein C-terminus binding | 5 | 1,037 | 0,67880409 | HRAS, ATF4, PLEKHB1, ITPR1, PSMD9 | 385 | 209 | 17446 | 1,084074 | 1 | 0,9993813 | 99,999995 |
| KEGG_PATHWAY | mmu04713:Circadian entrainment | 3 | 0,622 | 0,73668974 | ADCY1, GNAS, ITPR1 | 205 | 98 | 7720 | 1,152812 | 1 | 0,9409558 | 99,999996 |
| KEGG_PATHWAY | mmu04723:Retrograde endocannabinoid signaling | 3 | 0,622 | 0,76142485 | ADCY1, ITPR1, ITPR2 | 205 | 103 | 7720 | 1,096851 | 1 | 0,9493792 | 99,999999 |
| KEGG_PATHWAY | mmu05161:Hepatitis B | 3 | 0,622 | 0,90271126 | HRAS, ATF4, MAP2K2 | 205 | 146 | 7720 | 0,773806 | 1 | 0,9891319 | 100 |
| Annotation | Enrichment Score: 0.8541699720748208 | | | | | | | | | | | |
| Category | Term | Count | % | PValue | Genes | List Total | Pop Hits | Pop Total | Enrichment | Bonferroni | Benjamini | FDR |
| UP_KEYWORDS | Serine esterase | 4 | 0,83 | 0,03267308 | BAAT, CES3A, CES2A, CES1E | 456 | 35 | 22680 | 5,684211 | 0,999975 | 0,2746599 | 36,113235 |
| INTERPRO | IPR019826:Carboxylesterase type B, active site | 3 | 0,622 | 0,07790861 | CES3A, CES2A, CES1E | 436 | 22 | 20594 | 6,440992 | 1 | 0,9387069 | 71,73098 |
| INTERPRO | IPR019819:Carboxylesterase type B, conserved site | 3 | 0,622 | 0,11062243 | CES3A, CES2A, CES1E | 436 | 27 | 20594 | 5,248216 | 1 | 0,9764183 | 83,895256 |
| INTERPRO | IPR002018:Carboxylesterase, type B | 3 | 0,622 | 0,11753702 | CES3A, CES2A, CES1E | 436 | 28 | 20594 | 5,06078 | 1 | 0,9789098 | 85,738824 |
| COG_ONTOLOGY | Lipid metabolism | 5 | 1,037 | 0,24388137 | ACSM2, CES3A, CES2A, ECHDC2, CES1E | 62 | 88 | 2126 | 1,948314 | 0,9962692 | 0,8449044 | 88,088734 |
| GOTERM_MF_DIRECT | GO:0052689~carboxylic ester hydrolase activity | 4 | 0,83 | 0,26398443 | BAAT, CES3A, CES2A, CES1E | 385 | 81 | 17446 | 2,237743 | 1 | 0,9343882 | 98,951736 |
| UP_SEQ_FEATURE | active site:Charge relay system | 6 | 1,245 | 0,4923341 | BAAT, F10, UOX, CES3A, CES1E, PCSK4 | 408 | 205 | 18012 | 1,292109 | 1 | 1 | 99,997977 |
| Annotation | Enrichment Score: 0.8528982029640096 | | | | | | | | | | | |
| Category | Term | Count | % | PValue | Genes | List Total | Pop Hits | Pop Total | Enrichment | Bonferroni | Benjamini | FDR |

| | | | | | | | | | | | | |
|---|---|---|---|---|---|---|---|---|---|---|---|---|
| GOTERM_BP_DIRECT | GO:0070534~protein K63-linked ubiquitination | 4 | 0,83 | 0,0490971 | ARIH2, RNF152, UBE2F, UBE2S | 394 | 38 | 18082 | 4,830884 | 1 | 0,8691718 | 57,510733 |
| GOTERM_MF_DIRECT | GO:0061630~ubiquitin protein ligase activity | 8 | 1,66 | 0,15128958 | CUL2, ARIH2, KCMF1, RNF152, UBE2F, CDC34, UBE2S, BRAP | 385 | 199 | 17446 | 1,82168 | 1 | 0,8425029 | 91,278726 |
| GOTERM_BP_DIRECT | GO:0000209~protein polyubiquitination | 5 | 1,037 | 0,18616217 | ARIH2, RNF152, CDC34, UBE2S, MARCH5 | 394 | 103 | 18082 | 2,227835 | 1 | 0,9871527 | 96,987009 |
| GOTERM_BP_DIRECT | GO:0070936~protein K48-linked ubiquitination | 3 | 0,622 | 0,28031796 | ARIH2, RNF152, CDC34 | 394 | 48 | 18082 | 2,868338 | 1 | 0,9968179 | 99,627473 |
| Annotation | Enrichment Score: 0.8474096214021858 | | | | | | | | | | | |
| Category | Term | Count | % | PValue | Genes | List Total | Pop Hits | Pop Total | Enrichment | Bonferroni | Benjamini | FDR |
| GOTERM_MF_DIRECT | GO:0015347~sodium-independent organic anion transmembrane transporter activity | 3 | 0,622 | 0,11840453 | SLCO1A1, SLC22A28, SLC22A30 | 385 | 27 | 17446 | 5,034921 | 1 | 0,8026654 | 84,650364 |
| GOTERM_BP_DIRECT | GO:0043252~sodium-independent organic anion transport | 3 | 0,622 | 0,12316843 | SLCO1A1, SLC22A28, SLC22A30 | 394 | 28 | 18082 | 4,91715 | 1 | 0,9638063 | 89,297583 |
| INTERPRO | IPR020846:Major facilitator superfamily domain | 6 | 1,245 | 0,19674488 | DIRC2, SLCO1A1, SLC22A28, SLC46A3, SLC19A1, SLC22A30 | 436 | 146 | 20594 | 1,941121 | 1 | 0,996871 | 96,704093 |
| Annotation | Enrichment Score: 0.7375605369735999 | | | | | | | | | | | |
| Category | Term | Count | % | PValue | Genes | List Total | Pop Hits | Pop Total | Enrichment | Bonferroni | Benjamini | FDR |
| UP_KEYWORDS | Transferase | 52 | 10,79 | 0,00145929 | POMT2, ALAS2, ATAT1, ELOVL3, PTDSS1, ADCK5, EGFR, IRAK1, CMAS, PHKG2, PIM3, CHPT1, UGT1A1, BRAP, ZDHHC14, NME2, DGAT1, NME1, RIPK1, MAPK4, UBE2S, GSTP2, GSTP1, KALRN, XYLB, GCNT4, COASY, MAPKAPK5, UGT2B1, CDC34, UGT3A2, ARIH2, UGT1A9, ACAA1B, | 456 | 1654 | 22680 | 1,563673 | 0,3724003 | 0,0242204 | 1,9504669 |
| GOTERM_MF_DIRECT | GO:0016740~transferase activity | 46 | 9,544 | 0,01526716 | MAPKAPK5, CDC34, UGT3A2, GSTM1, POMT2, UGT1A9, ALAS2, ATAT1, ELOVL3, PTDSS1, ACAA1B, ADCK5, AATK, B4GALNT1, EGFR, IRAK1, GSTA4, MAP2K2, CMAS, PHKG2, PIM3, UGT1A1, CHPT1, ZDHHC14, NME2, DGAT1, BAAT, NME1, RIPK1, MAPK4, MAPK15, NAA60, NTRK2, | 385 | 1472 | 17446 | 1,416071 | 0,9999392 | 0,3020102 | 20,450406 |
| UP_KEYWORDS | ATP-binding | 39 | 8,091 | 0,02546239 | FARS2, CLPB, ASNS, CDC34, HSPA1B, HSPA2, NUBP1, ABCB10, AATK, MARS, EGFR, IRAK1, MAP2K2, PHKG2, OLA1, UBE2F, ATP1A1, PIM3, ABCB6, AK6, ABCG2, ABCG8, ACSM2, NME2, VCP, NME1, RIPK1, MAPK4, MAPK15, NTRK2, UBE2S, | 456 | 1363 | 22680 | 1,423138 | 0,9997329 | 0,2398631 | 29,382154 |
| UP_KEYWORDS | Nucleotide-binding | 47 | 9,751 | 0,03782832 | MAPKAPK5, CLPB, FARS2, EIF5, ASNS, CDC34, HSPA1B, HSPA2, NUBP1, ABCB10, TUBG1, AATK, MOCS1, MARS, EGFR, IRAK1, RAP2A, MAP2K2, PHKG2, OLA1, UBE2F, ATP1A1, PIM3, ABCB6, AK6, ABCG2, ABCG8, ACSM2, NME2, VCP, NME1, | 456 | 1754 | 22680 | 1,332743 | 0,9999955 | 0,2894426 | 40,555865 |

| | | | | | | | | | | | | |
|---|---|---|---|---|---|---|---|---|---|---|---|---|
| GOTERM_BP_DIRECT | GO:0046777~protein autophosphorylation | 9 | 1,867 | 0,04729888 | EGFR, IRAK1, NME2, MAPKAPK5, RIPK1, MAPK15, NTRK2, PIM3, AATK | 394 | 183 | 18082 | 2,257053 | 1 | 0,8710837 | 56,123787 |
| GOTERM_MF_DIRECT | GO:0005524~ATP binding | 41 | 8,506 | 0,123736 | FARS2, CLPB, ASNS, CDC34, HSPA1B, HSPA2, NUBP1, P2RY1, ABCB10, AATK, MARS, EGFR, IRAK1, MAP2K2, PHKG2, OLA1, UBE2F, ATP1A1, PIM3, ABCB6, AK6, ABCG2, ABCG8, ACSM2, NME2, VCP, NME1, RIPK1, MAPK4, MAPK15, | 385 | 1507 | 17446 | 1,232837 | 1 | 0,8111777 | 85,97438 |
| UP_KEYWORDS | Kinase | 20 | 4,149 | 0,12413992 | MAPKAPK5, PHKG2, PIM3, AK6, NME2, NME1, MAPK4, RIPK1, NTRK2, MAPK15, ADCK5, AATK, XYLB, KALRN | 456 | 707 | 22680 | 1,406983 | 1 | 0,570727 | 83,267765 |
| GOTERM_MF_DIRECT | GO:0000166~nucleotide binding | 50 | 10,37 | 0,16613168 | NUBP1, ABCB10, TUBG1, PABPN1, EGFR, RAP2A, IRAK1, PHKG2, OLA1, UBE2F, PIM3, BRAP, NME2, NME1, RIPK1, MAPK4, CELF2, GNAS, UBE2S, SLC27A2, KALRN, XYLB, COASY, MAPKAPK5, FARS2, ZCRB1, ASNS, CDC34, HSPA2, MARS, AATK, MOCS1, MAP2K2, ATP1A1, ABCB6, ABCG2, | 385 | 1936 | 17446 | 1,170307 | 1 | 0,8567337 | 93,291321 |
| GOTERM_BP_DIRECT | GO:0016310~phosphorylation | 18 | 3,734 | 0,18375236 | MAPKAPK5, PHKG2, PIM3, NME2, NME1, MAPK4, RIPK1, MAPK15, NTRK2, ADCK5, AATK, | 394 | 612 | 18082 | 1,349806 | 1 | 0,9882893 | 96,83168 |
| GOTERM_MF_DIRECT | GO:0016301~kinase activity | 19 | 3,942 | 0,23211438 | MAPKAPK5, PHKG2, PIM3, NME2, NME1, MAPK4, RIPK1, MAPK15, NTRK2, ADCK5, AATK, | 385 | 674 | 17446 | 1,277406 | 1 | 0,9168747 | 98,031027 |
| GOTERM_MF_DIRECT | GO:0004674~protein serine/threonine kinase activity | 13 | 2,697 | 0,23262967 | RIPK1, MAPK4, MAPK15, PIM3, ADCK5, AATK, NRBP2, KALRN | 385 | 428 | 17446 | 1,376368 | 1 | 0,9143132 | 98,050585 |
| UP_SEQ_FEATURE | active site:Proton acceptor | 20 | 4,149 | 0,25680148 | MAPKAPK5, PHKG2, UNG, HSD3B5, PIM3, SIRT2, RIPK1, MAPK4, NTRK2, MAPK15, TXNRD3, H2-KE6, ACAA1B, AATK, KALRN | 408 | 710 | 18012 | 1,243579 | 1 | 0,9999999 | 99,119018 |
| UP_KEYWORDS | Serine/threonine-protein kinase | 11 | 2,282 | 0,28775793 | IRAK1, MAP2K2, MAPKAPK5, PHKG2, RIPK1, MAPK4, MAPK15, PIM3, ADCK5, AATK, KALRN | 456 | 405 | 22680 | 1,350877 | 1 | 0,7638564 | 98,971447 |
| INTERPRO | IPR001245:Serine-threonine/tyrosine-protein kinase catalytic domain | 5 | 1,037 | 0,3382153 | EGFR, IRAK1, RIPK1, NTRK2, AATK | 436 | 139 | 20594 | 1,699063 | 1 | 0,9999026 | 99,838772 |
| UP_SEQ_FEATURE | binding site:ATP | 16 | 3,32 | 0,34186283 | PIM3, ACSM2, NME2, NME1, MAPK4, RIPK1, MAPK15, NTRK2, AATK, MARS, KALRN | 408 | 583 | 18012 | 1,211583 | 1 | 1 | 99,873136 |
| UP_SEQ_FEATURE | domain:Protein kinase | 14 | 2,905 | 0,35502507 | PIM3, MAPK4, RIPK1, MAPK15, NTRK2, ADCK5, KALRN, NRBP2, AATK | 408 | 502 | 18012 | 1,231193 | 1 | 1 | 99,90807 |
| INTERPRO | IPR011009:Protein kinase-like domain | 14 | 2,905 | 0,39113997 | PIM3, MAPK4, RIPK1, MAPK15, NTRK2, ADCK5, KALRN, NRBP2, AATK | 436 | 556 | 20594 | 1,189344 | 1 | 0,9999744 | 99,955986 |
| INTERPRO | IPR000719:Protein kinase, catalytic domain | 13 | 2,697 | 0,40401139 | PIM3, MAPK4, RIPK1, MAPK15, NTRK2, KALRN, NRBP2, AATK | 436 | 515 | 20594 | 1,192313 | 1 | 0,999979 | 99,968446 |
| SMART | SM00220:S_TKc | 8 | 1,66 | 0,43202558 | IRAK1, MAP2K2, MAPKAPK5, PHKG2, RIPK1, MAPK4, PIM3, KALRN | 172 | 380 | 10425 | 1,27601 | 1 | 0,9998953 | 99,893523 |
| INTERPRO | IPR017441:Protein kinase, ATP binding site | 10 | 2,075 | 0,44779695 | EGFR, IRAK1, MAP2K2, PHKG2, MAPK4, MAPK15, NTRK2, PIM3, AATK, KALRN | 436 | 394 | 20594 | 1,198831 | 1 | 0,9999904 | 99,990387 |
| GOTERM_MF_DIRECT | GO:0004713~protein tyrosine kinase activity | 4 | 0,83 | 0,49972759 | EGFR, MAP2K2, NTRK2, AATK | 385 | 121 | 17446 | 1,497993 | 1 | 0,9937911 | 99,996636 |

| | | | | | | | | | | | | |
|---|---|---|---|---|---|---|---|---|---|---|---|---|
| UP_SEQ_FEATURE | nucleotide phosphate-binding region:ATP | 23 | 4,772 | 0,51305558 | MAPKAPK5, PHKG2, CLPB, OLA1, PIM3, ABCB6, ABCG2, ACSM2, VCP, NUBP1, RIPK1, MAPK4, NTRK2, MAPK15, ABCB10, AATK, KALRN | 408 | 963 | 18012 | 1,054395 | 1 | 1 | 99,998959 |
| GOTERM_MF_DIRECT | GO:0004672~protein kinase activity | 13 | 2,697 | 0,5226582 | PIM3, MAPK4, RIPK1, MAPK15, NTRK2, KALRN, NRBP2, AATK | 385 | 531 | 17446 | 1,109389 | 1 | 0,9950219 | 99,998326 |
| INTERPRO | IPR008271:Serine/threonine-protein kinase, active site | 8 | 1,66 | 0,55819647 | IRAK1, MAP2K2, MAPKAPK5, PHKG2, RIPK1, MAPK4, PIM3, KALRN | 436 | 333 | 20594 | 1,134749 | 1 | 0,9999992 | 99,999702 |
| GOTERM_BP_DIRECT | GO:0006468~protein phosphorylation | 13 | 2,697 | 0,60077607 | PIM3, MAPK4, RIPK1, MAPK15, NTRK2, KALRN, NRBP2, AATK | 394 | 576 | 18082 | 1,035789 | 1 | 0,9999817 | 99,999983 |
| INTERPRO | IPR008266:Tyrosine-protein kinase, active site | 3 | 0,622 | 0,62178352 | EGFR, NTRK2, AATK | 436 | 99 | 20594 | 1,431332 | 1 | 0,9999999 | 99,999974 |
| Annotation | Enrichment Score: 0.7307673789648991 | | | | | | | | | | | |
| Category | Term | Count | % | PValue | Genes | List Total | Pop Hits | Pop Total | Enrichment | Bonferroni | Benjamini | FDR |
| GOTERM_CC_DIRECT | GO:0000139~Golgi membrane | 15 | 3,112 | 0,0548855 | CYP2E1, ABCB6, CHPT1, GBA2, 1190002N15RIK, PLEKHB1, NAA60, CNIH1, HPD | 424 | 403 | 19662 | 1,726029 | 1 | 0,5403372 | 53,912911 |
| GOTERM_CC_DIRECT | GO:0005794~Golgi apparatus | 32 | 6,639 | 0,15645229 | UBAC1, CRYL1, ATAT1, PLA2G12A, SERPINA1C, B4GALNT1, HPD, EGFR, FZD8, H2-Q10, MAP2K2, TMEM5, ATP1A1, ABCB6, CHPT1, GBA2, 1190002N15RIK, NAA60, DYM, GNAS, CTSC, | 424 | 1190 | 19662 | 1,246995 | 1 | 0,8158374 | 90,316723 |
| UP_KEYWORDS | Golgi apparatus | 14 | 2,905 | 0,74792542 | ABCB6, CHPT1, GBA2, 1190002N15RIK, NAA60, DYM, CNIH1, B4GALNT1 | 456 | 753 | 22680 | 0,924722 | 1 | 0,9703037 | 99,999999 |
| Annotation | Enrichment Score: 0.6769601775053186 | | | | | | | | | | | |
| Category | Term | Count | % | PValue | Genes | List Total | Pop Hits | Pop Total | Enrichment | Bonferroni | Benjamini | FDR |
| UP_KEYWORDS | Lysosome | 9 | 1,867 | 0,12948063 | PLEKHF1, MARCH2, DIRC2, RNF152, SMPD1, ARSA, ACP5, MCOLN1, CTSC | 456 | 249 | 22680 | 1,797717 | 1 | 0,5659554 | 84,592735 |
| GOTERM_CC_DIRECT | GO:0005764~lysosome | 11 | 2,282 | 0,17813826 | PLEKHF1, MARCH2, DIRC2, RNF152, SMPD1, ARSA, MT1, ACP5, MCOLN1, CTSC, RAB38 | 424 | 331 | 19662 | 1,541085 | 1 | 0,8424876 | 93,22661 |
| KEGG_PATHWAY | mmu04142:Lysosome | 5 | 1,037 | 0,40379176 | SMPD1, ARSA, ACP5, MCOLN1, CTSC | 205 | 122 | 7720 | 1,543383 | 1 | 0,7852032 | 99,86741 |
| Annotation | Enrichment Score: 0.623067793875792 | | | | | | | | | | | |
| Category | Term | Count | % | PValue | Genes | List Total | Pop Hits | Pop Total | Enrichment | Bonferroni | Benjamini | FDR |
| GOTERM_BP_DIRECT | GO:0045669~positive regulation of osteoblast differentiation | 4 | 0,83 | 0,18392781 | BMP4, CEBPB, GNAS, BMP7 | 394 | 68 | 18082 | 2,699612 | 1 | 0,9877099 | 96,843239 |
| GOTERM_BP_DIRECT | GO:0051216~cartilage development | 4 | 0,83 | 0,26381849 | BMP4, LECT1, GNAS, BMP7 | 394 | 82 | 18082 | 2,238702 | 1 | 0,9961824 | 99,452321 |
| GOTERM_CC_DIRECT | GO:0031982~vesicle | 6 | 1,245 | 0,27851206 | BMP4, GNAS, RAB38, BMP7, GSTP2, GSTP1 | 424 | 164 | 19662 | 1,69656 | 1 | 0,9079117 | 98,866255 |

| Annotation | Enrichment Score: 0.6158581393639151 | | | | | | | | | | | |
|---|---|---|---|---|---|---|---|---|---|---|---|---|
| Category | Term | Count | % | PValue | Genes | List Total | Pop Hits | Pop Total | Enrichment | Bonferroni | Benjamini | FDR |
| INTERPRO | dependent transferase, major region, subdomain 2 | 3 | 0,622 | 0,19131248 | ETNPPL, ALAS2, PSAT1 | 436 | 38 | 20594 | 3,728996 | 1 | 0,9968457 | 96,339248 |
| INTERPRO | IPR015424:Pyridoxal phosphate-dependent transferase | 3 | 0,622 | 0,230127 | ETNPPL, ALAS2, PSAT1 | 436 | 43 | 20594 | 3,295392 | 1 | 0,9987538 | 98,298484 |
| INTERPRO | dependent transferase, major region, subdomain 1 | 3 | 0,622 | 0,230127 | ETNPPL, ALAS2, PSAT1 | 436 | 43 | 20594 | 3,295392 | 1 | 0,9987538 | 98,298484 |
| UP_KEYWORDS | Pyridoxal phosphate | 3 | 0,622 | 0,33953848 | ETNPPL, ALAS2, PSAT1 | 456 | 60 | 22680 | 2,486842 | 1 | 0,8047862 | 99,628392 |
| Annotation | Enrichment Score: 0.5815472555650827 | | | | | | | | | | | |
| Category | Term | Count | % | PValue | Genes | List Total | Pop Hits | Pop Total | Enrichment | Bonferroni | Benjamini | FDR |
| KEGG_PATHWAY | mmu05030:Cocaine addiction | 4 | 0,83 | 0,13895772 | BDNF, ATF4, MAOA, GNAS | 205 | 49 | 7720 | 3,074166 | 1 | 0,5251736 | 85,291574 |
| KEGG_PATHWAY | mmu04728:Dopaminergic synapse | 7 | 1,452 | 0,14343964 | ATF4, PPP2R5B, MAOA, GNAS, ITPR1, PPP2R2A, ITPR2 | 205 | 134 | 7720 | 1,967237 | 1 | 0,5297011 | 86,242838 |
| KEGG_PATHWAY | mmu05034:Alcoholism | 7 | 1,452 | 0,44463559 | BDNF, HRAS, HIST1H2BC, ATF4, MAOA, NTRK2, GNAS | 205 | 202 | 7720 | 1,304999 | 1 | 0,8103813 | 99,946585 |
| KEGG_PATHWAY | mmu05031:Amphetamine addiction | 3 | 0,622 | 0,53242065 | ATF4, MAOA, GNAS | 205 | 67 | 7720 | 1,686203 | 1 | 0,8338171 | 99,994107 |
| Annotation | Enrichment Score: 0.568365909202043 | | | | | | | | | | | |
| Category | Term | Count | % | PValue | Genes | List Total | Pop Hits | Pop Total | Enrichment | Bonferroni | Benjamini | FDR |
| UP_KEYWORDS | Peroxisome | 5 | 1,037 | 0,16817047 | BAAT, UOX, PHYH, ACAA1B, SLC27A2 | 456 | 107 | 22680 | 2,324152 | 1 | 0,6562845 | 91,655216 |
| GOTERM_CC_DIRECT | GO:0005777~peroxisome | 5 | 1,037 | 0,31152598 | BAAT, UOX, PHYH, ACAA1B, SLC27A2 | 424 | 131 | 19662 | 1,769948 | 1 | 0,9309323 | 99,403825 |
| KEGG_PATHWAY | mmu04146:Peroxisome | 4 | 0,83 | 0,3764075 | BAAT, PHYH, ACAA1B, SLC27A2 | 205 | 83 | 7720 | 1,814869 | 1 | 0,7680997 | 99,764293 |
| Annotation | Enrichment Score: 0.5563880175973055 | | | | | | | | | | | |
| Category | Term | Count | % | PValue | Genes | List Total | Pop Hits | Pop Total | Enrichment | Bonferroni | Benjamini | FDR |
| KEGG_PATHWAY | mmu04912:GnRH signaling pathway | 8 | 1,66 | 0,00845519 | EGFR, HRAS, ADCY1, ATF4, MAP2K2, GNAS, ITPR1, ITPR2 | 205 | 88 | 7720 | 3,423503 | 0,856937 | 0,114435 | 10,307602 |
| KEGG_PATHWAY | mmu04540:Gap junction | 7 | 1,452 | 0,02587746 | EGFR, HRAS, ADCY1, MAP2K2, GNAS, ITPR1, ITPR2 | 205 | 86 | 7720 | 3,06523 | 0,997531 | 0,2213277 | 28,530119 |
| KEGG_PATHWAY | mmu04720:Long-term potentiation | 6 | 1,245 | 0,02984439 | HRAS, ADCY1, ATF4, MAP2K2, ITPR1, ITPR2 | 205 | 66 | 7720 | 3,423503 | 0,9990302 | 0,2423524 | 32,1705 |

| | | | | | | | | | | | | |
|---|---|---|---|---|---|---|---|---|---|---|---|---|
| BIOCARTA | m_erkPathway:Erk1/Erk2 Mapk Signaling pathway | 4 | 0,83 | 0,04219162 | EGFR, HRAS, MAP2K2, GNAS | 32 | 33 | 1289 | 4,882576 | 0,9929686 | 0,7104253 | 38,800429 |
| GOTERM_BP_DIRECT | GO:0070374~positive regulation of ERK1 and ERK2 cascade | 8 | 1,66 | 0,11671329 | NOX4, EGFR, BMP4, HRAS, MAP2K2, P2RY1, CCL9, FGF21 | 394 | 188 | 18082 | 1,952911 | 1 | 0,9637746 | 87,876142 |
| BIOCARTA | activation of JNK Pathway via Pyk2 dependent signaling | 3 | 0,622 | 0,18688109 | EGFR, HRAS, MAP2K2 | 32 | 33 | 1289 | 3,661932 | 1 | 0,9288838 | 90,524947 |
| KEGG_PATHWAY | mmu04921:Oxytocin signaling pathway | 7 | 1,452 | 0,23953149 | EGFR, HRAS, ADCY1, MAP2K2, GNAS, ITPR1, ITPR2 | 205 | 158 | 7720 | 1,668416 | 1 | 0,6736323 | 97,004479 |
| KEGG_PATHWAY | mmu05218:Melanoma | 4 | 0,83 | 0,28919035 | EGFR, HRAS, MAP2K2, FGF21 | 205 | 71 | 7720 | 2,121608 | 1 | 0,7223694 | 98,738934 |
| KEGG_PATHWAY | mmu05219:Bladder cancer | 3 | 0,622 | 0,29534314 | EGFR, HRAS, MAP2K2 | 205 | 41 | 7720 | 2,755503 | 1 | 0,7255279 | 98,871851 |
| KEGG_PATHWAY | mmu05213:Endometrial cancer | 3 | 0,622 | 0,40134158 | EGFR, HRAS, MAP2K2 | 205 | 52 | 7720 | 2,172608 | 1 | 0,7868892 | 99,860257 |
| KEGG_PATHWAY | mmu05215:Prostate cancer | 4 | 0,83 | 0,41228573 | EGFR, HRAS, ATF4, MAP2K2 | 205 | 88 | 7720 | 1,711752 | 1 | 0,789963 | 99,889675 |
| KEGG_PATHWAY | mmu05223:Non-small cell lung cancer | 3 | 0,622 | 0,43811502 | EGFR, HRAS, MAP2K2 | 205 | 56 | 7720 | 2,017422 | 1 | 0,8079715 | 99,937969 |
| KEGG_PATHWAY | mmu05205:Proteoglycans in cancer | 7 | 1,452 | 0,4545031 | EGFR, FZD8, HRAS, MAP2K2, RPS6, ITPR1, ITPR2 | 205 | 203 | 7720 | 1,29857 | 1 | 0,8083779 | 99,957546 |
| KEGG_PATHWAY | mmu04015:Rap1 signaling pathway | 7 | 1,452 | 0,50042306 | EGFR, HRAS, ADCY1, MAP2K2, P2RY1, GNAS, FGF21 | 205 | 214 | 7720 | 1,231821 | 1 | 0,8256035 | 99,986239 |
| KEGG_PATHWAY | mmu05231:Choline metabolism in cancer | 4 | 0,83 | 0,50200113 | EGFR, HRAS, MAP2K2, CHPT1 | 205 | 101 | 7720 | 1,491427 | 1 | 0,8236555 | 99,986786 |
| KEGG_PATHWAY | mmu05230:Central carbon metabolism in cancer | 3 | 0,622 | 0,50777455 | EGFR, HRAS, MAP2K2 | 205 | 64 | 7720 | 1,765244 | 1 | 0,8254196 | 99,98862 |
| KEGG_PATHWAY | mmu05214:Glioma | 3 | 0,622 | 0,51608307 | EGFR, HRAS, MAP2K2 | 205 | 65 | 7720 | 1,738086 | 1 | 0,8261672 | 99,99085 |
| KEGG_PATHWAY | mmu04151:PI3K-Akt signaling pathway | 10 | 2,075 | 0,58594613 | EGFR, IFNAR2, HRAS, ATF4, PPP2R5B, MAP2K2, COL27A1, FGF21, RPS6, PPP2R2A | 205 | 351 | 7720 | 1,072893 | 1 | 0,8726018 | 99,998759 |
| KEGG_PATHWAY | mmu04012:ErbB signaling pathway | 3 | 0,622 | 0,67459768 | EGFR, HRAS, MAP2K2 | 205 | 87 | 7720 | 1,29857 | 1 | 0,9215671 | 99,999943 |
| KEGG_PATHWAY | mmu04068:FoxO signaling pathway | 4 | 0,83 | 0,69283876 | EGFR, HRAS, MAP2K2, GADD45G | 205 | 134 | 7720 | 1,124135 | 1 | 0,929354 | 99,999973 |
| KEGG_PATHWAY | mmu04014:Ras signaling pathway | 6 | 1,245 | 0,72940861 | EGFR, HRAS, MAP2K2, PLA2G12A, FGF21, BRAP | 205 | 229 | 7720 | 0,986687 | 1 | 0,9406277 | 99,999995 |
| KEGG_PATHWAY | mmu05161:Hepatitis B | 3 | 0,622 | 0,90271126 | HRAS, ATF4, MAP2K2 | 205 | 146 | 7720 | 0,773806 | 1 | 0,9891319 | 100 |
| KEGG_PATHWAY | mmu04810:Regulation of actin cytoskeleton | 4 | 0,83 | 0,92624267 | EGFR, HRAS, MAP2K2, FGF21 | 205 | 214 | 7720 | 0,703898 | 1 | 0,9928013 | 100 |
| KEGG_PATHWAY | mmu05206:MicroRNAs in cancer | 3 | 0,622 | 0,99528073 | EGFR, HRAS, MAP2K2 | 205 | 276 | 7720 | 0,409332 | 1 | 0,9999142 | 100 |
| Annotation | Enrichment Score: 0.5414303114985964 | | | | | | | | | | | |

| Category | Term | Count | % | PValue | Genes | List Total | Pop Hits | Pop Total | Enrichment | Bonferroni | Benjamini | FDR |
|---|---|---|---|---|---|---|---|---|---|---|---|---|
| INTERPRO | IPR023313:Ubiquitin-conjugating enzyme, active site | 3 | 0,622 | 0,11753702 | UBE2F, CDC34, UBE2S | 436 | 28 | 20594 | 5,06078 | 1 | 0,9789098 | 85,738824 |
| GOTERM_MF_DIRECT | GO:0061630~ubiquitin protein ligase activity | 8 | 1,66 | 0,15128958 | CUL2, ARIH2, KCMF1, RNF152, UBE2F, CDC34, UBE2S, BRAP | 385 | 199 | 17446 | 1,82168 | 1 | 0,8425029 | 91,278726 |
| INTERPRO | IPR000608:Ubiquitin-conjugating enzyme, E2 | 3 | 0,622 | 0,230127 | UBE2F, CDC34, UBE2S | 436 | 43 | 20594 | 3,295392 | 1 | 0,9987538 | 98,298484 |
| INTERPRO | IPR016135:Ubiquitin-conjugating enzyme/RWD-like | 3 | 0,622 | 0,33968636 | UBE2F, CDC34, UBE2S | 436 | 57 | 20594 | 2,485997 | 1 | 0,9998838 | 99,844265 |
| UP_SEQ_FEATURE | active site:Glycyl thioester intermediate | 3 | 0,622 | 0,55609903 | UBE2F, CDC34, UBE2S | 408 | 82 | 18012 | 1,615136 | 1 | 1 | 99,999762 |
| KEGG_PATHWAY | mmu04120:Ubiquitin mediated proteolysis | 4 | 0,83 | 0,72985504 | CUL2, UBE2F, CDC34, UBE2S | 205 | 142 | 7720 | 1,060804 | 1 | 0,9392549 | 99,999995 |
| Annotation | Enrichment Score: 0.5188778129491886 | | | | | | | | | | | |
| Category | Term | Count | % | PValue | Genes | List Total | Pop Hits | Pop Total | Enrichment | Bonferroni | Benjamini | FDR |
| GOTERM_MF_DIRECT | GO:0061630~ubiquitin protein ligase activity | 8 | 1,66 | 0,15128958 | CUL2, ARIH2, KCMF1, RNF152, UBE2F, CDC34, UBE2S, BRAP | 385 | 199 | 17446 | 1,82168 | 1 | 0,8425029 | 91,278726 |
| UP_KEYWORDS | Ubl conjugation pathway | 16 | 3,32 | 0,28050844 | BRAP, CUL2, MARCH2, ARIH2, ATG4D, MAP1LC3A, KCMF1, VCP, RNF152, FBXO31, | 456 | 631 | 22680 | 1,261156 | 1 | 0,7627414 | 98,820906 |
| GOTERM_MF_DIRECT | GO:0004842~ubiquitin-protein transferase activity | 10 | 2,075 | 0,29329886 | MARCH2, ARIH2, RNF152, FBXO31, CDC34, UBE2S, CYHR1, MARCH5, TRAF4, BRAP | 385 | 326 | 17446 | 1,390009 | 1 | 0,950247 | 99,427232 |
| GOTERM_BP_DIRECT | GO:0016567~protein ubiquitination | 8 | 1,66 | 0,67518604 | MARCH2, ARIH2, KCMF1, VCP, CDC34, UBAC1, TRAF4, BRAP | 394 | 362 | 18082 | 1,014219 | 1 | 0,9999969 | 100 |
| Annotation | Enrichment Score: 0.4272335627352353 | | | | | | | | | | | |
| Category | Term | Count | % | PValue | Genes | List Total | Pop Hits | Pop Total | Enrichment | Bonferroni | Benjamini | FDR |
| GOTERM_MF_DIRECT | GO:0008270~zinc ion binding | 29 | 6,017 | 0,20077086 | MT1, GLO1, ZPR1, CYHR1, TRAF4, CGRRF1, MICAL2, PHF10, DNAJC21, CSRP2, MARCH5, SIRT2, BRAP, MARCH2, ZDHHC14, NR1I3, KCMF1, ZFP598, RNF152, ZFAND2B, CAR8, SMPD1 | 385 | 1075 | 17446 | 1,222432 | 1 | 0,8902512 | 96,43039 |
| UP_KEYWORDS | Zinc | 44 | 9,129 | 0,46783434 | RPL37, CPEB1, MCM10, ZC3H8, DCT, ARIH2, TIMM9, DTNB, ZFP354A, GLO1, ZPR1, CYHR1, TRAF4, CGRRF1, MICAL2, ZFP703, KLF16, PHF10, DNAJC21, IRF2BP2, CSRP2, MARCH5, SIRT2, BRAP, ZFP592, PLEKHF1, SALL2, MARCH2, NR1I3, | 456 | 2099 | 22680 | 1,042602 | 1 | 0,8824269 | 99,979821 |
| UP_KEYWORDS | Zinc-finger | 32 | 6,639 | 0,55655302 | ZC3H8, ARIH2, DTNB, ZFP354A, ZPR1, CYHR1, TRAF4, CGRRF1, ZFP703, KLF16, PHF10, DNAJC21, IRF2BP2, MARCH5, BRAP, ZFP592, PLEKHF1, SALL2, MARCH2, NR1I3, ZFP598, | 456 | 1565 | 22680 | 1,016983 | 1 | 0,9174428 | 99,998276 |

| Annotation | Enrichment Score: 0.412360225472114 | | | | | | | | | | | |
|---|---|---|---|---|---|---|---|---|---|---|---|---|
| Category | Term | Count | % | PValue | Genes | List Total | Pop Hits | Pop Total | Enrichment | Bonferroni | Benjamini | FDR |
| GOTERM_BP_DIRECT | GO:0060548~negative regulation of cell death | 4 | 0,83 | 0,25209017 | BMP4, BDNF, SRSF6, BMP7 | 394 | 80 | 18082 | 2,29467 | 1 | 0,9956784 | 99,283475 |
| GOTERM_BP_DIRECT | GO:0001657~ureteric bud development | 3 | 0,622 | 0,28031796 | BMP4, BDNF, BMP7 | 394 | 48 | 18082 | 2,868338 | 1 | 0,9968179 | 99,627473 |
| GOTERM_MF_DIRECT | GO:0008083~growth factor activity | 5 | 1,037 | 0,3962288 | BMP4, BDNF, OSGIN1, FGF21, BMP7 | 385 | 145 | 17446 | 1,562562 | 1 | 0,9794038 | 99,944876 |
| UP_KEYWORDS | Growth factor | 4 | 0,83 | 0,47018785 | BMP4, BDNF, FGF21, BMP7 | 456 | 127 | 22680 | 1,566515 | 1 | 0,881524 | 99,980992 |
| GOTERM_BP_DIRECT | GO:0045666~positive regulation of neuron differentiation | 3 | 0,622 | 0,65883186 | BMP4, BDNF, BMP7 | 394 | 103 | 18082 | 1,336701 | 1 | 0,9999954 | 99,999999 |
| Annotation | Enrichment Score: 0.4007112964564447 | | | | | | | | | | | |
| Category | Term | Count | % | PValue | Genes | List Total | Pop Hits | Pop Total | Enrichment | Bonferroni | Benjamini | FDR |
| UP_KEYWORDS | Disulfide bond | 71 | 14,73 | 0,16961342 | CSPG5, BDNF, TIMM9, CES1E, MUP14, MUP13, MUP12, EGFR, F10, COLEC12, SLC3A1, MIEN1, C8G, IFNAR2, GRM8, PDGFRL, TXNRD3, SUSD4, CTSC, KALRN, GCNT4, RBP4, CPM, C9, TMX4, CCL9, MME, ACP5, SFTPA1, GREM2, GCGR, SLCO1A1, LECT1, TSPAN33, COL27A1, P2RY1, PDILT, GLO1, PLTP, GLRX, B4GALNT1, NOX4, | 456 | 3124 | 22680 | 1,130383 | 1 | 0,6531145 | 91,848361 |
| UP_KEYWORDS | Glycoprotein | 85 | 17,63 | 0,1873222 | HP, CSPG5, SLC52A2, TTR, POMT2, DIRC2, BDNF, TMEM57, SERPINE2, H13, SERPINA6, ELOVL3, CREG1, CES1E, TSKU, EGFR, F10, COLEC12, SLC3A1, CYP1A2, UGT1A1, C8G, IFNAR2, SERPINA3K, GRM8, PDGFRL, SUSD4, CTSC, GNAS, FRAS1, GCNT4, CPM, C9, SLC38A2, SERPINA12, MME, ACP5, TMEM219, SFTPA1, NRN1, SLC19A1, GREM2, GCGR, UGT3A2, DCT, SLCO1A1, UGT1A9, LECT1, TSPAN33, COL27A1, SERPINA1C, P2RY1, PDILT, PCSK4, PLTP, B4GALNT1, NOX4, BMP4, | 456 | 3815 | 22680 | 1,10816 | 1 | 0,6742019 | 93,905048 |

| | | | | | | | | | | | | |
|---|---|---|---|---|---|---|---|---|---|---|---|---|
| UP_KEYWORDS | Signal | 93 | 19,29 | 0,49453247 | ILDR2, CSPG5, RDH9, MTHFD2, TTR, BDNF, SERPINE2, FXN, SERPINA6, CREG1, CES1E, MUP14, TSKU, MUP13, MUP12, EGFR, F10, CYP2C29, FGF21, UGT1A1, C8G, IFNAR2, SERPINA3K, 1190002N15RIK, GRM8, PDGFRL, SUSD4, CTSC, GNAS, FRAS1, RBP4, CPM, C9, CYP2C44, SERPINA12, TMX4, MAPKAPK5, UGT2B1, FAM98C, CCL9, TMEM219, ACP5, SFTPA1, NRN1, GCGR, GREM2, TMCC1, UGT3A2, DCT, UGT1A9, COL27A1, PLA2G12A, SERPINA1C, PDILT, PCSK6, PCSK4, PLTP, MOCS1, AATK, | 456 | 4543 | 22680 | 1,018166 | 1 | 0,8890228 | 99,989922 |
| UP_SEQ_FEATURE | signal peptide | 72 | 14,94 | 0,49920914 | BDNF, SERPINE2, SERPINA6, CREG1, CES1E, MUP14, TSKU, MUP13, MUP12, EGFR, F10, FGF21, UGT1A1, C8G, IFNAR2, SERPINA3K, 1190002N15RIK, GRM8, PDGFRL, SUSD4, CTSC, FRAS1, RBP4, CPM, C9, SERPINA12, TMX4, CCL9, ACP5, SFTPA1, NRN1, GREM2, GCGR, UGT3A2, DCT, UGT1A9, COL27A1, PLA2G12A, SERPINA1C, PDILT, PCSK4, PLTP, MUP5, BMP4, FZD8, MUP7, | 408 | 3124 | 18012 | 1,017474 | 1 | 1 | 99,998373 |
| UP_SEQ_FEATURE | disulfide bond | 56 | 11,62 | 0,62010477 | CES1E, MUP14, MUP13, MUP12, EGFR, F10, COLEC12, MIEN1, C8G, IFNAR2, PDGFRL, TXNRD3, SUSD4, CTSC, KALRN, RBP4, CPM, C9, TMX4, CCL9, ACP5, MME, SFTPA1, GREM2, GCGR, LECT1, P2RY1, PLTP, GLRX, B4GALNT1, BMP4, MUP5, FZD8, MUP7, H2-Q10, MUP1, | 408 | 2510 | 18012 | 0,984954 | 1 | 1 | 99,99998 |
| UP_SEQ_FEATURE | glycosylation site:N-linked (GlcNAc...) | 75 | 15,56 | 0,81047633 | CSPG5, TTR, POMT2, DIRC2, BDNF, TMEM57, SERPINE2, H13, SERPINA6, ELOVL3, CREG1, CES1E, TSKU, EGFR, F10, COLEC12, UGT1A1, C8G, IFNAR2, SERPINA3K, GRM8, PDGFRL, SUSD4, CTSC, FRAS1, CPM, SLC38A2, C9, SERPINA12, MME, ACP5, TMEM219, SFTPA1, SLC19A1, GREM2, GCGR, UGT3A2, DCT, SLCO1A1, UGT1A9, LECT1, TSPAN33, COL27A1, SERPINA1C, P2RY1, PDILT, PCSK4, PLTP, B4GALNT1, NOX4, BMP4, | 408 | 3563 | 18012 | 0,929281 | 1 | 1 | 100 |
| Annotation | Enrichment Score: 0.3847473444305742 | | | | | | | | | | | |
| Category | Term | Count | % | PValue | Genes | List Total | Pop Hits | Pop Total | Enrichment | Bonferroni | Benjamini | FDR |
| SMART | SM00132:LIM | 3 | 0,622 | 0,34324762 | LMO4, MICAL2, CSRP2 | 172 | 74 | 10425 | 2,457181 | 1 | 0,9999405 | 99,382739 |
| UP_KEYWORDS | LIM domain | 3 | 0,622 | 0,43890485 | LMO4, MICAL2, CSRP2 | 456 | 74 | 22680 | 2,016358 | 1 | 0,8710344 | 99,958792 |
| INTERPRO | IPR001781:Zinc finger, LIM-type | 3 | 0,622 | 0,46535036 | LMO4, MICAL2, CSRP2 | 436 | 74 | 20594 | 1,91489 | 1 | 0,9999934 | 99,994188 |

| | | | | | | | | | | | | |
|---|---|---|---|---|---|---|---|---|---|---|---|---|
| Annotation | Enrichment Score: 0.37161557192561706 | | | | | | | | | | | |
| Category | Term | Count | % | PValue | Genes | List Total | Pop Hits | Pop Total | Enrichment | Bonferroni | Benjamini | FDR |
| SMART | SM00353:HLH | 4 | 0,83 | 0,26978591 | EPAS1, HES6, TCF24, MYCN | 172 | 110 | 10425 | 2,204017 | 1 | 0,9997943 | 97,773209 |
| UP_SEQ_FEATURE | DNA-binding region:Basic motif | 6 | 1,245 | 0,27782084 | MAFF, ATF4, CEBPB, EPAS1, HES6, MYCN | 408 | 156 | 18012 | 1,697964 | 1 | 0,9999999 | 99,442412 |
| INTERPRO | IPR011598:Myc-type, basic helix-loop-helix (bHLH) domain | 4 | 0,83 | 0,42819892 | EPAS1, HES6, TCF24, MYCN | 436 | 113 | 20594 | 1,671998 | 1 | 0,9999851 | 99,98345 |
| GOTERM_MF_DIRECT | GO:0046983~protein dimerization activity | 5 | 1,037 | 0,60490958 | EPAS1, POLR1D, HES6, MYCN, ABCG2 | 385 | 190 | 17446 | 1,192481 | 1 | 0,9982866 | 99,999899 |
| UP_SEQ_FEATURE | domain:Helix-loop-helix motif | 3 | 0,622 | 0,71417071 | EPAS1, HES6, MYCN | 408 | 110 | 18012 | 1,204011 | 1 | 1 | 100 |
| Annotation | Enrichment Score: 0.3186777899172587 | | | | | | | | | | | |
| Category | Term | Count | % | PValue | Genes | List Total | Pop Hits | Pop Total | Enrichment | Bonferroni | Benjamini | FDR |
| GOTERM_MF_DIRECT | GO:0004842~ubiquitin-protein transferase activity | 10 | 2,075 | 0,29329886 | MARCH2, ARIH2, RNF152, FBXO31, CDC34, UBE2S, CYHR1, MARCH5, TRAF4, BRAP | 385 | 326 | 17446 | 1,390009 | 1 | 0,950247 | 99,427232 |
| INTERPRO | IPR013083:Zinc finger, RING/FYVE/PHD-type | 11 | 2,282 | 0,471198 | PLEKHF1, MARCH2, ARIH2, CGRRF1, ZFP598, RNF152, PHF10, CYHR1, MARCH5, TRAF4, BRAP | 436 | 445 | 20594 | 1,167581 | 1 | 0,9999931 | 99,995103 |
| SMART | SM00184:RING | 5 | 1,037 | 0,53761048 | ARIH2, CGRRF1, RNF152, TRAF4, BRAP | 172 | 234 | 10425 | 1,295095 | 1 | 0,99997 | 99,99116 |
| INTERPRO | IPR001841:Zinc finger, RING-type | 7 | 1,452 | 0,56382151 | ARIH2, CGRRF1, ZFP598, RNF152, CYHR1, TRAF4, BRAP | 436 | 286 | 20594 | 1,156076 | 1 | 0,9999992 | 99,999756 |
| UP_SEQ_FEATURE | zinc finger region:RING-type | 5 | 1,037 | 0,60881855 | CGRRF1, ZFP598, RNF152, TRAF4, BRAP | 408 | 186 | 18012 | 1,186749 | 1 | 1 | 99,999968 |
| Annotation | Enrichment Score: 0.3004060776250061 | | | | | | | | | | | |
| Category | Term | Count | % | PValue | Genes | List Total | Pop Hits | Pop Total | Enrichment | Bonferroni | Benjamini | FDR |
| SMART | SM00338:BRLZ | 3 | 0,622 | 0,20373707 | MAFF, ATF4, CEBPB | 172 | 51 | 10425 | 3,565321 | 1 | 0,9993774 | 93,650577 |
| UP_SEQ_FEATURE | DNA-binding region:Basic motif | 6 | 1,245 | 0,27782084 | MAFF, ATF4, CEBPB, EPAS1, HES6, MYCN | 408 | 156 | 18012 | 1,697964 | 1 | 0,9999999 | 99,442412 |
| INTERPRO | IPR004827:Basic-leucine zipper domain | 3 | 0,622 | 0,31641021 | MAFF, ATF4, CEBPB | 436 | 54 | 20594 | 2,624108 | 1 | 0,9998383 | 99,732862 |
| UP_SEQ_FEATURE | domain:Leucine-zipper | 4 | 0,83 | 0,44243573 | MAFF, ATF4, CEBPB, MYCN | 408 | 108 | 18012 | 1,635076 | 1 | 1 | 99,990983 |
| GOTERM_MF_DIRECT | GO:0008134~transcription factor binding | 8 | 1,66 | 0,62998429 | ATF4, CEBPB, EPAS1, LMO4, PPARG, CREG1, HES6, SIRT2 | 385 | 342 | 17446 | 1,059983 | 1 | 0,9986446 | 99,999962 |
| GOTERM_CC_DIRECT | GO:0005667~transcription factor complex | 5 | 1,037 | 0,82935107 | ATF4, EPAS1, LMO4, CREG1, HES6 | 424 | 267 | 19662 | 0,868402 | 1 | 0,9994663 | 100 |

| | | | | | | | | | | | | |
|---|---|---|---|---|---|---|---|---|---|---|---|---|
| GOTERM_MF_DIRECT | GO:0043565~sequence-specific DNA binding | 9 | 1,867 | 0,97074991 | SALL2, MAFF, NR1I3, ATF4, CEBPB, EPAS1, PPARG, SP5, ZC3H8 | 385 | 633 | 17446 | 0,644279 | 1 | 1 | 100 |
| GOTERM_MF_DIRECT | activity, RNA polymerase II core promoter proximal region sequence-specific binding | 3 | 0,622 | 0,98319649 | ATF4, CEBPB, EPAS1 | 385 | 270 | 17446 | 0,503492 | 1 | 1 | 100 |
| Annotation | Enrichment Score: 0.2670876281533303 | | | | | | | | | | | |
| Category | Term | Count | % | PValue | Genes | List Total | Pop Hits | Pop Total | Enrichment | Bonferroni | Benjamini | FDR |
| GOTERM_CC_DIRECT | GO:0014069~postsynaptic density | 8 | 1,66 | 0,25677331 | DLGAP1, P2RY1, NTRK2, ATP1A1, CPEB1, DTNBP1, ITPR1, KALRN | 424 | 239 | 19662 | 1,552222 | 1 | 0,910588 | 98,296148 |
| GOTERM_CC_DIRECT | GO:0045211~postsynaptic membrane | 5 | 1,037 | 0,70660726 | DLGAP1, P2RY1, NTRK2, CPEB1, DTNBP1 | 424 | 222 | 19662 | 1,044429 | 1 | 0,9966582 | 99,999995 |
| UP_KEYWORDS | Postsynaptic cell membrane | 3 | 0,622 | 0,87098192 | DLGAP1, CPEB1, DTNBP1 | 456 | 176 | 22680 | 0,847787 | 1 | 0,9926396 | 100 |
| Annotation | Enrichment Score: 0.25680835851693207 | | | | | | | | | | | |
| Category | Term | Count | % | PValue | Genes | List Total | Pop Hits | Pop Total | Enrichment | Bonferroni | Benjamini | FDR |
| GOTERM_MF_DIRECT | GO:0008201~heparin binding | 5 | 1,037 | 0,42595716 | BMP4, SERPINE2, PCSK6, BMP7, GREM2 | 385 | 151 | 17446 | 1,500473 | 1 | 0,9852973 | 99,973983 |
| UP_KEYWORDS | Cytokine | 5 | 1,037 | 0,57180258 | BMP4, NAMPT, CCL9, BMP7, GREM2 | 456 | 200 | 22680 | 1,243421 | 1 | 0,916446 | 99,998925 |
| GOTERM_MF_DIRECT | GO:0005125~cytokine activity | 5 | 1,037 | 0,696567 | BMP4, NAMPT, CCL9, BMP7, GREM2 | 385 | 214 | 17446 | 1,058745 | 1 | 0,9995002 | 99,999998 |
| Annotation | Enrichment Score: 0.24945961855098295 | | | | | | | | | | | |
| Category | Term | Count | % | PValue | Genes | List Total | Pop Hits | Pop Total | Enrichment | Bonferroni | Benjamini | FDR |
| UP_KEYWORDS | Prenylation | 6 | 1,245 | 0,208421 | RAP2A, HRAS, PTP4A1, RAB22A, RAB38, MIEN1 | 456 | 157 | 22680 | 1,900771 | 1 | 0,6995746 | 95,725547 |
| UP_SEQ_FEATURE | short sequence motif:Effector region | 4 | 0,83 | 0,39418406 | RAP2A, HRAS, RAB22A, RAB38 | 408 | 100 | 18012 | 1,765882 | 1 | 1 | 99,966134 |
| GOTERM_MF_DIRECT | GO:0005525~GTP binding | 10 | 2,075 | 0,47189602 | RAP2A, HRAS, NME1, EIF5, RAB22A, OLA1, GNAS, RAB38, TUBG1, MOCS1 | 385 | 383 | 17446 | 1,183141 | 1 | 0,9912582 | 99,992474 |
| UP_KEYWORDS | GTP-binding | 8 | 1,66 | 0,50581431 | RAP2A, HRAS, EIF5, RAB22A, GNAS, RAB38, TUBG1, MOCS1 | 456 | 332 | 22680 | 1,198478 | 1 | 0,8944375 | 99,992567 |
| INTERPRO | IPR001806:Small GTPase superfamily | 4 | 0,83 | 0,54049666 | RAP2A, HRAS, RAB22A, RAB38 | 436 | 134 | 20594 | 1,409969 | 1 | 0,9999991 | 99,999451 |
| GOTERM_MF_DIRECT | GO:0003924~GTPase activity | 5 | 1,037 | 0,67880409 | RAP2A, HRAS, RAB22A, GNAS, TUBG1 | 385 | 209 | 17446 | 1,084074 | 1 | 0,9993813 | 99,999995 |
| INTERPRO | IPR005225:Small GTP-binding protein domain | 4 | 0,83 | 0,68046959 | RAP2A, HRAS, RAB22A, RAB38 | 436 | 165 | 20594 | 1,145065 | 1 | 1 | 99,999998 |
| UP_SEQ_FEATURE | nucleotide phosphate-binding region:GTP | 7 | 1,452 | 0,73047322 | RAP2A, HRAS, EIF5, RAB22A, GNAS, RAB38, TUBG1 | 408 | 319 | 18012 | 0,968744 | 1 | 1 | 100 |

| | | | | | | | | | | | | |
|---|---|---|---|---|---|---|---|---|---|---|---|---|
| UP_KEYWORDS | Palmitate | 6 | 1,245 | 0,73252492 | RAP2A, ZDHHC14, HRAS, AVPR1A, GNAS, RAB38 | 456 | 304 | 22680 | 0,981648 | 1 | 0,9690892 | 99,999998 |
| UP_SEQ_FEATURE | lipid moiety-binding region:S-palmitoyl cysteine | 4 | 0,83 | 0,7731661 | HRAS, AVPR1A, GNAS, RAB38 | 408 | 179 | 18012 | 0,986526 | 1 | 1 | 100 |
| GOTERM_BP_DIRECT | GO:0007264~small GTPase mediated signal transduction | 4 | 0,83 | 0,8900626 | RAP2A, HRAS, RAB22A, RAB38 | 394 | 236 | 18082 | 0,777854 | 1 | 1 | 100 |
| Annotation | Enrichment Score: 0.24006061240135165 | | | | | | | | | | | |
| Category | Term | Count | % | PValue | Genes | List Total | Pop Hits | Pop Total | Enrichment | Bonferroni | Benjamini | FDR |
| KEGG_PATHWAY | mmu04923:Regulation of lipolysis in adipocytes | 3 | 0,622 | 0,44711806 | ADCY1, PDE3B, GNAS | 205 | 57 | 7720 | 1,982028 | 1 | 0,808903 | 99,949565 |
| KEGG_PATHWAY | mmu04024:cAMP signaling pathway | 6 | 1,245 | 0,60013783 | BDNF, ADCY1, MAP2K2, PDE3B, GNAS, ATP1A1 | 205 | 197 | 7720 | 1,146961 | 1 | 0,8800041 | 99,999206 |
| KEGG_PATHWAY | mmu05032:Morphine addiction | 3 | 0,622 | 0,70981463 | ADCY1, PDE3B, GNAS | 205 | 93 | 7720 | 1,214792 | 1 | 0,9344077 | 99,999987 |
| Annotation | Enrichment Score: 0.22474018757475697 | | | | | | | | | | | |
| Category | Term | Count | % | PValue | Genes | List Total | Pop Hits | Pop Total | Enrichment | Bonferroni | Benjamini | FDR |
| UP_KEYWORDS | Cell cycle | 15 | 3,112 | 0,38498977 | MARVELD1, RCC2, HJURP, PTP4A1, MAPK4, SDCCAG3, FBXO31, PELO, UBE2S | 456 | 626 | 22680 | 1,191777 | 1 | 0,8352214 | 99,857956 |
| GOTERM_BP_DIRECT | GO:0007049~cell cycle | 15 | 3,112 | 0,46238464 | MARVELD1, RCC2, HJURP, PTP4A1, MAPK4, SDCCAG3, FBXO31, PELO, UBE2S | 394 | 614 | 18082 | 1,121174 | 1 | 0,9998182 | 99,997384 |
| GOTERM_CC_DIRECT | GO:0030496~midbody | 4 | 0,83 | 0,53246376 | RAP2A, RCC2, SDCCAG3, SIRT2 | 424 | 130 | 19662 | 1,426851 | 1 | 0,9857758 | 99,997056 |
| UP_KEYWORDS | Cell division | 6 | 1,245 | 0,8699068 | SPC25, RCC2, SDCCAG3, PELO, UBE2S, SIRT2 | 456 | 372 | 22680 | 0,802207 | 1 | 0,9927648 | 100 |
| GOTERM_BP_DIRECT | GO:0051301~cell division | 6 | 1,245 | 0,9121866 | SPC25, RCC2, SDCCAG3, PELO, UBE2S, SIRT2 | 394 | 374 | 18082 | 0,736258 | 1 | 1 | 100 |
| Annotation | Enrichment Score: 0.21084253538217115 | | | | | | | | | | | |
| Category | Term | Count | % | PValue | Genes | List Total | Pop Hits | Pop Total | Enrichment | Bonferroni | Benjamini | FDR |
| SMART | SM00326:SH3 | 5 | 1,037 | 0,40532814 | MPP6, SH3D19, BIN1, ARHGAP26, KALRN | 172 | 197 | 10425 | 1,538337 | 1 | 0,9999135 | 99,814372 |
| UP_KEYWORDS | SH3 domain | 5 | 1,037 | 0,60276835 | MPP6, SH3D19, BIN1, ARHGAP26, KALRN | 456 | 208 | 22680 | 1,195597 | 1 | 0,9295789 | 99,999609 |
| INTERPRO | IPR001452:Src homology-3 domain | 5 | 1,037 | 0,65795491 | MPP6, SH3D19, BIN1, ARHGAP26, KALRN | 436 | 212 | 20594 | 1,114008 | 1 | 1 | 99,999994 |
| UP_SEQ_FEATURE | domain:SH3 | 3 | 0,622 | 0,8922294 | MPP6, BIN1, ARHGAP26 | 408 | 166 | 18012 | 0,797838 | 1 | 1 | 100 |
| Annotation | Enrichment Score: 0.158405790825931 | | | | | | | | | | | |

| Category | Term | Count | % | PValue | Genes | List Total | Pop Hits | Pop Total | Enrichment | Bonferroni | Benjamini | FDR |
|---|---|---|---|---|---|---|---|---|---|---|---|---|
| SMART | SM00233:PH | 6 | 1,245 | 0,40098042 | PLEKHF1, PLEKHB1, TIAM2, RAPH1, ARHGAP26, KALRN | 172 | 253 | 10425 | 1,437402 | 1 | 0,9999689 | 99,797267 |
| INTERPRO | IPR001849:Pleckstrin homology domain | 6 | 1,245 | 0,65181602 | PLEKHF1, PLEKHB1, TIAM2, RAPH1, ARHGAP26, KALRN | 436 | 262 | 20594 | 1,081693 | 1 | 1 | 99,999993 |
| INTERPRO | IPR011993:Pleckstrin homology-like domain | 6 | 1,245 | 0,93545529 | PLEKHF1, PLEKHB1, TIAM2, RAPH1, ARHGAP26, KALRN | 436 | 409 | 20594 | 0,692919 | 1 | 1 | 100 |
| UP_SEQ_FEATURE | domain:PH | 3 | 0,622 | 0,95083596 | PLEKHF1, PLEKHB1, ARHGAP26 | 408 | 208 | 18012 | 0,636736 | 1 | 1 | 100 |
| Annotation | Enrichment Score: 0.13317467887300968 | | | | | | | | | | | |
| Category | Term | Count | % | PValue | Genes | List Total | Pop Hits | Pop Total | Enrichment | Bonferroni | Benjamini | FDR |
| UP_SEQ_FEATURE | domain:PDZ | 3 | 0,622 | 0,66926357 | TIAM2, MPP6, PSMD9 | 408 | 101 | 18012 | 1,311299 | 1 | 1 | 99,999998 |
| SMART | SM00228:PDZ | 3 | 0,622 | 0,7092306 | TIAM2, MPP6, PSMD9 | 172 | 150 | 10425 | 1,212209 | 1 | 0,9999994 | 99,999968 |
| INTERPRO | IPR001478:PDZ domain | 3 | 0,622 | 0,83963651 | TIAM2, MPP6, PSMD9 | 436 | 154 | 20594 | 0,920142 | 1 | 1 | 100 |
| Annotation | Enrichment Score: 0.12842985119150016 | | | | | | | | | | | |
| Category | Term | Count | % | PValue | Genes | List Total | Pop Hits | Pop Total | Enrichment | Bonferroni | Benjamini | FDR |
| GOTERM_BP_DIRECT | GO:0055085~transmembrane transport | 9 | 1,867 | 0,5359209 | KCNC3, SLC25A23, ABCB10, SLC46A3, ABCB6, SLC22A30, ITPR1, ABCG2, ITPR2 | 394 | 364 | 18082 | 1,134727 | 1 | 0,999934 | 99,999785 |
| INTERPRO | IPR005821:Ion transport domain | 3 | 0,622 | 0,66378447 | KCNC3, ITPR1, ITPR2 | 436 | 107 | 20594 | 1,324316 | 1 | 1 | 99,999996 |
| GOTERM_MF_DIRECT | GO:0005216~ion channel activity | 3 | 0,622 | 0,89142923 | KCNC3, ITPR1, ITPR2 | 385 | 170 | 17446 | 0,799664 | 1 | 0,9999963 | 100 |
| UP_KEYWORDS | Ion channel | 4 | 0,83 | 0,96619757 | KCNC3, MCOLN1, ITPR1, ITPR2 | 456 | 336 | 22680 | 0,592105 | 1 | 0,999256 | 100 |
| Annotation | Enrichment Score: 0.11872447060469955 | | | | | | | | | | | |
| Category | Term | Count | % | PValue | Genes | List Total | Pop Hits | Pop Total | Enrichment | Bonferroni | Benjamini | FDR |
| UP_KEYWORDS | Calcium transport | 3 | 0,622 | 0,55947722 | MCOLN1, ITPR1, ITPR2 | 456 | 93 | 22680 | 1,604414 | 1 | 0,913192 | 99,998423 |
| GOTERM_BP_DIRECT | GO:0006816~calcium ion transport | 3 | 0,622 | 0,81466347 | MCOLN1, ITPR1, ITPR2 | 394 | 141 | 18082 | 0,976455 | 1 | 1 | 100 |
| UP_KEYWORDS | Ion channel | 4 | 0,83 | 0,96619757 | KCNC3, MCOLN1, ITPR1, ITPR2 | 456 | 336 | 22680 | 0,592105 | 1 | 0,999256 | 100 |
| Annotation | Enrichment Score: 0.11791248213035709 | | | | | | | | | | | |

| Category | Term | Count | % | PValue | Genes | List Total | Pop Hits | Pop Total | Enrichment | Bonferroni | Benjamini | FDR |
|---|---|---|---|---|---|---|---|---|---|---|---|---|
| SMART | SM00360:RRM | 5 | 1,037 | 0,5204367 | PABPN1, SRSF6, ZCRB1, CELF2, CPEB1 | 172 | 229 | 10425 | 1,323373 | 1 | 0,9999801 | 99,986257 |
| UP_KEYWORDS | RNA-binding | 13 | 2,697 | 0,54774248 | MRPL18, RPL11, RPL37, STRBP, CPEB1, ZC3H8, MARS | 456 | 601 | 22680 | 1,075839 | 1 | 0,9143555 | 99,997752 |
| INTERPRO | IPR012677:Nucleotide-binding, alpha-beta plait | 6 | 1,245 | 0,71084949 | PABPN1, SRSF6, ZCRB1, CELF2, CPEB1, BRAP | 436 | 281 | 20594 | 1,008554 | 1 | 1 | 100 |
| UP_KEYWORDS | mRNA processing | 6 | 1,245 | 0,74028668 | PABPN1, SRSF6, ZCRB1, CELF2, CPEB1, ZPR1 | 456 | 307 | 22680 | 0,972056 | 1 | 0,9696965 | 99,999999 |
| INTERPRO | IPR000504:RNA recognition motif domain | 5 | 1,037 | 0,75213042 | PABPN1, SRSF6, ZCRB1, CELF2, CPEB1 | 436 | 241 | 20594 | 0,979957 | 1 | 1 | 100 |
| GOTERM_BP_DIRECT | GO:0006397~mRNA processing | 6 | 1,245 | 0,83230393 | PABPN1, SRSF6, ZCRB1, CELF2, CPEB1, ZPR1 | 394 | 322 | 18082 | 0,855157 | 1 | 1 | 100 |
| UP_KEYWORDS | mRNA splicing | 3 | 0,622 | 0,95511408 | SRSF6, ZCRB1, ZPR1 | 456 | 240 | 22680 | 0,621711 | 1 | 0,9990156 | 100 |
| GOTERM_BP_DIRECT | GO:0008380~RNA splicing | 3 | 0,622 | 0,96891175 | SRSF6, ZCRB1, ZPR1 | 394 | 241 | 18082 | 0,571287 | 1 | 1 | 100 |
| GOTERM_MF_DIRECT | GO:0003676~nucleic acid binding | 14 | 2,905 | 0,99942718 | ENDOD1, CPEB1, ZFP592, SALL2, RPS18, SRSF6, SP5, CELF2, ZFP354A | 385 | 1237 | 17446 | 0,512854 | 1 | 1 | 100 |
| Annotation | Enrichment Score: 0.10884808017769458 | | | | | | | | | | | |
| Category | Term | Count | % | PValue | Genes | List Total | Pop Hits | Pop Total | Enrichment | Bonferroni | Benjamini | FDR |
| GOTERM_CC_DIRECT | GO:0072562~blood microparticle | 5 | 1,037 | 0,32125723 | SERPINA3K, C9, HSPA2, HP, C8G | 424 | 133 | 19662 | 1,743332 | 1 | 0,9341377 | 99,509617 |
| UP_KEYWORDS | Innate immunity | 3 | 0,622 | 0,95587001 | IRAK1, C9, C8G | 456 | 241 | 22680 | 0,619131 | 1 | 0,9990053 | 100 |
| UP_KEYWORDS | Immunity | 5 | 1,037 | 0,9615033 | IRAK1, H2-Q10, C9, HP, C8G | 456 | 401 | 22680 | 0,62016 | 1 | 0,9991886 | 100 |
| GOTERM_BP_DIRECT | GO:0002376~immune system process | 5 | 1,037 | 0,96863867 | IRAK1, H2-Q10, C9, HP, C8G | 394 | 383 | 18082 | 0,599131 | 1 | 1 | 100 |
| GOTERM_BP_DIRECT | GO:0045087~innate immune response | 3 | 0,622 | 0,99860853 | IRAK1, C9, C8G | 394 | 400 | 18082 | 0,344201 | 1 | 1 | 100 |
| Annotation | Enrichment Score: 0.10633895882794644 | | | | | | | | | | | |
| Category | Term | Count | % | PValue | Genes | List Total | Pop Hits | Pop Total | Enrichment | Bonferroni | Benjamini | FDR |
| UP_SEQ_FEATURE | zinc finger region:C2H2-type 7 | 4 | 0,83 | 0,56063067 | SALL2, GTF3A, ZFP354A, ZFP592 | 408 | 129 | 18012 | 1,368901 | 1 | 1 | 99,999798 |
| UP_SEQ_FEATURE | zinc finger region:C2H2-type 9 | 3 | 0,622 | 0,66926357 | GTF3A, ZFP354A, ZFP592 | 408 | 101 | 18012 | 1,311299 | 1 | 1 | 99,999998 |
| UP_SEQ_FEATURE | zinc finger region:C2H2-type 6 | 4 | 0,83 | 0,67115212 | SALL2, GTF3A, ZFP354A, ZFP592 | 408 | 152 | 18012 | 1,161765 | 1 | 1 | 99,999998 |

| | | | | | | | | | | | | |
|---|---|---|---|---|---|---|---|---|---|---|---|---|
| UP_SEQ_FEATURE | zinc finger region:C2H2-type 3 | 6 | 1,245 | 0,70591091 | SALL2, GTF3A, KLF16, SP5, ZFP354A, ZFP511 | 408 | 261 | 18012 | 1,014875 | 1 | 1 | 100 |
| UP_SEQ_FEATURE | zinc finger region:C2H2-type 1 | 6 | 1,245 | 0,71217923 | SALL2, GTF3A, KLF16, SP5, ZFP354A, ZFP511 | 408 | 263 | 18012 | 1,007157 | 1 | 1 | 100 |
| UP_SEQ_FEATURE | zinc finger region:C2H2-type 2 | 6 | 1,245 | 0,72743719 | SALL2, GTF3A, KLF16, SP5, ZFP354A, ZFP511 | 408 | 268 | 18012 | 0,988367 | 1 | 1 | 100 |
| UP_SEQ_FEATURE | zinc finger region:C2H2-type 8 | 3 | 0,622 | 0,75385691 | GTF3A, ZFP354A, ZFP592 | 408 | 119 | 18012 | 1,112951 | 1 | 1 | 100 |
| SMART | SM00355:ZnF_C2H2 | 10 | 2,075 | 0,81042032 | SALL2, GTF3A, KCMF1, ZFP598, KLF16, SP5, ZFP354A, DNAJC21, ZFP511, ZFP592 | 172 | 693 | 10425 | 0,87461 | 1 | 0,9999999 | 100 |
| UP_SEQ_FEATURE | zinc finger region:C2H2-type 4 | 4 | 0,83 | 0,86741081 | SALL2, GTF3A, ZFP354A, ZFP592 | 408 | 215 | 18012 | 0,821341 | 1 | 1 | 100 |
| UP_SEQ_FEATURE | zinc finger region:C2H2-type 5 | 3 | 0,622 | 0,93588565 | SALL2, GTF3A, ZFP354A | 408 | 194 | 18012 | 0,682686 | 1 | 1 | 100 |
| INTERPRO | IPR007087:Zinc finger, C2H2 | 11 | 2,282 | 0,94810849 | SALL2, GTF3A, KCMF1, ZFP598, ZFP703, KLF16, SP5, ZFP354A, DNAJC21, ZFP511, ZFP592 | 436 | 731 | 20594 | 0,710771 | 1 | 1 | 100 |
| INTERPRO | IPR015880:Zinc finger, C2H2-like | 10 | 2,075 | 0,9590081 | SALL2, GTF3A, KCMF1, ZFP598, KLF16, SP5, ZFP354A, DNAJC21, ZFP511, ZFP592 | 436 | 693 | 20594 | 0,681587 | 1 | 1 | 100 |
| INTERPRO | IPR013087:Zinc finger C2H2-type/integrase DNA-binding domain | 6 | 1,245 | 0,9982439 | SALL2, GTF3A, ZFP703, KLF16, SP5, ZFP354A | 436 | 652 | 20594 | 0,434668 | 1 | 1 | 100 |
| Annotation | Enrichment Score: 0.09137817452296763 | | | | | | | | | | | |
| Category | Term | Count | % | PValue | Genes | List Total | Pop Hits | Pop Total | Enrichment | Bonferroni | Benjamini | FDR |
| GOTERM_CC_DIRECT | GO:0000775~chromosome, centromeric region | 5 | 1,037 | 0,36033584 | SPC25, RCC2, HJURP, CBX3, CBX1 | 424 | 141 | 19662 | 1,64442 | 1 | 0,9482949 | 99,782664 |
| UP_KEYWORDS | Centromere | 3 | 0,622 | 0,75323218 | SPC25, RCC2, HJURP | 456 | 134 | 22680 | 1,113511 | 1 | 0,9710646 | 99,999999 |
| UP_KEYWORDS | Cell division | 6 | 1,245 | 0,8699068 | SPC25, RCC2, SDCCAG3, PELO, UBE2S, SIRT2 | 456 | 372 | 22680 | 0,802207 | 1 | 0,9927648 | 100 |
| GOTERM_BP_DIRECT | GO:0051301~cell division | 6 | 1,245 | 0,9121866 | SPC25, RCC2, SDCCAG3, PELO, UBE2S, SIRT2 | 394 | 374 | 18082 | 0,736258 | 1 | 1 | 100 |
| GOTERM_BP_DIRECT | GO:0007067~mitotic nuclear division | 4 | 0,83 | 0,9421917 | SPC25, RCC2, RPS6, SIRT2 | 394 | 277 | 18082 | 0,662721 | 1 | 1 | 100 |
| UP_KEYWORDS | Chromosome | 5 | 1,037 | 0,96559498 | SPC25, HIST1H2BC, RCC2, HJURP, SIRT2 | 456 | 409 | 22680 | 0,60803 | 1 | 0,9993326 | 100 |
| UP_KEYWORDS | Mitosis | 3 | 0,622 | 0,96756285 | SPC25, RCC2, SIRT2 | 456 | 259 | 22680 | 0,576102 | 1 | 0,9992848 | 100 |
| GOTERM_CC_DIRECT | GO:0005694~chromosome | 4 | 0,83 | 0,97986989 | SPC25, RCC2, HJURP, SIRT2 | 424 | 344 | 19662 | 0,539217 | 1 | 0,9999994 | 100 |
| Annotation | Enrichment Score: 0.07646915209679844 | | | | | | | | | | | |
| Category | Term | Count | % | PValue | Genes | List Total | Pop Hits | Pop Total | Enrichment | Bonferroni | Benjamini | FDR |

| | | | | | | | | | | | | |
|---|---|---|---|---|---|---|---|---|---|---|---|---|
| GOTERM_MF_DIRECT | GO:0004252~serine-type endopeptidase activity | 5 | 1,037 | 0,6895472 | F10, CTSC, HP, PCSK6, PCSK4 | 385 | 212 | 17446 | 1,068733 | 1 | 0,9994641 | 99,999997 |
| UP_KEYWORDS | Thiol protease | 3 | 0,622 | 0,75323218 | ATG4D, FAM188A, CTSC | 456 | 134 | 22680 | 1,113511 | 1 | 0,9710646 | 99,999999 |
| GOTERM_MF_DIRECT | GO:0008234~cysteine-type peptidase activity | 3 | 0,622 | 0,82021477 | ATG4D, FAM188A, CTSC | 385 | 141 | 17446 | 0,964134 | 1 | 0,9999558 | 100 |
| UP_KEYWORDS | Protease | 9 | 1,867 | 0,85422827 | CPM, F10, ATG4D, H13, MME, FAM188A, CTSC, PCSK6, PCSK4 | 456 | 542 | 22680 | 0,825889 | 1 | 0,9911329 | 100 |
| UP_KEYWORDS | Serine protease | 3 | 0,622 | 0,86467724 | F10, PCSK6, PCSK4 | 456 | 173 | 22680 | 0,862489 | 1 | 0,9923301 | 100 |
| GOTERM_BP_DIRECT | GO:0006508~proteolysis | 10 | 2,075 | 0,88964979 | CPM, F10, ATG4D, H13, SHFM1, MME, FAM188A, CTSC, HP, PCSK4 | 394 | 582 | 18082 | 0,788546 | 1 | 1 | 100 |
| UP_KEYWORDS | Zymogen | 3 | 0,622 | 0,92936421 | F10, CTSC, PCSK4 | 456 | 213 | 22680 | 0,700519 | 1 | 0,9977168 | 100 |
| GOTERM_MF_DIRECT | GO:0008233~peptidase activity | 8 | 1,66 | 0,9397089 | CPM, F10, ATG4D, H13, MME, FAM188A, CTSC, PCSK4 | 385 | 516 | 17446 | 0,702547 | 1 | 0,9999998 | 100 |
| Annotation | Enrichment Score: 0.05409512342443709 | | | | | | | | | | | |
| Category | Term | Count | % | PValue | Genes | List Total | Pop Hits | Pop Total | Enrichment | Bonferroni | Benjamini | FDR |
| UP_KEYWORDS | Ion transport | 11 | 2,282 | 0,79863524 | SLC25A28, SLC25A37, ATP1A1, MCOLN1, ITPR1, ITPR2 | 456 | 619 | 22680 | 0,883853 | 1 | 0,9815755 | 100 |
| GOTERM_BP_DIRECT | GO:0006811~ion transport | 10 | 2,075 | 0,89186726 | SLCO1A1, ATP5J2, KCNC3, SLC38A2, FXN, SLC25A28, ATP1A1, MCOLN1, ITPR1, ITPR2 | 394 | 584 | 18082 | 0,785846 | 1 | 1 | 100 |
| UP_KEYWORDS | Ion channel | 4 | 0,83 | 0,96619757 | KCNC3, MCOLN1, ITPR1, ITPR2 | 456 | 336 | 22680 | 0,592105 | 1 | 0,999256 | 100 |
| Annotation | Enrichment Score: 0.048623327283356596 | | | | | | | | | | | |
| Category | Term | Count | % | PValue | Genes | List Total | Pop Hits | Pop Total | Enrichment | Bonferroni | Benjamini | FDR |
| GOTERM_BP_DIRECT | transcription from RNA polymerase II promoter | 17 | 3,527 | 0,52660483 | HES6, SIRT2, ZC3H8, UXT, SALL2, RPS14, MAPK15, HOPX, ZFP354A, URI1, SUDS3 | 394 | 729 | 18082 | 1,070216 | 1 | 0,9999257 | 99,999699 |
| UP_KEYWORDS | Transcription regulation | 30 | 6,224 | 0,91173164 | ZFP354A, WAC, MN1, MAFF, CEBPB, EPAS1, ZFP703, KLF16, GTF2IRD2, PHF10, IRF2BP2, TLE2, HES6, SIRT2, ZFP592, SALL2, UXT, NR1I3, ATF4, | 456 | 1799 | 22680 | 0,829408 | 1 | 0,9964897 | 100 |
| UP_KEYWORDS | Transcription | 31 | 6,432 | 0,91426355 | ZFP354A, WAC, MN1, MAFF, CEBPB, EPAS1, POLR1D, ZFP703, KLF16, GTF2IRD2, PHF10, IRF2BP2, TLE2, HES6, SIRT2, ZFP592, SALL2, UXT, NR1I3, ATF4, HOPX, SP5, ZFP511, URI1, SUDS3 | 456 | 1859 | 22680 | 0,829393 | 1 | 0,9965809 | 100 |
| GOTERM_MF_DIRECT | GO:0003700~transcription factor activity, sequence-specific DNA binding | 14 | 2,905 | 0,95569454 | HES6, MYCN, ZC3H8, SALL2, ATF4, NR1I3, TSC22D2, ZFP354A | 385 | 883 | 17446 | 0,71846 | 1 | 1 | 100 |

| | | | | | | | | | | | | |
|---|---|---|---|---|---|---|---|---|---|---|---|---|
| GOTERM_BP_DIRECT | GO:0006351~transcription, DNA-templated | 31 | 6,432 | 0,97614271 | ZFP354A, WAC, MN1, MAFF, CEBPB, EPAS1, POLR1D, ZFP703, KLF16, GTF2IRD2, PHF10, IRF2BP2, TLE2, HES6, SIRT2, ZFP592, SALL2, UXT, NR1I3, ATF4, HOPX, SP5, ZFP511, URI1, SUDS3 | 394 | 1885 | 18082 | 0,754746 | 1 | 1 | 100 |
| UP_KEYWORDS | DNA-binding | 19 | 3,942 | 0,99824606 | PPARG, KLF16, GTF2IRD2, MCM10, MYCN, ZFP592, SALL2, NR1I3, ATF4, HJURP, SP5, | 456 | 1604 | 22680 | 0,589152 | 1 | 0,999998 | 100 |
| GOTERM_BP_DIRECT | GO:0006355~regulation of transcription, DNA-templated | 32 | 6,639 | 0,9990167 | CREG1, ZFP354A, WAC, MN1, MAFF, CEBPB, EPAS1, ZFP703, KLF16, GTF2IRD2, PHF10, IRF2BP2, TLE2, HES6, SIRT2, ZFP592, MYCN, SALL2, UXT, NR1I3, ATF4, HOPX, SP5, ZFP511, | 394 | 2279 | 18082 | 0,644401 | 1 | 1 | 100 |
| GOTERM_MF_DIRECT | GO:0003677~DNA binding | 22 | 4,564 | 0,99987874 | POLR1D, PPARG, KLF16, GTF2IRD2, HES6, MCM10, MYCN, ZFP592, SALL2, ATF4, NR1I3, HJURP, HOPX, SP5, ZFP354A, STRBP, ZFP511 | 385 | 1847 | 17446 | 0,539748 | 1 | 1 | 100 |
| Annotation | Enrichment Score: 0.04549558150251625 | | | | | | | | | | | |
| Category | Term | Count | % | PValue | Genes | List Total | Pop Hits | Pop Total | Enrichment | Bonferroni | Benjamini | FDR |
| SMART | SM00408:IGc2 | 4 | 0,83 | 0,76323317 | LRIT2, CADM4, NTRK2, KALRN | 172 | 242 | 10425 | 1,001826 | 1 | 0,9999998 | 99,999997 |
| UP_SEQ_FEATURE | domain:Ig-like C2-type 1 | 3 | 0,622 | 0,80275219 | CADM4, PDGFRL, NTRK2 | 408 | 132 | 18012 | 1,003342 | 1 | 1 | 100 |
| UP_SEQ_FEATURE | domain:Ig-like C2-type 2 | 3 | 0,622 | 0,80613183 | CADM4, PDGFRL, NTRK2 | 408 | 133 | 18012 | 0,995798 | 1 | 1 | 100 |
| INTERPRO | IPR003598:Immunoglobulin subtype 2 | 4 | 0,83 | 0,88847157 | LRIT2, CADM4, NTRK2, KALRN | 436 | 242 | 20594 | 0,780726 | 1 | 1 | 100 |
| SMART | SM00409:IG | 6 | 1,245 | 0,93204673 | LRIT2, CADM4, PDGFRL, NTRK2, ILDR2, KALRN | 172 | 518 | 10425 | 0,702052 | 1 | 1 | 100 |
| UP_KEYWORDS | Immunoglobulin domain | 6 | 1,245 | 0,96605612 | LRIT2, CADM4, PDGFRL, NTRK2, ILDR2, KALRN | 456 | 481 | 22680 | 0,620418 | 1 | 0,9992848 | 100 |
| INTERPRO | IPR003599:Immunoglobulin subtype | 6 | 1,245 | 0,98594042 | LRIT2, CADM4, PDGFRL, NTRK2, ILDR2, KALRN | 436 | 518 | 20594 | 0,547111 | 1 | 1 | 100 |
| INTERPRO | IPR007110:Immunoglobulin-like domain | 7 | 1,452 | 0,99992391 | LRIT2, H2-Q10, CADM4, PDGFRL, NTRK2, ILDR2, KALRN | 436 | 920 | 20594 | 0,359389 | 1 | 1 | 100 |
| INTERPRO | IPR013783:Immunoglobulin-like fold | 8 | 1,66 | 0,99998564 | IFNAR2, LRIT2, H2-Q10, CADM4, PDGFRL, NTRK2, ILDR2, KALRN | 436 | 1099 | 20594 | 0,343832 | 1 | 1 | 100 |
| Annotation | Enrichment Score: 0.03691428153852507 | | | | | | | | | | | |
| Category | Term | Count | % | PValue | Genes | List Total | Pop Hits | Pop Total | Enrichment | Bonferroni | Benjamini | FDR |
| GOTERM_CC_DIRECT | GO:0045202~synapse | 9 | 1,867 | 0,85404613 | EGFR, DLGAP1, TMEM57, DTNB, MME, HSPB1, CPEB1, NRN1, DTNBP1 | 424 | 505 | 19662 | 0,826443 | 1 | 0,999602 | 100 |
| UP_KEYWORDS | Postsynaptic cell membrane | 3 | 0,622 | 0,87098192 | DLGAP1, CPEB1, DTNBP1 | 456 | 176 | 22680 | 0,847787 | 1 | 0,9926396 | 100 |

| | | | | | | | | | | | | |
|---|---|---|---|---|---|---|---|---|---|---|---|---|
| UP_KEYWORDS | Synapse | 5 | 1,037 | 0,92982184 | DLGAP1, KCNC3, CPEB1, NRN1, DTNBP1 | 456 | 357 | 22680 | 0,696594 | 1 | 0,9976504 | 100 |
| UP_KEYWORDS | Cell junction | 9 | 1,867 | 0,95667458 | NOX4, DLGAP1, KCNC3, ATAT1, CPEB1, NRN1, DTNBP1, TRAF4, ARHGAP26 | 456 | 661 | 22680 | 0,677204 | 1 | 0,9989981 | 100 |
| GOTERM_CC_DIRECT | GO:0030054~cell junction | 9 | 1,867 | 0,98803913 | NOX4, NAMPT, DLGAP1, ATAT1, CPEB1, NRN1, DTNBP1, TRAF4, ARHGAP26 | 424 | 718 | 19662 | 0,581273 | 1 | 0,9999999 | 100 |
| Annotation | Enrichment Score: 0.023689826284511124 | | | | | | | | | | | |
| Category | Term | Count | % | PValue | Genes | List Total | Pop Hits | Pop Total | Enrichment | Bonferroni | Benjamini | FDR |
| GOTERM_BP_DIRECT | GO:0007166~cell surface receptor signaling pathway | 4 | 0,83 | 0,85814839 | EGFR, FZD8, TSPAN33, GCGR | 394 | 219 | 18082 | 0,838236 | 1 | 1 | 100 |
| GOTERM_MF_DIRECT | GO:0004888~transmembrane signaling receptor activity | 3 | 0,622 | 0,99318344 | EGFR, FZD8, GCGR | 385 | 316 | 17446 | 0,430199 | 1 | 1 | 100 |
| GOTERM_MF_DIRECT | GO:0004871~signal transducer activity | 7 | 1,452 | 0,99618193 | EGFR, FZD8, GRM8, P2RY1, AVPR1A, GNAS, GCGR | 385 | 648 | 17446 | 0,489506 | 1 | 1 | 100 |
| Annotation | Enrichment Score: 0.012629600068504103 | | | | | | | | | | | |
| Category | Term | Count | % | PValue | Genes | List Total | Pop Hits | Pop Total | Enrichment | Bonferroni | Benjamini | FDR |
| SMART | SM00320:WD40 | 3 | 0,622 | 0,93192261 | WDR72, TLE2, PPP2R2A | 172 | 262 | 10425 | 0,694013 | 1 | 1 | 100 |
| UP_KEYWORDS | WD repeat | 3 | 0,622 | 0,965846 | WDR72, TLE2, PPP2R2A | 456 | 256 | 22680 | 0,582854 | 1 | 0,9993098 | 100 |
| INTERPRO | IPR001680:WD40 repeat | 3 | 0,622 | 0,9762853 | WDR72, TLE2, PPP2R2A | 436 | 263 | 20594 | 0,53879 | 1 | 1 | 100 |
| INTERPRO | IPR017986:WD40-repeat-containing domain | 3 | 0,622 | 0,98952912 | WDR72, TLE2, PPP2R2A | 436 | 307 | 20594 | 0,461569 | 1 | 1 | 100 |
| INTERPRO | IPR015943:WD40/YVTN repeat-like-containing domain | 3 | 0,622 | 0,994395 | WDR72, TLE2, PPP2R2A | 436 | 340 | 20594 | 0,41677 | 1 | 1 | 100 |
| Annotation | Enrichment Score: 0.008462694806630536 | | | | | | | | | | | |
| Category | Term | Count | % | PValue | Genes | List Total | Pop Hits | Pop Total | Enrichment | Bonferroni | Benjamini | FDR |
| INTERPRO | IPR001611:Leucine-rich repeat | 3 | 0,622 | 0,97448197 | LRIT2, NTRK2, TSKU | 436 | 259 | 20594 | 0,547111 | 1 | 1 | 100 |
| UP_KEYWORDS | Leucine-rich repeat | 3 | 0,622 | 0,97540514 | LRIT2, NTRK2, TSKU | 456 | 275 | 22680 | 0,542584 | 1 | 0,9995803 | 100 |
| UP_SEQ_FEATURE | repeat:LRR 1 | 3 | 0,622 | 0,98649568 | LRIT2, NTRK2, TSKU | 408 | 274 | 18012 | 0,483362 | 1 | 1 | 100 |
| UP_SEQ_FEATURE | repeat:LRR 2 | 3 | 0,622 | 0,98649568 | LRIT2, NTRK2, TSKU | 408 | 274 | 18012 | 0,483362 | 1 | 1 | 100 |
| Annotation | Enrichment Score: | | | | | | | | | | | |

| Category | Term | Count | % | PValue | Genes | List Total | Pop Hits | Pop Total | Enrichment | Bonferroni | Benjamini | FDR |
|---|---|---|---|---|---|---|---|---|---|---|---|---|
| KEGG_PATHWAY | mmu04080:Neuroactive ligand-receptor interaction | 4 | 0,83 | 0,98279965 | GRM8, P2RY1, AVPR1A, GCGR | 205 | 285 | 7720 | 0,528541 | 1 | 0,9992625 | 100 |
| GOTERM_MF_DIRECT | GO:0004871~signal transducer activity | 7 | 1,452 | 0,99618193 | EGFR, FZD8, GRM8, P2RY1, AVPR1A, GNAS, GCGR | 385 | 648 | 17446 | 0,489506 | 1 | 1 | 100 |
| GOTERM_BP_DIRECT | GO:0007186~G-protein coupled receptor signaling pathway | 8 | 1,66 | 1 | FZD8, GDE1, GRM8, P2RY1, CCL9, AVPR1A, GNAS, GCGR | 394 | 1706 | 18082 | 0,215209 | 1 | 1 | 100 |
| UP_KEYWORDS | Transducer | 6 | 1,245 | 1 | FZD8, GRM8, P2RY1, AVPR1A, GNAS, GCGR | 456 | 1801 | 22680 | 0,165697 | 1 | 1 | 100 |
| UP_KEYWORDS | G-protein coupled receptor | 5 | 1,037 | 1 | FZD8, GRM8, P2RY1, AVPR1A, GCGR | 456 | 1741 | 22680 | 0,14284 | 1 | 1 | 100 |
| GOTERM_MF_DIRECT | GO:0004930~G-protein coupled receptor activity | 5 | 1,037 | 1 | FZD8, GRM8, P2RY1, AVPR1A, GCGR | 385 | 1749 | 17446 | 0,129543 | 1 | 1 | 100 |
