## Supplementary material for "The mouse HP1 proteins are essential for preventing liver tumorigenesis": suppl table 4

**Supplementary Table 4** : HP1-dependent genes belonging to the IFNγ response pathway

| gene name | log2Fold Change | padj |
|---|---|---|
| Rarres1 | -2,58820339 | 1,25E-14 |
| Colec12 | -2,23875669 | 1,12E-18 |
| Pcp4l1 | -1,70183437 | 3,98E-09 |
| Gadd45g | -1,65639913 | 0,00012264 |
| Ugt1a9 | -1,60897243 | 1,69E-07 |
| Arhgap26 | -1,5123532 | 6,77E-08 |
| Cpeb1 | -1,49357446 | 0,00026173 |
| Pdgfrl | -1,3855974 | 0,00487983 |
| 1190005I06Ri | -1,31648695 | 0,00853929 |
| Rbp4 | -1,30412565 | 1,91E-07 |
| Mthfd2 | -1,2801751 | 0,00185791 |
| Zdhhc14 | -1,27576213 | 0,00024428 |
| Mycn | -1,25871188 | 0,01732341 |
| Mcm10 | -1,22624317 | 6,77E-07 |
| Pparg | -1,21175115 | 0,00019636 |
| Endod1 | -1,17842219 | 0,00031757 |
| Gde1 | -1,16737678 | 6,11E-07 |
| Tspan33 | -1,16276925 | 0,00320933 |
| Pnpla1 | -1,15077743 | 0,00177917 |
| Hspa1b | -1,1450821 | 0,01770805 |
| Psat1 | -1,07953495 | 0,0106979 |
| Hspa2 | -1,07166411 | 0,01782915 |
| Glrx | -1,06707139 | 1,11E-06 |
| Asns | -1,05715195 | 0,00310925 |
| Tmem41a | -1,03145592 | 0,00290452 |
| Timm9 | -1,01568928 | 5,43E-08 |
| Osgin1 | -0,98145616 | 0,03345025 |
| Hist1h2bc | -0,93943657 | 0,03452248 |
| Ccl9 | -0,93941191 | 0,00411738 |
| Hp | -0,93267427 | 0,0167057 |
| Creg1 | -0,91506452 | 0,00191577 |
| Ung | -0,90871977 | 0,03133445 |
| Tmx4 | -0,89204561 | 0,00198819 |
| Slc35g1 | -0,88713997 | 0,01730065 |
| Slc19a1 | -0,88292497 | 0,00523511 |
| Cebpb | -0,87244466 | 0,00029838 |
| Paqr7 | -0,86852436 | 0,00245525 |
| Mt1 | -0,86811881 | 0,02814528 |
| Acp5 | -0,85822856 | 0,00241393 |
| Plekhf1 | -0,84650749 | 0,03704255 |
| Cryl1 | -0,83256098 | 0,00028678 |
| Cyp27a1 | -0,82969688 | 1,30E-05 |
| 1190002N15R | -0,82350252 | 0,00446713 |
| H2-Q10 | -0,79835742 | 0,00687099 |
| P2ry1 | -0,7815947 | 0,00895751 |

| | | |
|---|---|---|
| Ogfrl1 | -0,7766678 | 0,00980375 |
| Txnrd3 | -0,77363288 | 0,01099393 |
| Fbxo31 | -0,77183504 | 0,02074126 |
| Ttc7b | -0,7717874 | 0,00638523 |
| Slc46a3 | -0,7587901 | 0,02233571 |
| F10 | -0,74811823 | 0,00011131 |
| Ifnar2 | -0,74406335 | 0,00096175 |
| Pim3 | -0,74105429 | 0,01052143 |
| Itpr2 | -0,7359572 | 0,0002099 |
| Cyb5b | -0,73290166 | 0,00016212 |
| Hopx | -0,72343786 | 0,00621831 |
| Fads1 | -0,70185047 | 0,00020878 |
| Ugt1a1 | -0,69834176 | 0,03045486 |
| Tubg1 | -0,69087749 | 0,04749377 |
| Pop5 | -0,6815505 | 0,01475585 |
| Dexi | -0,66414674 | 0,03496857 |
| Noc4l | -0,6512646 | 0,04067035 |
| Tmcc1 | -0,64747816 | 0,03185868 |
| Slc25a37 | -0,64197636 | 0,00444528 |
| Ivns1abp | -0,63793089 | 0,02161066 |
| Gtf3a | -0,63774466 | 0,0304599 |
| Nop56 | -0,62475514 | 0,0039222 |
| Nampt | -0,62109888 | 0,01468309 |
| H13 | -0,62077046 | 0,011961 |
| Raph1 | -0,60769717 | 0,04473236 |
| Dtnbp1 | -0,60003768 | 0,0261511 |
| Aox1 | -0,5982157 | 0,02261104 |
| Tm7sf2 | -0,59165146 | 0,03976649 |
| Dnajc21 | -0,58890097 | 0,04711692 |
| Mcoln1 | -0,57761993 | 0,0184435 |
| Gstm1 | -0,57110923 | 0,01070874 |
| Sephs2 | -0,55798417 | 0,00853929 |
| Itpr1 | -0,55772336 | 0,0065641 |
| Irf2bp2 | -0,54805448 | 0,03699135 |
| March5 | -0,54549909 | 0,03419988 |
| Rpl24 | -0,54351307 | 0,00799431 |
| Ubac1 | -0,54342552 | 0,04378035 |
| Epas1 | -0,54233826 | 0,02894033 |
| Phyhd1 | -0,5324508 | 0,00809896 |
| Cyhr1 | -0,53051275 | 0,0167057 |
| Cmas | -0,525202 | 0,02112 |
| Ctsc | -0,52448658 | 0,01239299 |
| Ephx1 | -0,52357156 | 0,02322637 |
| Atf4 | -0,51171658 | 0,04841236 |
| Ahcy | -0,51023241 | 0,0136709 |
| Mafb | -0,49545789 | 0,34042993 |
| Cnih1 | -0,48839165 | 0,02588276 |
| Rcl1 | -0,47657125 | 0,02055361 |
| Nme1 | -0,47063056 | 0,03815082 |
| Ripk1 | -0,45954349 | 0,04796617 |

| | | |
|---|---|---|
| Coa5 | -0,45487103 | 0,04796617 |
| C4b | -0,33183117 | 0,28943071 |
| H2-Q5 | -0,25552403 | 0,6561026 |
| Megf8 | -0,24756742 | 0,64094861 |
| H2-K1 | -0,04148323 | 0,94310814 |
| Psme2 | 0,01719254 | 0,97217752 |
| Sp2 | 0,0689235 | 0,93530955 |
| Lmod1 | 0,07065146 | 0,94298204 |
| Mndal | 0,0809936 | 0,89601344 |
| Plxdc2 | 0,15296749 | 0,85888091 |
| Mnda | 0,17943096 | 0,82005173 |
| Mansc4 | 0,26693371 | 0,71718844 |
| Smc4 | 0,38787658 | 0,16869853 |
| Sdc1 | 0,50099946 | 0,04847907 |
| Cr1l | 0,52871866 | 0,02239576 |
| Ehd1 | 0,55071086 | 0,01048884 |
| Acad9 | 0,57140446 | 0,01721033 |
| Il18bp | 0,59615315 | 0,01304052 |
| Cd93 | 0,60790734 | 0,0348796 |
| H2-Q4 | 0,60993443 | 0,013577 |
| Amigo2 | 0,62722274 | 0,03349693 |
| P2ry13 | 0,6306553 | 0,03894412 |
| Gls | 0,63393932 | 0,03766085 |
| Sema4a | 0,63692349 | 0,04015343 |
| Id2 | 0,64282705 | 0,04433029 |
| Cdkn2c | 0,65517877 | 0,04448139 |
| Mid1ip1 | 0,65959427 | 0,04218888 |
| Alcam | 0,66360332 | 0,0073558 |
| Tlr7 | 0,66746166 | 0,04448139 |
| Klf6 | 0,67101833 | 0,01424369 |
| Herc6 | 0,67216083 | 0,0227596 |
| Kctd12 | 0,67549388 | 0,0264677 |
| Tstd3 | 0,68329683 | 0,02790779 |
| Myh10 | 0,68354377 | 0,04290116 |
| Tuft1 | 0,69919673 | 0,03196404 |
| Papss2 | 0,70890841 | 0,03452248 |
| Tra2a | 0,72148214 | 0,02522906 |
| Dab2ip | 0,73119228 | 0,03992849 |
| Slc43a2 | 0,73195738 | 0,02763346 |
| Fam111a | 0,74291378 | 0,01462021 |
| Trim21 | 0,74533454 | 0,02336772 |
| Calcrl | 0,74634014 | 0,04497473 |
| Tlr3 | 0,7565764 | 0,0247151 |
| Sgcb | 0,76145295 | 0,04647205 |
| Pygb | 0,76383879 | 0,01710272 |
| Parp2 | 0,77239395 | 0,03437957 |
| Ggct | 0,77262725 | 0,00980375 |
| Csrp1 | 0,77462962 | 0,00074725 |
| Ccr5 | 0,78375313 | 0,02031294 |
| Slc13a3 | 0,7981032 | 0,02066408 |

| | | |
|---|---|---|
| Casp12 | 0,80508165 | 0,01711967 |
| Tap1 | 0,80705725 | 0,00325509 |
| Dtx4 | 0,80729853 | 0,01870738 |
| Pstpip2 | 0,81360617 | 0,00393493 |
| Fancl | 0,82349452 | 0,02079577 |
| Mfge8 | 0,82374524 | 0,01624477 |
| Gbp2 | 0,82374586 | 0,00154028 |
| Pola1 | 0,83064945 | 0,01158574 |
| Fam129a | 0,83829881 | 0,04792174 |
| Rps6ka3 | 0,83986643 | 0,00134202 |
| Far1 | 0,84067575 | 0,03741436 |
| Clec7a | 0,85378584 | 0,01153743 |
| Nmi | 0,85676665 | 0,01470357 |
| Rhoc | 0,85839599 | 0,00072567 |
| Uba7 | 0,86902827 | 0,00487983 |
| H2-T22 | 0,87248652 | 0,06046715 |
| Vcam1 | 0,87259343 | 0,0095288 |
| Oxct1 | 0,87876751 | 0,00207396 |
| Rapgef5 | 0,87881336 | 0,00948245 |
| Gbp6 | 0,88348215 | 0,01932871 |
| Pck2 | 0,88493332 | 0,00635183 |
| Thra | 0,89426957 | 0,01568138 |
| Cd300lb | 0,89457401 | 0,03121424 |
| Lbh | 0,89860482 | 0,00410946 |
| Stk10 | 0,90038143 | 0,01782737 |
| Oas1a | 0,90043983 | 0,0271498 |
| Ube2l6 | 0,90648508 | 0,00019738 |
| Gbp10 | 0,91081093 | 0,02894033 |
| Kcne3 | 0,91388778 | 0,04349069 |
| Nab2 | 0,9227626 | 0,01722972 |
| Nmral1 | 0,92906036 | 0,00100707 |
| Ifi205 | 0,93243436 | 0,03518608 |
| Anxa2 | 0,95301039 | 5,09E-05 |
| Mycl | 0,95390866 | 0,0167057 |
| Gbp3 | 0,95405666 | 0,00055754 |
| Ppic | 0,95590494 | 0,00247095 |
| S100a4 | 0,97076433 | 0,03196404 |
| Parp11 | 0,97361317 | 0,00307499 |
| Rfx5 | 0,98827867 | 0,00121684 |
| Lrp11 | 0,99099492 | 0,04104012 |
| Mpeg1 | 0,99217958 | 2,30E-06 |
| Qpct | 1,00263426 | 0,00149235 |
| Tnfsf10 | 1,00320243 | 0,0054377 |
| Serpinb1a | 1,00937982 | 0,01871619 |
| Cxcl10 | 1,00979164 | 0,00221525 |
| H2-T10 | 1,01682912 | 0,0089777 |
| Gpr35 | 1,02626455 | 0,03349307 |
| Lnpep | 1,03030822 | 0,00403005 |
| Serpinb8 | 1,03051598 | 0,01124887 |
| Tarm1 | 1,03883376 | 0,04774951 |

| | | |
|---|---|---|
| Cnp | 1,03958649 | 0,0001948 |
| Tlr8 | 1,04021266 | 0,02580831 |
| Emp2 | 1,04559696 | 9,04E-06 |
| Selp | 1,04887273 | 0,02673341 |
| Zik1 | 1,05077199 | 0,04036475 |
| Serpina3i | 1,08091166 | 0,04869175 |
| Il1rn | 1,0828239 | 0,00238712 |
| Slc8a1 | 1,08462704 | 0,00502003 |
| Sorl1 | 1,08473487 | 0,04395424 |
| Ifit2 | 1,08946555 | 3,82E-05 |
| Gm2a | 1,09096726 | 2,85E-08 |
| Ly6e | 1,0995098 | 5,79E-05 |
| Gbp7 | 1,10166018 | 0,00085201 |
| Msantd3 | 1,10622361 | 0,04843577 |
| Uap1l1 | 1,10650365 | 1,01E-05 |
| Esr1 | 1,10952672 | 0,00592376 |
| S100a11 | 1,11251529 | 0,00035788 |
| Cdkn1a | 1,11806939 | 0,0078156 |
| Trim30d | 1,12149285 | 0,00185791 |
| Ccl2 | 1,12187135 | 0,01678878 |
| Dcaf4 | 1,13079062 | 0,0130671 |
| Mmp12 | 1,13880394 | 0,03196404 |
| March1 | 1,13989604 | 0,00068222 |
| Ttc30a2 | 1,14977597 | 0,03766085 |
| Aph1b | 1,15001811 | 0,00176928 |
| Ccr2 | 1,15083302 | 0,00258817 |
| Spp1 | 1,15433709 | 0,00241197 |
| Blnk | 1,16163764 | 0,01040626 |
| Trim56 | 1,17612756 | 0,00888503 |
| Zfp558 | 1,17614455 | 0,01054773 |
| Tgtp1 | 1,18126617 | 0,00692865 |
| 9930111J21Ri | 1,18244523 | 0,02449172 |
| Bcl2l14 | 1,19718007 | 0,02470927 |
| Mgst2 | 1,19977256 | 0,02748339 |
| Trim30a | 1,20567027 | 1,93E-06 |
| Osbpl3 | 1,21190454 | 0,00639622 |
| Cxcl14 | 1,21613773 | 0,00973158 |
| Plek | 1,21921211 | 2,67E-05 |
| Ptgr1 | 1,22166518 | 9,36E-06 |
| Psmb9 | 1,24398482 | 3,37E-06 |
| H2-Q2 | 1,25798614 | 0,00584597 |
| Gbp9 | 1,26423459 | 5,16E-05 |
| Tnfrsf22 | 1,26750527 | 0,01711967 |
| Src | 1,28680343 | 0,00146611 |
| Emp1 | 1,28989286 | 0,0019087 |
| Mlkl | 1,29501832 | 0,00020936 |
| Clic5 | 1,29722036 | 0,00813123 |
| Pak1 | 1,30179128 | 6,53E-06 |
| Nptx1 | 1,31052455 | 0,01013733 |
| Dhrs9 | 1,31142308 | 0,00543764 |

| | | |
|---|---|---|
| Cd244 | 1,31295132 | 0,00900022 |
| Prrg1 | 1,31976587 | 0,00262287 |
| Ocln | 1,33269398 | 4,57E-05 |
| Samd9l | 1,36311404 | 7,30E-06 |
| Timp1 | 1,36802799 | 0,00790976 |
| Greb1l | 1,38103932 | 0,00339396 |
| Zfp369 | 1,40189964 | 0,00185791 |
| C4a | 1,4048805 | 0,00025132 |
| Oasl1 | 1,43333246 | 0,00018136 |
| Senp7 | 1,45373823 | 8,93E-09 |
| Gpc2 | 1,46606097 | 0,00282807 |
| Gbp8 | 1,50996705 | 3,89E-06 |
| Stambpl1 | 1,55435046 | 0,00029838 |
| Inhba | 1,55642354 | 5,52E-05 |
| Zbp1 | 1,56026149 | 1,91E-06 |
| Abcd2 | 1,66513024 | 6,02E-07 |
| Parp16 | 1,68410567 | 4,85E-08 |
| Aldh1b1 | 1,69026274 | 1,19E-19 |
| Atp6v0d2 | 1,70187248 | 0,00041284 |
| Ifi44 | 1,75007843 | 3,53E-12 |
| Csf2rb | 1,77340069 | 2,34E-08 |
| Fam129b | 1,83033336 | 2,71E-09 |
| Afp | 1,91285528 | 1,99E-09 |
| Ubd | 1,91328307 | 7,73E-07 |
| Ly6a | 2,01931764 | 6,60E-10 |
| Zfp763 | 2,08571082 | 6,50E-14 |
| Rad51b | 2,09694892 | 1,20E-08 |
| Rab27a | 2,12253967 | 3,69E-11 |
| Ky | 2,64538207 | 4,77E-10 |
| Prelid2 | 2,67312861 | 5,44E-13 |
| Stap1 | 2,78458005 | 3,16E-17 |
| Ipcef1 | 2,93291288 | 3,00E-14 |
| Dpy19l3 | 3,24953358 | 3,35E-41 |
| D330045A20R | 3,37492815 | 1,31E-18 |
| Napb | 3,6406772 | 3,45E-34 |
| 2010005H15R | 4,18295311 | 1,92E-52 |
