## Supplementary material for "The mouse HP1 proteins are essential for preventing liver tumorigenesis": suppl table 5

**Supplementary Table 5** : HP1-dependent genes with liver specific functions

| gene name | log2FoldChange | padj | Drug | Lipid | Steroid |
|---|---|---|---|---|---|
| Cyp1a2 | -1,25 | 3,65E-12 | 1 | 1 | 1 |
| Ugt1a1 | -0,70 | 0,0305 | 1 | 0 | 1 |
| Ugt1a9 | -1,61 | 1,69E-07 | 1 | 0 | 1 |
| Ugt2b1 | -1,01 | 7,57E-05 | 1 | 0 | 1 |
| Ugt2b35 | -0,78 | 0,0268 | 1 | 0 | 1 |
| Ugt2b5 | -1,20 | 7,55E-06 | 1 | 0 | 1 |
| Cyp2e1 | -0,94 | 7,81E-05 | 1 | 0 | 1 |
| Hsd17b2 | -0,62 | 0,0040 | 0 | 1 | 1 |
| H2-Ke6 | -0,67 | 0,0025 | 0 | 1 | 1 |
| Srd5a2 | 1,01 | 0,0383 | 0 | 1 | 1 |
| Hsd3b5 | -1,39 | 0,0057 | 0 | 0 | 1 |
| Cyp3a25 | -0,85 | 0,0022 | 0 | 0 | 1 |
| Cyp2b10 | 4,06 | 1,41E-38 | 0 | 0 | 1 |
| Cyp2b13 | 1,80 | 0,0001 | 0 | 0 | 1 |
| Cyp2b9 | 2,68 | 4,51E-10 | 0 | 0 | 1 |
| Cyp2d12 | 1,38 | 0,0005 | 0 | 0 | 1 |
| Cyp2d26 | -0,95 | 2,57E-05 | 0 | 0 | 1 |
| Cyp2d40 | -1,42 | 6,83E-06 | 0 | 0 | 1 |
| Cyp2c29 | -2,60 | 7,65E-25 | 0 | 0 | 1 |
| Cyp2c44 | -2,20 | 2,18E-20 | 0 | 0 | 1 |
| Aox1 | -0,60 | 0,0226 | 1 | 0 | 0 |
| Fmo2 | 0,79 | 0,0431 | 1 | 0 | 0 |
| Fmo3 | 1,18 | 0,0224 | 1 | 0 | 0 |
| Gstm1 | -0,57 | 0,0107 | 1 | 0 | 0 |
| Mgst2 | 1,20 | 0,0275 | 1 | 0 | 0 |
| Maoa | -1,02 | 0,0008 | 1 | 0 | 0 |
| Gsta4 | -1,08 | 3,57E-05 | 1 | 0 | 0 |
| Gsto2 | 1,25 | 0,0197 | 1 | 0 | 0 |
| Gstp1 | -1,11 | 7,57E-05 | 1 | 0 | 0 |
| Gstp2 | -1,80 | 3,84E-09 | 1 | 0 | 0 |
| Il1rn | 1,08 | 0,0024 | 0 | 1 | 0 |
| Slc27a2 | -0,68 | 0,0171 | 0 | 1 | 0 |
| Pla2g12a | -0,98 | 0,0003 | 0 | 1 | 0 |
| Acnat2 | 1,17 | 0,0004 | 0 | 1 | 0 |
| Baat | -0,71 | 0,0236 | 0 | 1 | 0 |
| Echdc2 | -1,14 | 0,0014 | 0 | 1 | 0 |
| Gba2 | -0,85 | 0,0136 | 0 | 1 | 0 |
| Agpat9 | 1,95 | 9,38E-09 | 0 | 1 | 0 |
| Sult1b1 | 0,89 | 0,0190 | 0 | 1 | 0 |
| Abhd2 | 1,64 | 5,76E-09 | 0 | 1 | 0 |
| Acsm2 | -2,05 | 9,66E-10 | 0 | 1 | 0 |
| Far1 | 0,84 | 0,0374 | 0 | 1 | 0 |
| Gde1 | -1,17 | 6,11E-07 | 0 | 1 | 0 |
| Mogat2 | 1,39 | 0,0076 | 0 | 1 | 0 |
| Neu3 | 1,30 | 0,0004 | 0 | 1 | 0 |

| | | | | | |
|---|---|---|---|---|---|
| Isyna1 | -1,01 | 0,0350 | 0 | 1 | 0 |
| Acaa1b | -0,97 | 0,0336 | 0 | 1 | 0 |
| Gpx1 | -0,91 | 0,0326 | 0 | 1 | 0 |
| Pltp | -0,77 | 0,0294 | 0 | 1 | 0 |
| Sorl1 | 1,08 | 0,0440 | 0 | 1 | 0 |
| Chpt1 | -0,91 | 0,0001 | 0 | 1 | 0 |
| Acaca | 0,77 | 0,0272 | 0 | 1 | 0 |
| Acly | 0,82 | 0,0477 | 0 | 1 | 0 |
| Gal3st1 | 1,29 | 0,0043 | 0 | 1 | 0 |
| Gm2a | 1,09 | 2,85E-08 | 0 | 1 | 0 |
| Cyp46a1 | -1,64 | 0,0006 | 0 | 1 | 0 |
| Sptlc2 | 0,55 | 0,0447 | 0 | 1 | 0 |
| Ptdss1 | -0,64 | 0,0037 | 0 | 1 | 0 |
| Sult4a1 | 1,23 | 0,0197 | 0 | 1 | 0 |
| Hrasls | 2,93 | 1,33E-12 | 0 | 1 | 0 |
| Cyp39a1 | -0,88 | 0,0016 | 0 | 1 | 0 |
| Pnpla1 | -1,15 | 0,0018 | 0 | 1 | 0 |
| Elovl3 | -1,80 | 5,39E-15 | 0 | 1 | 0 |
| Fads1 | -0,70 | 0,0002 | 0 | 1 | 0 |
| Fads3 | 1,25 | 0,0006 | 0 | 1 | 0 |
| Lipo1 | 2,19 | 2,56E-17 | 0 | 1 | 0 |
| Pnliprp1 | 1,72 | 0,0003 | 0 | 1 | 0 |
| Tm7sf2 | -0,59 | 0,0398 | 0 | 1 | 0 |
| Ebp | -0,44 | 0,0461 | 0 | 1 | 0 |
| Mid1ip1 | 0,66 | 0,0422 | 0 | 1 | 0 |
