## Supplementary material for "The mouse HP1 proteins are essential for preventing liver tumorigenesis": suppl table 6

**Supplementary Table6**: genes with liver-specific expression according to the Tissue Specific Gene Expression and Regulation software (bioinfo.wilmer.jhu.edu/tiger)

| Symbol | baseMean | log2FoldChange | padj | Name | Chr | Strand |
|---|---|---|---|---|---|---|
| Agxt | 4310,378042 | 0,392971381 | 0,22527304 | alanine-glyoxylate aminotransferase | Chr1 | + |
| Aox1 | 1356,637989 | -0,598215697 | 0,022611041 | aldehyde oxidase 1 | Chr1 | + |
| Apcs | 3990,997066 | -0,044462603 | 0,945242233 | serum amyloid P-component | Chr1 | - |
| Apoa2 | 231294,2226 | -0,380168207 | 0,100885232 | apolipoprotein A-II | Chr1 | + |
| Cfh | 23300,31731 | -0,090317453 | 0,824000947 | complement component factor h | Chr1 | - |
| Col4a4 | 6,344384275 | 0,929660346 | 0,118052347 | collagen, type IV, alpha 4 | Chr1 | - |
| Cox5b | 3359,659769 | -0,22551435 | 0,788139622 | cytochrome c oxidase subunit Vb | Chr1 | + |
| Cps1 | 71826,86218 | -0,062296149 | 0,910948764 | carbamoyl-phosphate synthetase 1 | Chr1 | + |
| Crp | 10280,08718 | -0,168655374 | 0,583107393 | C-reactive protein, pentraxin-related | Chr1 | + |
| F13b | 5882,59283 | 0,326742237 | 0,402107622 | coagulation factor XIII, beta subunit | Chr1 | + |
| Fmo3 | 93,2763624 | 1,176646157 | 0,022395757 | flavin containing monooxygenase 3 | Chr1 | - |
| Il18r1 | 7,889858639 | -0,194846344 | 0,816025345 | interleukin 18 receptor 1 | Chr1 | + |
| Kcnt2 | 51,59229875 | 1,748179568 | 1,49E-05 | potassium channel, subfamily T, member 2 | Chr1 | + |
| Nr1i3 | 2027,588205 | -0,116569165 | 0,735365797 | nuclear receptor subfamily 1, group I, member 3 | Chr1 | + |
| Nr5a2 | 487,4929756 | 0,199761313 | 0,672205411 | nuclear receptor subfamily 5, group A, member 2 | Chr1 | - |
| Serpinc1 | 40791,14302 | 0,007064936 | 0,985797359 | serine (or cysteine) peptidase inhibitor, clade C (antithrombin), member 1 | Chr1 | + |
| Spp2 | 5667,511934 | 0,570410458 | 0,026196265 | secreted phosphoprotein 2 | Chr1 | + |
| Vil1 | 6,468875243 | 1,444840875 | 0,004663891 | villin 1 | Chr1 | + |
| C8g | 11292,22827 | -0,58682036 | 0,019663776 | complement component 8, gamma polypeptide | chr2 | - |
| F2 | 29920,1693 | -0,137097359 | 0,66780976 | coagulation factor II | chr2 | - |
| Gatm | 192,9860689 | 0,12978007 | 0,756319626 | glycine amidinotransferase (L-arginine:glycine amidinotransferase) | chr2 | - |
| Hao1 | 8744,825805 | -0,245613946 | 0,523898331 | hydroxyacid oxidase 1, liver | chr2 | - |
| Hnf4a | 7444,341009 | -0,408240078 | 0,109872624 | hepatic nuclear factor 4, alpha | chr2 | + |
| Il1b | 93,15456913 | 0,516940787 | 0,407602327 | interleukin 1 beta | chr2 | - |
| Itih2 | 21023,81469 | 0,23984246 | 0,397488617 | inter-alpha trypsin inhibitor, heavy chain 2 | chr2 | - |
| Lbp | 1708,60158 | -0,442706711 | 0,405102408 | lipopolysaccharide binding protein | chr2 | + |
| Aadac | 24155,27232 | -0,152110238 | 0,74146296 | arylacetamide deacetylase | chr3 | + |
| Adh4 | 2997,209697 | 0,241675563 | 0,589361744 | alcohol dehydrogenase 4 (class II), pi polypeptide | chr3 | + |
| Cryz | 2156,544675 | 0,293106753 | 0,361776578 | crystallin, zeta | chr3 | + |
| Cth | 8197,786062 | 0,176978166 | 0,672205411 | cystathionase (cystathionine gamma-lyase) | chr3 | - |
| Fga | 76667,08936 | -0,354458816 | 0,380154872 | fibrinogen alpha chain | chr3 | + |
| Fgb | 125579,2312 | -0,263619072 | 0,471563545 | fibrinogen beta chain | chr3 | - |
| Fgg | 89157,86322 | -0,298376732 | 0,348682374 | fibrinogen gamma chain | chr3 | + |
| Fmo5 | 12176,99942 | -0,648571501 | 0,073430016 | flavin containing monooxygenase 5 | chr3 | + |
| Hao2 | 31,27098371 | 1,980919168 | 7,57E-06 | hydroxyacid oxidase 2 | chr3 | - |
| Hfe2 | 865,3305975 | 0,100489653 | 0,793250646 | hemochromatosis type 2 (juvenile) | chr3 | + |

| | | | | | | |
|---|---|---|---|---|---|---|
| Hmgcs2 | 43049,43225 | 0,005189625 | 0,993369412 | 3-hydroxy-3-methylglutaryl-Coenzyme A synthase 2 | chr3 | + |
| Hps3 | 150,645189 | 0,497795871 | 0,211590547 | HPS3, biogenesis of lysosomal organelles complex 2 subunit 1 | chr3 | - |
| Mttp | 5907,157878 | 0,102574344 | 0,764714786 | microsomal triglyceride transfer protein | chr3 | - |
| Pklr | 4207,719837 | 0,672681684 | 0,104289738 | pyruvate kinase liver and red blood cell | chr3 | + |
| Rfxap | 102,1839249 | -0,474152974 | 0,223515497 | regulatory factor X-associated protein | chr3 | - |
| Tdo2 | 41327,9658 | -0,136691483 | 0,724652712 | tryptophan 2,3-dioxygenase | chr3 | - |
| Tm4sf4 | 2192,93019 | 0,126419128 | 0,737668661 | transmembrane 4 superfamily member 4 | chr3 | + |
| Aldob | 102792,0541 | -0,194182977 | 0,536431087 | aldolase B, fructose-bisphosphate | chr4 | - |
| Ambp | 43298,87819 | -0,162294728 | 0,637633057 | alpha 1 microglobulin/bikunin | chr4 | - |
| Angptl3 | 17937,07589 | 0,027303576 | 0,969048579 | angiopoietin-like 3 | chr4 | + |
| Baat | 4701,103712 | -0,707896278 | 0,023606121 | bile acid-Coenzyme A: amino acid N-acyltransferase | chr4 | - |
| C8a | 22713,28438 | 0,068432502 | 0,893831725 | complement component 8, alpha polypeptide | chr4 | - |
| C8b | 9190,635536 | -0,459338941 | 0,252617603 | complement component 8, beta polypeptide | chr4 | + |
| Cyp7a1 | 1407,020254 | 0,267646379 | 0,716169774 | cytochrome P450, family 7, subfamily a, polypeptide 1 | chr4 | - |
| Dio1 | 5025,176522 | -0,222136029 | 0,480124139 | deiodinase, iodothyronine, type I | chr4 | - |
| Masp2 | 5503,198073 | 0,025554566 | 0,952400398 | mannan-binding lectin serine peptidase 2 | chr4 | + |
| Nr0b2 | 304,6447185 | -0,247937432 | 0,764363417 | nuclear receptor subfamily 0, group B, member 2 | chr4 | + |
| Orm1 | 27310,59566 | -0,228707356 | 0,596219566 | orosomucoid 1 | chr4 | + |
| Orm2 | 823,6597845 | 0,511339555 | 0,451815929 | orosomucoid 2 | chr4 | + |
| Afm | 5622,890971 | -0,082522961 | 0,834675238 | afamin | chr5 | + |
| Afp | 65,33735904 | 1,912855277 | 1,99E-09 | alpha fetoprotein | chr5 | + |
| Alb | 2284647,37 | -0,108761562 | 0,798875954 | albumin | chr5 | + |
| Azgp1 | 44719,9991 | -0,293065521 | 0,264796388 | alpha-2-glycoprotein 1, zinc | chr5 | + |
| Cxcl2 | 5,715098119 | 0,422228601 | 0,559428875 | chemokine (C-X-C motif) ligand 2 | chr5 | + |
| Gc | 190075,3807 | -0,036711914 | 0,925034844 | vitamin D binding protein | chr5 | - |
| Hpd | 25785,81549 | -0,462573348 | 0,044747609 | 4-hydroxyphenylpyruvic acid dioxygenase | chr5 | - |
| Khk | 8052,930513 | 0,249872845 | 0,618723125 | ketohexokinase | chr5 | + |
| Ppargc1a | 205,3902206 | 0,588567561 | 0,105299254 | peroxisome proliferative activated receptor, gamma, coactivator 1 alpha | chr5 | - |
| Sds | 2664,960019 | -0,328749815 | 0,583782269 | serine dehydratase | chr5 | + |
| Tfr2 | 4387,025083 | -0,412660156 | 0,140524285 | transferrin receptor 2 | chr5 | + |
| A2m | 8,923418562 | -0,571007993 | 0,391907358 | alpha-2-macroglobulin | chr6 | + |
| Akr1d1 | 4271,097059 | 0,604774775 | 0,196626688 | aldo-keto reductase family 1, member D1 | chr6 | + |
| Fabp1 | 185249,4478 | -0,92549447 | 0,018952788 | fatty acid binding protein 1, liver | chr6 | + |
| Gys2 | 6479,112557 | -0,456057938 | 0,220525617 | glycogen synthase 2 | chr6 | - |
| Pon1 | 28512,73209 | -0,06581205 | 0,906333899 | paraoxonase 1 | chr6 | - |
| Pon3 | 1484,579152 | -0,143870921 | 0,704334789 | paraoxonase 3 | chr6 | - |
| Pzp | 105546,05 | -0,33635865 | 0,286763659 | PZP, alpha-2-macroglobulin like | chr6 | - |
| Uroc1 | 5772,267661 | 0,033344167 | 0,935762352 | urocanase domain containing 1 | chr6 | + |
| Abhd2 | 260,2455392 | 1,63604243 | 5,76E-09 | abhydrolase domain containing 2 | chr7 | + |
| Acadsb | 3133,205665 | -0,076607739 | 0,857773796 | acyl-Coenzyme A dehydrogenase, short/branched chain | chr7 | + |
| Acsm2 | 51,19588206 | -2,04591194 | 9,66E-10 | acyl-CoA synthetase medium-chain family member 2 | chr7 | + |
| Apoc1 | 108463,865 | -0,270352797 | 0,354665719 | apolipoprotein C-I | chr7 | - |
| Apoc2 | 9384,567849 | 0,335226982 | 0,293840783 | apolipoprotein C-II | chr7 | - |
| Apoc4 | 20741,83067 | -0,327475328 | 0,40070132 | apolipoprotein C-IV | chr7 | - |
| Cyp2e1 | 96588,97949 | -0,93971779 | 7,81E-05 | cytochrome P450, family 2, subfamily e, polypeptide 1 | chr7 | + |
| Hpn | 3839,008148 | -0,348406233 | 0,355134088 | hepsin | chr7 | - |
| Hpx | 88,05711311 | 0,435968168 | 0,319080738 | hemopexin | chr7 | - |
| Pcsk6 | 908,4042866 | -0,914678166 | 2,45E-05 | proprotein convertase subtilisin/kexin type 6 | chr7 | + |
| Saa1 | 5275,386127 | -0,431581971 | 0,518617521 | serum amyloid A 1 | chr7 | - |
| Saa2 | 2638,095039 | -0,30807436 | 0,648706509 | serum amyloid A 2 | chr7 | + |
| Saa4 | 8073,532904 | -0,8365031 | 0,079609563 | serum amyloid A 4 | chr7 | - |
| Slc27a5 | 19921,76371 | -0,282342419 | 0,29082501 | solute carrier family 27 (fatty acid transporter), member 5 | chr7 | - |
| Agt | 8988,579081 | -0,062314453 | 0,887518458 | angiotensinogen (serpin peptidase inhibitor, clade A, member 8) | chr8 | - |
| F11 | 1392,198202 | -0,543294197 | 0,076519476 | coagulation factor XI | chr8 | - |
| Fgl1 | 12403,97511 | 0,014267934 | 0,986661276 | fibrinogen-like protein 1 | chr8 | - |

| | | | | | | |
|---|---|---|---|---|---|---|
| Hp | 73195,35223 | -0,932674272 | 0,016705702 | haptoglobin | chr8 | - |
| Klkb1 | 3967,806813 | -0,093484388 | 0,820554805 | kallikrein B, plasma 1 | chr8 | - |
| Nat2 | 501,7427077 | 0,246097579 | 0,510690791 | N-acetyltransferase 2 (arylamine N-acetyltransferase) | chr8 | + |
| Nek1 | 122,7568758 | 0,427217929 | 0,331535469 | NIMA (never in mitosis gene a)-related expressed kinase 1 | chr8 | + |
| Papd5 | 604,6891684 | -0,199267073 | 0,572283367 | PAP associated domain containing 5 | chr8 | + |
| Proz | 4288,847332 | 0,200788482 | 0,500489603 | protein Z, vitamin K-dependent plasma glycoprotein | chr8 | + |
| Tat | 30695,35108 | -0,482357753 | 0,314000124 | tyrosine aminotransferase | chr8 | + |
| Apoa1 | 169709,2782 | -0,241130988 | 0,505223498 | apolipoprotein A-I | chr9 | + |
| Apoc3 | 60866,54061 | -0,098033011 | 0,765061508 | apolipoprotein C-III | chr9 | - |
| Aqp9 | 6818,271841 | -0,266546169 | 0,483638076 | aquaporin 9 | chr9 | - |
| Gsta1 | 8,573414703 | -0,620208675 | 0,346086865 | glutathione S-transferase, alpha 1 (Ya) | chr9 | + |
| Gsta2 | 136,6322176 | -0,658903163 | 0,184748288 | glutathione S-transferase, alpha 2 (Yc2) | chr9 | - |
| Lipc | 5380,295338 | 0,164657262 | 0,664082981 | lipase, hepatic | chr9 | - |
| Mst1 | 1989,520007 | -0,243837453 | 0,455499332 | macrophage stimulating 1 (hepatocyte growth factor-like) | chr9 | + |
| Nedd4 | 3034,891081 | 0,209813672 | 0,420273711 | neural precursor cell expressed, developmentally down-regulated 4 | chr9 | + |
| Nnmt | 4065,615431 | 0,089860928 | 0,895663283 | nicotinamide N-methyltransferase | chr9 | - |
| Oaf | 4419,3025 | -0,097586613 | 0,823949825 | out at first homolog | chr9 | - |
| Aldh8a1 | 6889,314539 | -0,029153858 | 0,947819654 | aldehyde dehydrogenase 8 family, member A1 | chr10 | + |
| Arg1 | 31191,94147 | -0,097865266 | 0,828120899 | arginase, liver | chr10 | - |
| Chst3 | 5,150348426 | 0,317931317 | 0,676716141 | carbohydrate (chondroitin 6/keratan) sulfotransferase 3 | chr10 | - |
| Gamt | 1499,786961 | 0,108741549 | 0,740596839 | guanidinoacetate methyltransferase | chr10 | - |
| Hal | 4285,832341 | 0,501409941 | 0,172087384 | histidine ammonia lyase | chr10 | + |
| Hsd17b6 | 4894,507674 | -1,034070262 | 0,067696785 | hydroxysteroid (17-beta) dehydrogenase 6 | chr10 | - |
| Inhbc | 1033,532768 | -0,670878007 | 0,135447688 | inhibin beta-C | chr10 | - |
| Nr1h4 | 2715,346051 | -0,055101702 | 0,91260409 | nuclear receptor subfamily 1, group H, member 4 | chr10 | - |
| Oit3 | 563,6602545 | 0,103806888 | 0,78044956 | oncoprotein induced transcript 3 | chr10 | - |
| Pah | 24058,84111 | -0,662581935 | 0,027397756 | phenylalanine hydroxylase | chr10 | + |
| Rdh16 | 2307,601244 | 0,05060823 | 0,935309554 | retinol dehydrogenase 16 | chr10 | + |
| Upb1 | 5955,965933 | -0,193461428 | 0,462967038 | ureidopropionase, beta | chr10 | + |
| Abca9 | 76,86052742 | 0,640769318 | 0,078316133 | ATP-binding cassette, sub-family A (ABC1), member 9 | chr11 | - |
| Apoh | 59548,0447 | 0,110415095 | 0,749603956 | apolipoprotein H | chr11 | + |
| Asgr1 | 11694,21403 | -0,247943899 | 0,35583041 | asialoglycoprotein receptor 1 | chr11 | + |
| Asgr2 | 3893,571955 | 0,127713756 | 0,666980604 | asialoglycoprotein receptor 2 | chr11 | + |
| Cd7 | 9,23267707 | 0,475507437 | 0,483930201 | CD7 antigen | chr11 | - |
| Fzd2 | 2,403858033 | -0,297109929 | NA | frizzled class receptor 2 | chr11 | + |
| G6pc | 9541,425668 | -0,423741075 | 0,553397688 | glucose-6-phosphatase, catalytic | chr11 | + |
| Krt10 | 50,75775318 | 0,01821677 | 0,980186108 | keratin 10 | chr11 | - |
| Nbr1 | 3877,197001 | 0,267764401 | 0,357345082 | neighbor of Brca1 gene 1 | chr11 | + |
| Pipox | 220,182909 | 0,305212899 | 0,470066395 | pipecolic acid oxidase | chr11 | - |
| Slc13a5 | 237,8092019 | 2,595002051 | 2,78E-12 | solute carrier family 13 (sodium-dependent citrate transporter), member 5 | chr11 | - |
| Slc4a1 | 2,508403914 | -0,282130188 | NA | solute carrier family 4 (anion exchanger), member 1 | chr11 | - |
| Vtn | 36687,12206 | -0,341739112 | 0,240427136 | vitronectin | chr11 | + |
| Apob | 97754,23377 | -0,421173809 | 0,118153597 | apolipoprotein B | chr12 | + |
| Serpina10 | 9392,877881 | -0,102745142 | 0,872774688 | serine (or cysteine) peptidase inhibitor, clade A (alpha-1 antiproteinase, antitrypsin), member 10 | chr12 | - |
| Serpina5 | 6,115707885 | -0,062884672 | 0,948314012 | serine (or cysteine) peptidase inhibitor, clade A, member 5 | chr12 | + |
| Serpina6 | 3116,735822 | -1,610950643 | 4,34E-11 | serine (or cysteine) peptidase inhibitor, clade A, member 6 | chr12 | - |
| Slc10a1 | 16112,51968 | -0,478467278 | 0,130822929 | solute carrier family 10 (sodium/bile acid cotransporter family), member 1 | chr12 | - |
| Bhmt | 57162,32616 | 0,908896108 | 0,024715096 | betaine-homocysteine methyltransferase | chr13 | - |
| Bhmt2 | 3956,540358 | 0,314822819 | 0,374761697 | betaine-homocysteine methyltransferase 2 | chr13 | - |
| Dmgdh | 7158,445326 | 0,179151563 | 0,58497677 | dimethylglycine dehydrogenase precursor | chr13 | + |
| F12 | 6723,444404 | -0,105095752 | 0,842925053 | coagulation factor XII (Hageman factor) | chr13 | - |

| | | | | | | |
|---|---|---|---|---|---|---|
| Lect2 | 4141,143682 | 0,332468931 | 0,383568565 | leukocyte cell-derived chemotaxin 2 | chr13 | - |
| Polr3g | 176,353346 | -0,510885727 | 0,140027516 | polymerase (RNA) III (DNA directed) polypeptide G | chr13 | - |
| Adra1a | 38,65281178 | -0,735534129 | 0,124458443 | adrenergic receptor, alpha 1a | chr14 | + |
| Ang | 8071,096287 | -0,651412501 | 0,098348401 | angiogenin, ribonuclease, RNase A family, 5 | chr14 | + |
| Cpb2 | 13986,29625 | -0,054213112 | 0,895594598 | carboxypeptidase B2 (plasma) | chr14 | + |
| Dnase1l3 | 906,207303 | -0,08267028 | 0,855769633 | deoxyribonuclease 1-like 3 | chr14 | - |
| Itih1 | 11463,2087 | -0,149857675 | 0,576265028 | inter-alpha trypsin inhibitor, heavy chain 1 | chr14 | - |
| Itih3 | 14760,99308 | 0,09335647 | 0,806012703 | inter-alpha trypsin inhibitor, heavy chain 3 | chr14 | - |
| Itih4 | 32690,31757 | 0,011401544 | 0,985849808 | inter alpha-trypsin inhibitor, heavy chain 4 | chr14 | + |
| Mat1a | 66856,712 | 0,466663554 | 0,13621142 | methionine adenosyltransferase I, alpha | chr14 | + |
| Rnase4 | 29721,33639 | -0,534577977 | 0,156728357 | ribonuclease, RNase A family 4 | chr14 | + |
| A1bg | 9,627075438 | 1,045058431 | 0,068097992 | alpha-1-B glycoprotein | chr15 | - |
| Agxt2 | 3241,865538 | 0,381244535 | 0,247584138 | alanine-glyoxylate aminotransferase 2 | chr15 | + |
| C6 | 3881,216575 | -0,569991365 | 0,235271103 | complement component 6 | chr15 | + |
| C9 | 15178,64109 | -1,359728005 | 1,29E-07 | complement component 9 | chr15 | + |
| Colec10 | 794,3796003 | 0,699177636 | 0,030454862 | collectin sub-family member 10 | chr15 | + |
| Dpys | 3110,369938 | -0,191239142 | 0,511697151 | dihydropyrimidinase | chr15 | - |
| Slc38a4 | 27061,20241 | -0,230258316 | 0,554660735 | solute carrier family 38, member 4 | chr15 | - |
| Tmprss6 | 6213,299598 | -0,117754804 | 0,783264136 | transmembrane serine protease 6 | chr15 | - |
| Abat | 3917,647613 | -0,133276359 | 0,756319626 | 4-aminobutyrate aminotransferase | chr16 | + |
| Ahsg | 245204,9226 | -0,014729266 | 0,974008692 | alpha-2-HS-glycoprotein | chr16 | + |
| Ehhadh | 6169,399133 | 0,409485766 | 0,437265048 | enoyl-Coenzyme A, hydratase/3-hydroxyacyl Coenzyme A dehydrogenase | chr16 | - |
| Fetub | 7196,39517 | -0,655889616 | 0,005350521 | fetuin beta | chr16 | + |
| Hgd | 13285,10159 | -0,552052108 | 0,030459905 | homogentisate 1, 2-dioxygenase | chr16 | + |
| Hrg | 11300,64188 | -0,370853027 | 0,15139513 | histidine-rich glycoprotein | chr16 | + |
| Kng1 | 50936,43207 | -0,265421364 | 0,386806294 | kininogen 1 | chr16 | + |
| Nr1i2 | 253,4181984 | -0,078444905 | 0,871983898 | nuclear receptor subfamily 1, group I, member 2 | chr16 | - |
| Serpind1 | 17281,94216 | 0,150421538 | 0,620918751 | serine (or cysteine) peptidase inhibitor, clade D, member 1 | chr16 | + |
| Apom | 7728,072982 | -0,086973437 | 0,814733578 | apolipoprotein M | chr17 | - |
| C3 | 83400,38506 | -0,196880022 | 0,583782269 | complement component 3 | chr17 | - |
| C4a | 427,7813857 | 1,404880496 | 0,000251317 | complement component 4A (Rodgers blood group) | chr17 | - |
| C4b | 11012,71832 | -0,331831172 | 0,289430713 | complement component 4B (Chido blood group) | chr17 | - |
| Gnmt | 19700,26138 | -0,255958654 | 0,653708291 | glycine N-methyltransferase | chr17 | - |
| Plg | 88,16462024 | 0,038489654 | 0,953751362 | plasminogen | chr17 | + |
| Slc22a1 | 4772,954103 | -0,441167258 | 0,106945732 | solute carrier family 22 (organic cation transporter), member 1 | chr17 | - |
| Slc22a3 | 66,93788498 | 0,179643654 | 0,772525785 | solute carrier family 22 (organic cation transporter), member 3 | chr17 | - |
| Slc22a7 | 961,9869562 | 0,719580837 | 0,030188319 | solute carrier family 22 (organic anion transporter), member 7 | chr17 | - |
| Proc | 9219,711203 | -0,385697994 | 0,204328779 | protein C | chr18 | - |
| Abcc2 | 3627,408112 | -0,017622833 | 0,975528401 | ATP-binding cassette, sub-family C (CFTR/MRP), member 2 | chr19 | + |
| Cpn1 | 3382,071658 | -0,086906448 | 0,859745168 | carboxypeptidase N, polypeptide 1 | chr19 | - |
| Frat2 | 39,57534469 | -0,464004946 | 0,421362686 | frequently rearranged in advanced T cell lymphomas 2 | chr19 | - |
| Glyat | 8632,21562 | -0,370401566 | 0,290819569 | glycine-N-acyltransferase | chr19 | + |
| Habp2 | 3239,5837 | -0,066185813 | 0,852923957 | hyaluronic acid binding protein 2 | chr19 | + |
| Pnliprp1 | 10,78775594 | 1,724825204 | 0,000314559 | pancreatic lipase related protein 1 | chr19 | + |
| Rbp4 | 40817,19012 | -1,304125645 | 1,91E-07 | retinol binding protein 4, plasma | chr19 | - |
| Alas2 | 2078,499095 | -0,805951322 | 0,016167446 | aminolevulinic acid synthase 2, erythroid | chrX | + |
| Dusp9 | 3,124626847 | 0,78378503 | 0,186292552 | dual specificity phosphatase 9 | chrX | + |
| F9 | 6895,991739 | 0,257839877 | 0,455467885 | coagulation factor IX | chrX | + |
| Mtm1 | 241,8177707 | 0,001360219 | 0,997729735 | X-linked myotubular myopathy gene 1 | chrX | + |
| Serpina7 | 615,936052 | -0,655103653 | 0,19521384 | serine (or cysteine) peptidase inhibitor, clade A (alpha-1 antiproteinase, antitrypsin), member 7 | chrX | - |
| Zxdb | 28,36580393 | -0,089428658 | 0,899463828 | zinc finger, X-linked, duplicated B | chrX | + |
