## Supplementary material for "The mouse HP1 proteins are essential for preventing liver tumorigenesis": suppl table 7

**Supplementary Table 7** : HP1-dependent repeats

| repeat | baseMean | log2FoldChange | padj | chromosome | start | end | family |
|---|---|---|---|---|---|---|---|
| RMER6A | 3,95 | -5,84 | 0,000870422 | chr10 | 63059617 | 63060330 | LTR/ERVK |
| RMER15 | 32,98 | -5,45 | 8,51E-11 | chr17 | 35476452 | 35476654 | LTR/ERVL |
| B1_Mus2 | 8,18 | -5,42 | 0,000782598 | chr19 | 5833516 | 5833638 | SINE/Alu |
| L1_Mm | 7,30 | -5,24 | 0,000539178 | chr13 | 4261557 | 4262228 | LINE/L1 |
| RMER17A2 | 2,40 | -5,12 | 0,014572378 | chr19 | 39328136 | 39328891 | LTR/ERVK |
| RLTR4_MM-int | 8,09 | -4,77 | 0,000792901 | chr15 | 76560514 | 76564037 | LTR/ERV1 |
| MER58B | 3,23 | -4,75 | 0,011449629 | chr18 | 9880852 | 9880950 | DNA/hAT-Charlie |
| MTC-int | 3,18 | -4,73 | 0,03510424 | chr11 | 16619510 | 16620370 | LTR/ERVL-MaLR |
| L1Md_A | 5,11 | -4,71 | 0,017744421 | chr12 | 104350610 | 104351590 | LINE/L1 |
| A-rich | 2,89 | -4,60 | 0,019042444 | chr12 | 37453617 | 37453668 | Low_complexity |
| RMER4B | 4,89 | -4,60 | 0,011037805 | chr16 | 37891814 | 37892263 | LTR/ERVK |
| L1MC3 | 2,78 | -4,53 | 0,02699166 | chr18 | 9881634 | 9881779 | LINE/L1 |
| RLTR9F | 2,79 | -4,53 | 0,031528522 | chr19 | 44025902 | 44026336 | LTR/ERVK |
| RLTRETN_Mm | 2,70 | -4,49 | 0,027917006 | chr18 | 9882147 | 9882472 | LTR/ERVK |
| MERVL_2A-int | 2,68 | -4,48 | 0,034133199 | chr19 | 8115363 | 8115659 | LTR/ERVL |
| MTC | 2,56 | -4,42 | 0,037509591 | chr13 | 98321981 | 98322265 | LTR/ERVL-MaLR |
| ORR1E | 4,29 | -4,41 | 0,010224369 | chr9 | 113128157 | 113128347 | LTR/ERVL-MaLR |
| B3A | 5,72 | -4,31 | 0,009878799 | chr6 | 121886838 | 121886967 | SINE/B2 |
| B3 | 2,40 | -4,31 | 0,046132983 | chr17 | 33774515 | 33774701 | SINE/B2 |
| L1MC3 | 5,52 | -4,26 | 0,003214307 | chr18 | 9881802 | 9881967 | LINE/L1 |
| MT2B1 | 3,52 | -4,14 | 0,032426789 | chr19 | 8115719 | 8116164 | LTR/ERVL |
| ORR1B2 | 6,81 | -4,12 | 0,009848954 | chr14 | 41140091 | 41140428 | LTR/ERVL-MaLR |
| B1_Mus1 | 9,77 | -3,85 | 0,001118207 | chr17 | 35475761 | 35475907 | SINE/Alu |
| L1MD | 4,29 | -3,79 | 0,02340491 | chr19 | 4043577 | 4043701 | LINE/L1 |
| Lx6 | 8,52 | -3,77 | 0,000907546 | chr13 | 4262799 | 4263199 | LINE/L1 |
| RLTR6_Mm | 7,04 | -3,68 | 0,004008783 | chr19 | 4045069 | 4045631 | LTR/ERV1 |
| RMER15 | 6,41 | -3,66 | 0,004322286 | chr10 | 63046386 | 63046480 | LTR/ERVL |
| PB1D9 | 6,33 | -3,63 | 0,032814054 | chr19 | 5834279 | 5834373 | SINE/Alu |
| Lx | 36,05 | -3,60 | 8,66E-11 | chr7 | 14396898 | 14398290 | LINE/L1 |
| PB1D10 | 14,55 | -3,58 | 0,000283123 | chr10 | 127988841 | 127988950 | SINE/Alu |
| MTA_Mm | 3,83 | -3,57 | 0,037431762 | chr4 | 86667135 | 86667529 | LTR/ERVL-MaLR |
| B3 | 15,35 | -3,45 | 1,49E-05 | chr17 | 35475318 | 35475504 | SINE/B2 |
| Lx | 13,20 | -3,35 | 1,30E-05 | chr7 | 119623280 | 119626761 | LINE/L1 |
| L1_Mus3 | 32,05 | -3,34 | 9,71E-09 | chr12 | 104348209 | 104350609 | LINE/L1 |
| B3 | 4,34 | -3,28 | 0,043256406 | chr11 | 106392311 | 106392493 | SINE/B2 |
| L1MA8 | 10,48 | -3,26 | 0,001174919 | chr5 | 87016554 | 87016950 | LINE/L1 |
| RLTR9D | 104,54 | -3,24 | 1,23E-15 | chr5 | 146113489 | 146113889 | LTR/ERVK |
| RLTR4_MM-int | 67,39 | -3,22 | 0,021990225 | chr11 | 6010614 | 6010759 | LTR/ERV1 |
| U8 | 5,98 | -3,19 | 0,038321408 | chr12 | 13163628 | 13163764 | snRNA |
| Lx9 | 11,81 | -3,18 | 0,003274204 | chr10 | 127989451 | 127989847 | LINE/L1 |
| L1Md_F2 | 8,63 | -3,12 | 0,002842823 | chr11 | 16630237 | 16632083 | LINE/L1 |
| (TGG)n | 7,01 | -3,11 | 0,007598171 | chr5 | 146095744 | 146095872 | Simple_repeat |
| MER5B | 7,34 | -3,09 | 0,009332059 | chr2 | 126592695 | 126592830 | DNA/hAT-Charlie |
| Lx8 | 19,29 | -3,07 | 7,86E-06 | chr5 | 146115586 | 146115974 | LINE/L1 |
| Lx | 9,65 | -3,03 | 0,009298495 | chr8 | 10858601 | 10859105 | LINE/L1 |
| RSINE1 | 146,79 | -2,96 | 7,46E-08 | chr15 | 4737073 | 4737153 | SINE/B4 |
| RLTR4_MM-int | 315,08 | -2,96 | 0,036213874 | chr11 | 6008754 | 6010616 | LTR/ERV1 |
| B2_Mm2 | 6,31 | -2,96 | 0,029362011 | chr6 | 83128973 | 83129163 | SINE/B2 |
| SSU-rRNA_Hsa | 12,44 | -2,96 | 0,01083526 | chr2 | 27121169 | 27121482 | rRNA |
| MTE2a | 18,77 | -2,95 | 9,44E-07 | chr5 | 146113992 | 146114353 | LTR/ERVL-MaLR |
| RMER17B | 10,85 | -2,93 | 0,00433108 | chr13 | 4257362 | 4258219 | LTR/ERVK |
| MMVL30-int | 65,45 | -2,90 | 2,69E-14 | chr17 | 35480336 | 35482627 | LTR/ERV1 |
| B1_Mur4 | 4,98 | -2,89 | 0,027616336 | chr4 | 104806754 | 104806881 | SINE/Alu |
| Lx | 8,96 | -2,87 | 0,020301221 | chr7 | 14398606 | 14399155 | LINE/L1 |
| Lx8 | 4,78 | -2,82 | 0,048787686 | chr15 | 85813573 | 85814032 | LINE/L1 |
| AT_rich | 10,01 | -2,82 | 0,012087126 | chr15 | 4737195 | 4737230 | |
| RLTR26C_MM | 5,46 | -2,79 | 0,028003169 | chr6 | 141691023 | 141691683 | LTR/ERVK |
| RMER6D | 9,23 | -2,78 | 0,006981237 | chr5 | 146094772 | 146095083 | LTR/ERVK |
| SSU-rRNA_Hsa | 9,11 | -2,77 | 0,047663178 | chr4 | 78773675 | 78774216 | rRNA |
| B1F | 5,90 | -2,75 | 0,022063905 | chr2 | 126593184 | 126593290 | SINE/Alu |

| | | | | | | | |
|---|---|---|---|---|---|---|---|
| RMER19C | 9,42 | -2,72 | 0,007369475 | chr11 | 120508417 | 120508922 | LTR/ERVK |
| MuRRS-int | 17,80 | -2,70 | 0,000588644 | chr2 | 126608834 | 126613401 | LTR/ERV1 |
| B3 | 6,69 | -2,70 | 0,01335349 | chr16 | 45955199 | 45955407 | SINE/B2 |
| MTA_Mm | 24,05 | -2,67 | 7,44E-05 | chr6 | 141901247 | 141901641 | LTR/ERVL-MaLR |
| B1_Mur3 | 6,05 | -2,67 | 0,044746919 | chr2 | 173142965 | 173143110 | SINE/Alu |
| MLT1A1 | 10,07 | -2,64 | 0,008811982 | chr6 | 149168300 | 149168365 | LTR/ERVL-MaLR |
| L1MC1 | 9,81 | -2,64 | 0,002126336 | chr3 | 119112297 | 119113288 | LINE/L1 |
| MLT1N2 | 28,02 | -2,63 | 6,57E-06 | chr17 | 85067642 | 85067821 | LTR/ERVL-MaLR |
| LSU-rRNA_Hsa | 1815,76 | -2,63 | 0,02547141 | chr8 | 15519775 | 15520124 | rRNA |
| L1MC1 | 10,90 | -2,62 | 0,00192959 | chr8 | 10907785 | 10907940 | LINE/L1 |
| MTC | 24,10 | -2,62 | 0,005739573 | chr10 | 127988260 | 127988652 | LTR/ERVL-MaLR |
| MTC | 6,89 | -2,62 | 0,013450959 | chr7 | 79446185 | 79446592 | LTR/ERVL-MaLR |
| L1_Mus3 | 286,71 | -2,61 | 3,35E-13 | chr6 | 141900334 | 141901246 | LINE/L1 |
| ORR1B1 | 16,69 | -2,59 | 0,006016625 | chr5 | 146104997 | 146105405 | LTR/ERVL-MaLR |
| Lx5c | 44,00 | -2,53 | 6,51E-06 | chr2 | 126591728 | 126592067 | LINE/L1 |
| B2_Mm1a | 12,46 | -2,53 | 0,02465358 | chr11 | 116169755 | 116169946 | SINE/B2 |
| B2_Mm2 | 23,56 | -2,49 | 0,000479299 | chr2 | 126595757 | 126595960 | SINE/B2 |
| RLTR20B5_MM | 5,88 | -2,49 | 0,035802445 | chr19 | 8114881 | 8115330 | LTR/ERVK |
| RLTR11A | 14,40 | -2,47 | 0,005113281 | chr15 | 75958424 | 75958932 | LTR/ERVK |
| LSU-rRNA_Hsa | 1937,30 | -2,47 | 0,031183865 | chr3 | 62159052 | 62159490 | rRNA |
| RMER6D | 22,55 | -2,47 | 0,001601407 | chr5 | 146095903 | 146096225 | LTR/ERVK |
| RSINE1 | 89,11 | -2,45 | 0,00013955 | chr15 | 4735618 | 4735774 | SINE/B4 |
| L1Md_F2 | 13,12 | -2,44 | 0,004624864 | chr6 | 149169991 | 149170483 | LINE/L1 |
| B2_Mm2 | 14,03 | -2,37 | 0,00586091 | chr10 | 70045222 | 70045332 | SINE/B2 |
| L1_Mur2 | 8,55 | -2,37 | 0,039995921 | chr11 | 16579370 | 16580426 | LINE/L1 |
| LSU-rRNA_Hsa | 2786,32 | -2,34 | 0,049775675 | chr9 | 3258790 | 3259525 | rRNA |
| RMER6C | 11,45 | -2,31 | 0,010762503 | chr19 | 44009230 | 44009756 | LTR/ERVK |
| MIR | 17,58 | -2,30 | 0,024427028 | chr4 | 60661330 | 60661403 | SINE/MIR |
| Lx8 | 16,10 | -2,30 | 0,005524551 | chr1 | 67105221 | 67105453 | LINE/L1 |
| Lx | 24,38 | -2,29 | 0,00029491 | chr7 | 119627192 | 119628983 | LINE/L1 |
| B1_Mus2 | 10,88 | -2,27 | 0,004368988 | chr5 | 123180304 | 123180449 | SINE/Alu |
| B4A | 9,15 | -2,27 | 0,015445104 | chr5 | 146115003 | 146115160 | SINE/B4 |
| RSINE1 | 7,29 | -2,27 | 0,049766997 | chr11 | 106198007 | 106198144 | SINE/B4 |
| (CAGAGA)n | 8,29 | -2,26 | 0,038163494 | chr13 | 91787708 | 91787761 | Simple_repeat |
| Lx9 | 10,18 | -2,26 | 0,009505502 | chr5 | 87015721 | 87016206 | LINE/L1 |
| L1_Mus3 | 788,28 | -2,25 | 6,74E-10 | chr12 | 104347895 | 104348154 | LINE/L1 |
| RLTR6_Mm | 143,58 | -2,25 | 8,11E-08 | chr2 | 75642372 | 75642945 | LTR/ERV1 |
| B1_Mus1 | 15,63 | -2,23 | 0,002132021 | chr18 | 46644868 | 46645015 | SINE/Alu |
| MIRb | 12,42 | -2,23 | 0,006977662 | chr1 | 67103226 | 67103351 | SINE/MIR |
| LSU-rRNA_Hsa | 17,61 | -2,21 | 0,019614648 | chr2 | 73867872 | 73868032 | rRNA |
| RLTR14 | 18,76 | -2,17 | 0,007639972 | chr18 | 46644791 | 46644867 | LTR/ERV1 |
| L1M5 | 11,64 | -2,15 | 0,027524063 | chr17 | 86703761 | 86704226 | LINE/L1 |
| L1MB5 | 15,90 | -2,12 | 0,022233061 | chr2 | 173136885 | 173137279 | LINE/L1 |
| L1MC1 | 16,51 | -2,11 | 0,002898163 | chr8 | 10907475 | 10907725 | LINE/L1 |
| B3A | 8,49 | -2,11 | 0,042203566 | chr10 | 89450977 | 89451076 | SINE/B2 |
| RSINE1 | 17,20 | -2,10 | 0,004491496 | chr2 | 173138083 | 173138218 | SINE/B4 |
| B2_Mm2 | 14,41 | -2,10 | 0,00950926 | chr8 | 109997268 | 109997413 | SINE/B2 |
| Tigger7 | 56,18 | -2,06 | 0,000252372 | chr18 | 46643766 | 46644017 | DNA/TcMar-Tigger |
| L2a | 18,48 | -2,06 | 0,001678237 | chr2 | 58783484 | 58783602 | LINE/L2 |
| MTD | 12,03 | -2,06 | 0,024102733 | chr13 | 34985723 | 34985913 | LTR/ERVL-MaLR |
| Lx3_Mus | 9,99 | -2,05 | 0,040404753 | chr1 | 140072071 | 140073834 | LINE/L1 |
| B2_Mm1a | 11,59 | -2,05 | 0,01692344 | chr12 | 80930730 | 80930917 | SINE/B2 |
| L1_Mus3 | 537,00 | -2,04 | 4,09E-08 | chr6 | 141901642 | 141903099 | LINE/L1 |
| RSINE1 | 19,02 | -2,04 | 0,002523212 | chr11 | 101375514 | 101375634 | SINE/B4 |
| MTD | 27,27 | -2,01 | 3,44E-05 | chr6 | 141946773 | 141947184 | LTR/ERVL-MaLR |
| Lx7 | 25,28 | -2,00 | 0,000500259 | chr5 | 87013798 | 87013877 | LINE/L1 |
| L1_Rod | 10,03 | -2,00 | 0,024422016 | chr1 | 139687811 | 139688258 | LINE/L1 |
| MTB_Mm | 11,74 | -1,99 | 0,046688072 | chr15 | 85740724 | 85741015 | LTR/ERVL-MaLR |
| SSU-rRNA_Hsa | 35,94 | -1,97 | 0,007369789 | chr10 | 82955117 | 82955391 | rRNA |
| B3A | 28,92 | -1,96 | 0,000335282 | chr15 | 3368091 | 3368165 | SINE/B2 |
| PB1D10 | 53,60 | -1,96 | 0,000337999 | chr6 | 149169113 | 149169218 | SINE/Alu |
| L1MdV_III | 30,48 | -1,95 | 0,000553846 | chr14 | 8057565 | 8059019 | LINE/L1 |
| RLTR14 | 70,17 | -1,95 | 3,44E-06 | chr18 | 46645016 | 46645392 | LTR/ERV1 |
| B2_Mm2 | 9,97 | -1,93 | 0,04590488 | chr18 | 27666719 | 27666894 | SINE/B2 |
| B3 | 70,06 | -1,93 | 0,003214307 | chr6 | 149168585 | 149168762 | SINE/B2 |
| LSU-rRNA_Hsa | 6361,51 | -1,90 | 0,000351014 | chr18 | 68692063 | 68692316 | rRNA |
| ORR1A2 | 125,17 | -1,89 | 8,37E-08 | chr15 | 82684706 | 82685021 | LTR/ERVL-MaLR |

| | | | | | | | |
|---|---|---|---|---|---|---|---|
| MTE2a | 40,33 | -1,88 | 0,000290755 | chr5 | 145976128 | 145976483 | LTR/ERVL-MaLR |
| Lx | 23,63 | -1,88 | 0,00204072 | chr14 | 8060976 | 8061560 | LINE/L1 |
| ORR1A2 | 13,75 | -1,88 | 0,023774988 | chr1 | 36418854 | 36419172 | LTR/ERVL-MaLR |
| ORR1A4 | 33,07 | -1,87 | 0,003141696 | chr13 | 24321717 | 24322040 | LTR/ERVL-MaLR |
| Lx7 | 11,41 | -1,85 | 0,037777968 | chr5 | 87013946 | 87014092 | LINE/L1 |
| B1_Mur4 | 37,27 | -1,85 | 0,009918549 | chr15 | 85737943 | 85738081 | SINE/Alu |
| RSINE1 | 21,75 | -1,85 | 0,004495357 | chr2 | 120602816 | 120602968 | SINE/B4 |
| B3 | 39,75 | -1,83 | 0,001671709 | chr3 | 130636854 | 130637007 | SINE/B2 |
| (A)n | 10,60 | -1,80 | 0,044183233 | chr11 | 96807371 | 96807396 | Simple_repeat |
| L1Md_F2 | 13,23 | -1,80 | 0,033857201 | chr6 | 122143305 | 122144303 | LINE/L1 |
| MTC | 24,97 | -1,79 | 0,043563098 | chr5 | 146000593 | 146000995 | LTR/ERVL-MaLR |
| B2_Mm2 | 11,47 | -1,79 | 0,044769696 | chr2 | 32413560 | 32413742 | SINE/B2 |
| PB1D7 | 50,83 | -1,79 | 4,05E-05 | chr19 | 4036195 | 4036289 | SINE/Alu |
| (GAAA)n | 22,75 | -1,78 | 0,011120116 | chr6 | 149168924 | 149169100 | Simple_repeat |
| Lx10 | 12,66 | -1,78 | 0,039193067 | chr10 | 128975821 | 128977028 | LINE/L1 |
| ID_B1 | 17,68 | -1,77 | 0,005453618 | chr4 | 150869049 | 150869252 | SINE/B4 |
| RSINE1 | 24,69 | -1,77 | 0,002049957 | chr1 | 67103066 | 67103189 | SINE/B4 |
| L1MC1 | 30,58 | -1,77 | 0,003075235 | chr8 | 10906826 | 10907328 | LINE/L1 |
| ID_B1 | 15,33 | -1,76 | 0,01391947 | chr2 | 120602610 | 120602814 | SINE/B4 |
| B3A | 12,68 | -1,76 | 0,018644692 | chr19 | 29019939 | 29020120 | SINE/B2 |
| LSU-rRNA_Hsa | 19,49 | -1,75 | 0,04335415 | chr8 | 85111864 | 85112078 | rRNA |
| L3 | 28,77 | -1,74 | 0,000922451 | chr12 | 72604450 | 72604565 | LINE/CR1 |
| B3 | 14,82 | -1,74 | 0,027524063 | chr10 | 128983421 | 128983590 | SINE/B2 |
| B2_Mm1a | 18,28 | -1,74 | 0,040340103 | chr11 | 116170407 | 116170561 | SINE/B2 |
| B1F1 | 18,14 | -1,72 | 0,029069429 | chr13 | 34980438 | 34980521 | SINE/Alu |
| (CAAAA)n | 13,32 | -1,72 | 0,023026582 | chr16 | 37634158 | 37634208 | Simple_repeat |
| Lx2 | 29,47 | -1,71 | 0,001205823 | chr5 | 87130853 | 87131225 | LINE/L1 |
| B1_Mus1 | 29,24 | -1,71 | 0,007262841 | chr18 | 46642836 | 46642971 | SINE/Alu |
| B3 | 20,85 | -1,71 | 0,002175606 | chr13 | 34988543 | 34988729 | SINE/B2 |
| ID_B1 | 33,39 | -1,69 | 0,012110937 | chr2 | 173147710 | 173147891 | SINE/B4 |
| LTR91 | 44,55 | -1,69 | 0,007139757 | chr16 | 24278714 | 24278873 | LTR |
| RLTR9D | 131,42 | -1,68 | 2,73E-06 | chr5 | 145976592 | 145976992 | LTR/ERVK |
| ORR1B1-int | 103,50 | -1,68 | 0,001341366 | chr9 | 43215534 | 43217140 | LTR/ERVL-MaLR |
| RSINE1 | 29,74 | -1,68 | 0,017689102 | chr6 | 149168428 | 149168567 | SINE/B4 |
| L1_Mus2 | 38,62 | -1,67 | 0,01279622 | chr7 | 119668813 | 119670163 | LINE/L1 |
| B2_Mm1t | 31,06 | -1,67 | 0,000700722 | chr17 | 84657607 | 84657774 | SINE/B2 |
| PB1D10 | 26,98 | -1,65 | 0,008390563 | chr6 | 113437457 | 113437581 | SINE/Alu |
| (GAAAA)n | 26,17 | -1,64 | 0,003085972 | chr16 | 43621902 | 43621955 | Simple_repeat |
| Lx9 | 81,24 | -1,64 | 3,44E-06 | chr11 | 16889083 | 16889577 | LINE/L1 |
| B4A | 235,88 | -1,64 | 4,41E-08 | chr8 | 114154499 | 114154773 | SINE/B4 |
| B2_Mm1a | 12,16 | -1,64 | 0,041534719 | chr2 | 126589437 | 126589644 | SINE/B2 |
| LSU-rRNA_Hsa | 1687,93 | -1,64 | 0,003008679 | chr16 | 35981658 | 35981821 | rRNA |
| L2c | 93,90 | -1,62 | 0,016668832 | chr17 | 23822843 | 23823594 | LINE/L2 |
| (ACATG)n | 23,69 | -1,62 | 0,041985213 | chr17 | 85066874 | 85067030 | Simple_repeat |
| LSU-rRNA_Hsa | 1818,19 | -1,61 | 0,003982147 | chr13 | 97190544 | 97190634 | rRNA |
| B1_Mur1 | 19,82 | -1,60 | 0,04725866 | chr15 | 85738692 | 85738839 | SINE/Alu |
| ID_B1 | 20,54 | -1,58 | 0,008612381 | chr5 | 123180103 | 123180190 | SINE/B4 |
| ORR1A4-int | 62,76 | -1,58 | 0,007572783 | chr13 | 24322074 | 24323015 | LTR/ERVL-MaLR |
| L1_Mus1 | 47,64 | -1,58 | 0,006607086 | chr13 | 24323728 | 24324157 | LINE/L1 |
| B1_Mur4 | 18,57 | -1,57 | 0,014983989 | chr7 | 19139709 | 19139855 | SINE/Alu |
| B2_Mm2 | 21,87 | -1,57 | 0,025825378 | chr17 | 84657408 | 84657522 | SINE/B2 |
| RLTR13D3A1 | 97,28 | -1,56 | 0,00228644 | chr15 | 82635520 | 82635800 | LTR/ERVK |
| Lx6 | 59,13 | -1,54 | 0,004886775 | chr14 | 21183922 | 21185125 | LINE/L1 |
| URR1B | 31,30 | -1,53 | 0,005341207 | chr7 | 27133728 | 27133897 | DNA/hAT-Charlie |
| B3 | 222,07 | -1,52 | 0,000186756 | chr1 | 58201688 | 58201899 | SINE/B2 |
| RLTR33 | 42,12 | -1,51 | 0,00228644 | chr7 | 141022621 | 141023387 | LTR/ERVK |
| (CAAA)n | 48,52 | -1,50 | 0,04667705 | chr8 | 109996462 | 109996488 | Simple_repeat |
| MTB-int | 43,76 | -1,50 | 0,006586862 | chr19 | 34665393 | 34665740 | LTR/ERVL-MaLR |
| L1MdFanc_I | 33,12 | -1,50 | 0,012795994 | chr5 | 87126547 | 87127063 | LINE/L1 |
| B2_Mm1a | 80,75 | -1,48 | 4,83E-06 | chr6 | 39311130 | 39311327 | SINE/B2 |
| Lx | 30,61 | -1,46 | 0,010270177 | chr15 | 9372769 | 9373815 | LINE/L1 |
| MIR | 52,32 | -1,46 | 0,015106641 | chr8 | 105056840 | 105056916 | SINE/MIR |
| ERVB7_2-LTR_MM | 28,33 | -1,45 | 0,016320086 | chr1 | 134433102 | 134433465 | LTR/ERVK |
| LTRIS_Mm | 30,54 | -1,45 | 0,002943696 | chr19 | 43840861 | 43841342 | LTR/ERV1 |
| RLTR13D6 | 16,73 | -1,45 | 0,04098205 | chr10 | 127880862 | 127881663 | LTR/ERVK |
| LSU-rRNA_Hsa | 198,42 | -1,44 | 0,020173713 | chr9 | 105819304 | 105819556 | rRNA |
| LSU-rRNA_Hsa | 50,49 | -1,42 | 0,015824937 | chr1 | 191120925 | 191121174 | rRNA |

| | | | | | | | |
|---|---|---|---|---|---|---|---|
| LSU-rRNA_Hsa | 988,14 | -1,42 | 0,008995228 | chr17 | 70963658 | 70963895 | rRNA |
| ORR1A4 | 110,91 | -1,41 | 0,005103961 | chr18 | 46631011 | 46631345 | LTR/ERVL-MaLR |
| (GAGAA)n | 26,55 | -1,40 | 0,010241928 | chr1 | 105361626 | 105361713 | Simple_repeat |
| L1MdFanc_II | 26,45 | -1,39 | 0,016913015 | chr1 | 172723355 | 172726652 | LINE/L1 |
| B1_Mus2 | 24,49 | -1,38 | 0,01279622 | chr7 | 45471376 | 45471501 | SINE/Alu |
| MIR3 | 39,28 | -1,38 | 0,012087126 | chr5 | 150592382 | 150592484 | SINE/MIR |
| GA-rich | 19,95 | -1,38 | 0,037057511 | chr6 | 10883344 | 10883420 | Low_complexity |
| MIR | 44,68 | -1,36 | 0,001349753 | chr5 | 123179915 | 123180015 | SINE/MIR |
| L1MdF_III | 80,00 | -1,35 | 0,005474719 | chr13 | 4439670 | 4442941 | LINE/L1 |
| MTB | 21,30 | -1,35 | 0,037029069 | chr19 | 34665741 | 34666118 | LTR/ERVL-MaLR |
| RMER10A | 125,67 | -1,35 | 0,001288837 | chr12 | 72605869 | 72606223 | LTR/ERVL |
| RLTR13D3A1 | 604,97 | -1,34 | 0,000117024 | chr15 | 82635948 | 82636628 | LTR/ERVK |
| L4_A_Mam | 52,59 | -1,34 | 0,035024734 | chr10 | 128977936 | 128978112 | LINE/RTE-X |
| RSINE1 | 27,16 | -1,33 | 0,023939441 | chr19 | 3387798 | 3387958 | SINE/B4 |
| MER5A | 21,19 | -1,32 | 0,048562236 | chr19 | 38121240 | 38121432 | DNA/hAT-Charlie |
| B2_Mm2 | 21,09 | -1,32 | 0,03163285 | chr11 | 77524603 | 77524775 | SINE/B2 |
| B3 | 78,98 | -1,30 | 0,001022536 | chr18 | 46641380 | 46641593 | SINE/B2 |
| LSU-rRNA_Hsa | 2286,11 | -1,30 | 0,015835057 | chr1 | 102628053 | 102628340 | rRNA |
| RMER19C | 31,39 | -1,28 | 0,016505987 | chr19 | 5914323 | 5914838 | LTR/ERVK |
| LSU-rRNA_Hsa | 1756,92 | -1,28 | 0,041973946 | chr1 | 136364889 | 136365168 | rRNA |
| IAPLTR4 | 60,44 | -1,27 | 0,030333422 | chr18 | 60309845 | 60310147 | LTR/ERVK |
| ORR1A2-int | 482,96 | -1,27 | 0,001744869 | chr15 | 82685028 | 82686602 | LTR/ERVL-MaLR |
| B1_Mur1 | 59,12 | -1,26 | 0,002057178 | chr10 | 88469667 | 88469812 | SINE/Alu |
| RLTR1D | 266,11 | -1,26 | 0,000337944 | chr7 | 119544868 | 119545406 | LTR/ERV1 |
| RMER19B2 | 154,96 | -1,26 | 0,00154445 | chr6 | 141687114 | 141687838 | LTR/ERVK |
| B4 | 104,82 | -1,26 | 0,012905068 | chr8 | 114154282 | 114154417 | SINE/B4 |
| L1_Mur2 | 124,68 | -1,26 | 0,004354023 | chr5 | 87317694 | 87317910 | LINE/L1 |
| (CAAAAA)n | 2822,55 | -1,25 | 0,029069429 | chr3 | 14872352 | 14872406 | Simple_repeat |
| MIRb | 24,20 | -1,24 | 0,049983301 | chr5 | 123179783 | 123179836 | SINE/MIR |
| MMVL30-int | 163,42 | -1,23 | 0,009112535 | chr2 | 75638672 | 75642371 | LTR/ERV1 |
| Lx2 | 42,31 | -1,22 | 0,011418932 | chr11 | 30924137 | 30925614 | LINE/L1 |
| RMER10B | 30,04 | -1,22 | 0,043229585 | chr5 | 127709015 | 127709434 | LTR/ERVL |
| L1M3 | 196,34 | -1,20 | 0,021954338 | chr3 | 83040410 | 83040576 | LINE/L1 |
| L1M2 | 32,76 | -1,20 | 0,035713944 | chr14 | 66080769 | 66081210 | LINE/L1 |
| MLT2F | 92,44 | -1,18 | 0,005416488 | chr17 | 35472469 | 35472587 | LTR/ERVL |
| Lx7 | 255,51 | -1,15 | 0,013310888 | chr13 | 23824407 | 23825279 | LINE/L1 |
| Lx5c | 163,18 | -1,15 | 0,005824224 | chr12 | 72606229 | 72606744 | LINE/L1 |
| B1F2 | 33,99 | -1,14 | 0,025634054 | chr8 | 93185291 | 93185412 | SINE/Alu |
| ORR1A2 | 397,29 | -1,12 | 0,000242453 | chr6 | 128482531 | 128482846 | LTR/ERVL-MaLR |
| ORR1D2 | 86,05 | -1,09 | 0,029284962 | chr5 | 135139624 | 135139807 | LTR/ERVL-MaLR |
| L1VL2 | 95,00 | -1,06 | 0,004256946 | chr3 | 10536277 | 10540163 | LINE/L1 |
| MLTR11A | 93,04 | -1,05 | 0,049763019 | chr5 | 135139448 | 135139617 | LTR/ERVK |
| (TTTTTG)n | 72,41 | -1,01 | 0,021416371 | chr13 | 24813773 | 24813802 | Simple_repeat |
| L1_Mus2 | 329,02 | -0,97 | 0,049577075 | chr14 | 18525961 | 18529145 | LINE/L1 |
| IAPLTR2_Mm | 89,81 | -0,97 | 0,017482449 | chr15 | 99106517 | 99107001 | LTR/ERVK |
| B2_Mm2 | 134,93 | -0,92 | 0,025313685 | chr1 | 13624848 | 13625041 | SINE/B2 |
| RLTR20A3_MM | 141,28 | -0,81 | 0,029639647 | chr1 | 21263652 | 21263920 | LTR/ERVK |
| L1_Mur3 | 1004,30 | -0,79 | 0,015206019 | chr3 | 74849401 | 74851277 | LINE/L1 |
| RSINE1 | 131,53 | -0,75 | 0,045884983 | chr10 | 63077707 | 63077858 | SINE/B4 |
| Lx | 557,40 | 0,76 | 0,045881998 | chr12 | 116504194 | 116510415 | LINE/L1 |
| B2_Mm2 | 74,50 | 0,89 | 0,022827454 | chr1 | 118350338 | 118350527 | SINE/B2 |
| RMER19A | 71,05 | 0,95 | 0,031527744 | chr3 | 100785880 | 100786500 | LTR/ERVK |
| RMER19B | 181,79 | 1,03 | 0,011958172 | chr17 | 35429133 | 35429844 | LTR/ERVK |
| MMSAT4 | 132,88 | 1,04 | 0,000971377 | chr17 | 22286643 | 22288522 | Satellite |
| MurSatRep1 | 55,21 | 1,05 | 0,030364436 | chr7 | 39566914 | 39567412 | Unknown |
| L1Md_F2 | 394,35 | 1,09 | 6,38E-05 | chrX | 10321050 | 10325274 | LINE/L1 |
| RMER17D2 | 51,71 | 1,11 | 0,035405283 | chr5 | 151430649 | 151431459 | LTR/ERVK |
| RMER19B | 44,70 | 1,12 | 0,04656417 | chr17 | 35384636 | 35385309 | LTR/ERVK |
| CT-rich | 262,80 | 1,15 | 0,011029277 | chr17 | 90567506 | 90567634 | |
| Lx9 | 72,95 | 1,15 | 0,034524203 | chr3 | 87001234 | 87001434 | LINE/L1 |
| ORR1A4 | 49,56 | 1,16 | 0,019614648 | chr7 | 42474306 | 42474641 | LTR/ERVL-MaLR |
| MurSatRep1 | 74,97 | 1,17 | 0,013254407 | chr10 | 82227405 | 82227996 | Unknown |
| MMSAT4 | 138,76 | 1,17 | 0,000269852 | chr7 | 42474815 | 42475841 | Satellite |
| IAP1-MM_I-int | 168,36 | 1,21 | 0,023637469 | chrX | 9329227 | 9331590 | LTR/ERVK |
| MurSatRep1 | 136,16 | 1,22 | 0,001067897 | chr7 | 39564671 | 39566865 | Unknown |
| MMVL30-int | 73,82 | 1,23 | 0,046883771 | chr14 | 32117142 | 32119968 | LTR/ERV1 |
| Lx6 | 54,61 | 1,26 | 0,011675225 | chr1 | 27903622 | 27904066 | LINE/L1 |

| | | | | | | | |
|---|---|---|---|---|---|---|---|
| MTB-int | 111,97 | 1,27 | 0,021289185 | chr2 | 177908249 | 177909225 | LTR/ERVL-MaLR |
| ORR1E | 45,33 | 1,29 | 0,010523451 | chr11 | 79258759 | 79259062 | LTR/ERVL-MaLR |
| Lx3C | 100,29 | 1,33 | 0,009660452 | chr1 | 27902574 | 27902719 | LINE/L1 |
| ORR1A4-int | 48,94 | 1,34 | 0,003942994 | chr13 | 74605221 | 74606364 | LTR/ERVL-MaLR |
| MurSatRep1 | 25,09 | 1,34 | 0,040395159 | chr5 | 104582393 | 104583114 | Unknown |
| MMSAT4 | 111,34 | 1,37 | 0,002692167 | chr4 | 147512487 | 147515105 | Satellite |
| MMVL30-int | 30,81 | 1,37 | 0,016894948 | chr4 | 146937407 | 146940954 | LTR/ERV1 |
| ORR1B2 | 33,06 | 1,38 | 0,030590511 | chr7 | 42473919 | 42474275 | LTR/ERVL-MaLR |
| L1ME3A | 55,18 | 1,40 | 0,021486477 | chr3 | 132644812 | 132645265 | LINE/L1 |
| MurSatRep1 | 25,84 | 1,46 | 0,025296713 | chr16 | 98008406 | 98010091 | Unknown |
| MMVL30-int | 18,58 | 1,51 | 0,039140827 | chr10 | 127871767 | 127872897 | LTR/ERV1 |
| ORR1A2 | 126,01 | 1,51 | 0,000482053 | chr10 | 95823922 | 95824232 | LTR/ERVL-MaLR |
| ID_B1 | 160,10 | 1,54 | 0,001937706 | chr7 | 14438471 | 14438652 | SINE/B4 |
| RMER1A | 39,31 | 1,56 | 0,00951232 | chr9 | 122898410 | 122898615 | Other |
| MT2B2 | 33,07 | 1,59 | 0,016232883 | chr4 | 12132159 | 12132714 | LTR/ERVL |
| RLTR11A | 25,76 | 1,61 | 0,012889454 | chr17 | 34996674 | 34997136 | LTR/ERVK |
| MurSatRep1 | 97,36 | 1,61 | 0,010224369 | chr2 | 176682352 | 176682607 | Unknown |
| MLTR18_MM | 28,15 | 1,65 | 0,006575531 | chr17 | 28454854 | 28455306 | LTR/ERVK |
| Lx2 | 99,76 | 1,67 | 1,76E-05 | chr17 | 80508406 | 80509589 | LINE/L1 |
| MurSatRep1 | 21,43 | 1,68 | 0,008941012 | chr5 | 110077968 | 110078732 | Unknown |
| MMVL30-int | 18,92 | 1,69 | 0,01919938 | chr10 | 127872938 | 127873861 | LTR/ERV1 |
| MIR3 | 17,16 | 1,69 | 0,042196166 | chr11 | 102406821 | 102406903 | SINE/MIR |
| RLTR20B5_MM | 50,42 | 1,72 | 0,003853974 | chr8 | 85611610 | 85612174 | LTR/ERVK |
| MMVL30-int | 14,83 | 1,74 | 0,047297452 | chr14 | 32116702 | 32116853 | LTR/ERV1 |
| ORR1B1 | 129,87 | 1,75 | 7,19E-08 | chr12 | 71361769 | 71362159 | LTR/ERVL-MaLR |
| L1MdMus_I | 32,55 | 1,83 | 0,000451092 | chr8 | 85610403 | 85611503 | LINE/L1 |
| ORR1A2 | 18,07 | 1,86 | 0,027685987 | chr13 | 21389832 | 21390101 | LTR/ERVL-MaLR |
| MTE-int | 44,14 | 1,86 | 6,68E-05 | chr7 | 67727064 | 67728320 | LTR/ERVL-MaLR |
| MMSAT4 | 24,67 | 1,91 | 0,024359186 | chr2 | 177906415 | 177906886 | Satellite |
| ORR1B1 | 93,38 | 1,92 | 4,94E-07 | chr10 | 95824326 | 95824551 | LTR/ERVL-MaLR |
| MMSAT4 | 33,24 | 1,92 | 0,013126117 | chr2 | 177905922 | 177906389 | Satellite |
| RMER5 | 15,13 | 1,96 | 0,034909248 | chr6 | 3988739 | 3989075 | LTR/ERV1 |
| RMER6A | 13,24 | 1,98 | 0,030594128 | chr5 | 45517907 | 45518625 | LTR/ERVK |
| L1M5 | 11,21 | 1,99 | 0,037931778 | chr2 | 65338240 | 65338547 | LINE/L1 |
| B2_Mm1t | 14,49 | 1,99 | 0,031959232 | chr8 | 95374893 | 95375082 | SINE/B2 |
| MLT1J | 12,23 | 1,99 | 0,026689044 | chr5 | 74358759 | 74358927 | LTR/ERVL-MaLR |
| RMER19B | 13,54 | 2,00 | 0,042886098 | chr2 | 27578098 | 27579119 | LTR/ERVK |
| Lx6 | 46,85 | 2,02 | 8,83E-06 | chr1 | 151240076 | 151241686 | LINE/L1 |
| MurSatRep1 | 13,69 | 2,05 | 0,046599593 | chr18 | 90592019 | 90592248 | Unknown |
| B3 | 29,41 | 2,05 | 0,00552977 | chr19 | 10047929 | 10048146 | SINE/B2 |
| B4A | 10,72 | 2,06 | 0,043936099 | chr9 | 121966938 | 121967161 | SINE/B4 |
| Lx6 | 16,55 | 2,11 | 0,007541672 | chr15 | 55960008 | 55960498 | LINE/L1 |
| RLTR14-int | 30,24 | 2,12 | 0,000335812 | chr7 | 42490071 | 42490177 | LTR/ERV1 |
| MER89 | 107,39 | 2,13 | 9,18E-09 | chr13 | 68228645 | 68229011 | LTR/ERV1 |
| MMSAT4 | 11,05 | 2,13 | 0,033967844 | chr13 | 62639787 | 62641268 | Satellite |
| ORR1A1-int | 16,73 | 2,13 | 0,002851634 | chr9 | 22202586 | 22203592 | LTR/ERVL-MaLR |
| MER89 | 12,40 | 2,14 | 0,025296713 | chr9 | 90543303 | 90543775 | LTR/ERV1 |
| RNERVK23-int | 14,40 | 2,14 | 0,007950685 | chr17 | 34998194 | 34998562 | LTR/ERVK |
| LTR16B2 | 30,07 | 2,14 | 0,00308824 | chr12 | 70231181 | 70231444 | LTR/ERVL |
| B1_Mus1 | 19,46 | 2,16 | 0,003085972 | chr9 | 122896182 | 122896306 | SINE/Alu |
| RLTR17D_Mm | 30,36 | 2,17 | 0,01101646 | chr19 | 7781055 | 7781706 | LTR/ERVK |
| RSINE1 | 12,71 | 2,22 | 0,035736599 | chr12 | 8830261 | 8830412 | SINE/B4 |
| Lx3_Mus | 11,91 | 2,23 | 0,032633507 | chr5 | 91281437 | 91282166 | LINE/L1 |
| ORR1D-int | 16,46 | 2,24 | 0,002387657 | chr13 | 74612429 | 74613212 | LTR/ERVL-MaLR |
| MMVL30-int | 71,20 | 2,29 | 7,21E-11 | chr17 | 21475317 | 21477755 | LTR/ERV1 |
| MurSatRep1 | 13,49 | 2,30 | 0,011368796 | chr18 | 90590873 | 90591145 | Unknown |
| ORR1A1 | 24,77 | 2,32 | 0,000347807 | chr5 | 74359578 | 74359919 | LTR/ERVL-MaLR |
| Lx4A | 12,84 | 2,34 | 0,028147529 | chr8 | 5031416 | 5031982 | LINE/L1 |
| B1_Mus2 | 25,29 | 2,37 | 0,000158712 | chr11 | 84147285 | 84147427 | SINE/Alu |
| Zaphod | 10,33 | 2,39 | 0,032633507 | chr11 | 79273647 | 79273930 | DNA/hAT-Tip100 |
| Lx8 | 19,04 | 2,42 | 0,031167731 | chr9 | 74519756 | 74519844 | LINE/L1 |
| ERVB4_1B-LTR_MM | 14,84 | 2,43 | 0,031499543 | chr5 | 8203747 | 8204182 | LTR/ERVK |
| MMVL30-int | 127,94 | 2,46 | 1,23E-09 | chr4 | 12133727 | 12137565 | LTR/ERV1 |
| RLTR48A | 13,90 | 2,51 | 0,006441371 | chr10 | 127873978 | 127874489 | LTR/ERV1 |
| ORR1A4 | 46,53 | 2,51 | 5,56E-07 | chr18 | 90592250 | 90592582 | LTR/ERVL-MaLR |
| ID_B1 | 45,95 | 2,52 | 4,76E-06 | chr19 | 4057466 | 4057504 | SINE/B4 |
| L1_Mus3 | 31,47 | 2,57 | 3,56E-05 | chr15 | 55958075 | 55959269 | LINE/L1 |

| | | | | | | | |
|---|---|---|---|---|---|---|---|
| IAP1-MM_I-int | 8,70 | 2,57 | 0,020576846 | chr13 | 117443563 | 117445202 | LTR/ERVK |
| MuLV-int | 28,63 | 2,57 | 0,000207624 | chr4 | 147219404 | 147226971 | LTR/ERV1 |
| (TCC)n | 17,36 | 2,58 | 0,001046212 | chr10 | 127879330 | 127879643 | Simple_repeat |
| RLTR6_Mm | 14,64 | 2,58 | 0,00243543 | chr13 | 68232872 | 68233439 | LTR/ERV1 |
| RLTR18-int | 13,50 | 2,59 | 0,020133786 | chr6 | 37540670 | 37540933 | LTR/ERVK |
| RLTR6-int | 10,62 | 2,62 | 0,018601615 | chr15 | 32870194 | 32877740 | LTR/ERV1 |
| L1ME4b | 9,31 | 2,65 | 0,028860763 | chr11 | 76906754 | 76906870 | LINE/L1 |
| MLT2F | 23,37 | 2,67 | 5,67E-05 | chr18 | 74280202 | 74280766 | LTR/ERVL |
| RMER19B | 9,17 | 2,68 | 0,022723011 | chr13 | 47171896 | 47172753 | LTR/ERVK |
| MurSatRep1 | 15,92 | 2,70 | 0,002842823 | chr8 | 129280788 | 129282476 | Unknown |
| Lx8 | 55,10 | 2,73 | 9,15E-05 | chr5 | 122661764 | 122662907 | LINE/L1 |
| MMSAT4 | 20,78 | 2,74 | 0,000485256 | chr18 | 90591087 | 90591392 | Satellite |
| MurSatRep1 | 8,24 | 2,77 | 0,025165377 | chr5 | 109881779 | 109882074 | Unknown |
| B2_Mm2 | 125,53 | 2,80 | 3,81E-16 | chr12 | 71362236 | 71362417 | SINE/B2 |
| MurSatRep1 | 62,37 | 2,81 | 1,09E-09 | chr9 | 124269966 | 124271716 | Unknown |
| L1MB7 | 18,75 | 2,82 | 0,011207168 | chr6 | 72749640 | 72750018 | LINE/L1 |
| MMSAT4 | 32,66 | 2,85 | 3,88E-06 | chr4 | 147178604 | 147180143 | Satellite |
| RLTR6-int | 15,37 | 2,85 | 0,004989997 | chr11 | 75357406 | 75365045 | LTR/ERV1 |
| RLTR6-int | 24,54 | 2,89 | 0,000171695 | chr11 | 121311593 | 121317617 | LTR/ERV1 |
| B2_Mm2 | 12,70 | 2,94 | 0,00666972 | chr5 | 86104522 | 86104705 | SINE/B2 |
| LTR16B | 13,11 | 2,95 | 0,001526234 | chr1 | 190075418 | 190075790 | LTR/ERVL |
| Tigger17b | 18,99 | 2,98 | 0,000145874 | chr18 | 74270504 | 74271020 | DNA/TcMar-Tigger |
| LTR16E2 | 7,72 | 2,99 | 0,030518769 | chrX | 14927976 | 14928282 | LTR/ERVL |
| RMER4B | 7,58 | 3,00 | 0,036448456 | chr15 | 55959697 | 55959955 | LTR/ERVK |
| B3A | 7,68 | 3,01 | 0,020990728 | chr11 | 84144759 | 84144943 | SINE/B2 |
| RLTR22_Mus | 11,60 | 3,02 | 0,002136501 | chr15 | 77742971 | 77743968 | LTR/ERVK |
| L1Md_F2 | 9,71 | 3,03 | 0,01083526 | chr6 | 35893481 | 35893897 | LINE/L1 |
| L1Md_F2 | 17,20 | 3,03 | 0,000292799 | chr7 | 14726665 | 14727081 | LINE/L1 |
| B4 | 42,95 | 3,04 | 3,71E-07 | chr4 | 148716273 | 148716537 | SINE/B4 |
| MurSatRep1 | 23,46 | 3,06 | 1,45E-05 | chr18 | 90588711 | 90589448 | Unknown |
| MT2C_Mm | 48,02 | 3,07 | 4,08E-09 | chr13 | 68231635 | 68232102 | LTR/ERVL |
| MTD | 14,18 | 3,09 | 0,000803436 | chr1 | 190089797 | 190090203 | LTR/ERVL-MaLR |
| MurSatRep1 | 67,24 | 3,11 | 2,54E-11 | chr18 | 90591348 | 90591982 | Unknown |
| MTE-int | 10,50 | 3,12 | 0,016934923 | chrX | 14967547 | 14967831 | LTR/ERVL-MaLR |
| MurSatRep1 | 8,24 | 3,14 | 0,021629376 | chr8 | 129282988 | 129283793 | Unknown |
| Lx3_Mus | 12,47 | 3,16 | 0,003802422 | chrX | 14912234 | 14913788 | LINE/L1 |
| MLT1A0 | 34,37 | 3,18 | 3,76E-06 | chr14 | 28697429 | 28697726 | LTR/ERVL-MaLR |
| MTC | 19,98 | 3,25 | 0,000132589 | chr5 | 90580703 | 90581030 | LTR/ERVL-MaLR |
| RSINE1 | 6,67 | 3,25 | 0,04381027 | chr1 | 93122898 | 93123060 | SINE/B4 |
| ORR1B1 | 6,66 | 3,26 | 0,029532493 | chr5 | 109883343 | 109883683 | LTR/ERVL-MaLR |
| Lx2B2 | 104,65 | 3,26 | 0,003951127 | chrX | 38962906 | 38963744 | LINE/L1 |
| Tigger17 | 11,29 | 3,26 | 0,001887224 | chr18 | 74269645 | 74269812 | DNA/TcMar-Tigger |
| MLTR14 | 11,16 | 3,27 | 0,002568044 | chr5 | 130855011 | 130855457 | LTR/ERV1 |
| MMETn-int | 6,90 | 3,31 | 0,029284962 | chr16 | 98031556 | 98035578 | LTR/ERVK |
| L1Md_F2 | 7,03 | 3,32 | 0,025416208 | chr14 | 11342221 | 11343626 | LINE/L1 |
| RMER6B | 9,70 | 3,36 | 0,005039031 | chr18 | 74270122 | 74270503 | LTR/ERVK |
| B4A | 274,17 | 3,36 | 1,10E-24 | chr8 | 5083819 | 5084062 | SINE/B4 |
| RLTR9A3B | 11,97 | 3,36 | 0,001227099 | chr9 | 22205989 | 22206332 | LTR/ERVK |
| L1Md_T | 9,55 | 3,37 | 0,005993847 | chr3 | 114714563 | 114720675 | LINE/L1 |
| ORR1A3-int | 16,87 | 3,38 | 8,10E-05 | chr19 | 4546565 | 4547168 | LTR/ERVL-MaLR |
| ORR1C2 | 19,65 | 3,41 | 1,36E-05 | chr13 | 68232595 | 68232828 | LTR/ERVL-MaLR |
| ORR1A4 | 10,68 | 3,51 | 0,00309154 | chr9 | 94477638 | 94477998 | LTR/ERVL-MaLR |
| ORR1A0 | 49,77 | 3,51 | 1,46E-10 | chr4 | 49488243 | 49488533 | LTR/ERVL-MaLR |
| MurSatRep1 | 68,81 | 3,51 | 2,25E-15 | chr9 | 124269376 | 124269924 | Unknown |
| MMSAT4 | 10,66 | 3,54 | 0,002619289 | chr13 | 62049079 | 62051245 | Satellite |
| MTE-int | 23,88 | 3,55 | 1,69E-06 | chr7 | 48724615 | 48724668 | LTR/ERVL-MaLR |
| L1_Mur3 | 21,84 | 3,55 | 1,33E-05 | chr1 | 151241833 | 151243152 | LINE/L1 |
| RLTR6-int | 8,34 | 3,58 | 0,011607939 | chr12 | 87935182 | 87942857 | LTR/ERV1 |
| MMVL30-int | 13,85 | 3,61 | 0,000362728 | chr7 | 48739226 | 48741569 | LTR/ERV1 |
| RLTR6-int | 30,71 | 3,61 | 3,49E-07 | chr13 | 20650545 | 20658238 | LTR/ERV1 |
| Lx4A | 70,09 | 3,63 | 1,14E-08 | chrX | 14926951 | 14927604 | LINE/L1 |
| MMVL30-int | 73,39 | 3,64 | 5,61E-13 | chr13 | 68233440 | 68235807 | LTR/ERV1 |
| MMSAT4 | 29,54 | 3,67 | 1,55E-07 | chr9 | 124268874 | 124269179 | Satellite |
| MurSatRep1 | 11,86 | 3,67 | 0,001853443 | chr12 | 21015459 | 21016121 | Unknown |
| MT2B1 | 14,77 | 3,68 | 0,000107128 | chr17 | 83852561 | 83852974 | LTR/ERVL |
| LTRIS_Mus | 32,18 | 3,69 | 2,15E-08 | chr7 | 14727114 | 14727696 | LTR/ERV1 |
| B4 | 9,41 | 3,77 | 0,010241928 | chr1 | 190089668 | 190089796 | SINE/B4 |

| | | | | | | | |
|---|---|---|---|---|---|---|---|
| MER77 | 12,41 | 3,77 | 0,00123544 | chr4 | 11611054 | 11611361 | LTR/ERVL |
| L1MdV_I | 19,29 | 3,80 | 1,43E-05 | chr12 | 20245093 | 20245503 | LINE/L1 |
| MER57C2 | 6,47 | 3,81 | 0,023423362 | chr17 | 83851727 | 83851920 | LTR/ERV1 |
| Lx | 6,56 | 3,86 | 0,035836128 | chr4 | 146480273 | 146480704 | LINE/L1 |
| L1_Mus3 | 10,38 | 3,91 | 0,01825762 | chr7 | 26221785 | 26222924 | LINE/L1 |
| ID_B1 | 90,05 | 3,92 | 8,53E-14 | chr9 | 65236045 | 65236261 | SINE/B4 |
| MMVL30-int | 13,78 | 3,94 | 0,000602867 | chr8 | 55564388 | 55566719 | LTR/ERV1 |
| RLTR1E_MM | 7,23 | 4,01 | 0,034120096 | chr4 | 62200266 | 62200839 | LTR/ERV1 |
| MurSatRep1 | 7,93 | 4,11 | 0,016232883 | chr12 | 23615601 | 23615744 | Unknown |
| PB1D11 | 16,09 | 4,12 | 0,00059969 | chr5 | 122661291 | 122661399 | SINE/Alu |
| L1MA6 | 4,24 | 4,13 | 0,048603213 | chr6 | 117268879 | 117269247 | LINE/L1 |
| MurSatRep1 | 27,89 | 4,15 | 3,60E-07 | chr4 | 147176775 | 147176879 | Unknown |
| IAP1-MM_I-int | 4,33 | 4,16 | 0,041410626 | chr7 | 7127324 | 7129685 | LTR/ERVK |
| Tigger5b | 24,43 | 4,16 | 4,43E-07 | chr13 | 81339947 | 81340346 | DNA/TcMar-Tigger |
| MERVL-int | 4,42 | 4,19 | 0,038471539 | chr4 | 49496605 | 49498939 | LTR/ERVL |
| MuRRS4-int | 4,55 | 4,23 | 0,038923925 | chr15 | 8235088 | 8236807 | LTR/ERV1 |
| Lx5 | 4,56 | 4,24 | 0,038076815 | chr19 | 52828331 | 52829733 | LINE/L1 |
| MIRb | 8,93 | 4,30 | 0,006019035 | chr14 | 35127804 | 35128031 | SINE/MIR |
| RLTR9A | 4,84 | 4,31 | 0,049263487 | chr4 | 62194938 | 62195253 | LTR/ERVK |
| L1MdF_III | 4,96 | 4,35 | 0,042382113 | chr19 | 8283708 | 8285370 | LINE/L1 |
| B3 | 9,49 | 4,39 | 0,004039724 | chr9 | 121966328 | 121966535 | SINE/B2 |
| RLTR6B_Mm | 105,95 | 4,42 | 2,99E-23 | chr18 | 74267226 | 74267774 | LTR/ERV1 |
| ETnERV2-int | 9,58 | 4,42 | 0,004225461 | chr4 | 62195257 | 62195565 | LTR/ERVK |
| L1_Mur3 | 2,62 | 4,45 | 0,049589484 | chr3 | 114665585 | 114666145 | LINE/L1 |
| RMER19B | 2,62 | 4,45 | 0,047977457 | chr2 | 166506669 | 166507487 | LTR/ERVK |
| RMER19B | 5,34 | 4,46 | 0,022449631 | chr18 | 74290041 | 74290533 | LTR/ERVK |
| MT2A | 2,63 | 4,47 | 0,049730265 | chr13 | 97378961 | 97379336 | LTR/ERVL |
| RLTR23 | 10,10 | 4,47 | 0,005846768 | chr5 | 77136576 | 77137014 | LTR/ERV1 |
| RMER5 | 2,65 | 4,48 | 0,049263487 | chr15 | 61053179 | 61053477 | LTR/ERV1 |
| MurSatRep1 | 10,12 | 4,48 | 0,003146651 | chr9 | 124268760 | 124268904 | Unknown |
| RLTR6B_Mm | 2,73 | 4,50 | 0,048787686 | chr10 | 47640058 | 47640611 | LTR/ERV1 |
| MT2C_Mm | 2,74 | 4,50 | 0,045721387 | chr4 | 49499149 | 49499612 | LTR/ERVL |
| Lx3A | 2,68 | 4,50 | 0,049730265 | chr10 | 47711381 | 47715545 | LINE/L1 |
| RLTR10D2 | 2,72 | 4,51 | 0,040380533 | chr6 | 59079243 | 59079599 | LTR/ERVK |
| RLTR6-int | 5,48 | 4,51 | 0,017416785 | chr10 | 6399323 | 6406972 | LTR/ERV1 |
| ID_B1 | 5,48 | 4,52 | 0,029069429 | chr10 | 8967496 | 8967711 | SINE/B4 |
| IAP1-MM_LTR | 2,73 | 4,52 | 0,048669827 | chr6 | 142233439 | 142233732 | LTR/ERVK |
| Lx4A | 2,76 | 4,53 | 0,040380533 | chr2 | 150340334 | 150340736 | LINE/L1 |
| IAPLTR3 | 2,75 | 4,53 | 0,049763019 | chrY | 5296562 | 5296854 | LTR/ERVK |
| L1_Mus1 | 2,79 | 4,54 | 0,041034803 | chr7 | 123697340 | 123699687 | LINE/L1 |
| ORR1B1-int | 15,77 | 4,54 | 0,000166678 | chr10 | 82046949 | 82047592 | LTR/ERVL-MaLR |
| MIRb | 63,59 | 4,57 | 7,03E-12 | chr8 | 72205989 | 72206042 | SINE/MIR |
| RLTR10D2 | 2,83 | 4,58 | 0,042660195 | chr14 | 87205465 | 87205804 | LTR/ERVK |
| Lx8b | 2,83 | 4,58 | 0,040574156 | chr7 | 48725654 | 48725851 | LINE/L1 |
| ORR1F | 2,88 | 4,59 | 0,03703367 | chr3 | 133945621 | 133945794 | LTR/ERVL-MaLR |
| L1_Mus3 | 2,84 | 4,59 | 0,043563098 | chr7 | 26197871 | 26198292 | LINE/L1 |
| Lx8 | 2,88 | 4,60 | 0,037584986 | chr6 | 118329752 | 118331080 | LINE/L1 |
| Lx3A | 2,92 | 4,60 | 0,035478248 | chr3 | 16517053 | 16518487 | LINE/L1 |
| RLTR6B_Mm | 2,95 | 4,61 | 0,035836128 | chr9 | 90931516 | 90932070 | LTR/ERV1 |
| RLTR6-int | 10,90 | 4,61 | 0,002082466 | chr5 | 58318097 | 58325707 | LTR/ERV1 |
| MMVL30-int | 2,93 | 4,62 | 0,045882499 | chrY | 40027157 | 40029150 | LTR/ERV1 |
| Lx6 | 2,90 | 4,62 | 0,044078436 | chr5 | 47838926 | 47839567 | LINE/L1 |
| RSINE1 | 2,90 | 4,62 | 0,043167106 | chr9 | 83113238 | 83113353 | SINE/B4 |
| MMSAT4 | 10,91 | 4,62 | 0,002260969 | chr9 | 124268622 | 124268759 | Satellite |
| MurSatRep1 | 2,98 | 4,62 | 0,033869244 | chr13 | 62169252 | 62170909 | Unknown |
| RMER17D2 | 3,01 | 4,63 | 0,047783156 | chr3 | 16463283 | 16464057 | LTR/ERVK |
| MurSatRep1 | 2,97 | 4,63 | 0,03044704 | chr12 | 19866264 | 19867343 | Unknown |
| RLTR31B2 | 2,97 | 4,63 | 0,030431189 | chr1 | 60624157 | 60624493 | LTR/ERVK |
| Lx | 6,03 | 4,64 | 0,014332528 | chr4 | 146480929 | 146481077 | LINE/L1 |
| RSINE1 | 3,01 | 4,64 | 0,043906878 | chr17 | 5839699 | 5839843 | SINE/B4 |
| RLTR6-int | 2,96 | 4,64 | 0,040395159 | chr2 | 25719497 | 25727040 | LTR/ERV1 |
| L1MdFanc_II | 3,02 | 4,64 | 0,034671018 | chr1 | 103550121 | 103550359 | LINE/L1 |
| ID_B1 | 3,04 | 4,65 | 0,037142275 | chr18 | 74271716 | 74271910 | SINE/B4 |
| ORR1A4 | 2,96 | 4,65 | 0,03703367 | chr17 | 83847712 | 83848039 | LTR/ERVL-MaLR |
| B4A | 3,07 | 4,66 | 0,035670276 | chr10 | 68963599 | 68963748 | SINE/B4 |
| MERV1_LTR | 3,01 | 4,66 | 0,03297021 | chr10 | 6363934 | 6364317 | LTR/ERV1 |
| MTA_Mm-int | 28,43 | 4,66 | 1,81E-07 | chr4 | 3087309 | 3088378 | LTR/ERVL-MaLR |

| | | | | | | | |
|---|---|---|---|---|---|---|---|
| MTE2a | 11,41 | 4,66 | 0,001564013 | chr14 | 45495702 | 45495821 | LTR/ERVL-MaLR |
| IAP1-MM_I-int | 3,02 | 4,67 | 0,045291357 | chr17 | 34544223 | 34544844 | LTR/ERVK |
| RMER17A | 11,52 | 4,67 | 0,001551306 | chr15 | 100345490 | 100346283 | LTR/ERVK |
| MMSAT4 | 3,08 | 4,68 | 0,038813863 | chr5 | 109479123 | 109479245 | Satellite |
| L1MA7 | 3,11 | 4,68 | 0,037067776 | chr12 | 30053695 | 30054199 | LINE/L1 |
| RLTR6-int | 34,13 | 4,68 | 8,26E-09 | chr2 | 84901752 | 84906844 | LTR/ERV1 |
| Lx8 | 3,07 | 4,68 | 0,034316349 | chr6 | 118332860 | 118333867 | LINE/L1 |
| MurSatRep1 | 47,48 | 4,68 | 0,012538544 | chr12 | 20899875 | 20900505 | Unknown |
| RLTR43C | 3,05 | 4,69 | 0,036921665 | chr8 | 72064466 | 72064812 | LTR/ERVK |
| B3A | 3,06 | 4,69 | 0,044122963 | chr6 | 71457522 | 71457675 | SINE/B2 |
| RLTR44-int | 3,10 | 4,70 | 0,029262596 | chr2 | 164523963 | 164524029 | LTR/ERVK |
| IAP1-MM_I-int | 6,16 | 4,70 | 0,014462017 | chr17 | 34545264 | 34546257 | LTR/ERVK |
| RMER13B | 3,06 | 4,70 | 0,037893496 | chr7 | 25937342 | 25937849 | LTR/ERVK |
| IAP1-MM_LTR | 3,09 | 4,70 | 0,029388487 | chr5 | 47833626 | 47833917 | LTR/ERVK |
| B4A | 3,13 | 4,71 | 0,039237856 | chr19 | 52833671 | 52833935 | SINE/B4 |
| MTC | 3,14 | 4,71 | 0,029712301 | chr15 | 60992652 | 60993027 | LTR/ERVL-MaLR |
| L1_Mus3 | 6,29 | 4,71 | 0,012889454 | chr19 | 8282179 | 8283601 | LINE/L1 |
| ORR1A2-int | 3,12 | 4,71 | 0,025780689 | chr10 | 47684346 | 47684736 | LTR/ERVL-MaLR |
| BGLII_B | 6,33 | 4,72 | 0,03297021 | chr4 | 108224887 | 108225111 | LTR/ERVK |
| RLTR6B_Mm | 3,15 | 4,72 | 0,048076392 | chr1 | 8341285 | 8341847 | LTR/ERV1 |
| MMERVK10D3_I-int | 3,16 | 4,73 | 0,030406419 | chr1 | 104404772 | 104407690 | LTR/ERVK |
| IAP1-MM_I-int | 3,16 | 4,73 | 0,025075424 | chr3 | 97379061 | 97379342 | LTR/ERVK |
| L1MB7 | 3,16 | 4,74 | 0,03042031 | chr1 | 90864154 | 90865293 | LINE/L1 |
| L2a | 29,39 | 4,74 | 2,77E-07 | chr8 | 72206656 | 72206705 | LINE/L2 |
| MER5B | 11,87 | 4,74 | 0,00188365 | chr9 | 70088682 | 70088817 | DNA/hAT-Charlie |
| ORR1E | 3,25 | 4,74 | 0,043056938 | chr18 | 8881804 | 8882015 | LTR/ERVL-MaLR |
| RLTR6-int | 3,22 | 4,75 | 0,031751463 | chr4 | 143679600 | 143687290 | LTR/ERV1 |
| PB1D9 | 3,20 | 4,75 | 0,023711296 | chr11 | 80159364 | 80159470 | SINE/Alu |
| IAP1-MM_I-int | 12,09 | 4,76 | 0,001327079 | chr1 | 194967681 | 194970228 | LTR/ERVK |
| Lx3A | 3,30 | 4,77 | 0,027240221 | chr3 | 16461117 | 16462126 | LINE/L1 |
| IAP1-MM_I-int | 3,31 | 4,77 | 0,025296713 | chr15 | 19648019 | 19648788 | LTR/ERVK |
| IAP1-MM_I-int | 3,27 | 4,78 | 0,048076392 | chr3 | 6558740 | 6558953 | LTR/ERVK |
| RLTR6-int | 3,28 | 4,78 | 0,021800958 | chr4 | 123844070 | 123851724 | LTR/ERV1 |
| MLT1C | 3,27 | 4,79 | 0,029420414 | chr7 | 25869624 | 25869703 | LTR/ERVL-MaLR |
| ORR1D1 | 3,27 | 4,79 | 0,02539554 | chr17 | 34531945 | 34532122 | LTR/ERVL-MaLR |
| IAP1-MM_I-int | 3,36 | 4,79 | 0,02613975 | chr6 | 90293035 | 90296024 | LTR/ERVK |
| ORR1A1-int | 3,31 | 4,79 | 0,020235138 | chr8 | 124093188 | 124093680 | LTR/ERVL-MaLR |
| RLTR1A2_MM | 3,32 | 4,79 | 0,02895041 | chr17 | 46356725 | 46357222 | LTR/ERV1 |
| MMVL30-int | 3,33 | 4,80 | 0,021448013 | chr4 | 143976144 | 143979844 | LTR/ERV1 |
| MurERV4_19-int | 3,37 | 4,80 | 0,023485783 | chr13 | 50509688 | 50510568 | LTR/ERVK |
| RLTR6-int | 3,30 | 4,80 | 0,03674958 | chr8 | 73246217 | 73253802 | LTR/ERV1 |
| MTC | 3,30 | 4,81 | 0,02692713 | chr1 | 136667177 | 136667407 | LTR/ERVL-MaLR |
| MT2C_Mm | 3,41 | 4,82 | 0,029262596 | chr6 | 92215678 | 92216143 | LTR/ERVL |
| RLTR6-int | 3,36 | 4,83 | 0,036544494 | chr13 | 77636012 | 77643640 | LTR/ERV1 |
| CT-rich | 3,41 | 4,83 | 0,019185567 | chr1 | 190088861 | 190088991 | |
| MMSAT4 | 3,42 | 4,83 | 0,017207234 | chr5 | 109478808 | 109479091 | Satellite |
| ID2 | 3,35 | 4,83 | 0,043664055 | chr7 | 29701387 | 29701473 | SINE/ID |
| B1_Mus1 | 3,43 | 4,84 | 0,016750297 | chr12 | 20898391 | 20898524 | SINE/Alu |
| RLTR6-int | 3,39 | 4,84 | 0,040627918 | chr7 | 42390865 | 42398572 | LTR/ERV1 |
| ORR1B2 | 2456,25 | 4,84 | 2,51E-08 | chr11 | 55123578 | 55123774 | LTR/ERVL-MaLR |
| L2b | 25,27 | 4,85 | 1,34E-06 | chr8 | 72206228 | 72206272 | LINE/L2 |
| B1_Mur2 | 3,45 | 4,85 | 0,015530101 | chr13 | 48886116 | 48886251 | SINE/Alu |
| B2_Mm2 | 3,49 | 4,85 | 0,016310341 | chr12 | 51308223 | 51308422 | SINE/B2 |
| MER44B | 6,95 | 4,85 | 0,008995228 | chr5 | 77136042 | 77136299 | DNA/TcMar-Tigger |
| B2_Mm2 | 3,49 | 4,86 | 0,023447597 | chr16 | 30953762 | 30953959 | SINE/B2 |
| RLTR6B_Mm | 3,48 | 4,86 | 0,018771479 | chr1 | 27737595 | 27738145 | LTR/ERV1 |
| MLT1F2 | 3,53 | 4,87 | 0,017471432 | chr6 | 71458406 | 71458758 | LTR/ERVL-MaLR |
| RLTR6B_Mm | 3,47 | 4,87 | 0,023116465 | chr11 | 75356838 | 75357405 | LTR/ERV1 |
| RLTR44B | 3,44 | 4,87 | 0,026490473 | chr7 | 25868293 | 25868690 | LTR/ERVK |
| RMER6A | 3,51 | 4,87 | 0,016820414 | chr5 | 109470834 | 109471550 | LTR/ERVK |
| Lx2 | 3,52 | 4,87 | 0,015088385 | chr7 | 24831507 | 24833409 | LINE/L1 |
| RLTR6-int | 3,48 | 4,87 | 0,039545032 | chr12 | 20709341 | 20716964 | LTR/ERV1 |
| MMERVK10D3_I-int | 3,54 | 4,88 | 0,024983771 | chr6 | 59078884 | 59079241 | LTR/ERVK |
| RMER1B | 78,45 | 4,89 | 4,53E-06 | chr14 | 55160009 | 55161037 | Other |
| MMVL30-int | 3,55 | 4,89 | 0,021458699 | chr4 | 145569278 | 145571263 | LTR/ERV1 |
| URR1A | 3,59 | 4,90 | 0,016206702 | chr13 | 40780025 | 40780192 | DNA/hAT-Charlie |
| MMERVK9E_I-int | 7,10 | 4,90 | 0,006058608 | chr9 | 22212094 | 22213829 | LTR/ERVK |

| | | | | | | | |
|---|---|---|---|---|---|---|---|
| RMER16-int | 26,38 | 4,91 | 1,36E-06 | chr4 | 62197385 | 62198705 | LTR/ERVK |
| IAPEY5_I-int | 3,62 | 4,91 | 0,016375933 | chr10 | 47612147 | 47612587 | LTR/ERVK |
| MurSatRep1 | 3,63 | 4,91 | 0,015297336 | chr12 | 18872420 | 18873055 | Unknown |
| RLTR6-int | 3,60 | 4,91 | 0,019022434 | chr9 | 4847273 | 4854976 | LTR/ERV1 |
| MERV1_I-int | 3,61 | 4,92 | 0,014197652 | chr10 | 6363453 | 6363933 | LTR/ERV1 |
| MTE2b | 3,62 | 4,92 | 0,013826609 | chrX | 18395835 | 18396108 | LTR/ERVL-MaLR |
| ORR1A0 | 373,10 | 4,92 | 0,000286307 | chr4 | 62012950 | 62013295 | LTR/ERVL-MaLR |
| MMVL30-int | 3,67 | 4,94 | 0,011822341 | chr14 | 44546705 | 44548892 | LTR/ERV1 |
| RLTR28 | 3,75 | 4,95 | 0,021448013 | chr2 | 25709093 | 25709645 | LTR/ERVL |
| (A)n | 3,72 | 4,96 | 0,011037805 | chr14 | 13949559 | 13949583 | Simple_repeat |
| Lx | 3,71 | 4,96 | 0,012094833 | chr10 | 6365928 | 6366764 | LINE/L1 |
| RSINE1 | 3,70 | 4,97 | 0,015012025 | chr10 | 95828209 | 95828348 | SINE/B4 |
| IAP1-MM_LTR | 3,72 | 4,97 | 0,020711852 | chr7 | 29877694 | 29877972 | LTR/ERVK |
| RLTR26_Mus | 49,22 | 4,97 | 2,66E-10 | chr7 | 26213117 | 26213738 | LTR/ERVK |
| B2_Mm1t | 7,55 | 4,98 | 0,004556001 | chr18 | 90578067 | 90578257 | SINE/B2 |
| B2_Mm2 | 3,77 | 4,98 | 0,010569861 | chr16 | 31399589 | 31399776 | SINE/B2 |
| RMER17A-int | 3,84 | 5,00 | 0,011073588 | chr15 | 100346544 | 100348057 | LTR/ERVK |
| Lx8b | 3,78 | 5,01 | 0,018771479 | chr7 | 29701713 | 29701829 | LINE/L1 |
| MMVL30-int | 3,92 | 5,01 | 0,017273213 | chr18 | 74292569 | 74293992 | LTR/ERV1 |
| Lx3C | 7,79 | 5,02 | 0,010569861 | chr2 | 177940053 | 177940389 | LINE/L1 |
| MuRRS4-int | 7,94 | 5,05 | 0,015269191 | chr17 | 34805500 | 34806327 | LTR/ERV1 |
| RodERV21-int | 3,92 | 5,05 | 0,013781477 | chr13 | 75605926 | 75606267 | LTR/ERV1 |
| ORR1B2 | 7,87 | 5,06 | 0,010230408 | chr13 | 35958694 | 35959114 | LTR/ERVL-MaLR |
| RMER19B | 4,08 | 5,07 | 0,01073364 | chr12 | 51291833 | 51292684 | LTR/ERVK |
| L1Md_F2 | 4,06 | 5,08 | 0,008442429 | chr7 | 84839436 | 84840423 | LINE/L1 |
| L1_Mus1 | 4,04 | 5,08 | 0,007511114 | chr10 | 103577673 | 103581013 | LINE/L1 |
| Lx2 | 4,09 | 5,09 | 0,009505502 | chrX | 120114130 | 120114642 | LINE/L1 |
| RLTR6B_Mm | 4,10 | 5,09 | 0,011073588 | chr12 | 18852803 | 18853363 | LTR/ERV1 |
| ORR1A4 | 4,10 | 5,10 | 0,007223335 | chr18 | 74239463 | 74239734 | LTR/ERVL-MaLR |
| L1MdV_I | 8,23 | 5,11 | 0,01139152 | chr13 | 89367979 | 89368707 | LINE/L1 |
| L1Md_F2 | 4,12 | 5,11 | 0,007753751 | chr19 | 36530027 | 36531021 | LINE/L1 |
| RMER10B | 4,17 | 5,12 | 0,009176447 | chr18 | 74238298 | 74238464 | LTR/ERVL |
| B1_Mus1 | 4,13 | 5,12 | 0,008460174 | chr16 | 56113401 | 56113548 | SINE/Alu |
| Lx2B | 4,19 | 5,12 | 0,006607086 | chr5 | 58607922 | 58608448 | LINE/L1 |
| ORR1A2 | 4,21 | 5,13 | 0,007922128 | chr6 | 141977429 | 141977756 | LTR/ERVL-MaLR |
| Lx6 | 4,14 | 5,13 | 0,021871681 | chr16 | 81414944 | 81415607 | LINE/L1 |
| ORR1B1 | 4,24 | 5,13 | 0,007447702 | chrX | 12408289 | 12408652 | LTR/ERVL-MaLR |
| MuRRS-int | 54,58 | 5,14 | 2,33E-13 | chr7 | 14727697 | 14728565 | LTR/ERV1 |
| IAP1-MM_I-int | 4,25 | 5,15 | 0,006451089 | chr1 | 50812546 | 50812786 | LTR/ERVK |
| L1_Mur3 | 4,23 | 5,16 | 0,01966406 | chr1 | 50824599 | 50825513 | LINE/L1 |
| L1Md_F2 | 4,34 | 5,17 | 0,005799372 | chr12 | 19861672 | 19864542 | LINE/L1 |
| RLTR10D | 4,38 | 5,17 | 0,016232883 | chr13 | 11613649 | 11614002 | LTR/ERVK |
| MTE-int | 4,29 | 5,17 | 0,006429101 | chr5 | 100213526 | 100214034 | LTR/ERVL-MaLR |
| IAP1-MM_I-int | 4,30 | 5,18 | 0,007463835 | chr1 | 50025281 | 50025858 | LTR/ERVK |
| MMSAT4 | 4,35 | 5,18 | 0,007139757 | chr13 | 61693845 | 61693979 | Satellite |
| IAP1-MM_LTR | 4,29 | 5,18 | 0,014188752 | chr10 | 94693157 | 94693433 | LTR/ERVK |
| IAP1-MM_I-int | 8,58 | 5,18 | 0,003580338 | chr19 | 43348588 | 43351006 | LTR/ERVK |
| B4 | 47,93 | 5,18 | 2,08E-12 | chr4 | 49492020 | 49492301 | SINE/B4 |
| RLTR6-int | 8,69 | 5,20 | 0,006451089 | chr4 | 3313587 | 3317499 | LTR/ERV1 |
| MT2A | 4,40 | 5,20 | 0,005993847 | chr5 | 73948765 | 73949230 | LTR/ERVL |
| RLTR6-int | 4,41 | 5,21 | 0,013155266 | chrX | 79067709 | 79075419 | LTR/ERV1 |
| RLTR6-int | 4,40 | 5,22 | 0,015460536 | chr14 | 40125462 | 40133075 | LTR/ERV1 |
| RLTR17 | 4,53 | 5,24 | 0,007511114 | chr10 | 73326615 | 73327178 | LTR/ERVK |
| Lx5 | 4,56 | 5,25 | 0,004734788 | chr19 | 54803723 | 54804434 | LINE/L1 |
| ORR1B1 | 4,57 | 5,25 | 0,007239298 | chr10 | 82046563 | 82046941 | LTR/ERVL-MaLR |
| B1_Mus1 | 9,13 | 5,25 | 0,003085972 | chr12 | 20637572 | 20637717 | SINE/Alu |
| RLTR51A_Mm | 4,50 | 5,25 | 0,008995228 | chr13 | 75605470 | 75605713 | LTR/ERVK? |
| MMSAT4 | 4,53 | 5,26 | 0,006361072 | chr9 | 124269294 | 124269519 | Satellite |
| RLTR6-int | 4,56 | 5,26 | 0,006083819 | chr9 | 18120159 | 18127810 | LTR/ERV1 |
| MMERVK9C_I-int | 4,62 | 5,27 | 0,005728603 | chr6 | 85880712 | 85883466 | LTR/ERVK |
| (CATATA)n | 4,63 | 5,29 | 0,004665877 | chr9 | 83126185 | 83126358 | Simple_repeat |
| ETnERV2-int | 17,30 | 5,29 | 0,000176882 | chr4 | 62195659 | 62196432 | LTR/ERVK |
| RLTR6-int | 4,64 | 5,29 | 0,005431826 | chr18 | 42476250 | 42483917 | LTR/ERV1 |
| MMSAT4 | 4,65 | 5,29 | 0,005680423 | chr12 | 20899194 | 20899401 | Satellite |
| Lx2A | 4,65 | 5,29 | 0,010615697 | chr5 | 58632362 | 58634445 | LINE/L1 |
| B1_Mus1 | 4,68 | 5,30 | 0,003942994 | chr14 | 13951388 | 13951530 | SINE/Alu |
| RLTR6-int | 4,68 | 5,30 | 0,005765783 | chr4 | 40651997 | 40659710 | LTR/ERV1 |

| | | | | | | | |
|---|---|---|---|---|---|---|---|
| RLTR13D2 | 4,69 | 5,31 | 0,004913865 | chr8 | 72059739 | 72060690 | LTR/ERVK |
| IAP1-MM_LTR | 4,76 | 5,31 | 0,003859558 | chr6 | 71035163 | 71035454 | LTR/ERVK |
| MMERVK10D3_I-int | 4,80 | 5,32 | 0,006523529 | chr6 | 59073679 | 59078736 | LTR/ERVK |
| MurSatRep1 | 4,73 | 5,32 | 0,004321139 | chr12 | 20900626 | 20900922 | Unknown |
| RLTR6-int | 17,71 | 5,32 | 0,000562196 | chr8 | 4959837 | 4967457 | LTR/ERV1 |
| RLTR6B_Mm | 4,83 | 5,32 | 0,003991062 | chr3 | 12584924 | 12585484 | LTR/ERV1 |
| IAP1-MM_I-int | 4,75 | 5,33 | 0,005824224 | chr10 | 94694168 | 94694588 | LTR/ERVK |
| MTEb | 4,85 | 5,33 | 0,005478946 | chr18 | 8906088 | 8906249 | LTR/ERVL-MaLR |
| IAPLTR3-int | 4,89 | 5,34 | 0,003305285 | chrY | 5296855 | 5297997 | LTR/ERVK |
| L1_Mus2 | 4,81 | 5,34 | 0,004772515 | chr1 | 147381193 | 147382410 | LINE/L1 |
| MLT1A | 4,93 | 5,35 | 0,003776019 | chr14 | 35104767 | 35105082 | LTR/ERVL-MaLR |
| IAP1-MM_LTR | 4,86 | 5,35 | 0,003745676 | chr9 | 109510414 | 109510692 | LTR/ERVK |
| MTEa | 4,94 | 5,36 | 0,008067767 | chr18 | 8906323 | 8906613 | LTR/ERVL-MaLR |
| ORR1F-int | 4,87 | 5,36 | 0,004413449 | chr3 | 133944988 | 133945551 | LTR/ERVL-MaLR |
| L1_Mur2 | 4,88 | 5,36 | 0,004842244 | chr1 | 103558618 | 103559982 | LINE/L1 |
| RMER19A | 4,91 | 5,36 | 0,003416941 | chr8 | 34437892 | 34438510 | LTR/ERVK |
| MurSatRep1 | 9,74 | 5,36 | 0,001193873 | chr9 | 124269121 | 124269324 | Unknown |
| RLTR6-int | 4,88 | 5,37 | 0,00396246 | chr6 | 10397379 | 10405064 | LTR/ERV1 |
| RodERV21-int | 4,92 | 5,38 | 0,004383978 | chr13 | 75610964 | 75612953 | LTR/ERV1 |
| IAP1-MM_LTR | 5,06 | 5,39 | 0,003735727 | chr4 | 8884663 | 8884953 | LTR/ERVK |
| B1_Mus1 | 5,11 | 5,41 | 0,00261289 | chr12 | 24069877 | 24070024 | SINE/Alu |
| IAP1-MM_I-int | 5,08 | 5,41 | 0,00461485 | chr1 | 161610679 | 161613634 | LTR/ERVK |
| AT_rich | 5,12 | 5,42 | 0,00236312 | chr14 | 13950718 | 13950780 | |
| RMER17B | 5,10 | 5,42 | 0,005406433 | chr7 | 26223938 | 26224874 | LTR/ERVK |
| L1MdF_I | 5,12 | 5,42 | 0,002645799 | chr7 | 84835113 | 84839435 | LINE/L1 |
| RLTR12BD_Mm | 5,20 | 5,44 | 0,002943696 | chr13 | 98717490 | 98717925 | LTR/ERVK |
| ORR1F-int | 5,14 | 5,44 | 0,003529873 | chr3 | 133944325 | 133944960 | LTR/ERVL-MaLR |
| L1MdV_I | 29,02 | 5,44 | 3,96E-06 | chr12 | 20899421 | 20899826 | LINE/L1 |
| IAP1-MM_I-int | 5,16 | 5,44 | 0,003160895 | chr7 | 34095924 | 34098966 | LTR/ERVK |
| MuRRS4-int | 5,28 | 5,46 | 0,002349129 | chr7 | 29740627 | 29742259 | LTR/ERV1 |
| L1_Mus3 | 10,36 | 5,46 | 0,002343453 | chr7 | 26220257 | 26221724 | LINE/L1 |
| RLTR6-int | 5,27 | 5,47 | 0,003480186 | chr12 | 51310300 | 51318003 | LTR/ERV1 |
| RMER19B | 5,29 | 5,48 | 0,00321787 | chr7 | 125099645 | 125100591 | LTR/ERVK |
| (CAAT)n | 5,30 | 5,49 | 0,007293787 | chr7 | 26210970 | 26211013 | Simple_repeat |
| RLTR30D2_MM | 5,45 | 5,50 | 0,003085972 | chr18 | 74236540 | 74236594 | LTR/ERV1 |
| RLTR6-int | 5,42 | 5,52 | 0,005832664 | chr7 | 11481625 | 11486985 | LTR/ERV1 |
| Lx2A1 | 5,47 | 5,52 | 0,003859558 | chr16 | 81427550 | 81428014 | LINE/L1 |
| RLTR6-int | 5,49 | 5,52 | 0,001619596 | chr9 | 90932071 | 90939701 | LTR/ERV1 |
| Tigger17b | 10,98 | 5,53 | 0,000737764 | chr18 | 74269841 | 74270121 | DNA/TcMar-Tigger |
| RMER19B | 5,50 | 5,53 | 0,002824004 | chr13 | 58442331 | 58443183 | LTR/ERVK |
| ORR1D1 | 5,50 | 5,54 | 0,002742691 | chr7 | 26225393 | 26225472 | LTR/ERVL-MaLR |
| L1_Mus3 | 5,59 | 5,54 | 0,001917711 | chrX | 120117662 | 120118092 | LINE/L1 |
| ORR1F | 5,62 | 5,55 | 0,002547376 | chr6 | 71464139 | 71464373 | LTR/ERVL-MaLR |
| RMER4B | 5,65 | 5,55 | 0,001854562 | chr10 | 119417474 | 119417574 | LTR/ERVK |
| L1_Mus1 | 5,65 | 5,57 | 0,003146651 | chr8 | 80529818 | 80530699 | LINE/L1 |
| MurSatRep1 | 5,77 | 5,58 | 0,001647984 | chr10 | 82110270 | 82110900 | Unknown |
| IAP1-MM_I-int | 5,71 | 5,58 | 0,003388002 | chr10 | 94694644 | 94694917 | LTR/ERVK |
| RMER16-int | 5,81 | 5,59 | 0,001917711 | chr4 | 62198796 | 62199125 | LTR/ERVK |
| (TCTA)n | 5,71 | 5,59 | 0,002402919 | chr7 | 26211085 | 26211262 | Simple_repeat |
| ORR1A0-int | 5,82 | 5,59 | 0,001757906 | chr10 | 27867989 | 27868404 | LTR/ERVL-MaLR |
| IAP1-MM_I-int | 5,78 | 5,60 | 0,001506156 | chr9 | 109514652 | 109514848 | LTR/ERVK |
| Lx3_Mus | 32,28 | 5,61 | 7,92E-07 | chr13 | 75613798 | 75615023 | LINE/L1 |
| RLTR20C2_MM | 5,87 | 5,61 | 0,001131704 | chr7 | 35722634 | 35722721 | LTR/ERVK |
| MERVL-int | 5,88 | 5,62 | 0,001263409 | chr4 | 49493178 | 49496172 | LTR/ERVL |
| MMSAT4 | 5,95 | 5,62 | 0,001553145 | chr13 | 62111517 | 62111656 | Satellite |
| B2_Mm2 | 5,91 | 5,63 | 0,001845915 | chr7 | 34086072 | 34086218 | SINE/B2 |
| ERVB3_1-LTR_MM | 5,97 | 5,64 | 0,001142172 | chr13 | 45522050 | 45522547 | LTR/ERVK |
| Lx5 | 5,96 | 5,65 | 0,001601407 | chr14 | 96117699 | 96118706 | LINE/L1 |
| MurSatRep1 | 6,07 | 5,65 | 0,001483814 | chr12 | 24071624 | 24072845 | Unknown |
| B1_Mus1 | 6,16 | 5,68 | 0,001551306 | chr18 | 74236595 | 74236742 | SINE/Alu |
| RLTR6B_Mm | 6,09 | 5,68 | 0,002372899 | chr10 | 47747263 | 47747816 | LTR/ERV1 |
| RLTR11A | 12,14 | 5,68 | 0,000527762 | chr5 | 90586505 | 90586995 | LTR/ERVK |
| B1_Mus1 | 6,19 | 5,69 | 0,000995831 | chr9 | 90093020 | 90093164 | SINE/Alu |
| ORR1A2 | 6,29 | 5,70 | 0,002568044 | chr9 | 122873786 | 122874095 | LTR/ERVL-MaLR |
| Charlie6 | 6,27 | 5,70 | 0,00180612 | chr6 | 71465022 | 71465631 | DNA/hAT-Charlie |
| L2a | 93,95 | 5,71 | 4,31E-08 | chr8 | 72206918 | 72207064 | LINE/L2 |
| L2a | 46,24 | 5,73 | 2,95E-09 | chr8 | 72206282 | 72206393 | LINE/L2 |

| | | | | | | | |
|---|---|---|---|---|---|---|---|
| RLTR11A | 6,43 | 5,73 | 0,000972488 | chr10 | 103533470 | 103533990 | LTR/ERVK |
| IAP-d-int | 6,51 | 5,75 | 0,001361303 | chr13 | 11612420 | 11612768 | LTR/ERVK |
| Lx | 6,48 | 5,77 | 0,000721971 | chr1 | 103583434 | 103583818 | LINE/L1 |
| ORR1A4 | 23,79 | 5,77 | 9,66E-06 | chr4 | 103496143 | 103496440 | LTR/ERVL-MaLR |
| RLTR6-int | 6,51 | 5,78 | 0,001286466 | chr5 | 6484001 | 6491710 | LTR/ERV1 |
| RMER16_Mm | 6,66 | 5,79 | 0,002785719 | chr9 | 122876128 | 122876654 | LTR/ERVK |
| Lx3_Mus | 6,64 | 5,79 | 0,000799652 | chr10 | 6287688 | 6289241 | LINE/L1 |
| L1Lx_III | 6,65 | 5,80 | 0,001780667 | chr3 | 6487377 | 6487901 | LINE/L1 |
| B3 | 6,62 | 5,81 | 0,000834162 | chr7 | 26015947 | 26016128 | SINE/B2 |
| IAP1-MM_LTR | 6,79 | 5,81 | 0,000836686 | chr3 | 97376053 | 97376350 | LTR/ERVK |
| RLTR12H | 6,70 | 5,82 | 0,001227099 | chr16 | 32588549 | 32589071 | LTR/ERVK |
| ORR1A0 | 6,76 | 5,82 | 0,000544107 | chr13 | 62112043 | 62112387 | LTR/ERVL-MaLR |
| Lx8 | 6,74 | 5,82 | 0,000786315 | chr15 | 61052713 | 61052868 | LINE/L1 |
| RLTR17 | 6,82 | 5,83 | 0,000628194 | chr5 | 73946183 | 73946846 | LTR/ERVK |
| ORR1B2 | 6,77 | 5,83 | 0,000756342 | chr1 | 35992001 | 35992292 | LTR/ERVL-MaLR |
| Lx10 | 6,84 | 5,84 | 0,00058126 | chr3 | 133958875 | 133959498 | LINE/L1 |
| Lx7 | 6,92 | 5,84 | 0,000557115 | chr10 | 119418755 | 119419541 | LINE/L1 |
| L1MdF_I | 6,91 | 5,86 | 0,000508235 | chr1 | 103543248 | 103548366 | LINE/L1 |
| Lx9 | 13,68 | 5,86 | 0,000407346 | chr8 | 113136258 | 113136640 | LINE/L1 |
| MuRRS-int | 6,91 | 5,86 | 0,000712755 | chr13 | 50311600 | 50316853 | LTR/ERV1 |
| MYSERV6-int | 6,94 | 5,87 | 0,000489672 | chr5 | 58682457 | 58684489 | LTR/ERVK |
| RLTR12A | 13,76 | 5,87 | 0,000190214 | chr1 | 50820298 | 50820815 | LTR/ERVK |
| IAP1-MM_I-int | 7,08 | 5,88 | 0,000512116 | chr16 | 36221558 | 36221828 | LTR/ERVK |
| IAP1-MM_I-int | 7,21 | 5,90 | 0,001268511 | chr10 | 82514930 | 82515228 | LTR/ERVK |
| MT2B | 7,14 | 5,90 | 0,00070216 | chr13 | 75624641 | 75625228 | LTR/ERVL |
| MT2B | 7,26 | 5,91 | 0,000418482 | chr19 | 4547163 | 4547520 | LTR/ERVL |
| ORR1D2 | 7,23 | 5,93 | 0,000604232 | chr6 | 71457963 | 71458241 | LTR/ERVL-MaLR |
| L1MdV_I | 7,40 | 5,94 | 0,000351014 | chr12 | 23780214 | 23780623 | LINE/L1 |
| ORR1A4 | 7,41 | 5,95 | 0,000520632 | chr5 | 109479373 | 109479622 | LTR/ERVL-MaLR |
| RLTR6B_Mm | 27,07 | 5,95 | 5,56E-06 | chr7 | 14730668 | 14731235 | LTR/ERV1 |
| MERVK26-int | 7,53 | 5,96 | 0,002048927 | chr9 | 122883012 | 122883537 | LTR/ERVK |
| MurSatRep1 | 7,59 | 5,97 | 0,000359783 | chr12 | 19406325 | 19407047 | Unknown |
| MERVK26-int | 7,59 | 5,98 | 0,001913789 | chr9 | 122880642 | 122881711 | LTR/ERVK |
| RLTR6-int | 7,57 | 5,99 | 0,000525606 | chr1 | 195350520 | 195358233 | LTR/ERV1 |
| MTA_Mm | 7,72 | 6,00 | 0,000478279 | chr4 | 3086915 | 3087308 | LTR/ERVL-MaLR |
| IAP1-MM_I-int | 7,70 | 6,00 | 0,000257786 | chr5 | 58577778 | 58578039 | LTR/ERVK |
| RMER19B2 | 7,82 | 6,01 | 0,000467729 | chr13 | 40549633 | 40550151 | LTR/ERVK |
| RLTR16B_MM | 7,80 | 6,03 | 0,000284126 | chr2 | 104558978 | 104559658 | LTR/ERVK |
| L1_Mur3 | 7,74 | 6,03 | 0,001174919 | chr1 | 50825705 | 50826713 | LINE/L1 |
| MERV1_LTR | 7,92 | 6,04 | 0,000298333 | chr18 | 74348728 | 74349170 | LTR/ERV1 |
| L1MB2 | 7,81 | 6,04 | 0,000380328 | chr4 | 11857757 | 11858272 | LINE/L1 |
| ORR1G | 7,99 | 6,06 | 0,000264707 | chr8 | 110444184 | 110444431 | LTR/ERVL-MaLR |
| ORR1B2 | 8,08 | 6,07 | 0,000196705 | chr17 | 83854217 | 83854572 | LTR/ERVL-MaLR |
| RLTR1B-int | 8,28 | 6,11 | 0,000180838 | chr14 | 13934553 | 13935985 | LTR/ERV1 |
| RLTR6B_Mm | 8,39 | 6,12 | 0,000509312 | chr6 | 10928324 | 10928883 | LTR/ERV1 |
| Lx3A | 8,50 | 6,14 | 0,000168609 | chr3 | 16462149 | 16463208 | LINE/L1 |
| Lx3A | 8,59 | 6,16 | 0,000167056 | chr3 | 4982350 | 4983387 | LINE/L1 |
| (T)n | 8,57 | 6,18 | 0,000168082 | chr7 | 29878154 | 29878180 | Simple_repeat |
| IAP1-MM_I-int | 8,65 | 6,18 | 0,000116652 | chr1 | 147366693 | 147366997 | LTR/ERVK |
| IAP1-MM_I-int | 8,84 | 6,21 | 0,000106298 | chr4 | 8886498 | 8886764 | LTR/ERVK |
| MurSatRep1 | 8,97 | 6,22 | 0,000161126 | chr5 | 109475748 | 109477768 | Unknown |
| ORR1D2 | 9,00 | 6,23 | 9,15E-05 | chr14 | 35144674 | 35145029 | LTR/ERVL-MaLR |
| (A)n | 9,01 | 6,23 | 9,19E-05 | chr10 | 103566568 | 103566588 | Simple_repeat |
| RLTR11D | 9,02 | 6,24 | 0,000341797 | chr7 | 34089646 | 34090150 | LTR/ERVK |
| RLTR6-int | 9,16 | 6,26 | 0,000166654 | chr6 | 13452608 | 13460198 | LTR/ERV1 |
| L1Lx_III | 9,14 | 6,26 | 0,000241947 | chr3 | 6451676 | 6452262 | LINE/L1 |
| Lx8 | 9,17 | 6,26 | 8,50E-05 | chr13 | 40779229 | 40779847 | LINE/L1 |
| Lx3_Mus | 9,13 | 6,26 | 0,000153148 | chr7 | 5745349 | 5748519 | LINE/L1 |
| MMERGLN-int | 9,21 | 6,28 | 0,000269867 | chr17 | 46350152 | 46356719 | LTR/ERV1 |
| MMERVK9E_I-int | 9,25 | 6,28 | 0,000111811 | chr7 | 5735345 | 5736104 | LTR/ERVK |
| RLTR9E | 9,30 | 6,28 | 9,73E-05 | chr7 | 5736105 | 5736512 | LTR/ERVK |
| MurSatRep1 | 9,42 | 6,28 | 0,000179195 | chr13 | 62106145 | 62107137 | Unknown |
| B1_Mus1 | 9,35 | 6,28 | 0,000105725 | chr14 | 13952167 | 13952306 | SINE/Alu |
| MMERVK9E_I-int | 9,59 | 6,31 | 8,45E-05 | chr4 | 62196796 | 62197232 | LTR/ERVK |
| BGLII | 9,51 | 6,32 | 6,59E-05 | chr1 | 147372529 | 147372923 | LTR/ERVK |
| RLTR6-int | 9,56 | 6,32 | 0,018771479 | chr6 | 104218707 | 104226420 | LTR/ERV1 |
| IAP-d-int | 9,84 | 6,36 | 5,70E-05 | chr13 | 11612756 | 11613645 | LTR/ERVK |

| | | | | | | | |
|---|---|---|---|---|---|---|---|
| IAP1-MM_I-int | 9,73 | 6,36 | 8,42E-05 | chr10 | 94693434 | 94694177 | LTR/ERVK |
| L1MD | 10,10 | 6,40 | 0,000119251 | chr6 | 71463257 | 71463744 | LINE/L1 |
| IAP1-MM_I-int | 10,48 | 6,45 | 0,000101878 | chr6 | 36141922 | 36144843 | LTR/ERVK |
| RLTR6-int | 10,44 | 6,45 | 0,027488102 | chr14 | 9415732 | 9423317 | LTR/ERV1 |
| RLTR9F | 10,69 | 6,47 | 4,05E-05 | chr17 | 36108474 | 36108905 | LTR/ERVK |
| MurSatRep1 | 10,78 | 6,49 | 3,37E-05 | chr12 | 18648635 | 18649269 | Unknown |
| IAP1-MM_LTR | 10,75 | 6,49 | 3,37E-05 | chr1 | 50816146 | 50816436 | LTR/ERVK |
| Lx | 10,73 | 6,49 | 5,97E-05 | chr4 | 50123672 | 50124239 | LINE/L1 |
| B1_Mus2 | 10,97 | 6,51 | 2,92E-05 | chr13 | 40776720 | 40776865 | SINE/Alu |
| IAP1-MM_LTR | 11,07 | 6,54 | 3,57E-05 | chr13 | 117440970 | 117441243 | LTR/ERVK |
| Lx7 | 11,37 | 6,56 | 2,40E-05 | chr7 | 24814508 | 24815142 | LINE/L1 |
| RLTR6-int | 11,79 | 6,63 | 0,000135053 | chr13 | 3173202 | 3180884 | LTR/ERV1 |
| IAP1-MM_I-int | 11,76 | 6,63 | 3,79E-05 | chr7 | 26006183 | 26006394 | LTR/ERVK |
| B4 | 11,87 | 6,64 | 1,67E-05 | chr14 | 15155562 | 15155800 | SINE/B4 |
| RLTR35B_MM | 11,96 | 6,64 | 1,42E-05 | chr1 | 50818207 | 50818654 | LTR/ERVL |
| RLTR17 | 12,08 | 6,66 | 8,35E-05 | chr13 | 97482567 | 97483206 | LTR/ERVK |
| RLTR6-int | 12,09 | 6,67 | 3,32E-05 | chrX | 120080225 | 120087908 | LTR/ERV1 |
| L2c | 45,14 | 6,69 | 1,99E-07 | chr8 | 72206099 | 72206146 | LINE/L2 |
| RLTR10D | 12,48 | 6,70 | 1,91E-05 | chr10 | 103507123 | 103507463 | LTR/ERVK |
| RLTR14-int | 12,50 | 6,70 | 7,93E-05 | chr18 | 74232319 | 74232673 | LTR/ERV1 |
| ORR1D1 | 12,53 | 6,71 | 5,99E-05 | chr6 | 71462690 | 71463072 | LTR/ERVL-MaLR |
| BGLII_B | 12,46 | 6,71 | 2,34E-05 | chr7 | 29901386 | 29901754 | LTR/ERVK |
| RMER19B | 12,57 | 6,72 | 1,69E-05 | chr7 | 34088741 | 34089426 | LTR/ERVK |
| IAP1-MM_I-int | 12,71 | 6,73 | 1,29E-05 | chrX | 120159232 | 120159490 | LTR/ERVK |
| Lx9 | 12,62 | 6,73 | 1,69E-05 | chr1 | 147387534 | 147388139 | LINE/L1 |
| L1ME3B | 12,85 | 6,74 | 2,58E-05 | chr10 | 85339579 | 85339873 | LINE/L1 |
| RLTR10D2 | 12,84 | 6,74 | 8,09E-06 | chr14 | 87201614 | 87201970 | LTR/ERVK |
| ORR1B2 | 13,22 | 6,79 | 7,15E-06 | chr1 | 50816972 | 50817316 | LTR/ERVL-MaLR |
| RLTR6-int | 13,28 | 6,80 | 4,01E-05 | chr3 | 67417161 | 67424750 | LTR/ERV1 |
| MYSERV-int | 13,55 | 6,82 | 5,68E-06 | chr3 | 104981797 | 104982515 | LTR/ERVK |
| ORR1A2-int | 13,63 | 6,82 | 3,09E-05 | chr6 | 141975794 | 141977428 | LTR/ERVL-MaLR |
| B4A | 14,07 | 6,87 | 4,25E-06 | chr1 | 147390766 | 147390803 | SINE/B4 |
| RMER3D-int | 14,07 | 6,88 | 3,99E-06 | chr1 | 147374787 | 147375691 | LTR/ERVK |
| ORR1A2-int | 14,25 | 6,88 | 1,91E-05 | chr9 | 122874100 | 122874920 | LTR/ERVL-MaLR |
| RLTR1A2_MM | 14,27 | 6,90 | 5,01E-06 | chr18 | 24391961 | 24392445 | LTR/ERV1 |
| Lx3_Mus | 52,12 | 6,91 | 1,32E-07 | chr7 | 26222921 | 26223937 | LINE/L1 |
| Lx7 | 14,33 | 6,91 | 1,30E-05 | chr1 | 50826773 | 50827760 | LINE/L1 |
| Lx | 28,10 | 6,91 | 2,46E-06 | chr5 | 58690058 | 58690564 | LINE/L1 |
| MERV1_LTR | 14,55 | 6,92 | 1,16E-05 | chr18 | 74230558 | 74230686 | LTR/ERV1 |
| MMERVK10D3_I-int | 14,94 | 6,96 | 4,08E-06 | chr14 | 87203192 | 87205249 | LTR/ERVK |
| IAP1-MM_I-int | 15,26 | 6,99 | 3,44E-06 | chr3 | 16453601 | 16456313 | LTR/ERVK |
| RLTR13B1 | 30,26 | 7,01 | 1,68E-06 | chr1 | 147199721 | 147200636 | LTR/ERVK |
| ORR1A2 | 15,61 | 7,03 | 3,24E-06 | chr14 | 87147180 | 87147476 | LTR/ERVL-MaLR |
| ORR1B1 | 15,91 | 7,06 | 3,88E-06 | chr7 | 125069748 | 125069980 | LTR/ERVL-MaLR |
| MMVL30-int | 840,10 | 7,09 | 1,11E-20 | chr12 | 19119487 | 19121933 | LTR/ERV1 |
| MLT1A0 | 16,31 | 7,10 | 1,72E-06 | chr1 | 50817331 | 50817655 | LTR/ERVL-MaLR |
| IAP1-MM_I-int | 16,36 | 7,10 | 2,09E-06 | chr6 | 142230779 | 142233438 | LTR/ERVK |
| MTE2a | 16,73 | 7,14 | 5,56E-06 | chr7 | 29899256 | 29899628 | LTR/ERVL-MaLR |
| ORR1B1 | 17,56 | 7,20 | 1,37E-06 | chr1 | 121345963 | 121346335 | LTR/ERVL-MaLR |
| LTR31 | 17,82 | 7,22 | 1,60E-06 | chr18 | 24392867 | 24392963 | LTR |
| MTC-int | 18,05 | 7,24 | 7,92E-07 | chr15 | 60993027 | 60993980 | LTR/ERVL-MaLR |
| IAP1-MM_I-int | 18,46 | 7,28 | 1,39E-06 | chr9 | 109510693 | 109514468 | LTR/ERVK |
| L1_Mus3 | 18,58 | 7,28 | 8,66E-07 | chr3 | 6487902 | 6491660 | LINE/L1 |
| B2_Mm1t | 18,72 | 7,30 | 8,14E-07 | chr7 | 29901907 | 29902122 | SINE/B2 |
| L1Md_F2 | 19,06 | 7,32 | 1,10E-06 | chr5 | 47848892 | 47849886 | LINE/L1 |
| IAP1-MM_I-int | 19,59 | 7,35 | 2,27E-06 | chr3 | 97376351 | 97379005 | LTR/ERVK |
| RLTR6_Mm | 72,43 | 7,38 | 4,08E-09 | chr12 | 19121934 | 19122607 | LTR/ERV1 |
| RLTR1B-int | 20,75 | 7,44 | 2,52E-07 | chr14 | 13932744 | 13932998 | LTR/ERV1 |
| RLTR12BD_Mm | 20,96 | 7,45 | 4,40E-07 | chr13 | 45524913 | 45525323 | LTR/ERVK |
| IAPEY4_I-int | 21,06 | 7,46 | 3,10E-07 | chr19 | 8314110 | 8318880 | LTR/ERVK |
| IAP1-MM_I-int | 21,23 | 7,48 | 2,71E-07 | chr7 | 26007603 | 26012143 | LTR/ERVK |
| B3A | 21,35 | 7,48 | 2,73E-07 | chr14 | 13952324 | 13952492 | SINE/B2 |
| (ACCCCC)n | 21,76 | 7,52 | 4,65E-07 | chr7 | 29901806 | 29901843 | Simple_repeat |
| RMER19B2 | 22,17 | 7,52 | 2,06E-06 | chr9 | 122882533 | 122883011 | LTR/ERVK |
| ORR1B2-int | 22,25 | 7,54 | 3,08E-07 | chr1 | 50816437 | 50816971 | LTR/ERVL-MaLR |
| RMER19B | 22,68 | 7,56 | 1,36E-06 | chr8 | 72075967 | 72076819 | LTR/ERVK |
| Lx | 22,70 | 7,57 | 1,17E-07 | chr3 | 4979482 | 4981104 | LINE/L1 |

| | | | | | | | |
|---|---|---|---|---|---|---|---|
| RLTR6-int | 22,77 | 7,58 | 5,89E-07 | chr1 | 27732563 | 27737594 | LTR/ERV1 |
| MERV1_LTR | 23,46 | 7,61 | 1,84E-07 | chr18 | 24392446 | 24392515 | LTR/ERV1 |
| L1MdV_I | 24,72 | 7,68 | 8,11E-08 | chr12 | 24070922 | 24071327 | LINE/L1 |
| MERVK26-int | 26,53 | 7,79 | 6,05E-05 | chr9 | 122885310 | 122885455 | LTR/ERVK |
| ORR1D1 | 26,63 | 7,80 | 3,33E-08 | chr5 | 58604216 | 58604579 | LTR/ERVL-MaLR |
| MMVL30-int | 53,32 | 7,83 | 3,82E-08 | chr12 | 19117853 | 19118426 | LTR/ERV1 |
| Lx | 27,68 | 7,85 | 8,48E-05 | chr2 | 61541251 | 61542005 | LINE/L1 |
| LTR31 | 27,77 | 7,85 | 6,13E-08 | chr18 | 24393407 | 24393420 | LTR |
| MTEb | 28,06 | 7,87 | 2,79E-08 | chr6 | 142235105 | 142235356 | LTR/ERVL-MaLR |
| B4A | 28,51 | 7,89 | 2,09E-08 | chr1 | 147390623 | 147390741 | SINE/B4 |
| MMERVK10D3_I-int | 30,05 | 7,97 | 2,04E-08 | chr14 | 87205246 | 87205418 | LTR/ERVK |
| RMER19B | 30,76 | 8,01 | 1,17E-08 | chr16 | 56113549 | 56114356 | LTR/ERVK |
| RLTR4_MM-int | 31,71 | 8,04 | 3,66E-08 | chr18 | 74235602 | 74235738 | LTR/ERV1 |
| IAP1-MM_I-int | 31,84 | 8,06 | 1,61E-08 | chr4 | 8884954 | 8886098 | LTR/ERVK |
| IAP1-MM_I-int | 32,06 | 8,07 | 1,62E-08 | chr6 | 142230391 | 142230712 | LTR/ERVK |
| MTB_Mm | 32,43 | 8,09 | 2,15E-08 | chr7 | 26225854 | 26226246 | LTR/ERVL-MaLR |
| Lx | 32,85 | 8,11 | 1,11E-08 | chr5 | 58606265 | 58607159 | LINE/L1 |
| ORR1A3 | 33,82 | 8,14 | 6,26E-09 | chr13 | 40776050 | 40776417 | LTR/ERVL-MaLR |
| IAP1-MM_I-int | 33,78 | 8,14 | 9,98E-09 | chr1 | 50815156 | 50816145 | LTR/ERVK |
| L1M2 | 33,99 | 8,15 | 8,05E-09 | chr1 | 147388420 | 147388516 | LINE/L1 |
| IAP1-MM_I-int | 34,08 | 8,15 | 7,61E-09 | chr6 | 71035455 | 71038203 | LTR/ERVK |
| IAP1-MM_I-int | 339,96 | 8,29 | 1,53E-27 | chrX | 86533245 | 86535906 | LTR/ERVK |
| Lx3_Mus | 37,51 | 8,30 | 6,84E-09 | chr7 | 26224875 | 26225042 | LINE/L1 |
| RLTR6-int | 37,96 | 8,31 | 8,40E-09 | chr6 | 129804456 | 129812065 | LTR/ERV1 |
| B3 | 39,82 | 8,38 | 1,35E-09 | chr14 | 13950464 | 13950675 | SINE/B2 |
| MLT1A0 | 40,30 | 8,40 | 1,40E-09 | chr13 | 45524602 | 45524890 | LTR/ERVL-MaLR |
| RLTR6-int | 41,07 | 8,42 | 1,24E-05 | chr10 | 47747817 | 47755559 | LTR/ERV1 |
| MT2C_Mm | 41,70 | 8,44 | 1,89E-09 | chr5 | 68502273 | 68502757 | LTR/ERVL |
| Lx9 | 42,20 | 8,47 | 1,57E-09 | chr16 | 56112116 | 56113296 | LINE/L1 |
| RLTR30D2_MM | 42,79 | 8,48 | 4,08E-09 | chr18 | 74236743 | 74237359 | LTR/ERV1 |
| L1M2 | 42,77 | 8,48 | 8,42E-10 | chr1 | 147389248 | 147389481 | LINE/L1 |
| L1_Mus3 | 44,27 | 8,53 | 6,26E-09 | chr7 | 26225043 | 26225391 | LINE/L1 |
| L1_Mus3 | 44,55 | 8,54 | 6,08E-10 | chrX | 120118097 | 120120563 | LINE/L1 |
| MTEa | 95,51 | 8,68 | 4,79E-10 | chr8 | 126885313 | 126885599 | LTR/ERVL-MaLR |
| L1_Mus3 | 189,91 | 8,77 | 2,26E-14 | chr1 | 147389670 | 147390364 | LINE/L1 |
| MMVL30-int | 117,49 | 8,98 | 6,66E-11 | chr12 | 19118682 | 19119399 | LTR/ERV1 |
| MMERVK10D3_I-int | 60,91 | 8,99 | 1,69E-10 | chr14 | 87201977 | 87202998 | LTR/ERVK |
| PB1D9 | 64,03 | 9,06 | 4,63E-11 | chr7 | 29878054 | 29878142 | SINE/Alu |
| IAP1-MM_I-int | 64,41 | 9,07 | 4,19E-11 | chrX | 86535962 | 86536228 | LTR/ERVK |
| L1MdF_III | 67,79 | 9,15 | 5,12E-11 | chr5 | 47843554 | 47848889 | LINE/L1 |
| RLTR14-int | 73,27 | 9,25 | 5,65E-11 | chr18 | 74236164 | 74236505 | LTR/ERV1 |
| IAP1-MM_LTR | 73,67 | 9,26 | 1,37E-11 | chrX | 86532956 | 86533244 | LTR/ERVK |
| IAPEY_LTR | 77,00 | 9,33 | 7,26E-12 | chr10 | 103567110 | 103567493 | LTR/ERVK |
| ORR1A3-int | 86,98 | 9,50 | 1,37E-12 | chr13 | 40775065 | 40776049 | LTR/ERVL-MaLR |
| MERVK26-int | 90,75 | 9,56 | 1,21E-08 | chr9 | 122884028 | 122884527 | LTR/ERVK |
| IAP1-MM_I-int | 132,17 | 10,11 | 5,74E-14 | chr13 | 117441244 | 117442302 | LTR/ERVK |
| RLTR6_Mm | 144,74 | 10,24 | 2,75E-14 | chr12 | 19117210 | 19117852 | LTR/ERV1 |
| RLTR10D | 306,06 | 10,36 | 3,72E-15 | chr3 | 88369595 | 88369986 | LTR/ERVK |
| RLTR24 | 237,94 | 10,96 | 1,99E-16 | chr18 | 24393456 | 24393925 | LTR/ERV1 |
