## Supplementary material for "The mouse HP1 proteins are essential for preventing liver tumorigenesis": suppl table 8

**Supplementary table 8** : repeats with increased expression upon loss of HP1 associated with genes up-regulated upon loss of HP1

| gene | repeats | repeat-ch | repeat-start | repeat-end | family-repeat | gene-start | gene-end |
|---|---|---|---|---|---|---|---|
| Cyp2a4 | (CAAT)n | chr7 | 26210970 | 26211013 | Simple | 26207192 | 26415088 |
| Cyp2b9 | (CAAT)n | chr7 | 26210970 | 26211013 | Simple | 26073409 | 26310660 |
| Zfp84 | (T)n | chr7 | 29878154 | 29878180 | Simple | 29668552 | 29881419 |
| Cyp2a4 | (TCTA)n | chr7 | 26211085 | 26211262 | Simple | 26207192 | 26415088 |
| Cyp2b9 | (TCTA)n | chr7 | 26211085 | 26211262 | Simple | 26073409 | 26310660 |
| 9030624G23Rik | B1_Mus1 | chr12 | 24069877 | 24070024 | SINE/Alu | 23943202 | 24197269 |
| Ccdc11 | B1_Mus1 | chr18 | 74236595 | 74236742 | SINE/Alu | 74183100 | 74459984 |
| Impg2 | B1_Mus1 | chr16 | 56113401 | 56113548 | SINE/Alu | 56104313 | 56373753 |
| Mbd1 | B1_Mus1 | chr18 | 74236595 | 74236742 | SINE/Alu | 74168288 | 74382684 |
| Senp7 | B1_Mus1 | chr16 | 56113401 | 56113548 | SINE/Alu | 55975409 | 56290011 |
| Zfp105 | B1_Mus1 | chr9 | 122896182 | 122896306 | SINE/Alu | 122823078 | 123031028 |
| Zfp445 | B1_Mus1 | chr9 | 122896182 | 122896306 | SINE/Alu | 122748910 | 122966006 |
| Zkscan7 | B1_Mus1 | chr9 | 122896182 | 122896306 | SINE/Alu | 122788471 | 122996124 |
| Acaca | B1_Mus2 | chr11 | 84147285 | 84147427 | SINE/Alu | 84095438 | 84501651 |
| Gm11437 | B1_Mus2 | chr11 | 84147285 | 84147427 | SINE/Alu | 84048361 | 84267476 |
| Mmp15 | B2_Mm1t | chr8 | 95374893 | 95375082 | SINE/B2 | 95252337 | 95474293 |
| Tmx3 | B2_Mm1t | chr18 | 90578067 | 90578257 | SINE/B2 | 90410154 | 90643267 |
| G2e3 | B2_Mm2 | chr12 | 51308223 | 51308422 | SINE/B2 | 51248230 | 51476986 |
| Stap1 | B2_Mm2 | chr5 | 86104522 | 86104705 | SINE/B2 | 85971828 | 86203993 |
| Uba6 | B2_Mm2 | chr5 | 86104522 | 86104705 | SINE/B2 | 86010730 | 86272743 |
| 1700048O20Rik | B3 | chr9 | 121966328 | 121966535 | SINE/B2 | 121837275 | 122047016 |
| Cyp2b10 | B3 | chr7 | 26015947 | 26016128 | SINE/B2 | 25797658 | 26026624 |
| Cyp2b13 | B3 | chr7 | 26015947 | 26016128 | SINE/B2 | 25961495 | 26196196 |
| Fads3 | B3 | chr19 | 10047929 | 10048146 | SINE/B2 | 9941548 | 10159671 |
| Fam198a | B3 | chr9 | 121966328 | 121966535 | SINE/B2 | 121850988 | 122080208 |
| Acaca | B3A | chr11 | 84144759 | 84144943 | SINE/B2 | 84095438 | 84501651 |
| Gm11437 | B3A | chr11 | 84144759 | 84144943 | SINE/B2 | 84048361 | 84267476 |
| Acnat2 | B4 | chr4 | 49492020 | 49492301 | SINE/B4 | 49279845 | 49508151 |
| 1700048O20Rik | B4A | chr9 | 121966938 | 121967161 | SINE/B4 | 121837275 | 122047016 |
| Fam198a | B4A | chr9 | 121966938 | 121967161 | SINE/B4 | 121850988 | 122080208 |
| Slc10a2 | B4A | chr8 | 5083819 | 5084062 | SINE/B4 | 4985623 | 5205232 |
| BC089597 | CT-rich | chr10 | 127879330 | 127879643 | Simple | 127766476 | 127977319 |
| Mup21 | ETnERV2-int | chr4 | 62195659 | 62196432 | LTR/ERVK | 62047832 | 62250841 |
| Mup21 | ETnERV2-int | chr4 | 62195257 | 62195565 | LTR/ERVK | 62047832 | 62250841 |
| 2010005H15Rik | IAP1-MM_I-int | chr16 | 36221558 | 36221828 | LTR/ERVK | 36121562 | 36357427 |
| BC048671 | IAP1-MM_I-int | chr6 | 90293035 | 90296024 | LTR/ERVK | 90201270 | 90405448 |
| Btnl4 | IAP1-MM_I-int | chr17 | 34545264 | 34546257 | LTR/ERVK | 34369042 | 34575937 |
| Btnl4 | IAP1-MM_I-int | chr17 | 34544223 | 34544844 | LTR/ERVK | 34369042 | 34575937 |
| Btnl5-ps | IAP1-MM_I-int | chr17 | 34545264 | 34546257 | LTR/ERVK | 34387406 | 34597429 |
| Btnl5-ps | IAP1-MM_I-int | chr17 | 34544223 | 34544844 | LTR/ERVK | 34387406 | 34597429 |
| Btnl6 | IAP1-MM_I-int | chr17 | 34545264 | 34546257 | LTR/ERVK | 34407935 | 34617352 |
| Btnl6 | IAP1-MM_I-int | chr17 | 34544223 | 34544844 | LTR/ERVK | 34407935 | 34617352 |
| Cd34 | IAP1-MM_I-int | chr1 | 194967681 | 194970228 | LTR/ERVK | 194838821 | 195061292 |
| Cd46 | IAP1-MM_I-int | chr1 | 194967681 | 194970228 | LTR/ERVK | 194941900 | 195192248 |
| Cep83 | IAP1-MM_I-int | chr10 | 94694644 | 94694917 | LTR/ERVK | 94588790 | 94890336 |
| Cep83 | IAP1-MM_I-int | chr10 | 94694168 | 94694588 | LTR/ERVK | 94588790 | 94890336 |
| Cep83 | IAP1-MM_I-int | chr10 | 94693434 | 94694177 | LTR/ERVK | 94588790 | 94890336 |
| Cep83os | IAP1-MM_I-int | chr10 | 94694644 | 94694917 | LTR/ERVK | 94573493 | 94788613 |
| Cep83os | IAP1-MM_I-int | chr10 | 94694168 | 94694588 | LTR/ERVK | 94573493 | 94788613 |
| Cep83os | IAP1-MM_I-int | chr10 | 94693434 | 94694177 | LTR/ERVK | 94573493 | 94788613 |
| Cyp2b10 | IAP1-MM_I-int | chr7 | 26007603 | 26012143 | LTR/ERVK | 25797658 | 26026624 |
| Cyp2b10 | IAP1-MM_I-int | chr7 | 26006183 | 26006394 | LTR/ERVK | 25797658 | 26026624 |
| Cyp2b13 | IAP1-MM_I-int | chr7 | 26007603 | 26012143 | LTR/ERVK | 25961495 | 26196196 |
| Cyp2b13 | IAP1-MM_I-int | chr7 | 26006183 | 26006394 | LTR/ERVK | 25961495 | 26196196 |
| Fbxw16 | IAP1-MM_I-int | chr9 | 109514652 | 109514848 | LTR/ERVK | 109332318 | 109549140 |
| Fbxw16 | IAP1-MM_I-int | chr9 | 109510693 | 109514468 | LTR/ERVK | 109332318 | 109549140 |
| Fbxw19 | IAP1-MM_I-int | chr9 | 109514652 | 109514848 | LTR/ERVK | 109378575 | 109595854 |
| Fbxw19 | IAP1-MM_I-int | chr9 | 109510693 | 109514468 | LTR/ERVK | 109378575 | 109595854 |
| Gm16897 | IAP1-MM_I-int | chr1 | 194967681 | 194970228 | LTR/ERVK | 194855759 | 195076959 |

| | | | | | | | |
|---|---|---|---|---|---|---|---|
| Zfp773 | IAP1-MM_I-int | chr7 | 7127324 | 7129685 | LTR/ERVK | 7030678 | 7236755 |
| Cep83 | IAP1-MM_LTR | chr10 | 94693157 | 94693433 | LTR/ERVK | 94588790 | 94890336 |
| Cep83os | IAP1-MM_LTR | chr10 | 94693157 | 94693433 | LTR/ERVK | 94573493 | 94788613 |
| Fbxw16 | IAP1-MM_LTR | chr9 | 109510414 | 109510692 | LTR/ERVK | 109332318 | 109549140 |
| Fbxw19 | IAP1-MM_LTR | chr9 | 109510414 | 109510692 | LTR/ERVK | 109378575 | 109595854 |
| Zfp84 | IAP1-MM_LTR | chr7 | 29877694 | 29877972 | LTR/ERVK | 29668552 | 29881419 |
| 1700055N04Rik | ID_B1 | chr19 | 4057466 | 4057504 | SINE/B4 | 3858808 | 4070438 |
| BC021614 | ID_B1 | chr19 | 4057466 | 4057504 | SINE/B4 | 3957487 | 4159294 |
| Ccdc11 | ID_B1 | chr18 | 74271716 | 74271910 | SINE/B4 | 74183100 | 74459984 |
| Mbd1 | ID_B1 | chr18 | 74271716 | 74271910 | SINE/B4 | 74168288 | 74382684 |
| Parp16 | ID_B1 | chr9 | 65236045 | 65236261 | SINE/B4 | 65114690 | 65339219 |
| Catsperg2 | ID2 | chr7 | 29701387 | 29701473 | SINE/ID | 29597219 | 29827015 |
| Zfp84 | ID2 | chr7 | 29701387 | 29701473 | SINE/ID | 29668552 | 29881419 |
| 9030624G23Rik | L1_Mus1 | chr12 | 24070922 | 24071327 | LINE/L1 | 23943202 | 24197269 |
| Cyp2a4 | L1_Mus3 | chr7 | 26225043 | 26225391 | LINE/L1 | 26207192 | 26415088 |
| Cyp2a4 | L1_Mus3 | chr7 | 26221785 | 26222924 | LINE/L1 | 26207192 | 26415088 |
| Cyp2a4 | L1_Mus3 | chr7 | 26220257 | 26221724 | LINE/L1 | 26207192 | 26415088 |
| Cyp2b9 | L1_Mus3 | chr7 | 26225043 | 26225391 | LINE/L1 | 26073409 | 26310660 |
| Cyp2b9 | L1_Mus3 | chr7 | 26221785 | 26222924 | LINE/L1 | 26073409 | 26310660 |
| Cyp2b9 | L1_Mus3 | chr7 | 26220257 | 26221724 | LINE/L1 | 26073409 | 26310660 |
| Cyp2b9 | L1_Mus3 | chr7 | 26197871 | 26198292 | LINE/L1 | 26073409 | 26310660 |
| Sntb1 | L1_Mus3 | chr15 | 55958075 | 55959269 | LINE/L1 | 55539154 | 56006949 |
| Tpo | L1MA7 | chr12 | 30053695 | 30054199 | LINE/L1 | 29954661 | 30232609 |
| Hectd2 | L1Md_F2 | chr19 | 36530027 | 36531021 | LINE/L1 | 36454639 | 36721135 |
| Gm5549 | L1ME3A | chr3 | 132644812 | 132645265 | LINE/L1 | 132530191 | 132744649 |
| Cib3 | L2a | chr8 | 72206918 | 72207064 | LINE/L2 | 72104335 | 72312837 |
| Cib3 | L2a | chr8 | 72206656 | 72206705 | LINE/L2 | 72104335 | 72312837 |
| Cib3 | L2a | chr8 | 72206282 | 72206393 | LINE/L2 | 72104335 | 72312837 |
| Tpm4 | L2a | chr8 | 72206918 | 72207064 | LINE/L2 | 72035292 | 72253129 |
| Tpm4 | L2a | chr8 | 72206656 | 72206705 | LINE/L2 | 72035292 | 72253129 |
| Tpm4 | L2a | chr8 | 72206282 | 72206393 | LINE/L2 | 72035292 | 72253129 |
| Cib3 | L2b | chr8 | 72206228 | 72206272 | LINE/L2 | 72104335 | 72312837 |
| Tpm4 | L2b | chr8 | 72206228 | 72206272 | LINE/L2 | 72035292 | 72253129 |
| Cib3 | L2c | chr8 | 72206099 | 72206146 | LINE/L2 | 72104335 | 72312837 |
| Tpm4 | L2c | chr8 | 72206099 | 72206146 | LINE/L2 | 72035292 | 72253129 |
| Rprd1a | LTR31 | chr18 | 24393407 | 24393420 | LTR | 24384962 | 24630204 |
| Rprd1a | LTR31 | chr18 | 24392867 | 24392963 | LTR | 24384962 | 24630204 |
| D430020J02Rik | Lx | chr12 | 116504194 | 116505161 | LINE/L1 | 116301947 | 116505161 |
| Gm13247 | Lx | chr4 | 146480929 | 146481077 | LINE/L1 | 146402000 | 146639395 |
| Gm13247 | Lx | chr4 | 146480273 | 146480704 | LINE/L1 | 146402000 | 146639395 |
| Gm13251 | Lx | chr4 | 146480929 | 146481077 | LINE/L1 | 146349030 | 146569440 |
| Gm13251 | Lx | chr4 | 146480273 | 146480704 | LINE/L1 | 146349030 | 146569440 |
| Ncam2 | Lx2A1 | chr16 | 81427550 | 81428014 | LINE/L1 | 81100697 | 81724287 |
| Cyp2a4 | Lx3_Mus | chr7 | 26224875 | 26225042 | LINE/L1 | 26207192 | 26415088 |
| Cyp2a4 | Lx3_Mus | chr7 | 26222921 | 26223937 | LINE/L1 | 26207192 | 26415088 |
| Cyp2b9 | Lx3_Mus | chr7 | 26224875 | 26225042 | LINE/L1 | 26073409 | 26310660 |
| Cyp2b9 | Lx3_Mus | chr7 | 26222921 | 26223937 | LINE/L1 | 26073409 | 26310660 |
| Gm14124 | Lx4A | chr2 | 150340334 | 150340736 | LINE/L1 | 150157517 | 150370300 |
| Slc10a2 | Lx4A | chr8 | 5031416 | 5031982 | LINE/L1 | 4985623 | 5205232 |
| Ncam2 | Lx6 | chr16 | 81414944 | 81415607 | LINE/L1 | 81100697 | 81724287 |
| Sntb1 | Lx6 | chr15 | 55960008 | 55960498 | LINE/L1 | 55539154 | 56006949 |
| 1700025F24Rik | Lx7 | chr10 | 119418755 | 119419541 | LINE/L1 | 119313444 | 119526864 |
| Grip1 | Lx7 | chr10 | 119418755 | 119419541 | LINE/L1 | 119354034 | 120187260 |
| Catsperg2 | Lx8b | chr7 | 29701713 | 29701829 | LINE/L1 | 29597219 | 29827015 |
| Zfp84 | Lx8b | chr7 | 29701713 | 29701829 | LINE/L1 | 29668552 | 29881419 |
| Impg2 | Lx9 | chr16 | 56112116 | 56113296 | LINE/L1 | 56104313 | 56373753 |
| Senp7 | Lx9 | chr16 | 56112116 | 56113296 | LINE/L1 | 55975409 | 56290011 |
| Fam81a | MER5B | chr9 | 70088682 | 70088817 | DNA/hAT-Charl | 69989310 | 70241557 |
| Ccdc11 | MERV1_LTR | chr18 | 74348728 | 74349170 | LTR/ERV1 | 74183100 | 74459984 |
| Ccdc11 | MERV1_LTR | chr18 | 74230558 | 74230686 | LTR/ERV1 | 74183100 | 74459984 |
| Mbd1 | MERV1_LTR | chr18 | 74348728 | 74349170 | LTR/ERV1 | 74168288 | 74382684 |
| Mbd1 | MERV1_LTR | chr18 | 74230558 | 74230686 | LTR/ERV1 | 74168288 | 74382684 |
| Rprd1a | MERV1_LTR | chr18 | 24392446 | 24392515 | LTR/ERV1 | 24384962 | 24630204 |

| | | | | | | | |
|---|---|---|---|---|---|---|---|
| Zfp105 | MERVK26-int | chr9 | 122885310 | 122885455 | LTR/ERVK | 122823078 | 123031028 |
| Zfp105 | MERVK26-int | chr9 | 122884028 | 122884527 | LTR/ERVK | 122823078 | 123031028 |
| Zfp105 | MERVK26-int | chr9 | 122883012 | 122883537 | LTR/ERVK | 122823078 | 123031028 |
| Zfp105 | MERVK26-int | chr9 | 122880642 | 122881711 | LTR/ERVK | 122823078 | 123031028 |
| Zfp445 | MERVK26-int | chr9 | 122885310 | 122885455 | LTR/ERVK | 122748910 | 122966006 |
| Zfp445 | MERVK26-int | chr9 | 122884028 | 122884527 | LTR/ERVK | 122748910 | 122966006 |
| Zfp445 | MERVK26-int | chr9 | 122883012 | 122883537 | LTR/ERVK | 122748910 | 122966006 |
| Zfp445 | MERVK26-int | chr9 | 122880642 | 122881711 | LTR/ERVK | 122748910 | 122966006 |
| Zkscan7 | MERVK26-int | chr9 | 122885310 | 122885455 | LTR/ERVK | 122788471 | 122996124 |
| Zkscan7 | MERVK26-int | chr9 | 122884028 | 122884527 | LTR/ERVK | 122788471 | 122996124 |
| Zkscan7 | MERVK26-int | chr9 | 122883012 | 122883537 | LTR/ERVK | 122788471 | 122996124 |
| Zkscan7 | MERVK26-int | chr9 | 122880642 | 122881711 | LTR/ERVK | 122788471 | 122996124 |
| Acnat2 | MERVL-int | chr4 | 49496605 | 49498939 | LTR/ERVL | 49279845 | 49508151 |
| Acnat2 | MERVL-int | chr4 | 49493178 | 49496172 | LTR/ERVL | 49279845 | 49508151 |
| Cib3 | MIRb | chr8 | 72205989 | 72206042 | SINE/MIR | 72104335 | 72312837 |
| Grid1 | MIRb | chr14 | 35127804 | 35128031 | SINE/MIR | 34720136 | 35681115 |
| Tpm4 | MIRb | chr8 | 72205989 | 72206042 | SINE/MIR | 72035292 | 72253129 |
| Grid1 | MLT1A | chr14 | 35104767 | 35105082 | LTR/ERVL-MaLI | 34720136 | 35681115 |
| Cyp2b10 | MLT1C | chr7 | 25869624 | 25869703 | LTR/ERVL-MaLI | 25797658 | 26026624 |
| Ccdc11 | MLT2F | chr18 | 74280202 | 74280766 | LTR/ERVL | 74183100 | 74459984 |
| Mbd1 | MLT2F | chr18 | 74280202 | 74280766 | LTR/ERVL | 74168288 | 74382684 |
| Slc22a7 | MMERGLN-int | chr17 | 46350152 | 46356719 | LTR/ERV1 | 46332185 | 46538477 |
| Diap3 | MMERVK10D3_I-int | chr14 | 87205246 | 87205418 | LTR/ERVK | 86556323 | 87241114 |
| Diap3 | MMERVK10D3_I-int | chr14 | 87203192 | 87205249 | LTR/ERVK | 86556323 | 87241114 |
| Diap3 | MMERVK10D3_I-int | chr14 | 87201977 | 87202998 | LTR/ERVK | 86556323 | 87241114 |
| Nat8 | MMERVK9C_I-int | chr6 | 85880712 | 85883466 | LTR/ERVK | 85730387 | 85931890 |
| 1810064F22Rik | MMERVK9E_I-int | chr9 | 22212094 | 22213829 | LTR/ERVK | 22106786 | 22313860 |
| Mup21 | MMERVK9E_I-int | chr4 | 62196796 | 62197232 | LTR/ERVK | 62047832 | 62250841 |
| Zfp809 | MMERVK9E_I-int | chr9 | 22212094 | 22213829 | LTR/ERVK | 22125703 | 22343354 |
| Zfp872 | MMERVK9E_I-int | chr9 | 22212094 | 22213829 | LTR/ERVK | 22088166 | 22302123 |
| 2010315B03Rik | MMSAT4 | chr9 | 124269294 | 124269519 | Satellite | 124191804 | 124412694 |
| 2010315B03Rik | MMSAT4 | chr9 | 124268874 | 124269179 | Satellite | 124191804 | 124412694 |
| 2010315B03Rik | MMSAT4 | chr9 | 124268622 | 124268759 | Satellite | 124191804 | 124412694 |
| 2610305D13Rik | MMSAT4 | chr4 | 147512487 | 147515105 | Satellite | 147511936 | 147742508 |
| 6720489N17Rik | MMSAT4 | chr13 | 62639787 | 62641268 | Satellite | 62503015 | 62724182 |
| BC052688 | MMSAT4 | chr13 | 61693845 | 61693979 | Satellite | 61671580 | 61887282 |
| Gm13152 | MMSAT4 | chr4 | 147512487 | 147515105 | Satellite | 147075866 | 147613420 |
| Gm13152 | MMSAT4 | chr4 | 147178604 | 147180143 | Satellite | 147075866 | 147613420 |
| Gm4944 | MMSAT4 | chr17 | 22286643 | 22288522 | Satellite | 22097265 | 22325614 |
| Tmx3 | MMSAT4 | chr18 | 90591087 | 90591392 | Satellite | 90410154 | 90643267 |
| Zfp808 | MMSAT4 | chr13 | 62111517 | 62111656 | Satellite | 62029890 | 62273936 |
| Zfp808 | MMSAT4 | chr13 | 62049079 | 62051245 | Satellite | 62029890 | 62273936 |
| Zfp934 | MMSAT4 | chr13 | 62639787 | 62641268 | Satellite | 62416797 | 62658599 |
| BC089597 | MMVL30-int | chr10 | 127872938 | 127873861 | LTR/ERV1 | 127766476 | 127977319 |
| BC089597 | MMVL30-int | chr10 | 127871767 | 127872897 | LTR/ERV1 | 127766476 | 127977319 |
| Ccdc11 | MMVL30-int | chr18 | 74292569 | 74293992 | LTR/ERV1 | 74183100 | 74459984 |
| Csrp3 | MMVL30-int | chr7 | 48739226 | 48741569 | LTR/ERV1 | 48730398 | 48948051 |
| Mbd1 | MMVL30-int | chr18 | 74292569 | 74293992 | LTR/ERV1 | 74168288 | 74382684 |
| Zfp53 | MMVL30-int | chr17 | 21475317 | 21477755 | LTR/ERV1 | 21388988 | 21610477 |
| Zfp677 | MMVL30-int | chr17 | 21475317 | 21477755 | LTR/ERV1 | 21283748 | 21499265 |
| Gm2373 | MT2A | chr13 | 97378961 | 97379336 | LTR/ERVL | 97334970 | 97597664 |
| Acnat2 | MT2C_Mm | chr4 | 49499149 | 49499612 | LTR/ERVL | 49279845 | 49508151 |
| Cyp2a4 | MTB_Mm | chr7 | 26225854 | 26226246 | LTR/ERVL-MaLI | 26207192 | 26415088 |
| Cyp2b9 | MTB_Mm | chr7 | 26225854 | 26226246 | LTR/ERVL-MaLI | 26073409 | 26310660 |
| Afp | MTC | chr5 | 90580703 | 90581030 | LTR/ERVL-MaLI | 90390714 | 90608907 |
| Gm19705 | MTC | chr1 | 136667177 | 136667407 | LTR/ERVL-MaLI | 136583396 | 136790805 |
| 2310034O05Rik | MTE-int | chr5 | 100213526 | 100214034 | LTR/ERVL-MaLI | 100110692 | 100318049 |
| Gm13152 | MuLV-int | chr4 | 147219404 | 147226971 | LTR/ERV1 | 147075866 | 147613420 |
| C4a | MuRRS4-int | chr17 | 34805500 | 34806327 | LTR/ERV1 | 34709092 | 34923454 |
| Catsperg2 | MuRRS4-int | chr7 | 29740627 | 29742259 | LTR/ERV1 | 29597219 | 29827015 |
| Zfp84 | MuRRS4-int | chr7 | 29740627 | 29742259 | LTR/ERV1 | 29668552 | 29881419 |
| 2010315B03Rik | MurSatRep1 | chr9 | 124269966 | 124271716 | Unknown | 124191804 | 124412694 |
| 2010315B03Rik | MurSatRep1 | chr9 | 124269376 | 124269924 | Unknown | 124191804 | 124412694 |

| | | | | | | | |
|---|---|---|---|---|---|---|---|
| 2010315B03Rik | MurSatRep1 | chr9 | 124269121 | 124269324 | Unknown | 124191804 | 124412694 |
| 2010315B03Rik | MurSatRep1 | chr9 | 124268760 | 124268904 | Unknown | 124191804 | 124412694 |
| 9030624G23Rik | MurSatRep1 | chr12 | 24071624 | 24072845 | Unknown | 23943202 | 24197269 |
| AU041133 | MurSatRep1 | chr10 | 82227405 | 82227996 | Unknown | 82028013 | 82253065 |
| AU041133 | MurSatRep1 | chr10 | 82110270 | 82110900 | Unknown | 82028013 | 82253065 |
| Gm13152 | MurSatRep1 | chr4 | 147176775 | 147176879 | Unknown | 147075866 | 147613420 |
| Tmx3 | MurSatRep1 | chr18 | 90592019 | 90592248 | Unknown | 90410154 | 90643267 |
| Tmx3 | MurSatRep1 | chr18 | 90591348 | 90591982 | Unknown | 90410154 | 90643267 |
| Tmx3 | MurSatRep1 | chr18 | 90590873 | 90591145 | Unknown | 90410154 | 90643267 |
| Tmx3 | MurSatRep1 | chr18 | 90588711 | 90589448 | Unknown | 90410154 | 90643267 |
| Zfp605 | MurSatRep1 | chr5 | 110077968 | 110078732 | Unknown | 110010092 | 110229794 |
| Zfp808 | MurSatRep1 | chr13 | 62169252 | 62170909 | Unknown | 62029890 | 62273936 |
| Zfp808 | MurSatRep1 | chr13 | 62106145 | 62107137 | Unknown | 62029890 | 62273936 |
| Zfp873 | MurSatRep1 | chr10 | 82110270 | 82110900 | Unknown | 81948127 | 82161586 |
| Zfp938 | MurSatRep1 | chr10 | 82227405 | 82227996 | Unknown | 82124856 | 82341275 |
| Acnat2 | ORR1A0 | chr4 | 49488243 | 49488533 | LTR/ERVL-MaLI | 49279845 | 49508151 |
| Zfp808 | ORR1A0 | chr13 | 62112043 | 62112387 | LTR/ERVL-MaLI | 62029890 | 62273936 |
| 1810064F22Rik | ORR1A1-int | chr9 | 22202586 | 22203592 | LTR/ERVL-MaLI | 22106786 | 22313860 |
| Zfp809 | ORR1A1-int | chr9 | 22202586 | 22203592 | LTR/ERVL-MaLI | 22125703 | 22343354 |
| Zfp872 | ORR1A1-int | chr9 | 22202586 | 22203592 | LTR/ERVL-MaLI | 22088166 | 22302123 |
| Diap3 | ORR1A2 | chr14 | 87147180 | 87147476 | LTR/ERVL-MaLI | 86556323 | 87241114 |
| Zfp105 | ORR1A2 | chr9 | 122873786 | 122874095 | LTR/ERVL-MaLI | 122823078 | 123031028 |
| Zfp445 | ORR1A2 | chr9 | 122873786 | 122874095 | LTR/ERVL-MaLI | 122748910 | 122966006 |
| Zkscan3 | ORR1A2 | chr13 | 21389832 | 21390101 | LTR/ERVL-MaLI | 21287004 | 21502755 |
| Zkscan4 | ORR1A2 | chr13 | 21389832 | 21390101 | LTR/ERVL-MaLI | 21378849 | 21585505 |
| Zkscan7 | ORR1A2 | chr9 | 122873786 | 122874095 | LTR/ERVL-MaLI | 122788471 | 122996124 |
| Zfp105 | ORR1A2-int | chr9 | 122874100 | 122874920 | LTR/ERVL-MaLI | 122823078 | 123031028 |
| Zfp445 | ORR1A2-int | chr9 | 122874100 | 122874920 | LTR/ERVL-MaLI | 122748910 | 122966006 |
| Zkscan7 | ORR1A2-int | chr9 | 122874100 | 122874920 | LTR/ERVL-MaLI | 122788471 | 122996124 |
| Ccdc11 | ORR1A4 | chr18 | 74239463 | 74239734 | LTR/ERVL-MaLI | 74183100 | 74459984 |
| Gm12718 | ORR1A4 | chr4 | 103496143 | 103496440 | LTR/ERVL-MaLI | 103282500 | 103592188 |
| Mbd1 | ORR1A4 | chr18 | 74239463 | 74239734 | LTR/ERVL-MaLI | 74168288 | 74382684 |
| Tmx3 | ORR1A4 | chr18 | 90592250 | 90592582 | LTR/ERVL-MaLI | 90410154 | 90643267 |
| AU041133 | ORR1B1 | chr10 | 82046563 | 82046941 | LTR/ERVL-MaLI | 82028013 | 82253065 |
| Zfp873 | ORR1B1 | chr10 | 82046563 | 82046941 | LTR/ERVL-MaLI | 81948127 | 82161586 |
| AU041133 | ORR1B1-int | chr10 | 82046949 | 82047592 | LTR/ERVL-MaLI | 82028013 | 82253065 |
| Zfp873 | ORR1B1-int | chr10 | 82046949 | 82047592 | LTR/ERVL-MaLI | 81948127 | 82161586 |
| Gm2a | ORR1B2 | chr11 | 55123578 | 55123774 | LTR/ERVL-MaLI | 54997985 | 55213028 |
| Rpp40 | ORR1B2 | chr13 | 35958694 | 35959114 | LTR/ERVL-MaLI | 35795104 | 36006347 |
| Btnl4 | ORR1D1 | chr17 | 34531945 | 34532122 | LTR/ERVL-MaLI | 34369042 | 34575937 |
| Btnl5-ps | ORR1D1 | chr17 | 34531945 | 34532122 | LTR/ERVL-MaLI | 34387406 | 34597429 |
| Btnl6 | ORR1D1 | chr17 | 34531945 | 34532122 | LTR/ERVL-MaLI | 34407935 | 34617352 |
| Cyp2a4 | ORR1D1 | chr7 | 26225393 | 26225472 | LTR/ERVL-MaLI | 26207192 | 26415088 |
| Cyp2b9 | ORR1D1 | chr7 | 26225393 | 26225472 | LTR/ERVL-MaLI | 26073409 | 26310660 |
| Grid1 | ORR1D2 | chr14 | 35144674 | 35145029 | LTR/ERVL-MaLI | 34720136 | 35681115 |
| Zfp84 | PB1D9 | chr7 | 29878054 | 29878142 | SINE/Alu | 29668552 | 29881419 |
| Bglap | RLTR10D | chr3 | 88369595 | 88369986 | LTR/ERVK | 88283495 | 88484466 |
| Bglap3 | RLTR10D | chr3 | 88369595 | 88369986 | LTR/ERVK | 88268617 | 88469720 |
| Sema4a | RLTR10D | chr3 | 88369595 | 88369986 | LTR/ERVK | 88335962 | 88561182 |
| Diap3 | RLTR10D2 | chr14 | 87205465 | 87205804 | LTR/ERVK | 86556323 | 87241114 |
| Diap3 | RLTR10D2 | chr14 | 87201614 | 87201970 | LTR/ERVK | 86556323 | 87241114 |
| Afp | RLTR11A | chr5 | 90586505 | 90586995 | LTR/ERVK | 90390714 | 90608907 |
| Tpm4 | RLTR13D2 | chr8 | 72059739 | 72060690 | LTR/ERVK | 72035292 | 72253129 |
| Ccdc11 | RLTR14-int | chr18 | 74236164 | 74236505 | LTR/ERV1 | 74183100 | 74459984 |
| Ccdc11 | RLTR14-int | chr18 | 74232319 | 74232673 | LTR/ERV1 | 74183100 | 74459984 |
| Mbd1 | RLTR14-int | chr18 | 74236164 | 74236505 | LTR/ERV1 | 74168288 | 74382684 |
| Mbd1 | RLTR14-int | chr18 | 74232319 | 74232673 | LTR/ERV1 | 74168288 | 74382684 |
| 4930500J02Rik | RLTR16B_MM | chr2 | 104558978 | 104559658 | LTR/ERVK | 104459184 | 104671429 |
| Gm2373 | RLTR17 | chr13 | 97482567 | 97483206 | LTR/ERVK | 97334970 | 97597664 |
| Slc22a26 | RLTR17D_Mm | chr19 | 7781055 | 7781706 | LTR/ERVK | 7681981 | 7902667 |
| Slc22a27 | RLTR17D_Mm | chr19 | 7781055 | 7781706 | LTR/ERVK | 7764389 | 8066027 |
| Rprd1a | RLTR1A2_MM | chr18 | 24391961 | 24392445 | LTR/ERV1 | 24384962 | 24630204 |
| Slc22a7 | RLTR1A2_MM | chr17 | 46356725 | 46357222 | LTR/ERV1 | 46332185 | 46538477 |

| | | | | | | | |
|---|---|---|---|---|---|---|---|
| Mup21 | RLTR1E_MM | chr4 | 62200266 | 62200839 | LTR/ERV1 | 62047832 | 62250841 |
| Dpy19l3 | RLTR20C2_MM | chr7 | 35722634 | 35722721 | LTR/ERVK | 35585500 | 35854454 |
| Rprd1a | RLTR24 | chr18 | 24393456 | 24393925 | LTR/ERV1 | 24384962 | 24630204 |
| Cyp2a4 | RLTR26_Mus | chr7 | 26213117 | 26213738 | LTR/ERVK | 26207192 | 26415088 |
| Cyp2b9 | RLTR26_Mus | chr7 | 26213117 | 26213738 | LTR/ERVK | 26073409 | 26310660 |
| Bmyc | RLTR28 | chr2 | 25709093 | 25709645 | LTR/ERVL | 25606879 | 25807719 |
| Obp2a | RLTR28 | chr2 | 25709093 | 25709645 | LTR/ERVL | 25600074 | 25803326 |
| Ccdc11 | RLTR30D2_MM | chr18 | 74236743 | 74237359 | LTR/ERV1 | 74183100 | 74459984 |
| Ccdc11 | RLTR30D2_MM | chr18 | 74236540 | 74236594 | LTR/ERV1 | 74183100 | 74459984 |
| Mbd1 | RLTR30D2_MM | chr18 | 74236743 | 74237359 | LTR/ERV1 | 74168288 | 74382684 |
| Mbd1 | RLTR30D2_MM | chr18 | 74236540 | 74236594 | LTR/ERV1 | 74168288 | 74382684 |
| Ccdc11 | RLTR4_MM-int | chr18 | 74235602 | 74235738 | LTR/ERV1 | 74183100 | 74459984 |
| Mbd1 | RLTR4_MM-int | chr18 | 74235602 | 74235738 | LTR/ERV1 | 74168288 | 74382684 |
| Tpm4 | RLTR43C | chr8 | 72064466 | 72064812 | LTR/ERVK | 72035292 | 72253129 |
| Cyp2b10 | RLTR44B | chr7 | 25868293 | 25868690 | LTR/ERVK | 25797658 | 26026624 |
| Dbndd2 | RLTR44-int | chr2 | 164523963 | 164524029 | LTR/ERVK | 164386140 | 164593323 |
| Wfdc2 | RLTR44-int | chr2 | 164523963 | 164524029 | LTR/ERVK | 164462716 | 164668506 |
| BC089597 | RLTR48A | chr10 | 127873978 | 127874489 | LTR/ERV1 | 127766476 | 127977319 |
| Ccdc11 | RLTR6B_Mm | chr18 | 74267226 | 74267774 | LTR/ERV1 | 74183100 | 74459984 |
| Mbd1 | RLTR6B_Mm | chr18 | 74267226 | 74267774 | LTR/ERV1 | 74168288 | 74382684 |
| Rpa1 | RLTR6B_Mm | chr11 | 75356838 | 75357405 | LTR/ERV1 | 75200259 | 75448383 |
| Smyd4 | RLTR6B_Mm | chr11 | 75356838 | 75357405 | LTR/ERV1 | 75248433 | 75505705 |
| Bmyc | RLTR6-int | chr2 | 25719497 | 25727040 | LTR/ERV1 | 25606879 | 25807719 |
| G2e3 | RLTR6-int | chr12 | 51310300 | 51318003 | LTR/ERV1 | 51248230 | 51476986 |
| Obp2a | RLTR6-int | chr2 | 25719497 | 25727040 | LTR/ERV1 | 25600074 | 25803326 |
| Rpa1 | RLTR6-int | chr11 | 75357406 | 75365045 | LTR/ERV1 | 75200259 | 75448383 |
| Smyd4 | RLTR6-int | chr11 | 75357406 | 75365045 | LTR/ERV1 | 75248433 | 75505705 |
| Ube2l6 | RLTR6-int | chr2 | 84901752 | 84906844 | LTR/ERV1 | 84698828 | 84910003 |
| Mup21 | RLTR9A | chr4 | 62194938 | 62195253 | LTR/ERVK | 62047832 | 62250841 |
| 1810064F22Rik | RLTR9A3B | chr9 | 22205989 | 22206332 | LTR/ERVK | 22106786 | 22313860 |
| Zfp809 | RLTR9A3B | chr9 | 22205989 | 22206332 | LTR/ERVK | 22125703 | 22343354 |
| Zfp872 | RLTR9A3B | chr9 | 22205989 | 22206332 | LTR/ERVK | 22088166 | 22302123 |
| C920025E04Rik | RLTR9F | chr17 | 36108474 | 36108905 | LTR/ERVK | 36009030 | 36211676 |
| H2-Bl | RLTR9F | chr17 | 36108474 | 36108905 | LTR/ERVK | 35980189 | 36184249 |
| H2-T10 | RLTR9F | chr17 | 36108474 | 36108905 | LTR/ERVK | 36015871 | 36221444 |
| Ccdc11 | RMER10B | chr18 | 74238298 | 74238464 | LTR/ERVL | 74183100 | 74459984 |
| Mbd1 | RMER10B | chr18 | 74238298 | 74238464 | LTR/ERVL | 74168288 | 74382684 |
| Cyp2b10 | RMER13B | chr7 | 25937342 | 25937849 | LTR/ERVK | 25797658 | 26026624 |
| Zfp105 | RMER16_Mm | chr9 | 122876128 | 122876654 | LTR/ERVK | 122823078 | 123031028 |
| Zfp445 | RMER16_Mm | chr9 | 122876128 | 122876654 | LTR/ERVK | 122748910 | 122966006 |
| Zkscan7 | RMER16_Mm | chr9 | 122876128 | 122876654 | LTR/ERVK | 122788471 | 122996124 |
| Mup21 | RMER16-int | chr4 | 62198796 | 62199125 | LTR/ERVK | 62047832 | 62250841 |
| Mup21 | RMER16-int | chr4 | 62197385 | 62198705 | LTR/ERVK | 62047832 | 62250841 |
| Mettl7a3 | RMER17A | chr15 | 100345490 | 100346283 | LTR/ERVK | 100234929 | 100440169 |
| Mettl7a3 | RMER17A-int | chr15 | 100346544 | 100348057 | LTR/ERVK | 100234929 | 100440169 |
| Cyp2a4 | RMER17B | chr7 | 26223938 | 26224874 | LTR/ERVK | 26207192 | 26415088 |
| Cyp2b9 | RMER17B | chr7 | 26223938 | 26224874 | LTR/ERVK | 26073409 | 26310660 |
| Vtcn1 | RMER19A | chr3 | 100785880 | 100786500 | LTR/ERVK | 100725459 | 100996922 |
| Ccdc11 | RMER19B | chr18 | 74290041 | 74290533 | LTR/ERVK | 74183100 | 74459984 |
| G2e3 | RMER19B | chr12 | 51291833 | 51292684 | LTR/ERVK | 51248230 | 51476986 |
| H2-Q1 | RMER19B | chr17 | 35384636 | 35385309 | LTR/ERVK | 35220558 | 35425099 |
| H2-Q2 | RMER19B | chr17 | 35429133 | 35429844 | LTR/ERVK | 35242333 | 35445722 |
| H2-Q2 | RMER19B | chr17 | 35384636 | 35385309 | LTR/ERVK | 35242333 | 35445722 |
| H2-Q4 | RMER19B | chr17 | 35429133 | 35429844 | LTR/ERVK | 35279617 | 35484674 |
| H2-Q4 | RMER19B | chr17 | 35384636 | 35385309 | LTR/ERVK | 35279617 | 35484674 |
| Impg2 | RMER19B | chr16 | 56113549 | 56114356 | LTR/ERVK | 56104313 | 56373753 |
| Mbd1 | RMER19B | chr18 | 74290041 | 74290533 | LTR/ERVK | 74168288 | 74382684 |
| Senp7 | RMER19B | chr16 | 56113549 | 56114356 | LTR/ERVK | 55975409 | 56290011 |
| Tpm4 | RMER19B | chr8 | 72075967 | 72076819 | LTR/ERVK | 72035292 | 72253129 |
| Zfp105 | RMER19B2 | chr9 | 122882533 | 122883011 | LTR/ERVK | 122823078 | 123031028 |
| Zfp445 | RMER19B2 | chr9 | 122882533 | 122883011 | LTR/ERVK | 122748910 | 122966006 |
| Zkscan7 | RMER19B2 | chr9 | 122882533 | 122883011 | LTR/ERVK | 122788471 | 122996124 |
| Zfp105 | RMER1A | chr9 | 122898410 | 122898615 | Other | 122823078 | 123031028 |

| | | | | | | | |
|---|---|---|---|---|---|---|---|
| Zfp445 | RMER1A | chr9 | 122898410 | 122898615 | Other | 122748910 | 122966006 |
| Zkscan7 | RMER1A | chr9 | 122898410 | 122898615 | Other | 122788471 | 122996124 |
| 1700025F24Rik | RMER4B | chr10 | 119417474 | 119417574 | LTR/ERVK | 119313444 | 119526864 |
| Grip1 | RMER4B | chr10 | 119417474 | 119417574 | LTR/ERVK | 119354034 | 120187260 |
| Sntb1 | RMER4B | chr15 | 55959697 | 55959955 | LTR/ERVK | 55539154 | 56006949 |
| Fam184b | RMER6A | chr5 | 45517907 | 45518625 | LTR/ERVK | 45429705 | 45739501 |
| Ccdc11 | RMER6B | chr18 | 74270122 | 74270503 | LTR/ERVK | 74183100 | 74459984 |
| Mbd1 | RMER6B | chr18 | 74270122 | 74270503 | LTR/ERVK | 74168288 | 74382684 |
| Sdc1 | RSINE1 | chr12 | 8830261 | 8830412 | SINE/B4 | 8671396 | 8893687 |
| Ccdc11 | Tigger17 | chr18 | 74269645 | 74269812 | DNA/TcMar-Tig | 74183100 | 74459984 |
| Mbd1 | Tigger17 | chr18 | 74269645 | 74269812 | DNA/TcMar-Tig | 74168288 | 74382684 |
| Ccdc11 | Tigger17b | chr18 | 74270504 | 74271020 | DNA/TcMar-Tig | 74183100 | 74459984 |
| Ccdc11 | Tigger17b | chr18 | 74269841 | 74270121 | DNA/TcMar-Tig | 74183100 | 74459984 |
| Mbd1 | Tigger17b | chr18 | 74270504 | 74271020 | DNA/TcMar-Tig | 74168288 | 74382684 |
| Mbd1 | Tigger17b | chr18 | 74269841 | 74270121 | DNA/TcMar-Tig | 74168288 | 74382684 |
| Gpr98 | Tigger5b | chr13 | 81339947 | 81340346 | DNA/TcMar-Tig | 80995068 | 81733144 |
