## Supplementary material for "The mouse HP1 proteins are essential for preventing liver tumorigenesis": suppl table 9

**Supplementary table 9** : repeats with decreased expression upon loss of HP1 associated with genes down-regulated u

| gene | repeats | repeat-ch | repeat-start | repeat-end | family-repeat | gene-start | gene-end |
|---|---|---|---|---|---|---|---|
| Colec12 | MER58B | chr18 | 9880852 | 9880950 | DNA/hAT-Charlie | 9607648 | 9977995 |
| Rbp4 | MER5A | chr19 | 38121240 | 38121432 | DNA/hAT-Charlie | 38016620 | 38225321 |
| Slc27a2 | MER5B | chr2 | 126592695 | 126592830 | DNA/hAT-Charlie | 126453024 | 126688243 |
| Cyp2f2 | URR1B | chr7 | 27133728 | 27133897 | DNA/hAT-Charlie | 27019955 | 27233660 |
| Akr1c19 | L1_Mm | chr13 | 4261557 | 4262228 | LINE/L1 | 4133740 | 4348359 |
| Marcksl1-ps4 | L1_Mm | chr13 | 4261557 | 4262228 | LINE/L1 | 4148735 | 4349777 |
| Akr1c12 | L1_Mm | chr13 | 4261557 | 4262228 | LINE/L1 | 4168172 | 4379399 |
| Akr1c13 | L1_Mm | chr13 | 4261557 | 4262228 | LINE/L1 | 4091187 | 4305603 |
| Ugt2b5 | L1_Mus1 | chr5 | 87126547 | 87127063 | LINE/L1 | 87024947 | 87240340 |
| Acsm2 | L1_Mus2 | chr7 | 119668813 | 119670163 | LINE/L1 | 119454340 | 119700694 |
| Serpina3k | L1_Mus3 | chr12 | 104347895 | 104348154 | LINE/L1 | 104238486 | 104445739 |
| Serpina3k | L1_Mus3 | chr12 | 104348209 | 104350609 | LINE/L1 | 104238486 | 104445739 |
| Slco1a1 | L1_Mus3 | chr6 | 141900334 | 141901246 | LINE/L1 | 141807281 | 142046962 |
| Slco1a1 | L1_Mus3 | chr6 | 141901642 | 141903099 | LINE/L1 | 141807281 | 142046962 |
| Gulo | L1M2 | chr14 | 66080769 | 66081210 | LINE/L1 | 65886787 | 66109210 |
| Epas1 | L1M5 | chr17 | 86703761 | 86704226 | LINE/L1 | 86653864 | 86933410 |
| Ugt2b1 | L1MA8 | chr5 | 87016554 | 87016950 | LINE/L1 | 86816639 | 87026503 |
| Ugt2b35 | L1MA8 | chr5 | 87016554 | 87016950 | LINE/L1 | 86900860 | 87113274 |
| Ctcflos | L1MB5 | chr2 | 173136885 | 173137279 | LINE/L1 | 173024749 | 173233204 |
| B930025P03Rik | L1MC1 | chr8 | 10906826 | 10907328 | LINE/L1 | 10770422 | 10982454 |
| 4833411C07Rik | L1MC1 | chr8 | 10906826 | 10907328 | LINE/L1 | 10799922 | 11002334 |
| B930025P03Rik | L1MC1 | chr8 | 10907475 | 10907725 | LINE/L1 | 10770422 | 10982454 |
| 4833411C07Rik | L1MC1 | chr8 | 10907475 | 10907725 | LINE/L1 | 10799922 | 11002334 |
| B930025P03Rik | L1MC1 | chr8 | 10907785 | 10907940 | LINE/L1 | 10770422 | 10982454 |
| 4833411C07Rik | L1MC1 | chr8 | 10907785 | 10907940 | LINE/L1 | 10799922 | 11002334 |
| Colec12 | L1MC3 | chr18 | 9881634 | 9881779 | LINE/L1 | 9607648 | 9977995 |
| Colec12 | L1MC3 | chr18 | 9881802 | 9881967 | LINE/L1 | 9607648 | 9977995 |
| Gstp2 | L1MD | chr19 | 4043577 | 4043701 | LINE/L1 | 3940288 | 4142221 |
| Gstp1 | L1MD | chr19 | 4043577 | 4043701 | LINE/L1 | 3935411 | 4137912 |
| Nudt8 | L1MD | chr19 | 4043577 | 4043701 | LINE/L1 | 3900580 | 4102102 |
| Serpina3k | L1Md_A | chr12 | 104350610 | 104351590 | LINE/L1 | 104238486 | 104445739 |
| Gm10319 | L1Md_F2 | chr6 | 122143305 | 122144303 | LINE/L1 | 122036628 | 122250958 |
| Ugt3a2 | Lx | chr15 | 9372769 | 9373815 | LINE/L1 | 9235598 | 9470870 |
| Acsm2 | Lx | chr7 | 119623280 | 119626761 | LINE/L1 | 119454340 | 119700694 |
| Pdilt | Lx | chr7 | 119623280 | 119623482 | LINE/L1 | 119386587 | 119623482 |
| Acsm2 | Lx | chr7 | 119627192 | 119628983 | LINE/L1 | 119454340 | 119700694 |
| B930025P03Rik | Lx | chr8 | 10858601 | 10859105 | LINE/L1 | 10770422 | 10982454 |
| 4833411C07Rik | Lx | chr8 | 10858601 | 10859105 | LINE/L1 | 10799922 | 11002334 |
| Ugt2b5 | Lx2 | chr5 | 87130853 | 87131225 | LINE/L1 | 87024947 | 87240340 |
| Slc27a2 | Lx5c | chr2 | 126591728 | 126592067 | LINE/L1 | 126453024 | 126688243 |
| Akr1c19 | Lx6 | chr13 | 4262799 | 4263199 | LINE/L1 | 4133740 | 4348359 |
| Marcksl1-ps4 | Lx6 | chr13 | 4262799 | 4263199 | LINE/L1 | 4148735 | 4349777 |
| Akr1c12 | Lx6 | chr13 | 4262799 | 4263199 | LINE/L1 | 4168172 | 4379399 |
| Akr1c13 | Lx6 | chr13 | 4262799 | 4263199 | LINE/L1 | 4091187 | 4305603 |
| Ugt2b1 | Lx7 | chr5 | 87013798 | 87013877 | LINE/L1 | 86816639 | 87026503 |
| Ugt2b35 | Lx7 | chr5 | 87013798 | 87013877 | LINE/L1 | 86900860 | 87113274 |
| Ugt2b1 | Lx7 | chr5 | 87013946 | 87014092 | LINE/L1 | 86816639 | 87026503 |
| Ugt2b35 | Lx7 | chr5 | 87013946 | 87014092 | LINE/L1 | 86900860 | 87113274 |
| Cyp3a59 | Lx8 | chr5 | 146115586 | 146115974 | LINE/L1 | 145979258 | 146213283 |
| Egfr | Lx9 | chr11 | 16889083 | 16889577 | LINE/L1 | 16652203 | 17013907 |
| Ugt2b1 | Lx9 | chr5 | 87015721 | 87016206 | LINE/L1 | 86816639 | 87026503 |
| Ugt2b35 | Lx9 | chr5 | 87015721 | 87016206 | LINE/L1 | 86900860 | 87113274 |
| Gm10768 | LTRIS_Mm | chr19 | 43840861 | 43841342 | LTR/ERV1 | 43738803 | 43940845 |
| H2-Q10 | MMVL30-i | chr17 | 35480336 | 35482627 | LTR/ERV1 | 35370089 | 35574563 |
| Slc27a2 | MuRRS-int | chr2 | 126608834 | 126613401 | LTR/ERV1 | 126453024 | 126688243 |
| Acsm2 | RLTR1D | chr7 | 119544868 | 119545406 | LTR/ERV1 | 119454340 | 119700694 |
| Pdilt | RLTR1D | chr7 | 119544868 | 119545406 | LTR/ERV1 | 119386587 | 119623482 |
| Kifc2 | RLTR4_MM | chr15 | 76560641 | 76564037 | LTR/ERV1 | 76560641 | 76768196 |
| Dgat1 | RLTR4_MM | chr15 | 76560514 | 76564037 | LTR/ERV1 | 76402015 | 76611818 |

| | | | | | | | |
|---|---|---|---|---|---|---|---|
| Cyhr1 | RLTR4_MM | chr15 | 76560514 | 76564037 | LTR/ERV1 | 76543395 | 76760208 |
| Adck5 | RLTR4_MM | chr15 | 76560514 | 76564037 | LTR/ERV1 | 76476359 | 76695811 |
| Slc52a2 | RLTR4_MM | chr15 | 76560514 | 76564037 | LTR/ERV1 | 76438943 | 76642130 |
| Gstp2 | RLTR6_Mm | chr19 | 4045069 | 4045631 | LTR/ERV1 | 3940288 | 4142221 |
| Gstp1 | RLTR6_Mm | chr19 | 4045069 | 4045631 | LTR/ERV1 | 3935411 | 4137912 |
| Nudt8 | RLTR6_Mm | chr19 | 4045069 | 4045631 | LTR/ERV1 | 3900580 | 4102102 |
| Mapk15 | RLTR11A | chr15 | 75958424 | 75958932 | LTR/ERVK | 75893769 | 76099153 |
| Cyp2d13 | RLTR13D3A | chr15 | 82635520 | 82635800 | LTR/ERVK | 82536750 | 82742045 |
| Cyp2d37-ps | RLTR13D3A | chr15 | 82635520 | 82635800 | LTR/ERVK | 82588750 | 82790058 |
| Cyp2d13 | RLTR13D3A | chr15 | 82635948 | 82636628 | LTR/ERVK | 82536750 | 82742045 |
| Cyp2d37-ps | RLTR13D3A | chr15 | 82635948 | 82636628 | LTR/ERVK | 82588750 | 82790058 |
| Rdh9 | RLTR13D6 | chr10 | 127880862 | 127881663 | LTR/ERVK | 127676405 | 127892697 |
| Gm4956 | RLTR20A3_ | chr1 | 21263652 | 21263920 | LTR/ERVK | 21185246 | 21398312 |
| Slc22a28 | RLTR20B5_ | chr19 | 8114881 | 8115330 | LTR/ERVK | 7962209 | 8231982 |
| Cyp3a25 | RLTR9D | chr5 | 145976592 | 145976992 | LTR/ERVK | 145877194 | 146109618 |
| Cyp3a59 | RLTR9D | chr5 | 146113489 | 146113889 | LTR/ERVK | 145979258 | 146213283 |
| Cyp2c44 | RLTR9F | chr19 | 44025902 | 44026336 | LTR/ERVK | 43905022 | 44129247 |
| Colec12 | RLTRETN_M | chr18 | 9882147 | 9882472 | LTR/ERVK | 9607648 | 9977995 |
| Cyp2c29 | RMER17A2 | chr19 | 39328136 | 39328891 | LTR/ERVK | 39187085 | 39430713 |
| Cyp2c53-ps | RMER17A2 | chr19 | 39328136 | 39328891 | LTR/ERVK | 39129254 | 39374737 |
| Akr1c19 | RMER17B | chr13 | 4257362 | 4258219 | LTR/ERVK | 4133740 | 4348359 |
| Marcksl1-ps4 | RMER17B | chr13 | 4257362 | 4258219 | LTR/ERVK | 4148735 | 4349777 |
| Akr1c12 | RMER17B | chr13 | 4257362 | 4258219 | LTR/ERVK | 4168172 | 4379399 |
| Akr1c13 | RMER17B | chr13 | 4257362 | 4258219 | LTR/ERVK | 4091187 | 4305603 |
| Gcgr | RMER19C | chr11 | 120508417 | 120508922 | LTR/ERVK | 120430727 | 120638984 |
| Pbld2 | RMER6A | chr10 | 63059617 | 63060330 | LTR/ERVK | 62924512 | 63158812 |
| Cyp2c44 | RMER6C | chr19 | 44009230 | 44009756 | LTR/ERVK | 43905022 | 44129247 |
| Cyp3a59 | RMER6D | chr5 | 146094772 | 146095083 | LTR/ERVK | 145979258 | 146213283 |
| Cyp3a25 | RMER6D | chr5 | 146094772 | 146095083 | LTR/ERVK | 145877194 | 146109618 |
| Cyp3a59 | RMER6D | chr5 | 146095903 | 146096225 | LTR/ERVK | 145979258 | 146213283 |
| Cyp3a25 | RMER6D | chr5 | 146095903 | 146096225 | LTR/ERVK | 145877194 | 146109618 |
| Slc22a28 | MERVL_2A | chr19 | 8115363 | 8115659 | LTR/ERVL | 7962209 | 8231982 |
| H2-Q10 | MLT2F | chr17 | 35472469 | 35472587 | LTR/ERVL | 35370089 | 35574563 |
| Slc22a28 | MT2B1 | chr19 | 8115719 | 8116164 | LTR/ERVL | 7962209 | 8231982 |
| Pbld2 | RMER15 | chr10 | 63046386 | 63046480 | LTR/ERVL | 62924512 | 63158812 |
| H2-Q10 | RMER15 | chr17 | 35476452 | 35476654 | LTR/ERVL | 35370089 | 35574563 |
| Slc3a1 | MLT1N2 | chr17 | 85067642 | 85067821 | LTR/ERVL-MaLR | 84928347 | 85164241 |
| Slco1a1 | MTA_Mm | chr6 | 141901247 | 141901641 | LTR/ERVL-MaLR | 141807281 | 142046962 |
| Cyp3a59 | MTC | chr5 | 146000593 | 146000995 | LTR/ERVL-MaLR | 145979258 | 146213283 |
| Cyp3a25 | MTC | chr5 | 146000593 | 146000995 | LTR/ERVL-MaLR | 145877194 | 146109618 |
| Slco1a1 | MTD | chr6 | 141946773 | 141947184 | LTR/ERVL-MaLR | 141807281 | 142046962 |
| Cyp3a25 | MTE2a | chr5 | 145976128 | 145976483 | LTR/ERVL-MaLR | 145877194 | 146109618 |
| Cyp3a59 | MTE2a | chr5 | 146113992 | 146114353 | LTR/ERVL-MaLR | 145979258 | 146213283 |
| Cyp2d13 | ORR1A2 | chr15 | 82684706 | 82685021 | LTR/ERVL-MaLR | 82536750 | 82742045 |
| Cyp2d37-ps | ORR1A2 | chr15 | 82684706 | 82685021 | LTR/ERVL-MaLR | 82588750 | 82790058 |
| Cyp2d40 | ORR1A2 | chr15 | 82684706 | 82685021 | LTR/ERVL-MaLR | 82659833 | 82864122 |
| BC048546 | ORR1A2 | chr6 | 128482531 | 128482846 | LTR/ERVL-MaLR | 128439822 | 128681606 |
| Cyp2d13 | ORR1A2-in | chr15 | 82685028 | 82686602 | LTR/ERVL-MaLR | 82536750 | 82742045 |
| Cyp2d37-ps | ORR1A2-in | chr15 | 82685028 | 82686602 | LTR/ERVL-MaLR | 82588750 | 82790058 |
| Cyp2d40 | ORR1A2-in | chr15 | 82685028 | 82686602 | LTR/ERVL-MaLR | 82659833 | 82864122 |
| Cyp3a59 | ORR1B1 | chr5 | 146104997 | 146105405 | LTR/ERVL-MaLR | 145979258 | 146213283 |
| Cyp3a25 | ORR1B1 | chr5 | 146104997 | 146105405 | LTR/ERVL-MaLR | 145877194 | 146109618 |
| Sftpa1 | ORR1B2 | chr14 | 41140091 | 41140428 | LTR/ERVL-MaLR | 41031788 | 41236373 |
| Cbx1 | (A)n | chr11 | 96807371 | 96807396 | Simple | 96689136 | 96908640 |
| Slc3a1 | (ACATG)n | chr17 | 85066874 | 85067030 | Simple | 84928347 | 85164241 |
| Hgd | (CAAAA)n | chr16 | 37634158 | 37634208 | Simple | 37480153 | 37732026 |
| Rnf152 | (GAGAA)n | chr1 | 105361626 | 105361713 | Simple | 105176917 | 105456710 |
| Cyp3a59 | (TGG)n | chr5 | 146095744 | 146095872 | Simple | 145979258 | 146213283 |
| Cyp3a25 | (TGG)n | chr5 | 146095744 | 146095872 | Simple | 145877194 | 146109618 |
| Gmnn | (TTTTTG)n | chr13 | 24813773 | 24813802 | Simple | 24651845 | 24861937 |
| Chpt1 | B1_Mur1 | chr10 | 88469667 | 88469812 | SINE/Alu | 88372813 | 88603970 |
| Ctcflos | B1_Mur3 | chr2 | 173142965 | 173143110 | SINE/Alu | 173024749 | 173233204 |
| H2-Q10 | B1_Mus1 | chr17 | 35475761 | 35475907 | SINE/Alu | 35370089 | 35574563 |

| | | | | | | | |
|---|---|---|---|---|---|---|---|
| Psmd9 | B1_Mus2 | chr5 | 123180304 | 123180449 | SINE/Alu | 123128190 | 123350125 |
| Hpd | B1_Mus2 | chr5 | 123180304 | 123180449 | SINE/Alu | 123071807 | 123282686 |
| 0610005C13Rik | B1_Mus2 | chr7 | 45471376 | 45471501 | SINE/Alu | 45467795 | 45675176 |
| Ftl1 | B1_Mus2 | chr7 | 45471376 | 45471501 | SINE/Alu | 45357944 | 45559886 |
| Slc27a2 | B1F | chr2 | 126593184 | 126593290 | SINE/Alu | 126453024 | 126688243 |
| Ces1e | B1F2 | chr8 | 93185291 | 93185412 | SINE/Alu | 93101218 | 93329619 |
| Gstp2 | PB1D7 | chr19 | 4036195 | 4036289 | SINE/Alu | 3940288 | 4142221 |
| Gstp1 | PB1D7 | chr19 | 4036195 | 4036289 | SINE/Alu | 3935411 | 4137912 |
| Nudt8 | PB1D7 | chr19 | 4036195 | 4036289 | SINE/Alu | 3900580 | 4102102 |
| Gm20337 | B2_Mm1a | chr12 | 80930730 | 80930917 | SINE/B2 | 80842652 | 81045399 |
| Slc27a2 | B2_Mm1a | chr2 | 126589437 | 126589644 | SINE/B2 | 126453024 | 126688243 |
| Abcg8 | B2_Mm1t | chr17 | 84657607 | 84657774 | SINE/B2 | 84576302 | 84800333 |
| Abcg8 | B2_Mm2 | chr17 | 84657408 | 84657522 | SINE/B2 | 84576302 | 84800333 |
| Slc27a2 | B2_Mm2 | chr2 | 126595757 | 126595960 | SINE/B2 | 126453024 | 126688243 |
| Aox1 | B3 | chr1 | 58201688 | 58201899 | SINE/B2 | 57929969 | 58206410 |
| March2 | B3 | chr17 | 33774515 | 33774701 | SINE/B2 | 33585692 | 33818677 |
| H2-Q10 | B3 | chr17 | 35475318 | 35475504 | SINE/B2 | 35370089 | 35574563 |
| Etnppl | B3 | chr3 | 130636854 | 130637007 | SINE/B2 | 130517448 | 130735750 |
| Rcl1 | B3A | chr19 | 29019939 | 29020120 | SINE/B2 | 29001375 | 29243843 |
| Mug1 | B3A | chr6 | 121886838 | 121886967 | SINE/B2 | 121738541 | 121989057 |
| Cyp3a59 | B4A | chr5 | 146115003 | 146115160 | SINE/B4 | 145979258 | 146213283 |
| Ctcflos | ID_B1 | chr2 | 173147710 | 173147891 | SINE/B4 | 173024749 | 173233204 |
| Errfi1 | ID_B1 | chr4 | 150869049 | 150869252 | SINE/B4 | 150755091 | 150968880 |
| Psmd9 | ID_B1 | chr5 | 123180103 | 123180190 | SINE/B4 | 123128190 | 123350125 |
| Hpd | ID_B1 | chr5 | 123180103 | 123180190 | SINE/B4 | 123071807 | 123282686 |
| Pbld2 | RSINE1 | chr10 | 63077707 | 63077858 | SINE/B4 | 62924512 | 63158812 |
| Mtl5 | RSINE1 | chr19 | 3387798 | 3387958 | SINE/B4 | 3288857 | 3507823 |
| Ctcflos | RSINE1 | chr2 | 173138083 | 173138218 | SINE/B4 | 173024749 | 173233204 |
| Mup12 | MIR | chr4 | 60661330 | 60661403 | SINE/MIR | 60637381 | 60841281 |
| Psmd9 | MIR | chr5 | 123179915 | 123180015 | SINE/MIR | 123128190 | 123350125 |
| Hpd | MIR | chr5 | 123179915 | 123180015 | SINE/MIR | 123071807 | 123282686 |
| Ces3a | MIR | chr8 | 105056840 | 105056916 | SINE/MIR | 104948599 | 105158414 |
| Psmd9 | MIRb | chr5 | 123179783 | 123179836 | SINE/MIR | 123128190 | 123350125 |
| Hpd | MIRb | chr5 | 123179783 | 123179836 | SINE/MIR | 123071807 | 123282686 |

pon loss of HP1
