## Supplementary material for "The mouse HP1 proteins are essential for preventing liver tumorigenesis": suppl table 10

**Supplementary table10** : list of the oligonucleotides used in this study

| | | | |
|---|---|---|---|
| cre | sense | Genotyping | ATTGCCTGCATTACCGGTC |
| cre | antisense | Genotyping | ATCAACGTTTTGTTTTCGGA |
| cbx5 | sense | Genotyping | CTCAAGCCTATGGCACTGCATATGC |
| cbx5 | antisense1 | Genotyping | AGGATAATGCCCTGTTTTCACCCAC |
| cbx5 | antisense2 | Genotyping | GATTCTCAGTTCGGTTGGCAG |
| cbx3 | sense | Genotyping | GAGTGGTGACTCTGGTAATTTGGTC |
| cbx3 | antisense1 | Genotyping | CCCTTTAAACTTCCTTGATCCAGTG |
| cbx3 | antisense2 | Genotyping | GTCCAAGTAACAGTTGTGGTGCTG |
| cbx1 | sense | Genotyping | CAGAAATTGTTCCCACACATTGTAG |
| cbx1 | antisense1 | Genotyping | CATGATGGTAATGTTTGCTAGGGGT |
| cbx1 | antisense2 | Genotyping | TCAGGCCGAGGGTCACTATCGAAGG |
| Cyp2c29 | sense | RT-qPCR | TTTTTCAGCCATTGGAAAGC |
| Cyp2c29 | antisense | RT-qPCR | TGGGCTCAAAGCCTACTGTC |
| Nox4 | sense | RT-qPCR | AACCTCAACTGCAGCCTCAT |
| Nox4 | antisense | RT-qPCR | GTTGAGGGCATTCACCAAGT |
| cyp2b10 | sense | RT-qPCR | CACACAGCATAACCACAGGC |
| cyp2b10 | antisense | RT-qPCR | TGCAGATGGACAGAGGAGG |
| Ifit2 | sense | RT-qPCR | TCTAACAGCTGTATCAGTGC |
| Ifit2 | antisense | RT-qPCR | CAGGGGAACCAGAGCATAGC |
| Rsl1 | sense | RT-qPCR | ACTCGTGCGTCTTTCTTAAGG |
| Rsl1 | antisense | RT-qPCR | GACAGCAAGACCCAGGAACA |
| zfp345 | sense | RT-qPCR | TGTAAGTCTCCGGCTCCTCT |
| zfp345 | antisense | RT-qPCR | ACTGTGGTATGCAAATCTTGG |
| zfp445 | sense | RT-qPCR | CGCAGCATCAGCATCTCTCT |
| zfp445 | antisense | RT-qPCR | TAGTTTCTGTGGTGGGCTCC |
| zkscan3 | sense | RT-qPCR | GTCAGAGTCACTCATGTGGGG |
| zkscan3 | antisense | RT-qPCR | TCCATCTGGGGGAAAGGACT |
| Trim28 | sense | RT-qPCR | TTGCTCACTCTGCCACGTGCTCCC |
| Trim28 | antisense | RT-qPCR | GCGAGCACGAATCAAGGTCAG |
| 261Rik | sense | RT-qPCR | GTGGTCAGGAAGGGATGGAA |
| 261Rik | antisense | RT-qPCR | ATGGAGACGTGGAAGGCAAC |
| Mbd1 | sense | RT-qPCR | TCGTTGGCGCCAGTGCCTGC |
| Mbd1 | antisense | RT-qPCR | AGCCTCTACCTGCTTGCTGC |
| Bglap3 | sense | RT-qPCR | CTGACAAAGCCTTCATGTCC |
| Bglap3 | antisense | RT-qPCR | TCAAGCTCACATAGCTCCC |
| majSatF | sense | RT-qPCR | ATATGTTGAGAAAACTGAAAATCACG |
| majSatR | antisense | RT-qPCR | CCTTCAGTGTGCATTTCTCATTTTTCAC |
