## Supplementary material for "The mouse HP1 proteins are essential for preventing liver tumorigenesis": suppl fig1

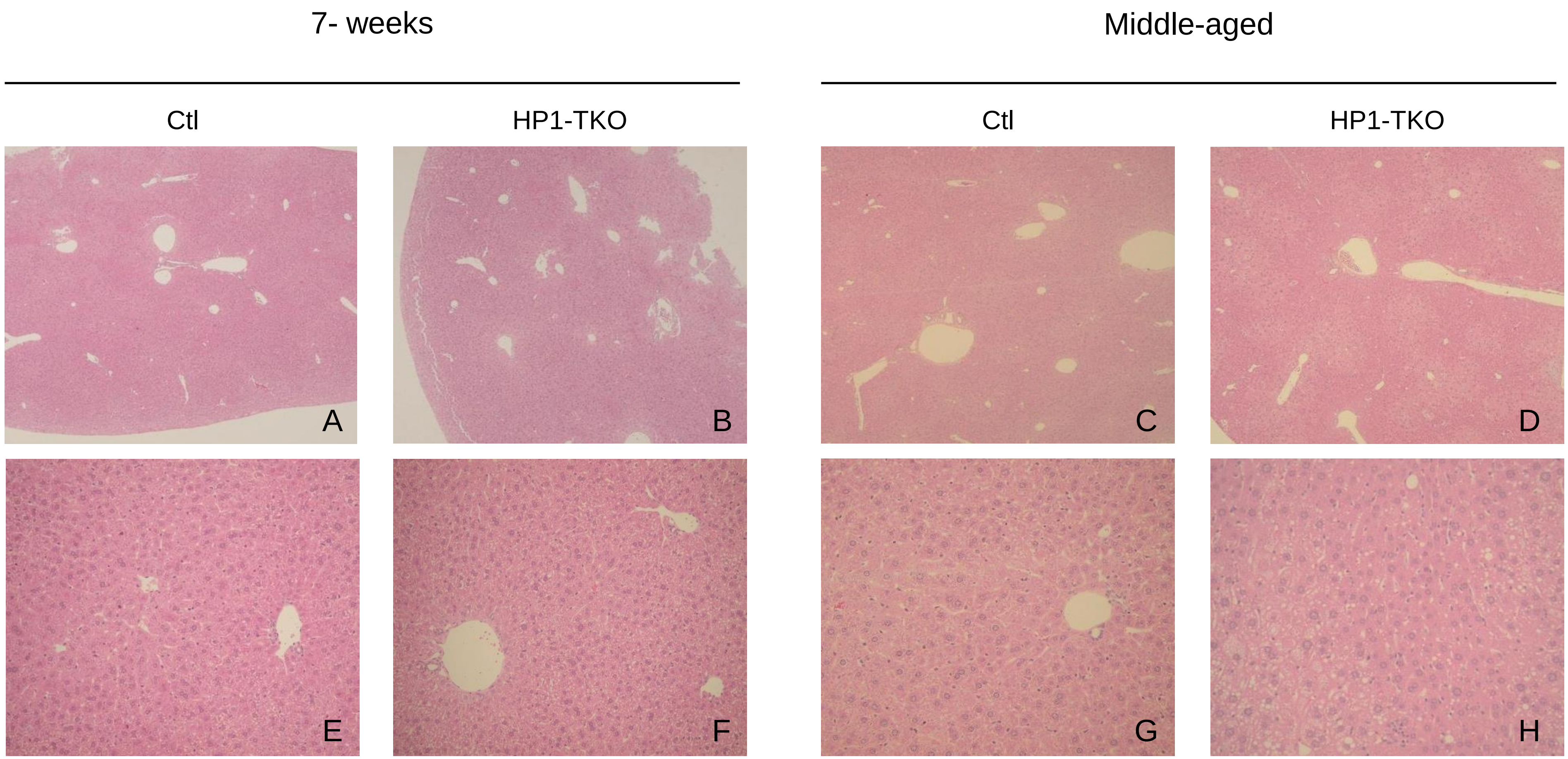


Supplementary figure 1 : The absence of HP1 in hepatocytes did not induce any significant histological alteration in the livers of young (7-week-old) and middle-aged (3-6-month-old) HP1-TKO mice. (A-D) low magnification, (E-H) high magnification.

Saksouk & Hajdari et al,
