## Supplementary material for "The mouse HP1 proteins are essential for preventing liver tumorigenesis": suppl fig2

A

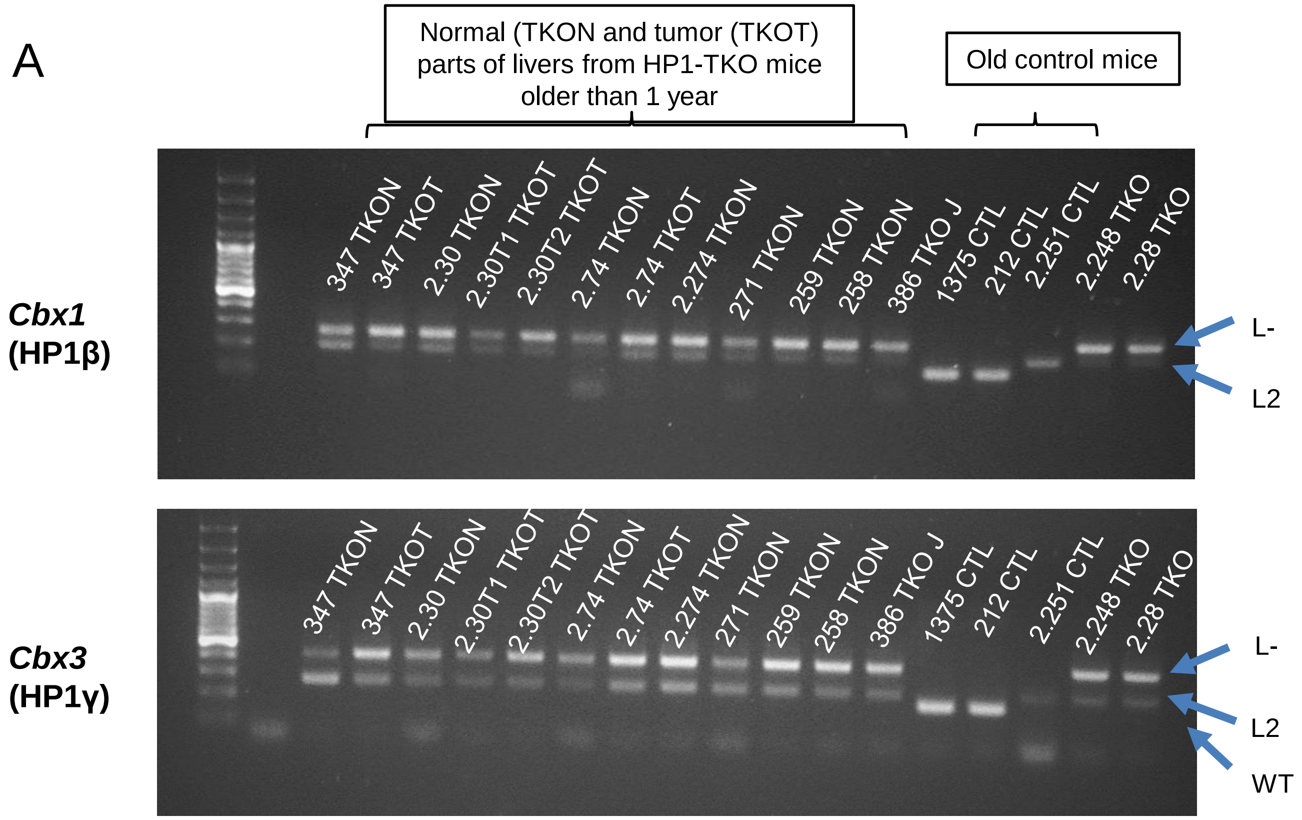


B

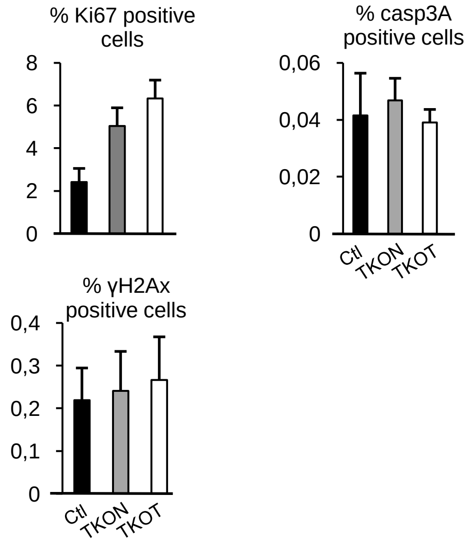


**Supplementary Figure 2**: (A) Excision of the *Cbx1* and *Cbx3* genes in the liver of old (> 1 year) HP1-TKO mice (TKON: normal part of liver; TKOT: tumor part of liver) compared with age-matched controls (CTL). (B) Quantification of Ki67-, caspase 3A- and γH2AX-positive cells on liver Tissue Micro Areas of old HP1-TKO mice and age-matched controls.
