## Supplementary material for "The mouse HP1 proteins are essential for preventing liver tumorigenesis": suppl methods

Supplementary methods

RNA-seq:

Total RNA from liver samples from 7-week-old control and HP1-KO mice was used for library preparation using the Illumina TruSeq Stranded mRNA Sample Preparation kit. Polyadenylated RNA was purified using magnetic oligo(dT) beads and then fragmented into small pieces using divalent cations at elevated temperature. The cleaved RNA fragments were copied into first-strand cDNA using reverse transcriptase and random primers in the presence of actinomycin D. Second-strand cDNA synthesis was performed using dUTP, instead of dTTP, resulting in blunt double-stranded cDNA fragments. A single 'A' nucleotide was added to the 3' ends of the blunt DNA fragments using a Klenow fragment (3' to 5'exo minus). The cDNA fragments were ligated to double-stranded TruSeq Universal adapters using T4 DNA Ligase. The ligated products were enriched by PCR amplification (98°C for 30sec, followed by 15 cycles of 98°C for 10sec, 60°C for 30sec, and 72°C for 30sec; finally, 72°C for 5min). Then, the excess of PCR primers was removed by purification using AMPure XP beads (Agencourt Biosciences Corporation). The quality and quantity of the final cDNA libraries were checked using a Fragment Analyzer (AATI) and the KAPA Library Quantification Kit (Roche), respectively. Libraries were equimolarly pooled and sequenced (50nt single read per lane) on a Hiseq2500 apparatus, according to the manufacturer's instructions. Image analysis and base calling were performed using the Illumina HiSeq Control software and the Illumina RTA software. Reads not mapping to rRNA sequences were mapped onto the mouse genome mm10 assembly using the Tophat mapper (v2.0.10 58) and the Bowtie2 (v2.1.0) aligner. Gene expression was quantified from uniquely aligned reads using HTSeq v0.5.4p3 59 and gene annotations from the Ensembl release 75.

Statistical analyses were performed using the method previously described [65] implemented in the DESeq2 Bioconductor library (v1.0.19), taking into account the batch effect. Adjustment for multiple testing was performed with the Hochberg and Benjamini method [66].

For repeat analysis, alignment positions were compared with repeats annotated in UCSC with RepeatMasker (rmsk table from mm10) and overlaps were kept relative to the strand.

Among the multiple mapped reads, only those mapping to the same repeat type were kept. Repeat types that were significantly different between conditions were identified with DESeq2 v1.0.19. Repeats found in exons were removed from the analysis because it was not possible to discriminate between differential gene and repeat expression.
